# Suprabasal cells retaining stem cell identity programs drive basal cell hyperplasia in eosinophilic esophagitis

**DOI:** 10.1101/2023.04.20.537495

**Authors:** Margarette H. Clevenger, Adam L. Karami, Dustin A. Carlson, Peter J. Kahrilas, Nirmala Gonsalves, John E. Pandolfino, Deborah R. Winter, Kelly A. Whelan, Marie-Pier Tétreault

## Abstract

Eosinophilic esophagitis (EoE) is an esophageal immune-mediated disease characterized by eosinophilic inflammation and epithelial remodeling, including basal cell hyperplasia (BCH) and loss of differentiation. Although BCH correlates with disease severity and with persistent symptoms in patients in histological remission, the molecular processes driving BCH remain poorly defined. Here, we demonstrate that despite the presence of BCH in all EoE patients examined, no increase in basal cell proportion was observed by scRNA-seq. Instead, EoE patients exhibited a reduced pool of *KRT15+ COL17A1+* quiescent cells, a modest increase in *KI67+* dividing epibasal cells, a substantial increase in *KRT13+ IVL+* suprabasal cells, and a loss of differentiated identity in superficial cells. Suprabasal and superficial cell populations demonstrated increased quiescent cell identity scoring in EoE with the enrichment of signaling pathways regulating pluripotency of stem cells. However, this was not paired with increased proliferation. Enrichment and trajectory analyses identified SOX2 and KLF5 as potential drivers of the increased quiescent identity and epithelial remodeling observed in EoE. Notably, these findings were not observed in GERD. Thus, our study demonstrates that BCH in EoE results from an expansion of non-proliferative cells that retain stem-like transcriptional programs while remaining committed to early differentiation.

## Introduction

Eosinophilic esophagitis (EoE) is a type 2 immune-mediated esophageal disease driven by the response to food allergens leading to eosinophilia and ultimately dysphagia, edema, esophageal stricture, and food impaction. Current management of EoE includes proton pump inhibitors, topical corticosteroids, diet elimination and dupilumab (1, 2). However, despite improvements in the treatment of EoE, a significant subset of patients experiences symptom recurrence or does not respond to these therapeutic options (3, 4), leading to poor prognosis, reduced quality of life, and substantial healthcare costs associated with the need for procedures and life-long therapy (5). Hence, current efforts to improve therapeutic options focus on the alleviation of symptoms and prevention of complications.

In the esophagus, primary protection against food antigens passing through the lumen is provided by the stratified squamous epithelial barrier. Esophageal epithelial cells (EEC) arise from the basal layer stem cells and undergo coordinated differentiation as they rise toward the lumen to ultimately desquamate. The terminal differentiation process occurring in the superficial compartment is intricately tied to the process of cornification, which contributes to the efficacy of the epithelial barrier. Upon damage to the epithelial barrier, a rapid restoration of epithelial homeostasis is achieved through balanced self-renewal and differentiation of stem/progenitor cells. Dysregulated inflammation, aberrant tissue repair mechanisms or failure to restore homeostasis will ultimately have pathological consequences.

Alterations to EEC are a primary driver of EoE (6) and include intraepithelial eosinophilic inflammation, basal cell hyperplasia (BCH), dilatation of intercellular space, and dysregulated terminal differentiation (7, 8). Histologically, BCH is the most prominent epithelial change in EoE and is defined by pathologists as an expansion of EEC within the basal zone (9). Despite the predominant incidence of BCH in EoE, the changes in the molecular and cellular identity occurring in BCH are largely unexplored. Highlighting the importance of understanding the role of BCH in EoE pathogenesis, BCH is linked to disease severity in EoE and correlates with persistent symptoms (odds ratio, 2.14; 95% CI, 1.03–4.42; *P* = .041) and endoscopic findings (odds ratio, 7.10; 95% CI, 3.12–16.18; *P* < .001) in patients in histologic remission (9). Thus, the molecular characterization of BCH is needed to improve the current understanding of symptom recurrence and persistent endoscopic findings in EoE. This will ultimately guide the development of novel therapeutic approaches for EoE, particularly for cases where reducing eosinophilic inflammation is not sufficient to restore epithelial tissue integrity or to improve clinical symptoms.

To address this gap in knowledge and investigate the molecular changes occurring in BCH, we performed scRNA-seq of esophageal mucosal biopsies from treatment-naive adult EoE patients and healthy controls. We uncovered that BCH in EoE mainly involves the expansion of early differentiated cells with non-proliferative stem cell identity. We identified the transcription factors and stem cell renewal regulators SOX2 and KLF5 as top predicted regulators of differentially expressed genes (DEGs) in these aberrant EEC in EoE. Elevated SOX2 and KLF5 levels and their downstream targets were confirmed in early differentiated EEC in EoE. Lastly, these changes were not observed in GERD.

## Results

### Characterization of esophageal mucosal cell populations in adult EoE

To define the single-cell transcriptomic landscape of the esophageal mucosa in EoE, proximal and distal esophageal mucosal biopsies were obtained from 6 adults with EoE along with 6 healthy controls (HC) (**Figure 1A**). Additional adjacent biopsies were collected and processed for histology (**Figure 1A**). To validate scRNA-seq findings using immunostaining, biopsies were also collected from 22 additional EoE subjects and 16 HC. Patient characteristics and demographics are summarized in **Table 1**. Freshly collected tissue specimens were digested to generate single-cell suspensions and sequenced using the 10X Genomics platform. After quality control filtering, integration was performed across all samples by reciprocal principal component analysis dimensional reduction. Following the calculation of Uniform Manifold Approximation and Projection for Dimension Reduction (UMAP) embeddings using the Seurat R package (10), unsupervised graph-based clustering was performed. Clusters were annotated based on established marker genes (**Figure 1B-C**) and transcriptional signatures (**Supplemental Figure 1A**).

**Figure 1.**
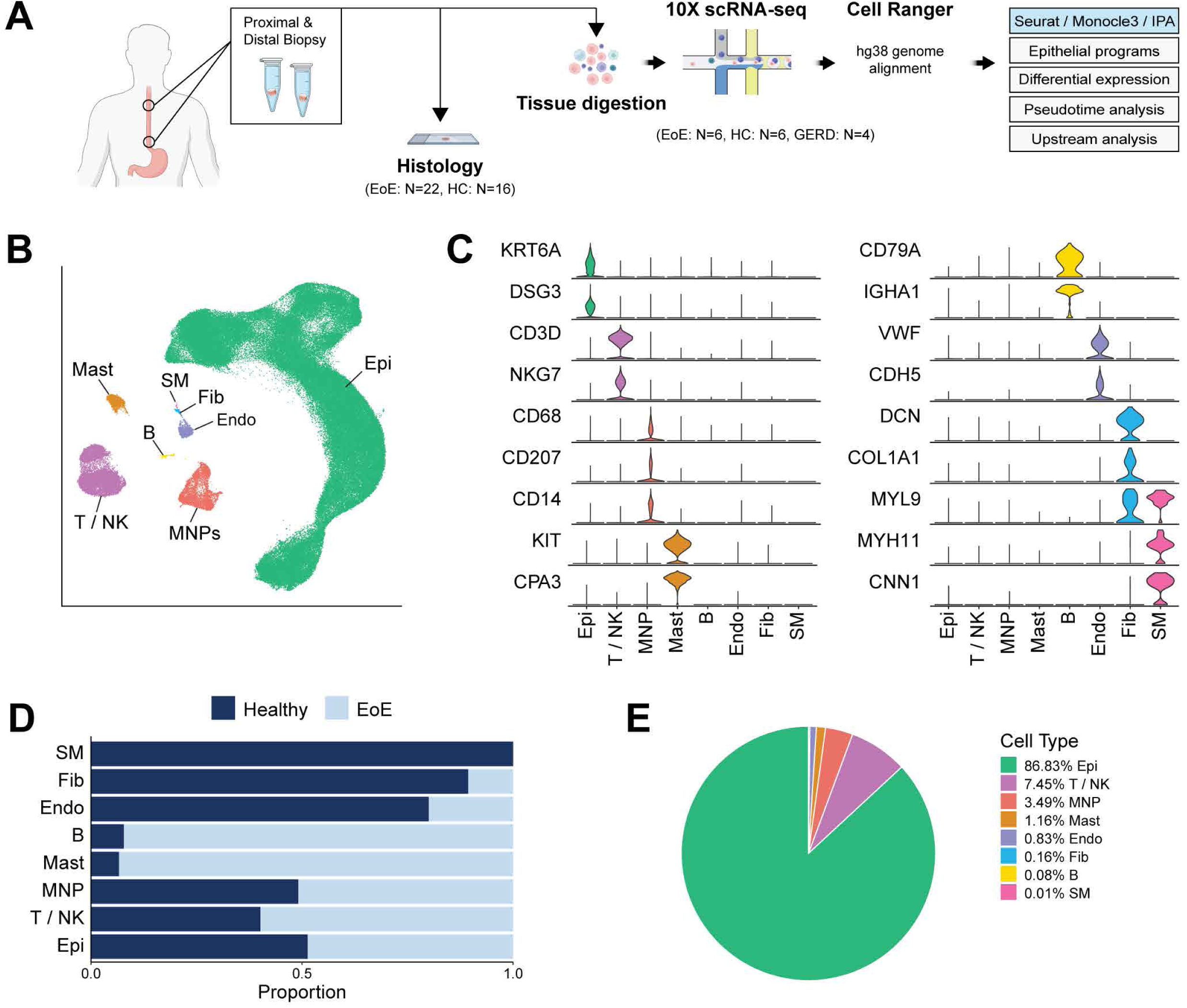
Single-cell transcriptomic landscape of esophageal mucosal cells in EoE and healthy subjects. **(A)** Schematic of study design. **(B)** UMAP embedding of the unsupervised clustering of cells from HC and EoE. **(C)** Violin plots illustrating the expression of established cell-type specific markers. **(D)** Bar plot displaying the frequency of each esophageal cell type in HC and EoE. **(E)** Pie chart demonstrating the proportion of each esophageal cell type. Epithelial cells (Epi), T cells/ natural killer cells (T/NK), mononuclear phagocytes (MNP), mast cells (Mast), B cells (B), endothelial cells (Endo), fibroblasts (Fib), smooth muscle (SM).

**Table 1:**
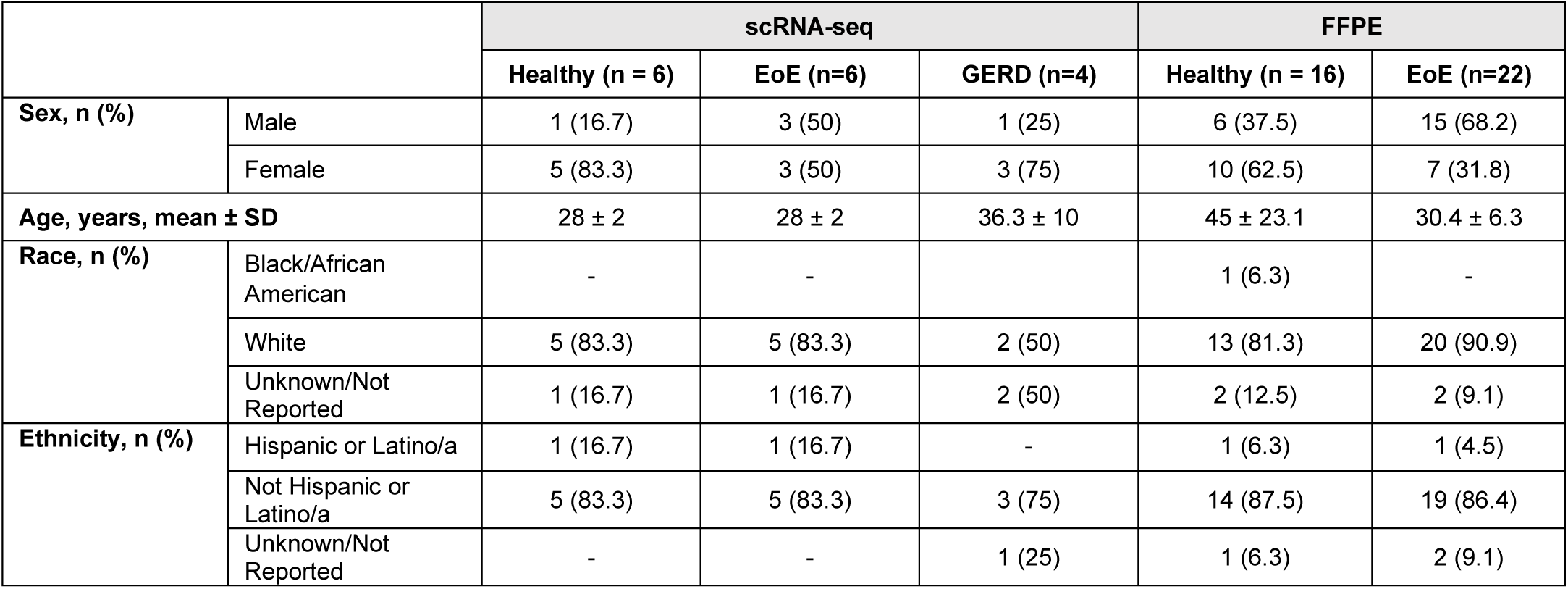
Patient demographics summary.

|  |  | scRNA-seq |  |  | FFPE |  |
| --- | --- | --- | --- | --- | --- | --- |
|  |  | Healthy (n = 6) | EoE (n=6) | GERD (n=4) | Healthy (n = 16) | EoE (n=22) |
| **Sex, n (%)** | Male | 1 (16.7) | 3 (50) | 1 (25) | 6 (37.5) | 15 (68.2) |
|  | Female | 5 (83.3) | 3 (50) | 3 (75) | 10 (62.5) | 7 (31.8) |
| **Age, years, mean ± SD** | | 28 ± 2 | 28 ± 2 | 36.3 ± 10 | 45 ± 23.1 | 30.4 ± 6.3 |
| **Race, n (%)** | Black/African American | - | - |  | 1 (6.3) | - |
|  | White | 5 (83.3) | 5 (83.3) | 2 (50) | 13 (81.3) | 20 (90.9) |
|  | Unknown/Not Reported | 1 (16.7) | 1 (16.7) | 2 (50) | 2 (12.5) | 2 (9.1) |
| **Ethnicity, n (%)** | Hispanic or Latino/a | 1 (16.7) | 1 (16.7) | - | 1 (6.3) | 1 (4.5) |
|  | Not Hispanic or Latino/a | 5 (83.3) | 5 (83.3) | 3 (75) | 14 (87.5) | 19 (86.4) |
|  | Unknown/Not Reported | - | - | 1 (25) | 1 (6.3) | 2 (9.1) |

Within the integrated dataset of 151,519 cells, we identified 7 major cell populations: epithelial cells (Epi) (*N* = 131,822), T cells and natural killer cells (T / NK) (*N* = 11,134), mononuclear phagocytes (MNP) (*N* = 5,211), mast cells (Mast) (*N* = 1,733), B cells (B) (*N* = 116), endothelial cells (Endo) (*N* = 1,239), fibroblasts (Fib) (*N* = 244) and smooth muscle cells (SM) (*N* = 20) (**Figure 1B**). Representative marker genes used for cell type annotation included *KRT6A* and *DSG3* (Epi), *CD3D*, and *NKG7* (T / NK), *CD68*, *CD207* and *CD14* (MNP), *KIT* and *CPA3* (Mast), *CD79A* and *IGHA1* (B), *VWF* and *CDH5* (Endo), *DCN*, *COL1A1* and *MYL9* (Fib), and *MYL9*, *MYH11*, and *CNN1* (SM) (**Figure 1C**). We obtained 85,745 cells were obtained from HC and 65,774 from EoE (**Supplemental Figure 1B**). Most major cell populations exhibited comparable representation from EoE and HC (**Figure 1D, Supplemental Figure 1C**), and EEC were the prominent cell type (**Figure 1E**).

### Defining EoE-associated transcriptomic alterations in EEC clusters

The prominent representation of EEC in our dataset (86.83%), a central contributor to EoE pathogenesis (11, 12), enabled high-resolution characterization of their transcriptional changes in EoE. EEC were re-integrated using anchors identified from healthy samples to ensure UMAP embeddings were assigned based on epithelial subtypes at homeostatic conditions, allowing the representation of EoE and HC EEC across each cluster (**Supplemental Figure 2A**). Ten epithelial clusters were identified via unsupervised graph-based clustering (**Figure 2A, Supplemental Figure 2B)**.

**Figure 2.**
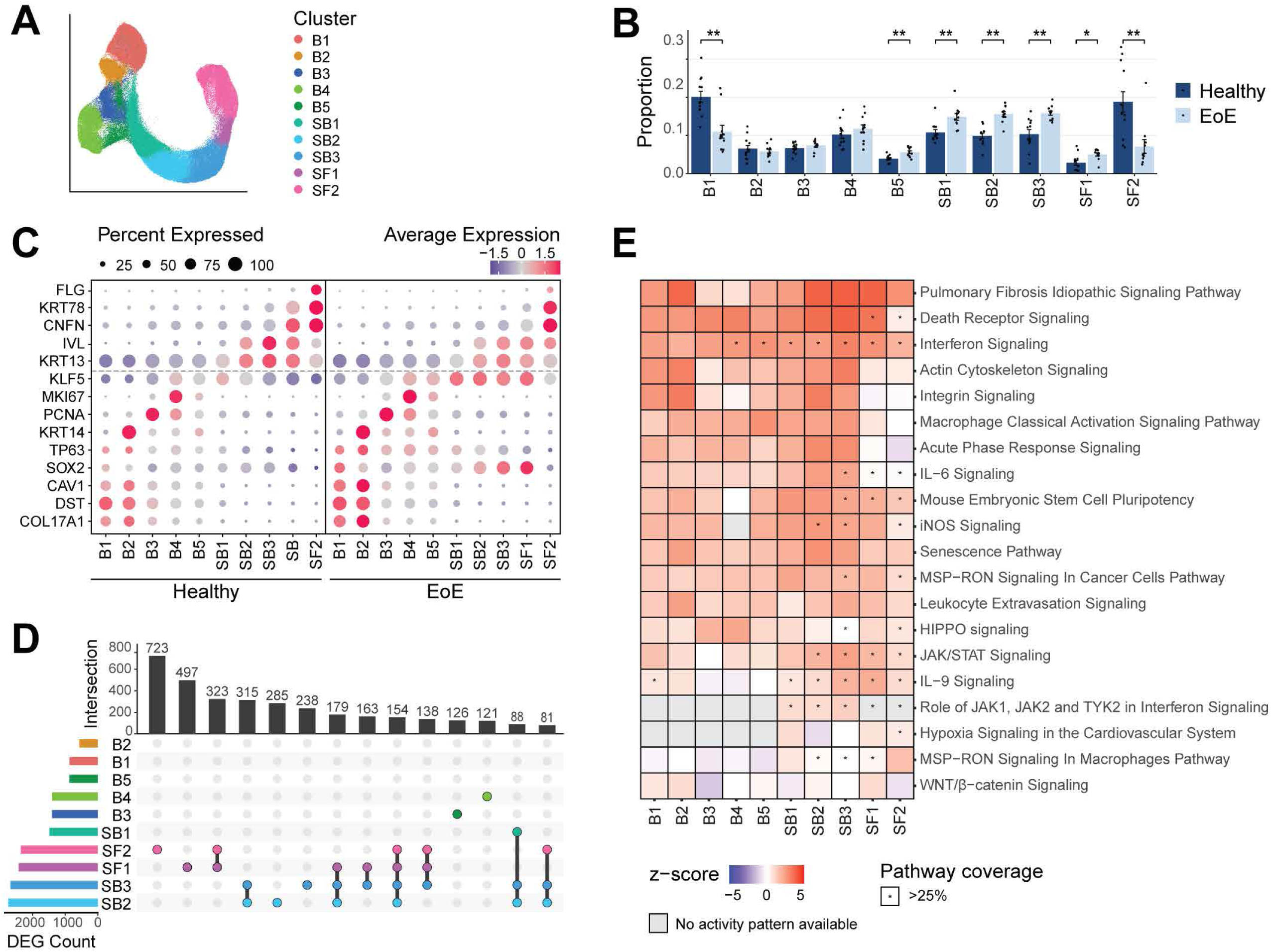
Identification and characterization of human EEC populations and their transcriptomic alterations in EoE. **(A)** UMAP of the sub-clustering of epithelial populations, colored by cluster. **(B)** Proportion of each epithelial cluster as a fraction of all epithelial cells for each disease condition. Data are expressed as mean +/- SEM and P values were determined using the Wilcoxon signed-ranked test with Benjamini & Hochberg adjustment for multiple comparisons. *P ≤ 0.05, **P ≤ 0.01. **(C)** Cluster average expression z-scores of basal and differentiated EEC markers in EoE and HC. Color gradient indicates average per-cluster gene expression, dot size indicates percent of cells per cluster that shows gene expression. **(D)** UpSet plot displaying the number of DEGs per EEC cluster, calculated between EoE and HC. **(E)** Ingenuity pathway analysis identified canonical pathways significantly altered in EEC populations in EoE compared to HC. Heatmap color intensity indicates activa­ tion z-score of each term; red denotes activation, blue denotes inhibition. Star indicates a pathway coverage of more than 25%. Basal (B), Suprabasal (SB), Superficial (SF).

The expression of established marker genes across the HC dataset was used to annotate the 10 epithelial clusters and the esophageal epithelial compartments (basal (B), suprabasal (SB), or superficial (SF)) (**Supplemental Figure 3A-B**). B1 and B2 exhibited high expression of quiescence markers *KRT15* and *DST* (7); B3 showed high expression of the S-phase marker *PCNA* (13, 14); B4 demonstrated predominant expression of the G2/mitosis marker *MKI67* (15); and B5 showed low *MKI67* expression, indicating recent cell cycle exit (**Supplemental Figure 3A-B**). Consistent with the literature, basal cells also expressed the transcription factors *SOX2* (16) and *TP63* (17–19) (**Supplemental Figure 3A-B**). Suprabasal clusters were defined based on the expression of *KRT13* (KRT13^high^) (20) *IVL* (21), and *SERPINB3* (7) (**Supplemental Figure 3A-B**).The early superficial marker *CNFN* (7, 22) and the late superficial markers *FLG* (23), *KRT78* (24), and *SPRR2D* (25) were used to identify superficial cell clusters (**Supplemental Figure 3A-B**). Cluster annotation was confirmed using transcriptional profiles of each HC cell cluster (**Supplemental Figure 4A**).

### Increased basal identity marker expression is observed in the differentiated epithelial compartment in EoE

We next investigated alterations in the relative representation of EEC clusters in EoE compared to HC. Broadly, we observed a decrease in quiescent cells (B1), an increased representation of suprabasal cells (SB1-SB3), and a substantial decrease in terminally differentiated EEC (SF2) (**Figure 2B**). We next examined changes in transcriptional profiles in epithelial clusters in EoE, focusing first on differentiation markers associated with basal, suprabasal, and superficial cell identity since EEC differentiation is dramatically affected in EoE (7, 11). We observed considerable downregulation of *FLG* in the terminally differentiated cluster SF2 and a decreased *CNFN* and *KRT78* expression in the early terminal differentiation cluster SF1 in EoE compared to HC (**Figure 2C**). However, the increased expression of *KRT13* and *IVL* occurring upon suprabasal commitment was still detected in the suprabasal clusters in EoE as compared to the basal compartment, but to slightly lower levels in EoE compared to HC (**Figure 2C**). Lastly, in addition to their expected expression in basal clusters, basal-associated genes including *SOX2*, *KLF5* and *TP63*, were also expressed throughout SB1-SF1 in EoE (**Figure 2C**). Thus, our analysis of differentiation across EEC in EoE revealed a more complex picture of the disruption of differentiation occurring in EoE than what has been previously reported; we now show a loss of terminal differentiation with a concurrent gain of basal-associated genes in differentiated cells.

We next performed differential expression testing between HC and EoE for each EEC cluster. SB2-SB3 clusters demonstrated the highest count of DEGs, followed by SF1-SF2 (**Figure 2D**). DEGs in SB2-SF2 also exhibited greater log2 fold-changes (logFC) compared to other clusters (**Supplemental Figure 4B**). Interestingly, the highest degree of distinct DEG overlaps was in suprabasal and superficial clusters (**Figure 2D**), indicating their critical role in EoE pathogenesis. Ingenuity Pathway Analysis (IPA) on DEGs for each cluster predicted the activation of various pathways, including some known to be involved in EoE, such as interferon signaling (26), senescence (27), integrin signaling (28), IL-6/IL-9 signaling (29) (**Figure 2E**). Supporting our findings that differentiated EEC express genes associated with stem cell identity in EoE, the Mouse Embryonic Stem Cell Pluripotency pathway, typically associated with epithelial progenitors in the basal compartment, was predicted to be activated across all epithelial clusters in EoE, with higher activation scores and pathway coverage in suprabasal and superficial clusters (**Figure 2E**).

### Evaluation of hyperproliferation in different epithelial compartments in EoE

We confirmed the presence of BCH in adjacent esophageal mucosal biopsies from scRNA-seq patients using EoEHSS criteria, which defines BCH as an increase in the percent of the epithelium occupied by the basal zone (level at which basal epithelial cell nuclei were separated at a distance ≥ the diameter of a basal cell nucleus) (30) (**Figure 3A-B, Supplemental Figure 5A-C)**. A median grade of 3 for BCH was observed in EoE, which replicates previous studies (**Supplemental Figure 5A**). To elucidate the impact of BCH on compartment-level dynamics in EoE, we combined the 10 epithelial clusters (**Figure 2A**) into compartments (Basal: B1-B5; Suprabasal: SB1-SB3; Superficial: SF1-SF2) using the expression of established markers (**Figure 3C, Supplemental Figure 3B, Supplemental Figure 5D**). Instead of observing basal compartment expansion in EoE, we detected an increase in differentiated suprabasal cells and a reduction of superficial cells in EoE (**Figure 3D**). This indicates that cells designated as basal in the histological evaluation of BCH may actually be differentiated suprabasal cells with atypical morphology.

**Figure 3.**
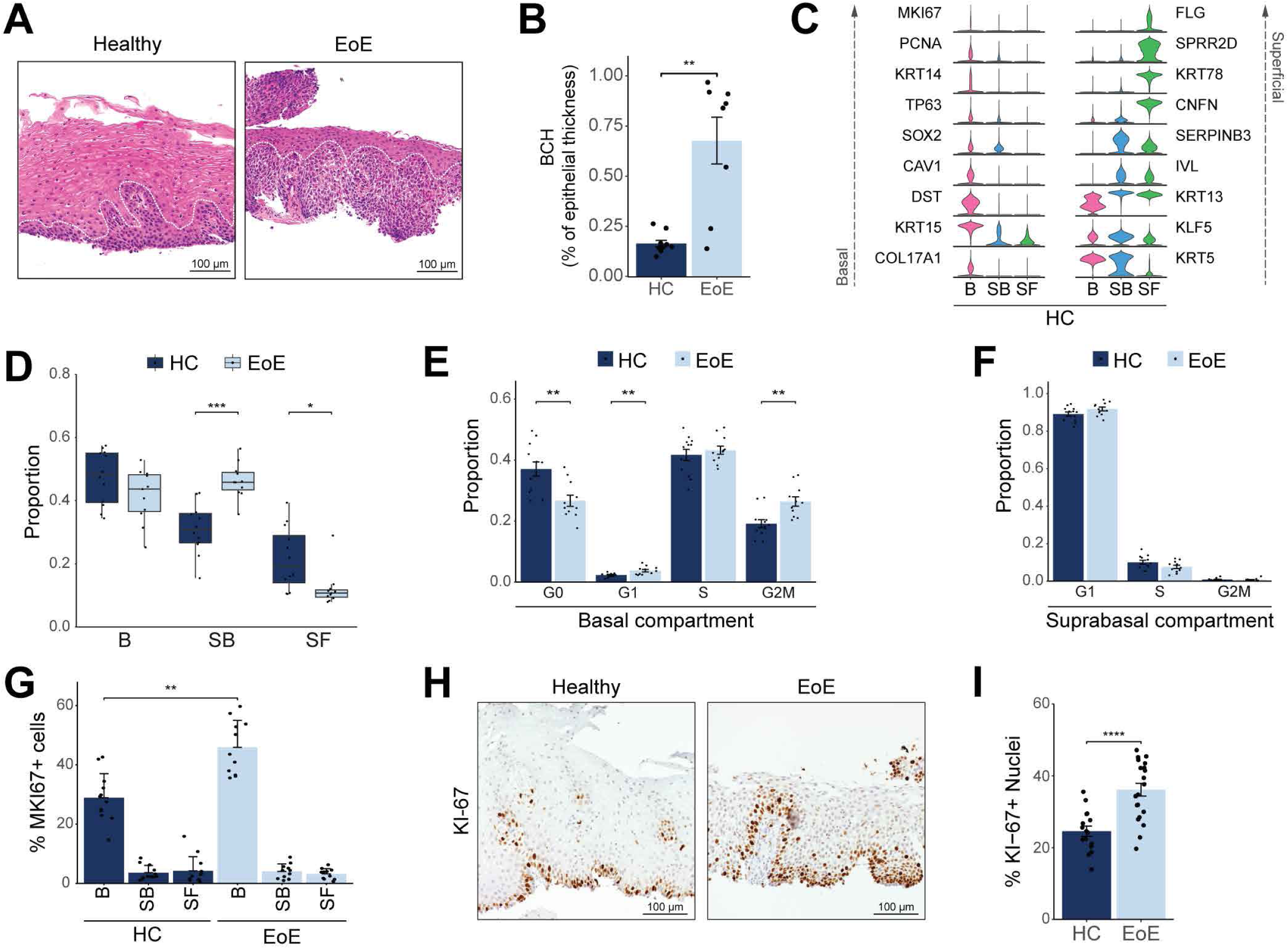
Determination of proliferation levels in different esophageal epithelial compartments in EoE. **(A)** H&E staining of HC (n = 10) and EoE (n = 8) esophageal mucosal sections from patients in the scRNA-seq cohort. The basal compartment in HC or BCH as defined by pathologists in EoE is outlined by the white dashed line. Scale bar, 100 µm. **(B)** Box plot showing the height of the epithelium occupied by the basal zone as a function of total thickness in H&E-stained sections from **(A). (C)** Violin plots displaying the expression of established marker genes for each EEC compartment in HC. **(D)** Box plot showing the proportion of each EEC compartment as a fraction of all epithelial cells for each disease condition. Boxes represent quartiles, whiskers indicate minima/maxima, and lines through each box denote median proportion. **(E-F)** Bar plots demonstrating the proportion of cells in each cell cycle phase in either the basal **(E)** or suprabasal **(F)** compartment **(G)** Bar plot showing the percent of cells expressing *MK/67* in each EEC compartment in EoE and HC. **(H)** Representative immunohistochemistry for Kl-67 in the esophageal epithelium of HC (n = 16) and EoE (n = 21). Scale bar, 100 µm. **(I)** Quantification of Kl-67+ cells from immunohistochemistry in HC and EoE. All indicated *P* values were determined using Wilcoxon signed-ranked test, with Benjamini & Hochberg adjustment for multiple comparisons specifically applied in **(D-G).** \**p ≤* 5 0.05, \*\**p ≤* 5 0.01, ***p *≤* 5 0.001, ****p *≤* 5 0.0001. Basal (B), Suprabasal (SB), Superficial (SF).

Because EoE has been associated with hyperproliferation (31, 32), we next examined proliferation rates in the basal and suprabasal compartments in EoE using published signatures (7, 33) (**Supplemental Figure 5E**). We detected a 7% increase in actively dividing G2/M cells in the basal compartment in EoE (**Figure 3E**), with no change in the suprabasal compartment (**Figure 3F**). To confirm this finding, we used a more permissive approach, quantifying the percentage of cells expressing the G2/M marker *MKI67* across each compartment, and observed a 17% increase in actively dividing basal cells and no difference in suprabasal or superficial cells in EoE compared to HC (**Figure 3G**).

The esophageal epithelial basal compartment contains distinct populations of cycling cells, including slow-cycling stem cells of the basal layer and faster-cycling epibasal cells (34) (**Supplemental Figure 3B**). To distinguish these, we subclustered quiescent and dividing clusters (B1-B5) and annotated populations based on the expression of *KRT13* (34), *DST* (35) and cell cycle markers (**Supplemental Figure 5F-G**). Population assignment was consistent with previous classification using high/low *PDPN* expression (7) (**Supplemental Figure 5H)**. Consistent with previous reports, relative proportions demonstrated an increase in dividing epibasal cells in EoE, as opposed to slow-cycling basal layer cells (7) (**Supplemental Figure 5I)**. Ki-67 immunostaining yielded similar findings, showing a 14.4% cell proliferation increase in EoE (**Figure 3H-I**). Taken together, our findings show that the expanded early differentiated EEC (SB1-SF1) retaining a basal-like identity in EoE are non-proliferative.

### Differential gene expression analysis reveals a dysfunctional differentiation process in EoE

To further characterize the expansion of early differentiated EEC expressing basal cell regulators in EoE, we investigated transcriptional changes in the different epithelial compartments. We conducted differential expression on all EEC in EoE compared to HC and performed pathway enrichment analysis on the hierarchical clustering of the resulting DEGs (**Figure 4A**). Decreased expression of cluster 4 genes associated with keratinocyte differentiation was observed in the superficial compartment in EoE (**Figure 4A**). Pathway enrichment analysis predicted terms associated with interferon signaling, regulation of cell activation, and cell-cell adhesion for cluster 2 genes increased in the basal and suprabasal compartments in EoE (**Figure 4A**). Increased expression of cluster 1 genes in the differentiated compartments in EoE were associated NABA matrisome associated, regulation of epithelial proliferation, and response to wounding (**Figure 4A**).

**Figure 4.**
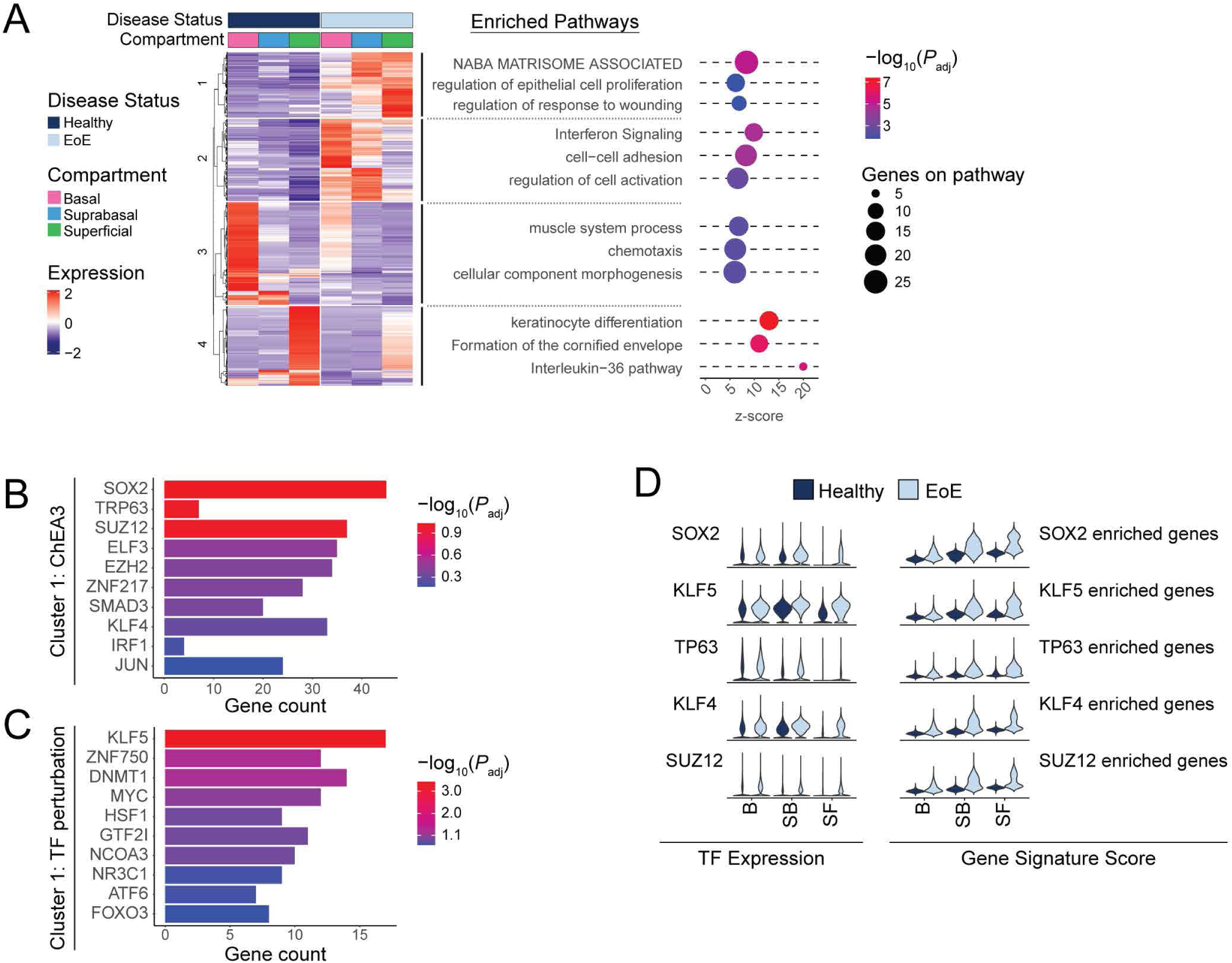
Transcriptional changes in the differentiated EEC compartment in EoE are driven by progenitor-associated transcription factors (TFs). (A) Heatmap of normalized log_2_ z-score expression of DEGs calculated between EoE and HC across all EEC (logFC >1 and FDR-adjusted *P* value < 0.05). Top hierarchical clusters are displayed with associated enriched pathways. Dot color intensity indicates –log_10_ of adjusted *P* value (P_adj_) for each term, x-axis indicates z-score for each term, dot size indicates number of DEG gene hits along each pathway. (B-C) TF analysis for genes upregulated in hierarchical cluster 1 is shown using the ChEA3 2022 database (B) or the TF perturbations followed by expression database (C). Color intensity indicates -log_10_ of *P*_adj_, x-axis indicates total DEG gene hits per term. (D) Violin plots displaying the expression of TFs identified in (B-C) and the gene signature scores of the DEGs associated with each TF in EEC compartments between HC and EoE. Basal (B), Suprabasal (SB), Superficial (SF).

Since our findings show that BCH in EoE consists primarily of differentiated cells with elevated expression of basal-associated transcription factors, we hypothesized that upregulated cluster 1 genes in the EoE differentiated compartment are key drivers of BCH. To identify upstream regulators of the DEGs in cluster 1, DEGs were used as input to Enrichr (36), which compiles updated databases of ChIP-seq experiments and the Gene Expression Omnibus (GEO) signature of DEGs following transcription factor perturbations. Enrichment analysis against these databases identified *SOX2*, *TP63*, and *KLF5*, three regulators of embryonic stem cells self-renewal (17, 18, 37), as top predicted transcription factors regulating cluster 1 DEGs (**Figure 4B-C**). Increased expression of *SOX2*, *TP63* and *KLF5* along with their downstream targets was confirmed in the differentiated compartments in EoE (**Figure 4D**).

To further explore this finding of differentiated cells in EoE that maintain active gene programs potentially conferring a stem cell identity, we developed two signatures representing genes that demonstrate preferential expression in either quiescent cells or superficial cells in HC to define a quiescent-basal-differentiation axis in human esophageal epithelium (**Supplemental Tables 1-2**). Violin plots and contour plots mapping the quiescent signature score (y-axis) and superficial signature score (x-axis) by disease condition showed a clear separation of the superficial compartment of HC from the basal and suprabasal compartments, defined by a high superficial score and minimal quiescent score (**Figure 5, Supplemental Figure 6**). However, in EoE, we observed a shift toward decreased superficial score and increased quiescent score in the superficial compartment, causing overlap with the suprabasal compartment (**Figure 5, Supplemental Figure 6**). Upon separation of the differentiated compartments into epithelial clusters, we observed increased quiescent identity in each differentiated cluster in EoE beginning at SB2, with the most dramatic shift in SF1 (**Figure 5, Supplemental Figure 6)**. This shift in the quiescent-basal-differentiation axis was consistent across all EoE patients, with few cells oriented in the terminally differentiated position in the lower right quadrant (**Figure 5**).

**Figure 5.**
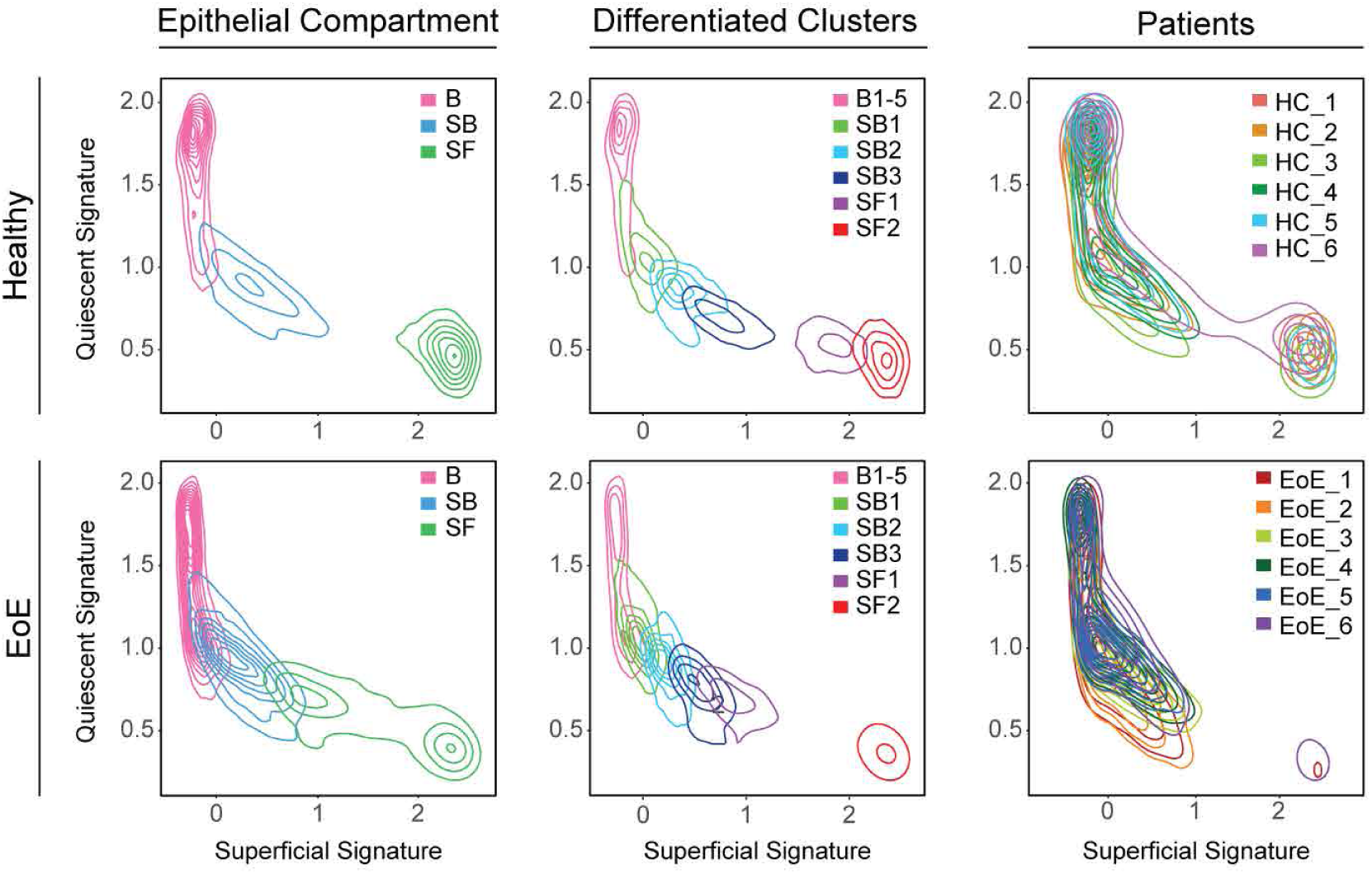
The quiescent-basal-differentiation transition is altered in the esophageal epithelium in EoE. Contour plots showing EEC from HC or EoE plotted along the quiescent gene signature (y-axis) and the superficial gene signature (x-axis). Line color indicates cell grouping by EEC compartment, with or without the differentiated clusters labeled, or patient origin. Basal (B), Suprabasal (SB), Superficial (SF).

To validate changes in the quiescent-basal-differentiation axis, we performed multispectral fluorescent staining of esophageal mucosal sections from HC and EoE using established markers of basal (KRT14, p63), suprabasal (IVL), and superficial (CNFN) cell identity (**Figure 6A-B**). For comparison, gene expression of each marker is shown across clusters in HC or EoE (**Figure 6C**). We confirmed the suprabasal identity marker IVL was appropriately expressed after exit from the basal compartment in EoE (**Figure 6B**). Next, EEC in EoE expressed p63 in the differentiated compartment (**Figure 6B**), compared to primarily basal-restricted expression in HC (**Figure 6A**). Finally, CNFN was restricted to the very top 1-2 layers of EEC in EoE compared to HC, confirming a delay in terminal differentiation (**Figure 6A-B**). Thus, our findings show that despite maintaining the correct spatial organization of suprabasal lineage commitment, most differentiated EEC in EoE retain a basal identity.

**Figure 6.**
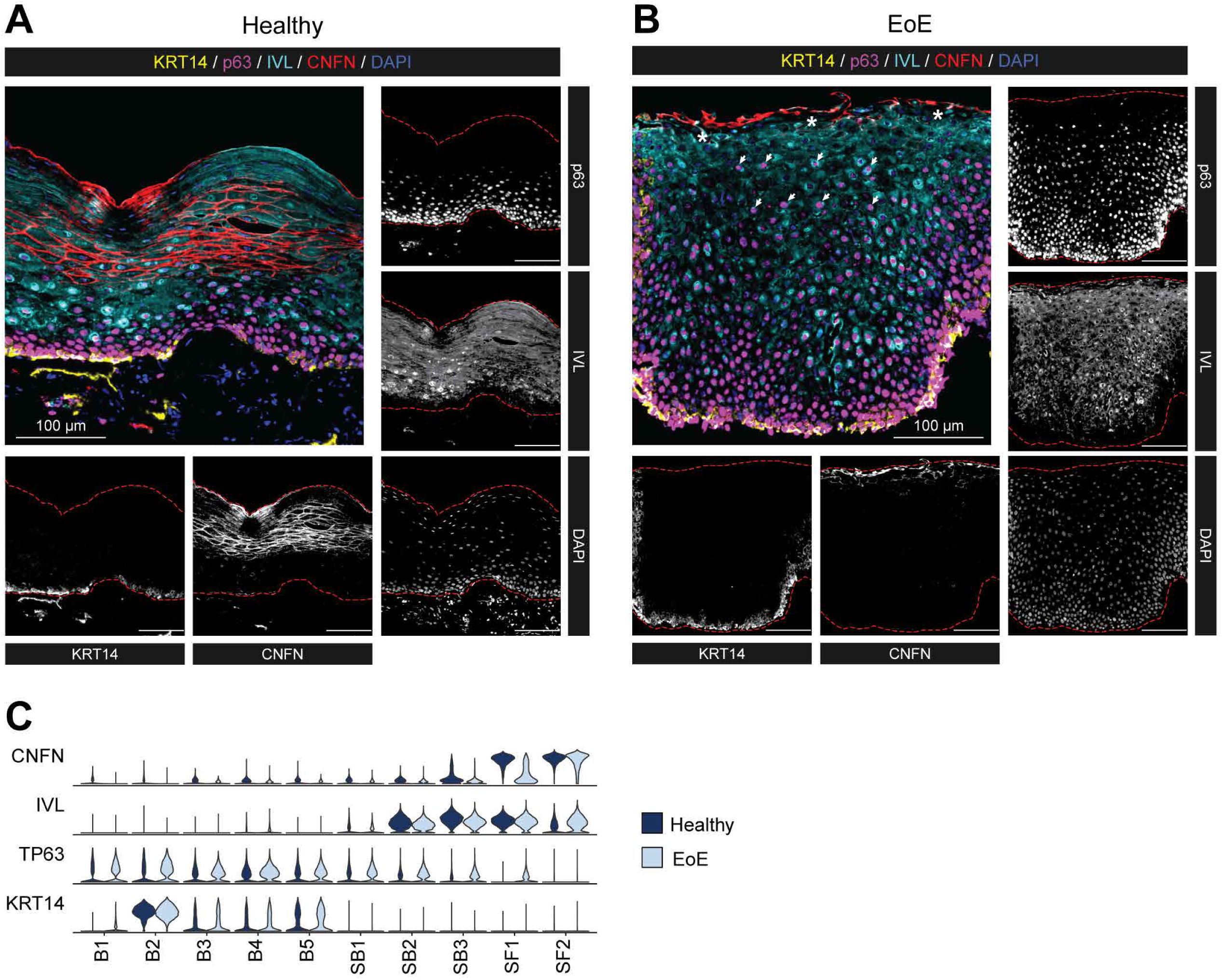
Validation of EoE-associated differentiation changes identified by scRNA-seq in esophageal tissue from HC and EoE. **(A-B)** Representative multispectral fluorescent tissue staining for markers of the epithelial basal (KRT14: yellow, p63: magenta), suprabasal (IVL: cyan), or superficial compartment (CNFN: red) in esophageal mucosaI biopsies of HC (n = 8) **(A)** or EoE (n = 14) (B). Scale bar, 100 µm. Arrows indicate p63+ IVL+ nuclei in the BCH expanded area, stars denote regions of CNFN loss, red dashed lines outline the epithelial area. (C) Violin plots displaying the expression of indicated markers of EEC compartments across epithelial clusters between EoE and HC. Basal (B), Suprabasal (SB), Superficial (SF).

### Pseudotemporal analysis confirms loss of terminal differentiation and a global differentiation shift toward basal identity in EEC in EoE

To examine differences in cell fate trajectories along the course of differentiation, we merged epithelial samples from HC and EoE patients for pseudotemporal analysis with Monocle3 (**Figure 7A, Supplemental Figure 7A**). We followed the previously established model for EEC ordering wherein S-phase cells (B3) are treated as root cells (22) (**Figure 7B, Supplemental Figure 7B**). In this model, EEC in S-phase of the cell cycle progress through G2/M phases (B4), and then are faced with the commitment decision (B5) to either return to the G0 quiescent reserve (B1-B2) or to commit to cellular differentiation (SB1-3, SF1-2) (**Figure 7C**). Trajectory analysis was performed on healthy and EoE conditions combined (**Figure 7B**) to allow direct comparison of pseudotime values between conditions. A severe decrease in late pseudotime peaks is observed in EEC in EoE, with a concentration of cells in an intermediate range of pseudotime values instead (**Figure 7D**). This shift in pseudotime value distribution was consistent across all EoE patients (**Supplemental Figure 7C**) and was not due to a decreased frequency of superficial cells (**Figure 7E**). The comparison of pseudotime densities between epithelial compartments revealed a marked shift in pseudotime density profiles in the suprabasal and superficial compartments in EoE, compared to HC (**Figure 7E**). Breakdown of the differentiated compartments into component clusters revealed a significant decrease in pseudotime values starting in SB3 in EoE compared to HC (**Figure 7F**). Hierarchical clustering of mean pseudotime values of each differentiated cluster demonstrated an 87% accuracy in distinguishing EoE from HC using the first two dendrogram nodes (**Supplemental Figure 7D**). This further demonstrates a global shift toward basal identity in the esophageal epithelial differentiated compartments in EoE.

**Figure 7.**
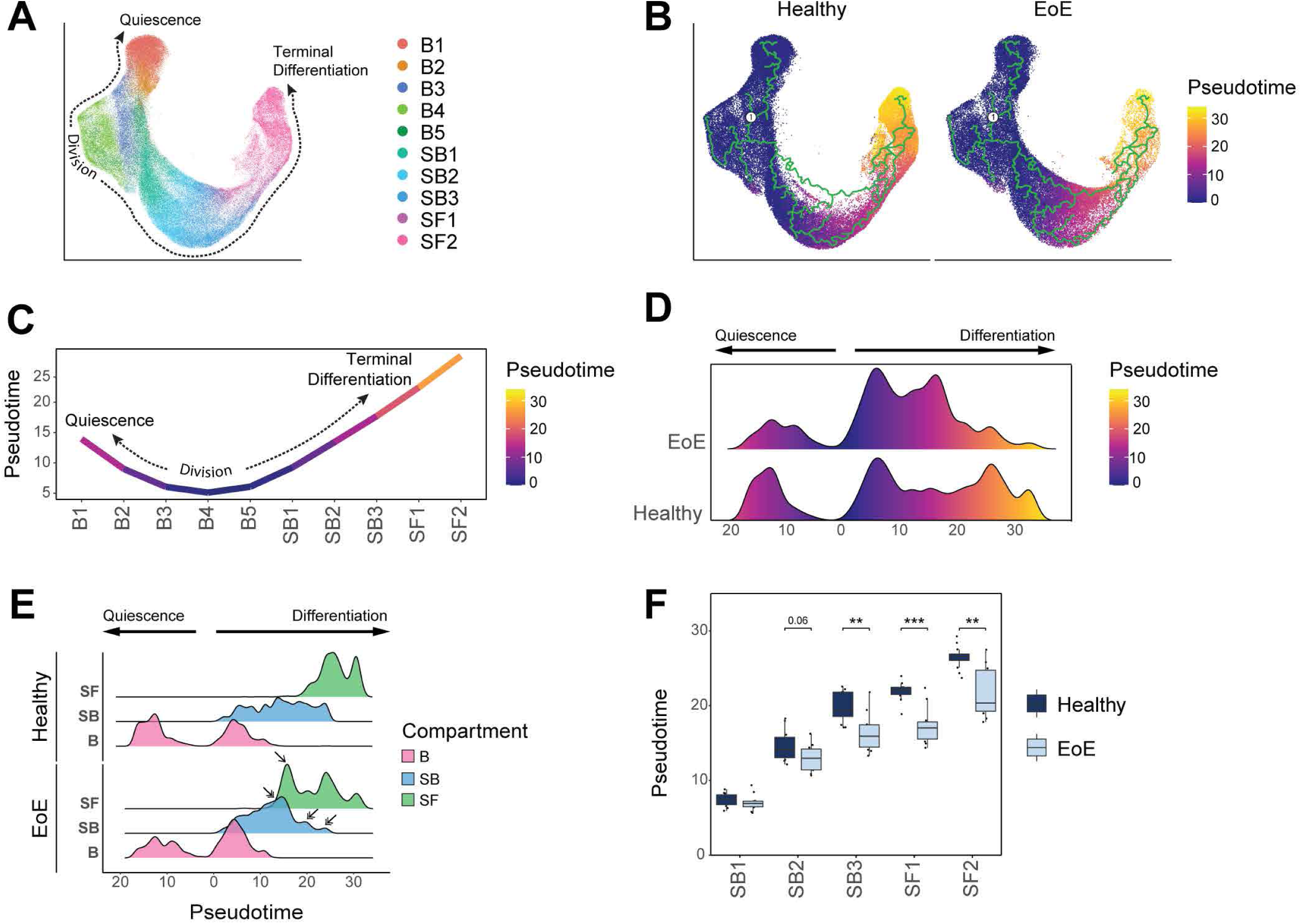
Pseudotemporal trajectory analysis of EEC demonstrates loss of terminal differentiation and global differentiation shift towards basal identity. (**A**) UMAP showing merged scRNA-seq datasets of EEC from EoE and HC, colored by integrated clusters. (**B**) Pseudotime trajectory analysis calculated using EoE and HC samples plotted on UMAP from (**A**), with S-phase cluster B3 as pseudotime origin. (**C**) Average pseudotime values across epithelial clusters. (**D**) Ridgeline plot illustrating the pseudotime value distribution between EEC in EoE and HC. Pseudotime begins at cells in S-phase and increases moving toward either quiescence or terminal differentiation. (**E**) Ridgeline plots depicting pseudotime value distribution between epithelial compartments in HC and EoE. Arrows indicate peaks with differential density between EoE and HC. (**F**) Box plot showing the distribution of pseudotime values per differentiated epithelial cluster in EoE or HC. Boxes indicate quartiles, whiskers indicate minima/maxima separated by 1.5 times the interquartile range, lines through each box indicate median pseudotime value. Indicated P values were determined using Wilcoxon signed-ranked test with Benjamini & Hochberg adjustment for multiple comparisons. \*\**P* ≤ 0.01, ***P ≤ 0.001. Basal (B), Suprabasal (SB), Superficial (SF).

Next, we identified gene modules that displayed EoE trajectory-dependent gene expression patterns (**Figure 8A-B, Supplemental Table 3**). Top terms from pathway enrichment analysis are shown for each in **Supplemental Figure 8A**. Modules 4, 5, 6 and 7 showed different expression patterns between EoE and HC (**Supplemental Figure 8B**). Modules 4, 5 and 6 were linked to EEC differentiation (**Supplemental Figure 8A**). Module 7 was particularly interesting as it contained genes with increased expression in all epithelial clusters in EoE (**Figure 8A, Supplemental Figure 8B-C, Supplemental Table 4**), with peak increase in SB2 (**Supplemental Figure 8D**). Pathway enrichment analysis of module 7 genes identified enriched terms associated with response to wounding, regulation of actin filament-based process, regulation of keratinocyte proliferation, and positive regulation of cell motility (**Figure 8C**). Mean module 7 gene signature scoring showed elevated expression along the differentiation trajectory in EoE as compared to HC, peaking at pseudotemporal values representing the differentiated clusters that showed earlier pseudotemporal identity in EoE (**Figure 8D**). Interestingly, *SOX2* and *KLF5* expression also increased across a similar pseudotemporal range in EoE compared to HC (**Figure 8D**). A higher percentage of EECs expressed overlapping *SOX2*, *KLF5*, and Module 7 signature scoring in EoE, with the highest level of coexpression within the same range of pseudotemporal values (**Figure 8D, Supplemental Figure 8E**), suggesting the regulation of module 7 genes by *SOX2* and *KLF5*. Unsupervised hierarchal clustering analysis showed two different expression patterns for module 7 genes in EoE: consistent increased expression throughout EEC (cluster 2) or increased expression in EEC peaking in clusters SB2-SF1 (cluster 1) (**Supplemental Figure 8F**). Notably, expression of module 7 genes in cluster 1 in EoE peaked in differentiated epithelial clusters showing aberrant *SOX2* and *KLF5* expression (**Supplemental Figure 8F**). Furthermore, over 47% of module 7 genes were known epithelial targets of either SOX2, KLF5, or the SOX2-KLF5 interaction (38–40) (**Figure 8E, Supplemental Table 5-6**) with known protein-protein interactions (**Supplemental Figure 9**). This supports our findings that *SOX2*, *KLF5* or their interaction play a prevalent role in controlling the upregulated gene programs observed in the differentiated compartments in EoE and in orchestrating disease-associated tissue remodeling.

**Figure 8.**
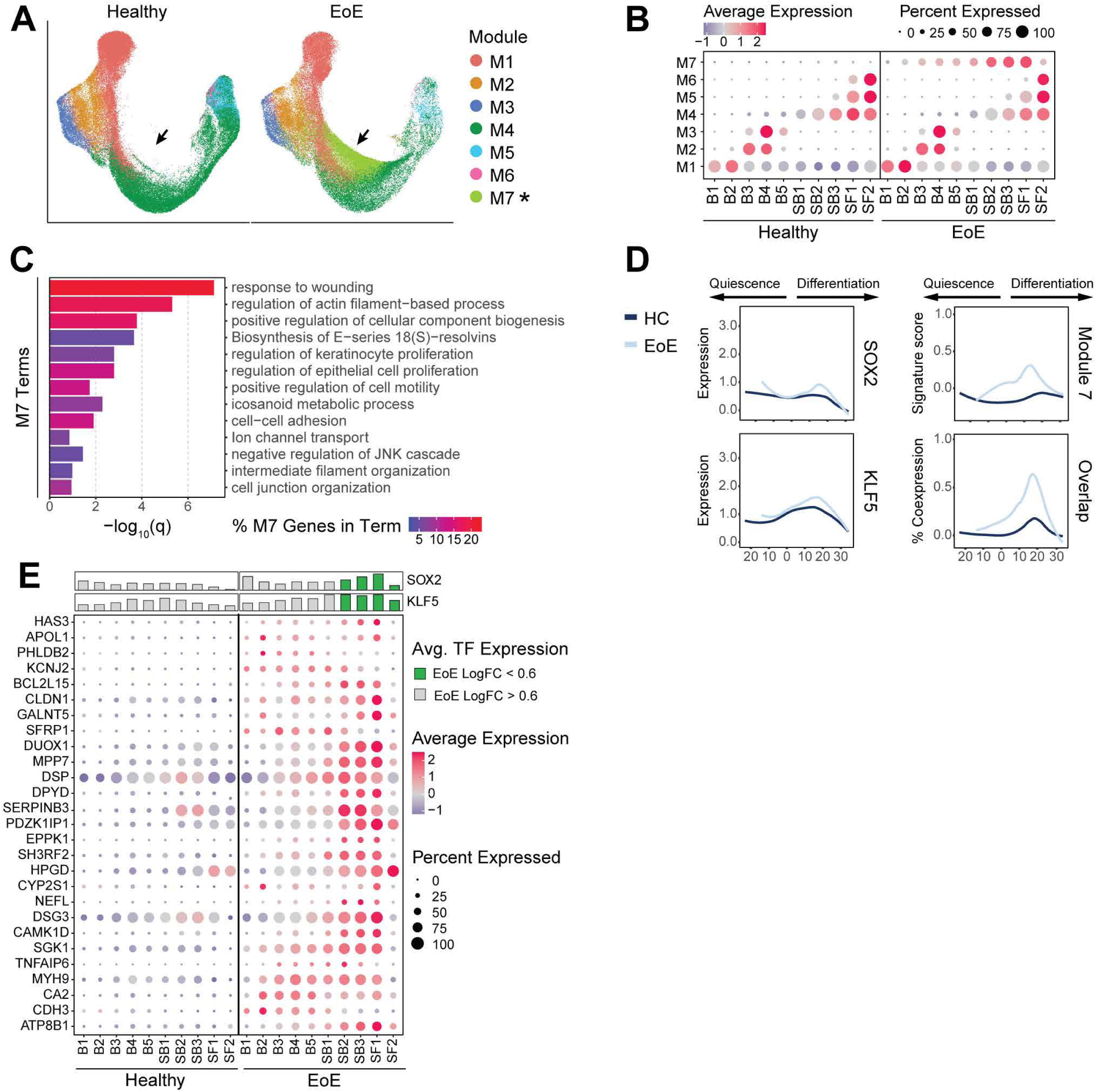
Pseudotemporal trajectory analysis reveals altered trajectory-dependent gene programs in EoE. **(A)** UMAP split by disease state, with cells colored by highest-scoring pseudotime-dependent module. Arrows indicate module 7 position. **(B)** Average signature score z-scores per module for epithelial clusters in HC or EoE. Color intensity indicates relative scoring. **(C)** Enriched pathways for module 7 genes ranked by *P* value. Color intensity indicates percentage of module 7 genes along each pathway. **(D)** The expression of SOX2, *KLF5* or module 7 genes across cells ordered by pseudotime in EoE and HC, as well as the percentage of cells expressing meaningful levels of all three conditions. Lines represent moving averages calculated by loess regression. **(E)** Dot plot displaying the average-expression z-scores of genes in module 7 regulated by SOX2 and/or KLF5 for each epithelial cluster in HC or EoE. For each cluster, mean log_2_ gene expression for SOX2 and *KLF5* is shown as a bar plot. Dot color gradient indicates average expression, bar color indicates whether logFC between EoE and HC clusters is above (green) or below (grey) 0.6-fold. Basal (B), Suprabasal (SB), Superficial (SF).

### SOX2 and KLF5 gene programs are altered in the esophageal epithelial differentiated compartments in EoE

In addition to an the observed coexpression of *SOX2* and *KLF5* in EoE, we found their expression to be elevated in a higher proportion of cells in the differentiated EEC clusters in EoE, compared to HC (**Figure 9A**, **Supplemental Figure 10A-B**). Immunohistochemistry confirmed increased nuclear expression of SOX2 and KLF5 in differentiated EEC in EoE (**Figure 9B-F**). Interestingly, KLF5 was recently identified as a SOX2 binding partner, and their interaction led to the acquisition of chromatin binding sites not observed with SOX2 or KLF5 alone (40). Published SOX2-, KLF5-, or SOX2/KLF5-regulated gene programs (38–40) were enriched across EEC clusters in EoE, particularly in differentiated clusters (**Figure 9G, Supplemental Figure 10C, Supplemental Table 5, 7**). 1,611 genes known to be regulated by SOX2 and/or KLF5 were significantly upregulated in EoE EEC (FDR *P* value < 0.05 and logFC > 0.25), 76.4% of which demonstrated the highest upregulation in the differentiated compartments (**Supplemental Table 7**). Additionally, 224 genes known to be co-regulated by the SOX2-KLF5 interaction had significantly altered expression in EoE (FDR *P* value < 0.05 and |logFC| > 0.25), and 86.7% of these were upregulated in EoE (**Supplemental Table 8**).

**Figure 9.**
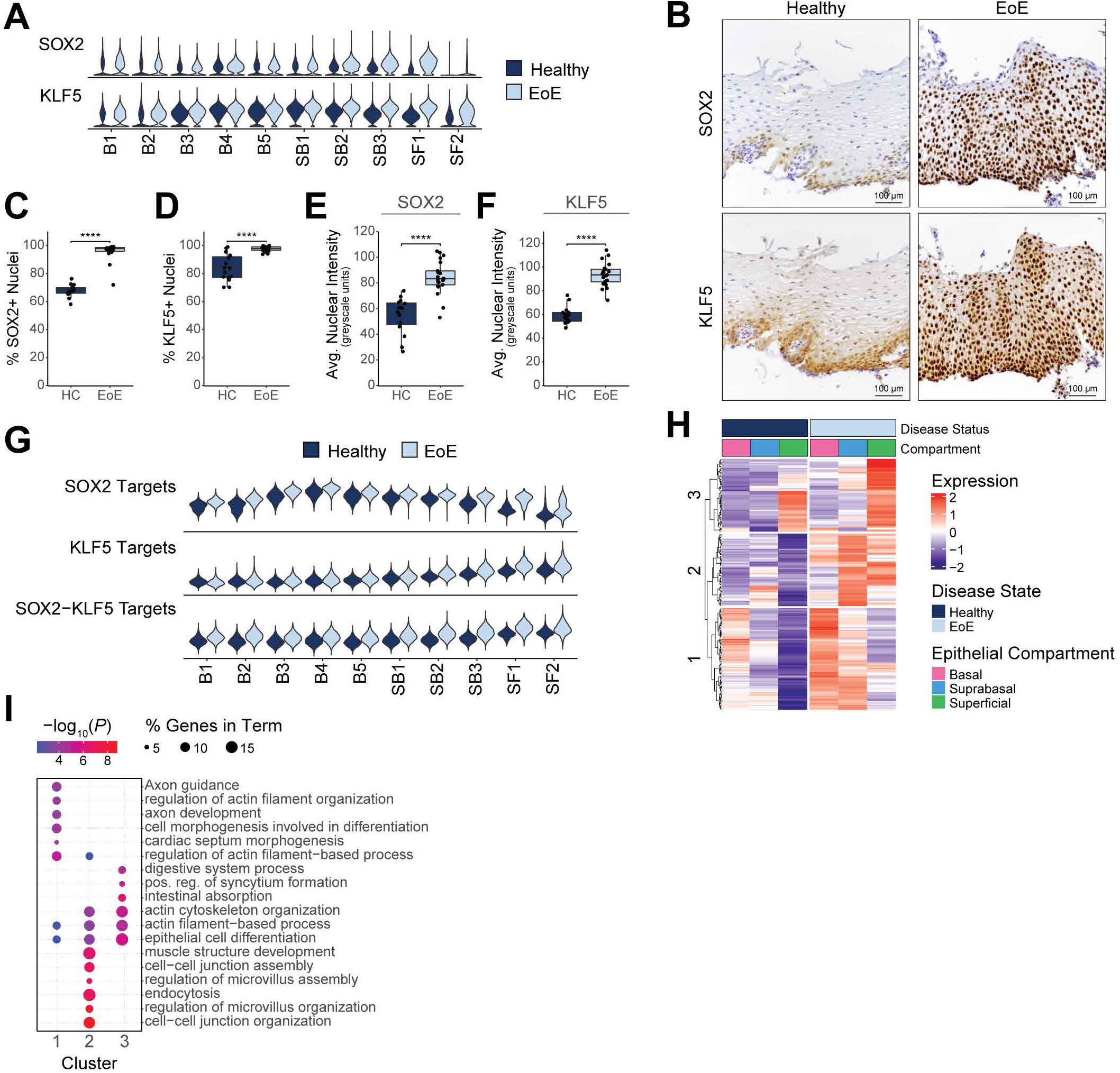
SOX2 and KLF5 gene programs are altered in the esophageal epithelial differentiated compartments in EoE. **(A)** Violin plots displaying *KLF5* and *SOX2* gene expression across epithelial clusters between EoE and HC. **(B)** lmmunohisto­ chemistry for indicated proteins in the esophageal epithelium in EoE (n = 21) or HC (n = 15). Scale bar, 100 µm. The positive staining for SOX2 **(C)** or KLF5 **(D)** out of total cells is quantified. The average nuclear intensity for SOX2 **(E)** or KLF5 **(F)** across all cells is quantified, measured in greyscale units. For **(C-F),** data are shown as mean ± SEM, *\*\*\*\*P ≤* 0.0001. **(G)** Violin plots showing gene signatures calculated from the transcriptional targets of SOX2, KLF5, or SOX2-KLF5 in epithelial clusters between EoE and HC. **(H)** Heatmap depicting the expression of SOX2-KLF5 coregulated genes that are significantly upregulated in EoE. The top hierarchal clusters are identified, with enriched terms for each shown in **(I).** Color indicates -log_10_(P value), dot size indicates percentage of genes along each term. Basal (B), Suprabasal (SB), Superficial (SF), positive (pos), regulation (reg).

We next performed unsupervised clustering analysis of SOX2-KLF5 co-regulated gene targets with increased expression in EoE. Cluster 1 genes displayed progressively reduced expression throughout the differentiated compartment in HC but showed increased expression in EoE (**Figure 9H**). Enrichment analysis revealed changes related to actin-filament based processes and cell morphogenesis associated with differentiation (**Figure 9I**). Increased cluster 2 genes expression was seen in the differentiated compartment in EoE, with relatively low expression in HC (**Figure 9H**). Cluster 2 genes were linked to pathways associated with cell-cell junction and actin cytoskeleton organization (**Figure 9I**). These findings demonstrate that dysregulated SOX2 and KLF5 expression and their downstream targets play a role in driving epithelial remodeling within the differentiated compartments in EoE.

Dysregulated differentiation and aberrant progenitor-regulating transcription factor signaling in EEC are unique to EoE and not observed in gastroesophageal reflux disease (GERD). EoE and GERD patients present with overlapping symptoms, such as heartburn and dysphagia (41), and undergo similar epithelial remodeling such as BCH (42). For this reason, we investigated whether transcriptomic changes observed in EoE were unique to the disease or were caused by acid reflux. We performed scRNA-seq on 4 GERD patients and imputed cell identities established in our human EEC dataset from HC and EoE onto GERD EEC. Differential expression was then calculated between EoE and HC for each epithelial compartment. LogFC from GERD compared to HC was calculated for each gene significantly changed in EoE (|logFC| > 0.5 and FDR adjusted *P* value < 0.05) for each compartment. As shown in Figure 10A, EEC from GERD and EoE only shared few genes changing in the same direction. Interestingly, most EoE DEGs in the basal and suprabasal compartments showed minimal change in GERD; whereas in the superficial compartment, 48% of DEGs displayed opposite changes in GERD (|logFC| > 0.5) (Figure 10A). Next, we compared known epithelial markers between HC, EoE and GERD. Contrasting the loss of terminal differentiation observed in EoE, GERD EEC showed the correct expression patterns of early (*KRT13*, *IVL*) and late differentiation markers (*CNFN, SPRR2D, FLG*, and *KRT78*) in the differentiated compartments (Figure 10B). To comprehensively compare the differentiated compartments in EoE and GERD, we first calculated differential expression between the differentiated compartments in EoE versus HC, and conducted hierarchical clustering of the DEGs obtained. Subsequently, we displayed DEGs as logFC between EoE and HC or logFC between GERD and HC for each epithelial compartment and performed pathway enrichment analysis on each hierarchical cluster (Figure 10C, Supplemental Figure 11A). Genes in clusters 2 and 3 were increased in the suprabasal and superficial compartments in EoE compared to HC but were not changed in GERD (Figure 10C). Enriched terms for these DEGs were associated with interferon and IL-4 signaling, histone modification, pluripotency of stem cells, cell junction organization and cytoskeleton organization (Supplemental Figure 11A). Similarly, cluster 4 genes related to keratinocyte differentiation showed decreased expression in the superficial compartment in EoE but were not decreased to the same extent in GERD (Supplemental Figure 11A, Figure 10C). We next compared changes along the quiescent-basal-differentiation axis between GERD, EoE, and HC by scoring EEC for quiescent and superficial gene signatures (Figure 10D). The superficial compartment in GERD demonstrated proper adoption of superficial cell identity and inhibition of basal cell identity, unlike in EoE (Figure 10D, Supplemental Figure 11B). Moreover, the changes observed in the quiescent and superficial cell identity in the differentiated clusters SB3-SF2 in EoE were not present in GERD (Figure 10D, Supplemental Figure 11B), which is consistent across GERD patients (Supplemental Figure 11C). Lastly, no aberrant *SOX2*, *KLF5*, *TP63* or *KLF4* expression was detected in the differentiated compartments in GERD, in contrast to EoE (Figure 10B, E; Supplemental Figure 11D). Hierarchical clustering of the main features identified in the suprabasal and superficial compartments in EoE was able to distinguish healthy and GERD from EoE patients with 93.3% accuracy at the top-level partition in the dendrogram (Supplemental Figure 12). Our findings demonstrate that loss of terminal differentiation, the shift toward basal cell identity in the differentiated compartment, and abnormal *SOX2* and/or *KLF5* expression are unique to EEC in EoE, and not a consequence of gastric reflux in these patients.

**Figure 10.**
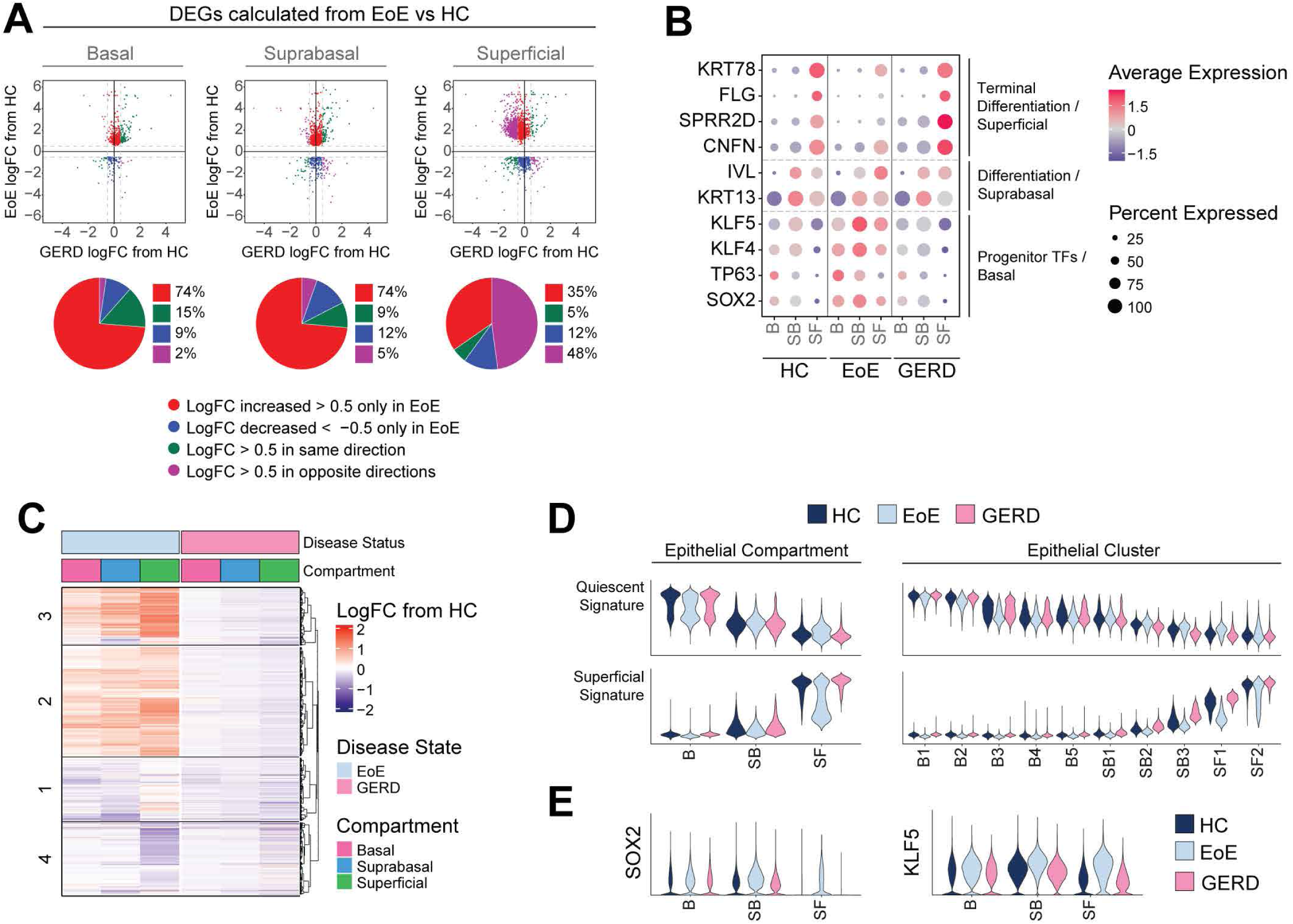
GERO does not share the esophageal epithelial transcriptional changes identified in EoE, nor does it exhibit a global shift toward basal-like identity. (**A**) Scatterplots of DEGs calculated per epithelial compartment between EoE and HC (llogFCI > 0.5 and FDR-ad­ justed P value < 0.05). Each gene is plotted as logFC in EoE (y-axis) or GERO (x-axis) from HC. Color indicates direction and magnitude of logFC in EoE and GERO. Red (logFC > 0.5 only in EoE), Blue (logFC < -0.5 only in EoE), Green (logFC > 0.5 in EoE and GERO or logFC < -0.5 in EoE and GERO), Purple (logFC > 0.5 in EoE and < -0.5 in GERO or logFC < -0.5 in EoE and > 0.5 in GERO). (**B**) EEC compartment average expression z-scores of known epithelial transcription factors and differentiation markers, compared between HC, EoE, and GERO. Color gradient indicates average per-cluster gene expression level, dot size indicates percent of cells per cluster that exhibit gene expression. (**C**) For DEGs calculated between EoE and HC across all differentiated EEC (llogFq > 0.25 and FDR-adjusted P value < 0.05), logFC from HC in EoE or GERO is visualized across EEC compartments, with top hierarchical clusters displayed. (**D**) Violin plots showing the scores for quiescent gene signature and superficial gene signature in either EEC compartments or clusters in HC, EoE, or GERO. (E) Violin plots showing the expression of *SOX2* or *KFL5* in EEC compartments in HC, EoE, or GERO. Basal (B), Suprabasal (SB), Superficial (SF).

## Discussion

Esophageal homeostasis relies on a careful balance between proliferation, differentiation, and cell death, which is critical for the maintenance of epithelial barrier function. This process is perturbed in EoE, resulting in Th2-mediated eosinophilic inflammation and epithelial remodeling including loss of differentiation and BCH (8). Clinical studies have highlighted the need to understand the role of BCH in EoE disease progression. Studies have identified a correlation between BCH and disease severity in EoE patients (9) and have shown that BCH occupies >66% of esophageal epithelial surface area in EoE (30). Even with treatment, BCH persists in approximately half of EoE patients and correlates with persistent symptoms and endoscopic findings in histologically inactive patients (9, 30). Because the molecular mechanisms driving BCH are poorly understood, we performed scRNA-seq on esophageal biopsies from adult EoE patients and healthy controls to investigate the cellular identities and transcriptional processes underlying BCH and altered epithelial differentiation in EoE.

BCH has been proposed to arise from the expansion of the basal compartment through a proliferative response (31, 43). However, despite histological confirmation of BCH across all EoE patients in the scRNA-seq dataset, we did not observe an expansion of the basal compartment by scRNA-seq. Instead, we found a decrease in quiescent basal cells, a substantial increase in suprabasal cells, and a decrease in superficial cells in EoE. Although the proportion of cells in the basal cell compartment was unchanged, we detected increased proliferation specifically driven by epibasal cells, but no increased proliferation in the basal layer or the suprabasal compartment. These findings are consistent with a recent human EoE single-cell transcriptome study that reported an increase in proliferating PDPN-negative cells in the epithelium of EoE subjects (7) as well as other scRNA-seq studies that showed a moderate increase in basal compartment proliferation, but no increase in suprabasal proliferation in human or experimental murine EoE (27, 44). However, this change in epibasal proliferation does not explain the presence of BCH in 65% of the total epithelial surface area in EoE patient biopsies from our scRNA-seq cohort. Instead, our findings show that BCH in EoE results from an expansion of non-proliferative suprabasal cells.

In addition to BCH, EoE is also characterized by a general loss of differentiation (7, 11). We found decreased expression of terminal differentiation markers in the superficial clusters in EoE, but our study further revealed a more refined understanding of the differentiation dynamics in EoE. We show that despite properly committing to early differentiation after exiting the basal compartment, differentiated EEC in EoE retain a basal-like identity. This finding is supported by earlier pseudotemporal identities and elevated expression of quiescence-associated genes in suprabasal and superficial epithelial clusters in EoE. Recently, a study investigating repair mechanisms in an intestinal injury model identified a transient cell population derived from transit amplifying cells that featured a transcriptional profile resembling regenerative stem cells, but lacked stem cell capacity (45). Although these cells harbored a stem-like transcriptional profile, they were differentiated and expressed lineage identity markers (45). The authors coined the term adaptive differentiation to describe this atypical differentiation process that occurred in response to tissue damage. We propose that the increased stem-like identity observed in the differentiated compartment of the esophageal epithelium in EoE reflects an adaptive differentiation process, triggered by a tissue-wide wound healing response to chronic inflammation in the EoE microenvironment. While adaptive differentiation may aid tissue repair in the intestine, it is critical to further investigate its role in EoE to determine whether it contributes to pathology in the context of chronic inflammation.

Enrichment analysis of genes upregulated in the differentiated compartment in EoE identified the transcription factors SOX2 and KLF5 as potential regulators of BCH, EoE disease progression, and the maintenance of stem-cell identity in differentiated EEC. SOX2 is involved in stem cell maintenance by suppressing differentiation gene programs and promoting self-renewal (17, 46, 47). Similarly, KLF5 regulates cell proliferation, migration, differentiation, and stemness (48–50). We confirmed increased, overlapping SOX2 and KLF5 expression in differentiated EEC in EoE and observed upregulation of many of their known target genes. A recent study found that SOX2 overexpression promoted SOX2-KLF5 binding, permitting the regulation of a distinct set of chromatin binding sites in esophageal squamous cell cancer (40). While this interaction was investigated in the context of cancer progression, it was also observed during the progression from normal to cancer, suggesting a potential physiological role in response to tissue injury (40, 51). Supporting this concept, pathway analysis of the SOX2/KLF5 interaction targets that showed increased expression in differentiated EEC in EoE identified terms associated with epithelial remodeling. Further research is necessary to confirm the role of the SOX2/KLF5 interaction in the injury response and the development of BCH/adaptive differentiation in EoE. Additionally, more studies are needed to clarify the upstream factors contributing to elevated SOX2 and KLF5 expression in differentiated EEC in EoE.

While our study focused on the interaction between SOX2 and KLF5, we cannot exclude the possibility that SOX2 also interacts with other factors in EoE. For example, in esophageal and lung squamous cell cancer cell lines, SOX2 and p63, another transcription factor upregulated in differentiated EEC in EoE, were shown to jointly occupy multiple genomic loci (52). Furthermore, the joint binding of p63, SOX2 and KLF5 was demonstrated to regulate chromatin accessibility, epigenetic modifications, and gene expression in ESCC (43). Further, SOX2 and KLF4 operate as a functional core in pluripotency induction across cells of different origins (53). Thus, additional investigations are needed to explore the interaction of SOX2 with other transcription factors predicted by our computational analyses in EoE.

Finally, considering the overlap in symptoms and histological presentation between EoE and GERD, such as BCH (41), it was crucial to determine whether our findings were exclusive to EoE or also applicable to GERD. Our results showed that the increased basal identity, the aberrant increase in SOX2 and KLF5 expression and the abnormal expression of other progenitor-regulating transcription factors observed in the differentiated compartment of EoE were not present in GERD patients. Consequently, these changes observed in EoE are not merely a response to gastric reflux. Interestingly, while GERD is a main risk factor for the development of esophageal cancer (54), epidemiological studies have found no association between EoE and esophageal cancer development, despite the presence of chronic inflammation (55). Further investigation into the changes in the cellular landscape of EEC in GERD may offer a better understanding of the differences between GERD and EoE and their distinct susceptibility to esophageal cancer progression.

In conclusion, our study revealed that BCH in EoE is an expansion of non-proliferative cells that commit to differentiation but retain a stem-like transcriptional program. The identification of SOX2 and KLF5 as potential master transcriptional regulators of this process offers valuable insight into the molecular mechanisms driving BCH and adaptive differentiation in EoE. Further investigation into mechanisms of epithelial remodeling and adaptive differentiation may not only advance our understanding of disease progression, but also open new avenues for therapeutic interventions, potentially improving patient outcomes in cases refractory to anti-inflammatory treatments.

## Methods

### Human specimen collection

Healthy controls met asymptomatic criteria including the lack of esophageal symptoms (heartburn, dysphagia, chest pain), history of tobacco use or alcohol dependency, body mass index greater than 30 kg/m^2^, or previous treatment with antacids or proton pump inhibitors. EoE patients were recruited at the primary visit contingent upon confirmed diagnosis and no history of steroid treatment. GERD patients were recruited at the primary visit contingent upon positive Bravo pH testing. Exclusion criteria for EoE and GERD included active severe esophagitis (Los Angeles esophagitis Grade C and above), evidence of mechanical obstruction due to peptic stricture (GERD), long-segment Barrett’s metaplasia, unstable medical illness with ongoing diagnostic work-up and treatment, current drug or alcohol abuse or dependency, current neurologic or cognitive impairment which would make the patient an unsuitable candidate for a research trial, severe mental illness, pregnancy and bleeding diathesis or need for anticoagulation that cannot be stopped for endoscopy. Biospecimen Reporting for Improved Study Quality data including age, sex, and race is detailed in **Table 1**.

### ScRNA-seq sample preparation, library preparation and sequencing

Esophageal mucosal biopsies from proximal and distal esophagus were processed immediately following collection and treated separately. Tissue was digested in Dispase (Corning, Corning, NY) diluted in HBSS containing 10 μM HEPES 10 μM and 10 μg/mL DNase I at 37 ⁰C for 15 min with 1500 rpm agitation, followed by digestion in 0.25% trypsin containing 10 μM HEPES and 10 μg/ml DNase I for 20 min at 37 ⁰C with agitation. The cell suspension was filtered through a 40 μm strainer followed by 12 min and 6 min centrifugation at 500g at 4⁰C. Resuspended pellets were filtered through a 40 μm flowmi filter (SP Bel-Art, Wayne, NJ), and measured for cell count and viability using the Cellometer Auto2000 (Nexcelom Bioscience, Lawrence, MA). All cell suspensions met an 85% minimum viability. 16,000 cells were loaded into the Chromium iX Controller (10X Genomics, Pleasanton, CA) on a Chromium Next GEM Chip G (10X Genomics) to capture ∼10,000 cells per sample and were processed for encapsulation according to the manufacturer’s protocol. The cDNA and library were generated using the Chromium Next GEM Single Cell 3’ Reagent Kits v3.1 (10X Genomics) and Dual Index Kit TT Set A (10X Genomics) according to the manufacturer’s manual. Quality control was performed by Agilent Bioanalyzer High Sensitivity DNA kit (Agilent Technologies) and Qubit DNA HS assay kit (Invitrogen, Waltham, MA) for qualitative and quantitative analysis, respectively. The multiplexed libraries were pooled and sequenced on Illumina Novaseq6000 sequencer (Illumina, San Diego, CA) with 100 cycle kits using the following read length: 28 bp Read1 for cell barcode and UMI, and 90 bp Read2 for transcript. Library preparation and sequencing was done at Northwestern University NU-seq facility core. The GRCh38 transcriptome was used as reference for alignment and feature counting using Cell Ranger (V4.0.0/6.0.0/6.1.0, 10X Genomics).

### Data filtering, integration, and clustering

Filtered matrix files were processed as Seurat objects in the Seurat R package 4.2.0 (56) with a minimum threshold of expression in ≥5 cells per gene. Each dataset was filtered to exclude cells with total gene counts <400 and total unique gene counts <100. Datasets were individually normalized, scaled, and processed to calculate variable features using Seurat’s SCTransform workflow. Stricter quality control filtering was performed across all samples to remove cell populations with low total counts of unique genes or cell populations with high mitochondrial gene percentage (mean >25%) following integration. Individual filtered samples were integrated using reverse principal component analysis dimensional reduction. Dimensionality reduction was performed followed by calculation of UMAP embeddings, nearest neighbors, and graph-based clustering. Clusters were annotated according to the expression of known cell-specific gene markers and confirmed against the transcriptional profiles identified by Seurat’s function FindAllMarkers.

### Epithelial cluster and compartment identification

Epithelial cells were subsetted and reintegrated on a per-sample basis using the Seurat integration pipeline described above. Integration anchors were calculated against HC samples as reference. Principal component analysis (PCA) was performed and the first 30 PCs were included for downstream analysis. Optimal clustering resolution of 0.5 was determined using Clustree (**Supplemental Figure 2A**). Epithelial clusters were annotated according to expression of known genes in HC as previously described (22) and confirmed against the transcriptional profiles identified by FindAllMarkers, performed on HC cells. Clusters were combined into parental epithelial compartments (Basal, Suprabasal, Superficial) based on the expression of established markers (**Supplemental Figure 3B**) (7, 22, 34, 35).

### Cell cycle and proliferation analysis

Seurat’s function CellCycleScoring (33) was used to assign the cell cycle phase of each cell. Cells exhibiting a weak predicted score for S and G2/M were classified as G0/G1 phase. Expression of the markers *KRT15* and *DST* identified B1 and B2 epithelial clusters as quiescent and distinguished G0 and G1 phase. Proliferation rates were calculated as the proportion of cells in different phases of the cell cycle or the percentage of cells expressing the G2/M phase marker *MKI67*. During the SCTransform workflow, cell cycle was not regressed, allowing EEC to cluster based on quiescence, S-phase, G2/M-phase, and progressive stages of differentiation, confirmed using the expression of marker genes for each stage. Cell proportion in each population was additionally used to assess proliferation rates.

### Detection of DEGs, gene expression analysis and gene set enrichment analysis

Identification of DEGs between cell clusters was performed using FindAllMarkers, with filtering for significantly upregulated genes with logFC > 0.25. For differential expression analysis comparing expression profiles between like cell identities across disease conditions, the per-sample population mean gene expression was calculated from the normalized RNA assay. Tested genes were filtered by a lower minimum percentage (min.pct) threshold of 5% to 10%, which is the percentage of cells expressing a given gene per cell group. The R package edgeR 3.36.0 (58–60) was utilized to create a DEGList object, followed by calculation of normalization factors and counts per million. The logFC was computed and significance was determined using the Wilcoxon rank-sum test, with false discovery rate (FDR) *P* value adjustment to correct for multiple comparisons. DEGs were filtered based on an FDR-adjusted *P* value < 0.05 and |log2FC| > 0.25, unless a more stringent threshold was specified. To visualize the percentage of cells expressing a gene across clusters, the percentage expression in each cluster was calculated using a minimum expression threshold to filter cells with negligible expression of the gene. Pathway enrichment analyses were performed on DEGs filtered for logFC and significance based on FDR adjusted *P* value, as mentioned above. The analysis of positively and negatively regulated DEGs was completed using the Ingenuity Pathway Analysis (IPA, Qiagen, Germantown, MD) software. For the analysis of DEGs changing in only one direction, pathway enrichment was performed with either Metascape or ClusterProfiler (61–63).

### Transcription factor analysis

Transcription factor (TF) analysis was performed using the R package Enrichr 3.1.0 (36, 64, 65). To identify upstream TFs that regulate input genes, DEGs were calculated across all EEC between disease conditions and filtered for |logFC| > 1 and FDR-adjusted *P* value < 0.05. Hierarchical clustering was performed on population z-scores of DEGs across healthy and EoE epithelial compartments. Relevant hierarchical clusters were selected and used as input for Enrichr analysis with either the ChEA3 2022 ChIP-seq database or the TF Perturbations followed by Expression GEO Signature database.

### Heatmap visualization, population z-score calculation and hierarchal clustering

Gene sets displayed in heatmaps, including gene sets incorporated from external sources, were confirmed as changed in EoE with differential expression testing filtered based on FDR-adjusted *P* value < 0.05 and minimum logFC threshold. To calculate population z-scores, average population expression values were derived from the normalized RNA assay and scaled by the mean and standard deviation calculated across all populations. All heatmaps show population z-scores unless otherwise indicated. Hierarchical clustering using the hclust function from the R Stats package 3.6.2 was performed on population z-scores, using the ward.D2 clustering method (66, 67) and the Pearson distance method (68, 69). Heatmaps were generated using the R package Complex Heatmap 2.10.0 (68, 69). To create heatmaps displaying logFC from HC cells, DEGs were identified by conducting differential expression analysis between EoE and HC cells. LogFC values were then calculated per epithelial cell compartment in either EoE or GERD relative to HC. For gene ordering, hierarchical clustering was performed on population z-scores calculated across HC and EoE compartments.

### Gene signature score analysis and functional analysis

Gene signatures were generated using Seurat’s function AddModuleScore. Quiescent and superficial gene signatures were defined using HC cells from our scRNA-seq dataset. Differential expression analysis was performed comparing either quiescent epithelial clusters (B1-B2) or superficial clusters (SF1-SF2) to the remaining epithelium. DEGs were filtered for FDR-adjusted *P* value < 0.05 and ranked by logFC, with the top 100 selected. Quiescent and superficial signature scores were plotted using the ggplot2 R package’s geom_density_2d function (70), with consistent binning applied across all compared conditions. TF-regulated gene signatures were identified via enrichment analysis (EnrichR) (36, 64, 65) or sourced from external experiments (**Supplemental Table 5**). The datasets included in this analysis were previously published (38–40, 71). Gene signatures were also calculated from co-expressed gene modules identified using Monocle3, which were also used as input to the stringDB R package (75) to infer protein-protein interactions.

### Pseudotime analysis

Pseudotime analysis was performed using the R package Monocle3 1.0.0 (72–74). Individual samples were log_2_ normalized, scaled, merged using Seurat’s merge function, dimensionally reduced, batch corrected using the fast mutual nearest neighbors (FMNN) method by individual sample, and UMAP embeddings were calculated. A CellDataSet object was created with normalized and scaled counts for 2000 variable genes and reduction feature loadings calculated by FMNN. Monocle3’s function learn_graph was used to infer a trajectory graph from the UMAP embeddings, with Euclidean distance ratio of 1, geodesic distance ratio of 0.5, and a minimum branch length of 10. Cells within the S-phase epithelial cluster B3 were assigned a root state of pseudotime 0. Increasing pseudotime values of cells committed to becoming quiescent are depicted to the left, and pseudotime values of cells committed to differentiation to the right on pseudotime axes.

For EoE samples only, Monocle3’s function graph_test was utilized to identify genes with differential expression along the trajectory. Identified genes were clustered into modules of co-expressed genes with corresponding gene signatures calculated. To determine the most represented module in each cell, each module gene signature was scaled and centered between -2 to 2 across all cells. Each cell was assigned to the module exhibiting the highest scaled scoring. To visualize the expression of genes or signatures across pseudotime-ordered cells, we plotted the gene expression or gene signature score for each cell and calculated local mean expression values using local weighted regression fitting of the data by the loess method. For the calculation of coexpression, cells were assessed on a binary basis for expression of all examined genes or gene signatures (value = 1) or expression of less than all or none of the examined genes or gene signatures (value = 0). The values were plotted for each cell ordered in pseudotime and local mean values were calculated using the loess method.

### Imputation of cell populations in the GERD scRNA-seq data from the EoE and HC scRNA-seq dataset

An imputation was performed on each cell in the processed GERD epithelial dataset to determine the analogous cell population in the integrated HC and EoE epithelial dataset using Seurat’s MapQuery function. Cluster labels were assigned based on the maximum prediction score for each of the query cells.

### Immunohistochemistry and scoring

Immunostaining was performed on formalin-fixed, paraffin-embedded esophageal mucosal biopsies as previously described (49). Briefly, heat-indued antigen retrieval was performed for 30 min in Buffer A (Electron Microscopy Sciences, pH 6). Tissue sections were blocked using 0.3% H*_2_*O_2_, streptavidin/biotin incubation and Starting Block blocking buffer (Thermo Fisher Scientific). Primary and secondary specific antibodies were added (**Supplemental Table 9**) and detection was performed as previously described (49). Images were acquired on a Nikon Eclipse Ci microscope with a Nikon DS-Ri2 camera and NIS Elements software. H&E staining was performed by the Robert H. Lurie Comprehensive Cancer Center Pathology Core. Image analysis was performed using Fiji software (76). H&E-stained slides were evaluated for BCH according to EoE-HSS (30). For staining quantification, positive cell fraction was calculated as the percentage of positively stained cells compared to the total cell count. For intensity quantification, nuclei were identified by thresholding, mask conversion, watershed segmentation, and particle analysis, followed by measurement of average inverted intensity (greyscale units) following background subtraction.

### Multispectral fluorescence staining and imaging

Multispectral fluorescence staining was performed using the Opal 6-Plex Detection kit (Akoya Biosciences, Marlborough, MA) using formalin-fixed, paraffin embedded tissue sections. Slides were baked at 60 °C for 15 min and deparaffinized with the Leica Bond Dewax solution (Leica Biosystems, Deer Park, IL), followed by heat-based antigen retrieval using Bond Epitope Retrieval Solution 1 (Leica Biosystems) for 30 min. Using the Leica Bond Rx™ Automated Stainer (Leica Biosystems), slides were incubated with primary antibodies followed by the appropriate secondary horseradish peroxidase-conjugated polymer. Incubation was next performed with a unique Opal dye permitting fluorophore covalent bonding to the horseradish polymer. Heat-based retrieval with Bond Epitope Retrieval 1 (Leica Biosystems) was finally performed for 20 min. Slides were subjected to sequential rounds of staining. Primary antibodies, concentrations, and associated fluorophores are detailed in **Supplemental Table 9**. Sections were counterstained with Spectral DAPI and mounted with ProLong Diamond Antifade Mountant (Thermo Fisher Scientific). Images were acquired using the Vectra3 microscope (Akoya Biosciences) and Phenochart Whole Slide Viewer (Akoya Biosciences). Post-acquisition image adjustments were performed using InForm Automated Image Analysis Software (Akoya Biosciences) and Fiji (76).

### Data and code availability

All data from this study or code not included within the article are available from the corresponding author upon reasonable request. The raw sequencing files and processed barcode and feature matrices are available at Gene Expression Omnibus (#GSE218607). Scripts are available at https://github.com/Tetreault-Lab/Tetreault-scRNA-Human_EoE_Esophagus-2023.

### Statistics

Statistical analyses were performed using R version 4.1.1. Descriptive statistics are displayed as mean ± standard error of the mean for continuous variables and frequency counts for categorical variables. For non-normally distributed continuous data, Wilcoxon rank-sum test was used. When testing multiple conditions, multiple comparison adjustment was employed.

## Human Study approval

Procedures using human tissue were performed by the Digestive Health Foundation Biorepository with approval from the Northwestern Institutional Review Board (study STU00208111). Written informed consent was received prior to participation.

## Authors contributions

MHC was involved in designing research studies, conducting experiments, acquiring data, analyzing data and writing the manuscript. ALK was involved in designing research studies and analyzing data. PJK, DAC, NG and JEP were involved in designing research studies and writing the manuscript. DRW and KAW were involved in designing research studies, analyzing data and writing the manuscript. MPT was involved in designing research studies, analyzing data, providing reagents and writing the manuscript.

## Acknowledgements

This work was supported by: NIH NIDDK P01 117824 and Digestive Health Foundation to MPT, PJK and JEP; NIH NHLBI F31HL147413 to MHC; NIH NIDDK R01DK121159 to KAW; by the Robert H. Lurie Comprehensive Cancer Center (NIH NCI CCSG P30 CA060553) through the Northwestern University Pathology Core and Facility and by the NU-Seq Core Facility (1S10OD025120). This research was supported in part through the computational resources and staff contributions provided for the Quest high performance computing facility at Northwestern University which is jointly supported by the Office of the Provost, the Office for Research, and Northwestern University Information Technology. This research was also supported in part through the computational resources and staff contributions provided by the Genomics Compute Cluster which is jointly supported by the Feinberg School of Medicine, the Center for Genetic Medicine, and Feinberg’s Department of Biochemistry and Molecular Genetics, the Office of the Provost, the Office for Research, and Northwestern Information Technology. The Genomics Compute Cluster is part of Quest, Northwestern University’s high performance computing facility.

