## Supplemental Figures for "Suprabasal cells retaining stem cell identity programs drive basal cell hyperplasia in eosinophilic esophagitis"

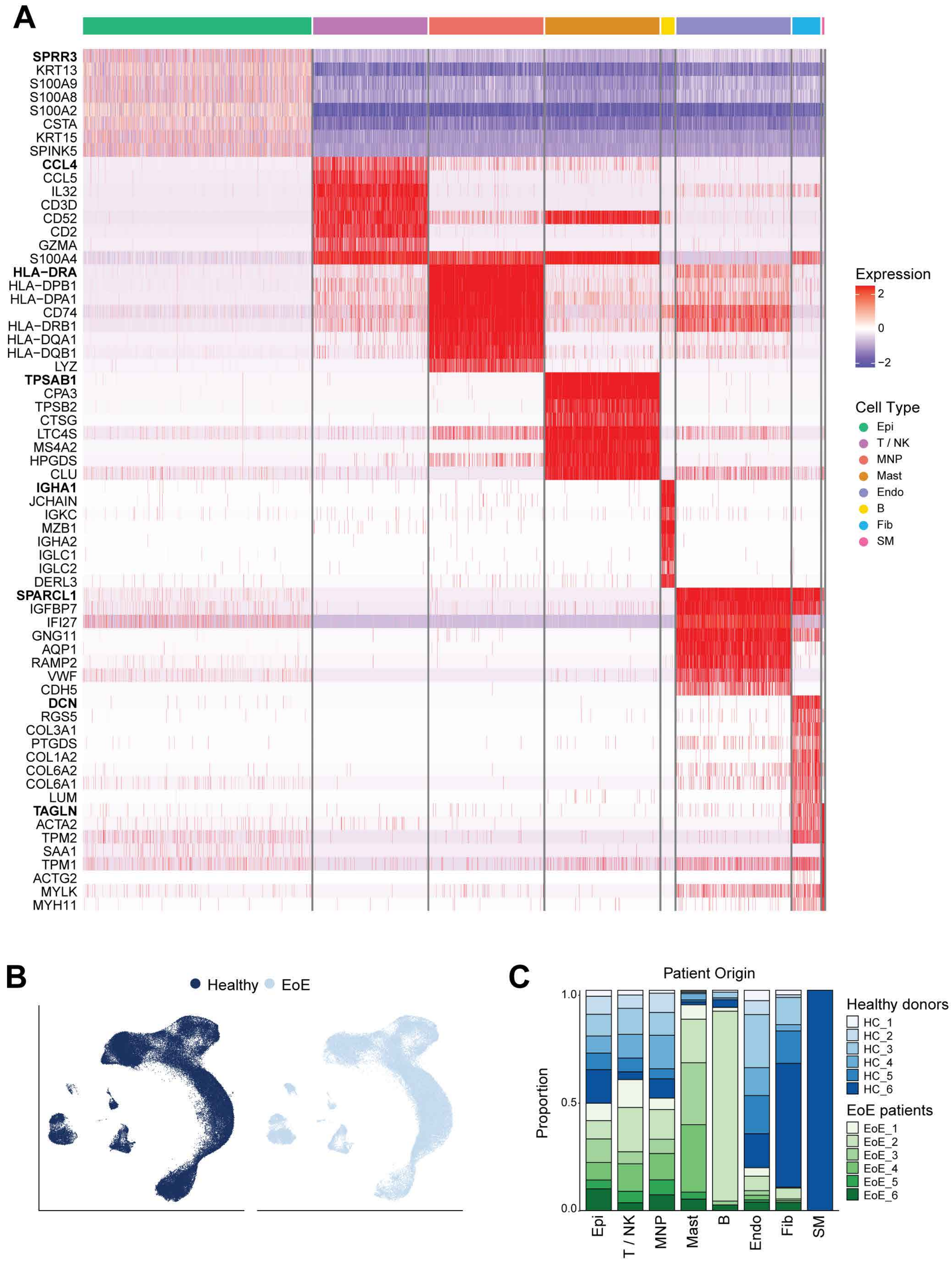


**Supplemental Figure 1. Characteristics of esophageal mucosal cells from EoE patients and healthy subjects.** (**A**) Heatmap of the the top 8 genes expressed per cell type. Ranked by average logFC, FDR < 0.05, row-normalized expression z-scores. (**B**) UMAP embedding of scRNA-seq data seperated by HC or EoE condition. (**C**) Bar plot depicting the frequency of each esophageal cell type in individual HC and EoE samples. Epithelial cells (Epi), T cells / natural killer cells (T / NK), mononuclear phagocytes (MNP), mast cells (Mast), B cells (B), endothelial cells (Endo), fibroblasts (Fib), or smooth muscle (SM).

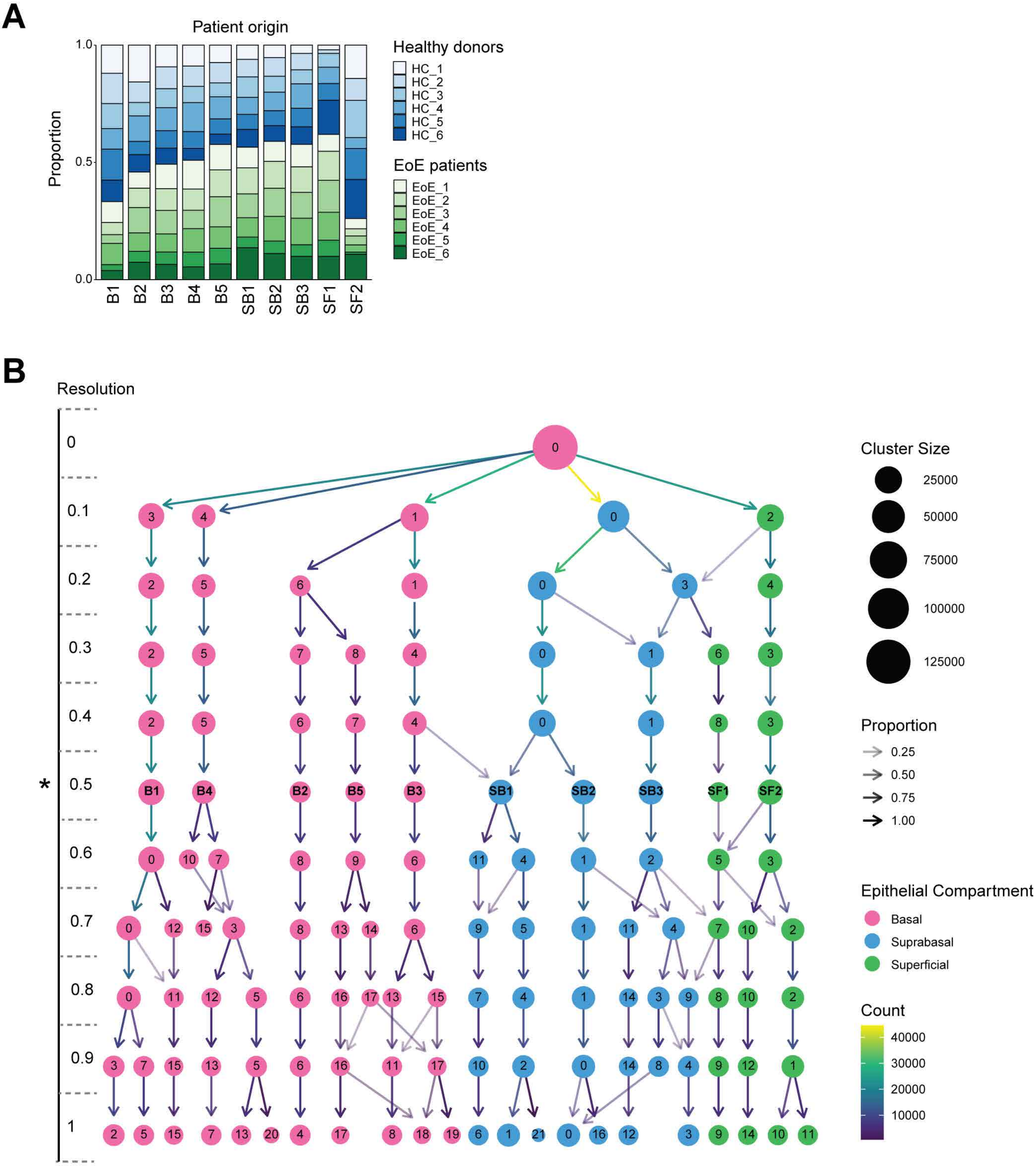


**Supplemental Figure 2. Characterization of human esophageal epithelial cell populations in HC and EoE.** (**A**) Bar plot depicting the frequency of each epithelial cluster in individual HC and EoE samples. (**B**) Clustering tree of shared nearest neighbor clustering on the epithelial dataset using 30 principal components across resolutions 0-1 with 0.1 increments. Dot size indicates cell count of each cluster, arrow transparency denotes proportion of cells moving to the indicated cluster in the next resolution increment as function of total cells in the originating cluster, arrow color indicates cell counts moving to the indicated cluster, dot color indicates epithelial compartment identity of the majority of cells in each cluster, star represents the selected resolution. Basal (B), Suprabasal (SB), Superficial (SF).

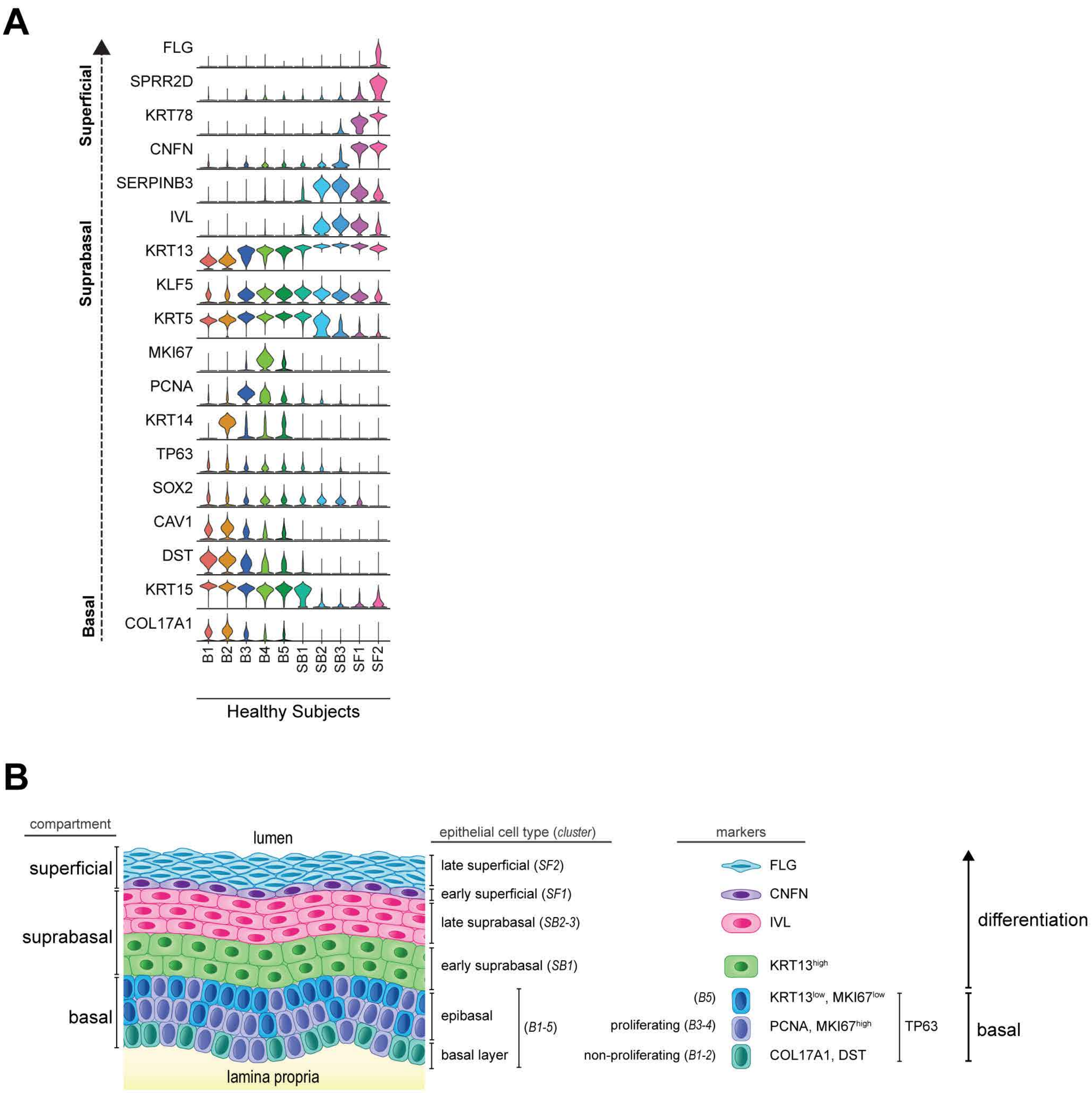


**Supplemental Figure 3. Annotation of epithelial cell clusters.** (**A**) Violin plot showing expression of known epithelial basal and differentiated markers across epithelial clusters in healthy controls. (**B**) Schematic summary of the three epithelial compartments of the adult esophagus, with their component epithelial clusters and corresponding gene markers. Basal (B), Suprabasal (SB), Superficial (SF).

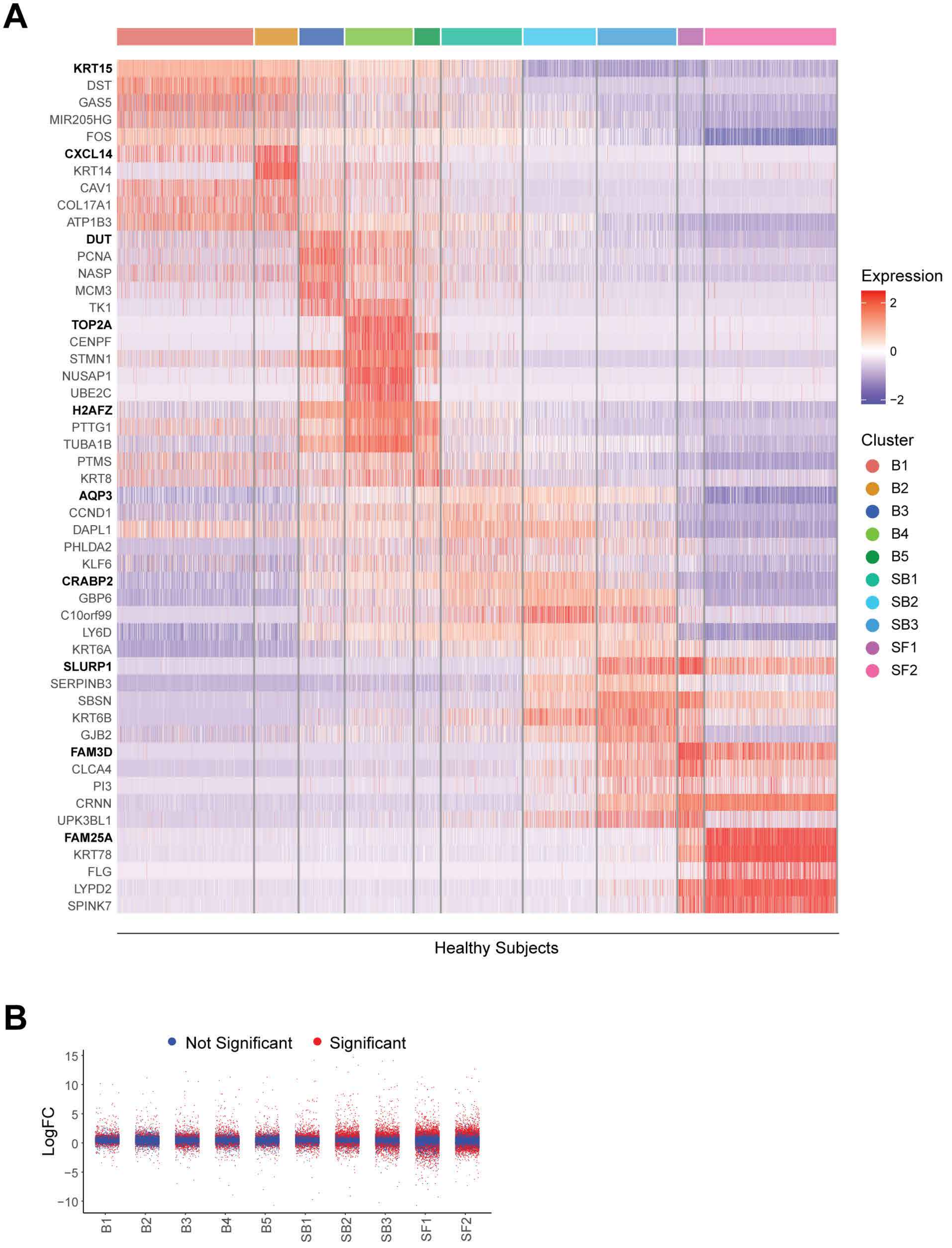


**Supplemental Figure 4. Transcriptional profile of epithelial cell clusters in healthy and EoE subjects.** (**A**) Heatmap of the the top 5 genes expressed per epithelial cluster. Ranked by average logFC, FDR < 0.05, row-normalized expression z-scores. (**B**) LogFC of DEGs in each epithelial population calculated in EoE subjects as compared to HC. Significantly changed genes (FDR < 0.05) are indicated in red. Basal (B), Suprabasal (SB), Superficial (SF).

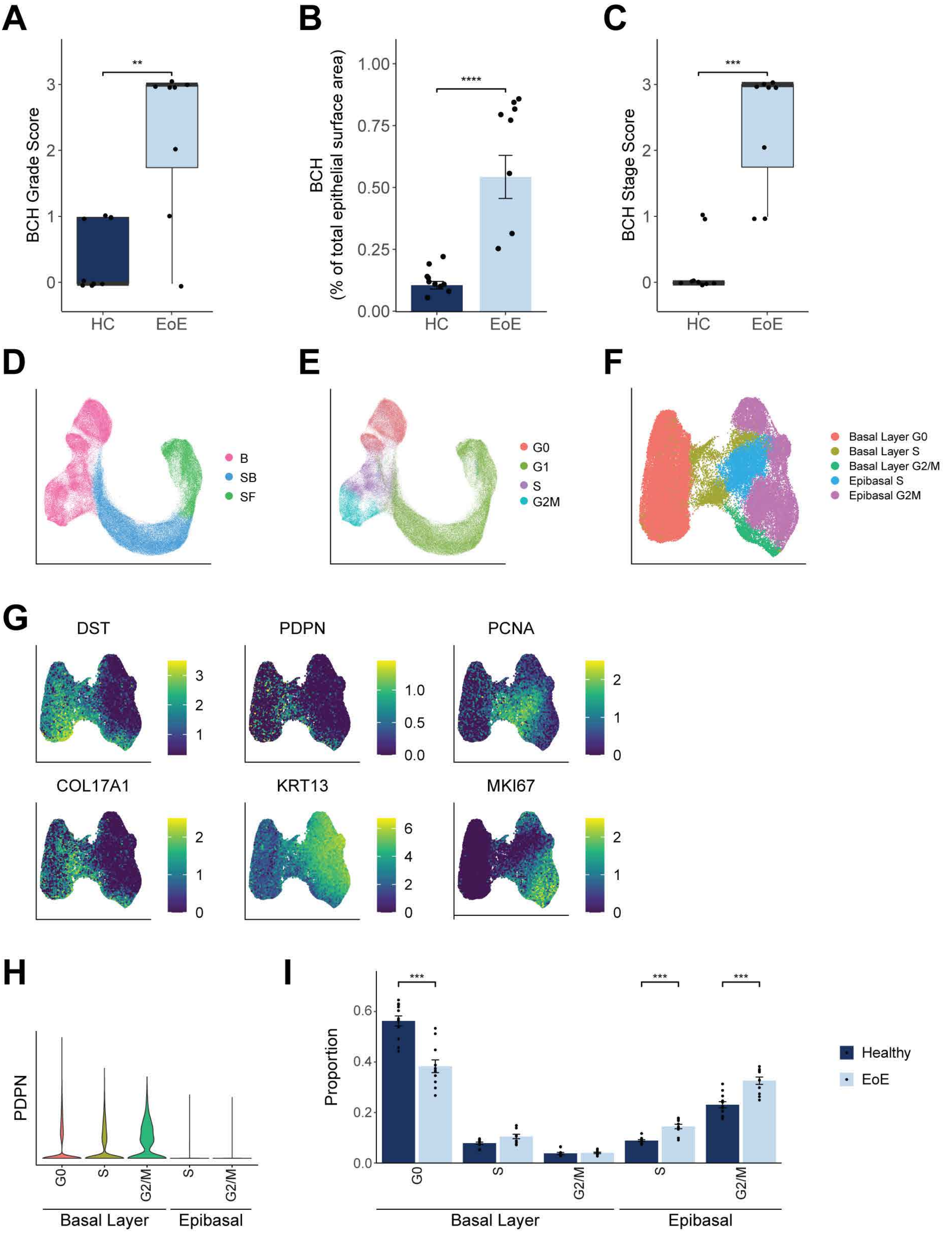


**Supplemental Figure 5. Examination of BCH and proliferation in the esophageal epithelial cell compartments in EoE.** (**A-C**) Quantification of BCH from HC and EoE esophageal epithelial sections from the scRNA-seq cohort shown in **Figure 3A**. BCH area is measured as defined in **Figure 3A**, underneath the dashed white lines. (**A**) Box plot showing the BCH grade score for HC and EoE epithelial sections. (**B**) Bar plot showing the proportion of the epithelium occupied by the basal zone as a function of surface area. (**C**) Box plot showing the BCH stage score for HC and EoE epithelial sections. (**D**) UMAP of the scRNA-seq dataset colored by epithelial cell compartment. (**E**) UMAP of the scRNA-seq dataset colored by cell cycle phase. (**F**) Sub-clustered UMAP of the quiescent and proliferating epithelial clusters (B1-B5) annotated according to basal layer or epibasal cell identity and cell cycle phase. (**G**) UMAP of the quiecent and proliferating epithelial clusters with each cell colored by the expression of epithelial markers *DST*, *COL17A1*, *PDPN*, *KRT13*, *PCNA*, or *MKI67*. (**H**) Violin plot showing *PDPN* expression across basal layer and epibasal populations within the basal cell compartment. (**I**) Bar plot showing the proportion of basal layer and epibasal cells in each phase of the cell cycle, as assigned in (**F**). All indicated *P* values were determined using Wilcoxon signed-ranked test, with Benjamini & Hochberg adjustment for multiple comparisons specifically applied in (I). ***P* ≤ 0.01, ****P* ≤ 0.001, *****P* ≤ 0.0001. Basal (B), Suprabasal (SB), Superficial (SF), Healthy Controls (HC).

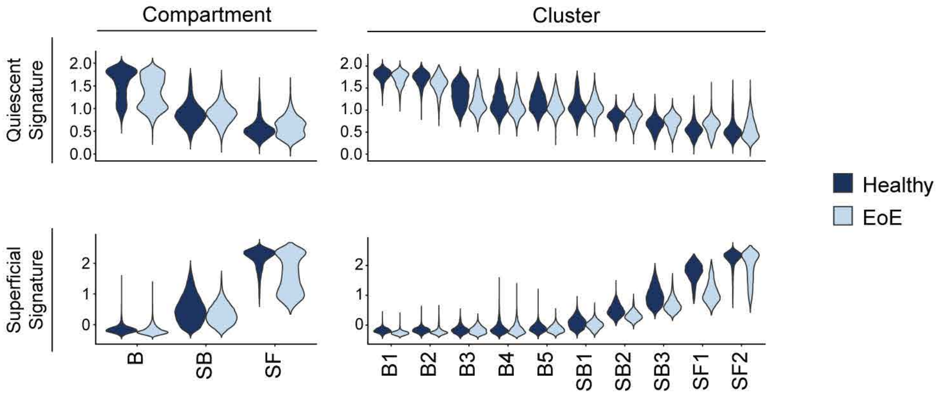


**Supplemental Figure 6. Alteration in the quiescent-basal-differentiation transition process in the esophageal epithelium of EoE.** Violin plots showing the scores for quiescent gene signature and superficial gene signature between disease states in either epithelial compartments or epithelial cell clusters. Basal (B), Suprabasal (SB), Superficial (SF).

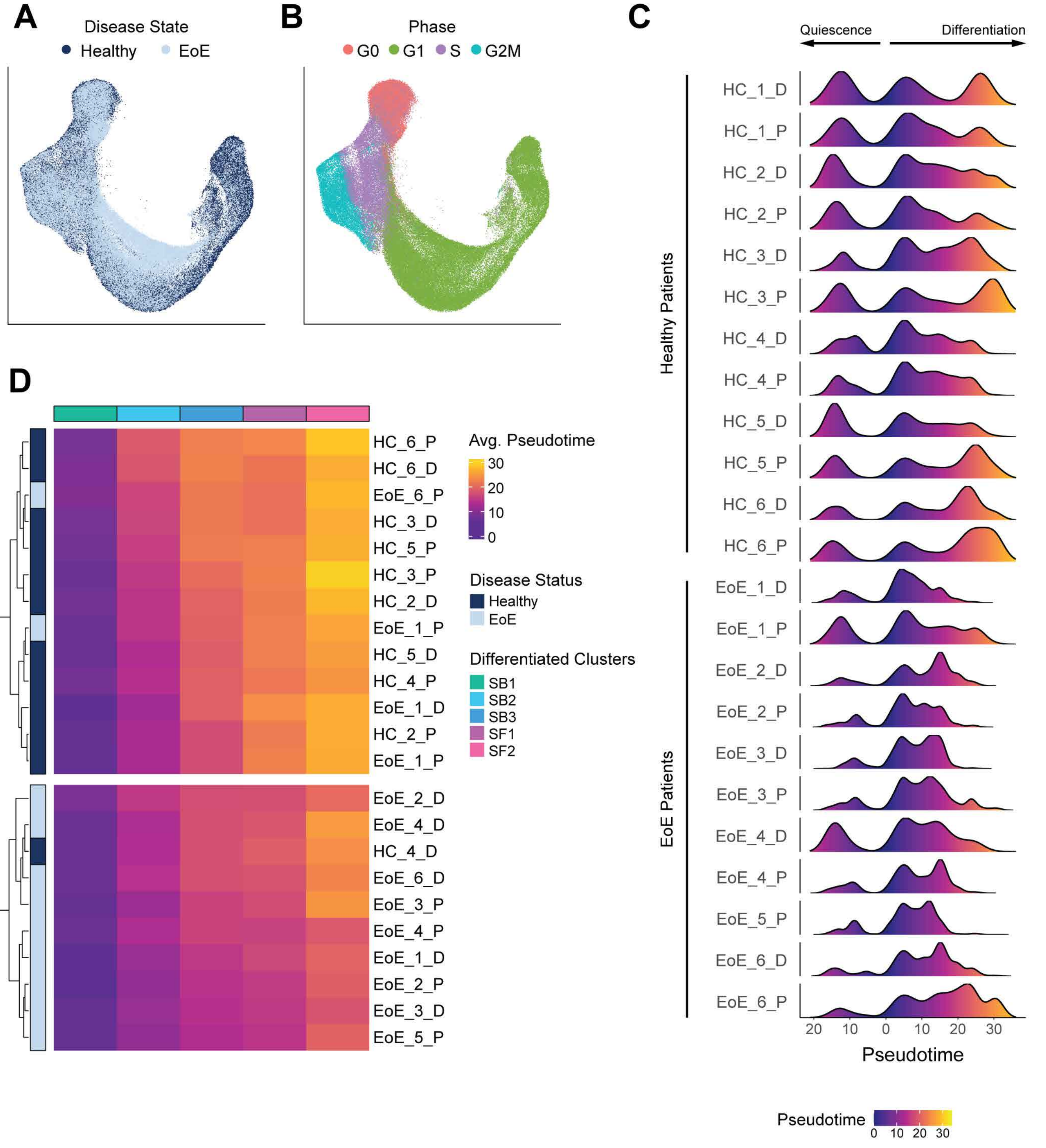


**Supplemental Figure 7. Average epithelial cluster pseudotime assignment can distinguish EoE from HC patient samples.** (**A**) UMAP showing merged scRNA-seq datasets of EEC from EoE and HC patients, colored by disease state (**A**) or cell cycle phase (**B**). (**C**) Ridgeline plots showing the distribution of pseudotime values between epithelia of individual healthy or EoE patient samples. (**D**) Heatmap of raw (non-scaled) average pseudotime values for each differentiated epithelial cluster in EoE and healthy patient samples, analyzed by hierarchical clustering. The two top-level hierarchical clusters are spacially separated for emphasis. Basal (B), Suprabasal (SB), Superficial (SF).

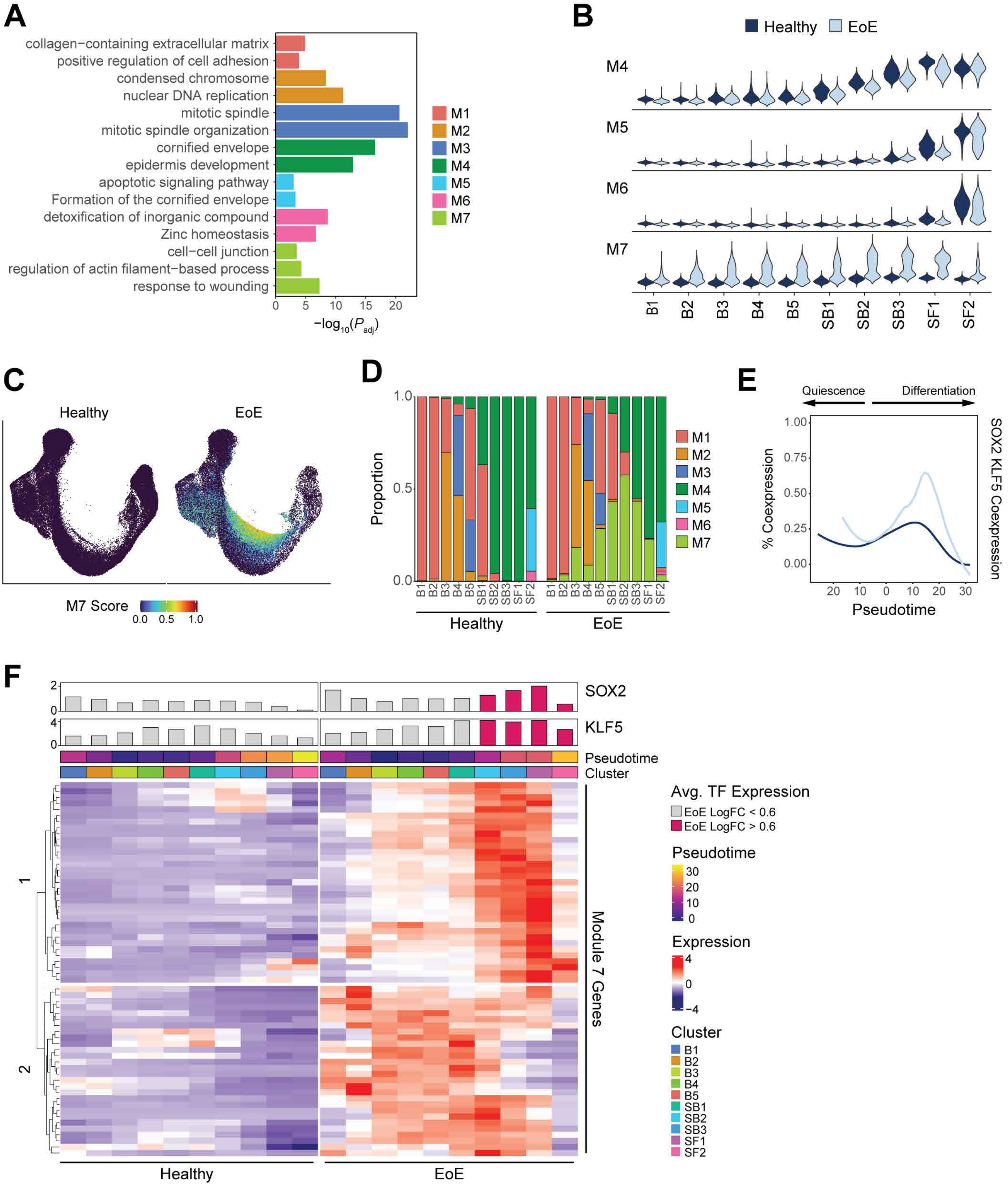


**Supplemental Figure 8. Trajectory-dependent gene programs altered in EoE mediate tissue remodeling and may be downstream of SOX2 and/or KLF5.** (**A**) Bar plot showing significantly enriched pathways as identified by gene set enrichment analysis for each EoE pseudotime-dependent gene module. Color indicates associated module. (**B**) Violin plots displaying the gene signature scoring of modules that exhibit meaningful change between EoE and HC epithelial clusters. (**C**) UMAP of the merged scRNA-seq data of esophageal epithelial cells from EoE and healthy patients colored by module 7 gene signature score. (**D**) Barplot showing epithelial clusters in EoE or HC broken down by cell proportions defined by highest-scoring pseudotime-dependent module. (**E**) The percentage of epithelial cells exhibiting coexpression of *SOX2* and *KLF5* is shown in HC and EoE patients across cells ordered by pseudotime. Lines represent moving averages calculated by loess regression across all cells. (**F**) Heatmap of genes in module 7 shown across epithelial clusters, separated by disease state. Shown as row-normalized expression z-scores, ordered by hierarchical clustering. For each cluster, mean $\log_2$ gene expression for *SOX2* and *KLF5* is shown in a top annotation as a bar plot. Bar color indicates whether logFC between EoE and HC clusters is above (magenta) or below (grey) 0.6-fold. Mean pseudotime value for each cluster is also depicted in top annotation. Basal (B), Suprabasal (SB), Superficial (SF), Module (M), Transcription Factor (TF), adjusted *P* value ($P_{adj}$).

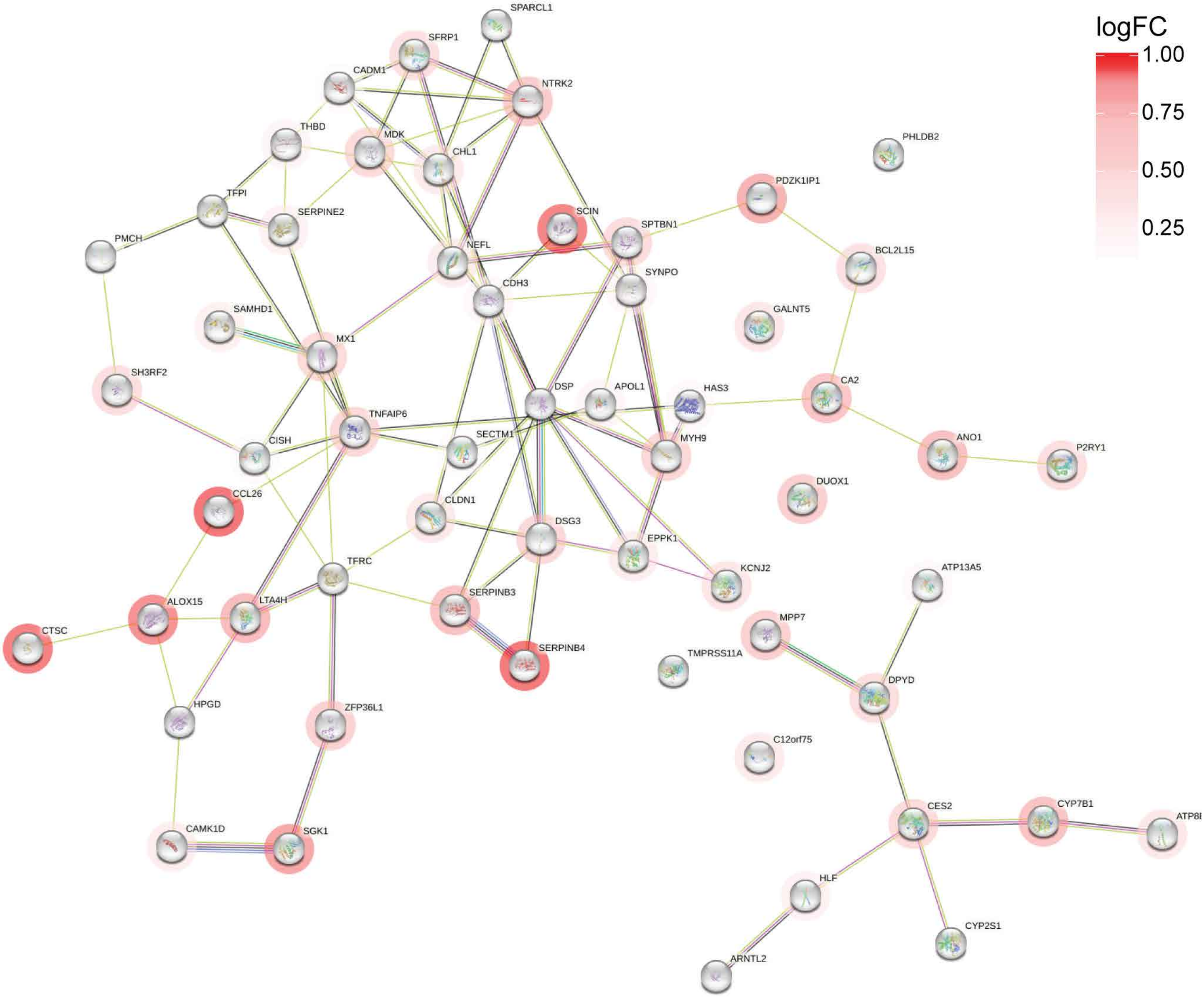


**Supplemental Figure 9. Protein-protein interactions in EoE-specific trajectory-dependent genes.** Known interactions are mapped between all module 7 genes. Lines indicate known interactions, halo color intensity indicates LogFC between differentiated cells in EoE versus HC.

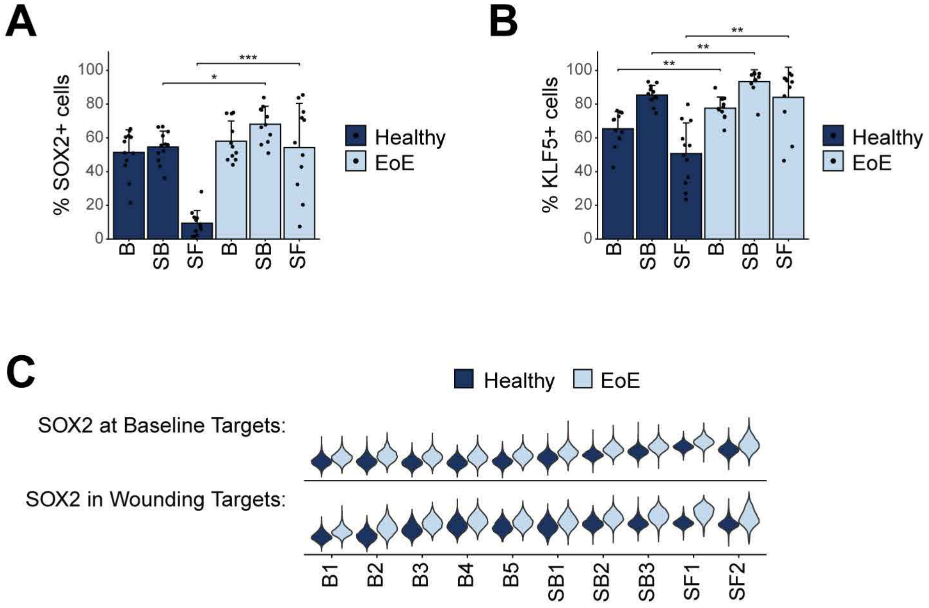


**Supplemental Figure 10. *SOX2*, *KLF5* and their transcriptional targets exhibit increased expression in the esophageal epithelial differentiated compartment in EoE.** (**A**) Bar plots showing the percentage of cells expressing meaningful levels of *SOX2* (A) or *KLF5* (**B**) in each epithelial compartment in either EoE or HC. Data are shown as mean ± SEM. Indicated *P* values were determined using Wilcoxon signed-ranked test with Benjamini & Hochberg adjustment for multiple comparisons. **P* ≤ 0.05, ***P* ≤ 0.01, ****P* ≤ 0.001. (**C**) Violin plots depicting the scoring of gene signatures of transcriptional targets of SOX2 at baseline or following wounding for each epithelial cluster. Basal (B), Suprabasal (SB), Superficial (SF). Details on SOX2 baseline dataset and SOX2 following wounding dataset are provided in Methods.

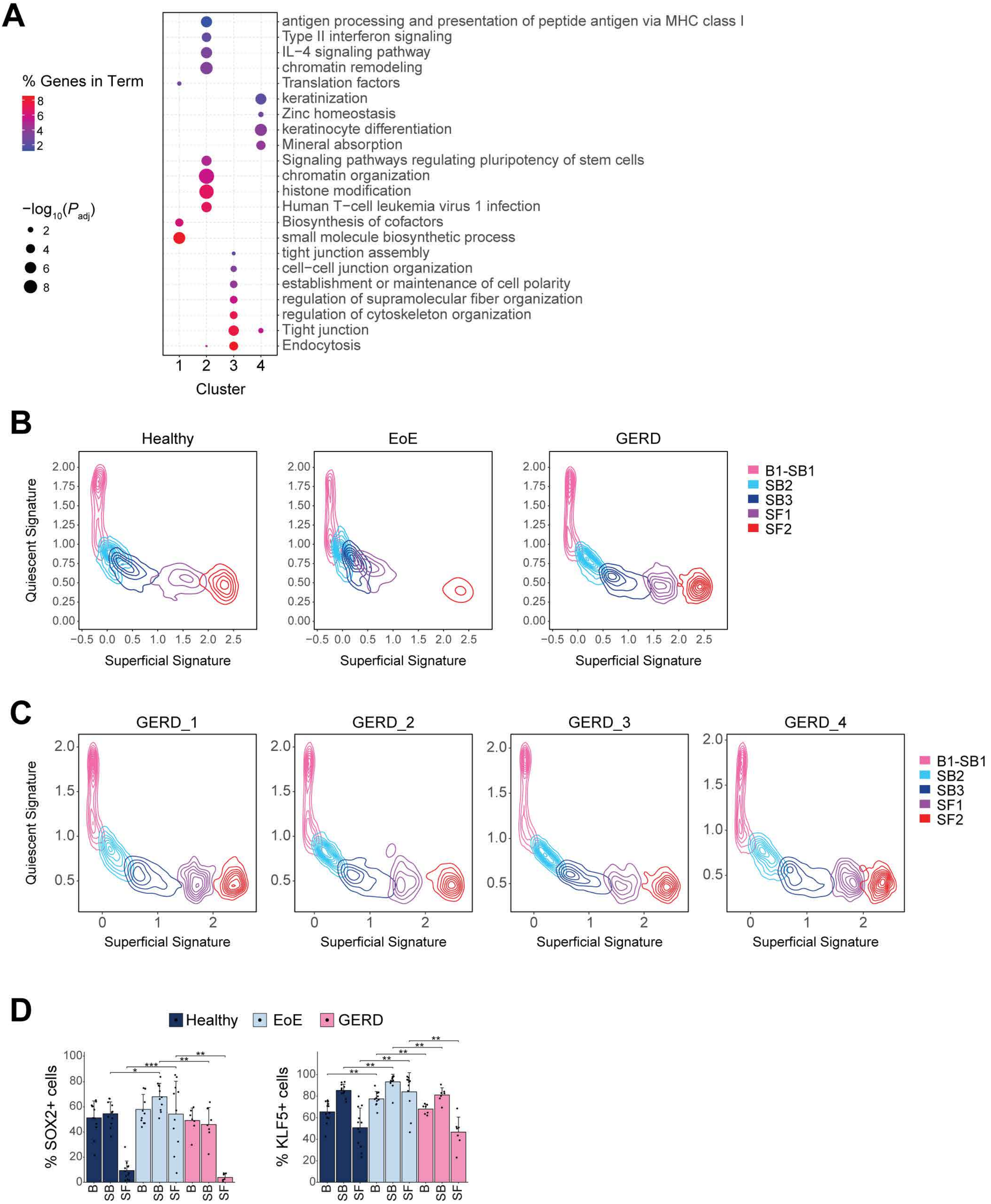


**Supplemental Figure 11. Differentiated esophageal epithelial compartments in GERD do not demonstrate increased basal-like identity or aberrant expression of basal transcriptional regulators.** (**A**) Top enriched terms for each hierarchical cluster identified in **Figure 10C**. Color indicates -$\log_{10}$ of adjusted *P* value ($P_{adj}$), dot size represents percentage of genes from each cluster along each term. (**B**) Contour plots showing epithelial cells from HC, EoE or GERD patients plotted along the quiescent gene signature (y-axis) and the superficial gene signature (x-axis). Line color shows cell grouping by indicated epithelial clusters. Clusters are visualized per individual GERD patient in (**C**). (**D**) Bar plots showing the percentage of cells expressing meaningful levels of *SOX2* or *KLF5* in each epithelial compartment in either EoE, GERD or HC. Data are shown as mean ± SEM. Indicated *P* values were determined using Wilcoxon signed-ranked test with Benjamini & Hochberg adjustment for multiple comparisons. *$P \leq 0.05$, **$P \leq 0.01$, ***$P \leq 0.001$. Basal (B), Suprabasal (SB), Superficial (SF).

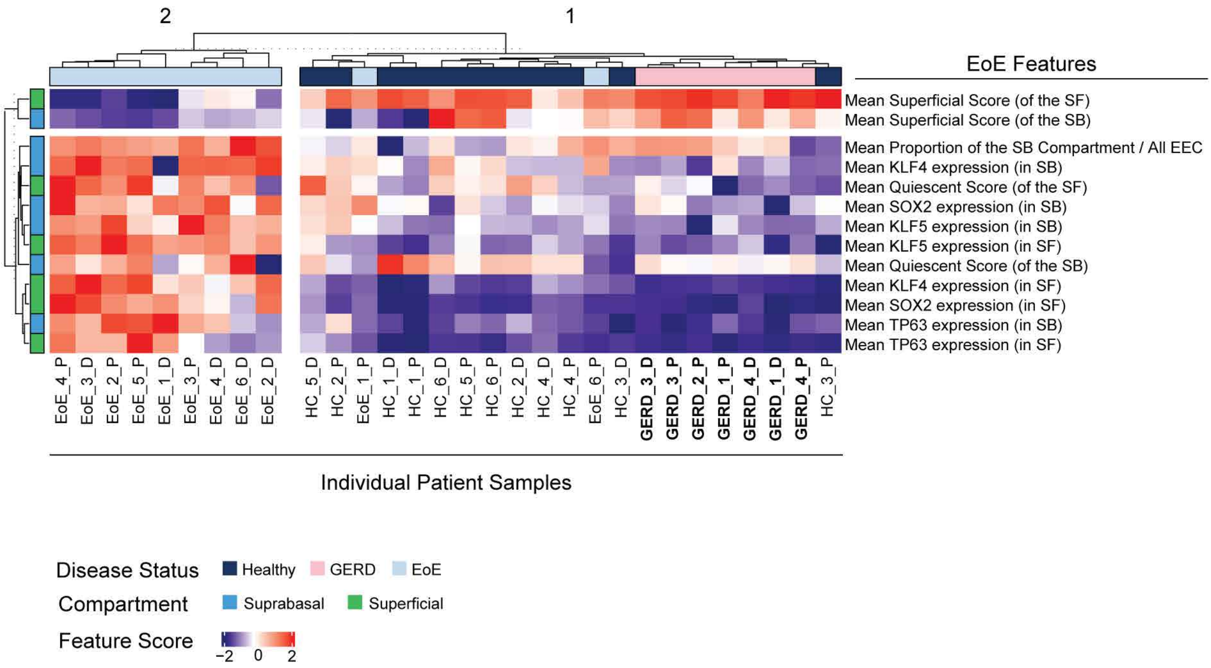


**Supplemental Figure 12. Suprabasal and superficial epithelial cell compartments in GERD patients do not demonstrate an impaired basal-differentiation transition.** Heatmap of identified EoE features calculated across suprabasal or superficial compartments for HC, EoE, or GERD patients, analyzed by hierarchical clustering. The two top-level hierarchical clusters are spacially separated for emphasis. Feature scoring is shown as row-normalized z-scores, top annotation indicates disease state, left annotation indicates epithelial cell compartment analyzed, columns represent individual patients. Suprabasal (SB), Superficial (SF).
