## Supplemental Tables for "Suprabasal cells retaining stem cell identity programs drive basal cell hyperplasia in eosinophilic esophagitis"

### Table of Contents


**Supplemental Table 1.** Genes included in the quiescent gene signature. …………………...pg 2

**Supplemental Table 2.** Genes included in the superficial gene signature. …………………...pg 5

**Supplemental Table 3.** Genes corresponding to co-expressed modules along the EoE epithelial pseudotemporal trajectory……………………………………………………..….…pg 8

**Supplemental Table 4.** Gene expression profile of module 7 genes in EoE vs healthy subjects. ……………………………………………………………………………………………pg 25

**Supplemental Table 5.** Identification of SOX2 and/or KLF5 downstream targets from Enrichr databases and literature-based analyses. ……………………………………………………...pg 27

**Supplemental Table 6.** Module 7 genes known to be regulated by SOX2-KLF5. …..............pg 167

**Supplemental Table 7.** Gene expression profile of upregulated EoE DEGs known to be regulated by SOX2 and/or KLF5. ......................................................................................…pg 168

**Supplemental Table 8.** Gene expression profile of EoE DEGs known to be regulated by the SOX2-KLF5 interaction.......................................................................................................…pg 191

**Supplemental Table 9.** List of antibodies used in the study. …………………………………pg 196

**Supplemental Table 1.** Genes included in the quiescent gene signature.

| | p_val | avg_log2FC | pct.1 | pct.2 | p_val_adj |
|---|---|---|---|---|---|
| DST | 0 | 2.829762397 | 0.937 | 0.211 | 0 |
| KRT15 | 0 | 2.749389646 | 1 | 0.685 | 0 |
| KRT19 | 0 | 2.027514439 | 0.99 | 0.662 | 0 |
| GAS5 | 0 | 1.991027262 | 0.727 | 0.641 | 0 |
| CXCL14 | 0 | 1.930840409 | 0.497 | 0.095 | 0 |
| MIR205HG | 0 | 1.681103048 | 0.728 | 0.686 | 0 |
| TXNIP | 0 | 1.679755517 | 0.787 | 0.396 | 0 |
| IL1R2 | 0 | 1.641511831 | 0.619 | 0.063 | 0 |
| IL18 | 0 | 1.616816891 | 0.9 | 0.729 | 0 |
| ZFP36L2 | 0 | 1.608479362 | 0.823 | 0.526 | 0 |
| IFITM3 | 0 | 1.606175204 | 0.858 | 0.54 | 0 |
| GPNMB | 0 | 1.602788604 | 0.741 | 0.217 | 0 |
| CAV1 | 0 | 1.57258403 | 0.693 | 0.127 | 0 |
| IGFBP5 | 0 | 1.564140033 | 0.586 | 0.14 | 0 |
| GPX4 | 0 | 1.556344306 | 0.955 | 0.852 | 0 |
| MT-ND2 | 0 | 1.541729919 | 0.95 | 0.884 | 0 |
| COL17A1 | 0 | 1.504913141 | 0.68 | 0.1 | 0 |
| MALAT1 | 0 | 1.497386787 | 0.975 | 0.92 | 0 |
| TSLP | 0 | 1.492823424 | 0.61 | 0.088 | 0 |
| FOS | 0 | 1.469149928 | 0.977 | 0.939 | 0 |
| FTH1 | 0 | 1.451440638 | 0.972 | 0.929 | 0 |
| SLC3A2 | 0 | 1.428585636 | 0.811 | 0.576 | 0 |
| MT-ND1 | 0 | 1.42571156 | 0.953 | 0.891 | 0 |
| MT-ND3 | 0 | 1.409695337 | 0.967 | 0.922 | 0 |
| MT-CYB | 0 | 1.374772617 | 0.979 | 0.933 | 0 |
| ATF3 | 0 | 1.369038577 | 0.828 | 0.673 | 0 |
| ATP1B3 | 0 | 1.315243129 | 0.871 | 0.741 | 0 |
| MT-ATP6 | 0 | 1.308089875 | 0.969 | 0.922 | 0 |
| SNHG8 | 0 | 1.296775833 | 0.926 | 0.798 | 0 |
| FHL2 | 0 | 1.288653373 | 0.659 | 0.194 | 0 |
| MT-CO3 | 0 | 1.262818148 | 0.977 | 0.944 | 0 |
| RPL34 | 0 | 1.252450795 | 1 | 0.994 | 0 |
| C12orf57 | 0 | 1.251320924 | 0.864 | 0.712 | 0 |
| ZFAS1 | 0 | 1.239408574 | 0.965 | 0.911 | 0 |
| CH25H | 0 | 1.196781019 | 0.496 | 0.051 | 0 |
| DLK2 | 0 | 1.177325313 | 0.524 | 0.032 | 0 |
| NOP53 | 0 | 1.162744015 | 0.822 | 0.724 | 0 |
| MT-ND5 | 0 | 1.162255222 | 0.889 | 0.81 | 0 |
| SNHG7 | 0 | 1.141464301 | 0.756 | 0.639 | 0 |
| ASS1 | 0 | 1.110706576 | 0.554 | 0.123 | 0 |
| TSC22D1 | 0 | 1.091835855 | 0.546 | 0.305 | 0 |
| NAP1L1 | 0 | 1.090727748 | 0.812 | 0.664 | 0 |
| CD81 | 0 | 1.085738797 | 0.753 | 0.571 | 0 |
| NPR3 | 0 | 1.085345049 | 0.547 | 0.171 | 0 |
| CTNNAL1 | 0 | 1.0784772 | 0.52 | 0.054 | 0 |

| | | | | | |
|---|---|---|---|---|---|
| THSD4 | 0 | 1.075858062 | 0.567 | 0.254 | 0 |
| SNHG29 | 0 | 1.070984512 | 0.734 | 0.772 | 0 |
| EEF1D | 0 | 1.069898077 | 0.98 | 0.945 | 0 |
| RPS27 | 0 | 1.058941417 | 1 | 0.994 | 0 |
| WNT4 | 0 | 1.051182329 | 0.588 | 0.419 | 0 |
| CIRBP | 0 | 1.048393598 | 0.85 | 0.799 | 0 |
| RACK1 | 0 | 1.044522976 | 1 | 0.99 | 0 |
| MT-ND4 | 0 | 1.044313515 | 0.963 | 0.917 | 0 |
| LAMB3 | 0 | 1.044175468 | 0.558 | 0.166 | 0 |
| RPL10 | 0 | 1.04270105 | 1 | 0.998 | 0 |
| RPL30 | 0 | 1.041514408 | 1 | 0.993 | 0 |
| ITGB1 | 0 | 1.038297217 | 0.623 | 0.389 | 0 |
| RPL5 | 0 | 1.027958819 | 1 | 0.992 | 0 |
| LIMA1 | 0 | 1.019869097 | 0.605 | 0.379 | 0 |
| EPB41L4A-AS1 | 0 | 1.015005509 | 0.646 | 0.384 | 0 |
| RPS9 | 0 | 0.997767959 | 1 | 0.993 | 0 |
| SLC1A3 | 0 | 0.989165554 | 0.48 | 0.093 | 0 |
| RPL13 | 0 | 0.986841391 | 1 | 0.998 | 0 |
| SLC7A1 | 0 | 0.985424125 | 0.624 | 0.378 | 0 |
| RPS4X | 0 | 0.977786324 | 1 | 0.997 | 0 |
| SLC7A5 | 0 | 0.959832268 | 0.522 | 0.109 | 0 |
| RPL12 | 0 | 0.954870183 | 1 | 0.996 | 0 |
| APP | 0 | 0.934912332 | 0.515 | 0.196 | 0 |
| RPS3 | 0 | 0.933890118 | 1 | 0.996 | 0 |
| EIF3E | 0 | 0.929264383 | 0.951 | 0.911 | 0 |
| ARPC1B | 0 | 0.92852849 | 0.531 | 0.218 | 0 |
| RPL35A | 0 | 0.926635559 | 1 | 0.992 | 0 |
| RPS5 | 0 | 0.917666457 | 1 | 0.991 | 0 |
| MT-CO1 | 0 | 0.910015039 | 0.988 | 0.958 | 0 |
| RPLP1 | 0 | 0.907581937 | 1 | 0.999 | 0 |
| NPM1 | 0 | 0.900359763 | 0.99 | 0.957 | 0 |
| RPL37 | 0 | 0.895612447 | 1 | 0.995 | 0 |
| RPL10A | 0 | 0.894123777 | 1 | 0.991 | 0 |
| PDLIM1 | 0 | 0.892139816 | 0.832 | 0.788 | 0 |
| MTRNR2L8 | 0 | 0.887226548 | 0.519 | 0.487 | 0 |
| RPS19 | 0 | 0.884909432 | 1 | 0.997 | 0 |
| JUNB | 0 | 0.884094259 | 0.855 | 0.863 | 0 |
| TP53AIP1 | 0 | 0.88224727 | 0.539 | 0.237 | 0 |
| MT2A | 0 | 0.876672684 | 0.81 | 0.535 | 0 |
| TIMP1 | 0 | 0.87575402 | 0.462 | 0.104 | 0 |
| RPS21 | 0 | 0.873160764 | 0.998 | 0.988 | 0 |
| HNRNPA1 | 0 | 0.872899675 | 0.92 | 0.885 | 0 |
| RPL3 | 0 | 0.869343675 | 1 | 0.996 | 0 |
| PNRC1 | 0 | 0.852584474 | 0.736 | 0.703 | 0 |
| RPS15A | 0 | 0.84947814 | 1 | 0.992 | 0 |
| MT-ND4L | 0 | 0.849387461 | 0.61 | 0.566 | 0 |
| RPL18A | 0 | 0.845159597 | 1 | 0.996 | 0 |

| | | | | | |
|---|---|---|---|---|---|
| BTF3 | 0 | 0.843251407 | 0.996 | 0.977 | 0 |
| RPL11 | 0 | 0.841695911 | 1 | 0.997 | 0 |
| KTN1 | 0 | 0.83970417 | 0.826 | 0.813 | 0 |
| RPL19 | 0 | 0.839507184 | 1 | 0.996 | 0 |
| POLR1D | 0 | 0.839246533 | 0.838 | 0.823 | 0 |
| MT-CO2 | 0 | 0.836466386 | 0.987 | 0.961 | 0 |
| EGR1 | 0 | 0.835157992 | 0.808 | 0.764 | 0 |
| RPL32 | 0 | 0.833150593 | 1 | 0.998 | 0 |

**Supplemental Table 2.** Genes included in the superficial gene signature.

| | p_val | avg_log2FC | pct.1 | pct.2 | p_val_adj |
|---|---|---|---|---|---|
| CRCT1 | 0 | 6.663212 | 0.998 | 0.407 | 0 |
| CNFN | 0 | 5.997242 | 1 | 0.526 | 0 |
| KRT78 | 0 | 5.298534 | 0.994 | 0.27 | 0 |
| SPRR2D | 0 | 5.277972 | 0.848 | 0.364 | 0 |
| LYPD2 | 0 | 5.240213 | 0.992 | 0.242 | 0 |
| FAM25A | 0 | 5.213557 | 0.968 | 0.284 | 0 |
| ECM1 | 0 | 5.109232 | 0.998 | 0.372 | 0 |
| MT1G | 0 | 5.034033 | 0.678 | 0.372 | 0 |
| SPINK7 | 0 | 4.961649 | 0.939 | 0.18 | 0 |
| SPRR2A | 0 | 4.84788 | 0.952 | 0.567 | 0 |
| TMPRSS11 | 0 | 4.762629 | 0.995 | 0.135 | 0 |
| FTH1 | 0 | 4.539188 | 0.999 | 0.952 | 0 |
| MAL | 0 | 4.438548 | 1 | 0.571 | 0 |
| HOPX | 0 | 4.340964 | 0.997 | 0.708 | 0 |
| CRISP3 | 0 | 4.325306 | 0.977 | 0.083 | 0 |
| FLG | 0 | 4.244267 | 0.598 | 0.052 | 0 |
| C15orf48 | 0 | 4.127274 | 0.971 | 0.131 | 0 |
| LCN2 | 0 | 4.051988 | 0.973 | 0.343 | 0 |
| ERO1A | 0 | 4.01261 | 0.981 | 0.314 | 0 |
| SPRR2E | 0 | 3.96558 | 0.749 | 0.288 | 0 |
| CRNN | 0 | 3.951264 | 0.999 | 0.386 | 0 |
| EMP1 | 0 | 3.933291 | 0.999 | 0.692 | 0 |
| PSCA | 0 | 3.913357 | 0.941 | 0.127 | 0 |
| S100P | 0 | 3.784702 | 0.879 | 0.116 | 0 |
| MUC21 | 0 | 3.777149 | 0.994 | 0.282 | 0 |
| ADIRF | 0 | 3.759468 | 0.97 | 0.538 | 0 |
| SPRR3 | 0 | 3.743638 | 1 | 0.859 | 0 |
| PHLDA1 | 0 | 3.551732 | 0.941 | 0.11 | 0 |
| PRSS27 | 0 | 3.295138 | 0.983 | 0.179 | 0 |
| SCEL | 0 | 3.236097 | 0.99 | 0.397 | 0 |
| MXD1 | 0 | 3.197818 | 0.97 | 0.203 | 0 |
| TGM3 | 0 | 3.127514 | 0.988 | 0.308 | 0 |
| CYSRT1 | 0 | 3.101654 | 0.981 | 0.343 | 0 |
| TMPRSS11 | 0 | 3.0563 | 0.976 | 0.197 | 0 |
| C4orf3 | 0 | 3.050302 | 0.993 | 0.75 | 0 |
| PPL | 0 | 2.852725 | 0.987 | 0.475 | 0 |
| MT2A | 0 | 2.842034 | 0.718 | 0.718 | 0 |
| MT1H | 0 | 2.835812 | 0.356 | 0.113 | 0 |
| LMO7 | 0 | 2.834769 | 0.954 | 0.297 | 0 |
| AIF1L | 0 | 2.79036 | 0.942 | 0.117 | 0 |
| SPRR1B | 0 | 2.708646 | 0.809 | 0.324 | 0 |
| CSTB | 0 | 2.708364 | 1 | 0.949 | 0 |
| SAMD9 | 0 | 2.691807 | 0.944 | 0.317 | 0 |
| SLURP2 | 0 | 2.682477 | 0.855 | 0.289 | 0 |
| NCCRP1 | 0 | 2.681409 | 0.976 | 0.211 | 0 |

| | | | | | |
|---|---|---|---|---|---|
| CAMK2N1 | 0 | 2.669605 | 0.91 | 0.027 | 0 |
| TTC9 | 0 | 2.669186 | 0.948 | 0.196 | 0 |
| EPS8L1 | 0 | 2.658087 | 0.956 | 0.113 | 0 |
| SPINK5 | 0 | 2.627968 | 0.999 | 0.692 | 0 |
| PPDPF | 0 | 2.562754 | 0.997 | 0.881 | 0 |
| TIMP2 | 0 | 2.549921 | 0.855 | 0.033 | 0 |
| RNASE7 | 0 | 2.530436 | 0.533 | 0.024 | 0 |
| IL1RN | 0 | 2.528854 | 0.993 | 0.576 | 0 |
| NIBAN2 | 0 | 2.474299 | 0.818 | 0.354 | 0 |
| DUSP5 | 0 | 2.450672 | 0.845 | 0.114 | 0 |
| SAT1 | 0 | 2.424409 | 0.988 | 0.789 | 0 |
| MT1M | 0 | 2.422127 | 0.292 | 0.094 | 0 |
| S100A9 | 0 | 2.421643 | 1 | 0.921 | 0 |
| MYL12A | 0 | 2.318397 | 0.99 | 0.844 | 0 |
| IL18 | 0 | 2.30016 | 0.983 | 0.784 | 0 |
| MUC1 | 0 | 2.283834 | 0.792 | 0.071 | 0 |
| TMOD3 | 0 | 2.271593 | 0.974 | 0.598 | 0 |
| C6orf132 | 0 | 2.250125 | 0.957 | 0.427 | 0 |
| FAM3D | 0 | 2.238508 | 0.9 | 0.195 | 0 |
| ELF3 | 0 | 2.224858 | 0.957 | 0.465 | 0 |
| KRT17 | 0 | 2.208477 | 0.553 | 0.312 | 0 |
| SH3BGRL2 | 0 | 2.205071 | 0.932 | 0.185 | 0 |
| SPNS2 | 0 | 2.190006 | 0.836 | 0.033 | 0 |
| KRT16 | 0 | 2.188358 | 0.754 | 0.32 | 0 |
| A2ML1 | 0 | 2.17596 | 0.973 | 0.392 | 0 |
| ANXA11 | 0 | 2.172568 | 0.972 | 0.567 | 0 |
| MT1E | 0 | 2.162944 | 0.577 | 0.616 | 0 |
| H2AFJ | 0 | 2.10223 | 0.976 | 0.626 | 0 |
| GCHFR | 0 | 2.049766 | 0.862 | 0.369 | 0 |
| SFTA2 | 0 | 2.045811 | 0.877 | 0.094 | 0 |
| HILPDA | 0 | 2.042208 | 0.575 | 0.27 | 0 |
| ADGRF1 | 0 | 1.991916 | 0.82 | 0.122 | 0 |
| ANXA1 | 0 | 1.954987 | 1 | 0.969 | 0 |
| MALL | 0 | 1.943558 | 0.973 | 0.472 | 0 |
| SLURP1 | 0 | 1.937649 | 0.918 | 0.313 | 0 |
| RAB5IF | 0 | 1.929411 | 0.949 | 0.625 | 0 |
| LGALS3 | 0 | 1.929329 | 0.995 | 0.937 | 0 |
| TAX1BP1 | 0 | 1.913211 | 0.967 | 0.576 | 0 |
| CLIC3 | 0 | 1.869309 | 0.959 | 0.415 | 0 |
| S100A8 | 0 | 1.853015 | 1 | 0.885 | 0 |
| PRSS3 | 0 | 1.82896 | 0.862 | 0.093 | 0 |
| CD59 | 0 | 1.819298 | 0.967 | 0.624 | 0 |
| CST6 | 0 | 1.818308 | 0.878 | 0.119 | 0 |
| GLUL | 0 | 1.809864 | 0.676 | 0.435 | 0 |
| HPGD | 0 | 1.807315 | 0.912 | 0.383 | 0 |
| KLK12 | 0 | 1.798359 | 0.635 | 0.175 | 0 |
| DHRS9 | 0 | 1.786528 | 0.711 | 0.018 | 0 |

| | | | | | |
|---|---|---|---|---|---|
| PLAC8 | 0 | 1.77181 | 0.734 | 0.2 | 0 |
| TMEM40 | 0 | 1.741066 | 0.963 | 0.511 | 0 |
| TPM4 | 0 | 1.723422 | 0.969 | 0.664 | 0 |
| SNCG | 0 | 1.687185 | 0.721 | 0.053 | 0 |
| RIOK3 | 0 | 1.652756 | 0.952 | 0.626 | 0 |
| YOD1 | 0 | 1.652293 | 0.693 | 0.078 | 0 |
| SDCBP2 | 0 | 1.650459 | 0.91 | 0.32 | 0 |
| CD24 | 0 | 1.63871 | 0.99 | 0.564 | 0 |

**Supplemental Table 3.** Genes corresponding to co-expressed modules along the EoE epithelial pseudotemporal trajectory.

| | gene | module | supermodule | dim_1 | dim_2 | status | p_value | morans_test_statistic | morans_I | q_value |
|---|---|---|---|---|---|---|---|---|---|---|
| 1 | AC020916. | 1 | 1 | 1.63928 | -2.4881 | OK | 0 | 251.3780328 | 0.254997 | 0 |
| 2 | AC036176. | 1 | 1 | 1.684324 | -2.50395 | OK | 0 | 126.1473819 | 0.127956 | 0 |
| 3 | ADH1C | 1 | 1 | 0.319839 | -3.03391 | OK | 0 | 106.8289895 | 0.108348 | 0 |
| 4 | ADM | 1 | 1 | 0.86305 | -3.03058 | OK | 0 | 143.3322682 | 0.14539 | 0 |
| 5 | APCDD1 | 1 | 1 | 0.538908 | -2.4998 | OK | 0 | 158.4539216 | 0.160729 | 0 |
| 6 | APOBEC3A | 1 | 1 | 2.50802 | -2.59125 | OK | 0 | 144.5728216 | 0.146645 | 0 |
| 7 | AREG | 1 | 1 | 2.392676 | -2.62937 | OK | 0 | 257.2458014 | 0.260948 | 0 |
| 8 | ASS1 | 1 | 1 | 0.560633 | -2.9207 | OK | 0 | 346.705388 | 0.351697 | 0 |
| 9 | BASP1 | 1 | 1 | 0.282518 | -3.29634 | OK | 0 | 237.0933783 | 0.240497 | 0 |
| 10 | BCAM | 1 | 1 | 0.46836 | -3.36526 | OK | 0 | 257.5000892 | 0.261195 | 0 |
| 11 | BHLHE40 | 1 | 1 | 2.382114 | -2.64325 | OK | 0 | 276.6513126 | 0.280635 | 0 |
| 12 | BMP2 | 1 | 1 | 0.852944 | -2.86915 | OK | 0 | 105.7401336 | 0.107248 | 0 |
| 13 | BST2 | 1 | 1 | -0.69009 | -2.77002 | OK | 0 | 152.2795412 | 0.154464 | 0 |
| 14 | BTG2 | 1 | 1 | 1.574787 | -2.56325 | OK | 0 | 277.813387 | 0.281814 | 0 |
| 15 | C1R | 1 | 1 | -0.58559 | -2.93039 | OK | 0 | 251.4742279 | 0.255083 | 0 |
| 16 | CALML3 | 1 | 1 | 1.173151 | -1.42515 | OK | 0 | 361.5015277 | 0.366709 | 0 |
| 17 | CASC19 | 1 | 1 | 0.111581 | -2.18274 | OK | 0 | 142.600573 | 0.144643 | 0 |
| 18 | CAV1 | 1 | 1 | 0.098602 | -2.92218 | OK | 0 | 514.6421679 | 0.522061 | 0 |
| 19 | CAVIN2 | 1 | 1 | 0.909655 | -2.9257 | OK | 0 | 119.7009119 | 0.121415 | 0 |
| 20 | CCDC80 | 1 | 1 | 0.513994 | -2.90377 | OK | 0 | 139.6095003 | 0.141599 | 0 |
| 21 | CCL20 | 1 | 1 | 1.002225 | -2.64332 | OK | 9.37E-292 | 36.48564812 | 0.036995 | 9.48E-292 |
| 22 | CCL4 | 1 | 1 | 1.999882 | -4.15636 | OK | 3.79E-152 | 26.2477909 | 0.026607 | 3.81E-152 |
| 23 | CCN1 | 1 | 1 | 0.771892 | -2.91044 | OK | 0 | 197.8381453 | 0.200674 | 0 |
| 24 | CCND1 | 1 | 1 | 0.636074 | -1.77022 | OK | 0 | 362.9119658 | 0.368142 | 0 |
| 25 | CCND2 | 1 | 1 | 0.367254 | -2.54339 | OK | 0 | 234.2296091 | 0.237594 | 0 |
| 26 | CD55 | 1 | 1 | 2.356376 | -2.35135 | OK | 0 | 370.4750918 | 0.375815 | 0 |
| 27 | CD74 | 1 | 1 | -0.65505 | -2.78553 | OK | 0 | 191.5775062 | 0.194331 | 0 |
| 28 | CDKN1A | 1 | 1 | 2.458907 | -2.55973 | OK | 0 | 264.6479424 | 0.268459 | 0 |
| 29 | CDKN2B | 1 | 1 | 2.322522 | -2.24507 | OK | 0 | 315.3491555 | 0.31989 | 0 |
| 30 | CFB | 1 | 1 | -0.67255 | -2.75234 | OK | 0 | 142.8735756 | 0.144923 | 0 |
| 31 | CH25H | 1 | 1 | -0.30336 | -2.86365 | OK | 0 | 243.1075023 | 0.246604 | 0 |
| 32 | CLDN4 | 1 | 1 | 2.610722 | -2.50706 | OK | 0 | 486.9148983 | 0.493936 | 0 |
| 33 | CLEC2B | 1 | 1 | 1.817906 | -4.08178 | OK | 0 | 134.2228228 | 0.136149 | 0 |
| 34 | CLU | 1 | 1 | -0.6151 | -2.91825 | OK | 0 | 115.1698537 | 0.116815 | 0 |
| 35 | COL17A1 | 1 | 1 | 0.254002 | -2.50034 | OK | 0 | 547.7495959 | 0.555644 | 0 |
| 36 | CRABP2 | 1 | 1 | 1.8071 | -1.92574 | OK | 0 | 365.6871792 | 0.370953 | 0 |
| 37 | CRIP1 | 1 | 1 | 1.011198 | -1.6993 | OK | 0 | 152.03995 | 0.154224 | 0 |
| 38 | CRIP2 | 1 | 1 | 1.081226 | -1.48333 | OK | 0 | 139.583941 | 0.141587 | 0 |
| 39 | CTNNAL1 | 1 | 1 | 0.117453 | -3.21299 | OK | 0 | 523.8488433 | 0.531391 | 0 |
| 40 | CTSG | 1 | 1 | 1.930303 | -4.23015 | OK | 0 | 69.62924232 | 0.070602 | 0 |
| 41 | CXCL14 | 1 | 1 | 1.252692 | -1.56485 | OK | 0 | 277.1710439 | 0.281144 | 0 |
| 42 | CXCL2 | 1 | 1 | 1.307683 | -1.7498 | OK | 2.79E-184 | 28.92629484 | 0.029325 | 2.81E-184 |
| 43 | CYP2E1 | 1 | 1 | -0.54738 | -2.79112 | OK | 0 | 67.68390842 | 0.068648 | 0 |
| 44 | CYP3A5 | 1 | 1 | 1.863549 | -2.0873 | OK | 0 | 103.0310223 | 0.104505 | 0 |
| 45 | DLK2 | 1 | 1 | 0.40801 | -3.39077 | OK | 0 | 412.9989488 | 0.418935 | 0 |
| 46 | DST | 1 | 1 | 0.461138 | -2.53634 | OK | 0 | 568.3045922 | 0.5765 | 0 |
| 47 | DUSP2 | 1 | 1 | 1.954673 | -2.62599 | OK | 0 | 51.81895849 | 0.052553 | 0 |
| 48 | EDN1 | 1 | 1 | 0.873766 | -3.03193 | OK | 0 | 127.9897084 | 0.129815 | 0 |
| 49 | EFNB2 | 1 | 1 | 2.486477 | -2.50172 | OK | 0 | 128.9859157 | 0.130835 | 0 |
| 50 | EGR3 | 1 | 1 | 1.452323 | -2.57406 | OK | 0 | 84.38707069 | 0.085588 | 0 |
| 51 | EIF1AY | 1 | 1 | -0.08711 | -2.36609 | OK | 0 | 70.75675467 | 0.071765 | 0 |
| 52 | EMP3 | 1 | 1 | -0.20937 | -3.07937 | OK | 0 | 256.9533955 | 0.260639 | 0 |
| 53 | EPHA2 | 1 | 1 | 2.656547 | -2.42386 | OK | 0 | 246.0033154 | 0.249544 | 0 |
| 54 | FABP5 | 1 | 1 | 0.751235 | -2.02107 | OK | 0 | 502.6097048 | 0.509848 | 0 |
| 55 | FKBP2 | 1 | 1 | 0.476846 | -3.51657 | OK | 0 | 53.25941835 | 0.054015 | 0 |
| 56 | FLNA | 1 | 1 | -0.13304 | -2.09958 | OK | 0 | 231.8223223 | 0.235158 | 0 |
| 57 | FST | 1 | 1 | 0.414913 | -3.38829 | OK | 0 | 125.6322119 | 0.12742 | 0 |
| 58 | FXYD5 | 1 | 1 | 0.292254 | -3.39987 | OK | 0 | 96.55240375 | 0.097933 | 0 |

| | | | | | | | | | | |
|---|---|---|---|---|---|---|---|---|---|---|
| 59 | G0S2 | 1 | 1 | -0.15712 | -2.87777 | OK | 0 | 140.1699029 | 0.142156 | 0 |
| 60 | GADD45A | 1 | 1 | 1.241321 | -2.99126 | OK | 0 | 196.1514741 | 0.198973 | 0 |
| 61 | GADD45B | 1 | 1 | 2.258333 | -1.48187 | OK | 0 | 105.6110451 | 0.107124 | 0 |
| 62 | GBP1 | 1 | 1 | 1.115467 | -2.94459 | OK | 0 | 142.4951207 | 0.144539 | 0 |
| 63 | GBP4 | 1 | 1 | -0.71441 | -2.42714 | OK | 0 | 95.03652978 | 0.096395 | 0 |
| 64 | GCLC | 1 | 1 | 0.342775 | -2.37627 | OK | 0 | 203.325739 | 0.206249 | 0 |
| 65 | GEM | 1 | 1 | 1.291327 | -1.65124 | OK | 0 | 116.9290194 | 0.118554 | 0 |
| 66 | GLDC | 1 | 1 | -0.27651 | -2.63336 | OK | 0 | 157.210424 | 0.159465 | 0 |
| 67 | GLUL | 1 | 1 | 0.434214 | -2.75789 | OK | 0 | 159.2299888 | 0.161518 | 0 |
| 68 | HBA2 | 1 | 1 | 2.493382 | -4.0692 | OK | 0.000385 | 3.363480976 | 0.003394 | 0.000385 |
| 69 | HBB | 1 | 1 | 2.377574 | -3.73461 | OK | 5.27E-50 | 14.82207236 | 0.015012 | 5.30E-50 |
| 70 | HBEGF | 1 | 1 | 2.164289 | -2.67934 | OK | 0 | 153.9177792 | 0.156129 | 0 |
| 71 | HES1 | 1 | 1 | 1.357425 | -2.38272 | OK | 0 | 201.3878748 | 0.204284 | 0 |
| 72 | HLA-DRA | 1 | 1 | -0.56867 | -2.83555 | OK | 9.69E-265 | 34.7403494 | 0.035227 | 9.80E-265 |
| 73 | ID1 | 1 | 1 | 2.581724 | -2.61791 | OK | 0 | 242.6811155 | 0.246175 | 0 |
| 74 | ID3 | 1 | 1 | 2.521021 | -2.49203 | OK | 0 | 241.5798604 | 0.245057 | 0 |
| 75 | ID4 | 1 | 1 | 1.217844 | -1.42026 | OK | 0 | 202.8877829 | 0.205806 | 0 |
| 76 | IFI27 | 1 | 1 | -0.67546 | -2.20074 | OK | 0 | 181.7270298 | 0.18434 | 0 |
| 77 | IFIT3 | 1 | 1 | -0.7039 | -2.72902 | OK | 0 | 74.26490012 | 0.075323 | 0 |
| 78 | IFITM1 | 1 | 1 | -0.57002 | -2.75697 | OK | 0 | 187.9036526 | 0.190596 | 0 |
| 79 | IFITM2 | 1 | 1 | 1.011108 | -1.78286 | OK | 0 | 74.91433667 | 0.075982 | 0 |
| 80 | IGFBP3 | 1 | 1 | 0.371005 | -2.70024 | OK | 0 | 306.9375209 | 0.31135 | 0 |
| 81 | IGFBP5 | 1 | 1 | 0.588682 | -2.71975 | OK | 0 | 160.1798855 | 0.162479 | 0 |
| 82 | IGFBP6 | 1 | 1 | 0.787968 | -2.10519 | OK | 0 | 262.7523086 | 0.266529 | 0 |
| 83 | IGFBP7 | 1 | 1 | -0.34624 | -2.96818 | OK | 0 | 352.2977457 | 0.357359 | 0 |
| 84 | IGHA1 | 1 | 1 | 2.557645 | -4.25112 | OK | 1.41E-126 | 23.90349071 | 0.024215 | 1.42E-126 |
| 85 | IGKC | 1 | 1 | 2.54377 | -4.25131 | OK | 2.44E-46 | 14.24399628 | 0.01443 | 2.45E-46 |
| 86 | IGLC1 | 1 | 1 | 2.556192 | -4.24673 | OK | 2.08E-45 | 14.09346196 | 0.014242 | 2.09E-45 |
| 87 | IGLC2 | 1 | 1 | 2.554935 | -4.25502 | OK | 5.26E-49 | 14.66684552 | 0.014848 | 5.28E-49 |
| 88 | IGLC3 | 1 | 1 | 2.555681 | -4.23277 | OK | 1.59E-16 | 8.165986867 | 0.008247 | 1.60E-16 |
| 89 | IL18 | 1 | 1 | 0.52498 | -3.26359 | OK | 0 | 296.1337585 | 0.300399 | 0 |
| 90 | IL1R2 | 1 | 1 | -0.23434 | -3.11895 | OK | 0 | 545.0326173 | 0.55288 | 0 |
| 91 | IRF1 | 1 | 1 | -0.60471 | -2.69317 | OK | 0 | 170.8912911 | 0.173347 | 0 |
| 92 | ISG15 | 1 | 1 | -0.66291 | -2.14224 | OK | 0 | 53.05617638 | 0.053809 | 0 |
| 93 | ITGA6 | 1 | 1 | 0.254387 | -2.40469 | OK | 0 | 303.19091 | 0.307557 | 0 |
| 94 | ITM2A | 1 | 1 | 1.483514 | -1.33433 | OK | 5.07E-44 | 13.86630512 | 0.01405 | 5.08E-44 |
| 95 | ITPKC | 1 | 1 | 2.68213 | -2.47037 | OK | 0 | 229.6816513 | 0.232987 | 0 |
| 96 | JCHAIN | 1 | 1 | 2.560954 | -4.2492 | OK | 3.61E-250 | 33.76159979 | 0.03417 | 3.65E-250 |
| 97 | KCNQ1OT1 | 1 | 1 | 1.962834 | -2.23346 | OK | 0 | 90.04412084 | 0.091329 | 0 |
| 98 | KLF2 | 1 | 1 | 2.378366 | -1.98028 | OK | 0 | 126.2714861 | 0.128081 | 0 |
| 99 | KLF6 | 1 | 1 | 1.352223 | -2.84309 | OK | 0 | 265.1055937 | 0.268921 | 0 |
| 100 | KLK5 | 1 | 1 | 0.547774 | -3.33708 | OK | 0 | 118.4480083 | 0.120138 | 0 |
| 101 | KRT14 | 1 | 1 | 1.203802 | -1.9339 | OK | 0 | 517.0799878 | 0.524537 | 0 |
| 102 | KRT17 | 1 | 1 | 0.385406 | -2.94431 | OK | 0 | 232.3568361 | 0.235701 | 0 |
| 103 | KRT31 | 1 | 1 | 0.948314 | -1.83795 | OK | 0 | 56.15630649 | 0.056952 | 0 |
| 104 | KRT8 | 1 | 1 | 1.223441 | -1.86004 | OK | 0 | 223.0145213 | 0.226224 | 0 |
| 105 | LAMB3 | 1 | 1 | 0.040066 | -2.46938 | OK | 0 | 415.3611254 | 0.421347 | 0 |
| 106 | LDLR | 1 | 1 | 2.208782 | -2.64574 | OK | 0 | 155.3873989 | 0.157619 | 0 |
| 107 | LGALS1 | 1 | 1 | 1.380708 | -1.75596 | OK | 2.14E-224 | 31.95732083 | 0.032402 | 2.16E-224 |
| 108 | LGALS7 | 1 | 1 | 1.090735 | -2.06942 | OK | 0 | 126.3137809 | 0.128127 | 0 |
| 109 | LGALSL | 1 | 1 | 2.31963 | -2.18426 | OK | 0 | 335.035367 | 0.339862 | 0 |
| 110 | LOXL4 | 1 | 1 | -0.19665 | -2.7509 | OK | 0 | 248.8545188 | 0.252411 | 0 |
| 111 | LSP1 | 1 | 1 | 1.118721 | -1.63978 | OK | 8.94E-25 | 10.21000348 | 0.010341 | 8.96E-25 |
| 112 | LTF | 1 | 1 | -0.5884 | -2.76523 | OK | 0 | 87.20869164 | 0.088434 | 0 |
| 113 | MAFF | 1 | 1 | 2.511036 | -2.52687 | OK | 0 | 211.3821363 | 0.214423 | 0 |
| 114 | MALAT1 | 1 | 1 | 1.5667 | -2.22478 | OK | 0 | 274.782913 | 0.278733 | 0 |
| 115 | MAP3K8 | 1 | 1 | 2.390412 | -2.41375 | OK | 0 | 135.6530637 | 0.137599 | 0 |
| 116 | MIDN | 1 | 1 | 2.452559 | -2.54166 | OK | 0 | 233.2363591 | 0.236593 | 0 |
| 117 | MT-ND4L | 1 | 1 | 1.018874 | -2.33358 | OK | 0 | 173.4250491 | 0.175918 | 0 |
| 118 | MTRNR2L8 | 1 | 1 | 0.833867 | -2.22614 | OK | 0 | 48.66179031 | 0.049352 | 0 |

| | | | | | | | | | | |
|---|---|---|---|---|---|---|---|---|---|---|
| 119 | MYADM | 1 | 1 | 1.113583 | -2.80532 | OK | 0 | 144.0347554 | 0.146098 | 0 |
| 120 | MYO1B | 1 | 1 | 0.028748 | -2.44418 | OK | 0 | 138.4852347 | 0.14047 | 0 |
| 121 | NCOA7 | 1 | 1 | 1.155427 | -2.83456 | OK | 0 | 337.5638755 | 0.342428 | 0 |
| 122 | NEAT1 | 1 | 1 | 1.836944 | -2.38097 | OK | 0 | 267.4669646 | 0.271314 | 0 |
| 123 | NEDD9 | 1 | 1 | 0.333455 | -2.76122 | OK | 0 | 193.8402198 | 0.196605 | 0 |
| 124 | NIBAN1 | 1 | 1 | -0.13468 | -2.07621 | OK | 0 | 123.085989 | 0.12485 | 0 |
| 125 | NPPC | 1 | 1 | 1.228183 | -1.53848 | OK | 0 | 114.497038 | 0.116054 | 0 |
| 126 | NR4A1 | 1 | 1 | 1.543684 | -2.59048 | OK | 0 | 169.8232228 | 0.172264 | 0 |
| 127 | NR4A2 | 1 | 1 | 1.544583 | -2.54334 | OK | 4.35E-307 | 37.44012295 | 0.037967 | 4.41E-307 |
| 128 | NRARP | 1 | 1 | 2.601736 | -2.46734 | OK | 0 | 394.009364 | 0.399689 | 0 |
| 129 | NSUN6 | 1 | 1 | 1.329912 | -2.37738 | OK | 0 | 49.45029147 | 0.05015 | 0 |
| 130 | OVOL1 | 1 | 1 | 2.476076 | -2.59022 | OK | 0 | 188.088242 | 0.190793 | 0 |
| 131 | PDPN | 1 | 1 | -0.19634 | -2.9963 | OK | 0 | 311.1397947 | 0.315608 | 0 |
| 132 | PEG3 | 1 | 1 | 0.799547 | -2.36057 | OK | 9.24E-221 | 31.69454113 | 0.032131 | 9.32E-221 |
| 133 | PER3 | 1 | 1 | 0.675756 | -1.9635 | OK | 0 | 132.3432152 | 0.134242 | 0 |
| 134 | PHGDH | 1 | 1 | 0.522621 | -3.54771 | OK | 0 | 233.1862418 | 0.236539 | 0 |
| 135 | PHLDA2 | 1 | 1 | 2.583079 | -2.49378 | OK | 0 | 345.8296751 | 0.350813 | 0 |
| 136 | PLAUR | 1 | 1 | 2.397017 | -2.61751 | OK | 0 | 90.32190452 | 0.091612 | 0 |
| 137 | PLK2 | 1 | 1 | 1.187502 | -2.94141 | OK | 0 | 172.1827367 | 0.174658 | 0 |
| 138 | PLK3 | 1 | 1 | 1.848076 | -2.37551 | OK | 0 | 89.0196517 | 0.090292 | 0 |
| 139 | POSTN | 1 | 1 | -0.36877 | -2.50106 | OK | 0 | 232.6168156 | 0.235962 | 0 |
| 140 | PRRX1 | 1 | 1 | -0.34054 | -2.66218 | OK | 0 | 223.6743965 | 0.226881 | 0 |
| 141 | PRSS23 | 1 | 1 | -0.11074 | -2.75069 | OK | 0 | 184.9629801 | 0.187603 | 0 |
| 142 | RAB11FIP1 | 1 | 1 | 2.403471 | -2.33391 | OK | 0 | 462.1266184 | 0.468791 | 0 |
| 143 | RAB31 | 1 | 1 | -0.02532 | -2.50432 | OK | 0 | 94.22632269 | 0.095572 | 0 |
| 144 | RAMP1 | 1 | 1 | 0.188345 | -3.31549 | OK | 0 | 317.2659924 | 0.32182 | 0 |
| 145 | RHOB | 1 | 1 | 2.242625 | -1.4497 | OK | 0 | 202.4356987 | 0.205347 | 0 |
| 146 | RPS4Y1 | 1 | 1 | -0.05598 | -2.47687 | OK | 0 | 186.392789 | 0.189073 | 0 |
| 147 | RRAD | 1 | 1 | 2.504522 | -2.52419 | OK | 0 | 279.0862894 | 0.283105 | 0 |
| 148 | S100A9 | 1 | 1 | -0.45281 | -2.63434 | OK | 0 | 389.1506354 | 0.394746 | 0 |
| 149 | SAA1 | 1 | 1 | -0.54209 | -2.95333 | OK | 0 | 186.5676271 | 0.189238 | 0 |
| 150 | SERPINB2 | 1 | 1 | 2.394515 | -2.29081 | OK | 0 | 359.4517273 | 0.364632 | 0 |
| 151 | SLC1A3 | 1 | 1 | 0.518801 | -2.61277 | OK | 0 | 347.6272537 | 0.352625 | 0 |
| 152 | SLC25A25 | 1 | 1 | 2.357609 | -2.66504 | OK | 0 | 123.5270162 | 0.125297 | 0 |
| 153 | SLC7A2 | 1 | 1 | 0.472764 | -2.38052 | OK | 0 | 111.0431031 | 0.112627 | 0 |
| 154 | SLC7A5 | 1 | 1 | 0.554562 | -2.59132 | OK | 0 | 357.6538348 | 0.3628 | 0 |
| 155 | SNHG17 | 1 | 1 | 0.759469 | -2.21123 | OK | 0 | 74.24134721 | 0.0753 | 0 |
| 156 | SOCS3 | 1 | 1 | -0.33121 | -2.88357 | OK | 0 | 140.3157356 | 0.142329 | 0 |
| 157 | SOD2 | 1 | 1 | -0.59301 | -2.74724 | OK | 0 | 188.8917606 | 0.191608 | 0 |
| 158 | SOX4 | 1 | 1 | 0.829218 | -2.55251 | OK | 0 | 98.85032033 | 0.100266 | 0 |
| 159 | SOX9 | 1 | 1 | 2.555352 | -2.57899 | OK | 0 | 98.2141115 | 0.099619 | 0 |
| 160 | SRGN | 1 | 1 | 1.931935 | -4.22501 | OK | 0 | 48.95780565 | 0.049641 | 0 |
| 161 | SYNPO2 | 1 | 1 | -0.11312 | -2.64356 | OK | 0 | 250.203614 | 0.253788 | 0 |
| 162 | SYT8 | 1 | 1 | 1.02998 | -1.7182 | OK | 0 | 81.57041209 | 0.082734 | 0 |
| 163 | TIMP1 | 1 | 1 | -0.36575 | -2.62806 | OK | 0 | 347.6863231 | 0.352696 | 0 |
| 164 | TINAGL1 | 1 | 1 | -0.21167 | -2.83272 | OK | 0 | 168.6046847 | 0.171013 | 0 |
| 165 | TNC | 1 | 1 | -0.15141 | -2.7058 | OK | 0 | 284.1738482 | 0.288246 | 0 |
| 166 | TNFAIP2 | 1 | 1 | -0.65859 | -2.53392 | OK | 0 | 197.2090193 | 0.200043 | 0 |
| 167 | TNFAIP3 | 1 | 1 | 1.433982 | -2.69384 | OK | 0 | 50.76613644 | 0.051486 | 0 |
| 168 | TNFRSF12A | 1 | 1 | 0.076905 | -2.818 | OK | 0 | 126.3962489 | 0.128204 | 0 |
| 169 | TPM2 | 1 | 1 | 0.403067 | -3.44387 | OK | 0 | 192.4588265 | 0.195213 | 0 |
| 170 | TPPP3 | 1 | 1 | 1.007442 | -1.70061 | OK | 0 | 89.45077119 | 0.090727 | 0 |
| 171 | TPSAB1 | 1 | 1 | 1.956884 | -4.24606 | OK | 0 | 105.468717 | 0.106962 | 0 |
| 172 | TPSB2 | 1 | 1 | 1.921124 | -4.24201 | OK | 0 | 113.88851 | 0.115504 | 0 |
| 173 | TRIB1 | 1 | 1 | 1.4767 | -2.83827 | OK | 0 | 77.35067692 | 0.078455 | 0 |
| 174 | TSLP | 1 | 1 | 0.266599 | -3.33594 | OK | 0 | 368.0925964 | 0.37339 | 0 |
| 175 | TUBB2A | 1 | 1 | 2.517156 | -2.00854 | OK | 0 | 217.7546552 | 0.220887 | 0 |
| 176 | TXNIP | 1 | 1 | 0.08213 | -2.86649 | OK | 0 | 247.3527351 | 0.250913 | 0 |
| 177 | UGCG | 1 | 1 | -0.69959 | -2.45222 | OK | 0 | 134.2644889 | 0.136191 | 0 |
| 178 | VIM | 1 | 1 | 1.89763 | -4.18436 | OK | 8.36E-207 | 30.66491519 | 0.031091 | 8.43E-207 |

| | | | | | | | | | | |
|---|---|---|---|---|---|---|---|---|---|---|
| 179 | WNT10A | 1 | 1 | 0.457502 | -3.26788 | OK | 0 | 389.7178211 | 0.395321 | 0 |
| 180 | XIST | 1 | 1 | 1.056281 | -2.09943 | OK | 0 | 71.0457907 | 0.072059 | 0 |
| 181 | ZNF750 | 1 | 1 | 2.368178 | -2.42324 | OK | 0 | 440.6977967 | 0.447052 | 0 |
| 182 | ASF1B | 2 | 2 | -4.33988 | -6.22398 | OK | 0 | 303.6197256 | 0.30797 | 0 |
| 183 | ATAD2 | 2 | 2 | -4.25714 | -5.88279 | OK | 0 | 332.4975271 | 0.337284 | 0 |
| 184 | ATAD5 | 2 | 2 | -4.15981 | -5.81475 | OK | 0 | 294.7481882 | 0.298979 | 0 |
| 185 | BRCA1 | 2 | 2 | -4.02936 | -5.89745 | OK | 0 | 324.5089025 | 0.329168 | 0 |
| 186 | BRCA2 | 2 | 2 | -4.24062 | -5.72768 | OK | 0 | 307.555001 | 0.311969 | 0 |
| 187 | CARHSP1 | 2 | 2 | -4.48334 | -6.19392 | OK | 0 | 270.5745973 | 0.274471 | 0 |
| 188 | CCDC34 | 2 | 2 | -5.14806 | -6.52291 | OK | 0 | 265.4406665 | 0.269262 | 0 |
| 189 | CDCA5 | 2 | 2 | -4.89061 | -5.89367 | OK | 0 | 322.045084 | 0.326657 | 0 |
| 190 | CDCA7 | 2 | 2 | -3.51188 | -6.09523 | OK | 0 | 291.3183535 | 0.295502 | 0 |
| 191 | CDT1 | 2 | 2 | -3.49152 | -6.16998 | OK | 0 | 302.2505479 | 0.306593 | 0 |
| 192 | CENPK | 2 | 2 | -4.80545 | -6.44827 | OK | 0 | 367.7205942 | 0.373007 | 0 |
| 193 | CENPM | 2 | 2 | -4.80206 | -6.44878 | OK | 0 | 383.4095938 | 0.388922 | 0 |
| 194 | CENPU | 2 | 2 | -4.12404 | -6.17607 | OK | 0 | 248.5664698 | 0.252142 | 0 |
| 195 | CEP78 | 2 | 2 | -4.31891 | -5.68348 | OK | 0 | 183.1694548 | 0.185795 | 0 |
| 196 | CHEK1 | 2 | 2 | -3.69022 | -6.05 | OK | 0 | 290.5412327 | 0.294715 | 0 |
| 197 | CLSPN | 2 | 2 | -4.09267 | -6.09342 | OK | 0 | 389.6023096 | 0.395193 | 0 |
| 198 | DEK | 2 | 2 | -3.67038 | -6.02709 | OK | 0 | 361.7305588 | 0.366943 | 0 |
| 199 | DHFR | 2 | 2 | -4.31338 | -6.14952 | OK | 0 | 336.6264085 | 0.341468 | 0 |
| 200 | DNMT1 | 2 | 2 | -3.74354 | -5.96857 | OK | 0 | 320.6282158 | 0.325245 | 0 |
| 201 | DUT | 2 | 2 | -3.7669 | -6.1359 | OK | 0 | 312.5193279 | 0.317021 | 0 |
| 202 | E2F8 | 2 | 2 | -4.82741 | -5.79589 | OK | 0 | 224.0968609 | 0.227316 | 0 |
| 203 | ESCO2 | 2 | 2 | -4.90131 | -5.85412 | OK | 0 | 417.5632557 | 0.42355 | 0 |
| 204 | EZH2 | 2 | 2 | -4.68819 | -5.78957 | OK | 0 | 212.581927 | 0.215637 | 0 |
| 205 | FAM111B | 2 | 2 | -4.00952 | -6.04591 | OK | 0 | 420.2372258 | 0.426279 | 0 |
| 206 | FANCI | 2 | 2 | -4.56759 | -5.80454 | OK | 0 | 306.0426272 | 0.310437 | 0 |
| 207 | FBXO5 | 2 | 2 | -4.94313 | -5.8549 | OK | 0 | 294.1621827 | 0.298379 | 0 |
| 208 | FEN1 | 2 | 2 | -3.99097 | -6.19048 | OK | 0 | 301.9577638 | 0.306297 | 0 |
| 209 | GINS2 | 2 | 2 | -3.50142 | -6.13664 | OK | 0 | 349.9647539 | 0.354994 | 0 |
| 210 | GMNN | 2 | 2 | -3.65876 | -6.06001 | OK | 0 | 260.1691145 | 0.263911 | 0 |
| 211 | HELLS | 2 | 2 | -3.72866 | -6.02014 | OK | 0 | 414.2059156 | 0.420169 | 0 |
| 212 | HES4 | 2 | 2 | -4.15843 | -6.23743 | OK | 0 | 260.8087447 | 0.264563 | 0 |
| 213 | HIST1H1A | 2 | 2 | 0.114293 | -8.62427 | OK | 0 | 371.44293 | 0.376778 | 0 |
| 214 | HIST1H1B | 2 | 2 | 0.127777 | -8.60943 | OK | 0 | 447.9944124 | 0.454436 | 0 |
| 215 | HIST1H1C | 2 | 2 | 0.106376 | -8.63327 | OK | 0 | 179.4587549 | 0.182038 | 0 |
| 216 | HIST1H1D | 2 | 2 | 0.135122 | -8.66808 | OK | 0 | 341.452203 | 0.346366 | 0 |
| 217 | HIST1H1E | 2 | 2 | 0.110582 | -8.62709 | OK | 0 | 229.9497091 | 0.233257 | 0 |
| 218 | HIST1H2AC | 2 | 2 | 0.18788 | -8.51691 | OK | 0 | 177.2708799 | 0.179818 | 0 |
| 219 | HIST1H2AG | 2 | 2 | 0.108157 | -8.63215 | OK | 0 | 308.5419311 | 0.312968 | 0 |
| 220 | HIST1H2BC | 2 | 2 | 0.111142 | -8.62646 | OK | 0 | 183.31419 | 0.185941 | 0 |
| 221 | HIST1H3B | 2 | 2 | 0.123162 | -8.60103 | OK | 0 | 388.4850417 | 0.394043 | 0 |
| 222 | HIST1H3D | 2 | 2 | 0.096466 | -8.64129 | OK | 0 | 281.4909056 | 0.285528 | 0 |
| 223 | HIST1H3G | 2 | 2 | 0.083851 | -8.65213 | OK | 0 | 343.3052112 | 0.348208 | 0 |
| 224 | HIST1H4C | 2 | 2 | 0.077953 | -8.66427 | OK | 0 | 330.5816847 | 0.335344 | 0 |
| 225 | HIST2H2AC | 2 | 2 | 0.103011 | -8.65129 | OK | 0 | 184.0538168 | 0.186697 | 0 |
| 226 | HSPH1 | 2 | 2 | -3.67822 | -5.32866 | OK | 0 | 186.3407804 | 0.18902 | 0 |
| 227 | LIG1 | 2 | 2 | -4.11585 | -6.01801 | OK | 0 | 237.5323514 | 0.240942 | 0 |
| 228 | LMNB1 | 2 | 2 | -4.49294 | -5.56294 | OK | 0 | 393.3574489 | 0.399018 | 0 |
| 229 | MCM3 | 2 | 2 | -3.46501 | -6.07193 | OK | 0 | 309.7711593 | 0.31423 | 0 |
| 230 | MCM4 | 2 | 2 | -3.66856 | -5.67707 | OK | 0 | 240.3590917 | 0.243817 | 0 |
| 231 | MCM5 | 2 | 2 | -3.44434 | -6.18908 | OK | 0 | 287.4815364 | 0.291616 | 0 |
| 232 | MCM6 | 2 | 2 | -3.47477 | -6.0325 | OK | 0 | 280.9989135 | 0.285035 | 0 |
| 233 | MCM7 | 2 | 2 | -3.52583 | -6.16576 | OK | 0 | 354.8152696 | 0.359923 | 0 |
| 234 | MELK | 2 | 2 | -5.14283 | -5.78189 | OK | 0 | 259.7973015 | 0.263523 | 0 |
| 235 | MND1 | 2 | 2 | -4.77889 | -6.44416 | OK | 0 | 321.7540889 | 0.326367 | 0 |
| 236 | MNS1 | 2 | 2 | -4.75841 | -6.36889 | OK | 0 | 303.8454906 | 0.308196 | 0 |
| 237 | MSH6 | 2 | 2 | -3.55908 | -5.85192 | OK | 0 | 223.4767841 | 0.226689 | 0 |
| 238 | MYBL2 | 2 | 2 | -4.73078 | -6.08976 | OK | 0 | 330.5514179 | 0.335289 | 0 |

| | | | | | | | | | | |
|---|---|---|---|---|---|---|---|---|---|---|
| 239 | NASP | 2 | 2 | -3.6002 | -6.05167 | OK | 0 | 322.7272722 | 0.327377 | 0 |
| 240 | NCAPD3 | 2 | 2 | -4.55137 | -5.6881 | OK | 0 | 246.7831849 | 0.250327 | 0 |
| 241 | NCAPG2 | 2 | 2 | -4.6771 | -5.68585 | OK | 0 | 256.4721136 | 0.260148 | 0 |
| 242 | ORC6 | 2 | 2 | -4.09723 | -6.13835 | OK | 0 | 363.8643327 | 0.369094 | 0 |
| 243 | PCLAF | 2 | 2 | -4.9571 | -6.52778 | OK | 0 | 448.9176111 | 0.455385 | 0 |
| 244 | PCNA | 2 | 2 | -3.6325 | -6.16611 | OK | 0 | 320.7812437 | 0.325402 | 0 |
| 245 | PKMYT1 | 2 | 2 | -4.58171 | -6.20377 | OK | 0 | 315.0700891 | 0.319584 | 0 |
| 246 | PPIF | 2 | 2 | -3.48611 | -6.15633 | OK | 0 | 120.3496948 | 0.122075 | 0 |
| 247 | PSIP1 | 2 | 2 | -3.75722 | -6.01724 | OK | 0 | 316.7354083 | 0.321295 | 0 |
| 248 | RAD51AP1 | 2 | 2 | -4.81258 | -6.3274 | OK | 0 | 345.7551801 | 0.350715 | 0 |
| 249 | RECQL | 2 | 2 | -3.74187 | -5.67753 | OK | 0 | 128.7470146 | 0.130593 | 0 |
| 250 | RFC3 | 2 | 2 | -3.86256 | -5.99625 | OK | 0 | 202.5201102 | 0.205419 | 0 |
| 251 | RIF1 | 2 | 2 | -3.734 | -5.47939 | OK | 0 | 203.6023775 | 0.20653 | 0 |
| 252 | RRM1 | 2 | 2 | -4.05986 | -5.81768 | OK | 0 | 281.6875114 | 0.285743 | 0 |
| 253 | RRM2 | 2 | 2 | -4.81254 | -6.01153 | OK | 0 | 477.6199408 | 0.484495 | 0 |
| 254 | SLC20A1 | 2 | 2 | -4.17883 | -5.66903 | OK | 0 | 113.0806074 | 0.114698 | 0 |
| 255 | SMC1A | 2 | 2 | -3.78537 | -5.43971 | OK | 0 | 208.891864 | 0.211896 | 0 |
| 256 | SMC2 | 2 | 2 | -4.43242 | -5.7005 | OK | 0 | 351.0626718 | 0.356119 | 0 |
| 257 | TCOF1 | 2 | 2 | -3.8235 | -5.51156 | OK | 0 | 197.243147 | 0.200075 | 0 |
| 258 | TK1 | 2 | 2 | -4.88838 | -6.52042 | OK | 0 | 432.2123114 | 0.438438 | 0 |
| 259 | TMEM106( | 2 | 2 | -4.82531 | -6.50557 | OK | 0 | 220.6734587 | 0.223848 | 0 |
| 260 | TMPO | 2 | 2 | -4.5418 | -5.50397 | OK | 0 | 294.0909353 | 0.298327 | 0 |
| 261 | TYMS | 2 | 2 | -4.60287 | -6.35712 | OK | 0 | 466.8802178 | 0.473602 | 0 |
| 262 | UBE2T | 2 | 2 | -5.24165 | -6.15643 | OK | 0 | 350.8187575 | 0.355866 | 0 |
| 263 | WDR76 | 2 | 2 | -3.75248 | -5.98112 | OK | 0 | 237.6101167 | 0.241015 | 0 |
| 264 | ZWINT | 2 | 2 | -4.90346 | -6.48976 | OK | 0 | 347.7647229 | 0.352768 | 0 |
| 265 | ANLN | 3 | 2 | -6.64805 | -5.71415 | OK | 0 | 475.0182829 | 0.48185 | 0 |
| 266 | ANP32E | 3 | 2 | -6.73942 | -6.17275 | OK | 0 | 250.3710827 | 0.253974 | 0 |
| 267 | ARHGAP11 | 3 | 2 | -6.77772 | -5.83861 | OK | 0 | 324.5657833 | 0.329221 | 0 |
| 268 | ARL6IP1 | 3 | 2 | -6.58939 | -6.87519 | OK | 0 | 225.2103818 | 0.228451 | 0 |
| 269 | ASPM | 3 | 2 | -7.02824 | -6.2946 | OK | 0 | 518.3509325 | 0.525808 | 0 |
| 270 | AURKA | 3 | 2 | -7.2686 | -5.79176 | OK | 0 | 334.7481773 | 0.339552 | 0 |
| 271 | AURKB | 3 | 2 | -6.9672 | -5.38816 | OK | 0 | 402.6692237 | 0.408447 | 0 |
| 272 | BIRC5 | 3 | 2 | -6.5156 | -6.71668 | OK | 0 | 477.3368896 | 0.484209 | 0 |
| 273 | BUB1 | 3 | 2 | -7.06104 | -6.22508 | OK | 0 | 380.6895813 | 0.386139 | 0 |
| 274 | BUB1B | 3 | 2 | -6.43377 | -5.48833 | OK | 0 | 354.5448231 | 0.359622 | 0 |
| 275 | CCNA2 | 3 | 2 | -6.79807 | -5.64315 | OK | 0 | 488.430511 | 0.495453 | 0 |
| 276 | CCNB1 | 3 | 2 | -6.61241 | -6.83441 | OK | 0 | 419.4382514 | 0.425475 | 0 |
| 277 | CCNB2 | 3 | 2 | -6.59044 | -6.82483 | OK | 0 | 427.368051 | 0.433513 | 0 |
| 278 | CDC20 | 3 | 2 | -6.624 | -6.81963 | OK | 0 | 440.6831851 | 0.44702 | 0 |
| 279 | CDCA3 | 3 | 2 | -7.24016 | -5.74895 | OK | 0 | 379.5626451 | 0.385006 | 0 |
| 280 | CDCA8 | 3 | 2 | -7.0953 | -5.58952 | OK | 0 | 398.6056673 | 0.404323 | 0 |
| 281 | CDK1 | 3 | 2 | -6.30193 | -5.3877 | OK | 0 | 456.769362 | 0.463342 | 0 |
| 282 | CDKN3 | 3 | 2 | -6.5829 | -6.83828 | OK | 0 | 449.2620406 | 0.455724 | 0 |
| 283 | CENPA | 3 | 2 | -7.28885 | -6.13209 | OK | 0 | 428.3067258 | 0.434442 | 0 |
| 284 | CENPE | 3 | 2 | -7.09678 | -6.29993 | OK | 0 | 450.2198086 | 0.45669 | 0 |
| 285 | CENPF | 3 | 2 | -6.81276 | -6.48649 | OK | 0 | 508.4613173 | 0.515784 | 0 |
| 286 | CENPN | 3 | 2 | -6.08699 | -6.71067 | OK | 0 | 265.6445725 | 0.26946 | 0 |
| 287 | CENPW | 3 | 2 | -6.13795 | -6.79454 | OK | 0 | 410.6286908 | 0.416546 | 0 |
| 288 | CEP55 | 3 | 2 | -6.93304 | -6.35802 | OK | 0 | 488.8382505 | 0.495863 | 0 |
| 289 | CKAP2 | 3 | 2 | -7.07195 | -5.88238 | OK | 0 | 273.5440429 | 0.27748 | 0 |
| 290 | CKAP2L | 3 | 2 | -6.44093 | -5.34519 | OK | 0 | 394.9006833 | 0.400563 | 0 |
| 291 | CKAP5 | 3 | 2 | -6.92619 | -6.09364 | OK | 0 | 173.8331852 | 0.176327 | 0 |
| 292 | CKS1B | 3 | 2 | -5.84114 | -6.27704 | OK | 0 | 315.6493946 | 0.320196 | 0 |
| 293 | CKS2 | 3 | 2 | -7.37503 | -6.04584 | OK | 0 | 265.0567637 | 0.268873 | 0 |
| 294 | DBF4 | 3 | 2 | -6.54748 | -5.65834 | OK | 0 | 220.4418124 | 0.223609 | 0 |
| 295 | DDX39A | 3 | 2 | -6.25706 | -6.82053 | OK | 0 | 233.5166373 | 0.236876 | 0 |
| 296 | DEPDC1 | 3 | 2 | -7.02457 | -5.82092 | OK | 0 | 429.0195123 | 0.435177 | 0 |
| 297 | DIAPH3 | 3 | 2 | -6.10346 | -5.51381 | OK | 0 | 383.3640675 | 0.388865 | 0 |
| 298 | DLGAP5 | 3 | 2 | -7.1365 | -6.33492 | OK | 0 | 509.9364777 | 0.517261 | 0 |

| | | | | | | | | | | |
|---|---|---|---|---|---|---|---|---|---|---|
| 299 | DTYMK | 3 | 2 | -6.04055 | -6.67561 | OK | 0 | 288.3997887 | 0.292552 | 0 |
| 300 | ECT2 | 3 | 2 | -6.86809 | -6.16992 | OK | 0 | 296.345316 | 0.300606 | 0 |
| 301 | FAM83D | 3 | 2 | -7.37405 | -5.93832 | OK | 0 | 178.0695124 | 0.180627 | 0 |
| 302 | FOXM1 | 3 | 2 | -6.52067 | -5.68305 | OK | 0 | 270.2818502 | 0.274154 | 0 |
| 303 | GTSE1 | 3 | 2 | -7.22138 | -5.75804 | OK | 0 | 459.0597043 | 0.465644 | 0 |
| 304 | H2AFX | 3 | 2 | -6.28617 | -5.26615 | OK | 0 | 263.2412019 | 0.267029 | 0 |
| 305 | H2AFZ | 3 | 2 | -6.00822 | -6.78725 | OK | 0 | 338.9789411 | 0.343862 | 0 |
| 306 | HJURP | 3 | 2 | -6.46623 | -5.38949 | OK | 0 | 334.6720879 | 0.339459 | 0 |
| 307 | HMGB2 | 3 | 2 | -5.92647 | -5.90951 | OK | 0 | 345.8882459 | 0.350872 | 0 |
| 308 | HMGB3 | 3 | 2 | -6.37366 | -6.78776 | OK | 0 | 237.7025782 | 0.241122 | 0 |
| 309 | HMMR | 3 | 2 | -7.17153 | -6.26332 | OK | 0 | 498.1264527 | 0.505283 | 0 |
| 310 | KIF11 | 3 | 2 | -6.35152 | -5.55997 | OK | 0 | 381.813916 | 0.387305 | 0 |
| 311 | KIF20B | 3 | 2 | -6.94495 | -6.16014 | OK | 0 | 377.0758759 | 0.382502 | 0 |
| 312 | KIF23 | 3 | 2 | -7.13769 | -5.70371 | OK | 0 | 443.0448743 | 0.4494 | 0 |
| 313 | KIF2C | 3 | 2 | -6.99928 | -5.81684 | OK | 0 | 400.3925918 | 0.40613 | 0 |
| 314 | KIFC1 | 3 | 2 | -6.39928 | -5.35402 | OK | 0 | 356.6972792 | 0.361812 | 0 |
| 315 | KNL1 | 3 | 2 | -6.75852 | -5.88556 | OK | 0 | 424.8288366 | 0.430926 | 0 |
| 316 | KNSTRN | 3 | 2 | -7.3572 | -6.13795 | OK | 0 | 257.8392919 | 0.261534 | 0 |
| 317 | KPNA2 | 3 | 2 | -7.38063 | -6.06982 | OK | 0 | 318.2757918 | 0.322856 | 0 |
| 318 | LMNB2 | 3 | 2 | -6.62174 | -5.8299 | OK | 0 | 244.4724342 | 0.247986 | 0 |
| 319 | MAD2L1 | 3 | 2 | -5.76078 | -6.2891 | OK | 0 | 414.4505642 | 0.420416 | 0 |
| 320 | MKI67 | 3 | 2 | -6.5938 | -5.80462 | OK | 0 | 526.9579538 | 0.534549 | 0 |
| 321 | MXD3 | 3 | 2 | -7.07385 | -5.51901 | OK | 0 | 290.628833 | 0.294789 | 0 |
| 322 | NCAPD2 | 3 | 2 | -6.68758 | -6.02272 | OK | 0 | 245.3189966 | 0.248842 | 0 |
| 323 | NCAPG | 3 | 2 | -6.3907 | -5.46391 | OK | 0 | 408.6318355 | 0.414496 | 0 |
| 324 | NDC80 | 3 | 2 | -6.37732 | -5.32053 | OK | 0 | 404.7600497 | 0.410566 | 0 |
| 325 | NUF2 | 3 | 2 | -6.94983 | -5.76908 | OK | 0 | 462.6247398 | 0.469268 | 0 |
| 326 | NUSAP1 | 3 | 2 | -6.49613 | -5.38855 | OK | 0 | 516.8984844 | 0.524342 | 0 |
| 327 | PBK | 3 | 2 | -6.3605 | -5.34012 | OK | 0 | 478.6983939 | 0.485578 | 0 |
| 328 | PHF19 | 3 | 2 | -6.14263 | -6.67714 | OK | 0 | 241.2201096 | 0.244688 | 0 |
| 329 | PLK1 | 3 | 2 | -7.37335 | -6.0216 | OK | 0 | 415.7317079 | 0.42169 | 0 |
| 330 | PRC1 | 3 | 2 | -6.78877 | -5.69713 | OK | 0 | 505.0432295 | 0.512314 | 0 |
| 331 | PRR11 | 3 | 2 | -7.0282 | -6.39204 | OK | 0 | 317.0958097 | 0.321633 | 0 |
| 332 | PTTG1 | 3 | 2 | -6.30087 | -6.88225 | OK | 0 | 389.5862866 | 0.395202 | 0 |
| 333 | RACGAP1 | 3 | 2 | -6.87837 | -5.82092 | OK | 0 | 265.8963449 | 0.269714 | 0 |
| 334 | SAP30 | 3 | 2 | -6.62856 | -5.38204 | OK | 0 | 197.6123397 | 0.200452 | 0 |
| 335 | SGO1 | 3 | 2 | -6.40308 | -5.54335 | OK | 0 | 382.9713219 | 0.388466 | 0 |
| 336 | SGO2 | 3 | 2 | -7.08072 | -5.92029 | OK | 0 | 380.4967547 | 0.38596 | 0 |
| 337 | SMC4 | 3 | 2 | -6.67579 | -5.8764 | OK | 0 | 306.5606786 | 0.310976 | 0 |
| 338 | SPC25 | 3 | 2 | -6.34006 | -5.34115 | OK | 0 | 383.6302199 | 0.389129 | 0 |
| 339 | SPDL1 | 3 | 2 | -6.87277 | -5.84299 | OK | 0 | 162.0347877 | 0.164355 | 0 |
| 340 | STMN1 | 3 | 2 | -5.82439 | -6.72014 | OK | 0 | 471.9225507 | 0.478727 | 0 |
| 341 | TACC3 | 3 | 2 | -6.43857 | -5.42185 | OK | 0 | 372.5025199 | 0.37785 | 0 |
| 342 | TOP2A | 3 | 2 | -6.92858 | -5.70179 | OK | 0 | 513.486736 | 0.520879 | 0 |
| 343 | TPX2 | 3 | 2 | -6.83594 | -5.78668 | OK | 0 | 506.3080196 | 0.513596 | 0 |
| 344 | TROAP | 3 | 2 | -6.52557 | -6.87157 | OK | 0 | 288.1826309 | 0.292316 | 0 |
| 345 | TTK | 3 | 2 | -6.79595 | -5.78075 | OK | 0 | 396.9420042 | 0.402639 | 0 |
| 346 | TUBA1B | 3 | 2 | -5.95699 | -6.75562 | OK | 0 | 321.286051 | 0.325914 | 0 |
| 347 | TUBB | 3 | 2 | -5.63881 | -6.60014 | OK | 0 | 298.4346713 | 0.302733 | 0 |
| 348 | UBE2C | 3 | 2 | -7.23741 | -5.70497 | OK | 0 | 441.0608542 | 0.447406 | 0 |
| 349 | UBE2S | 3 | 2 | -7.35072 | -5.93888 | OK | 0 | 200.7814907 | 0.203669 | 0 |
| 350 | A2ML1 | 4 | 1 | 1.891713 | -0.12064 | OK | 0 | 578.0125791 | 0.58635 | 0 |
| 351 | A4GALT | 4 | 1 | 0.016242 | 0.500441 | OK | 0 | 285.3128117 | 0.289414 | 0 |
| 352 | AC005532. | 4 | 1 | 2.785486 | 1.009048 | OK | 0 | 280.409881 | 0.284419 | 0 |
| 353 | AC008397. | 4 | 1 | 0.830541 | 1.249374 | OK | 0 | 143.2849463 | 0.145339 | 0 |
| 354 | AC068587. | 4 | 1 | 0.675013 | 1.432837 | OK | 0 | 196.7178579 | 0.199541 | 0 |
| 355 | ACPP | 4 | 1 | 2.342656 | 0.278526 | OK | 0 | 199.6201526 | 0.202487 | 0 |
| 356 | AGR2 | 4 | 1 | 2.474897 | 1.62014 | OK | 8.10E-113 | 22.54169602 | 0.022852 | 8.14E-113 |
| 357 | AIFM2 | 4 | 1 | 2.378798 | 2.138588 | OK | 0 | 100.4310324 | 0.101865 | 0 |
| 358 | AKR1B10 | 4 | 1 | 2.818513 | -0.77481 | OK | 0 | 225.3844686 | 0.228625 | 0 |

| | | | | | | | | | | |
|---|---|---|---|---|---|---|---|---|---|---|
| 359 | AKR1C1 | 4 | 1 | 2.833771 | -0.82888 | OK | 0 | 84.04776588 | 0.085246 | 0 |
| 360 | AKR1C2 | 4 | 1 | 2.859662 | -0.81994 | OK | 0 | 107.320345 | 0.108855 | 0 |
| 361 | AKR1C3 | 4 | 1 | 2.872497 | -0.79995 | OK | 0 | 115.1200906 | 0.116766 | 0 |
| 362 | AL355312.4 | 4 | 1 | 2.920897 | 2.80667 | OK | 0 | 164.3823196 | 0.166712 | 0 |
| 363 | ALDH1A1 | 4 | 1 | 1.694271 | -0.67138 | OK | 0 | 166.3984131 | 0.16878 | 0 |
| 364 | ALDH3B2 | 4 | 1 | 0.922371 | 0.04813 | OK | 0 | 343.2903912 | 0.348235 | 0 |
| 365 | ALOX12 | 4 | 1 | 2.882485 | 1.17424 | OK | 0 | 235.8135734 | 0.239173 | 0 |
| 366 | ALOX15B | 4 | 1 | 2.391708 | -0.47437 | OK | 0 | 68.19855221 | 0.069166 | 0 |
| 367 | ANKRD35 | 4 | 1 | 2.310583 | -0.383 | OK | 0 | 145.9227264 | 0.148013 | 0 |
| 368 | ANXA1 | 4 | 1 | 2.045873 | 1.193991 | OK | 0 | 250.8113796 | 0.254392 | 0 |
| 369 | ANXA3 | 4 | 1 | 0.921872 | 0.816581 | OK | 0 | 380.7749084 | 0.386263 | 0 |
| 370 | ASPG | 4 | 1 | 1.754522 | 1.836863 | OK | 0 | 436.7374864 | 0.443033 | 0 |
| 371 | ATP13A4 | 4 | 1 | 0.249626 | 0.977228 | OK | 0 | 259.4040441 | 0.263128 | 0 |
| 372 | ATP1B1 | 4 | 1 | 1.334709 | 0.925744 | OK | 0 | 286.7203277 | 0.29085 | 0 |
| 373 | BEX4 | 4 | 1 | 2.87767 | 1.555563 | OK | 0 | 131.258638 | 0.133141 | 0 |
| 374 | BICDL2 | 4 | 1 | 1.032993 | 1.797529 | OK | 0 | 418.8785116 | 0.424916 | 0 |
| 375 | BNIP3 | 4 | 1 | 2.943675 | 1.725236 | OK | 0 | 216.0187422 | 0.219119 | 0 |
| 376 | C10orf99 | 4 | 1 | 2.477624 | -0.38263 | OK | 0 | 300.3851886 | 0.30471 | 0 |
| 377 | C15orf48 | 4 | 1 | 2.983682 | 2.79249 | OK | 0 | 499.9754921 | 0.507174 | 0 |
| 378 | C4orf3 | 4 | 1 | 2.075654 | 1.675033 | OK | 0 | 349.1688992 | 0.354198 | 0 |
| 379 | C8orf82 | 4 | 1 | 2.740574 | -0.10305 | OK | 0 | 129.4587772 | 0.131316 | 0 |
| 380 | CALB2 | 4 | 1 | 2.74079 | 2.808747 | OK | 0 | 230.5555349 | 0.233783 | 0 |
| 381 | CALML5 | 4 | 1 | 1.214829 | 1.378088 | OK | 0 | 329.1750199 | 0.333914 | 0 |
| 382 | CAPN14 | 4 | 1 | 0.013119 | 0.054206 | OK | 0 | 428.6675591 | 0.434848 | 0 |
| 383 | CASTOR1 | 4 | 1 | 2.523008 | 0.100629 | OK | 0 | 217.3463446 | 0.220467 | 0 |
| 384 | CBR3 | 4 | 1 | 2.661851 | -0.33671 | OK | 0 | 226.9621347 | 0.230228 | 0 |
| 385 | CCNYL1 | 4 | 1 | 2.590217 | 2.890558 | OK | 0 | 88.93180731 | 0.090196 | 0 |
| 386 | CD24 | 4 | 1 | 1.887668 | 1.186092 | OK | 0 | 620.4954414 | 0.629444 | 0 |
| 387 | CDA | 4 | 1 | 2.855691 | 1.064485 | OK | 0 | 294.1648014 | 0.298381 | 0 |
| 388 | CDH26 | 4 | 1 | 0.053599 | 0.601023 | OK | 0 | 532.2038556 | 0.539874 | 0 |
| 389 | CEACAM1 | 4 | 1 | 0.515895 | 1.404806 | OK | 0 | 320.9092895 | 0.325517 | 0 |
| 390 | CEACAM5 | 4 | 1 | 0.372261 | 1.135503 | OK | 0 | 568.2811714 | 0.576472 | 0 |
| 391 | CEACAM6 | 4 | 1 | 0.283144 | 1.206619 | OK | 0 | 452.4860501 | 0.459007 | 0 |
| 392 | CES1 | 4 | 1 | 2.3606 | -0.42995 | OK | 0 | 246.4902038 | 0.250017 | 0 |
| 393 | CFD | 4 | 1 | 1.187713 | 1.825721 | OK | 0 | 122.640353 | 0.124399 | 0 |
| 394 | CGN | 4 | 1 | 0.209067 | 0.820116 | OK | 0 | 194.8267738 | 0.197619 | 0 |
| 395 | CITED4 | 4 | 1 | 2.740356 | -0.24382 | OK | 0 | 325.7651487 | 0.330459 | 0 |
| 396 | CLCA4 | 4 | 1 | 0.761373 | 0.817989 | OK | 0 | 482.5257374 | 0.489483 | 0 |
| 397 | CLDN10 | 4 | 1 | 2.529443 | 2.208148 | OK | 0 | 250.1852554 | 0.253675 | 0 |
| 398 | CLIC3 | 4 | 1 | 1.230177 | 1.439811 | OK | 0 | 614.1757048 | 0.623037 | 0 |
| 399 | CNFN | 4 | 1 | 2.98389 | 2.639575 | OK | 0 | 409.1930547 | 0.415089 | 0 |
| 400 | CNN3 | 4 | 1 | 2.288359 | 0.575354 | OK | 0 | 357.4356518 | 0.362586 | 0 |
| 401 | COX7A1 | 4 | 1 | 2.97478 | 3.300049 | OK | 0 | 40.77811439 | 0.041349 | 0 |
| 402 | CPVL | 4 | 1 | 1.721601 | -0.70395 | OK | 0 | 100.6883141 | 0.102125 | 0 |
| 403 | CRCT1 | 4 | 1 | 3.082563 | 2.963947 | OK | 0 | 534.7643131 | 0.542463 | 0 |
| 404 | CRISP3 | 4 | 1 | 2.897492 | 2.875373 | OK | 0 | 735.9159145 | 0.746439 | 0 |
| 405 | CRNN | 4 | 1 | 2.821413 | 1.232228 | OK | 0 | 654.9954888 | 0.664442 | 0 |
| 406 | CRYAB | 4 | 1 | 2.911975 | 1.436168 | OK | 0 | 233.3577469 | 0.236711 | 0 |
| 407 | CST3 | 4 | 1 | 1.679118 | 1.142904 | OK | 0 | 226.8616732 | 0.230121 | 0 |
| 408 | CST6 | 4 | 1 | 2.639604 | 2.887928 | OK | 0 | 511.3496995 | 0.518696 | 0 |
| 409 | CSTA | 4 | 1 | 2.12954 | 1.450359 | OK | 0 | 337.1288839 | 0.341971 | 0 |
| 410 | CSTB | 4 | 1 | 2.192769 | 1.327328 | OK | 0 | 319.268586 | 0.32384 | 0 |
| 411 | CTNNBIP1 | 4 | 1 | 2.674345 | -0.17579 | OK | 0 | 319.16856 | 0.323767 | 0 |
| 412 | CTTNBP2 | 4 | 1 | 1.952348 | -0.34842 | OK | 0 | 341.4477042 | 0.346365 | 0 |
| 413 | CWH43 | 4 | 1 | 2.460269 | 0.201596 | OK | 0 | 285.178642 | 0.289266 | 0 |
| 414 | CXCL17 | 4 | 1 | 0.921783 | 1.222796 | OK | 0 | 477.262265 | 0.484145 | 0 |
| 415 | CXXC5 | 4 | 1 | 0.79263 | 1.695409 | OK | 0 | 230.1887361 | 0.233499 | 0 |
| 416 | CYP2C18 | 4 | 1 | 1.678719 | -0.16493 | OK | 0 | 187.6605934 | 0.190353 | 0 |
| 417 | CYP4B1 | 4 | 1 | 1.479162 | -0.63646 | OK | 0 | 267.3179709 | 0.271166 | 0 |
| 418 | CYP51A1 | 4 | 1 | -0.95081 | 0.937862 | OK | 0 | 198.7919872 | 0.201651 | 0 |

| | | | | | | | | | | |
|---|---|---|---|---|---|---|---|---|---|---|
| 419 | CYSRT1 | 4 | 1 | 1.584584 | 2.066393 | OK | 0 | 625.8978119 | 0.634926 | 0 |
| 420 | DDAH2 | 4 | 1 | 0.487227 | 0.875252 | OK | 0 | 209.6080141 | 0.212623 | 0 |
| 421 | DEFB1 | 4 | 1 | 1.550191 | -0.67266 | OK | 0 | 305.7592893 | 0.310164 | 0 |
| 422 | DEGS2 | 4 | 1 | 2.481654 | -0.16399 | OK | 0 | 370.6941318 | 0.37603 | 0 |
| 423 | DIO2 | 4 | 1 | 0.31495 | 1.053826 | OK | 0 | 444.5257296 | 0.450924 | 0 |
| 424 | DMKN | 4 | 1 | 1.002554 | 1.290453 | OK | 0 | 417.0286926 | 0.423041 | 0 |
| 425 | DNASE1L3 | 4 | 1 | 2.687181 | -0.6936 | OK | 0 | 60.31481784 | 0.06117 | 0 |
| 426 | DSC2 | 4 | 1 | 1.595103 | -0.39286 | OK | 0 | 493.7029115 | 0.500822 | 0 |
| 427 | DSG1 | 4 | 1 | 2.346275 | -0.18419 | OK | 0 | 214.1713862 | 0.21721 | 0 |
| 428 | DYNLT3 | 4 | 1 | 1.46437 | 0.097176 | OK | 0 | 459.416003 | 0.466041 | 0 |
| 429 | ECM1 | 4 | 1 | 3.085808 | 2.952061 | OK | 0 | 518.096523 | 0.525563 | 0 |
| 430 | EGLN3 | 4 | 1 | 0.061429 | 0.815331 | OK | 0 | 245.6607025 | 0.249195 | 0 |
| 431 | EHD3 | 4 | 1 | 0.093974 | 1.610521 | OK | 0 | 191.7791492 | 0.194533 | 0 |
| 432 | ELL2 | 4 | 1 | 1.705955 | -0.29746 | OK | 0 | 209.4195313 | 0.212432 | 0 |
| 433 | ELOVL4 | 4 | 1 | 0.739471 | 0.677094 | OK | 0 | 223.2936189 | 0.226502 | 0 |
| 434 | EMP1 | 4 | 1 | 2.82942 | 2.286886 | OK | 0 | 443.7120214 | 0.450108 | 0 |
| 435 | ENDOU | 4 | 1 | 2.547215 | 0.610316 | OK | 0 | 334.4786733 | 0.339278 | 0 |
| 436 | EPB41L3 | 4 | 1 | 2.862269 | 1.611585 | OK | 0 | 257.5623751 | 0.261212 | 0 |
| 437 | EPHX3 | 4 | 1 | 1.341057 | 1.16126 | OK | 0 | 291.4868824 | 0.295684 | 0 |
| 438 | EREG | 4 | 1 | 2.331188 | 1.988232 | OK | 0 | 235.7198365 | 0.239058 | 0 |
| 439 | ETNK2 | 4 | 1 | 2.509976 | -0.27409 | OK | 0 | 134.7491731 | 0.136673 | 0 |
| 440 | EVPL | 4 | 1 | 0.337948 | 0.063773 | OK | 0 | 378.3322097 | 0.383785 | 0 |
| 441 | F3 | 4 | 1 | 2.918667 | 1.333382 | OK | 0 | 114.480609 | 0.116097 | 0 |
| 442 | FAM25A | 4 | 1 | 2.948397 | 3.376667 | OK | 0 | 608.867759 | 0.617613 | 0 |
| 443 | FAM3B | 4 | 1 | 0.796359 | 0.809433 | OK | 0 | 584.4285613 | 0.592859 | 0 |
| 444 | FAM3D | 4 | 1 | 2.300569 | 1.917214 | OK | 0 | 663.9994596 | 0.673564 | 0 |
| 445 | FAM83A | 4 | 1 | 0.172454 | 1.219733 | OK | 0 | 286.1997994 | 0.29032 | 0 |
| 446 | FETUB | 4 | 1 | 0.283222 | 0.810892 | OK | 0 | 475.6023852 | 0.482433 | 0 |
| 447 | FGF11 | 4 | 1 | -0.05823 | 0.448319 | OK | 0 | 155.6722978 | 0.157902 | 0 |
| 448 | FGFBP1 | 4 | 1 | -0.96586 | 0.937649 | OK | 0 | 309.3057201 | 0.313762 | 0 |
| 449 | FMO2 | 4 | 1 | -0.15245 | 0.761232 | OK | 0 | 177.0366765 | 0.179581 | 0 |
| 450 | FNDC4 | 4 | 1 | 2.759022 | 2.182998 | OK | 0 | 306.9385248 | 0.311293 | 0 |
| 451 | FUOM | 4 | 1 | 1.061342 | 1.679072 | OK | 0 | 103.8095339 | 0.105296 | 0 |
| 452 | GABRP | 4 | 1 | 0.087931 | 0.249211 | OK | 0 | 375.323666 | 0.380732 | 0 |
| 453 | GADD45G | 4 | 1 | 2.479284 | -0.25969 | OK | 0 | 98.44566144 | 0.09985 | 0 |
| 454 | GBP6 | 4 | 1 | 1.611603 | -0.21593 | OK | 0 | 512.5143507 | 0.519904 | 0 |
| 455 | GCNT3 | 4 | 1 | 0.260955 | 0.952298 | OK | 0 | 404.6933837 | 0.410509 | 0 |
| 456 | GDPD3 | 4 | 1 | 2.593174 | 0.719751 | OK | 0 | 469.1828156 | 0.475937 | 0 |
| 457 | GJB2 | 4 | 1 | 1.37193 | 0.681958 | OK | 0 | 355.6477572 | 0.360773 | 0 |
| 458 | GJB6 | 4 | 1 | 1.207641 | 0.704521 | OK | 0 | 402.2779436 | 0.408077 | 0 |
| 459 | GLTP | 4 | 1 | 1.380558 | 0.891863 | OK | 0 | 304.6894249 | 0.309078 | 0 |
| 460 | GNG4 | 4 | 1 | 2.07548 | 1.869417 | OK | 0 | 431.5150323 | 0.437705 | 0 |
| 461 | GPR160 | 4 | 1 | 0.062666 | 0.886411 | OK | 0 | 167.3407471 | 0.169732 | 0 |
| 462 | GPX2 | 4 | 1 | 2.793055 | -0.80877 | OK | 0 | 133.2438973 | 0.135154 | 0 |
| 463 | GPX3 | 4 | 1 | 2.613266 | 0.909512 | OK | 0 | 283.5345129 | 0.287605 | 0 |
| 464 | GRHL3 | 4 | 1 | 1.21195 | -0.14088 | OK | 0 | 251.2249215 | 0.25484 | 0 |
| 465 | GSN | 4 | 1 | 1.752859 | 0.796848 | OK | 0 | 376.292767 | 0.381714 | 0 |
| 466 | GSTA1 | 4 | 1 | 1.779266 | -0.62722 | OK | 0 | 137.3851127 | 0.139339 | 0 |
| 467 | GSTM3 | 4 | 1 | 2.777193 | -0.85057 | OK | 0 | 217.7896179 | 0.220923 | 0 |
| 468 | GSTT2B | 4 | 1 | 2.679172 | 2.768602 | OK | 0 | 42.74445998 | 0.043344 | 0 |
| 469 | HIF1A | 4 | 1 | 1.945038 | 0.138906 | OK | 0 | 149.890068 | 0.152043 | 0 |
| 470 | HMGCS1 | 4 | 1 | -0.91552 | 0.911791 | OK | 0 | 275.885265 | 0.279858 | 0 |
| 471 | HMOX2 | 4 | 1 | 1.807945 | 2.239631 | OK | 0 | 215.2434171 | 0.21834 | 0 |
| 472 | HOPX | 4 | 1 | 2.457014 | 1.924053 | OK | 0 | 471.7445903 | 0.478547 | 0 |
| 473 | HPSE | 4 | 1 | 2.156444 | 2.156605 | OK | 0 | 81.75021888 | 0.082913 | 0 |
| 474 | HS3ST1 | 4 | 1 | -0.01955 | 0.565166 | OK | 0 | 308.3170718 | 0.312757 | 0 |
| 475 | HSPA1A | 4 | 1 | 2.025709 | -0.88918 | OK | 0 | 220.5169297 | 0.223689 | 0 |
| 476 | HSPA1B | 4 | 1 | 1.987977 | -0.8351 | OK | 0 | 166.0934935 | 0.168481 | 0 |
| 477 | IDI1 | 4 | 1 | -0.99348 | 0.983613 | OK | 0 | 154.2160302 | 0.156431 | 0 |
| 478 | IFFO2 | 4 | 1 | 0.368857 | -0.03894 | OK | 0 | 285.3620682 | 0.28947 | 0 |

| | | | | | | | | | | |
|---|---|---|---|---|---|---|---|---|---|---|
| 479 | IFI35 | 4 | 1 | -0.25795 | 0.058423 | OK | 0 | 174.3475072 | 0.176852 | 0 |
| 480 | IGFL1 | 4 | 1 | 2.898487 | 1.440987 | OK | 0 | 176.6862462 | 0.179208 | 0 |
| 481 | IL12A-AS1 | 4 | 1 | 2.602433 | 2.423913 | OK | 0 | 377.3523034 | 0.3827 | 0 |
| 482 | INSIG1 | 4 | 1 | -0.88381 | 0.945471 | OK | 0 | 142.6558754 | 0.144701 | 0 |
| 483 | ISG20 | 4 | 1 | 2.134494 | 1.835485 | OK | 0 | 260.467267 | 0.264207 | 0 |
| 484 | IVL | 4 | 1 | 2.328467 | 0.443246 | OK | 0 | 548.5438079 | 0.556455 | 0 |
| 485 | KALRN | 4 | 1 | 0.312571 | 0.790947 | OK | 0 | 192.7387168 | 0.195502 | 0 |
| 486 | KCNK7 | 4 | 1 | 2.536252 | -0.57966 | OK | 0 | 292.2925825 | 0.296501 | 0 |
| 487 | KIAA1211L | 4 | 1 | 1.704933 | -0.27034 | OK | 0 | 173.7964577 | 0.176292 | 0 |
| 488 | KIF21A | 4 | 1 | 1.416156 | 0.667404 | OK | 0 | 240.6225903 | 0.244085 | 0 |
| 489 | KLK10 | 4 | 1 | 0.531004 | 1.157841 | OK | 0 | 303.5932559 | 0.307967 | 0 |
| 490 | KLK11 | 4 | 1 | 1.636212 | 0.602342 | OK | 0 | 442.831086 | 0.449215 | 0 |
| 491 | KLK12 | 4 | 1 | 0.345473 | 1.598014 | OK | 0 | 623.3935802 | 0.632373 | 0 |
| 492 | KLK13 | 4 | 1 | 0.474616 | 1.377214 | OK | 0 | 495.5938624 | 0.502741 | 0 |
| 493 | KLK6 | 4 | 1 | 0.396939 | 1.916518 | OK | 0 | 157.4716702 | 0.1597 | 0 |
| 494 | KLK7 | 4 | 1 | 0.513065 | 1.071659 | OK | 0 | 394.0594654 | 0.399727 | 0 |
| 495 | KLK8 | 4 | 1 | 0.59466 | 1.212487 | OK | 0 | 184.9105103 | 0.187569 | 0 |
| 496 | KRT24 | 4 | 1 | 0.514017 | 1.510684 | OK | 0 | 279.6687924 | 0.283669 | 0 |
| 497 | KRT4 | 4 | 1 | 1.337342 | -0.10778 | OK | 0 | 510.4046644 | 0.517761 | 0 |
| 498 | KRT6A | 4 | 1 | 2.29938 | -0.22425 | OK | 0 | 489.094413 | 0.496141 | 0 |
| 499 | KRT6B | 4 | 1 | 2.369385 | -0.19288 | OK | 0 | 379.5929138 | 0.385064 | 0 |
| 500 | KRT6C | 4 | 1 | 2.39509 | -0.19643 | OK | 0 | 295.7432313 | 0.3 | 0 |
| 501 | KRT78 | 4 | 1 | 3.040286 | 3.064029 | OK | 0 | 676.0771032 | 0.6858 | 0 |
| 502 | KRTDAP | 4 | 1 | 0.817564 | 1.374744 | OK | 0 | 86.4545668 | 0.087676 | 0 |
| 503 | LCN2 | 4 | 1 | 0.287852 | 2.102612 | OK | 0 | 406.9989418 | 0.412865 | 0 |
| 504 | LGALS7B | 4 | 1 | 0.617082 | -0.05204 | OK | 0 | 297.1630148 | 0.301438 | 0 |
| 505 | LINC02487 | 4 | 1 | 2.338763 | 2.180424 | OK | 0 | 313.4188888 | 0.317835 | 0 |
| 506 | LIPH | 4 | 1 | 0.696611 | 1.137707 | OK | 0 | 328.1725215 | 0.332871 | 0 |
| 507 | LPCAT4 | 4 | 1 | 0.042643 | 0.44259 | OK | 0 | 224.577213 | 0.227806 | 0 |
| 508 | LRRC8A | 4 | 1 | 0.365403 | 0.177683 | OK | 0 | 245.6056258 | 0.249141 | 0 |
| 509 | LY6G6C | 4 | 1 | 0.562589 | 1.142949 | OK | 0 | 270.3565806 | 0.274245 | 0 |
| 510 | LYPD2 | 4 | 1 | 2.941634 | 2.830833 | OK | 0 | 625.1571043 | 0.634153 | 0 |
| 511 | LYPD3 | 4 | 1 | 1.610132 | 0.84233 | OK | 0 | 476.2371706 | 0.483103 | 0 |
| 512 | MAF | 4 | 1 | 1.601965 | -0.42772 | OK | 0 | 206.0983223 | 0.209063 | 0 |
| 513 | MAFB | 4 | 1 | 2.128156 | -0.67614 | OK | 0 | 232.9020533 | 0.236254 | 0 |
| 514 | MAL | 4 | 1 | 2.821852 | 1.165321 | OK | 0 | 597.542185 | 0.606161 | 0 |
| 515 | MAL2 | 4 | 1 | 1.590067 | 0.476338 | OK | 0 | 473.1410088 | 0.479962 | 0 |
| 516 | MBOAT2 | 4 | 1 | 0.428182 | 0.634142 | OK | 0 | 367.9772419 | 0.373278 | 0 |
| 517 | ME1 | 4 | 1 | 1.418652 | 0.551196 | OK | 0 | 214.381397 | 0.217458 | 0 |
| 518 | MGLL | 4 | 1 | 2.435697 | -0.08866 | OK | 0 | 514.0350502 | 0.521448 | 0 |
| 519 | MGST1 | 4 | 1 | 2.816962 | -0.86092 | OK | 0 | 145.4212036 | 0.14751 | 0 |
| 520 | MIR210HG | 4 | 1 | 2.529175 | 1.486825 | OK | 0 | 188.879432 | 0.19159 | 0 |
| 521 | MROH6 | 4 | 1 | 0.175719 | 1.77124 | OK | 0 | 189.8738339 | 0.192598 | 0 |
| 522 | MSMO1 | 4 | 1 | -0.96475 | 0.951216 | OK | 0 | 167.6398132 | 0.170048 | 0 |
| 523 | MT1X | 4 | 1 | 1.249869 | -0.95343 | OK | 0 | 419.4349072 | 0.425482 | 0 |
| 524 | MUC15 | 4 | 1 | 1.499438 | -0.15659 | OK | 0 | 391.3346429 | 0.396974 | 0 |
| 525 | MUC21 | 4 | 1 | 2.914217 | 1.215436 | OK | 0 | 699.7974673 | 0.709878 | 0 |
| 526 | MUC4 | 4 | 1 | 0.076539 | 0.971408 | OK | 0 | 241.1563831 | 0.244621 | 0 |
| 527 | MYH14 | 4 | 1 | 0.457549 | 0.138987 | OK | 0 | 368.9494007 | 0.374265 | 0 |
| 528 | NAGK | 4 | 1 | 2.319376 | 0.315768 | OK | 0 | 393.0135785 | 0.398679 | 0 |
| 529 | NBEAL2 | 4 | 1 | 0.38379 | 0.043907 | OK | 0 | 277.5770228 | 0.281573 | 0 |
| 530 | NDUFA4L2 | 4 | 1 | 2.797015 | 1.523151 | OK | 0 | 276.55743 | 0.280534 | 0 |
| 531 | NQO1 | 4 | 1 | 2.804524 | -0.76223 | OK | 0 | 235.9907708 | 0.239387 | 0 |
| 532 | NUAK2 | 4 | 1 | 0.500038 | 0.875809 | OK | 0 | 134.9488258 | 0.13688 | 0 |
| 533 | NUCB2 | 4 | 1 | 2.366095 | 0.279266 | OK | 0 | 442.5090941 | 0.448889 | 0 |
| 534 | NUDT8 | 4 | 1 | 2.789125 | -0.1283 | OK | 0 | 362.8969509 | 0.368127 | 0 |
| 535 | OAS1 | 4 | 1 | 1.002852 | 0.851389 | OK | 0 | 234.2392558 | 0.23761 | 0 |
| 536 | PALMD | 4 | 1 | 2.44858 | 0.162384 | OK | 0 | 173.6776309 | 0.176173 | 0 |
| 537 | PAQR8 | 4 | 1 | 2.650169 | 0.643744 | OK | 0 | 176.6039156 | 0.179127 | 0 |
| 538 | PCDH1 | 4 | 1 | 0.242435 | 1.470965 | OK | 0 | 248.4782395 | 0.252048 | 0 |

| | | | | | | | | | | |
|---|---|---|---|---|---|---|---|---|---|---|
| 539 | PDLIM2 | 4 | 1 | 2.020938 | 1.288282 | OK | 0 | 267.415289 | 0.271266 | 0 |
| 540 | PGD | 4 | 1 | 0.832945 | -0.03596 | OK | 0 | 352.5557302 | 0.357636 | 0 |
| 541 | PHLDB3 | 4 | 1 | 0.85946 | 1.569552 | OK | 0 | 104.6262452 | 0.106123 | 0 |
| 542 | PI3 | 4 | 1 | 0.651783 | 1.354279 | OK | 0 | 407.2748079 | 0.413141 | 0 |
| 543 | PIM1 | 4 | 1 | 2.146766 | 1.915428 | OK | 0 | 263.4145094 | 0.267207 | 0 |
| 544 | PLAC8 | 4 | 1 | 0.144597 | 0.950613 | OK | 0 | 419.7971349 | 0.425848 | 0 |
| 545 | PLBD1 | 4 | 1 | 1.078558 | 1.830584 | OK | 0 | 281.9677579 | 0.286028 | 0 |
| 546 | PMM1 | 4 | 1 | 2.155952 | 1.748419 | OK | 0 | 207.2343136 | 0.210215 | 0 |
| 547 | PPM1K | 4 | 1 | 1.83032 | -0.58099 | OK | 0 | 138.4886736 | 0.140474 | 0 |
| 548 | PPP1R3C | 4 | 1 | 0.153254 | 1.807265 | OK | 0 | 237.4020019 | 0.240803 | 0 |
| 549 | PRSS3 | 4 | 1 | 0.542852 | 1.180968 | OK | 0 | 629.5136468 | 0.638585 | 0 |
| 550 | PTGR1 | 4 | 1 | 2.571143 | -0.5481 | OK | 0 | 281.6081875 | 0.285664 | 0 |
| 551 | PTN | 4 | 1 | 1.697821 | -0.68535 | OK | 0 | 326.1350535 | 0.330834 | 0 |
| 552 | RAB10 | 4 | 1 | 1.839725 | 0.852698 | OK | 0 | 370.9534977 | 0.376298 | 0 |
| 553 | RAET1E | 4 | 1 | 1.454728 | 0.26924 | OK | 0 | 328.9915155 | 0.333727 | 0 |
| 554 | RAET1L | 4 | 1 | 0.805305 | 1.064348 | OK | 0 | 355.554163 | 0.360668 | 0 |
| 555 | RASGRP1 | 4 | 1 | -0.11482 | 0.029162 | OK | 0 | 246.8680743 | 0.250418 | 0 |
| 556 | RBP1 | 4 | 1 | 2.246384 | 1.950141 | OK | 0 | 66.85399463 | 0.067804 | 0 |
| 557 | RBP7 | 4 | 1 | 2.573333 | 1.235962 | OK | 0 | 209.4662368 | 0.212477 | 0 |
| 558 | RHCG | 4 | 1 | 1.694526 | 1.071362 | OK | 0 | 628.5327879 | 0.637594 | 0 |
| 559 | RHOD | 4 | 1 | 1.926361 | 1.449176 | OK | 0 | 296.7577027 | 0.301032 | 0 |
| 560 | RTKN2 | 4 | 1 | 2.170628 | 2.034021 | OK | 0 | 206.497866 | 0.209466 | 0 |
| 561 | S100A12 | 4 | 1 | 2.697557 | 2.427675 | OK | 0 | 288.4841636 | 0.292622 | 0 |
| 562 | S100A14 | 4 | 1 | 2.043817 | 1.279993 | OK | 0 | 107.595503 | 0.109127 | 0 |
| 563 | S100A4 | 4 | 1 | 1.69841 | 1.330493 | OK | 0 | 78.06581567 | 0.079181 | 0 |
| 564 | S100A6 | 4 | 1 | 2.081936 | 1.355508 | OK | 0 | 175.1576173 | 0.177668 | 0 |
| 565 | S100A7 | 4 | 1 | 0.506301 | 1.987516 | OK | 0 | 182.4658658 | 0.185082 | 0 |
| 566 | S100A8 | 4 | 1 | 2.121369 | -1.02203 | OK | 0 | 350.5692893 | 0.355615 | 0 |
| 567 | SAMD5 | 4 | 1 | 1.96612 | -0.2933 | OK | 0 | 212.2935678 | 0.215333 | 0 |
| 568 | SBSN | 4 | 1 | 1.09376 | 1.207814 | OK | 0 | 506.6516845 | 0.513957 | 0 |
| 569 | SCD | 4 | 1 | -0.84 | 0.868824 | OK | 0 | 209.145983 | 0.212153 | 0 |
| 570 | SCGB1A1 | 4 | 1 | 1.448553 | 0.937721 | OK | 0 | 85.4408255 | 0.086658 | 0 |
| 571 | SCUBE2 | 4 | 1 | 0.044077 | 0.279219 | OK | 0 | 170.772135 | 0.17322 | 0 |
| 572 | SDR16C5 | 4 | 1 | 1.681411 | 0.284152 | OK | 0 | 386.9208303 | 0.392498 | 0 |
| 573 | SERPINB1 | 4 | 1 | 2.124381 | 0.752238 | OK | 0 | 461.2169169 | 0.467864 | 0 |
| 574 | SERPINB11 | 4 | 1 | 2.617615 | 0.656965 | OK | 0 | 75.04807329 | 0.07611 | 0 |
| 575 | SERPINB6 | 4 | 1 | 1.261897 | 1.348104 | OK | 0 | 302.9232283 | 0.307287 | 0 |
| 576 | SF3B3 | 4 | 1 | 2.726534 | 1.47483 | OK | 0 | 169.2326086 | 0.171664 | 0 |
| 577 | SFTA2 | 4 | 1 | 2.84756 | 2.707777 | OK | 0 | 484.0419175 | 0.490961 | 0 |
| 578 | SH3GL1 | 4 | 1 | 0.254033 | 1.882952 | OK | 0 | 270.8452574 | 0.274745 | 0 |
| 579 | SHROOM3 | 4 | 1 | 2.253306 | 2.040335 | OK | 0 | 238.2262222 | 0.241646 | 0 |
| 580 | SLC26A2 | 4 | 1 | 1.112813 | -0.32135 | OK | 0 | 257.2039339 | 0.260906 | 0 |
| 581 | SLC35C1 | 4 | 1 | 0.980234 | 1.880277 | OK | 0 | 194.9546313 | 0.197755 | 0 |
| 582 | SLPI | 4 | 1 | 0.859083 | 1.300873 | OK | 0 | 544.0574276 | 0.551905 | 0 |
| 583 | SLURP1 | 4 | 1 | 2.829994 | 1.155083 | OK | 0 | 594.8192823 | 0.603392 | 0 |
| 584 | SMAGP | 4 | 1 | 1.520575 | 1.925748 | OK | 0 | 349.3645672 | 0.354399 | 0 |
| 585 | SMIM5 | 4 | 1 | 1.426348 | 1.577802 | OK | 0 | 487.5480726 | 0.494577 | 0 |
| 586 | SNX31 | 4 | 1 | 0.59056 | 0.757567 | OK | 0 | 219.0628636 | 0.222205 | 0 |
| 587 | SPAG17 | 4 | 1 | 1.737239 | -0.26933 | OK | 0 | 227.284024 | 0.230549 | 0 |
| 588 | SPINK5 | 4 | 1 | 1.724913 | 0.030584 | OK | 0 | 579.7515218 | 0.588112 | 0 |
| 589 | SPINK7 | 4 | 1 | 3.047699 | 2.812878 | OK | 0 | 605.8485447 | 0.614539 | 0 |
| 590 | SPRR1A | 4 | 1 | 1.754708 | 1.922407 | OK | 0 | 392.3951057 | 0.398041 | 0 |
| 591 | SPRR1B | 4 | 1 | 0.537958 | 1.829924 | OK | 0 | 378.2818572 | 0.383733 | 0 |
| 592 | SPRR2A | 4 | 1 | 0.230627 | 2.248157 | OK | 0 | 310.9490761 | 0.315425 | 0 |
| 593 | SPRR2D | 4 | 1 | 0.228177 | 2.289025 | OK | 0 | 378.9342593 | 0.384366 | 0 |
| 594 | SPRR2E | 4 | 1 | 0.292071 | 2.122066 | OK | 0 | 232.6424019 | 0.235981 | 0 |
| 595 | SPRR2F | 4 | 1 | 0.327571 | 2.066921 | OK | 0 | 45.12483153 | 0.045674 | 0 |
| 596 | SPRR3 | 4 | 1 | 0.452652 | 0.557017 | OK | 0 | 489.8141973 | 0.496877 | 0 |
| 597 | SQLE | 4 | 1 | -0.96074 | 0.928281 | OK | 0 | 238.350283 | 0.241781 | 0 |
| 598 | SRPX2 | 4 | 1 | 2.283137 | 2.374033 | OK | 0 | 261.7557817 | 0.265484 | 0 |

| | | | | | | | | | | |
|---|---|---|---|---|---|---|---|---|---|---|
| 599 | ST3GAL4 | 4 | 1 | 2.348349 | 2.099585 | OK | 0 | 460.4004572 | 0.466986 | 0 |
| 600 | STN1 | 4 | 1 | 1.917301 | 2.140529 | OK | 0 | 313.9247032 | 0.318446 | 0 |
| 601 | SUCO | 4 | 1 | 2.346721 | 0.310317 | OK | 0 | 66.78028391 | 0.067731 | 0 |
| 602 | SULT2B1 | 4 | 1 | 1.283355 | 0.90419 | OK | 0 | 550.0032649 | 0.557936 | 0 |
| 603 | TGM1 | 4 | 1 | 1.230725 | 0.828541 | OK | 0 | 437.7703088 | 0.44408 | 0 |
| 604 | TGM3 | 4 | 1 | 2.873058 | 1.111881 | OK | 0 | 660.8776151 | 0.670402 | 0 |
| 605 | TM7SF2 | 4 | 1 | 0.997985 | 1.042426 | OK | 0 | 508.0459673 | 0.515371 | 0 |
| 606 | TMEM154 | 4 | 1 | 1.304725 | 0.400205 | OK | 0 | 331.3103783 | 0.336084 | 0 |
| 607 | TMEM184/ | 4 | 1 | 0.453562 | 0.09951 | OK | 0 | 257.9570182 | 0.261668 | 0 |
| 608 | TMEM265 | 4 | 1 | 1.326204 | 1.658581 | OK | 0 | 382.3852645 | 0.387895 | 0 |
| 609 | TMEM45B | 4 | 1 | 0.345424 | 0.973879 | OK | 0 | 457.9010382 | 0.464497 | 0 |
| 610 | TMEM79 | 4 | 1 | 1.57528 | 0.764211 | OK | 0 | 280.1790486 | 0.284214 | 0 |
| 611 | TMPRSS11 | 4 | 1 | 2.823402 | 2.631305 | OK | 0 | 713.8195537 | 0.724078 | 0 |
| 612 | TP53I3 | 4 | 1 | 2.734518 | 0.933128 | OK | 0 | 274.306019 | 0.278255 | 0 |
| 613 | TPRG1 | 4 | 1 | 2.007654 | 1.882388 | OK | 0 | 530.754327 | 0.538401 | 0 |
| 614 | TRNP1 | 4 | 1 | 0.038939 | 0.50024 | OK | 0 | 420.8161194 | 0.426882 | 0 |
| 615 | TSPAN3 | 4 | 1 | -0.11122 | 0.357103 | OK | 0 | 278.3696823 | 0.282379 | 0 |
| 616 | TST | 4 | 1 | 1.293028 | 1.491884 | OK | 0 | 360.9137332 | 0.366115 | 0 |
| 617 | TUBA4A | 4 | 1 | 2.469003 | 0.029911 | OK | 0 | 300.8033343 | 0.305136 | 0 |
| 618 | UPK1B | 4 | 1 | 0.215961 | 0.890304 | OK | 0 | 147.9459828 | 0.150053 | 0 |
| 619 | UPK3BL1 | 4 | 1 | 1.404048 | 1.797196 | OK | 0 | 359.6832323 | 0.364866 | 0 |
| 620 | VAT1 | 4 | 1 | 0.079908 | 1.560359 | OK | 0 | 294.9174343 | 0.299164 | 0 |
| 621 | WNK4 | 4 | 1 | 1.570086 | -0.22962 | OK | 0 | 226.5163455 | 0.229769 | 0 |
| 622 | WNT5A | 4 | 1 | 1.877451 | -0.10791 | OK | 0 | 161.7628873 | 0.164081 | 0 |
| 623 | ZBED2 | 4 | 1 | 2.91416 | 1.823195 | OK | 0 | 136.8986165 | 0.138827 | 0 |
| 624 | ZDHHC21 | 4 | 1 | 1.375249 | 0.227309 | OK | 0 | 77.28935721 | 0.078391 | 0 |
| 625 | ABLIM1 | 5 | 1 | -0.21718 | 3.794879 | OK | 0 | 327.5709265 | 0.332291 | 0 |
| 626 | ABLIM3 | 5 | 1 | 1.503048 | 4.188348 | OK | 0 | 422.3491115 | 0.428383 | 0 |
| 627 | AC004816. | 5 | 1 | 1.851445 | 3.969704 | OK | 0 | 241.5058012 | 0.244973 | 0 |
| 628 | AC006333. | 5 | 1 | 2.135271 | 4.528072 | OK | 0 | 96.67960308 | 0.098057 | 0 |
| 629 | ACADM | 5 | 1 | 1.650184 | 3.311115 | OK | 0 | 177.5233061 | 0.180075 | 0 |
| 630 | ADGRF1 | 5 | 1 | 0.548775 | 3.839283 | OK | 0 | 319.8729142 | 0.324472 | 0 |
| 631 | ADIPOR1 | 5 | 1 | 1.811903 | 3.611346 | OK | 0 | 169.9591125 | 0.172402 | 0 |
| 632 | ADIRF | 5 | 1 | 2.665995 | 3.774622 | OK | 0 | 311.1721144 | 0.315655 | 0 |
| 633 | AFDN | 5 | 1 | -0.8059 | 3.313125 | OK | 0 | 299.8386758 | 0.304158 | 0 |
| 634 | AGFG2 | 5 | 1 | 0.393171 | 3.175922 | OK | 0 | 118.2191598 | 0.119908 | 0 |
| 635 | AIF1L | 5 | 1 | 2.70897 | 3.52031 | OK | 0 | 559.0065887 | 0.567029 | 0 |
| 636 | AL121761. | 5 | 1 | 0.752995 | 2.716315 | OK | 0 | 390.2613009 | 0.395876 | 0 |
| 637 | AL121933.: | 5 | 1 | 0.371199 | 2.828874 | OK | 0 | 116.2867347 | 0.117875 | 0 |
| 638 | AL133453. | 5 | 1 | 0.284853 | 3.984752 | OK | 0 | 164.2049455 | 0.166556 | 0 |
| 639 | AL512274. | 5 | 1 | -0.38873 | 3.327168 | OK | 0 | 202.180458 | 0.205084 | 0 |
| 640 | AL627171.: | 5 | 1 | 0.66012 | 3.449998 | OK | 2.57E-119 | 23.19471124 | 0.023516 | 2.59E-119 |
| 641 | ALDH1A3 | 5 | 1 | 0.683516 | 4.196034 | OK | 0 | 343.992798 | 0.348919 | 0 |
| 642 | ANGPTL4 | 5 | 1 | 0.859333 | 2.25081 | OK | 0 | 101.3838166 | 0.102834 | 0 |
| 643 | ANXA11 | 5 | 1 | 1.525562 | 3.458153 | OK | 0 | 232.8567725 | 0.236208 | 0 |
| 644 | ANXA9 | 5 | 1 | 1.158595 | 3.677791 | OK | 0 | 569.6644374 | 0.577825 | 0 |
| 645 | AP000866. | 5 | 1 | 2.281986 | 4.074777 | OK | 0 | 282.3087951 | 0.286275 | 0 |
| 646 | ARHGAP27 | 5 | 1 | 0.6042 | 3.512483 | OK | 0 | 161.8326417 | 0.164152 | 0 |
| 647 | ARHGDIB | 5 | 1 | 1.870289 | 3.002141 | OK | 0 | 101.4788249 | 0.102926 | 0 |
| 648 | ATP6V1C2 | 5 | 1 | 0.852707 | 4.168924 | OK | 0 | 489.3704892 | 0.496324 | 0 |
| 649 | AVPI1 | 5 | 1 | 1.252371 | 5.156239 | OK | 0 | 86.16357186 | 0.087393 | 0 |
| 650 | B3GNT3 | 5 | 1 | 0.959854 | 3.224969 | OK | 0 | 409.9786668 | 0.415811 | 0 |
| 651 | B3GNT8 | 5 | 1 | 0.836993 | 2.476716 | OK | 0 | 131.5788502 | 0.133463 | 0 |
| 652 | B4GALT5 | 5 | 1 | 0.114756 | 3.200645 | OK | 0 | 121.3325945 | 0.123071 | 0 |
| 653 | BAIAP2L2 | 5 | 1 | 2.360196 | 3.156896 | OK | 0 | 204.8559365 | 0.207764 | 0 |
| 654 | BCAS1 | 5 | 1 | 0.791617 | 2.966411 | OK | 0 | 226.2373063 | 0.229458 | 0 |
| 655 | BICDL1 | 5 | 1 | 1.411277 | 4.532831 | OK | 0 | 89.13088503 | 0.090403 | 0 |
| 656 | BX255925. | 5 | 1 | 1.477151 | 2.671067 | OK | 0 | 227.5627156 | 0.230835 | 0 |
| 657 | C15orf62 | 5 | 1 | 1.008478 | 3.177142 | OK | 0 | 272.4061085 | 0.276294 | 0 |
| 658 | C1orf116 | 5 | 1 | -0.06089 | 3.797162 | OK | 0 | 260.1860847 | 0.263932 | 0 |

| | | | | | | | | | | |
|---|---|---|---|---|---|---|---|---|---|---|
| 659 | C5orf66-AS | 5 | 1 | 2.320604 | 4.171162 | OK | 0 | 177.4219913 | 0.179972 | 0 |
| 660 | C6orf132 | 5 | 1 | 1.550416 | 4.281843 | OK | 0 | 324.951936 | 0.329634 | 0 |
| 661 | CAMK2N1 | 5 | 1 | 1.366101 | 3.570694 | OK | 0 | 720.0195302 | 0.73036 | 0 |
| 662 | CAPN5 | 5 | 1 | 0.659942 | 3.134356 | OK | 0 | 206.50282 | 0.209451 | 0 |
| 663 | CASP8 | 5 | 1 | 1.744439 | 4.656468 | OK | 0 | 51.36895228 | 0.052097 | 0 |
| 664 | CCDC127 | 5 | 1 | 1.84141 | 3.403882 | OK | 0 | 88.04063806 | 0.089296 | 0 |
| 665 | CCNG2 | 5 | 1 | 0.089775 | 3.817275 | OK | 0 | 309.511388 | 0.313961 | 0 |
| 666 | CD59 | 5 | 1 | 1.786743 | 2.361181 | OK | 0 | 285.6603327 | 0.289774 | 0 |
| 667 | CD68 | 5 | 1 | 1.048224 | 2.470703 | OK | 0 | 146.6964098 | 0.148798 | 0 |
| 668 | CDC42EP5 | 5 | 1 | 1.02867 | 2.805127 | OK | 0 | 119.8495075 | 0.121568 | 0 |
| 669 | CDKN2AIP | 5 | 1 | -0.60176 | 3.547562 | OK | 0 | 181.2291134 | 0.183835 | 0 |
| 670 | CEACAM7 | 5 | 1 | 0.550713 | 2.842703 | OK | 0 | 121.9883098 | 0.123644 | 0 |
| 671 | CFL2 | 5 | 1 | 2.030293 | 3.209031 | OK | 1.09E-290 | 36.41840935 | 0.036928 | 1.10E-290 |
| 672 | CKB | 5 | 1 | 2.001853 | 3.755931 | OK | 0 | 120.3262199 | 0.122052 | 0 |
| 673 | CLIP1 | 5 | 1 | -0.85947 | 2.905122 | OK | 0 | 309.4371327 | 0.313895 | 0 |
| 674 | CPEB4 | 5 | 1 | -0.2317 | 3.266992 | OK | 0 | 169.5389228 | 0.171973 | 0 |
| 675 | CPPED1 | 5 | 1 | 2.085446 | 3.735493 | OK | 0 | 163.1978539 | 0.165542 | 0 |
| 676 | CRYM | 5 | 1 | 1.917867 | 4.557876 | OK | 0 | 364.347147 | 0.369392 | 0 |
| 677 | CTSV | 5 | 1 | 1.107496 | 3.798141 | OK | 0 | 224.6085663 | 0.227823 | 0 |
| 678 | CTTNBP2N | 5 | 1 | -0.71848 | 3.646636 | OK | 0 | 153.6084369 | 0.155814 | 0 |
| 679 | CYTOR | 5 | 1 | 1.116177 | 3.802751 | OK | 0 | 142.4010132 | 0.144432 | 0 |
| 680 | DDIT4 | 5 | 1 | 0.896958 | 2.290771 | OK | 0 | 66.00636315 | 0.066947 | 0 |
| 681 | DENND10 | 5 | 1 | 1.980604 | 2.828616 | OK | 0 | 144.8079838 | 0.146887 | 0 |
| 682 | DHRS9 | 5 | 1 | 1.585833 | 4.68469 | OK | 0 | 693.6118164 | 0.703454 | 0 |
| 683 | DNAJA4 | 5 | 1 | -0.55824 | 2.998199 | OK | 0 | 74.40412268 | 0.075464 | 0 |
| 684 | DNAJC5 | 5 | 1 | 1.229144 | 3.947332 | OK | 0 | 138.0881593 | 0.140069 | 0 |
| 685 | DOCK9 | 5 | 1 | -0.41522 | 3.633717 | OK | 0 | 144.7966763 | 0.146873 | 0 |
| 686 | DTX2 | 5 | 1 | 0.69056 | 3.525753 | OK | 0 | 121.0784942 | 0.122814 | 0 |
| 687 | DUOX2 | 5 | 1 | 0.207683 | 2.870212 | OK | 0 | 90.99370595 | 0.092289 | 0 |
| 688 | DUOXA2 | 5 | 1 | 0.234776 | 2.687604 | OK | 0 | 97.45812282 | 0.098786 | 0 |
| 689 | DUSP4 | 5 | 1 | 1.816091 | 4.581932 | OK | 0 | 385.3263131 | 0.390708 | 0 |
| 690 | ELF3 | 5 | 1 | 1.00227 | 2.629482 | OK | 0 | 306.605412 | 0.311022 | 0 |
| 691 | ELF3-AS1 | 5 | 1 | 1.94126 | 4.355634 | OK | 0 | 278.6263531 | 0.282592 | 0 |
| 692 | EPS8L1 | 5 | 1 | 0.754542 | 3.745923 | OK | 0 | 529.1657014 | 0.536786 | 0 |
| 693 | EPS8L2 | 5 | 1 | 1.287297 | 3.232457 | OK | 0 | 231.9668079 | 0.235305 | 0 |
| 694 | ESAM | 5 | 1 | 1.521073 | 4.614385 | OK | 0 | 512.8087214 | 0.520027 | 0 |
| 695 | FAM107B | 5 | 1 | 1.112191 | 4.82698 | OK | 0 | 177.8900624 | 0.18043 | 0 |
| 696 | FBXO32 | 5 | 1 | 1.164076 | 4.145256 | OK | 0 | 223.7851275 | 0.227002 | 0 |
| 697 | FBXO34 | 5 | 1 | -0.49435 | 3.499713 | OK | 0 | 138.5374189 | 0.140525 | 0 |
| 698 | FBXW11 | 5 | 1 | -0.57365 | 3.765322 | OK | 0 | 83.83945136 | 0.085037 | 0 |
| 699 | FCHO2 | 5 | 1 | -0.79349 | 3.428331 | OK | 0 | 219.8072272 | 0.222969 | 0 |
| 700 | FRMD4B | 5 | 1 | -0.86873 | 2.61828 | OK | 0 | 234.6813625 | 0.238059 | 0 |
| 701 | FTH1 | 5 | 1 | 2.290753 | 4.189312 | OK | 0 | 204.1897415 | 0.20712 | 0 |
| 702 | FUT3 | 5 | 1 | 0.442447 | 3.280182 | OK | 0 | 413.1890585 | 0.419138 | 0 |
| 703 | FUT6 | 5 | 1 | 0.545535 | 3.329131 | OK | 0 | 382.9857088 | 0.38848 | 0 |
| 704 | GABARAPL | 5 | 1 | 2.326746 | 4.353869 | OK | 0 | 237.2393712 | 0.240654 | 0 |
| 705 | GALNT1 | 5 | 1 | -0.6475 | 3.675255 | OK | 0 | 263.3999098 | 0.267192 | 0 |
| 706 | GCC2 | 5 | 1 | -0.07907 | 3.961014 | OK | 0 | 100.5941233 | 0.102034 | 0 |
| 707 | GCHFR | 5 | 1 | 1.234853 | 3.585375 | OK | 0 | 226.9033018 | 0.230168 | 0 |
| 708 | GFOD2 | 5 | 1 | 0.210803 | 3.698658 | OK | 0 | 99.32999089 | 0.100749 | 0 |
| 709 | GMDS | 5 | 1 | 2.136051 | 2.48588 | OK | 0 | 239.7198196 | 0.243167 | 0 |
| 710 | GNA15 | 5 | 1 | 1.357638 | 3.187749 | OK | 0 | 264.9053798 | 0.268719 | 0 |
| 711 | GPRC5A | 5 | 1 | 1.185082 | 4.240656 | OK | 0 | 212.3383391 | 0.215374 | 0 |
| 712 | GPT2 | 5 | 1 | 1.065449 | 3.264035 | OK | 0 | 174.4665032 | 0.176973 | 0 |
| 713 | GRHL1 | 5 | 1 | -0.45495 | 2.886757 | OK | 0 | 353.7192684 | 0.358814 | 0 |
| 714 | H19 | 5 | 1 | 1.840775 | 2.769955 | OK | 0 | 137.5867092 | 0.139557 | 0 |
| 715 | H1F0 | 5 | 1 | 0.761058 | 2.470833 | OK | 0 | 189.7791318 | 0.192508 | 0 |
| 716 | H2AFJ | 5 | 1 | 2.400451 | 4.163906 | OK | 0 | 269.0300459 | 0.272904 | 0 |
| 717 | HEBP2 | 5 | 1 | 2.182239 | 3.286149 | OK | 0 | 362.6946454 | 0.367919 | 0 |
| 718 | HLCS | 5 | 1 | 2.068586 | 4.209117 | OK | 0 | 85.73277037 | 0.086951 | 0 |

| | | | | | | | | | | |
|---|---|---|---|---|---|---|---|---|---|---|
| 719 | HMOX1 | 5 | 1 | 0.779808 | 4.198033 | OK | 0 | 448.5879042 | 0.454981 | 0 |
| 720 | HOTAIRM1 | 5 | 1 | 2.213121 | 4.560225 | OK | 0 | 95.55791247 | 0.096925 | 0 |
| 721 | HS3ST6 | 5 | 1 | 2.336584 | 4.240201 | OK | 0 | 144.1704474 | 0.146236 | 0 |
| 722 | HSPB8 | 5 | 1 | 0.30416 | 3.347195 | OK | 0 | 271.3592445 | 0.275262 | 0 |
| 723 | IL1A | 5 | 1 | 1.546709 | 4.849163 | OK | 0 | 257.9598712 | 0.261593 | 0 |
| 724 | IL36A | 5 | 1 | 1.322486 | 4.879153 | OK | 0 | 602.4112865 | 0.610955 | 0 |
| 725 | IL36RN | 5 | 1 | 0.264174 | 2.664308 | OK | 0 | 221.0506508 | 0.224128 | 0 |
| 726 | ITSN2 | 5 | 1 | -0.66939 | 3.280333 | OK | 0 | 203.8113867 | 0.206742 | 0 |
| 727 | KATNBL1 | 5 | 1 | 1.925369 | 3.441823 | OK | 0 | 164.5362015 | 0.1669 | 0 |
| 728 | KAZN | 5 | 1 | 1.359249 | 3.553072 | OK | 0 | 247.0795528 | 0.250635 | 0 |
| 729 | KIFC3 | 5 | 1 | 1.571456 | 4.702668 | OK | 0 | 263.9422913 | 0.267695 | 0 |
| 730 | KRT23 | 5 | 1 | 0.871363 | 2.97825 | OK | 0 | 194.875347 | 0.19761 | 0 |
| 731 | KRT7 | 5 | 1 | 1.646416 | 2.87513 | OK | 0 | 200.9638865 | 0.203845 | 0 |
| 732 | KRT80 | 5 | 1 | -0.29523 | 2.774517 | OK | 0 | 433.1171235 | 0.439351 | 0 |
| 733 | LEXM | 5 | 1 | 1.272289 | 3.287549 | OK | 0 | 311.254817 | 0.315634 | 0 |
| 734 | LGALS3 | 5 | 1 | 2.16017 | 4.488221 | OK | 0 | 222.8252583 | 0.226024 | 0 |
| 735 | LINC02303 | 5 | 1 | 2.070167 | 3.185497 | OK | 0 | 360.8859439 | 0.365871 | 0 |
| 736 | LIPE-AS1 | 5 | 1 | 2.196977 | 3.699196 | OK | 0 | 105.8625952 | 0.107358 | 0 |
| 737 | LMO2 | 5 | 1 | 0.924346 | 2.913144 | OK | 0 | 290.3298077 | 0.294458 | 0 |
| 738 | LMO7 | 5 | 1 | -0.66431 | 3.485337 | OK | 0 | 335.8673351 | 0.340707 | 0 |
| 739 | LPIN2 | 5 | 1 | 0.891192 | 4.190341 | OK | 0 | 253.9798394 | 0.257589 | 0 |
| 740 | LRG1 | 5 | 1 | 0.588316 | 2.907019 | OK | 0 | 63.62816472 | 0.064533 | 0 |
| 741 | LY6K | 5 | 1 | 2.176342 | 4.376997 | OK | 0 | 109.3602321 | 0.110926 | 0 |
| 742 | MAB21L4 | 5 | 1 | 1.804627 | 2.646532 | OK | 0 | 455.2922248 | 0.461857 | 0 |
| 743 | MACC1 | 5 | 1 | -0.33058 | 3.607851 | OK | 0 | 166.5992575 | 0.168991 | 0 |
| 744 | MALL | 5 | 1 | -0.33501 | 3.737777 | OK | 0 | 505.074935 | 0.512359 | 0 |
| 745 | MAP1LC3A | 5 | 1 | 2.415849 | 4.099238 | OK | 0 | 233.0291241 | 0.236371 | 0 |
| 746 | MAPK3 | 5 | 1 | 1.143363 | 3.356911 | OK | 0 | 183.2077446 | 0.18584 | 0 |
| 747 | MAPK8 | 5 | 1 | 1.438809 | 4.940138 | OK | 0 | 77.64106605 | 0.078747 | 0 |
| 748 | 5-Mar | 5 | 1 | 2.121731 | 4.037055 | OK | 0 | 186.8426922 | 0.189529 | 0 |
| 749 | MDFIC | 5 | 1 | 1.650656 | 3.576622 | OK | 0 | 93.29667592 | 0.09463 | 0 |
| 750 | METRNL | 5 | 1 | 1.661726 | 3.070854 | OK | 0 | 465.5651301 | 0.472278 | 0 |
| 751 | MGAT1 | 5 | 1 | 1.345755 | 3.920425 | OK | 0 | 152.810058 | 0.155004 | 0 |
| 752 | MPRIP | 5 | 1 | 1.50643 | 4.165703 | OK | 0 | 145.1902765 | 0.147274 | 0 |
| 753 | MPZL3 | 5 | 1 | 0.423167 | 3.243093 | OK | 0 | 186.8303627 | 0.1895 | 0 |
| 754 | MTERF4 | 5 | 1 | 2.188768 | 3.515887 | OK | 0 | 73.91733955 | 0.074972 | 0 |
| 755 | MUC1 | 5 | 1 | 0.063736 | 3.389034 | OK | 0 | 432.0390421 | 0.438262 | 0 |
| 756 | MUC20 | 5 | 1 | 1.590833 | 2.764943 | OK | 0 | 325.1346702 | 0.329816 | 0 |
| 757 | MUC22 | 5 | 1 | 1.379609 | 4.635846 | OK | 0 | 304.9171991 | 0.309142 | 0 |
| 758 | MXD1 | 5 | 1 | 0.794135 | 4.421854 | OK | 0 | 351.7035145 | 0.356769 | 0 |
| 759 | MYEOV | 5 | 1 | 2.141989 | 3.434928 | OK | 0 | 254.0325606 | 0.25768 | 0 |
| 760 | MYL12A | 5 | 1 | 2.239586 | 3.458912 | OK | 0 | 297.2749322 | 0.301554 | 0 |
| 761 | MYO5B | 5 | 1 | -0.29807 | 2.8949 | OK | 0 | 232.9744069 | 0.236323 | 0 |
| 762 | MYO6 | 5 | 1 | -0.58755 | 4.048232 | OK | 0 | 214.7233112 | 0.217812 | 0 |
| 763 | NAMPT | 5 | 1 | -0.50679 | 3.149124 | OK | 0 | 299.2756443 | 0.303587 | 0 |
| 764 | NAPRT | 5 | 1 | 1.497659 | 3.194599 | OK | 0 | 262.3857703 | 0.266164 | 0 |
| 765 | NCCRP1 | 5 | 1 | 1.198091 | 2.270749 | OK | 0 | 630.7482515 | 0.639842 | 0 |
| 766 | NDRG2 | 5 | 1 | 0.029267 | 3.900654 | OK | 0 | 271.247373 | 0.275152 | 0 |
| 767 | NIBAN2 | 5 | 1 | 1.78778 | 2.769645 | OK | 0 | 359.8871766 | 0.365074 | 0 |
| 768 | NT5C2 | 5 | 1 | -0.06407 | 3.433455 | OK | 0 | 191.4711457 | 0.194223 | 0 |
| 769 | PCBP1-AS1 | 5 | 1 | 2.084336 | 3.004049 | OK | 0 | 92.19801988 | 0.093511 | 0 |
| 770 | PDCD6 | 5 | 1 | 2.024525 | 3.328223 | OK | 0 | 280.1072271 | 0.28414 | 0 |
| 771 | PHACTR4 | 5 | 1 | -0.50647 | 3.799765 | OK | 0 | 159.9734864 | 0.162272 | 0 |
| 772 | PHLDA1 | 5 | 1 | 0.952496 | 4.285964 | OK | 0 | 463.6616327 | 0.47033 | 0 |
| 773 | PLEKHM1 | 5 | 1 | 0.317324 | 3.406489 | OK | 0 | 113.0244293 | 0.114641 | 0 |
| 774 | PLIN3 | 5 | 1 | 1.379777 | 4.410369 | OK | 0 | 177.4883491 | 0.18004 | 0 |
| 775 | PLSCR1 | 5 | 1 | 1.587247 | 3.427526 | OK | 0 | 141.7868003 | 0.143822 | 0 |
| 776 | PORCN | 5 | 1 | 1.56907 | 4.42071 | OK | 0 | 63.14920094 | 0.064041 | 0 |
| 777 | PPDPF | 5 | 1 | 2.374848 | 3.995792 | OK | 0 | 187.5298512 | 0.19022 | 0 |
| 778 | PPP1R14B- | 5 | 1 | 1.82749 | 2.408816 | OK | 0 | 171.1926783 | 0.173651 | 0 |

| | | | | | | | | | | |
|---|---|---|---|---|---|---|---|---|---|---|
| 779 | PRDM1 | 5 | 1 | -0.24971 | 2.779587 | OK | 0 | 346.0109725 | 0.35099 | 0 |
| 780 | PRSS22 | 5 | 1 | 0.868889 | 2.496705 | OK | 0 | 284.1912347 | 0.288283 | 0 |
| 781 | PRSS27 | 5 | 1 | 1.380203 | 2.34829 | OK | 0 | 644.7511215 | 0.654044 | 0 |
| 782 | PRSS8 | 5 | 1 | 0.802625 | 3.794003 | OK | 0 | 116.5088726 | 0.118179 | 0 |
| 783 | PSCA | 5 | 1 | 2.35008 | 4.001967 | OK | 0 | 578.3721199 | 0.586677 | 0 |
| 784 | PTK6 | 5 | 1 | 0.200677 | 3.203274 | OK | 0 | 341.5986134 | 0.346519 | 0 |
| 785 | PWWP2B | 5 | 1 | 1.057336 | 3.463561 | OK | 0 | 82.89897147 | 0.084081 | 0 |
| 786 | RAB5IF | 5 | 1 | 2.120733 | 2.949411 | OK | 0 | 356.1051098 | 0.361236 | 0 |
| 787 | RAB9A | 5 | 1 | 2.194157 | 4.534154 | OK | 0 | 166.2163694 | 0.168605 | 0 |
| 788 | RANBP9 | 5 | 1 | -0.24236 | 3.762839 | OK | 0 | 202.9141436 | 0.205831 | 0 |
| 789 | RFK | 5 | 1 | 1.717212 | 4.748087 | OK | 0 | 120.8421224 | 0.122573 | 0 |
| 790 | RHOF | 5 | 1 | 1.457303 | 3.943709 | OK | 0 | 222.8249627 | 0.225977 | 0 |
| 791 | RNF223 | 5 | 1 | 1.050098 | 4.393172 | OK | 0 | 340.3687653 | 0.345252 | 0 |
| 792 | RNF39 | 5 | 1 | 1.393121 | 4.611402 | OK | 0 | 119.0689491 | 0.120769 | 0 |
| 793 | S100P | 5 | 1 | 0.621184 | 2.987616 | OK | 0 | 484.7854691 | 0.491768 | 0 |
| 794 | SAMD9 | 5 | 1 | 0.108711 | 3.801657 | OK | 0 | 311.5740276 | 0.316063 | 0 |
| 795 | SAMD9L | 5 | 1 | 0.061001 | 3.980484 | OK | 0 | 49.42289159 | 0.050122 | 0 |
| 796 | SAT1 | 5 | 1 | 1.854859 | 2.707714 | OK | 0 | 187.8987578 | 0.190599 | 0 |
| 797 | SCEL | 5 | 1 | -0.19999 | 3.733634 | OK | 0 | 581.4354464 | 0.589823 | 0 |
| 798 | SCNN1A | 5 | 1 | -0.00666 | 3.139956 | OK | 0 | 258.2221932 | 0.261939 | 0 |
| 799 | SDCBP2 | 5 | 1 | 2.066228 | 3.755651 | OK | 0 | 422.0514287 | 0.428135 | 0 |
| 800 | SDHA | 5 | 1 | 1.821548 | 3.494744 | OK | 0 | 136.0052107 | 0.137957 | 0 |
| 801 | SEC24C | 5 | 1 | 0.61069 | 3.386855 | OK | 0 | 88.05993781 | 0.089317 | 0 |
| 802 | SERINC2 | 5 | 1 | 1.80268 | 3.990841 | OK | 0 | 193.086298 | 0.195862 | 0 |
| 803 | SERPINB13 | 5 | 1 | -0.93278 | 2.497324 | OK | 0 | 412.3513152 | 0.418295 | 0 |
| 804 | SGTA | 5 | 1 | 1.079404 | 3.86509 | OK | 0 | 71.36433426 | 0.072381 | 0 |
| 805 | SH3BGRL2 | 5 | 1 | 0.390353 | 4.000653 | OK | 0 | 472.1057588 | 0.478901 | 0 |
| 806 | SLC16A3 | 5 | 1 | 1.170061 | 3.080048 | OK | 0 | 192.9056967 | 0.195649 | 0 |
| 807 | SLC25A23 | 5 | 1 | 0.346444 | 3.508048 | OK | 0 | 179.6251488 | 0.182207 | 0 |
| 808 | SLC9A3R2 | 5 | 1 | 2.348631 | 4.31608 | OK | 0 | 84.21810082 | 0.085418 | 0 |
| 809 | SLK | 5 | 1 | -0.81044 | 3.252818 | OK | 0 | 321.1278672 | 0.325755 | 0 |
| 810 | SLURP2 | 5 | 1 | 1.894557 | 2.564852 | OK | 0 | 602.1832332 | 0.610864 | 0 |
| 811 | SMIM22 | 5 | 1 | 1.180317 | 2.798591 | OK | 0 | 84.16330774 | 0.085354 | 0 |
| 812 | SNCG | 5 | 1 | 2.554073 | 3.84925 | OK | 0 | 289.5755367 | 0.293724 | 0 |
| 813 | SNX24 | 5 | 1 | 2.420009 | 3.822438 | OK | 0 | 207.5905829 | 0.210576 | 0 |
| 814 | SNX8 | 5 | 1 | 1.358911 | 3.326992 | OK | 0 | 97.14405305 | 0.098528 | 0 |
| 815 | SORBS2 | 5 | 1 | 0.783658 | 2.88426 | OK | 0 | 61.31761275 | 0.062186 | 0 |
| 816 | SORT1 | 5 | 1 | -0.33217 | 2.948587 | OK | 0 | 431.0424572 | 0.437252 | 0 |
| 817 | SPAG1 | 5 | 1 | 0.931965 | 3.899994 | OK | 0 | 231.9243405 | 0.235246 | 0 |
| 818 | SPECC1 | 5 | 1 | 1.182206 | 4.404376 | OK | 0 | 215.8631206 | 0.218955 | 0 |
| 819 | SPINT1 | 5 | 1 | 1.243959 | 3.191832 | OK | 0 | 258.9356485 | 0.262664 | 0 |
| 820 | SPNS2 | 5 | 1 | 1.044174 | 4.720157 | OK | 0 | 761.2272175 | 0.772107 | 0 |
| 821 | SPRR2B | 5 | 1 | 0.264666 | 2.428786 | OK | 0 | 93.96744768 | 0.095105 | 0 |
| 822 | SPSB3 | 5 | 1 | 2.275677 | 4.124184 | OK | 0 | 134.1963623 | 0.136122 | 0 |
| 823 | SRGAP1 | 5 | 1 | 0.717617 | 3.138111 | OK | 0 | 162.3382386 | 0.164657 | 0 |
| 824 | SRSF5 | 5 | 1 | 1.515917 | 3.191329 | OK | 0 | 263.4900155 | 0.267284 | 0 |
| 825 | STK39 | 5 | 1 | -0.45692 | 2.962277 | OK | 0 | 272.3405255 | 0.27626 | 0 |
| 826 | STRADB | 5 | 1 | 1.968378 | 4.183064 | OK | 0 | 105.2190489 | 0.106721 | 0 |
| 827 | SYAP1 | 5 | 1 | -0.46349 | 3.983084 | OK | 0 | 161.8119811 | 0.164137 | 0 |
| 828 | TAX1BP1 | 5 | 1 | -0.75498 | 3.994061 | OK | 0 | 344.2932369 | 0.349254 | 0 |
| 829 | TGFA | 5 | 1 | 0.100981 | 3.936362 | OK | 0 | 102.6043849 | 0.104069 | 0 |
| 830 | TGFB1 | 5 | 1 | 0.698172 | 2.55398 | OK | 0 | 130.7056169 | 0.132578 | 0 |
| 831 | TIMP2 | 5 | 1 | 2.07942 | 4.107532 | OK | 0 | 764.7726239 | 0.775709 | 0 |
| 832 | TJP1 | 5 | 1 | -0.87765 | 3.114618 | OK | 0 | 339.8626254 | 0.34476 | 0 |
| 833 | TJP3 | 5 | 1 | 0.947805 | 2.869461 | OK | 0 | 221.3523421 | 0.224532 | 0 |
| 834 | TMEM127 | 5 | 1 | 0.665525 | 3.56113 | OK | 0 | 103.4900656 | 0.10497 | 0 |
| 835 | TMEM40 | 5 | 1 | 2.52876 | 3.960896 | OK | 0 | 485.0987622 | 0.492094 | 0 |
| 836 | TMEM8A | 5 | 1 | 1.090903 | 3.311615 | OK | 0 | 104.2874945 | 0.105775 | 0 |
| 837 | TMPRSS11 | 5 | 1 | -0.06506 | 2.814024 | OK | 0 | 302.1537244 | 0.306503 | 0 |
| 838 | TMPRSS11 | 5 | 1 | 0.417922 | 3.994252 | OK | 0 | 634.0384728 | 0.643171 | 0 |

| | | | | | | | | | | |
|---|---|---|---|---|---|---|---|---|---|---|
| 839 | TMPRSS2 | 5 | 1 | 0.472113 | 4.025514 | OK | 0 | 389.1246995 | 0.394696 | 0 |
| 840 | TP53INP2 | 5 | 1 | 0.531974 | 3.888636 | OK | 0 | 304.4155496 | 0.308773 | 0 |
| 841 | TRIP10 | 5 | 1 | 0.530346 | 3.708187 | OK | 0 | 328.0358406 | 0.33276 | 0 |
| 842 | TSPAN6 | 5 | 1 | 1.738747 | 2.806931 | OK | 0 | 263.4586299 | 0.26725 | 0 |
| 843 | TTC22 | 5 | 1 | 0.462347 | 3.264755 | OK | 0 | 177.7604196 | 0.180313 | 0 |
| 844 | TTC9 | 5 | 1 | 0.878832 | 3.969417 | OK | 0 | 435.4616317 | 0.441737 | 0 |
| 845 | TTYH1 | 5 | 1 | 1.150912 | 3.766236 | OK | 0 | 344.0595602 | 0.34894 | 0 |
| 846 | UACA | 5 | 1 | -0.84764 | 3.453666 | OK | 0 | 191.9155807 | 0.194675 | 0 |
| 847 | UBE2D1 | 5 | 1 | 2.084018 | 3.717723 | OK | 0 | 123.5960563 | 0.125368 | 0 |
| 848 | UBL3 | 5 | 1 | -0.26175 | 3.669374 | OK | 0 | 216.1318563 | 0.219241 | 0 |
| 849 | UPK3B | 5 | 1 | 2.369172 | 3.07044 | OK | 0 | 434.2684569 | 0.440492 | 0 |
| 850 | VASN | 5 | 1 | 0.761317 | 3.646885 | OK | 0 | 306.1249153 | 0.310486 | 0 |
| 851 | VSIG10 | 5 | 1 | 1.646806 | 3.884756 | OK | 0 | 101.6243571 | 0.103076 | 0 |
| 852 | VSIG10L | 5 | 1 | 0.533208 | 3.115912 | OK | 0 | 408.1557926 | 0.414023 | 0 |
| 853 | VSIG2 | 5 | 1 | 2.261975 | 3.558542 | OK | 0 | 609.2245601 | 0.617912 | 0 |
| 854 | WDR26 | 5 | 1 | -0.37237 | 3.645668 | OK | 0 | 158.0215038 | 0.16029 | 0 |
| 855 | WDR45B | 5 | 1 | -0.68458 | 3.321651 | OK | 0 | 179.6375217 | 0.18222 | 0 |
| 856 | YPEL3 | 5 | 1 | 2.103713 | 4.032508 | OK | 0 | 169.9387242 | 0.172381 | 0 |
| 857 | ZFAND2A | 5 | 1 | 2.303498 | 4.45271 | OK | 7.89E-303 | 37.17746599 | 0.037701 | 8.00E-303 |
| 858 | ZFAND5 | 5 | 1 | 0.997223 | 4.390746 | OK | 0 | 220.1789921 | 0.223347 | 0 |
| 859 | ZNF101 | 5 | 1 | 2.071027 | 4.206877 | OK | 0 | 91.93897505 | 0.093244 | 0 |
| 860 | ZNF185 | 5 | 1 | -0.11033 | 3.275476 | OK | 0 | 389.9671791 | 0.395588 | 0 |
| 861 | ZNRF1 | 5 | 1 | 2.361413 | 4.01704 | OK | 0 | 218.7315085 | 0.221879 | 0 |
| 862 | AC023157. | 6 | 1 | 0.654477 | 5.484818 | OK | 0 | 104.1856252 | 0.105675 | 0 |
| 863 | ACOX1 | 6 | 1 | -0.45764 | 4.56818 | OK | 0 | 217.4989344 | 0.220627 | 0 |
| 864 | ADIPOR2 | 6 | 1 | 0.631732 | 4.765007 | OK | 0 | 179.564683 | 0.182145 | 0 |
| 865 | AMOTL2 | 6 | 1 | -0.26919 | 4.415826 | OK | 0 | 69.07158248 | 0.070055 | 0 |
| 866 | ANKRD13A | 6 | 1 | -0.16876 | 4.462662 | OK | 0 | 113.9198441 | 0.115552 | 0 |
| 867 | ANKRD37 | 6 | 1 | 0.94784 | 5.604868 | OK | 0 | 167.7151696 | 0.170125 | 0 |
| 868 | ARHGAP10 | 6 | 1 | 0.234525 | 4.402782 | OK | 0 | 207.7814854 | 0.210769 | 0 |
| 869 | ARL4C | 6 | 1 | 0.452183 | 4.74883 | OK | 0 | 180.6920978 | 0.183289 | 0 |
| 870 | ASCC2 | 6 | 1 | 0.639094 | 5.42503 | OK | 0 | 123.0272127 | 0.124791 | 0 |
| 871 | ATG9B | 6 | 1 | 0.574758 | 4.585546 | OK | 0 | 537.9230503 | 0.545572 | 0 |
| 872 | BPGM | 6 | 1 | 0.162323 | 5.505266 | OK | 0 | 48.0966126 | 0.048777 | 0 |
| 873 | C18orf25 | 6 | 1 | 0.468652 | 5.148888 | OK | 0 | 201.5397077 | 0.204433 | 0 |
| 874 | CDKL5 | 6 | 1 | -0.20618 | 4.554295 | OK | 0 | 91.96738186 | 0.093275 | 0 |
| 875 | CFLAR | 6 | 1 | 0.583019 | 5.199415 | OK | 0 | 142.281627 | 0.144324 | 0 |
| 876 | CGNL1 | 6 | 1 | 0.820019 | 4.883241 | OK | 0 | 545.4976824 | 0.553242 | 0 |
| 877 | CHAC1 | 6 | 1 | -0.01431 | 5.09707 | OK | 0 | 280.486403 | 0.284308 | 0 |
| 878 | CITED2 | 6 | 1 | 0.685853 | 5.244548 | OK | 0 | 129.654138 | 0.131514 | 0 |
| 879 | CNST | 6 | 1 | 0.580458 | 5.115193 | OK | 0 | 117.2043621 | 0.118879 | 0 |
| 880 | CSNK1E | 6 | 1 | 0.114267 | 4.935921 | OK | 0 | 129.6477081 | 0.131506 | 0 |
| 881 | DKK1 | 6 | 1 | 0.65997 | 5.518457 | OK | 0 | 336.7215333 | 0.341411 | 0 |
| 882 | DUSP14 | 6 | 1 | 0.223446 | 5.654048 | OK | 0 | 102.7899874 | 0.104261 | 0 |
| 883 | DUSP5 | 6 | 1 | 0.543382 | 5.326819 | OK | 0 | 331.9651945 | 0.336738 | 0 |
| 884 | ERO1A | 6 | 1 | 0.795767 | 4.57673 | OK | 0 | 347.3011844 | 0.352302 | 0 |
| 885 | FAM214A | 6 | 1 | 0.253163 | 4.863839 | OK | 0 | 191.4079091 | 0.194155 | 0 |
| 886 | FLG | 6 | 1 | 0.21221 | 5.657109 | OK | 0 | 325.6668067 | 0.330273 | 0 |
| 887 | GNE | 6 | 1 | -0.18452 | 4.501211 | OK | 0 | 98.67681552 | 0.100075 | 0 |
| 888 | GPAT3 | 6 | 1 | -0.10053 | 4.944178 | OK | 0 | 312.9460905 | 0.317343 | 0 |
| 889 | GRPEL2 | 6 | 1 | 0.849469 | 5.273012 | OK | 0 | 311.9734578 | 0.316444 | 0 |
| 890 | HCG22 | 6 | 1 | 0.667227 | 5.208042 | OK | 0 | 389.00955 | 0.394543 | 0 |
| 891 | HILPDA | 6 | 1 | 0.87843 | 5.501628 | OK | 0 | 250.1113345 | 0.253702 | 0 |
| 892 | IL1RN | 6 | 1 | 0.58372 | 5.044149 | OK | 0 | 450.6357138 | 0.457133 | 0 |
| 893 | ITGB8 | 6 | 1 | -0.11459 | 4.204758 | OK | 0 | 136.1836871 | 0.138138 | 0 |
| 894 | KPRP | 6 | 1 | 0.71485 | 5.559824 | OK | 0 | 276.705257 | 0.280506 | 0 |
| 895 | KRT16 | 6 | 1 | -0.2535 | 5.099516 | OK | 0 | 250.291273 | 0.253895 | 0 |
| 896 | KRTAP3-2 | 6 | 1 | 1.176404 | 6.71268 | OK | 0 | 183.2759151 | 0.185607 | 0 |
| 897 | LNX1 | 6 | 1 | 0.6567 | 4.788741 | OK | 0 | 185.9705633 | 0.188637 | 0 |
| 898 | LPIN1 | 6 | 1 | 0.49636 | 4.964568 | OK | 0 | 466.4110305 | 0.473084 | 0 |

| | | | | | | | | | | |
|---|---|---|---|---|---|---|---|---|---|---|
| 899 | MACO1 | 6 | 1 | 0.359179 | 4.90566 | OK | 0 | 114.9008094 | 0.116547 | 0 |
| 900 | MCU | 6 | 1 | 0.35528 | 4.623194 | OK | 0 | 116.4249278 | 0.118089 | 0 |
| 901 | MT1A | 6 | 1 | 1.203329 | 6.768424 | OK | 0 | 76.09390949 | 0.07693 | 0 |
| 902 | MT1E | 6 | 1 | 1.213834 | 6.769074 | OK | 0 | 390.9570623 | 0.396593 | 0 |
| 903 | MT1F | 6 | 1 | 1.210291 | 6.774556 | OK | 0 | 72.21828667 | 0.07324 | 0 |
| 904 | MT1G | 6 | 1 | 1.228783 | 6.795896 | OK | 0 | 285.1050767 | 0.2892 | 0 |
| 905 | MT1H | 6 | 1 | 1.194084 | 6.752085 | OK | 0 | 279.0143532 | 0.282987 | 0 |
| 906 | MT1M | 6 | 1 | 1.237857 | 6.802794 | OK | 0 | 161.0983657 | 0.16339 | 0 |
| 907 | MT2A | 6 | 1 | 1.250069 | 6.821646 | OK | 0 | 466.5079858 | 0.473235 | 0 |
| 908 | MYZAP | 6 | 1 | 0.263878 | 4.836772 | OK | 0 | 320.8284676 | 0.32542 | 0 |
| 909 | NABP1 | 6 | 1 | 0.434566 | 5.238608 | OK | 0 | 127.0786783 | 0.1289 | 0 |
| 910 | NEDD4L | 6 | 1 | -0.07737 | 4.443242 | OK | 0 | 82.95396745 | 0.084137 | 0 |
| 911 | NR1D1 | 6 | 1 | 1.081408 | 6.453282 | OK | 0 | 96.46358663 | 0.097841 | 0 |
| 912 | OXSR1 | 6 | 1 | -0.25546 | 4.587452 | OK | 0 | 137.6766898 | 0.139652 | 0 |
| 913 | PADI1 | 6 | 1 | 0.139826 | 4.762465 | OK | 0 | 435.3303383 | 0.441577 | 0 |
| 914 | PARD6B | 6 | 1 | -0.1064 | 4.56817 | OK | 0 | 138.7384022 | 0.140726 | 0 |
| 915 | PDP1 | 6 | 1 | -0.10933 | 4.360188 | OK | 0 | 65.35033446 | 0.06628 | 0 |
| 916 | PDZD2 | 6 | 1 | -0.12244 | 4.343046 | OK | 0 | 78.1201569 | 0.079233 | 0 |
| 917 | PELI1 | 6 | 1 | -0.10783 | 4.880102 | OK | 0 | 150.4246262 | 0.152583 | 0 |
| 918 | PHACTR2 | 6 | 1 | -0.07845 | 4.995728 | OK | 0 | 203.3841078 | 0.206309 | 0 |
| 919 | PPL | 6 | 1 | -0.50209 | 4.450413 | OK | 0 | 426.5544047 | 0.432704 | 0 |
| 920 | PTPN3 | 6 | 1 | -0.28792 | 4.71844 | OK | 0 | 81.25068811 | 0.08241 | 0 |
| 921 | QSOX1 | 6 | 1 | 0.291849 | 5.440802 | OK | 0 | 294.215965 | 0.298448 | 0 |
| 922 | RASGEF1B | 6 | 1 | 0.600836 | 5.473722 | OK | 0 | 135.6035204 | 0.137539 | 0 |
| 923 | RIOK3 | 6 | 1 | -0.62499 | 4.121132 | OK | 0 | 330.2011143 | 0.334958 | 0 |
| 924 | RMND5A | 6 | 1 | 0.270898 | 4.99914 | OK | 0 | 151.8914784 | 0.154072 | 0 |
| 925 | RNASE7 | 6 | 1 | 0.201578 | 5.575142 | OK | 0 | 602.393195 | 0.610916 | 0 |
| 926 | RNF169 | 6 | 1 | 0.384669 | 4.670601 | OK | 0 | 131.59774 | 0.133485 | 0 |
| 927 | SASH1 | 6 | 1 | -0.40658 | 4.257826 | OK | 0 | 231.7463291 | 0.235081 | 0 |
| 928 | SEC14L1 | 6 | 1 | 0.381333 | 4.559777 | OK | 0 | 122.4094454 | 0.124162 | 0 |
| 929 | SECISBP2L | 6 | 1 | -0.36416 | 4.571112 | OK | 0 | 174.6378579 | 0.177145 | 0 |
| 930 | SESN2 | 6 | 1 | 0.709718 | 5.434245 | OK | 0 | 176.021871 | 0.178534 | 0 |
| 931 | SH3D19 | 6 | 1 | -0.3114 | 4.144282 | OK | 0 | 99.98079513 | 0.101411 | 0 |
| 932 | SLC20A2 | 6 | 1 | -0.28897 | 4.951072 | OK | 0 | 112.7618792 | 0.114377 | 0 |
| 933 | SNX9 | 6 | 1 | 0.589619 | 5.014684 | OK | 0 | 213.368937 | 0.216437 | 0 |
| 934 | SYNPO2L | 6 | 1 | 0.628094 | 5.540041 | OK | 0 | 228.9929963 | 0.232066 | 0 |
| 935 | TCP11L2 | 6 | 1 | 0.793559 | 4.927032 | OK | 0 | 275.0539928 | 0.279 | 0 |
| 936 | TEAD1 | 6 | 1 | -0.36468 | 4.406013 | OK | 0 | 137.5743625 | 0.139547 | 0 |
| 937 | TICAM1 | 6 | 1 | 0.624626 | 4.888031 | OK | 0 | 157.9724732 | 0.160237 | 0 |
| 938 | TMOD3 | 6 | 1 | -0.64278 | 4.180426 | OK | 0 | 357.3484758 | 0.362498 | 0 |
| 939 | TNIP1 | 6 | 1 | -0.39878 | 4.572251 | OK | 0 | 186.0050496 | 0.188679 | 0 |
| 940 | TP53BP2 | 6 | 1 | 0.026254 | 4.778004 | OK | 0 | 106.268608 | 0.107789 | 0 |
| 941 | TRIM2 | 6 | 1 | 0.020364 | 4.50175 | OK | 0 | 176.6294491 | 0.179154 | 0 |
| 942 | TUFT1 | 6 | 1 | -0.25342 | 4.955034 | OK | 0 | 107.4780284 | 0.109016 | 0 |
| 943 | UBE2G1 | 6 | 1 | 0.755913 | 4.739602 | OK | 0 | 158.3813099 | 0.160656 | 0 |
| 944 | UPP1 | 6 | 1 | 0.987655 | 5.478302 | OK | 0 | 276.4968601 | 0.280478 | 0 |
| 945 | USP53 | 6 | 1 | -0.2943 | 4.19818 | OK | 0 | 110.5725745 | 0.112156 | 0 |
| 946 | USP6NL | 6 | 1 | -0.14036 | 4.18733 | OK | 0 | 222.892655 | 0.226096 | 0 |
| 947 | VPS37B | 6 | 1 | -0.27223 | 4.80607 | OK | 0 | 195.324401 | 0.198132 | 0 |
| 948 | WWC1 | 6 | 1 | -0.35907 | 4.546819 | OK | 0 | 144.8576233 | 0.146932 | 0 |
| 949 | YOD1 | 6 | 1 | 0.466948 | 5.4359 | OK | 0 | 269.6401227 | 0.27351 | 0 |
| 950 | ZNF365 | 6 | 1 | 0.819224 | 5.448735 | OK | 0 | 540.6312193 | 0.548272 | 0 |
| 951 | ZNF426 | 6 | 1 | 0.16132 | 5.103724 | OK | 0 | 153.103905 | 0.155296 | 0 |
| 952 | ZNF430 | 6 | 1 | 0.226499 | 5.644871 | OK | 0 | 64.31879868 | 0.065234 | 0 |
| 953 | ZNF431 | 6 | 1 | 0.287737 | 5.576567 | OK | 0 | 177.384566 | 0.179932 | 0 |
| 954 | ZNF714 | 6 | 1 | 0.213168 | 5.632364 | OK | 0 | 66.0553414 | 0.066993 | 0 |
| 955 | ZNF92 | 6 | 1 | 0.212482 | 5.639435 | OK | 0 | 78.66965378 | 0.079788 | 0 |
| 956 | ALOX15 | 7 | 1 | -0.32891 | -0.23817 | OK | 0 | 434.6651069 | 0.44093 | 0 |
| 957 | ANO1 | 7 | 1 | -0.14253 | -0.50269 | OK | 0 | 320.4936555 | 0.325111 | 0 |
| 958 | APOL1 | 7 | 1 | -0.09545 | -1.11637 | OK | 0 | 136.483189 | 0.13844 | 0 |

| | | | | | | | | | | |
|---|---|---|---|---|---|---|---|---|---|---|
| 959 | ARNTL2 | 7 | 1 | -0.19849 | -0.43845 | OK | 0 | 264.2569026 | 0.268061 | 0 |
| 960 | ATP13A5 | 7 | 1 | -0.51693 | -0.85023 | OK | 0 | 130.64077 | 0.132511 | 0 |
| 961 | ATP8B1 | 7 | 1 | 0.089923 | -0.57546 | OK | 0 | 230.211679 | 0.233524 | 0 |
| 962 | BCL2L15 | 7 | 1 | -0.49916 | -0.51159 | OK | 0 | 219.2263121 | 0.222379 | 0 |
| 963 | C12orf75 | 7 | 1 | -0.56691 | -1.34929 | OK | 0 | 160.5192575 | 0.162825 | 0 |
| 964 | CA2 | 7 | 1 | -0.10917 | -1.40183 | OK | 0 | 236.2899464 | 0.239691 | 0 |
| 965 | CADM1 | 7 | 1 | -0.27633 | -0.56155 | OK | 0 | 118.3391683 | 0.120033 | 0 |
| 966 | CAMK1D | 7 | 1 | -0.07629 | -0.47949 | OK | 0 | 217.4202527 | 0.220547 | 0 |
| 967 | CCL26 | 7 | 1 | -0.31755 | -0.54385 | OK | 0 | 302.0231403 | 0.306374 | 0 |
| 968 | CDH3 | 7 | 1 | -0.30856 | -1.40471 | OK | 0 | 180.9037785 | 0.183503 | 0 |
| 969 | CES2 | 7 | 1 | 0.44769 | -0.48062 | OK | 0 | 470.9342532 | 0.477725 | 0 |
| 970 | CHL1 | 7 | 1 | -0.38052 | -0.41476 | OK | 0 | 237.1249703 | 0.240535 | 0 |
| 971 | CISH | 7 | 1 | -0.40716 | -1.21551 | OK | 0 | 69.58338323 | 0.070575 | 0 |
| 972 | CLDN1 | 7 | 1 | 0.034547 | -0.95149 | OK | 0 | 201.4444818 | 0.204342 | 0 |
| 973 | CTSC | 7 | 1 | -0.54689 | -1.06939 | OK | 0 | 293.4937114 | 0.297721 | 0 |
| 974 | CYP2S1 | 7 | 1 | -0.07855 | -0.88485 | OK | 0 | 150.6675981 | 0.152829 | 0 |
| 975 | CYP7B1 | 7 | 1 | -0.4931 | -0.6963 | OK | 0 | 294.3818033 | 0.298622 | 0 |
| 976 | DPYD | 7 | 1 | -0.29173 | -0.409 | OK | 0 | 253.6263682 | 0.257277 | 0 |
| 977 | DSG3 | 7 | 1 | 0.633453 | -0.60171 | OK | 0 | 372.6045303 | 0.377972 | 0 |
| 978 | DSP | 7 | 1 | 0.566945 | -0.72709 | OK | 0 | 236.2433509 | 0.239636 | 0 |
| 979 | DUOX1 | 7 | 1 | 0.201838 | -0.24161 | OK | 0 | 326.2683787 | 0.330969 | 0 |
| 980 | EPPK1 | 7 | 1 | -0.34352 | -0.37426 | OK | 0 | 266.1956611 | 0.270024 | 0 |
| 981 | GALNT5 | 7 | 1 | -0.06889 | -0.77017 | OK | 0 | 214.1991008 | 0.21728 | 0 |
| 982 | HAS3 | 7 | 1 | -0.10233 | -0.90178 | OK | 0 | 143.95017 | 0.146014 | 0 |
| 983 | HLF | 7 | 1 | -0.34972 | -1.28568 | OK | 0 | 174.4748259 | 0.176982 | 0 |
| 984 | HPGD | 7 | 1 | -0.61451 | -0.86886 | OK | 0 | 354.9179683 | 0.360032 | 0 |
| 985 | KCNJ2 | 7 | 1 | -0.45649 | -1.24272 | OK | 0 | 130.4389326 | 0.13231 | 0 |
| 986 | LINC01497 | 7 | 1 | -0.58644 | -1.01596 | OK | 0 | 193.505593 | 0.196287 | 0 |
| 987 | LTA4H | 7 | 1 | 0.05077 | -0.24902 | OK | 0 | 348.1073431 | 0.353124 | 0 |
| 988 | MDK | 7 | 1 | -0.54927 | -1.18398 | OK | 0 | 188.5971158 | 0.191309 | 0 |
| 989 | MPP7 | 7 | 1 | 0.16097 | -0.52633 | OK | 0 | 364.668985 | 0.369925 | 0 |
| 990 | MX1 | 7 | 1 | 0.046814 | -0.70669 | OK | 0 | 268.5388615 | 0.272406 | 0 |
| 991 | MYH9 | 7 | 1 | 0.247705 | -0.85357 | OK | 0 | 257.7729451 | 0.261484 | 0 |
| 992 | NEFL | 7 | 1 | -0.29065 | -0.31323 | OK | 0 | 177.4029801 | 0.179948 | 0 |
| 993 | NTRK2 | 7 | 1 | -0.44555 | -1.27409 | OK | 0 | 213.3491183 | 0.216419 | 0 |
| 994 | P2RY1 | 7 | 1 | 0.238221 | -0.6107 | OK | 0 | 239.3458131 | 0.242791 | 0 |
| 995 | PDZK1IP1 | 7 | 1 | -0.33246 | 0.007324 | OK | 0 | 292.720704 | 0.296937 | 0 |
| 996 | PHLDB2 | 7 | 1 | -0.07042 | -1.44813 | OK | 0 | 99.34597926 | 0.100766 | 0 |
| 997 | PMCH | 7 | 1 | -0.59368 | -0.71376 | OK | 0 | 68.39893734 | 0.069363 | 0 |
| 998 | SAMHD1 | 7 | 1 | -0.54264 | -0.71986 | OK | 0 | 227.8986842 | 0.231177 | 0 |
| 999 | SCIN | 7 | 1 | -0.49259 | -0.70821 | OK | 0 | 469.4131307 | 0.476182 | 0 |
| 1000 | SECTM1 | 7 | 1 | -0.24964 | -0.44844 | OK | 0 | 136.333547 | 0.138287 | 0 |
| 1001 | SERPINB3 | 7 | 1 | -0.68084 | -0.75916 | OK | 0 | 480.0374551 | 0.486956 | 0 |
| 1002 | SERPINB4 | 7 | 1 | -0.6339 | -0.65821 | OK | 0 | 367.927037 | 0.37323 | 0 |
| 1003 | SERPINE2 | 7 | 1 | -0.59254 | -0.91515 | OK | 0 | 174.5971759 | 0.177105 | 0 |
| 1004 | SFRP1 | 7 | 1 | -0.55747 | -1.05463 | OK | 0 | 216.6401506 | 0.219756 | 0 |
| 1005 | SGK1 | 7 | 1 | -0.50208 | -1.14599 | OK | 0 | 252.2459678 | 0.255878 | 0 |
| 1006 | SH3RF2 | 7 | 1 | -0.07761 | -0.70377 | OK | 0 | 244.2873354 | 0.247804 | 0 |
| 1007 | SLC26A4-A | 7 | 1 | -0.39774 | -0.35791 | OK | 0 | 218.6808982 | 0.221818 | 0 |
| 1008 | SPARCL1 | 7 | 1 | -0.47782 | -1.06132 | OK | 0 | 78.91544755 | 0.080039 | 0 |
| 1009 | SPTBN1 | 7 | 1 | 0.286668 | -0.71725 | OK | 0 | 247.5870723 | 0.251151 | 0 |
| 1010 | SYNPO | 7 | 1 | -0.29803 | -1.18115 | OK | 0 | 122.6287639 | 0.124385 | 0 |
| 1011 | TFPI | 7 | 1 | -0.34551 | -1.25913 | OK | 0 | 66.9720114 | 0.067924 | 0 |
| 1012 | TFRC | 7 | 1 | 0.40516 | -0.7634 | OK | 0 | 175.4500203 | 0.177971 | 0 |
| 1013 | THBD | 7 | 1 | 0.039195 | -1.08419 | OK | 0 | 199.7149552 | 0.202586 | 0 |
| 1014 | TMPRSS11. | 7 | 1 | 0.105847 | -0.24942 | OK | 0 | 361.7085279 | 0.366921 | 0 |
| 1015 | TNFAIP6 | 7 | 1 | -0.58827 | -0.73305 | OK | 0 | 208.3303277 | 0.211322 | 0 |
| 1016 | ZFP36L1 | 7 | 1 | 0.786903 | -0.87909 | OK | 0 | 378.064425 | 0.38351 | 0 |

**Supplemental Table 4.** Gene expression profile of module 7 genes in EoE vs healthy subjects.

| | gene | meanEoE | meanHC | log2FC | status | p_value | morans_test_statistic | morans_I | q_value |
|---|---|---|---|---|---|---|---|---|---|
| 1 | TNFAIP6 | 0.966075 | 3.85E-05 | 14.61634 | OK | 0 | 208.3303277 | 0.211322 | 0 |
| 2 | ALOX15 | 1.961654 | 0.000725 | 11.4022 | OK | 0 | 434.6651069 | 0.44093 | 0 |
| 3 | PMCH | 0.202487 | 0.000257 | 9.619501 | OK | 0 | 68.39893734 | 0.069363 | 0 |
| 4 | CCL26 | 6.859651 | 0.016739 | 8.67874 | OK | 0 | 302.0231403 | 0.306374 | 0 |
| 5 | SLC26A4-A | 0.120048 | 0.000587 | 7.676937 | OK | 0 | 218.6808982 | 0.221818 | 0 |
| 6 | ATP13A5 | 0.157916 | 0.001202 | 7.03789 | OK | 0 | 130.64077 | 0.132511 | 0 |
| 7 | ANO1 | 0.836764 | 0.025489 | 5.036852 | OK | 0 | 320.4936555 | 0.325111 | 0 |
| 8 | LINC01497 | 0.661007 | 0.028314 | 4.545071 | OK | 0 | 193.505593 | 0.196287 | 0 |
| 9 | EPPK1 | 0.265399 | 0.015432 | 4.104191 | OK | 0 | 266.1956611 | 0.270024 | 0 |
| 10 | CA2 | 1.371843 | 0.080025 | 4.099519 | OK | 0 | 236.2899464 | 0.239691 | 0 |
| 11 | SERPINB4 | 6.673203 | 0.427246 | 3.965242 | OK | 0 | 367.927037 | 0.37323 | 0 |
| 12 | NEFL | 0.280174 | 0.018846 | 3.894011 | OK | 0 | 177.4029801 | 0.179948 | 0 |
| 13 | SYNPO | 0.235893 | 0.016821 | 3.809789 | OK | 0 | 122.6287639 | 0.124385 | 0 |
| 14 | SCIN | 3.688674 | 0.275633 | 3.742284 | OK | 0 | 469.4131307 | 0.476182 | 0 |
| 15 | BCL2L15 | 0.386922 | 0.030523 | 3.664075 | OK | 0 | 219.2263121 | 0.222379 | 0 |
| 16 | SFRP1 | 0.617312 | 0.057587 | 3.422193 | OK | 0 | 216.6401506 | 0.219756 | 0 |
| 17 | CTSC | 3.435824 | 0.360298 | 3.253396 | OK | 0 | 293.4937114 | 0.297721 | 0 |
| 18 | CYP7B1 | 0.957797 | 0.11481 | 3.060474 | OK | 0 | 294.3818033 | 0.298622 | 0 |
| 19 | KCNJ2 | 0.580258 | 0.073126 | 2.988245 | OK | 0 | 130.4389326 | 0.13231 | 0 |
| 20 | SGK1 | 2.223951 | 0.3163 | 2.813758 | OK | 0 | 252.2459678 | 0.255878 | 0 |
| 21 | MDK | 0.882212 | 0.130983 | 2.751741 | OK | 0 | 188.5971158 | 0.191309 | 0 |
| 22 | NTRK2 | 0.948013 | 0.143536 | 2.723496 | OK | 0 | 213.3491183 | 0.216419 | 0 |
| 23 | CISH | 0.225675 | 0.035959 | 2.649824 | OK | 0 | 69.58338323 | 0.070575 | 0 |
| 24 | CADM1 | 0.144856 | 0.024142 | 2.584993 | OK | 0 | 118.3391683 | 0.120033 | 0 |
| 25 | HAS3 | 0.200476 | 0.034917 | 2.52142 | OK | 0 | 143.95017 | 0.146014 | 0 |
| 26 | SECTM1 | 0.169999 | 0.031297 | 2.441448 | OK | 0 | 136.333547 | 0.138287 | 0 |
| 27 | PHLDB2 | 0.153398 | 0.029796 | 2.36408 | OK | 0 | 99.34597926 | 0.100766 | 0 |
| 28 | APOL1 | 0.285596 | 0.056621 | 2.33457 | OK | 0 | 136.483189 | 0.13844 | 0 |
| 29 | CHL1 | 0.33855 | 0.071509 | 2.243166 | OK | 0 | 237.1249703 | 0.240535 | 0 |
| 30 | DPYD | 0.493135 | 0.106768 | 2.2075 | OK | 0 | 253.6263682 | 0.257277 | 0 |
| 31 | DUOX1 | 1.171431 | 0.253832 | 2.206326 | OK | 0 | 326.2683787 | 0.330969 | 0 |
| 32 | PDZK1IP1 | 3.354051 | 0.747063 | 2.166603 | OK | 0 | 292.720704 | 0.296937 | 0 |
| 33 | CAMK1D | 0.299013 | 0.067236 | 2.152896 | OK | 0 | 217.4202527 | 0.220547 | 0 |
| 34 | SH3RF2 | 0.653009 | 0.161891 | 2.01208 | OK | 0 | 244.2873354 | 0.247804 | 0 |
| 35 | C12orf75 | 0.453379 | 0.11281 | 2.006827 | OK | 0 | 160.5192575 | 0.162825 | 0 |
| 36 | GALNT5 | 0.476173 | 0.130597 | 1.866365 | OK | 0 | 214.1991008 | 0.21728 | 0 |
| 37 | SAMHD1 | 0.34603 | 0.096709 | 1.839171 | OK | 0 | 227.8986842 | 0.231177 | 0 |
| 38 | LTA4H | 1.99693 | 0.570936 | 1.806382 | OK | 0 | 348.1073431 | 0.353124 | 0 |
| 39 | MPP7 | 1.046935 | 0.307843 | 1.765906 | OK | 0 | 364.668985 | 0.369925 | 0 |
| 40 | SERPINB3 | 17.48953 | 5.324917 | 1.715661 | OK | 0 | 480.0374551 | 0.486956 | 0 |
| 41 | SERPINE2 | 0.356731 | 0.108897 | 1.711879 | OK | 0 | 174.5971759 | 0.177105 | 0 |
| 42 | MX1 | 0.797846 | 0.246848 | 1.692489 | OK | 0 | 268.5388615 | 0.272406 | 0 |
| 43 | MYH9 | 1.216838 | 0.379624 | 1.680495 | OK | 0 | 257.7729451 | 0.261484 | 0 |
| 44 | ATP8B1 | 0.488133 | 0.158052 | 1.626876 | OK | 0 | 230.211679 | 0.233524 | 0 |
| 45 | P2RY1 | 0.65632 | 0.213077 | 1.623026 | OK | 0 | 239.3458131 | 0.242791 | 0 |
| 46 | SPTBN1 | 0.839629 | 0.273835 | 1.616447 | OK | 0 | 247.5870723 | 0.251151 | 0 |
| 47 | CDH3 | 0.359101 | 0.118118 | 1.604159 | OK | 0 | 180.9037785 | 0.183503 | 0 |
| 48 | THBD | 0.404966 | 0.134632 | 1.588778 | OK | 0 | 199.7149552 | 0.202586 | 0 |
| 49 | HPGD | 3.934351 | 1.313569 | 1.582633 | OK | 0 | 354.9179683 | 0.360032 | 0 |
| 50 | TFPI | 0.120224 | 0.041968 | 1.518378 | OK | 0 | 66.9720114 | 0.067924 | 0 |
| 51 | HLF | 0.498671 | 0.178091 | 1.48547 | OK | 0 | 174.4748259 | 0.176982 | 0 |

| | | | | | | | | | |
|---|---|---|---|---|---|---|---|---|---|
| 52 | CLDN1 | 0.710254 | 0.256736 | 1.468051 | OK | 0 | 201.4444818 | 0.204342 | 0 |
| 53 | SPARCL1 | 0.139903 | 0.054756 | 1.353335 | OK | 0 | 78.91544755 | 0.080039 | 0 |
| 54 | CYP2S1 | 0.247008 | 0.099059 | 1.318203 | OK | 0 | 150.6675981 | 0.152829 | 0 |
| 55 | TMPRSS11/ | 0.897483 | 0.377531 | 1.249291 | OK | 0 | 361.7085279 | 0.366921 | 0 |
| 56 | CES2 | 2.138067 | 0.953795 | 1.164556 | OK | 0 | 470.9342532 | 0.477725 | 0 |
| 57 | DSG3 | 4.166981 | 1.902058 | 1.131441 | OK | 0 | 372.6045303 | 0.377972 | 0 |
| 58 | ARNTL2 | 0.553917 | 0.277875 | 0.995237 | OK | 0 | 264.2569026 | 0.268061 | 0 |
| 59 | DSP | 15.03454 | 7.578231 | 0.988348 | OK | 0 | 236.2433509 | 0.239636 | 0 |
| 60 | TFRC | 0.392033 | 0.209336 | 0.905155 | OK | 0 | 175.4500203 | 0.177971 | 0 |
| 61 | ZFP36L1 | 5.059103 | 2.971625 | 0.76763 | OK | 0 | 378.064425 | 0.38351 | 0 |

**Supplemental Table 5.** Identification of SOX2 and/or KLF5 downstream targets from Enrichr databases and literature-based analyses.

| | gene | source | regulator |
|---|---|---|---|
| 1 | CACNA2D4 | Enrichr CHEA_2022: SOX2 20726797 ChIP-Seq SW620 Human | SOX2 |
| 2 | KIAA1586 | Enrichr CHEA_2022: SOX2 20726797 ChIP-Seq SW620 Human | SOX2 |
| 3 | ISL1 | Enrichr CHEA_2022: SOX2 20726797 ChIP-Seq SW620 Human | SOX2 |
| 4 | LARP6 | Enrichr CHEA_2022: SOX2 20726797 ChIP-Seq SW620 Human | SOX2 |
| 5 | L1CAM | Enrichr CHEA_2022: SOX2 20726797 ChIP-Seq SW620 Human | SOX2 |
| 6 | SPINT2 | Enrichr CHEA_2022: SOX2 20726797 ChIP-Seq SW620 Human | SOX2 |
| 7 | ASPHD1 | Enrichr CHEA_2022: SOX2 20726797 ChIP-Seq SW620 Human | SOX2 |
| 8 | SPINT1 | Enrichr CHEA_2022: SOX2 20726797 ChIP-Seq SW620 Human | SOX2 |
| 9 | FEZ2 | Enrichr CHEA_2022: SOX2 20726797 ChIP-Seq SW620 Human | SOX2 |
| 10 | SCO1 | Enrichr CHEA_2022: SOX2 20726797 ChIP-Seq SW620 Human | SOX2 |
| 11 | ATF3 | Enrichr CHEA_2022: SOX2 20726797 ChIP-Seq SW620 Human | SOX2 |
| 12 | SAMD9 | Enrichr CHEA_2022: SOX2 20726797 ChIP-Seq SW620 Human | SOX2 |
| 13 | HPGD | Enrichr CHEA_2022: SOX2 20726797 ChIP-Seq SW620 Human | SOX2 |
| 14 | PON3 | Enrichr CHEA_2022: SOX2 20726797 ChIP-Seq SW620 Human | SOX2 |
| 15 | CRABP2 | Enrichr CHEA_2022: SOX2 20726797 ChIP-Seq SW620 Human | SOX2 |
| 16 | TFPI | Enrichr CHEA_2022: SOX2 20726797 ChIP-Seq SW620 Human | SOX2 |
| 17 | UACA | Enrichr CHEA_2022: SOX2 20726797 ChIP-Seq SW620 Human | SOX2 |
| 18 | SPDEF | Enrichr CHEA_2022: SOX2 20726797 ChIP-Seq SW620 Human | SOX2 |
| 19 | TFPT | Enrichr CHEA_2022: SOX2 20726797 ChIP-Seq SW620 Human | SOX2 |
| 20 | DPYSL3 | Enrichr CHEA_2022: SOX2 20726797 ChIP-Seq SW620 Human | SOX2 |
| 21 | ZNF19 | Enrichr CHEA_2022: SOX2 20726797 ChIP-Seq SW620 Human | SOX2 |
| 22 | JAG1 | Enrichr CHEA_2022: SOX2 20726797 ChIP-Seq SW620 Human | SOX2 |
| 23 | NEBL | Enrichr CHEA_2022: SOX2 20726797 ChIP-Seq SW620 Human | SOX2 |
| 24 | ATP6AP2 | Enrichr CHEA_2022: SOX2 20726797 ChIP-Seq SW620 Human | SOX2 |
| 25 | ATP6AP1 | Enrichr CHEA_2022: SOX2 20726797 ChIP-Seq SW620 Human | SOX2 |
| 26 | NR2F2 | Enrichr CHEA_2022: SOX2 20726797 ChIP-Seq SW620 Human | SOX2 |
| 27 | PBX3 | Enrichr CHEA_2022: SOX2 20726797 ChIP-Seq SW620 Human | SOX2 |
| 28 | VPS13A | Enrichr CHEA_2022: SOX2 20726797 ChIP-Seq SW620 Human | SOX2 |
| 29 | SUPV3L1 | Enrichr CHEA_2022: SOX2 20726797 ChIP-Seq SW620 Human | SOX2 |
| 30 | VANGL1 | Enrichr CHEA_2022: SOX2 20726797 ChIP-Seq SW620 Human | SOX2 |
| 31 | BCR | Enrichr CHEA_2022: SOX2 20726797 ChIP-Seq SW620 Human | SOX2 |
| 32 | PHF20L1 | Enrichr CHEA_2022: SOX2 20726797 ChIP-Seq SW620 Human | SOX2 |
| 33 | PPFIBP2 | Enrichr CHEA_2022: SOX2 20726797 ChIP-Seq SW620 Human | SOX2 |
| 34 | FAM111A | Enrichr CHEA_2022: SOX2 20726797 ChIP-Seq SW620 Human | SOX2 |
| 35 | PLSCR1 | Enrichr CHEA_2022: SOX2 20726797 ChIP-Seq SW620 Human | SOX2 |
| 36 | PKIA | Enrichr CHEA_2022: SOX2 20726797 ChIP-Seq SW620 Human | SOX2 |
| 37 | SFXN3 | Enrichr CHEA_2022: SOX2 20726797 ChIP-Seq SW620 Human | SOX2 |
| 38 | CHST5 | Enrichr CHEA_2022: SOX2 20726797 ChIP-Seq SW620 Human | SOX2 |
| 39 | PKIG | Enrichr CHEA_2022: SOX2 20726797 ChIP-Seq SW620 Human | SOX2 |
| 40 | ATF4 | Enrichr CHEA_2022: SOX2 20726797 ChIP-Seq SW620 Human | SOX2 |
| 41 | ATF5 | Enrichr CHEA_2022: SOX2 20726797 ChIP-Seq SW620 Human | SOX2 |
| 42 | CHST4 | Enrichr CHEA_2022: SOX2 20726797 ChIP-Seq SW620 Human | SOX2 |
| 43 | CDCA2 | Enrichr CHEA_2022: SOX2 20726797 ChIP-Seq SW620 Human | SOX2 |
| 44 | PDGFC | Enrichr CHEA_2022: SOX2 20726797 ChIP-Seq SW620 Human | SOX2 |
| 45 | ZNF22 | Enrichr CHEA_2022: SOX2 20726797 ChIP-Seq SW620 Human | SOX2 |
| 46 | GATA6 | Enrichr CHEA_2022: SOX2 20726797 ChIP-Seq SW620 Human | SOX2 |
| 47 | PDGFA | Enrichr CHEA_2022: SOX2 20726797 ChIP-Seq SW620 Human | SOX2 |
| 48 | F11R | Enrichr CHEA_2022: SOX2 20726797 ChIP-Seq SW620 Human | SOX2 |
| 49 | GATA3 | Enrichr CHEA_2022: SOX2 20726797 ChIP-Seq SW620 Human | SOX2 |
| 50 | ALKBH8 | Enrichr CHEA_2022: SOX2 20726797 ChIP-Seq SW620 Human | SOX2 |
| 51 | GATA2 | Enrichr CHEA_2022: SOX2 20726797 ChIP-Seq SW620 Human | SOX2 |
| 52 | ZNF26 | Enrichr CHEA_2022: SOX2 20726797 ChIP-Seq SW620 Human | SOX2 |
| 53 | ALKBH6 | Enrichr CHEA_2022: SOX2 20726797 ChIP-Seq SW620 Human | SOX2 |
| 54 | EPB41L1 | Enrichr CHEA_2022: SOX2 20726797 ChIP-Seq SW620 Human | SOX2 |
| 55 | ZC3H12A | Enrichr CHEA_2022: SOX2 20726797 ChIP-Seq SW620 Human | SOX2 |
| 56 | EPB41L2 | Enrichr CHEA_2022: SOX2 20726797 ChIP-Seq SW620 Human | SOX2 |
| 57 | FAM110C | Enrichr CHEA_2022: SOX2 20726797 ChIP-Seq SW620 Human | SOX2 |
| 58 | NKX3-1 | Enrichr CHEA_2022: SOX2 20726797 ChIP-Seq SW620 Human | SOX2 |
| 59 | PIP5K1B | Enrichr CHEA_2022: SOX2 20726797 ChIP-Seq SW620 Human | SOX2 |

| | | | |
|---|---|---|---|
| 60 | PRKACB | Enrichr CHEA_2022: SOX2 20726797 ChIP-Seq SW620 Human | SOX2 |
| 61 | ZNF488 | Enrichr CHEA_2022: SOX2 20726797 ChIP-Seq SW620 Human | SOX2 |
| 62 | DUSP5 | Enrichr CHEA_2022: SOX2 20726797 ChIP-Seq SW620 Human | SOX2 |
| 63 | FUCA1 | Enrichr CHEA_2022: SOX2 20726797 ChIP-Seq SW620 Human | SOX2 |
| 64 | JUN | Enrichr CHEA_2022: SOX2 20726797 ChIP-Seq SW620 Human | SOX2 |
| 65 | DUSP4 | Enrichr CHEA_2022: SOX2 20726797 ChIP-Seq SW620 Human | SOX2 |
| 66 | JUP | Enrichr CHEA_2022: SOX2 20726797 ChIP-Seq SW620 Human | SOX2 |
| 67 | FN1 | Enrichr CHEA_2022: SOX2 20726797 ChIP-Seq SW620 Human | SOX2 |
| 68 | DCBLD2 | Enrichr CHEA_2022: SOX2 20726797 ChIP-Seq SW620 Human | SOX2 |
| 69 | TMEFF1 | Enrichr CHEA_2022: SOX2 20726797 ChIP-Seq SW620 Human | SOX2 |
| 70 | SLC7A5 | Enrichr CHEA_2022: SOX2 20726797 ChIP-Seq SW620 Human | SOX2 |
| 71 | CENPF | Enrichr CHEA_2022: SOX2 20726797 ChIP-Seq SW620 Human | SOX2 |
| 72 | SLC7A7 | Enrichr CHEA_2022: SOX2 20726797 ChIP-Seq SW620 Human | SOX2 |
| 73 | SLC7A8 | Enrichr CHEA_2022: SOX2 20726797 ChIP-Seq SW620 Human | SOX2 |
| 74 | DPYD | Enrichr CHEA_2022: SOX2 20726797 ChIP-Seq SW620 Human | SOX2 |
| 75 | FXYD6 | Enrichr CHEA_2022: SOX2 20726797 ChIP-Seq SW620 Human | SOX2 |
| 76 | RAPGEF4 | Enrichr CHEA_2022: SOX2 20726797 ChIP-Seq SW620 Human | SOX2 |
| 77 | RAPGEF5 | Enrichr CHEA_2022: SOX2 20726797 ChIP-Seq SW620 Human | SOX2 |
| 78 | LIMS2 | Enrichr CHEA_2022: SOX2 20726797 ChIP-Seq SW620 Human | SOX2 |
| 79 | TFCP2L1 | Enrichr CHEA_2022: SOX2 20726797 ChIP-Seq SW620 Human | SOX2 |
| 80 | ZC3HAV1 | Enrichr CHEA_2022: SOX2 20726797 ChIP-Seq SW620 Human | SOX2 |
| 81 | SLC7A1 | Enrichr CHEA_2022: SOX2 20726797 ChIP-Seq SW620 Human | SOX2 |
| 82 | HOXC13 | Enrichr CHEA_2022: SOX2 20726797 ChIP-Seq SW620 Human | SOX2 |
| 83 | SLC7A2 | Enrichr CHEA_2022: SOX2 20726797 ChIP-Seq SW620 Human | SOX2 |
| 84 | HOXC10 | Enrichr CHEA_2022: SOX2 20726797 ChIP-Seq SW620 Human | SOX2 |
| 85 | HLCS | Enrichr CHEA_2022: SOX2 20726797 ChIP-Seq SW620 Human | SOX2 |
| 86 | SHH | Enrichr CHEA_2022: SOX2 20726797 ChIP-Seq SW620 Human | SOX2 |
| 87 | PANX1 | Enrichr CHEA_2022: SOX2 20726797 ChIP-Seq SW620 Human | SOX2 |
| 88 | DLGAP1 | Enrichr CHEA_2022: SOX2 20726797 ChIP-Seq SW620 Human | SOX2 |
| 89 | ZNF468 | Enrichr CHEA_2022: SOX2 20726797 ChIP-Seq SW620 Human | SOX2 |
| 90 | SLC18A2 | Enrichr CHEA_2022: SOX2 20726797 ChIP-Seq SW620 Human | SOX2 |
| 91 | PCMTD2 | Enrichr CHEA_2022: SOX2 20726797 ChIP-Seq SW620 Human | SOX2 |
| 92 | EGLN2 | Enrichr CHEA_2022: SOX2 20726797 ChIP-Seq SW620 Human | SOX2 |
| 93 | ATAD2 | Enrichr CHEA_2022: SOX2 20726797 ChIP-Seq SW620 Human | SOX2 |
| 94 | CDKL5 | Enrichr CHEA_2022: SOX2 20726797 ChIP-Seq SW620 Human | SOX2 |
| 95 | PPARA | Enrichr CHEA_2022: SOX2 20726797 ChIP-Seq SW620 Human | SOX2 |
| 96 | DGAT1 | Enrichr CHEA_2022: SOX2 20726797 ChIP-Seq SW620 Human | SOX2 |
| 97 | SNX25 | Enrichr CHEA_2022: SOX2 20726797 ChIP-Seq SW620 Human | SOX2 |
| 98 | BGN | Enrichr CHEA_2022: SOX2 20726797 ChIP-Seq SW620 Human | SOX2 |
| 99 | ADAM10 | Enrichr CHEA_2022: SOX2 20726797 ChIP-Seq SW620 Human | SOX2 |
| 100 | SLC39A14 | Enrichr CHEA_2022: SOX2 20726797 ChIP-Seq SW620 Human | SOX2 |
| 101 | DAB2IP | Enrichr CHEA_2022: SOX2 20726797 ChIP-Seq SW620 Human | SOX2 |
| 102 | MYBBP1A | Enrichr CHEA_2022: SOX2 20726797 ChIP-Seq SW620 Human | SOX2 |
| 103 | ADAM17 | Enrichr CHEA_2022: SOX2 20726797 ChIP-Seq SW620 Human | SOX2 |
| 104 | LARS2 | Enrichr CHEA_2022: SOX2 20726797 ChIP-Seq SW620 Human | SOX2 |
| 105 | APEX2 | Enrichr CHEA_2022: SOX2 20726797 ChIP-Seq SW620 Human | SOX2 |
| 106 | ETNK1 | Enrichr CHEA_2022: SOX2 20726797 ChIP-Seq SW620 Human | SOX2 |
| 107 | PPARG | Enrichr CHEA_2022: SOX2 20726797 ChIP-Seq SW620 Human | SOX2 |
| 108 | TSR1 | Enrichr CHEA_2022: SOX2 20726797 ChIP-Seq SW620 Human | SOX2 |
| 109 | LGR6 | Enrichr CHEA_2022: SOX2 20726797 ChIP-Seq SW620 Human | SOX2 |
| 110 | PPARD | Enrichr CHEA_2022: SOX2 20726797 ChIP-Seq SW620 Human | SOX2 |
| 111 | LGR4 | Enrichr CHEA_2022: SOX2 20726797 ChIP-Seq SW620 Human | SOX2 |
| 112 | OAT | Enrichr CHEA_2022: SOX2 20726797 ChIP-Seq SW620 Human | SOX2 |
| 113 | MTMR10 | Enrichr CHEA_2022: SOX2 20726797 ChIP-Seq SW620 Human | SOX2 |
| 114 | SNX10 | Enrichr CHEA_2022: SOX2 20726797 ChIP-Seq SW620 Human | SOX2 |
| 115 | SMPD3 | Enrichr CHEA_2022: SOX2 20726797 ChIP-Seq SW620 Human | SOX2 |
| 116 | WRN | Enrichr CHEA_2022: SOX2 20726797 ChIP-Seq SW620 Human | SOX2 |
| 117 | FGFR1OP2 | Enrichr CHEA_2022: SOX2 20726797 ChIP-Seq SW620 Human | SOX2 |
| 118 | PLAGL1 | Enrichr CHEA_2022: SOX2 20726797 ChIP-Seq SW620 Human | SOX2 |
| 119 | AGR2 | Enrichr CHEA_2022: SOX2 20726797 ChIP-Seq SW620 Human | SOX2 |
| 120 | NUDT13 | Enrichr CHEA_2022: SOX2 20726797 ChIP-Seq SW620 Human | SOX2 |

| | | | |
|---|---|---|---|
| 121 | FADS1 | Enrichr CHEA_2022: SOX2 20726797 ChIP-Seq SW620 Human | SOX2 |
| 122 | MAP3K1 | Enrichr CHEA_2022: SOX2 20726797 ChIP-Seq SW620 Human | SOX2 |
| 123 | BIK | Enrichr CHEA_2022: SOX2 20726797 ChIP-Seq SW620 Human | SOX2 |
| 124 | CORO2A | Enrichr CHEA_2022: SOX2 20726797 ChIP-Seq SW620 Human | SOX2 |
| 125 | FBXO17 | Enrichr CHEA_2022: SOX2 20726797 ChIP-Seq SW620 Human | SOX2 |
| 126 | STX8 | Enrichr CHEA_2022: SOX2 20726797 ChIP-Seq SW620 Human | SOX2 |
| 127 | ATP11C | Enrichr CHEA_2022: SOX2 20726797 ChIP-Seq SW620 Human | SOX2 |
| 128 | SDCBP2 | Enrichr CHEA_2022: SOX2 20726797 ChIP-Seq SW620 Human | SOX2 |
| 129 | WT1 | Enrichr CHEA_2022: SOX2 20726797 ChIP-Seq SW620 Human | SOX2 |
| 130 | MFNG | Enrichr CHEA_2022: SOX2 20726797 ChIP-Seq SW620 Human | SOX2 |
| 131 | LY6E | Enrichr CHEA_2022: SOX2 20726797 ChIP-Seq SW620 Human | SOX2 |
| 132 | PDCD4 | Enrichr CHEA_2022: SOX2 20726797 ChIP-Seq SW620 Human | SOX2 |
| 133 | PKP4 | Enrichr CHEA_2022: SOX2 20726797 ChIP-Seq SW620 Human | SOX2 |
| 134 | IL6ST | Enrichr CHEA_2022: SOX2 20726797 ChIP-Seq SW620 Human | SOX2 |
| 135 | AGRN | Enrichr CHEA_2022: SOX2 20726797 ChIP-Seq SW620 Human | SOX2 |
| 136 | FRK | Enrichr CHEA_2022: SOX2 20726797 ChIP-Seq SW620 Human | SOX2 |
| 137 | ZNF430 | Enrichr CHEA_2022: SOX2 20726797 ChIP-Seq SW620 Human | SOX2 |
| 138 | NIP7 | Enrichr CHEA_2022: SOX2 20726797 ChIP-Seq SW620 Human | SOX2 |
| 139 | PDCD6 | Enrichr CHEA_2022: SOX2 20726797 ChIP-Seq SW620 Human | SOX2 |
| 140 | TSPAN31 | Enrichr CHEA_2022: SOX2 20726797 ChIP-Seq SW620 Human | SOX2 |
| 141 | ATP10B | Enrichr CHEA_2022: SOX2 20726797 ChIP-Seq SW620 Human | SOX2 |
| 142 | PLOD2 | Enrichr CHEA_2022: SOX2 20726797 ChIP-Seq SW620 Human | SOX2 |
| 143 | FRY | Enrichr CHEA_2022: SOX2 20726797 ChIP-Seq SW620 Human | SOX2 |
| 144 | EGFR | Enrichr CHEA_2022: SOX2 20726797 ChIP-Seq SW620 Human | SOX2 |
| 145 | LAPTM4A | Enrichr CHEA_2022: SOX2 20726797 ChIP-Seq SW620 Human | SOX2 |
| 146 | STX5 | Enrichr CHEA_2022: SOX2 20726797 ChIP-Seq SW620 Human | SOX2 |
| 147 | MAP3K8 | Enrichr CHEA_2022: SOX2 20726797 ChIP-Seq SW620 Human | SOX2 |
| 148 | ZNF420 | Enrichr CHEA_2022: SOX2 20726797 ChIP-Seq SW620 Human | SOX2 |
| 149 | NEK6 | Enrichr CHEA_2022: SOX2 20726797 ChIP-Seq SW620 Human | SOX2 |
| 150 | BNIP3L | Enrichr CHEA_2022: SOX2 20726797 ChIP-Seq SW620 Human | SOX2 |
| 151 | SLC9A3R1 | Enrichr CHEA_2022: SOX2 20726797 ChIP-Seq SW620 Human | SOX2 |
| 152 | PIAS3 | Enrichr CHEA_2022: SOX2 20726797 ChIP-Seq SW620 Human | SOX2 |
| 153 | FBXO31 | Enrichr CHEA_2022: SOX2 20726797 ChIP-Seq SW620 Human | SOX2 |
| 154 | FBXO32 | Enrichr CHEA_2022: SOX2 20726797 ChIP-Seq SW620 Human | SOX2 |
| 155 | FBXO30 | Enrichr CHEA_2022: SOX2 20726797 ChIP-Seq SW620 Human | SOX2 |
| 156 | LCN2 | Enrichr CHEA_2022: SOX2 20726797 ChIP-Seq SW620 Human | SOX2 |
| 157 | BMP2K | Enrichr CHEA_2022: SOX2 20726797 ChIP-Seq SW620 Human | SOX2 |
| 158 | MNAT1 | Enrichr CHEA_2022: SOX2 20726797 ChIP-Seq SW620 Human | SOX2 |
| 159 | SLC27A5 | Enrichr CHEA_2022: SOX2 20726797 ChIP-Seq SW620 Human | SOX2 |
| 160 | OSBPL1A | Enrichr CHEA_2022: SOX2 20726797 ChIP-Seq SW620 Human | SOX2 |
| 161 | KCNE3 | Enrichr CHEA_2022: SOX2 20726797 ChIP-Seq SW620 Human | SOX2 |
| 162 | COL17A1 | Enrichr CHEA_2022: SOX2 20726797 ChIP-Seq SW620 Human | SOX2 |
| 163 | EIF4EBP1 | Enrichr CHEA_2022: SOX2 20726797 ChIP-Seq SW620 Human | SOX2 |
| 164 | LY75 | Enrichr CHEA_2022: SOX2 20726797 ChIP-Seq SW620 Human | SOX2 |
| 165 | MSI2 | Enrichr CHEA_2022: SOX2 20726797 ChIP-Seq SW620 Human | SOX2 |
| 166 | NHSL1 | Enrichr CHEA_2022: SOX2 20726797 ChIP-Seq SW620 Human | SOX2 |
| 167 | MED31 | Enrichr CHEA_2022: SOX2 20726797 ChIP-Seq SW620 Human | SOX2 |
| 168 | FNTA | Enrichr CHEA_2022: SOX2 20726797 ChIP-Seq SW620 Human | SOX2 |
| 169 | UROD | Enrichr CHEA_2022: SOX2 20726797 ChIP-Seq SW620 Human | SOX2 |
| 170 | NEK1 | Enrichr CHEA_2022: SOX2 20726797 ChIP-Seq SW620 Human | SOX2 |
| 171 | NEK2 | Enrichr CHEA_2022: SOX2 20726797 ChIP-Seq SW620 Human | SOX2 |
| 172 | NFATC2IP | Enrichr CHEA_2022: SOX2 20726797 ChIP-Seq SW620 Human | SOX2 |
| 173 | SLC2A4RG | Enrichr CHEA_2022: SOX2 20726797 ChIP-Seq SW620 Human | SOX2 |
| 174 | ITGA5 | Enrichr CHEA_2022: SOX2 20726797 ChIP-Seq SW620 Human | SOX2 |
| 175 | ITGA3 | Enrichr CHEA_2022: SOX2 20726797 ChIP-Seq SW620 Human | SOX2 |
| 176 | TNFRSF19 | Enrichr CHEA_2022: SOX2 20726797 ChIP-Seq SW620 Human | SOX2 |
| 177 | TNFRSF1B | Enrichr CHEA_2022: SOX2 20726797 ChIP-Seq SW620 Human | SOX2 |
| 178 | TMBIM1 | Enrichr CHEA_2022: SOX2 20726797 ChIP-Seq SW620 Human | SOX2 |
| 179 | DBNDD2 | Enrichr CHEA_2022: SOX2 20726797 ChIP-Seq SW620 Human | SOX2 |
| 180 | TNFRSF21 | Enrichr CHEA_2022: SOX2 20726797 ChIP-Seq SW620 Human | SOX2 |
| 181 | LCP1 | Enrichr CHEA_2022: SOX2 20726797 ChIP-Seq SW620 Human | SOX2 |

| | | | |
|---|---|---|---|
| 182 | TMBIM4 | Enrichr CHEA_2022: SOX2 20726797 ChIP-Seq SW620 Human | SOX2 |
| 183 | PCCA | Enrichr CHEA_2022: SOX2 20726797 ChIP-Seq SW620 Human | SOX2 |
| 184 | SP5 | Enrichr CHEA_2022: SOX2 20726797 ChIP-Seq SW620 Human | SOX2 |
| 185 | OGT | Enrichr CHEA_2022: SOX2 20726797 ChIP-Seq SW620 Human | SOX2 |
| 186 | KCNK1 | Enrichr CHEA_2022: SOX2 20726797 ChIP-Seq SW620 Human | SOX2 |
| 187 | BCAR3 | Enrichr CHEA_2022: SOX2 20726797 ChIP-Seq SW620 Human | SOX2 |
| 188 | MCFD2 | Enrichr CHEA_2022: SOX2 20726797 ChIP-Seq SW620 Human | SOX2 |
| 189 | SHMT2 | Enrichr CHEA_2022: SOX2 20726797 ChIP-Seq SW620 Human | SOX2 |
| 190 | MLLT3 | Enrichr CHEA_2022: SOX2 20726797 ChIP-Seq SW620 Human | SOX2 |
| 191 | SPATA7 | Enrichr CHEA_2022: SOX2 20726797 ChIP-Seq SW620 Human | SOX2 |
| 192 | MLLT6 | Enrichr CHEA_2022: SOX2 20726797 ChIP-Seq SW620 Human | SOX2 |
| 193 | ALAD | Enrichr CHEA_2022: SOX2 20726797 ChIP-Seq SW620 Human | SOX2 |
| 194 | RDH13 | Enrichr CHEA_2022: SOX2 20726797 ChIP-Seq SW620 Human | SOX2 |
| 195 | KIAA1522 | Enrichr CHEA_2022: SOX2 20726797 ChIP-Seq SW620 Human | SOX2 |
| 196 | BOK | Enrichr CHEA_2022: SOX2 20726797 ChIP-Seq SW620 Human | SOX2 |
| 197 | BCAS1 | Enrichr CHEA_2022: SOX2 20726797 ChIP-Seq SW620 Human | SOX2 |
| 198 | NADK | Enrichr CHEA_2022: SOX2 20726797 ChIP-Seq SW620 Human | SOX2 |
| 199 | MBIP | Enrichr CHEA_2022: SOX2 20726797 ChIP-Seq SW620 Human | SOX2 |
| 200 | STYK1 | Enrichr CHEA_2022: SOX2 20726797 ChIP-Seq SW620 Human | SOX2 |
| 201 | GZMB | Enrichr CHEA_2022: SOX2 20726797 ChIP-Seq SW620 Human | SOX2 |
| 202 | MID1IP1 | Enrichr CHEA_2022: SOX2 20726797 ChIP-Seq SW620 Human | SOX2 |
| 203 | MYO10 | Enrichr CHEA_2022: SOX2 20726797 ChIP-Seq SW620 Human | SOX2 |
| 204 | RNASET2 | Enrichr CHEA_2022: SOX2 20726797 ChIP-Seq SW620 Human | SOX2 |
| 205 | CABLES1 | Enrichr CHEA_2022: SOX2 20726797 ChIP-Seq SW620 Human | SOX2 |
| 206 | TM6SF1 | Enrichr CHEA_2022: SOX2 20726797 ChIP-Seq SW620 Human | SOX2 |
| 207 | TBX3 | Enrichr CHEA_2022: SOX2 20726797 ChIP-Seq SW620 Human | SOX2 |
| 208 | CFDP1 | Enrichr CHEA_2022: SOX2 20726797 ChIP-Seq SW620 Human | SOX2 |
| 209 | MYO1C | Enrichr CHEA_2022: SOX2 20726797 ChIP-Seq SW620 Human | SOX2 |
| 210 | TSPAN14 | Enrichr CHEA_2022: SOX2 20726797 ChIP-Seq SW620 Human | SOX2 |
| 211 | MYO1B | Enrichr CHEA_2022: SOX2 20726797 ChIP-Seq SW620 Human | SOX2 |
| 212 | TRIM36 | Enrichr CHEA_2022: SOX2 20726797 ChIP-Seq SW620 Human | SOX2 |
| 213 | KCTD15 | Enrichr CHEA_2022: SOX2 20726797 ChIP-Seq SW620 Human | SOX2 |
| 214 | KCTD12 | Enrichr CHEA_2022: SOX2 20726797 ChIP-Seq SW620 Human | SOX2 |
| 215 | TRIM32 | Enrichr CHEA_2022: SOX2 20726797 ChIP-Seq SW620 Human | SOX2 |
| 216 | PIK3AP1 | Enrichr CHEA_2022: SOX2 20726797 ChIP-Seq SW620 Human | SOX2 |
| 217 | TRIM34 | Enrichr CHEA_2022: SOX2 20726797 ChIP-Seq SW620 Human | SOX2 |
| 218 | PRPS2 | Enrichr CHEA_2022: SOX2 20726797 ChIP-Seq SW620 Human | SOX2 |
| 219 | SRC | Enrichr CHEA_2022: SOX2 20726797 ChIP-Seq SW620 Human | SOX2 |
| 220 | MSLN | Enrichr CHEA_2022: SOX2 20726797 ChIP-Seq SW620 Human | SOX2 |
| 221 | TRIM31 | Enrichr CHEA_2022: SOX2 20726797 ChIP-Seq SW620 Human | SOX2 |
| 222 | ITGB5 | Enrichr CHEA_2022: SOX2 20726797 ChIP-Seq SW620 Human | SOX2 |
| 223 | SRI | Enrichr CHEA_2022: SOX2 20726797 ChIP-Seq SW620 Human | SOX2 |
| 224 | TSPAN12 | Enrichr CHEA_2022: SOX2 20726797 ChIP-Seq SW620 Human | SOX2 |
| 225 | NRGN | Enrichr CHEA_2022: SOX2 20726797 ChIP-Seq SW620 Human | SOX2 |
| 226 | ZNRF1 | Enrichr CHEA_2022: SOX2 20726797 ChIP-Seq SW620 Human | SOX2 |
| 227 | ZNRF2 | Enrichr CHEA_2022: SOX2 20726797 ChIP-Seq SW620 Human | SOX2 |
| 228 | SRR | Enrichr CHEA_2022: SOX2 20726797 ChIP-Seq SW620 Human | SOX2 |
| 229 | HHEX | Enrichr CHEA_2022: SOX2 20726797 ChIP-Seq SW620 Human | SOX2 |
| 230 | KIAA1549 | Enrichr CHEA_2022: SOX2 20726797 ChIP-Seq SW620 Human | SOX2 |
| 231 | TRIM29 | Enrichr CHEA_2022: SOX2 20726797 ChIP-Seq SW620 Human | SOX2 |
| 232 | PTER | Enrichr CHEA_2022: SOX2 20726797 ChIP-Seq SW620 Human | SOX2 |
| 233 | IGF2BP1 | Enrichr CHEA_2022: SOX2 20726797 ChIP-Seq SW620 Human | SOX2 |
| 234 | ITGB8 | Enrichr CHEA_2022: SOX2 20726797 ChIP-Seq SW620 Human | SOX2 |
| 235 | ITGAV | Enrichr CHEA_2022: SOX2 20726797 ChIP-Seq SW620 Human | SOX2 |
| 236 | FYN | Enrichr CHEA_2022: SOX2 20726797 ChIP-Seq SW620 Human | SOX2 |
| 237 | ARL5B | Enrichr CHEA_2022: SOX2 20726797 ChIP-Seq SW620 Human | SOX2 |
| 238 | SMARCA1 | Enrichr CHEA_2022: SOX2 20726797 ChIP-Seq SW620 Human | SOX2 |
| 239 | ACTN4 | Enrichr CHEA_2022: SOX2 20726797 ChIP-Seq SW620 Human | SOX2 |
| 240 | FBXO3 | Enrichr CHEA_2022: SOX2 20726797 ChIP-Seq SW620 Human | SOX2 |
| 241 | ARL14 | Enrichr CHEA_2022: SOX2 20726797 ChIP-Seq SW620 Human | SOX2 |
| 242 | SMARCA2 | Enrichr CHEA_2022: SOX2 20726797 ChIP-Seq SW620 Human | SOX2 |

| | | | |
|---|---|---|---|
| 243 | PPA2 | Enrichr CHEA_2022: SOX2 20726797 ChIP-Seq SW620 Human | SOX2 |
| 244 | SOAT1 | Enrichr CHEA_2022: SOX2 20726797 ChIP-Seq SW620 Human | SOX2 |
| 245 | NINJ2 | Enrichr CHEA_2022: SOX2 20726797 ChIP-Seq SW620 Human | SOX2 |
| 246 | MT1G | Enrichr CHEA_2022: SOX2 20726797 ChIP-Seq SW620 Human | SOX2 |
| 247 | MT1H | Enrichr CHEA_2022: SOX2 20726797 ChIP-Seq SW620 Human | SOX2 |
| 248 | TRIM16 | Enrichr CHEA_2022: SOX2 20726797 ChIP-Seq SW620 Human | SOX2 |
| 249 | CCDC82 | Enrichr CHEA_2022: SOX2 20726797 ChIP-Seq SW620 Human | SOX2 |
| 250 | RIN3 | Enrichr CHEA_2022: SOX2 20726797 ChIP-Seq SW620 Human | SOX2 |
| 251 | MT1F | Enrichr CHEA_2022: SOX2 20726797 ChIP-Seq SW620 Human | SOX2 |
| 252 | SLC23A3 | Enrichr CHEA_2022: SOX2 20726797 ChIP-Seq SW620 Human | SOX2 |
| 253 | EPAS1 | Enrichr CHEA_2022: SOX2 20726797 ChIP-Seq SW620 Human | SOX2 |
| 254 | TAGLN | Enrichr CHEA_2022: SOX2 20726797 ChIP-Seq SW620 Human | SOX2 |
| 255 | CCDC69 | Enrichr CHEA_2022: SOX2 20726797 ChIP-Seq SW620 Human | SOX2 |
| 256 | MYCBP2 | Enrichr CHEA_2022: SOX2 20726797 ChIP-Seq SW620 Human | SOX2 |
| 257 | CPNE8 | Enrichr CHEA_2022: SOX2 20726797 ChIP-Seq SW620 Human | SOX2 |
| 258 | IL27RA | Enrichr CHEA_2022: SOX2 20726797 ChIP-Seq SW620 Human | SOX2 |
| 259 | STS | Enrichr CHEA_2022: SOX2 20726797 ChIP-Seq SW620 Human | SOX2 |
| 260 | CDC23 | Enrichr CHEA_2022: SOX2 20726797 ChIP-Seq SW620 Human | SOX2 |
| 261 | BSG | Enrichr CHEA_2022: SOX2 20726797 ChIP-Seq SW620 Human | SOX2 |
| 262 | CSNK1D | Enrichr CHEA_2022: SOX2 20726797 ChIP-Seq SW620 Human | SOX2 |
| 263 | PEG10 | Enrichr CHEA_2022: SOX2 20726797 ChIP-Seq SW620 Human | SOX2 |
| 264 | STK38L | Enrichr CHEA_2022: SOX2 20726797 ChIP-Seq SW620 Human | SOX2 |
| 265 | CCDC77 | Enrichr CHEA_2022: SOX2 20726797 ChIP-Seq SW620 Human | SOX2 |
| 266 | E2F8 | Enrichr CHEA_2022: SOX2 20726797 ChIP-Seq SW620 Human | SOX2 |
| 267 | EGR3 | Enrichr CHEA_2022: SOX2 20726797 ChIP-Seq SW620 Human | SOX2 |
| 268 | TGDS | Enrichr CHEA_2022: SOX2 20726797 ChIP-Seq SW620 Human | SOX2 |
| 269 | MYO5A | Enrichr CHEA_2022: SOX2 20726797 ChIP-Seq SW620 Human | SOX2 |
| 270 | RAB27B | Enrichr CHEA_2022: SOX2 20726797 ChIP-Seq SW620 Human | SOX2 |
| 271 | MYO5B | Enrichr CHEA_2022: SOX2 20726797 ChIP-Seq SW620 Human | SOX2 |
| 272 | CSNK1E | Enrichr CHEA_2022: SOX2 20726797 ChIP-Seq SW620 Human | SOX2 |
| 273 | PTK6 | Enrichr CHEA_2022: SOX2 20726797 ChIP-Seq SW620 Human | SOX2 |
| 274 | EXT1 | Enrichr CHEA_2022: SOX2 20726797 ChIP-Seq SW620 Human | SOX2 |
| 275 | PTK2 | Enrichr CHEA_2022: SOX2 20726797 ChIP-Seq SW620 Human | SOX2 |
| 276 | CNKSR3 | Enrichr CHEA_2022: SOX2 20726797 ChIP-Seq SW620 Human | SOX2 |
| 277 | BTC | Enrichr CHEA_2022: SOX2 20726797 ChIP-Seq SW620 Human | SOX2 |
| 278 | TLCD1 | Enrichr CHEA_2022: SOX2 20726797 ChIP-Seq SW620 Human | SOX2 |
| 279 | FBXL2 | Enrichr CHEA_2022: SOX2 20726797 ChIP-Seq SW620 Human | SOX2 |
| 280 | SDC1 | Enrichr CHEA_2022: SOX2 20726797 ChIP-Seq SW620 Human | SOX2 |
| 281 | MON1B | Enrichr CHEA_2022: SOX2 20726797 ChIP-Seq SW620 Human | SOX2 |
| 282 | APOBEC3B | Enrichr CHEA_2022: SOX2 20726797 ChIP-Seq SW620 Human | SOX2 |
| 283 | ATP6V1B2 | Enrichr CHEA_2022: SOX2 20726797 ChIP-Seq SW620 Human | SOX2 |
| 284 | APOBEC3C | Enrichr CHEA_2022: SOX2 20726797 ChIP-Seq SW620 Human | SOX2 |
| 285 | LTV1 | Enrichr CHEA_2022: SOX2 20726797 ChIP-Seq SW620 Human | SOX2 |
| 286 | SLC45A4 | Enrichr CHEA_2022: SOX2 20726797 ChIP-Seq SW620 Human | SOX2 |
| 287 | WFDC2 | Enrichr CHEA_2022: SOX2 20726797 ChIP-Seq SW620 Human | SOX2 |
| 288 | MT1X | Enrichr CHEA_2022: SOX2 20726797 ChIP-Seq SW620 Human | SOX2 |
| 289 | ICAM2 | Enrichr CHEA_2022: SOX2 20726797 ChIP-Seq SW620 Human | SOX2 |
| 290 | LGALS4 | Enrichr CHEA_2022: SOX2 20726797 ChIP-Seq SW620 Human | SOX2 |
| 291 | MT2A | Enrichr CHEA_2022: SOX2 20726797 ChIP-Seq SW620 Human | SOX2 |
| 292 | CCDC92 | Enrichr CHEA_2022: SOX2 20726797 ChIP-Seq SW620 Human | SOX2 |
| 293 | LGALS1 | Enrichr CHEA_2022: SOX2 20726797 ChIP-Seq SW620 Human | SOX2 |
| 294 | REG4 | Enrichr CHEA_2022: SOX2 20726797 ChIP-Seq SW620 Human | SOX2 |
| 295 | TNFSF10 | Enrichr CHEA_2022: SOX2 20726797 ChIP-Seq SW620 Human | SOX2 |
| 296 | LGALS8 | Enrichr CHEA_2022: SOX2 20726797 ChIP-Seq SW620 Human | SOX2 |
| 297 | CCDC96 | Enrichr CHEA_2022: SOX2 20726797 ChIP-Seq SW620 Human | SOX2 |
| 298 | SPA17 | Enrichr CHEA_2022: SOX2 20726797 ChIP-Seq SW620 Human | SOX2 |
| 299 | GABBR1 | Enrichr CHEA_2022: SOX2 20726797 ChIP-Seq SW620 Human | SOX2 |
| 300 | IL15 | Enrichr CHEA_2022: SOX2 20726797 ChIP-Seq SW620 Human | SOX2 |
| 301 | SLC10A3 | Enrichr CHEA_2022: SOX2 20726797 ChIP-Seq SW620 Human | SOX2 |
| 302 | GAB2 | Enrichr CHEA_2022: SOX2 20726797 ChIP-Seq SW620 Human | SOX2 |
| 303 | GAB1 | Enrichr CHEA_2022: SOX2 20726797 ChIP-Seq SW620 Human | SOX2 |

| | | | |
|---|---|---|---|
| 304 | KLHL3 | Enrichr CHEA_2022: SOX2 20726797 ChIP-Seq SW620 Human | SOX2 |
| 305 | KLHL5 | Enrichr CHEA_2022: SOX2 20726797 ChIP-Seq SW620 Human | SOX2 |
| 306 | ZHX1 | Enrichr CHEA_2022: SOX2 20726797 ChIP-Seq SW620 Human | SOX2 |
| 307 | RAP2A | Enrichr CHEA_2022: SOX2 20726797 ChIP-Seq SW620 Human | SOX2 |
| 308 | CTTNBP2NL | Enrichr CHEA_2022: SOX2 20726797 ChIP-Seq SW620 Human | SOX2 |
| 309 | BST2 | Enrichr CHEA_2022: SOX2 20726797 ChIP-Seq SW620 Human | SOX2 |
| 310 | DEPDC7 | Enrichr CHEA_2022: SOX2 20726797 ChIP-Seq SW620 Human | SOX2 |
| 311 | KITLG | Enrichr CHEA_2022: SOX2 20726797 ChIP-Seq SW620 Human | SOX2 |
| 312 | ARHGEF4 | Enrichr CHEA_2022: SOX2 20726797 ChIP-Seq SW620 Human | SOX2 |
| 313 | ARHGEF3 | Enrichr CHEA_2022: SOX2 20726797 ChIP-Seq SW620 Human | SOX2 |
| 314 | FJX1 | Enrichr CHEA_2022: SOX2 20726797 ChIP-Seq SW620 Human | SOX2 |
| 315 | RANBP10 | Enrichr CHEA_2022: SOX2 20726797 ChIP-Seq SW620 Human | SOX2 |
| 316 | NRN1 | Enrichr CHEA_2022: SOX2 20726797 ChIP-Seq SW620 Human | SOX2 |
| 317 | NOXO1 | Enrichr CHEA_2022: SOX2 20726797 ChIP-Seq SW620 Human | SOX2 |
| 318 | PPCS | Enrichr CHEA_2022: SOX2 20726797 ChIP-Seq SW620 Human | SOX2 |
| 319 | TFB1M | Enrichr CHEA_2022: SOX2 20726797 ChIP-Seq SW620 Human | SOX2 |
| 320 | KALRN | Enrichr CHEA_2022: SOX2 20726797 ChIP-Seq SW620 Human | SOX2 |
| 321 | PPP3CA | Enrichr CHEA_2022: SOX2 20726797 ChIP-Seq SW620 Human | SOX2 |
| 322 | SEC14L4 | Enrichr CHEA_2022: SOX2 20726797 ChIP-Seq SW620 Human | SOX2 |
| 323 | RRAS | Enrichr CHEA_2022: SOX2 20726797 ChIP-Seq SW620 Human | SOX2 |
| 324 | SMYD3 | Enrichr CHEA_2022: SOX2 20726797 ChIP-Seq SW620 Human | SOX2 |
| 325 | HTATIP2 | Enrichr CHEA_2022: SOX2 20726797 ChIP-Seq SW620 Human | SOX2 |
| 326 | PHGDH | Enrichr CHEA_2022: SOX2 20726797 ChIP-Seq SW620 Human | SOX2 |
| 327 | NFIL3 | Enrichr CHEA_2022: SOX2 20726797 ChIP-Seq SW620 Human | SOX2 |
| 328 | PCK2 | Enrichr CHEA_2022: SOX2 20726797 ChIP-Seq SW620 Human | SOX2 |
| 329 | LYAR | Enrichr CHEA_2022: SOX2 20726797 ChIP-Seq SW620 Human | SOX2 |
| 330 | CAP2 | Enrichr CHEA_2022: SOX2 20726797 ChIP-Seq SW620 Human | SOX2 |
| 331 | GAL | Enrichr CHEA_2022: SOX2 20726797 ChIP-Seq SW620 Human | SOX2 |
| 332 | TPST2 | Enrichr CHEA_2022: SOX2 20726797 ChIP-Seq SW620 Human | SOX2 |
| 333 | TTC9 | Enrichr CHEA_2022: SOX2 20726797 ChIP-Seq SW620 Human | SOX2 |
| 334 | TTC8 | Enrichr CHEA_2022: SOX2 20726797 ChIP-Seq SW620 Human | SOX2 |
| 335 | KCNQ1 | Enrichr CHEA_2022: SOX2 20726797 ChIP-Seq SW620 Human | SOX2 |
| 336 | TPST1 | Enrichr CHEA_2022: SOX2 20726797 ChIP-Seq SW620 Human | SOX2 |
| 337 | TCEA2 | Enrichr CHEA_2022: SOX2 20726797 ChIP-Seq SW620 Human | SOX2 |
| 338 | TACC1 | Enrichr CHEA_2022: SOX2 20726797 ChIP-Seq SW620 Human | SOX2 |
| 339 | PNPLA3 | Enrichr CHEA_2022: SOX2 20726797 ChIP-Seq SW620 Human | SOX2 |
| 340 | NRP2 | Enrichr CHEA_2022: SOX2 20726797 ChIP-Seq SW620 Human | SOX2 |
| 341 | CAPG | Enrichr CHEA_2022: SOX2 20726797 ChIP-Seq SW620 Human | SOX2 |
| 342 | NUDT4 | Enrichr CHEA_2022: SOX2 20726797 ChIP-Seq SW620 Human | SOX2 |
| 343 | SPAG16 | Enrichr CHEA_2022: SOX2 20726797 ChIP-Seq SW620 Human | SOX2 |
| 344 | HERPUD2 | Enrichr CHEA_2022: SOX2 20726797 ChIP-Seq SW620 Human | SOX2 |
| 345 | AKAP13 | Enrichr CHEA_2022: SOX2 20726797 ChIP-Seq SW620 Human | SOX2 |
| 346 | AKAP11 | Enrichr CHEA_2022: SOX2 20726797 ChIP-Seq SW620 Human | SOX2 |
| 347 | PCM1 | Enrichr CHEA_2022: SOX2 20726797 ChIP-Seq SW620 Human | SOX2 |
| 348 | AKAP12 | Enrichr CHEA_2022: SOX2 20726797 ChIP-Seq SW620 Human | SOX2 |
| 349 | FAM3C | Enrichr CHEA_2022: SOX2 20726797 ChIP-Seq SW620 Human | SOX2 |
| 350 | FAM3B | Enrichr CHEA_2022: SOX2 20726797 ChIP-Seq SW620 Human | SOX2 |
| 351 | FAM3D | Enrichr CHEA_2022: SOX2 20726797 ChIP-Seq SW620 Human | SOX2 |
| 352 | RARRES1 | Enrichr CHEA_2022: SOX2 20726797 ChIP-Seq SW620 Human | SOX2 |
| 353 | SPINK5 | Enrichr CHEA_2022: SOX2 20726797 ChIP-Seq SW620 Human | SOX2 |
| 354 | HS3ST1 | Enrichr CHEA_2022: SOX2 20726797 ChIP-Seq SW620 Human | SOX2 |
| 355 | UBE2H | Enrichr CHEA_2022: SOX2 20726797 ChIP-Seq SW620 Human | SOX2 |
| 356 | MORC4 | Enrichr CHEA_2022: SOX2 20726797 ChIP-Seq SW620 Human | SOX2 |
| 357 | FAM3A | Enrichr CHEA_2022: SOX2 20726797 ChIP-Seq SW620 Human | SOX2 |
| 358 | CACHD1 | Enrichr CHEA_2022: SOX2 20726797 ChIP-Seq SW620 Human | SOX2 |
| 359 | DNAJB14 | Enrichr CHEA_2022: SOX2 20726797 ChIP-Seq SW620 Human | SOX2 |
| 360 | BSPRY | Enrichr CHEA_2022: SOX2 20726797 ChIP-Seq SW620 Human | SOX2 |
| 361 | CDKAL1 | Enrichr CHEA_2022: SOX2 20726797 ChIP-Seq SW620 Human | SOX2 |
| 362 | YPEL2 | Enrichr CHEA_2022: SOX2 20726797 ChIP-Seq SW620 Human | SOX2 |
| 363 | WDFY2 | Enrichr CHEA_2022: SOX2 20726797 ChIP-Seq SW620 Human | SOX2 |
| 364 | DEGS2 | Enrichr CHEA_2022: SOX2 20726797 ChIP-Seq SW620 Human | SOX2 |

| | | | |
|---|---|---|---|
| 365 | WDFY3 | Enrichr CHEA_2022: SOX2 20726797 ChIP-Seq SW620 Human | SOX2 |
| 366 | OTUD1 | Enrichr CHEA_2022: SOX2 20726797 ChIP-Seq SW620 Human | SOX2 |
| 367 | PPIC | Enrichr CHEA_2022: SOX2 20726797 ChIP-Seq SW620 Human | SOX2 |
| 368 | PPID | Enrichr CHEA_2022: SOX2 20726797 ChIP-Seq SW620 Human | SOX2 |
| 369 | GDA | Enrichr CHEA_2022: SOX2 20726797 ChIP-Seq SW620 Human | SOX2 |
| 370 | NAPA | Enrichr CHEA_2022: SOX2 20726797 ChIP-Seq SW620 Human | SOX2 |
| 371 | SCNN1A | Enrichr CHEA_2022: SOX2 20726797 ChIP-Seq SW620 Human | SOX2 |
| 372 | PRDM1 | Enrichr CHEA_2022: SOX2 20726797 ChIP-Seq SW620 Human | SOX2 |
| 373 | LOXL4 | Enrichr CHEA_2022: SOX2 20726797 ChIP-Seq SW620 Human | SOX2 |
| 374 | LOXL2 | Enrichr CHEA_2022: SOX2 20726797 ChIP-Seq SW620 Human | SOX2 |
| 375 | LOXL3 | Enrichr CHEA_2022: SOX2 20726797 ChIP-Seq SW620 Human | SOX2 |
| 376 | HSD17B1 | Enrichr CHEA_2022: SOX2 20726797 ChIP-Seq SW620 Human | SOX2 |
| 377 | HSD17B2 | Enrichr CHEA_2022: SOX2 20726797 ChIP-Seq SW620 Human | SOX2 |
| 378 | MSX1 | Enrichr CHEA_2022: SOX2 20726797 ChIP-Seq SW620 Human | SOX2 |
| 379 | TBC1D16 | Enrichr CHEA_2022: SOX2 20726797 ChIP-Seq SW620 Human | SOX2 |
| 380 | TBC1D17 | Enrichr CHEA_2022: SOX2 20726797 ChIP-Seq SW620 Human | SOX2 |
| 381 | DENND2D | Enrichr CHEA_2022: SOX2 20726797 ChIP-Seq SW620 Human | SOX2 |
| 382 | PCYT1B | Enrichr CHEA_2022: SOX2 20726797 ChIP-Seq SW620 Human | SOX2 |
| 383 | SYT1 | Enrichr CHEA_2022: SOX2 20726797 ChIP-Seq SW620 Human | SOX2 |
| 384 | TIMM44 | Enrichr CHEA_2022: SOX2 20726797 ChIP-Seq SW620 Human | SOX2 |
| 385 | NAT1 | Enrichr CHEA_2022: SOX2 20726797 ChIP-Seq SW620 Human | SOX2 |
| 386 | CCNB1IP1 | Enrichr CHEA_2022: SOX2 20726797 ChIP-Seq SW620 Human | SOX2 |
| 387 | FGF18 | Enrichr CHEA_2022: SOX2 20726797 ChIP-Seq SW620 Human | SOX2 |
| 388 | ZNF91 | Enrichr CHEA_2022: SOX2 20726797 ChIP-Seq SW620 Human | SOX2 |
| 389 | GNB2 | Enrichr CHEA_2022: SOX2 20726797 ChIP-Seq SW620 Human | SOX2 |
| 390 | MKRN2 | Enrichr CHEA_2022: SOX2 20726797 ChIP-Seq SW620 Human | SOX2 |
| 391 | GNB5 | Enrichr CHEA_2022: SOX2 20726797 ChIP-Seq SW620 Human | SOX2 |
| 392 | GNAS | Enrichr CHEA_2022: SOX2 20726797 ChIP-Seq SW620 Human | SOX2 |
| 393 | MUC20 | Enrichr CHEA_2022: SOX2 20726797 ChIP-Seq SW620 Human | SOX2 |
| 394 | TCF3 | Enrichr CHEA_2022: SOX2 20726797 ChIP-Seq SW620 Human | SOX2 |
| 395 | TSNARE1 | Enrichr CHEA_2022: SOX2 20726797 ChIP-Seq SW620 Human | SOX2 |
| 396 | VIL1 | Enrichr CHEA_2022: SOX2 20726797 ChIP-Seq SW620 Human | SOX2 |
| 397 | DYRK4 | Enrichr CHEA_2022: SOX2 20726797 ChIP-Seq SW620 Human | SOX2 |
| 398 | DYRK3 | Enrichr CHEA_2022: SOX2 20726797 ChIP-Seq SW620 Human | SOX2 |
| 399 | DYRK2 | Enrichr CHEA_2022: SOX2 20726797 ChIP-Seq SW620 Human | SOX2 |
| 400 | SLC41A2 | Enrichr CHEA_2022: SOX2 20726797 ChIP-Seq SW620 Human | SOX2 |
| 401 | PLAU | Enrichr CHEA_2022: SOX2 20726797 ChIP-Seq SW620 Human | SOX2 |
| 402 | HTRA1 | Enrichr CHEA_2022: SOX2 20726797 ChIP-Seq SW620 Human | SOX2 |
| 403 | ETS1 | Enrichr CHEA_2022: SOX2 20726797 ChIP-Seq SW620 Human | SOX2 |
| 404 | ETS2 | Enrichr CHEA_2022: SOX2 20726797 ChIP-Seq SW620 Human | SOX2 |
| 405 | DAPP1 | Enrichr CHEA_2022: SOX2 20726797 ChIP-Seq SW620 Human | SOX2 |
| 406 | IGF1R | Enrichr CHEA_2022: SOX2 20726797 ChIP-Seq SW620 Human | SOX2 |
| 407 | TGM2 | Enrichr CHEA_2022: SOX2 20726797 ChIP-Seq SW620 Human | SOX2 |
| 408 | CNN3 | Enrichr CHEA_2022: SOX2 20726797 ChIP-Seq SW620 Human | SOX2 |
| 409 | ARFGEF1 | Enrichr CHEA_2022: SOX2 20726797 ChIP-Seq SW620 Human | SOX2 |
| 410 | GGH | Enrichr CHEA_2022: SOX2 20726797 ChIP-Seq SW620 Human | SOX2 |
| 411 | CAV1 | Enrichr CHEA_2022: SOX2 20726797 ChIP-Seq SW620 Human | SOX2 |
| 412 | NAV1 | Enrichr CHEA_2022: SOX2 20726797 ChIP-Seq SW620 Human | SOX2 |
| 413 | ETV1 | Enrichr CHEA_2022: SOX2 20726797 ChIP-Seq SW620 Human | SOX2 |
| 414 | NAV2 | Enrichr CHEA_2022: SOX2 20726797 ChIP-Seq SW620 Human | SOX2 |
| 415 | ETV5 | Enrichr CHEA_2022: SOX2 20726797 ChIP-Seq SW620 Human | SOX2 |
| 416 | TTC17 | Enrichr CHEA_2022: SOX2 20726797 ChIP-Seq SW620 Human | SOX2 |
| 417 | ETV6 | Enrichr CHEA_2022: SOX2 20726797 ChIP-Seq SW620 Human | SOX2 |
| 418 | TAPBP | Enrichr CHEA_2022: SOX2 20726797 ChIP-Seq SW620 Human | SOX2 |
| 419 | EPN3 | Enrichr CHEA_2022: SOX2 20726797 ChIP-Seq SW620 Human | SOX2 |
| 420 | ELF2 | Enrichr CHEA_2022: SOX2 20726797 ChIP-Seq SW620 Human | SOX2 |
| 421 | ELF3 | Enrichr CHEA_2022: SOX2 20726797 ChIP-Seq SW620 Human | SOX2 |
| 422 | MANSC1 | Enrichr CHEA_2022: SOX2 20726797 ChIP-Seq SW620 Human | SOX2 |
| 423 | CUEDC1 | Enrichr CHEA_2022: SOX2 20726797 ChIP-Seq SW620 Human | SOX2 |
| 424 | FBP1 | Enrichr CHEA_2022: SOX2 20726797 ChIP-Seq SW620 Human | SOX2 |
| 425 | SHANK2 | Enrichr CHEA_2022: SOX2 20726797 ChIP-Seq SW620 Human | SOX2 |

| | | | |
|---|---|---|---|
| 426 | TTC26 | Enrichr CHEA_2022: SOX2 20726797 ChIP-Seq SW620 Human | SOX2 |
| 427 | NNMT | Enrichr CHEA_2022: SOX2 20726797 ChIP-Seq SW620 Human | SOX2 |
| 428 | TNFAIP8 | Enrichr CHEA_2022: SOX2 20726797 ChIP-Seq SW620 Human | SOX2 |
| 429 | TNFAIP2 | Enrichr CHEA_2022: SOX2 20726797 ChIP-Seq SW620 Human | SOX2 |
| 430 | PITPNC1 | Enrichr CHEA_2022: SOX2 20726797 ChIP-Seq SW620 Human | SOX2 |
| 431 | AGPAT5 | Enrichr CHEA_2022: SOX2 20726797 ChIP-Seq SW620 Human | SOX2 |
| 432 | CA2 | Enrichr CHEA_2022: SOX2 20726797 ChIP-Seq SW620 Human | SOX2 |
| 433 | IRAK1 | Enrichr CHEA_2022: SOX2 20726797 ChIP-Seq SW620 Human | SOX2 |
| 434 | LPXN | Enrichr CHEA_2022: SOX2 20726797 ChIP-Seq SW620 Human | SOX2 |
| 435 | ATF7IP | Enrichr CHEA_2022: SOX2 20726797 ChIP-Seq SW620 Human | SOX2 |
| 436 | DAPK1 | Enrichr CHEA_2022: SOX2 20726797 ChIP-Seq SW620 Human | SOX2 |
| 437 | ERCC1 | Enrichr CHEA_2022: SOX2 20726797 ChIP-Seq SW620 Human | SOX2 |
| 438 | GALE | Enrichr CHEA_2022: SOX2 20726797 ChIP-Seq SW620 Human | SOX2 |
| 439 | KRAS | Enrichr CHEA_2022: SOX2 20726797 ChIP-Seq SW620 Human | SOX2 |
| 440 | TRIB2 | Enrichr CHEA_2022: SOX2 20726797 ChIP-Seq SW620 Human | SOX2 |
| 441 | TRIB3 | Enrichr CHEA_2022: SOX2 20726797 ChIP-Seq SW620 Human | SOX2 |
| 442 | SIGIRR | Enrichr CHEA_2022: SOX2 20726797 ChIP-Seq SW620 Human | SOX2 |
| 443 | TRIB1 | Enrichr CHEA_2022: SOX2 20726797 ChIP-Seq SW620 Human | SOX2 |
| 444 | RND3 | Enrichr CHEA_2022: SOX2 20726797 ChIP-Seq SW620 Human | SOX2 |
| 445 | NR3C1 | Enrichr CHEA_2022: SOX2 20726797 ChIP-Seq SW620 Human | SOX2 |
| 446 | POLD4 | Enrichr CHEA_2022: SOX2 20726797 ChIP-Seq SW620 Human | SOX2 |
| 447 | ACOXL | Enrichr CHEA_2022: SOX2 20726797 ChIP-Seq SW620 Human | SOX2 |
| 448 | OXCT1 | Enrichr CHEA_2022: SOX2 20726797 ChIP-Seq SW620 Human | SOX2 |
| 449 | DIP2C | Enrichr CHEA_2022: SOX2 20726797 ChIP-Seq SW620 Human | SOX2 |
| 450 | HOMER3 | Enrichr CHEA_2022: SOX2 20726797 ChIP-Seq SW620 Human | SOX2 |
| 451 | PCGF6 | Enrichr CHEA_2022: SOX2 20726797 ChIP-Seq SW620 Human | SOX2 |
| 452 | HUS1 | Enrichr CHEA_2022: SOX2 20726797 ChIP-Seq SW620 Human | SOX2 |
| 453 | NUP93 | Enrichr CHEA_2022: SOX2 20726797 ChIP-Seq SW620 Human | SOX2 |
| 454 | CAPN12 | Enrichr CHEA_2022: SOX2 20726797 ChIP-Seq SW620 Human | SOX2 |
| 455 | DEPDC1B | Enrichr CHEA_2022: SOX2 20726797 ChIP-Seq SW620 Human | SOX2 |
| 456 | TEF | Enrichr CHEA_2022: SOX2 20726797 ChIP-Seq SW620 Human | SOX2 |
| 457 | RALB | Enrichr CHEA_2022: SOX2 20726797 ChIP-Seq SW620 Human | SOX2 |
| 458 | ZNF395 | Enrichr CHEA_2022: SOX2 20726797 ChIP-Seq SW620 Human | SOX2 |
| 459 | ADARB1 | Enrichr CHEA_2022: SOX2 20726797 ChIP-Seq SW620 Human | SOX2 |
| 460 | GLS | Enrichr CHEA_2022: SOX2 20726797 ChIP-Seq SW620 Human | SOX2 |
| 461 | ASCL2 | Enrichr CHEA_2022: SOX2 20726797 ChIP-Seq SW620 Human | SOX2 |
| 462 | ADCY7 | Enrichr CHEA_2022: SOX2 20726797 ChIP-Seq SW620 Human | SOX2 |
| 463 | GNG2 | Enrichr CHEA_2022: SOX2 20726797 ChIP-Seq SW620 Human | SOX2 |
| 464 | APOL6 | Enrichr CHEA_2022: SOX2 20726797 ChIP-Seq SW620 Human | SOX2 |
| 465 | GNG4 | Enrichr CHEA_2022: SOX2 20726797 ChIP-Seq SW620 Human | SOX2 |
| 466 | FLRT3 | Enrichr CHEA_2022: SOX2 20726797 ChIP-Seq SW620 Human | SOX2 |
| 467 | WDR72 | Enrichr CHEA_2022: SOX2 20726797 ChIP-Seq SW620 Human | SOX2 |
| 468 | ALOX5 | Enrichr CHEA_2022: SOX2 20726797 ChIP-Seq SW620 Human | SOX2 |
| 469 | ST3GAL5 | Enrichr CHEA_2022: SOX2 20726797 ChIP-Seq SW620 Human | SOX2 |
| 470 | PRSS8 | Enrichr CHEA_2022: SOX2 20726797 ChIP-Seq SW620 Human | SOX2 |
| 471 | LUC7L | Enrichr CHEA_2022: SOX2 20726797 ChIP-Seq SW620 Human | SOX2 |
| 472 | NUP88 | Enrichr CHEA_2022: SOX2 20726797 ChIP-Seq SW620 Human | SOX2 |
| 473 | ST3GAL1 | Enrichr CHEA_2022: SOX2 20726797 ChIP-Seq SW620 Human | SOX2 |
| 474 | PRSS3 | Enrichr CHEA_2022: SOX2 20726797 ChIP-Seq SW620 Human | SOX2 |
| 475 | APOL1 | Enrichr CHEA_2022: SOX2 20726797 ChIP-Seq SW620 Human | SOX2 |
| 476 | FDFT1 | Enrichr CHEA_2022: SOX2 20726797 ChIP-Seq SW620 Human | SOX2 |
| 477 | ST3GAL4 | Enrichr CHEA_2022: SOX2 20726797 ChIP-Seq SW620 Human | SOX2 |
| 478 | TAF9B | Enrichr CHEA_2022: SOX2 20726797 ChIP-Seq SW620 Human | SOX2 |
| 479 | PRKCH | Enrichr CHEA_2022: SOX2 20726797 ChIP-Seq SW620 Human | SOX2 |
| 480 | NDFIP1 | Enrichr CHEA_2022: SOX2 20726797 ChIP-Seq SW620 Human | SOX2 |
| 481 | PHC3 | Enrichr CHEA_2022: SOX2 20726797 ChIP-Seq SW620 Human | SOX2 |
| 482 | ICMT | Enrichr CHEA_2022: SOX2 20726797 ChIP-Seq SW620 Human | SOX2 |
| 483 | WDR59 | Enrichr CHEA_2022: SOX2 20726797 ChIP-Seq SW620 Human | SOX2 |
| 484 | STARD13 | Enrichr CHEA_2022: SOX2 20726797 ChIP-Seq SW620 Human | SOX2 |
| 485 | STARD10 | Enrichr CHEA_2022: SOX2 20726797 ChIP-Seq SW620 Human | SOX2 |
| 486 | PLK2 | Enrichr CHEA_2022: SOX2 20726797 ChIP-Seq SW620 Human | SOX2 |

| | | | |
|---|---|---|---|
| 487 | PRKCE | Enrichr CHEA_2022: SOX2 20726797 ChIP-Seq SW620 Human | SOX2 |
| 488 | PARD3 | Enrichr CHEA_2022: SOX2 20726797 ChIP-Seq SW620 Human | SOX2 |
| 489 | PRKCA | Enrichr CHEA_2022: SOX2 20726797 ChIP-Seq SW620 Human | SOX2 |
| 490 | TERF2 | Enrichr CHEA_2022: SOX2 20726797 ChIP-Seq SW620 Human | SOX2 |
| 491 | ELL2 | Enrichr CHEA_2022: SOX2 20726797 ChIP-Seq SW620 Human | SOX2 |
| 492 | ELL3 | Enrichr CHEA_2022: SOX2 20726797 ChIP-Seq SW620 Human | SOX2 |
| 493 | COQ10A | Enrichr CHEA_2022: SOX2 20726797 ChIP-Seq SW620 Human | SOX2 |
| 494 | EHD2 | Enrichr CHEA_2022: SOX2 20726797 ChIP-Seq SW620 Human | SOX2 |
| 495 | SQLE | Enrichr CHEA_2022: SOX2 20726797 ChIP-Seq SW620 Human | SOX2 |
| 496 | GJB3 | Enrichr CHEA_2022: SOX2 20726797 ChIP-Seq SW620 Human | SOX2 |
| 497 | GJB2 | Enrichr CHEA_2022: SOX2 20726797 ChIP-Seq SW620 Human | SOX2 |
| 498 | TCN1 | Enrichr CHEA_2022: SOX2 20726797 ChIP-Seq SW620 Human | SOX2 |
| 499 | SNAI2 | Enrichr CHEA_2022: SOX2 20726797 ChIP-Seq SW620 Human | SOX2 |
| 500 | PRKD2 | Enrichr CHEA_2022: SOX2 20726797 ChIP-Seq SW620 Human | SOX2 |
| 501 | GNE | Enrichr CHEA_2022: SOX2 20726797 ChIP-Seq SW620 Human | SOX2 |
| 502 | CFB | Enrichr CHEA_2022: SOX2 20726797 ChIP-Seq SW620 Human | SOX2 |
| 503 | FANK1 | Enrichr CHEA_2022: SOX2 20726797 ChIP-Seq SW620 Human | SOX2 |
| 504 | CFH | Enrichr CHEA_2022: SOX2 20726797 ChIP-Seq SW620 Human | SOX2 |
| 505 | PHYHIPL | Enrichr CHEA_2022: SOX2 20726797 ChIP-Seq SW620 Human | SOX2 |
| 506 | GNS | Enrichr CHEA_2022: SOX2 20726797 ChIP-Seq SW620 Human | SOX2 |
| 507 | ELK3 | Enrichr CHEA_2022: SOX2 20726797 ChIP-Seq SW620 Human | SOX2 |
| 508 | PLAC8 | Enrichr CHEA_2022: SOX2 20726797 ChIP-Seq SW620 Human | SOX2 |
| 509 | NUP62 | Enrichr CHEA_2022: SOX2 20726797 ChIP-Seq SW620 Human | SOX2 |
| 510 | AP1S3 | Enrichr CHEA_2022: SOX2 20726797 ChIP-Seq SW620 Human | SOX2 |
| 511 | CEP72 | Enrichr CHEA_2022: SOX2 20726797 ChIP-Seq SW620 Human | SOX2 |
| 512 | AP1S2 | Enrichr CHEA_2022: SOX2 20726797 ChIP-Seq SW620 Human | SOX2 |
| 513 | PROM2 | Enrichr CHEA_2022: SOX2 20726797 ChIP-Seq SW620 Human | SOX2 |
| 514 | PROM1 | Enrichr CHEA_2022: SOX2 20726797 ChIP-Seq SW620 Human | SOX2 |
| 515 | ZNF367 | Enrichr CHEA_2022: SOX2 20726797 ChIP-Seq SW620 Human | SOX2 |
| 516 | SHPRH | Enrichr CHEA_2022: SOX2 20726797 ChIP-Seq SW620 Human | SOX2 |
| 517 | PHF10 | Enrichr CHEA_2022: SOX2 20726797 ChIP-Seq SW620 Human | SOX2 |
| 518 | PHF13 | Enrichr CHEA_2022: SOX2 20726797 ChIP-Seq SW620 Human | SOX2 |
| 519 | GRTP1 | Enrichr CHEA_2022: SOX2 20726797 ChIP-Seq SW620 Human | SOX2 |
| 520 | CGN | Enrichr CHEA_2022: SOX2 20726797 ChIP-Seq SW620 Human | SOX2 |
| 521 | TCP1 | Enrichr CHEA_2022: SOX2 20726797 ChIP-Seq SW620 Human | SOX2 |
| 522 | NFATC3 | Enrichr CHEA_2022: SOX2 20726797 ChIP-Seq SW620 Human | SOX2 |
| 523 | ASH1L | Enrichr CHEA_2022: SOX2 20726797 ChIP-Seq SW620 Human | SOX2 |
| 524 | WDR35 | Enrichr CHEA_2022: SOX2 20726797 ChIP-Seq SW620 Human | SOX2 |
| 525 | TUBB2B | Enrichr CHEA_2022: SOX2 20726797 ChIP-Seq SW620 Human | SOX2 |
| 526 | HLA-DQB1 | Enrichr CHEA_2022: SOX2 20726797 ChIP-Seq SW620 Human | SOX2 |
| 527 | PHF14 | Enrichr CHEA_2022: SOX2 20726797 ChIP-Seq SW620 Human | SOX2 |
| 528 | RARA | Enrichr CHEA_2022: SOX2 20726797 ChIP-Seq SW620 Human | SOX2 |
| 529 | MGLL | Enrichr CHEA_2022: SOX2 20726797 ChIP-Seq SW620 Human | SOX2 |
| 530 | GAS6 | Enrichr CHEA_2022: SOX2 20726797 ChIP-Seq SW620 Human | SOX2 |
| 531 | BTG3 | Enrichr CHEA_2022: SOX2 20726797 ChIP-Seq SW620 Human | SOX2 |
| 532 | BTG2 | Enrichr CHEA_2022: SOX2 20726797 ChIP-Seq SW620 Human | SOX2 |
| 533 | BTG1 | Enrichr CHEA_2022: SOX2 20726797 ChIP-Seq SW620 Human | SOX2 |
| 534 | PRDM11 | Enrichr CHEA_2022: SOX2 20726797 ChIP-Seq SW620 Human | SOX2 |
| 535 | TIMP4 | Enrichr CHEA_2022: SOX2 20726797 ChIP-Seq SW620 Human | SOX2 |
| 536 | PBK | Enrichr CHEA_2022: SOX2 20726797 ChIP-Seq SW620 Human | SOX2 |
| 537 | PLLP | Enrichr CHEA_2022: SOX2 20726797 ChIP-Seq SW620 Human | SOX2 |
| 538 | TIMP2 | Enrichr CHEA_2022: SOX2 20726797 ChIP-Seq SW620 Human | SOX2 |
| 539 | TEAD2 | Enrichr CHEA_2022: SOX2 20726797 ChIP-Seq SW620 Human | SOX2 |
| 540 | TEAD3 | Enrichr CHEA_2022: SOX2 20726797 ChIP-Seq SW620 Human | SOX2 |
| 541 | AP1M2 | Enrichr CHEA_2022: SOX2 20726797 ChIP-Seq SW620 Human | SOX2 |
| 542 | TEAD4 | Enrichr CHEA_2022: SOX2 20726797 ChIP-Seq SW620 Human | SOX2 |
| 543 | CEP57 | Enrichr CHEA_2022: SOX2 20726797 ChIP-Seq SW620 Human | SOX2 |
| 544 | FARP2 | Enrichr CHEA_2022: SOX2 20726797 ChIP-Seq SW620 Human | SOX2 |
| 545 | CDKN2C | Enrichr CHEA_2022: SOX2 20726797 ChIP-Seq SW620 Human | SOX2 |
| 546 | MBNL2 | Enrichr CHEA_2022: SOX2 20726797 ChIP-Seq SW620 Human | SOX2 |
| 547 | YTHDF3 | Enrichr CHEA_2022: SOX2 20726797 ChIP-Seq SW620 Human | SOX2 |

| | | | |
|---|---|---|---|
| 548 | MGMT | Enrichr CHEA_2022: SOX2 20726797 ChIP-Seq SW620 Human | SOX2 |
| 549 | CDKN2B | Enrichr CHEA_2022: SOX2 20726797 ChIP-Seq SW620 Human | SOX2 |
| 550 | G0S2 | Enrichr CHEA_2022: SOX2 20726797 ChIP-Seq SW620 Human | SOX2 |
| 551 | NR1H3 | Enrichr CHEA_2022: SOX2 20726797 ChIP-Seq SW620 Human | SOX2 |
| 552 | ELP3 | Enrichr CHEA_2022: SOX2 20726797 ChIP-Seq SW620 Human | SOX2 |
| 553 | NFIA | Enrichr CHEA_2022: SOX2 20726797 ChIP-Seq SW620 Human | SOX2 |
| 554 | DHRS2 | Enrichr CHEA_2022: SOX2 20726797 ChIP-Seq SW620 Human | SOX2 |
| 555 | DHRS3 | Enrichr CHEA_2022: SOX2 20726797 ChIP-Seq SW620 Human | SOX2 |
| 556 | FARP1 | Enrichr CHEA_2022: SOX2 20726797 ChIP-Seq SW620 Human | SOX2 |
| 557 | ZNF33B | Enrichr CHEA_2022: SOX2 20726797 ChIP-Seq SW620 Human | SOX2 |
| 558 | DHRS9 | Enrichr CHEA_2022: SOX2 20726797 ChIP-Seq SW620 Human | SOX2 |
| 559 | NFIB | Enrichr CHEA_2022: SOX2 20726797 ChIP-Seq SW620 Human | SOX2 |
| 560 | PLEKHO1 | Enrichr CHEA_2022: SOX2 20726797 ChIP-Seq SW620 Human | SOX2 |
| 561 | SMOC2 | Enrichr CHEA_2022: SOX2 20726797 ChIP-Seq SW620 Human | SOX2 |
| 562 | MAP1B | Enrichr CHEA_2022: SOX2 20726797 ChIP-Seq SW620 Human | SOX2 |
| 563 | NUCB2 | Enrichr CHEA_2022: SOX2 20726797 ChIP-Seq SW620 Human | SOX2 |
| 564 | ZNF331 | Enrichr CHEA_2022: SOX2 20726797 ChIP-Seq SW620 Human | SOX2 |
| 565 | PHF6 | Enrichr CHEA_2022: SOX2 20726797 ChIP-Seq SW620 Human | SOX2 |
| 566 | CDKN1C | Enrichr CHEA_2022: SOX2 20726797 ChIP-Seq SW620 Human | SOX2 |
| 567 | GRN | Enrichr CHEA_2022: SOX2 20726797 ChIP-Seq SW620 Human | SOX2 |
| 568 | PYGB | Enrichr CHEA_2022: SOX2 20726797 ChIP-Seq SW620 Human | SOX2 |
| 569 | SETD7 | Enrichr CHEA_2022: SOX2 20726797 ChIP-Seq SW620 Human | SOX2 |
| 570 | PYGL | Enrichr CHEA_2022: SOX2 20726797 ChIP-Seq SW620 Human | SOX2 |
| 571 | STON2 | Enrichr CHEA_2022: SOX2 20726797 ChIP-Seq SW620 Human | SOX2 |
| 572 | PDF | Enrichr CHEA_2022: SOX2 20726797 ChIP-Seq SW620 Human | SOX2 |
| 573 | PPP1R2 | Enrichr CHEA_2022: SOX2 20726797 ChIP-Seq SW620 Human | SOX2 |
| 574 | UBL3 | Enrichr CHEA_2022: SOX2 20726797 ChIP-Seq SW620 Human | SOX2 |
| 575 | ASH2L | Enrichr CHEA_2022: SOX2 20726797 ChIP-Seq SW620 Human | SOX2 |
| 576 | CKB | Enrichr CHEA_2022: SOX2 20726797 ChIP-Seq SW620 Human | SOX2 |
| 577 | SLC17A5 | Enrichr CHEA_2022: SOX2 20726797 ChIP-Seq SW620 Human | SOX2 |
| 578 | GSN | Enrichr CHEA_2022: SOX2 20726797 ChIP-Seq SW620 Human | SOX2 |
| 579 | TLE4 | Enrichr CHEA_2022: SOX2 20726797 ChIP-Seq SW620 Human | SOX2 |
| 580 | TLE1 | Enrichr CHEA_2022: SOX2 20726797 ChIP-Seq SW620 Human | SOX2 |
| 581 | TPM2 | Enrichr CHEA_2022: SOX2 20726797 ChIP-Seq SW620 Human | SOX2 |
| 582 | VRK2 | Enrichr CHEA_2022: SOX2 20726797 ChIP-Seq SW620 Human | SOX2 |
| 583 | VRK3 | Enrichr CHEA_2022: SOX2 20726797 ChIP-Seq SW620 Human | SOX2 |
| 584 | HIPK2 | Enrichr CHEA_2022: SOX2 20726797 ChIP-Seq SW620 Human | SOX2 |
| 585 | RAB30 | Enrichr CHEA_2022: SOX2 20726797 ChIP-Seq SW620 Human | SOX2 |
| 586 | NRBP2 | Enrichr CHEA_2022: SOX2 20726797 ChIP-Seq SW620 Human | SOX2 |
| 587 | FAM91A1 | Enrichr CHEA_2022: SOX2 20726797 ChIP-Seq SW620 Human | SOX2 |
| 588 | SLC7A6OS | Enrichr CHEA_2022: SOX2 20726797 ChIP-Seq SW620 Human | SOX2 |
| 589 | RAB31 | Enrichr CHEA_2022: SOX2 20726797 ChIP-Seq SW620 Human | SOX2 |
| 590 | PSAT1 | Enrichr CHEA_2022: SOX2 20726797 ChIP-Seq SW620 Human | SOX2 |
| 591 | PLXNB2 | Enrichr CHEA_2022: SOX2 20726797 ChIP-Seq SW620 Human | SOX2 |
| 592 | CSPG5 | Enrichr CHEA_2022: SOX2 20726797 ChIP-Seq SW620 Human | SOX2 |
| 593 | CNIH3 | Enrichr CHEA_2022: SOX2 20726797 ChIP-Seq SW620 Human | SOX2 |
| 594 | TCTA | Enrichr CHEA_2022: SOX2 20726797 ChIP-Seq SW620 Human | SOX2 |
| 595 | PPIL4 | Enrichr CHEA_2022: SOX2 20726797 ChIP-Seq SW620 Human | SOX2 |
| 596 | FKBP7 | Enrichr CHEA_2022: SOX2 20726797 ChIP-Seq SW620 Human | SOX2 |
| 597 | GOLGA8A | Enrichr CHEA_2022: SOX2 20726797 ChIP-Seq SW620 Human | SOX2 |
| 598 | GTF2I | Enrichr CHEA_2022: SOX2 20726797 ChIP-Seq SW620 Human | SOX2 |
| 599 | RNMT | Enrichr CHEA_2022: SOX2 20726797 ChIP-Seq SW620 Human | SOX2 |
| 600 | PROS1 | Enrichr CHEA_2022: SOX2 20726797 ChIP-Seq SW620 Human | SOX2 |
| 601 | TCEAL1 | Enrichr CHEA_2022: SOX2 20726797 ChIP-Seq SW620 Human | SOX2 |
| 602 | PAPSS1 | Enrichr CHEA_2022: SOX2 20726797 ChIP-Seq SW620 Human | SOX2 |
| 603 | PAPSS2 | Enrichr CHEA_2022: SOX2 20726797 ChIP-Seq SW620 Human | SOX2 |
| 604 | FGD5 | Enrichr CHEA_2022: SOX2 20726797 ChIP-Seq SW620 Human | SOX2 |
| 605 | TCEAL8 | Enrichr CHEA_2022: SOX2 20726797 ChIP-Seq SW620 Human | SOX2 |
| 606 | ABLIM1 | Enrichr CHEA_2022: SOX2 20726797 ChIP-Seq SW620 Human | SOX2 |
| 607 | ABLIM3 | Enrichr CHEA_2022: SOX2 20726797 ChIP-Seq SW620 Human | SOX2 |
| 608 | CLCC1 | Enrichr CHEA_2022: SOX2 20726797 ChIP-Seq SW620 Human | SOX2 |

| | | | |
|---|---|---|---|
| 609 | TCEAL4 | Enrichr CHEA_2022: SOX2 20726797 ChIP-Seq SW620 Human | SOX2 |
| 610 | RAB25 | Enrichr CHEA_2022: SOX2 20726797 ChIP-Seq SW620 Human | SOX2 |
| 611 | SLC39A6 | Enrichr CHEA_2022: SOX2 20726797 ChIP-Seq SW620 Human | SOX2 |
| 612 | TXNL4B | Enrichr CHEA_2022: SOX2 20726797 ChIP-Seq SW620 Human | SOX2 |
| 613 | ZNF302 | Enrichr CHEA_2022: SOX2 20726797 ChIP-Seq SW620 Human | SOX2 |
| 614 | SIAE | Enrichr CHEA_2022: SOX2 20726797 ChIP-Seq SW620 Human | SOX2 |
| 615 | KCMF1 | Enrichr CHEA_2022: SOX2 20726797 ChIP-Seq SW620 Human | SOX2 |
| 616 | SERF1A | Enrichr CHEA_2022: SOX2 20726797 ChIP-Seq SW620 Human | SOX2 |
| 617 | TRAPPC2 | Enrichr CHEA_2022: SOX2 20726797 ChIP-Seq SW620 Human | SOX2 |
| 618 | OSBPL5 | Enrichr CHEA_2022: SOX2 20726797 ChIP-Seq SW620 Human | SOX2 |
| 619 | FRMD4B | Enrichr CHEA_2022: SOX2 20726797 ChIP-Seq SW620 Human | SOX2 |
| 620 | TFDP1 | Enrichr CHEA_2022: SOX2 20726797 ChIP-Seq SW620 Human | SOX2 |
| 621 | FRMD4A | Enrichr CHEA_2022: SOX2 20726797 ChIP-Seq SW620 Human | SOX2 |
| 622 | TPP2 | Enrichr CHEA_2022: SOX2 20726797 ChIP-Seq SW620 Human | SOX2 |
| 623 | LHX6 | Enrichr CHEA_2022: SOX2 20726797 ChIP-Seq SW620 Human | SOX2 |
| 624 | RAB3D | Enrichr CHEA_2022: SOX2 20726797 ChIP-Seq SW620 Human | SOX2 |
| 625 | SLC26A2 | Enrichr CHEA_2022: SOX2 20726797 ChIP-Seq SW620 Human | SOX2 |
| 626 | CBFB | Enrichr CHEA_2022: SOX2 20726797 ChIP-Seq SW620 Human | SOX2 |
| 627 | NPEPPS | Enrichr CHEA_2022: SOX2 20726797 ChIP-Seq SW620 Human | SOX2 |
| 628 | CDC14B | Enrichr CHEA_2022: SOX2 20726797 ChIP-Seq SW620 Human | SOX2 |
| 629 | TM7SF2 | Enrichr CHEA_2022: SOX2 20726797 ChIP-Seq SW620 Human | SOX2 |
| 630 | TM7SF3 | Enrichr CHEA_2022: SOX2 20726797 ChIP-Seq SW620 Human | SOX2 |
| 631 | PLXNA3 | Enrichr CHEA_2022: SOX2 20726797 ChIP-Seq SW620 Human | SOX2 |
| 632 | PI3 | Enrichr CHEA_2022: SOX2 20726797 ChIP-Seq SW620 Human | SOX2 |
| 633 | PLXNA2 | Enrichr CHEA_2022: SOX2 20726797 ChIP-Seq SW620 Human | SOX2 |
| 634 | PAPOLA | Enrichr CHEA_2022: SOX2 20726797 ChIP-Seq SW620 Human | SOX2 |
| 635 | PLXNA1 | Enrichr CHEA_2022: SOX2 20726797 ChIP-Seq SW620 Human | SOX2 |
| 636 | PLS3 | Enrichr CHEA_2022: SOX2 20726797 ChIP-Seq SW620 Human | SOX2 |
| 637 | MAP2K6 | Enrichr CHEA_2022: SOX2 20726797 ChIP-Seq SW620 Human | SOX2 |
| 638 | CDADC1 | Enrichr CHEA_2022: SOX2 20726797 ChIP-Seq SW620 Human | SOX2 |
| 639 | RFC4 | Enrichr CHEA_2022: SOX2 20726797 ChIP-Seq SW620 Human | SOX2 |
| 640 | XCL1 | Enrichr CHEA_2022: SOX2 20726797 ChIP-Seq SW620 Human | SOX2 |
| 641 | KBTBD11 | Enrichr CHEA_2022: SOX2 20726797 ChIP-Seq SW620 Human | SOX2 |
| 642 | IRAK1BP1 | Enrichr CHEA_2022: SOX2 20726797 ChIP-Seq SW620 Human | SOX2 |
| 643 | DAB2 | Enrichr CHEA_2022: SOX2 20726797 ChIP-Seq SW620 Human | SOX2 |
| 644 | MFAP2 | Enrichr CHEA_2022: SOX2 20726797 ChIP-Seq SW620 Human | SOX2 |
| 645 | ESPL1 | Enrichr CHEA_2022: SOX2 20726797 ChIP-Seq SW620 Human | SOX2 |
| 646 | BCL6 | Enrichr CHEA_2022: SOX2 20726797 ChIP-Seq SW620 Human | SOX2 |
| 647 | DNAJC6 | Enrichr CHEA_2022: SOX2 20726797 ChIP-Seq SW620 Human | SOX2 |
| 648 | DNAJC5 | Enrichr CHEA_2022: SOX2 20726797 ChIP-Seq SW620 Human | SOX2 |
| 649 | PIR | Enrichr CHEA_2022: SOX2 20726797 ChIP-Seq SW620 Human | SOX2 |
| 650 | XCL2 | Enrichr CHEA_2022: SOX2 20726797 ChIP-Seq SW620 Human | SOX2 |
| 651 | LAT | Enrichr CHEA_2022: SOX2 20726797 ChIP-Seq SW620 Human | SOX2 |
| 652 | BMPR2 | Enrichr CHEA_2022: SOX2 20726797 ChIP-Seq SW620 Human | SOX2 |
| 653 | CCDC25 | Enrichr CHEA_2022: SOX2 20726797 ChIP-Seq SW620 Human | SOX2 |
| 654 | HMMR | Enrichr CHEA_2022: SOX2 20726797 ChIP-Seq SW620 Human | SOX2 |
| 655 | ZC3H8 | Enrichr CHEA_2022: SOX2 20726797 ChIP-Seq SW620 Human | SOX2 |
| 656 | UBR1 | Enrichr CHEA_2022: SOX2 20726797 ChIP-Seq SW620 Human | SOX2 |
| 657 | CYP3A5 | Enrichr CHEA_2022: SOX2 20726797 ChIP-Seq SW620 Human | SOX2 |
| 658 | CYP3A7 | Enrichr CHEA_2022: SOX2 20726797 ChIP-Seq SW620 Human | SOX2 |
| 659 | LBH | Enrichr CHEA_2022: SOX2 20726797 ChIP-Seq SW620 Human | SOX2 |
| 660 | PSTPIP2 | Enrichr CHEA_2022: SOX2 20726797 ChIP-Seq SW620 Human | SOX2 |
| 661 | P2RY2 | Enrichr CHEA_2022: SOX2 20726797 ChIP-Seq SW620 Human | SOX2 |
| 662 | DVL2 | Enrichr CHEA_2022: SOX2 20726797 ChIP-Seq SW620 Human | SOX2 |
| 663 | PHACTR3 | Enrichr CHEA_2022: SOX2 20726797 ChIP-Seq SW620 Human | SOX2 |
| 664 | PHACTR2 | Enrichr CHEA_2022: SOX2 20726797 ChIP-Seq SW620 Human | SOX2 |
| 665 | DNAJB9 | Enrichr CHEA_2022: SOX2 20726797 ChIP-Seq SW620 Human | SOX2 |
| 666 | SERAC1 | Enrichr CHEA_2022: SOX2 20726797 ChIP-Seq SW620 Human | SOX2 |
| 667 | EMILIN2 | Enrichr CHEA_2022: SOX2 20726797 ChIP-Seq SW620 Human | SOX2 |
| 668 | CCDC34 | Enrichr CHEA_2022: SOX2 20726797 ChIP-Seq SW620 Human | SOX2 |
| 669 | TPPP | Enrichr CHEA_2022: SOX2 20726797 ChIP-Seq SW620 Human | SOX2 |

| | | | |
|---|---|---|---|
| 670 | CEP170 | Enrichr CHEA_2022: SOX2 20726797 ChIP-Seq SW620 Human | SOX2 |
| 671 | ADSL | Enrichr CHEA_2022: SOX2 20726797 ChIP-Seq SW620 Human | SOX2 |
| 672 | PLEKHG6 | Enrichr CHEA_2022: SOX2 20726797 ChIP-Seq SW620 Human | SOX2 |
| 673 | KRR1 | Enrichr CHEA_2022: SOX2 20726797 ChIP-Seq SW620 Human | SOX2 |
| 674 | COBL | Enrichr CHEA_2022: SOX2 20726797 ChIP-Seq SW620 Human | SOX2 |
| 675 | FBXL16 | Enrichr CHEA_2022: SOX2 20726797 ChIP-Seq SW620 Human | SOX2 |
| 676 | LEMD1 | Enrichr CHEA_2022: SOX2 20726797 ChIP-Seq SW620 Human | SOX2 |
| 677 | LCK | Enrichr CHEA_2022: SOX2 20726797 ChIP-Seq SW620 Human | SOX2 |
| 678 | P2RX5 | Enrichr CHEA_2022: SOX2 20726797 ChIP-Seq SW620 Human | SOX2 |
| 679 | RAB15 | Enrichr CHEA_2022: SOX2 20726797 ChIP-Seq SW620 Human | SOX2 |
| 680 | APOC1 | Enrichr CHEA_2022: SOX2 20726797 ChIP-Seq SW620 Human | SOX2 |
| 681 | KIAA0586 | Enrichr CHEA_2022: SOX2 20726797 ChIP-Seq SW620 Human | SOX2 |
| 682 | DNAJA4 | Enrichr CHEA_2022: SOX2 20726797 ChIP-Seq SW620 Human | SOX2 |
| 683 | POLR1D | Enrichr CHEA_2022: SOX2 20726797 ChIP-Seq SW620 Human | SOX2 |
| 684 | APOC2 | Enrichr CHEA_2022: SOX2 20726797 ChIP-Seq SW620 Human | SOX2 |
| 685 | POLR1E | Enrichr CHEA_2022: SOX2 20726797 ChIP-Seq SW620 Human | SOX2 |
| 686 | CCDC66 | Enrichr CHEA_2022: SOX2 20726797 ChIP-Seq SW620 Human | SOX2 |
| 687 | PLEKHG1 | Enrichr CHEA_2022: SOX2 20726797 ChIP-Seq SW620 Human | SOX2 |
| 688 | HDAC4 | Enrichr CHEA_2022: SOX2 20726797 ChIP-Seq SW620 Human | SOX2 |
| 689 | CEBPD | Enrichr CHEA_2022: SOX2 20726797 ChIP-Seq SW620 Human | SOX2 |
| 690 | MCM8 | Enrichr CHEA_2022: SOX2 20726797 ChIP-Seq SW620 Human | SOX2 |
| 691 | WDR4 | Enrichr CHEA_2022: SOX2 20726797 ChIP-Seq SW620 Human | SOX2 |
| 692 | TTL | Enrichr CHEA_2022: SOX2 20726797 ChIP-Seq SW620 Human | SOX2 |
| 693 | THSD1 | Enrichr CHEA_2022: SOX2 20726797 ChIP-Seq SW620 Human | SOX2 |
| 694 | NOB1 | Enrichr CHEA_2022: SOX2 20726797 ChIP-Seq SW620 Human | SOX2 |
| 695 | TTK | Enrichr CHEA_2022: SOX2 20726797 ChIP-Seq SW620 Human | SOX2 |
| 696 | HDAC9 | Enrichr CHEA_2022: SOX2 20726797 ChIP-Seq SW620 Human | SOX2 |
| 697 | SYNE2 | Enrichr CHEA_2022: SOX2 20726797 ChIP-Seq SW620 Human | SOX2 |
| 698 | THSD4 | Enrichr CHEA_2022: SOX2 20726797 ChIP-Seq SW620 Human | SOX2 |
| 699 | SCRN2 | Enrichr CHEA_2022: SOX2 20726797 ChIP-Seq SW620 Human | SOX2 |
| 700 | SCRN3 | Enrichr CHEA_2022: SOX2 20726797 ChIP-Seq SW620 Human | SOX2 |
| 701 | UGT1A6 | Enrichr CHEA_2022: SOX2 20726797 ChIP-Seq SW620 Human | SOX2 |
| 702 | CCDC51 | Enrichr CHEA_2022: SOX2 20726797 ChIP-Seq SW620 Human | SOX2 |
| 703 | UGT1A8 | Enrichr CHEA_2022: SOX2 20726797 ChIP-Seq SW620 Human | SOX2 |
| 704 | IFRD1 | Enrichr CHEA_2022: SOX2 20726797 ChIP-Seq SW620 Human | SOX2 |
| 705 | IL1R2 | Enrichr CHEA_2022: SOX2 20726797 ChIP-Seq SW620 Human | SOX2 |
| 706 | KRT7 | Enrichr CHEA_2022: SOX2 20726797 ChIP-Seq SW620 Human | SOX2 |
| 707 | CRNKL1 | Enrichr CHEA_2022: SOX2 20726797 ChIP-Seq SW620 Human | SOX2 |
| 708 | IL23A | Enrichr CHEA_2022: SOX2 20726797 ChIP-Seq SW620 Human | SOX2 |
| 709 | KATNAL1 | Enrichr CHEA_2022: SOX2 20726797 ChIP-Seq SW620 Human | SOX2 |
| 710 | NEIL2 | Enrichr CHEA_2022: SOX2 20726797 ChIP-Seq SW620 Human | SOX2 |
| 711 | OBSL1 | Enrichr CHEA_2022: SOX2 20726797 ChIP-Seq SW620 Human | SOX2 |
| 712 | DCP1B | Enrichr CHEA_2022: SOX2 20726797 ChIP-Seq SW620 Human | SOX2 |
| 713 | IFITM1 | Enrichr CHEA_2022: SOX2 20726797 ChIP-Seq SW620 Human | SOX2 |
| 714 | CBLC | Enrichr CHEA_2022: SOX2 20726797 ChIP-Seq SW620 Human | SOX2 |
| 715 | CALML4 | Enrichr CHEA_2022: SOX2 20726797 ChIP-Seq SW620 Human | SOX2 |
| 716 | AQP5 | Enrichr CHEA_2022: SOX2 20726797 ChIP-Seq SW620 Human | SOX2 |
| 717 | AQP3 | Enrichr CHEA_2022: SOX2 20726797 ChIP-Seq SW620 Human | SOX2 |
| 718 | APCDD1 | Enrichr CHEA_2022: SOX2 20726797 ChIP-Seq SW620 Human | SOX2 |
| 719 | MTHFD1L | Enrichr CHEA_2022: SOX2 20726797 ChIP-Seq SW620 Human | SOX2 |
| 720 | STXBP6 | Enrichr CHEA_2022: SOX2 20726797 ChIP-Seq SW620 Human | SOX2 |
| 721 | RFFL | Enrichr CHEA_2022: SOX2 20726797 ChIP-Seq SW620 Human | SOX2 |
| 722 | HLA-E | Enrichr CHEA_2022: SOX2 20726797 ChIP-Seq SW620 Human | SOX2 |
| 723 | VASP | Enrichr CHEA_2022: SOX2 20726797 ChIP-Seq SW620 Human | SOX2 |
| 724 | VNN1 | Enrichr CHEA_2022: SOX2 20726797 ChIP-Seq SW620 Human | SOX2 |
| 725 | SMTN | Enrichr CHEA_2022: SOX2 20726797 ChIP-Seq SW620 Human | SOX2 |
| 726 | SPRR1A | Enrichr CHEA_2022: SOX2 20726797 ChIP-Seq SW620 Human | SOX2 |
| 727 | RPS23 | Enrichr CHEA_2022: SOX2 20726797 ChIP-Seq SW620 Human | SOX2 |
| 728 | SIM2 | Enrichr CHEA_2022: SOX2 20726797 ChIP-Seq SW620 Human | SOX2 |
| 729 | SPRR1B | Enrichr CHEA_2022: SOX2 20726797 ChIP-Seq SW620 Human | SOX2 |
| 730 | CFTR | Enrichr CHEA_2022: SOX2 20726797 ChIP-Seq SW620 Human | SOX2 |

| | | | |
|---|---|---|---|
| 731 | IL1RN | Enrichr CHEA_2022: SOX2 20726797 ChIP-Seq SW620 Human | SOX2 |
| 732 | MAGED2 | Enrichr CHEA_2022: SOX2 20726797 ChIP-Seq SW620 Human | SOX2 |
| 733 | DCUN1D4 | Enrichr CHEA_2022: SOX2 20726797 ChIP-Seq SW620 Human | SOX2 |
| 734 | FAM89A | Enrichr CHEA_2022: SOX2 20726797 ChIP-Seq SW620 Human | SOX2 |
| 735 | PPL | Enrichr CHEA_2022: SOX2 20726797 ChIP-Seq SW620 Human | SOX2 |
| 736 | IFIT2 | Enrichr CHEA_2022: SOX2 20726797 ChIP-Seq SW620 Human | SOX2 |
| 737 | IFIT1 | Enrichr CHEA_2022: SOX2 20726797 ChIP-Seq SW620 Human | SOX2 |
| 738 | TRPM2 | Enrichr CHEA_2022: SOX2 20726797 ChIP-Seq SW620 Human | SOX2 |
| 739 | PDK4 | Enrichr CHEA_2022: SOX2 20726797 ChIP-Seq SW620 Human | SOX2 |
| 740 | PDK1 | Enrichr CHEA_2022: SOX2 20726797 ChIP-Seq SW620 Human | SOX2 |
| 741 | RALGPS2 | Enrichr CHEA_2022: SOX2 20726797 ChIP-Seq SW620 Human | SOX2 |
| 742 | TRPM4 | Enrichr CHEA_2022: SOX2 20726797 ChIP-Seq SW620 Human | SOX2 |
| 743 | PDK3 | Enrichr CHEA_2022: SOX2 20726797 ChIP-Seq SW620 Human | SOX2 |
| 744 | PSENEN | Enrichr CHEA_2022: SOX2 20726797 ChIP-Seq SW620 Human | SOX2 |
| 745 | EDN1 | Enrichr CHEA_2022: SOX2 20726797 ChIP-Seq SW620 Human | SOX2 |
| 746 | HEG1 | Enrichr CHEA_2022: SOX2 20726797 ChIP-Seq SW620 Human | SOX2 |
| 747 | UBL4A | Enrichr CHEA_2022: SOX2 20726797 ChIP-Seq SW620 Human | SOX2 |
| 748 | MLF1 | Enrichr CHEA_2022: SOX2 20726797 ChIP-Seq SW620 Human | SOX2 |
| 749 | MTSS1 | Enrichr CHEA_2022: SOX2 20726797 ChIP-Seq SW620 Human | SOX2 |
| 750 | CLDN11 | Enrichr CHEA_2022: SOX2 20726797 ChIP-Seq SW620 Human | SOX2 |
| 751 | KBTBD7 | Enrichr CHEA_2022: SOX2 20726797 ChIP-Seq SW620 Human | SOX2 |
| 752 | P4HA2 | Enrichr CHEA_2022: SOX2 20726797 ChIP-Seq SW620 Human | SOX2 |
| 753 | VWA1 | Enrichr CHEA_2022: SOX2 20726797 ChIP-Seq SW620 Human | SOX2 |
| 754 | TLR4 | Enrichr CHEA_2022: SOX2 20726797 ChIP-Seq SW620 Human | SOX2 |
| 755 | RAB7B | Enrichr CHEA_2022: SOX2 20726797 ChIP-Seq SW620 Human | SOX2 |
| 756 | SLC44A4 | Enrichr CHEA_2022: SOX2 20726797 ChIP-Seq SW620 Human | SOX2 |
| 757 | CITED1 | Enrichr CHEA_2022: SOX2 20726797 ChIP-Seq SW620 Human | SOX2 |
| 758 | RTN4R | Enrichr CHEA_2022: SOX2 20726797 ChIP-Seq SW620 Human | SOX2 |
| 759 | SLC44A2 | Enrichr CHEA_2022: SOX2 20726797 ChIP-Seq SW620 Human | SOX2 |
| 760 | FAM43A | Enrichr CHEA_2022: SOX2 20726797 ChIP-Seq SW620 Human | SOX2 |
| 761 | CAB39L | Enrichr CHEA_2022: SOX2 20726797 ChIP-Seq SW620 Human | SOX2 |
| 762 | SLC44A1 | Enrichr CHEA_2022: SOX2 20726797 ChIP-Seq SW620 Human | SOX2 |
| 763 | CHRAC1 | Enrichr CHEA_2022: SOX2 20726797 ChIP-Seq SW620 Human | SOX2 |
| 764 | SLC22A15 | Enrichr CHEA_2022: SOX2 20726797 ChIP-Seq SW620 Human | SOX2 |
| 765 | UGCG | Enrichr CHEA_2022: SOX2 20726797 ChIP-Seq SW620 Human | SOX2 |
| 766 | SLC22A18 | Enrichr CHEA_2022: SOX2 20726797 ChIP-Seq SW620 Human | SOX2 |
| 767 | C1QBP | Enrichr CHEA_2022: SOX2 20726797 ChIP-Seq SW620 Human | SOX2 |
| 768 | TMED3 | Enrichr CHEA_2022: SOX2 20726797 ChIP-Seq SW620 Human | SOX2 |
| 769 | TMED8 | Enrichr CHEA_2022: SOX2 20726797 ChIP-Seq SW620 Human | SOX2 |
| 770 | TMED5 | Enrichr CHEA_2022: SOX2 20726797 ChIP-Seq SW620 Human | SOX2 |
| 771 | MYEF2 | Enrichr CHEA_2022: SOX2 20726797 ChIP-Seq SW620 Human | SOX2 |
| 772 | PARP4 | Enrichr CHEA_2022: SOX2 20726797 ChIP-Seq SW620 Human | SOX2 |
| 773 | CBR1 | Enrichr CHEA_2022: SOX2 20726797 ChIP-Seq SW620 Human | SOX2 |
| 774 | PARP2 | Enrichr CHEA_2022: SOX2 20726797 ChIP-Seq SW620 Human | SOX2 |
| 775 | METTL3 | Enrichr CHEA_2022: SOX2 20726797 ChIP-Seq SW620 Human | SOX2 |
| 776 | PROX1 | Enrichr CHEA_2022: SOX2 20726797 ChIP-Seq SW620 Human | SOX2 |
| 777 | SORBS1 | Enrichr CHEA_2022: SOX2 20726797 ChIP-Seq SW620 Human | SOX2 |
| 778 | QPCT | Enrichr CHEA_2022: SOX2 20726797 ChIP-Seq SW620 Human | SOX2 |
| 779 | SORBS2 | Enrichr CHEA_2022: SOX2 20726797 ChIP-Seq SW620 Human | SOX2 |
| 780 | RAB11A | Enrichr CHEA_2022: SOX2 20726797 ChIP-Seq SW620 Human | SOX2 |
| 781 | GFRA3 | Enrichr CHEA_2022: SOX2 20726797 ChIP-Seq SW620 Human | SOX2 |
| 782 | LIMA1 | Enrichr CHEA_2022: SOX2 20726797 ChIP-Seq SW620 Human | SOX2 |
| 783 | PARP9 | Enrichr CHEA_2022: SOX2 20726797 ChIP-Seq SW620 Human | SOX2 |
| 784 | PDP2 | Enrichr CHEA_2022: SOX2 20726797 ChIP-Seq SW620 Human | SOX2 |
| 785 | ALDH5A1 | Enrichr CHEA_2022: SOX2 20726797 ChIP-Seq SW620 Human | SOX2 |
| 786 | GDPD1 | Enrichr CHEA_2022: SOX2 20726797 ChIP-Seq SW620 Human | SOX2 |
| 787 | RAB9A | Enrichr CHEA_2022: SOX2 20726797 ChIP-Seq SW620 Human | SOX2 |
| 788 | CSTB | Enrichr CHEA_2022: SOX2 20726797 ChIP-Seq SW620 Human | SOX2 |
| 789 | GBE1 | Enrichr CHEA_2022: SOX2 20726797 ChIP-Seq SW620 Human | SOX2 |
| 790 | SIPA1L3 | Enrichr CHEA_2022: SOX2 20726797 ChIP-Seq SW620 Human | SOX2 |
| 791 | CST7 | Enrichr CHEA_2022: SOX2 20726797 ChIP-Seq SW620 Human | SOX2 |

| | | | |
|---|---|---|---|
| 792 | ROBO1 | Enrichr CHEA_2022: SOX2 20726797 ChIP-Seq SW620 Human | SOX2 |
| 793 | CST6 | Enrichr CHEA_2022: SOX2 20726797 ChIP-Seq SW620 Human | SOX2 |
| 794 | NKAP | Enrichr CHEA_2022: SOX2 20726797 ChIP-Seq SW620 Human | SOX2 |
| 795 | RPAIN | Enrichr CHEA_2022: SOX2 20726797 ChIP-Seq SW620 Human | SOX2 |
| 796 | SIPA1L1 | Enrichr CHEA_2022: SOX2 20726797 ChIP-Seq SW620 Human | SOX2 |
| 797 | SIPA1L2 | Enrichr CHEA_2022: SOX2 20726797 ChIP-Seq SW620 Human | SOX2 |
| 798 | SOSTDC1 | Enrichr CHEA_2022: SOX2 20726797 ChIP-Seq SW620 Human | SOX2 |
| 799 | LMO2 | Enrichr CHEA_2022: SOX2 20726797 ChIP-Seq SW620 Human | SOX2 |
| 800 | GABRP | Enrichr CHEA_2022: SOX2 20726797 ChIP-Seq SW620 Human | SOX2 |
| 801 | LMO4 | Enrichr CHEA_2022: SOX2 20726797 ChIP-Seq SW620 Human | SOX2 |
| 802 | TRPV2 | Enrichr CHEA_2022: SOX2 20726797 ChIP-Seq SW620 Human | SOX2 |
| 803 | GRHL2 | Enrichr CHEA_2022: SOX2 20726797 ChIP-Seq SW620 Human | SOX2 |
| 804 | ATP1B1 | Enrichr CHEA_2022: SOX2 20726797 ChIP-Seq SW620 Human | SOX2 |
| 805 | TATDN1 | Enrichr CHEA_2022: SOX2 20726797 ChIP-Seq SW620 Human | SOX2 |
| 806 | AIG1 | Enrichr CHEA_2022: SOX2 20726797 ChIP-Seq SW620 Human | SOX2 |
| 807 | GRHL1 | Enrichr CHEA_2022: SOX2 20726797 ChIP-Seq SW620 Human | SOX2 |
| 808 | BAMBI | Enrichr CHEA_2022: SOX2 20726797 ChIP-Seq SW620 Human | SOX2 |
| 809 | FANCD2 | Enrichr CHEA_2022: SOX2 20726797 ChIP-Seq SW620 Human | SOX2 |
| 810 | TRPV6 | Enrichr CHEA_2022: SOX2 20726797 ChIP-Seq SW620 Human | SOX2 |
| 811 | CRYL1 | Enrichr CHEA_2022: SOX2 20726797 ChIP-Seq SW620 Human | SOX2 |
| 812 | CHMP4B | Enrichr CHEA_2022: SOX2 20726797 ChIP-Seq SW620 Human | SOX2 |
| 813 | MYADM | Enrichr CHEA_2022: SOX2 20726797 ChIP-Seq SW620 Human | SOX2 |
| 814 | BRWD1 | Enrichr CHEA_2022: SOX2 20726797 ChIP-Seq SW620 Human | SOX2 |
| 815 | LRP10 | Enrichr CHEA_2022: SOX2 20726797 ChIP-Seq SW620 Human | SOX2 |
| 816 | MPHOSPH6 | Enrichr CHEA_2022: SOX2 20726797 ChIP-Seq SW620 Human | SOX2 |
| 817 | XPR1 | Enrichr CHEA_2022: SOX2 20726797 ChIP-Seq SW620 Human | SOX2 |
| 818 | PVR | Enrichr CHEA_2022: SOX2 20726797 ChIP-Seq SW620 Human | SOX2 |
| 819 | FBLN1 | Enrichr CHEA_2022: SOX2 20726797 ChIP-Seq SW620 Human | SOX2 |
| 820 | NKD1 | Enrichr CHEA_2022: SOX2 20726797 ChIP-Seq SW620 Human | SOX2 |
| 821 | COMT | Enrichr CHEA_2022: SOX2 20726797 ChIP-Seq SW620 Human | SOX2 |
| 822 | CNDP2 | Enrichr CHEA_2022: SOX2 20726797 ChIP-Seq SW620 Human | SOX2 |
| 823 | HOXA13 | Enrichr CHEA_2022: SOX2 20726797 ChIP-Seq SW620 Human | SOX2 |
| 824 | LMNA | Enrichr CHEA_2022: SOX2 20726797 ChIP-Seq SW620 Human | SOX2 |
| 825 | DOCK11 | Enrichr CHEA_2022: SOX2 20726797 ChIP-Seq SW620 Human | SOX2 |
| 826 | NT5E | Enrichr CHEA_2022: SOX2 20726797 ChIP-Seq SW620 Human | SOX2 |
| 827 | PURB | Enrichr CHEA_2022: SOX2 20726797 ChIP-Seq SW620 Human | SOX2 |
| 828 | KISS1R | Enrichr CHEA_2022: SOX2 20726797 ChIP-Seq SW620 Human | SOX2 |
| 829 | GAS2L1 | Enrichr CHEA_2022: SOX2 20726797 ChIP-Seq SW620 Human | SOX2 |
| 830 | EIF2S3 | Enrichr CHEA_2022: SOX2 20726797 ChIP-Seq SW620 Human | SOX2 |
| 831 | EIF2S1 | Enrichr CHEA_2022: SOX2 20726797 ChIP-Seq SW620 Human | SOX2 |
| 832 | DIAPH1 | Enrichr CHEA_2022: SOX2 20726797 ChIP-Seq SW620 Human | SOX2 |
| 833 | EFNA4 | Enrichr CHEA_2022: SOX2 20726797 ChIP-Seq SW620 Human | SOX2 |
| 834 | PRPF8 | Enrichr CHEA_2022: SOX2 20726797 ChIP-Seq SW620 Human | SOX2 |
| 835 | COL6A1 | Enrichr CHEA_2022: SOX2 20726797 ChIP-Seq SW620 Human | SOX2 |
| 836 | DDAH1 | Enrichr CHEA_2022: SOX2 20726797 ChIP-Seq SW620 Human | SOX2 |
| 837 | CDH13 | Enrichr CHEA_2022: SOX2 20726797 ChIP-Seq SW620 Human | SOX2 |
| 838 | FGFR1 | Enrichr CHEA_2022: SOX2 20726797 ChIP-Seq SW620 Human | SOX2 |
| 839 | CHMP2A | Enrichr CHEA_2022: SOX2 20726797 ChIP-Seq SW620 Human | SOX2 |
| 840 | FGFR3 | Enrichr CHEA_2022: SOX2 20726797 ChIP-Seq SW620 Human | SOX2 |
| 841 | ZCCHC2 | Enrichr CHEA_2022: SOX2 20726797 ChIP-Seq SW620 Human | SOX2 |
| 842 | FGFR2 | Enrichr CHEA_2022: SOX2 20726797 ChIP-Seq SW620 Human | SOX2 |
| 843 | PXK | Enrichr CHEA_2022: SOX2 20726797 ChIP-Seq SW620 Human | SOX2 |
| 844 | LZIC | Enrichr CHEA_2022: SOX2 20726797 ChIP-Seq SW620 Human | SOX2 |
| 845 | RASEF | Enrichr CHEA_2022: SOX2 20726797 ChIP-Seq SW620 Human | SOX2 |
| 846 | SPPL2A | Enrichr CHEA_2022: SOX2 20726797 ChIP-Seq SW620 Human | SOX2 |
| 847 | RBMX2 | Enrichr CHEA_2022: SOX2 20726797 ChIP-Seq SW620 Human | SOX2 |
| 848 | EME2 | Enrichr CHEA_2022: SOX2 20726797 ChIP-Seq SW620 Human | SOX2 |
| 849 | PMAIP1 | Enrichr CHEA_2022: SOX2 20726797 ChIP-Seq SW620 Human | SOX2 |
| 850 | MLKL | Enrichr CHEA_2022: SOX2 20726797 ChIP-Seq SW620 Human | SOX2 |
| 851 | SACS | Enrichr CHEA_2022: SOX2 20726797 ChIP-Seq SW620 Human | SOX2 |
| 852 | CYP2J2 | Enrichr CHEA_2022: SOX2 20726797 ChIP-Seq SW620 Human | SOX2 |

| | | | |
|---|---|---|---|
| 853 | CBX6 | Enrichr CHEA_2022: SOX2 20726797 ChIP-Seq SW620 Human | SOX2 |
| 854 | CBX4 | Enrichr CHEA_2022: SOX2 20726797 ChIP-Seq SW620 Human | SOX2 |
| 855 | CBX2 | Enrichr CHEA_2022: SOX2 20726797 ChIP-Seq SW620 Human | SOX2 |
| 856 | PARVB | Enrichr CHEA_2022: SOX2 20726797 ChIP-Seq SW620 Human | SOX2 |
| 857 | IQCE | Enrichr CHEA_2022: SOX2 20726797 ChIP-Seq SW620 Human | SOX2 |
| 858 | OAS1 | Enrichr CHEA_2022: SOX2 20726797 ChIP-Seq SW620 Human | SOX2 |
| 859 | DNM1 | Enrichr CHEA_2022: SOX2 20726797 ChIP-Seq SW620 Human | SOX2 |
| 860 | RBKS | Enrichr CHEA_2022: SOX2 20726797 ChIP-Seq SW620 Human | SOX2 |
| 861 | RASA1 | Enrichr CHEA_2022: SOX2 20726797 ChIP-Seq SW620 Human | SOX2 |
| 862 | RABEP1 | Enrichr CHEA_2022: SOX2 20726797 ChIP-Seq SW620 Human | SOX2 |
| 863 | RASA4 | Enrichr CHEA_2022: SOX2 20726797 ChIP-Seq SW620 Human | SOX2 |
| 864 | OAS3 | Enrichr CHEA_2022: SOX2 20726797 ChIP-Seq SW620 Human | SOX2 |
| 865 | AKT1S1 | Enrichr CHEA_2022: SOX2 20726797 ChIP-Seq SW620 Human | SOX2 |
| 866 | BLMH | Enrichr CHEA_2022: SOX2 20726797 ChIP-Seq SW620 Human | SOX2 |
| 867 | DNASE2 | Enrichr CHEA_2022: SOX2 20726797 ChIP-Seq SW620 Human | SOX2 |
| 868 | TERF2IP | Enrichr CHEA_2022: SOX2 20726797 ChIP-Seq SW620 Human | SOX2 |
| 869 | TRPS1 | Enrichr CHEA_2022: SOX2 20726797 ChIP-Seq SW620 Human | SOX2 |
| 870 | SLC40A1 | Enrichr CHEA_2022: SOX2 20726797 ChIP-Seq SW620 Human | SOX2 |
| 871 | ERICH1 | Enrichr CHEA_2022: SOX2 20726797 ChIP-Seq SW620 Human | SOX2 |
| 872 | FAM83H | Enrichr CHEA_2022: SOX2 20726797 ChIP-Seq SW620 Human | SOX2 |
| 873 | WNT11 | Enrichr CHEA_2022: SOX2 20726797 ChIP-Seq SW620 Human | SOX2 |
| 874 | CASP6 | Enrichr CHEA_2022: SOX2 20726797 ChIP-Seq SW620 Human | SOX2 |
| 875 | CASP7 | Enrichr CHEA_2022: SOX2 20726797 ChIP-Seq SW620 Human | SOX2 |
| 876 | CASP4 | Enrichr CHEA_2022: SOX2 20726797 ChIP-Seq SW620 Human | SOX2 |
| 877 | SNAPC5 | Enrichr CHEA_2022: SOX2 20726797 ChIP-Seq SW620 Human | SOX2 |
| 878 | FAM83A | Enrichr CHEA_2022: SOX2 20726797 ChIP-Seq SW620 Human | SOX2 |
| 879 | MYEOV | Enrichr CHEA_2022: SOX2 20726797 ChIP-Seq SW620 Human | SOX2 |
| 880 | SNAPC1 | Enrichr CHEA_2022: SOX2 20726797 ChIP-Seq SW620 Human | SOX2 |
| 881 | LONRF1 | Enrichr CHEA_2022: SOX2 20726797 ChIP-Seq SW620 Human | SOX2 |
| 882 | FAM83B | Enrichr CHEA_2022: SOX2 20726797 ChIP-Seq SW620 Human | SOX2 |
| 883 | RHBDF1 | Enrichr CHEA_2022: SOX2 20726797 ChIP-Seq SW620 Human | SOX2 |
| 884 | IL15RA | Enrichr CHEA_2022: SOX2 20726797 ChIP-Seq SW620 Human | SOX2 |
| 885 | TMEM30B | Enrichr CHEA_2022: SOX2 20726797 ChIP-Seq SW620 Human | SOX2 |
| 886 | PLAUR | Enrichr CHEA_2022: SOX2 20726797 ChIP-Seq SW620 Human | SOX2 |
| 887 | LSR | Enrichr CHEA_2022: SOX2 20726797 ChIP-Seq SW620 Human | SOX2 |
| 888 | ANXA10 | Enrichr CHEA_2022: SOX2 20726797 ChIP-Seq SW620 Human | SOX2 |
| 889 | TIAM1 | Enrichr CHEA_2022: SOX2 20726797 ChIP-Seq SW620 Human | SOX2 |
| 890 | MLPH | Enrichr CHEA_2022: SOX2 20726797 ChIP-Seq SW620 Human | SOX2 |
| 891 | TRAPPC6A | Enrichr CHEA_2022: SOX2 20726797 ChIP-Seq SW620 Human | SOX2 |
| 892 | ALDH1A1 | Enrichr CHEA_2022: SOX2 20726797 ChIP-Seq SW620 Human | SOX2 |
| 893 | PPP2R2C | Enrichr CHEA_2022: SOX2 20726797 ChIP-Seq SW620 Human | SOX2 |
| 894 | ALDH1A3 | Enrichr CHEA_2022: SOX2 20726797 ChIP-Seq SW620 Human | SOX2 |
| 895 | DDIT4 | Enrichr CHEA_2022: SOX2 20726797 ChIP-Seq SW620 Human | SOX2 |
| 896 | DDIT3 | Enrichr CHEA_2022: SOX2 20726797 ChIP-Seq SW620 Human | SOX2 |
| 897 | CLOCK | Enrichr CHEA_2022: SOX2 20726797 ChIP-Seq SW620 Human | SOX2 |
| 898 | SIX1 | Enrichr CHEA_2022: SOX2 20726797 ChIP-Seq SW620 Human | SOX2 |
| 899 | THBS1 | Enrichr CHEA_2022: SOX2 20726797 ChIP-Seq SW620 Human | SOX2 |
| 900 | OASL | Enrichr CHEA_2022: SOX2 20726797 ChIP-Seq SW620 Human | SOX2 |
| 901 | GNA14 | Enrichr CHEA_2022: SOX2 20726797 ChIP-Seq SW620 Human | SOX2 |
| 902 | GNA15 | Enrichr CHEA_2022: SOX2 20726797 ChIP-Seq SW620 Human | SOX2 |
| 903 | HEPH | Enrichr CHEA_2022: SOX2 20726797 ChIP-Seq SW620 Human | SOX2 |
| 904 | ARHGDIB | Enrichr CHEA_2022: SOX2 20726797 ChIP-Seq SW620 Human | SOX2 |
| 905 | LEPROTL1 | Enrichr CHEA_2022: SOX2 20726797 ChIP-Seq SW620 Human | SOX2 |
| 906 | RPS6KA2 | Enrichr CHEA_2022: SOX2 20726797 ChIP-Seq SW620 Human | SOX2 |
| 907 | B3GALNT1 | Enrichr CHEA_2022: SOX2 20726797 ChIP-Seq SW620 Human | SOX2 |
| 908 | GDAP1 | Enrichr CHEA_2022: SOX2 20726797 ChIP-Seq SW620 Human | SOX2 |
| 909 | WDHD1 | Enrichr CHEA_2022: SOX2 20726797 ChIP-Seq SW620 Human | SOX2 |
| 910 | PSMB9 | Enrichr CHEA_2022: SOX2 20726797 ChIP-Seq SW620 Human | SOX2 |
| 911 | EML4 | Enrichr CHEA_2022: SOX2 20726797 ChIP-Seq SW620 Human | SOX2 |
| 912 | MYL5 | Enrichr CHEA_2022: SOX2 20726797 ChIP-Seq SW620 Human | SOX2 |
| 913 | RBP4 | Enrichr CHEA_2022: SOX2 20726797 ChIP-Seq SW620 Human | SOX2 |

| | | | |
|---|---|---|---|
| 914 | RBP1 | Enrichr CHEA_2022: SOX2 20726797 ChIP-Seq SW620 Human | SOX2 |
| 915 | LETMD1 | Enrichr CHEA_2022: SOX2 20726797 ChIP-Seq SW620 Human | SOX2 |
| 916 | LEF1 | Enrichr CHEA_2022: SOX2 20726797 ChIP-Seq SW620 Human | SOX2 |
| 917 | CGGBP1 | Enrichr CHEA_2022: SOX2 20726797 ChIP-Seq SW620 Human | SOX2 |
| 918 | PDXK | Enrichr CHEA_2022: SOX2 20726797 ChIP-Seq SW620 Human | SOX2 |
| 919 | ZNF274 | Enrichr CHEA_2022: SOX2 20726797 ChIP-Seq SW620 Human | SOX2 |
| 920 | ZNF273 | Enrichr CHEA_2022: SOX2 20726797 ChIP-Seq SW620 Human | SOX2 |
| 921 | IER5L | Enrichr CHEA_2022: SOX2 20726797 ChIP-Seq SW620 Human | SOX2 |
| 922 | FZD10 | Enrichr CHEA_2022: SOX2 20726797 ChIP-Seq SW620 Human | SOX2 |
| 923 | ARAF | Enrichr CHEA_2022: SOX2 20726797 ChIP-Seq SW620 Human | SOX2 |
| 924 | CLDN3 | Enrichr CHEA_2022: SOX2 20726797 ChIP-Seq SW620 Human | SOX2 |
| 925 | CLDN1 | Enrichr CHEA_2022: SOX2 20726797 ChIP-Seq SW620 Human | SOX2 |
| 926 | KRTCAP3 | Enrichr CHEA_2022: SOX2 20726797 ChIP-Seq SW620 Human | SOX2 |
| 927 | PREX1 | Enrichr CHEA_2022: SOX2 20726797 ChIP-Seq SW620 Human | SOX2 |
| 928 | DNTTIP1 | Enrichr CHEA_2022: SOX2 20726797 ChIP-Seq SW620 Human | SOX2 |
| 929 | SNCG | Enrichr CHEA_2022: SOX2 20726797 ChIP-Seq SW620 Human | SOX2 |
| 930 | STAT6 | Enrichr CHEA_2022: SOX2 20726797 ChIP-Seq SW620 Human | SOX2 |
| 931 | RABAC1 | Enrichr CHEA_2022: SOX2 20726797 ChIP-Seq SW620 Human | SOX2 |
| 932 | GBP3 | Enrichr CHEA_2022: SOX2 20726797 ChIP-Seq SW620 Human | SOX2 |
| 933 | GBP2 | Enrichr CHEA_2022: SOX2 20726797 ChIP-Seq SW620 Human | SOX2 |
| 934 | HES6 | Enrichr CHEA_2022: SOX2 20726797 ChIP-Seq SW620 Human | SOX2 |
| 935 | NGFR | Enrichr CHEA_2022: SOX2 20726797 ChIP-Seq SW620 Human | SOX2 |
| 936 | MYH7B | Enrichr CHEA_2022: SOX2 20726797 ChIP-Seq SW620 Human | SOX2 |
| 937 | KLF12 | Enrichr CHEA_2022: SOX2 20726797 ChIP-Seq SW620 Human | SOX2 |
| 938 | ZNF264 | Enrichr CHEA_2022: SOX2 20726797 ChIP-Seq SW620 Human | SOX2 |
| 939 | GABARAPL1 | Enrichr CHEA_2022: SOX2 20726797 ChIP-Seq SW620 Human | SOX2 |
| 940 | FIBCD1 | Enrichr CHEA_2022: SOX2 20726797 ChIP-Seq SW620 Human | SOX2 |
| 941 | LATS2 | Enrichr CHEA_2022: SOX2 20726797 ChIP-Seq SW620 Human | SOX2 |
| 942 | MLLT11 | Enrichr CHEA_2022: SOX2 20726797 ChIP-Seq SW620 Human | SOX2 |
| 943 | IL22RA1 | Enrichr CHEA_2022: SOX2 20726797 ChIP-Seq SW620 Human | SOX2 |
| 944 | PRKAR1B | Enrichr CHEA_2022: SOX2 20726797 ChIP-Seq SW620 Human | SOX2 |
| 945 | VAMP1 | Enrichr CHEA_2022: SOX2 20726797 ChIP-Seq SW620 Human | SOX2 |
| 946 | PADI2 | Enrichr CHEA_2022: SOX2 20726797 ChIP-Seq SW620 Human | SOX2 |
| 947 | VAMP3 | Enrichr CHEA_2022: SOX2 20726797 ChIP-Seq SW620 Human | SOX2 |
| 948 | MCPH1 | Enrichr CHEA_2022: SOX2 20726797 ChIP-Seq SW620 Human | SOX2 |
| 949 | PADI1 | Enrichr CHEA_2022: SOX2 20726797 ChIP-Seq SW620 Human | SOX2 |
| 950 | LXN | Enrichr CHEA_2022: SOX2 20726797 ChIP-Seq SW620 Human | SOX2 |
| 951 | SCHIP1 | Enrichr CHEA_2022: SOX2 20726797 ChIP-Seq SW620 Human | SOX2 |
| 952 | UXS1 | Enrichr CHEA_2022: SOX2 20726797 ChIP-Seq SW620 Human | SOX2 |
| 953 | MRPL13 | Enrichr CHEA_2022: SOX2 20726797 ChIP-Seq SW620 Human | SOX2 |
| 954 | TMEM92 | Enrichr CHEA_2022: SOX2 20726797 ChIP-Seq SW620 Human | SOX2 |
| 955 | MYLK | Enrichr CHEA_2022: SOX2 20726797 ChIP-Seq SW620 Human | SOX2 |
| 956 | LIPG | Enrichr CHEA_2022: SOX2 20726797 ChIP-Seq SW620 Human | SOX2 |
| 957 | NETO2 | Enrichr CHEA_2022: SOX2 20726797 ChIP-Seq SW620 Human | SOX2 |
| 958 | GRPEL2 | Enrichr CHEA_2022: SOX2 20726797 ChIP-Seq SW620 Human | SOX2 |
| 959 | IL20RA | Enrichr CHEA_2022: SOX2 20726797 ChIP-Seq SW620 Human | SOX2 |
| 960 | KCNMB4 | Enrichr CHEA_2022: SOX2 20726797 ChIP-Seq SW620 Human | SOX2 |
| 961 | ZBTB7C | Enrichr CHEA_2022: SOX2 20726797 ChIP-Seq SW620 Human | SOX2 |
| 962 | AMZ2 | Enrichr CHEA_2022: SOX2 20726797 ChIP-Seq SW620 Human | SOX2 |
| 963 | RHBDL2 | Enrichr CHEA_2022: SOX2 20726797 ChIP-Seq SW620 Human | SOX2 |
| 964 | ANKIB1 | Enrichr CHEA_2022: SOX2 20726797 ChIP-Seq SW620 Human | SOX2 |
| 965 | HACE1 | Enrichr CHEA_2022: SOX2 20726797 ChIP-Seq SW620 Human | SOX2 |
| 966 | UPF2 | Enrichr CHEA_2022: SOX2 20726797 ChIP-Seq SW620 Human | SOX2 |
| 967 | LYN | Enrichr CHEA_2022: SOX2 20726797 ChIP-Seq SW620 Human | SOX2 |
| 968 | FGFBP1 | Enrichr CHEA_2022: SOX2 20726797 ChIP-Seq SW620 Human | SOX2 |
| 969 | RIPK3 | Enrichr CHEA_2022: SOX2 20726797 ChIP-Seq SW620 Human | SOX2 |
| 970 | NSUN4 | Enrichr CHEA_2022: SOX2 20726797 ChIP-Seq SW620 Human | SOX2 |
| 971 | EMP3 | Enrichr CHEA_2022: SOX2 20726797 ChIP-Seq SW620 Human | SOX2 |
| 972 | FAHD2A | Enrichr CHEA_2022: SOX2 20726797 ChIP-Seq SW620 Human | SOX2 |
| 973 | TMCO4 | Enrichr CHEA_2022: SOX2 20726797 ChIP-Seq SW620 Human | SOX2 |
| 974 | TMC5 | Enrichr CHEA_2022: SOX2 20726797 ChIP-Seq SW620 Human | SOX2 |

| | | | |
|---|---|---|---|
| 975 | LYZ | Enrichr CHEA_2022: SOX2 20726797 ChIP-Seq SW620 Human | SOX2 |
| 976 | TMC4 | Enrichr CHEA_2022: SOX2 20726797 ChIP-Seq SW620 Human | SOX2 |
| 977 | GPRC5A | Enrichr CHEA_2022: SOX2 20726797 ChIP-Seq SW620 Human | SOX2 |
| 978 | LETM2 | Enrichr CHEA_2022: SOX2 20726797 ChIP-Seq SW620 Human | SOX2 |
| 979 | PIGA | Enrichr CHEA_2022: SOX2 20726797 ChIP-Seq SW620 Human | SOX2 |
| 980 | PIGK | Enrichr CHEA_2022: SOX2 20726797 ChIP-Seq SW620 Human | SOX2 |
| 981 | ZNF239 | Enrichr CHEA_2022: SOX2 20726797 ChIP-Seq SW620 Human | SOX2 |
| 982 | ZNF232 | Enrichr CHEA_2022: SOX2 20726797 ChIP-Seq SW620 Human | SOX2 |
| 983 | ATF7IP2 | Enrichr CHEA_2022: SOX2 20726797 ChIP-Seq SW620 Human | SOX2 |
| 984 | ACAT1 | Enrichr CHEA_2022: SOX2 20726797 ChIP-Seq SW620 Human | SOX2 |
| 985 | IQGAP1 | Enrichr CHEA_2022: SOX2 20726797 ChIP-Seq SW620 Human | SOX2 |
| 986 | LITAF | Enrichr CHEA_2022: SOX2 20726797 ChIP-Seq SW620 Human | SOX2 |
| 987 | IQGAP2 | Enrichr CHEA_2022: SOX2 20726797 ChIP-Seq SW620 Human | SOX2 |
| 988 | MRPL38 | Enrichr CHEA_2022: SOX2 20726797 ChIP-Seq SW620 Human | SOX2 |
| 989 | LRRC1 | Enrichr CHEA_2022: SOX2 20726797 ChIP-Seq SW620 Human | SOX2 |
| 990 | AREG | Enrichr CHEA_2022: SOX2 20726797 ChIP-Seq SW620 Human | SOX2 |
| 991 | LRRC34 | Enrichr CHEA_2022: SOX2 20726797 ChIP-Seq SW620 Human | SOX2 |
| 992 | HSD11B2 | Enrichr CHEA_2022: SOX2 20726797 ChIP-Seq SW620 Human | SOX2 |
| 993 | UGT8 | Enrichr CHEA_2022: SOX2 20726797 ChIP-Seq SW620 Human | SOX2 |
| 994 | MYO6 | Enrichr CHEA_2022: SOX2 20726797 ChIP-Seq SW620 Human | SOX2 |
| 995 | PACSIN3 | Enrichr CHEA_2022: SOX2 20726797 ChIP-Seq SW620 Human | SOX2 |
| 996 | SH3BGRL3 | Enrichr CHEA_2022: SOX2 20726797 ChIP-Seq SW620 Human | SOX2 |
| 997 | STAP2 | Enrichr CHEA_2022: SOX2 20726797 ChIP-Seq SW620 Human | SOX2 |
| 998 | CTBS | Enrichr CHEA_2022: SOX2 20726797 ChIP-Seq SW620 Human | SOX2 |
| 999 | CGNL1 | Enrichr CHEA_2022: SOX2 20726797 ChIP-Seq SW620 Human | SOX2 |
| 1000 | ELOVL1 | Enrichr CHEA_2022: SOX2 20726797 ChIP-Seq SW620 Human | SOX2 |
| 1001 | GPR37 | Enrichr CHEA_2022: SOX2 20726797 ChIP-Seq SW620 Human | SOX2 |
| 1002 | NANOS1 | Enrichr CHEA_2022: SOX2 20726797 ChIP-Seq SW620 Human | SOX2 |
| 1003 | AADAT | Enrichr CHEA_2022: SOX2 20726797 ChIP-Seq SW620 Human | SOX2 |
| 1004 | CSNK2A2 | Enrichr CHEA_2022: SOX2 20726797 ChIP-Seq SW620 Human | SOX2 |
| 1005 | ELOVL5 | Enrichr CHEA_2022: SOX2 20726797 ChIP-Seq SW620 Human | SOX2 |
| 1006 | ARG2 | Enrichr CHEA_2022: SOX2 20726797 ChIP-Seq SW620 Human | SOX2 |
| 1007 | STAT3 | Enrichr CHEA_2022: SOX2 20726797 ChIP-Seq SW620 Human | SOX2 |
| 1008 | TMEM62 | Enrichr CHEA_2022: SOX2 20726797 ChIP-Seq SW620 Human | SOX2 |
| 1009 | TRNT1 | Enrichr CHEA_2022: SOX2 20726797 ChIP-Seq SW620 Human | SOX2 |
| 1010 | ELOVL7 | Enrichr CHEA_2022: SOX2 20726797 ChIP-Seq SW620 Human | SOX2 |
| 1011 | RNASE4 | Enrichr CHEA_2022: SOX2 20726797 ChIP-Seq SW620 Human | SOX2 |
| 1012 | TMEM65 | Enrichr CHEA_2022: SOX2 20726797 ChIP-Seq SW620 Human | SOX2 |
| 1013 | LRG1 | Enrichr CHEA_2022: SOX2 20726797 ChIP-Seq SW620 Human | SOX2 |
| 1014 | F5 | Enrichr CHEA_2022: SOX2 20726797 ChIP-Seq SW620 Human | SOX2 |
| 1015 | FABP5 | Enrichr CHEA_2022: SOX2 20726797 ChIP-Seq SW620 Human | SOX2 |
| 1016 | MRPL52 | Enrichr CHEA_2022: SOX2 20726797 ChIP-Seq SW620 Human | SOX2 |
| 1017 | AADAC | Enrichr CHEA_2022: SOX2 20726797 ChIP-Seq SW620 Human | SOX2 |
| 1018 | PELI1 | Enrichr CHEA_2022: SOX2 20726797 ChIP-Seq SW620 Human | SOX2 |
| 1019 | GOPC | Enrichr CHEA_2022: SOX2 20726797 ChIP-Seq SW620 Human | SOX2 |
| 1020 | ZNF697 | Enrichr CHEA_2022: SOX2 20726797 ChIP-Seq SW620 Human | SOX2 |
| 1021 | ZNF211 | Enrichr CHEA_2022: SOX2 20726797 ChIP-Seq SW620 Human | SOX2 |
| 1022 | PIGV | Enrichr CHEA_2022: SOX2 20726797 ChIP-Seq SW620 Human | SOX2 |
| 1023 | RNASEL | Enrichr CHEA_2022: SOX2 20726797 ChIP-Seq SW620 Human | SOX2 |
| 1024 | QPRT | Enrichr CHEA_2022: SOX2 20726797 ChIP-Seq SW620 Human | SOX2 |
| 1025 | F13A1 | Enrichr CHEA_2022: SOX2 20726797 ChIP-Seq SW620 Human | SOX2 |
| 1026 | TMEM51 | Enrichr CHEA_2022: SOX2 20726797 ChIP-Seq SW620 Human | SOX2 |
| 1027 | TMEM54 | Enrichr CHEA_2022: SOX2 20726797 ChIP-Seq SW620 Human | SOX2 |
| 1028 | JPH1 | Enrichr CHEA_2022: SOX2 20726797 ChIP-Seq SW620 Human | SOX2 |
| 1029 | PTGS1 | Enrichr CHEA_2022: SOX2 20726797 ChIP-Seq SW620 Human | SOX2 |
| 1030 | PTGS2 | Enrichr CHEA_2022: SOX2 20726797 ChIP-Seq SW620 Human | SOX2 |
| 1031 | NSDHL | Enrichr CHEA_2022: SOX2 20726797 ChIP-Seq SW620 Human | SOX2 |
| 1032 | HEY1 | Enrichr CHEA_2022: SOX2 20726797 ChIP-Seq SW620 Human | SOX2 |
| 1033 | TMEM47 | Enrichr CHEA_2022: SOX2 20726797 ChIP-Seq SW620 Human | SOX2 |
| 1034 | ZNF207 | Enrichr CHEA_2022: SOX2 20726797 ChIP-Seq SW620 Human | SOX2 |
| 1035 | MYLIP | Enrichr CHEA_2022: SOX2 20726797 ChIP-Seq SW620 Human | SOX2 |

| | | | |
|---|---|---|---|
| 1036 | AGTPBP1 | Enrichr CHEA_2022: SOX2 20726797 ChIP-Seq SW620 Human | SOX2 |
| 1037 | SEC24B | Enrichr CHEA_2022: SOX2 20726797 ChIP-Seq SW620 Human | SOX2 |
| 1038 | GK | Enrichr CHEA_2022: SOX2 20726797 ChIP-Seq SW620 Human | SOX2 |
| 1039 | S100A3 | Enrichr CHEA_2022: SOX2 20726797 ChIP-Seq SW620 Human | SOX2 |
| 1040 | BRPF3 | Enrichr CHEA_2022: SOX2 20726797 ChIP-Seq SW620 Human | SOX2 |
| 1041 | S100A2 | Enrichr CHEA_2022: SOX2 20726797 ChIP-Seq SW620 Human | SOX2 |
| 1042 | CRYBB2 | Enrichr CHEA_2022: SOX2 20726797 ChIP-Seq SW620 Human | SOX2 |
| 1043 | ZBTB10 | Enrichr CHEA_2022: SOX2 20726797 ChIP-Seq SW620 Human | SOX2 |
| 1044 | SAP30 | Enrichr CHEA_2022: SOX2 20726797 ChIP-Seq SW620 Human | SOX2 |
| 1045 | MAPKAPK3 | Enrichr CHEA_2022: SOX2 20726797 ChIP-Seq SW620 Human | SOX2 |
| 1046 | RIMBP2 | Enrichr CHEA_2022: SOX2 20726797 ChIP-Seq SW620 Human | SOX2 |
| 1047 | SQSTM1 | Enrichr CHEA_2022: SOX2 20726797 ChIP-Seq SW620 Human | SOX2 |
| 1048 | SEC24D | Enrichr CHEA_2022: SOX2 20726797 ChIP-Seq SW620 Human | SOX2 |
| 1049 | ROR1 | Enrichr CHEA_2022: SOX2 20726797 ChIP-Seq SW620 Human | SOX2 |
| 1050 | UTRN | Enrichr CHEA_2022: SOX2 20726797 ChIP-Seq SW620 Human | SOX2 |
| 1051 | DSC2 | Enrichr CHEA_2022: SOX2 20726797 ChIP-Seq SW620 Human | SOX2 |
| 1052 | UGT1A10 | Enrichr CHEA_2022: SOX2 20726797 ChIP-Seq SW620 Human | SOX2 |
| 1053 | HDDC2 | Enrichr CHEA_2022: SOX2 20726797 ChIP-Seq SW620 Human | SOX2 |
| 1054 | TMF1 | Enrichr CHEA_2022: SOX2 20726797 ChIP-Seq SW620 Human | SOX2 |
| 1055 | UBE2D3 | Enrichr CHEA_2022: SOX2 20726797 ChIP-Seq SW620 Human | SOX2 |
| 1056 | HP | Enrichr CHEA_2022: SOX2 20726797 ChIP-Seq SW620 Human | SOX2 |
| 1057 | UCP2 | Enrichr CHEA_2022: SOX2 20726797 ChIP-Seq SW620 Human | SOX2 |
| 1058 | DEFB1 | Enrichr CHEA_2022: SOX2 20726797 ChIP-Seq SW620 Human | SOX2 |
| 1059 | CXCL16 | Enrichr CHEA_2022: SOX2 20726797 ChIP-Seq SW620 Human | SOX2 |
| 1060 | SH3BP2 | Enrichr CHEA_2022: SOX2 20726797 ChIP-Seq SW620 Human | SOX2 |
| 1061 | HAS2 | Enrichr CHEA_2022: SOX2 20726797 ChIP-Seq SW620 Human | SOX2 |
| 1062 | SH3BP5 | Enrichr CHEA_2022: SOX2 20726797 ChIP-Seq SW620 Human | SOX2 |
| 1063 | HAS3 | Enrichr CHEA_2022: SOX2 20726797 ChIP-Seq SW620 Human | SOX2 |
| 1064 | SLC16A9 | Enrichr CHEA_2022: SOX2 20726797 ChIP-Seq SW620 Human | SOX2 |
| 1065 | SLC16A4 | Enrichr CHEA_2022: SOX2 20726797 ChIP-Seq SW620 Human | SOX2 |
| 1066 | PITX2 | Enrichr CHEA_2022: SOX2 20726797 ChIP-Seq SW620 Human | SOX2 |
| 1067 | SLC16A5 | Enrichr CHEA_2022: SOX2 20726797 ChIP-Seq SW620 Human | SOX2 |
| 1068 | ATP9A | Enrichr CHEA_2022: SOX2 20726797 ChIP-Seq SW620 Human | SOX2 |
| 1069 | SLC16A6 | Enrichr CHEA_2022: SOX2 20726797 ChIP-Seq SW620 Human | SOX2 |
| 1070 | SLC38A2 | Enrichr CHEA_2022: SOX2 20726797 ChIP-Seq SW620 Human | SOX2 |
| 1071 | TNFRSF10B | Enrichr CHEA_2022: SOX2 20726797 ChIP-Seq SW620 Human | SOX2 |
| 1072 | FRMD6 | Enrichr CHEA_2022: SOX2 20726797 ChIP-Seq SW620 Human | SOX2 |
| 1073 | CDC42EP5 | Enrichr CHEA_2022: SOX2 20726797 ChIP-Seq SW620 Human | SOX2 |
| 1074 | CDC42EP4 | Enrichr CHEA_2022: SOX2 20726797 ChIP-Seq SW620 Human | SOX2 |
| 1075 | BIRC5 | Enrichr CHEA_2022: SOX2 20726797 ChIP-Seq SW620 Human | SOX2 |
| 1076 | FSCN1 | Enrichr CHEA_2022: SOX2 20726797 ChIP-Seq SW620 Human | SOX2 |
| 1077 | CDC42EP2 | Enrichr CHEA_2022: SOX2 20726797 ChIP-Seq SW620 Human | SOX2 |
| 1078 | ZNF655 | Enrichr CHEA_2022: SOX2 20726797 ChIP-Seq SW620 Human | SOX2 |
| 1079 | ZNF654 | Enrichr CHEA_2022: SOX2 20726797 ChIP-Seq SW620 Human | SOX2 |
| 1080 | LRRC57 | Enrichr CHEA_2022: SOX2 20726797 ChIP-Seq SW620 Human | SOX2 |
| 1081 | PFKFB4 | Enrichr CHEA_2022: SOX2 20726797 ChIP-Seq SW620 Human | SOX2 |
| 1082 | GPR87 | Enrichr CHEA_2022: SOX2 20726797 ChIP-Seq SW620 Human | SOX2 |
| 1083 | TACSTD2 | Enrichr CHEA_2022: SOX2 20726797 ChIP-Seq SW620 Human | SOX2 |
| 1084 | SAT2 | Enrichr CHEA_2022: SOX2 20726797 ChIP-Seq SW620 Human | SOX2 |
| 1085 | HNMT | Enrichr CHEA_2022: SOX2 20726797 ChIP-Seq SW620 Human | SOX2 |
| 1086 | GLIS2 | Enrichr CHEA_2022: SOX2 20726797 ChIP-Seq SW620 Human | SOX2 |
| 1087 | PORCN | Enrichr CHEA_2022: SOX2 20726797 ChIP-Seq SW620 Human | SOX2 |
| 1088 | SESN2 | Enrichr CHEA_2022: SOX2 20726797 ChIP-Seq SW620 Human | SOX2 |
| 1089 | SESN3 | Enrichr CHEA_2022: SOX2 20726797 ChIP-Seq SW620 Human | SOX2 |
| 1090 | LEPR | Enrichr CHEA_2022: SOX2 20726797 ChIP-Seq SW620 Human | SOX2 |
| 1091 | MBP | Enrichr CHEA_2022: SOX2 20726797 ChIP-Seq SW620 Human | SOX2 |
| 1092 | SLC38A5 | Enrichr CHEA_2022: SOX2 20726797 ChIP-Seq SW620 Human | SOX2 |
| 1093 | DGCR2 | Enrichr CHEA_2022: SOX2 20726797 ChIP-Seq SW620 Human | SOX2 |
| 1094 | CYB5B | Enrichr CHEA_2022: SOX2 20726797 ChIP-Seq SW620 Human | SOX2 |
| 1095 | SETDB2 | Enrichr CHEA_2022: SOX2 20726797 ChIP-Seq SW620 Human | SOX2 |
| 1096 | MCC | Enrichr CHEA_2022: SOX2 20726797 ChIP-Seq SW620 Human | SOX2 |

| | | | |
|---|---|---|---|
| 1097 | CYB5A | Enrichr CHEA_2022: SOX2 20726797 ChIP-Seq SW620 Human | SOX2 |
| 1098 | QKI | Enrichr CHEA_2022: SOX2 20726797 ChIP-Seq SW620 Human | SOX2 |
| 1099 | UST | Enrichr CHEA_2022: SOX2 20726797 ChIP-Seq SW620 Human | SOX2 |
| 1100 | AAK1 | Enrichr CHEA_2022: SOX2 20726797 ChIP-Seq SW620 Human | SOX2 |
| 1101 | SLC25A1 | Enrichr CHEA_2022: SOX2 20726797 ChIP-Seq SW620 Human | SOX2 |
| 1102 | SLC25A4 | Enrichr CHEA_2022: SOX2 20726797 ChIP-Seq SW620 Human | SOX2 |
| 1103 | RAP1GAP | Enrichr CHEA_2022: SOX2 20726797 ChIP-Seq SW620 Human | SOX2 |
| 1104 | PELP1 | Enrichr CHEA_2022: SOX2 20726797 ChIP-Seq SW620 Human | SOX2 |
| 1105 | FCHO2 | Enrichr CHEA_2022: SOX2 20726797 ChIP-Seq SW620 Human | SOX2 |
| 1106 | DSP | Enrichr CHEA_2022: SOX2 20726797 ChIP-Seq SW620 Human | SOX2 |
| 1107 | NDUFA8 | Enrichr CHEA_2022: SOX2 20726797 ChIP-Seq SW620 Human | SOX2 |
| 1108 | SEC22A | Enrichr CHEA_2022: SOX2 20726797 ChIP-Seq SW620 Human | SOX2 |
| 1109 | DST | Enrichr CHEA_2022: SOX2 20726797 ChIP-Seq SW620 Human | SOX2 |
| 1110 | NDUFA3 | Enrichr CHEA_2022: SOX2 20726797 ChIP-Seq SW620 Human | SOX2 |
| 1111 | LYSMD2 | Enrichr CHEA_2022: SOX2 20726797 ChIP-Seq SW620 Human | SOX2 |
| 1112 | TPD52L1 | Enrichr CHEA_2022: SOX2 20726797 ChIP-Seq SW620 Human | SOX2 |
| 1113 | PPM1A | Enrichr CHEA_2022: SOX2 20726797 ChIP-Seq SW620 Human | SOX2 |
| 1114 | ZNF618 | Enrichr CHEA_2022: SOX2 20726797 ChIP-Seq SW620 Human | SOX2 |
| 1115 | MET | Enrichr CHEA_2022: SOX2 20726797 ChIP-Seq SW620 Human | SOX2 |
| 1116 | SEC22C | Enrichr CHEA_2022: SOX2 20726797 ChIP-Seq SW620 Human | SOX2 |
| 1117 | SLC25A14 | Enrichr CHEA_2022: SOX2 20726797 ChIP-Seq SW620 Human | SOX2 |
| 1118 | EPHB6 | Enrichr CHEA_2022: SOX2 20726797 ChIP-Seq SW620 Human | SOX2 |
| 1119 | FHL1 | Enrichr CHEA_2022: SOX2 20726797 ChIP-Seq SW620 Human | SOX2 |
| 1120 | PIK3CD | Enrichr CHEA_2022: SOX2 20726797 ChIP-Seq SW620 Human | SOX2 |
| 1121 | PTPRZ1 | Enrichr CHEA_2022: SOX2 20726797 ChIP-Seq SW620 Human | SOX2 |
| 1122 | SLITRK6 | Enrichr CHEA_2022: SOX2 20726797 ChIP-Seq SW620 Human | SOX2 |
| 1123 | UPP1 | Enrichr CHEA_2022: SOX2 20726797 ChIP-Seq SW620 Human | SOX2 |
| 1124 | EPHB2 | Enrichr CHEA_2022: SOX2 20726797 ChIP-Seq SW620 Human | SOX2 |
| 1125 | EPHB4 | Enrichr CHEA_2022: SOX2 20726797 ChIP-Seq SW620 Human | SOX2 |
| 1126 | ACVR1 | Enrichr CHEA_2022: SOX2 20726797 ChIP-Seq SW620 Human | SOX2 |
| 1127 | ZFP1 | Enrichr CHEA_2022: SOX2 20726797 ChIP-Seq SW620 Human | SOX2 |
| 1128 | SLC12A2 | Enrichr CHEA_2022: SOX2 20726797 ChIP-Seq SW620 Human | SOX2 |
| 1129 | PLEKHA6 | Enrichr CHEA_2022: SOX2 20726797 ChIP-Seq SW620 Human | SOX2 |
| 1130 | TM4SF4 | Enrichr CHEA_2022: SOX2 20726797 ChIP-Seq SW620 Human | SOX2 |
| 1131 | EREG | Enrichr CHEA_2022: SOX2 20726797 ChIP-Seq SW620 Human | SOX2 |
| 1132 | PPP1R3D | Enrichr CHEA_2022: SOX2 20726797 ChIP-Seq SW620 Human | SOX2 |
| 1133 | IFI27 | Enrichr CHEA_2022: SOX2 20726797 ChIP-Seq SW620 Human | SOX2 |
| 1134 | LACTB2 | Enrichr CHEA_2022: SOX2 20726797 ChIP-Seq SW620 Human | SOX2 |
| 1135 | MXRA7 | Enrichr CHEA_2022: SOX2 20726797 ChIP-Seq SW620 Human | SOX2 |
| 1136 | COL9A3 | Enrichr CHEA_2022: SOX2 20726797 ChIP-Seq SW620 Human | SOX2 |
| 1137 | ALPK3 | Enrichr CHEA_2022: SOX2 20726797 ChIP-Seq SW620 Human | SOX2 |
| 1138 | EPHA1 | Enrichr CHEA_2022: SOX2 20726797 ChIP-Seq SW620 Human | SOX2 |
| 1139 | EPHA4 | Enrichr CHEA_2022: SOX2 20726797 ChIP-Seq SW620 Human | SOX2 |
| 1140 | TMEM87A | Enrichr CHEA_2022: SOX2 20726797 ChIP-Seq SW620 Human | SOX2 |
| 1141 | SATB1 | Enrichr CHEA_2022: SOX2 20726797 ChIP-Seq SW620 Human | SOX2 |
| 1142 | KDELR2 | Enrichr CHEA_2022: SOX2 20726797 ChIP-Seq SW620 Human | SOX2 |
| 1143 | PEX11A | Enrichr CHEA_2022: SOX2 20726797 ChIP-Seq SW620 Human | SOX2 |
| 1144 | PEX11B | Enrichr CHEA_2022: SOX2 20726797 ChIP-Seq SW620 Human | SOX2 |
| 1145 | LRP4 | Enrichr CHEA_2022: SOX2 20726797 ChIP-Seq SW620 Human | SOX2 |
| 1146 | PRCP | Enrichr CHEA_2022: SOX2 20726797 ChIP-Seq SW620 Human | SOX2 |
| 1147 | CYB5D1 | Enrichr CHEA_2022: SOX2 20726797 ChIP-Seq SW620 Human | SOX2 |
| 1148 | UBE2L6 | Enrichr CHEA_2022: SOX2 20726797 ChIP-Seq SW620 Human | SOX2 |
| 1149 | LRP8 | Enrichr CHEA_2022: SOX2 20726797 ChIP-Seq SW620 Human | SOX2 |
| 1150 | NTN4 | Enrichr CHEA_2022: SOX2 20726797 ChIP-Seq SW620 Human | SOX2 |
| 1151 | CYB5D2 | Enrichr CHEA_2022: SOX2 20726797 ChIP-Seq SW620 Human | SOX2 |
| 1152 | BAIAP2L1 | Enrichr CHEA_2022: SOX2 20726797 ChIP-Seq SW620 Human | SOX2 |
| 1153 | RAB11FIP1 | Enrichr CHEA_2022: SOX2 20726797 ChIP-Seq SW620 Human | SOX2 |
| 1154 | EXOSC6 | Enrichr CHEA_2022: SOX2 20726797 ChIP-Seq SW620 Human | SOX2 |
| 1155 | PARD6B | Enrichr CHEA_2022: SOX2 20726797 ChIP-Seq SW620 Human | SOX2 |
| 1156 | BAIAP2L2 | Enrichr CHEA_2022: SOX2 20726797 ChIP-Seq SW620 Human | SOX2 |
| 1157 | CHORDC1 | Enrichr CHEA_2022: SOX2 20726797 ChIP-Seq SW620 Human | SOX2 |

| | | | |
|---|---|---|---|
| 1158 | TGOLN2 | Enrichr CHEA_2022: SOX2 20726797 ChIP-Seq SW620 Human | SOX2 |
| 1159 | POF1B | Enrichr CHEA_2022: SOX2 20726797 ChIP-Seq SW620 Human | SOX2 |
| 1160 | RAB11FIP4 | Enrichr CHEA_2022: SOX2 20726797 ChIP-Seq SW620 Human | SOX2 |
| 1161 | WWTR1 | Enrichr CHEA_2022: SOX2 20726797 ChIP-Seq SW620 Human | SOX2 |
| 1162 | PAQR5 | Enrichr CHEA_2022: SOX2 20726797 ChIP-Seq SW620 Human | SOX2 |
| 1163 | FOXF2 | Enrichr CHEA_2022: SOX2 20726797 ChIP-Seq SW620 Human | SOX2 |
| 1164 | ST6GAL1 | Enrichr CHEA_2022: SOX2 20726797 ChIP-Seq SW620 Human | SOX2 |
| 1165 | PAQR3 | Enrichr CHEA_2022: SOX2 20726797 ChIP-Seq SW620 Human | SOX2 |
| 1166 | PLA2G4A | Enrichr CHEA_2022: SOX2 20726797 ChIP-Seq SW620 Human | SOX2 |
| 1167 | HSPA12A | Enrichr CHEA_2022: SOX2 20726797 ChIP-Seq SW620 Human | SOX2 |
| 1168 | IFI44 | Enrichr CHEA_2022: SOX2 20726797 ChIP-Seq SW620 Human | SOX2 |
| 1169 | PIK3R1 | Enrichr CHEA_2022: SOX2 20726797 ChIP-Seq SW620 Human | SOX2 |
| 1170 | TAGLN2 | Enrichr CHEA_2022: SOX2 20726797 ChIP-Seq SW620 Human | SOX2 |
| 1171 | RGL2 | Enrichr CHEA_2022: SOX2 20726797 ChIP-Seq SW620 Human | SOX2 |
| 1172 | RGL1 | Enrichr CHEA_2022: SOX2 20726797 ChIP-Seq SW620 Human | SOX2 |
| 1173 | SPON2 | Enrichr CHEA_2022: SOX2 20726797 ChIP-Seq SW620 Human | SOX2 |
| 1174 | IFI35 | Enrichr CHEA_2022: SOX2 20726797 ChIP-Seq SW620 Human | SOX2 |
| 1175 | PRF1 | Enrichr CHEA_2022: SOX2 20726797 ChIP-Seq SW620 Human | SOX2 |
| 1176 | BRCC3 | Enrichr CHEA_2022: SOX2 20726797 ChIP-Seq SW620 Human | SOX2 |
| 1177 | PELO | Enrichr CHEA_2022: SOX2 20726797 ChIP-Seq SW620 Human | SOX2 |
| 1178 | RASSF3 | Enrichr CHEA_2022: SOX2 20726797 ChIP-Seq SW620 Human | SOX2 |
| 1179 | RASSF5 | Enrichr CHEA_2022: SOX2 20726797 ChIP-Seq SW620 Human | SOX2 |
| 1180 | RASSF6 | Enrichr CHEA_2022: SOX2 20726797 ChIP-Seq SW620 Human | SOX2 |
| 1181 | MICAL2 | Enrichr CHEA_2022: SOX2 20726797 ChIP-Seq SW620 Human | SOX2 |
| 1182 | TSPAN8 | Enrichr CHEA_2022: SOX2 20726797 ChIP-Seq SW620 Human | SOX2 |
| 1183 | ANKH | Enrichr CHEA_2022: SOX2 20726797 ChIP-Seq SW620 Human | SOX2 |
| 1184 | ARSJ | Enrichr CHEA_2022: SOX2 20726797 ChIP-Seq SW620 Human | SOX2 |
| 1185 | TSPAN6 | Enrichr CHEA_2022: SOX2 20726797 ChIP-Seq SW620 Human | SOX2 |
| 1186 | ZBED2 | Enrichr CHEA_2022: SOX2 20726797 ChIP-Seq SW620 Human | SOX2 |
| 1187 | KIF21B | Enrichr CHEA_2022: SOX2 20726797 ChIP-Seq SW620 Human | SOX2 |
| 1188 | TSPAN3 | Enrichr CHEA_2022: SOX2 20726797 ChIP-Seq SW620 Human | SOX2 |
| 1189 | CD14 | Enrichr CHEA_2022: SOX2 20726797 ChIP-Seq SW620 Human | SOX2 |
| 1190 | TSPAN1 | Enrichr CHEA_2022: SOX2 20726797 ChIP-Seq SW620 Human | SOX2 |
| 1191 | ACAD11 | Enrichr CHEA_2022: SOX2 20726797 ChIP-Seq SW620 Human | SOX2 |
| 1192 | PGBD1 | Enrichr CHEA_2022: SOX2 20726797 ChIP-Seq SW620 Human | SOX2 |
| 1193 | ARSD | Enrichr CHEA_2022: SOX2 20726797 ChIP-Seq SW620 Human | SOX2 |
| 1194 | CTSB | Enrichr CHEA_2022: SOX2 20726797 ChIP-Seq SW620 Human | SOX2 |
| 1195 | NBPF3 | Enrichr CHEA_2022: SOX2 20726797 ChIP-Seq SW620 Human | SOX2 |
| 1196 | FOXD1 | Enrichr CHEA_2022: SOX2 20726797 ChIP-Seq SW620 Human | SOX2 |
| 1197 | TNFRSF12A | Enrichr CHEA_2022: SOX2 20726797 ChIP-Seq SW620 Human | SOX2 |
| 1198 | MMEL1 | Enrichr CHEA_2022: SOX2 20726797 ChIP-Seq SW620 Human | SOX2 |
| 1199 | KLK10 | Enrichr CHEA_2022: SOX2 20726797 ChIP-Seq SW620 Human | SOX2 |
| 1200 | KLK11 | Enrichr CHEA_2022: SOX2 20726797 ChIP-Seq SW620 Human | SOX2 |
| 1201 | RGMA | Enrichr CHEA_2022: SOX2 20726797 ChIP-Seq SW620 Human | SOX2 |
| 1202 | ENAH | Enrichr CHEA_2022: SOX2 20726797 ChIP-Seq SW620 Human | SOX2 |
| 1203 | NR5A2 | Enrichr CHEA_2022: SOX2 20726797 ChIP-Seq SW620 Human | SOX2 |
| 1204 | PTPRF | Enrichr CHEA_2022: SOX2 20726797 ChIP-Seq SW620 Human | SOX2 |
| 1205 | SERGEF | Enrichr CHEA_2022: SOX2 20726797 ChIP-Seq SW620 Human | SOX2 |
| 1206 | TH | Enrichr CHEA_2022: SOX2 20726797 ChIP-Seq SW620 Human | SOX2 |
| 1207 | PTPRB | Enrichr CHEA_2022: SOX2 20726797 ChIP-Seq SW620 Human | SOX2 |
| 1208 | VSNL1 | Enrichr CHEA_2022: SOX2 20726797 ChIP-Seq SW620 Human | SOX2 |
| 1209 | ID2 | Enrichr CHEA_2022: SOX2 20726797 ChIP-Seq SW620 Human | SOX2 |
| 1210 | MKX | Enrichr CHEA_2022: SOX2 20726797 ChIP-Seq SW620 Human | SOX2 |
| 1211 | CD24 | Enrichr CHEA_2022: SOX2 20726797 ChIP-Seq SW620 Human | SOX2 |
| 1212 | FOXC1 | Enrichr CHEA_2022: SOX2 20726797 ChIP-Seq SW620 Human | SOX2 |
| 1213 | CETN2 | Enrichr CHEA_2022: SOX2 20726797 ChIP-Seq SW620 Human | SOX2 |
| 1214 | PTPRO | Enrichr CHEA_2022: SOX2 20726797 ChIP-Seq SW620 Human | SOX2 |
| 1215 | INTS10 | Enrichr CHEA_2022: SOX2 20726797 ChIP-Seq SW620 Human | SOX2 |
| 1216 | TMPRSS4 | Enrichr CHEA_2022: SOX2 20726797 ChIP-Seq SW620 Human | SOX2 |
| 1217 | CTSZ | Enrichr CHEA_2022: SOX2 20726797 ChIP-Seq SW620 Human | SOX2 |
| 1218 | KIF15 | Enrichr CHEA_2022: SOX2 20726797 ChIP-Seq SW620 Human | SOX2 |

| | | | |
|---|---|---|---|
| 1219 | KIF14 | Enrichr CHEA_2022: SOX2 20726797 ChIP-Seq SW620 Human | SOX2 |
| 1220 | HNF4G | Enrichr CHEA_2022: SOX2 20726797 ChIP-Seq SW620 Human | SOX2 |
| 1221 | TMPRSS3 | Enrichr CHEA_2022: SOX2 20726797 ChIP-Seq SW620 Human | SOX2 |
| 1222 | PCSK9 | Enrichr CHEA_2022: SOX2 20726797 ChIP-Seq SW620 Human | SOX2 |
| 1223 | NEDD9 | Enrichr CHEA_2022: SOX2 20726797 ChIP-Seq SW620 Human | SOX2 |
| 1224 | PCSK6 | Enrichr CHEA_2022: SOX2 20726797 ChIP-Seq SW620 Human | SOX2 |
| 1225 | CTSS | Enrichr CHEA_2022: SOX2 20726797 ChIP-Seq SW620 Human | SOX2 |
| 1226 | PTPRH | Enrichr CHEA_2022: SOX2 20726797 ChIP-Seq SW620 Human | SOX2 |
| 1227 | ACACA | Enrichr CHEA_2022: SOX2 20726797 ChIP-Seq SW620 Human | SOX2 |
| 1228 | SNX5 | Enrichr CHEA_2022: SOX2 20726797 ChIP-Seq SW620 Human | SOX2 |
| 1229 | SOCS2 | Enrichr CHEA_2022: SOX2 20726797 ChIP-Seq SW620 Human | SOX2 |
| 1230 | SNX1 | Enrichr CHEA_2022: SOX2 20726797 ChIP-Seq SW620 Human | SOX2 |
| 1231 | IDS | Enrichr CHEA_2022: SOX2 20726797 ChIP-Seq SW620 Human | SOX2 |
| 1232 | CTSL | Enrichr CHEA_2022: SOX2 20726797 ChIP-Seq SW620 Human | SOX2 |
| 1233 | STK39 | Enrichr CHEA_2022: SOX2 20726797 ChIP-Seq SW620 Human | SOX2 |
| 1234 | CTSH | Enrichr CHEA_2022: SOX2 20726797 ChIP-Seq SW620 Human | SOX2 |
| 1235 | LMCD1 | Enrichr CHEA_2022: SOX2 20726797 ChIP-Seq SW620 Human | SOX2 |
| 1236 | SNX9 | Enrichr CHEA_2022: SOX2 20726797 ChIP-Seq SW620 Human | SOX2 |
| 1237 | CTSD | Enrichr CHEA_2022: SOX2 20726797 ChIP-Seq SW620 Human | SOX2 |
| 1238 | SOCS5 | Enrichr CHEA_2022: SOX2 20726797 ChIP-Seq SW620 Human | SOX2 |
| 1239 | SOCS6 | Enrichr CHEA_2022: SOX2 20726797 ChIP-Seq SW620 Human | SOX2 |
| 1240 | MMD | Enrichr CHEA_2022: SOX2 20726797 ChIP-Seq SW620 Human | SOX2 |
| 1241 | BCAP31 | Enrichr CHEA_2022: SOX2 20726797 ChIP-Seq SW620 Human | SOX2 |
| 1242 | MME | Enrichr CHEA_2022: SOX2 20726797 ChIP-Seq SW620 Human | SOX2 |
| 1243 | AP2B1 | Enrichr CHEA_2022: SOX2 20726797 ChIP-Seq SW620 Human | SOX2 |
| 1244 | UBE2E2 | Enrichr CHEA_2022: SOX2 20726797 ChIP-Seq SW620 Human | SOX2 |
| 1245 | MIB1 | Enrichr CHEA_2022: SOX2 20726797 ChIP-Seq SW620 Human | SOX2 |
| 1246 | FKBP1B | Enrichr CHEA_2022: SOX2 20726797 ChIP-Seq SW620 Human | SOX2 |
| 1247 | NEDD1 | Enrichr CHEA_2022: SOX2 20726797 ChIP-Seq SW620 Human | SOX2 |
| 1248 | STK17B | Enrichr CHEA_2022: SOX2 20726797 ChIP-Seq SW620 Human | SOX2 |
| 1249 | CCNE2 | Enrichr CHEA_2022: SOX2 20726797 ChIP-Seq SW620 Human | SOX2 |
| 1250 | CCNE1 | Enrichr CHEA_2022: SOX2 20726797 ChIP-Seq SW620 Human | SOX2 |
| 1251 | EPPK1 | Enrichr CHEA_2022: SOX2 20726797 ChIP-Seq SW620 Human | SOX2 |
| 1252 | IL17D | Enrichr CHEA_2022: SOX2 20726797 ChIP-Seq SW620 Human | SOX2 |
| 1253 | CD44 | Enrichr CHEA_2022: SOX2 20726797 ChIP-Seq SW620 Human | SOX2 |
| 1254 | NPDC1 | Enrichr CHEA_2022: SOX2 20726797 ChIP-Seq SW620 Human | SOX2 |
| 1255 | DBN1 | Enrichr CHEA_2022: SOX2 20726797 ChIP-Seq SW620 Human | SOX2 |
| 1256 | FOXA2 | Enrichr CHEA_2022: SOX2 20726797 ChIP-Seq SW620 Human | SOX2 |
| 1257 | ERRFI1 | Enrichr CHEA_2022: SOX2 20726797 ChIP-Seq SW620 Human | SOX2 |
| 1258 | FOXA1 | Enrichr CHEA_2022: SOX2 20726797 ChIP-Seq SW620 Human | SOX2 |
| 1259 | FKBP11 | Enrichr CHEA_2022: SOX2 20726797 ChIP-Seq SW620 Human | SOX2 |
| 1260 | SLC43A3 | Enrichr CHEA_2022: SOX2 20726797 ChIP-Seq SW620 Human | SOX2 |
| 1261 | SLC43A1 | Enrichr CHEA_2022: SOX2 20726797 ChIP-Seq SW620 Human | SOX2 |
| 1262 | FKBP14 | Enrichr CHEA_2022: SOX2 20726797 ChIP-Seq SW620 Human | SOX2 |
| 1263 | AKR1B1 | Enrichr CHEA_2022: SOX2 20726797 ChIP-Seq SW620 Human | SOX2 |
| 1264 | FHOD3 | Enrichr CHEA_2022: SOX2 20726797 ChIP-Seq SW620 Human | SOX2 |
| 1265 | WASL | Enrichr CHEA_2022: SOX2 20726797 ChIP-Seq SW620 Human | SOX2 |
| 1266 | DERL1 | Enrichr CHEA_2022: SOX2 20726797 ChIP-Seq SW620 Human | SOX2 |
| 1267 | ZDHHC2 | Enrichr CHEA_2022: SOX2 20726797 ChIP-Seq SW620 Human | SOX2 |
| 1268 | CRIP1 | Enrichr CHEA_2022: SOX2 20726797 ChIP-Seq SW620 Human | SOX2 |
| 1269 | CRIP2 | Enrichr CHEA_2022: SOX2 20726797 ChIP-Seq SW620 Human | SOX2 |
| 1270 | SCML1 | Enrichr CHEA_2022: SOX2 20726797 ChIP-Seq SW620 Human | SOX2 |
| 1271 | PGRMC2 | Enrichr CHEA_2022: SOX2 20726797 ChIP-Seq SW620 Human | SOX2 |
| 1272 | SPTLC2 | Enrichr CHEA_2022: SOX2 20726797 ChIP-Seq SW620 Human | SOX2 |
| 1273 | CAMK2N1 | Enrichr CHEA_2022: SOX2 20726797 ChIP-Seq SW620 Human | SOX2 |
| 1274 | SERTAD2 | Enrichr CHEA_2022: SOX2 20726797 ChIP-Seq SW620 Human | SOX2 |
| 1275 | SERTAD3 | Enrichr CHEA_2022: SOX2 20726797 ChIP-Seq SW620 Human | SOX2 |
| 1276 | ENC1 | Enrichr CHEA_2022: SOX2 20726797 ChIP-Seq SW620 Human | SOX2 |
| 1277 | SERTAD1 | Enrichr CHEA_2022: SOX2 20726797 ChIP-Seq SW620 Human | SOX2 |
| 1278 | CD59 | Enrichr CHEA_2022: SOX2 20726797 ChIP-Seq SW620 Human | SOX2 |
| 1279 | CD55 | Enrichr CHEA_2022: SOX2 20726797 ChIP-Seq SW620 Human | SOX2 |

| | | | |
|---|---|---|---|
| 1280 | NPNT | Enrichr CHEA_2022: SOX2 20726797 ChIP-Seq SW620 Human | SOX2 |
| 1281 | G6PD | Enrichr CHEA_2022: SOX2 20726797 ChIP-Seq SW620 Human | SOX2 |
| 1282 | CRIM1 | Enrichr CHEA_2022: SOX2 20726797 ChIP-Seq SW620 Human | SOX2 |
| 1283 | MID1 | Enrichr CHEA_2022: SOX2 20726797 ChIP-Seq SW620 Human | SOX2 |
| 1284 | TANC2 | Enrichr CHEA_2022: SOX2 20726797 ChIP-Seq SW620 Human | SOX2 |
| 1285 | XK | Enrichr CHEA_2022: SOX2 20726797 ChIP-Seq SW620 Human | SOX2 |
| 1286 | IVNS1ABP | Enrichr CHEA_2022: SOX2 20726797 ChIP-Seq SW620 Human | SOX2 |
| 1287 | CD82 | Enrichr CHEA_2022: SOX2 20726797 ChIP-Seq SW620 Human | SOX2 |
| 1288 | SLC1A1 | Enrichr CHEA_2022: SOX2 20726797 ChIP-Seq SW620 Human | SOX2 |
| 1289 | AURKB | Enrichr CHEA_2022: SOX2 20726797 ChIP-Seq SW620 Human | SOX2 |
| 1290 | RXRA | Enrichr CHEA_2022: SOX2 20726797 ChIP-Seq SW620 Human | SOX2 |
| 1291 | RAB40B | Enrichr CHEA_2022: SOX2 20726797 ChIP-Seq SW620 Human | SOX2 |
| 1292 | SH3TC2 | Enrichr CHEA_2022: SOX2 20726797 ChIP-Seq SW620 Human | SOX2 |
| 1293 | HSF2 | Enrichr CHEA_2022: SOX2 20726797 ChIP-Seq SW620 Human | SOX2 |
| 1294 | NPHP3 | Enrichr CHEA_2022: SOX2 20726797 ChIP-Seq SW620 Human | SOX2 |
| 1295 | CCRL2 | Enrichr CHEA_2022: SOX2 20726797 ChIP-Seq SW620 Human | SOX2 |
| 1296 | GPC4 | Enrichr CHEA_2022: SOX2 20726797 ChIP-Seq SW620 Human | SOX2 |
| 1297 | PAK6 | Enrichr CHEA_2022: SOX2 20726797 ChIP-Seq SW620 Human | SOX2 |
| 1298 | MICB | Enrichr CHEA_2022: SOX2 20726797 ChIP-Seq SW620 Human | SOX2 |
| 1299 | SLC30A7 | Enrichr CHEA_2022: SOX2 20726797 ChIP-Seq SW620 Human | SOX2 |
| 1300 | INHBB | Enrichr CHEA_2022: SOX2 20726797 ChIP-Seq SW620 Human | SOX2 |
| 1301 | ISG15 | Enrichr CHEA_2022: SOX2 20726797 ChIP-Seq SW620 Human | SOX2 |
| 1302 | SSH3 | Enrichr CHEA_2022: SOX2 20726797 ChIP-Seq SW620 Human | SOX2 |
| 1303 | PAN3 | Enrichr CHEA_2022: SOX2 20726797 ChIP-Seq SW620 Human | SOX2 |
| 1304 | QDPR | Enrichr CHEA_2022: SOX2 20726797 ChIP-Seq SW620 Human | SOX2 |
| 1305 | PROCR | Enrichr CHEA_2022: SOX2 20726797 ChIP-Seq SW620 Human | SOX2 |
| 1306 | GEMIN7 | Enrichr CHEA_2022: SOX2 20726797 ChIP-Seq SW620 Human | SOX2 |
| 1307 | TAB3 | Enrichr CHEA_2022: SOX2 20726797 ChIP-Seq SW620 Human | SOX2 |
| 1308 | GCLM | Enrichr CHEA_2022: SOX2 20726797 ChIP-Seq SW620 Human | SOX2 |
| 1309 | BLVRB | Enrichr CHEA_2022: SOX2 20726797 ChIP-Seq SW620 Human | SOX2 |
| 1310 | ARHGEF10 | Enrichr CHEA_2022: SOX2 20726797 ChIP-Seq SW620 Human | SOX2 |
| 1311 | LRRFIP1 | Enrichr CHEA_2022: SOX2 20726797 ChIP-Seq SW620 Human | SOX2 |
| 1312 | KLHL15 | Enrichr CHEA_2022: SOX2 20726797 ChIP-Seq SW620 Human | SOX2 |
| 1313 | WWC1 | Enrichr CHEA_2022: SOX2 20726797 ChIP-Seq SW620 Human | SOX2 |
| 1314 | WWC2 | Enrichr CHEA_2022: SOX2 20726797 ChIP-Seq SW620 Human | SOX2 |
| 1315 | RGS3 | Enrichr CHEA_2022: SOX2 20726797 ChIP-Seq SW620 Human | SOX2 |
| 1316 | CALD1 | Enrichr CHEA_2022: SOX2 20726797 ChIP-Seq SW620 Human | SOX2 |
| 1317 | RGS2 | Enrichr CHEA_2022: SOX2 20726797 ChIP-Seq SW620 Human | SOX2 |
| 1318 | NDEL1 | Enrichr CHEA_2022: SOX2 20726797 ChIP-Seq SW620 Human | SOX2 |
| 1319 | CD99 | Enrichr CHEA_2022: SOX2 20726797 ChIP-Seq SW620 Human | SOX2 |
| 1320 | MMP7 | Enrichr CHEA_2022: SOX2 20726797 ChIP-Seq SW620 Human | SOX2 |
| 1321 | GALNT6 | Enrichr CHEA_2022: SOX2 20726797 ChIP-Seq SW620 Human | SOX2 |
| 1322 | GALNT5 | Enrichr CHEA_2022: SOX2 20726797 ChIP-Seq SW620 Human | SOX2 |
| 1323 | GALNT4 | Enrichr CHEA_2022: SOX2 20726797 ChIP-Seq SW620 Human | SOX2 |
| 1324 | GALNT3 | Enrichr CHEA_2022: SOX2 20726797 ChIP-Seq SW620 Human | SOX2 |
| 1325 | HMGA2 | Enrichr CHEA_2022: SOX2 20726797 ChIP-Seq SW620 Human | SOX2 |
| 1326 | GULP1 | Enrichr CHEA_2022: SOX2 20726797 ChIP-Seq SW620 Human | SOX2 |
| 1327 | BMP7 | Enrichr CHEA_2022: SOX2 20726797 ChIP-Seq SW620 Human | SOX2 |
| 1328 | BMP5 | Enrichr CHEA_2022: SOX2 20726797 ChIP-Seq SW620 Human | SOX2 |
| 1329 | IRF1 | Enrichr CHEA_2022: SOX2 20726797 ChIP-Seq SW620 Human | SOX2 |
| 1330 | BMP4 | Enrichr CHEA_2022: SOX2 20726797 ChIP-Seq SW620 Human | SOX2 |
| 1331 | BMP2 | Enrichr CHEA_2022: SOX2 20726797 ChIP-Seq SW620 Human | SOX2 |
| 1332 | MMRN2 | Enrichr CHEA_2022: SOX2 20726797 ChIP-Seq SW620 Human | SOX2 |
| 1333 | RBM20 | Enrichr CHEA_2022: SOX2 20726797 ChIP-Seq SW620 Human | SOX2 |
| 1334 | IRF6 | Enrichr CHEA_2022: SOX2 20726797 ChIP-Seq SW620 Human | SOX2 |
| 1335 | IRF7 | Enrichr CHEA_2022: SOX2 20726797 ChIP-Seq SW620 Human | SOX2 |
| 1336 | TMEM45A | Enrichr CHEA_2022: SOX2 20726797 ChIP-Seq SW620 Human | SOX2 |
| 1337 | TMEM45B | Enrichr CHEA_2022: SOX2 20726797 ChIP-Seq SW620 Human | SOX2 |
| 1338 | PSMD11 | Enrichr CHEA_2022: SOX2 20726797 ChIP-Seq SW620 Human | SOX2 |
| 1339 | RTN3 | Enrichr CHEA_2022: SOX2 20726797 ChIP-Seq SW620 Human | SOX2 |
| 1340 | CHD2 | Enrichr CHEA_2022: SOX2 20726797 ChIP-Seq SW620 Human | SOX2 |

| | | | |
|---|---|---|---|
| 1341 | FUT2 | Enrichr CHEA_2022: SOX2 20726797 ChIP-Seq SW620 Human | SOX2 |
| 1342 | AFF1 | Enrichr CHEA_2022: SOX2 20726797 ChIP-Seq SW620 Human | SOX2 |
| 1343 | PDLIM1 | Enrichr CHEA_2022: SOX2 20726797 ChIP-Seq SW620 Human | SOX2 |
| 1344 | CALB1 | Enrichr CHEA_2022: SOX2 20726797 ChIP-Seq SW620 Human | SOX2 |
| 1345 | PDLIM4 | Enrichr CHEA_2022: SOX2 20726797 ChIP-Seq SW620 Human | SOX2 |
| 1346 | MARVELD1 | Enrichr CHEA_2022: SOX2 20726797 ChIP-Seq SW620 Human | SOX2 |
| 1347 | CALB2 | Enrichr CHEA_2022: SOX2 20726797 ChIP-Seq SW620 Human | SOX2 |
| 1348 | FUT8 | Enrichr CHEA_2022: SOX2 20726797 ChIP-Seq SW620 Human | SOX2 |
| 1349 | ISOC2 | Enrichr CHEA_2022: SOX2 20726797 ChIP-Seq SW620 Human | SOX2 |
| 1350 | PDLIM5 | Enrichr CHEA_2022: SOX2 20726797 ChIP-Seq SW620 Human | SOX2 |
| 1351 | WNT3 | Enrichr CHEA_2022: SOX2 20726797 ChIP-Seq SW620 Human | SOX2 |
| 1352 | DDX17 | Enrichr CHEA_2022: SOX2 20726797 ChIP-Seq SW620 Human | SOX2 |
| 1353 | TBC1D9 | Enrichr CHEA_2022: SOX2 20726797 ChIP-Seq SW620 Human | SOX2 |
| 1354 | ATP2B4 | Enrichr CHEA_2022: SOX2 20726797 ChIP-Seq SW620 Human | SOX2 |
| 1355 | FNDC3A | Enrichr CHEA_2022: SOX2 20726797 ChIP-Seq SW620 Human | SOX2 |
| 1356 | FNDC3B | Enrichr CHEA_2022: SOX2 20726797 ChIP-Seq SW620 Human | SOX2 |
| 1357 | ATP2B1 | Enrichr CHEA_2022: SOX2 20726797 ChIP-Seq SW620 Human | SOX2 |
| 1358 | TBC1D1 | Enrichr CHEA_2022: SOX2 20726797 ChIP-Seq SW620 Human | SOX2 |
| 1359 | TBC1D2 | Enrichr CHEA_2022: SOX2 20726797 ChIP-Seq SW620 Human | SOX2 |
| 1360 | PTHLH | Enrichr CHEA_2022: SOX2 20726797 ChIP-Seq SW620 Human | SOX2 |
| 1361 | SBK1 | Enrichr CHEA_2022: SOX2 20726797 ChIP-Seq SW620 Human | SOX2 |
| 1362 | TBC1D4 | Enrichr CHEA_2022: SOX2 20726797 ChIP-Seq SW620 Human | SOX2 |
| 1363 | TBC1D5 | Enrichr CHEA_2022: SOX2 20726797 ChIP-Seq SW620 Human | SOX2 |
| 1364 | LNX2 | Enrichr CHEA_2022: SOX2 20726797 ChIP-Seq SW620 Human | SOX2 |
| 1365 | FAM50A | Enrichr CHEA_2022: SOX2 20726797 ChIP-Seq SW620 Human | SOX2 |
| 1366 | LNX1 | Enrichr CHEA_2022: SOX2 20726797 ChIP-Seq SW620 Human | SOX2 |
| 1367 | S100P | Enrichr CHEA_2022: SOX2 20726797 ChIP-Seq SW620 Human | SOX2 |
| 1368 | CHURC1 | Enrichr CHEA_2022: SOX2 20726797 ChIP-Seq SW620 Human | SOX2 |
| 1369 | ZP3 | Enrichr CHEA_2022: SOX2 20726797 ChIP-Seq SW620 Human | SOX2 |
| 1370 | CLUAP1 | Enrichr CHEA_2022: SOX2 20726797 ChIP-Seq SW620 Human | SOX2 |
| 1371 | SCARB2 | Enrichr CHEA_2022: SOX2 20726797 ChIP-Seq SW620 Human | SOX2 |
| 1372 | DCTN6 | Enrichr CHEA_2022: SOX2 20726797 ChIP-Seq SW620 Human | SOX2 |
| 1373 | DDX28 | Enrichr CHEA_2022: SOX2 20726797 ChIP-Seq SW620 Human | SOX2 |
| 1374 | LAMA3 | Enrichr CHEA_2022: SOX2 20726797 ChIP-Seq SW620 Human | SOX2 |
| 1375 | MVP | Enrichr CHEA_2022: SOX2 20726797 ChIP-Seq SW620 Human | SOX2 |
| 1376 | ZNF292 | Enrichr CHEA_2022: SOX2 20726797 ChIP-Seq SW620 Human | SOX2 |
| 1377 | CRMP1 | Enrichr CHEA_2022: SOX2 20726797 ChIP-Seq SW620 Human | SOX2 |
| 1378 | ATP2A3 | Enrichr CHEA_2022: SOX2 20726797 ChIP-Seq SW620 Human | SOX2 |
| 1379 | ATP2A2 | Enrichr CHEA_2022: SOX2 20726797 ChIP-Seq SW620 Human | SOX2 |
| 1380 | RPGR | Enrichr CHEA_2022: SOX2 20726797 ChIP-Seq SW620 Human | SOX2 |
| 1381 | CLN6 | Enrichr CHEA_2022: SOX2 20726797 ChIP-Seq SW620 Human | SOX2 |
| 1382 | CLMN | Enrichr CHEA_2022: SOX2 20726797 ChIP-Seq SW620 Human | SOX2 |
| 1383 | EPS8L1 | Enrichr CHEA_2022: SOX2 20726797 ChIP-Seq SW620 Human | SOX2 |
| 1384 | SCN5A | Enrichr CHEA_2022: SOX2 20726797 ChIP-Seq SW620 Human | SOX2 |
| 1385 | EPS8L2 | Enrichr CHEA_2022: SOX2 20726797 ChIP-Seq SW620 Human | SOX2 |
| 1386 | NPTN | Enrichr CHEA_2022: SOX2 20726797 ChIP-Seq SW620 Human | SOX2 |
| 1387 | ACOT9 | Enrichr CHEA_2022: SOX2 20726797 ChIP-Seq SW620 Human | SOX2 |
| 1388 | MX1 | Enrichr CHEA_2022: SOX2 20726797 ChIP-Seq SW620 Human | SOX2 |
| 1389 | GLCE | Enrichr CHEA_2022: SOX2 20726797 ChIP-Seq SW620 Human | SOX2 |
| 1390 | RPL23 | Enrichr CHEA_2022: SOX2 20726797 ChIP-Seq SW620 Human | SOX2 |
| 1391 | SYTL2 | Enrichr CHEA_2022: SOX2 20726797 ChIP-Seq SW620 Human | SOX2 |
| 1392 | UPF3B | Enrichr CHEA_2022: SOX2 20726797 ChIP-Seq SW620 Human | SOX2 |
| 1393 | CHFR | Enrichr CHEA_2022: SOX2 20726797 ChIP-Seq SW620 Human | SOX2 |
| 1394 | DYNC1LI1 | Enrichr CHEA_2022: SOX2 20726797 ChIP-Seq SW620 Human | SOX2 |
| 1395 | GALNS | Enrichr CHEA_2022: SOX2 20726797 ChIP-Seq SW620 Human | SOX2 |
| 1396 | CPEB2 | Enrichr CHEA_2022: SOX2 20726797 ChIP-Seq SW620 Human | SOX2 |
| 1397 | GPR160 | Enrichr CHEA_2022: SOX2 20726797 ChIP-Seq SW620 Human | SOX2 |
| 1398 | ASB9 | Enrichr CHEA_2022: SOX2 20726797 ChIP-Seq SW620 Human | SOX2 |
| 1399 | TRIP6 | Enrichr CHEA_2022: SOX2 20726797 ChIP-Seq SW620 Human | SOX2 |
| 1400 | VIM | Enrichr CHEA_2022: SOX2 20726797 ChIP-Seq SW620 Human | SOX2 |
| 1401 | HSPA1A | Enrichr CHEA_2022: SOX2 20726797 ChIP-Seq SW620 Human | SOX2 |

| | | | |
|---|---|---|---|
| 1402 | ASF1A | Enrichr CHEA_2022: SOX2 20726797 ChIP-Seq SW620 Human | SOX2 |
| 1403 | ACOT2 | Enrichr CHEA_2022: SOX2 20726797 ChIP-Seq SW620 Human | SOX2 |
| 1404 | ASB4 | Enrichr CHEA_2022: SOX2 20726797 ChIP-Seq SW620 Human | SOX2 |
| 1405 | CPVL | Enrichr CHEA_2022: SOX2 20726797 ChIP-Seq SW620 Human | SOX2 |
| 1406 | HSPA1B | Enrichr CHEA_2022: SOX2 20726797 ChIP-Seq SW620 Human | SOX2 |
| 1407 | PARG | Enrichr CHEA_2022: SOX2 20726797 ChIP-Seq SW620 Human | SOX2 |
| 1408 | SF3B3 | Enrichr CHEA_2022: SOX2 20726797 ChIP-Seq SW620 Human | SOX2 |
| 1409 | EHF | Enrichr CHEA_2022: SOX2 20726797 ChIP-Seq SW620 Human | SOX2 |
| 1410 | MAML3 | Enrichr CHEA_2022: SOX2 20726797 ChIP-Seq SW620 Human | SOX2 |
| 1411 | AHNAK | Enrichr CHEA_2022: SOX2 20726797 ChIP-Seq SW620 Human | SOX2 |
| 1412 | SERPINE1 | Enrichr CHEA_2022: SOX2 20726797 ChIP-Seq SW620 Human | SOX2 |
| 1413 | THYN1 | Enrichr CHEA_2022: SOX2 20726797 ChIP-Seq SW620 Human | SOX2 |
| 1414 | ASAH1 | Enrichr CHEA_2022: SOX2 20726797 ChIP-Seq SW620 Human | SOX2 |
| 1415 | PCMT1 | Enrichr CHEA_2022: SOX2 20726797 ChIP-Seq SW620 Human | SOX2 |
| 1416 | CREB3L1 | Enrichr CHEA_2022: SOX2 20726797 ChIP-Seq SW620 Human | SOX2 |
| 1417 | MYB | Enrichr CHEA_2022: SOX2 20726797 ChIP-Seq SW620 Human | SOX2 |
| 1418 | CHST13 | Enrichr CHEA_2022: SOX2 20726797 ChIP-Seq SW620 Human | SOX2 |
| 1419 | CHST11 | Enrichr CHEA_2022: SOX2 20726797 ChIP-Seq SW620 Human | SOX2 |
| 1420 | S100A14 | Enrichr CHEA_2022: SOX2 20726797 ChIP-Seq SW620 Human | SOX2 |
| 1421 | SERPINF1 | Enrichr CHEA_2022: SOX2 20726797 ChIP-Seq SW620 Human | SOX2 |
| 1422 | HOXA2 | Enrichr CHEA_2022: SOX2 20726797 ChIP-Seq SW620 Human | SOX2 |
| 1423 | CYP1B1 | Enrichr CHEA_2022: SOX2 20726797 ChIP-Seq SW620 Human | SOX2 |
| 1424 | HOXA7 | Enrichr CHEA_2022: SOX2 20726797 ChIP-Seq SW620 Human | SOX2 |
| 1425 | HOXA5 | Enrichr CHEA_2022: SOX2 20726797 ChIP-Seq SW620 Human | SOX2 |
| 1426 | F8A1 | Enrichr CHEA_2022: SOX2 20726797 ChIP-Seq SW620 Human | SOX2 |
| 1427 | WNT10A | Enrichr CHEA_2022: SOX2 20726797 ChIP-Seq SW620 Human | SOX2 |
| 1428 | FZD3 | Enrichr CHEA_2022: SOX2 20726797 ChIP-Seq SW620 Human | SOX2 |
| 1429 | FZD6 | Enrichr CHEA_2022: SOX2 20726797 ChIP-Seq SW620 Human | SOX2 |
| 1430 | ST14 | Enrichr CHEA_2022: SOX2 20726797 ChIP-Seq SW620 Human | SOX2 |
| 1431 | RAB3IP | Enrichr CHEA_2022: SOX2 20726797 ChIP-Seq SW620 Human | SOX2 |
| 1432 | FZD7 | Enrichr CHEA_2022: SOX2 20726797 ChIP-Seq SW620 Human | SOX2 |
| 1433 | SLC16A10 | Enrichr CHEA_2022: SOX2 20726797 ChIP-Seq SW620 Human | SOX2 |
| 1434 | GPR137B | Enrichr CHEA_2022: SOX2 20726797 ChIP-Seq SW620 Human | SOX2 |
| 1435 | SULT2B1 | Enrichr CHEA_2022: SOX2 20726797 ChIP-Seq SW620 Human | SOX2 |
| 1436 | SLC6A8 | Enrichr CHEA_2022: SOX2 20726797 ChIP-Seq SW620 Human | SOX2 |
| 1437 | BIN3 | Enrichr CHEA_2022: SOX2 20726797 ChIP-Seq SW620 Human | SOX2 |
| 1438 | HOXB3 | Enrichr CHEA_2022: SOX2 20726797 ChIP-Seq SW620 Human | SOX2 |
| 1439 | SSPN | Enrichr CHEA_2022: SOX2 20726797 ChIP-Seq SW620 Human | SOX2 |
| 1440 | HBP1 | Enrichr CHEA_2022: SOX2 20726797 ChIP-Seq SW620 Human | SOX2 |
| 1441 | HOXB9 | Enrichr CHEA_2022: SOX2 20726797 ChIP-Seq SW620 Human | SOX2 |
| 1442 | HSPA4L | Enrichr CHEA_2022: SOX2 20726797 ChIP-Seq SW620 Human | SOX2 |
| 1443 | RPL5 | Enrichr CHEA_2022: SOX2 20726797 ChIP-Seq SW620 Human | SOX2 |
| 1444 | CDC42SE2 | Enrichr CHEA_2022: SOX2 20726797 ChIP-Seq SW620 Human | SOX2 |
| 1445 | ZNF134 | Enrichr CHEA_2022: SOX2 20726797 ChIP-Seq SW620 Human | SOX2 |
| 1446 | STMN3 | Enrichr CHEA_2022: SOX2 20726797 ChIP-Seq SW620 Human | SOX2 |
| 1447 | ENO3 | Enrichr CHEA_2022: SOX2 20726797 ChIP-Seq SW620 Human | SOX2 |
| 1448 | PRDX4 | Enrichr CHEA_2022: SOX2 20726797 ChIP-Seq SW620 Human | SOX2 |
| 1449 | ZXDB | Enrichr CHEA_2022: SOX2 20726797 ChIP-Seq SW620 Human | SOX2 |
| 1450 | STMN1 | Enrichr CHEA_2022: SOX2 20726797 ChIP-Seq SW620 Human | SOX2 |
| 1451 | RPL37 | Enrichr CHEA_2022: SOX2 20726797 ChIP-Seq SW620 Human | SOX2 |
| 1452 | PLCG1 | Enrichr CHEA_2022: SOX2 20726797 ChIP-Seq SW620 Human | SOX2 |
| 1453 | IER5 | Enrichr CHEA_2022: SOX2 20726797 ChIP-Seq SW620 Human | SOX2 |
| 1454 | ZNF124 | Enrichr CHEA_2022: SOX2 20726797 ChIP-Seq SW620 Human | SOX2 |
| 1455 | BLCAP | Enrichr CHEA_2022: SOX2 20726797 ChIP-Seq SW620 Human | SOX2 |
| 1456 | ANXA4 | Enrichr CHEA_2022: SOX2 20726797 ChIP-Seq SW620 Human | SOX2 |
| 1457 | CCL20 | Enrichr CHEA_2022: SOX2 20726797 ChIP-Seq SW620 Human | SOX2 |
| 1458 | ANXA5 | Enrichr CHEA_2022: SOX2 20726797 ChIP-Seq SW620 Human | SOX2 |
| 1459 | TESC | Enrichr CHEA_2022: SOX2 20726797 ChIP-Seq SW620 Human | SOX2 |
| 1460 | PAX6 | Enrichr CHEA_2022: SOX2 20726797 ChIP-Seq SW620 Human | SOX2 |
| 1461 | TIMM22 | Enrichr CHEA_2022: SOX2 20726797 ChIP-Seq SW620 Human | SOX2 |
| 1462 | CARD6 | Enrichr CHEA_2022: SOX2 20726797 ChIP-Seq SW620 Human | SOX2 |

| | | | |
|---|---|---|---|
| 1463 | ITPKA | Enrichr CHEA_2022: SOX2 20726797 ChIP-Seq SW620 Human | SOX2 |
| 1464 | GIPC2 | Enrichr CHEA_2022: SOX2 20726797 ChIP-Seq SW620 Human | SOX2 |
| 1465 | JDP2 | Enrichr CHEA_2022: SOX2 20726797 ChIP-Seq SW620 Human | SOX2 |
| 1466 | RNF141 | Enrichr CHEA_2022: SOX2 20726797 ChIP-Seq SW620 Human | SOX2 |
| 1467 | REN | Enrichr CHEA_2022: SOX2 20726797 ChIP-Seq SW620 Human | SOX2 |
| 1468 | ADA | Enrichr CHEA_2022: SOX2 20726797 ChIP-Seq SW620 Human | SOX2 |
| 1469 | SERPINA1 | Enrichr CHEA_2022: SOX2 20726797 ChIP-Seq SW620 Human | SOX2 |
| 1470 | CLTA | Enrichr CHEA_2022: SOX2 20726797 ChIP-Seq SW620 Human | SOX2 |
| 1471 | GLG1 | Enrichr CHEA_2022: SOX2 20726797 ChIP-Seq SW620 Human | SOX2 |
| 1472 | RNF157 | Enrichr CHEA_2022: SOX2 20726797 ChIP-Seq SW620 Human | SOX2 |
| 1473 | CSRP2 | Enrichr CHEA_2022: SOX2 20726797 ChIP-Seq SW620 Human | SOX2 |
| 1474 | TRIM4 | Enrichr CHEA_2022: SOX2 20726797 ChIP-Seq SW620 Human | SOX2 |
| 1475 | GPRASP2 | Enrichr CHEA_2022: SOX2 20726797 ChIP-Seq SW620 Human | SOX2 |
| 1476 | ZNF589 | Enrichr CHEA_2022: SOX2 20726797 ChIP-Seq SW620 Human | SOX2 |
| 1477 | TIGD2 | Enrichr CHEA_2022: SOX2 20726797 ChIP-Seq SW620 Human | SOX2 |
| 1478 | ZNF587 | Enrichr CHEA_2022: SOX2 20726797 ChIP-Seq SW620 Human | SOX2 |
| 1479 | ZNF586 | Enrichr CHEA_2022: SOX2 20726797 ChIP-Seq SW620 Human | SOX2 |
| 1480 | WWOX | Enrichr CHEA_2022: SOX2 20726797 ChIP-Seq SW620 Human | SOX2 |
| 1481 | SERPINB5 | Enrichr CHEA_2022: SOX2 20726797 ChIP-Seq SW620 Human | SOX2 |
| 1482 | SF3A3 | Enrichr CHEA_2022: SOX2 20726797 ChIP-Seq SW620 Human | SOX2 |
| 1483 | SERPINB3 | Enrichr CHEA_2022: SOX2 20726797 ChIP-Seq SW620 Human | SOX2 |
| 1484 | IFNGR1 | Enrichr CHEA_2022: SOX2 20726797 ChIP-Seq SW620 Human | SOX2 |
| 1485 | TMOD3 | Enrichr CHEA_2022: SOX2 20726797 ChIP-Seq SW620 Human | SOX2 |
| 1486 | WWP1 | Enrichr CHEA_2022: SOX2 20726797 ChIP-Seq SW620 Human | SOX2 |
| 1487 | LAPTM5 | Enrichr CHEA_2022: SOX2 20726797 ChIP-Seq SW620 Human | SOX2 |
| 1488 | EN2 | Enrichr CHEA_2022: SOX2 20726797 ChIP-Seq SW620 Human | SOX2 |
| 1489 | ADAT1 | Enrichr CHEA_2022: SOX2 20726797 ChIP-Seq SW620 Human | SOX2 |
| 1490 | GTF2H2 | Enrichr CHEA_2022: SOX2 20726797 ChIP-Seq SW620 Human | SOX2 |
| 1491 | NMNAT3 | Enrichr CHEA_2022: SOX2 20726797 ChIP-Seq SW620 Human | SOX2 |
| 1492 | TTC7B | Enrichr CHEA_2022: SOX2 20726797 ChIP-Seq SW620 Human | SOX2 |
| 1493 | SERPINB8 | Enrichr CHEA_2022: SOX2 20726797 ChIP-Seq SW620 Human | SOX2 |
| 1494 | SULF2 | Enrichr CHEA_2022: SOX2 20726797 ChIP-Seq SW620 Human | SOX2 |
| 1495 | TMEM154 | Enrichr CHEA_2022: SOX2 20726797 ChIP-Seq SW620 Human | SOX2 |
| 1496 | COL4A2 | Enrichr CHEA_2022: SOX2 20726797 ChIP-Seq SW620 Human | SOX2 |
| 1497 | RFWD3 | Enrichr CHEA_2022: SOX2 20726797 ChIP-Seq SW620 Human | SOX2 |
| 1498 | MSRB3 | Enrichr CHEA_2022: SOX2 20726797 ChIP-Seq SW620 Human | SOX2 |
| 1499 | NOTUM | Enrichr CHEA_2022: SOX2 20726797 ChIP-Seq SW620 Human | SOX2 |
| 1500 | HBEGF | Enrichr CHEA_2022: SOX2 20726797 ChIP-Seq SW620 Human | SOX2 |
| 1501 | BSCL2 | Enrichr CHEA_2022: SOX2 20726797 ChIP-Seq SW620 Human | SOX2 |
| 1502 | ZNF573 | Enrichr CHEA_2022: SOX2 20726797 ChIP-Seq SW620 Human | SOX2 |
| 1503 | CCL14 | Enrichr CHEA_2022: SOX2 20726797 ChIP-Seq SW620 Human | SOX2 |
| 1504 | PFAS | Enrichr CHEA_2022: SOX2 20726797 ChIP-Seq SW620 Human | SOX2 |
| 1505 | ZNF572 | Enrichr CHEA_2022: SOX2 20726797 ChIP-Seq SW620 Human | SOX2 |
| 1506 | OGG1 | Enrichr CHEA_2022: SOX2 20726797 ChIP-Seq SW620 Human | SOX2 |
| 1507 | TGFB1I1 | Enrichr CHEA_2022: SOX2 20726797 ChIP-Seq SW620 Human | SOX2 |
| 1508 | TMEM141 | Enrichr CHEA_2022: SOX2 20726797 ChIP-Seq SW620 Human | SOX2 |
| 1509 | PRSS23 | Enrichr CHEA_2022: SOX2 20726797 ChIP-Seq SW620 Human | SOX2 |
| 1510 | RNF135 | Enrichr CHEA_2022: SOX2 20726797 ChIP-Seq SW620 Human | SOX2 |
| 1511 | FAM102B | Enrichr CHEA_2022: SOX2 20726797 ChIP-Seq SW620 Human | SOX2 |
| 1512 | RNF139 | Enrichr CHEA_2022: SOX2 20726797 ChIP-Seq SW620 Human | SOX2 |
| 1513 | CHN2 | Enrichr CHEA_2022: SOX2 20726797 ChIP-Seq SW620 Human | SOX2 |
| 1514 | METTL7A | Enrichr CHEA_2022: SOX2 20726797 ChIP-Seq SW620 Human | SOX2 |
| 1515 | CHN1 | Enrichr CHEA_2022: SOX2 20726797 ChIP-Seq SW620 Human | SOX2 |
| 1516 | GCNT3 | Enrichr CHEA_2022: SOX2 20726797 ChIP-Seq SW620 Human | SOX2 |
| 1517 | ANXA9 | Enrichr CHEA_2022: SOX2 20726797 ChIP-Seq SW620 Human | SOX2 |
| 1518 | RIOK2 | Enrichr CHEA_2022: SOX2 20726797 ChIP-Seq SW620 Human | SOX2 |
| 1519 | ANKRD50 | Enrichr CHEA_2022: SOX2 20726797 ChIP-Seq SW620 Human | SOX2 |
| 1520 | ZNF567 | Enrichr CHEA_2022: SOX2 20726797 ChIP-Seq SW620 Human | SOX2 |
| 1521 | BRD7 | Enrichr CHEA_2022: SOX2 20726797 ChIP-Seq SW620 Human | SOX2 |
| 1522 | TNKS1BP1 | Enrichr CHEA_2022: SOX2 20726797 ChIP-Seq SW620 Human | SOX2 |
| 1523 | ZNF565 | Enrichr CHEA_2022: SOX2 20726797 ChIP-Seq SW620 Human | SOX2 |

| | | | |
|---|---|---|---|
| 1524 | RNF43 | Enrichr CHEA_2022: SOX2 20726797 ChIP-Seq SW620 Human | SOX2 |
| 1525 | SEMA4B | Enrichr CHEA_2022: SOX2 20726797 ChIP-Seq SW620 Human | SOX2 |
| 1526 | PHLDA1 | Enrichr CHEA_2022: SOX2 20726797 ChIP-Seq SW620 Human | SOX2 |
| 1527 | BRF2 | Enrichr CHEA_2022: SOX2 20726797 ChIP-Seq SW620 Human | SOX2 |
| 1528 | SEMA4F | Enrichr CHEA_2022: SOX2 20726797 ChIP-Seq SW620 Human | SOX2 |
| 1529 | SEMA4C | Enrichr CHEA_2022: SOX2 20726797 ChIP-Seq SW620 Human | SOX2 |
| 1530 | PLCL2 | Enrichr CHEA_2022: SOX2 20726797 ChIP-Seq SW620 Human | SOX2 |
| 1531 | SEMA4D | Enrichr CHEA_2022: SOX2 20726797 ChIP-Seq SW620 Human | SOX2 |
| 1532 | GPT2 | Enrichr CHEA_2022: SOX2 20726797 ChIP-Seq SW620 Human | SOX2 |
| 1533 | FAM24B | Enrichr CHEA_2022: SOX2 20726797 ChIP-Seq SW620 Human | SOX2 |
| 1534 | CYBRD1 | Enrichr CHEA_2022: SOX2 20726797 ChIP-Seq SW620 Human | SOX2 |
| 1535 | SEMA4G | Enrichr CHEA_2022: SOX2 20726797 ChIP-Seq SW620 Human | SOX2 |
| 1536 | CNOT7 | Enrichr CHEA_2022: SOX2 20726797 ChIP-Seq SW620 Human | SOX2 |
| 1537 | SYPL1 | Enrichr CHEA_2022: SOX2 20726797 ChIP-Seq SW620 Human | SOX2 |
| 1538 | RNF183 | Enrichr CHEA_2022: SOX2 20726797 ChIP-Seq SW620 Human | SOX2 |
| 1539 | TUSC1 | Enrichr CHEA_2022: SOX2 20726797 ChIP-Seq SW620 Human | SOX2 |
| 1540 | ANKRD22 | Enrichr CHEA_2022: SOX2 20726797 ChIP-Seq SW620 Human | SOX2 |
| 1541 | ZNF559 | Enrichr CHEA_2022: SOX2 20726797 ChIP-Seq SW620 Human | SOX2 |
| 1542 | TNIK | Enrichr CHEA_2022: SOX2 20726797 ChIP-Seq SW620 Human | SOX2 |
| 1543 | SLC28A3 | Enrichr CHEA_2022: SOX2 20726797 ChIP-Seq SW620 Human | SOX2 |
| 1544 | RNF32 | Enrichr CHEA_2022: SOX2 20726797 ChIP-Seq SW620 Human | SOX2 |
| 1545 | DGKH | Enrichr CHEA_2022: SOX2 20726797 ChIP-Seq SW620 Human | SOX2 |
| 1546 | ZNF552 | Enrichr CHEA_2022: SOX2 20726797 ChIP-Seq SW620 Human | SOX2 |
| 1547 | ZNF551 | Enrichr CHEA_2022: SOX2 20726797 ChIP-Seq SW620 Human | SOX2 |
| 1548 | COL16A1 | Enrichr CHEA_2022: SOX2 20726797 ChIP-Seq SW620 Human | SOX2 |
| 1549 | PHLDB2 | Enrichr CHEA_2022: SOX2 20726797 ChIP-Seq SW620 Human | SOX2 |
| 1550 | ZNF550 | Enrichr CHEA_2022: SOX2 20726797 ChIP-Seq SW620 Human | SOX2 |
| 1551 | ANKRD37 | Enrichr CHEA_2022: SOX2 20726797 ChIP-Seq SW620 Human | SOX2 |
| 1552 | AHR | Enrichr CHEA_2022: SOX2 20726797 ChIP-Seq SW620 Human | SOX2 |
| 1553 | METRNL | Enrichr CHEA_2022: SOX2 20726797 ChIP-Seq SW620 Human | SOX2 |
| 1554 | TANK | Enrichr CHEA_2022: SOX2 20726797 ChIP-Seq SW620 Human | SOX2 |
| 1555 | PYCARD | Enrichr CHEA_2022: SOX2 20726797 ChIP-Seq SW620 Human | SOX2 |
| 1556 | AP3M2 | Enrichr CHEA_2022: SOX2 20726797 ChIP-Seq SW620 Human | SOX2 |
| 1557 | ZFP36 | Enrichr CHEA_2022: SOX2 20726797 ChIP-Seq SW620 Human | SOX2 |
| 1558 | TUBB6 | Enrichr CHEA_2022: SOX2 20726797 ChIP-Seq SW620 Human | SOX2 |
| 1559 | ZFP30 | Enrichr CHEA_2022: SOX2 20726797 ChIP-Seq SW620 Human | SOX2 |
| 1560 | GRB14 | Enrichr CHEA_2022: SOX2 20726797 ChIP-Seq SW620 Human | SOX2 |
| 1561 | ZNF548 | Enrichr CHEA_2022: SOX2 20726797 ChIP-Seq SW620 Human | SOX2 |
| 1562 | ACSS1 | Enrichr CHEA_2022: SOX2 20726797 ChIP-Seq SW620 Human | SOX2 |
| 1563 | IGFBP2 | Enrichr CHEA_2022: SOX2 20726797 ChIP-Seq SW620 Human | SOX2 |
| 1564 | BIVM | Enrichr CHEA_2022: SOX2 20726797 ChIP-Seq SW620 Human | SOX2 |
| 1565 | F2R | Enrichr CHEA_2022: SOX2 20726797 ChIP-Seq SW620 Human | SOX2 |
| 1566 | RNF170 | Enrichr CHEA_2022: SOX2 20726797 ChIP-Seq SW620 Human | SOX2 |
| 1567 | ADRA2A | Enrichr CHEA_2022: SOX2 20726797 ChIP-Seq SW620 Human | SOX2 |
| 1568 | DUSP23 | Enrichr CHEA_2022: SOX2 20726797 ChIP-Seq SW620 Human | SOX2 |
| 1569 | ARMCX3 | Enrichr CHEA_2022: SOX2 20726797 ChIP-Seq SW620 Human | SOX2 |
| 1570 | TRAF5 | Enrichr CHEA_2022: SOX2 20726797 ChIP-Seq SW620 Human | SOX2 |
| 1571 | RHEB | Enrichr CHEA_2022: SOX2 20726797 ChIP-Seq SW620 Human | SOX2 |
| 1572 | TRAF4 | Enrichr CHEA_2022: SOX2 20726797 ChIP-Seq SW620 Human | SOX2 |
| 1573 | ARMCX6 | Enrichr CHEA_2022: SOX2 20726797 ChIP-Seq SW620 Human | SOX2 |
| 1574 | DGKQ | Enrichr CHEA_2022: SOX2 20726797 ChIP-Seq SW620 Human | SOX2 |
| 1575 | ILDR1 | Enrichr CHEA_2022: SOX2 20726797 ChIP-Seq SW620 Human | SOX2 |
| 1576 | ERF | Enrichr CHEA_2022: SOX2 20726797 ChIP-Seq SW620 Human | SOX2 |
| 1577 | ESAM | Enrichr CHEA_2022: SOX2 20726797 ChIP-Seq SW620 Human | SOX2 |
| 1578 | RLN2 | Enrichr CHEA_2022: SOX2 20726797 ChIP-Seq SW620 Human | SOX2 |
| 1579 | ZNF530 | Enrichr CHEA_2022: SOX2 20726797 ChIP-Seq SW620 Human | SOX2 |
| 1580 | IRS1 | Enrichr CHEA_2022: SOX2 20726797 ChIP-Seq SW620 Human | SOX2 |
| 1581 | FMR1 | Enrichr CHEA_2022: SOX2 20726797 ChIP-Seq SW620 Human | SOX2 |
| 1582 | BRI3 | Enrichr CHEA_2022: SOX2 20726797 ChIP-Seq SW620 Human | SOX2 |
| 1583 | AK5 | Enrichr CHEA_2022: SOX2 20726797 ChIP-Seq SW620 Human | SOX2 |
| 1584 | MYOM3 | Enrichr CHEA_2022: SOX2 20726797 ChIP-Seq SW620 Human | SOX2 |

| | | | |
|---|---|---|---|
| 1585 | ABHD12 | Enrichr CHEA_2022: SOX2 20726797 ChIP-Seq SW620 Human | SOX2 |
| 1586 | BAG4 | Enrichr CHEA_2022: SOX2 20726797 ChIP-Seq SW620 Human | SOX2 |
| 1587 | ABHD10 | Enrichr CHEA_2022: SOX2 20726797 ChIP-Seq SW620 Human | SOX2 |
| 1588 | ZNF529 | Enrichr CHEA_2022: SOX2 20726797 ChIP-Seq SW620 Human | SOX2 |
| 1589 | ANKRD10 | Enrichr CHEA_2022: SOX2 20726797 ChIP-Seq SW620 Human | SOX2 |
| 1590 | SLC37A1 | Enrichr CHEA_2022: SOX2 20726797 ChIP-Seq SW620 Human | SOX2 |
| 1591 | GPX3 | Enrichr CHEA_2022: SOX2 20726797 ChIP-Seq SW620 Human | SOX2 |
| 1592 | GPX2 | Enrichr CHEA_2022: SOX2 20726797 ChIP-Seq SW620 Human | SOX2 |
| 1593 | DTX3L | Enrichr CHEA_2022: SOX2 20726797 ChIP-Seq SW620 Human | SOX2 |
| 1594 | GPX7 | Enrichr CHEA_2022: SOX2 20726797 ChIP-Seq SW620 Human | SOX2 |
| 1595 | SMARCE1 | Enrichr CHEA_2022: SOX2 20726797 ChIP-Seq SW620 Human | SOX2 |
| 1596 | CFLAR | Enrichr CHEA_2022: SOX2 20726797 ChIP-Seq SW620 Human | SOX2 |
| 1597 | ASNS | Enrichr CHEA_2022: SOX2 20726797 ChIP-Seq SW620 Human | SOX2 |
| 1598 | PTPN13 | Enrichr CHEA_2022: SOX2 20726797 ChIP-Seq SW620 Human | SOX2 |
| 1599 | CDC42BPA | Enrichr CHEA_2022: SOX2 20726797 ChIP-Seq SW620 Human | SOX2 |
| 1600 | TPCN2 | Enrichr CHEA_2022: SOX2 20726797 ChIP-Seq SW620 Human | SOX2 |
| 1601 | CUL4A | Enrichr CHEA_2022: SOX2 20726797 ChIP-Seq SW620 Human | SOX2 |
| 1602 | SERPINI1 | Enrichr CHEA_2022: SOX2 20726797 ChIP-Seq SW620 Human | SOX2 |
| 1603 | NES | Enrichr CHEA_2022: SOX2 20726797 ChIP-Seq SW620 Human | SOX2 |
| 1604 | SERINC2 | Enrichr CHEA_2022: SOX2 20726797 ChIP-Seq SW620 Human | SOX2 |
| 1605 | ZNF512 | Enrichr CHEA_2022: SOX2 20726797 ChIP-Seq SW620 Human | SOX2 |
| 1606 | EIF4E2 | Enrichr CHEA_2022: SOX2 20726797 ChIP-Seq SW620 Human | SOX2 |
| 1607 | METAP1 | Enrichr CHEA_2022: SOX2 20726797 ChIP-Seq SW620 Human | SOX2 |
| 1608 | METAP2 | Enrichr CHEA_2022: SOX2 20726797 ChIP-Seq SW620 Human | SOX2 |
| 1609 | SERINC5 | Enrichr CHEA_2022: SOX2 20726797 ChIP-Seq SW620 Human | SOX2 |
| 1610 | TNKS | Enrichr CHEA_2022: SOX2 20726797 ChIP-Seq SW620 Human | SOX2 |
| 1611 | ITPR2 | Enrichr CHEA_2022: SOX2 20726797 ChIP-Seq SW620 Human | SOX2 |
| 1612 | ITPR3 | Enrichr CHEA_2022: SOX2 20726797 ChIP-Seq SW620 Human | SOX2 |
| 1613 | DUSP14 | Enrichr CHEA_2022: SOX2 20726797 ChIP-Seq SW620 Human | SOX2 |
| 1614 | RAD21 | Enrichr CHEA_2022: SOX2 20726797 ChIP-Seq SW620 Human | SOX2 |
| 1615 | PSAP | Enrichr CHEA_2022: SOX2 20726797 ChIP-Seq SW620 Human | SOX2 |
| 1616 | WASF1 | Enrichr CHEA_2022: SOX2 20726797 ChIP-Seq SW620 Human | SOX2 |
| 1617 | SKIL | Enrichr CHEA_2022: SOX2 20726797 ChIP-Seq SW620 Human | SOX2 |
| 1618 | WASF3 | Enrichr CHEA_2022: SOX2 20726797 ChIP-Seq SW620 Human | SOX2 |
| 1619 | IRX2 | Enrichr CHEA_2022: SOX2 20726797 ChIP-Seq SW620 Human | SOX2 |
| 1620 | SPAG7 | Enrichr CHEA_2022: SOX2 20726797 ChIP-Seq SW620 Human | SOX2 |
| 1621 | SPAG4 | Enrichr CHEA_2022: SOX2 20726797 ChIP-Seq SW620 Human | SOX2 |
| 1622 | AMT | Enrichr CHEA_2022: SOX2 20726797 ChIP-Seq SW620 Human | SOX2 |
| 1623 | MND1 | Enrichr CHEA_2022: SOX2 20726797 ChIP-Seq SW620 Human | SOX2 |
| 1624 | RAD50 | Enrichr CHEA_2022: SOX2 20726797 ChIP-Seq SW620 Human | SOX2 |
| 1625 | RNF20 | Enrichr CHEA_2022: SOX2 20726797 ChIP-Seq SW620 Human | SOX2 |
| 1626 | KLF2 | Enrichr CHEA_2022: SOX2 20726797 ChIP-Seq SW620 Human | SOX2 |
| 1627 | ZFP90 | Enrichr CHEA_2022: SOX2 20726797 ChIP-Seq SW620 Human | SOX2 |
| 1628 | RAD52 | Enrichr CHEA_2022: SOX2 20726797 ChIP-Seq SW620 Human | SOX2 |
| 1629 | MGAT4B | Enrichr CHEA_2022: SOX2 20726797 ChIP-Seq SW620 Human | SOX2 |
| 1630 | ASPH | Enrichr CHEA_2022: SOX2 20726797 ChIP-Seq SW620 Human | SOX2 |
| 1631 | CD109 | Enrichr CHEA_2022: SOX2 20726797 ChIP-Seq SW620 Human | SOX2 |
| 1632 | EVL | Enrichr CHEA_2022: SOX2 20726797 ChIP-Seq SW620 Human | SOX2 |
| 1633 | CLEC2D | Enrichr CHEA_2022: SOX2 20726797 ChIP-Seq SW620 Human | SOX2 |
| 1634 | SMPDL3B | Enrichr CHEA_2022: SOX2 20726797 ChIP-Seq SW620 Human | SOX2 |
| 1635 | FOLR1 | Enrichr CHEA_2022: SOX2 20726797 ChIP-Seq SW620 Human | SOX2 |
| 1636 | SEMA3A | Enrichr CHEA_2022: SOX2 20726797 ChIP-Seq SW620 Human | SOX2 |
| 1637 | LSP1 | Enrichr CHEA_2022: SOX2 20726797 ChIP-Seq SW620 Human | SOX2 |
| 1638 | SEMA3C | Enrichr CHEA_2022: SOX2 20726797 ChIP-Seq SW620 Human | SOX2 |
| 1639 | SEMA3F | Enrichr CHEA_2022: SOX2 20726797 ChIP-Seq SW620 Human | SOX2 |
| 1640 | SGCE | Enrichr CHEA_2022: SOX2 20726797 ChIP-Seq SW620 Human | SOX2 |
| 1641 | OPA1 | Enrichr CHEA_2022: SOX2 20726797 ChIP-Seq SW620 Human | SOX2 |
| 1642 | OPA3 | Enrichr CHEA_2022: SOX2 20726797 ChIP-Seq SW620 Human | SOX2 |
| 1643 | NHS | Enrichr CHEA_2022: SOX2 20726797 ChIP-Seq SW620 Human | SOX2 |
| 1644 | PTPRN2 | Enrichr CHEA_2022: SOX2 20726797 ChIP-Seq SW620 Human | SOX2 |
| 1645 | CDON | Enrichr CHEA_2022: SOX2 20726797 ChIP-Seq SW620 Human | SOX2 |

| | | | |
|---|---|---|---|
| 1646 | SS18L1 | Enrichr CHEA_2022: SOX2 20726797 ChIP-Seq SW620 Human | SOX2 |
| 1647 | SP110 | Enrichr CHEA_2022: SOX2 20726797 ChIP-Seq SW620 Human | SOX2 |
| 1648 | ACSL1 | Enrichr CHEA_2022: SOX2 20726797 ChIP-Seq SW620 Human | SOX2 |
| 1649 | RUNX1 | Enrichr CHEA_2022: SOX2 20726797 ChIP-Seq SW620 Human | SOX2 |
| 1650 | RECQL | Enrichr CHEA_2022: SOX2 20726797 ChIP-Seq SW620 Human | SOX2 |
| 1651 | ACSL5 | Enrichr CHEA_2022: SOX2 20726797 ChIP-Seq SW620 Human | SOX2 |
| 1652 | RUNX3 | Enrichr CHEA_2022: SOX2 20726797 ChIP-Seq SW620 Human | SOX2 |
| 1653 | MTUS1 | Enrichr CHEA_2022: SOX2 20726797 ChIP-Seq SW620 Human | SOX2 |
| 1654 | RUNX2 | Enrichr CHEA_2022: SOX2 20726797 ChIP-Seq SW620 Human | SOX2 |
| 1655 | MALL | Enrichr CHEA_2022: SOX2 20726797 ChIP-Seq SW620 Human | SOX2 |
| 1656 | RHEBL1 | Enrichr CHEA_2022: SOX2 20726797 ChIP-Seq SW620 Human | SOX2 |
| 1657 | XRN2 | Enrichr CHEA_2022: SOX2 20726797 ChIP-Seq SW620 Human | SOX2 |
| 1658 | RRAGD | Enrichr CHEA_2022: SOX2 20726797 ChIP-Seq SW620 Human | SOX2 |
| 1659 | TMCC1 | Enrichr CHEA_2022: SOX2 20726797 ChIP-Seq SW620 Human | SOX2 |
| 1660 | GUF1 | Enrichr CHEA_2022: SOX2 20726797 ChIP-Seq SW620 Human | SOX2 |
| 1661 | TFAM | Enrichr CHEA_2022: SOX2 20726797 ChIP-Seq SW620 Human | SOX2 |
| 1662 | PFN2 | Enrichr CHEA_2022: SOX2 20726797 ChIP-Seq SW620 Human | SOX2 |
| 1663 | SPRR3 | Enrichr CHEA_2022: SOX2 20726797 ChIP-Seq SW620 Human | SOX2 |
| 1664 | GLS2 | Enrichr CHEA_2022: SOX2 20726797 ChIP-Seq SW620 Human | SOX2 |
| 1665 | CMAS | Enrichr CHEA_2022: SOX2 20726797 ChIP-Seq SW620 Human | SOX2 |
| 1666 | VIPR1 | Enrichr CHEA_2022: SOX2 20726797 ChIP-Seq SW620 Human | SOX2 |
| 1667 | GLRX | Enrichr CHEA_2022: SOX2 20726797 ChIP-Seq SW620 Human | SOX2 |
| 1668 | RAD54B | Enrichr CHEA_2022: SOX2 20726797 ChIP-Seq SW620 Human | SOX2 |
| 1669 | ANTXR2 | Enrichr CHEA_2022: SOX2 20726797 ChIP-Seq SW620 Human | SOX2 |
| 1670 | CARHSP1 | Enrichr CHEA_2022: SOX2 20726797 ChIP-Seq SW620 Human | SOX2 |
| 1671 | ANTXR1 | Enrichr CHEA_2022: SOX2 20726797 ChIP-Seq SW620 Human | SOX2 |
| 1672 | CHAC1 | Enrichr CHEA_2022: SOX2 20726797 ChIP-Seq SW620 Human | SOX2 |
| 1673 | MAL2 | Enrichr CHEA_2022: SOX2 20726797 ChIP-Seq SW620 Human | SOX2 |
| 1674 | RABGAP1L | Enrichr CHEA_2022: SOX2 20726797 ChIP-Seq SW620 Human | SOX2 |
| 1675 | AQR | Enrichr CHEA_2022: SOX2 20726797 ChIP-Seq SW620 Human | SOX2 |
| 1676 | LIX1L | Enrichr CHEA_2022: SOX2 20726797 ChIP-Seq SW620 Human | SOX2 |
| 1677 | GADD45B | Enrichr CHEA_2022: SOX2 20726797 ChIP-Seq SW620 Human | SOX2 |
| 1678 | GADD45A | Enrichr CHEA_2022: SOX2 20726797 ChIP-Seq SW620 Human | SOX2 |
| 1679 | SLC11A2 | Enrichr CHEA_2022: SOX2 20726797 ChIP-Seq SW620 Human | SOX2 |
| 1680 | ACADSB | Enrichr CHEA_2022: SOX2 20726797 ChIP-Seq SW620 Human | SOX2 |
| 1681 | OVOL1 | Enrichr CHEA_2022: SOX2 20726797 ChIP-Seq SW620 Human | SOX2 |
| 1682 | IL17RD | Enrichr CHEA_2022: SOX2 20726797 ChIP-Seq SW620 Human | SOX2 |
| 1683 | PARP14 | Enrichr CHEA_2022: SOX2 20726797 ChIP-Seq SW620 Human | SOX2 |
| 1684 | INPP4B | Enrichr CHEA_2022: SOX2 20726797 ChIP-Seq SW620 Human | SOX2 |
| 1685 | PARP12 | Enrichr CHEA_2022: SOX2 20726797 ChIP-Seq SW620 Human | SOX2 |
| 1686 | TSHZ1 | Enrichr CHEA_2022: SOX2 20726797 ChIP-Seq SW620 Human | SOX2 |
| 1687 | TRMT12 | Enrichr CHEA_2022: SOX2 20726797 ChIP-Seq SW620 Human | SOX2 |
| 1688 | ITM2C | Enrichr CHEA_2022: SOX2 20726797 ChIP-Seq SW620 Human | SOX2 |
| 1689 | CDS1 | Enrichr CHEA_2022: SOX2 20726797 ChIP-Seq SW620 Human | SOX2 |
| 1690 | BEX2 | Enrichr CHEA_2022: SOX2 20726797 ChIP-Seq SW620 Human | SOX2 |
| 1691 | ACSM3 | Enrichr CHEA_2022: SOX2 20726797 ChIP-Seq SW620 Human | SOX2 |
| 1692 | MGST2 | Enrichr CHEA_2022: SOX2 20726797 ChIP-Seq SW620 Human | SOX2 |
| 1693 | DIXDC1 | Enrichr CHEA_2022: SOX2 20726797 ChIP-Seq SW620 Human | SOX2 |
| 1694 | EPM2AIP1 | Enrichr CHEA_2022: SOX2 20726797 ChIP-Seq SW620 Human | SOX2 |
| 1695 | SOX2 | Enrichr CHEA_2022: SOX2 20726797 ChIP-Seq SW620 Human | SOX2 |
| 1696 | INPP5F | Enrichr CHEA_2022: SOX2 20726797 ChIP-Seq SW620 Human | SOX2 |
| 1697 | ASL | Enrichr CHEA_2022: SOX2 20726797 ChIP-Seq SW620 Human | SOX2 |
| 1698 | ARL6IP5 | Enrichr CHEA_2022: SOX2 20726797 ChIP-Seq SW620 Human | SOX2 |
| 1699 | TREX1 | Enrichr CHEA_2022: SOX2 20726797 ChIP-Seq SW620 Human | SOX2 |
| 1700 | GLUL | Enrichr CHEA_2022: SOX2 20726797 ChIP-Seq SW620 Human | SOX2 |
| 1701 | GOLGA7 | Enrichr CHEA_2022: SOX2 20726797 ChIP-Seq SW620 Human | SOX2 |
| 1702 | TNS4 | Enrichr CHEA_2022: SOX2 20726797 ChIP-Seq SW620 Human | SOX2 |
| 1703 | SOX8 | Enrichr CHEA_2022: SOX2 20726797 ChIP-Seq SW620 Human | SOX2 |
| 1704 | TNS3 | Enrichr CHEA_2022: SOX2 20726797 ChIP-Seq SW620 Human | SOX2 |
| 1705 | CDR2 | Enrichr CHEA_2022: SOX2 20726797 ChIP-Seq SW620 Human | SOX2 |
| 1706 | NMB | Enrichr CHEA_2022: SOX2 20726797 ChIP-Seq SW620 Human | SOX2 |

| | | | |
|---|---|---|---|
| 1707 | TSC22D2 | Enrichr CHEA_2022: SOX2 20726797 ChIP-Seq SW620 Human | SOX2 |
| 1708 | NMI | Enrichr CHEA_2022: SOX2 20726797 ChIP-Seq SW620 Human | SOX2 |
| 1709 | NME3 | Enrichr CHEA_2022: SOX2 20726797 ChIP-Seq SW620 Human | SOX2 |
| 1710 | TFEB | Enrichr CHEA_2022: SOX2 20726797 ChIP-Seq SW620 Human | SOX2 |
| 1711 | KRT13 | Enrichr CHEA_2022: SOX2 20726797 ChIP-Seq SW620 Human | SOX2 |
| 1712 | NME4 | Enrichr CHEA_2022: SOX2 20726797 ChIP-Seq SW620 Human | SOX2 |
| 1713 | TUBE1 | Enrichr CHEA_2022: SOX2 20726797 ChIP-Seq SW620 Human | SOX2 |
| 1714 | GNG12 | Enrichr CHEA_2022: SOX2 20726797 ChIP-Seq SW620 Human | SOX2 |
| 1715 | PDZK1IP1 | Enrichr CHEA_2022: SOX2 20726797 ChIP-Seq SW620 Human | SOX2 |
| 1716 | RAD51C | Enrichr CHEA_2022: SOX2 20726797 ChIP-Seq SW620 Human | SOX2 |
| 1717 | RHOQ | Enrichr CHEA_2022: SOX2 20726797 ChIP-Seq SW620 Human | SOX2 |
| 1718 | CEACAM1 | Enrichr CHEA_2022: SOX2 20726797 ChIP-Seq SW620 Human | SOX2 |
| 1719 | NME7 | Enrichr CHEA_2022: SOX2 20726797 ChIP-Seq SW620 Human | SOX2 |
| 1720 | R3HCC1 | Enrichr CHEA_2022: SOX2 20726797 ChIP-Seq SW620 Human | SOX2 |
| 1721 | KRT17 | Enrichr CHEA_2022: SOX2 20726797 ChIP-Seq SW620 Human | SOX2 |
| 1722 | RPUSD3 | Enrichr CHEA_2022: SOX2 20726797 ChIP-Seq SW620 Human | SOX2 |
| 1723 | CEACAM6 | Enrichr CHEA_2022: SOX2 20726797 ChIP-Seq SW620 Human | SOX2 |
| 1724 | RPUSD4 | Enrichr CHEA_2022: SOX2 20726797 ChIP-Seq SW620 Human | SOX2 |
| 1725 | ATM | Enrichr CHEA_2022: SOX2 20726797 ChIP-Seq SW620 Human | SOX2 |
| 1726 | TFF3 | Enrichr CHEA_2022: SOX2 20726797 ChIP-Seq SW620 Human | SOX2 |
| 1727 | CAMK1 | Enrichr CHEA_2022: SOX2 20726797 ChIP-Seq SW620 Human | SOX2 |
| 1728 | TFF1 | Enrichr CHEA_2022: SOX2 20726797 ChIP-Seq SW620 Human | SOX2 |
| 1729 | SGK2 | Enrichr CHEA_2022: SOX2 20726797 ChIP-Seq SW620 Human | SOX2 |
| 1730 | AKR1B10 | Enrichr CHEA_2022: SOX2 20726797 ChIP-Seq SW620 Human | SOX2 |
| 1731 | RHOU | Enrichr CHEA_2022: SOX2 20726797 ChIP-Seq SW620 Human | SOX2 |
| 1732 | TRAM2 | Enrichr CHEA_2022: SOX2 20726797 ChIP-Seq SW620 Human | SOX2 |
| 1733 | CLIC5 | Enrichr CHEA_2022: SOX2 20726797 ChIP-Seq SW620 Human | SOX2 |
| 1734 | CLIC4 | Enrichr CHEA_2022: SOX2 20726797 ChIP-Seq SW620 Human | SOX2 |
| 1735 | MAOB | Enrichr CHEA_2022: SOX2 20726797 ChIP-Seq SW620 Human | SOX2 |
| 1736 | MAOA | Enrichr CHEA_2022: SOX2 20726797 ChIP-Seq SW620 Human | SOX2 |
| 1737 | IGSF3 | Enrichr CHEA_2022: SOX2 20726797 ChIP-Seq SW620 Human | SOX2 |
| 1738 | KLK6 | Enrichr CHEA_2022: SOX2 20726797 ChIP-Seq SW620 Human | SOX2 |
| 1739 | STC2 | Enrichr CHEA_2022: SOX2 20726797 ChIP-Seq SW620 Human | SOX2 |
| 1740 | VLDLR | Enrichr CHEA_2022: SOX2 20726797 ChIP-Seq SW620 Human | SOX2 |
| 1741 | KRT20 | Enrichr CHEA_2022: SOX2 20726797 ChIP-Seq SW620 Human | SOX2 |
| 1742 | NNT | Enrichr CHEA_2022: SOX2 20726797 ChIP-Seq SW620 Human | SOX2 |
| 1743 | HINT3 | Enrichr CHEA_2022: SOX2 20726797 ChIP-Seq SW620 Human | SOX2 |
| 1744 | PITRM1 | Enrichr CHEA_2022: SOX2 20726797 ChIP-Seq SW620 Human | SOX2 |
| 1745 | CLIC3 | Enrichr CHEA_2022: SOX2 20726797 ChIP-Seq SW620 Human | SOX2 |
| 1746 | ARNT2 | Enrichr CHEA_2022: SOX2 20726797 ChIP-Seq SW620 Human | SOX2 |
| 1747 | PRMT7 | Enrichr CHEA_2022: SOX2 20726797 ChIP-Seq SW620 Human | SOX2 |
| 1748 | PRMT5 | Enrichr CHEA_2022: SOX2 20726797 ChIP-Seq SW620 Human | SOX2 |
| 1749 | SCD5 | Enrichr CHEA_2022: SOX2 20726797 ChIP-Seq SW620 Human | SOX2 |
| 1750 | RSL1D1 | Enrichr CHEA_2022: SOX2 20726797 ChIP-Seq SW620 Human | SOX2 |
| 1751 | TJP2 | Enrichr CHEA_2022: SOX2 20726797 ChIP-Seq SW620 Human | SOX2 |
| 1752 | TJP1 | Enrichr CHEA_2022: SOX2 20726797 ChIP-Seq SW620 Human | SOX2 |
| 1753 | DNASE1L1 | Enrichr CHEA_2022: SOX2 20726797 ChIP-Seq SW620 Human | SOX2 |
| 1754 | TJP3 | Enrichr CHEA_2022: SOX2 20726797 ChIP-Seq SW620 Human | SOX2 |
| 1755 | DCP2 | Enrichr CHEA_2022: SOX2 20726797 ChIP-Seq SW620 Human | SOX2 |
| 1756 | GDI1 | Enrichr CHEA_2022: SOX2 20726797 ChIP-Seq SW620 Human | SOX2 |
| 1757 | PRIM1 | Enrichr CHEA_2022: SOX2 20726797 ChIP-Seq SW620 Human | SOX2 |
| 1758 | HCCS | Enrichr CHEA_2022: SOX2 20726797 ChIP-Seq SW620 Human | SOX2 |
| 1759 | TRAK2 | Enrichr CHEA_2022: SOX2 20726797 ChIP-Seq SW620 Human | SOX2 |
| 1760 | TRAK1 | Enrichr CHEA_2022: SOX2 20726797 ChIP-Seq SW620 Human | SOX2 |
| 1761 | GYS1 | Enrichr CHEA_2022: SOX2 20726797 ChIP-Seq SW620 Human | SOX2 |
| 1762 | NPW | Enrichr CHEA_2022: SOX2 20726797 ChIP-Seq SW620 Human | SOX2 |
| 1763 | MAP4K4 | Enrichr CHEA_2022: SOX2 20726797 ChIP-Seq SW620 Human | SOX2 |
| 1764 | GDF11 | Enrichr CHEA_2022: SOX2 20726797 ChIP-Seq SW620 Human | SOX2 |
| 1765 | CA13 | Enrichr CHEA_2022: SOX2 20726797 ChIP-Seq SW620 Human | SOX2 |
| 1766 | BTNL9 | Enrichr CHEA_2022: SOX2 20726797 ChIP-Seq SW620 Human | SOX2 |
| 1767 | CA12 | Enrichr CHEA_2022: SOX2 20726797 ChIP-Seq SW620 Human | SOX2 |

| | | | |
|---|---|---|---|
| 1768 | TGFB2 | Enrichr CHEA_2022: SOX2 20726797 ChIP-Seq SW620 Human | SOX2 |
| 1769 | KLRC4 | Enrichr CHEA_2022: SOX2 20726797 ChIP-Seq SW620 Human | SOX2 |
| 1770 | EPDR1 | Enrichr CHEA_2022: SOX2 20726797 ChIP-Seq SW620 Human | SOX2 |
| 1771 | ARID3A | Enrichr CHEA_2022: SOX2 20726797 ChIP-Seq SW620 Human | SOX2 |
| 1772 | ARHGAP27 | Enrichr CHEA_2022: SOX2 20726797 ChIP-Seq SW620 Human | SOX2 |
| 1773 | BATF | Enrichr CHEA_2022: SOX2 20726797 ChIP-Seq SW620 Human | SOX2 |
| 1774 | ZNFX1 | Enrichr CHEA_2022: SOX2 20726797 ChIP-Seq SW620 Human | SOX2 |
| 1775 | GNPDA2 | Enrichr CHEA_2022: SOX2 20726797 ChIP-Seq SW620 Human | SOX2 |
| 1776 | AXL | Enrichr CHEA_2022: SOX2 20726797 ChIP-Seq SW620 Human | SOX2 |
| 1777 | TMEM139 | Enrichr CHEA_2022: SOX2 20726797 ChIP-Seq SW620 Human | SOX2 |
| 1778 | FAS | Enrichr CHEA_2022: SOX2 20726797 ChIP-Seq SW620 Human | SOX2 |
| 1779 | ELMO3 | Enrichr CHEA_2022: SOX2 20726797 ChIP-Seq SW620 Human | SOX2 |
| 1780 | ABCG1 | Enrichr CHEA_2022: SOX2 20726797 ChIP-Seq SW620 Human | SOX2 |
| 1781 | GOLT1A | Enrichr CHEA_2022: SOX2 20726797 ChIP-Seq SW620 Human | SOX2 |
| 1782 | SCG2 | Enrichr CHEA_2022: SOX2 20726797 ChIP-Seq SW620 Human | SOX2 |
| 1783 | STEAP2 | Enrichr CHEA_2022: SOX2 20726797 ChIP-Seq SW620 Human | SOX2 |
| 1784 | ABCG2 | Enrichr CHEA_2022: SOX2 20726797 ChIP-Seq SW620 Human | SOX2 |
| 1785 | FAIM | Enrichr CHEA_2022: SOX2 20726797 ChIP-Seq SW620 Human | SOX2 |
| 1786 | STEAP1 | Enrichr CHEA_2022: SOX2 20726797 ChIP-Seq SW620 Human | SOX2 |
| 1787 | GALNT12 | Enrichr CHEA_2022: SOX2 20726797 ChIP-Seq SW620 Human | SOX2 |
| 1788 | GALNT14 | Enrichr CHEA_2022: SOX2 20726797 ChIP-Seq SW620 Human | SOX2 |
| 1789 | SLFN5 | Enrichr CHEA_2022: SOX2 20726797 ChIP-Seq SW620 Human | SOX2 |
| 1790 | HDLBP | Enrichr CHEA_2022: SOX2 20726797 ChIP-Seq SW620 Human | SOX2 |
| 1791 | IFI6 | Enrichr CHEA_2022: SOX2 20726797 ChIP-Seq SW620 Human | SOX2 |
| 1792 | MANEA | Enrichr CHEA_2022: SOX2 20726797 ChIP-Seq SW620 Human | SOX2 |
| 1793 | GALNT10 | Enrichr CHEA_2022: SOX2 20726797 ChIP-Seq SW620 Human | SOX2 |
| 1794 | ZFP36L2 | Enrichr CHEA_2022: SOX2 20726797 ChIP-Seq SW620 Human | SOX2 |
| 1795 | SCEL | Enrichr CHEA_2022: SOX2 20726797 ChIP-Seq SW620 Human | SOX2 |
| 1796 | KRT6A | Enrichr CHEA_2022: SOX2 20726797 ChIP-Seq SW620 Human | SOX2 |
| 1797 | TMEM125 | Enrichr CHEA_2022: SOX2 20726797 ChIP-Seq SW620 Human | SOX2 |
| 1798 | MYH14 | Enrichr CHEA_2022: SOX2 20726797 ChIP-Seq SW620 Human | SOX2 |
| 1799 | MYH10 | Enrichr CHEA_2022: SOX2 20726797 ChIP-Seq SW620 Human | SOX2 |
| 1800 | CLASP2 | Enrichr CHEA_2022: SOX2 20726797 ChIP-Seq SW620 Human | SOX2 |
| 1801 | OPN3 | Enrichr CHEA_2022: SOX2 20726797 ChIP-Seq SW620 Human | SOX2 |
| 1802 | USP47 | Enrichr CHEA_2022: SOX2 20726797 ChIP-Seq SW620 Human | SOX2 |
| 1803 | SMAD3 | Enrichr CHEA_2022: SOX2 20726797 ChIP-Seq SW620 Human | SOX2 |
| 1804 | PDIA3 | Enrichr CHEA_2022: SOX2 20726797 ChIP-Seq SW620 Human | SOX2 |
| 1805 | SMAD5 | Enrichr CHEA_2022: SOX2 20726797 ChIP-Seq SW620 Human | SOX2 |
| 1806 | ARID5B | Enrichr CHEA_2022: SOX2 20726797 ChIP-Seq SW620 Human | SOX2 |
| 1807 | DNAJC12 | Enrichr CHEA_2022: SOX2 20726797 ChIP-Seq SW620 Human | SOX2 |
| 1808 | ASPSCR1 | Enrichr CHEA_2022: SOX2 20726797 ChIP-Seq SW620 Human | SOX2 |
| 1809 | CTPS2 | Enrichr CHEA_2022: SOX2 20726797 ChIP-Seq SW620 Human | SOX2 |
| 1810 | CDYL | Enrichr CHEA_2022: SOX2 20726797 ChIP-Seq SW620 Human | SOX2 |
| 1811 | SMAD7 | Enrichr CHEA_2022: SOX2 20726797 ChIP-Seq SW620 Human | SOX2 |
| 1812 | UPK3B | Enrichr CHEA_2022: SOX2 20726797 ChIP-Seq SW620 Human | SOX2 |
| 1813 | SMAD6 | Enrichr CHEA_2022: SOX2 20726797 ChIP-Seq SW620 Human | SOX2 |
| 1814 | DNAJC18 | Enrichr CHEA_2022: SOX2 20726797 ChIP-Seq SW620 Human | SOX2 |
| 1815 | SDCBP | Enrichr CHEA_2022: SOX2 20726797 ChIP-Seq SW620 Human | SOX2 |
| 1816 | PDIA6 | Enrichr CHEA_2022: SOX2 20726797 ChIP-Seq SW620 Human | SOX2 |
| 1817 | ALDH3A2 | Enrichr CHEA_2022: SOX2 20726797 ChIP-Seq SW620 Human | SOX2 |
| 1818 | PREPL | Enrichr CHEA_2022: SOX2 20726797 ChIP-Seq SW620 Human | SOX2 |
| 1819 | DLG4 | Enrichr CHEA_2022: SOX2 20726797 ChIP-Seq SW620 Human | SOX2 |
| 1820 | CHMP7 | Enrichr CHEA_2022: SOX2 20726797 ChIP-Seq SW620 Human | SOX2 |
| 1821 | ATP6V1A | Enrichr CHEA_2022: SOX2 20726797 ChIP-Seq SW620 Human | SOX2 |
| 1822 | DOCK4 | Enrichr CHEA_2022: SOX2 20726797 ChIP-Seq SW620 Human | SOX2 |
| 1823 | USP54 | Enrichr CHEA_2022: SOX2 20726797 ChIP-Seq SW620 Human | SOX2 |
| 1824 | SH3KBP1 | Enrichr CHEA_2022: SOX2 20726797 ChIP-Seq SW620 Human | SOX2 |
| 1825 | SLC2A3 | Enrichr CHEA_2022: SOX2 20726797 ChIP-Seq SW620 Human | SOX2 |
| 1826 | ARHGAP18 | Enrichr CHEA_2022: SOX2 20726797 ChIP-Seq SW620 Human | SOX2 |
| 1827 | PLCXD2 | Enrichr CHEA_2022: SOX2 20726797 ChIP-Seq SW620 Human | SOX2 |
| 1828 | MYL6B | Enrichr CHEA_2022: SOX2 20726797 ChIP-Seq SW620 Human | SOX2 |

| | | | |
|---|---|---|---|
| 1829 | ARHGAP23 | Enrichr CHEA_2022: SOX2 20726797 ChIP-Seq SW620 Human | SOX2 |
| 1830 | PODXL | Enrichr CHEA_2022: SOX2 20726797 ChIP-Seq SW620 Human | SOX2 |
| 1831 | MAN2A2 | Enrichr CHEA_2022: SOX2 20726797 ChIP-Seq SW620 Human | SOX2 |
| 1832 | TMEM107 | Enrichr CHEA_2022: SOX2 20726797 ChIP-Seq SW620 Human | SOX2 |
| 1833 | ALDH3B1 | Enrichr CHEA_2022: SOX2 20726797 ChIP-Seq SW620 Human | SOX2 |
| 1834 | COBLL1 | Enrichr CHEA_2022: SOX2 20726797 ChIP-Seq SW620 Human | SOX2 |
| 1835 | PSPH | Enrichr CHEA_2022: SOX2 20726797 ChIP-Seq SW620 Human | SOX2 |
| 1836 | USP25 | Enrichr CHEA_2022: SOX2 20726797 ChIP-Seq SW620 Human | SOX2 |
| 1837 | ABCC5 | Enrichr CHEA_2022: SOX2 20726797 ChIP-Seq SW620 Human | SOX2 |
| 1838 | ABCC3 | Enrichr CHEA_2022: SOX2 20726797 ChIP-Seq SW620 Human | SOX2 |
| 1839 | HSPA5 | Enrichr CHEA_2022: SOX2 20726797 ChIP-Seq SW620 Human | SOX2 |
| 1840 | SMURF1 | Enrichr CHEA_2022: SOX2 20726797 ChIP-Seq SW620 Human | SOX2 |
| 1841 | HSPA2 | Enrichr CHEA_2022: SOX2 20726797 ChIP-Seq SW620 Human | SOX2 |
| 1842 | NFKBIZ | Enrichr CHEA_2022: SOX2 20726797 ChIP-Seq SW620 Human | SOX2 |
| 1843 | TGFBR3 | Enrichr CHEA_2022: SOX2 20726797 ChIP-Seq SW620 Human | SOX2 |
| 1844 | NAP1L5 | Enrichr CHEA_2022: SOX2 20726797 ChIP-Seq SW620 Human | SOX2 |
| 1845 | DDX19A | Enrichr CHEA_2022: SOX2 20726797 ChIP-Seq SW620 Human | SOX2 |
| 1846 | DDX19B | Enrichr CHEA_2022: SOX2 20726797 ChIP-Seq SW620 Human | SOX2 |
| 1847 | MEIS2 | Enrichr CHEA_2022: SOX2 20726797 ChIP-Seq SW620 Human | SOX2 |
| 1848 | MNS1 | Enrichr CHEA_2022: SOX2 20726797 ChIP-Seq SW620 Human | SOX2 |
| 1849 | PRTFDC1 | Enrichr CHEA_2022: SOX2 20726797 ChIP-Seq SW620 Human | SOX2 |
| 1850 | B4GALT1 | Enrichr CHEA_2022: SOX2 20726797 ChIP-Seq SW620 Human | SOX2 |
| 1851 | MCTP1 | Enrichr CHEA_2022: SOX2 20726797 ChIP-Seq SW620 Human | SOX2 |
| 1852 | ABCB1 | Enrichr CHEA_2022: SOX2 20726797 ChIP-Seq SW620 Human | SOX2 |
| 1853 | NID1 | Enrichr CHEA_2022: SOX2 20726797 ChIP-Seq SW620 Human | SOX2 |
| 1854 | MTIF3 | Enrichr CHEA_2022: SOX2 20726797 ChIP-Seq SW620 Human | SOX2 |
| 1855 | CXCL1 | Enrichr CHEA_2022: SOX2 20726797 ChIP-Seq SW620 Human | SOX2 |
| 1856 | DTX4 | Enrichr CHEA_2022: SOX2 20726797 ChIP-Seq SW620 Human | SOX2 |
| 1857 | ELAVL2 | Enrichr CHEA_2022: SOX2 20726797 ChIP-Seq SW620 Human | SOX2 |
| 1858 | CXCL3 | Enrichr CHEA_2022: SOX2 20726797 ChIP-Seq SW620 Human | SOX2 |
| 1859 | MOSPD1 | Enrichr CHEA_2022: SOX2 20726797 ChIP-Seq SW620 Human | SOX2 |
| 1860 | CXCL6 | Enrichr CHEA_2022: SOX2 20726797 ChIP-Seq SW620 Human | SOX2 |
| 1861 | DPP4 | Enrichr CHEA_2022: SOX2 20726797 ChIP-Seq SW620 Human | SOX2 |
| 1862 | DPP7 | Enrichr CHEA_2022: SOX2 20726797 ChIP-Seq SW620 Human | SOX2 |
| 1863 | NRIP1 | Enrichr CHEA_2022: SOX2 20726797 ChIP-Seq SW620 Human | SOX2 |
| 1864 | NRIP3 | Enrichr CHEA_2022: SOX2 20726797 ChIP-Seq SW620 Human | SOX2 |
| 1865 | ERBB2 | Enrichr CHEA_2022: SOX2 20726797 ChIP-Seq SW620 Human | SOX2 |
| 1866 | PDE4A | Enrichr CHEA_2022: SOX2 20726797 ChIP-Seq SW620 Human | SOX2 |
| 1867 | USP31 | Enrichr CHEA_2022: SOX2 20726797 ChIP-Seq SW620 Human | SOX2 |
| 1868 | TCFL5 | Enrichr CHEA_2022: SOX2 20726797 ChIP-Seq SW620 Human | SOX2 |
| 1869 | ZNF185 | Enrichr CHEA_2022: SOX2 20726797 ChIP-Seq SW620 Human | SOX2 |
| 1870 | PDE4DIP | Enrichr CHEA_2022: SOX2 20726797 ChIP-Seq SW620 Human | SOX2 |
| 1871 | RWDD1 | Enrichr CHEA_2022: SOX2 20726797 ChIP-Seq SW620 Human | SOX2 |
| 1872 | PDE4D | Enrichr CHEA_2022: SOX2 20726797 ChIP-Seq SW620 Human | SOX2 |
| 1873 | ZNF182 | Enrichr CHEA_2022: SOX2 20726797 ChIP-Seq SW620 Human | SOX2 |
| 1874 | KCTD1 | Enrichr CHEA_2022: SOX2 20726797 ChIP-Seq SW620 Human | SOX2 |
| 1875 | KCTD6 | Enrichr CHEA_2022: SOX2 20726797 ChIP-Seq SW620 Human | SOX2 |
| 1876 | SGPP2 | Enrichr CHEA_2022: SOX2 20726797 ChIP-Seq SW620 Human | SOX2 |
| 1877 | CLDN7 | Enrichr CHEA_2022: SOX2 20726797 ChIP-Seq SW620 Human | SOX2 |
| 1878 | FMNL2 | Enrichr CHEA_2022: SOX2 20726797 ChIP-Seq SW620 Human | SOX2 |
| 1879 | CLDN4 | Enrichr CHEA_2022: SOX2 20726797 ChIP-Seq SW620 Human | SOX2 |
| 1880 | PALLD | Enrichr CHEA_2022: SOX2 20726797 ChIP-Seq SW620 Human | SOX2 |
| 1881 | COL8A1 | Enrichr CHEA_2022: SOX2 20726797 ChIP-Seq SW620 Human | SOX2 |
| 1882 | HKDC1 | Enrichr CHEA_2022: SOX2 20726797 ChIP-Seq SW620 Human | SOX2 |
| 1883 | VDAC3 | Enrichr CHEA_2022: SOX2 20726797 ChIP-Seq SW620 Human | SOX2 |
| 1884 | SPRY2 | Enrichr CHEA_2022: SOX2 20726797 ChIP-Seq SW620 Human | SOX2 |
| 1885 | SSBP2 | Enrichr CHEA_2022: SOX2 20726797 ChIP-Seq SW620 Human | SOX2 |
| 1886 | B4GALT6 | Enrichr CHEA_2022: SOX2 20726797 ChIP-Seq SW620 Human | SOX2 |
| 1887 | CAMK2D | Enrichr CHEA_2022: SOX2 20726797 ChIP-Seq SW620 Human | SOX2 |
| 1888 | TSEN34 | Enrichr CHEA_2022: SOX2 20726797 ChIP-Seq SW620 Human | SOX2 |
| 1889 | STEAP4 | Enrichr CHEA_2022: SOX2 20726797 ChIP-Seq SW620 Human | SOX2 |

| | | | |
|---|---|---|---|
| 1890 | ZNF174 | Enrichr CHEA_2022: SOX2 20726797 ChIP-Seq SW620 Human | SOX2 |
| 1891 | USP11 | Enrichr CHEA_2022: SOX2 20726797 ChIP-Seq SW620 Human | SOX2 |
| 1892 | USP12 | Enrichr CHEA_2022: SOX2 20726797 ChIP-Seq SW620 Human | SOX2 |
| 1893 | USP13 | Enrichr CHEA_2022: SOX2 20726797 ChIP-Seq SW620 Human | SOX2 |
| 1894 | CD163L1 | Enrichr CHEA_2022: SOX2 20726797 ChIP-Seq SW620 Human | SOX2 |
| 1895 | HSPD1 | Enrichr CHEA_2022: SOX2 20726797 ChIP-Seq SW620 Human | SOX2 |
| 1896 | USP18 | Enrichr CHEA_2022: SOX2 20726797 ChIP-Seq SW620 Human | SOX2 |
| 1897 | SH3YL1 | Enrichr CHEA_2022: SOX2 20726797 ChIP-Seq SW620 Human | SOX2 |
| 1898 | NT5DC1 | Enrichr CHEA_2022: SOX2 20726797 ChIP-Seq SW620 Human | SOX2 |
| 1899 | PRRG4 | Enrichr CHEA_2022: SOX2 20726797 ChIP-Seq SW620 Human | SOX2 |
| 1900 | POLB | Enrichr CHEA_2022: SOX2 20726797 ChIP-Seq SW620 Human | SOX2 |
| 1901 | VSIG2 | Enrichr CHEA_2022: SOX2 20726797 ChIP-Seq SW620 Human | SOX2 |
| 1902 | VSIG1 | Enrichr CHEA_2022: SOX2 20726797 ChIP-Seq SW620 Human | SOX2 |
| 1903 | CCNJL | Enrichr TF Perturbations followed by expression: KLF5 20726797 ChIP-Seq SW620 Human | KLF5 |
| 1904 | RBFOX3 | Enrichr TF Perturbations followed by expression: KLF5 20726797 ChIP-Seq SW620 Human | KLF5 |
| 1905 | SP110 | Enrichr TF Perturbations followed by expression: KLF5 20726797 ChIP-Seq SW620 Human | KLF5 |
| 1906 | HOMER2 | Enrichr TF Perturbations followed by expression: KLF5 20726797 ChIP-Seq SW620 Human | KLF5 |
| 1907 | KRT4 | Enrichr TF Perturbations followed by expression: KLF5 20726797 ChIP-Seq SW620 Human | KLF5 |
| 1908 | ARL14 | Enrichr TF Perturbations followed by expression: KLF5 20726797 ChIP-Seq SW620 Human | KLF5 |
| 1909 | CKMT1A | Enrichr TF Perturbations followed by expression: KLF5 20726797 ChIP-Seq SW620 Human | KLF5 |
| 1910 | KRT7 | Enrichr TF Perturbations followed by expression: KLF5 20726797 ChIP-Seq SW620 Human | KLF5 |
| 1911 | L1CAM | Enrichr TF Perturbations followed by expression: KLF5 20726797 ChIP-Seq SW620 Human | KLF5 |
| 1912 | MALL | Enrichr TF Perturbations followed by expression: KLF5 20726797 ChIP-Seq SW620 Human | KLF5 |
| 1913 | CAPN13 | Enrichr TF Perturbations followed by expression: KLF5 20726797 ChIP-Seq SW620 Human | KLF5 |
| 1914 | LGALS12 | Enrichr TF Perturbations followed by expression: KLF5 20726797 ChIP-Seq SW620 Human | KLF5 |
| 1915 | MYL9 | Enrichr TF Perturbations followed by expression: KLF5 20726797 ChIP-Seq SW620 Human | KLF5 |
| 1916 | VIP | Enrichr TF Perturbations followed by expression: KLF5 20726797 ChIP-Seq SW620 Human | KLF5 |
| 1917 | EPHA4 | Enrichr TF Perturbations followed by expression: KLF5 20726797 ChIP-Seq SW620 Human | KLF5 |
| 1918 | CHIC1 | Enrichr TF Perturbations followed by expression: KLF5 20726797 ChIP-Seq SW620 Human | KLF5 |
| 1919 | SYNPO2 | Enrichr TF Perturbations followed by expression: KLF5 20726797 ChIP-Seq SW620 Human | KLF5 |
| 1920 | MST1R | Enrichr TF Perturbations followed by expression: KLF5 20726797 ChIP-Seq SW620 Human | KLF5 |
| 1921 | CRACR2A | Enrichr TF Perturbations followed by expression: KLF5 20726797 ChIP-Seq SW620 Human | KLF5 |
| 1922 | MUC1 | Enrichr TF Perturbations followed by expression: KLF5 20726797 ChIP-Seq SW620 Human | KLF5 |
| 1923 | STRIP2 | Enrichr TF Perturbations followed by expression: KLF5 20726797 ChIP-Seq SW620 Human | KLF5 |
| 1924 | GNG3 | Enrichr TF Perturbations followed by expression: KLF5 20726797 ChIP-Seq SW620 Human | KLF5 |
| 1925 | CREB3L4 | Enrichr TF Perturbations followed by expression: KLF5 20726797 ChIP-Seq SW620 Human | KLF5 |
| 1926 | SV2C | Enrichr TF Perturbations followed by expression: KLF5 20726797 ChIP-Seq SW620 Human | KLF5 |
| 1927 | GMDS | Enrichr TF Perturbations followed by expression: KLF5 20726797 ChIP-Seq SW620 Human | KLF5 |
| 1928 | SPOCK3 | Enrichr TF Perturbations followed by expression: KLF5 20726797 ChIP-Seq SW620 Human | KLF5 |
| 1929 | S100A14 | Enrichr TF Perturbations followed by expression: KLF5 20726797 ChIP-Seq SW620 Human | KLF5 |
| 1930 | SUSD4 | Enrichr TF Perturbations followed by expression: KLF5 20726797 ChIP-Seq SW620 Human | KLF5 |
| 1931 | SPOCK2 | Enrichr TF Perturbations followed by expression: KLF5 20726797 ChIP-Seq SW620 Human | KLF5 |
| 1932 | MUC4 | Enrichr TF Perturbations followed by expression: KLF5 20726797 ChIP-Seq SW620 Human | KLF5 |
| 1933 | SLC12A8 | Enrichr TF Perturbations followed by expression: KLF5 20726797 ChIP-Seq SW620 Human | KLF5 |
| 1934 | PPARGC1B | Enrichr TF Perturbations followed by expression: KLF5 20726797 ChIP-Seq SW620 Human | KLF5 |
| 1935 | FZD3 | Enrichr TF Perturbations followed by expression: KLF5 20726797 ChIP-Seq SW620 Human | KLF5 |
| 1936 | PAQR6 | Enrichr TF Perturbations followed by expression: KLF5 20726797 ChIP-Seq SW620 Human | KLF5 |
| 1937 | RAB27A | Enrichr TF Perturbations followed by expression: KLF5 20726797 ChIP-Seq SW620 Human | KLF5 |
| 1938 | CMBL | Enrichr TF Perturbations followed by expression: KLF5 20726797 ChIP-Seq SW620 Human | KLF5 |
| 1939 | ANK2 | Enrichr TF Perturbations followed by expression: KLF5 20726797 ChIP-Seq SW620 Human | KLF5 |
| 1940 | FA2H | Enrichr TF Perturbations followed by expression: KLF5 20726797 ChIP-Seq SW620 Human | KLF5 |
| 1941 | CNKSR2 | Enrichr TF Perturbations followed by expression: KLF5 20726797 ChIP-Seq SW620 Human | KLF5 |
| 1942 | POR | Enrichr TF Perturbations followed by expression: KLF5 20726797 ChIP-Seq SW620 Human | KLF5 |
| 1943 | RIPPLY3 | Enrichr TF Perturbations followed by expression: KLF5 20726797 ChIP-Seq SW620 Human | KLF5 |
| 1944 | PPP1R1B | Enrichr TF Perturbations followed by expression: KLF5 20726797 ChIP-Seq SW620 Human | KLF5 |
| 1945 | SNHG11 | Enrichr TF Perturbations followed by expression: KLF5 20726797 ChIP-Seq SW620 Human | KLF5 |
| 1946 | SPATS2L | Enrichr TF Perturbations followed by expression: KLF5 20726797 ChIP-Seq SW620 Human | KLF5 |
| 1947 | IGSF9B | Enrichr TF Perturbations followed by expression: KLF5 20726797 ChIP-Seq SW620 Human | KLF5 |
| 1948 | XKR4 | Enrichr TF Perturbations followed by expression: KLF5 20726797 ChIP-Seq SW620 Human | KLF5 |
| 1949 | PTGES | Enrichr TF Perturbations followed by expression: KLF5 20726797 ChIP-Seq SW620 Human | KLF5 |
| 1950 | GLDC | Enrichr TF Perturbations followed by expression: KLF5 20726797 ChIP-Seq SW620 Human | KLF5 |

| | | | |
|---|---|---|---|
| 1951 | ACSM3 | Enrichr TF Perturbations followed by expression: KLF5 20726797 ChIP-Seq SW620 Human | KLF5 |
| 1952 | NECAB1 | Enrichr TF Perturbations followed by expression: KLF5 20726797 ChIP-Seq SW620 Human | KLF5 |
| 1953 | HHIP | Enrichr TF Perturbations followed by expression: KLF5 20726797 ChIP-Seq SW620 Human | KLF5 |
| 1954 | NAPEPLD | Enrichr TF Perturbations followed by expression: KLF5 20726797 ChIP-Seq SW620 Human | KLF5 |
| 1955 | WFDC2 | Enrichr TF Perturbations followed by expression: KLF5 20726797 ChIP-Seq SW620 Human | KLF5 |
| 1956 | RAPGEFL1 | Enrichr TF Perturbations followed by expression: KLF5 20726797 ChIP-Seq SW620 Human | KLF5 |
| 1957 | SESTD1 | Enrichr TF Perturbations followed by expression: KLF5 20726797 ChIP-Seq SW620 Human | KLF5 |
| 1958 | GATA2 | Enrichr TF Perturbations followed by expression: KLF5 20726797 ChIP-Seq SW620 Human | KLF5 |
| 1959 | SOX2 | Enrichr TF Perturbations followed by expression: KLF5 20726797 ChIP-Seq SW620 Human | KLF5 |
| 1960 | TSPAN8 | Enrichr TF Perturbations followed by expression: KLF5 20726797 ChIP-Seq SW620 Human | KLF5 |
| 1961 | REG4 | Enrichr TF Perturbations followed by expression: KLF5 20726797 ChIP-Seq SW620 Human | KLF5 |
| 1962 | FFAR4 | Enrichr TF Perturbations followed by expression: KLF5 20726797 ChIP-Seq SW620 Human | KLF5 |
| 1963 | TNFSF10 | Enrichr TF Perturbations followed by expression: KLF5 20726797 ChIP-Seq SW620 Human | KLF5 |
| 1964 | CD14 | Enrichr TF Perturbations followed by expression: KLF5 20726797 ChIP-Seq SW620 Human | KLF5 |
| 1965 | TSPAN2 | Enrichr TF Perturbations followed by expression: KLF5 20726797 ChIP-Seq SW620 Human | KLF5 |
| 1966 | PROM1 | Enrichr TF Perturbations followed by expression: KLF5 20726797 ChIP-Seq SW620 Human | KLF5 |
| 1967 | LRRC26 | Enrichr TF Perturbations followed by expression: KLF5 20726797 ChIP-Seq SW620 Human | KLF5 |
| 1968 | PCP4L1 | Enrichr TF Perturbations followed by expression: KLF5 20726797 ChIP-Seq SW620 Human | KLF5 |
| 1969 | GSDMC | Enrichr TF Perturbations followed by expression: KLF5 20726797 ChIP-Seq SW620 Human | KLF5 |
| 1970 | TESC | Enrichr TF Perturbations followed by expression: KLF5 20726797 ChIP-Seq SW620 Human | KLF5 |
| 1971 | GP2 | Enrichr TF Perturbations followed by expression: KLF5 20726797 ChIP-Seq SW620 Human | KLF5 |
| 1972 | TMC5 | Enrichr TF Perturbations followed by expression: KLF5 20726797 ChIP-Seq SW620 Human | KLF5 |
| 1973 | SYN2 | Enrichr TF Perturbations followed by expression: KLF5 20726797 ChIP-Seq SW620 Human | KLF5 |
| 1974 | GPRC5A | Enrichr TF Perturbations followed by expression: KLF5 20726797 ChIP-Seq SW620 Human | KLF5 |
| 1975 | TTLL7 | Enrichr TF Perturbations followed by expression: KLF5 20726797 ChIP-Seq SW620 Human | KLF5 |
| 1976 | TOX3 | Enrichr TF Perturbations followed by expression: KLF5 20726797 ChIP-Seq SW620 Human | KLF5 |
| 1977 | TPPP3 | Enrichr TF Perturbations followed by expression: KLF5 20726797 ChIP-Seq SW620 Human | KLF5 |
| 1978 | ARHGEF4 | Enrichr TF Perturbations followed by expression: KLF5 20726797 ChIP-Seq SW620 Human | KLF5 |
| 1979 | TFF3 | Enrichr TF Perturbations followed by expression: KLF5 20726797 ChIP-Seq SW620 Human | KLF5 |
| 1980 | RET | Enrichr TF Perturbations followed by expression: KLF5 20726797 ChIP-Seq SW620 Human | KLF5 |
| 1981 | KLK1 | Enrichr TF Perturbations followed by expression: KLF5 20726797 ChIP-Seq SW620 Human | KLF5 |
| 1982 | TMPRSS4 | Enrichr TF Perturbations followed by expression: KLF5 20726797 ChIP-Seq SW620 Human | KLF5 |
| 1983 | NR1I3 | Enrichr TF Perturbations followed by expression: KLF5 20726797 ChIP-Seq SW620 Human | KLF5 |
| 1984 | TMPRSS2 | Enrichr TF Perturbations followed by expression: KLF5 20726797 ChIP-Seq SW620 Human | KLF5 |
| 1985 | HNF4G | Enrichr TF Perturbations followed by expression: KLF5 20726797 ChIP-Seq SW620 Human | KLF5 |
| 1986 | SLC7A14 | Enrichr TF Perturbations followed by expression: KLF5 20726797 ChIP-Seq SW620 Human | KLF5 |
| 1987 | HEPACAM2 | Enrichr TF Perturbations followed by expression: KLF5 20726797 ChIP-Seq SW620 Human | KLF5 |
| 1988 | HIF3A | Enrichr TF Perturbations followed by expression: KLF5 20726797 ChIP-Seq SW620 Human | KLF5 |
| 1989 | KRT23 | Enrichr TF Perturbations followed by expression: KLF5 20726797 ChIP-Seq SW620 Human | KLF5 |
| 1990 | KIF19 | Enrichr TF Perturbations followed by expression: KLF5 20726797 ChIP-Seq SW620 Human | KLF5 |
| 1991 | ACACB | Enrichr TF Perturbations followed by expression: KLF5 20726797 ChIP-Seq SW620 Human | KLF5 |
| 1992 | HSD11B2 | Enrichr TF Perturbations followed by expression: KLF5 20726797 ChIP-Seq SW620 Human | KLF5 |
| 1993 | SULT1C2 | Enrichr TF Perturbations followed by expression: KLF5 20726797 ChIP-Seq SW620 Human | KLF5 |
| 1994 | UCHL1 | Enrichr TF Perturbations followed by expression: KLF5 20726797 ChIP-Seq SW620 Human | KLF5 |
| 1995 | MECOM | Enrichr TF Perturbations followed by expression: KLF5 20726797 ChIP-Seq SW620 Human | KLF5 |
| 1996 | RNF150 | Enrichr TF Perturbations followed by expression: KLF5 20726797 ChIP-Seq SW620 Human | KLF5 |
| 1997 | QSOX1 | Enrichr TF Perturbations followed by expression: KLF5 20726797 ChIP-Seq SW620 Human | KLF5 |
| 1998 | PLLP | Enrichr TF Perturbations followed by expression: KLF5 20726797 ChIP-Seq SW620 Human | KLF5 |
| 1999 | RAB6B | Enrichr TF Perturbations followed by expression: KLF5 20726797 ChIP-Seq SW620 Human | KLF5 |
| 2000 | CTSE | Enrichr TF Perturbations followed by expression: KLF5 20726797 ChIP-Seq SW620 Human | KLF5 |
| 2001 | DGAT2 | Enrichr TF Perturbations followed by expression: KLF5 20726797 ChIP-Seq SW620 Human | KLF5 |
| 2002 | GSTO1 | Enrichr TF Perturbations followed by expression: KLF5 20726797 ChIP-Seq SW620 Human | KLF5 |
| 2003 | ANO9 | Enrichr TF Perturbations followed by expression: KLF5 20726797 ChIP-Seq SW620 Human | KLF5 |
| 2004 | SYP | Enrichr TF Perturbations followed by expression: KLF5 20726797 ChIP-Seq SW620 Human | KLF5 |
| 2005 | SORBS1 | Enrichr TF Perturbations followed by expression: KLF5 20726797 ChIP-Seq SW620 Human | KLF5 |
| 2006 | SLC35G2 | Enrichr TF Perturbations followed by expression: KLF5 20726797 ChIP-Seq SW620 Human | KLF5 |
| 2007 | F3 | Enrichr TF Perturbations followed by expression: KLF5 20726797 ChIP-Seq SW620 Human | KLF5 |
| 2008 | CLIP4 | Enrichr TF Perturbations followed by expression: KLF5 20726797 ChIP-Seq SW620 Human | KLF5 |
| 2009 | DHRS9 | Enrichr TF Perturbations followed by expression: KLF5 20726797 ChIP-Seq SW620 Human | KLF5 |
| 2010 | KCNQ1 | Enrichr TF Perturbations followed by expression: KLF5 20726797 ChIP-Seq SW620 Human | KLF5 |
| 2011 | FABP7 | Enrichr TF Perturbations followed by expression: KLF5 20726797 ChIP-Seq SW620 Human | KLF5 |

| | | | |
|---|---|---|---|
| 2012 | PPARG | Enrichr TF Perturbations followed by expression: KLF5 20726797 ChIP-Seq SW620 Human | KLF5 |
| 2013 | B3GNT3 | Enrichr TF Perturbations followed by expression: KLF5 20726797 ChIP-Seq SW620 Human | KLF5 |
| 2014 | ENGASE | Enrichr TF Perturbations followed by expression: KLF5 20726797 ChIP-Seq SW620 Human | KLF5 |
| 2015 | SERPINB11 | Enrichr TF Perturbations followed by expression: KLF5 20726797 ChIP-Seq SW620 Human | KLF5 |
| 2016 | PRSS27 | Enrichr TF Perturbations followed by expression: KLF5 20726797 ChIP-Seq SW620 Human | KLF5 |
| 2017 | SH3RF2 | Enrichr TF Perturbations followed by expression: KLF5 20726797 ChIP-Seq SW620 Human | KLF5 |
| 2018 | SMPD3 | Enrichr TF Perturbations followed by expression: KLF5 20726797 ChIP-Seq SW620 Human | KLF5 |
| 2019 | ADAM28 | Enrichr TF Perturbations followed by expression: KLF5 20726797 ChIP-Seq SW620 Human | KLF5 |
| 2020 | SULT1E1 | Enrichr TF Perturbations followed by expression: KLF5 20726797 ChIP-Seq SW620 Human | KLF5 |
| 2021 | NEFL | Enrichr TF Perturbations followed by expression: KLF5 20726797 ChIP-Seq SW620 Human | KLF5 |
| 2022 | MKRN2OS | Enrichr TF Perturbations followed by expression: KLF5 20726797 ChIP-Seq SW620 Human | KLF5 |
| 2023 | FAM3B | Enrichr TF Perturbations followed by expression: KLF5 20726797 ChIP-Seq SW620 Human | KLF5 |
| 2024 | ENPP2 | Enrichr TF Perturbations followed by expression: KLF5 20726797 ChIP-Seq SW620 Human | KLF5 |
| 2025 | GPA33 | Enrichr TF Perturbations followed by expression: KLF5 20726797 ChIP-Seq SW620 Human | KLF5 |
| 2026 | MGAT3 | Enrichr TF Perturbations followed by expression: KLF5 20726797 ChIP-Seq SW620 Human | KLF5 |
| 2027 | GSN | Enrichr TF Perturbations followed by expression: KLF5 20726797 ChIP-Seq SW620 Human | KLF5 |
| 2028 | ABCG8 | Enrichr TF Perturbations followed by expression: KLF5 20726797 ChIP-Seq SW620 Human | KLF5 |
| 2029 | GABRP | Enrichr TF Perturbations followed by expression: KLF5 20726797 ChIP-Seq SW620 Human | KLF5 |
| 2030 | NPNT | Enrichr TF Perturbations followed by expression: KLF5 20726797 ChIP-Seq SW620 Human | KLF5 |
| 2031 | GOLM1 | Enrichr TF Perturbations followed by expression: KLF5 20726797 ChIP-Seq SW620 Human | KLF5 |
| 2032 | AGT | Enrichr TF Perturbations followed by expression: KLF5 20726797 ChIP-Seq SW620 Human | KLF5 |
| 2033 | TC2N | Enrichr TF Perturbations followed by expression: KLF5 20726797 ChIP-Seq SW620 Human | KLF5 |
| 2034 | ERN2 | Enrichr TF Perturbations followed by expression: KLF5 20726797 ChIP-Seq SW620 Human | KLF5 |
| 2035 | OLFM4 | Enrichr TF Perturbations followed by expression: KLF5 20726797 ChIP-Seq SW620 Human | KLF5 |
| 2036 | PCP4 | Enrichr TF Perturbations followed by expression: KLF5 20726797 ChIP-Seq SW620 Human | KLF5 |
| 2037 | GSTA4 | Enrichr TF Perturbations followed by expression: KLF5 20726797 ChIP-Seq SW620 Human | KLF5 |
| 2038 | GSTA3 | Enrichr TF Perturbations followed by expression: KLF5 20726797 ChIP-Seq SW620 Human | KLF5 |
| 2039 | S100A6 | Enrichr TF Perturbations followed by expression: KLF5 20726797 ChIP-Seq SW620 Human | KLF5 |
| 2040 | LY6D | Enrichr TF Perturbations followed by expression: KLF5 20726797 ChIP-Seq SW620 Human | KLF5 |
| 2041 | ABCG5 | Enrichr TF Perturbations followed by expression: KLF5 20726797 ChIP-Seq SW620 Human | KLF5 |
| 2042 | CHRNA3 | Enrichr TF Perturbations followed by expression: KLF5 20726797 ChIP-Seq SW620 Human | KLF5 |
| 2043 | TNXB | Enrichr TF Perturbations followed by expression: KLF5 20726797 ChIP-Seq SW620 Human | KLF5 |
| 2044 | DGKB | Enrichr TF Perturbations followed by expression: KLF5 20726797 ChIP-Seq SW620 Human | KLF5 |
| 2045 | SCNN1A | Enrichr TF Perturbations followed by expression: KLF5 20726797 ChIP-Seq SW620 Human | KLF5 |
| 2046 | BEST2 | Enrichr TF Perturbations followed by expression: KLF5 20726797 ChIP-Seq SW620 Human | KLF5 |
| 2047 | CAPN8 | Enrichr TF Perturbations followed by expression: KLF5 20726797 ChIP-Seq SW620 Human | KLF5 |
| 2048 | ARHGAP40 | Enrichr TF Perturbations followed by expression: KLF5 20726797 ChIP-Seq SW620 Human | KLF5 |
| 2049 | ASB11 | Enrichr TF Perturbations followed by expression: KLF5 20726797 ChIP-Seq SW620 Human | KLF5 |
| 2050 | CAPN5 | Enrichr TF Perturbations followed by expression: KLF5 20726797 ChIP-Seq SW620 Human | KLF5 |
| 2051 | GPC3 | Enrichr TF Perturbations followed by expression: KLF5 20726797 ChIP-Seq SW620 Human | KLF5 |
| 2052 | PDE8B | Enrichr TF Perturbations followed by expression: KLF5 20726797 ChIP-Seq SW620 Human | KLF5 |
| 2053 | MYH11 | Enrichr TF Perturbations followed by expression: KLF5 20726797 ChIP-Seq SW620 Human | KLF5 |
| 2054 | KCNH2 | Enrichr TF Perturbations followed by expression: KLF5 20726797 ChIP-Seq SW620 Human | KLF5 |
| 2055 | SLC39A4 | Enrichr TF Perturbations followed by expression: KLF5 20726797 ChIP-Seq SW620 Human | KLF5 |
| 2056 | IGFBP2 | Enrichr TF Perturbations followed by expression: KLF5 20726797 ChIP-Seq SW620 Human | KLF5 |
| 2057 | LYNX1 | Enrichr TF Perturbations followed by expression: KLF5 20726797 ChIP-Seq SW620 Human | KLF5 |
| 2058 | PSAPL1 | Enrichr TF Perturbations followed by expression: KLF5 20726797 ChIP-Seq SW620 Human | KLF5 |
| 2059 | SYT1 | Enrichr TF Perturbations followed by expression: KLF5 20726797 ChIP-Seq SW620 Human | KLF5 |
| 2060 | B3GALT5 | Enrichr TF Perturbations followed by expression: KLF5 20726797 ChIP-Seq SW620 Human | KLF5 |
| 2061 | CRNDE | Enrichr TF Perturbations followed by expression: KLF5 20726797 ChIP-Seq SW620 Human | KLF5 |
| 2062 | SYT8 | Enrichr TF Perturbations followed by expression: KLF5 20726797 ChIP-Seq SW620 Human | KLF5 |
| 2063 | ADRA2A | Enrichr TF Perturbations followed by expression: KLF5 20726797 ChIP-Seq SW620 Human | KLF5 |
| 2064 | TMIE | Enrichr TF Perturbations followed by expression: KLF5 20726797 ChIP-Seq SW620 Human | KLF5 |
| 2065 | REEP6 | Enrichr TF Perturbations followed by expression: KLF5 20726797 ChIP-Seq SW620 Human | KLF5 |
| 2066 | MAL | Enrichr TF Perturbations followed by expression: KLF5 20726797 ChIP-Seq SW620 Human | KLF5 |
| 2067 | HPSE | Enrichr TF Perturbations followed by expression: KLF5 20726797 ChIP-Seq SW620 Human | KLF5 |
| 2068 | SPIRE2 | Enrichr TF Perturbations followed by expression: KLF5 20726797 ChIP-Seq SW620 Human | KLF5 |
| 2069 | MUC20 | Enrichr TF Perturbations followed by expression: KLF5 20726797 ChIP-Seq SW620 Human | KLF5 |
| 2070 | CDH19 | Enrichr TF Perturbations followed by expression: KLF5 20726797 ChIP-Seq SW620 Human | KLF5 |
| 2071 | FGFR3 | Enrichr TF Perturbations followed by expression: KLF5 20726797 ChIP-Seq SW620 Human | KLF5 |
| 2072 | CERS3 | Enrichr TF Perturbations followed by expression: KLF5 20726797 ChIP-Seq SW620 Human | KLF5 |

| | | | |
|---|---|---|---|
| 2073 | RAB3C | Enrichr TF Perturbations followed by expression: KLF5 20726797 ChIP-Seq SW620 Human | KLF5 |
| 2074 | KCNE3 | Enrichr TF Perturbations followed by expression: KLF5 20726797 ChIP-Seq SW620 Human | KLF5 |
| 2075 | KCNE4 | Enrichr TF Perturbations followed by expression: KLF5 20726797 ChIP-Seq SW620 Human | KLF5 |
| 2076 | UPK2 | Enrichr TF Perturbations followed by expression: KLF5 20726797 ChIP-Seq SW620 Human | KLF5 |
| 2077 | TACSTD2 | Enrichr TF Perturbations followed by expression: KLF5 20726797 ChIP-Seq SW620 Human | KLF5 |
| 2078 | FAM107A | Enrichr TF Perturbations followed by expression: KLF5 20726797 ChIP-Seq SW620 Human | KLF5 |
| 2079 | PPP1R9A | Enrichr TF Perturbations followed by expression: KLF5 20726797 ChIP-Seq SW620 Human | KLF5 |
| 2080 | GPR83 | Enrichr TF Perturbations followed by expression: KLF5 20726797 ChIP-Seq SW620 Human | KLF5 |
| 2081 | PLCXD3 | Enrichr TF Perturbations followed by expression: KLF5 20726797 ChIP-Seq SW620 Human | KLF5 |
| 2082 | ELFN1 | Enrichr TF Perturbations followed by expression: KLF5 20726797 ChIP-Seq SW620 Human | KLF5 |
| 2083 | ACTG2 | Enrichr TF Perturbations followed by expression: KLF5 20726797 ChIP-Seq SW620 Human | KLF5 |
| 2084 | EFEMP1 | Enrichr TF Perturbations followed by expression: KLF5 20726797 ChIP-Seq SW620 Human | KLF5 |
| 2085 | CNN1 | Enrichr TF Perturbations followed by expression: KLF5 20726797 ChIP-Seq SW620 Human | KLF5 |
| 2086 | ALDH3B2 | Enrichr TF Perturbations followed by expression: KLF5 20726797 ChIP-Seq SW620 Human | KLF5 |
| 2087 | B3GALT2 | Enrichr TF Perturbations followed by expression: KLF5 20726797 ChIP-Seq SW620 Human | KLF5 |
| 2088 | CHAD | Enrichr TF Perturbations followed by expression: KLF5 20726797 ChIP-Seq SW620 Human | KLF5 |
| 2089 | LEPR | Enrichr TF Perturbations followed by expression: KLF5 20726797 ChIP-Seq SW620 Human | KLF5 |
| 2090 | PRODH | Enrichr TF Perturbations followed by expression: KLF5 20726797 ChIP-Seq SW620 Human | KLF5 |
| 2091 | EFHD1 | Enrichr TF Perturbations followed by expression: KLF5 20726797 ChIP-Seq SW620 Human | KLF5 |
| 2092 | SH3GL2 | Enrichr TF Perturbations followed by expression: KLF5 20726797 ChIP-Seq SW620 Human | KLF5 |
| 2093 | SLC37A1 | Enrichr TF Perturbations followed by expression: KLF5 20726797 ChIP-Seq SW620 Human | KLF5 |
| 2094 | GALNT7 | Enrichr TF Perturbations followed by expression: KLF5 20726797 ChIP-Seq SW620 Human | KLF5 |
| 2095 | GALNT5 | Enrichr TF Perturbations followed by expression: KLF5 20726797 ChIP-Seq SW620 Human | KLF5 |
| 2096 | HMGCS2 | Enrichr TF Perturbations followed by expression: KLF5 20726797 ChIP-Seq SW620 Human | KLF5 |
| 2097 | GALNT4 | Enrichr TF Perturbations followed by expression: KLF5 20726797 ChIP-Seq SW620 Human | KLF5 |
| 2098 | PEG3 | Enrichr TF Perturbations followed by expression: KLF5 20726797 ChIP-Seq SW620 Human | KLF5 |
| 2099 | MYOCD | Enrichr TF Perturbations followed by expression: KLF5 20726797 ChIP-Seq SW620 Human | KLF5 |
| 2100 | CADPS | Enrichr TF Perturbations followed by expression: KLF5 20726797 ChIP-Seq SW620 Human | KLF5 |
| 2101 | TNFRSF19 | Enrichr TF Perturbations followed by expression: KLF5 20726797 ChIP-Seq SW620 Human | KLF5 |
| 2102 | INMT | Enrichr TF Perturbations followed by expression: KLF5 20726797 ChIP-Seq SW620 Human | KLF5 |
| 2103 | ELF3 | Enrichr TF Perturbations followed by expression: KLF5 20726797 ChIP-Seq SW620 Human | KLF5 |
| 2104 | TNIP3 | Enrichr TF Perturbations followed by expression: KLF5 20726797 ChIP-Seq SW620 Human | KLF5 |
| 2105 | GNAO1 | Enrichr TF Perturbations followed by expression: KLF5 20726797 ChIP-Seq SW620 Human | KLF5 |
| 2106 | DSG3 | Enrichr TF Perturbations followed by expression: KLF5 20726797 ChIP-Seq SW620 Human | KLF5 |
| 2107 | SOWAHA | Enrichr TF Perturbations followed by expression: KLF5 20726797 ChIP-Seq SW620 Human | KLF5 |
| 2108 | SNAP25 | Enrichr TF Perturbations followed by expression: KLF5 20726797 ChIP-Seq SW620 Human | KLF5 |
| 2109 | ST6GALNAC1 | Enrichr TF Perturbations followed by expression: KLF5 20726797 ChIP-Seq SW620 Human | KLF5 |
| 2110 | ST6GALNAC2 | Enrichr TF Perturbations followed by expression: KLF5 20726797 ChIP-Seq SW620 Human | KLF5 |
| 2111 | RTN1 | Enrichr TF Perturbations followed by expression: KLF5 20726797 ChIP-Seq SW620 Human | KLF5 |
| 2112 | SMIM6 | Enrichr TF Perturbations followed by expression: KLF5 20726797 ChIP-Seq SW620 Human | KLF5 |
| 2113 | CRYBG2 | Enrichr TF Perturbations followed by expression: KLF5 20726797 ChIP-Seq SW620 Human | KLF5 |
| 2114 | TTC22 | Enrichr TF Perturbations followed by expression: KLF5 20726797 ChIP-Seq SW620 Human | KLF5 |
| 2115 | TNFAIP8 | Enrichr TF Perturbations followed by expression: KLF5 20726797 ChIP-Seq SW620 Human | KLF5 |
| 2116 | NPR3 | Enrichr TF Perturbations followed by expression: KLF5 20726797 ChIP-Seq SW620 Human | KLF5 |
| 2117 | ITPR2 | Enrichr TF Perturbations followed by expression: KLF5 20726797 ChIP-Seq SW620 Human | KLF5 |
| 2118 | FBXO44 | Enrichr TF Perturbations followed by expression: KLF5 20726797 ChIP-Seq SW620 Human | KLF5 |
| 2119 | RASGEF1A | Enrichr TF Perturbations followed by expression: KLF5 20726797 ChIP-Seq SW620 Human | KLF5 |
| 2120 | FUT8 | Enrichr TF Perturbations followed by expression: KLF5 20726797 ChIP-Seq SW620 Human | KLF5 |
| 2121 | UPK1B | Enrichr TF Perturbations followed by expression: KLF5 20726797 ChIP-Seq SW620 Human | KLF5 |
| 2122 | RDH16 | Enrichr TF Perturbations followed by expression: KLF5 20726797 ChIP-Seq SW620 Human | KLF5 |
| 2123 | CPXM2 | Enrichr TF Perturbations followed by expression: KLF5 20726797 ChIP-Seq SW620 Human | KLF5 |
| 2124 | SYMPK | Enrichr TF Perturbations followed by expression: KLF5 20726797 ChIP-Seq SW620 Human | KLF5 |
| 2125 | CASP1 | Enrichr TF Perturbations followed by expression: KLF5 20726797 ChIP-Seq SW620 Human | KLF5 |
| 2126 | FAM83E | Enrichr TF Perturbations followed by expression: KLF5 20726797 ChIP-Seq SW620 Human | KLF5 |
| 2127 | PRPH | Enrichr TF Perturbations followed by expression: KLF5 20726797 ChIP-Seq SW620 Human | KLF5 |
| 2128 | WNT4 | Enrichr TF Perturbations followed by expression: KLF5 20726797 ChIP-Seq SW620 Human | KLF5 |
| 2129 | HGFAC | Enrichr TF Perturbations followed by expression: KLF5 20726797 ChIP-Seq SW620 Human | KLF5 |
| 2130 | TBX1 | Enrichr TF Perturbations followed by expression: KLF5 20726797 ChIP-Seq SW620 Human | KLF5 |
| 2131 | CAMK1D | Enrichr TF Perturbations followed by expression: KLF5 20726797 ChIP-Seq SW620 Human | KLF5 |
| 2132 | MUC5B | Enrichr TF Perturbations followed by expression: KLF5 20726797 ChIP-Seq SW620 Human | KLF5 |
| 2133 | STYK1 | Enrichr TF Perturbations followed by expression: KLF5 20726797 ChIP-Seq SW620 Human | KLF5 |

| | | | |
|---|---|---|---|
| 2134 | DCXR | Enrichr TF Perturbations followed by expression: KLF5 20726797 ChIP-Seq SW620 Human | KLF5 |
| 2135 | SYT16 | Enrichr TF Perturbations followed by expression: KLF5 20726797 ChIP-Seq SW620 Human | KLF5 |
| 2136 | MCOLN2 | Enrichr TF Perturbations followed by expression: KLF5 20726797 ChIP-Seq SW620 Human | KLF5 |
| 2137 | KLF5 | Enrichr TF Perturbations followed by expression: KLF5 20726797 ChIP-Seq SW620 Human | KLF5 |
| 2138 | HTR3A | Enrichr TF Perturbations followed by expression: KLF5 20726797 ChIP-Seq SW620 Human | KLF5 |
| 2139 | HSH2D | Enrichr TF Perturbations followed by expression: KLF5 20726797 ChIP-Seq SW620 Human | KLF5 |
| 2140 | TST | Enrichr TF Perturbations followed by expression: KLF5 20726797 ChIP-Seq SW620 Human | KLF5 |
| 2141 | P2RX1 | Enrichr TF Perturbations followed by expression: KLF5 20726797 ChIP-Seq SW620 Human | KLF5 |
| 2142 | LTF | Enrichr TF Perturbations followed by expression: KLF5 20726797 ChIP-Seq SW620 Human | KLF5 |
| 2143 | ASPA | Enrichr TF Perturbations followed by expression: KLF5 20726797 ChIP-Seq SW620 Human | KLF5 |
| 2144 | SLC24A3 | Enrichr TF Perturbations followed by expression: KLF5 20726797 ChIP-Seq SW620 Human | KLF5 |
| 2145 | SH2D4A | Enrichr TF Perturbations followed by expression: KLF5 20726797 ChIP-Seq SW620 Human | KLF5 |
| 2146 | MSLN | Enrichr TF Perturbations followed by expression: KLF5 20726797 ChIP-Seq SW620 Human | KLF5 |
| 2147 | TRIM31 | Enrichr TF Perturbations followed by expression: KLF5 20726797 ChIP-Seq SW620 Human | KLF5 |
| 2148 | PSCA | Enrichr TF Perturbations followed by expression: KLF5 20726797 ChIP-Seq SW620 Human | KLF5 |
| 2149 | SEMA3C | Enrichr TF Perturbations followed by expression: KLF5 20726797 ChIP-Seq SW620 Human | KLF5 |
| 2150 | ATP2A3 | Enrichr TF Perturbations followed by expression: KLF5 20726797 ChIP-Seq SW620 Human | KLF5 |
| 2151 | HSD17B14 | Enrichr TF Perturbations followed by expression: KLF5 20726797 ChIP-Seq SW620 Human | KLF5 |
| 2152 | CLCN2 | Enrichr TF Perturbations followed by expression: KLF5 20726797 ChIP-Seq SW620 Human | KLF5 |
| 2153 | TPRN | Enrichr TF Perturbations followed by expression: KLF5 20726797 ChIP-Seq SW620 Human | KLF5 |
| 2154 | PTPRZ1 | Enrichr TF Perturbations followed by expression: KLF5 20726797 ChIP-Seq SW620 Human | KLF5 |
| 2155 | NT5DC3 | Enrichr TF Perturbations followed by expression: KLF5 20726797 ChIP-Seq SW620 Human | KLF5 |
| 2156 | VSIG1 | Enrichr TF Perturbations followed by expression: KLF5 20726797 ChIP-Seq SW620 Human | KLF5 |
| 2157 | NOS1 | Enrichr TF Perturbations followed by expression: KLF5 20726797 ChIP-Seq SW620 Human | KLF5 |
| 2158 | DLGAP1 | SOX2-KLF5 coregulated genes from PMID: 33972779 | SOX2 + KLF5 |
| 2159 | SLC35E4 | SOX2-KLF5 coregulated genes from PMID: 33972779 | SOX2 + KLF5 |
| 2160 | USP12 | SOX2-KLF5 coregulated genes from PMID: 33972779 | SOX2 + KLF5 |
| 2161 | CARD10 | SOX2-KLF5 coregulated genes from PMID: 33972779 | SOX2 + KLF5 |
| 2162 | KLF5 | SOX2-KLF5 coregulated genes from PMID: 33972779 | SOX2 + KLF5 |
| 2163 | CUBN | SOX2-KLF5 coregulated genes from PMID: 33972779 | SOX2 + KLF5 |
| 2164 | HS3ST1 | SOX2-KLF5 coregulated genes from PMID: 33972779 | SOX2 + KLF5 |
| 2165 | TMEM47 | SOX2-KLF5 coregulated genes from PMID: 33972779 | SOX2 + KLF5 |
| 2166 | RUNX1 | SOX2-KLF5 coregulated genes from PMID: 33972779 | SOX2 + KLF5 |
| 2167 | JAG1 | SOX2-KLF5 coregulated genes from PMID: 33972779 | SOX2 + KLF5 |
| 2168 | PLCB1 | SOX2-KLF5 coregulated genes from PMID: 33972779 | SOX2 + KLF5 |
| 2169 | PCDH17 | SOX2-KLF5 coregulated genes from PMID: 33972779 | SOX2 + KLF5 |
| 2170 | KCTD1 | SOX2-KLF5 coregulated genes from PMID: 33972779 | SOX2 + KLF5 |
| 2171 | DSG3 | SOX2-KLF5 coregulated genes from PMID: 33972779 | SOX2 + KLF5 |
| 2172 | BMP2 | SOX2-KLF5 coregulated genes from PMID: 33972779 | SOX2 + KLF5 |
| 2173 | PTGER2 | SOX2-KLF5 coregulated genes from PMID: 33972779 | SOX2 + KLF5 |
| 2174 | SLC5A3 | SOX2-KLF5 coregulated genes from PMID: 33972779 | SOX2 + KLF5 |
| 2175 | AKAP6 | SOX2-KLF5 coregulated genes from PMID: 33972779 | SOX2 + KLF5 |
| 2176 | SH3GL2 | SOX2-KLF5 coregulated genes from PMID: 33972779 | SOX2 + KLF5 |
| 2177 | TTC39C | SOX2-KLF5 coregulated genes from PMID: 33972779 | SOX2 + KLF5 |
| 2178 | NRIP1 | SOX2-KLF5 coregulated genes from PMID: 33972779 | SOX2 + KLF5 |
| 2179 | OCIAD2 | SOX2-KLF5 coregulated genes from PMID: 33972779 | SOX2 + KLF5 |
| 2180 | KIAA1328 | SOX2-KLF5 coregulated genes from PMID: 33972779 | SOX2 + KLF5 |
| 2181 | ETS2 | SOX2-KLF5 coregulated genes from PMID: 33972779 | SOX2 + KLF5 |
| 2182 | SLC1A1 | SOX2-KLF5 coregulated genes from PMID: 33972779 | SOX2 + KLF5 |
| 2183 | LMO7 | SOX2-KLF5 coregulated genes from PMID: 33972779 | SOX2 + KLF5 |
| 2184 | BTG3 | SOX2-KLF5 coregulated genes from PMID: 33972779 | SOX2 + KLF5 |
| 2185 | DACH1 | SOX2-KLF5 coregulated genes from PMID: 33972779 | SOX2 + KLF5 |
| 2186 | MIPEP | SOX2-KLF5 coregulated genes from PMID: 33972779 | SOX2 + KLF5 |
| 2187 | FERMT2 | SOX2-KLF5 coregulated genes from PMID: 33972779 | SOX2 + KLF5 |
| 2188 | SLIT2 | SOX2-KLF5 coregulated genes from PMID: 33972779 | SOX2 + KLF5 |
| 2189 | MAPRE2 | SOX2-KLF5 coregulated genes from PMID: 33972779 | SOX2 + KLF5 |
| 2190 | FUNDC1 | SOX2-KLF5 coregulated genes from PMID: 33972779 | SOX2 + KLF5 |
| 2191 | NFATC2 | SOX2-KLF5 coregulated genes from PMID: 33972779 | SOX2 + KLF5 |
| 2192 | TXNDC16 | SOX2-KLF5 coregulated genes from PMID: 33972779 | SOX2 + KLF5 |
| 2193 | ALG13 | SOX2-KLF5 coregulated genes from PMID: 33972779 | SOX2 + KLF5 |
| 2194 | RAB27B | SOX2-KLF5 coregulated genes from PMID: 33972779 | SOX2 + KLF5 |

| | | | |
|---|---|---|---|
| 2195 | SPATA13 | SOX2-KLF5 coregulated genes from PMID: 33972779 | SOX2 + KLF5 |
| 2196 | NOVA1 | SOX2-KLF5 coregulated genes from PMID: 33972779 | SOX2 + KLF5 |
| 2197 | BACE2 | SOX2-KLF5 coregulated genes from PMID: 33972779 | SOX2 + KLF5 |
| 2198 | CYYR1 | SOX2-KLF5 coregulated genes from PMID: 33972779 | SOX2 + KLF5 |
| 2199 | FUT8 | SOX2-KLF5 coregulated genes from PMID: 33972779 | SOX2 + KLF5 |
| 2200 | TRMT2B | SOX2-KLF5 coregulated genes from PMID: 33972779 | SOX2 + KLF5 |
| 2201 | SIK1B | SOX2-KLF5 coregulated genes from PMID: 33972779 | SOX2 + KLF5 |
| 2202 | SIK1 | SOX2-KLF5 coregulated genes from PMID: 33972779 | SOX2 + KLF5 |
| 2203 | CNTLN | SOX2-KLF5 coregulated genes from PMID: 33972779 | SOX2 + KLF5 |
| 2204 | LINGO2 | SOX2-KLF5 coregulated genes from PMID: 33972779 | SOX2 + KLF5 |
| 2205 | APBB2 | SOX2-KLF5 coregulated genes from PMID: 33972779 | SOX2 + KLF5 |
| 2206 | SERPINB2 | SOX2-KLF5 coregulated genes from PMID: 33972779 | SOX2 + KLF5 |
| 2207 | SHROOM2 | SOX2-KLF5 coregulated genes from PMID: 33972779 | SOX2 + KLF5 |
| 2208 | GNA14 | SOX2-KLF5 coregulated genes from PMID: 33972779 | SOX2 + KLF5 |
| 2209 | TULP3 | SOX2-KLF5 coregulated genes from PMID: 33972779 | SOX2 + KLF5 |
| 2210 | VOPP1 | SOX2-KLF5 coregulated genes from PMID: 33972779 | SOX2 + KLF5 |
| 2211 | DOCK11 | SOX2-KLF5 coregulated genes from PMID: 33972779 | SOX2 + KLF5 |
| 2212 | SETBP1 | SOX2-KLF5 coregulated genes from PMID: 33972779 | SOX2 + KLF5 |
| 2213 | GCNT3 | SOX2-KLF5 coregulated genes from PMID: 33972779 | SOX2 + KLF5 |
| 2214 | OSBPL3 | SOX2-KLF5 coregulated genes from PMID: 33972779 | SOX2 + KLF5 |
| 2215 | ANXA11 | SOX2-KLF5 coregulated genes from PMID: 33972779 | SOX2 + KLF5 |
| 2216 | GREB1L | SOX2-KLF5 coregulated genes from PMID: 33972779 | SOX2 + KLF5 |
| 2217 | KLF3 | SOX2-KLF5 coregulated genes from PMID: 33972779 | SOX2 + KLF5 |
| 2218 | RSU1 | SOX2-KLF5 coregulated genes from PMID: 33972779 | SOX2 + KLF5 |
| 2219 | ATRNL1 | SOX2-KLF5 coregulated genes from PMID: 33972779 | SOX2 + KLF5 |
| 2220 | B4GALNT3 | SOX2-KLF5 coregulated genes from PMID: 33972779 | SOX2 + KLF5 |
| 2221 | MLLT3 | SOX2-KLF5 coregulated genes from PMID: 33972779 | SOX2 + KLF5 |
| 2222 | UGT8 | SOX2-KLF5 coregulated genes from PMID: 33972779 | SOX2 + KLF5 |
| 2223 | SHROOM3 | SOX2-KLF5 coregulated genes from PMID: 33972779 | SOX2 + KLF5 |
| 2224 | TMTC1 | SOX2-KLF5 coregulated genes from PMID: 33972779 | SOX2 + KLF5 |
| 2225 | DCLK2 | SOX2-KLF5 coregulated genes from PMID: 33972779 | SOX2 + KLF5 |
| 2226 | DUSP14 | SOX2-KLF5 coregulated genes from PMID: 33972779 | SOX2 + KLF5 |
| 2227 | SAMD4A | SOX2-KLF5 coregulated genes from PMID: 33972779 | SOX2 + KLF5 |
| 2228 | NDUFA4 | SOX2-KLF5 coregulated genes from PMID: 33972779 | SOX2 + KLF5 |
| 2229 | TGFBR1 | SOX2-KLF5 coregulated genes from PMID: 33972779 | SOX2 + KLF5 |
| 2230 | APLN | SOX2-KLF5 coregulated genes from PMID: 33972779 | SOX2 + KLF5 |
| 2231 | CALCB | SOX2-KLF5 coregulated genes from PMID: 33972779 | SOX2 + KLF5 |
| 2232 | LMTK2 | SOX2-KLF5 coregulated genes from PMID: 33972779 | SOX2 + KLF5 |
| 2233 | FMN1 | SOX2-KLF5 coregulated genes from PMID: 33972779 | SOX2 + KLF5 |
| 2234 | TNS4 | SOX2-KLF5 coregulated genes from PMID: 33972779 | SOX2 + KLF5 |
| 2235 | TNFAIP2 | SOX2-KLF5 coregulated genes from PMID: 33972779 | SOX2 + KLF5 |
| 2236 | MOXD1 | SOX2-KLF5 coregulated genes from PMID: 33972779 | SOX2 + KLF5 |
| 2237 | MACROD2 | SOX2-KLF5 coregulated genes from PMID: 33972779 | SOX2 + KLF5 |
| 2238 | LNX1 | SOX2-KLF5 coregulated genes from PMID: 33972779 | SOX2 + KLF5 |
| 2239 | ETV1 | SOX2-KLF5 coregulated genes from PMID: 33972779 | SOX2 + KLF5 |
| 2240 | CCNG2 | SOX2-KLF5 coregulated genes from PMID: 33972779 | SOX2 + KLF5 |
| 2241 | ST8SIA6 | SOX2-KLF5 coregulated genes from PMID: 33972779 | SOX2 + KLF5 |
| 2242 | RNF150 | SOX2-KLF5 coregulated genes from PMID: 33972779 | SOX2 + KLF5 |
| 2243 | STARD13 | SOX2-KLF5 coregulated genes from PMID: 33972779 | SOX2 + KLF5 |
| 2244 | IPMK | SOX2-KLF5 coregulated genes from PMID: 33972779 | SOX2 + KLF5 |
| 2245 | NCOA7 | SOX2-KLF5 coregulated genes from PMID: 33972779 | SOX2 + KLF5 |
| 2246 | ATP8B4 | SOX2-KLF5 coregulated genes from PMID: 33972779 | SOX2 + KLF5 |
| 2247 | PLIN2 | SOX2-KLF5 coregulated genes from PMID: 33972779 | SOX2 + KLF5 |
| 2248 | AHNAK | SOX2-KLF5 coregulated genes from PMID: 33972779 | SOX2 + KLF5 |
| 2249 | RAB11FIP2 | SOX2-KLF5 coregulated genes from PMID: 33972779 | SOX2 + KLF5 |
| 2250 | STK38 | SOX2-KLF5 coregulated genes from PMID: 33972779 | SOX2 + KLF5 |
| 2251 | ARMC2 | SOX2-KLF5 coregulated genes from PMID: 33972779 | SOX2 + KLF5 |
| 2252 | AVPI1 | SOX2-KLF5 coregulated genes from PMID: 33972779 | SOX2 + KLF5 |
| 2253 | TNIP3 | SOX2-KLF5 coregulated genes from PMID: 33972779 | SOX2 + KLF5 |
| 2254 | MYO10 | SOX2-KLF5 coregulated genes from PMID: 33972779 | SOX2 + KLF5 |
| 2255 | EPDR1 | SOX2-KLF5 coregulated genes from PMID: 33972779 | SOX2 + KLF5 |

| | | | |
|---|---|---|---|
| 2256 | RASSF6 | SOX2-KLF5 coregulated genes from PMID: 33972779 | SOX2 + KLF5 |
| 2257 | BMP6 | SOX2-KLF5 coregulated genes from PMID: 33972779 | SOX2 + KLF5 |
| 2258 | RUNX2 | SOX2-KLF5 coregulated genes from PMID: 33972779 | SOX2 + KLF5 |
| 2259 | PAPPA | SOX2-KLF5 coregulated genes from PMID: 33972779 | SOX2 + KLF5 |
| 2260 | FOXF2 | SOX2-KLF5 coregulated genes from PMID: 33972779 | SOX2 + KLF5 |
| 2261 | APCDD1 | SOX2-KLF5 coregulated genes from PMID: 33972779 | SOX2 + KLF5 |
| 2262 | IL15 | SOX2-KLF5 coregulated genes from PMID: 33972779 | SOX2 + KLF5 |
| 2263 | FAM171A1 | SOX2-KLF5 coregulated genes from PMID: 33972779 | SOX2 + KLF5 |
| 2264 | GRID2 | SOX2-KLF5 coregulated genes from PMID: 33972779 | SOX2 + KLF5 |
| 2265 | PCDH7 | SOX2-KLF5 coregulated genes from PMID: 33972779 | SOX2 + KLF5 |
| 2266 | FARP1 | SOX2-KLF5 coregulated genes from PMID: 33972779 | SOX2 + KLF5 |
| 2267 | TMPRSS11D | SOX2-KLF5 coregulated genes from PMID: 33972779 | SOX2 + KLF5 |
| 2268 | XPNPEP2 | SOX2-KLF5 coregulated genes from PMID: 33972779 | SOX2 + KLF5 |
| 2269 | CTBP2 | SOX2-KLF5 coregulated genes from PMID: 33972779 | SOX2 + KLF5 |
| 2270 | SUSD1 | SOX2-KLF5 coregulated genes from PMID: 33972779 | SOX2 + KLF5 |
| 2271 | ITGB8 | SOX2-KLF5 coregulated genes from PMID: 33972779 | SOX2 + KLF5 |
| 2272 | MICAL3 | SOX2-KLF5 coregulated genes from PMID: 33972779 | SOX2 + KLF5 |
| 2273 | BCAT1 | SOX2-KLF5 coregulated genes from PMID: 33972779 | SOX2 + KLF5 |
| 2274 | MYOF | SOX2-KLF5 coregulated genes from PMID: 33972779 | SOX2 + KLF5 |
| 2275 | SDCBP2 | SOX2-KLF5 coregulated genes from PMID: 33972779 | SOX2 + KLF5 |
| 2276 | MARCKS | SOX2-KLF5 coregulated genes from PMID: 33972779 | SOX2 + KLF5 |
| 2277 | PC | SOX2-KLF5 coregulated genes from PMID: 33972779 | SOX2 + KLF5 |
| 2278 | PKIB | SOX2-KLF5 coregulated genes from PMID: 33972779 | SOX2 + KLF5 |
| 2279 | ATP8B1 | SOX2-KLF5 coregulated genes from PMID: 33972779 | SOX2 + KLF5 |
| 2280 | APBB1IP | SOX2-KLF5 coregulated genes from PMID: 33972779 | SOX2 + KLF5 |
| 2281 | LAMA3 | SOX2-KLF5 coregulated genes from PMID: 33972779 | SOX2 + KLF5 |
| 2282 | TSPAN14 | SOX2-KLF5 coregulated genes from PMID: 33972779 | SOX2 + KLF5 |
| 2283 | BACH2 | SOX2-KLF5 coregulated genes from PMID: 33972779 | SOX2 + KLF5 |
| 2284 | MKX | SOX2-KLF5 coregulated genes from PMID: 33972779 | SOX2 + KLF5 |
| 2285 | RRP12 | SOX2-KLF5 coregulated genes from PMID: 33972779 | SOX2 + KLF5 |
| 2286 | SULT1B1 | SOX2-KLF5 coregulated genes from PMID: 33972779 | SOX2 + KLF5 |
| 2287 | PIM1 | SOX2-KLF5 coregulated genes from PMID: 33972779 | SOX2 + KLF5 |
| 2288 | AQP11 | SOX2-KLF5 coregulated genes from PMID: 33972779 | SOX2 + KLF5 |
| 2289 | ZNF395 | SOX2-KLF5 coregulated genes from PMID: 33972779 | SOX2 + KLF5 |
| 2290 | DIRAS2 | SOX2-KLF5 coregulated genes from PMID: 33972779 | SOX2 + KLF5 |
| 2291 | RNF24 | SOX2-KLF5 coregulated genes from PMID: 33972779 | SOX2 + KLF5 |
| 2292 | EDN1 | SOX2-KLF5 coregulated genes from PMID: 33972779 | SOX2 + KLF5 |
| 2293 | SH3RF1 | SOX2-KLF5 coregulated genes from PMID: 33972779 | SOX2 + KLF5 |
| 2294 | FNBP1 | SOX2-KLF5 coregulated genes from PMID: 33972779 | SOX2 + KLF5 |
| 2295 | OPHN1 | SOX2-KLF5 coregulated genes from PMID: 33972779 | SOX2 + KLF5 |
| 2296 | AGR3 | SOX2-KLF5 coregulated genes from PMID: 33972779 | SOX2 + KLF5 |
| 2297 | HTRA1 | SOX2-KLF5 coregulated genes from PMID: 33972779 | SOX2 + KLF5 |
| 2298 | GFRA2 | SOX2-KLF5 coregulated genes from PMID: 33972779 | SOX2 + KLF5 |
| 2299 | TNC | SOX2-KLF5 coregulated genes from PMID: 33972779 | SOX2 + KLF5 |
| 2300 | CTSZ | SOX2-KLF5 coregulated genes from PMID: 33972779 | SOX2 + KLF5 |
| 2301 | TSPAN5 | SOX2-KLF5 coregulated genes from PMID: 33972779 | SOX2 + KLF5 |
| 2302 | SERPINB5 | SOX2-KLF5 coregulated genes from PMID: 33972779 | SOX2 + KLF5 |
| 2303 | LPCAT1 | SOX2-KLF5 coregulated genes from PMID: 33972779 | SOX2 + KLF5 |
| 2304 | DCLK1 | SOX2-KLF5 coregulated genes from PMID: 33972779 | SOX2 + KLF5 |
| 2305 | PPP3CA | SOX2-KLF5 coregulated genes from PMID: 33972779 | SOX2 + KLF5 |
| 2306 | SLC7A11 | SOX2-KLF5 coregulated genes from PMID: 33972779 | SOX2 + KLF5 |
| 2307 | TLE1 | SOX2-KLF5 coregulated genes from PMID: 33972779 | SOX2 + KLF5 |
| 2308 | ANK3 | SOX2-KLF5 coregulated genes from PMID: 33972779 | SOX2 + KLF5 |
| 2309 | PIP5K1B | SOX2-KLF5 coregulated genes from PMID: 33972779 | SOX2 + KLF5 |
| 2310 | GRIK1 | SOX2-KLF5 coregulated genes from PMID: 33972779 | SOX2 + KLF5 |
| 2311 | TADA2A | SOX2-KLF5 coregulated genes from PMID: 33972779 | SOX2 + KLF5 |
| 2312 | PARM1 | SOX2-KLF5 coregulated genes from PMID: 33972779 | SOX2 + KLF5 |
| 2313 | PRRG4 | SOX2-KLF5 coregulated genes from PMID: 33972779 | SOX2 + KLF5 |
| 2314 | SOD3 | SOX2-KLF5 coregulated genes from PMID: 33972779 | SOX2 + KLF5 |
| 2315 | FAM83B | SOX2-KLF5 coregulated genes from PMID: 33972779 | SOX2 + KLF5 |
| 2316 | FAM53B | SOX2-KLF5 coregulated genes from PMID: 33972779 | SOX2 + KLF5 |

| | | | |
|---|---|---|---|
| 2317 | RAB11FIP1 | SOX2-KLF5 coregulated genes from PMID: 33972779 | SOX2 + KLF5 |
| 2318 | CYTH3 | SOX2-KLF5 coregulated genes from PMID: 33972779 | SOX2 + KLF5 |
| 2319 | ENKUR | SOX2-KLF5 coregulated genes from PMID: 33972779 | SOX2 + KLF5 |
| 2320 | NRP1 | SOX2-KLF5 coregulated genes from PMID: 33972779 | SOX2 + KLF5 |
| 2321 | OSBPL2 | SOX2-KLF5 coregulated genes from PMID: 33972779 | SOX2 + KLF5 |
| 2322 | SEMA5A | SOX2-KLF5 coregulated genes from PMID: 33972779 | SOX2 + KLF5 |
| 2323 | RCBTB2 | SOX2-KLF5 coregulated genes from PMID: 33972779 | SOX2 + KLF5 |
| 2324 | FOXQ1 | SOX2-KLF5 coregulated genes from PMID: 33972779 | SOX2 + KLF5 |
| 2325 | RAP1GAP2 | SOX2-KLF5 coregulated genes from PMID: 33972779 | SOX2 + KLF5 |
| 2326 | ARL4A | SOX2-KLF5 coregulated genes from PMID: 33972779 | SOX2 + KLF5 |
| 2327 | KLF4 | SOX2-KLF5 coregulated genes from PMID: 33972779 | SOX2 + KLF5 |
| 2328 | CHST3 | SOX2-KLF5 coregulated genes from PMID: 33972779 | SOX2 + KLF5 |
| 2329 | SLC16A7 | SOX2-KLF5 coregulated genes from PMID: 33972779 | SOX2 + KLF5 |
| 2330 | GPR158 | SOX2-KLF5 coregulated genes from PMID: 33972779 | SOX2 + KLF5 |
| 2331 | ARHGAP10 | SOX2-KLF5 coregulated genes from PMID: 33972779 | SOX2 + KLF5 |
| 2332 | ABCC4 | SOX2-KLF5 coregulated genes from PMID: 33972779 | SOX2 + KLF5 |
| 2333 | ZFAND5 | SOX2-KLF5 coregulated genes from PMID: 33972779 | SOX2 + KLF5 |
| 2334 | RIN2 | SOX2-KLF5 coregulated genes from PMID: 33972779 | SOX2 + KLF5 |
| 2335 | DSC3 | SOX2-KLF5 coregulated genes from PMID: 33972779 | SOX2 + KLF5 |
| 2336 | SOX21 | SOX2-KLF5 coregulated genes from PMID: 33972779 | SOX2 + KLF5 |
| 2337 | FRAT2 | SOX2-KLF5 coregulated genes from PMID: 33972779 | SOX2 + KLF5 |
| 2338 | ANXA1 | SOX2-KLF5 coregulated genes from PMID: 33972779 | SOX2 + KLF5 |
| 2339 | FCHSD2 | SOX2-KLF5 coregulated genes from PMID: 33972779 | SOX2 + KLF5 |
| 2340 | TMPRSS11E | SOX2-KLF5 coregulated genes from PMID: 33972779 | SOX2 + KLF5 |
| 2341 | SLITRK6 | SOX2-KLF5 coregulated genes from PMID: 33972779 | SOX2 + KLF5 |
| 2342 | PALLD | SOX2-KLF5 coregulated genes from PMID: 33972779 | SOX2 + KLF5 |
| 2343 | ZNF451 | SOX2-KLF5 coregulated genes from PMID: 33972779 | SOX2 + KLF5 |
| 2344 | PDE1C | SOX2-KLF5 coregulated genes from PMID: 33972779 | SOX2 + KLF5 |
| 2345 | EFNB2 | SOX2-KLF5 coregulated genes from PMID: 33972779 | SOX2 + KLF5 |
| 2346 | COL15A1 | SOX2-KLF5 coregulated genes from PMID: 33972779 | SOX2 + KLF5 |
| 2347 | KLHL32 | SOX2-KLF5 coregulated genes from PMID: 33972779 | SOX2 + KLF5 |
| 2348 | ASAH1 | SOX2-KLF5 coregulated genes from PMID: 33972779 | SOX2 + KLF5 |
| 2349 | CHIC1 | SOX2-KLF5 coregulated genes from PMID: 33972779 | SOX2 + KLF5 |
| 2350 | JUP | SOX2-KLF5 coregulated genes from PMID: 33972779 | SOX2 + KLF5 |
| 2351 | NDFIP2 | SOX2-KLF5 coregulated genes from PMID: 33972779 | SOX2 + KLF5 |
| 2352 | POLM | SOX2-KLF5 coregulated genes from PMID: 33972779 | SOX2 + KLF5 |
| 2353 | TMEM211 | SOX2-KLF5 coregulated genes from PMID: 33972779 | SOX2 + KLF5 |
| 2354 | SLC1A4 | SOX2-KLF5 coregulated genes from PMID: 33972779 | SOX2 + KLF5 |
| 2355 | TSPAN8 | SOX2-KLF5 coregulated genes from PMID: 33972779 | SOX2 + KLF5 |
| 2356 | TRIP10 | SOX2-KLF5 coregulated genes from PMID: 33972779 | SOX2 + KLF5 |
| 2357 | PCDH18 | SOX2-KLF5 coregulated genes from PMID: 33972779 | SOX2 + KLF5 |
| 2358 | PPP1CB | SOX2-KLF5 coregulated genes from PMID: 33972779 | SOX2 + KLF5 |
| 2359 | CA10 | SOX2-KLF5 coregulated genes from PMID: 33972779 | SOX2 + KLF5 |
| 2360 | SYNE2 | SOX2-KLF5 coregulated genes from PMID: 33972779 | SOX2 + KLF5 |
| 2361 | CECR2 | SOX2-KLF5 coregulated genes from PMID: 33972779 | SOX2 + KLF5 |
| 2362 | DOCK9 | SOX2-KLF5 coregulated genes from PMID: 33972779 | SOX2 + KLF5 |
| 2363 | SULF1 | SOX2-KLF5 coregulated genes from PMID: 33972779 | SOX2 + KLF5 |
| 2364 | WDR72 | SOX2-KLF5 coregulated genes from PMID: 33972779 | SOX2 + KLF5 |
| 2365 | RAP2A | SOX2-KLF5 coregulated genes from PMID: 33972779 | SOX2 + KLF5 |
| 2366 | KIRREL3 | SOX2-KLF5 coregulated genes from PMID: 33972779 | SOX2 + KLF5 |
| 2367 | EXOC4 | SOX2-KLF5 coregulated genes from PMID: 33972779 | SOX2 + KLF5 |
| 2368 | LY75 | SOX2-KLF5 coregulated genes from PMID: 33972779 | SOX2 + KLF5 |
| 2369 | LAMA4 | SOX2-KLF5 coregulated genes from PMID: 33972779 | SOX2 + KLF5 |
| 2370 | IGSF11 | SOX2-KLF5 coregulated genes from PMID: 33972779 | SOX2 + KLF5 |
| 2371 | FRAS1 | SOX2-KLF5 coregulated genes from PMID: 33972779 | SOX2 + KLF5 |
| 2372 | DENND1A | SOX2-KLF5 coregulated genes from PMID: 33972779 | SOX2 + KLF5 |
| 2373 | MAPK10 | SOX2-KLF5 coregulated genes from PMID: 33972779 | SOX2 + KLF5 |
| 2374 | TGM3 | SOX2-KLF5 coregulated genes from PMID: 33972779 | SOX2 + KLF5 |
| 2375 | TMPRSS4 | SOX2-KLF5 coregulated genes from PMID: 33972779 | SOX2 + KLF5 |
| 2376 | SNRPN | SOX2-KLF5 coregulated genes from PMID: 33972779 | SOX2 + KLF5 |
| 2377 | DPYSL3 | SOX2-KLF5 coregulated genes from PMID: 33972779 | SOX2 + KLF5 |

| | | | |
|---|---|---|---|
| 2378 | C6orf136 | SOX2-KLF5 coregulated genes from PMID: 33972779 | SOX2 + KLF5 |
| 2379 | BNC2 | SOX2-KLF5 coregulated genes from PMID: 33972779 | SOX2 + KLF5 |
| 2380 | TANK | SOX2-KLF5 coregulated genes from PMID: 33972779 | SOX2 + KLF5 |
| 2381 | TANC1 | SOX2-KLF5 coregulated genes from PMID: 33972779 | SOX2 + KLF5 |
| 2382 | NOL10 | SOX2-KLF5 coregulated genes from PMID: 33972779 | SOX2 + KLF5 |
| 2383 | AKAP12 | SOX2-KLF5 coregulated genes from PMID: 33972779 | SOX2 + KLF5 |
| 2384 | SEMA6D | SOX2-KLF5 coregulated genes from PMID: 33972779 | SOX2 + KLF5 |
| 2385 | RIMS2 | SOX2-KLF5 coregulated genes from PMID: 33972779 | SOX2 + KLF5 |
| 2386 | GDNF | SOX2-KLF5 coregulated genes from PMID: 33972779 | SOX2 + KLF5 |
| 2387 | FIGN | SOX2-KLF5 coregulated genes from PMID: 33972779 | SOX2 + KLF5 |
| 2388 | WFDC1 | SOX2-KLF5 coregulated genes from PMID: 33972779 | SOX2 + KLF5 |
| 2389 | SIL1 | SOX2-KLF5 coregulated genes from PMID: 33972779 | SOX2 + KLF5 |
| 2390 | BARX2 | SOX2-KLF5 coregulated genes from PMID: 33972779 | SOX2 + KLF5 |
| 2391 | WNT7A | SOX2-KLF5 coregulated genes from PMID: 33972779 | SOX2 + KLF5 |
| 2392 | MACC1 | SOX2-KLF5 coregulated genes from PMID: 33972779 | SOX2 + KLF5 |
| 2393 | CITED2 | SOX2-KLF5 coregulated genes from PMID: 33972779 | SOX2 + KLF5 |
| 2394 | FBXO16 | SOX2-KLF5 coregulated genes from PMID: 33972779 | SOX2 + KLF5 |
| 2395 | TGFBI | SOX2-KLF5 coregulated genes from PMID: 33972779 | SOX2 + KLF5 |
| 2396 | ASB1 | SOX2-KLF5 coregulated genes from PMID: 33972779 | SOX2 + KLF5 |
| 2397 | ATP8A1 | SOX2-KLF5 coregulated genes from PMID: 33972779 | SOX2 + KLF5 |
| 2398 | CCDC85A | SOX2-KLF5 coregulated genes from PMID: 33972779 | SOX2 + KLF5 |
| 2399 | OSGIN1 | SOX2-KLF5 coregulated genes from PMID: 33972779 | SOX2 + KLF5 |
| 2400 | WNK2 | SOX2-KLF5 coregulated genes from PMID: 33972779 | SOX2 + KLF5 |
| 2401 | THUMPD2 | SOX2-KLF5 coregulated genes from PMID: 33972779 | SOX2 + KLF5 |
| 2402 | SLC8A1 | SOX2-KLF5 coregulated genes from PMID: 33972779 | SOX2 + KLF5 |
| 2403 | N4BP1 | SOX2-KLF5 coregulated genes from PMID: 33972779 | SOX2 + KLF5 |
| 2404 | PRKCH | SOX2-KLF5 coregulated genes from PMID: 33972779 | SOX2 + KLF5 |
| 2405 | PLS3 | SOX2-KLF5 coregulated genes from PMID: 33972779 | SOX2 + KLF5 |
| 2406 | PPIF | SOX2-KLF5 coregulated genes from PMID: 33972779 | SOX2 + KLF5 |
| 2407 | AIG1 | SOX2-KLF5 coregulated genes from PMID: 33972779 | SOX2 + KLF5 |
| 2408 | VCL | SOX2-KLF5 coregulated genes from PMID: 33972779 | SOX2 + KLF5 |
| 2409 | NRXN3 | SOX2-KLF5 coregulated genes from PMID: 33972779 | SOX2 + KLF5 |
| 2410 | GSS | SOX2-KLF5 coregulated genes from PMID: 33972779 | SOX2 + KLF5 |
| 2411 | ARHGAP26 | SOX2-KLF5 coregulated genes from PMID: 33972779 | SOX2 + KLF5 |
| 2412 | RPS6KA3 | SOX2-KLF5 coregulated genes from PMID: 33972779 | SOX2 + KLF5 |
| 2413 | SLC16A9 | SOX2-KLF5 coregulated genes from PMID: 33972779 | SOX2 + KLF5 |
| 2414 | RGS9 | SOX2-KLF5 coregulated genes from PMID: 33972779 | SOX2 + KLF5 |
| 2415 | RERE | SOX2-KLF5 coregulated genes from PMID: 33972779 | SOX2 + KLF5 |
| 2416 | FBLN2 | SOX2-KLF5 coregulated genes from PMID: 33972779 | SOX2 + KLF5 |
| 2417 | MEIS1 | SOX2-KLF5 coregulated genes from PMID: 33972779 | SOX2 + KLF5 |
| 2418 | LY6E | SOX2-KLF5 coregulated genes from PMID: 33972779 | SOX2 + KLF5 |
| 2419 | COL8A1 | SOX2-KLF5 coregulated genes from PMID: 33972779 | SOX2 + KLF5 |
| 2420 | KCMF1 | SOX2-KLF5 coregulated genes from PMID: 33972779 | SOX2 + KLF5 |
| 2421 | PARP12 | SOX2-KLF5 coregulated genes from PMID: 33972779 | SOX2 + KLF5 |
| 2422 | SLCO5A1 | SOX2-KLF5 coregulated genes from PMID: 33972779 | SOX2 + KLF5 |
| 2423 | SMPD3 | SOX2-KLF5 coregulated genes from PMID: 33972779 | SOX2 + KLF5 |
| 2424 | SLC4A4 | SOX2-KLF5 coregulated genes from PMID: 33972779 | SOX2 + KLF5 |
| 2425 | FHL2 | SOX2-KLF5 coregulated genes from PMID: 33972779 | SOX2 + KLF5 |
| 2426 | SLC44A3 | SOX2-KLF5 coregulated genes from PMID: 33972779 | SOX2 + KLF5 |
| 2427 | SYN2 | SOX2-KLF5 coregulated genes from PMID: 33972779 | SOX2 + KLF5 |
| 2428 | GLI3 | SOX2-KLF5 coregulated genes from PMID: 33972779 | SOX2 + KLF5 |
| 2429 | TRIM24 | SOX2-KLF5 coregulated genes from PMID: 33972779 | SOX2 + KLF5 |
| 2430 | ENPP2 | SOX2-KLF5 coregulated genes from PMID: 33972779 | SOX2 + KLF5 |
| 2431 | RFX3 | SOX2-KLF5 coregulated genes from PMID: 33972779 | SOX2 + KLF5 |
| 2432 | SCARA5 | SOX2-KLF5 coregulated genes from PMID: 33972779 | SOX2 + KLF5 |
| 2433 | EMB | SOX2-KLF5 coregulated genes from PMID: 33972779 | SOX2 + KLF5 |
| 2434 | MMP2 | SOX2-KLF5 coregulated genes from PMID: 33972779 | SOX2 + KLF5 |
| 2435 | PGM2L1 | SOX2-KLF5 coregulated genes from PMID: 33972779 | SOX2 + KLF5 |
| 2436 | MGAT4A | SOX2-KLF5 coregulated genes from PMID: 33972779 | SOX2 + KLF5 |
| 2437 | FSD1L | SOX2-KLF5 coregulated genes from PMID: 33972779 | SOX2 + KLF5 |
| 2438 | PLEKHH1 | SOX2-KLF5 coregulated genes from PMID: 33972779 | SOX2 + KLF5 |

| | | | |
|---|---|---|---|
| 2439 | FLRT2 | SOX2-KLF5 coregulated genes from PMID: 33972779 | SOX2 + KLF5 |
| 2440 | AMOT | SOX2-KLF5 coregulated genes from PMID: 33972779 | SOX2 + KLF5 |
| 2441 | FAM81A | SOX2-KLF5 coregulated genes from PMID: 33972779 | SOX2 + KLF5 |
| 2442 | DNMBP | SOX2-KLF5 coregulated genes from PMID: 33972779 | SOX2 + KLF5 |
| 2443 | EPHB6 | SOX2-KLF5 coregulated genes from PMID: 33972779 | SOX2 + KLF5 |
| 2444 | STXBP1 | SOX2-KLF5 coregulated genes from PMID: 33972779 | SOX2 + KLF5 |
| 2445 | NEBL | SOX2-KLF5 coregulated genes from PMID: 33972779 | SOX2 + KLF5 |
| 2446 | IGF2R | SOX2-KLF5 coregulated genes from PMID: 33972779 | SOX2 + KLF5 |
| 2447 | ZFHX3 | SOX2-KLF5 coregulated genes from PMID: 33972779 | SOX2 + KLF5 |
| 2448 | TLE4 | SOX2-KLF5 coregulated genes from PMID: 33972779 | SOX2 + KLF5 |
| 2449 | GJB5 | SOX2-KLF5 coregulated genes from PMID: 33972779 | SOX2 + KLF5 |
| 2450 | KLF12 | SOX2-KLF5 coregulated genes from PMID: 33972779 | SOX2 + KLF5 |
| 2451 | RAI14 | SOX2-KLF5 coregulated genes from PMID: 33972779 | SOX2 + KLF5 |
| 2452 | PBRM1 | SOX2-KLF5 coregulated genes from PMID: 33972779 | SOX2 + KLF5 |
| 2453 | ILVBL | SOX2-KLF5 coregulated genes from PMID: 33972779 | SOX2 + KLF5 |
| 2454 | PCTP | SOX2-KLF5 coregulated genes from PMID: 33972779 | SOX2 + KLF5 |
| 2455 | SOX5 | SOX2-KLF5 coregulated genes from PMID: 33972779 | SOX2 + KLF5 |
| 2456 | NCOR2 | SOX2-KLF5 coregulated genes from PMID: 33972779 | SOX2 + KLF5 |
| 2457 | RAPGEF5 | SOX2-KLF5 coregulated genes from PMID: 33972779 | SOX2 + KLF5 |
| 2458 | PHACTR4 | SOX2-KLF5 coregulated genes from PMID: 33972779 | SOX2 + KLF5 |
| 2459 | ZNF365 | SOX2-KLF5 coregulated genes from PMID: 33972779 | SOX2 + KLF5 |
| 2460 | ARFGAP3 | SOX2-KLF5 coregulated genes from PMID: 33972779 | SOX2 + KLF5 |
| 2461 | PACS1 | SOX2-KLF5 coregulated genes from PMID: 33972779 | SOX2 + KLF5 |
| 2462 | COL4A6 | SOX2-KLF5 coregulated genes from PMID: 33972779 | SOX2 + KLF5 |
| 2463 | IL20RB | SOX2-KLF5 coregulated genes from PMID: 33972779 | SOX2 + KLF5 |
| 2464 | MREG | SOX2-KLF5 coregulated genes from PMID: 33972779 | SOX2 + KLF5 |
| 2465 | TMEM200A | SOX2-KLF5 coregulated genes from PMID: 33972779 | SOX2 + KLF5 |
| 2466 | PKD2 | SOX2-KLF5 coregulated genes from PMID: 33972779 | SOX2 + KLF5 |
| 2467 | SOCS2 | SOX2-KLF5 coregulated genes from PMID: 33972779 | SOX2 + KLF5 |
| 2468 | MYH9 | SOX2-KLF5 coregulated genes from PMID: 33972779 | SOX2 + KLF5 |
| 2469 | ETS1 | SOX2-KLF5 coregulated genes from PMID: 33972779 | SOX2 + KLF5 |
| 2470 | CDK14 | SOX2-KLF5 coregulated genes from PMID: 33972779 | SOX2 + KLF5 |
| 2471 | NFKBIA | SOX2-KLF5 coregulated genes from PMID: 33972779 | SOX2 + KLF5 |
| 2472 | GUCY1A2 | SOX2-KLF5 coregulated genes from PMID: 33972779 | SOX2 + KLF5 |
| 2473 | DTNBP1 | SOX2-KLF5 coregulated genes from PMID: 33972779 | SOX2 + KLF5 |
| 2474 | MAL2 | SOX2-KLF5 coregulated genes from PMID: 33972779 | SOX2 + KLF5 |
| 2475 | PDZRN3 | SOX2-KLF5 coregulated genes from PMID: 33972779 | SOX2 + KLF5 |
| 2476 | MRAS | SOX2-KLF5 coregulated genes from PMID: 33972779 | SOX2 + KLF5 |
| 2477 | B3GAT1 | SOX2-KLF5 coregulated genes from PMID: 33972779 | SOX2 + KLF5 |
| 2478 | ADD2 | SOX2-KLF5 coregulated genes from PMID: 33972779 | SOX2 + KLF5 |
| 2479 | SGMS1 | SOX2-KLF5 coregulated genes from PMID: 33972779 | SOX2 + KLF5 |
| 2480 | IL1A | SOX2-KLF5 coregulated genes from PMID: 33972779 | SOX2 + KLF5 |
| 2481 | RANBP17 | SOX2-KLF5 coregulated genes from PMID: 33972779 | SOX2 + KLF5 |
| 2482 | NUDT4 | SOX2-KLF5 coregulated genes from PMID: 33972779 | SOX2 + KLF5 |
| 2483 | TXNDC12 | SOX2-KLF5 coregulated genes from PMID: 33972779 | SOX2 + KLF5 |
| 2484 | PEBP1 | SOX2-KLF5 coregulated genes from PMID: 33972779 | SOX2 + KLF5 |
| 2485 | TMC1 | SOX2-KLF5 coregulated genes from PMID: 33972779 | SOX2 + KLF5 |
| 2486 | RBMS3 | SOX2-KLF5 coregulated genes from PMID: 33972779 | SOX2 + KLF5 |
| 2487 | LIPT2 | SOX2-KLF5 coregulated genes from PMID: 33972779 | SOX2 + KLF5 |
| 2488 | CDH6 | SOX2-KLF5 coregulated genes from PMID: 33972779 | SOX2 + KLF5 |
| 2489 | FAM3C | SOX2-KLF5 coregulated genes from PMID: 33972779 | SOX2 + KLF5 |
| 2490 | GHR | SOX2-KLF5 coregulated genes from PMID: 33972779 | SOX2 + KLF5 |
| 2491 | ALDH3A1 | SOX2-KLF5 coregulated genes from PMID: 33972779 | SOX2 + KLF5 |
| 2492 | EYA1 | SOX2-KLF5 coregulated genes from PMID: 33972779 | SOX2 + KLF5 |
| 2493 | LAMB1 | SOX2-KLF5 coregulated genes from PMID: 33972779 | SOX2 + KLF5 |
| 2494 | CDH11 | SOX2-KLF5 coregulated genes from PMID: 33972779 | SOX2 + KLF5 |
| 2495 | ESYT3 | SOX2-KLF5 coregulated genes from PMID: 33972779 | SOX2 + KLF5 |
| 2496 | BAIAP2L1 | SOX2-KLF5 coregulated genes from PMID: 33972779 | SOX2 + KLF5 |
| 2497 | CCDC88C | SOX2-KLF5 coregulated genes from PMID: 33972779 | SOX2 + KLF5 |
| 2498 | HMGCLL1 | SOX2-KLF5 coregulated genes from PMID: 33972779 | SOX2 + KLF5 |
| 2499 | CMYA5 | SOX2-KLF5 coregulated genes from PMID: 33972779 | SOX2 + KLF5 |

| | | | |
|---|---|---|---|
| 2500 | TOX3 | SOX2-KLF5 coregulated genes from PMID: 33972779 | SOX2 + KLF5 |
| 2501 | HS6ST1 | SOX2-KLF5 coregulated genes from PMID: 33972779 | SOX2 + KLF5 |
| 2502 | MMP16 | SOX2-KLF5 coregulated genes from PMID: 33972779 | SOX2 + KLF5 |
| 2503 | DMD | SOX2-KLF5 coregulated genes from PMID: 33972779 | SOX2 + KLF5 |
| 2504 | ARID5B | SOX2-KLF5 coregulated genes from PMID: 33972779 | SOX2 + KLF5 |
| 2505 | SYT17 | SOX2-KLF5 coregulated genes from PMID: 33972779 | SOX2 + KLF5 |
| 2506 | PI15 | SOX2-KLF5 coregulated genes from PMID: 33972779 | SOX2 + KLF5 |
| 2507 | EEA1 | SOX2-KLF5 coregulated genes from PMID: 33972779 | SOX2 + KLF5 |
| 2508 | PRKD1 | SOX2-KLF5 coregulated genes from PMID: 33972779 | SOX2 + KLF5 |
| 2509 | MCC | SOX2-KLF5 coregulated genes from PMID: 33972779 | SOX2 + KLF5 |
| 2510 | TRIM2 | SOX2-KLF5 coregulated genes from PMID: 33972779 | SOX2 + KLF5 |
| 2511 | B4GALNT4 | SOX2-KLF5 coregulated genes from PMID: 33972779 | SOX2 + KLF5 |
| 2512 | MPPED2 | SOX2-KLF5 coregulated genes from PMID: 33972779 | SOX2 + KLF5 |
| 2513 | GIPC2 | SOX2-KLF5 coregulated genes from PMID: 33972779 | SOX2 + KLF5 |
| 2514 | TEK | SOX2-KLF5 coregulated genes from PMID: 33972779 | SOX2 + KLF5 |
| 2515 | ROBO2 | SOX2-KLF5 coregulated genes from PMID: 33972779 | SOX2 + KLF5 |
| 2516 | TJP2 | SOX2-KLF5 coregulated genes from PMID: 33972779 | SOX2 + KLF5 |
| 2517 | KCNN2 | SOX2-KLF5 coregulated genes from PMID: 33972779 | SOX2 + KLF5 |
| 2518 | NEGR1 | SOX2-KLF5 coregulated genes from PMID: 33972779 | SOX2 + KLF5 |
| 2519 | GPHN | SOX2-KLF5 coregulated genes from PMID: 33972779 | SOX2 + KLF5 |
| 2520 | RNF19A | SOX2-KLF5 coregulated genes from PMID: 33972779 | SOX2 + KLF5 |
| 2521 | FAM181B | SOX2-KLF5 coregulated genes from PMID: 33972779 | SOX2 + KLF5 |
| 2522 | TM4SF1 | SOX2-KLF5 coregulated genes from PMID: 33972779 | SOX2 + KLF5 |
| 2523 | UGCG | SOX2-KLF5 coregulated genes from PMID: 33972779 | SOX2 + KLF5 |
| 2524 | ATP10B | SOX2-KLF5 coregulated genes from PMID: 33972779 | SOX2 + KLF5 |
| 2525 | FKBP5 | SOX2-KLF5 coregulated genes from PMID: 33972779 | SOX2 + KLF5 |
| 2526 | TGM7 | SOX2-KLF5 coregulated genes from PMID: 33972779 | SOX2 + KLF5 |
| 2527 | PLA1A | SOX2-KLF5 coregulated genes from PMID: 33972779 | SOX2 + KLF5 |
| 2528 | NHLRC2 | SOX2-KLF5 coregulated genes from PMID: 33972779 | SOX2 + KLF5 |
| 2529 | NDUFS4 | SOX2-KLF5 coregulated genes from PMID: 33972779 | SOX2 + KLF5 |
| 2530 | AGPAT4 | SOX2-KLF5 coregulated genes from PMID: 33972779 | SOX2 + KLF5 |
| 2531 | TULP4 | SOX2-KLF5 coregulated genes from PMID: 33972779 | SOX2 + KLF5 |
| 2532 | EHF | SOX2-KLF5 coregulated genes from PMID: 33972779 | SOX2 + KLF5 |
| 2533 | SLC9A2 | SOX2-KLF5 coregulated genes from PMID: 33972779 | SOX2 + KLF5 |
| 2534 | TNFSF15 | SOX2-KLF5 coregulated genes from PMID: 33972779 | SOX2 + KLF5 |
| 2535 | GRM8 | SOX2-KLF5 coregulated genes from PMID: 33972779 | SOX2 + KLF5 |
| 2536 | NUDT7 | SOX2-KLF5 coregulated genes from PMID: 33972779 | SOX2 + KLF5 |
| 2537 | PARD3 | SOX2-KLF5 coregulated genes from PMID: 33972779 | SOX2 + KLF5 |
| 2538 | SPRY1 | SOX2-KLF5 coregulated genes from PMID: 33972779 | SOX2 + KLF5 |
| 2539 | EPHA3 | SOX2-KLF5 coregulated genes from PMID: 33972779 | SOX2 + KLF5 |
| 2540 | PLB1 | SOX2-KLF5 coregulated genes from PMID: 33972779 | SOX2 + KLF5 |
| 2541 | ANKH | SOX2-KLF5 coregulated genes from PMID: 33972779 | SOX2 + KLF5 |
| 2542 | DNAJC6 | SOX2-KLF5 coregulated genes from PMID: 33972779 | SOX2 + KLF5 |
| 2543 | VEGFC | SOX2-KLF5 coregulated genes from PMID: 33972779 | SOX2 + KLF5 |
| 2544 | QKI | SOX2-KLF5 coregulated genes from PMID: 33972779 | SOX2 + KLF5 |
| 2545 | ADM | SOX2-KLF5 coregulated genes from PMID: 33972779 | SOX2 + KLF5 |
| 2546 | PODXL | SOX2-KLF5 coregulated genes from PMID: 33972779 | SOX2 + KLF5 |
| 2547 | PPP4R4 | SOX2-KLF5 coregulated genes from PMID: 33972779 | SOX2 + KLF5 |
| 2548 | EPCAM | SOX2-KLF5 coregulated genes from PMID: 33972779 | SOX2 + KLF5 |
| 2549 | PTER | SOX2-KLF5 coregulated genes from PMID: 33972779 | SOX2 + KLF5 |
| 2550 | CEP68 | SOX2-KLF5 coregulated genes from PMID: 33972779 | SOX2 + KLF5 |
| 2551 | CRADD | SOX2-KLF5 coregulated genes from PMID: 33972779 | SOX2 + KLF5 |
| 2552 | PTGR1 | SOX2-KLF5 coregulated genes from PMID: 33972779 | SOX2 + KLF5 |
| 2553 | EPGN | SOX2-KLF5 coregulated genes from PMID: 33972779 | SOX2 + KLF5 |
| 2554 | BTG1 | SOX2-KLF5 coregulated genes from PMID: 33972779 | SOX2 + KLF5 |
| 2555 | ARMCX1 | SOX2-KLF5 coregulated genes from PMID: 33972779 | SOX2 + KLF5 |
| 2556 | KLHL8 | SOX2-KLF5 coregulated genes from PMID: 33972779 | SOX2 + KLF5 |
| 2557 | ALDH5A1 | SOX2-KLF5 coregulated genes from PMID: 33972779 | SOX2 + KLF5 |
| 2558 | CAMK4 | SOX2-KLF5 coregulated genes from PMID: 33972779 | SOX2 + KLF5 |
| 2559 | ARG1 | SOX2-KLF5 coregulated genes from PMID: 33972779 | SOX2 + KLF5 |
| 2560 | NEURL1B | SOX2-KLF5 coregulated genes from PMID: 33972779 | SOX2 + KLF5 |

| | | | |
|---|---|---|---|
| 2561 | FAM110C | SOX2-KLF5 coregulated genes from PMID: 33972779 | SOX2 + KLF5 |
| 2562 | SLC25A25 | SOX2-KLF5 coregulated genes from PMID: 33972779 | SOX2 + KLF5 |
| 2563 | ANKRD6 | SOX2-KLF5 coregulated genes from PMID: 33972779 | SOX2 + KLF5 |
| 2564 | MAP3K8 | SOX2-KLF5 coregulated genes from PMID: 33972779 | SOX2 + KLF5 |
| 2565 | ALCAM | SOX2-KLF5 coregulated genes from PMID: 33972779 | SOX2 + KLF5 |
| 2566 | JPH1 | SOX2-KLF5 coregulated genes from PMID: 33972779 | SOX2 + KLF5 |
| 2567 | DNAJC21 | SOX2-KLF5 coregulated genes from PMID: 33972779 | SOX2 + KLF5 |
| 2568 | SESTD1 | SOX2-KLF5 coregulated genes from PMID: 33972779 | SOX2 + KLF5 |
| 2569 | SRL | SOX2-KLF5 coregulated genes from PMID: 33972779 | SOX2 + KLF5 |
| 2570 | ARHGEF12 | SOX2-KLF5 coregulated genes from PMID: 33972779 | SOX2 + KLF5 |
| 2571 | SORBS2 | SOX2-KLF5 coregulated genes from PMID: 33972779 | SOX2 + KLF5 |
| 2572 | EFEMP1 | SOX2-KLF5 coregulated genes from PMID: 33972779 | SOX2 + KLF5 |
| 2573 | MBNL3 | SOX2-KLF5 coregulated genes from PMID: 33972779 | SOX2 + KLF5 |
| 2574 | KHDRBS3 | SOX2-KLF5 coregulated genes from PMID: 33972779 | SOX2 + KLF5 |
| 2575 | MFAP3L | SOX2-KLF5 coregulated genes from PMID: 33972779 | SOX2 + KLF5 |
| 2576 | MEGF9 | SOX2-KLF5 coregulated genes from PMID: 33972779 | SOX2 + KLF5 |
| 2577 | GLIS3 | SOX2-KLF5 coregulated genes from PMID: 33972779 | SOX2 + KLF5 |
| 2578 | PAIP2B | SOX2-KLF5 coregulated genes from PMID: 33972779 | SOX2 + KLF5 |
| 2579 | CX3CL1 | SOX2-KLF5 coregulated genes from PMID: 33972779 | SOX2 + KLF5 |
| 2580 | PFN2 | SOX2-KLF5 coregulated genes from PMID: 33972779 | SOX2 + KLF5 |
| 2581 | MBOAT2 | SOX2-KLF5 coregulated genes from PMID: 33972779 | SOX2 + KLF5 |
| 2582 | TBC1D30 | SOX2-KLF5 coregulated genes from PMID: 33972779 | SOX2 + KLF5 |
| 2583 | NUAK2 | SOX2-KLF5 coregulated genes from PMID: 33972779 | SOX2 + KLF5 |
| 2584 | IL6R | SOX2-KLF5 coregulated genes from PMID: 33972779 | SOX2 + KLF5 |
| 2585 | B3GNT2 | SOX2-KLF5 coregulated genes from PMID: 33972779 | SOX2 + KLF5 |
| 2586 | PBX1 | SOX2-KLF5 coregulated genes from PMID: 33972779 | SOX2 + KLF5 |
| 2587 | TBCA | SOX2-KLF5 coregulated genes from PMID: 33972779 | SOX2 + KLF5 |
| 2588 | SETD5 | SOX2-KLF5 coregulated genes from PMID: 33972779 | SOX2 + KLF5 |
| 2589 | KLHL29 | SOX2-KLF5 coregulated genes from PMID: 33972779 | SOX2 + KLF5 |
| 2590 | DGKG | SOX2-KLF5 coregulated genes from PMID: 33972779 | SOX2 + KLF5 |
| 2591 | LITAF | SOX2-KLF5 coregulated genes from PMID: 33972779 | SOX2 + KLF5 |
| 2592 | ST14 | SOX2-KLF5 coregulated genes from PMID: 33972779 | SOX2 + KLF5 |
| 2593 | RBPJ | SOX2-KLF5 coregulated genes from PMID: 33972779 | SOX2 + KLF5 |
| 2594 | ADAM12 | SOX2-KLF5 coregulated genes from PMID: 33972779 | SOX2 + KLF5 |
| 2595 | SLC4A3 | SOX2-KLF5 coregulated genes from PMID: 33972779 | SOX2 + KLF5 |
| 2596 | CCDC141 | SOX2-KLF5 coregulated genes from PMID: 33972779 | SOX2 + KLF5 |
| 2597 | IDH2 | SOX2-KLF5 coregulated genes from PMID: 33972779 | SOX2 + KLF5 |
| 2598 | TNRC18 | SOX2-KLF5 coregulated genes from PMID: 33972779 | SOX2 + KLF5 |
| 2599 | CDH13 | SOX2-KLF5 coregulated genes from PMID: 33972779 | SOX2 + KLF5 |
| 2600 | MPZL1 | SOX2-KLF5 coregulated genes from PMID: 33972779 | SOX2 + KLF5 |
| 2601 | TXNRD1 | SOX2-KLF5 coregulated genes from PMID: 33972779 | SOX2 + KLF5 |
| 2602 | TANC2 | SOX2-KLF5 coregulated genes from PMID: 33972779 | SOX2 + KLF5 |
| 2603 | RGS4 | SOX2-KLF5 coregulated genes from PMID: 33972779 | SOX2 + KLF5 |
| 2604 | IL10 | SOX2-KLF5 coregulated genes from PMID: 33972779 | SOX2 + KLF5 |
| 2605 | KRT19 | SOX2-KLF5 coregulated genes from PMID: 33972779 | SOX2 + KLF5 |
| 2606 | ARHGAP18 | SOX2-KLF5 coregulated genes from PMID: 33972779 | SOX2 + KLF5 |
| 2607 | MEGF11 | SOX2-KLF5 coregulated genes from PMID: 33972779 | SOX2 + KLF5 |
| 2608 | KIRREL1 | SOX2-KLF5 coregulated genes from PMID: 33972779 | SOX2 + KLF5 |
| 2609 | SLC13A2 | SOX2-KLF5 coregulated genes from PMID: 33972779 | SOX2 + KLF5 |
| 2610 | ALOX12 | SOX2-KLF5 coregulated genes from PMID: 33972779 | SOX2 + KLF5 |
| 2611 | SHISA6 | SOX2-KLF5 coregulated genes from PMID: 33972779 | SOX2 + KLF5 |
| 2612 | SMOC2 | SOX2-KLF5 coregulated genes from PMID: 33972779 | SOX2 + KLF5 |
| 2613 | NRXN1 | SOX2-KLF5 coregulated genes from PMID: 33972779 | SOX2 + KLF5 |
| 2614 | RGL1 | SOX2-KLF5 coregulated genes from PMID: 33972779 | SOX2 + KLF5 |
| 2615 | SOX6 | SOX2-KLF5 coregulated genes from PMID: 33972779 | SOX2 + KLF5 |
| 2616 | CTNND2 | SOX2-KLF5 coregulated genes from PMID: 33972779 | SOX2 + KLF5 |
| 2617 | BLNK | SOX2-KLF5 coregulated genes from PMID: 33972779 | SOX2 + KLF5 |
| 2618 | RARRES1 | SOX2-KLF5 coregulated genes from PMID: 33972779 | SOX2 + KLF5 |
| 2619 | NMNAT2 | SOX2-KLF5 coregulated genes from PMID: 33972779 | SOX2 + KLF5 |
| 2620 | PPL | SOX2-KLF5 coregulated genes from PMID: 33972779 | SOX2 + KLF5 |
| 2621 | CADPS2 | SOX2-KLF5 coregulated genes from PMID: 33972779 | SOX2 + KLF5 |

| | | | |
|---|---|---|---|
| 2622 | EFNA5 | SOX2-KLF5 coregulated genes from PMID: 33972779 | SOX2 + KLF5 |
| 2623 | UTRN | SOX2-KLF5 coregulated genes from PMID: 33972779 | SOX2 + KLF5 |
| 2624 | PKP1 | SOX2-KLF5 coregulated genes from PMID: 33972779 | SOX2 + KLF5 |
| 2625 | KCNJ2 | SOX2-KLF5 coregulated genes from PMID: 33972779 | SOX2 + KLF5 |
| 2626 | NUDT4B | SOX2-KLF5 coregulated genes from PMID: 33972779 | SOX2 + KLF5 |
| 2627 | ATP1B1 | SOX2-KLF5 coregulated genes from PMID: 33972779 | SOX2 + KLF5 |
| 2628 | FAM43A | SOX2-KLF5 coregulated genes from PMID: 33972779 | SOX2 + KLF5 |
| 2629 | ADRB1 | SOX2-KLF5 coregulated genes from PMID: 33972779 | SOX2 + KLF5 |
| 2630 | LYPD6B | SOX2-KLF5 coregulated genes from PMID: 33972779 | SOX2 + KLF5 |
| 2631 | TNK2 | SOX2-KLF5 coregulated genes from PMID: 33972779 | SOX2 + KLF5 |
| 2632 | LY96 | SOX2-KLF5 coregulated genes from PMID: 33972779 | SOX2 + KLF5 |
| 2633 | OTOP1 | SOX2-KLF5 coregulated genes from PMID: 33972779 | SOX2 + KLF5 |
| 2634 | HDAC9 | SOX2-KLF5 coregulated genes from PMID: 33972779 | SOX2 + KLF5 |
| 2635 | IQGAP1 | SOX2-KLF5 coregulated genes from PMID: 33972779 | SOX2 + KLF5 |
| 2636 | PLD1 | SOX2-KLF5 coregulated genes from PMID: 33972779 | SOX2 + KLF5 |
| 2637 | SMPDL3B | SOX2-KLF5 coregulated genes from PMID: 33972779 | SOX2 + KLF5 |
| 2638 | KCNK1 | SOX2-KLF5 coregulated genes from PMID: 33972779 | SOX2 + KLF5 |
| 2639 | IRAK2 | SOX2-KLF5 coregulated genes from PMID: 33972779 | SOX2 + KLF5 |
| 2640 | TMEM161B | SOX2-KLF5 coregulated genes from PMID: 33972779 | SOX2 + KLF5 |
| 2641 | RC3H1 | SOX2-KLF5 coregulated genes from PMID: 33972779 | SOX2 + KLF5 |
| 2642 | HHAT | SOX2-KLF5 coregulated genes from PMID: 33972779 | SOX2 + KLF5 |
| 2643 | IRS1 | SOX2-KLF5 coregulated genes from PMID: 33972779 | SOX2 + KLF5 |
| 2644 | CNIH4 | SOX2-KLF5 coregulated genes from PMID: 33972779 | SOX2 + KLF5 |
| 2645 | ITGA6 | SOX2-KLF5 coregulated genes from PMID: 33972779 | SOX2 + KLF5 |
| 2646 | RAB43 | SOX2-KLF5 coregulated genes from PMID: 33972779 | SOX2 + KLF5 |
| 2647 | EPHA7 | SOX2-KLF5 coregulated genes from PMID: 33972779 | SOX2 + KLF5 |
| 2648 | CFTR | SOX2-KLF5 coregulated genes from PMID: 33972779 | SOX2 + KLF5 |
| 2649 | ADAM9 | SOX2-KLF5 coregulated genes from PMID: 33972779 | SOX2 + KLF5 |
| 2650 | SPRY4 | SOX2-KLF5 coregulated genes from PMID: 33972779 | SOX2 + KLF5 |
| 2651 | ST3GAL3 | SOX2-KLF5 coregulated genes from PMID: 33972779 | SOX2 + KLF5 |
| 2652 | BTG2 | SOX2-KLF5 coregulated genes from PMID: 33972779 | SOX2 + KLF5 |
| 2653 | NFKBIZ | SOX2-KLF5 coregulated genes from PMID: 33972779 | SOX2 + KLF5 |
| 2654 | PGBD1 | SOX2-KLF5 coregulated genes from PMID: 33972779 | SOX2 + KLF5 |
| 2655 | LPCAT4 | SOX2-KLF5 coregulated genes from PMID: 33972779 | SOX2 + KLF5 |
| 2656 | RFFL | SOX2-KLF5 coregulated genes from PMID: 33972779 | SOX2 + KLF5 |
| 2657 | NOS1AP | SOX2-KLF5 coregulated genes from PMID: 33972779 | SOX2 + KLF5 |
| 2658 | CLSTN2 | SOX2-KLF5 coregulated genes from PMID: 33972779 | SOX2 + KLF5 |
| 2659 | CLDN4 | SOX2-KLF5 coregulated genes from PMID: 33972779 | SOX2 + KLF5 |
| 2660 | SLC25A24 | SOX2-KLF5 coregulated genes from PMID: 33972779 | SOX2 + KLF5 |
| 2661 | GGH | SOX2-KLF5 coregulated genes from PMID: 33972779 | SOX2 + KLF5 |
| 2662 | ROCK2 | SOX2-KLF5 coregulated genes from PMID: 33972779 | SOX2 + KLF5 |
| 2663 | ARHGAP20 | SOX2-KLF5 coregulated genes from PMID: 33972779 | SOX2 + KLF5 |
| 2664 | DNM3 | SOX2-KLF5 coregulated genes from PMID: 33972779 | SOX2 + KLF5 |
| 2665 | GRIP1 | SOX2-KLF5 coregulated genes from PMID: 33972779 | SOX2 + KLF5 |
| 2666 | CHMP2B | SOX2-KLF5 coregulated genes from PMID: 33972779 | SOX2 + KLF5 |
| 2667 | CAPN2 | SOX2-KLF5 coregulated genes from PMID: 33972779 | SOX2 + KLF5 |
| 2668 | ELF3 | SOX2-KLF5 coregulated genes from PMID: 33972779 | SOX2 + KLF5 |
| 2669 | BMPR2 | SOX2-KLF5 coregulated genes from PMID: 33972779 | SOX2 + KLF5 |
| 2670 | EIF4G3 | SOX2-KLF5 coregulated genes from PMID: 33972779 | SOX2 + KLF5 |
| 2671 | DSP | SOX2-KLF5 coregulated genes from PMID: 33972779 | SOX2 + KLF5 |
| 2672 | ERC1 | SOX2-KLF5 coregulated genes from PMID: 33972779 | SOX2 + KLF5 |
| 2673 | QSOX1 | SOX2-KLF5 coregulated genes from PMID: 33972779 | SOX2 + KLF5 |
| 2674 | KRT20 | SOX2-KLF5 coregulated genes from PMID: 33972779 | SOX2 + KLF5 |
| 2675 | ACOX1 | SOX2-KLF5 coregulated genes from PMID: 33972779 | SOX2 + KLF5 |
| 2676 | RASGRP1 | SOX2-KLF5 coregulated genes from PMID: 33972779 | SOX2 + KLF5 |
| 2677 | HMCN1 | SOX2-KLF5 coregulated genes from PMID: 33972779 | SOX2 + KLF5 |
| 2678 | TNFAIP3 | SOX2-KLF5 coregulated genes from PMID: 33972779 | SOX2 + KLF5 |
| 2679 | NGEF | SOX2-KLF5 coregulated genes from PMID: 33972779 | SOX2 + KLF5 |
| 2680 | ATP10A | SOX2-KLF5 coregulated genes from PMID: 33972779 | SOX2 + KLF5 |
| 2681 | LMNA | SOX2-KLF5 coregulated genes from PMID: 33972779 | SOX2 + KLF5 |
| 2682 | C3orf52 | SOX2-KLF5 coregulated genes from PMID: 33972779 | SOX2 + KLF5 |

| | | | |
|---|---|---|---|
| 2683 | DST | SOX2-KLF5 coregulated genes from PMID: 33972779 | SOX2 + KLF5 |
| 2684 | SGMS2 | SOX2-KLF5 coregulated genes from PMID: 33972779 | SOX2 + KLF5 |
| 2685 | FAM189B | SOX2-KLF5 coregulated genes from PMID: 33972779 | SOX2 + KLF5 |
| 2686 | EZR | SOX2-KLF5 coregulated genes from PMID: 33972779 | SOX2 + KLF5 |
| 2687 | CASQ2 | SOX2-KLF5 coregulated genes from PMID: 33972779 | SOX2 + KLF5 |
| 2688 | CSGALNACT1 | SOX2-KLF5 coregulated genes from PMID: 33972779 | SOX2 + KLF5 |
| 2689 | ZNF362 | SOX2-KLF5 coregulated genes from PMID: 33972779 | SOX2 + KLF5 |
| 2690 | ACAD9 | SOX2-KLF5 coregulated genes from PMID: 33972779 | SOX2 + KLF5 |
| 2691 | CDC42EP3 | SOX2-KLF5 coregulated genes from PMID: 33972779 | SOX2 + KLF5 |
| 2692 | LIMA1 | SOX2-KLF5 coregulated genes from PMID: 33972779 | SOX2 + KLF5 |
| 2693 | NR4A1 | SOX2-KLF5 coregulated genes from PMID: 33972779 | SOX2 + KLF5 |
| 2694 | NRP2 | SOX2-KLF5 coregulated genes from PMID: 33972779 | SOX2 + KLF5 |
| 2695 | ODC1 | SOX2-KLF5 coregulated genes from PMID: 33972779 | SOX2 + KLF5 |
| 2696 | ACVR1 | SOX2-KLF5 coregulated genes from PMID: 33972779 | SOX2 + KLF5 |
| 2697 | FAM126A | SOX2-KLF5 coregulated genes from PMID: 33972779 | SOX2 + KLF5 |
| 2698 | IFIH1 | SOX2-KLF5 coregulated genes from PMID: 33972779 | SOX2 + KLF5 |
| 2699 | SYTL3 | SOX2-KLF5 coregulated genes from PMID: 33972779 | SOX2 + KLF5 |
| 2700 | SMG7 | SOX2-KLF5 coregulated genes from PMID: 33972779 | SOX2 + KLF5 |
| 2701 | SGK1 | SOX2-KLF5 coregulated genes from PMID: 33972779 | SOX2 + KLF5 |
| 2702 | IL18RAP | SOX2-KLF5 coregulated genes from PMID: 33972779 | SOX2 + KLF5 |
| 2703 | LFNG | SOX2-KLF5 coregulated genes from PMID: 33972779 | SOX2 + KLF5 |
| 2704 | SLC4A7 | SOX2-KLF5 coregulated genes from PMID: 33972779 | SOX2 + KLF5 |
| 2705 | C3orf70 | SOX2-KLF5 coregulated genes from PMID: 33972779 | SOX2 + KLF5 |
| 2706 | LAMC2 | SOX2-KLF5 coregulated genes from PMID: 33972779 | SOX2 + KLF5 |
| 2707 | EIF1 | SOX2-KLF5 coregulated genes from PMID: 33972779 | SOX2 + KLF5 |
| 2708 | SFXN3 | SOX2-KLF5 coregulated genes from PMID: 33972779 | SOX2 + KLF5 |
| 2709 | MBNL1 | SOX2-KLF5 coregulated genes from PMID: 33972779 | SOX2 + KLF5 |
| 2710 | NHSL1 | SOX2-KLF5 coregulated genes from PMID: 33972779 | SOX2 + KLF5 |
| 2711 | KIT | SOX2-KLF5 coregulated genes from PMID: 33972779 | SOX2 + KLF5 |
| 2712 | GBP2 | SOX2-KLF5 coregulated genes from PMID: 33972779 | SOX2 + KLF5 |
| 2713 | PLD5 | SOX2-KLF5 coregulated genes from PMID: 33972779 | SOX2 + KLF5 |
| 2714 | STC2 | SOX2-KLF5 coregulated genes from PMID: 33972779 | SOX2 + KLF5 |
| 2715 | RGS5 | SOX2-KLF5 coregulated genes from PMID: 33972779 | SOX2 + KLF5 |
| 2716 | ASPA | SOX2-KLF5 coregulated genes from PMID: 33972779 | SOX2 + KLF5 |
| 2717 | GBP5 | SOX2-KLF5 coregulated genes from PMID: 33972779 | SOX2 + KLF5 |
| 2718 | RPRM | SOX2-KLF5 coregulated genes from PMID: 33972779 | SOX2 + KLF5 |
| 2719 | FOXP1 | SOX2-KLF5 coregulated genes from PMID: 33972779 | SOX2 + KLF5 |
| 2720 | AMIGO1 | SOX2-KLF5 coregulated genes from PMID: 33972779 | SOX2 + KLF5 |
| 2721 | PDE4DIP | SOX2-KLF5 coregulated genes from PMID: 33972779 | SOX2 + KLF5 |
| 2722 | SNRNP48 | SOX2-KLF5 coregulated genes from PMID: 33972779 | SOX2 + KLF5 |
| 2723 | PXDN | SOX2-KLF5 coregulated genes from PMID: 33972779 | SOX2 + KLF5 |
| 2724 | SULT2B1 | SOX2-KLF5 coregulated genes from PMID: 33972779 | SOX2 + KLF5 |
| 2725 | CRYZ | SOX2-KLF5 coregulated genes from PMID: 33972779 | SOX2 + KLF5 |
| 2726 | THBS2 | SOX2-KLF5 coregulated genes from PMID: 33972779 | SOX2 + KLF5 |
| 2727 | STAC | SOX2-KLF5 coregulated genes from PMID: 33972779 | SOX2 + KLF5 |
| 2728 | SLC6A6 | SOX2-KLF5 coregulated genes from PMID: 33972779 | SOX2 + KLF5 |
| 2729 | SPPL3 | SOX2-KLF5 coregulated genes from PMID: 33972779 | SOX2 + KLF5 |
| 2730 | LPGAT1 | SOX2-KLF5 coregulated genes from PMID: 33972779 | SOX2 + KLF5 |
| 2731 | CLDN1 | SOX2-KLF5 coregulated genes from PMID: 33972779 | SOX2 + KLF5 |
| 2732 | PDLIM4 | SOX2-KLF5 coregulated genes from PMID: 33972779 | SOX2 + KLF5 |
| 2733 | EPN2 | SOX2-KLF5 coregulated genes from PMID: 33972779 | SOX2 + KLF5 |
| 2734 | JUN | SOX2-KLF5 coregulated genes from PMID: 33972779 | SOX2 + KLF5 |
| 2735 | TNIP1 | SOX2-KLF5 coregulated genes from PMID: 33972779 | SOX2 + KLF5 |
| 2736 | S1PR1 | SOX2-KLF5 coregulated genes from PMID: 33972779 | SOX2 + KLF5 |
| 2737 | CD9 | SOX2-KLF5 coregulated genes from PMID: 33972779 | SOX2 + KLF5 |
| 2738 | ANGPT1 | SOX2-KLF5 coregulated genes from PMID: 33972779 | SOX2 + KLF5 |
| 2739 | SLC6A2 | SOX2-KLF5 coregulated genes from PMID: 33972779 | SOX2 + KLF5 |
| 2740 | RBFOX1 | SOX2-KLF5 coregulated genes from PMID: 33972779 | SOX2 + KLF5 |
| 2741 | DGKI | SOX2-KLF5 coregulated genes from PMID: 33972779 | SOX2 + KLF5 |
| 2742 | ATXN1 | SOX2-KLF5 coregulated genes from PMID: 33972779 | SOX2 + KLF5 |
| 2743 | SOX13 | SOX2-KLF5 coregulated genes from PMID: 33972779 | SOX2 + KLF5 |

| | | | |
|---|---|---|---|
| 2744 | VCAM1 | SOX2-KLF5 coregulated genes from PMID: 33972779 | SOX2 + KLF5 |
| 2745 | TFCP2L1 | SOX2-KLF5 coregulated genes from PMID: 33972779 | SOX2 + KLF5 |
| 2746 | LRRC8B | SOX2-KLF5 coregulated genes from PMID: 33972779 | SOX2 + KLF5 |
| 2747 | MACF1 | SOX2-KLF5 coregulated genes from PMID: 33972779 | SOX2 + KLF5 |
| 2748 | TP63 | SOX2-KLF5 coregulated genes from PMID: 33972779 | SOX2 + KLF5 |
| 2749 | DUSP1 | SOX2-KLF5 coregulated genes from PMID: 33972779 | SOX2 + KLF5 |
| 2750 | SLC16A11 | SOX2-KLF5 coregulated genes from PMID: 33972779 | SOX2 + KLF5 |
| 2751 | PLS1 | SOX2-KLF5 coregulated genes from PMID: 33972779 | SOX2 + KLF5 |
| 2752 | ABCB11 | SOX2-KLF5 coregulated genes from PMID: 33972779 | SOX2 + KLF5 |
| 2753 | HS3ST3A1 | SOX2-KLF5 coregulated genes from PMID: 33972779 | SOX2 + KLF5 |
| 2754 | ANGPTL4 | SOX2-KLF5 coregulated genes from PMID: 33972779 | SOX2 + KLF5 |
| 2755 | HRH1 | SOX2-KLF5 coregulated genes from PMID: 33972779 | SOX2 + KLF5 |
| 2756 | GNB4 | SOX2-KLF5 coregulated genes from PMID: 33972779 | SOX2 + KLF5 |
| 2757 | CNN3 | SOX2-KLF5 coregulated genes from PMID: 33972779 | SOX2 + KLF5 |
| 2758 | CREG1 | SOX2-KLF5 coregulated genes from PMID: 33972779 | SOX2 + KLF5 |
| 2759 | RNF19B | SOX2-KLF5 coregulated genes from PMID: 33972779 | SOX2 + KLF5 |
| 2760 | PDE4D | SOX2-KLF5 coregulated genes from PMID: 33972779 | SOX2 + KLF5 |
| 2761 | MDFIC | SOX2-KLF5 coregulated genes from PMID: 33972779 | SOX2 + KLF5 |
| 2762 | BCL2L15 | SOX2-KLF5 coregulated genes from PMID: 33972779 | SOX2 + KLF5 |
| 2763 | TRIO | SOX2-KLF5 coregulated genes from PMID: 33972779 | SOX2 + KLF5 |
| 2764 | LPAR3 | SOX2-KLF5 coregulated genes from PMID: 33972779 | SOX2 + KLF5 |
| 2765 | GOLIM4 | SOX2-KLF5 coregulated genes from PMID: 33972779 | SOX2 + KLF5 |
| 2766 | MYH14 | SOX2-KLF5 coregulated genes from PMID: 33972779 | SOX2 + KLF5 |
| 2767 | VIM | SOX2-KLF5 coregulated genes from PMID: 33972779 | SOX2 + KLF5 |
| 2768 | COL4A3 | SOX2-KLF5 coregulated genes from PMID: 33972779 | SOX2 + KLF5 |
| 2769 | PTPRJ | SOX2-KLF5 coregulated genes from PMID: 33972779 | SOX2 + KLF5 |
| 2770 | NWD1 | SOX2-KLF5 coregulated genes from PMID: 33972779 | SOX2 + KLF5 |
| 2771 | MNDA | SOX2-KLF5 coregulated genes from PMID: 33972779 | SOX2 + KLF5 |
| 2772 | SLC25A10 | SOX2-KLF5 coregulated genes from PMID: 33972779 | SOX2 + KLF5 |
| 2773 | PRMT6 | SOX2-KLF5 coregulated genes from PMID: 33972779 | SOX2 + KLF5 |
| 2774 | PTGS2 | SOX2-KLF5 coregulated genes from PMID: 33972779 | SOX2 + KLF5 |
| 2775 | CCL20 | SOX2-KLF5 coregulated genes from PMID: 33972779 | SOX2 + KLF5 |
| 2776 | RARB | SOX2-KLF5 coregulated genes from PMID: 33972779 | SOX2 + KLF5 |
| 2777 | TWF2 | SOX2-KLF5 coregulated genes from PMID: 33972779 | SOX2 + KLF5 |
| 2778 | DHRS9 | SOX2-KLF5 coregulated genes from PMID: 33972779 | SOX2 + KLF5 |
| 2779 | SH3PXD2A | SOX2-KLF5 coregulated genes from PMID: 33972779 | SOX2 + KLF5 |
| 2780 | PLEKHM3 | SOX2-KLF5 coregulated genes from PMID: 33972779 | SOX2 + KLF5 |
| 2781 | AFF1 | SOX2-KLF5 coregulated genes from PMID: 33972779 | SOX2 + KLF5 |
| 2782 | TRIM33 | SOX2-KLF5 coregulated genes from PMID: 33972779 | SOX2 + KLF5 |
| 2783 | LRIG1 | SOX2-KLF5 coregulated genes from PMID: 33972779 | SOX2 + KLF5 |
| 2784 | SIPA1L2 | SOX2-KLF5 coregulated genes from PMID: 33972779 | SOX2 + KLF5 |
| 2785 | ABR | SOX2-KLF5 coregulated genes from PMID: 33972779 | SOX2 + KLF5 |
| 2786 | NUAK1 | SOX2-KLF5 coregulated genes from PMID: 33972779 | SOX2 + KLF5 |
| 2787 | RAP2B | SOX2-KLF5 coregulated genes from PMID: 33972779 | SOX2 + KLF5 |
| 2788 | KRT8 | SOX2-KLF5 coregulated genes from PMID: 33972779 | SOX2 + KLF5 |
| 2789 | SEMA6A | SOX2-KLF5 coregulated genes from PMID: 33972779 | SOX2 + KLF5 |
| 2790 | SBF2 | SOX2-KLF5 coregulated genes from PMID: 33972779 | SOX2 + KLF5 |
| 2791 | LRRC8C | SOX2-KLF5 coregulated genes from PMID: 33972779 | SOX2 + KLF5 |
| 2792 | IQGAP2 | SOX2-KLF5 coregulated genes from PMID: 33972779 | SOX2 + KLF5 |
| 2793 | BMP2 | SOX2 regulated genes from PMID: 20726797 | SOX2 |
| 2794 | BMP4 | SOX2 regulated genes from PMID: 20726797 | SOX2 |
| 2795 | BMPR2 | SOX2 regulated genes from PMID: 20726797 | SOX2 |
| 2796 | PCSK6 | SOX2 regulated genes from PMID: 20726797 | SOX2 |
| 2797 | SMAD5 | SOX2 regulated genes from PMID: 20726797 | SOX2 |
| 2798 | SMAD6 | SOX2 regulated genes from PMID: 20726797 | SOX2 |
| 2799 | SMAD7 | SOX2 regulated genes from PMID: 20726797 | SOX2 |
| 2800 | SMURF1 | SOX2 regulated genes from PMID: 20726797 | SOX2 |
| 2801 | IGF1R | SOX2 regulated genes from PMID: 20726797 | SOX2 |
| 2802 | GNA15 | SOX2 regulated genes from PMID: 20726797 | SOX2 |
| 2803 | TGFB2 | SOX2 regulated genes from PMID: 20726797 | SOX2 |
| 2804 | RGS3 | SOX2 regulated genes from PMID: 20726797 | SOX2 |

| | | | |
|---|---|---|---|
| 2805 | FGFR3 | SOX2 regulated genes from PMID: 20726797 | SOX2 |
| 2806 | IL6ST | SOX2 regulated genes from PMID: 20726797 | SOX2 |
| 2807 | EREG | SOX2 regulated genes from PMID: 20726797 | SOX2 |
| 2808 | SOCS5 | SOX2 regulated genes from PMID: 20726797 | SOX2 |
| 2809 | IRAK1 | SOX2 regulated genes from PMID: 20726797 | SOX2 |
| 2810 | PIK3R1 | SOX2 regulated genes from PMID: 20726797 | SOX2 |
| 2811 | EDN1 | SOX2 regulated genes from PMID: 20726797 | SOX2 |
| 2812 | OPN3 | SOX2 regulated genes from PMID: 20726797 | SOX2 |
| 2813 | PYCARD | SOX2 regulated genes from PMID: 20726797 | SOX2 |
| 2814 | SIGIRR | SOX2 regulated genes from PMID: 20726797 | SOX2 |
| 2815 | CD24 | SOX2 regulated genes from PMID: 20726797 | SOX2 |
| 2816 | AGRN | SOX2 regulated genes from PMID: 20726797 | SOX2 |
| 2817 | RHOQ | SOX2 regulated genes from PMID: 20726797 | SOX2 |
| 2818 | IRS1 | SOX2 regulated genes from PMID: 20726797 | SOX2 |
| 2819 | MAML3 | SOX2 regulated genes from PMID: 20726797 | SOX2 |
| 2820 | GNB2 | SOX2 regulated genes from PMID: 20726797 | SOX2 |
| 2821 | COL16A1 | SOX2 regulated genes from PMID: 20726797 | SOX2 |
| 2822 | MTSS1 | SOX2 regulated genes from PMID: 20726797 | SOX2 |
| 2823 | TRIB1 | SOX2 regulated genes from PMID: 20726797 | SOX2 |
| 2824 | PTK2 | SOX2 regulated genes from PMID: 20726797 | SOX2 |
| 2825 | IFITM1 | SOX2 regulated genes from PMID: 20726797 | SOX2 |
| 2826 | ACVR1 | SOX2 regulated genes from PMID: 20726797 | SOX2 |
| 2827 | CCRL2 | SOX2 regulated genes from PMID: 20726797 | SOX2 |
| 2828 | TACSTD2 | SOX2 regulated genes from PMID: 20726797 | SOX2 |
| 2829 | PTPRF | SOX2 regulated genes from PMID: 20726797 | SOX2 |
| 2830 | SMAD3 | SOX2 regulated genes from PMID: 20726797 | SOX2 |
| 2831 | DGKQ | SOX2 regulated genes from PMID: 20726797 | SOX2 |
| 2832 | SRC | SOX2 regulated genes from PMID: 20726797 | SOX2 |
| 2833 | EGFR | SOX2 regulated genes from PMID: 20726797 | SOX2 |
| 2834 | DGKH | SOX2 regulated genes from PMID: 20726797 | SOX2 |
| 2835 | KLK6 | SOX2 regulated genes from PMID: 20726797 | SOX2 |
| 2836 | GAB1 | SOX2 regulated genes from PMID: 20726797 | SOX2 |
| 2837 | VIPR1 | SOX2 regulated genes from PMID: 20726797 | SOX2 |
| 2838 | PARD3 | SOX2 regulated genes from PMID: 20726797 | SOX2 |
| 2839 | CD14 | SOX2 regulated genes from PMID: 20726797 | SOX2 |
| 2840 | ROR1 | SOX2 regulated genes from PMID: 20726797 | SOX2 |
| 2841 | ADAM17 | SOX2 regulated genes from PMID: 20726797 | SOX2 |
| 2842 | ADAM10 | SOX2 regulated genes from PMID: 20726797 | SOX2 |
| 2843 | NUP62 | SOX2 regulated genes from PMID: 20726797 | SOX2 |
| 2844 | LEPR | SOX2 regulated genes from PMID: 20726797 | SOX2 |
| 2845 | P2RY2 | SOX2 regulated genes from PMID: 20726797 | SOX2 |
| 2846 | AFAP1L2 | SOX2 regulated genes from PMID: 20726797 | SOX2 |
| 2847 | AREG | SOX2 regulated genes from PMID: 20726797 | SOX2 |
| 2848 | HPGD | SOX2 regulated genes from PMID: 20726797 | SOX2 |
| 2849 | STAT3 | SOX2 regulated genes from PMID: 20726797 | SOX2 |
| 2850 | CXCL1 | SOX2 regulated genes from PMID: 20726797 | SOX2 |
| 2851 | GPR37 | SOX2 regulated genes from PMID: 20726797 | SOX2 |
| 2852 | CD59 | SOX2 regulated genes from PMID: 20726797 | SOX2 |
| 2853 | LY6E | SOX2 regulated genes from PMID: 20726797 | SOX2 |
| 2854 | CBLC | SOX2 regulated genes from PMID: 20726797 | SOX2 |
| 2855 | PSENEN | SOX2 regulated genes from PMID: 20726797 | SOX2 |
| 2856 | THBS1 | SOX2 regulated genes from PMID: 20726797 | SOX2 |
| 2857 | DEFB1 | SOX2 regulated genes from PMID: 20726797 | SOX2 |
| 2858 | ADRA2A | SOX2 regulated genes from PMID: 20726797 | SOX2 |
| 2859 | RGS2 | SOX2 regulated genes from PMID: 20726797 | SOX2 |
| 2860 | ITPR3 | SOX2 regulated genes from PMID: 20726797 | SOX2 |
| 2861 | RBP4 | SOX2 regulated genes from PMID: 20726797 | SOX2 |
| 2862 | HSD17B2 | SOX2 regulated genes from PMID: 20726797 | SOX2 |
| 2863 | RXRA | SOX2 regulated genes from PMID: 20726797 | SOX2 |
| 2864 | AQP3 | SOX2 regulated genes from PMID: 20726797 | SOX2 |
| 2865 | RARA | SOX2 regulated genes from PMID: 20726797 | SOX2 |

| | | | |
|---|---|---|---|
| 2866 | TRIM16 | SOX2 regulated genes from PMID: 20726797 | SOX2 |
| 2867 | ELF1 | SOX2 regulated genes in Wounding from PMID: 30772301 | SOX2 |
| 2868 | FLRT3 | SOX2 regulated genes in Wounding from PMID: 30772301 | SOX2 |
| 2869 | SUV39H2 | SOX2 regulated genes in Wounding from PMID: 30772301 | SOX2 |
| 2870 | PSTPIP2 | SOX2 regulated genes in Wounding from PMID: 30772301 | SOX2 |
| 2871 | TTC39B | SOX2 regulated genes in Wounding from PMID: 30772301 | SOX2 |
| 2872 | EXOC5 | SOX2 regulated genes in Wounding from PMID: 30772301 | SOX2 |
| 2873 | SLITRK4 | SOX2 regulated genes in Wounding from PMID: 30772301 | SOX2 |
| 2874 | TFAP2C | SOX2 regulated genes in Wounding from PMID: 30772301 | SOX2 |
| 2875 | AHCY | SOX2 regulated genes in Wounding from PMID: 30772301 | SOX2 |
| 2876 | OSBP2 | SOX2 regulated genes in Wounding from PMID: 30772301 | SOX2 |
| 2877 | IL13RA1 | SOX2 regulated genes in Wounding from PMID: 30772301 | SOX2 |
| 2878 | ADGRA3 | SOX2 regulated genes in Wounding from PMID: 30772301 | SOX2 |
| 2879 | PBDC1 | SOX2 regulated genes in Wounding from PMID: 30772301 | SOX2 |
| 2880 | SMOX | SOX2 regulated genes in Wounding from PMID: 30772301 | SOX2 |
| 2881 | TCEAL5 | SOX2 regulated genes in Wounding from PMID: 30772301 | SOX2 |
| 2882 | FBXO7 | SOX2 regulated genes in Wounding from PMID: 30772301 | SOX2 |
| 2883 | SLC39A6 | SOX2 regulated genes in Wounding from PMID: 30772301 | SOX2 |
| 2884 | YES1 | SOX2 regulated genes in Wounding from PMID: 30772301 | SOX2 |
| 2885 | RGS6 | SOX2 regulated genes in Wounding from PMID: 30772301 | SOX2 |
| 2886 | TMEM50B | SOX2 regulated genes in Wounding from PMID: 30772301 | SOX2 |
| 2887 | GAREM1 | SOX2 regulated genes in Wounding from PMID: 30772301 | SOX2 |
| 2888 | SLC37A1 | SOX2 regulated genes in Wounding from PMID: 30772301 | SOX2 |
| 2889 | FAM3B | SOX2 regulated genes in Wounding from PMID: 30772301 | SOX2 |
| 2890 | CCT8 | SOX2 regulated genes in Wounding from PMID: 30772301 | SOX2 |
| 2891 | ASCC2 | SOX2 regulated genes in Wounding from PMID: 30772301 | SOX2 |
| 2892 | TPTEP2-CSNK1E | SOX2 regulated genes in Wounding from PMID: 30772301 | SOX2 |
| 2893 | CSNK1E | SOX2 regulated genes in Wounding from PMID: 30772301 | SOX2 |
| 2894 | KLF6 | SOX2 regulated genes in Wounding from PMID: 30772301 | SOX2 |
| 2895 | BTG3 | SOX2 regulated genes in Wounding from PMID: 30772301 | SOX2 |
| 2896 | LURAP1L | SOX2 regulated genes in Wounding from PMID: 30772301 | SOX2 |
| 2897 | DSG3 | SOX2 regulated genes in Wounding from PMID: 30772301 | SOX2 |
| 2898 | MAFF | SOX2 regulated genes in Wounding from PMID: 30772301 | SOX2 |
| 2899 | HSPA13 | SOX2 regulated genes in Wounding from PMID: 30772301 | SOX2 |
| 2900 | WDR1 | SOX2 regulated genes in Wounding from PMID: 30772301 | SOX2 |
| 2901 | PSPC1 | SOX2 regulated genes in Wounding from PMID: 30772301 | SOX2 |
| 2902 | RAE1 | SOX2 regulated genes in Wounding from PMID: 30772301 | SOX2 |
| 2903 | EIF2S2 | SOX2 regulated genes in Wounding from PMID: 30772301 | SOX2 |
| 2904 | GTPBP6 | SOX2 regulated genes in Wounding from PMID: 30772301 | SOX2 |
| 2905 | PRELID3B | SOX2 regulated genes in Wounding from PMID: 30772301 | SOX2 |
| 2906 | UNC79 | SOX2 regulated genes in Wounding from PMID: 30772301 | SOX2 |
| 2907 | ZBTB7C | SOX2 regulated genes in Wounding from PMID: 30772301 | SOX2 |
| 2908 | LTBP2 | SOX2 regulated genes in Wounding from PMID: 30772301 | SOX2 |
| 2909 | DEPDC5 | SOX2 regulated genes in Wounding from PMID: 30772301 | SOX2 |
| 2910 | TMEM30B | SOX2 regulated genes in Wounding from PMID: 30772301 | SOX2 |
| 2911 | PCSK2 | SOX2 regulated genes in Wounding from PMID: 30772301 | SOX2 |
| 2912 | NUDT11 | SOX2 regulated genes in Wounding from PMID: 30772301 | SOX2 |
| 2913 | BRWD1 | SOX2 regulated genes in Wounding from PMID: 30772301 | SOX2 |
| 2914 | BTBD7 | SOX2 regulated genes in Wounding from PMID: 30772301 | SOX2 |
| 2915 | DSTN | SOX2 regulated genes in Wounding from PMID: 30772301 | SOX2 |
| 2916 | SAMD10 | SOX2 regulated genes in Wounding from PMID: 30772301 | SOX2 |
| 2917 | XKRX | SOX2 regulated genes in Wounding from PMID: 30772301 | SOX2 |
| 2918 | MTCL1 | SOX2 regulated genes in Wounding from PMID: 30772301 | SOX2 |
| 2919 | NAA30 | SOX2 regulated genes in Wounding from PMID: 30772301 | SOX2 |
| 2920 | SCEL | SOX2 regulated genes in Wounding from PMID: 30772301 | SOX2 |
| 2921 | RRP1 | SOX2 regulated genes in Wounding from PMID: 30772301 | SOX2 |
| 2922 | CCDC85C | SOX2 regulated genes in Wounding from PMID: 30772301 | SOX2 |
| 2923 | RNF152 | SOX2 regulated genes in Wounding from PMID: 30772301 | SOX2 |
| 2924 | CLDN14 | SOX2 regulated genes in Wounding from PMID: 30772301 | SOX2 |
| 2925 | PLEKHD1 | SOX2 regulated genes in Wounding from PMID: 30772301 | SOX2 |
| 2926 | OSM | SOX2 regulated genes in Wounding from PMID: 30772301 | SOX2 |

| | | | |
|---|---|---|---|
| 2927 | RNF17 | SOX2 regulated genes in Wounding from PMID: 30772301 | SOX2 |
| 2928 | TMEM33 | SOX2 regulated genes in Wounding from PMID: 30772301 | SOX2 |
| 2929 | TRMT5 | SOX2 regulated genes in Wounding from PMID: 30772301 | SOX2 |
| 2930 | BMP2 | SOX2 regulated genes in Wounding from PMID: 30772301 | SOX2 |
| 2931 | SRSF5 | SOX2 regulated genes in Wounding from PMID: 30772301 | SOX2 |
| 2932 | LHFPL1 | SOX2 regulated genes in Wounding from PMID: 30772301 | SOX2 |
| 2933 | CDKN2B | SOX2 regulated genes in Wounding from PMID: 30772301 | SOX2 |
| 2934 | TXNL1 | SOX2 regulated genes in Wounding from PMID: 30772301 | SOX2 |
| 2935 | DHX15 | SOX2 regulated genes in Wounding from PMID: 30772301 | SOX2 |
| 2936 | POF1B | SOX2 regulated genes in Wounding from PMID: 30772301 | SOX2 |
| 2937 | NDP | SOX2 regulated genes in Wounding from PMID: 30772301 | SOX2 |
| 2938 | MTAP | SOX2 regulated genes in Wounding from PMID: 30772301 | SOX2 |
| 2939 | RIPK4 | SOX2 regulated genes in Wounding from PMID: 30772301 | SOX2 |
| 2940 | EFNB1 | SOX2 regulated genes in Wounding from PMID: 30772301 | SOX2 |
| 2941 | ACAA2 | SOX2 regulated genes in Wounding from PMID: 30772301 | SOX2 |
| 2942 | PDE9A | SOX2 regulated genes in Wounding from PMID: 30772301 | SOX2 |
| 2943 | LIMK2 | SOX2 regulated genes in Wounding from PMID: 30772301 | SOX2 |
| 2944 | CTIF | SOX2 regulated genes in Wounding from PMID: 30772301 | SOX2 |
| 2945 | SYNGR1 | SOX2 regulated genes in Wounding from PMID: 30772301 | SOX2 |
| 2946 | PMEPA1 | SOX2 regulated genes in Wounding from PMID: 30772301 | SOX2 |
| 2947 | HSPH1 | SOX2 regulated genes in Wounding from PMID: 30772301 | SOX2 |
| 2948 | USP16 | SOX2 regulated genes in Wounding from PMID: 30772301 | SOX2 |
| 2949 | C21orf91 | SOX2 regulated genes in Wounding from PMID: 30772301 | SOX2 |
| 2950 | RAB27B | SOX2 regulated genes in Wounding from PMID: 30772301 | SOX2 |
| 2951 | CORIN | SOX2 regulated genes in Wounding from PMID: 30772301 | SOX2 |
| 2952 | EIF4ENIF1 | SOX2 regulated genes in Wounding from PMID: 30772301 | SOX2 |
| 2953 | TGIF1 | SOX2 regulated genes in Wounding from PMID: 30772301 | SOX2 |
| 2954 | MYO5B | SOX2 regulated genes in Wounding from PMID: 30772301 | SOX2 |
| 2955 | SLAIN1 | SOX2 regulated genes in Wounding from PMID: 30772301 | SOX2 |
| 2956 | MSX1 | SOX2 regulated genes in Wounding from PMID: 30772301 | SOX2 |
| 2957 | GTF2F2 | SOX2 regulated genes in Wounding from PMID: 30772301 | SOX2 |
| 2958 | JAG1 | SOX2 regulated genes in Wounding from PMID: 30772301 | SOX2 |
| 2959 | SLC25A17 | SOX2 regulated genes in Wounding from PMID: 30772301 | SOX2 |
| 2960 | CDKL5 | SOX2 regulated genes in Wounding from PMID: 30772301 | SOX2 |
| 2961 | BRWD3 | SOX2 regulated genes in Wounding from PMID: 30772301 | SOX2 |
| 2962 | SMAD2 | SOX2 regulated genes in Wounding from PMID: 30772301 | SOX2 |
| 2963 | TEC | SOX2 regulated genes in Wounding from PMID: 30772301 | SOX2 |
| 2964 | HOPX | SOX2 regulated genes in Wounding from PMID: 30772301 | SOX2 |
| 2965 | SDC4 | SOX2 regulated genes in Wounding from PMID: 30772301 | SOX2 |
| 2966 | ATRN | SOX2 regulated genes in Wounding from PMID: 30772301 | SOX2 |
| 2967 | COL18A1 | SOX2 regulated genes in Wounding from PMID: 30772301 | SOX2 |
| 2968 | PLA2G3 | SOX2 regulated genes in Wounding from PMID: 30772301 | SOX2 |
| 2969 | ETS2 | SOX2 regulated genes in Wounding from PMID: 30772301 | SOX2 |
| 2970 | TBC1D25 | SOX2 regulated genes in Wounding from PMID: 30772301 | SOX2 |
| 2971 | DSG4 | SOX2 regulated genes in Wounding from PMID: 30772301 | SOX2 |
| 2972 | CCDC68 | SOX2 regulated genes in Wounding from PMID: 30772301 | SOX2 |
| 2973 | RUNX1 | SOX2 regulated genes in Wounding from PMID: 30772301 | SOX2 |
| 2974 | S100G | SOX2 regulated genes in Wounding from PMID: 30772301 | SOX2 |
| 2975 | EYA2 | SOX2 regulated genes in Wounding from PMID: 30772301 | SOX2 |
| 2976 | CPLX1 | SOX2 regulated genes in Wounding from PMID: 30772301 | SOX2 |
| 2977 | JADE3 | SOX2 regulated genes in Wounding from PMID: 30772301 | SOX2 |
| 2978 | RNF4 | SOX2 regulated genes in Wounding from PMID: 30772301 | SOX2 |
| 2979 | MORC4 | SOX2 regulated genes in Wounding from PMID: 30772301 | SOX2 |
| 2980 | SYCP2 | SOX2 regulated genes in Wounding from PMID: 30772301 | SOX2 |
| 2981 | PPP2R5E | SOX2 regulated genes in Wounding from PMID: 30772301 | SOX2 |
| 2982 | RBBP8NL | SOX2 regulated genes in Wounding from PMID: 30772301 | SOX2 |
| 2983 | CSTB | SOX2 regulated genes in Wounding from PMID: 30772301 | SOX2 |
| 2984 | TP53INP2 | SOX2 regulated genes in Wounding from PMID: 30772301 | SOX2 |
| 2985 | BMP7 | SOX2 regulated genes in Wounding from PMID: 30772301 | SOX2 |
| 2986 | CLDN8 | SOX2 regulated genes in Wounding from PMID: 30772301 | SOX2 |
| 2987 | RPRD1A | SOX2 regulated genes in Wounding from PMID: 30772301 | SOX2 |

| | | | |
|---|---|---|---|
| 2988 | ATP9A | SOX2 regulated genes in Wounding from PMID: 30772301 | SOX2 |
| 2989 | SYN3 | SOX2 regulated genes in Wounding from PMID: 30772301 | SOX2 |
| 2990 | PCGF3 | SOX2 regulated genes in Wounding from PMID: 30772301 | SOX2 |
| 2991 | SEC23B | SOX2 regulated genes in Wounding from PMID: 30772301 | SOX2 |
| 2992 | PCBP3 | SOX2 regulated genes in Wounding from PMID: 30772301 | SOX2 |
| 2993 | PCNX4 | SOX2 regulated genes in Wounding from PMID: 30772301 | SOX2 |
| 2994 | PDRG1 | SOX2 regulated genes in Wounding from PMID: 30772301 | SOX2 |
| 2995 | RGP1 | SOX2 regulated genes in Wounding from PMID: 30772301 | SOX2 |
| 2996 | OSBPL3 | SOX2 regulated genes in Wounding from PMID: 30772301 | SOX2 |
| 2997 | PTPN3 | SOX2 regulated genes in Wounding from PMID: 30772301 | SOX2 |
| 2998 | DNASE1L2 | SOX2 regulated genes in Wounding from PMID: 30772301 | SOX2 |
| 2999 | TNFRSF13C | SOX2 regulated genes in Wounding from PMID: 30772301 | SOX2 |
| 3000 | RNPS1 | SOX2 regulated genes in Wounding from PMID: 30772301 | SOX2 |
| 3001 | SPEF1 | SOX2 regulated genes in Wounding from PMID: 30772301 | SOX2 |
| 3002 | RASL10A | SOX2 regulated genes in Wounding from PMID: 30772301 | SOX2 |
| 3003 | OSER1 | SOX2 regulated genes in Wounding from PMID: 30772301 | SOX2 |
| 3004 | TES | SOX2 regulated genes in Wounding from PMID: 30772301 | SOX2 |
| 3005 | SPIDR | SOX2 regulated genes in Wounding from PMID: 30772301 | SOX2 |
| 3006 | CAGE1 | SOX2 regulated genes in Wounding from PMID: 30772301 | SOX2 |
| 3007 | PNP | SOX2 regulated genes in Wounding from PMID: 30772301 | SOX2 |
| 3008 | ARID3A | SOX2 regulated genes in Wounding from PMID: 30772301 | SOX2 |
| 3009 | BNIP3L | SOX2 regulated genes in Wounding from PMID: 30772301 | SOX2 |
| 3010 | CSPG4 | SOX2 regulated genes in Wounding from PMID: 30772301 | SOX2 |
| 3011 | PRRG4 | SOX2 regulated genes in Wounding from PMID: 30772301 | SOX2 |
| 3012 | PDAP1 | SOX2 regulated genes in Wounding from PMID: 30772301 | SOX2 |
| 3013 | CELSR1 | SOX2 regulated genes in Wounding from PMID: 30772301 | SOX2 |
| 3014 | PJA1 | SOX2 regulated genes in Wounding from PMID: 30772301 | SOX2 |
| 3015 | COL4A5 | SOX2 regulated genes in Wounding from PMID: 30772301 | SOX2 |
| 3016 | AIMP2 | SOX2 regulated genes in Wounding from PMID: 30772301 | SOX2 |
| 3017 | DUSP9 | SOX2 regulated genes in Wounding from PMID: 30772301 | SOX2 |
| 3018 | MICALL2 | SOX2 regulated genes in Wounding from PMID: 30772301 | SOX2 |
| 3019 | FRYL | SOX2 regulated genes in Wounding from PMID: 30772301 | SOX2 |
| 3020 | CD2AP | SOX2 regulated genes in Wounding from PMID: 30772301 | SOX2 |
| 3021 | ITGA5 | SOX2 regulated genes in Wounding from PMID: 30772301 | SOX2 |
| 3022 | C8orf48 | SOX2 regulated genes in Wounding from PMID: 30772301 | SOX2 |
| 3023 | GSDMA | SOX2 regulated genes in Wounding from PMID: 30772301 | SOX2 |
| 3024 | BBOX1 | SOX2 regulated genes in Wounding from PMID: 30772301 | SOX2 |
| 3025 | ANKRD50 | SOX2 regulated genes in Wounding from PMID: 30772301 | SOX2 |
| 3026 | PARM1 | SOX2 regulated genes in Wounding from PMID: 30772301 | SOX2 |
| 3027 | SMTN | SOX2 regulated genes in Wounding from PMID: 30772301 | SOX2 |
| 3028 | PARD6B | SOX2 regulated genes in Wounding from PMID: 30772301 | SOX2 |
| 3029 | SLC39A2 | SOX2 regulated genes in Wounding from PMID: 30772301 | SOX2 |
| 3030 | TIAM1 | SOX2 regulated genes in Wounding from PMID: 30772301 | SOX2 |
| 3031 | TAX1BP3 | SOX2 regulated genes in Wounding from PMID: 30772301 | SOX2 |
| 3032 | RBM12 | SOX2 regulated genes in Wounding from PMID: 30772301 | SOX2 |
| 3033 | QSOX2 | SOX2 regulated genes in Wounding from PMID: 30772301 | SOX2 |
| 3034 | STAR | SOX2 regulated genes in Wounding from PMID: 30772301 | SOX2 |
| 3035 | JHY | SOX2 regulated genes in Wounding from PMID: 30772301 | SOX2 |
| 3036 | CSNK2A3 | SOX2 regulated genes in Wounding from PMID: 30772301 | SOX2 |
| 3037 | CSNK2A1 | SOX2 regulated genes in Wounding from PMID: 30772301 | SOX2 |
| 3038 | TADA2B | SOX2 regulated genes in Wounding from PMID: 30772301 | SOX2 |
| 3039 | CTBP2 | SOX2 regulated genes in Wounding from PMID: 30772301 | SOX2 |
| 3040 | SLC38A2 | SOX2 regulated genes in Wounding from PMID: 30772301 | SOX2 |
| 3041 | CARMIL1 | SOX2 regulated genes in Wounding from PMID: 30772301 | SOX2 |
| 3042 | FAM83G | SOX2 regulated genes in Wounding from PMID: 30772301 | SOX2 |
| 3043 | LMX1B | SOX2 regulated genes in Wounding from PMID: 30772301 | SOX2 |
| 3044 | LAMA1 | SOX2 regulated genes in Wounding from PMID: 30772301 | SOX2 |
| 3045 | GLS2 | SOX2 regulated genes in Wounding from PMID: 30772301 | SOX2 |
| 3046 | NUP58 | SOX2 regulated genes in Wounding from PMID: 30772301 | SOX2 |
| 3047 | GCOM1 | SOX2 regulated genes in Wounding from PMID: 30772301 | SOX2 |
| 3048 | SERPINA3 | SOX2 regulated genes in Wounding from PMID: 30772301 | SOX2 |

| | | | |
|---|---|---|---|
| 3049 | AMOTL1 | SOX2 regulated genes in Wounding from PMID: 30772301 | SOX2 |
| 3050 | PIP5K1B | SOX2 regulated genes in Wounding from PMID: 30772301 | SOX2 |
| 3051 | DSC2 | SOX2 regulated genes in Wounding from PMID: 30772301 | SOX2 |
| 3052 | PTPN13 | SOX2 regulated genes in Wounding from PMID: 30772301 | SOX2 |
| 3053 | THNSL1 | SOX2 regulated genes in Wounding from PMID: 30772301 | SOX2 |
| 3054 | ORMDL3 | SOX2 regulated genes in Wounding from PMID: 30772301 | SOX2 |
| 3055 | GJA1 | SOX2 regulated genes in Wounding from PMID: 30772301 | SOX2 |
| 3056 | CASK | SOX2 regulated genes in Wounding from PMID: 30772301 | SOX2 |
| 3057 | NCBP1 | SOX2 regulated genes in Wounding from PMID: 30772301 | SOX2 |
| 3058 | HNRNPK | SOX2 regulated genes in Wounding from PMID: 30772301 | SOX2 |
| 3059 | DIO3 | SOX2 regulated genes in Wounding from PMID: 30772301 | SOX2 |
| 3060 | PPIL1 | SOX2 regulated genes in Wounding from PMID: 30772301 | SOX2 |
| 3061 | TLE1 | SOX2 regulated genes in Wounding from PMID: 30772301 | SOX2 |
| 3062 | DLGAP4 | SOX2 regulated genes in Wounding from PMID: 30772301 | SOX2 |
| 3063 | CYB5R4 | SOX2 regulated genes in Wounding from PMID: 30772301 | SOX2 |
| 3064 | TBX1 | SOX2 regulated genes in Wounding from PMID: 30772301 | SOX2 |
| 3065 | RNGTT | SOX2 regulated genes in Wounding from PMID: 30772301 | SOX2 |
| 3066 | MYO1D | SOX2 regulated genes in Wounding from PMID: 30772301 | SOX2 |
| 3067 | FMN1 | SOX2 regulated genes in Wounding from PMID: 30772301 | SOX2 |
| 3068 | POLR2M | SOX2 regulated genes in Wounding from PMID: 30772301 | SOX2 |
| 3069 | EIF2B2 | SOX2 regulated genes in Wounding from PMID: 30772301 | SOX2 |
| 3070 | ABHD13 | SOX2 regulated genes in Wounding from PMID: 30772301 | SOX2 |
| 3071 | ARHGEF17 | SOX2 regulated genes in Wounding from PMID: 30772301 | SOX2 |
| 3072 | CCT6A | SOX2 regulated genes in Wounding from PMID: 30772301 | SOX2 |
| 3073 | SMARCA5 | SOX2 regulated genes in Wounding from PMID: 30772301 | SOX2 |
| 3074 | SPIRE1 | SOX2 regulated genes in Wounding from PMID: 30772301 | SOX2 |
| 3075 | DCLK2 | SOX2 regulated genes in Wounding from PMID: 30772301 | SOX2 |
| 3076 | PRAG1 | SOX2 regulated genes in Wounding from PMID: 30772301 | SOX2 |
| 3077 | APLN | SOX2 regulated genes in Wounding from PMID: 30772301 | SOX2 |
| 3078 | GGT1 | SOX2 regulated genes in Wounding from PMID: 30772301 | SOX2 |
| 3079 | NEDD9 | SOX2 regulated genes in Wounding from PMID: 30772301 | SOX2 |
| 3080 | ERI1 | SOX2 regulated genes in Wounding from PMID: 30772301 | SOX2 |
| 3081 | PFN1 | SOX2 regulated genes in Wounding from PMID: 30772301 | SOX2 |
| 3082 | PGAP3 | SOX2 regulated genes in Wounding from PMID: 30772301 | SOX2 |
| 3083 | BCR | SOX2 regulated genes in Wounding from PMID: 30772301 | SOX2 |
| 3084 | ATP12A | SOX2 regulated genes in Wounding from PMID: 30772301 | SOX2 |
| 3085 | PROCR | SOX2 regulated genes in Wounding from PMID: 30772301 | SOX2 |
| 3086 | SEMA7A | SOX2 regulated genes in Wounding from PMID: 30772301 | SOX2 |
| 3087 | EFNB2 | SOX2 regulated genes in Wounding from PMID: 30772301 | SOX2 |
| 3088 | TMEM154 | SOX2 regulated genes in Wounding from PMID: 30772301 | SOX2 |
| 3089 | ANXA1 | SOX2 regulated genes in Wounding from PMID: 30772301 | SOX2 |
| 3090 | SRSF6 | SOX2 regulated genes in Wounding from PMID: 30772301 | SOX2 |
| 3091 | ARHGDIG | SOX2 regulated genes in Wounding from PMID: 30772301 | SOX2 |
| 3092 | FOXQ1 | SOX2 regulated genes in Wounding from PMID: 30772301 | SOX2 |
| 3093 | MBOAT1 | SOX2 regulated genes in Wounding from PMID: 30772301 | SOX2 |
| 3094 | GSPT1 | SOX2 regulated genes in Wounding from PMID: 30772301 | SOX2 |
| 3095 | GPR180 | SOX2 regulated genes in Wounding from PMID: 30772301 | SOX2 |
| 3096 | TMEM98 | SOX2 regulated genes in Wounding from PMID: 30772301 | SOX2 |
| 3097 | LRRC75B | SOX2 regulated genes in Wounding from PMID: 30772301 | SOX2 |
| 3098 | LPIN3 | SOX2 regulated genes in Wounding from PMID: 30772301 | SOX2 |
| 3099 | ATP6V1E1 | SOX2 regulated genes in Wounding from PMID: 30772301 | SOX2 |
| 3100 | TMPRSS2 | SOX2 regulated genes in Wounding from PMID: 30772301 | SOX2 |
| 3101 | ABRACL | SOX2 regulated genes in Wounding from PMID: 30772301 | SOX2 |
| 3102 | ITPRIP | SOX2 regulated genes in Wounding from PMID: 30772301 | SOX2 |
| 3103 | CPNE8 | SOX2 regulated genes in Wounding from PMID: 30772301 | SOX2 |
| 3104 | EBPL | SOX2 regulated genes in Wounding from PMID: 30772301 | SOX2 |
| 3105 | MYO10 | SOX2 regulated genes in Wounding from PMID: 30772301 | SOX2 |
| 3106 | SPIN1 | SOX2 regulated genes in Wounding from PMID: 30772301 | SOX2 |
| 3107 | SCML4 | SOX2 regulated genes in Wounding from PMID: 30772301 | SOX2 |
| 3108 | PARD6G | SOX2 regulated genes in Wounding from PMID: 30772301 | SOX2 |
| 3109 | CDC6 | SOX2 regulated genes in Wounding from PMID: 30772301 | SOX2 |

| | | | |
|---|---|---|---|
| 3110 | TBC1D2 | SOX2 regulated genes in Wounding from PMID: 30772301 | SOX2 |
| 3111 | CDKL2 | SOX2 regulated genes in Wounding from PMID: 30772301 | SOX2 |
| 3112 | HIRA | SOX2 regulated genes in Wounding from PMID: 30772301 | SOX2 |
| 3113 | CNN2 | SOX2 regulated genes in Wounding from PMID: 30772301 | SOX2 |
| 3114 | GATAD1 | SOX2 regulated genes in Wounding from PMID: 30772301 | SOX2 |
| 3115 | PORCN | SOX2 regulated genes in Wounding from PMID: 30772301 | SOX2 |
| 3116 | SENP6 | SOX2 regulated genes in Wounding from PMID: 30772301 | SOX2 |
| 3117 | DIS3 | SOX2 regulated genes in Wounding from PMID: 30772301 | SOX2 |
| 3118 | HBS1L | SOX2 regulated genes in Wounding from PMID: 30772301 | SOX2 |
| 3119 | HSBP1L1 | SOX2 regulated genes in Wounding from PMID: 30772301 | SOX2 |
| 3120 | CXXC4 | SOX2 regulated genes in Wounding from PMID: 30772301 | SOX2 |
| 3121 | CREB3 | SOX2 regulated genes in Wounding from PMID: 30772301 | SOX2 |
| 3122 | FEM1B | SOX2 regulated genes in Wounding from PMID: 30772301 | SOX2 |
| 3123 | STRAP | SOX2 regulated genes in Wounding from PMID: 30772301 | SOX2 |
| 3124 | TADA2A | SOX2 regulated genes in Wounding from PMID: 30772301 | SOX2 |
| 3125 | ITPR3 | SOX2 regulated genes in Wounding from PMID: 30772301 | SOX2 |
| 3126 | MYBL2 | SOX2 regulated genes in Wounding from PMID: 30772301 | SOX2 |
| 3127 | SYT7 | SOX2 regulated genes in Wounding from PMID: 30772301 | SOX2 |
| 3128 | PLXNA3 | SOX2 regulated genes in Wounding from PMID: 30772301 | SOX2 |
| 3129 | FAM20C | SOX2 regulated genes in Wounding from PMID: 30772301 | SOX2 |
| 3130 | SEC14L2 | SOX2 regulated genes in Wounding from PMID: 30772301 | SOX2 |
| 3131 | RWDD4 | SOX2 regulated genes in Wounding from PMID: 30772301 | SOX2 |
| 3132 | BICD2 | SOX2 regulated genes in Wounding from PMID: 30772301 | SOX2 |
| 3133 | HSP90AA1 | SOX2 regulated genes in Wounding from PMID: 30772301 | SOX2 |
| 3134 | FARP1 | SOX2 regulated genes in Wounding from PMID: 30772301 | SOX2 |
| 3135 | KREMEN1 | SOX2 regulated genes in Wounding from PMID: 30772301 | SOX2 |
| 3136 | SMAP1 | SOX2 regulated genes in Wounding from PMID: 30772301 | SOX2 |
| 3137 | CAAP1 | SOX2 regulated genes in Wounding from PMID: 30772301 | SOX2 |
| 3138 | MMP9 | SOX2 regulated genes in Wounding from PMID: 30772301 | SOX2 |
| 3139 | RHOV | SOX2 regulated genes in Wounding from PMID: 30772301 | SOX2 |
| 3140 | THBS1 | SOX2 regulated genes in Wounding from PMID: 30772301 | SOX2 |
| 3141 | FANCF | SOX2 regulated genes in Wounding from PMID: 30772301 | SOX2 |
| 3142 | PCDH19 | SOX2 regulated genes in Wounding from PMID: 30772301 | SOX2 |
| 3143 | BDKRB1 | SOX2 regulated genes in Wounding from PMID: 30772301 | SOX2 |
| 3144 | COG6 | SOX2 regulated genes in Wounding from PMID: 30772301 | SOX2 |
| 3145 | PAWR | SOX2 regulated genes in Wounding from PMID: 30772301 | SOX2 |
| 3146 | DUSP14 | SOX2 regulated genes in Wounding from PMID: 30772301 | SOX2 |
| 3147 | NCKAP5L | SOX2 regulated genes in Wounding from PMID: 30772301 | SOX2 |
| 3148 | GP1BB | SOX2 regulated genes in Wounding from PMID: 30772301 | SOX2 |
| 3149 | CCDC120 | SOX2 regulated genes in Wounding from PMID: 30772301 | SOX2 |
| 3150 | HSP90AB1 | SOX2 regulated genes in Wounding from PMID: 30772301 | SOX2 |
| 3151 | CCDC25 | SOX2 regulated genes in Wounding from PMID: 30772301 | SOX2 |
| 3152 | SEC63 | SOX2 regulated genes in Wounding from PMID: 30772301 | SOX2 |
| 3153 | CCDC116 | SOX2 regulated genes in Wounding from PMID: 30772301 | SOX2 |
| 3154 | ENTPD7 | SOX2 regulated genes in Wounding from PMID: 30772301 | SOX2 |
| 3155 | LNX1 | SOX2 regulated genes in Wounding from PMID: 30772301 | SOX2 |
| 3156 | SMAD7 | SOX2 regulated genes in Wounding from PMID: 30772301 | SOX2 |
| 3157 | ACE2 | SOX2 regulated genes in Wounding from PMID: 30772301 | SOX2 |
| 3158 | ARID2 | SOX2 regulated genes in Wounding from PMID: 30772301 | SOX2 |
| 3159 | ARHGAP24 | SOX2 regulated genes in Wounding from PMID: 30772301 | SOX2 |
| 3160 | DEPDC7 | SOX2 regulated genes in Wounding from PMID: 30772301 | SOX2 |
| 3161 | CLNS1A | SOX2 regulated genes in Wounding from PMID: 30772301 | SOX2 |
| 3162 | GSTCD | SOX2 regulated genes in Wounding from PMID: 30772301 | SOX2 |
| 3163 | DERA | SOX2 regulated genes in Wounding from PMID: 30772301 | SOX2 |
| 3164 | PRICKLE1 | SOX2 regulated genes in Wounding from PMID: 30772301 | SOX2 |
| 3165 | CAMSAP1 | SOX2 regulated genes in Wounding from PMID: 30772301 | SOX2 |
| 3166 | RERG | SOX2 regulated genes in Wounding from PMID: 30772301 | SOX2 |
| 3167 | GFOD1 | SOX2 regulated genes in Wounding from PMID: 30772301 | SOX2 |
| 3168 | TMPRSS11E | SOX2 regulated genes in Wounding from PMID: 30772301 | SOX2 |
| 3169 | MORC2 | SOX2 regulated genes in Wounding from PMID: 30772301 | SOX2 |
| 3170 | NIPSNAP1 | SOX2 regulated genes in Wounding from PMID: 30772301 | SOX2 |

| | | | |
|---|---|---|---|
| 3171 | GMEB2 | SOX2 regulated genes in Wounding from PMID: 30772301 | SOX2 |
| 3172 | FOXE1 | SOX2 regulated genes in Wounding from PMID: 30772301 | SOX2 |
| 3173 | RUNX2 | SOX2 regulated genes in Wounding from PMID: 30772301 | SOX2 |
| 3174 | C10orf88 | SOX2 regulated genes in Wounding from PMID: 30772301 | SOX2 |
| 3175 | DPH6 | SOX2 regulated genes in Wounding from PMID: 30772301 | SOX2 |
| 3176 | TBC1D10A | SOX2 regulated genes in Wounding from PMID: 30772301 | SOX2 |
| 3177 | RFLNB | SOX2 regulated genes in Wounding from PMID: 30772301 | SOX2 |
| 3178 | MINPP1 | SOX2 regulated genes in Wounding from PMID: 30772301 | SOX2 |
| 3179 | CD109 | SOX2 regulated genes in Wounding from PMID: 30772301 | SOX2 |
| 3180 | MTRR | SOX2 regulated genes in Wounding from PMID: 30772301 | SOX2 |
| 3181 | NXN | SOX2 regulated genes in Wounding from PMID: 30772301 | SOX2 |
| 3182 | SMOC1 | SOX2 regulated genes in Wounding from PMID: 30772301 | SOX2 |
| 3183 | TCF7L2 | SOX2 regulated genes in Wounding from PMID: 30772301 | SOX2 |
| 3184 | CRELD2 | SOX2 regulated genes in Wounding from PMID: 30772301 | SOX2 |
| 3185 | ATP6V1B2 | SOX2 regulated genes in Wounding from PMID: 30772301 | SOX2 |
| 3186 | CISD2 | SOX2 regulated genes in Wounding from PMID: 30772301 | SOX2 |
| 3187 | TMPRSS11F | SOX2 regulated genes in Wounding from PMID: 30772301 | SOX2 |
| 3188 | GCLC | SOX2 regulated genes in Wounding from PMID: 30772301 | SOX2 |
| 3189 | SPATA18 | SOX2 regulated genes in Wounding from PMID: 30772301 | SOX2 |
| 3190 | GBA2 | SOX2 regulated genes in Wounding from PMID: 30772301 | SOX2 |
| 3191 | CPXM2 | SOX2 regulated genes in Wounding from PMID: 30772301 | SOX2 |
| 3192 | RNF182 | SOX2 regulated genes in Wounding from PMID: 30772301 | SOX2 |
| 3193 | CDK5R1 | SOX2 regulated genes in Wounding from PMID: 30772301 | SOX2 |
| 3194 | AFAP1L2 | SOX2 regulated genes in Wounding from PMID: 30772301 | SOX2 |
| 3195 | PDLIM1 | SOX2 regulated genes in Wounding from PMID: 30772301 | SOX2 |
| 3196 | HDAC2 | SOX2 regulated genes in Wounding from PMID: 30772301 | SOX2 |
| 3197 | KCNE1 | SOX2 regulated genes in Wounding from PMID: 30772301 | SOX2 |
| 3198 | RCN2 | SOX2 regulated genes in Wounding from PMID: 30772301 | SOX2 |
| 3199 | PRSS12 | SOX2 regulated genes in Wounding from PMID: 30772301 | SOX2 |
| 3200 | CLU | SOX2 regulated genes in Wounding from PMID: 30772301 | SOX2 |
| 3201 | MICAL3 | SOX2 regulated genes in Wounding from PMID: 30772301 | SOX2 |
| 3202 | CYP39A1 | SOX2 regulated genes in Wounding from PMID: 30772301 | SOX2 |
| 3203 | SLC15A1 | SOX2 regulated genes in Wounding from PMID: 30772301 | SOX2 |
| 3204 | SLC25A37 | SOX2 regulated genes in Wounding from PMID: 30772301 | SOX2 |
| 3205 | SERPINB5 | SOX2 regulated genes in Wounding from PMID: 30772301 | SOX2 |
| 3206 | SMAD6 | SOX2 regulated genes in Wounding from PMID: 30772301 | SOX2 |
| 3207 | HDDC2 | SOX2 regulated genes in Wounding from PMID: 30772301 | SOX2 |
| 3208 | PAQR8 | SOX2 regulated genes in Wounding from PMID: 30772301 | SOX2 |
| 3209 | TCFL5 | SOX2 regulated genes in Wounding from PMID: 30772301 | SOX2 |
| 3210 | HACE1 | SOX2 regulated genes in Wounding from PMID: 30772301 | SOX2 |
| 3211 | LRAT | SOX2 regulated genes in Wounding from PMID: 30772301 | SOX2 |
| 3212 | CCDC169 | SOX2 regulated genes in Wounding from PMID: 30772301 | SOX2 |
| 3213 | RASSF6 | SOX2 regulated genes in Wounding from PMID: 30772301 | SOX2 |
| 3214 | ENTPD8 | SOX2 regulated genes in Wounding from PMID: 30772301 | SOX2 |
| 3215 | STYK1 | SOX2 regulated genes in Wounding from PMID: 30772301 | SOX2 |
| 3216 | RANGAP1 | SOX2 regulated genes in Wounding from PMID: 30772301 | SOX2 |
| 3217 | DDX27 | SOX2 regulated genes in Wounding from PMID: 30772301 | SOX2 |
| 3218 | PLCXD1 | SOX2 regulated genes in Wounding from PMID: 30772301 | SOX2 |
| 3219 | SLC6A8 | SOX2 regulated genes in Wounding from PMID: 30772301 | SOX2 |
| 3220 | SHROOM3 | SOX2 regulated genes in Wounding from PMID: 30772301 | SOX2 |
| 3221 | RANBP1 | SOX2 regulated genes in Wounding from PMID: 30772301 | SOX2 |
| 3222 | RIOK1 | SOX2 regulated genes in Wounding from PMID: 30772301 | SOX2 |
| 3223 | FOXC1 | SOX2 regulated genes in Wounding from PMID: 30772301 | SOX2 |
| 3224 | AK7 | SOX2 regulated genes in Wounding from PMID: 30772301 | SOX2 |
| 3225 | CWH43 | SOX2 regulated genes in Wounding from PMID: 30772301 | SOX2 |
| 3226 | PROSER2 | SOX2 regulated genes in Wounding from PMID: 30772301 | SOX2 |
| 3227 | PALLD | SOX2 regulated genes in Wounding from PMID: 30772301 | SOX2 |
| 3228 | FAM83C | SOX2 regulated genes in Wounding from PMID: 30772301 | SOX2 |
| 3229 | TDRD3 | SOX2 regulated genes in Wounding from PMID: 30772301 | SOX2 |
| 3230 | MAPK11 | SOX2 regulated genes in Wounding from PMID: 30772301 | SOX2 |
| 3231 | LAMA3 | SOX2 regulated genes in Wounding from PMID: 30772301 | SOX2 |

| | | | |
|---|---|---|---|
| 3232 | AJUBA | SOX2 regulated genes in Wounding from PMID: 30772301 | SOX2 |
| 3233 | SERPINB13 | SOX2 regulated genes in Wounding from PMID: 30772301 | SOX2 |
| 3234 | ABTB2 | SOX2 regulated genes in Wounding from PMID: 30772301 | SOX2 |
| 3235 | DBNDD2 | SOX2 regulated genes in Wounding from PMID: 30772301 | SOX2 |
| 3236 | TM9SF3 | SOX2 regulated genes in Wounding from PMID: 30772301 | SOX2 |
| 3237 | MORF4L2 | SOX2 regulated genes in Wounding from PMID: 30772301 | SOX2 |
| 3238 | MAPK1 | SOX2 regulated genes in Wounding from PMID: 30772301 | SOX2 |
| 3239 | DLST | SOX2 regulated genes in Wounding from PMID: 30772301 | SOX2 |
| 3240 | SWI5 | SOX2 regulated genes in Wounding from PMID: 30772301 | SOX2 |
| 3241 | MAGOHB | SOX2 regulated genes in Wounding from PMID: 30772301 | SOX2 |
| 3242 | RAD23B | SOX2 regulated genes in Wounding from PMID: 30772301 | SOX2 |
| 3243 | GPRIN3 | SOX2 regulated genes in Wounding from PMID: 30772301 | SOX2 |
| 3244 | IRX4 | SOX2 regulated genes in Wounding from PMID: 30772301 | SOX2 |
| 3245 | BMP6 | SOX2 regulated genes in Wounding from PMID: 30772301 | SOX2 |
| 3246 | NAA35 | SOX2 regulated genes in Wounding from PMID: 30772301 | SOX2 |
| 3247 | NUS1 | SOX2 regulated genes in Wounding from PMID: 30772301 | SOX2 |
| 3248 | SUCLA2 | SOX2 regulated genes in Wounding from PMID: 30772301 | SOX2 |
| 3249 | DSG2 | SOX2 regulated genes in Wounding from PMID: 30772301 | SOX2 |
| 3250 | CPNE5 | SOX2 regulated genes in Wounding from PMID: 30772301 | SOX2 |
| 3251 | CRK | SOX2 regulated genes in Wounding from PMID: 30772301 | SOX2 |
| 3252 | RCAN1 | SOX2 regulated genes in Wounding from PMID: 30772301 | SOX2 |
| 3253 | FLNA | SOX2 regulated genes in Wounding from PMID: 30772301 | SOX2 |
| 3254 | FJX1 | SOX2 regulated genes in Wounding from PMID: 30772301 | SOX2 |
| 3255 | B4GALNT3 | SOX2 regulated genes in Wounding from PMID: 30772301 | SOX2 |
| 3256 | CA12 | SOX2 regulated genes in Wounding from PMID: 30772301 | SOX2 |
| 3257 | SNX12 | SOX2 regulated genes in Wounding from PMID: 30772301 | SOX2 |
| 3258 | KCTD9 | SOX2 regulated genes in Wounding from PMID: 30772301 | SOX2 |
| 3259 | MEOX1 | SOX2 regulated genes in Wounding from PMID: 30772301 | SOX2 |
| 3260 | FERMT1 | SOX2 regulated genes in Wounding from PMID: 30772301 | SOX2 |
| 3261 | EDN1 | SOX2 regulated genes in Wounding from PMID: 30772301 | SOX2 |
| 3262 | PLP2 | SOX2 regulated genes in Wounding from PMID: 30772301 | SOX2 |
| 3263 | FAM199X | SOX2 regulated genes in Wounding from PMID: 30772301 | SOX2 |
| 3264 | L3MBTL1 | SOX2 regulated genes in Wounding from PMID: 30772301 | SOX2 |
| 3265 | SPRYD3 | SOX2 regulated genes in Wounding from PMID: 30772301 | SOX2 |
| 3266 | TFIP11 | SOX2 regulated genes in Wounding from PMID: 30772301 | SOX2 |
| 3267 | GOPC | SOX2 regulated genes in Wounding from PMID: 30772301 | SOX2 |
| 3268 | TMEFF1 | SOX2 regulated genes in Wounding from PMID: 30772301 | SOX2 |
| 3269 | DSC3 | SOX2 regulated genes in Wounding from PMID: 30772301 | SOX2 |
| 3270 | NFIL3 | SOX2 regulated genes in Wounding from PMID: 30772301 | SOX2 |
| 3271 | NUDT15 | SOX2 regulated genes in Wounding from PMID: 30772301 | SOX2 |
| 3272 | ARHGAP1 | SOX2 regulated genes in Wounding from PMID: 30772301 | SOX2 |
| 3273 | ST8SIA6 | SOX2 regulated genes in Wounding from PMID: 30772301 | SOX2 |
| 3274 | SLC24A3 | SOX2 regulated genes in Wounding from PMID: 30772301 | SOX2 |
| 3275 | PAK3 | SOX2 regulated genes in Wounding from PMID: 30772301 | SOX2 |
| 3276 | PLAGL2 | SOX2 regulated genes in Wounding from PMID: 30772301 | SOX2 |
| 3277 | LGR4 | SOX2 regulated genes in Wounding from PMID: 30772301 | SOX2 |
| 3278 | FHOD3 | SOX2 regulated genes in Wounding from PMID: 30772301 | SOX2 |
| 3279 | SNU13 | SOX2 regulated genes in Wounding from PMID: 30772301 | SOX2 |
| 3280 | KDM1B | SOX2 regulated genes in Wounding from PMID: 30772301 | SOX2 |
| 3281 | GOLGA7B | SOX2 regulated genes in Wounding from PMID: 30772301 | SOX2 |
| 3282 | BTBD9 | SOX2 regulated genes in Wounding from PMID: 30772301 | SOX2 |
| 3283 | CHAC1 | SOX2 regulated genes in Wounding from PMID: 30772301 | SOX2 |
| 3284 | ID4 | SOX2 regulated genes in Wounding from PMID: 30772301 | SOX2 |
| 3285 | PPDPF | SOX2 regulated genes in Wounding from PMID: 30772301 | SOX2 |
| 3286 | CSTF3 | SOX2 regulated genes in Wounding from PMID: 30772301 | SOX2 |
| 3287 | RAP1GAP2 | SOX2 regulated genes in Wounding from PMID: 30772301 | SOX2 |
| 3288 | LMAN1L | SOX2 regulated genes in Wounding from PMID: 30772301 | SOX2 |
| 3289 | ANG | SOX2 regulated genes in Wounding from PMID: 30772301 | SOX2 |
| 3290 | MED27 | SOX2 regulated genes in Wounding from PMID: 30772301 | SOX2 |
| 3291 | EIF3A | SOX2 regulated genes in Wounding from PMID: 30772301 | SOX2 |
| 3292 | LMTK2 | SOX2 regulated genes in Wounding from PMID: 30772301 | SOX2 |

| | | | |
|---|---|---|---|
| 3293 | SERPINE1 | SOX2 regulated genes in Wounding from PMID: 30772301 | SOX2 |
| 3294 | ENDOD1 | SOX2 regulated genes in Wounding from PMID: 30772301 | SOX2 |
| 3295 | DLG5 | SOX2 regulated genes in Wounding from PMID: 30772301 | SOX2 |
| 3296 | ARIH1 | SOX2 regulated genes in Wounding from PMID: 30772301 | SOX2 |
| 3297 | SF3A1 | SOX2 regulated genes in Wounding from PMID: 30772301 | SOX2 |
| 3298 | HSPA4L | SOX2 regulated genes in Wounding from PMID: 30772301 | SOX2 |
| 3299 | PPP1R3B | SOX2 regulated genes in Wounding from PMID: 30772301 | SOX2 |
| 3300 | PPP2R1B | SOX2 regulated genes in Wounding from PMID: 30772301 | SOX2 |
| 3301 | NEDD4L | SOX2 regulated genes in Wounding from PMID: 30772301 | SOX2 |
| 3302 | IPPK | SOX2 regulated genes in Wounding from PMID: 30772301 | SOX2 |
| 3303 | THOC1 | SOX2 regulated genes in Wounding from PMID: 30772301 | SOX2 |
| 3304 | OVOL2 | SOX2 regulated genes in Wounding from PMID: 30772301 | SOX2 |
| 3305 | LEPROTL1 | SOX2 regulated genes in Wounding from PMID: 30772301 | SOX2 |
| 3306 | TCEAL3 | SOX2 regulated genes in Wounding from PMID: 30772301 | SOX2 |
| 3307 | SLC30A9 | SOX2 regulated genes in Wounding from PMID: 30772301 | SOX2 |
| 3308 | CHIC2 | SOX2 regulated genes in Wounding from PMID: 30772301 | SOX2 |
| 3309 | DSG1 | SOX2 regulated genes in Wounding from PMID: 30772301 | SOX2 |
| 3310 | PRDM1 | SOX2 regulated genes in Wounding from PMID: 30772301 | SOX2 |
| 3311 | LIN7C | SOX2 regulated genes in Wounding from PMID: 30772301 | SOX2 |
| 3312 | COL17A1 | SOX2 regulated genes in Wounding from PMID: 30772301 | SOX2 |
| 3313 | GCH1 | SOX2 regulated genes in Wounding from PMID: 30772301 | SOX2 |
| 3314 | LONRF1 | SOX2 regulated genes in Wounding from PMID: 30772301 | SOX2 |
| 3315 | EVC2 | SOX2 regulated genes in Wounding from PMID: 30772301 | SOX2 |
| 3316 | MUC15 | SOX2 regulated genes in Wounding from PMID: 30772301 | SOX2 |
| 3317 | MYO1E | SOX2 regulated genes in Wounding from PMID: 30772301 | SOX2 |
| 3318 | BTC | SOX2 regulated genes in Wounding from PMID: 30772301 | SOX2 |
| 3319 | BCORL1 | SOX2 regulated genes in Wounding from PMID: 30772301 | SOX2 |
| 3320 | PDCL | SOX2 regulated genes in Wounding from PMID: 30772301 | SOX2 |
| 3321 | C15orf62 | SOX2 regulated genes in Wounding from PMID: 30772301 | SOX2 |
| 3322 | CRNKL1 | SOX2 regulated genes in Wounding from PMID: 30772301 | SOX2 |
| 3323 | RRP7A | SOX2 regulated genes in Wounding from PMID: 30772301 | SOX2 |
| 3324 | PLEKHA7 | SOX2 regulated genes in Wounding from PMID: 30772301 | SOX2 |
| 3325 | COBL | SOX2 regulated genes in Wounding from PMID: 30772301 | SOX2 |
| 3326 | BCOR | SOX2 regulated genes in Wounding from PMID: 30772301 | SOX2 |
| 3327 | GPAT3 | SOX2 regulated genes in Wounding from PMID: 30772301 | SOX2 |
| 3328 | CTDP1 | SOX2 regulated genes in Wounding from PMID: 30772301 | SOX2 |
| 3329 | DHX32 | SOX2 regulated genes in Wounding from PMID: 30772301 | SOX2 |
| 3330 | SLK | SOX2 regulated genes in Wounding from PMID: 30772301 | SOX2 |
| 3331 | CSTF2 | SOX2 regulated genes in Wounding from PMID: 30772301 | SOX2 |
| 3332 | LIG3 | SOX2 regulated genes in Wounding from PMID: 30772301 | SOX2 |
| 3333 | EEF1G | SOX2 regulated genes in Wounding from PMID: 30772301 | SOX2 |
| 3334 | CFAP46 | SOX2 regulated genes in Wounding from PMID: 30772301 | SOX2 |
| 3335 | PPP1R14C | SOX2 regulated genes in Wounding from PMID: 30772301 | SOX2 |
| 3336 | STK26 | SOX2 regulated genes in Wounding from PMID: 30772301 | SOX2 |
| 3337 | MEI1 | SOX2 regulated genes in Wounding from PMID: 30772301 | SOX2 |
| 3338 | TLL1 | SOX2 regulated genes in Wounding from PMID: 30772301 | SOX2 |
| 3339 | TMTC4 | SOX2 regulated genes in Wounding from PMID: 30772301 | SOX2 |
| 3340 | GTF2E2 | SOX2 regulated genes in Wounding from PMID: 30772301 | SOX2 |
| 3341 | RHOD | SOX2 regulated genes in Wounding from PMID: 30772301 | SOX2 |
| 3342 | HEPHL1 | SOX2 regulated genes in Wounding from PMID: 30772301 | SOX2 |
| 3343 | ERVFRD-1 | SOX2 regulated genes in Wounding from PMID: 30772301 | SOX2 |
| 3344 | RAB11FIP1 | SOX2 regulated genes in Wounding from PMID: 30772301 | SOX2 |
| 3345 | MAPRE1 | SOX2 regulated genes in Wounding from PMID: 30772301 | SOX2 |
| 3346 | CDCA4 | SOX2 regulated genes in Wounding from PMID: 30772301 | SOX2 |
| 3347 | USP22 | SOX2 regulated genes in Wounding from PMID: 30772301 | SOX2 |
| 3348 | TOP1 | SOX2 regulated genes in Wounding from PMID: 30772301 | SOX2 |
| 3349 | TSPAN14 | SOX2 regulated genes in Wounding from PMID: 30772301 | SOX2 |
| 3350 | TTPAL | SOX2 regulated genes in Wounding from PMID: 30772301 | SOX2 |
| 3351 | VPS4B | SOX2 regulated genes in Wounding from PMID: 30772301 | SOX2 |
| 3352 | XPNPEP1 | SOX2 regulated genes in Wounding from PMID: 30772301 | SOX2 |
| 3353 | ZMYM5 | SOX2 regulated genes in Wounding from PMID: 30772301 | SOX2 |

| | | | |
|---|---|---|---|
| 3354 | WIPF2 | SOX2 regulated genes in Wounding from PMID: 30772301 | SOX2 |
| 3355 | UTP14A | SOX2 regulated genes in Wounding from PMID: 30772301 | SOX2 |
| 3356 | ZNF407 | SOX2 regulated genes in Wounding from PMID: 30772301 | SOX2 |
| 3357 | TOLLIP | SOX2 regulated genes in Wounding from PMID: 30772301 | SOX2 |
| 3358 | USP6 | SOX2 regulated genes in Wounding from PMID: 30772301 | SOX2 |
| 3359 | TOM1 | SOX2 regulated genes in Wounding from PMID: 30772301 | SOX2 |
| 3360 | TRPM6 | SOX2 regulated genes in Wounding from PMID: 30772301 | SOX2 |
| 3361 | ZFAND2A | SOX2 regulated genes in Wounding from PMID: 30772301 | SOX2 |
| 3362 | UBE2A | SOX2 regulated genes in Wounding from PMID: 30772301 | SOX2 |
| 3363 | TUBB6 | SOX2 regulated genes in Wounding from PMID: 30772301 | SOX2 |
| 3364 | TRIM29 | SOX2 regulated genes in Wounding from PMID: 30772301 | SOX2 |
| 3365 | ZGPAT | SOX2 regulated genes in Wounding from PMID: 30772301 | SOX2 |
| 3366 | ZMYND19 | SOX2 regulated genes in Wounding from PMID: 30772301 | SOX2 |
| 3367 | YWHAH | SOX2 regulated genes in Wounding from PMID: 30772301 | SOX2 |
| 3368 | TULP3 | SOX2 regulated genes in Wounding from PMID: 30772301 | SOX2 |
| 3369 | TRIL | SOX2 regulated genes in Wounding from PMID: 30772301 | SOX2 |
| 3370 | TRAPPC8 | SOX2 regulated genes in Wounding from PMID: 30772301 | SOX2 |
| 3371 | ZNF532 | SOX2 regulated genes in Wounding from PMID: 30772301 | SOX2 |
| 3372 | YAP1 | SOX2 regulated genes in Wounding from PMID: 30772301 | SOX2 |
| 3373 | ULK3 | SOX2 regulated genes in Wounding from PMID: 30772301 | SOX2 |
| 3374 | TXNDC5 | SOX2 regulated genes in Wounding from PMID: 30772301 | SOX2 |
| 3375 | WBP4 | SOX2 regulated genes in Wounding from PMID: 30772301 | SOX2 |
| 3376 | ZDHHC13 | SOX2 regulated genes in Wounding from PMID: 30772301 | SOX2 |
| 3377 | TSPAN9 | SOX2 regulated genes in Wounding from PMID: 30772301 | SOX2 |
| 3378 | TSPAN6 | SOX2 regulated genes in Wounding from PMID: 30772301 | SOX2 |
| 3379 | UNC5B | SOX2 regulated genes in Wounding from PMID: 30772301 | SOX2 |
| 3380 | UMODL1 | SOX2 regulated genes in Wounding from PMID: 30772301 | SOX2 |
| 3381 | TXNDC17 | SOX2 regulated genes in Wounding from PMID: 30772301 | SOX2 |
| 3382 | TOMM22 | SOX2 regulated genes in Wounding from PMID: 30772301 | SOX2 |
| 3383 | ZDHHC9 | SOX2 regulated genes in Wounding from PMID: 30772301 | SOX2 |
| 3384 | ZNRF2 | SOX2 regulated genes in Wounding from PMID: 30772301 | SOX2 |
| 3385 | AFAP1 | SOX2 regulated genes in Wounding from PMID: 30772301 | SOX2 |
| 3386 | CCDC112 | SOX2 regulated genes in Wounding from PMID: 30772301 | SOX2 |
| 3387 | SYNE2 | SOX2 regulated genes in Wounding from PMID: 30772301 | SOX2 |
| 3388 | FHDC1 | SOX2 regulated genes in Wounding from PMID: 30772301 | SOX2 |
| 3389 | CCNC | SOX2 regulated genes in Wounding from PMID: 30772301 | SOX2 |
| 3390 | SOCS5 | SOX2 regulated genes in Wounding from PMID: 30772301 | SOX2 |
| 3391 | NAP1L4 | SOX2 regulated genes in Wounding from PMID: 30772301 | SOX2 |
| 3392 | ITGA3 | SOX2 regulated genes in Wounding from PMID: 30772301 | SOX2 |
| 3393 | CLIC3 | SOX2 regulated genes in Wounding from PMID: 30772301 | SOX2 |
| 3394 | SERPINA1 | SOX2 regulated genes in Wounding from PMID: 30772301 | SOX2 |
| 3395 | AEBP2 | SOX2 regulated genes in Wounding from PMID: 30772301 | SOX2 |
| 3396 | EFCAB1 | SOX2 regulated genes in Wounding from PMID: 30772301 | SOX2 |
| 3397 | MSANTD3 | SOX2 regulated genes in Wounding from PMID: 30772301 | SOX2 |
| 3398 | DNAJB13 | SOX2 regulated genes in Wounding from PMID: 30772301 | SOX2 |
| 3399 | NCS1 | SOX2 regulated genes in Wounding from PMID: 30772301 | SOX2 |
| 3400 | NOTCH1 | SOX2 regulated genes in Wounding from PMID: 30772301 | SOX2 |
| 3401 | NUP62CL | SOX2 regulated genes in Wounding from PMID: 30772301 | SOX2 |
| 3402 | SMARCE1 | SOX2 regulated genes in Wounding from PMID: 30772301 | SOX2 |
| 3403 | GDPD5 | SOX2 regulated genes in Wounding from PMID: 30772301 | SOX2 |
| 3404 | CD151 | SOX2 regulated genes in Wounding from PMID: 30772301 | SOX2 |
| 3405 | MGAT4A | SOX2 regulated genes in Wounding from PMID: 30772301 | SOX2 |
| 3406 | FAM135A | SOX2 regulated genes in Wounding from PMID: 30772301 | SOX2 |
| 3407 | BEGAIN | SOX2 regulated genes in Wounding from PMID: 30772301 | SOX2 |
| 3408 | CDK17 | SOX2 regulated genes in Wounding from PMID: 30772301 | SOX2 |
| 3409 | BRAF | SOX2 regulated genes in Wounding from PMID: 30772301 | SOX2 |
| 3410 | ZNF597 | SOX2 regulated genes in Wounding from PMID: 30772301 | SOX2 |
| 3411 | UTP14C | SOX2 regulated genes in Wounding from PMID: 30772301 | SOX2 |
| 3412 | KRTAP9-4 | SOX2 regulated genes in Wounding from PMID: 30772301 | SOX2 |
| 3413 | EFL1 | SOX2 regulated genes in Wounding from PMID: 30772301 | SOX2 |
| 3414 | RHBDL2 | SOX2 regulated genes in Wounding from PMID: 30772301 | SOX2 |

| | | | |
|---|---|---|---|
| 3415 | UBE2N | SOX2 regulated genes in Wounding from PMID: 30772301 | SOX2 |
| 3416 | WAPL | SOX2 regulated genes in Wounding from PMID: 30772301 | SOX2 |
| 3417 | CTNNB1 | SOX2 regulated genes in Wounding from PMID: 30772301 | SOX2 |
| 3418 | SHB | SOX2 regulated genes in Wounding from PMID: 30772301 | SOX2 |
| 3419 | MYH9 | SOX2 regulated genes in Wounding from PMID: 30772301 | SOX2 |
| 3420 | SERINC2 | SOX2 regulated genes in Wounding from PMID: 30772301 | SOX2 |
| 3421 | ZNF664 | SOX2 regulated genes in Wounding from PMID: 30772301 | SOX2 |
| 3422 | OSBPL5 | SOX2 regulated genes in Wounding from PMID: 30772301 | SOX2 |
| 3423 | VPS35 | SOX2 regulated genes in Wounding from PMID: 30772301 | SOX2 |
| 3424 | SNX18 | SOX2 regulated genes in Wounding from PMID: 30772301 | SOX2 |
| 3425 | ATP11A | SOX2 regulated genes in Wounding from PMID: 30772301 | SOX2 |
| 3426 | PRSS8 | SOX2 regulated genes in Wounding from PMID: 30772301 | SOX2 |
| 3427 | CPEB4 | SOX2 regulated genes in Wounding from PMID: 30772301 | SOX2 |
| 3428 | PLEK2 | SOX2 regulated genes in Wounding from PMID: 30772301 | SOX2 |
| 3429 | BICDL2 | SOX2 regulated genes in Wounding from PMID: 30772301 | SOX2 |
| 3430 | LEO1 | SOX2 regulated genes in Wounding from PMID: 30772301 | SOX2 |
| 3431 | NOCT | SOX2 regulated genes in Wounding from PMID: 30772301 | SOX2 |
| 3432 | UBAP1 | SOX2 regulated genes in Wounding from PMID: 30772301 | SOX2 |
| 3433 | RAB23 | SOX2 regulated genes in Wounding from PMID: 30772301 | SOX2 |
| 3434 | GUCA1B | SOX2 regulated genes in Wounding from PMID: 30772301 | SOX2 |
| 3435 | ZDHHC5 | SOX2 regulated genes in Wounding from PMID: 30772301 | SOX2 |
| 3436 | BTBD11 | SOX2 regulated genes in Wounding from PMID: 30772301 | SOX2 |
| 3437 | NCOR2 | SOX2 regulated genes in Wounding from PMID: 30772301 | SOX2 |
| 3438 | TMEM51 | SOX2 regulated genes in Wounding from PMID: 30772301 | SOX2 |
| 3439 | TGFA | SOX2 regulated genes in Wounding from PMID: 30772301 | SOX2 |
| 3440 | LPAR6 | SOX2 regulated genes in Wounding from PMID: 30772301 | SOX2 |
| 3441 | MPHOSPH6 | SOX2 regulated genes in Wounding from PMID: 30772301 | SOX2 |
| 3442 | MICB | SOX2 regulated genes in Wounding from PMID: 30772301 | SOX2 |
| 3443 | SLC39A4 | SOX2 regulated genes in Wounding from PMID: 30772301 | SOX2 |
| 3444 | TMEM151A | SOX2 regulated genes in Wounding from PMID: 30772301 | SOX2 |
| 3445 | TPM4 | SOX2 regulated genes in Wounding from PMID: 30772301 | SOX2 |
| 3446 | DGKA | SOX2 regulated genes in Wounding from PMID: 30772301 | SOX2 |
| 3447 | TGM7 | SOX2 regulated genes in Wounding from PMID: 30772301 | SOX2 |
| 3448 | DDX17 | SOX2 regulated genes in Wounding from PMID: 30772301 | SOX2 |
| 3449 | ZNF365 | SOX2 regulated genes in Wounding from PMID: 30772301 | SOX2 |
| 3450 | KAZN | SOX2 regulated genes in Wounding from PMID: 30772301 | SOX2 |
| 3451 | CD2BP2 | SOX2 regulated genes in Wounding from PMID: 30772301 | SOX2 |
| 3452 | BMP3 | SOX2 regulated genes in Wounding from PMID: 30772301 | SOX2 |
| 3453 | GNL2 | SOX2 regulated genes in Wounding from PMID: 30772301 | SOX2 |
| 3454 | SFR1 | SOX2 regulated genes in Wounding from PMID: 30772301 | SOX2 |
| 3455 | AREG | SOX2 regulated genes in Wounding from PMID: 30772301 | SOX2 |
| 3456 | DUSP5 | SOX2 regulated genes in Wounding from PMID: 30772301 | SOX2 |
| 3457 | CUX1 | SOX2 regulated genes in Wounding from PMID: 30772301 | SOX2 |
| 3458 | CPE | SOX2 regulated genes in Wounding from PMID: 30772301 | SOX2 |
| 3459 | ESRP2 | SOX2 regulated genes in Wounding from PMID: 30772301 | SOX2 |
| 3460 | CA9 | SOX2 regulated genes in Wounding from PMID: 30772301 | SOX2 |
| 3461 | PRKCH | SOX2 regulated genes in Wounding from PMID: 30772301 | SOX2 |
| 3462 | PAK1 | SOX2 regulated genes in Wounding from PMID: 30772301 | SOX2 |
| 3463 | CPA4 | SOX2 regulated genes in Wounding from PMID: 30772301 | SOX2 |
| 3464 | SNAI2 | SOX2 regulated genes in Wounding from PMID: 30772301 | SOX2 |
| 3465 | DNAJB6 | SOX2 regulated genes in Wounding from PMID: 30772301 | SOX2 |
| 3466 | BAG2 | SOX2 regulated genes in Wounding from PMID: 30772301 | SOX2 |
| 3467 | KRT72 | SOX2 regulated genes in Wounding from PMID: 30772301 | SOX2 |
| 3468 | DDX18 | SOX2 regulated genes in Wounding from PMID: 30772301 | SOX2 |
| 3469 | PUM3 | SOX2 regulated genes in Wounding from PMID: 30772301 | SOX2 |
| 3470 | REPS1 | SOX2 regulated genes in Wounding from PMID: 30772301 | SOX2 |
| 3471 | CDK4 | SOX2 regulated genes in Wounding from PMID: 30772301 | SOX2 |
| 3472 | FBXW8 | SOX2 regulated genes in Wounding from PMID: 30772301 | SOX2 |
| 3473 | HAS2 | SOX2 regulated genes in Wounding from PMID: 30772301 | SOX2 |
| 3474 | SYNJ2 | SOX2 regulated genes in Wounding from PMID: 30772301 | SOX2 |
| 3475 | FHL2 | SOX2 regulated genes in Wounding from PMID: 30772301 | SOX2 |

| | | | |
|---|---|---|---|
| 3476 | GALR3 | SOX2 regulated genes in Wounding from PMID: 30772301 | SOX2 |
| 3477 | PPP1R37 | SOX2 regulated genes in Wounding from PMID: 30772301 | SOX2 |
| 3478 | UBE2Z | SOX2 regulated genes in Wounding from PMID: 30772301 | SOX2 |
| 3479 | RIOX2 | SOX2 regulated genes in Wounding from PMID: 30772301 | SOX2 |
| 3480 | MAPKAPK3 | SOX2 regulated genes in Wounding from PMID: 30772301 | SOX2 |
| 3481 | STRBP | SOX2 regulated genes in Wounding from PMID: 30772301 | SOX2 |
| 3482 | TENM4 | SOX2 regulated genes in Wounding from PMID: 30772301 | SOX2 |
| 3483 | PARD3 | SOX2 regulated genes in Wounding from PMID: 30772301 | SOX2 |
| 3484 | ST3GAL2 | SOX2 regulated genes in Wounding from PMID: 30772301 | SOX2 |
| 3485 | TRIP6 | SOX2 regulated genes in Wounding from PMID: 30772301 | SOX2 |
| 3486 | KLC2 | SOX2 regulated genes in Wounding from PMID: 30772301 | SOX2 |
| 3487 | SRD5A2 | SOX2 regulated genes in Wounding from PMID: 30772301 | SOX2 |
| 3488 | STX1A | SOX2 regulated genes in Wounding from PMID: 30772301 | SOX2 |
| 3489 | SLC5A5 | SOX2 regulated genes in Wounding from PMID: 30772301 | SOX2 |
| 3490 | SCX | SOX2 regulated genes in Wounding from PMID: 30772301 | SOX2 |
| 3491 | TGM6 | SOX2 regulated genes in Wounding from PMID: 30772301 | SOX2 |
| 3492 | TMC7 | SOX2 regulated genes in Wounding from PMID: 30772301 | SOX2 |
| 3493 | SIPA1L3 | SOX2 regulated genes in Wounding from PMID: 30772301 | SOX2 |
| 3494 | OTUD6B | SOX2 regulated genes in Wounding from PMID: 30772301 | SOX2 |
| 3495 | TMEM163 | SOX2 regulated genes in Wounding from PMID: 30772301 | SOX2 |
| 3496 | GPSM1 | SOX2 regulated genes in Wounding from PMID: 30772301 | SOX2 |
| 3497 | FSCN1 | SOX2 regulated genes in Wounding from PMID: 30772301 | SOX2 |
| 3498 | ATP6V1C2 | SOX2 regulated genes in Wounding from PMID: 30772301 | SOX2 |
| 3499 | NDFIP2 | SOX2 regulated genes in Wounding from PMID: 30772301 | SOX2 |
| 3500 | EEF1AKMT1 | SOX2 regulated genes in Wounding from PMID: 30772301 | SOX2 |
| 3501 | SKIL | SOX2 regulated genes in Wounding from PMID: 30772301 | SOX2 |
| 3502 | USP20 | SOX2 regulated genes in Wounding from PMID: 30772301 | SOX2 |
| 3503 | SMDT1 | SOX2 regulated genes in Wounding from PMID: 30772301 | SOX2 |
| 3504 | IFITM10 | SOX2 regulated genes in Wounding from PMID: 30772301 | SOX2 |
| 3505 | RAB18 | SOX2 regulated genes in Wounding from PMID: 30772301 | SOX2 |
| 3506 | STARD4 | SOX2 regulated genes in Wounding from PMID: 30772301 | SOX2 |
| 3507 | ANKRD6 | SOX2 regulated genes in Wounding from PMID: 30772301 | SOX2 |
| 3508 | EGR4 | SOX2 regulated genes in Wounding from PMID: 30772301 | SOX2 |
| 3509 | FZD6 | SOX2 regulated genes in Wounding from PMID: 30772301 | SOX2 |
| 3510 | CRB3 | SOX2 regulated genes in Wounding from PMID: 30772301 | SOX2 |
| 3511 | CDKN2AIP | SOX2 regulated genes in Wounding from PMID: 30772301 | SOX2 |
| 3512 | PHLDB2 | SOX2 regulated genes in Wounding from PMID: 30772301 | SOX2 |
| 3513 | TMTC3 | SOX2 regulated genes in Wounding from PMID: 30772301 | SOX2 |
| 3514 | EPPK1 | SOX2 regulated genes in Wounding from PMID: 30772301 | SOX2 |
| 3515 | SLC7A6 | SOX2 regulated genes in Wounding from PMID: 30772301 | SOX2 |
| 3516 | LAP3 | SOX2 regulated genes in Wounding from PMID: 30772301 | SOX2 |
| 3517 | ARHGEF16 | SOX2 regulated genes in Wounding from PMID: 30772301 | SOX2 |
| 3518 | SSRP1 | SOX2 regulated genes in Wounding from PMID: 30772301 | SOX2 |
| 3519 | MSX2 | SOX2 regulated genes in Wounding from PMID: 30772301 | SOX2 |
| 3520 | BTBD10 | SOX2 regulated genes in Wounding from PMID: 30772301 | SOX2 |
| 3521 | LRRFIP2 | SOX2 regulated genes in Wounding from PMID: 30772301 | SOX2 |
| 3522 | GGACT | SOX2 regulated genes in Wounding from PMID: 30772301 | SOX2 |
| 3523 | SLC30A4 | SOX2 regulated genes in Wounding from PMID: 30772301 | SOX2 |
| 3524 | SMYD5 | SOX2 regulated genes in Wounding from PMID: 30772301 | SOX2 |
| 3525 | TUBB4B | SOX2 regulated genes in Wounding from PMID: 30772301 | SOX2 |
| 3526 | STIM2 | SOX2 regulated genes in Wounding from PMID: 30772301 | SOX2 |
| 3527 | EIF4A1 | SOX2 regulated genes in Wounding from PMID: 30772301 | SOX2 |
| 3528 | ANPEP | SOX2 regulated genes in Wounding from PMID: 30772301 | SOX2 |
| 3529 | SERINC5 | SOX2 regulated genes in Wounding from PMID: 30772301 | SOX2 |
| 3530 | SCRG1 | SOX2 regulated genes in Wounding from PMID: 30772301 | SOX2 |
| 3531 | SH3BGRL2 | SOX2 regulated genes in Wounding from PMID: 30772301 | SOX2 |
| 3532 | HYAL2 | SOX2 regulated genes in Wounding from PMID: 30772301 | SOX2 |
| 3533 | GDE1 | SOX2 regulated genes in Wounding from PMID: 30772301 | SOX2 |
| 3534 | CHD7 | SOX2 regulated genes in Wounding from PMID: 30772301 | SOX2 |
| 3535 | ARL1 | SOX2 regulated genes in Wounding from PMID: 30772301 | SOX2 |
| 3536 | NIM1K | SOX2 regulated genes in Wounding from PMID: 30772301 | SOX2 |

| | | | |
|---|---|---|---|
| 3537 | EIF3CL | SOX2 regulated genes in Wounding from PMID: 30772301 | SOX2 |
| 3538 | EIF3C | SOX2 regulated genes in Wounding from PMID: 30772301 | SOX2 |
| 3539 | SURF6 | SOX2 regulated genes in Wounding from PMID: 30772301 | SOX2 |
| 3540 | ESRP1 | SOX2 regulated genes in Wounding from PMID: 30772301 | SOX2 |
| 3541 | NAA15 | SOX2 regulated genes in Wounding from PMID: 30772301 | SOX2 |
| 3542 | SMURF1 | SOX2 regulated genes in Wounding from PMID: 30772301 | SOX2 |
| 3543 | FTSJ3 | SOX2 regulated genes in Wounding from PMID: 30772301 | SOX2 |
| 3544 | RABL2B | SOX2 regulated genes in Wounding from PMID: 30772301 | SOX2 |
| 3545 | LPAR2 | SOX2 regulated genes in Wounding from PMID: 30772301 | SOX2 |
| 3546 | ZCCHC7 | SOX2 regulated genes in Wounding from PMID: 30772301 | SOX2 |
| 3547 | KLHL25 | SOX2 regulated genes in Wounding from PMID: 30772301 | SOX2 |
| 3548 | GRIPAP1 | SOX2 regulated genes in Wounding from PMID: 30772301 | SOX2 |
| 3549 | NBN | SOX2 regulated genes in Wounding from PMID: 30772301 | SOX2 |
| 3550 | LGALS3 | SOX2 regulated genes in Wounding from PMID: 30772301 | SOX2 |
| 3551 | TMF1 | SOX2 regulated genes in Wounding from PMID: 30772301 | SOX2 |
| 3552 | DZIP1L | SOX2 regulated genes in Wounding from PMID: 30772301 | SOX2 |
| 3553 | TUBB | SOX2 regulated genes in Wounding from PMID: 30772301 | SOX2 |
| 3554 | ZNF274 | SOX2 regulated genes in Wounding from PMID: 30772301 | SOX2 |
| 3555 | GDNF | SOX2 regulated genes in Wounding from PMID: 30772301 | SOX2 |
| 3556 | MIA-RAB4B | SOX2 regulated genes in Wounding from PMID: 30772301 | SOX2 |
| 3557 | MIA | SOX2 regulated genes in Wounding from PMID: 30772301 | SOX2 |
| 3558 | VPS36 | SOX2 regulated genes in Wounding from PMID: 30772301 | SOX2 |
| 3559 | MATN3 | SOX2 regulated genes in Wounding from PMID: 30772301 | SOX2 |
| 3560 | DCBLD1 | SOX2 regulated genes in Wounding from PMID: 30772301 | SOX2 |
| 3561 | CLDN12 | SOX2 regulated genes in Wounding from PMID: 30772301 | SOX2 |
| 3562 | CST6 | SOX2 regulated genes in Wounding from PMID: 30772301 | SOX2 |
| 3563 | EIF2S3 | SOX2 regulated genes in Wounding from PMID: 30772301 | SOX2 |
| 3564 | MAGEL2 | SOX2 regulated genes in Wounding from PMID: 30772301 | SOX2 |
| 3565 | GALNT14 | SOX2 regulated genes in Wounding from PMID: 30772301 | SOX2 |
| 3566 | YKT6 | SOX2 regulated genes in Wounding from PMID: 30772301 | SOX2 |
| 3567 | WNK2 | SOX2 regulated genes in Wounding from PMID: 30772301 | SOX2 |
| 3568 | TUBB2A | SOX2 regulated genes in Wounding from PMID: 30772301 | SOX2 |
| 3569 | EOGT | SOX2 regulated genes in Wounding from PMID: 30772301 | SOX2 |
| 3570 | ERCC6 | SOX2 regulated genes in Wounding from PMID: 30772301 | SOX2 |
| 3571 | ZDHHC21 | SOX2 regulated genes in Wounding from PMID: 30772301 | SOX2 |
| 3572 | TUBA1B | SOX2 regulated genes in Wounding from PMID: 30772301 | SOX2 |
| 3573 | GDF7 | SOX2 regulated genes in Wounding from PMID: 30772301 | SOX2 |
| 3574 | IL6 | SOX2 regulated genes in Wounding from PMID: 30772301 | SOX2 |
| 3575 | PADI4 | SOX2 regulated genes in Wounding from PMID: 30772301 | SOX2 |
| 3576 | THSD1 | SOX2 regulated genes in Wounding from PMID: 30772301 | SOX2 |
| 3577 | EIF2S1 | SOX2 regulated genes in Wounding from PMID: 30772301 | SOX2 |
| 3578 | RHPN2 | SOX2 regulated genes in Wounding from PMID: 30772301 | SOX2 |
| 3579 | ZC3H15 | SOX2 regulated genes in Wounding from PMID: 30772301 | SOX2 |
| 3580 | FAM156A | SOX2 regulated genes in Wounding from PMID: 30772301 | SOX2 |
| 3581 | UHRF1BP1L | SOX2 regulated genes in Wounding from PMID: 30772301 | SOX2 |
| 3582 | TAB3 | SOX2 regulated genes in Wounding from PMID: 30772301 | SOX2 |
| 3583 | TAF4 | SOX2 regulated genes in Wounding from PMID: 30772301 | SOX2 |
| 3584 | PPP1CA | SOX2 regulated genes in Wounding from PMID: 30772301 | SOX2 |
| 3585 | CYP21A2 | SOX2 regulated genes in Wounding from PMID: 30772301 | SOX2 |
| 3586 | CDSN | SOX2 regulated genes in Wounding from PMID: 30772301 | SOX2 |
| 3587 | TUBB2B | SOX2 regulated genes in Wounding from PMID: 30772301 | SOX2 |
| 3588 | BHLHB9 | SOX2 regulated genes in Wounding from PMID: 30772301 | SOX2 |
| 3589 | RNF19A | SOX2 regulated genes in Wounding from PMID: 30772301 | SOX2 |
| 3590 | SPATS2 | SOX2 regulated genes in Wounding from PMID: 30772301 | SOX2 |
| 3591 | ARHGEF26 | SOX2 regulated genes in Wounding from PMID: 30772301 | SOX2 |
| 3592 | NAB2 | SOX2 regulated genes in Wounding from PMID: 30772301 | SOX2 |
| 3593 | CABLES2 | SOX2 regulated genes in Wounding from PMID: 30772301 | SOX2 |
| 3594 | CREG2 | SOX2 regulated genes in Wounding from PMID: 30772301 | SOX2 |
| 3595 | MTHFD2L | SOX2 regulated genes in Wounding from PMID: 30772301 | SOX2 |
| 3596 | RNF185 | SOX2 regulated genes in Wounding from PMID: 30772301 | SOX2 |
| 3597 | PKP3 | SOX2 regulated genes in Wounding from PMID: 30772301 | SOX2 |

| | | | |
|---|---|---|---|
| 3598 | GNL3L | SOX2 regulated genes in Wounding from PMID: 30772301 | SOX2 |
| 3599 | ATF4 | SOX2 regulated genes in Wounding from PMID: 30772301 | SOX2 |
| 3600 | GJB6 | SOX2 regulated genes in Wounding from PMID: 30772301 | SOX2 |
| 3601 | CRIP2 | SOX2 regulated genes in Wounding from PMID: 30772301 | SOX2 |
| 3602 | MAEA | SOX2 regulated genes in Wounding from PMID: 30772301 | SOX2 |
| 3603 | USP38 | SOX2 regulated genes in Wounding from PMID: 30772301 | SOX2 |
| 3604 | COL4A6 | SOX2 regulated genes in Wounding from PMID: 30772301 | SOX2 |
| 3605 | ILDR1 | SOX2 regulated genes in Wounding from PMID: 30772301 | SOX2 |
| 3606 | MET | SOX2 regulated genes in Wounding from PMID: 30772301 | SOX2 |
| 3607 | MRPL48 | SOX2 regulated genes in Wounding from PMID: 30772301 | SOX2 |
| 3608 | ZNF507 | SOX2 regulated genes in Wounding from PMID: 30772301 | SOX2 |
| 3609 | OXSR1 | SOX2 regulated genes in Wounding from PMID: 30772301 | SOX2 |
| 3610 | PTBP3 | SOX2 regulated genes in Wounding from PMID: 30772301 | SOX2 |
| 3611 | GIPC1 | SOX2 regulated genes in Wounding from PMID: 30772301 | SOX2 |
| 3612 | ADD2 | SOX2 regulated genes in Wounding from PMID: 30772301 | SOX2 |
| 3613 | FBXW7 | SOX2 regulated genes in Wounding from PMID: 30772301 | SOX2 |
| 3614 | TRAPPC13 | SOX2 regulated genes in Wounding from PMID: 30772301 | SOX2 |
| 3615 | MARK3 | SOX2 regulated genes in Wounding from PMID: 30772301 | SOX2 |
| 3616 | HNRNPA2B1 | SOX2 regulated genes in Wounding from PMID: 30772301 | SOX2 |
| 3617 | ZBTB44 | SOX2 regulated genes in Wounding from PMID: 30772301 | SOX2 |
| 3618 | BAMBI | SOX2 regulated genes in Wounding from PMID: 30772301 | SOX2 |
| 3619 | CEACAM21 | SOX2 regulated genes in Wounding from PMID: 30772301 | SOX2 |
| 3620 | WASF1 | SOX2 regulated genes in Wounding from PMID: 30772301 | SOX2 |
| 3621 | CTNS | SOX2 regulated genes in Wounding from PMID: 30772301 | SOX2 |
| 3622 | PELO | SOX2 regulated genes in Wounding from PMID: 30772301 | SOX2 |
| 3623 | UBE4B | SOX2 regulated genes in Wounding from PMID: 30772301 | SOX2 |
| 3624 | PDZD7 | SOX2 regulated genes in Wounding from PMID: 30772301 | SOX2 |
| 3625 | CNKSR1 | SOX2 regulated genes in Wounding from PMID: 30772301 | SOX2 |
| 3626 | GALT | SOX2 regulated genes in Wounding from PMID: 30772301 | SOX2 |
| 3627 | METAP2 | SOX2 regulated genes in Wounding from PMID: 30772301 | SOX2 |
| 3628 | PLSCR3 | SOX2 regulated genes in Wounding from PMID: 30772301 | SOX2 |
| 3629 | ABCE1 | SOX2 regulated genes in Wounding from PMID: 30772301 | SOX2 |
| 3630 | RAB6A | SOX2 regulated genes in Wounding from PMID: 30772301 | SOX2 |
| 3631 | CMTM8 | SOX2 regulated genes in Wounding from PMID: 30772301 | SOX2 |
| 3632 | FCHSD1 | SOX2 regulated genes in Wounding from PMID: 30772301 | SOX2 |
| 3633 | CRIM1 | SOX2 regulated genes in Wounding from PMID: 30772301 | SOX2 |
| 3634 | JUP | SOX2 regulated genes in Wounding from PMID: 30772301 | SOX2 |
| 3635 | GRHL2 | SOX2 regulated genes in Wounding from PMID: 30772301 | SOX2 |
| 3636 | HSPA1A | SOX2 regulated genes in Wounding from PMID: 30772301 | SOX2 |
| 3637 | HSPA1B | SOX2 regulated genes in Wounding from PMID: 30772301 | SOX2 |
| 3638 | NEBL | SOX2 regulated genes in Wounding from PMID: 30772301 | SOX2 |
| 3639 | FBLIM1 | SOX2 regulated genes in Wounding from PMID: 30772301 | SOX2 |
| 3640 | SHISA2 | SOX2 regulated genes in Wounding from PMID: 30772301 | SOX2 |
| 3641 | PPFIA1 | SOX2 regulated genes in Wounding from PMID: 30772301 | SOX2 |
| 3642 | UCP2 | SOX2 regulated genes in Wounding from PMID: 30772301 | SOX2 |
| 3643 | COPS5 | SOX2 regulated genes in Wounding from PMID: 30772301 | SOX2 |
| 3644 | DAPK2 | SOX2 regulated genes in Wounding from PMID: 30772301 | SOX2 |
| 3645 | ESYT3 | SOX2 regulated genes in Wounding from PMID: 30772301 | SOX2 |
| 3646 | CDK12 | SOX2 regulated genes in Wounding from PMID: 30772301 | SOX2 |
| 3647 | BASP1 | SOX2 regulated genes in Wounding from PMID: 30772301 | SOX2 |
| 3648 | MAP3K5 | SOX2 regulated genes in Wounding from PMID: 30772301 | SOX2 |
| 3649 | AP3M1 | SOX2 regulated genes in Wounding from PMID: 30772301 | SOX2 |
| 3650 | CNPY4 | SOX2 regulated genes in Wounding from PMID: 30772301 | SOX2 |
| 3651 | PFKP | SOX2 regulated genes in Wounding from PMID: 30772301 | SOX2 |
| 3652 | PXDC1 | SOX2 regulated genes in Wounding from PMID: 30772301 | SOX2 |
| 3653 | EHD3 | SOX2 regulated genes in Wounding from PMID: 30772301 | SOX2 |
| 3654 | EXOSC3 | SOX2 regulated genes in Wounding from PMID: 30772301 | SOX2 |
| 3655 | BCL2L2 | SOX2 regulated genes in Wounding from PMID: 30772301 | SOX2 |
| 3656 | PPP4C | SOX2 regulated genes in Wounding from PMID: 30772301 | SOX2 |
| 3657 | SRF | SOX2 regulated genes in Wounding from PMID: 30772301 | SOX2 |
| 3658 | BCAR3 | SOX2 regulated genes in Wounding from PMID: 30772301 | SOX2 |

| | | | |
|---|---|---|---|
| 3659 | ATP8A2 | SOX2 regulated genes in Wounding from PMID: 30772301 | SOX2 |
| 3660 | RCN1 | SOX2 regulated genes in Wounding from PMID: 30772301 | SOX2 |
| 3661 | ALCAM | SOX2 regulated genes in Wounding from PMID: 30772301 | SOX2 |
| 3662 | HYAL4 | SOX2 regulated genes in Wounding from PMID: 30772301 | SOX2 |
| 3663 | RFLNA | SOX2 regulated genes in Wounding from PMID: 30772301 | SOX2 |
| 3664 | TNIK | SOX2 regulated genes in Wounding from PMID: 30772301 | SOX2 |
| 3665 | DMTF1 | SOX2 regulated genes in Wounding from PMID: 30772301 | SOX2 |
| 3666 | RIN1 | SOX2 regulated genes in Wounding from PMID: 30772301 | SOX2 |
| 3667 | CALML3 | SOX2 regulated genes in Wounding from PMID: 30772301 | SOX2 |
| 3668 | MREG | SOX2 regulated genes in Wounding from PMID: 30772301 | SOX2 |
| 3669 | YWHAG | SOX2 regulated genes in Wounding from PMID: 30772301 | SOX2 |
| 3670 | GPC2 | SOX2 regulated genes in Wounding from PMID: 30772301 | SOX2 |
| 3671 | UBE2D3 | SOX2 regulated genes in Wounding from PMID: 30772301 | SOX2 |
| 3672 | FZD1 | SOX2 regulated genes in Wounding from PMID: 30772301 | SOX2 |
| 3673 | BARX2 | SOX2 regulated genes in Wounding from PMID: 30772301 | SOX2 |
| 3674 | SLC22A23 | SOX2 regulated genes in Wounding from PMID: 30772301 | SOX2 |
| 3675 | TNKS1BP1 | SOX2 regulated genes in Wounding from PMID: 30772301 | SOX2 |
| 3676 | WNT10B | SOX2 regulated genes in Wounding from PMID: 30772301 | SOX2 |
| 3677 | CCR4 | SOX2 regulated genes in Wounding from PMID: 30772301 | SOX2 |
| 3678 | DAAM1 | SOX2 regulated genes in Wounding from PMID: 30772301 | SOX2 |
| 3679 | HCFC2 | SOX2 regulated genes in Wounding from PMID: 30772301 | SOX2 |
| 3680 | MTERF4 | SOX2 regulated genes in Wounding from PMID: 30772301 | SOX2 |
| 3681 | ATP10B | SOX2 regulated genes in Wounding from PMID: 30772301 | SOX2 |
| 3682 | TMPRSS13 | SOX2 regulated genes in Wounding from PMID: 30772301 | SOX2 |
| 3683 | TACSTD2 | SOX2 regulated genes in Wounding from PMID: 30772301 | SOX2 |
| 3684 | TRPV6 | SOX2 regulated genes in Wounding from PMID: 30772301 | SOX2 |
| 3685 | TMPRSS7 | SOX2 regulated genes in Wounding from PMID: 30772301 | SOX2 |
| 3686 | TTLL10 | SOX2 regulated genes in Wounding from PMID: 30772301 | SOX2 |
| 3687 | TRIB1 | SOX2 regulated genes in Wounding from PMID: 30772301 | SOX2 |
| 3688 | SCNN1G | SOX2 regulated genes in Wounding from PMID: 30772301 | SOX2 |
| 3689 | CHMP3 | SOX2 regulated genes in Wounding from PMID: 30772301 | SOX2 |
| 3690 | RNF103-CHMP3 | SOX2 regulated genes in Wounding from PMID: 30772301 | SOX2 |
| 3691 | BPGM | SOX2 regulated genes in Wounding from PMID: 30772301 | SOX2 |
| 3692 | PRRC2B | SOX2 regulated genes in Wounding from PMID: 30772301 | SOX2 |
| 3693 | ALDH3A2 | SOX2 regulated genes in Wounding from PMID: 30772301 | SOX2 |
| 3694 | SLC25A13 | SOX2 regulated genes in Wounding from PMID: 30772301 | SOX2 |
| 3695 | BCL7A | SOX2 regulated genes in Wounding from PMID: 30772301 | SOX2 |
| 3696 | SERTAD1 | SOX2 regulated genes in Wounding from PMID: 30772301 | SOX2 |
| 3697 | GPNMB | SOX2 regulated genes in Wounding from PMID: 30772301 | SOX2 |
| 3698 | SAE1 | SOX2 regulated genes in Wounding from PMID: 30772301 | SOX2 |
| 3699 | DNAJC6 | SOX2 regulated genes in Wounding from PMID: 30772301 | SOX2 |
| 3700 | STOX1 | SOX2 regulated genes in Wounding from PMID: 30772301 | SOX2 |
| 3701 | DYNC1LI2 | SOX2 regulated genes in Wounding from PMID: 30772301 | SOX2 |
| 3702 | ZMYM2 | SOX2 regulated genes in Wounding from PMID: 30772301 | SOX2 |
| 3703 | RABGEF1 | SOX2 regulated genes in Wounding from PMID: 30772301 | SOX2 |
| 3704 | STBD1 | SOX2 regulated genes in Wounding from PMID: 30772301 | SOX2 |
| 3705 | INTS6 | SOX2 regulated genes in Wounding from PMID: 30772301 | SOX2 |
| 3706 | AP1AR | SOX2 regulated genes in Wounding from PMID: 30772301 | SOX2 |
| 3707 | DDX51 | SOX2 regulated genes in Wounding from PMID: 30772301 | SOX2 |
| 3708 | PI4K2B | SOX2 regulated genes in Wounding from PMID: 30772301 | SOX2 |
| 3709 | PUF60 | SOX2 regulated genes in Wounding from PMID: 30772301 | SOX2 |
| 3710 | SCT | SOX2 regulated genes in Wounding from PMID: 30772301 | SOX2 |
| 3711 | ANAPC2 | SOX2 regulated genes in Wounding from PMID: 30772301 | SOX2 |
| 3712 | CYP4F12 | SOX2 regulated genes in Wounding from PMID: 30772301 | SOX2 |
| 3713 | CDCP1 | SOX2 regulated genes in Wounding from PMID: 30772301 | SOX2 |
| 3714 | TJP2 | SOX2 regulated genes in Wounding from PMID: 30772301 | SOX2 |
| 3715 | VCP | SOX2 regulated genes in Wounding from PMID: 30772301 | SOX2 |
| 3716 | PDZRN3 | SOX2 regulated genes in Wounding from PMID: 30772301 | SOX2 |
| 3717 | SRC | SOX2 regulated genes in Wounding from PMID: 30772301 | SOX2 |
| 3718 | CA6 | SOX2 regulated genes in Wounding from PMID: 30772301 | SOX2 |
| 3719 | LMNB1 | SOX2 regulated genes in Wounding from PMID: 30772301 | SOX2 |

| | | | |
|---|---|---|---|
| 3720 | SH3YL1 | SOX2 regulated genes in Wounding from PMID: 30772301 | SOX2 |
| 3721 | JMY | SOX2 regulated genes in Wounding from PMID: 30772301 | SOX2 |
| 3722 | NUMB | SOX2 regulated genes in Wounding from PMID: 30772301 | SOX2 |
| 3723 | TOB1 | SOX2 regulated genes in Wounding from PMID: 30772301 | SOX2 |
| 3724 | GIPC2 | SOX2 regulated genes in Wounding from PMID: 30772301 | SOX2 |
| 3725 | FOXK2 | SOX2 regulated genes in Wounding from PMID: 30772301 | SOX2 |
| 3726 | IMP4 | SOX2 regulated genes in Wounding from PMID: 30772301 | SOX2 |
| 3727 | KATNBL1 | SOX2 regulated genes in Wounding from PMID: 30772301 | SOX2 |
| 3728 | HSPA14 | SOX2 regulated genes in Wounding from PMID: 30772301 | SOX2 |
| 3729 | NFE2L3 | SOX2 regulated genes in Wounding from PMID: 30772301 | SOX2 |
| 3730 | YWHAZ | SOX2 regulated genes in Wounding from PMID: 30772301 | SOX2 |
| 3731 | PTPRF | SOX2 regulated genes in Wounding from PMID: 30772301 | SOX2 |
| 3732 | SLC38A5 | SOX2 regulated genes in Wounding from PMID: 30772301 | SOX2 |
| 3733 | ACP7 | SOX2 regulated genes in Wounding from PMID: 30772301 | SOX2 |
| 3734 | KCTD4 | SOX2 regulated genes in Wounding from PMID: 30772301 | SOX2 |
| 3735 | HEPH | SOX2 regulated genes in Wounding from PMID: 30772301 | SOX2 |
| 3736 | ETF1 | SOX2 regulated genes in Wounding from PMID: 30772301 | SOX2 |
| 3737 | FLRT2 | SOX2 regulated genes in Wounding from PMID: 30772301 | SOX2 |
| 3738 | GCNT4 | SOX2 regulated genes in Wounding from PMID: 30772301 | SOX2 |
| 3739 | NRBP2 | SOX2 regulated genes in Wounding from PMID: 30772301 | SOX2 |
| 3740 | PNLDC1 | SOX2 regulated genes in Wounding from PMID: 30772301 | SOX2 |
| 3741 | WHRN | SOX2 regulated genes in Wounding from PMID: 30772301 | SOX2 |
| 3742 | TMEM64 | SOX2 regulated genes in Wounding from PMID: 30772301 | SOX2 |
| 3743 | TRO | SOX2 regulated genes in Wounding from PMID: 30772301 | SOX2 |
| 3744 | PPARD | SOX2 regulated genes in Wounding from PMID: 30772301 | SOX2 |
| 3745 | SH3D19 | SOX2 regulated genes in Wounding from PMID: 30772301 | SOX2 |
| 3746 | SRSF12 | SOX2 regulated genes in Wounding from PMID: 30772301 | SOX2 |
| 3747 | AGAP3 | SOX2 regulated genes in Wounding from PMID: 30772301 | SOX2 |
| 3748 | C5orf46 | SOX2 regulated genes in Wounding from PMID: 30772301 | SOX2 |
| 3749 | SYT17 | SOX2 regulated genes in Wounding from PMID: 30772301 | SOX2 |
| 3750 | PLAUR | SOX2 regulated genes in Wounding from PMID: 30772301 | SOX2 |
| 3751 | SNN | SOX2 regulated genes in Wounding from PMID: 30772301 | SOX2 |
| 3752 | UBAP2 | SOX2 regulated genes in Wounding from PMID: 30772301 | SOX2 |
| 3753 | MFSD13A | SOX2 regulated genes in Wounding from PMID: 30772301 | SOX2 |
| 3754 | EPB41L4A | SOX2 regulated genes in Wounding from PMID: 30772301 | SOX2 |
| 3755 | PREP | SOX2 regulated genes in Wounding from PMID: 30772301 | SOX2 |
| 3756 | TSSK1B | SOX2 regulated genes in Wounding from PMID: 30772301 | SOX2 |
| 3757 | ATP6V0A4 | SOX2 regulated genes in Wounding from PMID: 30772301 | SOX2 |
| 3758 | LXN | SOX2 regulated genes in Wounding from PMID: 30772301 | SOX2 |
| 3759 | DUSP6 | SOX2 regulated genes in Wounding from PMID: 30772301 | SOX2 |
| 3760 | SATB2 | SOX2 regulated genes in Wounding from PMID: 30772301 | SOX2 |
| 3761 | GORASP2 | SOX2 regulated genes in Wounding from PMID: 30772301 | SOX2 |
| 3762 | KDM5C | SOX2 regulated genes in Wounding from PMID: 30772301 | SOX2 |
| 3763 | DGKZ | SOX2 regulated genes in Wounding from PMID: 30772301 | SOX2 |
| 3764 | OTUB2 | SOX2 regulated genes in Wounding from PMID: 30772301 | SOX2 |
| 3765 | EEA1 | SOX2 regulated genes in Wounding from PMID: 30772301 | SOX2 |
| 3766 | VSIG10 | SOX2 regulated genes in Wounding from PMID: 30772301 | SOX2 |
| 3767 | ALDH1A3 | SOX2 regulated genes in Wounding from PMID: 30772301 | SOX2 |
| 3768 | USP32 | SOX2 regulated genes in Wounding from PMID: 30772301 | SOX2 |
| 3769 | ANKLE1 | SOX2 regulated genes in Wounding from PMID: 30772301 | SOX2 |
| 3770 | AKIRIN2 | SOX2 regulated genes in Wounding from PMID: 30772301 | SOX2 |
| 3771 | SAMD5 | SOX2 regulated genes in Wounding from PMID: 30772301 | SOX2 |
| 3772 | DYNLT1 | SOX2 regulated genes in Wounding from PMID: 30772301 | SOX2 |
| 3773 | PTK2 | SOX2 regulated genes in Wounding from PMID: 30772301 | SOX2 |
| 3774 | CROT | SOX2 regulated genes in Wounding from PMID: 30772301 | SOX2 |
| 3775 | XBP1 | SOX2 regulated genes in Wounding from PMID: 30772301 | SOX2 |
| 3776 | PTK2B | SOX2 regulated genes in Wounding from PMID: 30772301 | SOX2 |
| 3777 | TCF7L1 | SOX2 regulated genes in Wounding from PMID: 30772301 | SOX2 |
| 3778 | CBX1 | SOX2 regulated genes in Wounding from PMID: 30772301 | SOX2 |
| 3779 | GABRP | SOX2 regulated genes in Wounding from PMID: 30772301 | SOX2 |
| 3780 | FZD2 | SOX2 regulated genes in Wounding from PMID: 30772301 | SOX2 |

| | | | |
|---|---|---|---|
| 3781 | BAIAP2L1 | SOX2 regulated genes in Wounding from PMID: 30772301 | SOX2 |
| 3782 | PINLYP | SOX2 regulated genes in Wounding from PMID: 30772301 | SOX2 |
| 3783 | STIP1 | SOX2 regulated genes in Wounding from PMID: 30772301 | SOX2 |
| 3784 | CLDND1 | SOX2 regulated genes in Wounding from PMID: 30772301 | SOX2 |
| 3785 | C1QTNF12 | SOX2 regulated genes in Wounding from PMID: 30772301 | SOX2 |
| 3786 | DYNLL1 | SOX2 regulated genes in Wounding from PMID: 30772301 | SOX2 |
| 3787 | SLCO4C1 | SOX2 regulated genes in Wounding from PMID: 30772301 | SOX2 |
| 3788 | GJB2 | SOX2 regulated genes in Wounding from PMID: 30772301 | SOX2 |
| 3789 | CCL27 | SOX2 regulated genes in Wounding from PMID: 30772301 | SOX2 |
| 3790 | GABARAPL2 | SOX2 regulated genes in Wounding from PMID: 30772301 | SOX2 |
| 3791 | CDC42BPG | SOX2 regulated genes in Wounding from PMID: 30772301 | SOX2 |
| 3792 | GLB1L2 | SOX2 regulated genes in Wounding from PMID: 30772301 | SOX2 |
| 3793 | SLC35A2 | SOX2 regulated genes in Wounding from PMID: 30772301 | SOX2 |
| 3794 | JPH4 | SOX2 regulated genes in Wounding from PMID: 30772301 | SOX2 |
| 3795 | STRN | SOX2 regulated genes in Wounding from PMID: 30772301 | SOX2 |
| 3796 | RET | SOX2 regulated genes in Wounding from PMID: 30772301 | SOX2 |
| 3797 | GPR37 | SOX2 regulated genes in Wounding from PMID: 30772301 | SOX2 |
| 3798 | BOK | SOX2 regulated genes in Wounding from PMID: 30772301 | SOX2 |
| 3799 | CTSF | SOX2 regulated genes in Wounding from PMID: 30772301 | SOX2 |
| 3800 | RTKN2 | SOX2 regulated genes in Wounding from PMID: 30772301 | SOX2 |
| 3801 | PEBP1 | SOX2 regulated genes in Wounding from PMID: 30772301 | SOX2 |
| 3802 | C3orf18 | SOX2 regulated genes in Wounding from PMID: 30772301 | SOX2 |
| 3803 | GAN | SOX2 regulated genes in Wounding from PMID: 30772301 | SOX2 |
| 3804 | EIF4A3 | SOX2 regulated genes in Wounding from PMID: 30772301 | SOX2 |
| 3805 | SOWAHB | SOX2 regulated genes in Wounding from PMID: 30772301 | SOX2 |
| 3806 | SUN2 | SOX2 regulated genes in Wounding from PMID: 30772301 | SOX2 |
| 3807 | GPC1 | SOX2 regulated genes in Wounding from PMID: 30772301 | SOX2 |
| 3808 | TDRD9 | SOX2 regulated genes in Wounding from PMID: 30772301 | SOX2 |
| 3809 | BCO1 | SOX2 regulated genes in Wounding from PMID: 30772301 | SOX2 |
| 3810 | EML4 | SOX2 regulated genes in Wounding from PMID: 30772301 | SOX2 |
| 3811 | THAP1 | SOX2 regulated genes in Wounding from PMID: 30772301 | SOX2 |
| 3812 | RGS9 | SOX2 regulated genes in Wounding from PMID: 30772301 | SOX2 |
| 3813 | TTLL7 | SOX2 regulated genes in Wounding from PMID: 30772301 | SOX2 |
| 3814 | IGF1R | SOX2 regulated genes in Wounding from PMID: 30772301 | SOX2 |
| 3815 | EPGN | SOX2 regulated genes in Wounding from PMID: 30772301 | SOX2 |
| 3816 | VASN | SOX2 regulated genes in Wounding from PMID: 30772301 | SOX2 |
| 3817 | CRISP3 | SOX2 regulated genes in Wounding from PMID: 30772301 | SOX2 |
| 3818 | EIF1AX | SOX2 regulated genes in Wounding from PMID: 30772301 | SOX2 |
| 3819 | HSPA2 | SOX2 regulated genes in Wounding from PMID: 30772301 | SOX2 |
| 3820 | BTG1 | SOX2 regulated genes in Wounding from PMID: 30772301 | SOX2 |
| 3821 | KIAA1191 | SOX2 regulated genes in Wounding from PMID: 30772301 | SOX2 |
| 3822 | TPM1 | SOX2 regulated genes in Wounding from PMID: 30772301 | SOX2 |
| 3823 | UNC45A | SOX2 regulated genes in Wounding from PMID: 30772301 | SOX2 |
| 3824 | EIF4G2 | SOX2 regulated genes in Wounding from PMID: 30772301 | SOX2 |
| 3825 | PPP1R36 | SOX2 regulated genes in Wounding from PMID: 30772301 | SOX2 |
| 3826 | DUSP2 | SOX2 regulated genes in Wounding from PMID: 30772301 | SOX2 |
| 3827 | C11orf52 | SOX2 regulated genes in Wounding from PMID: 30772301 | SOX2 |
| 3828 | TMEM62 | SOX2 regulated genes in Wounding from PMID: 30772301 | SOX2 |
| 3829 | CXCL14 | SOX2 regulated genes in Wounding from PMID: 30772301 | SOX2 |
| 3830 | CAMTA1 | SOX2 regulated genes in Wounding from PMID: 30772301 | SOX2 |
| 3831 | SERTAD2 | SOX2 regulated genes in Wounding from PMID: 30772301 | SOX2 |
| 3832 | PLS3 | SOX2 regulated genes in Wounding from PMID: 30772301 | SOX2 |
| 3833 | SIAH2 | SOX2 regulated genes in Wounding from PMID: 30772301 | SOX2 |
| 3834 | NOL10 | SOX2 regulated genes in Wounding from PMID: 30772301 | SOX2 |
| 3835 | ZNF827 | SOX2 regulated genes in Wounding from PMID: 30772301 | SOX2 |
| 3836 | ITGB6 | SOX2 regulated genes in Wounding from PMID: 30772301 | SOX2 |
| 3837 | EIF1AD | SOX2 regulated genes in Wounding from PMID: 30772301 | SOX2 |
| 3838 | PCYOX1 | SOX2 regulated genes in Wounding from PMID: 30772301 | SOX2 |
| 3839 | IGF2BP1 | SOX2 regulated genes in Wounding from PMID: 30772301 | SOX2 |
| 3840 | FADS3 | SOX2 regulated genes in Wounding from PMID: 30772301 | SOX2 |
| 3841 | SMARCC1 | SOX2 regulated genes in Wounding from PMID: 30772301 | SOX2 |

| | | | |
|---|---|---|---|
| 3842 | MT4 | SOX2 regulated genes in Wounding from PMID: 30772301 | SOX2 |
| 3843 | ATXN2 | SOX2 regulated genes in Wounding from PMID: 30772301 | SOX2 |
| 3844 | FAM47E | SOX2 regulated genes in Wounding from PMID: 30772301 | SOX2 |
| 3845 | STK39 | SOX2 regulated genes in Wounding from PMID: 30772301 | SOX2 |
| 3846 | ANO9 | SOX2 regulated genes in Wounding from PMID: 30772301 | SOX2 |
| 3847 | POLH | SOX2 regulated genes in Wounding from PMID: 30772301 | SOX2 |
| 3848 | TNFAIP1 | SOX2 regulated genes in Wounding from PMID: 30772301 | SOX2 |
| 3849 | FAM107B | SOX2 regulated genes in Wounding from PMID: 30772301 | SOX2 |
| 3850 | DENR | SOX2 regulated genes in Wounding from PMID: 30772301 | SOX2 |
| 3851 | PPP2R5B | SOX2 regulated genes in Wounding from PMID: 30772301 | SOX2 |
| 3852 | CCNO | SOX2 regulated genes in Wounding from PMID: 30772301 | SOX2 |
| 3853 | HSD17B14 | SOX2 regulated genes in Wounding from PMID: 30772301 | SOX2 |
| 3854 | LY6D | SOX2 regulated genes in Wounding from PMID: 30772301 | SOX2 |
| 3855 | CSRNP1 | SOX2 regulated genes in Wounding from PMID: 30772301 | SOX2 |
| 3856 | DLX2 | SOX2 regulated genes in Wounding from PMID: 30772301 | SOX2 |
| 3857 | CA2 | SOX2 regulated genes in Wounding from PMID: 30772301 | SOX2 |
| 3858 | SLC35F6 | SOX2 regulated genes in Wounding from PMID: 30772301 | SOX2 |
| 3859 | CARM1 | SOX2 regulated genes in Wounding from PMID: 30772301 | SOX2 |
| 3860 | PSMD12 | SOX2 regulated genes in Wounding from PMID: 30772301 | SOX2 |
| 3861 | EDF1 | SOX2 regulated genes in Wounding from PMID: 30772301 | SOX2 |
| 3862 | PLCXD2 | SOX2 regulated genes in Wounding from PMID: 30772301 | SOX2 |
| 3863 | DDX19A | SOX2 regulated genes in Wounding from PMID: 30772301 | SOX2 |
| 3864 | PURB | SOX2 regulated genes in Wounding from PMID: 30772301 | SOX2 |
| 3865 | SEC23IP | SOX2 regulated genes in Wounding from PMID: 30772301 | SOX2 |
| 3866 | HIF1A | SOX2 regulated genes in Wounding from PMID: 30772301 | SOX2 |
| 3867 | UBA2 | SOX2 regulated genes in Wounding from PMID: 30772301 | SOX2 |
| 3868 | ALG9 | SOX2 regulated genes in Wounding from PMID: 30772301 | SOX2 |
| 3869 | FBP1 | SOX2 regulated genes in Wounding from PMID: 30772301 | SOX2 |
| 3870 | N4BP3 | SOX2 regulated genes in Wounding from PMID: 30772301 | SOX2 |
| 3871 | CSNK1G3 | SOX2 regulated genes in Wounding from PMID: 30772301 | SOX2 |
| 3872 | PLEKHG3 | SOX2 regulated genes in Wounding from PMID: 30772301 | SOX2 |
| 3873 | PPP2R1A | SOX2 regulated genes in Wounding from PMID: 30772301 | SOX2 |
| 3874 | BOP1 | SOX2 regulated genes in Wounding from PMID: 30772301 | SOX2 |
| 3875 | ZNF185 | SOX2 regulated genes in Wounding from PMID: 30772301 | SOX2 |
| 3876 | ZBTB1 | SOX2 regulated genes in Wounding from PMID: 30772301 | SOX2 |
| 3877 | OPLAH | SOX2 regulated genes in Wounding from PMID: 30772301 | SOX2 |
| 3878 | KRT80 | SOX2 regulated genes in Wounding from PMID: 30772301 | SOX2 |
| 3879 | TRPM4 | SOX2 regulated genes in Wounding from PMID: 30772301 | SOX2 |
| 3880 | MAP3K20 | SOX2 regulated genes in Wounding from PMID: 30772301 | SOX2 |
| 3881 | OSGIN1 | SOX2 regulated genes in Wounding from PMID: 30772301 | SOX2 |
| 3882 | GPC3 | SOX2 regulated genes in Wounding from PMID: 30772301 | SOX2 |
| 3883 | POLR2J3 | SOX2 regulated genes in Wounding from PMID: 30772301 | SOX2 |
| 3884 | UPK3BL1 | SOX2 regulated genes in Wounding from PMID: 30772301 | SOX2 |
| 3885 | UPK3BL2 | SOX2 regulated genes in Wounding from PMID: 30772301 | SOX2 |
| 3886 | PLCH2 | SOX2 regulated genes in Wounding from PMID: 30772301 | SOX2 |
| 3887 | CNIH2 | SOX2 regulated genes in Wounding from PMID: 30772301 | SOX2 |
| 3888 | CRYM | SOX2 regulated genes in Wounding from PMID: 30772301 | SOX2 |
| 3889 | EPS8L1 | SOX2 regulated genes in Wounding from PMID: 30772301 | SOX2 |
| 3890 | LCN12 | SOX2 regulated genes in Wounding from PMID: 30772301 | SOX2 |
| 3891 | GTF2H2 | SOX2 regulated genes in Wounding from PMID: 30772301 | SOX2 |
| 3892 | SLC20A2 | SOX2 regulated genes in Wounding from PMID: 30772301 | SOX2 |
| 3893 | SCAPER | SOX2 regulated genes in Wounding from PMID: 30772301 | SOX2 |
| 3894 | SLC38A3 | SOX2 regulated genes in Wounding from PMID: 30772301 | SOX2 |
| 3895 | SH2D4A | SOX2 regulated genes in Wounding from PMID: 30772301 | SOX2 |
| 3896 | NAE1 | SOX2 regulated genes in Wounding from PMID: 30772301 | SOX2 |
| 3897 | MYCL | SOX2 regulated genes in Wounding from PMID: 30772301 | SOX2 |
| 3898 | YWHAQ | SOX2 regulated genes in Wounding from PMID: 30772301 | SOX2 |
| 3899 | NEK5 | SOX2 regulated genes in Wounding from PMID: 30772301 | SOX2 |
| 3900 | PTPRN2 | SOX2 regulated genes in Wounding from PMID: 30772301 | SOX2 |
| 3901 | TAGLN3 | SOX2 regulated genes in Wounding from PMID: 30772301 | SOX2 |
| 3902 | METAP1 | SOX2 regulated genes in Wounding from PMID: 30772301 | SOX2 |

| | | | |
|---|---|---|---|
| 3903 | PLEKHG1 | SOX2 regulated genes in Wounding from PMID: 30772301 | SOX2 |
| 3904 | POLD4 | SOX2 regulated genes in Wounding from PMID: 30772301 | SOX2 |
| 3905 | CLDN3 | SOX2 regulated genes in Wounding from PMID: 30772301 | SOX2 |
| 3906 | SRRT | SOX2 regulated genes in Wounding from PMID: 30772301 | SOX2 |
| 3907 | UNC93A | SOX2 regulated genes in Wounding from PMID: 30772301 | SOX2 |
| 3908 | EIF4E | SOX2 regulated genes in Wounding from PMID: 30772301 | SOX2 |
| 3909 | PXYLP1 | SOX2 regulated genes in Wounding from PMID: 30772301 | SOX2 |
| 3910 | EPCAM | SOX2 regulated genes in Wounding from PMID: 30772301 | SOX2 |
| 3911 | WRNIP1 | SOX2 regulated genes in Wounding from PMID: 30772301 | SOX2 |
| 3912 | LATS1 | SOX2 regulated genes in Wounding from PMID: 30772301 | SOX2 |
| 3913 | PAPSS2 | SOX2 regulated genes in Wounding from PMID: 30772301 | SOX2 |
| 3914 | ANKH | SOX2 regulated genes in Wounding from PMID: 30772301 | SOX2 |
| 3915 | PPP1CB | SOX2 regulated genes in Wounding from PMID: 30772301 | SOX2 |
| 3916 | C4orf36 | SOX2 regulated genes in Wounding from PMID: 30772301 | SOX2 |
| 3917 | CLASRP | SOX2 regulated genes in Wounding from PMID: 30772301 | SOX2 |
| 3918 | DHX16 | SOX2 regulated genes in Wounding from PMID: 30772301 | SOX2 |
| 3919 | PSCA | SOX2 regulated genes in Wounding from PMID: 30772301 | SOX2 |
| 3920 | NECAB1 | SOX2 regulated genes in Wounding from PMID: 30772301 | SOX2 |
| 3921 | CKAP4 | SOX2 regulated genes in Wounding from PMID: 30772301 | SOX2 |
| 3922 | BNC2 | SOX2 regulated genes in Wounding from PMID: 30772301 | SOX2 |
| 3923 | PDZD2 | SOX2 regulated genes in Wounding from PMID: 30772301 | SOX2 |
| 3924 | ENOX1 | SOX2 regulated genes in Wounding from PMID: 30772301 | SOX2 |
| 3925 | PRKCI | SOX2 regulated genes in Wounding from PMID: 30772301 | SOX2 |
| 3926 | XRCC1 | SOX2 regulated genes in Wounding from PMID: 30772301 | SOX2 |
| 3927 | MPRIP | SOX2 regulated genes in Wounding from PMID: 30772301 | SOX2 |
| 3928 | BMPR1B | SOX2 regulated genes in Wounding from PMID: 30772301 | SOX2 |
| 3929 | CCK | SOX2 regulated genes in Wounding from PMID: 30772301 | SOX2 |
| 3930 | LHX2 | SOX2 regulated genes in Wounding from PMID: 30772301 | SOX2 |
| 3931 | SP6 | SOX2 regulated genes in Wounding from PMID: 30772301 | SOX2 |
| 3932 | MFSD6 | SOX2 regulated genes in Wounding from PMID: 30772301 | SOX2 |
| 3933 | TOM1L1 | SOX2 regulated genes in Wounding from PMID: 30772301 | SOX2 |
| 3934 | FAM83H | SOX2 regulated genes in Wounding from PMID: 30772301 | SOX2 |
| 3935 | AFF4 | SOX2 regulated genes in Wounding from PMID: 30772301 | SOX2 |
| 3936 | PIP4K2C | SOX2 regulated genes in Wounding from PMID: 30772301 | SOX2 |
| 3937 | RNF144B | SOX2 regulated genes in Wounding from PMID: 30772301 | SOX2 |
| 3938 | RBM15B | SOX2 regulated genes in Wounding from PMID: 30772301 | SOX2 |
| 3939 | LGALS12 | SOX2 regulated genes in Wounding from PMID: 30772301 | SOX2 |
| 3940 | ATP6V1G1 | SOX2 regulated genes in Wounding from PMID: 30772301 | SOX2 |
| 3941 | SCARB2 | SOX2 regulated genes in Wounding from PMID: 30772301 | SOX2 |
| 3942 | INTS8 | SOX2 regulated genes in Wounding from PMID: 30772301 | SOX2 |
| 3943 | MTFR1 | SOX2 regulated genes in Wounding from PMID: 30772301 | SOX2 |
| 3944 | AHI1 | SOX2 regulated genes in Wounding from PMID: 30772301 | SOX2 |
| 3945 | PGAP2 | SOX2 regulated genes in Wounding from PMID: 30772301 | SOX2 |
| 3946 | PHRF1 | SOX2 regulated genes in Wounding from PMID: 30772301 | SOX2 |
| 3947 | MYC | SOX2 regulated genes in Wounding from PMID: 30772301 | SOX2 |
| 3948 | PPRC1 | SOX2 regulated genes in Wounding from PMID: 30772301 | SOX2 |
| 3949 | SIGMAR1 | SOX2 regulated genes in Wounding from PMID: 30772301 | SOX2 |
| 3950 | CLIC4 | SOX2 regulated genes in Wounding from PMID: 30772301 | SOX2 |
| 3951 | ARNTL | SOX2 regulated genes in Wounding from PMID: 30772301 | SOX2 |
| 3952 | CADM4 | SOX2 regulated genes in Wounding from PMID: 30772301 | SOX2 |
| 3953 | GARNL3 | SOX2 regulated genes in Wounding from PMID: 30772301 | SOX2 |
| 3954 | SPINK5 | SOX2 regulated genes in Wounding from PMID: 30772301 | SOX2 |
| 3955 | PPP1R10 | SOX2 regulated genes in Wounding from PMID: 30772301 | SOX2 |
| 3956 | FFAR3 | SOX2 regulated genes in Wounding from PMID: 30772301 | SOX2 |
| 3957 | SEC16A | SOX2 regulated genes in Wounding from PMID: 30772301 | SOX2 |
| 3958 | BCAR1 | SOX2 regulated genes in Wounding from PMID: 30772301 | SOX2 |
| 3959 | SC5D | SOX2 regulated genes in Wounding from PMID: 30772301 | SOX2 |
| 3960 | HOMER2 | SOX2 regulated genes in Wounding from PMID: 30772301 | SOX2 |
| 3961 | MTHFSD | SOX2 regulated genes in Wounding from PMID: 30772301 | SOX2 |
| 3962 | RIPK2 | SOX2 regulated genes in Wounding from PMID: 30772301 | SOX2 |
| 3963 | RPL27 | SOX2 regulated genes in Wounding from PMID: 30772301 | SOX2 |

| | | | |
|---|---|---|---|
| 3964 | MACC1 | SOX2 regulated genes in Wounding from PMID: 30772301 | SOX2 |
| 3965 | PAK4 | SOX2 regulated genes in Wounding from PMID: 30772301 | SOX2 |
| 3966 | FMNL2 | SOX2 regulated genes in Wounding from PMID: 30772301 | SOX2 |
| 3967 | LTV1 | SOX2 regulated genes in Wounding from PMID: 30772301 | SOX2 |
| 3968 | FBXO42 | SOX2 regulated genes in Wounding from PMID: 30772301 | SOX2 |
| 3969 | PRKCA | SOX2 regulated genes in Wounding from PMID: 30772301 | SOX2 |
| 3970 | CMAS | SOX2 regulated genes in Wounding from PMID: 30772301 | SOX2 |
| 3971 | CTNNA1 | SOX2 regulated genes in Wounding from PMID: 30772301 | SOX2 |
| 3972 | KLHL32 | SOX2 regulated genes in Wounding from PMID: 30772301 | SOX2 |
| 3973 | SNTB1 | SOX2 regulated genes in Wounding from PMID: 30772301 | SOX2 |
| 3974 | ANKRD27 | SOX2 regulated genes in Wounding from PMID: 30772301 | SOX2 |
| 3975 | PHACTR4 | SOX2 regulated genes in Wounding from PMID: 30772301 | SOX2 |
| 3976 | SORCS2 | SOX2 regulated genes in Wounding from PMID: 30772301 | SOX2 |
| 3977 | SLC9A2 | SOX2 regulated genes in Wounding from PMID: 30772301 | SOX2 |
| 3978 | AIG1 | SOX2 regulated genes in Wounding from PMID: 30772301 | SOX2 |
| 3979 | CALD1 | SOX2 regulated genes in Wounding from PMID: 30772301 | SOX2 |
| 3980 | LRP8 | SOX2 regulated genes in Wounding from PMID: 30772301 | SOX2 |
| 3981 | PEX26 | SOX2 regulated genes in Wounding from PMID: 30772301 | SOX2 |
| 3982 | MICAL1 | SOX2 regulated genes in Wounding from PMID: 30772301 | SOX2 |
| 3983 | HSPB1 | SOX2 regulated genes in Wounding from PMID: 30772301 | SOX2 |
| 3984 | AQP9 | SOX2 regulated genes in Wounding from PMID: 30772301 | SOX2 |
| 3985 | ZNF428 | SOX2 regulated genes in Wounding from PMID: 30772301 | SOX2 |
| 3986 | DUSP8 | SOX2 regulated genes in Wounding from PMID: 30772301 | SOX2 |
| 3987 | UBLCP1 | SOX2 regulated genes in Wounding from PMID: 30772301 | SOX2 |
| 3988 | PERP | SOX2 regulated genes in Wounding from PMID: 30772301 | SOX2 |
| 3989 | CCT2 | SOX2 regulated genes in Wounding from PMID: 30772301 | SOX2 |
| 3990 | ATAD2B | SOX2 regulated genes in Wounding from PMID: 30772301 | SOX2 |
| 3991 | NCK2 | SOX2 regulated genes in Wounding from PMID: 30772301 | SOX2 |
| 3992 | PLB1 | SOX2 regulated genes in Wounding from PMID: 30772301 | SOX2 |
| 3993 | DACT2 | SOX2 regulated genes in Wounding from PMID: 30772301 | SOX2 |
| 3994 | PPP2CA | SOX2 regulated genes in Wounding from PMID: 30772301 | SOX2 |
| 3995 | KRT73 | SOX2 regulated genes in Wounding from PMID: 30772301 | SOX2 |
| 3996 | RAB5A | SOX2 regulated genes in Wounding from PMID: 30772301 | SOX2 |
| 3997 | CD82 | SOX2 regulated genes in Wounding from PMID: 30772301 | SOX2 |
| 3998 | ITPKC | SOX2 regulated genes in Wounding from PMID: 30772301 | SOX2 |
| 3999 | NUP62 | SOX2 regulated genes in Wounding from PMID: 30772301 | SOX2 |
| 4000 | GJA3 | SOX2 regulated genes in Wounding from PMID: 30772301 | SOX2 |
| 4001 | FBXO30 | SOX2 regulated genes in Wounding from PMID: 30772301 | SOX2 |
| 4002 | RNF6 | SOX2 regulated genes in Wounding from PMID: 30772301 | SOX2 |
| 4003 | ARHGEF19 | SOX2 regulated genes in Wounding from PMID: 30772301 | SOX2 |
| 4004 | TSPAN8 | SOX2 regulated genes in Wounding from PMID: 30772301 | SOX2 |
| 4005 | DLX3 | SOX2 regulated genes in Wounding from PMID: 30772301 | SOX2 |
| 4006 | KRT75 | SOX2 regulated genes in Wounding from PMID: 30772301 | SOX2 |
| 4007 | MAP3K9 | SOX2 regulated genes in Wounding from PMID: 30772301 | SOX2 |
| 4008 | NECTIN1 | SOX2 regulated genes in Wounding from PMID: 30772301 | SOX2 |
| 4009 | SUDS3 | SOX2 regulated genes in Wounding from PMID: 30772301 | SOX2 |
| 4010 | MSI1 | SOX2 regulated genes in Wounding from PMID: 30772301 | SOX2 |
| 4011 | SARNP | SOX2 regulated genes in Wounding from PMID: 30772301 | SOX2 |
| 4012 | ASH2L | SOX2 regulated genes in Wounding from PMID: 30772301 | SOX2 |
| 4013 | EPB41L4B | SOX2 regulated genes in Wounding from PMID: 30772301 | SOX2 |
| 4014 | ERCC1 | SOX2 regulated genes in Wounding from PMID: 30772301 | SOX2 |
| 4015 | IER3 | SOX2 regulated genes in Wounding from PMID: 30772301 | SOX2 |
| 4016 | TFAP2A | SOX2 regulated genes in Wounding from PMID: 30772301 | SOX2 |
| 4017 | PRSS53 | SOX2 regulated genes in Wounding from PMID: 30772301 | SOX2 |
| 4018 | CDH6 | SOX2 regulated genes in Wounding from PMID: 30772301 | SOX2 |
| 4019 | CAV2 | SOX2 regulated genes in Wounding from PMID: 30772301 | SOX2 |
| 4020 | BCAS1 | SOX2 regulated genes in Wounding from PMID: 30772301 | SOX2 |
| 4021 | BDKRB2 | SOX2 regulated genes in Wounding from PMID: 30772301 | SOX2 |
| 4022 | MYCBP | SOX2 regulated genes in Wounding from PMID: 30772301 | SOX2 |
| 4023 | STOX2 | SOX2 regulated genes in Wounding from PMID: 30772301 | SOX2 |
| 4024 | NUP107 | SOX2 regulated genes in Wounding from PMID: 30772301 | SOX2 |

| | | | |
|---|---|---|---|
| 4025 | RAI14 | SOX2 regulated genes in Wounding from PMID: 30772301 | SOX2 |
| 4026 | TMEM74 | SOX2 regulated genes in Wounding from PMID: 30772301 | SOX2 |
| 4027 | NOXA1 | SOX2 regulated genes in Wounding from PMID: 30772301 | SOX2 |
| 4028 | TMEM115 | SOX2 regulated genes in Wounding from PMID: 30772301 | SOX2 |
| 4029 | UBA3 | SOX2 regulated genes in Wounding from PMID: 30772301 | SOX2 |
| 4030 | INHBB | SOX2 regulated genes in Wounding from PMID: 30772301 | SOX2 |
| 4031 | EPHA2 | SOX2 regulated genes in Wounding from PMID: 30772301 | SOX2 |
| 4032 | CPT1A | SOX2 regulated genes in Wounding from PMID: 30772301 | SOX2 |
| 4033 | UBXN2A | SOX2 regulated genes in Wounding from PMID: 30772301 | SOX2 |
| 4034 | MICALL1 | SOX2 regulated genes in Wounding from PMID: 30772301 | SOX2 |
| 4035 | CPM | SOX2 regulated genes in Wounding from PMID: 30772301 | SOX2 |
| 4036 | STEAP1B | SOX2 regulated genes in Wounding from PMID: 30772301 | SOX2 |
| 4037 | BAG3 | SOX2 regulated genes in Wounding from PMID: 30772301 | SOX2 |
| 4038 | DCAF12 | SOX2 regulated genes in Wounding from PMID: 30772301 | SOX2 |
| 4039 | CTNNBIP1 | SOX2 regulated genes in Wounding from PMID: 30772301 | SOX2 |
| 4040 | KDM8 | SOX2 regulated genes in Wounding from PMID: 30772301 | SOX2 |
| 4041 | MANF | SOX2 regulated genes in Wounding from PMID: 30772301 | SOX2 |
| 4042 | PITPNB | SOX2 regulated genes in Wounding from PMID: 30772301 | SOX2 |
| 4043 | APOL3 | SOX2 regulated genes in Wounding from PMID: 30772301 | SOX2 |
| 4044 | APOL4 | SOX2 regulated genes in Wounding from PMID: 30772301 | SOX2 |
| 4045 | APOL2 | SOX2 regulated genes in Wounding from PMID: 30772301 | SOX2 |
| 4046 | APOL1 | SOX2 regulated genes in Wounding from PMID: 30772301 | SOX2 |
| 4047 | RCOR2 | SOX2 regulated genes in Wounding from PMID: 30772301 | SOX2 |
| 4048 | ARPP19 | SOX2 regulated genes in Wounding from PMID: 30772301 | SOX2 |
| 4049 | ECHDC2 | SOX2 regulated genes in Wounding from PMID: 30772301 | SOX2 |
| 4050 | HAPLN4 | SOX2 regulated genes in Wounding from PMID: 30772301 | SOX2 |
| 4051 | PPIF | SOX2 regulated genes in Wounding from PMID: 30772301 | SOX2 |
| 4052 | SRBD1 | SOX2 regulated genes in Wounding from PMID: 30772301 | SOX2 |
| 4053 | LRRC42 | SOX2 regulated genes in Wounding from PMID: 30772301 | SOX2 |
| 4054 | ESM1 | SOX2 regulated genes in Wounding from PMID: 30772301 | SOX2 |
| 4055 | PITRM1 | SOX2 regulated genes in Wounding from PMID: 30772301 | SOX2 |
| 4056 | METAP1D | SOX2 regulated genes in Wounding from PMID: 30772301 | SOX2 |
| 4057 | MAP2K3 | SOX2 regulated genes in Wounding from PMID: 30772301 | SOX2 |
| 4058 | RFXANK | SOX2 regulated genes in Wounding from PMID: 30772301 | SOX2 |
| 4059 | TP53 | SOX2 regulated genes in Wounding from PMID: 30772301 | SOX2 |
| 4060 | NR2C2AP | SOX2 regulated genes in Wounding from PMID: 30772301 | SOX2 |
| 4061 | AGAP1 | SOX2 regulated genes in Wounding from PMID: 30772301 | SOX2 |
| 4062 | SNAI3 | SOX2 regulated genes in Wounding from PMID: 30772301 | SOX2 |
| 4063 | NCBP2L | SOX2 regulated genes in Wounding from PMID: 30772301 | SOX2 |
| 4064 | SYCE1L | SOX2 regulated genes in Wounding from PMID: 30772301 | SOX2 |
| 4065 | TENM2 | SOX2 regulated genes in Wounding from PMID: 30772301 | SOX2 |
| 4066 | DYNC1LI1 | SOX2 regulated genes in Wounding from PMID: 30772301 | SOX2 |
| 4067 | HOXC13 | SOX2 regulated genes in Wounding from PMID: 30772301 | SOX2 |
| 4068 | LEF1 | SOX2 regulated genes in Wounding from PMID: 30772301 | SOX2 |
| 4069 | STAM2 | SOX2 regulated genes in Wounding from PMID: 30772301 | SOX2 |
| 4070 | EPB41L5 | SOX2 regulated genes in Wounding from PMID: 30772301 | SOX2 |
| 4071 | METTL1 | SOX2 regulated genes in Wounding from PMID: 30772301 | SOX2 |
| 4072 | MYZAP | SOX2 regulated genes in Wounding from PMID: 30772301 | SOX2 |
| 4073 | LMO4 | SOX2 regulated genes in Wounding from PMID: 30772301 | SOX2 |
| 4074 | MSRB3 | SOX2 regulated genes in Wounding from PMID: 30772301 | SOX2 |
| 4075 | DMBT1 | SOX2 regulated genes in Wounding from PMID: 30772301 | SOX2 |
| 4076 | UPK1B | SOX2 regulated genes in Wounding from PMID: 30772301 | SOX2 |
| 4077 | MYRF | SOX2 regulated genes in Wounding from PMID: 30772301 | SOX2 |
| 4078 | FASTKD5 | SOX2 regulated genes in Wounding from PMID: 30772301 | SOX2 |
| 4079 | PROM2 | SOX2 regulated genes in Wounding from PMID: 30772301 | SOX2 |
| 4080 | FRMPD1 | SOX2 regulated genes in Wounding from PMID: 30772301 | SOX2 |
| 4081 | KANK1 | SOX2 regulated genes in Wounding from PMID: 30772301 | SOX2 |
| 4082 | RHCG | SOX2 regulated genes in Wounding from PMID: 30772301 | SOX2 |
| 4083 | GRPEL2 | SOX2 regulated genes in Wounding from PMID: 30772301 | SOX2 |
| 4084 | MXI1 | SOX2 regulated genes in Wounding from PMID: 30772301 | SOX2 |
| 4085 | SGTB | SOX2 regulated genes in Wounding from PMID: 30772301 | SOX2 |

| | | | |
|---|---|---|---|
| 4086 | WNK4 | SOX2 regulated genes in Wounding from PMID: 30772301 | SOX2 |
| 4087 | MUS81 | SOX2 regulated genes in Wounding from PMID: 30772301 | SOX2 |
| 4088 | EIF3J | SOX2 regulated genes in Wounding from PMID: 30772301 | SOX2 |
| 4089 | VCL | SOX2 regulated genes in Wounding from PMID: 30772301 | SOX2 |
| 4090 | HOXB6 | SOX2 regulated genes in Wounding from PMID: 30772301 | SOX2 |
| 4091 | KMT5B | SOX2 regulated genes in Wounding from PMID: 30772301 | SOX2 |
| 4092 | SRPRA | SOX2 regulated genes in Wounding from PMID: 30772301 | SOX2 |
| 4093 | RBM18 | SOX2 regulated genes in Wounding from PMID: 30772301 | SOX2 |
| 4094 | HSPA5 | SOX2 regulated genes in Wounding from PMID: 30772301 | SOX2 |
| 4095 | USO1 | SOX2 regulated genes in Wounding from PMID: 30772301 | SOX2 |
| 4096 | PHTF2 | SOX2 regulated genes in Wounding from PMID: 30772301 | SOX2 |
| 4097 | SERPINA11 | SOX2 regulated genes in Wounding from PMID: 30772301 | SOX2 |
| 4098 | PDCL3 | SOX2 regulated genes in Wounding from PMID: 30772301 | SOX2 |
| 4099 | PCYOX1L | SOX2 regulated genes in Wounding from PMID: 30772301 | SOX2 |
| 4100 | SLCO5A1 | SOX2 regulated genes in Wounding from PMID: 30772301 | SOX2 |
| 4101 | SLC27A1 | SOX2 regulated genes in Wounding from PMID: 30772301 | SOX2 |
| 4102 | AK4 | SOX2 regulated genes in Wounding from PMID: 30772301 | SOX2 |
| 4103 | IL1B | SOX2 regulated genes in Wounding from PMID: 30772301 | SOX2 |
| 4104 | MPZL2 | SOX2 regulated genes in Wounding from PMID: 30772301 | SOX2 |
| 4105 | MAS1 | SOX2 regulated genes in Wounding from PMID: 30772301 | SOX2 |
| 4106 | TNFRSF11B | SOX2 regulated genes in Wounding from PMID: 30772301 | SOX2 |
| 4107 | TECTA | SOX2 regulated genes in Wounding from PMID: 30772301 | SOX2 |
| 4108 | CAPN1 | SOX2 regulated genes in Wounding from PMID: 30772301 | SOX2 |
| 4109 | CNKSR3 | SOX2 regulated genes in Wounding from PMID: 30772301 | SOX2 |
| 4110 | UBE2H | SOX2 regulated genes in Wounding from PMID: 30772301 | SOX2 |
| 4111 | SIAH1 | SOX2 regulated genes in Wounding from PMID: 30772301 | SOX2 |
| 4112 | TRIM2 | SOX2 regulated genes in Wounding from PMID: 30772301 | SOX2 |
| 4113 | CD164L2 | SOX2 regulated genes in Wounding from PMID: 30772301 | SOX2 |
| 4114 | ATP1B3 | SOX2 regulated genes in Wounding from PMID: 30772301 | SOX2 |
| 4115 | SEC24C | SOX2 regulated genes in Wounding from PMID: 30772301 | SOX2 |
| 4116 | CTTN | SOX2 regulated genes in Wounding from PMID: 30772301 | SOX2 |
| 4117 | IFFO2 | SOX2 regulated genes in Wounding from PMID: 30772301 | SOX2 |
| 4118 | GDPD3 | SOX2 regulated genes in Wounding from PMID: 30772301 | SOX2 |
| 4119 | PRICKLE2 | SOX2 regulated genes in Wounding from PMID: 30772301 | SOX2 |
| 4120 | GCSH | SOX2 regulated genes in Wounding from PMID: 30772301 | SOX2 |
| 4121 | COQ5 | SOX2 regulated genes in Wounding from PMID: 30772301 | SOX2 |
| 4122 | OAF | SOX2 regulated genes in Wounding from PMID: 30772301 | SOX2 |
| 4123 | PWWP2B | SOX2 regulated genes in Wounding from PMID: 30772301 | SOX2 |
| 4124 | HADHB | SOX2 regulated genes in Wounding from PMID: 30772301 | SOX2 |
| 4125 | PLEKHH1 | SOX2 regulated genes in Wounding from PMID: 30772301 | SOX2 |
| 4126 | ELL2 | SOX2 regulated genes in Wounding from PMID: 30772301 | SOX2 |
| 4127 | PSORS1C2 | SOX2 regulated genes in Wounding from PMID: 30772301 | SOX2 |
| 4128 | ABHD17C | SOX2 regulated genes in Wounding from PMID: 30772301 | SOX2 |
| 4129 | ELF5 | SOX2 regulated genes in Wounding from PMID: 30772301 | SOX2 |
| 4130 | TM4SF1 | SOX2 regulated genes in Wounding from PMID: 30772301 | SOX2 |
| 4131 | EIF3L | SOX2 regulated genes in Wounding from PMID: 30772301 | SOX2 |
| 4132 | EIF3D | SOX2 regulated genes in Wounding from PMID: 30772301 | SOX2 |
| 4133 | C10orf95 | SOX2 regulated genes in Wounding from PMID: 30772301 | SOX2 |
| 4134 | LYPD5 | SOX2 regulated genes in Wounding from PMID: 30772301 | SOX2 |
| 4135 | C11orf80 | SOX2 regulated genes in Wounding from PMID: 30772301 | SOX2 |
| 4136 | MAPK3 | SOX2 regulated genes in Wounding from PMID: 30772301 | SOX2 |
| 4137 | PTK7 | SOX2 regulated genes in Wounding from PMID: 30772301 | SOX2 |
| 4138 | INHBA | SOX2 regulated genes in Wounding from PMID: 30772301 | SOX2 |
| 4139 | SNIP1 | SOX2 regulated genes in Wounding from PMID: 30772301 | SOX2 |
| 4140 | EIF5AL1 | SOX2 regulated genes in Wounding from PMID: 30772301 | SOX2 |
| 4141 | RAB21 | SOX2 regulated genes in Wounding from PMID: 30772301 | SOX2 |
| 4142 | ACTN1 | SOX2 regulated genes in Wounding from PMID: 30772301 | SOX2 |
| 4143 | NIPAL1 | SOX2 regulated genes in Wounding from PMID: 30772301 | SOX2 |
| 4144 | ICMT | SOX2 regulated genes in Wounding from PMID: 30772301 | SOX2 |
| 4145 | SARDH | SOX2 regulated genes in Wounding from PMID: 30772301 | SOX2 |
| 4146 | RDH10 | SOX2 regulated genes in Wounding from PMID: 30772301 | SOX2 |

| | | | |
|---|---|---|---|
| 4147 | KIF26A | SOX2 regulated genes in Wounding from PMID: 30772301 | SOX2 |
| 4148 | RAB15 | SOX2 regulated genes in Wounding from PMID: 30772301 | SOX2 |
| 4149 | LINC02210-CRHR1 | SOX2 regulated genes in Wounding from PMID: 30772301 | SOX2 |
| 4150 | TCP1 | SOX2 regulated genes in Wounding from PMID: 30772301 | SOX2 |
| 4151 | SOX9 | SOX2 regulated genes in Wounding from PMID: 30772301 | SOX2 |
| 4152 | FGFR3 | SOX2 regulated genes in Wounding from PMID: 30772301 | SOX2 |
| 4153 | ANXA8 | SOX2 regulated genes in Wounding from PMID: 30772301 | SOX2 |
| 4154 | ANXA8L1 | SOX2 regulated genes in Wounding from PMID: 30772301 | SOX2 |
| 4155 | PTPRK | SOX2 regulated genes in Wounding from PMID: 30772301 | SOX2 |
| 4156 | SEC11A | SOX2 regulated genes in Wounding from PMID: 30772301 | SOX2 |
| 4157 | PTGR1 | SOX2 regulated genes in Wounding from PMID: 30772301 | SOX2 |
| 4158 | CNTFR | SOX2 regulated genes in Wounding from PMID: 30772301 | SOX2 |
| 4159 | ACTB | SOX2 regulated genes in Wounding from PMID: 30772301 | SOX2 |
| 4160 | TMEM39A | SOX2 regulated genes in Wounding from PMID: 30772301 | SOX2 |
| 4161 | SFRP1 | SOX2 regulated genes in Wounding from PMID: 30772301 | SOX2 |
| 4162 | EPHB6 | SOX2 regulated genes in Wounding from PMID: 30772301 | SOX2 |
| 4163 | PIGH | SOX2 regulated genes in Wounding from PMID: 30772301 | SOX2 |
| 4164 | ZFAND6 | SOX2 regulated genes in Wounding from PMID: 30772301 | SOX2 |
| 4165 | NECTIN3 | SOX2 regulated genes in Wounding from PMID: 30772301 | SOX2 |
| 4166 | DOK4 | SOX2 regulated genes in Wounding from PMID: 30772301 | SOX2 |
| 4167 | NRARP | SOX2 regulated genes in Wounding from PMID: 30772301 | SOX2 |
| 4168 | NTN4 | SOX2 regulated genes in Wounding from PMID: 30772301 | SOX2 |
| 4169 | TSFM | SOX2 regulated genes in Wounding from PMID: 30772301 | SOX2 |
| 4170 | CXXC5 | SOX2 regulated genes in Wounding from PMID: 30772301 | SOX2 |
| 4171 | COPB1 | SOX2 regulated genes in Wounding from PMID: 30772301 | SOX2 |
| 4172 | SCNN1B | SOX2 regulated genes in Wounding from PMID: 30772301 | SOX2 |
| 4173 | FAF2 | SOX2 regulated genes in Wounding from PMID: 30772301 | SOX2 |
| 4174 | ACTR1A | SOX2 regulated genes in Wounding from PMID: 30772301 | SOX2 |
| 4175 | CLCF1 | SOX2 regulated genes in Wounding from PMID: 30772301 | SOX2 |
| 4176 | CPEB2 | SOX2 regulated genes in Wounding from PMID: 30772301 | SOX2 |
| 4177 | MAL | SOX2 regulated genes in Wounding from PMID: 30772301 | SOX2 |
| 4178 | RALA | SOX2 regulated genes in Wounding from PMID: 30772301 | SOX2 |
| 4179 | MOB1B | SOX2 regulated genes in Wounding from PMID: 30772301 | SOX2 |
| 4180 | UBA1 | SOX2 regulated genes in Wounding from PMID: 30772301 | SOX2 |
| 4181 | EVA1A | SOX2 regulated genes in Wounding from PMID: 30772301 | SOX2 |
| 4182 | VGLL3 | SOX2 regulated genes in Wounding from PMID: 30772301 | SOX2 |
| 4183 | LBH | SOX2 regulated genes in Wounding from PMID: 30772301 | SOX2 |
| 4184 | C6orf132 | SOX2 regulated genes in Wounding from PMID: 30772301 | SOX2 |
| 4185 | CAND1 | SOX2 regulated genes in Wounding from PMID: 30772301 | SOX2 |
| 4186 | NUP35 | SOX2 regulated genes in Wounding from PMID: 30772301 | SOX2 |
| 4187 | SNRPA | SOX2 regulated genes in Wounding from PMID: 30772301 | SOX2 |
| 4188 | FOSL1 | SOX2 regulated genes in Wounding from PMID: 30772301 | SOX2 |
| 4189 | PLPP5 | SOX2 regulated genes in Wounding from PMID: 30772301 | SOX2 |
| 4190 | DLX4 | SOX2 regulated genes in Wounding from PMID: 30772301 | SOX2 |
| 4191 | BRAP | SOX2 regulated genes in Wounding from PMID: 30772301 | SOX2 |
| 4192 | HS6ST1 | SOX2 regulated genes in Wounding from PMID: 30772301 | SOX2 |
| 4193 | GPC4 | SOX2 regulated genes in Wounding from PMID: 30772301 | SOX2 |
| 4194 | DPYSL5 | SOX2 regulated genes in Wounding from PMID: 30772301 | SOX2 |
| 4195 | PPFIA3 | SOX2 regulated genes in Wounding from PMID: 30772301 | SOX2 |
| 4196 | MMP1 | SOX2 regulated genes in Wounding from PMID: 30772301 | SOX2 |
| 4197 | ATP6V1A | SOX2 regulated genes in Wounding from PMID: 30772301 | SOX2 |
| 4198 | SMNDC1 | SOX2 regulated genes in Wounding from PMID: 30772301 | SOX2 |
| 4199 | TMEM121 | SOX2 regulated genes in Wounding from PMID: 30772301 | SOX2 |
| 4200 | GJB4 | SOX2 regulated genes in Wounding from PMID: 30772301 | SOX2 |
| 4201 | HNRNPAB | SOX2 regulated genes in Wounding from PMID: 30772301 | SOX2 |
| 4202 | BSPRY | SOX2 regulated genes in Wounding from PMID: 30772301 | SOX2 |
| 4203 | BRD1 | SOX2 regulated genes in Wounding from PMID: 30772301 | SOX2 |
| 4204 | SUFU | SOX2 regulated genes in Wounding from PMID: 30772301 | SOX2 |
| 4205 | TLE2 | SOX2 regulated genes in Wounding from PMID: 30772301 | SOX2 |
| 4206 | ARHGAP39 | SOX2 regulated genes in Wounding from PMID: 30772301 | SOX2 |
| 4207 | EFNA2 | SOX2 regulated genes in Wounding from PMID: 30772301 | SOX2 |

| | | | |
|---|---|---|---|
| 4208 | ZNF821 | SOX2 regulated genes in Wounding from PMID: 30772301 | SOX2 |
| 4209 | FGF19 | SOX2 regulated genes in Wounding from PMID: 30772301 | SOX2 |
| 4210 | IL11RA | SOX2 regulated genes in Wounding from PMID: 30772301 | SOX2 |
| 4211 | MYADM | SOX2 regulated genes in Wounding from PMID: 30772301 | SOX2 |
| 4212 | UPF1 | SOX2 regulated genes in Wounding from PMID: 30772301 | SOX2 |
| 4213 | PLCD3 | SOX2 regulated genes in Wounding from PMID: 30772301 | SOX2 |
| 4214 | PRDX2 | SOX2 regulated genes in Wounding from PMID: 30772301 | SOX2 |
| 4215 | ATP2B4 | SOX2 regulated genes in Wounding from PMID: 30772301 | SOX2 |
| 4216 | MRTO4 | SOX2 regulated genes in Wounding from PMID: 30772301 | SOX2 |
| 4217 | UNC119B | SOX2 regulated genes in Wounding from PMID: 30772301 | SOX2 |
| 4218 | PLA2G2F | SOX2 regulated genes in Wounding from PMID: 30772301 | SOX2 |
| 4219 | ODC1 | SOX2 regulated genes in Wounding from PMID: 30772301 | SOX2 |
| 4220 | FLNB | SOX2 regulated genes in Wounding from PMID: 30772301 | SOX2 |
| 4221 | SPTSSB | SOX2 regulated genes in Wounding from PMID: 30772301 | SOX2 |
| 4222 | MAP6D1 | SOX2 regulated genes in Wounding from PMID: 30772301 | SOX2 |
| 4223 | YOD1 | SOX2 regulated genes in Wounding from PMID: 30772301 | SOX2 |
| 4224 | PDIA4 | SOX2 regulated genes in Wounding from PMID: 30772301 | SOX2 |
| 4225 | GNG4 | SOX2 regulated genes in Wounding from PMID: 30772301 | SOX2 |
| 4226 | MTA2 | SOX2 regulated genes in Wounding from PMID: 30772301 | SOX2 |
| 4227 | B3GNT3 | SOX2 regulated genes in Wounding from PMID: 30772301 | SOX2 |
| 4228 | KRT17 | SOX2 regulated genes in Wounding from PMID: 30772301 | SOX2 |
| 4229 | DDB1 | SOX2 regulated genes in Wounding from PMID: 30772301 | SOX2 |
| 4230 | CGN | SOX2 regulated genes in Wounding from PMID: 30772301 | SOX2 |
| 4231 | CD44 | SOX2 regulated genes in Wounding from PMID: 30772301 | SOX2 |
| 4232 | CHTF8 | SOX2 regulated genes in Wounding from PMID: 30772301 | SOX2 |
| 4233 | VAMP8 | SOX2 regulated genes in Wounding from PMID: 30772301 | SOX2 |
| 4234 | POR | SOX2 regulated genes in Wounding from PMID: 30772301 | SOX2 |
| 4235 | ARHGAP21 | SOX2 regulated genes in Wounding from PMID: 30772301 | SOX2 |
| 4236 | SOX13 | SOX2 regulated genes in Wounding from PMID: 30772301 | SOX2 |
| 4237 | COBLL1 | SOX2 regulated genes in Wounding from PMID: 30772301 | SOX2 |
| 4238 | ARL15 | SOX2 regulated genes in Wounding from PMID: 30772301 | SOX2 |
| 4239 | CCSER1 | SOX2 regulated genes in Wounding from PMID: 30772301 | SOX2 |
| 4240 | CHDH | SOX2 regulated genes in Wounding from PMID: 30772301 | SOX2 |
| 4241 | PAFAH1B1 | SOX2 regulated genes in Wounding from PMID: 30772301 | SOX2 |
| 4242 | SNTB2 | SOX2 regulated genes in Wounding from PMID: 30772301 | SOX2 |
| 4243 | RNF40 | SOX2 regulated genes in Wounding from PMID: 30772301 | SOX2 |
| 4244 | SS18L2 | SOX2 regulated genes in Wounding from PMID: 30772301 | SOX2 |
| 4245 | KLK10 | SOX2 regulated genes in Wounding from PMID: 30772301 | SOX2 |
| 4246 | CYB561D1 | SOX2 regulated genes in Wounding from PMID: 30772301 | SOX2 |
| 4247 | MYO18A | SOX2 regulated genes in Wounding from PMID: 30772301 | SOX2 |
| 4248 | CDH3 | SOX2 regulated genes in Wounding from PMID: 30772301 | SOX2 |
| 4249 | HNRNPM | SOX2 regulated genes in Wounding from PMID: 30772301 | SOX2 |
| 4250 | ZNF608 | SOX2 regulated genes in Wounding from PMID: 30772301 | SOX2 |
| 4251 | ATG4B | SOX2 regulated genes in Wounding from PMID: 30772301 | SOX2 |
| 4252 | CDK18 | SOX2 regulated genes in Wounding from PMID: 30772301 | SOX2 |
| 4253 | HDAC5 | SOX2 regulated genes in Wounding from PMID: 30772301 | SOX2 |
| 4254 | SNX7 | SOX2 regulated genes in Wounding from PMID: 30772301 | SOX2 |
| 4255 | METTL16 | SOX2 regulated genes in Wounding from PMID: 30772301 | SOX2 |
| 4256 | GBP6 | SOX2 regulated genes in Wounding from PMID: 30772301 | SOX2 |
| 4257 | KPNA4 | SOX2 regulated genes in Wounding from PMID: 30772301 | SOX2 |
| 4258 | CSNK1A1 | SOX2 regulated genes in Wounding from PMID: 30772301 | SOX2 |
| 4259 | BLZF1 | SOX2 regulated genes in Wounding from PMID: 30772301 | SOX2 |
| 4260 | PADI1 | SOX2 regulated genes in Wounding from PMID: 30772301 | SOX2 |
| 4261 | TLN2 | SOX2 regulated genes in Wounding from PMID: 30772301 | SOX2 |
| 4262 | KRTAP2-3 | SOX2 regulated genes in Wounding from PMID: 30772301 | SOX2 |
| 4263 | KRTAP4-11 | SOX2 regulated genes in Wounding from PMID: 30772301 | SOX2 |
| 4264 | KRTAP4-8 | SOX2 regulated genes in Wounding from PMID: 30772301 | SOX2 |
| 4265 | CAPN12 | SOX2 regulated genes in Wounding from PMID: 30772301 | SOX2 |
| 4266 | SPAG9 | SOX2 regulated genes in Wounding from PMID: 30772301 | SOX2 |
| 4267 | TJP1 | SOX2 regulated genes in Wounding from PMID: 30772301 | SOX2 |
| 4268 | C1orf52 | SOX2 regulated genes in Wounding from PMID: 30772301 | SOX2 |

| | | | |
|---|---|---|---|
| 4269 | APLP2 | SOX2 regulated genes in Wounding from PMID: 30772301 | SOX2 |
| 4270 | PTPN14 | SOX2 regulated genes in Wounding from PMID: 30772301 | SOX2 |
| 4271 | ACBD3 | SOX2 regulated genes in Wounding from PMID: 30772301 | SOX2 |
| 4272 | PDE4B | SOX2 regulated genes in Wounding from PMID: 30772301 | SOX2 |
| 4273 | PSPN | SOX2 regulated genes in Wounding from PMID: 30772301 | SOX2 |
| 4274 | HYOU1 | SOX2 regulated genes in Wounding from PMID: 30772301 | SOX2 |
| 4275 | CREB3L4 | SOX2 regulated genes in Wounding from PMID: 30772301 | SOX2 |
| 4276 | EDN2 | SOX2 regulated genes in Wounding from PMID: 30772301 | SOX2 |
| 4277 | CHFR | SOX2 regulated genes in Wounding from PMID: 30772301 | SOX2 |
| 4278 | WDR12 | SOX2 regulated genes in Wounding from PMID: 30772301 | SOX2 |
| 4279 | SLC40A1 | SOX2 regulated genes in Wounding from PMID: 30772301 | SOX2 |
| 4280 | MICA | SOX2 regulated genes in Wounding from PMID: 30772301 | SOX2 |
| 4281 | CDV3 | SOX2 regulated genes in Wounding from PMID: 30772301 | SOX2 |
| 4282 | FZD7 | SOX2 regulated genes in Wounding from PMID: 30772301 | SOX2 |
| 4283 | CALR | SOX2 regulated genes in Wounding from PMID: 30772301 | SOX2 |
| 4284 | SPRR3 | SOX2 regulated genes in Wounding from PMID: 30772301 | SOX2 |
| 4285 | TRAPPC4 | SOX2 regulated genes in Wounding from PMID: 30772301 | SOX2 |
| 4286 | CLEC2D | SOX2 regulated genes in Wounding from PMID: 30772301 | SOX2 |
| 4287 | DENND2C | SOX2 regulated genes in Wounding from PMID: 30772301 | SOX2 |
| 4288 | FXYD4 | SOX2 regulated genes in Wounding from PMID: 30772301 | SOX2 |
| 4289 | MAST4 | SOX2 regulated genes in Wounding from PMID: 30772301 | SOX2 |
| 4290 | MEN1 | SOX2 regulated genes in Wounding from PMID: 30772301 | SOX2 |
| 4291 | PRDM4 | SOX2 regulated genes in Wounding from PMID: 30772301 | SOX2 |
| 4292 | BAHD1 | SOX2 regulated genes in Wounding from PMID: 30772301 | SOX2 |
| 4293 | UBQLN4 | SOX2 regulated genes in Wounding from PMID: 30772301 | SOX2 |
| 4294 | CELSR2 | SOX2 regulated genes in Wounding from PMID: 30772301 | SOX2 |
| 4295 | TSEN2 | SOX2 regulated genes in Wounding from PMID: 30772301 | SOX2 |
| 4296 | PRPF31 | SOX2 regulated genes in Wounding from PMID: 30772301 | SOX2 |
| 4297 | EPS8L2 | SOX2 regulated genes in Wounding from PMID: 30772301 | SOX2 |
| 4298 | IGSF9 | SOX2 regulated genes in Wounding from PMID: 30772301 | SOX2 |
| 4299 | SLC16A6 | SOX2 regulated genes in Wounding from PMID: 30772301 | SOX2 |
| 4300 | GTF2B | SOX2 regulated genes in Wounding from PMID: 30772301 | SOX2 |
| 4301 | SH3TC2 | SOX2 regulated genes in Wounding from PMID: 30772301 | SOX2 |
| 4302 | EPHA1 | SOX2 regulated genes in Wounding from PMID: 30772301 | SOX2 |
| 4303 | MLF1 | SOX2 regulated genes in Wounding from PMID: 30772301 | SOX2 |
| 4304 | RUVBL2 | SOX2 regulated genes in Wounding from PMID: 30772301 | SOX2 |
| 4305 | MAP7 | SOX2 regulated genes in Wounding from PMID: 30772301 | SOX2 |
| 4306 | MYO19 | SOX2 regulated genes in Wounding from PMID: 30772301 | SOX2 |
| 4307 | SF1 | SOX2 regulated genes in Wounding from PMID: 30772301 | SOX2 |
| 4308 | CTTNBP2 | SOX2 regulated genes in Wounding from PMID: 30772301 | SOX2 |
| 4309 | KDF1 | SOX2 regulated genes in Wounding from PMID: 30772301 | SOX2 |
| 4310 | PPL | SOX2 regulated genes in Wounding from PMID: 30772301 | SOX2 |
| 4311 | EFHD1 | SOX2 regulated genes in Wounding from PMID: 30772301 | SOX2 |
| 4312 | GSTM3 | SOX2 regulated genes in Wounding from PMID: 30772301 | SOX2 |
| 4313 | TIAM2 | SOX2 regulated genes in Wounding from PMID: 30772301 | SOX2 |
| 4314 | CMPK1 | SOX2 regulated genes in Wounding from PMID: 30772301 | SOX2 |
| 4315 | KRTAP1-5 | SOX2 regulated genes in Wounding from PMID: 30772301 | SOX2 |
| 4316 | KRTAP1-3 | SOX2 regulated genes in Wounding from PMID: 30772301 | SOX2 |
| 4317 | KRTAP1-1 | SOX2 regulated genes in Wounding from PMID: 30772301 | SOX2 |
| 4318 | DNAJA4 | SOX2 regulated genes in Wounding from PMID: 30772301 | SOX2 |
| 4319 | ERC2 | SOX2 regulated genes in Wounding from PMID: 30772301 | SOX2 |
| 4320 | PPFIBP2 | SOX2 regulated genes in Wounding from PMID: 30772301 | SOX2 |
| 4321 | INTS4 | SOX2 regulated genes in Wounding from PMID: 30772301 | SOX2 |
| 4322 | S100A6 | SOX2 regulated genes in Wounding from PMID: 30772301 | SOX2 |
| 4323 | MOB4 | SOX2 regulated genes in Wounding from PMID: 30772301 | SOX2 |
| 4324 | KLC1 | SOX2 regulated genes in Wounding from PMID: 30772301 | SOX2 |
| 4325 | LSR | SOX2 regulated genes in Wounding from PMID: 30772301 | SOX2 |
| 4326 | CAPN8 | SOX2 regulated genes in Wounding from PMID: 30772301 | SOX2 |
| 4327 | NHSL1 | SOX2 regulated genes in Wounding from PMID: 30772301 | SOX2 |
| 4328 | HES2 | SOX2 regulated genes in Wounding from PMID: 30772301 | SOX2 |
| 4329 | S100A16 | SOX2 regulated genes in Wounding from PMID: 30772301 | SOX2 |

| | | | |
|---|---|---|---|
| 4330 | PUM1 | SOX2 regulated genes in Wounding from PMID: 30772301 | SOX2 |
| 4331 | PLXNA2 | SOX2 regulated genes in Wounding from PMID: 30772301 | SOX2 |
| 4332 | PKD1L3 | SOX2 regulated genes in Wounding from PMID: 30772301 | SOX2 |
| 4333 | UBE2G1 | SOX2 regulated genes in Wounding from PMID: 30772301 | SOX2 |
| 4334 | NDEL1 | SOX2 regulated genes in Wounding from PMID: 30772301 | SOX2 |
| 4335 | SGMS2 | SOX2 regulated genes in Wounding from PMID: 30772301 | SOX2 |
| 4336 | IQGAP1 | SOX2 regulated genes in Wounding from PMID: 30772301 | SOX2 |
| 4337 | NRDC | SOX2 regulated genes in Wounding from PMID: 30772301 | SOX2 |
| 4338 | CSGALNACT1 | SOX2 regulated genes in Wounding from PMID: 30772301 | SOX2 |
| 4339 | LRRC8E | SOX2 regulated genes in Wounding from PMID: 30772301 | SOX2 |
| 4340 | GMCL1 | SOX2 regulated genes in Wounding from PMID: 30772301 | SOX2 |
| 4341 | WNT11 | SOX2 regulated genes in Wounding from PMID: 30772301 | SOX2 |
| 4342 | GALNT3 | SOX2 regulated genes in Wounding from PMID: 30772301 | SOX2 |
| 4343 | PDLIM7 | SOX2 regulated genes in Wounding from PMID: 30772301 | SOX2 |
| 4344 | WDR26 | SOX2 regulated genes in Wounding from PMID: 30772301 | SOX2 |
| 4345 | OCLN | SOX2 regulated genes in Wounding from PMID: 30772301 | SOX2 |
| 4346 | FARP2 | SOX2 regulated genes in Wounding from PMID: 30772301 | SOX2 |
| 4347 | RGS12 | SOX2 regulated genes in Wounding from PMID: 30772301 | SOX2 |
| 4348 | TM4SF4 | SOX2 regulated genes in Wounding from PMID: 30772301 | SOX2 |
| 4349 | CLIP1 | SOX2 regulated genes in Wounding from PMID: 30772301 | SOX2 |
| 4350 | ADGRL3 | SOX2 regulated genes in Wounding from PMID: 30772301 | SOX2 |
| 4351 | KRT34 | SOX2 regulated genes in Wounding from PMID: 30772301 | SOX2 |
| 4352 | NKAPL | SOX2 regulated genes in Wounding from PMID: 30772301 | SOX2 |
| 4353 | BHLHE40 | SOX2 regulated genes in Wounding from PMID: 30772301 | SOX2 |
| 4354 | NIPAL3 | SOX2 regulated genes in Wounding from PMID: 30772301 | SOX2 |
| 4355 | KALRN | SOX2 regulated genes in Wounding from PMID: 30772301 | SOX2 |
| 4356 | ETV2 | SOX2 regulated genes in Wounding from PMID: 30772301 | SOX2 |
| 4357 | B3GALNT2 | SOX2 regulated genes in Wounding from PMID: 30772301 | SOX2 |
| 4358 | GTF2I | SOX2 regulated genes in Wounding from PMID: 30772301 | SOX2 |
| 4359 | HNRNPC | SOX2 regulated genes in Wounding from PMID: 30772301 | SOX2 |
| 4360 | HNRNPCL1 | SOX2 regulated genes in Wounding from PMID: 30772301 | SOX2 |
| 4361 | HNRNPCL2 | SOX2 regulated genes in Wounding from PMID: 30772301 | SOX2 |
| 4362 | LRIG1 | SOX2 regulated genes in Wounding from PMID: 30772301 | SOX2 |
| 4363 | TRIM33 | SOX2 regulated genes in Wounding from PMID: 30772301 | SOX2 |
| 4364 | SPNS2 | SOX2 regulated genes in Wounding from PMID: 30772301 | SOX2 |
| 4365 | HAS3 | SOX2 regulated genes in Wounding from PMID: 30772301 | SOX2 |
| 4366 | INSIG2 | SOX2 regulated genes in Wounding from PMID: 30772301 | SOX2 |
| 4367 | BNIPL | SOX2 regulated genes in Wounding from PMID: 30772301 | SOX2 |
| 4368 | CETN3 | SOX2 regulated genes in Wounding from PMID: 30772301 | SOX2 |
| 4369 | TRAK2 | SOX2 regulated genes in Wounding from PMID: 30772301 | SOX2 |
| 4370 | MAST1 | SOX2 regulated genes in Wounding from PMID: 30772301 | SOX2 |
| 4371 | NECTIN4 | SOX2 regulated genes in Wounding from PMID: 30772301 | SOX2 |
| 4372 | ACTN4 | SOX2 regulated genes in Wounding from PMID: 30772301 | SOX2 |
| 4373 | TNFAIP6 | SOX2 regulated genes in Wounding from PMID: 30772301 | SOX2 |
| 4374 | TMBIM1 | SOX2 regulated genes in Wounding from PMID: 30772301 | SOX2 |
| 4375 | CRNN | SOX2 regulated genes in Wounding from PMID: 30772301 | SOX2 |
| 4376 | SMIM5 | SOX2 regulated genes in Wounding from PMID: 30772301 | SOX2 |
| 4377 | SIRT6 | SOX2 regulated genes in Wounding from PMID: 30772301 | SOX2 |
| 4378 | FAM167A | SOX2 regulated genes in Wounding from PMID: 30772301 | SOX2 |
| 4379 | DNAJB2 | SOX2 regulated genes in Wounding from PMID: 30772301 | SOX2 |
| 4380 | GOLT1B | SOX2 regulated genes in Wounding from PMID: 30772301 | SOX2 |
| 4381 | KLHL18 | SOX2 regulated genes in Wounding from PMID: 30772301 | SOX2 |
| 4382 | ZYX | SOX2 regulated genes in Wounding from PMID: 30772301 | SOX2 |
| 4383 | ATXN1 | SOX2 regulated genes in Wounding from PMID: 30772301 | SOX2 |
| 4384 | ELAVL1 | SOX2 regulated genes in Wounding from PMID: 30772301 | SOX2 |
| 4385 | ACOT1 | SOX2 regulated genes in Wounding from PMID: 30772301 | SOX2 |
| 4386 | ACOT2 | SOX2 regulated genes in Wounding from PMID: 30772301 | SOX2 |
| 4387 | NCKAP5 | SOX2 regulated genes in Wounding from PMID: 30772301 | SOX2 |
| 4388 | DSP | SOX2 regulated genes in Wounding from PMID: 30772301 | SOX2 |
| 4389 | PFDN1 | SOX2 regulated genes in Wounding from PMID: 30772301 | SOX2 |
| 4390 | TET3 | SOX2 regulated genes in Wounding from PMID: 30772301 | SOX2 |

| | | | |
|---|---|---|---|
| 4391 | KRT32 | SOX2 regulated genes in Wounding from PMID: 30772301 | SOX2 |
| 4392 | TMEM120B | SOX2 regulated genes in Wounding from PMID: 30772301 | SOX2 |
| 4393 | CDK2AP1 | SOX2 regulated genes in Wounding from PMID: 30772301 | SOX2 |
| 4394 | PRR7 | SOX2 regulated genes in Wounding from PMID: 30772301 | SOX2 |
| 4395 | TMEM184A | SOX2 regulated genes in Wounding from PMID: 30772301 | SOX2 |
| 4396 | SMUG1 | SOX2 regulated genes in Wounding from PMID: 30772301 | SOX2 |
| 4397 | RAP1GAP | SOX2 regulated genes in Wounding from PMID: 30772301 | SOX2 |
| 4398 | FURIN | SOX2 regulated genes in Wounding from PMID: 30772301 | SOX2 |
| 4399 | PTPN11 | SOX2 regulated genes in Wounding from PMID: 30772301 | SOX2 |
| 4400 | EIF2B3 | SOX2 regulated genes in Wounding from PMID: 30772301 | SOX2 |
| 4401 | AKAP9 | SOX2 regulated genes in Wounding from PMID: 30772301 | SOX2 |
| 4402 | SERTAD4 | SOX2 regulated genes in Wounding from PMID: 30772301 | SOX2 |
| 4403 | SNRPD2 | SOX2 regulated genes in Wounding from PMID: 30772301 | SOX2 |
| 4404 | SMIM7 | SOX2 regulated genes in Wounding from PMID: 30772301 | SOX2 |
| 4405 | MYL6 | SOX2 regulated genes in Wounding from PMID: 30772301 | SOX2 |
| 4406 | TLX1 | SOX2 regulated genes in Wounding from PMID: 30772301 | SOX2 |
| 4407 | TPPP | SOX2 regulated genes in Wounding from PMID: 30772301 | SOX2 |
| 4408 | KRTCAP3 | SOX2 regulated genes in Wounding from PMID: 30772301 | SOX2 |
| 4409 | TBCD | SOX2 regulated genes in Wounding from PMID: 30772301 | SOX2 |
| 4410 | PIGM | SOX2 regulated genes in Wounding from PMID: 30772301 | SOX2 |
| 4411 | FZD3 | SOX2 regulated genes in Wounding from PMID: 30772301 | SOX2 |
| 4412 | RNF222 | SOX2 regulated genes in Wounding from PMID: 30772301 | SOX2 |
| 4413 | LINGO1 | SOX2 regulated genes in Wounding from PMID: 30772301 | SOX2 |
| 4414 | SPACA6 | SOX2 regulated genes in Wounding from PMID: 30772301 | SOX2 |
| 4415 | LMNB2 | SOX2 regulated genes in Wounding from PMID: 30772301 | SOX2 |
| 4416 | NAPEPLD | SOX2 regulated genes in Wounding from PMID: 30772301 | SOX2 |
| 4417 | ABCF3 | SOX2 regulated genes in Wounding from PMID: 30772301 | SOX2 |
| 4418 | ALG8 | SOX2 regulated genes in Wounding from PMID: 30772301 | SOX2 |
| 4419 | TMED2 | SOX2 regulated genes in Wounding from PMID: 30772301 | SOX2 |
| 4420 | HES1 | SOX2 regulated genes in Wounding from PMID: 30772301 | SOX2 |
| 4421 | SFPQ | SOX2 regulated genes in Wounding from PMID: 30772301 | SOX2 |
| 4422 | MARVELD3 | SOX2 regulated genes in Wounding from PMID: 30772301 | SOX2 |
| 4423 | PYROXD1 | SOX2 regulated genes in Wounding from PMID: 30772301 | SOX2 |
| 4424 | NUAK1 | SOX2 regulated genes in Wounding from PMID: 30772301 | SOX2 |
| 4425 | LTBP4 | SOX2 regulated genes in Wounding from PMID: 30772301 | SOX2 |
| 4426 | VANGL2 | SOX2 regulated genes in Wounding from PMID: 30772301 | SOX2 |
| 4427 | EFNA3 | SOX2 regulated genes in Wounding from PMID: 30772301 | SOX2 |
| 4428 | JMJD4 | SOX2 regulated genes in Wounding from PMID: 30772301 | SOX2 |
| 4429 | CALM1 | SOX2 regulated genes in Wounding from PMID: 30772301 | SOX2 |
| 4430 | PRKCZ | SOX2 regulated genes in Wounding from PMID: 30772301 | SOX2 |
| 4431 | DDR1 | SOX2 regulated genes in Wounding from PMID: 30772301 | SOX2 |
| 4432 | SNRPA1 | SOX2 regulated genes in Wounding from PMID: 30772301 | SOX2 |
| 4433 | ARHGEF28 | SOX2 regulated genes in Wounding from PMID: 30772301 | SOX2 |
| 4434 | PHF23 | SOX2 regulated genes in Wounding from PMID: 30772301 | SOX2 |
| 4435 | SLC2A1 | SOX2 regulated genes in Wounding from PMID: 30772301 | SOX2 |
| 4436 | CNTNAP1 | SOX2 regulated genes in Wounding from PMID: 30772301 | SOX2 |
| 4437 | VDR | SOX2 regulated genes in Wounding from PMID: 30772301 | SOX2 |
| 4438 | LMNA | SOX2 regulated genes in Wounding from PMID: 30772301 | SOX2 |
| 4439 | GPRC5D | SOX2 regulated genes in Wounding from PMID: 30772301 | SOX2 |
| 4440 | COPB2 | SOX2 regulated genes in Wounding from PMID: 30772301 | SOX2 |
| 4441 | EXOC7 | SOX2 regulated genes in Wounding from PMID: 30772301 | SOX2 |
| 4442 | GAS8 | SOX2 regulated genes in Wounding from PMID: 30772301 | SOX2 |
| 4443 | GLI2 | SOX2 regulated genes in Wounding from PMID: 30772301 | SOX2 |
| 4444 | AFDN | SOX2 regulated genes in Wounding from PMID: 30772301 | SOX2 |
| 4445 | THEM4 | SOX2 regulated genes in Wounding from PMID: 30772301 | SOX2 |
| 4446 | ST14 | SOX2 regulated genes in Wounding from PMID: 30772301 | SOX2 |
| 4447 | RRM2 | SOX2 regulated genes in Wounding from PMID: 30772301 | SOX2 |
| 4448 | RABL2A | SOX2 regulated genes in Wounding from PMID: 30772301 | SOX2 |
| 4449 | PPM1G | SOX2 regulated genes in Wounding from PMID: 30772301 | SOX2 |
| 4450 | KIF21A | SOX2 regulated genes in Wounding from PMID: 30772301 | SOX2 |
| 4451 | NPM2 | SOX2 regulated genes in Wounding from PMID: 30772301 | SOX2 |

| | | | |
|---|---|---|---|
| 4452 | EIF2S3B | SOX2 regulated genes in Wounding from PMID: 30772301 | SOX2 |
| 4453 | KRT8 | SOX2 regulated genes in Wounding from PMID: 30772301 | SOX2 |
| 4454 | NKD2 | SOX2 regulated genes in Wounding from PMID: 30772301 | SOX2 |
| 4455 | NUDT21 | SOX2 regulated genes in Wounding from PMID: 30772301 | SOX2 |
| 4456 | SYT13 | SOX2 regulated genes in Wounding from PMID: 30772301 | SOX2 |
| 4457 | TNN | SOX2 regulated genes in Wounding from PMID: 30772301 | SOX2 |
| 4458 | XXYLT1 | SOX2 regulated genes in Wounding from PMID: 30772301 | SOX2 |
| 4459 | LMTK3 | SOX2 regulated genes in Wounding from PMID: 30772301 | SOX2 |
| 4460 | THUMPD3 | SOX2 regulated genes in Wounding from PMID: 30772301 | SOX2 |
| 4461 | FOXJ1 | SOX2 regulated genes in Wounding from PMID: 30772301 | SOX2 |
| 4462 | ENAH | SOX2 regulated genes in Wounding from PMID: 30772301 | SOX2 |
| 4463 | ARHGAP32 | SOX2 regulated genes in Wounding from PMID: 30772301 | SOX2 |
| 4464 | CAPN2 | SOX2 regulated genes in Wounding from PMID: 30772301 | SOX2 |
| 4465 | TMEM54 | SOX2 regulated genes in Wounding from PMID: 30772301 | SOX2 |
| 4466 | VPS45 | SOX2 regulated genes in Wounding from PMID: 30772301 | SOX2 |
| 4467 | FAM8A1 | SOX2 regulated genes in Wounding from PMID: 30772301 | SOX2 |
| 4468 | TMEM231 | SOX2 regulated genes in Wounding from PMID: 30772301 | SOX2 |
| 4469 | S100A14 | SOX2 regulated genes in Wounding from PMID: 30772301 | SOX2 |
| 4470 | ARHGEF5 | SOX2 regulated genes in Wounding from PMID: 30772301 | SOX2 |
| 4471 | PPP1R2 | SOX2 regulated genes in Wounding from PMID: 30772301 | SOX2 |
| 4472 | PPP1R2B | SOX2 regulated genes in Wounding from PMID: 30772301 | SOX2 |
| 4473 | RAB3IP | SOX2 regulated genes in Wounding from PMID: 30772301 | SOX2 |
| 4474 | TAX1BP1 | SOX2 regulated genes in Wounding from PMID: 30772301 | SOX2 |
| 4475 | PM20D1 | SOX2 regulated genes in Wounding from PMID: 30772301 | SOX2 |
| 4476 | EMP1 | SOX2 regulated genes in Wounding from PMID: 30772301 | SOX2 |
| 4477 | FZD10 | SOX2 regulated genes in Wounding from PMID: 30772301 | SOX2 |
| 4478 | UBE2O | SOX2 regulated genes in Wounding from PMID: 30772301 | SOX2 |
| 4479 | FAM98A | SOX2 regulated genes in Wounding from PMID: 30772301 | SOX2 |
| 4480 | CNN3 | SOX2 regulated genes in Wounding from PMID: 30772301 | SOX2 |
| 4481 | STK16 | SOX2 regulated genes in Wounding from PMID: 30772301 | SOX2 |
| 4482 | FBRS | SOX2 regulated genes in Wounding from PMID: 30772301 | SOX2 |
| 4483 | TCERG1 | SOX2 regulated genes in Wounding from PMID: 30772301 | SOX2 |
| 4484 | FAM136A | SOX2 regulated genes in Wounding from PMID: 30772301 | SOX2 |
| 4485 | ANP32E | SOX2 regulated genes in Wounding from PMID: 30772301 | SOX2 |
| 4486 | NEMF | SOX2 regulated genes in Wounding from PMID: 30772301 | SOX2 |
| 4487 | ZNF326 | SOX2 regulated genes in Wounding from PMID: 30772301 | SOX2 |
| 4488 | MTUS1 | SOX2 regulated genes in Wounding from PMID: 30772301 | SOX2 |
| 4489 | ADGRG1 | SOX2 regulated genes in Wounding from PMID: 30772301 | SOX2 |
| 4490 | NBEAL2 | SOX2 regulated genes in Wounding from PMID: 30772301 | SOX2 |
| 4491 | SIPA1L2 | SOX2 regulated genes in Wounding from PMID: 30772301 | SOX2 |
| 4492 | LLGL1 | SOX2 regulated genes in Wounding from PMID: 30772301 | SOX2 |
| 4493 | SPG7 | SOX2 regulated genes in Wounding from PMID: 30772301 | SOX2 |
| 4494 | MTX2 | SOX2 regulated genes in Wounding from PMID: 30772301 | SOX2 |
| 4495 | SHKBP1 | SOX2 regulated genes in Wounding from PMID: 30772301 | SOX2 |
| 4496 | FNBP1L | SOX2 regulated genes in Wounding from PMID: 30772301 | SOX2 |
| 4497 | BAZ1A | SOX2 regulated genes in Wounding from PMID: 30772301 | SOX2 |
| 4498 | KCTD3 | SOX2 regulated genes in Wounding from PMID: 30772301 | SOX2 |
| 4499 | SLC6A2 | SOX2 regulated genes in Wounding from PMID: 30772301 | SOX2 |
| 4500 | VGLL4 | SOX2 regulated genes in Wounding from PMID: 30772301 | SOX2 |
| 4501 | OST4 | SOX2 regulated genes in Wounding from PMID: 30772301 | SOX2 |
| 4502 | ARHGAP20 | SOX2 regulated genes in Wounding from PMID: 30772301 | SOX2 |
| 4503 | DNAJB1 | SOX2 regulated genes in Wounding from PMID: 30772301 | SOX2 |
| 4504 | EPHX4 | SOX2 regulated genes in Wounding from PMID: 30772301 | SOX2 |
| 4505 | SIKE1 | SOX2 regulated genes in Wounding from PMID: 30772301 | SOX2 |
| 4506 | CD9 | SOX2 regulated genes in Wounding from PMID: 30772301 | SOX2 |
| 4507 | DNAJC11 | SOX2 regulated genes in Wounding from PMID: 30772301 | SOX2 |
| 4508 | KRT40 | SOX2 regulated genes in Wounding from PMID: 30772301 | SOX2 |
| 4509 | CNOT9 | SOX2 regulated genes in Wounding from PMID: 30772301 | SOX2 |
| 4510 | MAP4 | SOX2 regulated genes in Wounding from PMID: 30772301 | SOX2 |
| 4511 | COG1 | SOX2 regulated genes in Wounding from PMID: 30772301 | SOX2 |
| 4512 | PTPN12 | SOX2 regulated genes in Wounding from PMID: 30772301 | SOX2 |

| | | | |
|---|---|---|---|
| 4513 | IL1F10 | SOX2 regulated genes in Wounding from PMID: 30772301 | SOX2 |
| 4514 | MAP7D1 | SOX2 regulated genes in Wounding from PMID: 30772301 | SOX2 |
| 4515 | SH3BP5L | SOX2 regulated genes in Wounding from PMID: 30772301 | SOX2 |
| 4516 | RHBDD2 | SOX2 regulated genes in Wounding from PMID: 30772301 | SOX2 |
| 4517 | EPHB2 | SOX2 regulated genes in Wounding from PMID: 30772301 | SOX2 |
| 4518 | DAPL1 | SOX2 regulated genes in Wounding from PMID: 30772301 | SOX2 |
| 4519 | ERBB2 | SOX2 regulated genes in Wounding from PMID: 30772301 | SOX2 |
| 4520 | SAMD4B | SOX2 regulated genes in Wounding from PMID: 30772301 | SOX2 |
| 4521 | LPCAT4 | SOX2 regulated genes in Wounding from PMID: 30772301 | SOX2 |
| 4522 | CDC14A | SOX2 regulated genes in Wounding from PMID: 30772301 | SOX2 |
| 4523 | WNT9A | SOX2 regulated genes in Wounding from PMID: 30772301 | SOX2 |
| 4524 | GTF2H3 | SOX2 regulated genes in Wounding from PMID: 30772301 | SOX2 |
| 4525 | HEATR3 | SOX2 regulated genes in Wounding from PMID: 30772301 | SOX2 |
| 4526 | PKP1 | SOX2 regulated genes in Wounding from PMID: 30772301 | SOX2 |
| 4527 | ANKRD13C | SOX2 regulated genes in Wounding from PMID: 30772301 | SOX2 |
| 4528 | ZC3H11A | SOX2 regulated genes in Wounding from PMID: 30772301 | SOX2 |
| 4529 | ZC3H11B | SOX2 regulated genes in Wounding from PMID: 30772301 | SOX2 |
| 4530 | LMBRD2 | SOX2 regulated genes in Wounding from PMID: 30772301 | SOX2 |
| 4531 | NECAB2 | SOX2 regulated genes in Wounding from PMID: 30772301 | SOX2 |
| 4532 | SFN | SOX2 regulated genes in Wounding from PMID: 30772301 | SOX2 |
| 4533 | GRHL1 | SOX2 regulated genes in Wounding from PMID: 30772301 | SOX2 |
| 4534 | MYBPC3 | SOX2 regulated genes in Wounding from PMID: 30772301 | SOX2 |
| 4535 | AMOTL2 | SOX2 regulated genes in Wounding from PMID: 30772301 | SOX2 |
| 4536 | PHLDA1 | SOX2 regulated genes in Wounding from PMID: 30772301 | SOX2 |
| 4537 | TRIM39 | SOX2 regulated genes in Wounding from PMID: 30772301 | SOX2 |
| 4538 | TUBA1C | SOX2 regulated genes in Wounding from PMID: 30772301 | SOX2 |
| 4539 | EML2 | SOX2 regulated genes in Wounding from PMID: 30772301 | SOX2 |
| 4540 | HNRNPH1 | SOX2 regulated genes in Wounding from PMID: 30772301 | SOX2 |
| 4541 | TRIM28 | SOX2 regulated genes in Wounding from PMID: 30772301 | SOX2 |
| 4542 | KRTAP3-1 | SOX2 regulated genes in Wounding from PMID: 30772301 | SOX2 |
| 4543 | KRT23 | SOX2 regulated genes in Wounding from PMID: 30772301 | SOX2 |
| 4544 | ERP44 | SOX2 regulated genes in Wounding from PMID: 30772301 | SOX2 |
| 4545 | F2RL1 | SOX2 regulated genes in Wounding from PMID: 30772301 | SOX2 |
| 4546 | SPRY4 | SOX2 regulated genes in Wounding from PMID: 30772301 | SOX2 |
| 4547 | DUOXA1 | SOX2 regulated genes in Wounding from PMID: 30772301 | SOX2 |
| 4548 | RAD17 | SOX2 regulated genes in Wounding from PMID: 30772301 | SOX2 |
| 4549 | STK11 | SOX2 regulated genes in Wounding from PMID: 30772301 | SOX2 |
| 4550 | USP6NL | SOX2 regulated genes in Wounding from PMID: 30772301 | SOX2 |
| 4551 | EPHB1 | SOX2 regulated genes in Wounding from PMID: 30772301 | SOX2 |
| 4552 | SNAP47 | SOX2 regulated genes in Wounding from PMID: 30772301 | SOX2 |
| 4553 | RBM14 | SOX2 regulated genes in Wounding from PMID: 30772301 | SOX2 |
| 4554 | MYT1L | SOX2 regulated genes in Wounding from PMID: 30772301 | SOX2 |
| 4555 | RABIF | SOX2 regulated genes in Wounding from PMID: 30772301 | SOX2 |
| 4556 | CDC42EP4 | SOX2 regulated genes in Wounding from PMID: 30772301 | SOX2 |
| 4557 | F2R | SOX2 regulated genes in Wounding from PMID: 30772301 | SOX2 |
| 4558 | CAMSAP3 | SOX2 regulated genes in Wounding from PMID: 30772301 | SOX2 |
| 4559 | KPRP | SOX2 regulated genes in Wounding from PMID: 30772301 | SOX2 |
| 4560 | THOC7 | SOX2 regulated genes in Wounding from PMID: 30772301 | SOX2 |
| 4561 | TRAPPC3 | SOX2 regulated genes in Wounding from PMID: 30772301 | SOX2 |
| 4562 | CDC25A | SOX2 regulated genes in Wounding from PMID: 30772301 | SOX2 |
| 4563 | OLFM2 | SOX2 regulated genes in Wounding from PMID: 30772301 | SOX2 |
| 4564 | WSB2 | SOX2 regulated genes in Wounding from PMID: 30772301 | SOX2 |
| 4565 | GPATCH4 | SOX2 regulated genes in Wounding from PMID: 30772301 | SOX2 |
| 4566 | PLK2 | SOX2 regulated genes in Wounding from PMID: 30772301 | SOX2 |
| 4567 | HMGXB3 | SOX2 regulated genes in Wounding from PMID: 30772301 | SOX2 |
| 4568 | PCDHB16 | SOX2 regulated genes in Wounding from PMID: 30772301 | SOX2 |
| 4569 | EIF4G1 | SOX2 regulated genes in Wounding from PMID: 30772301 | SOX2 |
| 4570 | VPS37B | SOX2 regulated genes in Wounding from PMID: 30772301 | SOX2 |
| 4571 | NPPC | SOX2 regulated genes in Wounding from PMID: 30772301 | SOX2 |
| 4572 | TAGLN2 | SOX2 regulated genes in Wounding from PMID: 30772301 | SOX2 |
| 4573 | P3H2 | SOX2 regulated genes in Wounding from PMID: 30772301 | SOX2 |

| | | | |
|---|---|---|---|
| 4574 | NPEPPS | SOX2 regulated genes in Wounding from PMID: 30772301 | SOX2 |
| 4575 | SEC22B | SOX2 regulated genes in Wounding from PMID: 30772301 | SOX2 |
| 4576 | SH3PXD2A | SOX2 regulated genes in Wounding from PMID: 30772301 | SOX2 |
| 4577 | KRT39 | SOX2 regulated genes in Wounding from PMID: 30772301 | SOX2 |
| 4578 | ALDH3B2 | SOX2 regulated genes in Wounding from PMID: 30772301 | SOX2 |
| 4579 | GRAMD2A | SOX2 regulated genes in Wounding from PMID: 30772301 | SOX2 |
| 4580 | PCBP1 | SOX2 regulated genes in Wounding from PMID: 30772301 | SOX2 |
| 4581 | HS2ST1 | SOX2 regulated genes in Wounding from PMID: 30772301 | SOX2 |
| 4582 | KRAS | SOX2 regulated genes in Wounding from PMID: 30772301 | SOX2 |
| 4583 | FAM83E | SOX2 regulated genes in Wounding from PMID: 30772301 | SOX2 |
| 4584 | PTHLH | SOX2 regulated genes in Wounding from PMID: 30772301 | SOX2 |
| 4585 | USP31 | SOX2 regulated genes in Wounding from PMID: 30772301 | SOX2 |
| 4586 | POLR3D | SOX2 regulated genes in Wounding from PMID: 30772301 | SOX2 |
| 4587 | S100A1 | SOX2 regulated genes in Wounding from PMID: 30772301 | SOX2 |
| 4588 | FUT1 | SOX2 regulated genes in Wounding from PMID: 30772301 | SOX2 |
| 4589 | NT5E | SOX2 regulated genes in Wounding from PMID: 30772301 | SOX2 |
| 4590 | SGK1 | SOX2 regulated genes in Wounding from PMID: 30772301 | SOX2 |
| 4591 | ZNF23 | SOX2 regulated genes in Wounding from PMID: 30772301 | SOX2 |
| 4592 | HMCN1 | SOX2 regulated genes in Wounding from PMID: 30772301 | SOX2 |
| 4593 | NECTIN2 | SOX2 regulated genes in Wounding from PMID: 30772301 | SOX2 |
| 4594 | EPN3 | SOX2 regulated genes in Wounding from PMID: 30772301 | SOX2 |
| 4595 | FOXO6 | SOX2 regulated genes in Wounding from PMID: 30772301 | SOX2 |
| 4596 | G3BP1 | SOX2 regulated genes in Wounding from PMID: 30772301 | SOX2 |
| 4597 | CAB39 | SOX2 regulated genes in Wounding from PMID: 30772301 | SOX2 |
| 4598 | PCBP4 | SOX2 regulated genes in Wounding from PMID: 30772301 | SOX2 |
| 4599 | THOC3 | SOX2 regulated genes in Wounding from PMID: 30772301 | SOX2 |
| 4600 | YAF2 | SOX2 regulated genes in Wounding from PMID: 30772301 | SOX2 |
| 4601 | GZMM | SOX2 regulated genes in Wounding from PMID: 30772301 | SOX2 |
| 4602 | CYP2S1 | SOX2 regulated genes in Wounding from PMID: 30772301 | SOX2 |
| 4603 | SUCO | SOX2 regulated genes in Wounding from PMID: 30772301 | SOX2 |
| 4604 | CAST | SOX2 regulated genes in Wounding from PMID: 30772301 | SOX2 |
| 4605 | RNF225 | SOX2 regulated genes in Wounding from PMID: 30772301 | SOX2 |
| 4606 | KLHL12 | SOX2 regulated genes in Wounding from PMID: 30772301 | SOX2 |
| 4607 | STON2 | SOX2 regulated genes in Wounding from PMID: 30772301 | SOX2 |
| 4608 | SLC20A1 | SOX2 regulated genes in Wounding from PMID: 30772301 | SOX2 |
| 4609 | SCNN1A | SOX2 regulated genes in Wounding from PMID: 30772301 | SOX2 |
| 4610 | NAPA | SOX2 regulated genes in Wounding from PMID: 30772301 | SOX2 |
| 4611 | TMEM107 | SOX2 regulated genes in Wounding from PMID: 30772301 | SOX2 |
| 4612 | FGF22 | SOX2 regulated genes in Wounding from PMID: 30772301 | SOX2 |
| 4613 | DUSP11 | SOX2 regulated genes in Wounding from PMID: 30772301 | SOX2 |
| 4614 | NOP58 | SOX2 regulated genes in Wounding from PMID: 30772301 | SOX2 |
| 4615 | MFN1 | SOX2 regulated genes in Wounding from PMID: 30772301 | SOX2 |
| 4616 | UCK2 | SOX2 regulated genes in Wounding from PMID: 30772301 | SOX2 |
| 4617 | RNF10 | SOX2 regulated genes in Wounding from PMID: 30772301 | SOX2 |
| 4618 | CLIP4 | SOX2 regulated genes in Wounding from PMID: 30772301 | SOX2 |
| 4619 | ZSWIM4 | SOX2 regulated genes in Wounding from PMID: 30772301 | SOX2 |
| 4620 | RAB10 | SOX2 regulated genes in Wounding from PMID: 30772301 | SOX2 |
| 4621 | KDM6B | SOX2 regulated genes in Wounding from PMID: 30772301 | SOX2 |
| 4622 | SPINK6 | SOX2 regulated genes in Wounding from PMID: 30772301 | SOX2 |
| 4623 | LEMD3 | SOX2 regulated genes in Wounding from PMID: 30772301 | SOX2 |
| 4624 | KDM1A | SOX2 regulated genes in Wounding from PMID: 30772301 | SOX2 |
| 4625 | TTYH2 | SOX2 regulated genes in Wounding from PMID: 30772301 | SOX2 |
| 4626 | ANAPC13 | SOX2 regulated genes in Wounding from PMID: 30772301 | SOX2 |
| 4627 | EPHB3 | SOX2 regulated genes in Wounding from PMID: 30772301 | SOX2 |
| 4628 | FBXO45 | SOX2 regulated genes in Wounding from PMID: 30772301 | SOX2 |
| 4629 | RHBG | SOX2 regulated genes in Wounding from PMID: 30772301 | SOX2 |
| 4630 | SEC13 | SOX2 regulated genes in Wounding from PMID: 30772301 | SOX2 |
| 4631 | MAPK6 | SOX2 regulated genes in Wounding from PMID: 30772301 | SOX2 |
| 4632 | ZNF750 | SOX2 regulated genes in Wounding from PMID: 30772301 | SOX2 |
| 4633 | CPS1 | SOX2 regulated genes in Wounding from PMID: 30772301 | SOX2 |
| 4634 | GOLGA7 | SOX2 regulated genes in Wounding from PMID: 30772301 | SOX2 |

| | | | |
|---|---|---|---|
| 4635 | CUL1 | SOX2 regulated genes in Wounding from PMID: 30772301 | SOX2 |
| 4636 | TRIM16 | SOX2 regulated genes in Wounding from PMID: 30772301 | SOX2 |
| 4637 | IRF6 | SOX2 regulated genes in Wounding from PMID: 30772301 | SOX2 |
| 4638 | LRRC20 | SOX2 regulated genes in Wounding from PMID: 30772301 | SOX2 |
| 4639 | RASSF1 | SOX2 regulated genes in Wounding from PMID: 30772301 | SOX2 |
| 4640 | INTS7 | SOX2 regulated genes in Wounding from PMID: 30772301 | SOX2 |
| 4641 | SPINT2 | SOX2 regulated genes in Wounding from PMID: 30772301 | SOX2 |
| 4642 | RYK | SOX2 regulated genes in Wounding from PMID: 30772301 | SOX2 |
| 4643 | SPTBN2 | SOX2 regulated genes in Wounding from PMID: 30772301 | SOX2 |
| 4644 | FPGT | SOX2 regulated genes in Wounding from PMID: 30772301 | SOX2 |
| 4645 | KRTAP5-8 | SOX2 regulated genes in Wounding from PMID: 30772301 | SOX2 |
| 4646 | NRAS | SOX2 regulated genes in Wounding from PMID: 30772301 | SOX2 |
| 4647 | WNT5A | SOX2 regulated genes in Wounding from PMID: 30772301 | SOX2 |
| 4648 | CDC42EP3 | SOX2 regulated genes in Wounding from PMID: 30772301 | SOX2 |
| 4649 | CUL3 | SOX2 regulated genes in Wounding from PMID: 30772301 | SOX2 |
| 4650 | CAMSAP2 | SOX2 regulated genes in Wounding from PMID: 30772301 | SOX2 |
| 4651 | ASCL4 | SOX2 regulated genes in Wounding from PMID: 30772301 | SOX2 |
| 4652 | NOTUM | SOX2 regulated genes in Wounding from PMID: 30772301 | SOX2 |
| 4653 | BBS5 | SOX2 regulated genes in Wounding from PMID: 30772301 | SOX2 |
| 4654 | GPA33 | SOX2 regulated genes in Wounding from PMID: 30772301 | SOX2 |
| 4655 | GNB1 | SOX2 regulated genes in Wounding from PMID: 30772301 | SOX2 |
| 4656 | CYP4F11 | SOX2 regulated genes in Wounding from PMID: 30772301 | SOX2 |
| 4657 | CYP4A11 | SOX2 regulated genes in Wounding from PMID: 30772301 | SOX2 |
| 4658 | MARVELD2 | SOX2 regulated genes in Wounding from PMID: 30772301 | SOX2 |
| 4659 | SDC1 | SOX2 regulated genes in Wounding from PMID: 30772301 | SOX2 |
| 4660 | KRT33A | SOX2 regulated genes in Wounding from PMID: 30772301 | SOX2 |
| 4661 | BMPR2 | SOX2 regulated genes in Wounding from PMID: 30772301 | SOX2 |
| 4662 | PATJ | SOX2 regulated genes in Wounding from PMID: 30772301 | SOX2 |
| 4663 | ARF1 | SOX2 regulated genes in Wounding from PMID: 30772301 | SOX2 |
| 4664 | TCF3 | SOX2 regulated genes in Wounding from PMID: 30772301 | SOX2 |
| 4665 | CORO1C | SOX2 regulated genes in Wounding from PMID: 30772301 | SOX2 |
| 4666 | EZR | SOX2 regulated genes in Wounding from PMID: 30772301 | SOX2 |
| 4667 | APPBP2 | SOX2 regulated genes in Wounding from PMID: 30772301 | SOX2 |
| 4668 | RBM12B | SOX2 regulated genes in Wounding from PMID: 30772301 | SOX2 |
| 4669 | RASA2 | SOX2 regulated genes in Wounding from PMID: 30772301 | SOX2 |
| 4670 | ATP1A1 | SOX2 regulated genes in Wounding from PMID: 30772301 | SOX2 |
| 4671 | DUOX1 | SOX2 regulated genes in Wounding from PMID: 30772301 | SOX2 |
| 4672 | LSM12 | SOX2 regulated genes in Wounding from PMID: 30772301 | SOX2 |
| 4673 | GPX2 | SOX2 regulated genes in Wounding from PMID: 30772301 | SOX2 |
| 4674 | GPR87 | SOX2 regulated genes in Wounding from PMID: 30772301 | SOX2 |
| 4675 | KRTAP3-3 | SOX2 regulated genes in Wounding from PMID: 30772301 | SOX2 |
| 4676 | KRTAP3-2 | SOX2 regulated genes in Wounding from PMID: 30772301 | SOX2 |
| 4677 | DUSP7 | SOX2 regulated genes in Wounding from PMID: 30772301 | SOX2 |
| 4678 | APLP1 | SOX2 regulated genes in Wounding from PMID: 30772301 | SOX2 |
| 4679 | GOLIM4 | SOX2 regulated genes in Wounding from PMID: 30772301 | SOX2 |
| 4680 | CTNND1 | SOX2 regulated genes in Wounding from PMID: 30772301 | SOX2 |
| 4681 | PLPPR2 | SOX2 regulated genes in Wounding from PMID: 30772301 | SOX2 |
| 4682 | FAF1 | SOX2 regulated genes in Wounding from PMID: 30772301 | SOX2 |
| 4683 | IFT20 | SOX2 regulated genes in Wounding from PMID: 30772301 | SOX2 |
| 4684 | LAMC2 | SOX2 regulated genes in Wounding from PMID: 30772301 | SOX2 |
| 4685 | ZNF7 | SOX2 regulated genes in Wounding from PMID: 30772301 | SOX2 |
| 4686 | FOXP4 | SOX2 regulated genes in Wounding from PMID: 30772301 | SOX2 |
| 4687 | RNF145 | SOX2 regulated genes in Wounding from PMID: 30772301 | SOX2 |
| 4688 | ITPKB | SOX2 regulated genes in Wounding from PMID: 30772301 | SOX2 |
| 4689 | ACVR1B | SOX2 regulated genes in Wounding from PMID: 30772301 | SOX2 |
| 4690 | IGSF3 | SOX2 regulated genes in Wounding from PMID: 30772301 | SOX2 |
| 4691 | PIK3R3 | SOX2 regulated genes in Wounding from PMID: 30772301 | SOX2 |
| 4692 | CBX2 | SOX2 regulated genes in Wounding from PMID: 30772301 | SOX2 |
| 4693 | USP7 | SOX2 regulated genes in Wounding from PMID: 30772301 | SOX2 |
| 4694 | KMT5A | SOX2 regulated genes in Wounding from PMID: 30772301 | SOX2 |
| 4695 | MMADHC | SOX2 regulated genes in Wounding from PMID: 30772301 | SOX2 |

| | | | |
|---|---|---|---|
| 4696 | RRP15 | SOX2 regulated genes in Wounding from PMID: 30772301 | SOX2 |
| 4697 | PRMT7 | SOX2 regulated genes in Wounding from PMID: 30772301 | SOX2 |
| 4698 | TSR1 | SOX2 regulated genes in Wounding from PMID: 30772301 | SOX2 |
| 4699 | HDAC1 | SOX2 regulated genes in Wounding from PMID: 30772301 | SOX2 |
| 4700 | OTOP2 | SOX2 regulated genes in Wounding from PMID: 30772301 | SOX2 |
| 4701 | EVPL | SOX2 regulated genes in Wounding from PMID: 30772301 | SOX2 |
| 4702 | IST1 | SOX2 regulated genes in Wounding from PMID: 30772301 | SOX2 |
| 4703 | ATP2A2 | SOX2 regulated genes in Wounding from PMID: 30772301 | SOX2 |
| 4704 | CITED4 | SOX2 regulated genes in Wounding from PMID: 30772301 | SOX2 |
| 4705 | ERBB3 | SOX2 regulated genes in Wounding from PMID: 30772301 | SOX2 |
| 4706 | CCDC92 | SOX2 regulated genes in Wounding from PMID: 30772301 | SOX2 |
| 4707 | KIFC2 | SOX2 regulated genes in Wounding from PMID: 30772301 | SOX2 |
| 4708 | ACTG1 | SOX2 regulated genes in Wounding from PMID: 30772301 | SOX2 |
| 4709 | CFTR | SOX2 regulated genes in Wounding from PMID: 30772301 | SOX2 |
| 4710 | TRIM59 | SOX2 regulated genes in Wounding from PMID: 30772301 | SOX2 |
| 4711 | AGBL5 | SOX2 regulated genes in Wounding from PMID: 30772301 | SOX2 |
| 4712 | B4GALT3 | SOX2 regulated genes in Wounding from PMID: 30772301 | SOX2 |
| 4713 | MAP4K4 | SOX2 regulated genes in Wounding from PMID: 30772301 | SOX2 |
| 4714 | TMEM65 | SOX2 regulated genes in Wounding from PMID: 30772301 | SOX2 |
| 4715 | RNF39 | SOX2 regulated genes in Wounding from PMID: 30772301 | SOX2 |
| 4716 | C6orf141 | SOX2 regulated genes in Wounding from PMID: 30772301 | SOX2 |
| 4717 | CTPS1 | SOX2 regulated genes in Wounding from PMID: 30772301 | SOX2 |
| 4718 | ATP2C2 | SOX2 regulated genes in Wounding from PMID: 30772301 | SOX2 |
| 4719 | ADGRL2 | SOX2 regulated genes in Wounding from PMID: 30772301 | SOX2 |
| 4720 | WDR75 | SOX2 regulated genes in Wounding from PMID: 30772301 | SOX2 |
| 4721 | LRP12 | SOX2 regulated genes in Wounding from PMID: 30772301 | SOX2 |
| 4722 | TMEM120A | SOX2 regulated genes in Wounding from PMID: 30772301 | SOX2 |
| 4723 | FER | SOX2 regulated genes in Wounding from PMID: 30772301 | SOX2 |
| 4724 | HSPA4 | SOX2 regulated genes in Wounding from PMID: 30772301 | SOX2 |
| 4725 | PTPN9 | SOX2 regulated genes in Wounding from PMID: 30772301 | SOX2 |
| 4726 | DVL3 | SOX2 regulated genes in Wounding from PMID: 30772301 | SOX2 |
| 4727 | FZD5 | SOX2 regulated genes in Wounding from PMID: 30772301 | SOX2 |
| 4728 | USP53 | SOX2 regulated genes in Wounding from PMID: 30772301 | SOX2 |
| 4729 | CHD1 | SOX2 regulated genes in Wounding from PMID: 30772301 | SOX2 |
| 4730 | KLF10 | SOX2 regulated genes in Wounding from PMID: 30772301 | SOX2 |
| 4731 | IGSF8 | SOX2 regulated genes in Wounding from PMID: 30772301 | SOX2 |
| 4732 | SYT8 | SOX2 regulated genes in Wounding from PMID: 30772301 | SOX2 |
| 4733 | PTBP1 | SOX2 regulated genes in Wounding from PMID: 30772301 | SOX2 |
| 4734 | WWC1 | SOX2 regulated genes in Wounding from PMID: 30772301 | SOX2 |
| 4735 | PITHD1 | SOX2 regulated genes in Wounding from PMID: 30772301 | SOX2 |
| 4736 | RBMXL1 | SOX2 regulated genes in Wounding from PMID: 30772301 | SOX2 |
| 4737 | RASGEF1B | SOX2 regulated genes in Wounding from PMID: 30772301 | SOX2 |
| 4738 | RNF25 | SOX2 regulated genes in Wounding from PMID: 30772301 | SOX2 |
| 4739 | PADI3 | SOX2 regulated genes in Wounding from PMID: 30772301 | SOX2 |
| 4740 | CSRP1 | SOX2 regulated genes in Wounding from PMID: 30772301 | SOX2 |
| 4741 | AGRN | SOX2 regulated genes in Wounding from PMID: 30772301 | SOX2 |
| 4742 | TMEM229B | SOX2 regulated genes in Wounding from PMID: 30772301 | SOX2 |
| 4743 | CHD4 | SOX2 regulated genes in Wounding from PMID: 30772301 | SOX2 |
| 4744 | CYTH2 | SOX2 regulated genes in Wounding from PMID: 30772301 | SOX2 |
| 4745 | SLC30A3 | SOX2 regulated genes in Wounding from PMID: 30772301 | SOX2 |
| 4746 | EPHA4 | SOX2 regulated genes in Wounding from PMID: 30772301 | SOX2 |
| 4747 | CRIP3 | SOX2 regulated genes in Wounding from PMID: 30772301 | SOX2 |
| 4748 | ACTR1B | SOX2 regulated genes in Wounding from PMID: 30772301 | SOX2 |
| 4749 | LY6G6D | SOX2 regulated genes in Wounding from PMID: 30772301 | SOX2 |
| 4750 | NGF | SOX2 regulated genes in Wounding from PMID: 30772301 | SOX2 |
| 4751 | ZRANB2 | SOX2 regulated genes in Wounding from PMID: 30772301 | SOX2 |
| 4752 | SSTR2 | SOX2 regulated genes in Wounding from PMID: 30772301 | SOX2 |
| 4753 | GALE | SOX2 regulated genes in Wounding from PMID: 30772301 | SOX2 |
| 4754 | TMEM68 | SOX2 regulated genes in Wounding from PMID: 30772301 | SOX2 |
| 4755 | GATA3 | SOX2 regulated genes in Wounding from PMID: 30772301 | SOX2 |
| 4756 | SPTY2D1 | SOX2 regulated genes in Wounding from PMID: 30772301 | SOX2 |

| | | | |
|---|---|---|---|
| 4757 | ZNF605 | SOX2 regulated genes in Wounding from PMID: 30772301 | SOX2 |
| 4758 | PRPF40B | SOX2 regulated genes in Wounding from PMID: 30772301 | SOX2 |
| 4759 | KLHL29 | SOX2 regulated genes in Wounding from PMID: 30772301 | SOX2 |
| 4760 | TUBA1A | SOX2 regulated genes in Wounding from PMID: 30772301 | SOX2 |
| 4761 | TAF7 | SOX2 regulated genes in Wounding from PMID: 30772301 | SOX2 |
| 4762 | CEP83 | SOX2 regulated genes in Wounding from PMID: 30772301 | SOX2 |
| 4763 | LLGL2 | SOX2 regulated genes in Wounding from PMID: 30772301 | SOX2 |
| 4764 | KRT79 | SOX2 regulated genes in Wounding from PMID: 30772301 | SOX2 |
| 4765 | MAP6 | SOX2 regulated genes in Wounding from PMID: 30772301 | SOX2 |
| 4766 | PLK3 | SOX2 regulated genes in Wounding from PMID: 30772301 | SOX2 |
| 4767 | ACSL3 | SOX2 regulated genes in Wounding from PMID: 30772301 | SOX2 |
| 4768 | EGR2 | SOX2 regulated genes in Wounding from PMID: 30772301 | SOX2 |
| 4769 | PALMD | SOX2 regulated genes in Wounding from PMID: 30772301 | SOX2 |
| 4770 | KIF21B | SOX2 regulated genes in Wounding from PMID: 30772301 | SOX2 |
| 4771 | TRNP1 | SOX2 regulated genes in Wounding from PMID: 30772301 | SOX2 |
| 4772 | NUDCD1 | SOX2 regulated genes in Wounding from PMID: 30772301 | SOX2 |
| 4773 | DYNC1I2 | SOX2 regulated genes in Wounding from PMID: 30772301 | SOX2 |
| 4774 | ANGPTL4 | SOX2 regulated genes in Wounding from PMID: 30772301 | SOX2 |
| 4775 | POLB | SOX2 regulated genes in Wounding from PMID: 30772301 | SOX2 |
| 4776 | TMEM139 | SOX2 regulated genes in Wounding from PMID: 30772301 | SOX2 |
| 4777 | ABCC1 | SOX2 regulated genes in Wounding from PMID: 30772301 | SOX2 |
| 4778 | KLHDC8A | SOX2 regulated genes in Wounding from PMID: 30772301 | SOX2 |
| 4779 | HMGCS2 | SOX2 regulated genes in Wounding from PMID: 30772301 | SOX2 |
| 4780 | OTUD7B | SOX2 regulated genes in Wounding from PMID: 30772301 | SOX2 |
| 4781 | LINS1 | SOX2 regulated genes in Wounding from PMID: 30772301 | SOX2 |
| 4782 | FLG2 | SOX2 regulated genes in Wounding from PMID: 30772301 | SOX2 |
| 4783 | CCDC106 | SOX2 regulated genes in Wounding from PMID: 30772301 | SOX2 |
| 4784 | ENC1 | SOX2 regulated genes in Wounding from PMID: 30772301 | SOX2 |
| 4785 | TPPP3 | SOX2 regulated genes in Wounding from PMID: 30772301 | SOX2 |
| 4786 | SYTL2 | SOX2 regulated genes in Wounding from PMID: 30772301 | SOX2 |
| 4787 | HRAS | SOX2 regulated genes in Wounding from PMID: 30772301 | SOX2 |
| 4788 | CCT7 | SOX2 regulated genes in Wounding from PMID: 30772301 | SOX2 |
| 4789 | BAHCC1 | SOX2 regulated genes in Wounding from PMID: 30772301 | SOX2 |
| 4790 | TUFT1 | SOX2 regulated genes in Wounding from PMID: 30772301 | SOX2 |
| 4791 | SHOC2 | SOX2 regulated genes in Wounding from PMID: 30772301 | SOX2 |
| 4792 | ARL13B | SOX2 regulated genes in Wounding from PMID: 30772301 | SOX2 |
| 4793 | SLC4A9 | SOX2 regulated genes in Wounding from PMID: 30772301 | SOX2 |
| 4794 | LYG2 | SOX2 regulated genes in Wounding from PMID: 30772301 | SOX2 |
| 4795 | IRGQ | SOX2 regulated genes in Wounding from PMID: 30772301 | SOX2 |
| 4796 | FXYD3 | SOX2 regulated genes in Wounding from PMID: 30772301 | SOX2 |
| 4797 | MIDN | SOX2 regulated genes in Wounding from PMID: 30772301 | SOX2 |
| 4798 | PTGS2 | SOX2 regulated genes in Wounding from PMID: 30772301 | SOX2 |
| 4799 | FSTL3 | SOX2 regulated genes in Wounding from PMID: 30772301 | SOX2 |
| 4800 | RNF223 | SOX2 regulated genes in Wounding from PMID: 30772301 | SOX2 |
| 4801 | GTF2H2C | SOX2 regulated genes in Wounding from PMID: 30772301 | SOX2 |
| 4802 | BMPR1A | SOX2 regulated genes in Wounding from PMID: 30772301 | SOX2 |
| 4803 | SETD5 | SOX2 regulated genes in Wounding from PMID: 30772301 | SOX2 |
| 4804 | S100A11 | SOX2 regulated genes in Wounding from PMID: 30772301 | SOX2 |
| 4805 | NT5C3A | SOX2 regulated genes in Wounding from PMID: 30772301 | SOX2 |
| 4806 | RAB7B | SOX2 regulated genes in Wounding from PMID: 30772301 | SOX2 |
| 4807 | HOOK2 | SOX2 regulated genes in Wounding from PMID: 30772301 | SOX2 |
| 4808 | TBC1D30 | SOX2 regulated genes in Wounding from PMID: 30772301 | SOX2 |
| 4809 | ZBTB17 | SOX2 regulated genes in Wounding from PMID: 30772301 | SOX2 |
| 4810 | ANGPT2 | SOX2 regulated genes in Wounding from PMID: 30772301 | SOX2 |
| 4811 | EBNA1BP2 | SOX2 regulated genes in Wounding from PMID: 30772301 | SOX2 |
| 4812 | CNOT2 | SOX2 regulated genes in Wounding from PMID: 30772301 | SOX2 |
| 4813 | PLXNB1 | SOX2 regulated genes in Wounding from PMID: 30772301 | SOX2 |
| 4814 | GALR2 | SOX2 regulated genes in Wounding from PMID: 30772301 | SOX2 |
| 4815 | EIF3B | SOX2 regulated genes in Wounding from PMID: 30772301 | SOX2 |
| 4816 | MAGI1 | SOX2 regulated genes in Wounding from PMID: 30772301 | SOX2 |
| 4817 | ANXA9 | SOX2 regulated genes in Wounding from PMID: 30772301 | SOX2 |

| | | | |
|---|---|---|---|
| 4818 | SPRR4 | SOX2 regulated genes in Wounding from PMID: 30772301 | SOX2 |
| 4819 | ERH | SOX2 regulated genes in Wounding from PMID: 30772301 | SOX2 |
| 4820 | HR | SOX2 regulated genes in Wounding from PMID: 30772301 | SOX2 |
| 4821 | CCNYL1 | SOX2 regulated genes in Wounding from PMID: 30772301 | SOX2 |
| 4822 | FAM89A | SOX2 regulated genes in Wounding from PMID: 30772301 | SOX2 |
| 4823 | KRTAP4-6 | SOX2 regulated genes in Wounding from PMID: 30772301 | SOX2 |
| 4824 | TCHH | SOX2 regulated genes in Wounding from PMID: 30772301 | SOX2 |
| 4825 | FAM221A | SOX2 regulated genes in Wounding from PMID: 30772301 | SOX2 |
| 4826 | ENPP1 | SOX2 regulated genes in Wounding from PMID: 30772301 | SOX2 |
| 4827 | KPNB1 | SOX2 regulated genes in Wounding from PMID: 30772301 | SOX2 |
| 4828 | MFF | SOX2 regulated genes in Wounding from PMID: 30772301 | SOX2 |
| 4829 | CDC42SE1 | SOX2 regulated genes in Wounding from PMID: 30772301 | SOX2 |
| 4830 | ALYREF | SOX2 regulated genes in Wounding from PMID: 30772301 | SOX2 |
| 4831 | RND3 | SOX2 regulated genes in Wounding from PMID: 30772301 | SOX2 |
| 4832 | TANC2 | SOX2 regulated genes in Wounding from PMID: 30772301 | SOX2 |
| 4833 | HDLBP | SOX2 regulated genes in Wounding from PMID: 30772301 | SOX2 |
| 4834 | C17orf80 | SOX2 regulated genes in Wounding from PMID: 30772301 | SOX2 |
| 4835 | CAP1 | SOX2 regulated genes in Wounding from PMID: 30772301 | SOX2 |
| 4836 | ATP6V0E1 | SOX2 regulated genes in Wounding from PMID: 30772301 | SOX2 |
| 4837 | PGM1 | SOX2 regulated genes in Wounding from PMID: 30772301 | SOX2 |
| 4838 | MAPRE3 | SOX2 regulated genes in Wounding from PMID: 30772301 | SOX2 |
| 4839 | C3orf52 | SOX2 regulated genes in Wounding from PMID: 30772301 | SOX2 |
| 4840 | TXNRD3 | SOX2 regulated genes in Wounding from PMID: 30772301 | SOX2 |
| 4841 | WIZ | SOX2 regulated genes in Wounding from PMID: 30772301 | SOX2 |
| 4842 | YWHAE | SOX2 regulated genes in Wounding from PMID: 30772301 | SOX2 |
| 4843 | CUL2 | SOX2 regulated genes in Wounding from PMID: 30772301 | SOX2 |
| 4844 | SMYD2 | SOX2 regulated genes in Wounding from PMID: 30772301 | SOX2 |
| 4845 | TMEM181 | SOX2 regulated genes in Wounding from PMID: 30772301 | SOX2 |
| 4846 | PDGFA | SOX2 regulated genes in Wounding from PMID: 30772301 | SOX2 |
| 4847 | CDH1 | SOX2 regulated genes in Wounding from PMID: 30772301 | SOX2 |
| 4848 | E2F3 | SOX2 regulated genes in Wounding from PMID: 30772301 | SOX2 |
| 4849 | SPECC1 | SOX2 regulated genes in Wounding from PMID: 30772301 | SOX2 |
| 4850 | GMPPB | SOX2 regulated genes in Wounding from PMID: 30772301 | SOX2 |
| 4851 | TATDN2 | SOX2 regulated genes in Wounding from PMID: 30772301 | SOX2 |
| 4852 | SEC22A | SOX2 regulated genes in Wounding from PMID: 30772301 | SOX2 |
| 4853 | PTGES3 | SOX2 regulated genes in Wounding from PMID: 30772301 | SOX2 |
| 4854 | CCT3 | SOX2 regulated genes in Wounding from PMID: 30772301 | SOX2 |
| 4855 | RAD23A | SOX2 regulated genes in Wounding from PMID: 30772301 | SOX2 |
| 4856 | LYPD6B | SOX2 regulated genes in Wounding from PMID: 30772301 | SOX2 |
| 4857 | DCLK3 | SOX2 regulated genes in Wounding from PMID: 30772301 | SOX2 |
| 4858 | GPRC5A | SOX2 regulated genes in Wounding from PMID: 30772301 | SOX2 |
| 4859 | PIK3CB | SOX2 regulated genes in Wounding from PMID: 30772301 | SOX2 |
| 4860 | BEND5 | SOX2 regulated genes in Wounding from PMID: 30772301 | SOX2 |
| 4861 | DPP9 | SOX2 regulated genes in Wounding from PMID: 30772301 | SOX2 |
| 4862 | SLC23A3 | SOX2 regulated genes in Wounding from PMID: 30772301 | SOX2 |
| 4863 | PAFAH1B3 | SOX2 regulated genes in Wounding from PMID: 30772301 | SOX2 |
| 4864 | TRIM5 | SOX2 regulated genes in Wounding from PMID: 30772301 | SOX2 |
| 4865 | PIP5K1A | SOX2 regulated genes in Wounding from PMID: 30772301 | SOX2 |
| 4866 | RECQL | SOX2 regulated genes in Wounding from PMID: 30772301 | SOX2 |
| 4867 | TNFRSF10B | SOX2 regulated genes in Wounding from PMID: 30772301 | SOX2 |
| 4868 | TNFRSF10C | SOX2 regulated genes in Wounding from PMID: 30772301 | SOX2 |
| 4869 | TNFRSF10D | SOX2 regulated genes in Wounding from PMID: 30772301 | SOX2 |
| 4870 | YME1L1 | SOX2 regulated genes in Wounding from PMID: 30772301 | SOX2 |
| 4871 | HBEGF | SOX2 regulated genes in Wounding from PMID: 30772301 | SOX2 |
| 4872 | RALGPS2 | SOX2 regulated genes in Wounding from PMID: 30772301 | SOX2 |
| 4873 | PDCD4 | SOX2 regulated genes in Wounding from PMID: 30772301 | SOX2 |
| 4874 | CNST | SOX2 regulated genes in Wounding from PMID: 30772301 | SOX2 |
| 4875 | HNRNPU | SOX2 regulated genes in Wounding from PMID: 30772301 | SOX2 |
| 4876 | ZNF746 | SOX2 regulated genes in Wounding from PMID: 30772301 | SOX2 |
| 4877 | TMEM69 | SOX2 regulated genes in Wounding from PMID: 30772301 | SOX2 |
| 4878 | WDR47 | SOX2 regulated genes in Wounding from PMID: 30772301 | SOX2 |

| | | | |
|---|---|---|---|
| 4879 | TUBA4A | SOX2 regulated genes in Wounding from PMID: 30772301 | SOX2 |
| 4880 | SYNRG | SOX2 regulated genes in Wounding from PMID: 30772301 | SOX2 |
| 4881 | HECTD1 | SOX2 regulated genes in Wounding from PMID: 30772301 | SOX2 |
| 4882 | FADS2 | SOX2 regulated genes in Wounding from PMID: 30772301 | SOX2 |
| 4883 | QRICH2 | SOX2 regulated genes in Wounding from PMID: 30772301 | SOX2 |
| 4884 | IWS1 | SOX2 regulated genes in Wounding from PMID: 30772301 | SOX2 |
| 4885 | SAXO2 | SOX2 regulated genes in Wounding from PMID: 30772301 | SOX2 |
| 4886 | IL11 | SOX2 regulated genes in Wounding from PMID: 30772301 | SOX2 |
| 4887 | DVL1 | SOX2 regulated genes in Wounding from PMID: 30772301 | SOX2 |
| 4888 | BHLHE41 | SOX2 regulated genes in Wounding from PMID: 30772301 | SOX2 |
| 4889 | ATP6V0A2 | SOX2 regulated genes in Wounding from PMID: 30772301 | SOX2 |
| 4890 | SPINT1 | SOX2 regulated genes in Wounding from PMID: 30772301 | SOX2 |
| 4891 | GCLM | SOX2 regulated genes in Wounding from PMID: 30772301 | SOX2 |
| 4892 | CNTN5 | SOX2 regulated genes in Wounding from PMID: 30772301 | SOX2 |
| 4893 | BZW2 | SOX2 regulated genes in Wounding from PMID: 30772301 | SOX2 |
| 4894 | ST7L | SOX2 regulated genes in Wounding from PMID: 30772301 | SOX2 |
| 4895 | ENOPH1 | SOX2 regulated genes in Wounding from PMID: 30772301 | SOX2 |
| 4896 | STX6 | SOX2 regulated genes in Wounding from PMID: 30772301 | SOX2 |
| 4897 | FAT1 | SOX2 regulated genes in Wounding from PMID: 30772301 | SOX2 |
| 4898 | SDR16C5 | SOX2 regulated genes in Wounding from PMID: 30772301 | SOX2 |
| 4899 | ZNF266 | SOX2 regulated genes in Wounding from PMID: 30772301 | SOX2 |
| 4900 | POLR2D | SOX2 regulated genes in Wounding from PMID: 30772301 | SOX2 |
| 4901 | CCDC91 | SOX2 regulated genes in Wounding from PMID: 30772301 | SOX2 |
| 4902 | PDPN | SOX2 regulated genes in Wounding from PMID: 30772301 | SOX2 |
| 4903 | VANGL1 | SOX2 regulated genes in Wounding from PMID: 30772301 | SOX2 |
| 4904 | KCNK12 | SOX2 regulated genes in Wounding from PMID: 30772301 | SOX2 |
| 4905 | ITGA6 | SOX2 regulated genes in Wounding from PMID: 30772301 | SOX2 |
| 4906 | GPR42 | SOX2 regulated genes in Wounding from PMID: 30772301 | SOX2 |
| 4907 | FILIP1L | SOX2 regulated genes in Wounding from PMID: 30772301 | SOX2 |
| 4908 | NDUFAF7 | SOX2 regulated genes in Wounding from PMID: 30772301 | SOX2 |
| 4909 | MPP7 | SOX2 regulated genes in Wounding from PMID: 30772301 | SOX2 |
| 4910 | MYH14 | SOX2 regulated genes in Wounding from PMID: 30772301 | SOX2 |
| 4911 | GTF2F1 | SOX2 regulated genes in Wounding from PMID: 30772301 | SOX2 |
| 4912 | DHRS9 | SOX2 regulated genes in Wounding from PMID: 30772301 | SOX2 |
| 4913 | MARK1 | SOX2 regulated genes in Wounding from PMID: 30772301 | SOX2 |
| 4914 | C12orf56 | SOX2 regulated genes in Wounding from PMID: 30772301 | SOX2 |
| 4915 | BNC1 | SOX2 regulated genes in Wounding from PMID: 30772301 | SOX2 |
| 4916 | POGK | SOX2 regulated genes in Wounding from PMID: 30772301 | SOX2 |
| 4917 | P4HB | SOX2 regulated genes in Wounding from PMID: 30772301 | SOX2 |
| 4918 | B3GNT8 | SOX2 regulated genes in Wounding from PMID: 30772301 | SOX2 |
| 4919 | METTL23 | SOX2 regulated genes in Wounding from PMID: 30772301 | SOX2 |
| 4920 | TMED5 | SOX2 regulated genes in Wounding from PMID: 30772301 | SOX2 |
| 4921 | SLC4A1 | SOX2 regulated genes in Wounding from PMID: 30772301 | SOX2 |
| 4922 | PKP2 | SOX2 regulated genes in Wounding from PMID: 30772301 | SOX2 |
| 4923 | GPSM2 | SOX2 regulated genes in Wounding from PMID: 30772301 | SOX2 |
| 4924 | ITGB4 | SOX2 regulated genes in Wounding from PMID: 30772301 | SOX2 |
| 4925 | CLASP1 | SOX2 regulated genes in Wounding from PMID: 30772301 | SOX2 |
| 4926 | SIRT1 | SOX2 regulated genes in Wounding from PMID: 30772301 | SOX2 |
| 4927 | CCHCR1 | SOX2 regulated genes in Wounding from PMID: 30772301 | SOX2 |
| 4928 | TSG101 | SOX2 regulated genes in Wounding from PMID: 30772301 | SOX2 |
| 4929 | MACF1 | SOX2 regulated genes in Wounding from PMID: 30772301 | SOX2 |
| 4930 | TRAF4 | SOX2 regulated genes in Wounding from PMID: 30772301 | SOX2 |
| 4931 | TXNRD1 | SOX2 regulated genes in Wounding from PMID: 30772301 | SOX2 |
| 4932 | ARHGAP22 | SOX2 regulated genes in Wounding from PMID: 30772301 | SOX2 |
| 4933 | INTS14 | SOX2 regulated genes in Wounding from PMID: 30772301 | SOX2 |
| 4934 | CLK3 | SOX2 regulated genes in Wounding from PMID: 30772301 | SOX2 |
| 4935 | EGR1 | SOX2 regulated genes in Wounding from PMID: 30772301 | SOX2 |
| 4936 | ECD | SOX2 regulated genes in Wounding from PMID: 30772301 | SOX2 |
| 4937 | MYH15 | SOX2 regulated genes in Wounding from PMID: 30772301 | SOX2 |
| 4938 | NSUN2 | SOX2 regulated genes in Wounding from PMID: 30772301 | SOX2 |
| 4939 | GEMIN4 | SOX2 regulated genes in Wounding from PMID: 30772301 | SOX2 |

| | | | |
|---|---|---|---|
| 4940 | LAD1 | SOX2 regulated genes in Wounding from PMID: 30772301 | SOX2 |
| 4941 | RBBP6 | SOX2 regulated genes in Wounding from PMID: 30772301 | SOX2 |
| 4942 | ATXN7L3 | SOX2 regulated genes in Wounding from PMID: 30772301 | SOX2 |
| 4943 | PPP2CB | SOX2 regulated genes in Wounding from PMID: 30772301 | SOX2 |
| 4944 | SPPL3 | SOX2 regulated genes in Wounding from PMID: 30772301 | SOX2 |
| 4945 | SNAPC3 | SOX2 regulated genes in Wounding from PMID: 30772301 | SOX2 |
| 4946 | FBXO46 | SOX2 regulated genes in Wounding from PMID: 30772301 | SOX2 |
| 4947 | PDXDC1 | SOX2 regulated genes in Wounding from PMID: 30772301 | SOX2 |
| 4948 | HGS | SOX2 regulated genes in Wounding from PMID: 30772301 | SOX2 |
| 4949 | STEAP1 | SOX2 regulated genes in Wounding from PMID: 30772301 | SOX2 |
| 4950 | KCTD5 | SOX2 regulated genes in Wounding from PMID: 30772301 | SOX2 |
| 4951 | PRSS22 | SOX2 regulated genes in Wounding from PMID: 30772301 | SOX2 |
| 4952 | PLXNA1 | SOX2 regulated genes in Wounding from PMID: 30772301 | SOX2 |
| 4953 | STRIP1 | SOX2 regulated genes in Wounding from PMID: 30772301 | SOX2 |
| 4954 | DLG1 | SOX2 regulated genes in Wounding from PMID: 30772301 | SOX2 |
| 4955 | TP63 | SOX2 regulated genes in Wounding from PMID: 30772301 | SOX2 |
| 4956 | USP43 | SOX2 regulated genes in Wounding from PMID: 30772301 | SOX2 |
| 4957 | MGST3 | SOX2 regulated genes in Wounding from PMID: 30772301 | SOX2 |
| 4958 | ANKRD13B | SOX2 regulated genes in Wounding from PMID: 30772301 | SOX2 |
| 4959 | MPZL1 | SOX2 regulated genes in Wounding from PMID: 30772301 | SOX2 |
| 4960 | SMURF2 | SOX2 regulated genes in Wounding from PMID: 30772301 | SOX2 |
| 4961 | PRPF3 | SOX2 regulated genes in Wounding from PMID: 30772301 | SOX2 |
| 4962 | RBM45 | SOX2 regulated genes in Wounding from PMID: 30772301 | SOX2 |
| 4963 | SLC30A1 | SOX2 regulated genes in Wounding from PMID: 30772301 | SOX2 |
| 4964 | SHH | SOX2 regulated genes in Wounding from PMID: 30772301 | SOX2 |
| 4965 | SLC7A8 | SOX2 regulated genes in Wounding from PMID: 30772301 | SOX2 |
| 4966 | PHACTR1 | SOX2 regulated genes in Wounding from PMID: 30772301 | SOX2 |
| 4967 | BPNT1 | SOX2 regulated genes in Wounding from PMID: 30772301 | SOX2 |
| 4968 | PCDHGB1 | SOX2 regulated genes in Wounding from PMID: 30772301 | SOX2 |
| 4969 | PSMD7 | SOX2 regulated genes in Wounding from PMID: 30772301 | SOX2 |
| 4970 | PELI1 | SOX2 regulated genes in Wounding from PMID: 30772301 | SOX2 |
| 4971 | ZNF800 | SOX2 regulated genes in Wounding from PMID: 30772301 | SOX2 |
| 4972 | FCAMR | SOX2 regulated genes in Wounding from PMID: 30772301 | SOX2 |
| 4973 | DDX1 | SOX2 regulated genes in Wounding from PMID: 30772301 | SOX2 |
| 4974 | PPIP5K1 | SOX2 regulated genes in Wounding from PMID: 30772301 | SOX2 |
| 4975 | RHOU | SOX2 regulated genes in Wounding from PMID: 30772301 | SOX2 |
| 4976 | UBA5 | SOX2 regulated genes in Wounding from PMID: 30772301 | SOX2 |
| 4977 | LRRFIP1 | SOX2 regulated genes in Wounding from PMID: 30772301 | SOX2 |
| 4978 | NIPAL4 | SOX2 regulated genes in Wounding from PMID: 30772301 | SOX2 |
| 4979 | PDK1 | SOX2 regulated genes in Wounding from PMID: 30772301 | SOX2 |
| 4980 | TMEM40 | SOX2 regulated genes in Wounding from PMID: 30772301 | SOX2 |
| 4981 | NME6 | SOX2 regulated genes in Wounding from PMID: 30772301 | SOX2 |
| 4982 | SLC27A6 | SOX2 regulated genes in Wounding from PMID: 30772301 | SOX2 |
| 4983 | PGAP1 | SOX2 regulated genes in Wounding from PMID: 30772301 | SOX2 |
| 4984 | KCTD11 | SOX2 regulated genes in Wounding from PMID: 30772301 | SOX2 |
| 4985 | POMGNT1 | SOX2 regulated genes in Wounding from PMID: 30772301 | SOX2 |
| 4986 | PRMT1 | SOX2 regulated genes in Wounding from PMID: 30772301 | SOX2 |
| 4987 | PDLIM4 | SOX2 regulated genes in Wounding from PMID: 30772301 | SOX2 |
| 4988 | CHMP4C | SOX2 regulated genes in Wounding from PMID: 30772301 | SOX2 |
| 4989 | SLC44A2 | SOX2 regulated genes in Wounding from PMID: 30772301 | SOX2 |
| 4990 | LUZP1 | SOX2 regulated genes in Wounding from PMID: 30772301 | SOX2 |
| 4991 | TWF1 | SOX2 regulated genes in Wounding from PMID: 30772301 | SOX2 |
| 4992 | CTSE | SOX2 regulated genes in Wounding from PMID: 30772301 | SOX2 |
| 4993 | CDC42BPA | SOX2 regulated genes in Wounding from PMID: 30772301 | SOX2 |
| 4994 | SSR2 | SOX2 regulated genes in Wounding from PMID: 30772301 | SOX2 |
| 4995 | LIMA1 | SOX2 regulated genes in Wounding from PMID: 30772301 | SOX2 |
| 4996 | EFNA4 | SOX2 regulated genes in Wounding from PMID: 30772301 | SOX2 |
| 4997 | GUCY2C | SOX2 regulated genes in Wounding from PMID: 30772301 | SOX2 |
| 4998 | ATP1B1 | SOX2 regulated genes in Wounding from PMID: 30772301 | SOX2 |
| 4999 | EFNA1 | SOX2 regulated genes in Wounding from PMID: 30772301 | SOX2 |
| 5000 | GTF2IRD1 | SOX2 regulated genes in Wounding from PMID: 30772301 | SOX2 |

| | | | |
|---|---|---|---|
| 5001 | PSME3 | SOX2 regulated genes in Wounding from PMID: 30772301 | SOX2 |
| 5002 | MKRN2OS | SOX2 regulated genes in Wounding from PMID: 30772301 | SOX2 |
| 5003 | B3GNT5 | SOX2 regulated genes in Wounding from PMID: 30772301 | SOX2 |
| 5004 | CNTN2 | SOX2 regulated genes in Wounding from PMID: 30772301 | SOX2 |
| 5005 | KLC3 | SOX2 regulated genes in Wounding from PMID: 30772301 | SOX2 |
| 5006 | HIP1R | SOX2 regulated genes in Wounding from PMID: 30772301 | SOX2 |
| 5007 | EFNA5 | SOX2 regulated genes in Wounding from PMID: 30772301 | SOX2 |
| 5008 | DAZAP1 | SOX2 regulated genes in Wounding from PMID: 30772301 | SOX2 |
| 5009 | PRKAR1A | SOX2 regulated genes in Wounding from PMID: 30772301 | SOX2 |
| 5010 | NAV2 | SOX2 regulated genes in Wounding from PMID: 30772301 | SOX2 |
| 5011 | RANBP10 | SOX2 regulated genes in Wounding from PMID: 30772301 | SOX2 |
| 5012 | PFN2 | SOX2 regulated genes in Wounding from PMID: 30772301 | SOX2 |
| 5013 | SNX16 | SOX2 regulated genes in Wounding from PMID: 30772301 | SOX2 |
| 5014 | NME1 | SOX2 regulated genes in Wounding from PMID: 30772301 | SOX2 |
| 5015 | FYTTD1 | SOX2 regulated genes in Wounding from PMID: 30772301 | SOX2 |
| 5016 | BEAN1 | SOX2 regulated genes in Wounding from PMID: 30772301 | SOX2 |
| 5017 | SORT1 | SOX2 regulated genes in Wounding from PMID: 30772301 | SOX2 |
| 5018 | NCMAP | SOX2 regulated genes in Wounding from PMID: 30772301 | SOX2 |
| 5019 | LTBP3 | SOX2 regulated genes in Wounding from PMID: 30772301 | SOX2 |
| 5020 | B3GNT2 | SOX2 regulated genes in Wounding from PMID: 30772301 | SOX2 |
| 5021 | CCDC124 | SOX2 regulated genes in Wounding from PMID: 30772301 | SOX2 |
| 5022 | CLDN4 | SOX2 regulated genes in Wounding from PMID: 30772301 | SOX2 |
| 5023 | SHF | SOX2 regulated genes in Wounding from PMID: 30772301 | SOX2 |
| 5024 | RPL29 | SOX2 regulated genes in Wounding from PMID: 30772301 | SOX2 |
| 5025 | COQ10B | SOX2 regulated genes in Wounding from PMID: 30772301 | SOX2 |
| 5026 | FAM187B | SOX2 regulated genes in Wounding from PMID: 30772301 | SOX2 |
| 5027 | PSMD8 | SOX2 regulated genes in Wounding from PMID: 30772301 | SOX2 |
| 5028 | GATAD2A | SOX2 regulated genes in Wounding from PMID: 30772301 | SOX2 |
| 5029 | TUBB3 | SOX2 regulated genes in Wounding from PMID: 30772301 | SOX2 |
| 5030 | BZW1 | SOX2 regulated genes in Wounding from PMID: 30772301 | SOX2 |
| 5031 | KDM5B | SOX2 regulated genes in Wounding from PMID: 30772301 | SOX2 |
| 5032 | TMEM222 | SOX2 regulated genes in Wounding from PMID: 30772301 | SOX2 |
| 5033 | HAUS8 | SOX2 regulated genes in Wounding from PMID: 30772301 | SOX2 |
| 5034 | IL20 | SOX2 regulated genes in Wounding from PMID: 30772301 | SOX2 |
| 5035 | TIGD5 | SOX2 regulated genes in Wounding from PMID: 30772301 | SOX2 |
| 5036 | FBXO28 | SOX2 regulated genes in Wounding from PMID: 30772301 | SOX2 |
| 5037 | VSIG8 | SOX2 regulated genes in Wounding from PMID: 30772301 | SOX2 |
| 5038 | FBXO38 | SOX2 regulated genes in Wounding from PMID: 30772301 | SOX2 |
| 5039 | CSNK1D | SOX2 regulated genes in Wounding from PMID: 30772301 | SOX2 |
| 5040 | FOXN1 | SOX2 regulated genes in Wounding from PMID: 30772301 | SOX2 |
| 5041 | WNT3A | SOX2 regulated genes in Wounding from PMID: 30772301 | SOX2 |
| 5042 | EAF1 | SOX2 regulated genes in Wounding from PMID: 30772301 | SOX2 |
| 5043 | TOP2B | SOX2 regulated genes in Wounding from PMID: 30772301 | SOX2 |
| 5044 | NGEF | SOX2 regulated genes in Wounding from PMID: 30772301 | SOX2 |
| 5045 | LRRC59 | SOX2 regulated genes in Wounding from PMID: 30772301 | SOX2 |
| 5046 | TTC13 | SOX2 regulated genes in Wounding from PMID: 30772301 | SOX2 |
| 5047 | WDR77 | SOX2 regulated genes in Wounding from PMID: 30772301 | SOX2 |
| 5048 | S100A3 | SOX2 regulated genes in Wounding from PMID: 30772301 | SOX2 |
| 5049 | GNAI3 | SOX2 regulated genes in Wounding from PMID: 30772301 | SOX2 |
| 5050 | KRT27 | SOX2 regulated genes in Wounding from PMID: 30772301 | SOX2 |
| 5051 | DIS3L | SOX2 regulated genes in Wounding from PMID: 30772301 | SOX2 |
| 5052 | GAB1 | SOX2 regulated genes in Wounding from PMID: 30772301 | SOX2 |
| 5053 | DDX5 | SOX2 regulated genes in Wounding from PMID: 30772301 | SOX2 |
| 5054 | UBTF | SOX2 regulated genes in Wounding from PMID: 30772301 | SOX2 |
| 5055 | TXLNA | SOX2 regulated genes in Wounding from PMID: 30772301 | SOX2 |
| 5056 | MEMO1 | SOX2 regulated genes in Wounding from PMID: 30772301 | SOX2 |
| 5057 | CRABP2 | SOX2 regulated genes in Wounding from PMID: 30772301 | SOX2 |
| 5058 | DEDD2 | SOX2 regulated genes in Wounding from PMID: 30772301 | SOX2 |
| 5059 | TPRG1 | SOX2 regulated genes in Wounding from PMID: 30772301 | SOX2 |
| 5060 | MYO6 | SOX2 regulated genes in Wounding from PMID: 30772301 | SOX2 |
| 5061 | HOOK1 | SOX2 regulated genes in Wounding from PMID: 30772301 | SOX2 |

| | | | |
|---|---|---|---|
| 5062 | SEMA6A | SOX2 regulated genes in Wounding from PMID: 30772301 | SOX2 |
| 5063 | TEPP | SOX2 regulated genes in Wounding from PMID: 30772301 | SOX2 |
| 5064 | SMAD3 | SOX2 regulated genes in Wounding from PMID: 30772301 | SOX2 |
| 5065 | REEP5 | SOX2 regulated genes in Wounding from PMID: 30772301 | SOX2 |
| 5066 | C1QL1 | SOX2 regulated genes in Wounding from PMID: 30772301 | SOX2 |
| 5067 | ECE1 | SOX2 regulated genes in Wounding from PMID: 30772301 | SOX2 |
| 5068 | HINFP | SOX2 regulated genes in Wounding from PMID: 30772301 | SOX2 |
| 5069 | KRT18 | SOX2 regulated genes in Wounding from PMID: 30772301 | SOX2 |
| 5070 | ARF6 | SOX2 regulated genes in Wounding from PMID: 30772301 | SOX2 |
| 5071 | GSK3B | SOX2 regulated genes in Wounding from PMID: 30772301 | SOX2 |
| 5072 | DUSP10 | SOX2 regulated genes in Wounding from PMID: 30772301 | SOX2 |
| 5073 | SEC61A1 | SOX2 regulated genes in Wounding from PMID: 30772301 | SOX2 |
| 5074 | FBXW11 | SOX2 regulated genes in Wounding from PMID: 30772301 | SOX2 |
| 5075 | SF3B4 | SOX2 regulated genes in Wounding from PMID: 30772301 | SOX2 |
| 5076 | ATP2C1 | SOX2 regulated genes in Wounding from PMID: 30772301 | SOX2 |
| 5077 | KRT36 | SOX2 regulated genes in Wounding from PMID: 30772301 | SOX2 |
| 5078 | HEYL | SOX2 regulated genes in Wounding from PMID: 30772301 | SOX2 |
| 5079 | PHC2 | SOX2 regulated genes in Wounding from PMID: 30772301 | SOX2 |
| 5080 | UBE2I | SOX2 regulated genes in Wounding from PMID: 30772301 | SOX2 |
| 5081 | KCNH1 | SOX2 regulated genes in Wounding from PMID: 30772301 | SOX2 |
| 5082 | PPP1R13L | SOX2 regulated genes in Wounding from PMID: 30772301 | SOX2 |
| 5083 | PPM1D | SOX2 regulated genes in Wounding from PMID: 30772301 | SOX2 |
| 5084 | ACP5 | SOX2 regulated genes in Wounding from PMID: 30772301 | SOX2 |
| 5085 | MRGPRX3 | SOX2 regulated genes in Wounding from PMID: 30772301 | SOX2 |
| 5086 | CHMP2B | SOX2 regulated genes in Wounding from PMID: 30772301 | SOX2 |
| 5087 | PIWIL2 | SOX2 regulated genes in Wounding from PMID: 30772301 | SOX2 |
| 5088 | ATP11B | SOX2 regulated genes in Wounding from PMID: 30772301 | SOX2 |
| 5089 | CFAP57 | SOX2 regulated genes in Wounding from PMID: 30772301 | SOX2 |
| 5090 | EIF5A | SOX2 regulated genes in Wounding from PMID: 30772301 | SOX2 |
| 5091 | TFG | SOX2 regulated genes in Wounding from PMID: 30772301 | SOX2 |
| 5092 | RAB34 | SOX2 regulated genes in Wounding from PMID: 30772301 | SOX2 |
| 5093 | ATOH7 | SOX2 regulated genes in Wounding from PMID: 30772301 | SOX2 |
| 5094 | KLK7 | SOX2 regulated genes in Wounding from PMID: 30772301 | SOX2 |
| 5095 | PLEKHG6 | SOX2 regulated genes in Wounding from PMID: 30772301 | SOX2 |
| 5096 | ARAP2 | SOX2 regulated genes in Wounding from PMID: 30772301 | SOX2 |
| 5097 | PIAS4 | SOX2 regulated genes in Wounding from PMID: 30772301 | SOX2 |
| 5098 | ABCB11 | SOX2 regulated genes in Wounding from PMID: 30772301 | SOX2 |
| 5099 | CLDN1 | SOX2 regulated genes in Wounding from PMID: 30772301 | SOX2 |
| 5100 | LAMB3 | SOX2 regulated genes in Wounding from PMID: 30772301 | SOX2 |
| 5101 | HAS1 | SOX2 regulated genes in Wounding from PMID: 30772301 | SOX2 |
| 5102 | VTCN1 | SOX2 regulated genes in Wounding from PMID: 30772301 | SOX2 |
| 5103 | UBE2F | SOX2 regulated genes in Wounding from PMID: 30772301 | SOX2 |
| 5104 | MUC1 | SOX2 regulated genes in Wounding from PMID: 30772301 | SOX2 |
| 5105 | TSC22D2 | SOX2 regulated genes in Wounding from PMID: 30772301 | SOX2 |
| 5106 | ACTL6A | SOX2 regulated genes in Wounding from PMID: 30772301 | SOX2 |
| 5107 | TNRC18 | SOX2 regulated genes in Wounding from PMID: 30772301 | SOX2 |
| 5108 | DCAKD | SOX2 regulated genes in Wounding from PMID: 30772301 | SOX2 |
| 5109 | LPAR3 | SOX2 regulated genes in Wounding from PMID: 30772301 | SOX2 |
| 5110 | NIF3L1 | SOX2 regulated genes in Wounding from PMID: 30772301 | SOX2 |
| 5111 | TAF13 | SOX2 regulated genes in Wounding from PMID: 30772301 | SOX2 |
| 5112 | PCDH1 | SOX2 regulated genes in Wounding from PMID: 30772301 | SOX2 |
| 5113 | RTN4 | SOX2 regulated genes in Wounding from PMID: 30772301 | SOX2 |
| 5114 | ZPR1 | SOX2 regulated genes in Wounding from PMID: 30772301 | SOX2 |
| 5115 | E2F2 | SOX2 regulated genes in Wounding from PMID: 30772301 | SOX2 |
| 5116 | GRAMD1C | SOX2 regulated genes in Wounding from PMID: 30772301 | SOX2 |
| 5117 | GCNT2 | SOX2 regulated genes in Wounding from PMID: 30772301 | SOX2 |
| 5118 | RGMA | SOX2 regulated genes in Wounding from PMID: 30772301 | SOX2 |
| 5119 | MXD1 | SOX2 regulated genes in Wounding from PMID: 30772301 | SOX2 |
| 5120 | MEX3A | SOX2 regulated genes in Wounding from PMID: 30772301 | SOX2 |
| 5121 | DYRK3 | SOX2 regulated genes in Wounding from PMID: 30772301 | SOX2 |
| 5122 | RNF11 | SOX2 regulated genes in Wounding from PMID: 30772301 | SOX2 |

| | | | |
|---|---|---|---|
| 5123 | USP2 | SOX2 regulated genes in Wounding from PMID: 30772301 | SOX2 |
| 5124 | FSCN2 | SOX2 regulated genes in Wounding from PMID: 30772301 | SOX2 |
| 5125 | PGBD1 | SOX2 regulated genes in Wounding from PMID: 30772301 | SOX2 |
| 5126 | IFNLR1 | SOX2 regulated genes in Wounding from PMID: 30772301 | SOX2 |
| 5127 | EMP2 | SOX2 regulated genes in Wounding from PMID: 30772301 | SOX2 |
| 5128 | ACOT7 | SOX2 regulated genes in Wounding from PMID: 30772301 | SOX2 |
| 5129 | NTF4 | SOX2 regulated genes in Wounding from PMID: 30772301 | SOX2 |
| 5130 | SLCO2A1 | SOX2 regulated genes in Wounding from PMID: 30772301 | SOX2 |
| 5131 | TSPAN1 | SOX2 regulated genes in Wounding from PMID: 30772301 | SOX2 |
| 5132 | HNRNPF | SOX2 regulated genes in Wounding from PMID: 30772301 | SOX2 |
| 5133 | MYO1B | SOX2 regulated genes in Wounding from PMID: 30772301 | SOX2 |
| 5134 | DHX29 | SOX2 regulated genes in Wounding from PMID: 30772301 | SOX2 |
| 5135 | HOMER3 | SOX2 regulated genes in Wounding from PMID: 30772301 | SOX2 |
| 5136 | RIMS1 | SOX2 regulated genes in Wounding from PMID: 30772301 | SOX2 |
| 5137 | F11R | SOX2 regulated genes in Wounding from PMID: 30772301 | SOX2 |
| 5138 | PHYHIP | SOX2 regulated genes in Wounding from PMID: 30772301 | SOX2 |
| 5139 | ERC1 | SOX2 regulated genes in Wounding from PMID: 30772301 | SOX2 |
| 5140 | SLC45A3 | SOX2 regulated genes in Wounding from PMID: 30772301 | SOX2 |
| 5141 | GPR12 | SOX2 regulated genes in Keratinocyte Overexpression from PMID: 30772301 | SOX2 |
| 5142 | ZNF512B | SOX2 regulated genes in Keratinocyte Overexpression from PMID: 30772301 | SOX2 |
| 5143 | PWP2 | SOX2 regulated genes in Keratinocyte Overexpression from PMID: 30772301 | SOX2 |
| 5144 | RAB27B | SOX2 regulated genes in Keratinocyte Overexpression from PMID: 30772301 | SOX2 |
| 5145 | MZT1 | SOX2 regulated genes in Keratinocyte Overexpression from PMID: 30772301 | SOX2 |
| 5146 | SYN3 | SOX2 regulated genes in Keratinocyte Overexpression from PMID: 30772301 | SOX2 |
| 5147 | USP12 | SOX2 regulated genes in Keratinocyte Overexpression from PMID: 30772301 | SOX2 |
| 5148 | ARL5B | SOX2 regulated genes in Keratinocyte Overexpression from PMID: 30772301 | SOX2 |
| 5149 | PCGF3 | SOX2 regulated genes in Keratinocyte Overexpression from PMID: 30772301 | SOX2 |
| 5150 | HORMAD2 | SOX2 regulated genes in Keratinocyte Overexpression from PMID: 30772301 | SOX2 |
| 5151 | LIPG | SOX2 regulated genes in Keratinocyte Overexpression from PMID: 30772301 | SOX2 |
| 5152 | YES1 | SOX2 regulated genes in Keratinocyte Overexpression from PMID: 30772301 | SOX2 |
| 5153 | UNC79 | SOX2 regulated genes in Keratinocyte Overexpression from PMID: 30772301 | SOX2 |
| 5154 | ZBTB21 | SOX2 regulated genes in Keratinocyte Overexpression from PMID: 30772301 | SOX2 |
| 5155 | GTPBP6 | SOX2 regulated genes in Keratinocyte Overexpression from PMID: 30772301 | SOX2 |
| 5156 | CPLX1 | SOX2 regulated genes in Keratinocyte Overexpression from PMID: 30772301 | SOX2 |
| 5157 | ATG2B | SOX2 regulated genes in Keratinocyte Overexpression from PMID: 30772301 | SOX2 |
| 5158 | SMOX | SOX2 regulated genes in Keratinocyte Overexpression from PMID: 30772301 | SOX2 |
| 5159 | CLDN17 | SOX2 regulated genes in Keratinocyte Overexpression from PMID: 30772301 | SOX2 |
| 5160 | DCAF4 | SOX2 regulated genes in Keratinocyte Overexpression from PMID: 30772301 | SOX2 |
| 5161 | MOCOS | SOX2 regulated genes in Keratinocyte Overexpression from PMID: 30772301 | SOX2 |
| 5162 | PBDC1 | SOX2 regulated genes in Keratinocyte Overexpression from PMID: 30772301 | SOX2 |
| 5163 | PRELID3B | SOX2 regulated genes in Keratinocyte Overexpression from PMID: 30772301 | SOX2 |
| 5164 | MEX3C | SOX2 regulated genes in Keratinocyte Overexpression from PMID: 30772301 | SOX2 |
| 5165 | EVA1C | SOX2 regulated genes in Keratinocyte Overexpression from PMID: 30772301 | SOX2 |
| 5166 | PSMA7 | SOX2 regulated genes in Keratinocyte Overexpression from PMID: 30772301 | SOX2 |
| 5167 | NUDT5 | SOX2 regulated genes in Keratinocyte Overexpression from PMID: 30772301 | SOX2 |
| 5168 | VBP1 | SOX2 regulated genes in Keratinocyte Overexpression from PMID: 30772301 | SOX2 |
| 5169 | SLC25A43 | SOX2 regulated genes in Keratinocyte Overexpression from PMID: 30772301 | SOX2 |
| 5170 | TTLL1 | SOX2 regulated genes in Keratinocyte Overexpression from PMID: 30772301 | SOX2 |
| 5171 | SIM2 | SOX2 regulated genes in Keratinocyte Overexpression from PMID: 30772301 | SOX2 |
| 5172 | RRP1B | SOX2 regulated genes in Keratinocyte Overexpression from PMID: 30772301 | SOX2 |
| 5173 | CEP76 | SOX2 regulated genes in Keratinocyte Overexpression from PMID: 30772301 | SOX2 |
| 5174 | PCNX4 | SOX2 regulated genes in Keratinocyte Overexpression from PMID: 30772301 | SOX2 |
| 5175 | ZFP64 | SOX2 regulated genes in Keratinocyte Overexpression from PMID: 30772301 | SOX2 |
| 5176 | TAF4B | SOX2 regulated genes in Keratinocyte Overexpression from PMID: 30772301 | SOX2 |
| 5177 | SKA1 | SOX2 regulated genes in Keratinocyte Overexpression from PMID: 30772301 | SOX2 |
| 5178 | PROSER1 | SOX2 regulated genes in Keratinocyte Overexpression from PMID: 30772301 | SOX2 |
| 5179 | TTC6 | SOX2 regulated genes in Keratinocyte Overexpression from PMID: 30772301 | SOX2 |
| 5180 | NFX1 | SOX2 regulated genes in Keratinocyte Overexpression from PMID: 30772301 | SOX2 |
| 5181 | POMP | SOX2 regulated genes in Keratinocyte Overexpression from PMID: 30772301 | SOX2 |
| 5182 | SCAF4 | SOX2 regulated genes in Keratinocyte Overexpression from PMID: 30772301 | SOX2 |
| 5183 | LINGO2 | SOX2 regulated genes in Keratinocyte Overexpression from PMID: 30772301 | SOX2 |

| | | | |
|---|---|---|---|
| 5184 | PPP6R2 | SOX2 regulated genes in Keratinocyte Overexpression from PMID: 30772301 | SOX2 |
| 5185 | SEC23B | SOX2 regulated genes in Keratinocyte Overexpression from PMID: 30772301 | SOX2 |
| 5186 | RALY | SOX2 regulated genes in Keratinocyte Overexpression from PMID: 30772301 | SOX2 |
| 5187 | PDE9A | SOX2 regulated genes in Keratinocyte Overexpression from PMID: 30772301 | SOX2 |
| 5188 | HAUS6 | SOX2 regulated genes in Keratinocyte Overexpression from PMID: 30772301 | SOX2 |
| 5189 | ANKRD9 | SOX2 regulated genes in Keratinocyte Overexpression from PMID: 30772301 | SOX2 |
| 5190 | RBX1 | SOX2 regulated genes in Keratinocyte Overexpression from PMID: 30772301 | SOX2 |
| 5191 | MNAT1 | SOX2 regulated genes in Keratinocyte Overexpression from PMID: 30772301 | SOX2 |
| 5192 | BEX3 | SOX2 regulated genes in Keratinocyte Overexpression from PMID: 30772301 | SOX2 |
| 5193 | PSPC1 | SOX2 regulated genes in Keratinocyte Overexpression from PMID: 30772301 | SOX2 |
| 5194 | BMP7 | SOX2 regulated genes in Keratinocyte Overexpression from PMID: 30772301 | SOX2 |
| 5195 | RPS21 | SOX2 regulated genes in Keratinocyte Overexpression from PMID: 30772301 | SOX2 |
| 5196 | SCEL | SOX2 regulated genes in Keratinocyte Overexpression from PMID: 30772301 | SOX2 |
| 5197 | PCDH8 | SOX2 regulated genes in Keratinocyte Overexpression from PMID: 30772301 | SOX2 |
| 5198 | PSIP1 | SOX2 regulated genes in Keratinocyte Overexpression from PMID: 30772301 | SOX2 |
| 5199 | NUFIP1 | SOX2 regulated genes in Keratinocyte Overexpression from PMID: 30772301 | SOX2 |
| 5200 | XKRX | SOX2 regulated genes in Keratinocyte Overexpression from PMID: 30772301 | SOX2 |
| 5201 | CSTB | SOX2 regulated genes in Keratinocyte Overexpression from PMID: 30772301 | SOX2 |
| 5202 | GTF3A | SOX2 regulated genes in Keratinocyte Overexpression from PMID: 30772301 | SOX2 |
| 5203 | CDKN2B | SOX2 regulated genes in Keratinocyte Overexpression from PMID: 30772301 | SOX2 |
| 5204 | CDC123 | SOX2 regulated genes in Keratinocyte Overexpression from PMID: 30772301 | SOX2 |
| 5205 | PAN3 | SOX2 regulated genes in Keratinocyte Overexpression from PMID: 30772301 | SOX2 |
| 5206 | HOPX | SOX2 regulated genes in Keratinocyte Overexpression from PMID: 30772301 | SOX2 |
| 5207 | TP53INP2 | SOX2 regulated genes in Keratinocyte Overexpression from PMID: 30772301 | SOX2 |
| 5208 | WDR1 | SOX2 regulated genes in Keratinocyte Overexpression from PMID: 30772301 | SOX2 |
| 5209 | GID8 | SOX2 regulated genes in Keratinocyte Overexpression from PMID: 30772301 | SOX2 |
| 5210 | POLR3F | SOX2 regulated genes in Keratinocyte Overexpression from PMID: 30772301 | SOX2 |
| 5211 | MAGEE2 | SOX2 regulated genes in Keratinocyte Overexpression from PMID: 30772301 | SOX2 |
| 5212 | MOK | SOX2 regulated genes in Keratinocyte Overexpression from PMID: 30772301 | SOX2 |
| 5213 | HMGN5 | SOX2 regulated genes in Keratinocyte Overexpression from PMID: 30772301 | SOX2 |
| 5214 | NUP50 | SOX2 regulated genes in Keratinocyte Overexpression from PMID: 30772301 | SOX2 |
| 5215 | MRPL39 | SOX2 regulated genes in Keratinocyte Overexpression from PMID: 30772301 | SOX2 |
| 5216 | RPRD1B | SOX2 regulated genes in Keratinocyte Overexpression from PMID: 30772301 | SOX2 |
| 5217 | DHX15 | SOX2 regulated genes in Keratinocyte Overexpression from PMID: 30772301 | SOX2 |
| 5218 | UPF2 | SOX2 regulated genes in Keratinocyte Overexpression from PMID: 30772301 | SOX2 |
| 5219 | CDC42BPB | SOX2 regulated genes in Keratinocyte Overexpression from PMID: 30772301 | SOX2 |
| 5220 | RPS4X | SOX2 regulated genes in Keratinocyte Overexpression from PMID: 30772301 | SOX2 |
| 5221 | CITED1 | SOX2 regulated genes in Keratinocyte Overexpression from PMID: 30772301 | SOX2 |
| 5222 | RNF152 | SOX2 regulated genes in Keratinocyte Overexpression from PMID: 30772301 | SOX2 |
| 5223 | RFC3 | SOX2 regulated genes in Keratinocyte Overexpression from PMID: 30772301 | SOX2 |
| 5224 | RBM17 | SOX2 regulated genes in Keratinocyte Overexpression from PMID: 30772301 | SOX2 |
| 5225 | SPATA2 | SOX2 regulated genes in Keratinocyte Overexpression from PMID: 30772301 | SOX2 |
| 5226 | KIN | SOX2 regulated genes in Keratinocyte Overexpression from PMID: 30772301 | SOX2 |
| 5227 | MIS18BP1 | SOX2 regulated genes in Keratinocyte Overexpression from PMID: 30772301 | SOX2 |
| 5228 | PRR14L | SOX2 regulated genes in Keratinocyte Overexpression from PMID: 30772301 | SOX2 |
| 5229 | HAUS1 | SOX2 regulated genes in Keratinocyte Overexpression from PMID: 30772301 | SOX2 |
| 5230 | CDC42EP1 | SOX2 regulated genes in Keratinocyte Overexpression from PMID: 30772301 | SOX2 |
| 5231 | CRYL1 | SOX2 regulated genes in Keratinocyte Overexpression from PMID: 30772301 | SOX2 |
| 5232 | AMER1 | SOX2 regulated genes in Keratinocyte Overexpression from PMID: 30772301 | SOX2 |
| 5233 | YY1 | SOX2 regulated genes in Keratinocyte Overexpression from PMID: 30772301 | SOX2 |
| 5234 | NOL12 | SOX2 regulated genes in Keratinocyte Overexpression from PMID: 30772301 | SOX2 |
| 5235 | PSMG1 | SOX2 regulated genes in Keratinocyte Overexpression from PMID: 30772301 | SOX2 |
| 5236 | SLC19A1 | SOX2 regulated genes in Keratinocyte Overexpression from PMID: 30772301 | SOX2 |
| 5237 | SLC25A17 | SOX2 regulated genes in Keratinocyte Overexpression from PMID: 30772301 | SOX2 |
| 5238 | BTG3 | SOX2 regulated genes in Keratinocyte Overexpression from PMID: 30772301 | SOX2 |
| 5239 | YBEY | SOX2 regulated genes in Keratinocyte Overexpression from PMID: 30772301 | SOX2 |
| 5240 | ISM1 | SOX2 regulated genes in Keratinocyte Overexpression from PMID: 30772301 | SOX2 |
| 5241 | SSTR1 | SOX2 regulated genes in Keratinocyte Overexpression from PMID: 30772301 | SOX2 |
| 5242 | CTNNBL1 | SOX2 regulated genes in Keratinocyte Overexpression from PMID: 30772301 | SOX2 |
| 5243 | DSG3 | SOX2 regulated genes in Keratinocyte Overexpression from PMID: 30772301 | SOX2 |
| 5244 | BTBD7 | SOX2 regulated genes in Keratinocyte Overexpression from PMID: 30772301 | SOX2 |

| | | | |
|---|---|---|---|
| 5245 | TUBAL3 | SOX2 regulated genes in Keratinocyte Overexpression from PMID: 30772301 | SOX2 |
| 5246 | ATP5F1A | SOX2 regulated genes in Keratinocyte Overexpression from PMID: 30772301 | SOX2 |
| 5247 | BTBD3 | SOX2 regulated genes in Keratinocyte Overexpression from PMID: 30772301 | SOX2 |
| 5248 | MAFF | SOX2 regulated genes in Keratinocyte Overexpression from PMID: 30772301 | SOX2 |
| 5249 | DOK5 | SOX2 regulated genes in Keratinocyte Overexpression from PMID: 30772301 | SOX2 |
| 5250 | U2AF1 | SOX2 regulated genes in Keratinocyte Overexpression from PMID: 30772301 | SOX2 |
| 5251 | NUDT11 | SOX2 regulated genes in Keratinocyte Overexpression from PMID: 30772301 | SOX2 |
| 5252 | DLG3 | SOX2 regulated genes in Keratinocyte Overexpression from PMID: 30772301 | SOX2 |
| 5253 | BRWD1 | SOX2 regulated genes in Keratinocyte Overexpression from PMID: 30772301 | SOX2 |
| 5254 | MORC4 | SOX2 regulated genes in Keratinocyte Overexpression from PMID: 30772301 | SOX2 |
| 5255 | SMU1 | SOX2 regulated genes in Keratinocyte Overexpression from PMID: 30772301 | SOX2 |
| 5256 | TSR2 | SOX2 regulated genes in Keratinocyte Overexpression from PMID: 30772301 | SOX2 |
| 5257 | FANCM | SOX2 regulated genes in Keratinocyte Overexpression from PMID: 30772301 | SOX2 |
| 5258 | DMRTA1 | SOX2 regulated genes in Keratinocyte Overexpression from PMID: 30772301 | SOX2 |
| 5259 | RBBP8NL | SOX2 regulated genes in Keratinocyte Overexpression from PMID: 30772301 | SOX2 |
| 5260 | TASP1 | SOX2 regulated genes in Keratinocyte Overexpression from PMID: 30772301 | SOX2 |
| 5261 | TECPR2 | SOX2 regulated genes in Keratinocyte Overexpression from PMID: 30772301 | SOX2 |
| 5262 | TFAP2C | SOX2 regulated genes in Keratinocyte Overexpression from PMID: 30772301 | SOX2 |
| 5263 | PRDM15 | SOX2 regulated genes in Keratinocyte Overexpression from PMID: 30772301 | SOX2 |
| 5264 | RUNX1 | SOX2 regulated genes in Keratinocyte Overexpression from PMID: 30772301 | SOX2 |
| 5265 | ERCC6L | SOX2 regulated genes in Keratinocyte Overexpression from PMID: 30772301 | SOX2 |
| 5266 | CHODL | SOX2 regulated genes in Keratinocyte Overexpression from PMID: 30772301 | SOX2 |
| 5267 | FGD1 | SOX2 regulated genes in Keratinocyte Overexpression from PMID: 30772301 | SOX2 |
| 5268 | RBM3 | SOX2 regulated genes in Keratinocyte Overexpression from PMID: 30772301 | SOX2 |
| 5269 | GPR137C | SOX2 regulated genes in Keratinocyte Overexpression from PMID: 30772301 | SOX2 |
| 5270 | SYTL5 | SOX2 regulated genes in Keratinocyte Overexpression from PMID: 30772301 | SOX2 |
| 5271 | HSPH1 | SOX2 regulated genes in Keratinocyte Overexpression from PMID: 30772301 | SOX2 |
| 5272 | CDKL5 | SOX2 regulated genes in Keratinocyte Overexpression from PMID: 30772301 | SOX2 |
| 5273 | GLRX5 | SOX2 regulated genes in Keratinocyte Overexpression from PMID: 30772301 | SOX2 |
| 5274 | SAMD10 | SOX2 regulated genes in Keratinocyte Overexpression from PMID: 30772301 | SOX2 |
| 5275 | KIF4A | SOX2 regulated genes in Keratinocyte Overexpression from PMID: 30772301 | SOX2 |
| 5276 | LRR1 | SOX2 regulated genes in Keratinocyte Overexpression from PMID: 30772301 | SOX2 |
| 5277 | POLE2 | SOX2 regulated genes in Keratinocyte Overexpression from PMID: 30772301 | SOX2 |
| 5278 | EPG5 | SOX2 regulated genes in Keratinocyte Overexpression from PMID: 30772301 | SOX2 |
| 5279 | DCUN1D4 | SOX2 regulated genes in Keratinocyte Overexpression from PMID: 30772301 | SOX2 |
| 5280 | SEC61A2 | SOX2 regulated genes in Keratinocyte Overexpression from PMID: 30772301 | SOX2 |
| 5281 | PKNOX1 | SOX2 regulated genes in Keratinocyte Overexpression from PMID: 30772301 | SOX2 |
| 5282 | FRMD4A | SOX2 regulated genes in Keratinocyte Overexpression from PMID: 30772301 | SOX2 |
| 5283 | CDH22 | SOX2 regulated genes in Keratinocyte Overexpression from PMID: 30772301 | SOX2 |
| 5284 | RBM11 | SOX2 regulated genes in Keratinocyte Overexpression from PMID: 30772301 | SOX2 |
| 5285 | SEH1L | SOX2 regulated genes in Keratinocyte Overexpression from PMID: 30772301 | SOX2 |
| 5286 | FAM110A | SOX2 regulated genes in Keratinocyte Overexpression from PMID: 30772301 | SOX2 |
| 5287 | C21orf58 | SOX2 regulated genes in Keratinocyte Overexpression from PMID: 30772301 | SOX2 |
| 5288 | HS3ST1 | SOX2 regulated genes in Keratinocyte Overexpression from PMID: 30772301 | SOX2 |
| 5289 | PFDN4 | SOX2 regulated genes in Keratinocyte Overexpression from PMID: 30772301 | SOX2 |
| 5290 | SEPHS1 | SOX2 regulated genes in Keratinocyte Overexpression from PMID: 30772301 | SOX2 |
| 5291 | TMX4 | SOX2 regulated genes in Keratinocyte Overexpression from PMID: 30772301 | SOX2 |
| 5292 | SPATA13 | SOX2 regulated genes in Keratinocyte Overexpression from PMID: 30772301 | SOX2 |
| 5293 | RCL1 | SOX2 regulated genes in Keratinocyte Overexpression from PMID: 30772301 | SOX2 |
| 5294 | BMP4 | SOX2 regulated genes in Keratinocyte Overexpression from PMID: 30772301 | SOX2 |
| 5295 | DEPDC5 | SOX2 regulated genes in Keratinocyte Overexpression from PMID: 30772301 | SOX2 |
| 5296 | PCNX1 | SOX2 regulated genes in Keratinocyte Overexpression from PMID: 30772301 | SOX2 |
| 5297 | ETS2 | SOX2 regulated genes in Keratinocyte Overexpression from PMID: 30772301 | SOX2 |
| 5298 | TPGS2 | SOX2 regulated genes in Keratinocyte Overexpression from PMID: 30772301 | SOX2 |
| 5299 | CDK5RAP1 | SOX2 regulated genes in Keratinocyte Overexpression from PMID: 30772301 | SOX2 |
| 5300 | LTN1 | SOX2 regulated genes in Keratinocyte Overexpression from PMID: 30772301 | SOX2 |
| 5301 | CFAP61 | SOX2 regulated genes in Keratinocyte Overexpression from PMID: 30772301 | SOX2 |
| 5302 | DONSON | SOX2 regulated genes in Keratinocyte Overexpression from PMID: 30772301 | SOX2 |
| 5303 | DIAPH3 | SOX2 regulated genes in Keratinocyte Overexpression from PMID: 30772301 | SOX2 |
| 5304 | KCTD1 | SOX2 regulated genes in Keratinocyte Overexpression from PMID: 30772301 | SOX2 |
| 5305 | LSM14B | SOX2 regulated genes in Keratinocyte Overexpression from PMID: 30772301 | SOX2 |

| | | | |
|---|---|---|---|
| 5306 | LHFPL6 | SOX2 regulated genes in Keratinocyte Overexpression from PMID: 30772301 | SOX2 |
| 5307 | PSMG2 | SOX2 regulated genes in Keratinocyte Overexpression from PMID: 30772301 | SOX2 |
| 5308 | ADRM1 | SOX2 regulated genes in Keratinocyte Overexpression from PMID: 30772301 | SOX2 |
| 5309 | MIPEP | SOX2 regulated genes in Keratinocyte Overexpression from PMID: 30772301 | SOX2 |
| 5310 | GTSE1 | SOX2 regulated genes in Keratinocyte Overexpression from PMID: 30772301 | SOX2 |
| 5311 | SLC37A1 | SOX2 regulated genes in Keratinocyte Overexpression from PMID: 30772301 | SOX2 |
| 5312 | TGIF1 | SOX2 regulated genes in Keratinocyte Overexpression from PMID: 30772301 | SOX2 |
| 5313 | RNF17 | SOX2 regulated genes in Keratinocyte Overexpression from PMID: 30772301 | SOX2 |
| 5314 | CENPC | SOX2 regulated genes in Keratinocyte Overexpression from PMID: 30772301 | SOX2 |
| 5315 | PSMC6 | SOX2 regulated genes in Keratinocyte Overexpression from PMID: 30772301 | SOX2 |
| 5316 | AHCY | SOX2 regulated genes in Keratinocyte Overexpression from PMID: 30772301 | SOX2 |
| 5317 | MYO5B | SOX2 regulated genes in Keratinocyte Overexpression from PMID: 30772301 | SOX2 |
| 5318 | NPC1 | SOX2 regulated genes in Keratinocyte Overexpression from PMID: 30772301 | SOX2 |
| 5319 | GRPEL1 | SOX2 regulated genes in Keratinocyte Overexpression from PMID: 30772301 | SOX2 |
| 5320 | PDXP | SOX2 regulated genes in Keratinocyte Overexpression from PMID: 30772301 | SOX2 |
| 5321 | RNGTT | SOX2 regulated genes in Keratinocyte Overexpression from PMID: 30772301 | SOX2 |
| 5322 | ESF1 | SOX2 regulated genes in Keratinocyte Overexpression from PMID: 30772301 | SOX2 |
| 5323 | QRFPR | SOX2 regulated genes in Keratinocyte Overexpression from PMID: 30772301 | SOX2 |
| 5324 | THTPA | SOX2 regulated genes in Keratinocyte Overexpression from PMID: 30772301 | SOX2 |
| 5325 | NKX1-2 | SOX2 regulated genes in Keratinocyte Overexpression from PMID: 30772301 | SOX2 |
| 5326 | C14orf93 | SOX2 regulated genes in Keratinocyte Overexpression from PMID: 30772301 | SOX2 |
| 5327 | MYO1D | SOX2 regulated genes in Keratinocyte Overexpression from PMID: 30772301 | SOX2 |
| 5328 | LASP1 | SOX2 regulated genes in Keratinocyte Overexpression from PMID: 30772301 | SOX2 |
| 5329 | DLGAP4 | SOX2 regulated genes in Keratinocyte Overexpression from PMID: 30772301 | SOX2 |
| 5330 | BCOR | SOX2 regulated genes in Keratinocyte Overexpression from PMID: 30772301 | SOX2 |
| 5331 | CYP17A1 | SOX2 regulated genes in Keratinocyte Overexpression from PMID: 30772301 | SOX2 |
| 5332 | FAM234B | SOX2 regulated genes in Keratinocyte Overexpression from PMID: 30772301 | SOX2 |
| 5333 | GPAT3 | SOX2 regulated genes in Keratinocyte Overexpression from PMID: 30772301 | SOX2 |
| 5334 | TNKS2 | SOX2 regulated genes in Keratinocyte Overexpression from PMID: 30772301 | SOX2 |
| 5335 | EFCAB11 | SOX2 regulated genes in Keratinocyte Overexpression from PMID: 30772301 | SOX2 |
| 5336 | GPATCH8 | SOX2 regulated genes in Keratinocyte Overexpression from PMID: 30772301 | SOX2 |
| 5337 | SUPT16H | SOX2 regulated genes in Keratinocyte Overexpression from PMID: 30772301 | SOX2 |
| 5338 | ZNF407 | SOX2 regulated genes in Keratinocyte Overexpression from PMID: 30772301 | SOX2 |
| 5339 | ATP5MF-PTCD1 | SOX2 regulated genes in Keratinocyte Overexpression from PMID: 30772301 | SOX2 |
| 5340 | TEX30 | SOX2 regulated genes in Keratinocyte Overexpression from PMID: 30772301 | SOX2 |
| 5341 | GRIA3 | SOX2 regulated genes in Keratinocyte Overexpression from PMID: 30772301 | SOX2 |
| 5342 | COPS4 | SOX2 regulated genes in Keratinocyte Overexpression from PMID: 30772301 | SOX2 |
| 5343 | TSSK4 | SOX2 regulated genes in Keratinocyte Overexpression from PMID: 30772301 | SOX2 |
| 5344 | INCENP | SOX2 regulated genes in Keratinocyte Overexpression from PMID: 30772301 | SOX2 |
| 5345 | LZTS1 | SOX2 regulated genes in Keratinocyte Overexpression from PMID: 30772301 | SOX2 |
| 5346 | SETBP1 | SOX2 regulated genes in Keratinocyte Overexpression from PMID: 30772301 | SOX2 |
| 5347 | RPL36A | SOX2 regulated genes in Keratinocyte Overexpression from PMID: 30772301 | SOX2 |
| 5348 | PLPP2 | SOX2 regulated genes in Keratinocyte Overexpression from PMID: 30772301 | SOX2 |
| 5349 | NF2 | SOX2 regulated genes in Keratinocyte Overexpression from PMID: 30772301 | SOX2 |
| 5350 | C18orf32 | SOX2 regulated genes in Keratinocyte Overexpression from PMID: 30772301 | SOX2 |
| 5351 | ABTB2 | SOX2 regulated genes in Keratinocyte Overexpression from PMID: 30772301 | SOX2 |
| 5352 | SERPINB13 | SOX2 regulated genes in Keratinocyte Overexpression from PMID: 30772301 | SOX2 |
| 5353 | MAP3K7 | SOX2 regulated genes in Keratinocyte Overexpression from PMID: 30772301 | SOX2 |
| 5354 | FRAT1 | SOX2 regulated genes in Keratinocyte Overexpression from PMID: 30772301 | SOX2 |
| 5355 | RNF167 | SOX2 regulated genes in Keratinocyte Overexpression from PMID: 30772301 | SOX2 |
| 5356 | TSPAN7 | SOX2 regulated genes in Keratinocyte Overexpression from PMID: 30772301 | SOX2 |
| 5357 | GUCD1 | SOX2 regulated genes in Keratinocyte Overexpression from PMID: 30772301 | SOX2 |
| 5358 | GSTCD | SOX2 regulated genes in Keratinocyte Overexpression from PMID: 30772301 | SOX2 |
| 5359 | ZBTB43 | SOX2 regulated genes in Keratinocyte Overexpression from PMID: 30772301 | SOX2 |
| 5360 | PPP2R2A | SOX2 regulated genes in Keratinocyte Overexpression from PMID: 30772301 | SOX2 |
| 5361 | NUDCD3 | SOX2 regulated genes in Keratinocyte Overexpression from PMID: 30772301 | SOX2 |
| 5362 | SERPINB11 | SOX2 regulated genes in Keratinocyte Overexpression from PMID: 30772301 | SOX2 |
| 5363 | SRXN1 | SOX2 regulated genes in Keratinocyte Overexpression from PMID: 30772301 | SOX2 |
| 5364 | LDLRAD3 | SOX2 regulated genes in Keratinocyte Overexpression from PMID: 30772301 | SOX2 |
| 5365 | NRBF2 | SOX2 regulated genes in Keratinocyte Overexpression from PMID: 30772301 | SOX2 |
| 5366 | MIEF1 | SOX2 regulated genes in Keratinocyte Overexpression from PMID: 30772301 | SOX2 |

| | | | |
|---|---|---|---|
| 5367 | LIG3 | SOX2 regulated genes in Keratinocyte Overexpression from PMID: 30772301 | SOX2 |
| 5368 | DLST | SOX2 regulated genes in Keratinocyte Overexpression from PMID: 30772301 | SOX2 |
| 5369 | ASPHD2 | SOX2 regulated genes in Keratinocyte Overexpression from PMID: 30772301 | SOX2 |
| 5370 | SPATC1L | SOX2 regulated genes in Keratinocyte Overexpression from PMID: 30772301 | SOX2 |
| 5371 | KIF1C | SOX2 regulated genes in Keratinocyte Overexpression from PMID: 30772301 | SOX2 |
| 5372 | PPIL2 | SOX2 regulated genes in Keratinocyte Overexpression from PMID: 30772301 | SOX2 |
| 5373 | PPP1R26 | SOX2 regulated genes in Keratinocyte Overexpression from PMID: 30772301 | SOX2 |
| 5374 | MED4 | SOX2 regulated genes in Keratinocyte Overexpression from PMID: 30772301 | SOX2 |
| 5375 | ZFYVE19 | SOX2 regulated genes in Keratinocyte Overexpression from PMID: 30772301 | SOX2 |
| 5376 | FRMD6 | SOX2 regulated genes in Keratinocyte Overexpression from PMID: 30772301 | SOX2 |
| 5377 | PDZD8 | SOX2 regulated genes in Keratinocyte Overexpression from PMID: 30772301 | SOX2 |
| 5378 | BACH2 | SOX2 regulated genes in Keratinocyte Overexpression from PMID: 30772301 | SOX2 |
| 5379 | AQP11 | SOX2 regulated genes in Keratinocyte Overexpression from PMID: 30772301 | SOX2 |
| 5380 | AP1S1 | SOX2 regulated genes in Keratinocyte Overexpression from PMID: 30772301 | SOX2 |
| 5381 | ANO3 | SOX2 regulated genes in Keratinocyte Overexpression from PMID: 30772301 | SOX2 |
| 5382 | CENPP | SOX2 regulated genes in Keratinocyte Overexpression from PMID: 30772301 | SOX2 |
| 5383 | POLR2M | SOX2 regulated genes in Keratinocyte Overexpression from PMID: 30772301 | SOX2 |
| 5384 | CEP44 | SOX2 regulated genes in Keratinocyte Overexpression from PMID: 30772301 | SOX2 |
| 5385 | ZNRF2 | SOX2 regulated genes in Keratinocyte Overexpression from PMID: 30772301 | SOX2 |
| 5386 | C9orf40 | SOX2 regulated genes in Keratinocyte Overexpression from PMID: 30772301 | SOX2 |
| 5387 | UBR7 | SOX2 regulated genes in Keratinocyte Overexpression from PMID: 30772301 | SOX2 |
| 5388 | TPD52L1 | SOX2 regulated genes in Keratinocyte Overexpression from PMID: 30772301 | SOX2 |
| 5389 | RSL1D1 | SOX2 regulated genes in Keratinocyte Overexpression from PMID: 30772301 | SOX2 |
| 5390 | LSM5 | SOX2 regulated genes in Keratinocyte Overexpression from PMID: 30772301 | SOX2 |
| 5391 | IRAK1BP1 | SOX2 regulated genes in Keratinocyte Overexpression from PMID: 30772301 | SOX2 |
| 5392 | ABT1 | SOX2 regulated genes in Keratinocyte Overexpression from PMID: 30772301 | SOX2 |
| 5393 | TSTD2 | SOX2 regulated genes in Keratinocyte Overexpression from PMID: 30772301 | SOX2 |
| 5394 | CLDN10 | SOX2 regulated genes in Keratinocyte Overexpression from PMID: 30772301 | SOX2 |
| 5395 | TOR2A | SOX2 regulated genes in Keratinocyte Overexpression from PMID: 30772301 | SOX2 |
| 5396 | MAPK1 | SOX2 regulated genes in Keratinocyte Overexpression from PMID: 30772301 | SOX2 |
| 5397 | VPS16 | SOX2 regulated genes in Keratinocyte Overexpression from PMID: 30772301 | SOX2 |
| 5398 | HAUS2 | SOX2 regulated genes in Keratinocyte Overexpression from PMID: 30772301 | SOX2 |
| 5399 | BUB3 | SOX2 regulated genes in Keratinocyte Overexpression from PMID: 30772301 | SOX2 |
| 5400 | PAK6 | SOX2 regulated genes in Keratinocyte Overexpression from PMID: 30772301 | SOX2 |
| 5401 | RCE1 | SOX2 regulated genes in Keratinocyte Overexpression from PMID: 30772301 | SOX2 |
| 5402 | YBX3 | SOX2 regulated genes in Keratinocyte Overexpression from PMID: 30772301 | SOX2 |
| 5403 | MORC2 | SOX2 regulated genes in Keratinocyte Overexpression from PMID: 30772301 | SOX2 |
| 5404 | PALM | SOX2 regulated genes in Keratinocyte Overexpression from PMID: 30772301 | SOX2 |
| 5405 | RHOD | SOX2 regulated genes in Keratinocyte Overexpression from PMID: 30772301 | SOX2 |
| 5406 | METTL14 | SOX2 regulated genes in Keratinocyte Overexpression from PMID: 30772301 | SOX2 |
| 5407 | NECAP1 | SOX2 regulated genes in Keratinocyte Overexpression from PMID: 30772301 | SOX2 |
| 5408 | AAR2 | SOX2 regulated genes in Keratinocyte Overexpression from PMID: 30772301 | SOX2 |
| 5409 | RBM26 | SOX2 regulated genes in Keratinocyte Overexpression from PMID: 30772301 | SOX2 |
| 5410 | UCHL3 | SOX2 regulated genes in Keratinocyte Overexpression from PMID: 30772301 | SOX2 |
| 5411 | BCL2L14 | SOX2 regulated genes in Keratinocyte Overexpression from PMID: 30772301 | SOX2 |
| 5412 | NONO | SOX2 regulated genes in Keratinocyte Overexpression from PMID: 30772301 | SOX2 |
| 5413 | USP14 | SOX2 regulated genes in Keratinocyte Overexpression from PMID: 30772301 | SOX2 |
| 5414 | EDC3 | SOX2 regulated genes in Keratinocyte Overexpression from PMID: 30772301 | SOX2 |
| 5415 | ETV4 | SOX2 regulated genes in Keratinocyte Overexpression from PMID: 30772301 | SOX2 |
| 5416 | NKX3-1 | SOX2 regulated genes in Keratinocyte Overexpression from PMID: 30772301 | SOX2 |
| 5417 | BCL2L1 | SOX2 regulated genes in Keratinocyte Overexpression from PMID: 30772301 | SOX2 |
| 5418 | MAGOHB | SOX2 regulated genes in Keratinocyte Overexpression from PMID: 30772301 | SOX2 |
| 5419 | UBE2K | SOX2 regulated genes in Keratinocyte Overexpression from PMID: 30772301 | SOX2 |
| 5420 | HARBI1 | SOX2 regulated genes in Keratinocyte Overexpression from PMID: 30772301 | SOX2 |
| 5421 | CAPN15 | SOX2 regulated genes in Keratinocyte Overexpression from PMID: 30772301 | SOX2 |
| 5422 | RPL23 | SOX2 regulated genes in Keratinocyte Overexpression from PMID: 30772301 | SOX2 |
| 5423 | LRP6 | SOX2 regulated genes in Keratinocyte Overexpression from PMID: 30772301 | SOX2 |
| 5424 | DSC1 | SOX2 regulated genes in Keratinocyte Overexpression from PMID: 30772301 | SOX2 |
| 5425 | PAPSS1 | SOX2 regulated genes in Keratinocyte Overexpression from PMID: 30772301 | SOX2 |
| 5426 | RBBP7 | SOX2 regulated genes in Keratinocyte Overexpression from PMID: 30772301 | SOX2 |
| 5427 | C7orf26 | SOX2 regulated genes in Keratinocyte Overexpression from PMID: 30772301 | SOX2 |

| | | | |
|---|---|---|---|
| 5428 | ID1 | SOX2 regulated genes in Keratinocyte Overexpression from PMID: 30772301 | SOX2 |
| 5429 | DSG2 | SOX2 regulated genes in Keratinocyte Overexpression from PMID: 30772301 | SOX2 |
| 5430 | MCM3 | SOX2 regulated genes in Keratinocyte Overexpression from PMID: 30772301 | SOX2 |
| 5431 | SREBF2 | SOX2 regulated genes in Keratinocyte Overexpression from PMID: 30772301 | SOX2 |
| 5432 | DNTTIP1 | SOX2 regulated genes in Keratinocyte Overexpression from PMID: 30772301 | SOX2 |
| 5433 | ZGPAT | SOX2 regulated genes in Keratinocyte Overexpression from PMID: 30772301 | SOX2 |
| 5434 | RBM39 | SOX2 regulated genes in Keratinocyte Overexpression from PMID: 30772301 | SOX2 |
| 5435 | NCOA5 | SOX2 regulated genes in Keratinocyte Overexpression from PMID: 30772301 | SOX2 |
| 5436 | POLR2F | SOX2 regulated genes in Keratinocyte Overexpression from PMID: 30772301 | SOX2 |
| 5437 | ITFG2 | SOX2 regulated genes in Keratinocyte Overexpression from PMID: 30772301 | SOX2 |
| 5438 | DUSP4 | SOX2 regulated genes in Keratinocyte Overexpression from PMID: 30772301 | SOX2 |
| 5439 | MCM8 | SOX2 regulated genes in Keratinocyte Overexpression from PMID: 30772301 | SOX2 |
| 5440 | RAB11FIP1 | SOX2 regulated genes in Keratinocyte Overexpression from PMID: 30772301 | SOX2 |
| 5441 | NEDD9 | SOX2 regulated genes in Keratinocyte Overexpression from PMID: 30772301 | SOX2 |
| 5442 | MIOS | SOX2 regulated genes in Keratinocyte Overexpression from PMID: 30772301 | SOX2 |
| 5443 | RUNX2 | SOX2 regulated genes in Keratinocyte Overexpression from PMID: 30772301 | SOX2 |
| 5444 | C10orf88 | SOX2 regulated genes in Keratinocyte Overexpression from PMID: 30772301 | SOX2 |
| 5445 | TRIM27 | SOX2 regulated genes in Keratinocyte Overexpression from PMID: 30772301 | SOX2 |
| 5446 | SOX21 | SOX2 regulated genes in Keratinocyte Overexpression from PMID: 30772301 | SOX2 |
| 5447 | GRIN3B | SOX2 regulated genes in Keratinocyte Overexpression from PMID: 30772301 | SOX2 |
| 5448 | NIPSNAP1 | SOX2 regulated genes in Keratinocyte Overexpression from PMID: 30772301 | SOX2 |
| 5449 | MRPL1 | SOX2 regulated genes in Keratinocyte Overexpression from PMID: 30772301 | SOX2 |
| 5450 | KDM5A | SOX2 regulated genes in Keratinocyte Overexpression from PMID: 30772301 | SOX2 |
| 5451 | PSMB5 | SOX2 regulated genes in Keratinocyte Overexpression from PMID: 30772301 | SOX2 |
| 5452 | RPA3 | SOX2 regulated genes in Keratinocyte Overexpression from PMID: 30772301 | SOX2 |
| 5453 | ASB6 | SOX2 regulated genes in Keratinocyte Overexpression from PMID: 30772301 | SOX2 |
| 5454 | CAAP1 | SOX2 regulated genes in Keratinocyte Overexpression from PMID: 30772301 | SOX2 |
| 5455 | TXNDC17 | SOX2 regulated genes in Keratinocyte Overexpression from PMID: 30772301 | SOX2 |
| 5456 | BRD3 | SOX2 regulated genes in Keratinocyte Overexpression from PMID: 30772301 | SOX2 |
| 5457 | IPPK | SOX2 regulated genes in Keratinocyte Overexpression from PMID: 30772301 | SOX2 |
| 5458 | TYMS | SOX2 regulated genes in Keratinocyte Overexpression from PMID: 30772301 | SOX2 |
| 5459 | NAPG | SOX2 regulated genes in Keratinocyte Overexpression from PMID: 30772301 | SOX2 |
| 5460 | ESCO1 | SOX2 regulated genes in Keratinocyte Overexpression from PMID: 30772301 | SOX2 |
| 5461 | THOC1 | SOX2 regulated genes in Keratinocyte Overexpression from PMID: 30772301 | SOX2 |
| 5462 | FAM186B | SOX2 regulated genes in Keratinocyte Overexpression from PMID: 30772301 | SOX2 |
| 5463 | STYK1 | SOX2 regulated genes in Keratinocyte Overexpression from PMID: 30772301 | SOX2 |
| 5464 | FAM98B | SOX2 regulated genes in Keratinocyte Overexpression from PMID: 30772301 | SOX2 |
| 5465 | SPECC1L | SOX2 regulated genes in Keratinocyte Overexpression from PMID: 30772301 | SOX2 |
| 5466 | DGCR2 | SOX2 regulated genes in Keratinocyte Overexpression from PMID: 30772301 | SOX2 |
| 5467 | SRGAP1 | SOX2 regulated genes in Keratinocyte Overexpression from PMID: 30772301 | SOX2 |
| 5468 | SLC38A2 | SOX2 regulated genes in Keratinocyte Overexpression from PMID: 30772301 | SOX2 |
| 5469 | PTAR1 | SOX2 regulated genes in Keratinocyte Overexpression from PMID: 30772301 | SOX2 |
| 5470 | MRPL41 | SOX2 regulated genes in Keratinocyte Overexpression from PMID: 30772301 | SOX2 |
| 5471 | GRID2 | SOX2 regulated genes in Keratinocyte Overexpression from PMID: 30772301 | SOX2 |
| 5472 | FLYWCH2 | SOX2 regulated genes in Keratinocyte Overexpression from PMID: 30772301 | SOX2 |
| 5473 | CENPQ | SOX2 regulated genes in Keratinocyte Overexpression from PMID: 30772301 | SOX2 |
| 5474 | NUDT16L1 | SOX2 regulated genes in Keratinocyte Overexpression from PMID: 30772301 | SOX2 |
| 5475 | NUPR2 | SOX2 regulated genes in Keratinocyte Overexpression from PMID: 30772301 | SOX2 |
| 5476 | ING2 | SOX2 regulated genes in Keratinocyte Overexpression from PMID: 30772301 | SOX2 |
| 5477 | TADA2B | SOX2 regulated genes in Keratinocyte Overexpression from PMID: 30772301 | SOX2 |
| 5478 | CPLX3 | SOX2 regulated genes in Keratinocyte Overexpression from PMID: 30772301 | SOX2 |
| 5479 | EIF6 | SOX2 regulated genes in Keratinocyte Overexpression from PMID: 30772301 | SOX2 |
| 5480 | CSE1L | SOX2 regulated genes in Keratinocyte Overexpression from PMID: 30772301 | SOX2 |
| 5481 | PRNP | SOX2 regulated genes in Keratinocyte Overexpression from PMID: 30772301 | SOX2 |
| 5482 | UROS | SOX2 regulated genes in Keratinocyte Overexpression from PMID: 30772301 | SOX2 |
| 5483 | CCDC120 | SOX2 regulated genes in Keratinocyte Overexpression from PMID: 30772301 | SOX2 |
| 5484 | FKBP4 | SOX2 regulated genes in Keratinocyte Overexpression from PMID: 30772301 | SOX2 |
| 5485 | ARHGAP23 | SOX2 regulated genes in Keratinocyte Overexpression from PMID: 30772301 | SOX2 |
| 5486 | CXCL6 | SOX2 regulated genes in Keratinocyte Overexpression from PMID: 30772301 | SOX2 |
| 5487 | TRABD | SOX2 regulated genes in Keratinocyte Overexpression from PMID: 30772301 | SOX2 |
| 5488 | RIOK1 | SOX2 regulated genes in Keratinocyte Overexpression from PMID: 30772301 | SOX2 |

| | | | |
|---|---|---|---|
| 5489 | HCFC1 | SOX2 regulated genes in Keratinocyte Overexpression from PMID: 30772301 | SOX2 |
| 5490 | SLC52A1 | SOX2 regulated genes in Keratinocyte Overexpression from PMID: 30772301 | SOX2 |
| 5491 | CRTAC1 | SOX2 regulated genes in Keratinocyte Overexpression from PMID: 30772301 | SOX2 |
| 5492 | APEX2 | SOX2 regulated genes in Keratinocyte Overexpression from PMID: 30772301 | SOX2 |
| 5493 | RASEF | SOX2 regulated genes in Keratinocyte Overexpression from PMID: 30772301 | SOX2 |
| 5494 | EGR3 | SOX2 regulated genes in Keratinocyte Overexpression from PMID: 30772301 | SOX2 |
| 5495 | TRIM29 | SOX2 regulated genes in Keratinocyte Overexpression from PMID: 30772301 | SOX2 |
| 5496 | DKK1 | SOX2 regulated genes in Keratinocyte Overexpression from PMID: 30772301 | SOX2 |
| 5497 | SAMD8 | SOX2 regulated genes in Keratinocyte Overexpression from PMID: 30772301 | SOX2 |
| 5498 | NUP58 | SOX2 regulated genes in Keratinocyte Overexpression from PMID: 30772301 | SOX2 |
| 5499 | SIVA1 | SOX2 regulated genes in Keratinocyte Overexpression from PMID: 30772301 | SOX2 |
| 5500 | LRFN4 | SOX2 regulated genes in Keratinocyte Overexpression from PMID: 30772301 | SOX2 |
| 5501 | UMODL1 | SOX2 regulated genes in Keratinocyte Overexpression from PMID: 30772301 | SOX2 |
| 5502 | POLR2E | SOX2 regulated genes in Keratinocyte Overexpression from PMID: 30772301 | SOX2 |
| 5503 | ACOT8 | SOX2 regulated genes in Keratinocyte Overexpression from PMID: 30772301 | SOX2 |
| 5504 | NEDD8 | SOX2 regulated genes in Keratinocyte Overexpression from PMID: 30772301 | SOX2 |
| 5505 | REM2 | SOX2 regulated genes in Keratinocyte Overexpression from PMID: 30772301 | SOX2 |
| 5506 | ZNF503 | SOX2 regulated genes in Keratinocyte Overexpression from PMID: 30772301 | SOX2 |
| 5507 | PRICKLE3 | SOX2 regulated genes in Keratinocyte Overexpression from PMID: 30772301 | SOX2 |
| 5508 | RHOT2 | SOX2 regulated genes in Keratinocyte Overexpression from PMID: 30772301 | SOX2 |
| 5509 | C15orf65 | SOX2 regulated genes in Keratinocyte Overexpression from PMID: 30772301 | SOX2 |
| 5510 | TFAM | SOX2 regulated genes in Keratinocyte Overexpression from PMID: 30772301 | SOX2 |
| 5511 | SLC26A8 | SOX2 regulated genes in Keratinocyte Overexpression from PMID: 30772301 | SOX2 |
| 5512 | PACSIN1 | SOX2 regulated genes in Keratinocyte Overexpression from PMID: 30772301 | SOX2 |
| 5513 | FANCF | SOX2 regulated genes in Keratinocyte Overexpression from PMID: 30772301 | SOX2 |
| 5514 | SH3RF1 | SOX2 regulated genes in Keratinocyte Overexpression from PMID: 30772301 | SOX2 |
| 5515 | OSBP | SOX2 regulated genes in Keratinocyte Overexpression from PMID: 30772301 | SOX2 |
| 5516 | NUP88 | SOX2 regulated genes in Keratinocyte Overexpression from PMID: 30772301 | SOX2 |
| 5517 | PCID2 | SOX2 regulated genes in Keratinocyte Overexpression from PMID: 30772301 | SOX2 |
| 5518 | SLCO4A1 | SOX2 regulated genes in Keratinocyte Overexpression from PMID: 30772301 | SOX2 |
| 5519 | LMX1B | SOX2 regulated genes in Keratinocyte Overexpression from PMID: 30772301 | SOX2 |
| 5520 | SMC2 | SOX2 regulated genes in Keratinocyte Overexpression from PMID: 30772301 | SOX2 |
| 5521 | FRMD8 | SOX2 regulated genes in Keratinocyte Overexpression from PMID: 30772301 | SOX2 |
| 5522 | CWH43 | SOX2 regulated genes in Keratinocyte Overexpression from PMID: 30772301 | SOX2 |
| 5523 | FNDC11 | SOX2 regulated genes in Keratinocyte Overexpression from PMID: 30772301 | SOX2 |
| 5524 | ABCB7 | SOX2 regulated genes in Keratinocyte Overexpression from PMID: 30772301 | SOX2 |
| 5525 | LPIN2 | SOX2 regulated genes in Keratinocyte Overexpression from PMID: 30772301 | SOX2 |
| 5526 | TMEM251 | SOX2 regulated genes in Keratinocyte Overexpression from PMID: 30772301 | SOX2 |
| 5527 | MIF | SOX2 regulated genes in Keratinocyte Overexpression from PMID: 30772301 | SOX2 |
| 5528 | MRPS2 | SOX2 regulated genes in Keratinocyte Overexpression from PMID: 30772301 | SOX2 |
| 5529 | TERT | SOX2 regulated genes in Keratinocyte Overexpression from PMID: 30772301 | SOX2 |
| 5530 | THNSL1 | SOX2 regulated genes in Keratinocyte Overexpression from PMID: 30772301 | SOX2 |
| 5531 | AK7 | SOX2 regulated genes in Keratinocyte Overexpression from PMID: 30772301 | SOX2 |
| 5532 | RBMX | SOX2 regulated genes in Keratinocyte Overexpression from PMID: 30772301 | SOX2 |
| 5533 | SLC30A9 | SOX2 regulated genes in Keratinocyte Overexpression from PMID: 30772301 | SOX2 |
| 5534 | CHIC2 | SOX2 regulated genes in Keratinocyte Overexpression from PMID: 30772301 | SOX2 |
| 5535 | SLC44A1 | SOX2 regulated genes in Keratinocyte Overexpression from PMID: 30772301 | SOX2 |
| 5536 | NGDN | SOX2 regulated genes in Keratinocyte Overexpression from PMID: 30772301 | SOX2 |
| 5537 | NEU3 | SOX2 regulated genes in Keratinocyte Overexpression from PMID: 30772301 | SOX2 |
| 5538 | DSG1 | SOX2 regulated genes in Keratinocyte Overexpression from PMID: 30772301 | SOX2 |
| 5539 | TAF1D | SOX2 regulated genes in Keratinocyte Overexpression from PMID: 30772301 | SOX2 |
| 5540 | TMEM144 | SOX2 regulated genes in Keratinocyte Overexpression from PMID: 30772301 | SOX2 |
| 5541 | NSFL1C | SOX2 regulated genes in Keratinocyte Overexpression from PMID: 30772301 | SOX2 |
| 5542 | TTC5 | SOX2 regulated genes in Keratinocyte Overexpression from PMID: 30772301 | SOX2 |
| 5543 | MRPL36 | SOX2 regulated genes in Keratinocyte Overexpression from PMID: 30772301 | SOX2 |
| 5544 | NCLN | SOX2 regulated genes in Keratinocyte Overexpression from PMID: 30772301 | SOX2 |
| 5545 | RBM20 | SOX2 regulated genes in Keratinocyte Overexpression from PMID: 30772301 | SOX2 |
| 5546 | FOXC1 | SOX2 regulated genes in Keratinocyte Overexpression from PMID: 30772301 | SOX2 |
| 5547 | URM1 | SOX2 regulated genes in Keratinocyte Overexpression from PMID: 30772301 | SOX2 |
| 5548 | TMEM255A | SOX2 regulated genes in Keratinocyte Overexpression from PMID: 30772301 | SOX2 |
| 5549 | PRPF19 | SOX2 regulated genes in Keratinocyte Overexpression from PMID: 30772301 | SOX2 |

| | | | |
|---|---|---|---|
| 5550 | CCDC116 | SOX2 regulated genes in Keratinocyte Overexpression from PMID: 30772301 | SOX2 |
| 5551 | RFXAP | SOX2 regulated genes in Keratinocyte Overexpression from PMID: 30772301 | SOX2 |
| 5552 | TDP1 | SOX2 regulated genes in Keratinocyte Overexpression from PMID: 30772301 | SOX2 |
| 5553 | LEPROTL1 | SOX2 regulated genes in Keratinocyte Overexpression from PMID: 30772301 | SOX2 |
| 5554 | RAB11FIP2 | SOX2 regulated genes in Keratinocyte Overexpression from PMID: 30772301 | SOX2 |
| 5555 | PDCL | SOX2 regulated genes in Keratinocyte Overexpression from PMID: 30772301 | SOX2 |
| 5556 | TDRD3 | SOX2 regulated genes in Keratinocyte Overexpression from PMID: 30772301 | SOX2 |
| 5557 | RND2 | SOX2 regulated genes in Keratinocyte Overexpression from PMID: 30772301 | SOX2 |
| 5558 | TRIM68 | SOX2 regulated genes in Keratinocyte Overexpression from PMID: 30772301 | SOX2 |
| 5559 | BMPER | SOX2 regulated genes in Keratinocyte Overexpression from PMID: 30772301 | SOX2 |
| 5560 | WFIKKN1 | SOX2 regulated genes in Keratinocyte Overexpression from PMID: 30772301 | SOX2 |
| 5561 | MAPK11 | SOX2 regulated genes in Keratinocyte Overexpression from PMID: 30772301 | SOX2 |
| 5562 | ZNF219 | SOX2 regulated genes in Keratinocyte Overexpression from PMID: 30772301 | SOX2 |
| 5563 | ARHGAP24 | SOX2 regulated genes in Keratinocyte Overexpression from PMID: 30772301 | SOX2 |
| 5564 | MUC15 | SOX2 regulated genes in Keratinocyte Overexpression from PMID: 30772301 | SOX2 |
| 5565 | BCORL1 | SOX2 regulated genes in Keratinocyte Overexpression from PMID: 30772301 | SOX2 |
| 5566 | NMRK1 | SOX2 regulated genes in Keratinocyte Overexpression from PMID: 30772301 | SOX2 |
| 5567 | EVC2 | SOX2 regulated genes in Keratinocyte Overexpression from PMID: 30772301 | SOX2 |
| 5568 | ARPC1A | SOX2 regulated genes in Keratinocyte Overexpression from PMID: 30772301 | SOX2 |
| 5569 | SHMT2 | SOX2 regulated genes in Keratinocyte Overexpression from PMID: 30772301 | SOX2 |
| 5570 | CDC5L | SOX2 regulated genes in Keratinocyte Overexpression from PMID: 30772301 | SOX2 |
| 5571 | TLE6 | SOX2 regulated genes in Keratinocyte Overexpression from PMID: 30772301 | SOX2 |
| 5572 | TEX9 | SOX2 regulated genes in Keratinocyte Overexpression from PMID: 30772301 | SOX2 |
| 5573 | PRPSAP2 | SOX2 regulated genes in Keratinocyte Overexpression from PMID: 30772301 | SOX2 |
| 5574 | ARGLU1 | SOX2 regulated genes in Keratinocyte Overexpression from PMID: 30772301 | SOX2 |
| 5575 | FASTKD3 | SOX2 regulated genes in Keratinocyte Overexpression from PMID: 30772301 | SOX2 |
| 5576 | SLTM | SOX2 regulated genes in Keratinocyte Overexpression from PMID: 30772301 | SOX2 |
| 5577 | PIM1 | SOX2 regulated genes in Keratinocyte Overexpression from PMID: 30772301 | SOX2 |
| 5578 | PLAA | SOX2 regulated genes in Keratinocyte Overexpression from PMID: 30772301 | SOX2 |
| 5579 | UBQLN1 | SOX2 regulated genes in Keratinocyte Overexpression from PMID: 30772301 | SOX2 |
| 5580 | ORMDL3 | SOX2 regulated genes in Keratinocyte Overexpression from PMID: 30772301 | SOX2 |
| 5581 | TBRG4 | SOX2 regulated genes in Keratinocyte Overexpression from PMID: 30772301 | SOX2 |
| 5582 | PORCN | SOX2 regulated genes in Keratinocyte Overexpression from PMID: 30772301 | SOX2 |
| 5583 | SNRPD1 | SOX2 regulated genes in Keratinocyte Overexpression from PMID: 30772301 | SOX2 |
| 5584 | PSMF1 | SOX2 regulated genes in Keratinocyte Overexpression from PMID: 30772301 | SOX2 |
| 5585 | MRPL50 | SOX2 regulated genes in Keratinocyte Overexpression from PMID: 30772301 | SOX2 |
| 5586 | SLC10A4 | SOX2 regulated genes in Keratinocyte Overexpression from PMID: 30772301 | SOX2 |
| 5587 | AIMP2 | SOX2 regulated genes in Keratinocyte Overexpression from PMID: 30772301 | SOX2 |
| 5588 | TRMT11 | SOX2 regulated genes in Keratinocyte Overexpression from PMID: 30772301 | SOX2 |
| 5589 | GATAD1 | SOX2 regulated genes in Keratinocyte Overexpression from PMID: 30772301 | SOX2 |
| 5590 | AARS2 | SOX2 regulated genes in Keratinocyte Overexpression from PMID: 30772301 | SOX2 |
| 5591 | RDM1 | SOX2 regulated genes in Keratinocyte Overexpression from PMID: 30772301 | SOX2 |
| 5592 | CBFA2T2 | SOX2 regulated genes in Keratinocyte Overexpression from PMID: 30772301 | SOX2 |
| 5593 | UBE2A | SOX2 regulated genes in Keratinocyte Overexpression from PMID: 30772301 | SOX2 |
| 5594 | C9orf85 | SOX2 regulated genes in Keratinocyte Overexpression from PMID: 30772301 | SOX2 |
| 5595 | COL4A5 | SOX2 regulated genes in Keratinocyte Overexpression from PMID: 30772301 | SOX2 |
| 5596 | GSDMA | SOX2 regulated genes in Keratinocyte Overexpression from PMID: 30772301 | SOX2 |
| 5597 | PRIM2 | SOX2 regulated genes in Keratinocyte Overexpression from PMID: 30772301 | SOX2 |
| 5598 | CBX6 | SOX2 regulated genes in Keratinocyte Overexpression from PMID: 30772301 | SOX2 |
| 5599 | RPGR | SOX2 regulated genes in Keratinocyte Overexpression from PMID: 30772301 | SOX2 |
| 5600 | P2RX5 | SOX2 regulated genes in Keratinocyte Overexpression from PMID: 30772301 | SOX2 |
| 5601 | NUDT15 | SOX2 regulated genes in Keratinocyte Overexpression from PMID: 30772301 | SOX2 |
| 5602 | PLK4 | SOX2 regulated genes in Keratinocyte Overexpression from PMID: 30772301 | SOX2 |
| 5603 | AXIN1 | SOX2 regulated genes in Keratinocyte Overexpression from PMID: 30772301 | SOX2 |
| 5604 | ARHGAP1 | SOX2 regulated genes in Keratinocyte Overexpression from PMID: 30772301 | SOX2 |
| 5605 | RPS24 | SOX2 regulated genes in Keratinocyte Overexpression from PMID: 30772301 | SOX2 |
| 5606 | BLMH | SOX2 regulated genes in Keratinocyte Overexpression from PMID: 30772301 | SOX2 |
| 5607 | TNC | SOX2 regulated genes in Keratinocyte Overexpression from PMID: 30772301 | SOX2 |
| 5608 | TUG1 | SOX2 regulated genes in Keratinocyte Overexpression from PMID: 30772301 | SOX2 |
| 5609 | NATD1 | SOX2 regulated genes in Keratinocyte Overexpression from PMID: 30772301 | SOX2 |
| 5610 | RNF20 | SOX2 regulated genes in Keratinocyte Overexpression from PMID: 30772301 | SOX2 |

| | | | |
|---|---|---|---|
| 5611 | HMG20A | SOX2 regulated genes in Keratinocyte Overexpression from PMID: 30772301 | SOX2 |
| 5612 | AVPI1 | SOX2 regulated genes in Keratinocyte Overexpression from PMID: 30772301 | SOX2 |
| 5613 | MRPL49 | SOX2 regulated genes in Keratinocyte Overexpression from PMID: 30772301 | SOX2 |
| 5614 | TSPAN9 | SOX2 regulated genes in Keratinocyte Overexpression from PMID: 30772301 | SOX2 |
| 5615 | MISP | SOX2 regulated genes in Keratinocyte Overexpression from PMID: 30772301 | SOX2 |
| 5616 | UPF3A | SOX2 regulated genes in Keratinocyte Overexpression from PMID: 30772301 | SOX2 |
| 5617 | TMTC1 | SOX2 regulated genes in Keratinocyte Overexpression from PMID: 30772301 | SOX2 |
| 5618 | ANLN | SOX2 regulated genes in Keratinocyte Overexpression from PMID: 30772301 | SOX2 |
| 5619 | PINX1 | SOX2 regulated genes in Keratinocyte Overexpression from PMID: 30772301 | SOX2 |
| 5620 | EXPH5 | SOX2 regulated genes in Keratinocyte Overexpression from PMID: 30772301 | SOX2 |
| 5621 | POLR3H | SOX2 regulated genes in Keratinocyte Overexpression from PMID: 30772301 | SOX2 |
| 5622 | ZSWIM1 | SOX2 regulated genes in Keratinocyte Overexpression from PMID: 30772301 | SOX2 |
| 5623 | SHPK | SOX2 regulated genes in Keratinocyte Overexpression from PMID: 30772301 | SOX2 |
| 5624 | RRP12 | SOX2 regulated genes in Keratinocyte Overexpression from PMID: 30772301 | SOX2 |
| 5625 | SNRPD3 | SOX2 regulated genes in Keratinocyte Overexpression from PMID: 30772301 | SOX2 |
| 5626 | LIME1 | SOX2 regulated genes in Keratinocyte Overexpression from PMID: 30772301 | SOX2 |
| 5627 | WIPI2 | SOX2 regulated genes in Keratinocyte Overexpression from PMID: 30772301 | SOX2 |
| 5628 | EFS | SOX2 regulated genes in Keratinocyte Overexpression from PMID: 30772301 | SOX2 |
| 5629 | DHRS1 | SOX2 regulated genes in Keratinocyte Overexpression from PMID: 30772301 | SOX2 |
| 5630 | EFHC1 | SOX2 regulated genes in Keratinocyte Overexpression from PMID: 30772301 | SOX2 |
| 5631 | TAB1 | SOX2 regulated genes in Keratinocyte Overexpression from PMID: 30772301 | SOX2 |
| 5632 | VAPA | SOX2 regulated genes in Keratinocyte Overexpression from PMID: 30772301 | SOX2 |
| 5633 | HSBP1L1 | SOX2 regulated genes in Keratinocyte Overexpression from PMID: 30772301 | SOX2 |
| 5634 | TMC5 | SOX2 regulated genes in Keratinocyte Overexpression from PMID: 30772301 | SOX2 |
| 5635 | CD2AP | SOX2 regulated genes in Keratinocyte Overexpression from PMID: 30772301 | SOX2 |
| 5636 | ROMO1 | SOX2 regulated genes in Keratinocyte Overexpression from PMID: 30772301 | SOX2 |
| 5637 | SLC15A1 | SOX2 regulated genes in Keratinocyte Overexpression from PMID: 30772301 | SOX2 |
| 5638 | XKR6 | SOX2 regulated genes in Keratinocyte Overexpression from PMID: 30772301 | SOX2 |
| 5639 | PAPPA | SOX2 regulated genes in Keratinocyte Overexpression from PMID: 30772301 | SOX2 |
| 5640 | KIF27 | SOX2 regulated genes in Keratinocyte Overexpression from PMID: 30772301 | SOX2 |
| 5641 | YAP1 | SOX2 regulated genes in Keratinocyte Overexpression from PMID: 30772301 | SOX2 |
| 5642 | R3HDM4 | SOX2 regulated genes in Keratinocyte Overexpression from PMID: 30772301 | SOX2 |
| 5643 | ZFAND2A | SOX2 regulated genes in Keratinocyte Overexpression from PMID: 30772301 | SOX2 |
| 5644 | SERGEF | SOX2 regulated genes in Keratinocyte Overexpression from PMID: 30772301 | SOX2 |
| 5645 | WDHD1 | SOX2 regulated genes in Keratinocyte Overexpression from PMID: 30772301 | SOX2 |
| 5646 | PALD1 | SOX2 regulated genes in Keratinocyte Overexpression from PMID: 30772301 | SOX2 |
| 5647 | RNF138 | SOX2 regulated genes in Keratinocyte Overexpression from PMID: 30772301 | SOX2 |
| 5648 | FRYL | SOX2 regulated genes in Keratinocyte Overexpression from PMID: 30772301 | SOX2 |
| 5649 | IPO8 | SOX2 regulated genes in Keratinocyte Overexpression from PMID: 30772301 | SOX2 |
| 5650 | SENP6 | SOX2 regulated genes in Keratinocyte Overexpression from PMID: 30772301 | SOX2 |
| 5651 | UTP14A | SOX2 regulated genes in Keratinocyte Overexpression from PMID: 30772301 | SOX2 |
| 5652 | PRPS1 | SOX2 regulated genes in Keratinocyte Overexpression from PMID: 30772301 | SOX2 |
| 5653 | RPH3AL | SOX2 regulated genes in Keratinocyte Overexpression from PMID: 30772301 | SOX2 |
| 5654 | CCAR1 | SOX2 regulated genes in Keratinocyte Overexpression from PMID: 30772301 | SOX2 |
| 5655 | INTS13 | SOX2 regulated genes in Keratinocyte Overexpression from PMID: 30772301 | SOX2 |
| 5656 | MTERF1 | SOX2 regulated genes in Keratinocyte Overexpression from PMID: 30772301 | SOX2 |
| 5657 | HDDC2 | SOX2 regulated genes in Keratinocyte Overexpression from PMID: 30772301 | SOX2 |
| 5658 | TCP11 | SOX2 regulated genes in Keratinocyte Overexpression from PMID: 30772301 | SOX2 |
| 5659 | ADNP | SOX2 regulated genes in Keratinocyte Overexpression from PMID: 30772301 | SOX2 |
| 5660 | SLC39A2 | SOX2 regulated genes in Keratinocyte Overexpression from PMID: 30772301 | SOX2 |
| 5661 | IPO4 | SOX2 regulated genes in Keratinocyte Overexpression from PMID: 30772301 | SOX2 |
| 5662 | FAM204A | SOX2 regulated genes in Keratinocyte Overexpression from PMID: 30772301 | SOX2 |
| 5663 | SDAD1 | SOX2 regulated genes in Keratinocyte Overexpression from PMID: 30772301 | SOX2 |
| 5664 | STK24 | SOX2 regulated genes in Keratinocyte Overexpression from PMID: 30772301 | SOX2 |
| 5665 | SLC34A2 | SOX2 regulated genes in Keratinocyte Overexpression from PMID: 30772301 | SOX2 |
| 5666 | RAP2C | SOX2 regulated genes in Keratinocyte Overexpression from PMID: 30772301 | SOX2 |
| 5667 | C7orf50 | SOX2 regulated genes in Keratinocyte Overexpression from PMID: 30772301 | SOX2 |
| 5668 | BARX1 | SOX2 regulated genes in Keratinocyte Overexpression from PMID: 30772301 | SOX2 |
| 5669 | ZDHHC16 | SOX2 regulated genes in Keratinocyte Overexpression from PMID: 30772301 | SOX2 |
| 5670 | PHF2 | SOX2 regulated genes in Keratinocyte Overexpression from PMID: 30772301 | SOX2 |
| 5671 | ARFGAP2 | SOX2 regulated genes in Keratinocyte Overexpression from PMID: 30772301 | SOX2 |

| | | | |
|---|---|---|---|
| 5672 | RPS27L | SOX2 regulated genes in Keratinocyte Overexpression from PMID: 30772301 | SOX2 |
| 5673 | PSMB7 | SOX2 regulated genes in Keratinocyte Overexpression from PMID: 30772301 | SOX2 |
| 5674 | FAU | SOX2 regulated genes in Keratinocyte Overexpression from PMID: 30772301 | SOX2 |
| 5675 | PRDM8 | SOX2 regulated genes in Keratinocyte Overexpression from PMID: 30772301 | SOX2 |
| 5676 | ANKRD50 | SOX2 regulated genes in Keratinocyte Overexpression from PMID: 30772301 | SOX2 |
| 5677 | ID4 | SOX2 regulated genes in Keratinocyte Overexpression from PMID: 30772301 | SOX2 |
| 5678 | SERPINB5 | SOX2 regulated genes in Keratinocyte Overexpression from PMID: 30772301 | SOX2 |
| 5679 | PARM1 | SOX2 regulated genes in Keratinocyte Overexpression from PMID: 30772301 | SOX2 |
| 5680 | RIBC1 | SOX2 regulated genes in Keratinocyte Overexpression from PMID: 30772301 | SOX2 |
| 5681 | MRPS31 | SOX2 regulated genes in Keratinocyte Overexpression from PMID: 30772301 | SOX2 |
| 5682 | IGF2BP3 | SOX2 regulated genes in Keratinocyte Overexpression from PMID: 30772301 | SOX2 |
| 5683 | U2AF1L5 | SOX2 regulated genes in Keratinocyte Overexpression from PMID: 30772301 | SOX2 |
| 5684 | TFAP2B | SOX2 regulated genes in Keratinocyte Overexpression from PMID: 30772301 | SOX2 |
| 5685 | E4F1 | SOX2 regulated genes in Keratinocyte Overexpression from PMID: 30772301 | SOX2 |
| 5686 | OPHN1 | SOX2 regulated genes in Keratinocyte Overexpression from PMID: 30772301 | SOX2 |
| 5687 | SOX12 | SOX2 regulated genes in Keratinocyte Overexpression from PMID: 30772301 | SOX2 |
| 5688 | CEP78 | SOX2 regulated genes in Keratinocyte Overexpression from PMID: 30772301 | SOX2 |
| 5689 | HOMEZ | SOX2 regulated genes in Keratinocyte Overexpression from PMID: 30772301 | SOX2 |
| 5690 | TRIL | SOX2 regulated genes in Keratinocyte Overexpression from PMID: 30772301 | SOX2 |
| 5691 | PEBP4 | SOX2 regulated genes in Keratinocyte Overexpression from PMID: 30772301 | SOX2 |
| 5692 | ARIH1 | SOX2 regulated genes in Keratinocyte Overexpression from PMID: 30772301 | SOX2 |
| 5693 | NKAP | SOX2 regulated genes in Keratinocyte Overexpression from PMID: 30772301 | SOX2 |
| 5694 | ASCC3 | SOX2 regulated genes in Keratinocyte Overexpression from PMID: 30772301 | SOX2 |
| 5695 | GPKOW | SOX2 regulated genes in Keratinocyte Overexpression from PMID: 30772301 | SOX2 |
| 5696 | BORCS5 | SOX2 regulated genes in Keratinocyte Overexpression from PMID: 30772301 | SOX2 |
| 5697 | SEC14L2 | SOX2 regulated genes in Keratinocyte Overexpression from PMID: 30772301 | SOX2 |
| 5698 | CCDC169 | SOX2 regulated genes in Keratinocyte Overexpression from PMID: 30772301 | SOX2 |
| 5699 | RWDD4 | SOX2 regulated genes in Keratinocyte Overexpression from PMID: 30772301 | SOX2 |
| 5700 | AFG3L2 | SOX2 regulated genes in Keratinocyte Overexpression from PMID: 30772301 | SOX2 |
| 5701 | UXT | SOX2 regulated genes in Keratinocyte Overexpression from PMID: 30772301 | SOX2 |
| 5702 | ZNHIT2 | SOX2 regulated genes in Keratinocyte Overexpression from PMID: 30772301 | SOX2 |
| 5703 | HSPA4L | SOX2 regulated genes in Keratinocyte Overexpression from PMID: 30772301 | SOX2 |
| 5704 | GON7 | SOX2 regulated genes in Keratinocyte Overexpression from PMID: 30772301 | SOX2 |
| 5705 | PCNA | SOX2 regulated genes in Keratinocyte Overexpression from PMID: 30772301 | SOX2 |
| 5706 | ENDOU | SOX2 regulated genes in Keratinocyte Overexpression from PMID: 30772301 | SOX2 |
| 5707 | AGR2 | SOX2 regulated genes in Keratinocyte Overexpression from PMID: 30772301 | SOX2 |
| 5708 | EIF2AK4 | SOX2 regulated genes in Keratinocyte Overexpression from PMID: 30772301 | SOX2 |
| 5709 | RTCB | SOX2 regulated genes in Keratinocyte Overexpression from PMID: 30772301 | SOX2 |
| 5710 | RPP40 | SOX2 regulated genes in Keratinocyte Overexpression from PMID: 30772301 | SOX2 |
| 5711 | KIAA1217 | SOX2 regulated genes in Keratinocyte Overexpression from PMID: 30772301 | SOX2 |
| 5712 | CDCA7L | SOX2 regulated genes in Keratinocyte Overexpression from PMID: 30772301 | SOX2 |
| 5713 | SCD | SOX2 regulated genes in Keratinocyte Overexpression from PMID: 30772301 | SOX2 |
| 5714 | CDC42EP2 | SOX2 regulated genes in Keratinocyte Overexpression from PMID: 30772301 | SOX2 |
| 5715 | FOXO1 | SOX2 regulated genes in Keratinocyte Overexpression from PMID: 30772301 | SOX2 |
| 5716 | ONECUT3 | SOX2 regulated genes in Keratinocyte Overexpression from PMID: 30772301 | SOX2 |
| 5717 | TCFL5 | SOX2 regulated genes in Keratinocyte Overexpression from PMID: 30772301 | SOX2 |
| 5718 | RPL35 | SOX2 regulated genes in Keratinocyte Overexpression from PMID: 30772301 | SOX2 |
| 5719 | HACE1 | SOX2 regulated genes in Keratinocyte Overexpression from PMID: 30772301 | SOX2 |
| 5720 | QSOX2 | SOX2 regulated genes in Keratinocyte Overexpression from PMID: 30772301 | SOX2 |
| 5721 | HNRNPD | SOX2 regulated genes in Keratinocyte Overexpression from PMID: 30772301 | SOX2 |
| 5722 | PLXNA3 | SOX2 regulated genes in Keratinocyte Overexpression from PMID: 30772301 | SOX2 |
| 5723 | UQCR10 | SOX2 regulated genes in Keratinocyte Overexpression from PMID: 30772301 | SOX2 |
| 5724 | AASDHPPT | SOX2 regulated genes in Keratinocyte Overexpression from PMID: 30772301 | SOX2 |
| 5725 | TXNL4A | SOX2 regulated genes in Keratinocyte Overexpression from PMID: 30772301 | SOX2 |
| 5726 | CEP295 | SOX2 regulated genes in Keratinocyte Overexpression from PMID: 30772301 | SOX2 |
| 5727 | ZNF280C | SOX2 regulated genes in Keratinocyte Overexpression from PMID: 30772301 | SOX2 |
| 5728 | TAX1BP3 | SOX2 regulated genes in Keratinocyte Overexpression from PMID: 30772301 | SOX2 |
| 5729 | ELP4 | SOX2 regulated genes in Keratinocyte Overexpression from PMID: 30772301 | SOX2 |
| 5730 | RPUSD2 | SOX2 regulated genes in Keratinocyte Overexpression from PMID: 30772301 | SOX2 |
| 5731 | EIF3A | SOX2 regulated genes in Keratinocyte Overexpression from PMID: 30772301 | SOX2 |
| 5732 | TSPAN14 | SOX2 regulated genes in Keratinocyte Overexpression from PMID: 30772301 | SOX2 |

| | | | |
|---|---|---|---|
| 5733 | HMGXB4 | SOX2 regulated genes in Keratinocyte Overexpression from PMID: 30772301 | SOX2 |
| 5734 | YTHDF1 | SOX2 regulated genes in Keratinocyte Overexpression from PMID: 30772301 | SOX2 |
| 5735 | SPEF1 | SOX2 regulated genes in Keratinocyte Overexpression from PMID: 30772301 | SOX2 |
| 5736 | JARID2 | SOX2 regulated genes in Keratinocyte Overexpression from PMID: 30772301 | SOX2 |
| 5737 | TXN | SOX2 regulated genes in Keratinocyte Overexpression from PMID: 30772301 | SOX2 |
| 5738 | OAT | SOX2 regulated genes in Keratinocyte Overexpression from PMID: 30772301 | SOX2 |
| 5739 | PRMT9 | SOX2 regulated genes in Keratinocyte Overexpression from PMID: 30772301 | SOX2 |
| 5740 | ATP12A | SOX2 regulated genes in Keratinocyte Overexpression from PMID: 30772301 | SOX2 |
| 5741 | OSER1 | SOX2 regulated genes in Keratinocyte Overexpression from PMID: 30772301 | SOX2 |
| 5742 | ZNF335 | SOX2 regulated genes in Keratinocyte Overexpression from PMID: 30772301 | SOX2 |
| 5743 | ZSWIM3 | SOX2 regulated genes in Keratinocyte Overexpression from PMID: 30772301 | SOX2 |
| 5744 | SFXN2 | SOX2 regulated genes in Keratinocyte Overexpression from PMID: 30772301 | SOX2 |
| 5745 | ZFAND5 | SOX2 regulated genes in Keratinocyte Overexpression from PMID: 30772301 | SOX2 |
| 5746 | ETNPPL | SOX2 regulated genes in Keratinocyte Overexpression from PMID: 30772301 | SOX2 |
| 5747 | RGP1 | SOX2 regulated genes in Keratinocyte Overexpression from PMID: 30772301 | SOX2 |
| 5748 | TRUB2 | SOX2 regulated genes in Keratinocyte Overexpression from PMID: 30772301 | SOX2 |
| 5749 | RCAN1 | SOX2 regulated genes in Keratinocyte Overexpression from PMID: 30772301 | SOX2 |
| 5750 | C11orf58 | SOX2 regulated genes in Keratinocyte Overexpression from PMID: 30772301 | SOX2 |
| 5751 | MAK16 | SOX2 regulated genes in Keratinocyte Overexpression from PMID: 30772301 | SOX2 |
| 5752 | TNFRSF13C | SOX2 regulated genes in Keratinocyte Overexpression from PMID: 30772301 | SOX2 |
| 5753 | ZNF330 | SOX2 regulated genes in Keratinocyte Overexpression from PMID: 30772301 | SOX2 |
| 5754 | FPGS | SOX2 regulated genes in Keratinocyte Overexpression from PMID: 30772301 | SOX2 |
| 5755 | DHX33 | SOX2 regulated genes in Keratinocyte Overexpression from PMID: 30772301 | SOX2 |
| 5756 | SAC3D1 | SOX2 regulated genes in Keratinocyte Overexpression from PMID: 30772301 | SOX2 |
| 5757 | RAB40C | SOX2 regulated genes in Keratinocyte Overexpression from PMID: 30772301 | SOX2 |
| 5758 | RFLNB | SOX2 regulated genes in Keratinocyte Overexpression from PMID: 30772301 | SOX2 |
| 5759 | MACROD2 | SOX2 regulated genes in Keratinocyte Overexpression from PMID: 30772301 | SOX2 |
| 5760 | VNN1 | SOX2 regulated genes in Keratinocyte Overexpression from PMID: 30772301 | SOX2 |
| 5761 | HIPK3 | SOX2 regulated genes in Keratinocyte Overexpression from PMID: 30772301 | SOX2 |
| 5762 | ARHGEF39 | SOX2 regulated genes in Keratinocyte Overexpression from PMID: 30772301 | SOX2 |
| 5763 | PGAP3 | SOX2 regulated genes in Keratinocyte Overexpression from PMID: 30772301 | SOX2 |
| 5764 | CCKBR | SOX2 regulated genes in Keratinocyte Overexpression from PMID: 30772301 | SOX2 |
| 5765 | BMS1 | SOX2 regulated genes in Keratinocyte Overexpression from PMID: 30772301 | SOX2 |
| 5766 | MED31 | SOX2 regulated genes in Keratinocyte Overexpression from PMID: 30772301 | SOX2 |
| 5767 | DCDC2 | SOX2 regulated genes in Keratinocyte Overexpression from PMID: 30772301 | SOX2 |
| 5768 | XRCC6 | SOX2 regulated genes in Keratinocyte Overexpression from PMID: 30772301 | SOX2 |
| 5769 | EGFL6 | SOX2 regulated genes in Keratinocyte Overexpression from PMID: 30772301 | SOX2 |
| 5770 | KCTD9 | SOX2 regulated genes in Keratinocyte Overexpression from PMID: 30772301 | SOX2 |
| 5771 | SNX12 | SOX2 regulated genes in Keratinocyte Overexpression from PMID: 30772301 | SOX2 |
| 5772 | NT5C3B | SOX2 regulated genes in Keratinocyte Overexpression from PMID: 30772301 | SOX2 |
| 5773 | BCCIP | SOX2 regulated genes in Keratinocyte Overexpression from PMID: 30772301 | SOX2 |
| 5774 | STUB1 | SOX2 regulated genes in Keratinocyte Overexpression from PMID: 30772301 | SOX2 |
| 5775 | INIP | SOX2 regulated genes in Keratinocyte Overexpression from PMID: 30772301 | SOX2 |
| 5776 | DDX3X | SOX2 regulated genes in Keratinocyte Overexpression from PMID: 30772301 | SOX2 |
| 5777 | ANKRD10 | SOX2 regulated genes in Keratinocyte Overexpression from PMID: 30772301 | SOX2 |
| 5778 | EHMT1 | SOX2 regulated genes in Keratinocyte Overexpression from PMID: 30772301 | SOX2 |
| 5779 | DICER1 | SOX2 regulated genes in Keratinocyte Overexpression from PMID: 30772301 | SOX2 |
| 5780 | MBOAT1 | SOX2 regulated genes in Keratinocyte Overexpression from PMID: 30772301 | SOX2 |
| 5781 | NR6A1 | SOX2 regulated genes in Keratinocyte Overexpression from PMID: 30772301 | SOX2 |
| 5782 | MDN1 | SOX2 regulated genes in Keratinocyte Overexpression from PMID: 30772301 | SOX2 |
| 5783 | CREB5 | SOX2 regulated genes in Keratinocyte Overexpression from PMID: 30772301 | SOX2 |
| 5784 | S1PR3 | SOX2 regulated genes in Keratinocyte Overexpression from PMID: 30772301 | SOX2 |
| 5785 | ASCC1 | SOX2 regulated genes in Keratinocyte Overexpression from PMID: 30772301 | SOX2 |
| 5786 | SOGA3 | SOX2 regulated genes in Keratinocyte Overexpression from PMID: 30772301 | SOX2 |
| 5787 | ARHGAP42 | SOX2 regulated genes in Keratinocyte Overexpression from PMID: 30772301 | SOX2 |
| 5788 | KIF6 | SOX2 regulated genes in Keratinocyte Overexpression from PMID: 30772301 | SOX2 |
| 5789 | GNB1L | SOX2 regulated genes in Keratinocyte Overexpression from PMID: 30772301 | SOX2 |
| 5790 | ALDH8A1 | SOX2 regulated genes in Keratinocyte Overexpression from PMID: 30772301 | SOX2 |
| 5791 | CDKAL1 | SOX2 regulated genes in Keratinocyte Overexpression from PMID: 30772301 | SOX2 |
| 5792 | SLC35F2 | SOX2 regulated genes in Keratinocyte Overexpression from PMID: 30772301 | SOX2 |
| 5793 | MID2 | SOX2 regulated genes in Keratinocyte Overexpression from PMID: 30772301 | SOX2 |

| | | | |
|---|---|---|---|
| 5794 | EBF4 | SOX2 regulated genes in Keratinocyte Overexpression from PMID: 30772301 | SOX2 |
| 5795 | UFD1 | SOX2 regulated genes in Keratinocyte Overexpression from PMID: 30772301 | SOX2 |
| 5796 | NUDT3 | SOX2 regulated genes in Keratinocyte Overexpression from PMID: 30772301 | SOX2 |
| 5797 | TRMT2A | SOX2 regulated genes in Keratinocyte Overexpression from PMID: 30772301 | SOX2 |
| 5798 | ZC3H14 | SOX2 regulated genes in Keratinocyte Overexpression from PMID: 30772301 | SOX2 |
| 5799 | ARFGEF2 | SOX2 regulated genes in Keratinocyte Overexpression from PMID: 30772301 | SOX2 |
| 5800 | COX4I2 | SOX2 regulated genes in Keratinocyte Overexpression from PMID: 30772301 | SOX2 |
| 5801 | SALL3 | SOX2 regulated genes in Keratinocyte Overexpression from PMID: 30772301 | SOX2 |
| 5802 | EFNB2 | SOX2 regulated genes in Keratinocyte Overexpression from PMID: 30772301 | SOX2 |
| 5803 | PMS2 | SOX2 regulated genes in Keratinocyte Overexpression from PMID: 30772301 | SOX2 |
| 5804 | SCLT1 | SOX2 regulated genes in Keratinocyte Overexpression from PMID: 30772301 | SOX2 |
| 5805 | TTF1 | SOX2 regulated genes in Keratinocyte Overexpression from PMID: 30772301 | SOX2 |
| 5806 | ITPRIP | SOX2 regulated genes in Keratinocyte Overexpression from PMID: 30772301 | SOX2 |
| 5807 | DSN1 | SOX2 regulated genes in Keratinocyte Overexpression from PMID: 30772301 | SOX2 |
| 5808 | CPNE8 | SOX2 regulated genes in Keratinocyte Overexpression from PMID: 30772301 | SOX2 |
| 5809 | RPAP3 | SOX2 regulated genes in Keratinocyte Overexpression from PMID: 30772301 | SOX2 |
| 5810 | CISD2 | SOX2 regulated genes in Keratinocyte Overexpression from PMID: 30772301 | SOX2 |
| 5811 | INTS1 | SOX2 regulated genes in Keratinocyte Overexpression from PMID: 30772301 | SOX2 |
| 5812 | METTL4 | SOX2 regulated genes in Keratinocyte Overexpression from PMID: 30772301 | SOX2 |
| 5813 | TMPRSS11F | SOX2 regulated genes in Keratinocyte Overexpression from PMID: 30772301 | SOX2 |
| 5814 | BNIP3L | SOX2 regulated genes in Keratinocyte Overexpression from PMID: 30772301 | SOX2 |
| 5815 | ADGRF1 | SOX2 regulated genes in Keratinocyte Overexpression from PMID: 30772301 | SOX2 |
| 5816 | SPRYD3 | SOX2 regulated genes in Keratinocyte Overexpression from PMID: 30772301 | SOX2 |
| 5817 | MRPL16 | SOX2 regulated genes in Keratinocyte Overexpression from PMID: 30772301 | SOX2 |
| 5818 | FANCC | SOX2 regulated genes in Keratinocyte Overexpression from PMID: 30772301 | SOX2 |
| 5819 | PLP2 | SOX2 regulated genes in Keratinocyte Overexpression from PMID: 30772301 | SOX2 |
| 5820 | WDR24 | SOX2 regulated genes in Keratinocyte Overexpression from PMID: 30772301 | SOX2 |
| 5821 | TCF7L2 | SOX2 regulated genes in Keratinocyte Overexpression from PMID: 30772301 | SOX2 |
| 5822 | LRP10 | SOX2 regulated genes in Keratinocyte Overexpression from PMID: 30772301 | SOX2 |
| 5823 | CRELD2 | SOX2 regulated genes in Keratinocyte Overexpression from PMID: 30772301 | SOX2 |
| 5824 | ABRACL | SOX2 regulated genes in Keratinocyte Overexpression from PMID: 30772301 | SOX2 |
| 5825 | NOL8 | SOX2 regulated genes in Keratinocyte Overexpression from PMID: 30772301 | SOX2 |
| 5826 | MTMR3 | SOX2 regulated genes in Keratinocyte Overexpression from PMID: 30772301 | SOX2 |
| 5827 | HHIP | SOX2 regulated genes in Keratinocyte Overexpression from PMID: 30772301 | SOX2 |
| 5828 | HAP1 | SOX2 regulated genes in Keratinocyte Overexpression from PMID: 30772301 | SOX2 |
| 5829 | CHKA | SOX2 regulated genes in Keratinocyte Overexpression from PMID: 30772301 | SOX2 |
| 5830 | CYP46A1 | SOX2 regulated genes in Keratinocyte Overexpression from PMID: 30772301 | SOX2 |
| 5831 | MBD2 | SOX2 regulated genes in Keratinocyte Overexpression from PMID: 30772301 | SOX2 |
| 5832 | TMEM98 | SOX2 regulated genes in Keratinocyte Overexpression from PMID: 30772301 | SOX2 |
| 5833 | FERMT1 | SOX2 regulated genes in Keratinocyte Overexpression from PMID: 30772301 | SOX2 |
| 5834 | SMCHD1 | SOX2 regulated genes in Keratinocyte Overexpression from PMID: 30772301 | SOX2 |
| 5835 | PLXNB2 | SOX2 regulated genes in Keratinocyte Overexpression from PMID: 30772301 | SOX2 |
| 5836 | MAD2L1 | SOX2 regulated genes in Keratinocyte Overexpression from PMID: 30772301 | SOX2 |
| 5837 | VTA1 | SOX2 regulated genes in Keratinocyte Overexpression from PMID: 30772301 | SOX2 |
| 5838 | CARNMT1 | SOX2 regulated genes in Keratinocyte Overexpression from PMID: 30772301 | SOX2 |
| 5839 | CELSR1 | SOX2 regulated genes in Keratinocyte Overexpression from PMID: 30772301 | SOX2 |
| 5840 | C5orf49 | SOX2 regulated genes in Keratinocyte Overexpression from PMID: 30772301 | SOX2 |
| 5841 | CDKL2 | SOX2 regulated genes in Keratinocyte Overexpression from PMID: 30772301 | SOX2 |
| 5842 | CNOT7 | SOX2 regulated genes in Keratinocyte Overexpression from PMID: 30772301 | SOX2 |
| 5843 | DSC3 | SOX2 regulated genes in Keratinocyte Overexpression from PMID: 30772301 | SOX2 |
| 5844 | BUD31 | SOX2 regulated genes in Keratinocyte Overexpression from PMID: 30772301 | SOX2 |
| 5845 | HDAC2 | SOX2 regulated genes in Keratinocyte Overexpression from PMID: 30772301 | SOX2 |
| 5846 | C20orf27 | SOX2 regulated genes in Keratinocyte Overexpression from PMID: 30772301 | SOX2 |
| 5847 | PITPNM3 | SOX2 regulated genes in Keratinocyte Overexpression from PMID: 30772301 | SOX2 |
| 5848 | SUZ12 | SOX2 regulated genes in Keratinocyte Overexpression from PMID: 30772301 | SOX2 |
| 5849 | TUBA8 | SOX2 regulated genes in Keratinocyte Overexpression from PMID: 30772301 | SOX2 |
| 5850 | ZCCHC3 | SOX2 regulated genes in Keratinocyte Overexpression from PMID: 30772301 | SOX2 |
| 5851 | DMRT1 | SOX2 regulated genes in Keratinocyte Overexpression from PMID: 30772301 | SOX2 |
| 5852 | RPS19BP1 | SOX2 regulated genes in Keratinocyte Overexpression from PMID: 30772301 | SOX2 |
| 5853 | OSBPL2 | SOX2 regulated genes in Keratinocyte Overexpression from PMID: 30772301 | SOX2 |
| 5854 | FABP7 | SOX2 regulated genes in Keratinocyte Overexpression from PMID: 30772301 | SOX2 |

| | | | |
|---|---|---|---|
| 5855 | PGP | SOX2 regulated genes in Keratinocyte Overexpression from PMID: 30772301 | SOX2 |
| 5856 | KRT6B | SOX2 regulated genes in Keratinocyte Overexpression from PMID: 30772301 | SOX2 |
| 5857 | KRT6C | SOX2 regulated genes in Keratinocyte Overexpression from PMID: 30772301 | SOX2 |
| 5858 | CPXM2 | SOX2 regulated genes in Keratinocyte Overexpression from PMID: 30772301 | SOX2 |
| 5859 | SCML4 | SOX2 regulated genes in Keratinocyte Overexpression from PMID: 30772301 | SOX2 |
| 5860 | PARD6G | SOX2 regulated genes in Keratinocyte Overexpression from PMID: 30772301 | SOX2 |
| 5861 | ZNF341 | SOX2 regulated genes in Keratinocyte Overexpression from PMID: 30772301 | SOX2 |
| 5862 | AFAP1L2 | SOX2 regulated genes in Keratinocyte Overexpression from PMID: 30772301 | SOX2 |
| 5863 | PIGW | SOX2 regulated genes in Keratinocyte Overexpression from PMID: 30772301 | SOX2 |
| 5864 | ASNS | SOX2 regulated genes in Keratinocyte Overexpression from PMID: 30772301 | SOX2 |
| 5865 | DNAAF5 | SOX2 regulated genes in Keratinocyte Overexpression from PMID: 30772301 | SOX2 |
| 5866 | GRB7 | SOX2 regulated genes in Keratinocyte Overexpression from PMID: 30772301 | SOX2 |
| 5867 | FKBP1A | SOX2 regulated genes in Keratinocyte Overexpression from PMID: 30772301 | SOX2 |
| 5868 | FKBP1C | SOX2 regulated genes in Keratinocyte Overexpression from PMID: 30772301 | SOX2 |
| 5869 | PRRG4 | SOX2 regulated genes in Keratinocyte Overexpression from PMID: 30772301 | SOX2 |
| 5870 | GBA2 | SOX2 regulated genes in Keratinocyte Overexpression from PMID: 30772301 | SOX2 |
| 5871 | CUL4B | SOX2 regulated genes in Keratinocyte Overexpression from PMID: 30772301 | SOX2 |
| 5872 | TFIP11 | SOX2 regulated genes in Keratinocyte Overexpression from PMID: 30772301 | SOX2 |
| 5873 | PLEKHS1 | SOX2 regulated genes in Keratinocyte Overexpression from PMID: 30772301 | SOX2 |
| 5874 | GOPC | SOX2 regulated genes in Keratinocyte Overexpression from PMID: 30772301 | SOX2 |
| 5875 | KDM2A | SOX2 regulated genes in Keratinocyte Overexpression from PMID: 30772301 | SOX2 |
| 5876 | SPIN1 | SOX2 regulated genes in Keratinocyte Overexpression from PMID: 30772301 | SOX2 |
| 5877 | PON2 | SOX2 regulated genes in Keratinocyte Overexpression from PMID: 30772301 | SOX2 |
| 5878 | MRPS17 | SOX2 regulated genes in Keratinocyte Overexpression from PMID: 30772301 | SOX2 |
| 5879 | KIAA0753 | SOX2 regulated genes in Keratinocyte Overexpression from PMID: 30772301 | SOX2 |
| 5880 | TSSC4 | SOX2 regulated genes in Keratinocyte Overexpression from PMID: 30772301 | SOX2 |
| 5881 | USP22 | SOX2 regulated genes in Keratinocyte Overexpression from PMID: 30772301 | SOX2 |
| 5882 | UBL4A | SOX2 regulated genes in Keratinocyte Overexpression from PMID: 30772301 | SOX2 |
| 5883 | CSPG4 | SOX2 regulated genes in Keratinocyte Overexpression from PMID: 30772301 | SOX2 |
| 5884 | FANCI | SOX2 regulated genes in Keratinocyte Overexpression from PMID: 30772301 | SOX2 |
| 5885 | PERP | SOX2 regulated genes in Keratinocyte Overexpression from PMID: 30772301 | SOX2 |
| 5886 | TP53RK | SOX2 regulated genes in Keratinocyte Overexpression from PMID: 30772301 | SOX2 |
| 5887 | COX8A | SOX2 regulated genes in Keratinocyte Overexpression from PMID: 30772301 | SOX2 |
| 5888 | TIMMDC1 | SOX2 regulated genes in Keratinocyte Overexpression from PMID: 30772301 | SOX2 |
| 5889 | BMP2K | SOX2 regulated genes in Keratinocyte Overexpression from PMID: 30772301 | SOX2 |
| 5890 | ACTR5 | SOX2 regulated genes in Keratinocyte Overexpression from PMID: 30772301 | SOX2 |
| 5891 | WDR59 | SOX2 regulated genes in Keratinocyte Overexpression from PMID: 30772301 | SOX2 |
| 5892 | POLR2K | SOX2 regulated genes in Keratinocyte Overexpression from PMID: 30772301 | SOX2 |
| 5893 | WRAP53 | SOX2 regulated genes in Keratinocyte Overexpression from PMID: 30772301 | SOX2 |
| 5894 | APOM | SOX2 regulated genes in Keratinocyte Overexpression from PMID: 30772301 | SOX2 |
| 5895 | NELFB | SOX2 regulated genes in Keratinocyte Overexpression from PMID: 30772301 | SOX2 |
| 5896 | KIF18A | SOX2 regulated genes in Keratinocyte Overexpression from PMID: 30772301 | SOX2 |
| 5897 | ZNF274 | SOX2 regulated genes in Keratinocyte Overexpression from PMID: 30772301 | SOX2 |
| 5898 | PNN | SOX2 regulated genes in Keratinocyte Overexpression from PMID: 30772301 | SOX2 |
| 5899 | GABRQ | SOX2 regulated genes in Keratinocyte Overexpression from PMID: 30772301 | SOX2 |
| 5900 | UBE2E1 | SOX2 regulated genes in Keratinocyte Overexpression from PMID: 30772301 | SOX2 |
| 5901 | MIA-RAB4B | SOX2 regulated genes in Keratinocyte Overexpression from PMID: 30772301 | SOX2 |
| 5902 | MIA | SOX2 regulated genes in Keratinocyte Overexpression from PMID: 30772301 | SOX2 |
| 5903 | RNF214 | SOX2 regulated genes in Keratinocyte Overexpression from PMID: 30772301 | SOX2 |
| 5904 | SERPIND1 | SOX2 regulated genes in Keratinocyte Overexpression from PMID: 30772301 | SOX2 |
| 5905 | ZNF426 | SOX2 regulated genes in Keratinocyte Overexpression from PMID: 30772301 | SOX2 |
| 5906 | DXO | SOX2 regulated genes in Keratinocyte Overexpression from PMID: 30772301 | SOX2 |
| 5907 | CD82 | SOX2 regulated genes in Keratinocyte Overexpression from PMID: 30772301 | SOX2 |
| 5908 | ITPKC | SOX2 regulated genes in Keratinocyte Overexpression from PMID: 30772301 | SOX2 |
| 5909 | NXPE4 | SOX2 regulated genes in Keratinocyte Overexpression from PMID: 30772301 | SOX2 |
| 5910 | TRAPPC11 | SOX2 regulated genes in Keratinocyte Overexpression from PMID: 30772301 | SOX2 |
| 5911 | CST6 | SOX2 regulated genes in Keratinocyte Overexpression from PMID: 30772301 | SOX2 |
| 5912 | SYMPK | SOX2 regulated genes in Keratinocyte Overexpression from PMID: 30772301 | SOX2 |
| 5913 | RWDD1 | SOX2 regulated genes in Keratinocyte Overexpression from PMID: 30772301 | SOX2 |
| 5914 | CHCHD6 | SOX2 regulated genes in Keratinocyte Overexpression from PMID: 30772301 | SOX2 |
| 5915 | YRDC | SOX2 regulated genes in Keratinocyte Overexpression from PMID: 30772301 | SOX2 |

| | | | |
|---|---|---|---|
| 5916 | FBXO30 | SOX2 regulated genes in Keratinocyte Overexpression from PMID: 30772301 | SOX2 |
| 5917 | POLR3B | SOX2 regulated genes in Keratinocyte Overexpression from PMID: 30772301 | SOX2 |
| 5918 | CLPX | SOX2 regulated genes in Keratinocyte Overexpression from PMID: 30772301 | SOX2 |
| 5919 | TPRA1 | SOX2 regulated genes in Keratinocyte Overexpression from PMID: 30772301 | SOX2 |
| 5920 | GALNT14 | SOX2 regulated genes in Keratinocyte Overexpression from PMID: 30772301 | SOX2 |
| 5921 | MYO5C | SOX2 regulated genes in Keratinocyte Overexpression from PMID: 30772301 | SOX2 |
| 5922 | ZNRF1 | SOX2 regulated genes in Keratinocyte Overexpression from PMID: 30772301 | SOX2 |
| 5923 | NUP214 | SOX2 regulated genes in Keratinocyte Overexpression from PMID: 30772301 | SOX2 |
| 5924 | WNK2 | SOX2 regulated genes in Keratinocyte Overexpression from PMID: 30772301 | SOX2 |
| 5925 | CBWD3 | SOX2 regulated genes in Keratinocyte Overexpression from PMID: 30772301 | SOX2 |
| 5926 | CBWD5 | SOX2 regulated genes in Keratinocyte Overexpression from PMID: 30772301 | SOX2 |
| 5927 | CBWD6 | SOX2 regulated genes in Keratinocyte Overexpression from PMID: 30772301 | SOX2 |
| 5928 | USP1 | SOX2 regulated genes in Keratinocyte Overexpression from PMID: 30772301 | SOX2 |
| 5929 | SLC27A5 | SOX2 regulated genes in Keratinocyte Overexpression from PMID: 30772301 | SOX2 |
| 5930 | RNF6 | SOX2 regulated genes in Keratinocyte Overexpression from PMID: 30772301 | SOX2 |
| 5931 | TMEM254 | SOX2 regulated genes in Keratinocyte Overexpression from PMID: 30772301 | SOX2 |
| 5932 | RPP30 | SOX2 regulated genes in Keratinocyte Overexpression from PMID: 30772301 | SOX2 |
| 5933 | RNASEH2C | SOX2 regulated genes in Keratinocyte Overexpression from PMID: 30772301 | SOX2 |
| 5934 | COMMD2 | SOX2 regulated genes in Keratinocyte Overexpression from PMID: 30772301 | SOX2 |
| 5935 | AEN | SOX2 regulated genes in Keratinocyte Overexpression from PMID: 30772301 | SOX2 |
| 5936 | NRG2 | SOX2 regulated genes in Keratinocyte Overexpression from PMID: 30772301 | SOX2 |
| 5937 | CENPU | SOX2 regulated genes in Keratinocyte Overexpression from PMID: 30772301 | SOX2 |
| 5938 | KCNMB4 | SOX2 regulated genes in Keratinocyte Overexpression from PMID: 30772301 | SOX2 |
| 5939 | MAP3K9 | SOX2 regulated genes in Keratinocyte Overexpression from PMID: 30772301 | SOX2 |
| 5940 | EPC1 | SOX2 regulated genes in Keratinocyte Overexpression from PMID: 30772301 | SOX2 |
| 5941 | GRTP1 | SOX2 regulated genes in Keratinocyte Overexpression from PMID: 30772301 | SOX2 |
| 5942 | NETO2 | SOX2 regulated genes in Keratinocyte Overexpression from PMID: 30772301 | SOX2 |
| 5943 | SEMA4D | SOX2 regulated genes in Keratinocyte Overexpression from PMID: 30772301 | SOX2 |
| 5944 | TMC1 | SOX2 regulated genes in Keratinocyte Overexpression from PMID: 30772301 | SOX2 |
| 5945 | SARNP | SOX2 regulated genes in Keratinocyte Overexpression from PMID: 30772301 | SOX2 |
| 5946 | RHPN1 | SOX2 regulated genes in Keratinocyte Overexpression from PMID: 30772301 | SOX2 |
| 5947 | ALG11 | SOX2 regulated genes in Keratinocyte Overexpression from PMID: 30772301 | SOX2 |
| 5948 | CTHRC1 | SOX2 regulated genes in Keratinocyte Overexpression from PMID: 30772301 | SOX2 |
| 5949 | EPB41L4B | SOX2 regulated genes in Keratinocyte Overexpression from PMID: 30772301 | SOX2 |
| 5950 | MAMDC4 | SOX2 regulated genes in Keratinocyte Overexpression from PMID: 30772301 | SOX2 |
| 5951 | TLK1 | SOX2 regulated genes in Keratinocyte Overexpression from PMID: 30772301 | SOX2 |
| 5952 | IER3 | SOX2 regulated genes in Keratinocyte Overexpression from PMID: 30772301 | SOX2 |
| 5953 | ENHO | SOX2 regulated genes in Keratinocyte Overexpression from PMID: 30772301 | SOX2 |
| 5954 | TAB3 | SOX2 regulated genes in Keratinocyte Overexpression from PMID: 30772301 | SOX2 |
| 5955 | MED22 | SOX2 regulated genes in Keratinocyte Overexpression from PMID: 30772301 | SOX2 |
| 5956 | DBR1 | SOX2 regulated genes in Keratinocyte Overexpression from PMID: 30772301 | SOX2 |
| 5957 | MPV17L | SOX2 regulated genes in Keratinocyte Overexpression from PMID: 30772301 | SOX2 |
| 5958 | IFT74 | SOX2 regulated genes in Keratinocyte Overexpression from PMID: 30772301 | SOX2 |
| 5959 | CYP1B1 | SOX2 regulated genes in Keratinocyte Overexpression from PMID: 30772301 | SOX2 |
| 5960 | TG | SOX2 regulated genes in Keratinocyte Overexpression from PMID: 30772301 | SOX2 |
| 5961 | CDSN | SOX2 regulated genes in Keratinocyte Overexpression from PMID: 30772301 | SOX2 |
| 5962 | PAIP2B | SOX2 regulated genes in Keratinocyte Overexpression from PMID: 30772301 | SOX2 |
| 5963 | DTYMK | SOX2 regulated genes in Keratinocyte Overexpression from PMID: 30772301 | SOX2 |
| 5964 | SPATA5L1 | SOX2 regulated genes in Keratinocyte Overexpression from PMID: 30772301 | SOX2 |
| 5965 | COPG2 | SOX2 regulated genes in Keratinocyte Overexpression from PMID: 30772301 | SOX2 |
| 5966 | ZBTB40 | SOX2 regulated genes in Keratinocyte Overexpression from PMID: 30772301 | SOX2 |
| 5967 | CTU2 | SOX2 regulated genes in Keratinocyte Overexpression from PMID: 30772301 | SOX2 |
| 5968 | NEIL1 | SOX2 regulated genes in Keratinocyte Overexpression from PMID: 30772301 | SOX2 |
| 5969 | MYCBP | SOX2 regulated genes in Keratinocyte Overexpression from PMID: 30772301 | SOX2 |
| 5970 | BDKRB2 | SOX2 regulated genes in Keratinocyte Overexpression from PMID: 30772301 | SOX2 |
| 5971 | BHLHB9 | SOX2 regulated genes in Keratinocyte Overexpression from PMID: 30772301 | SOX2 |
| 5972 | CHCHD2 | SOX2 regulated genes in Keratinocyte Overexpression from PMID: 30772301 | SOX2 |
| 5973 | POLR2L | SOX2 regulated genes in Keratinocyte Overexpression from PMID: 30772301 | SOX2 |
| 5974 | SNRNP200 | SOX2 regulated genes in Keratinocyte Overexpression from PMID: 30772301 | SOX2 |
| 5975 | CALCOCO1 | SOX2 regulated genes in Keratinocyte Overexpression from PMID: 30772301 | SOX2 |
| 5976 | NUP107 | SOX2 regulated genes in Keratinocyte Overexpression from PMID: 30772301 | SOX2 |

| | | | |
|---|---|---|---|
| 5977 | NDE1 | SOX2 regulated genes in Keratinocyte Overexpression from PMID: 30772301 | SOX2 |
| 5978 | SKIV2L | SOX2 regulated genes in Keratinocyte Overexpression from PMID: 30772301 | SOX2 |
| 5979 | PRSS36 | SOX2 regulated genes in Keratinocyte Overexpression from PMID: 30772301 | SOX2 |
| 5980 | RNF19A | SOX2 regulated genes in Keratinocyte Overexpression from PMID: 30772301 | SOX2 |
| 5981 | SOD2 | SOX2 regulated genes in Keratinocyte Overexpression from PMID: 30772301 | SOX2 |
| 5982 | FXN | SOX2 regulated genes in Keratinocyte Overexpression from PMID: 30772301 | SOX2 |
| 5983 | NOXA1 | SOX2 regulated genes in Keratinocyte Overexpression from PMID: 30772301 | SOX2 |
| 5984 | EXOSC6 | SOX2 regulated genes in Keratinocyte Overexpression from PMID: 30772301 | SOX2 |
| 5985 | TRPV3 | SOX2 regulated genes in Keratinocyte Overexpression from PMID: 30772301 | SOX2 |
| 5986 | TMEM115 | SOX2 regulated genes in Keratinocyte Overexpression from PMID: 30772301 | SOX2 |
| 5987 | FBXO43 | SOX2 regulated genes in Keratinocyte Overexpression from PMID: 30772301 | SOX2 |
| 5988 | PDIA2 | SOX2 regulated genes in Keratinocyte Overexpression from PMID: 30772301 | SOX2 |
| 5989 | HAUS7 | SOX2 regulated genes in Keratinocyte Overexpression from PMID: 30772301 | SOX2 |
| 5990 | IL17D | SOX2 regulated genes in Keratinocyte Overexpression from PMID: 30772301 | SOX2 |
| 5991 | RAD52 | SOX2 regulated genes in Keratinocyte Overexpression from PMID: 30772301 | SOX2 |
| 5992 | INHBB | SOX2 regulated genes in Keratinocyte Overexpression from PMID: 30772301 | SOX2 |
| 5993 | FAM184B | SOX2 regulated genes in Keratinocyte Overexpression from PMID: 30772301 | SOX2 |
| 5994 | TAPT1 | SOX2 regulated genes in Keratinocyte Overexpression from PMID: 30772301 | SOX2 |
| 5995 | UBXN2A | SOX2 regulated genes in Keratinocyte Overexpression from PMID: 30772301 | SOX2 |
| 5996 | SEMA4G | SOX2 regulated genes in Keratinocyte Overexpression from PMID: 30772301 | SOX2 |
| 5997 | CPT1A | SOX2 regulated genes in Keratinocyte Overexpression from PMID: 30772301 | SOX2 |
| 5998 | BRMS1 | SOX2 regulated genes in Keratinocyte Overexpression from PMID: 30772301 | SOX2 |
| 5999 | ARHGEF26 | SOX2 regulated genes in Keratinocyte Overexpression from PMID: 30772301 | SOX2 |
| 6000 | PTCD1 | SOX2 regulated genes in Keratinocyte Overexpression from PMID: 30772301 | SOX2 |
| 6001 | CDKN1A | SOX2 regulated genes in Keratinocyte Overexpression from PMID: 30772301 | SOX2 |
| 6002 | CABLES2 | SOX2 regulated genes in Keratinocyte Overexpression from PMID: 30772301 | SOX2 |
| 6003 | RBM22 | SOX2 regulated genes in Keratinocyte Overexpression from PMID: 30772301 | SOX2 |
| 6004 | MTHFD2L | SOX2 regulated genes in Keratinocyte Overexpression from PMID: 30772301 | SOX2 |
| 6005 | DTX3 | SOX2 regulated genes in Keratinocyte Overexpression from PMID: 30772301 | SOX2 |
| 6006 | SMARCAD1 | SOX2 regulated genes in Keratinocyte Overexpression from PMID: 30772301 | SOX2 |
| 6007 | DRD4 | SOX2 regulated genes in Keratinocyte Overexpression from PMID: 30772301 | SOX2 |
| 6008 | KLK1 | SOX2 regulated genes in Keratinocyte Overexpression from PMID: 30772301 | SOX2 |
| 6009 | LAMB1 | SOX2 regulated genes in Keratinocyte Overexpression from PMID: 30772301 | SOX2 |
| 6010 | RNF185 | SOX2 regulated genes in Keratinocyte Overexpression from PMID: 30772301 | SOX2 |
| 6011 | POLR1D | SOX2 regulated genes in Keratinocyte Overexpression from PMID: 30772301 | SOX2 |
| 6012 | GLE1 | SOX2 regulated genes in Keratinocyte Overexpression from PMID: 30772301 | SOX2 |
| 6013 | CBY1 | SOX2 regulated genes in Keratinocyte Overexpression from PMID: 30772301 | SOX2 |
| 6014 | SPOPL | SOX2 regulated genes in Keratinocyte Overexpression from PMID: 30772301 | SOX2 |
| 6015 | MASTL | SOX2 regulated genes in Keratinocyte Overexpression from PMID: 30772301 | SOX2 |
| 6016 | POLR2B | SOX2 regulated genes in Keratinocyte Overexpression from PMID: 30772301 | SOX2 |
| 6017 | RBM27 | SOX2 regulated genes in Keratinocyte Overexpression from PMID: 30772301 | SOX2 |
| 6018 | KATNA1 | SOX2 regulated genes in Keratinocyte Overexpression from PMID: 30772301 | SOX2 |
| 6019 | PALB2 | SOX2 regulated genes in Keratinocyte Overexpression from PMID: 30772301 | SOX2 |
| 6020 | MCRS1 | SOX2 regulated genes in Keratinocyte Overexpression from PMID: 30772301 | SOX2 |
| 6021 | GPR3 | SOX2 regulated genes in Keratinocyte Overexpression from PMID: 30772301 | SOX2 |
| 6022 | MANF | SOX2 regulated genes in Keratinocyte Overexpression from PMID: 30772301 | SOX2 |
| 6023 | QRSL1 | SOX2 regulated genes in Keratinocyte Overexpression from PMID: 30772301 | SOX2 |
| 6024 | SH3RF2 | SOX2 regulated genes in Keratinocyte Overexpression from PMID: 30772301 | SOX2 |
| 6025 | RCOR2 | SOX2 regulated genes in Keratinocyte Overexpression from PMID: 30772301 | SOX2 |
| 6026 | HIPK4 | SOX2 regulated genes in Keratinocyte Overexpression from PMID: 30772301 | SOX2 |
| 6027 | NUP205 | SOX2 regulated genes in Keratinocyte Overexpression from PMID: 30772301 | SOX2 |
| 6028 | NT5DC2 | SOX2 regulated genes in Keratinocyte Overexpression from PMID: 30772301 | SOX2 |
| 6029 | ARPP19 | SOX2 regulated genes in Keratinocyte Overexpression from PMID: 30772301 | SOX2 |
| 6030 | RNF7 | SOX2 regulated genes in Keratinocyte Overexpression from PMID: 30772301 | SOX2 |
| 6031 | MAGOH | SOX2 regulated genes in Keratinocyte Overexpression from PMID: 30772301 | SOX2 |
| 6032 | ECHDC2 | SOX2 regulated genes in Keratinocyte Overexpression from PMID: 30772301 | SOX2 |
| 6033 | MET | SOX2 regulated genes in Keratinocyte Overexpression from PMID: 30772301 | SOX2 |
| 6034 | MRPL52 | SOX2 regulated genes in Keratinocyte Overexpression from PMID: 30772301 | SOX2 |
| 6035 | MRPL48 | SOX2 regulated genes in Keratinocyte Overexpression from PMID: 30772301 | SOX2 |
| 6036 | YPEL3 | SOX2 regulated genes in Keratinocyte Overexpression from PMID: 30772301 | SOX2 |
| 6037 | PPIF | SOX2 regulated genes in Keratinocyte Overexpression from PMID: 30772301 | SOX2 |

| | | | |
|---|---|---|---|
| 6038 | SRBD1 | SOX2 regulated genes in Keratinocyte Overexpression from PMID: 30772301 | SOX2 |
| 6039 | RANBP3 | SOX2 regulated genes in Keratinocyte Overexpression from PMID: 30772301 | SOX2 |
| 6040 | ZNF507 | SOX2 regulated genes in Keratinocyte Overexpression from PMID: 30772301 | SOX2 |
| 6041 | NHLRC2 | SOX2 regulated genes in Keratinocyte Overexpression from PMID: 30772301 | SOX2 |
| 6042 | OXSR1 | SOX2 regulated genes in Keratinocyte Overexpression from PMID: 30772301 | SOX2 |
| 6043 | MRRF | SOX2 regulated genes in Keratinocyte Overexpression from PMID: 30772301 | SOX2 |
| 6044 | ESM1 | SOX2 regulated genes in Keratinocyte Overexpression from PMID: 30772301 | SOX2 |
| 6045 | PIGO | SOX2 regulated genes in Keratinocyte Overexpression from PMID: 30772301 | SOX2 |
| 6046 | UTP3 | SOX2 regulated genes in Keratinocyte Overexpression from PMID: 30772301 | SOX2 |
| 6047 | CBX4 | SOX2 regulated genes in Keratinocyte Overexpression from PMID: 30772301 | SOX2 |
| 6048 | C6orf226 | SOX2 regulated genes in Keratinocyte Overexpression from PMID: 30772301 | SOX2 |
| 6049 | PKMYT1 | SOX2 regulated genes in Keratinocyte Overexpression from PMID: 30772301 | SOX2 |
| 6050 | ERICH5 | SOX2 regulated genes in Keratinocyte Overexpression from PMID: 30772301 | SOX2 |
| 6051 | FBXW7 | SOX2 regulated genes in Keratinocyte Overexpression from PMID: 30772301 | SOX2 |
| 6052 | CENPI | SOX2 regulated genes in Keratinocyte Overexpression from PMID: 30772301 | SOX2 |
| 6053 | CDKL3 | SOX2 regulated genes in Keratinocyte Overexpression from PMID: 30772301 | SOX2 |
| 6054 | CCT4 | SOX2 regulated genes in Keratinocyte Overexpression from PMID: 30772301 | SOX2 |
| 6055 | TRAPPC13 | SOX2 regulated genes in Keratinocyte Overexpression from PMID: 30772301 | SOX2 |
| 6056 | INSIG1 | SOX2 regulated genes in Keratinocyte Overexpression from PMID: 30772301 | SOX2 |
| 6057 | PRC1 | SOX2 regulated genes in Keratinocyte Overexpression from PMID: 30772301 | SOX2 |
| 6058 | HIPK2 | SOX2 regulated genes in Keratinocyte Overexpression from PMID: 30772301 | SOX2 |
| 6059 | FSBP | SOX2 regulated genes in Keratinocyte Overexpression from PMID: 30772301 | SOX2 |
| 6060 | PSMB1 | SOX2 regulated genes in Keratinocyte Overexpression from PMID: 30772301 | SOX2 |
| 6061 | GCAT | SOX2 regulated genes in Keratinocyte Overexpression from PMID: 30772301 | SOX2 |
| 6062 | ZBTB44 | SOX2 regulated genes in Keratinocyte Overexpression from PMID: 30772301 | SOX2 |
| 6063 | ORC6 | SOX2 regulated genes in Keratinocyte Overexpression from PMID: 30772301 | SOX2 |
| 6064 | DERL1 | SOX2 regulated genes in Keratinocyte Overexpression from PMID: 30772301 | SOX2 |
| 6065 | NUTF2 | SOX2 regulated genes in Keratinocyte Overexpression from PMID: 30772301 | SOX2 |
| 6066 | UBE4B | SOX2 regulated genes in Keratinocyte Overexpression from PMID: 30772301 | SOX2 |
| 6067 | YLPM1 | SOX2 regulated genes in Keratinocyte Overexpression from PMID: 30772301 | SOX2 |
| 6068 | GALT | SOX2 regulated genes in Keratinocyte Overexpression from PMID: 30772301 | SOX2 |
| 6069 | NR2C2AP | SOX2 regulated genes in Keratinocyte Overexpression from PMID: 30772301 | SOX2 |
| 6070 | RFWD3 | SOX2 regulated genes in Keratinocyte Overexpression from PMID: 30772301 | SOX2 |
| 6071 | GINS2 | SOX2 regulated genes in Keratinocyte Overexpression from PMID: 30772301 | SOX2 |
| 6072 | ETAA1 | SOX2 regulated genes in Keratinocyte Overexpression from PMID: 30772301 | SOX2 |
| 6073 | PDCD2L | SOX2 regulated genes in Keratinocyte Overexpression from PMID: 30772301 | SOX2 |
| 6074 | NCBP2L | SOX2 regulated genes in Keratinocyte Overexpression from PMID: 30772301 | SOX2 |
| 6075 | CHP1 | SOX2 regulated genes in Keratinocyte Overexpression from PMID: 30772301 | SOX2 |
| 6076 | BRPF3 | SOX2 regulated genes in Keratinocyte Overexpression from PMID: 30772301 | SOX2 |
| 6077 | KRTAP9-4 | SOX2 regulated genes in Keratinocyte Overexpression from PMID: 30772301 | SOX2 |
| 6078 | XPO4 | SOX2 regulated genes in Keratinocyte Overexpression from PMID: 30772301 | SOX2 |
| 6079 | DDA1 | SOX2 regulated genes in Keratinocyte Overexpression from PMID: 30772301 | SOX2 |
| 6080 | DOCK3 | SOX2 regulated genes in Keratinocyte Overexpression from PMID: 30772301 | SOX2 |
| 6081 | ENO4 | SOX2 regulated genes in Keratinocyte Overexpression from PMID: 30772301 | SOX2 |
| 6082 | TUSC2 | SOX2 regulated genes in Keratinocyte Overexpression from PMID: 30772301 | SOX2 |
| 6083 | SPEN | SOX2 regulated genes in Keratinocyte Overexpression from PMID: 30772301 | SOX2 |
| 6084 | DHFR | SOX2 regulated genes in Keratinocyte Overexpression from PMID: 30772301 | SOX2 |
| 6085 | MED6 | SOX2 regulated genes in Keratinocyte Overexpression from PMID: 30772301 | SOX2 |
| 6086 | KANK4 | SOX2 regulated genes in Keratinocyte Overexpression from PMID: 30772301 | SOX2 |
| 6087 | FUS | SOX2 regulated genes in Keratinocyte Overexpression from PMID: 30772301 | SOX2 |
| 6088 | SGF29 | SOX2 regulated genes in Keratinocyte Overexpression from PMID: 30772301 | SOX2 |
| 6089 | EPB41L5 | SOX2 regulated genes in Keratinocyte Overexpression from PMID: 30772301 | SOX2 |
| 6090 | STAM2 | SOX2 regulated genes in Keratinocyte Overexpression from PMID: 30772301 | SOX2 |
| 6091 | FEM1C | SOX2 regulated genes in Keratinocyte Overexpression from PMID: 30772301 | SOX2 |
| 6092 | CKAP2 | SOX2 regulated genes in Keratinocyte Overexpression from PMID: 30772301 | SOX2 |
| 6093 | SLC25A25 | SOX2 regulated genes in Keratinocyte Overexpression from PMID: 30772301 | SOX2 |
| 6094 | OXNAD1 | SOX2 regulated genes in Keratinocyte Overexpression from PMID: 30772301 | SOX2 |
| 6095 | PPP6R1 | SOX2 regulated genes in Keratinocyte Overexpression from PMID: 30772301 | SOX2 |
| 6096 | TMEM86B | SOX2 regulated genes in Keratinocyte Overexpression from PMID: 30772301 | SOX2 |
| 6097 | AGMAT | SOX2 regulated genes in Keratinocyte Overexpression from PMID: 30772301 | SOX2 |
| 6098 | DYNLRB2 | SOX2 regulated genes in Keratinocyte Overexpression from PMID: 30772301 | SOX2 |

| | | | |
|---|---|---|---|
| 6099 | HYLS1 | SOX2 regulated genes in Keratinocyte Overexpression from PMID: 30772301 | SOX2 |
| 6100 | ZNF398 | SOX2 regulated genes in Keratinocyte Overexpression from PMID: 30772301 | SOX2 |
| 6101 | CMSS1 | SOX2 regulated genes in Keratinocyte Overexpression from PMID: 30772301 | SOX2 |
| 6102 | CMTM8 | SOX2 regulated genes in Keratinocyte Overexpression from PMID: 30772301 | SOX2 |
| 6103 | PDCD7 | SOX2 regulated genes in Keratinocyte Overexpression from PMID: 30772301 | SOX2 |
| 6104 | MAP3K10 | SOX2 regulated genes in Keratinocyte Overexpression from PMID: 30772301 | SOX2 |
| 6105 | CCDC65 | SOX2 regulated genes in Keratinocyte Overexpression from PMID: 30772301 | SOX2 |
| 6106 | RDX | SOX2 regulated genes in Keratinocyte Overexpression from PMID: 30772301 | SOX2 |
| 6107 | NOA1 | SOX2 regulated genes in Keratinocyte Overexpression from PMID: 30772301 | SOX2 |
| 6108 | RPA1 | SOX2 regulated genes in Keratinocyte Overexpression from PMID: 30772301 | SOX2 |
| 6109 | DOC2A | SOX2 regulated genes in Keratinocyte Overexpression from PMID: 30772301 | SOX2 |
| 6110 | LARGE2 | SOX2 regulated genes in Keratinocyte Overexpression from PMID: 30772301 | SOX2 |
| 6111 | GABRE | SOX2 regulated genes in Keratinocyte Overexpression from PMID: 30772301 | SOX2 |
| 6112 | MED13 | SOX2 regulated genes in Keratinocyte Overexpression from PMID: 30772301 | SOX2 |
| 6113 | IRF2BP1 | SOX2 regulated genes in Keratinocyte Overexpression from PMID: 30772301 | SOX2 |
| 6114 | RBM19 | SOX2 regulated genes in Keratinocyte Overexpression from PMID: 30772301 | SOX2 |
| 6115 | AMTN | SOX2 regulated genes in Keratinocyte Overexpression from PMID: 30772301 | SOX2 |
| 6116 | PROM2 | SOX2 regulated genes in Keratinocyte Overexpression from PMID: 30772301 | SOX2 |
| 6117 | JUP | SOX2 regulated genes in Keratinocyte Overexpression from PMID: 30772301 | SOX2 |
| 6118 | CTR9 | SOX2 regulated genes in Keratinocyte Overexpression from PMID: 30772301 | SOX2 |
| 6119 | FRMPD1 | SOX2 regulated genes in Keratinocyte Overexpression from PMID: 30772301 | SOX2 |
| 6120 | SLC5A8 | SOX2 regulated genes in Keratinocyte Overexpression from PMID: 30772301 | SOX2 |
| 6121 | HSPA1A | SOX2 regulated genes in Keratinocyte Overexpression from PMID: 30772301 | SOX2 |
| 6122 | HSPA1B | SOX2 regulated genes in Keratinocyte Overexpression from PMID: 30772301 | SOX2 |
| 6123 | PNOC | SOX2 regulated genes in Keratinocyte Overexpression from PMID: 30772301 | SOX2 |
| 6124 | GRHL2 | SOX2 regulated genes in Keratinocyte Overexpression from PMID: 30772301 | SOX2 |
| 6125 | KANK1 | SOX2 regulated genes in Keratinocyte Overexpression from PMID: 30772301 | SOX2 |
| 6126 | ARFGEF1 | SOX2 regulated genes in Keratinocyte Overexpression from PMID: 30772301 | SOX2 |
| 6127 | NOL7 | SOX2 regulated genes in Keratinocyte Overexpression from PMID: 30772301 | SOX2 |
| 6128 | NAF1 | SOX2 regulated genes in Keratinocyte Overexpression from PMID: 30772301 | SOX2 |
| 6129 | RHCG | SOX2 regulated genes in Keratinocyte Overexpression from PMID: 30772301 | SOX2 |
| 6130 | ATP6V1C1 | SOX2 regulated genes in Keratinocyte Overexpression from PMID: 30772301 | SOX2 |
| 6131 | ZNF212 | SOX2 regulated genes in Keratinocyte Overexpression from PMID: 30772301 | SOX2 |
| 6132 | MAD1L1 | SOX2 regulated genes in Keratinocyte Overexpression from PMID: 30772301 | SOX2 |
| 6133 | GRPEL2 | SOX2 regulated genes in Keratinocyte Overexpression from PMID: 30772301 | SOX2 |
| 6134 | UBE3C | SOX2 regulated genes in Keratinocyte Overexpression from PMID: 30772301 | SOX2 |
| 6135 | TMEM147 | SOX2 regulated genes in Keratinocyte Overexpression from PMID: 30772301 | SOX2 |
| 6136 | C16orf87 | SOX2 regulated genes in Keratinocyte Overexpression from PMID: 30772301 | SOX2 |
| 6137 | DYNLT1 | SOX2 regulated genes in Keratinocyte Overexpression from PMID: 30772301 | SOX2 |
| 6138 | NRF1 | SOX2 regulated genes in Keratinocyte Overexpression from PMID: 30772301 | SOX2 |
| 6139 | CRHBP | SOX2 regulated genes in Keratinocyte Overexpression from PMID: 30772301 | SOX2 |
| 6140 | SGTB | SOX2 regulated genes in Keratinocyte Overexpression from PMID: 30772301 | SOX2 |
| 6141 | IDI2 | SOX2 regulated genes in Keratinocyte Overexpression from PMID: 30772301 | SOX2 |
| 6142 | TIMM17B | SOX2 regulated genes in Keratinocyte Overexpression from PMID: 30772301 | SOX2 |
| 6143 | MATN4 | SOX2 regulated genes in Keratinocyte Overexpression from PMID: 30772301 | SOX2 |
| 6144 | COL12A1 | SOX2 regulated genes in Keratinocyte Overexpression from PMID: 30772301 | SOX2 |
| 6145 | FBXO32 | SOX2 regulated genes in Keratinocyte Overexpression from PMID: 30772301 | SOX2 |
| 6146 | COX6A1 | SOX2 regulated genes in Keratinocyte Overexpression from PMID: 30772301 | SOX2 |
| 6147 | EIF3J | SOX2 regulated genes in Keratinocyte Overexpression from PMID: 30772301 | SOX2 |
| 6148 | SOWAHA | SOX2 regulated genes in Keratinocyte Overexpression from PMID: 30772301 | SOX2 |
| 6149 | VCL | SOX2 regulated genes in Keratinocyte Overexpression from PMID: 30772301 | SOX2 |
| 6150 | PHF20L1 | SOX2 regulated genes in Keratinocyte Overexpression from PMID: 30772301 | SOX2 |
| 6151 | RAPGEF4 | SOX2 regulated genes in Keratinocyte Overexpression from PMID: 30772301 | SOX2 |
| 6152 | METTL17 | SOX2 regulated genes in Keratinocyte Overexpression from PMID: 30772301 | SOX2 |
| 6153 | SCRIB | SOX2 regulated genes in Keratinocyte Overexpression from PMID: 30772301 | SOX2 |
| 6154 | SHISA2 | SOX2 regulated genes in Keratinocyte Overexpression from PMID: 30772301 | SOX2 |
| 6155 | PBX2 | SOX2 regulated genes in Keratinocyte Overexpression from PMID: 30772301 | SOX2 |
| 6156 | SYTL1 | SOX2 regulated genes in Keratinocyte Overexpression from PMID: 30772301 | SOX2 |
| 6157 | RNF103 | SOX2 regulated genes in Keratinocyte Overexpression from PMID: 30772301 | SOX2 |
| 6158 | KMT5B | SOX2 regulated genes in Keratinocyte Overexpression from PMID: 30772301 | SOX2 |
| 6159 | SMG9 | SOX2 regulated genes in Keratinocyte Overexpression from PMID: 30772301 | SOX2 |

| | | | |
|---|---|---|---|
| 6160 | ALX1 | SOX2 regulated genes in Keratinocyte Overexpression from PMID: 30772301 | SOX2 |
| 6161 | IFT57 | SOX2 regulated genes in Keratinocyte Overexpression from PMID: 30772301 | SOX2 |
| 6162 | LRRC74A | SOX2 regulated genes in Keratinocyte Overexpression from PMID: 30772301 | SOX2 |
| 6163 | CPT2 | SOX2 regulated genes in Keratinocyte Overexpression from PMID: 30772301 | SOX2 |
| 6164 | NUDT14 | SOX2 regulated genes in Keratinocyte Overexpression from PMID: 30772301 | SOX2 |
| 6165 | COPS5 | SOX2 regulated genes in Keratinocyte Overexpression from PMID: 30772301 | SOX2 |
| 6166 | LANCL1 | SOX2 regulated genes in Keratinocyte Overexpression from PMID: 30772301 | SOX2 |
| 6167 | TTC9C | SOX2 regulated genes in Keratinocyte Overexpression from PMID: 30772301 | SOX2 |
| 6168 | COX18 | SOX2 regulated genes in Keratinocyte Overexpression from PMID: 30772301 | SOX2 |
| 6169 | RSPH9 | SOX2 regulated genes in Keratinocyte Overexpression from PMID: 30772301 | SOX2 |
| 6170 | SARS2 | SOX2 regulated genes in Keratinocyte Overexpression from PMID: 30772301 | SOX2 |
| 6171 | FIS1 | SOX2 regulated genes in Keratinocyte Overexpression from PMID: 30772301 | SOX2 |
| 6172 | POLE3 | SOX2 regulated genes in Keratinocyte Overexpression from PMID: 30772301 | SOX2 |
| 6173 | SSNA1 | SOX2 regulated genes in Keratinocyte Overexpression from PMID: 30772301 | SOX2 |
| 6174 | GPAA1 | SOX2 regulated genes in Keratinocyte Overexpression from PMID: 30772301 | SOX2 |
| 6175 | MRE11 | SOX2 regulated genes in Keratinocyte Overexpression from PMID: 30772301 | SOX2 |
| 6176 | UBE3A | SOX2 regulated genes in Keratinocyte Overexpression from PMID: 30772301 | SOX2 |
| 6177 | SPOP | SOX2 regulated genes in Keratinocyte Overexpression from PMID: 30772301 | SOX2 |
| 6178 | POC5 | SOX2 regulated genes in Keratinocyte Overexpression from PMID: 30772301 | SOX2 |
| 6179 | HSPA5 | SOX2 regulated genes in Keratinocyte Overexpression from PMID: 30772301 | SOX2 |
| 6180 | PDCL3 | SOX2 regulated genes in Keratinocyte Overexpression from PMID: 30772301 | SOX2 |
| 6181 | PSMA2 | SOX2 regulated genes in Keratinocyte Overexpression from PMID: 30772301 | SOX2 |
| 6182 | ALDH5A1 | SOX2 regulated genes in Keratinocyte Overexpression from PMID: 30772301 | SOX2 |
| 6183 | MRPS23 | SOX2 regulated genes in Keratinocyte Overexpression from PMID: 30772301 | SOX2 |
| 6184 | TFB1M | SOX2 regulated genes in Keratinocyte Overexpression from PMID: 30772301 | SOX2 |
| 6185 | PANK3 | SOX2 regulated genes in Keratinocyte Overexpression from PMID: 30772301 | SOX2 |
| 6186 | CDK12 | SOX2 regulated genes in Keratinocyte Overexpression from PMID: 30772301 | SOX2 |
| 6187 | CCDC60 | SOX2 regulated genes in Keratinocyte Overexpression from PMID: 30772301 | SOX2 |
| 6188 | DAPK2 | SOX2 regulated genes in Keratinocyte Overexpression from PMID: 30772301 | SOX2 |
| 6189 | LAS1L | SOX2 regulated genes in Keratinocyte Overexpression from PMID: 30772301 | SOX2 |
| 6190 | DCTD | SOX2 regulated genes in Keratinocyte Overexpression from PMID: 30772301 | SOX2 |
| 6191 | PCYOX1L | SOX2 regulated genes in Keratinocyte Overexpression from PMID: 30772301 | SOX2 |
| 6192 | NUDCD2 | SOX2 regulated genes in Keratinocyte Overexpression from PMID: 30772301 | SOX2 |
| 6193 | MCM6 | SOX2 regulated genes in Keratinocyte Overexpression from PMID: 30772301 | SOX2 |
| 6194 | CPA5 | SOX2 regulated genes in Keratinocyte Overexpression from PMID: 30772301 | SOX2 |
| 6195 | HES5 | SOX2 regulated genes in Keratinocyte Overexpression from PMID: 30772301 | SOX2 |
| 6196 | APOL3 | SOX2 regulated genes in Keratinocyte Overexpression from PMID: 30772301 | SOX2 |
| 6197 | APOL4 | SOX2 regulated genes in Keratinocyte Overexpression from PMID: 30772301 | SOX2 |
| 6198 | APOL2 | SOX2 regulated genes in Keratinocyte Overexpression from PMID: 30772301 | SOX2 |
| 6199 | APOL1 | SOX2 regulated genes in Keratinocyte Overexpression from PMID: 30772301 | SOX2 |
| 6200 | MPZL2 | SOX2 regulated genes in Keratinocyte Overexpression from PMID: 30772301 | SOX2 |
| 6201 | SPRED2 | SOX2 regulated genes in Keratinocyte Overexpression from PMID: 30772301 | SOX2 |
| 6202 | RPEL1 | SOX2 regulated genes in Keratinocyte Overexpression from PMID: 30772301 | SOX2 |
| 6203 | RPE | SOX2 regulated genes in Keratinocyte Overexpression from PMID: 30772301 | SOX2 |
| 6204 | CFL1 | SOX2 regulated genes in Keratinocyte Overexpression from PMID: 30772301 | SOX2 |
| 6205 | DUSP19 | SOX2 regulated genes in Keratinocyte Overexpression from PMID: 30772301 | SOX2 |
| 6206 | CCDC15 | SOX2 regulated genes in Keratinocyte Overexpression from PMID: 30772301 | SOX2 |
| 6207 | MAPK8IP1 | SOX2 regulated genes in Keratinocyte Overexpression from PMID: 30772301 | SOX2 |
| 6208 | CAPN1 | SOX2 regulated genes in Keratinocyte Overexpression from PMID: 30772301 | SOX2 |
| 6209 | TECTA | SOX2 regulated genes in Keratinocyte Overexpression from PMID: 30772301 | SOX2 |
| 6210 | ELP3 | SOX2 regulated genes in Keratinocyte Overexpression from PMID: 30772301 | SOX2 |
| 6211 | PPAT | SOX2 regulated genes in Keratinocyte Overexpression from PMID: 30772301 | SOX2 |
| 6212 | RFX3 | SOX2 regulated genes in Keratinocyte Overexpression from PMID: 30772301 | SOX2 |
| 6213 | FAM227A | SOX2 regulated genes in Keratinocyte Overexpression from PMID: 30772301 | SOX2 |
| 6214 | MRPL18 | SOX2 regulated genes in Keratinocyte Overexpression from PMID: 30772301 | SOX2 |
| 6215 | TRIM2 | SOX2 regulated genes in Keratinocyte Overexpression from PMID: 30772301 | SOX2 |
| 6216 | ASAP2 | SOX2 regulated genes in Keratinocyte Overexpression from PMID: 30772301 | SOX2 |
| 6217 | SIAH1 | SOX2 regulated genes in Keratinocyte Overexpression from PMID: 30772301 | SOX2 |
| 6218 | ATP1B3 | SOX2 regulated genes in Keratinocyte Overexpression from PMID: 30772301 | SOX2 |
| 6219 | CYB5D1 | SOX2 regulated genes in Keratinocyte Overexpression from PMID: 30772301 | SOX2 |
| 6220 | SRF | SOX2 regulated genes in Keratinocyte Overexpression from PMID: 30772301 | SOX2 |

| | | | |
|---|---|---|---|
| 6221 | CPSF1 | SOX2 regulated genes in Keratinocyte Overexpression from PMID: 30772301 | SOX2 |
| 6222 | DTWD2 | SOX2 regulated genes in Keratinocyte Overexpression from PMID: 30772301 | SOX2 |
| 6223 | SEC24C | SOX2 regulated genes in Keratinocyte Overexpression from PMID: 30772301 | SOX2 |
| 6224 | GNE | SOX2 regulated genes in Keratinocyte Overexpression from PMID: 30772301 | SOX2 |
| 6225 | PTCH1 | SOX2 regulated genes in Keratinocyte Overexpression from PMID: 30772301 | SOX2 |
| 6226 | ZNF282 | SOX2 regulated genes in Keratinocyte Overexpression from PMID: 30772301 | SOX2 |
| 6227 | MTOR | SOX2 regulated genes in Keratinocyte Overexpression from PMID: 30772301 | SOX2 |
| 6228 | SUPT5H | SOX2 regulated genes in Keratinocyte Overexpression from PMID: 30772301 | SOX2 |
| 6229 | ATP8A2 | SOX2 regulated genes in Keratinocyte Overexpression from PMID: 30772301 | SOX2 |
| 6230 | GDPD3 | SOX2 regulated genes in Keratinocyte Overexpression from PMID: 30772301 | SOX2 |
| 6231 | IFFO2 | SOX2 regulated genes in Keratinocyte Overexpression from PMID: 30772301 | SOX2 |
| 6232 | RCN1 | SOX2 regulated genes in Keratinocyte Overexpression from PMID: 30772301 | SOX2 |
| 6233 | MBTD1 | SOX2 regulated genes in Keratinocyte Overexpression from PMID: 30772301 | SOX2 |
| 6234 | GCSH | SOX2 regulated genes in Keratinocyte Overexpression from PMID: 30772301 | SOX2 |
| 6235 | SLBP | SOX2 regulated genes in Keratinocyte Overexpression from PMID: 30772301 | SOX2 |
| 6236 | DOCK9 | SOX2 regulated genes in Keratinocyte Overexpression from PMID: 30772301 | SOX2 |
| 6237 | COQ5 | SOX2 regulated genes in Keratinocyte Overexpression from PMID: 30772301 | SOX2 |
| 6238 | POLR2G | SOX2 regulated genes in Keratinocyte Overexpression from PMID: 30772301 | SOX2 |
| 6239 | WDR20 | SOX2 regulated genes in Keratinocyte Overexpression from PMID: 30772301 | SOX2 |
| 6240 | RBM28 | SOX2 regulated genes in Keratinocyte Overexpression from PMID: 30772301 | SOX2 |
| 6241 | MRPS30 | SOX2 regulated genes in Keratinocyte Overexpression from PMID: 30772301 | SOX2 |
| 6242 | PWWP2B | SOX2 regulated genes in Keratinocyte Overexpression from PMID: 30772301 | SOX2 |
| 6243 | AP4S1 | SOX2 regulated genes in Keratinocyte Overexpression from PMID: 30772301 | SOX2 |
| 6244 | ALCAM | SOX2 regulated genes in Keratinocyte Overexpression from PMID: 30772301 | SOX2 |
| 6245 | JMJD7-PLA2G4B | SOX2 regulated genes in Keratinocyte Overexpression from PMID: 30772301 | SOX2 |
| 6246 | HYAL4 | SOX2 regulated genes in Keratinocyte Overexpression from PMID: 30772301 | SOX2 |
| 6247 | NUBP1 | SOX2 regulated genes in Keratinocyte Overexpression from PMID: 30772301 | SOX2 |
| 6248 | HADHB | SOX2 regulated genes in Keratinocyte Overexpression from PMID: 30772301 | SOX2 |
| 6249 | SPINK9 | SOX2 regulated genes in Keratinocyte Overexpression from PMID: 30772301 | SOX2 |
| 6250 | CCP110 | SOX2 regulated genes in Keratinocyte Overexpression from PMID: 30772301 | SOX2 |
| 6251 | FAAP100 | SOX2 regulated genes in Keratinocyte Overexpression from PMID: 30772301 | SOX2 |
| 6252 | JAKMIP2 | SOX2 regulated genes in Keratinocyte Overexpression from PMID: 30772301 | SOX2 |
| 6253 | DMTF1 | SOX2 regulated genes in Keratinocyte Overexpression from PMID: 30772301 | SOX2 |
| 6254 | NDOR1 | SOX2 regulated genes in Keratinocyte Overexpression from PMID: 30772301 | SOX2 |
| 6255 | UBE3D | SOX2 regulated genes in Keratinocyte Overexpression from PMID: 30772301 | SOX2 |
| 6256 | ANAPC1 | SOX2 regulated genes in Keratinocyte Overexpression from PMID: 30772301 | SOX2 |
| 6257 | CALML3 | SOX2 regulated genes in Keratinocyte Overexpression from PMID: 30772301 | SOX2 |
| 6258 | HSPA8 | SOX2 regulated genes in Keratinocyte Overexpression from PMID: 30772301 | SOX2 |
| 6259 | SF3B5 | SOX2 regulated genes in Keratinocyte Overexpression from PMID: 30772301 | SOX2 |
| 6260 | FOSL2 | SOX2 regulated genes in Keratinocyte Overexpression from PMID: 30772301 | SOX2 |
| 6261 | MREG | SOX2 regulated genes in Keratinocyte Overexpression from PMID: 30772301 | SOX2 |
| 6262 | DNPH1 | SOX2 regulated genes in Keratinocyte Overexpression from PMID: 30772301 | SOX2 |
| 6263 | GPC2 | SOX2 regulated genes in Keratinocyte Overexpression from PMID: 30772301 | SOX2 |
| 6264 | MTMR2 | SOX2 regulated genes in Keratinocyte Overexpression from PMID: 30772301 | SOX2 |
| 6265 | TOP2A | SOX2 regulated genes in Keratinocyte Overexpression from PMID: 30772301 | SOX2 |
| 6266 | WDR70 | SOX2 regulated genes in Keratinocyte Overexpression from PMID: 30772301 | SOX2 |
| 6267 | PEX6 | SOX2 regulated genes in Keratinocyte Overexpression from PMID: 30772301 | SOX2 |
| 6268 | ADAT2 | SOX2 regulated genes in Keratinocyte Overexpression from PMID: 30772301 | SOX2 |
| 6269 | LYPD5 | SOX2 regulated genes in Keratinocyte Overexpression from PMID: 30772301 | SOX2 |
| 6270 | PTPN1 | SOX2 regulated genes in Keratinocyte Overexpression from PMID: 30772301 | SOX2 |
| 6271 | C10orf95 | SOX2 regulated genes in Keratinocyte Overexpression from PMID: 30772301 | SOX2 |
| 6272 | TNKS1BP1 | SOX2 regulated genes in Keratinocyte Overexpression from PMID: 30772301 | SOX2 |
| 6273 | SLC22A23 | SOX2 regulated genes in Keratinocyte Overexpression from PMID: 30772301 | SOX2 |
| 6274 | CACFD1 | SOX2 regulated genes in Keratinocyte Overexpression from PMID: 30772301 | SOX2 |
| 6275 | SUGP1 | SOX2 regulated genes in Keratinocyte Overexpression from PMID: 30772301 | SOX2 |
| 6276 | ZNF786 | SOX2 regulated genes in Keratinocyte Overexpression from PMID: 30772301 | SOX2 |
| 6277 | EHD1 | SOX2 regulated genes in Keratinocyte Overexpression from PMID: 30772301 | SOX2 |
| 6278 | ARG1 | SOX2 regulated genes in Keratinocyte Overexpression from PMID: 30772301 | SOX2 |
| 6279 | WNT10B | SOX2 regulated genes in Keratinocyte Overexpression from PMID: 30772301 | SOX2 |
| 6280 | LIG1 | SOX2 regulated genes in Keratinocyte Overexpression from PMID: 30772301 | SOX2 |
| 6281 | HILPDA | SOX2 regulated genes in Keratinocyte Overexpression from PMID: 30772301 | SOX2 |

| | | | |
|---|---|---|---|
| 6282 | RABGAP1 | SOX2 regulated genes in Keratinocyte Overexpression from PMID: 30772301 | SOX2 |
| 6283 | EDAR | SOX2 regulated genes in Keratinocyte Overexpression from PMID: 30772301 | SOX2 |
| 6284 | HADHA | SOX2 regulated genes in Keratinocyte Overexpression from PMID: 30772301 | SOX2 |
| 6285 | ADM | SOX2 regulated genes in Keratinocyte Overexpression from PMID: 30772301 | SOX2 |
| 6286 | TMCO3 | SOX2 regulated genes in Keratinocyte Overexpression from PMID: 30772301 | SOX2 |
| 6287 | HNF4G | SOX2 regulated genes in Keratinocyte Overexpression from PMID: 30772301 | SOX2 |
| 6288 | SNIP1 | SOX2 regulated genes in Keratinocyte Overexpression from PMID: 30772301 | SOX2 |
| 6289 | KBTBD6 | SOX2 regulated genes in Keratinocyte Overexpression from PMID: 30772301 | SOX2 |
| 6290 | KBTBD7 | SOX2 regulated genes in Keratinocyte Overexpression from PMID: 30772301 | SOX2 |
| 6291 | PAX1 | SOX2 regulated genes in Keratinocyte Overexpression from PMID: 30772301 | SOX2 |
| 6292 | TOMM6 | SOX2 regulated genes in Keratinocyte Overexpression from PMID: 30772301 | SOX2 |
| 6293 | ATP10B | SOX2 regulated genes in Keratinocyte Overexpression from PMID: 30772301 | SOX2 |
| 6294 | KAT5 | SOX2 regulated genes in Keratinocyte Overexpression from PMID: 30772301 | SOX2 |
| 6295 | MTERF4 | SOX2 regulated genes in Keratinocyte Overexpression from PMID: 30772301 | SOX2 |
| 6296 | MAP3K15 | SOX2 regulated genes in Keratinocyte Overexpression from PMID: 30772301 | SOX2 |
| 6297 | NIPAL1 | SOX2 regulated genes in Keratinocyte Overexpression from PMID: 30772301 | SOX2 |
| 6298 | PUS10 | SOX2 regulated genes in Keratinocyte Overexpression from PMID: 30772301 | SOX2 |
| 6299 | ANAPC5 | SOX2 regulated genes in Keratinocyte Overexpression from PMID: 30772301 | SOX2 |
| 6300 | RDH10 | SOX2 regulated genes in Keratinocyte Overexpression from PMID: 30772301 | SOX2 |
| 6301 | NOP14 | SOX2 regulated genes in Keratinocyte Overexpression from PMID: 30772301 | SOX2 |
| 6302 | SPATA20 | SOX2 regulated genes in Keratinocyte Overexpression from PMID: 30772301 | SOX2 |
| 6303 | TACSTD2 | SOX2 regulated genes in Keratinocyte Overexpression from PMID: 30772301 | SOX2 |
| 6304 | MED29 | SOX2 regulated genes in Keratinocyte Overexpression from PMID: 30772301 | SOX2 |
| 6305 | DMAP1 | SOX2 regulated genes in Keratinocyte Overexpression from PMID: 30772301 | SOX2 |
| 6306 | JOSD2 | SOX2 regulated genes in Keratinocyte Overexpression from PMID: 30772301 | SOX2 |
| 6307 | TMPRSS13 | SOX2 regulated genes in Keratinocyte Overexpression from PMID: 30772301 | SOX2 |
| 6308 | KIF26A | SOX2 regulated genes in Keratinocyte Overexpression from PMID: 30772301 | SOX2 |
| 6309 | TRPV6 | SOX2 regulated genes in Keratinocyte Overexpression from PMID: 30772301 | SOX2 |
| 6310 | CDKN3 | SOX2 regulated genes in Keratinocyte Overexpression from PMID: 30772301 | SOX2 |
| 6311 | TTLL10 | SOX2 regulated genes in Keratinocyte Overexpression from PMID: 30772301 | SOX2 |
| 6312 | RTN4IP1 | SOX2 regulated genes in Keratinocyte Overexpression from PMID: 30772301 | SOX2 |
| 6313 | TMEM35A | SOX2 regulated genes in Keratinocyte Overexpression from PMID: 30772301 | SOX2 |
| 6314 | TRIB1 | SOX2 regulated genes in Keratinocyte Overexpression from PMID: 30772301 | SOX2 |
| 6315 | IZUMO1 | SOX2 regulated genes in Keratinocyte Overexpression from PMID: 30772301 | SOX2 |
| 6316 | TMEM134 | SOX2 regulated genes in Keratinocyte Overexpression from PMID: 30772301 | SOX2 |
| 6317 | ABCB8 | SOX2 regulated genes in Keratinocyte Overexpression from PMID: 30772301 | SOX2 |
| 6318 | YEATS4 | SOX2 regulated genes in Keratinocyte Overexpression from PMID: 30772301 | SOX2 |
| 6319 | RAD54B | SOX2 regulated genes in Keratinocyte Overexpression from PMID: 30772301 | SOX2 |
| 6320 | SCUBE1 | SOX2 regulated genes in Keratinocyte Overexpression from PMID: 30772301 | SOX2 |
| 6321 | OGDHL | SOX2 regulated genes in Keratinocyte Overexpression from PMID: 30772301 | SOX2 |
| 6322 | SCNN1G | SOX2 regulated genes in Keratinocyte Overexpression from PMID: 30772301 | SOX2 |
| 6323 | CEBPB | SOX2 regulated genes in Keratinocyte Overexpression from PMID: 30772301 | SOX2 |
| 6324 | RAD51C | SOX2 regulated genes in Keratinocyte Overexpression from PMID: 30772301 | SOX2 |
| 6325 | SERPINA1 | SOX2 regulated genes in Keratinocyte Overexpression from PMID: 30772301 | SOX2 |
| 6326 | NIPAL2 | SOX2 regulated genes in Keratinocyte Overexpression from PMID: 30772301 | SOX2 |
| 6327 | PTPRK | SOX2 regulated genes in Keratinocyte Overexpression from PMID: 30772301 | SOX2 |
| 6328 | B3GNT4 | SOX2 regulated genes in Keratinocyte Overexpression from PMID: 30772301 | SOX2 |
| 6329 | ALDH3A2 | SOX2 regulated genes in Keratinocyte Overexpression from PMID: 30772301 | SOX2 |
| 6330 | KITLG | SOX2 regulated genes in Keratinocyte Overexpression from PMID: 30772301 | SOX2 |
| 6331 | PRRC2B | SOX2 regulated genes in Keratinocyte Overexpression from PMID: 30772301 | SOX2 |
| 6332 | HAT1 | SOX2 regulated genes in Keratinocyte Overexpression from PMID: 30772301 | SOX2 |
| 6333 | NDST1 | SOX2 regulated genes in Keratinocyte Overexpression from PMID: 30772301 | SOX2 |
| 6334 | SLC25A13 | SOX2 regulated genes in Keratinocyte Overexpression from PMID: 30772301 | SOX2 |
| 6335 | FAM102A | SOX2 regulated genes in Keratinocyte Overexpression from PMID: 30772301 | SOX2 |
| 6336 | CCND2 | SOX2 regulated genes in Keratinocyte Overexpression from PMID: 30772301 | SOX2 |
| 6337 | PAFAH1B2 | SOX2 regulated genes in Keratinocyte Overexpression from PMID: 30772301 | SOX2 |
| 6338 | UBE2E3 | SOX2 regulated genes in Keratinocyte Overexpression from PMID: 30772301 | SOX2 |
| 6339 | ZNF169 | SOX2 regulated genes in Keratinocyte Overexpression from PMID: 30772301 | SOX2 |
| 6340 | SLC4A2 | SOX2 regulated genes in Keratinocyte Overexpression from PMID: 30772301 | SOX2 |
| 6341 | SLC22A1 | SOX2 regulated genes in Keratinocyte Overexpression from PMID: 30772301 | SOX2 |
| 6342 | SFRP1 | SOX2 regulated genes in Keratinocyte Overexpression from PMID: 30772301 | SOX2 |

| | | | |
|---|---|---|---|
| 6343 | E2F5 | SOX2 regulated genes in Keratinocyte Overexpression from PMID: 30772301 | SOX2 |
| 6344 | CHRM5 | SOX2 regulated genes in Keratinocyte Overexpression from PMID: 30772301 | SOX2 |
| 6345 | GTF3C1 | SOX2 regulated genes in Keratinocyte Overexpression from PMID: 30772301 | SOX2 |
| 6346 | NXT1 | SOX2 regulated genes in Keratinocyte Overexpression from PMID: 30772301 | SOX2 |
| 6347 | TULP1 | SOX2 regulated genes in Keratinocyte Overexpression from PMID: 30772301 | SOX2 |
| 6348 | EPHB6 | SOX2 regulated genes in Keratinocyte Overexpression from PMID: 30772301 | SOX2 |
| 6349 | ATOH8 | SOX2 regulated genes in Keratinocyte Overexpression from PMID: 30772301 | SOX2 |
| 6350 | HAUS5 | SOX2 regulated genes in Keratinocyte Overexpression from PMID: 30772301 | SOX2 |
| 6351 | RABGEF1 | SOX2 regulated genes in Keratinocyte Overexpression from PMID: 30772301 | SOX2 |
| 6352 | ZMYM2 | SOX2 regulated genes in Keratinocyte Overexpression from PMID: 30772301 | SOX2 |
| 6353 | STBD1 | SOX2 regulated genes in Keratinocyte Overexpression from PMID: 30772301 | SOX2 |
| 6354 | ZDHHC6 | SOX2 regulated genes in Keratinocyte Overexpression from PMID: 30772301 | SOX2 |
| 6355 | MAD2L1BP | SOX2 regulated genes in Keratinocyte Overexpression from PMID: 30772301 | SOX2 |
| 6356 | PSMB2 | SOX2 regulated genes in Keratinocyte Overexpression from PMID: 30772301 | SOX2 |
| 6357 | DDX51 | SOX2 regulated genes in Keratinocyte Overexpression from PMID: 30772301 | SOX2 |
| 6358 | MED7 | SOX2 regulated genes in Keratinocyte Overexpression from PMID: 30772301 | SOX2 |
| 6359 | NECTIN3 | SOX2 regulated genes in Keratinocyte Overexpression from PMID: 30772301 | SOX2 |
| 6360 | KRT6A | SOX2 regulated genes in Keratinocyte Overexpression from PMID: 30772301 | SOX2 |
| 6361 | CNOT6 | SOX2 regulated genes in Keratinocyte Overexpression from PMID: 30772301 | SOX2 |
| 6362 | ZBTB39 | SOX2 regulated genes in Keratinocyte Overexpression from PMID: 30772301 | SOX2 |
| 6363 | STAG2 | SOX2 regulated genes in Keratinocyte Overexpression from PMID: 30772301 | SOX2 |
| 6364 | ZSCAN25 | SOX2 regulated genes in Keratinocyte Overexpression from PMID: 30772301 | SOX2 |
| 6365 | RHOG | SOX2 regulated genes in Keratinocyte Overexpression from PMID: 30772301 | SOX2 |
| 6366 | CYP4F12 | SOX2 regulated genes in Keratinocyte Overexpression from PMID: 30772301 | SOX2 |
| 6367 | KAT2B | SOX2 regulated genes in Keratinocyte Overexpression from PMID: 30772301 | SOX2 |
| 6368 | ANAPC2 | SOX2 regulated genes in Keratinocyte Overexpression from PMID: 30772301 | SOX2 |
| 6369 | FRAS1 | SOX2 regulated genes in Keratinocyte Overexpression from PMID: 30772301 | SOX2 |
| 6370 | SLX1B | SOX2 regulated genes in Keratinocyte Overexpression from PMID: 30772301 | SOX2 |
| 6371 | SLX1A | SOX2 regulated genes in Keratinocyte Overexpression from PMID: 30772301 | SOX2 |
| 6372 | CKAP2L | SOX2 regulated genes in Keratinocyte Overexpression from PMID: 30772301 | SOX2 |
| 6373 | CDCP1 | SOX2 regulated genes in Keratinocyte Overexpression from PMID: 30772301 | SOX2 |
| 6374 | NRARP | SOX2 regulated genes in Keratinocyte Overexpression from PMID: 30772301 | SOX2 |
| 6375 | PNO1 | SOX2 regulated genes in Keratinocyte Overexpression from PMID: 30772301 | SOX2 |
| 6376 | VCP | SOX2 regulated genes in Keratinocyte Overexpression from PMID: 30772301 | SOX2 |
| 6377 | HECW2 | SOX2 regulated genes in Keratinocyte Overexpression from PMID: 30772301 | SOX2 |
| 6378 | ABHD17B | SOX2 regulated genes in Keratinocyte Overexpression from PMID: 30772301 | SOX2 |
| 6379 | DNTTIP2 | SOX2 regulated genes in Keratinocyte Overexpression from PMID: 30772301 | SOX2 |
| 6380 | LIPM | SOX2 regulated genes in Keratinocyte Overexpression from PMID: 30772301 | SOX2 |
| 6381 | HOXB8 | SOX2 regulated genes in Keratinocyte Overexpression from PMID: 30772301 | SOX2 |
| 6382 | PARN | SOX2 regulated genes in Keratinocyte Overexpression from PMID: 30772301 | SOX2 |
| 6383 | SCUBE3 | SOX2 regulated genes in Keratinocyte Overexpression from PMID: 30772301 | SOX2 |
| 6384 | RALGAPA1 | SOX2 regulated genes in Keratinocyte Overexpression from PMID: 30772301 | SOX2 |
| 6385 | TIMM10 | SOX2 regulated genes in Keratinocyte Overexpression from PMID: 30772301 | SOX2 |
| 6386 | ANAPC10 | SOX2 regulated genes in Keratinocyte Overexpression from PMID: 30772301 | SOX2 |
| 6387 | CXXC5 | SOX2 regulated genes in Keratinocyte Overexpression from PMID: 30772301 | SOX2 |
| 6388 | CTCF | SOX2 regulated genes in Keratinocyte Overexpression from PMID: 30772301 | SOX2 |
| 6389 | EEF1A1 | SOX2 regulated genes in Keratinocyte Overexpression from PMID: 30772301 | SOX2 |
| 6390 | UACA | SOX2 regulated genes in Keratinocyte Overexpression from PMID: 30772301 | SOX2 |
| 6391 | FAF2 | SOX2 regulated genes in Keratinocyte Overexpression from PMID: 30772301 | SOX2 |
| 6392 | SUB1 | SOX2 regulated genes in Keratinocyte Overexpression from PMID: 30772301 | SOX2 |
| 6393 | SCNN1B | SOX2 regulated genes in Keratinocyte Overexpression from PMID: 30772301 | SOX2 |
| 6394 | WAC | SOX2 regulated genes in Keratinocyte Overexpression from PMID: 30772301 | SOX2 |
| 6395 | SNUPN | SOX2 regulated genes in Keratinocyte Overexpression from PMID: 30772301 | SOX2 |
| 6396 | USP34 | SOX2 regulated genes in Keratinocyte Overexpression from PMID: 30772301 | SOX2 |
| 6397 | SKIDA1 | SOX2 regulated genes in Keratinocyte Overexpression from PMID: 30772301 | SOX2 |
| 6398 | BICD1 | SOX2 regulated genes in Keratinocyte Overexpression from PMID: 30772301 | SOX2 |
| 6399 | TOB1 | SOX2 regulated genes in Keratinocyte Overexpression from PMID: 30772301 | SOX2 |
| 6400 | TANGO6 | SOX2 regulated genes in Keratinocyte Overexpression from PMID: 30772301 | SOX2 |
| 6401 | BAALC | SOX2 regulated genes in Keratinocyte Overexpression from PMID: 30772301 | SOX2 |
| 6402 | FOXK2 | SOX2 regulated genes in Keratinocyte Overexpression from PMID: 30772301 | SOX2 |
| 6403 | FOXA3 | SOX2 regulated genes in Keratinocyte Overexpression from PMID: 30772301 | SOX2 |

| | | | |
|---|---|---|---|
| 6404 | MKRN1 | SOX2 regulated genes in Keratinocyte Overexpression from PMID: 30772301 | SOX2 |
| 6405 | GRM7 | SOX2 regulated genes in Keratinocyte Overexpression from PMID: 30772301 | SOX2 |
| 6406 | PHPT1 | SOX2 regulated genes in Keratinocyte Overexpression from PMID: 30772301 | SOX2 |
| 6407 | CPEB2 | SOX2 regulated genes in Keratinocyte Overexpression from PMID: 30772301 | SOX2 |
| 6408 | YWHAZ | SOX2 regulated genes in Keratinocyte Overexpression from PMID: 30772301 | SOX2 |
| 6409 | RPP25L | SOX2 regulated genes in Keratinocyte Overexpression from PMID: 30772301 | SOX2 |
| 6410 | TMEM11 | SOX2 regulated genes in Keratinocyte Overexpression from PMID: 30772301 | SOX2 |
| 6411 | MRPL11 | SOX2 regulated genes in Keratinocyte Overexpression from PMID: 30772301 | SOX2 |
| 6412 | ZNF276 | SOX2 regulated genes in Keratinocyte Overexpression from PMID: 30772301 | SOX2 |
| 6413 | RALA | SOX2 regulated genes in Keratinocyte Overexpression from PMID: 30772301 | SOX2 |
| 6414 | NUP188 | SOX2 regulated genes in Keratinocyte Overexpression from PMID: 30772301 | SOX2 |
| 6415 | PTPRF | SOX2 regulated genes in Keratinocyte Overexpression from PMID: 30772301 | SOX2 |
| 6416 | MEST | SOX2 regulated genes in Keratinocyte Overexpression from PMID: 30772301 | SOX2 |
| 6417 | POLD4 | SOX2 regulated genes in Keratinocyte Overexpression from PMID: 30772301 | SOX2 |
| 6418 | SLITRK5 | SOX2 regulated genes in Keratinocyte Overexpression from PMID: 30772301 | SOX2 |
| 6419 | N4BP2L2 | SOX2 regulated genes in Keratinocyte Overexpression from PMID: 30772301 | SOX2 |
| 6420 | ACP7 | SOX2 regulated genes in Keratinocyte Overexpression from PMID: 30772301 | SOX2 |
| 6421 | HBP1 | SOX2 regulated genes in Keratinocyte Overexpression from PMID: 30772301 | SOX2 |
| 6422 | LRRTM2 | SOX2 regulated genes in Keratinocyte Overexpression from PMID: 30772301 | SOX2 |
| 6423 | RRN3 | SOX2 regulated genes in Keratinocyte Overexpression from PMID: 30772301 | SOX2 |
| 6424 | HEPH | SOX2 regulated genes in Keratinocyte Overexpression from PMID: 30772301 | SOX2 |
| 6425 | MOB1B | SOX2 regulated genes in Keratinocyte Overexpression from PMID: 30772301 | SOX2 |
| 6426 | BRCC3 | SOX2 regulated genes in Keratinocyte Overexpression from PMID: 30772301 | SOX2 |
| 6427 | OPA3 | SOX2 regulated genes in Keratinocyte Overexpression from PMID: 30772301 | SOX2 |
| 6428 | UBA1 | SOX2 regulated genes in Keratinocyte Overexpression from PMID: 30772301 | SOX2 |
| 6429 | FBXO10 | SOX2 regulated genes in Keratinocyte Overexpression from PMID: 30772301 | SOX2 |
| 6430 | MPZL3 | SOX2 regulated genes in Keratinocyte Overexpression from PMID: 30772301 | SOX2 |
| 6431 | ORMDL2 | SOX2 regulated genes in Keratinocyte Overexpression from PMID: 30772301 | SOX2 |
| 6432 | PRR3 | SOX2 regulated genes in Keratinocyte Overexpression from PMID: 30772301 | SOX2 |
| 6433 | TSPAN3 | SOX2 regulated genes in Keratinocyte Overexpression from PMID: 30772301 | SOX2 |
| 6434 | CBLN3 | SOX2 regulated genes in Keratinocyte Overexpression from PMID: 30772301 | SOX2 |
| 6435 | LETM1 | SOX2 regulated genes in Keratinocyte Overexpression from PMID: 30772301 | SOX2 |
| 6436 | RC3H2 | SOX2 regulated genes in Keratinocyte Overexpression from PMID: 30772301 | SOX2 |
| 6437 | CX3CL1 | SOX2 regulated genes in Keratinocyte Overexpression from PMID: 30772301 | SOX2 |
| 6438 | FLRT2 | SOX2 regulated genes in Keratinocyte Overexpression from PMID: 30772301 | SOX2 |
| 6439 | GEN1 | SOX2 regulated genes in Keratinocyte Overexpression from PMID: 30772301 | SOX2 |
| 6440 | BCL11A | SOX2 regulated genes in Keratinocyte Overexpression from PMID: 30772301 | SOX2 |
| 6441 | TMEM178A | SOX2 regulated genes in Keratinocyte Overexpression from PMID: 30772301 | SOX2 |
| 6442 | EXOSC7 | SOX2 regulated genes in Keratinocyte Overexpression from PMID: 30772301 | SOX2 |
| 6443 | LBH | SOX2 regulated genes in Keratinocyte Overexpression from PMID: 30772301 | SOX2 |
| 6444 | COPS7A | SOX2 regulated genes in Keratinocyte Overexpression from PMID: 30772301 | SOX2 |
| 6445 | RBMX2 | SOX2 regulated genes in Keratinocyte Overexpression from PMID: 30772301 | SOX2 |
| 6446 | NRBP2 | SOX2 regulated genes in Keratinocyte Overexpression from PMID: 30772301 | SOX2 |
| 6447 | KLRG2 | SOX2 regulated genes in Keratinocyte Overexpression from PMID: 30772301 | SOX2 |
| 6448 | DPH3 | SOX2 regulated genes in Keratinocyte Overexpression from PMID: 30772301 | SOX2 |
| 6449 | VPS37D | SOX2 regulated genes in Keratinocyte Overexpression from PMID: 30772301 | SOX2 |
| 6450 | PLG | SOX2 regulated genes in Keratinocyte Overexpression from PMID: 30772301 | SOX2 |
| 6451 | C1QTNF4 | SOX2 regulated genes in Keratinocyte Overexpression from PMID: 30772301 | SOX2 |
| 6452 | CSRP2 | SOX2 regulated genes in Keratinocyte Overexpression from PMID: 30772301 | SOX2 |
| 6453 | EFCAB7 | SOX2 regulated genes in Keratinocyte Overexpression from PMID: 30772301 | SOX2 |
| 6454 | TEAD2 | SOX2 regulated genes in Keratinocyte Overexpression from PMID: 30772301 | SOX2 |
| 6455 | HERC4 | SOX2 regulated genes in Keratinocyte Overexpression from PMID: 30772301 | SOX2 |
| 6456 | GLRX3 | SOX2 regulated genes in Keratinocyte Overexpression from PMID: 30772301 | SOX2 |
| 6457 | SLC16A8 | SOX2 regulated genes in Keratinocyte Overexpression from PMID: 30772301 | SOX2 |
| 6458 | NFIB | SOX2 regulated genes in Keratinocyte Overexpression from PMID: 30772301 | SOX2 |
| 6459 | NUP35 | SOX2 regulated genes in Keratinocyte Overexpression from PMID: 30772301 | SOX2 |
| 6460 | PPARD | SOX2 regulated genes in Keratinocyte Overexpression from PMID: 30772301 | SOX2 |
| 6461 | GLA | SOX2 regulated genes in Keratinocyte Overexpression from PMID: 30772301 | SOX2 |
| 6462 | SRSF12 | SOX2 regulated genes in Keratinocyte Overexpression from PMID: 30772301 | SOX2 |
| 6463 | COPS6 | SOX2 regulated genes in Keratinocyte Overexpression from PMID: 30772301 | SOX2 |
| 6464 | SH3D19 | SOX2 regulated genes in Keratinocyte Overexpression from PMID: 30772301 | SOX2 |

| | | | |
|---|---|---|---|
| 6465 | SYCP3 | SOX2 regulated genes in Keratinocyte Overexpression from PMID: 30772301 | SOX2 |
| 6466 | GZF1 | SOX2 regulated genes in Keratinocyte Overexpression from PMID: 30772301 | SOX2 |
| 6467 | AGAP3 | SOX2 regulated genes in Keratinocyte Overexpression from PMID: 30772301 | SOX2 |
| 6468 | JAG2 | SOX2 regulated genes in Keratinocyte Overexpression from PMID: 30772301 | SOX2 |
| 6469 | TLE4 | SOX2 regulated genes in Keratinocyte Overexpression from PMID: 30772301 | SOX2 |
| 6470 | SNX33 | SOX2 regulated genes in Keratinocyte Overexpression from PMID: 30772301 | SOX2 |
| 6471 | ALKBH4 | SOX2 regulated genes in Keratinocyte Overexpression from PMID: 30772301 | SOX2 |
| 6472 | GPC4 | SOX2 regulated genes in Keratinocyte Overexpression from PMID: 30772301 | SOX2 |
| 6473 | BRAP | SOX2 regulated genes in Keratinocyte Overexpression from PMID: 30772301 | SOX2 |
| 6474 | HS6ST1 | SOX2 regulated genes in Keratinocyte Overexpression from PMID: 30772301 | SOX2 |
| 6475 | MPHOSPH10 | SOX2 regulated genes in Keratinocyte Overexpression from PMID: 30772301 | SOX2 |
| 6476 | KIAA0586 | SOX2 regulated genes in Keratinocyte Overexpression from PMID: 30772301 | SOX2 |
| 6477 | HSF1 | SOX2 regulated genes in Keratinocyte Overexpression from PMID: 30772301 | SOX2 |
| 6478 | DPYSL5 | SOX2 regulated genes in Keratinocyte Overexpression from PMID: 30772301 | SOX2 |
| 6479 | ADO | SOX2 regulated genes in Keratinocyte Overexpression from PMID: 30772301 | SOX2 |
| 6480 | PDIK1L | SOX2 regulated genes in Keratinocyte Overexpression from PMID: 30772301 | SOX2 |
| 6481 | PPFIA3 | SOX2 regulated genes in Keratinocyte Overexpression from PMID: 30772301 | SOX2 |
| 6482 | TSSK1B | SOX2 regulated genes in Keratinocyte Overexpression from PMID: 30772301 | SOX2 |
| 6483 | DUSP6 | SOX2 regulated genes in Keratinocyte Overexpression from PMID: 30772301 | SOX2 |
| 6484 | PPIL4 | SOX2 regulated genes in Keratinocyte Overexpression from PMID: 30772301 | SOX2 |
| 6485 | SMNDC1 | SOX2 regulated genes in Keratinocyte Overexpression from PMID: 30772301 | SOX2 |
| 6486 | SOSTDC1 | SOX2 regulated genes in Keratinocyte Overexpression from PMID: 30772301 | SOX2 |
| 6487 | NUP98 | SOX2 regulated genes in Keratinocyte Overexpression from PMID: 30772301 | SOX2 |
| 6488 | TMEM121 | SOX2 regulated genes in Keratinocyte Overexpression from PMID: 30772301 | SOX2 |
| 6489 | GJB4 | SOX2 regulated genes in Keratinocyte Overexpression from PMID: 30772301 | SOX2 |
| 6490 | RPL37 | SOX2 regulated genes in Keratinocyte Overexpression from PMID: 30772301 | SOX2 |
| 6491 | MAP9 | SOX2 regulated genes in Keratinocyte Overexpression from PMID: 30772301 | SOX2 |
| 6492 | MTTP | SOX2 regulated genes in Keratinocyte Overexpression from PMID: 30772301 | SOX2 |
| 6493 | IQCK | SOX2 regulated genes in Keratinocyte Overexpression from PMID: 30772301 | SOX2 |
| 6494 | PPP2R5D | SOX2 regulated genes in Keratinocyte Overexpression from PMID: 30772301 | SOX2 |
| 6495 | NUP37 | SOX2 regulated genes in Keratinocyte Overexpression from PMID: 30772301 | SOX2 |
| 6496 | MLLT10 | SOX2 regulated genes in Keratinocyte Overexpression from PMID: 30772301 | SOX2 |
| 6497 | FNTB | SOX2 regulated genes in Keratinocyte Overexpression from PMID: 30772301 | SOX2 |
| 6498 | PPP1R15A | SOX2 regulated genes in Keratinocyte Overexpression from PMID: 30772301 | SOX2 |
| 6499 | PNPLA1 | SOX2 regulated genes in Keratinocyte Overexpression from PMID: 30772301 | SOX2 |
| 6500 | RPL22L1 | SOX2 regulated genes in Keratinocyte Overexpression from PMID: 30772301 | SOX2 |
| 6501 | NRN1L | SOX2 regulated genes in Keratinocyte Overexpression from PMID: 30772301 | SOX2 |
| 6502 | EEFSEC | SOX2 regulated genes in Keratinocyte Overexpression from PMID: 30772301 | SOX2 |
| 6503 | PPID | SOX2 regulated genes in Keratinocyte Overexpression from PMID: 30772301 | SOX2 |
| 6504 | ARMC10 | SOX2 regulated genes in Keratinocyte Overexpression from PMID: 30772301 | SOX2 |
| 6505 | ZBTB45 | SOX2 regulated genes in Keratinocyte Overexpression from PMID: 30772301 | SOX2 |
| 6506 | NXPE2 | SOX2 regulated genes in Keratinocyte Overexpression from PMID: 30772301 | SOX2 |
| 6507 | PSMD14 | SOX2 regulated genes in Keratinocyte Overexpression from PMID: 30772301 | SOX2 |
| 6508 | KIF20B | SOX2 regulated genes in Keratinocyte Overexpression from PMID: 30772301 | SOX2 |
| 6509 | TMEM161A | SOX2 regulated genes in Keratinocyte Overexpression from PMID: 30772301 | SOX2 |
| 6510 | CCDC112 | SOX2 regulated genes in Keratinocyte Overexpression from PMID: 30772301 | SOX2 |
| 6511 | PRRG3 | SOX2 regulated genes in Keratinocyte Overexpression from PMID: 30772301 | SOX2 |
| 6512 | PUS3 | SOX2 regulated genes in Keratinocyte Overexpression from PMID: 30772301 | SOX2 |
| 6513 | SYNE2 | SOX2 regulated genes in Keratinocyte Overexpression from PMID: 30772301 | SOX2 |
| 6514 | ZBTB12 | SOX2 regulated genes in Keratinocyte Overexpression from PMID: 30772301 | SOX2 |
| 6515 | REXO4 | SOX2 regulated genes in Keratinocyte Overexpression from PMID: 30772301 | SOX2 |
| 6516 | WDR31 | SOX2 regulated genes in Keratinocyte Overexpression from PMID: 30772301 | SOX2 |
| 6517 | AP1S3 | SOX2 regulated genes in Keratinocyte Overexpression from PMID: 30772301 | SOX2 |
| 6518 | GTF3C5 | SOX2 regulated genes in Keratinocyte Overexpression from PMID: 30772301 | SOX2 |
| 6519 | PTK2 | SOX2 regulated genes in Keratinocyte Overexpression from PMID: 30772301 | SOX2 |
| 6520 | LRFN1 | SOX2 regulated genes in Keratinocyte Overexpression from PMID: 30772301 | SOX2 |
| 6521 | CCNC | SOX2 regulated genes in Keratinocyte Overexpression from PMID: 30772301 | SOX2 |
| 6522 | PLA2G12A | SOX2 regulated genes in Keratinocyte Overexpression from PMID: 30772301 | SOX2 |
| 6523 | ARID1B | SOX2 regulated genes in Keratinocyte Overexpression from PMID: 30772301 | SOX2 |
| 6524 | SOCS5 | SOX2 regulated genes in Keratinocyte Overexpression from PMID: 30772301 | SOX2 |
| 6525 | FANCA | SOX2 regulated genes in Keratinocyte Overexpression from PMID: 30772301 | SOX2 |

| | | | |
|---|---|---|---|
| 6526 | NAXD | SOX2 regulated genes in Keratinocyte Overexpression from PMID: 30772301 | SOX2 |
| 6527 | CLIC3 | SOX2 regulated genes in Keratinocyte Overexpression from PMID: 30772301 | SOX2 |
| 6528 | ABHD10 | SOX2 regulated genes in Keratinocyte Overexpression from PMID: 30772301 | SOX2 |
| 6529 | INSM2 | SOX2 regulated genes in Keratinocyte Overexpression from PMID: 30772301 | SOX2 |
| 6530 | MYCBPAP | SOX2 regulated genes in Keratinocyte Overexpression from PMID: 30772301 | SOX2 |
| 6531 | TEAD3 | SOX2 regulated genes in Keratinocyte Overexpression from PMID: 30772301 | SOX2 |
| 6532 | PTK2B | SOX2 regulated genes in Keratinocyte Overexpression from PMID: 30772301 | SOX2 |
| 6533 | TCF7L1 | SOX2 regulated genes in Keratinocyte Overexpression from PMID: 30772301 | SOX2 |
| 6534 | CBX1 | SOX2 regulated genes in Keratinocyte Overexpression from PMID: 30772301 | SOX2 |
| 6535 | MDGA1 | SOX2 regulated genes in Keratinocyte Overexpression from PMID: 30772301 | SOX2 |
| 6536 | CLDND1 | SOX2 regulated genes in Keratinocyte Overexpression from PMID: 30772301 | SOX2 |
| 6537 | CENPH | SOX2 regulated genes in Keratinocyte Overexpression from PMID: 30772301 | SOX2 |
| 6538 | PLEKHF2 | SOX2 regulated genes in Keratinocyte Overexpression from PMID: 30772301 | SOX2 |
| 6539 | DNAJB13 | SOX2 regulated genes in Keratinocyte Overexpression from PMID: 30772301 | SOX2 |
| 6540 | LUC7L3 | SOX2 regulated genes in Keratinocyte Overexpression from PMID: 30772301 | SOX2 |
| 6541 | ARHGEF1 | SOX2 regulated genes in Keratinocyte Overexpression from PMID: 30772301 | SOX2 |
| 6542 | PLOD2 | SOX2 regulated genes in Keratinocyte Overexpression from PMID: 30772301 | SOX2 |
| 6543 | NPRL2 | SOX2 regulated genes in Keratinocyte Overexpression from PMID: 30772301 | SOX2 |
| 6544 | METTL2B | SOX2 regulated genes in Keratinocyte Overexpression from PMID: 30772301 | SOX2 |
| 6545 | TUBE1 | SOX2 regulated genes in Keratinocyte Overexpression from PMID: 30772301 | SOX2 |
| 6546 | C1QTNF12 | SOX2 regulated genes in Keratinocyte Overexpression from PMID: 30772301 | SOX2 |
| 6547 | BORCS8 | SOX2 regulated genes in Keratinocyte Overexpression from PMID: 30772301 | SOX2 |
| 6548 | SLCO4C1 | SOX2 regulated genes in Keratinocyte Overexpression from PMID: 30772301 | SOX2 |
| 6549 | LIG4 | SOX2 regulated genes in Keratinocyte Overexpression from PMID: 30772301 | SOX2 |
| 6550 | MEPCE | SOX2 regulated genes in Keratinocyte Overexpression from PMID: 30772301 | SOX2 |
| 6551 | CYFIP2 | SOX2 regulated genes in Keratinocyte Overexpression from PMID: 30772301 | SOX2 |
| 6552 | CD151 | SOX2 regulated genes in Keratinocyte Overexpression from PMID: 30772301 | SOX2 |
| 6553 | LYRM4 | SOX2 regulated genes in Keratinocyte Overexpression from PMID: 30772301 | SOX2 |
| 6554 | TP53I11 | SOX2 regulated genes in Keratinocyte Overexpression from PMID: 30772301 | SOX2 |
| 6555 | CTBP1 | SOX2 regulated genes in Keratinocyte Overexpression from PMID: 30772301 | SOX2 |
| 6556 | ARL6 | SOX2 regulated genes in Keratinocyte Overexpression from PMID: 30772301 | SOX2 |
| 6557 | LRWD1 | SOX2 regulated genes in Keratinocyte Overexpression from PMID: 30772301 | SOX2 |
| 6558 | HDHD3 | SOX2 regulated genes in Keratinocyte Overexpression from PMID: 30772301 | SOX2 |
| 6559 | ZC3HAV1L | SOX2 regulated genes in Keratinocyte Overexpression from PMID: 30772301 | SOX2 |
| 6560 | STAM | SOX2 regulated genes in Keratinocyte Overexpression from PMID: 30772301 | SOX2 |
| 6561 | URI1 | SOX2 regulated genes in Keratinocyte Overexpression from PMID: 30772301 | SOX2 |
| 6562 | MRPL2 | SOX2 regulated genes in Keratinocyte Overexpression from PMID: 30772301 | SOX2 |
| 6563 | GABARAPL2 | SOX2 regulated genes in Keratinocyte Overexpression from PMID: 30772301 | SOX2 |
| 6564 | RARG | SOX2 regulated genes in Keratinocyte Overexpression from PMID: 30772301 | SOX2 |
| 6565 | NXF1 | SOX2 regulated genes in Keratinocyte Overexpression from PMID: 30772301 | SOX2 |
| 6566 | VWA7 | SOX2 regulated genes in Keratinocyte Overexpression from PMID: 30772301 | SOX2 |
| 6567 | BBS12 | SOX2 regulated genes in Keratinocyte Overexpression from PMID: 30772301 | SOX2 |
| 6568 | SESN2 | SOX2 regulated genes in Keratinocyte Overexpression from PMID: 30772301 | SOX2 |
| 6569 | PYURF | SOX2 regulated genes in Keratinocyte Overexpression from PMID: 30772301 | SOX2 |
| 6570 | FAM166A | SOX2 regulated genes in Keratinocyte Overexpression from PMID: 30772301 | SOX2 |
| 6571 | GLB1L2 | SOX2 regulated genes in Keratinocyte Overexpression from PMID: 30772301 | SOX2 |
| 6572 | PSMC2 | SOX2 regulated genes in Keratinocyte Overexpression from PMID: 30772301 | SOX2 |
| 6573 | NSMCE4A | SOX2 regulated genes in Keratinocyte Overexpression from PMID: 30772301 | SOX2 |
| 6574 | CLUAP1 | SOX2 regulated genes in Keratinocyte Overexpression from PMID: 30772301 | SOX2 |
| 6575 | JMJD7 | SOX2 regulated genes in Keratinocyte Overexpression from PMID: 30772301 | SOX2 |
| 6576 | CDC42BPG | SOX2 regulated genes in Keratinocyte Overexpression from PMID: 30772301 | SOX2 |
| 6577 | MEA1 | SOX2 regulated genes in Keratinocyte Overexpression from PMID: 30772301 | SOX2 |
| 6578 | CDK17 | SOX2 regulated genes in Keratinocyte Overexpression from PMID: 30772301 | SOX2 |
| 6579 | ATAD5 | SOX2 regulated genes in Keratinocyte Overexpression from PMID: 30772301 | SOX2 |
| 6580 | DDX19B | SOX2 regulated genes in Keratinocyte Overexpression from PMID: 30772301 | SOX2 |
| 6581 | CCDC189 | SOX2 regulated genes in Keratinocyte Overexpression from PMID: 30772301 | SOX2 |
| 6582 | ADAMTS13 | SOX2 regulated genes in Keratinocyte Overexpression from PMID: 30772301 | SOX2 |
| 6583 | EMC4 | SOX2 regulated genes in Keratinocyte Overexpression from PMID: 30772301 | SOX2 |
| 6584 | ZNF597 | SOX2 regulated genes in Keratinocyte Overexpression from PMID: 30772301 | SOX2 |
| 6585 | CEP131 | SOX2 regulated genes in Keratinocyte Overexpression from PMID: 30772301 | SOX2 |
| 6586 | KLHL2 | SOX2 regulated genes in Keratinocyte Overexpression from PMID: 30772301 | SOX2 |

| | | | |
|---|---|---|---|
| 6587 | FRA10AC1 | SOX2 regulated genes in Keratinocyte Overexpression from PMID: 30772301 | SOX2 |
| 6588 | RPS6KA3 | SOX2 regulated genes in Keratinocyte Overexpression from PMID: 30772301 | SOX2 |
| 6589 | PSMA3 | SOX2 regulated genes in Keratinocyte Overexpression from PMID: 30772301 | SOX2 |
| 6590 | CDC26 | SOX2 regulated genes in Keratinocyte Overexpression from PMID: 30772301 | SOX2 |
| 6591 | SPSB3 | SOX2 regulated genes in Keratinocyte Overexpression from PMID: 30772301 | SOX2 |
| 6592 | GPATCH1 | SOX2 regulated genes in Keratinocyte Overexpression from PMID: 30772301 | SOX2 |
| 6593 | TPO | SOX2 regulated genes in Keratinocyte Overexpression from PMID: 30772301 | SOX2 |
| 6594 | RUFY3 | SOX2 regulated genes in Keratinocyte Overexpression from PMID: 30772301 | SOX2 |
| 6595 | PRPF4 | SOX2 regulated genes in Keratinocyte Overexpression from PMID: 30772301 | SOX2 |
| 6596 | STRN | SOX2 regulated genes in Keratinocyte Overexpression from PMID: 30772301 | SOX2 |
| 6597 | LMNTD2 | SOX2 regulated genes in Keratinocyte Overexpression from PMID: 30772301 | SOX2 |
| 6598 | NCAPG2 | SOX2 regulated genes in Keratinocyte Overexpression from PMID: 30772301 | SOX2 |
| 6599 | CEP170B | SOX2 regulated genes in Keratinocyte Overexpression from PMID: 30772301 | SOX2 |
| 6600 | XPO5 | SOX2 regulated genes in Keratinocyte Overexpression from PMID: 30772301 | SOX2 |
| 6601 | BOK | SOX2 regulated genes in Keratinocyte Overexpression from PMID: 30772301 | SOX2 |
| 6602 | UTP20 | SOX2 regulated genes in Keratinocyte Overexpression from PMID: 30772301 | SOX2 |
| 6603 | CRYBB3 | SOX2 regulated genes in Keratinocyte Overexpression from PMID: 30772301 | SOX2 |
| 6604 | ARF5 | SOX2 regulated genes in Keratinocyte Overexpression from PMID: 30772301 | SOX2 |
| 6605 | CTSF | SOX2 regulated genes in Keratinocyte Overexpression from PMID: 30772301 | SOX2 |
| 6606 | AVEN | SOX2 regulated genes in Keratinocyte Overexpression from PMID: 30772301 | SOX2 |
| 6607 | SFTPD | SOX2 regulated genes in Keratinocyte Overexpression from PMID: 30772301 | SOX2 |
| 6608 | GNMT | SOX2 regulated genes in Keratinocyte Overexpression from PMID: 30772301 | SOX2 |
| 6609 | ZMYND8 | SOX2 regulated genes in Keratinocyte Overexpression from PMID: 30772301 | SOX2 |
| 6610 | TRIM32 | SOX2 regulated genes in Keratinocyte Overexpression from PMID: 30772301 | SOX2 |
| 6611 | CENPK | SOX2 regulated genes in Keratinocyte Overexpression from PMID: 30772301 | SOX2 |
| 6612 | ZSCAN22 | SOX2 regulated genes in Keratinocyte Overexpression from PMID: 30772301 | SOX2 |
| 6613 | C3orf18 | SOX2 regulated genes in Keratinocyte Overexpression from PMID: 30772301 | SOX2 |
| 6614 | AGO2 | SOX2 regulated genes in Keratinocyte Overexpression from PMID: 30772301 | SOX2 |
| 6615 | NPHP4 | SOX2 regulated genes in Keratinocyte Overexpression from PMID: 30772301 | SOX2 |
| 6616 | IRX6 | SOX2 regulated genes in Keratinocyte Overexpression from PMID: 30772301 | SOX2 |
| 6617 | MRPS10 | SOX2 regulated genes in Keratinocyte Overexpression from PMID: 30772301 | SOX2 |
| 6618 | OIP5 | SOX2 regulated genes in Keratinocyte Overexpression from PMID: 30772301 | SOX2 |
| 6619 | CEP55 | SOX2 regulated genes in Keratinocyte Overexpression from PMID: 30772301 | SOX2 |
| 6620 | GAPVD1 | SOX2 regulated genes in Keratinocyte Overexpression from PMID: 30772301 | SOX2 |
| 6621 | GPC1 | SOX2 regulated genes in Keratinocyte Overexpression from PMID: 30772301 | SOX2 |
| 6622 | SSC4D | SOX2 regulated genes in Keratinocyte Overexpression from PMID: 30772301 | SOX2 |
| 6623 | UBE2N | SOX2 regulated genes in Keratinocyte Overexpression from PMID: 30772301 | SOX2 |
| 6624 | UBN2 | SOX2 regulated genes in Keratinocyte Overexpression from PMID: 30772301 | SOX2 |
| 6625 | CDC40 | SOX2 regulated genes in Keratinocyte Overexpression from PMID: 30772301 | SOX2 |
| 6626 | YPEL5 | SOX2 regulated genes in Keratinocyte Overexpression from PMID: 30772301 | SOX2 |
| 6627 | NEK4 | SOX2 regulated genes in Keratinocyte Overexpression from PMID: 30772301 | SOX2 |
| 6628 | TIMM9 | SOX2 regulated genes in Keratinocyte Overexpression from PMID: 30772301 | SOX2 |
| 6629 | TRIM9 | SOX2 regulated genes in Keratinocyte Overexpression from PMID: 30772301 | SOX2 |
| 6630 | DDX31 | SOX2 regulated genes in Keratinocyte Overexpression from PMID: 30772301 | SOX2 |
| 6631 | MYH9 | SOX2 regulated genes in Keratinocyte Overexpression from PMID: 30772301 | SOX2 |
| 6632 | TTPA | SOX2 regulated genes in Keratinocyte Overexpression from PMID: 30772301 | SOX2 |
| 6633 | PARG | SOX2 regulated genes in Keratinocyte Overexpression from PMID: 30772301 | SOX2 |
| 6634 | OTOA | SOX2 regulated genes in Keratinocyte Overexpression from PMID: 30772301 | SOX2 |
| 6635 | RGS9 | SOX2 regulated genes in Keratinocyte Overexpression from PMID: 30772301 | SOX2 |
| 6636 | MYRFL | SOX2 regulated genes in Keratinocyte Overexpression from PMID: 30772301 | SOX2 |
| 6637 | SALL2 | SOX2 regulated genes in Keratinocyte Overexpression from PMID: 30772301 | SOX2 |
| 6638 | TTLL7 | SOX2 regulated genes in Keratinocyte Overexpression from PMID: 30772301 | SOX2 |
| 6639 | POU5F2 | SOX2 regulated genes in Keratinocyte Overexpression from PMID: 30772301 | SOX2 |
| 6640 | NDUFAF4 | SOX2 regulated genes in Keratinocyte Overexpression from PMID: 30772301 | SOX2 |
| 6641 | IGF1R | SOX2 regulated genes in Keratinocyte Overexpression from PMID: 30772301 | SOX2 |
| 6642 | VASN | SOX2 regulated genes in Keratinocyte Overexpression from PMID: 30772301 | SOX2 |
| 6643 | CASKIN1 | SOX2 regulated genes in Keratinocyte Overexpression from PMID: 30772301 | SOX2 |
| 6644 | HERPUD2 | SOX2 regulated genes in Keratinocyte Overexpression from PMID: 30772301 | SOX2 |
| 6645 | OXCT2 | SOX2 regulated genes in Keratinocyte Overexpression from PMID: 30772301 | SOX2 |
| 6646 | SLC4A10 | SOX2 regulated genes in Keratinocyte Overexpression from PMID: 30772301 | SOX2 |
| 6647 | APPL2 | SOX2 regulated genes in Keratinocyte Overexpression from PMID: 30772301 | SOX2 |

| | | | |
|---|---|---|---|
| 6648 | OMD | SOX2 regulated genes in Keratinocyte Overexpression from PMID: 30772301 | SOX2 |
| 6649 | WNK3 | SOX2 regulated genes in Keratinocyte Overexpression from PMID: 30772301 | SOX2 |
| 6650 | SNX18 | SOX2 regulated genes in Keratinocyte Overexpression from PMID: 30772301 | SOX2 |
| 6651 | DPM2 | SOX2 regulated genes in Keratinocyte Overexpression from PMID: 30772301 | SOX2 |
| 6652 | KIAA1191 | SOX2 regulated genes in Keratinocyte Overexpression from PMID: 30772301 | SOX2 |
| 6653 | RFC2 | SOX2 regulated genes in Keratinocyte Overexpression from PMID: 30772301 | SOX2 |
| 6654 | BTG1 | SOX2 regulated genes in Keratinocyte Overexpression from PMID: 30772301 | SOX2 |
| 6655 | ATP11A | SOX2 regulated genes in Keratinocyte Overexpression from PMID: 30772301 | SOX2 |
| 6656 | MPLKIP | SOX2 regulated genes in Keratinocyte Overexpression from PMID: 30772301 | SOX2 |
| 6657 | SPNS1 | SOX2 regulated genes in Keratinocyte Overexpression from PMID: 30772301 | SOX2 |
| 6658 | SAP18 | SOX2 regulated genes in Keratinocyte Overexpression from PMID: 30772301 | SOX2 |
| 6659 | JRK | SOX2 regulated genes in Keratinocyte Overexpression from PMID: 30772301 | SOX2 |
| 6660 | CAPN13 | SOX2 regulated genes in Keratinocyte Overexpression from PMID: 30772301 | SOX2 |
| 6661 | EIF4G2 | SOX2 regulated genes in Keratinocyte Overexpression from PMID: 30772301 | SOX2 |
| 6662 | RNFT1 | SOX2 regulated genes in Keratinocyte Overexpression from PMID: 30772301 | SOX2 |
| 6663 | COQ7 | SOX2 regulated genes in Keratinocyte Overexpression from PMID: 30772301 | SOX2 |
| 6664 | DNAAF2 | SOX2 regulated genes in Keratinocyte Overexpression from PMID: 30772301 | SOX2 |
| 6665 | BRIX1 | SOX2 regulated genes in Keratinocyte Overexpression from PMID: 30772301 | SOX2 |
| 6666 | PAF1 | SOX2 regulated genes in Keratinocyte Overexpression from PMID: 30772301 | SOX2 |
| 6667 | BFSP2 | SOX2 regulated genes in Keratinocyte Overexpression from PMID: 30772301 | SOX2 |
| 6668 | PPP1R36 | SOX2 regulated genes in Keratinocyte Overexpression from PMID: 30772301 | SOX2 |
| 6669 | ARMC1 | SOX2 regulated genes in Keratinocyte Overexpression from PMID: 30772301 | SOX2 |
| 6670 | C11orf52 | SOX2 regulated genes in Keratinocyte Overexpression from PMID: 30772301 | SOX2 |
| 6671 | CEMIP | SOX2 regulated genes in Keratinocyte Overexpression from PMID: 30772301 | SOX2 |
| 6672 | CPEB4 | SOX2 regulated genes in Keratinocyte Overexpression from PMID: 30772301 | SOX2 |
| 6673 | ZNF593 | SOX2 regulated genes in Keratinocyte Overexpression from PMID: 30772301 | SOX2 |
| 6674 | PRSS8 | SOX2 regulated genes in Keratinocyte Overexpression from PMID: 30772301 | SOX2 |
| 6675 | HDAC11 | SOX2 regulated genes in Keratinocyte Overexpression from PMID: 30772301 | SOX2 |
| 6676 | PHF14 | SOX2 regulated genes in Keratinocyte Overexpression from PMID: 30772301 | SOX2 |
| 6677 | CETN2 | SOX2 regulated genes in Keratinocyte Overexpression from PMID: 30772301 | SOX2 |
| 6678 | TBRG1 | SOX2 regulated genes in Keratinocyte Overexpression from PMID: 30772301 | SOX2 |
| 6679 | ERCC3 | SOX2 regulated genes in Keratinocyte Overexpression from PMID: 30772301 | SOX2 |
| 6680 | CDK5 | SOX2 regulated genes in Keratinocyte Overexpression from PMID: 30772301 | SOX2 |
| 6681 | EYA3 | SOX2 regulated genes in Keratinocyte Overexpression from PMID: 30772301 | SOX2 |
| 6682 | CAMTA1 | SOX2 regulated genes in Keratinocyte Overexpression from PMID: 30772301 | SOX2 |
| 6683 | POP7 | SOX2 regulated genes in Keratinocyte Overexpression from PMID: 30772301 | SOX2 |
| 6684 | HNRNPLL | SOX2 regulated genes in Keratinocyte Overexpression from PMID: 30772301 | SOX2 |
| 6685 | ZFP92 | SOX2 regulated genes in Keratinocyte Overexpression from PMID: 30772301 | SOX2 |
| 6686 | TMEM63B | SOX2 regulated genes in Keratinocyte Overexpression from PMID: 30772301 | SOX2 |
| 6687 | ZNF473 | SOX2 regulated genes in Keratinocyte Overexpression from PMID: 30772301 | SOX2 |
| 6688 | TNFAIP8 | SOX2 regulated genes in Keratinocyte Overexpression from PMID: 30772301 | SOX2 |
| 6689 | NLN | SOX2 regulated genes in Keratinocyte Overexpression from PMID: 30772301 | SOX2 |
| 6690 | CNGA1 | SOX2 regulated genes in Keratinocyte Overexpression from PMID: 30772301 | SOX2 |
| 6691 | LDB1 | SOX2 regulated genes in Keratinocyte Overexpression from PMID: 30772301 | SOX2 |
| 6692 | UBAP1 | SOX2 regulated genes in Keratinocyte Overexpression from PMID: 30772301 | SOX2 |
| 6693 | TFAP4 | SOX2 regulated genes in Keratinocyte Overexpression from PMID: 30772301 | SOX2 |
| 6694 | YTHDC1 | SOX2 regulated genes in Keratinocyte Overexpression from PMID: 30772301 | SOX2 |
| 6695 | SIAH2 | SOX2 regulated genes in Keratinocyte Overexpression from PMID: 30772301 | SOX2 |
| 6696 | PLS3 | SOX2 regulated genes in Keratinocyte Overexpression from PMID: 30772301 | SOX2 |
| 6697 | LZTR1 | SOX2 regulated genes in Keratinocyte Overexpression from PMID: 30772301 | SOX2 |
| 6698 | ZNF827 | SOX2 regulated genes in Keratinocyte Overexpression from PMID: 30772301 | SOX2 |
| 6699 | NAP1L1 | SOX2 regulated genes in Keratinocyte Overexpression from PMID: 30772301 | SOX2 |
| 6700 | CRISPLD1 | SOX2 regulated genes in Keratinocyte Overexpression from PMID: 30772301 | SOX2 |
| 6701 | EIF3F | SOX2 regulated genes in Keratinocyte Overexpression from PMID: 30772301 | SOX2 |
| 6702 | KIF23 | SOX2 regulated genes in Keratinocyte Overexpression from PMID: 30772301 | SOX2 |
| 6703 | RRP36 | SOX2 regulated genes in Keratinocyte Overexpression from PMID: 30772301 | SOX2 |
| 6704 | TRIM31 | SOX2 regulated genes in Keratinocyte Overexpression from PMID: 30772301 | SOX2 |
| 6705 | TAF6L | SOX2 regulated genes in Keratinocyte Overexpression from PMID: 30772301 | SOX2 |
| 6706 | ITGB6 | SOX2 regulated genes in Keratinocyte Overexpression from PMID: 30772301 | SOX2 |
| 6707 | ARMCX5 | SOX2 regulated genes in Keratinocyte Overexpression from PMID: 30772301 | SOX2 |
| 6708 | BTBD11 | SOX2 regulated genes in Keratinocyte Overexpression from PMID: 30772301 | SOX2 |

| | | | |
|---|---|---|---|
| 6709 | EIF1AD | SOX2 regulated genes in Keratinocyte Overexpression from PMID: 30772301 | SOX2 |
| 6710 | ACRBP | SOX2 regulated genes in Keratinocyte Overexpression from PMID: 30772301 | SOX2 |
| 6711 | METTL3 | SOX2 regulated genes in Keratinocyte Overexpression from PMID: 30772301 | SOX2 |
| 6712 | IGF2BP1 | SOX2 regulated genes in Keratinocyte Overexpression from PMID: 30772301 | SOX2 |
| 6713 | GPR176 | SOX2 regulated genes in Keratinocyte Overexpression from PMID: 30772301 | SOX2 |
| 6714 | PDZD4 | SOX2 regulated genes in Keratinocyte Overexpression from PMID: 30772301 | SOX2 |
| 6715 | TRMO | SOX2 regulated genes in Keratinocyte Overexpression from PMID: 30772301 | SOX2 |
| 6716 | KDM4B | SOX2 regulated genes in Keratinocyte Overexpression from PMID: 30772301 | SOX2 |
| 6717 | FAIM2 | SOX2 regulated genes in Keratinocyte Overexpression from PMID: 30772301 | SOX2 |
| 6718 | DMGDH | SOX2 regulated genes in Keratinocyte Overexpression from PMID: 30772301 | SOX2 |
| 6719 | ZNF101 | SOX2 regulated genes in Keratinocyte Overexpression from PMID: 30772301 | SOX2 |
| 6720 | MT4 | SOX2 regulated genes in Keratinocyte Overexpression from PMID: 30772301 | SOX2 |
| 6721 | ATXN2 | SOX2 regulated genes in Keratinocyte Overexpression from PMID: 30772301 | SOX2 |
| 6722 | XRCC2 | SOX2 regulated genes in Keratinocyte Overexpression from PMID: 30772301 | SOX2 |
| 6723 | CNOT11 | SOX2 regulated genes in Keratinocyte Overexpression from PMID: 30772301 | SOX2 |
| 6724 | RAVER2 | SOX2 regulated genes in Keratinocyte Overexpression from PMID: 30772301 | SOX2 |
| 6725 | UBE2V2 | SOX2 regulated genes in Keratinocyte Overexpression from PMID: 30772301 | SOX2 |
| 6726 | MPHOSPH6 | SOX2 regulated genes in Keratinocyte Overexpression from PMID: 30772301 | SOX2 |
| 6727 | THRAP3 | SOX2 regulated genes in Keratinocyte Overexpression from PMID: 30772301 | SOX2 |
| 6728 | FAM47E | SOX2 regulated genes in Keratinocyte Overexpression from PMID: 30772301 | SOX2 |
| 6729 | STK39 | SOX2 regulated genes in Keratinocyte Overexpression from PMID: 30772301 | SOX2 |
| 6730 | TPM4 | SOX2 regulated genes in Keratinocyte Overexpression from PMID: 30772301 | SOX2 |
| 6731 | TDRD12 | SOX2 regulated genes in Keratinocyte Overexpression from PMID: 30772301 | SOX2 |
| 6732 | VAMP3 | SOX2 regulated genes in Keratinocyte Overexpression from PMID: 30772301 | SOX2 |
| 6733 | ALDH18A1 | SOX2 regulated genes in Keratinocyte Overexpression from PMID: 30772301 | SOX2 |
| 6734 | ZDHHC24 | SOX2 regulated genes in Keratinocyte Overexpression from PMID: 30772301 | SOX2 |
| 6735 | CDCA7 | SOX2 regulated genes in Keratinocyte Overexpression from PMID: 30772301 | SOX2 |
| 6736 | PIGT | SOX2 regulated genes in Keratinocyte Overexpression from PMID: 30772301 | SOX2 |
| 6737 | EMD | SOX2 regulated genes in Keratinocyte Overexpression from PMID: 30772301 | SOX2 |
| 6738 | WDR18 | SOX2 regulated genes in Keratinocyte Overexpression from PMID: 30772301 | SOX2 |
| 6739 | SLC52A2 | SOX2 regulated genes in Keratinocyte Overexpression from PMID: 30772301 | SOX2 |
| 6740 | DENR | SOX2 regulated genes in Keratinocyte Overexpression from PMID: 30772301 | SOX2 |
| 6741 | GLT1D1 | SOX2 regulated genes in Keratinocyte Overexpression from PMID: 30772301 | SOX2 |
| 6742 | LCAT | SOX2 regulated genes in Keratinocyte Overexpression from PMID: 30772301 | SOX2 |
| 6743 | CRYBG3 | SOX2 regulated genes in Keratinocyte Overexpression from PMID: 30772301 | SOX2 |
| 6744 | TMEM102 | SOX2 regulated genes in Keratinocyte Overexpression from PMID: 30772301 | SOX2 |
| 6745 | C2orf49 | SOX2 regulated genes in Keratinocyte Overexpression from PMID: 30772301 | SOX2 |
| 6746 | CASC3 | SOX2 regulated genes in Keratinocyte Overexpression from PMID: 30772301 | SOX2 |
| 6747 | SLC35B1 | SOX2 regulated genes in Keratinocyte Overexpression from PMID: 30772301 | SOX2 |
| 6748 | MAPK13 | SOX2 regulated genes in Keratinocyte Overexpression from PMID: 30772301 | SOX2 |
| 6749 | JOSD1 | SOX2 regulated genes in Keratinocyte Overexpression from PMID: 30772301 | SOX2 |
| 6750 | ASIC1 | SOX2 regulated genes in Keratinocyte Overexpression from PMID: 30772301 | SOX2 |
| 6751 | PCSK1 | SOX2 regulated genes in Keratinocyte Overexpression from PMID: 30772301 | SOX2 |
| 6752 | CEACAM21 | SOX2 regulated genes in Keratinocyte Overexpression from PMID: 30772301 | SOX2 |
| 6753 | UTP6 | SOX2 regulated genes in Keratinocyte Overexpression from PMID: 30772301 | SOX2 |
| 6754 | TOMM70 | SOX2 regulated genes in Keratinocyte Overexpression from PMID: 30772301 | SOX2 |
| 6755 | LCOR | SOX2 regulated genes in Keratinocyte Overexpression from PMID: 30772301 | SOX2 |
| 6756 | KIF24 | SOX2 regulated genes in Keratinocyte Overexpression from PMID: 30772301 | SOX2 |
| 6757 | CCDC61 | SOX2 regulated genes in Keratinocyte Overexpression from PMID: 30772301 | SOX2 |
| 6758 | RRM1 | SOX2 regulated genes in Keratinocyte Overexpression from PMID: 30772301 | SOX2 |
| 6759 | TRPV5 | SOX2 regulated genes in Keratinocyte Overexpression from PMID: 30772301 | SOX2 |
| 6760 | NEMP1 | SOX2 regulated genes in Keratinocyte Overexpression from PMID: 30772301 | SOX2 |
| 6761 | SBF1 | SOX2 regulated genes in Keratinocyte Overexpression from PMID: 30772301 | SOX2 |
| 6762 | CAPN11 | SOX2 regulated genes in Keratinocyte Overexpression from PMID: 30772301 | SOX2 |
| 6763 | WNT6 | SOX2 regulated genes in Keratinocyte Overexpression from PMID: 30772301 | SOX2 |
| 6764 | SPOCK3 | SOX2 regulated genes in Keratinocyte Overexpression from PMID: 30772301 | SOX2 |
| 6765 | CTRL | SOX2 regulated genes in Keratinocyte Overexpression from PMID: 30772301 | SOX2 |
| 6766 | SMO | SOX2 regulated genes in Keratinocyte Overexpression from PMID: 30772301 | SOX2 |
| 6767 | KNOP1 | SOX2 regulated genes in Keratinocyte Overexpression from PMID: 30772301 | SOX2 |
| 6768 | DNAJC21 | SOX2 regulated genes in Keratinocyte Overexpression from PMID: 30772301 | SOX2 |
| 6769 | CARM1 | SOX2 regulated genes in Keratinocyte Overexpression from PMID: 30772301 | SOX2 |

| | | | |
|---|---|---|---|
| 6770 | NUSAP1 | SOX2 regulated genes in Keratinocyte Overexpression from PMID: 30772301 | SOX2 |
| 6771 | PSMD12 | SOX2 regulated genes in Keratinocyte Overexpression from PMID: 30772301 | SOX2 |
| 6772 | NFKBIB | SOX2 regulated genes in Keratinocyte Overexpression from PMID: 30772301 | SOX2 |
| 6773 | PPP4R4 | SOX2 regulated genes in Keratinocyte Overexpression from PMID: 30772301 | SOX2 |
| 6774 | ESRP2 | SOX2 regulated genes in Keratinocyte Overexpression from PMID: 30772301 | SOX2 |
| 6775 | PEX7 | SOX2 regulated genes in Keratinocyte Overexpression from PMID: 30772301 | SOX2 |
| 6776 | IYD | SOX2 regulated genes in Keratinocyte Overexpression from PMID: 30772301 | SOX2 |
| 6777 | BCDIN3D | SOX2 regulated genes in Keratinocyte Overexpression from PMID: 30772301 | SOX2 |
| 6778 | NCKAP1 | SOX2 regulated genes in Keratinocyte Overexpression from PMID: 30772301 | SOX2 |
| 6779 | ILVBL | SOX2 regulated genes in Keratinocyte Overexpression from PMID: 30772301 | SOX2 |
| 6780 | NEK6 | SOX2 regulated genes in Keratinocyte Overexpression from PMID: 30772301 | SOX2 |
| 6781 | PRKCH | SOX2 regulated genes in Keratinocyte Overexpression from PMID: 30772301 | SOX2 |
| 6782 | POLD2 | SOX2 regulated genes in Keratinocyte Overexpression from PMID: 30772301 | SOX2 |
| 6783 | ZNF740 | SOX2 regulated genes in Keratinocyte Overexpression from PMID: 30772301 | SOX2 |
| 6784 | NIPBL | SOX2 regulated genes in Keratinocyte Overexpression from PMID: 30772301 | SOX2 |
| 6785 | CPA4 | SOX2 regulated genes in Keratinocyte Overexpression from PMID: 30772301 | SOX2 |
| 6786 | SNAI2 | SOX2 regulated genes in Keratinocyte Overexpression from PMID: 30772301 | SOX2 |
| 6787 | SMYD4 | SOX2 regulated genes in Keratinocyte Overexpression from PMID: 30772301 | SOX2 |
| 6788 | CDPF1 | SOX2 regulated genes in Keratinocyte Overexpression from PMID: 30772301 | SOX2 |
| 6789 | DNAJB6 | SOX2 regulated genes in Keratinocyte Overexpression from PMID: 30772301 | SOX2 |
| 6790 | DCAF10 | SOX2 regulated genes in Keratinocyte Overexpression from PMID: 30772301 | SOX2 |
| 6791 | CLSPN | SOX2 regulated genes in Keratinocyte Overexpression from PMID: 30772301 | SOX2 |
| 6792 | CFDP1 | SOX2 regulated genes in Keratinocyte Overexpression from PMID: 30772301 | SOX2 |
| 6793 | EYA1 | SOX2 regulated genes in Keratinocyte Overexpression from PMID: 30772301 | SOX2 |
| 6794 | UBN1 | SOX2 regulated genes in Keratinocyte Overexpression from PMID: 30772301 | SOX2 |
| 6795 | ATXN7L3B | SOX2 regulated genes in Keratinocyte Overexpression from PMID: 30772301 | SOX2 |
| 6796 | GALNT6 | SOX2 regulated genes in Keratinocyte Overexpression from PMID: 30772301 | SOX2 |
| 6797 | TTC23 | SOX2 regulated genes in Keratinocyte Overexpression from PMID: 30772301 | SOX2 |
| 6798 | TCP11L2 | SOX2 regulated genes in Keratinocyte Overexpression from PMID: 30772301 | SOX2 |
| 6799 | CHCHD4 | SOX2 regulated genes in Keratinocyte Overexpression from PMID: 30772301 | SOX2 |
| 6800 | CEP89 | SOX2 regulated genes in Keratinocyte Overexpression from PMID: 30772301 | SOX2 |
| 6801 | FBXW8 | SOX2 regulated genes in Keratinocyte Overexpression from PMID: 30772301 | SOX2 |
| 6802 | DMRTA2 | SOX2 regulated genes in Keratinocyte Overexpression from PMID: 30772301 | SOX2 |
| 6803 | KAT7 | SOX2 regulated genes in Keratinocyte Overexpression from PMID: 30772301 | SOX2 |
| 6804 | NCAPH2 | SOX2 regulated genes in Keratinocyte Overexpression from PMID: 30772301 | SOX2 |
| 6805 | PPP2R1A | SOX2 regulated genes in Keratinocyte Overexpression from PMID: 30772301 | SOX2 |
| 6806 | SAFB2 | SOX2 regulated genes in Keratinocyte Overexpression from PMID: 30772301 | SOX2 |
| 6807 | IGF2R | SOX2 regulated genes in Keratinocyte Overexpression from PMID: 30772301 | SOX2 |
| 6808 | AOC2 | SOX2 regulated genes in Keratinocyte Overexpression from PMID: 30772301 | SOX2 |
| 6809 | EXOSC9 | SOX2 regulated genes in Keratinocyte Overexpression from PMID: 30772301 | SOX2 |
| 6810 | IGSF11 | SOX2 regulated genes in Keratinocyte Overexpression from PMID: 30772301 | SOX2 |
| 6811 | ALDH4A1 | SOX2 regulated genes in Keratinocyte Overexpression from PMID: 30772301 | SOX2 |
| 6812 | SEMA3F | SOX2 regulated genes in Keratinocyte Overexpression from PMID: 30772301 | SOX2 |
| 6813 | ZBTB1 | SOX2 regulated genes in Keratinocyte Overexpression from PMID: 30772301 | SOX2 |
| 6814 | MRPL54 | SOX2 regulated genes in Keratinocyte Overexpression from PMID: 30772301 | SOX2 |
| 6815 | RPL22 | SOX2 regulated genes in Keratinocyte Overexpression from PMID: 30772301 | SOX2 |
| 6816 | KRT80 | SOX2 regulated genes in Keratinocyte Overexpression from PMID: 30772301 | SOX2 |
| 6817 | SOX5 | SOX2 regulated genes in Keratinocyte Overexpression from PMID: 30772301 | SOX2 |
| 6818 | GTF3C4 | SOX2 regulated genes in Keratinocyte Overexpression from PMID: 30772301 | SOX2 |
| 6819 | TRPM4 | SOX2 regulated genes in Keratinocyte Overexpression from PMID: 30772301 | SOX2 |
| 6820 | PAXIP1 | SOX2 regulated genes in Keratinocyte Overexpression from PMID: 30772301 | SOX2 |
| 6821 | AARSD1 | SOX2 regulated genes in Keratinocyte Overexpression from PMID: 30772301 | SOX2 |
| 6822 | ZRANB3 | SOX2 regulated genes in Keratinocyte Overexpression from PMID: 30772301 | SOX2 |
| 6823 | GPR173 | SOX2 regulated genes in Keratinocyte Overexpression from PMID: 30772301 | SOX2 |
| 6824 | HSPA1L | SOX2 regulated genes in Keratinocyte Overexpression from PMID: 30772301 | SOX2 |
| 6825 | TMEM170A | SOX2 regulated genes in Keratinocyte Overexpression from PMID: 30772301 | SOX2 |
| 6826 | NOC3L | SOX2 regulated genes in Keratinocyte Overexpression from PMID: 30772301 | SOX2 |
| 6827 | MTRF1L | SOX2 regulated genes in Keratinocyte Overexpression from PMID: 30772301 | SOX2 |
| 6828 | ATF1 | SOX2 regulated genes in Keratinocyte Overexpression from PMID: 30772301 | SOX2 |
| 6829 | MED18 | SOX2 regulated genes in Keratinocyte Overexpression from PMID: 30772301 | SOX2 |
| 6830 | GPC3 | SOX2 regulated genes in Keratinocyte Overexpression from PMID: 30772301 | SOX2 |

| | | | |
|---|---|---|---|
| 6831 | SYBU | SOX2 regulated genes in Keratinocyte Overexpression from PMID: 30772301 | SOX2 |
| 6832 | BBS1 | SOX2 regulated genes in Keratinocyte Overexpression from PMID: 30772301 | SOX2 |
| 6833 | SUPT4H1 | SOX2 regulated genes in Keratinocyte Overexpression from PMID: 30772301 | SOX2 |
| 6834 | TSSK6 | SOX2 regulated genes in Keratinocyte Overexpression from PMID: 30772301 | SOX2 |
| 6835 | UTP14C | SOX2 regulated genes in Keratinocyte Overexpression from PMID: 30772301 | SOX2 |
| 6836 | PLCH2 | SOX2 regulated genes in Keratinocyte Overexpression from PMID: 30772301 | SOX2 |
| 6837 | UBR5 | SOX2 regulated genes in Keratinocyte Overexpression from PMID: 30772301 | SOX2 |
| 6838 | EDC4 | SOX2 regulated genes in Keratinocyte Overexpression from PMID: 30772301 | SOX2 |
| 6839 | LSM8 | SOX2 regulated genes in Keratinocyte Overexpression from PMID: 30772301 | SOX2 |
| 6840 | LCN12 | SOX2 regulated genes in Keratinocyte Overexpression from PMID: 30772301 | SOX2 |
| 6841 | MBNL3 | SOX2 regulated genes in Keratinocyte Overexpression from PMID: 30772301 | SOX2 |
| 6842 | NAE1 | SOX2 regulated genes in Keratinocyte Overexpression from PMID: 30772301 | SOX2 |
| 6843 | SOX10 | SOX2 regulated genes in Keratinocyte Overexpression from PMID: 30772301 | SOX2 |
| 6844 | STRN4 | SOX2 regulated genes in Keratinocyte Overexpression from PMID: 30772301 | SOX2 |
| 6845 | SLC38A3 | SOX2 regulated genes in Keratinocyte Overexpression from PMID: 30772301 | SOX2 |
| 6846 | RIOX2 | SOX2 regulated genes in Keratinocyte Overexpression from PMID: 30772301 | SOX2 |
| 6847 | HELQ | SOX2 regulated genes in Keratinocyte Overexpression from PMID: 30772301 | SOX2 |
| 6848 | CD200R1 | SOX2 regulated genes in Keratinocyte Overexpression from PMID: 30772301 | SOX2 |
| 6849 | EXOSC8 | SOX2 regulated genes in Keratinocyte Overexpression from PMID: 30772301 | SOX2 |
| 6850 | FANCE | SOX2 regulated genes in Keratinocyte Overexpression from PMID: 30772301 | SOX2 |
| 6851 | NEK5 | SOX2 regulated genes in Keratinocyte Overexpression from PMID: 30772301 | SOX2 |
| 6852 | YWHAQ | SOX2 regulated genes in Keratinocyte Overexpression from PMID: 30772301 | SOX2 |
| 6853 | TENM4 | SOX2 regulated genes in Keratinocyte Overexpression from PMID: 30772301 | SOX2 |
| 6854 | AMMECR1 | SOX2 regulated genes in Keratinocyte Overexpression from PMID: 30772301 | SOX2 |
| 6855 | STRBP | SOX2 regulated genes in Keratinocyte Overexpression from PMID: 30772301 | SOX2 |
| 6856 | METAP1 | SOX2 regulated genes in Keratinocyte Overexpression from PMID: 30772301 | SOX2 |
| 6857 | DNAJC2 | SOX2 regulated genes in Keratinocyte Overexpression from PMID: 30772301 | SOX2 |
| 6858 | HAUS3 | SOX2 regulated genes in Keratinocyte Overexpression from PMID: 30772301 | SOX2 |
| 6859 | PAQR5 | SOX2 regulated genes in Keratinocyte Overexpression from PMID: 30772301 | SOX2 |
| 6860 | CECR2 | SOX2 regulated genes in Keratinocyte Overexpression from PMID: 30772301 | SOX2 |
| 6861 | SPDL1 | SOX2 regulated genes in Keratinocyte Overexpression from PMID: 30772301 | SOX2 |
| 6862 | ABLIM2 | SOX2 regulated genes in Keratinocyte Overexpression from PMID: 30772301 | SOX2 |
| 6863 | PIN1 | SOX2 regulated genes in Keratinocyte Overexpression from PMID: 30772301 | SOX2 |
| 6864 | PHF6 | SOX2 regulated genes in Keratinocyte Overexpression from PMID: 30772301 | SOX2 |
| 6865 | CTDSPL | SOX2 regulated genes in Keratinocyte Overexpression from PMID: 30772301 | SOX2 |
| 6866 | SERHL2 | SOX2 regulated genes in Keratinocyte Overexpression from PMID: 30772301 | SOX2 |
| 6867 | NOL11 | SOX2 regulated genes in Keratinocyte Overexpression from PMID: 30772301 | SOX2 |
| 6868 | PIGU | SOX2 regulated genes in Keratinocyte Overexpression from PMID: 30772301 | SOX2 |
| 6869 | STX1A | SOX2 regulated genes in Keratinocyte Overexpression from PMID: 30772301 | SOX2 |
| 6870 | SDK2 | SOX2 regulated genes in Keratinocyte Overexpression from PMID: 30772301 | SOX2 |
| 6871 | SRD5A2 | SOX2 regulated genes in Keratinocyte Overexpression from PMID: 30772301 | SOX2 |
| 6872 | BOLA2B | SOX2 regulated genes in Keratinocyte Overexpression from PMID: 30772301 | SOX2 |
| 6873 | COPS3 | SOX2 regulated genes in Keratinocyte Overexpression from PMID: 30772301 | SOX2 |
| 6874 | C9orf152 | SOX2 regulated genes in Keratinocyte Overexpression from PMID: 30772301 | SOX2 |
| 6875 | UNC93A | SOX2 regulated genes in Keratinocyte Overexpression from PMID: 30772301 | SOX2 |
| 6876 | MROH7 | SOX2 regulated genes in Keratinocyte Overexpression from PMID: 30772301 | SOX2 |
| 6877 | SCX | SOX2 regulated genes in Keratinocyte Overexpression from PMID: 30772301 | SOX2 |
| 6878 | ZC3H4 | SOX2 regulated genes in Keratinocyte Overexpression from PMID: 30772301 | SOX2 |
| 6879 | RPL37A | SOX2 regulated genes in Keratinocyte Overexpression from PMID: 30772301 | SOX2 |
| 6880 | ADCK2 | SOX2 regulated genes in Keratinocyte Overexpression from PMID: 30772301 | SOX2 |
| 6881 | SIPA1L3 | SOX2 regulated genes in Keratinocyte Overexpression from PMID: 30772301 | SOX2 |
| 6882 | TMC7 | SOX2 regulated genes in Keratinocyte Overexpression from PMID: 30772301 | SOX2 |
| 6883 | TMEM171 | SOX2 regulated genes in Keratinocyte Overexpression from PMID: 30772301 | SOX2 |
| 6884 | OCM | SOX2 regulated genes in Keratinocyte Overexpression from PMID: 30772301 | SOX2 |
| 6885 | DYRK2 | SOX2 regulated genes in Keratinocyte Overexpression from PMID: 30772301 | SOX2 |
| 6886 | PLAG1 | SOX2 regulated genes in Keratinocyte Overexpression from PMID: 30772301 | SOX2 |
| 6887 | DYNLL1 | SOX2 regulated genes in Keratinocyte Overexpression from PMID: 30772301 | SOX2 |
| 6888 | PTGES2 | SOX2 regulated genes in Keratinocyte Overexpression from PMID: 30772301 | SOX2 |
| 6889 | PLA2G4C | SOX2 regulated genes in Keratinocyte Overexpression from PMID: 30772301 | SOX2 |
| 6890 | PXYLP1 | SOX2 regulated genes in Keratinocyte Overexpression from PMID: 30772301 | SOX2 |
| 6891 | RIMBP3C | SOX2 regulated genes in Keratinocyte Overexpression from PMID: 30772301 | SOX2 |

| | | | |
|---|---|---|---|
| 6892 | NDFIP2 | SOX2 regulated genes in Keratinocyte Overexpression from PMID: 30772301 | SOX2 |
| 6893 | RSPO3 | SOX2 regulated genes in Keratinocyte Overexpression from PMID: 30772301 | SOX2 |
| 6894 | TMEM185A | SOX2 regulated genes in Keratinocyte Overexpression from PMID: 30772301 | SOX2 |
| 6895 | LSM3 | SOX2 regulated genes in Keratinocyte Overexpression from PMID: 30772301 | SOX2 |
| 6896 | ATP6V1H | SOX2 regulated genes in Keratinocyte Overexpression from PMID: 30772301 | SOX2 |
| 6897 | EPCAM | SOX2 regulated genes in Keratinocyte Overexpression from PMID: 30772301 | SOX2 |
| 6898 | EEF1AKMT1 | SOX2 regulated genes in Keratinocyte Overexpression from PMID: 30772301 | SOX2 |
| 6899 | LATS1 | SOX2 regulated genes in Keratinocyte Overexpression from PMID: 30772301 | SOX2 |
| 6900 | PPP4R3A | SOX2 regulated genes in Keratinocyte Overexpression from PMID: 30772301 | SOX2 |
| 6901 | APBA3 | SOX2 regulated genes in Keratinocyte Overexpression from PMID: 30772301 | SOX2 |
| 6902 | SERTAD3 | SOX2 regulated genes in Keratinocyte Overexpression from PMID: 30772301 | SOX2 |
| 6903 | PPP1CB | SOX2 regulated genes in Keratinocyte Overexpression from PMID: 30772301 | SOX2 |
| 6904 | MCM4 | SOX2 regulated genes in Keratinocyte Overexpression from PMID: 30772301 | SOX2 |
| 6905 | ODF2L | SOX2 regulated genes in Keratinocyte Overexpression from PMID: 30772301 | SOX2 |
| 6906 | CASP8AP2 | SOX2 regulated genes in Keratinocyte Overexpression from PMID: 30772301 | SOX2 |
| 6907 | SKIL | SOX2 regulated genes in Keratinocyte Overexpression from PMID: 30772301 | SOX2 |
| 6908 | FUBP3 | SOX2 regulated genes in Keratinocyte Overexpression from PMID: 30772301 | SOX2 |
| 6909 | MTBP | SOX2 regulated genes in Keratinocyte Overexpression from PMID: 30772301 | SOX2 |
| 6910 | PSCA | SOX2 regulated genes in Keratinocyte Overexpression from PMID: 30772301 | SOX2 |
| 6911 | VPS33B | SOX2 regulated genes in Keratinocyte Overexpression from PMID: 30772301 | SOX2 |
| 6912 | RAB3B | SOX2 regulated genes in Keratinocyte Overexpression from PMID: 30772301 | SOX2 |
| 6913 | CLDN34 | SOX2 regulated genes in Keratinocyte Overexpression from PMID: 30772301 | SOX2 |
| 6914 | LRRC45 | SOX2 regulated genes in Keratinocyte Overexpression from PMID: 30772301 | SOX2 |
| 6915 | HMMR | SOX2 regulated genes in Keratinocyte Overexpression from PMID: 30772301 | SOX2 |
| 6916 | RAC3 | SOX2 regulated genes in Keratinocyte Overexpression from PMID: 30772301 | SOX2 |
| 6917 | NECAB1 | SOX2 regulated genes in Keratinocyte Overexpression from PMID: 30772301 | SOX2 |
| 6918 | ZNF248 | SOX2 regulated genes in Keratinocyte Overexpression from PMID: 30772301 | SOX2 |
| 6919 | PSMA1 | SOX2 regulated genes in Keratinocyte Overexpression from PMID: 30772301 | SOX2 |
| 6920 | CKAP4 | SOX2 regulated genes in Keratinocyte Overexpression from PMID: 30772301 | SOX2 |
| 6921 | LGALSL | SOX2 regulated genes in Keratinocyte Overexpression from PMID: 30772301 | SOX2 |
| 6922 | ARMC5 | SOX2 regulated genes in Keratinocyte Overexpression from PMID: 30772301 | SOX2 |
| 6923 | PSMD11 | SOX2 regulated genes in Keratinocyte Overexpression from PMID: 30772301 | SOX2 |
| 6924 | FBXL6 | SOX2 regulated genes in Keratinocyte Overexpression from PMID: 30772301 | SOX2 |
| 6925 | CYC1 | SOX2 regulated genes in Keratinocyte Overexpression from PMID: 30772301 | SOX2 |
| 6926 | RPL31 | SOX2 regulated genes in Keratinocyte Overexpression from PMID: 30772301 | SOX2 |
| 6927 | SMDT1 | SOX2 regulated genes in Keratinocyte Overexpression from PMID: 30772301 | SOX2 |
| 6928 | TALDO1 | SOX2 regulated genes in Keratinocyte Overexpression from PMID: 30772301 | SOX2 |
| 6929 | ZFP37 | SOX2 regulated genes in Keratinocyte Overexpression from PMID: 30772301 | SOX2 |
| 6930 | SNF8 | SOX2 regulated genes in Keratinocyte Overexpression from PMID: 30772301 | SOX2 |
| 6931 | CFAP54 | SOX2 regulated genes in Keratinocyte Overexpression from PMID: 30772301 | SOX2 |
| 6932 | ACAT1 | SOX2 regulated genes in Keratinocyte Overexpression from PMID: 30772301 | SOX2 |
| 6933 | STARD4 | SOX2 regulated genes in Keratinocyte Overexpression from PMID: 30772301 | SOX2 |
| 6934 | SAAL1 | SOX2 regulated genes in Keratinocyte Overexpression from PMID: 30772301 | SOX2 |
| 6935 | PRPF18 | SOX2 regulated genes in Keratinocyte Overexpression from PMID: 30772301 | SOX2 |
| 6936 | BMPR1B | SOX2 regulated genes in Keratinocyte Overexpression from PMID: 30772301 | SOX2 |
| 6937 | ANKRD6 | SOX2 regulated genes in Keratinocyte Overexpression from PMID: 30772301 | SOX2 |
| 6938 | LHX2 | SOX2 regulated genes in Keratinocyte Overexpression from PMID: 30772301 | SOX2 |
| 6939 | NELFA | SOX2 regulated genes in Keratinocyte Overexpression from PMID: 30772301 | SOX2 |
| 6940 | TMEM129 | SOX2 regulated genes in Keratinocyte Overexpression from PMID: 30772301 | SOX2 |
| 6941 | C4orf46 | SOX2 regulated genes in Keratinocyte Overexpression from PMID: 30772301 | SOX2 |
| 6942 | MCC | SOX2 regulated genes in Keratinocyte Overexpression from PMID: 30772301 | SOX2 |
| 6943 | TBCC | SOX2 regulated genes in Keratinocyte Overexpression from PMID: 30772301 | SOX2 |
| 6944 | RAB37 | SOX2 regulated genes in Keratinocyte Overexpression from PMID: 30772301 | SOX2 |
| 6945 | EGR4 | SOX2 regulated genes in Keratinocyte Overexpression from PMID: 30772301 | SOX2 |
| 6946 | CHSY3 | SOX2 regulated genes in Keratinocyte Overexpression from PMID: 30772301 | SOX2 |
| 6947 | ZNF2 | SOX2 regulated genes in Keratinocyte Overexpression from PMID: 30772301 | SOX2 |
| 6948 | FZD6 | SOX2 regulated genes in Keratinocyte Overexpression from PMID: 30772301 | SOX2 |
| 6949 | NELFE | SOX2 regulated genes in Keratinocyte Overexpression from PMID: 30772301 | SOX2 |
| 6950 | TRMT10A | SOX2 regulated genes in Keratinocyte Overexpression from PMID: 30772301 | SOX2 |
| 6951 | DSCC1 | SOX2 regulated genes in Keratinocyte Overexpression from PMID: 30772301 | SOX2 |
| 6952 | DNAH17 | SOX2 regulated genes in Keratinocyte Overexpression from PMID: 30772301 | SOX2 |

| | | | |
|---|---|---|---|
| 6953 | CRB3 | SOX2 regulated genes in Keratinocyte Overexpression from PMID: 30772301 | SOX2 |
| 6954 | CREBBP | SOX2 regulated genes in Keratinocyte Overexpression from PMID: 30772301 | SOX2 |
| 6955 | PMPCA | SOX2 regulated genes in Keratinocyte Overexpression from PMID: 30772301 | SOX2 |
| 6956 | C11orf94 | SOX2 regulated genes in Keratinocyte Overexpression from PMID: 30772301 | SOX2 |
| 6957 | BARD1 | SOX2 regulated genes in Keratinocyte Overexpression from PMID: 30772301 | SOX2 |
| 6958 | DNAJA2 | SOX2 regulated genes in Keratinocyte Overexpression from PMID: 30772301 | SOX2 |
| 6959 | TPBG | SOX2 regulated genes in Keratinocyte Overexpression from PMID: 30772301 | SOX2 |
| 6960 | RANBP17 | SOX2 regulated genes in Keratinocyte Overexpression from PMID: 30772301 | SOX2 |
| 6961 | KIAA1958 | SOX2 regulated genes in Keratinocyte Overexpression from PMID: 30772301 | SOX2 |
| 6962 | MSH3 | SOX2 regulated genes in Keratinocyte Overexpression from PMID: 30772301 | SOX2 |
| 6963 | PDCD5 | SOX2 regulated genes in Keratinocyte Overexpression from PMID: 30772301 | SOX2 |
| 6964 | ATN1 | SOX2 regulated genes in Keratinocyte Overexpression from PMID: 30772301 | SOX2 |
| 6965 | PIP4K2C | SOX2 regulated genes in Keratinocyte Overexpression from PMID: 30772301 | SOX2 |
| 6966 | DEF6 | SOX2 regulated genes in Keratinocyte Overexpression from PMID: 30772301 | SOX2 |
| 6967 | TMTC3 | SOX2 regulated genes in Keratinocyte Overexpression from PMID: 30772301 | SOX2 |
| 6968 | PHLDB2 | SOX2 regulated genes in Keratinocyte Overexpression from PMID: 30772301 | SOX2 |
| 6969 | LGALS12 | SOX2 regulated genes in Keratinocyte Overexpression from PMID: 30772301 | SOX2 |
| 6970 | ZC3HC1 | SOX2 regulated genes in Keratinocyte Overexpression from PMID: 30772301 | SOX2 |
| 6971 | DCAF13 | SOX2 regulated genes in Keratinocyte Overexpression from PMID: 30772301 | SOX2 |
| 6972 | CHIC1 | SOX2 regulated genes in Keratinocyte Overexpression from PMID: 30772301 | SOX2 |
| 6973 | TP53BP1 | SOX2 regulated genes in Keratinocyte Overexpression from PMID: 30772301 | SOX2 |
| 6974 | SBK1 | SOX2 regulated genes in Keratinocyte Overexpression from PMID: 30772301 | SOX2 |
| 6975 | GPLD1 | SOX2 regulated genes in Keratinocyte Overexpression from PMID: 30772301 | SOX2 |
| 6976 | AHI1 | SOX2 regulated genes in Keratinocyte Overexpression from PMID: 30772301 | SOX2 |
| 6977 | NHS | SOX2 regulated genes in Keratinocyte Overexpression from PMID: 30772301 | SOX2 |
| 6978 | IFT80 | SOX2 regulated genes in Keratinocyte Overexpression from PMID: 30772301 | SOX2 |
| 6979 | PAM16 | SOX2 regulated genes in Keratinocyte Overexpression from PMID: 30772301 | SOX2 |
| 6980 | ZCCHC10 | SOX2 regulated genes in Keratinocyte Overexpression from PMID: 30772301 | SOX2 |
| 6981 | SLF1 | SOX2 regulated genes in Keratinocyte Overexpression from PMID: 30772301 | SOX2 |
| 6982 | MCMBP | SOX2 regulated genes in Keratinocyte Overexpression from PMID: 30772301 | SOX2 |
| 6983 | OSGIN2 | SOX2 regulated genes in Keratinocyte Overexpression from PMID: 30772301 | SOX2 |
| 6984 | XYLB | SOX2 regulated genes in Keratinocyte Overexpression from PMID: 30772301 | SOX2 |
| 6985 | BTBD10 | SOX2 regulated genes in Keratinocyte Overexpression from PMID: 30772301 | SOX2 |
| 6986 | CLIC4 | SOX2 regulated genes in Keratinocyte Overexpression from PMID: 30772301 | SOX2 |
| 6987 | GEMIN6 | SOX2 regulated genes in Keratinocyte Overexpression from PMID: 30772301 | SOX2 |
| 6988 | MRPL46 | SOX2 regulated genes in Keratinocyte Overexpression from PMID: 30772301 | SOX2 |
| 6989 | ATP7A | SOX2 regulated genes in Keratinocyte Overexpression from PMID: 30772301 | SOX2 |
| 6990 | PPM1B | SOX2 regulated genes in Keratinocyte Overexpression from PMID: 30772301 | SOX2 |
| 6991 | GADD45A | SOX2 regulated genes in Keratinocyte Overexpression from PMID: 30772301 | SOX2 |
| 6992 | WTAP | SOX2 regulated genes in Keratinocyte Overexpression from PMID: 30772301 | SOX2 |
| 6993 | GRIK4 | SOX2 regulated genes in Keratinocyte Overexpression from PMID: 30772301 | SOX2 |
| 6994 | MED26 | SOX2 regulated genes in Keratinocyte Overexpression from PMID: 30772301 | SOX2 |
| 6995 | PREX2 | SOX2 regulated genes in Keratinocyte Overexpression from PMID: 30772301 | SOX2 |
| 6996 | GGACT | SOX2 regulated genes in Keratinocyte Overexpression from PMID: 30772301 | SOX2 |
| 6997 | ARNTL | SOX2 regulated genes in Keratinocyte Overexpression from PMID: 30772301 | SOX2 |
| 6998 | SLC30A4 | SOX2 regulated genes in Keratinocyte Overexpression from PMID: 30772301 | SOX2 |
| 6999 | SUGT1 | SOX2 regulated genes in Keratinocyte Overexpression from PMID: 30772301 | SOX2 |
| 7000 | PPWD1 | SOX2 regulated genes in Keratinocyte Overexpression from PMID: 30772301 | SOX2 |
| 7001 | RASSF7 | SOX2 regulated genes in Keratinocyte Overexpression from PMID: 30772301 | SOX2 |
| 7002 | TUBB4B | SOX2 regulated genes in Keratinocyte Overexpression from PMID: 30772301 | SOX2 |
| 7003 | MND1 | SOX2 regulated genes in Keratinocyte Overexpression from PMID: 30772301 | SOX2 |
| 7004 | ZFYVE28 | SOX2 regulated genes in Keratinocyte Overexpression from PMID: 30772301 | SOX2 |
| 7005 | TTC26 | SOX2 regulated genes in Keratinocyte Overexpression from PMID: 30772301 | SOX2 |
| 7006 | GARNL3 | SOX2 regulated genes in Keratinocyte Overexpression from PMID: 30772301 | SOX2 |
| 7007 | PRPF6 | SOX2 regulated genes in Keratinocyte Overexpression from PMID: 30772301 | SOX2 |
| 7008 | KDM4A | SOX2 regulated genes in Keratinocyte Overexpression from PMID: 30772301 | SOX2 |
| 7009 | SPATA5 | SOX2 regulated genes in Keratinocyte Overexpression from PMID: 30772301 | SOX2 |
| 7010 | SYT1 | SOX2 regulated genes in Keratinocyte Overexpression from PMID: 30772301 | SOX2 |
| 7011 | DUS4L | SOX2 regulated genes in Keratinocyte Overexpression from PMID: 30772301 | SOX2 |
| 7012 | MRPS9 | SOX2 regulated genes in Keratinocyte Overexpression from PMID: 30772301 | SOX2 |
| 7013 | ERMARD | SOX2 regulated genes in Keratinocyte Overexpression from PMID: 30772301 | SOX2 |

| | | | |
|---|---|---|---|
| 7014 | NSMAF | SOX2 regulated genes in Keratinocyte Overexpression from PMID: 30772301 | SOX2 |
| 7015 | CACNG4 | SOX2 regulated genes in Keratinocyte Overexpression from PMID: 30772301 | SOX2 |
| 7016 | CEP126 | SOX2 regulated genes in Keratinocyte Overexpression from PMID: 30772301 | SOX2 |
| 7017 | ZFHX2 | SOX2 regulated genes in Keratinocyte Overexpression from PMID: 30772301 | SOX2 |
| 7018 | TMEM132A | SOX2 regulated genes in Keratinocyte Overexpression from PMID: 30772301 | SOX2 |
| 7019 | AOC1 | SOX2 regulated genes in Keratinocyte Overexpression from PMID: 30772301 | SOX2 |
| 7020 | NDUFAF8 | SOX2 regulated genes in Keratinocyte Overexpression from PMID: 30772301 | SOX2 |
| 7021 | HYAL2 | SOX2 regulated genes in Keratinocyte Overexpression from PMID: 30772301 | SOX2 |
| 7022 | MSH5 | SOX2 regulated genes in Keratinocyte Overexpression from PMID: 30772301 | SOX2 |
| 7023 | FFAR3 | SOX2 regulated genes in Keratinocyte Overexpression from PMID: 30772301 | SOX2 |
| 7024 | C16orf74 | SOX2 regulated genes in Keratinocyte Overexpression from PMID: 30772301 | SOX2 |
| 7025 | METTL26 | SOX2 regulated genes in Keratinocyte Overexpression from PMID: 30772301 | SOX2 |
| 7026 | SC5D | SOX2 regulated genes in Keratinocyte Overexpression from PMID: 30772301 | SOX2 |
| 7027 | MIP | SOX2 regulated genes in Keratinocyte Overexpression from PMID: 30772301 | SOX2 |
| 7028 | SEC16A | SOX2 regulated genes in Keratinocyte Overexpression from PMID: 30772301 | SOX2 |
| 7029 | ZBTB8B | SOX2 regulated genes in Keratinocyte Overexpression from PMID: 30772301 | SOX2 |
| 7030 | DNAJC17 | SOX2 regulated genes in Keratinocyte Overexpression from PMID: 30772301 | SOX2 |
| 7031 | MACC1 | SOX2 regulated genes in Keratinocyte Overexpression from PMID: 30772301 | SOX2 |
| 7032 | FBXO34 | SOX2 regulated genes in Keratinocyte Overexpression from PMID: 30772301 | SOX2 |
| 7033 | ARID4A | SOX2 regulated genes in Keratinocyte Overexpression from PMID: 30772301 | SOX2 |
| 7034 | FBXO42 | SOX2 regulated genes in Keratinocyte Overexpression from PMID: 30772301 | SOX2 |
| 7035 | ESRP1 | SOX2 regulated genes in Keratinocyte Overexpression from PMID: 30772301 | SOX2 |
| 7036 | FMNL2 | SOX2 regulated genes in Keratinocyte Overexpression from PMID: 30772301 | SOX2 |
| 7037 | RPL30 | SOX2 regulated genes in Keratinocyte Overexpression from PMID: 30772301 | SOX2 |
| 7038 | ZNF213 | SOX2 regulated genes in Keratinocyte Overexpression from PMID: 30772301 | SOX2 |
| 7039 | SUPV3L1 | SOX2 regulated genes in Keratinocyte Overexpression from PMID: 30772301 | SOX2 |
| 7040 | ZNF334 | SOX2 regulated genes in Keratinocyte Overexpression from PMID: 30772301 | SOX2 |
| 7041 | LPAR2 | SOX2 regulated genes in Keratinocyte Overexpression from PMID: 30772301 | SOX2 |
| 7042 | ZCCHC7 | SOX2 regulated genes in Keratinocyte Overexpression from PMID: 30772301 | SOX2 |
| 7043 | SYP | SOX2 regulated genes in Keratinocyte Overexpression from PMID: 30772301 | SOX2 |
| 7044 | CTNNA1 | SOX2 regulated genes in Keratinocyte Overexpression from PMID: 30772301 | SOX2 |
| 7045 | ANKRD27 | SOX2 regulated genes in Keratinocyte Overexpression from PMID: 30772301 | SOX2 |
| 7046 | LCMT1 | SOX2 regulated genes in Keratinocyte Overexpression from PMID: 30772301 | SOX2 |
| 7047 | SNTB1 | SOX2 regulated genes in Keratinocyte Overexpression from PMID: 30772301 | SOX2 |
| 7048 | KLHL32 | SOX2 regulated genes in Keratinocyte Overexpression from PMID: 30772301 | SOX2 |
| 7049 | STX5 | SOX2 regulated genes in Keratinocyte Overexpression from PMID: 30772301 | SOX2 |
| 7050 | IGHMBP2 | SOX2 regulated genes in Keratinocyte Overexpression from PMID: 30772301 | SOX2 |
| 7051 | PLD6 | SOX2 regulated genes in Keratinocyte Overexpression from PMID: 30772301 | SOX2 |
| 7052 | GNA11 | SOX2 regulated genes in Keratinocyte Overexpression from PMID: 30772301 | SOX2 |
| 7053 | TTL | SOX2 regulated genes in Keratinocyte Overexpression from PMID: 30772301 | SOX2 |
| 7054 | SLC25A46 | SOX2 regulated genes in Keratinocyte Overexpression from PMID: 30772301 | SOX2 |
| 7055 | GLI1 | SOX2 regulated genes in Keratinocyte Overexpression from PMID: 30772301 | SOX2 |
| 7056 | RBM10 | SOX2 regulated genes in Keratinocyte Overexpression from PMID: 30772301 | SOX2 |
| 7057 | THAP11 | SOX2 regulated genes in Keratinocyte Overexpression from PMID: 30772301 | SOX2 |
| 7058 | DCTN3 | SOX2 regulated genes in Keratinocyte Overexpression from PMID: 30772301 | SOX2 |
| 7059 | CALD1 | SOX2 regulated genes in Keratinocyte Overexpression from PMID: 30772301 | SOX2 |
| 7060 | FAM193B | SOX2 regulated genes in Keratinocyte Overexpression from PMID: 30772301 | SOX2 |
| 7061 | AIG1 | SOX2 regulated genes in Keratinocyte Overexpression from PMID: 30772301 | SOX2 |
| 7062 | SLC9A2 | SOX2 regulated genes in Keratinocyte Overexpression from PMID: 30772301 | SOX2 |
| 7063 | TMEM268 | SOX2 regulated genes in Keratinocyte Overexpression from PMID: 30772301 | SOX2 |
| 7064 | HAGH | SOX2 regulated genes in Keratinocyte Overexpression from PMID: 30772301 | SOX2 |
| 7065 | NUP210 | SOX2 regulated genes in Keratinocyte Overexpression from PMID: 30772301 | SOX2 |
| 7066 | LRP8 | SOX2 regulated genes in Keratinocyte Overexpression from PMID: 30772301 | SOX2 |
| 7067 | RBM42 | SOX2 regulated genes in Keratinocyte Overexpression from PMID: 30772301 | SOX2 |
| 7068 | PTGER4 | SOX2 regulated genes in Keratinocyte Overexpression from PMID: 30772301 | SOX2 |
| 7069 | MSH2 | SOX2 regulated genes in Keratinocyte Overexpression from PMID: 30772301 | SOX2 |
| 7070 | YIF1A | SOX2 regulated genes in Keratinocyte Overexpression from PMID: 30772301 | SOX2 |
| 7071 | TUFM | SOX2 regulated genes in Keratinocyte Overexpression from PMID: 30772301 | SOX2 |
| 7072 | MICAL1 | SOX2 regulated genes in Keratinocyte Overexpression from PMID: 30772301 | SOX2 |
| 7073 | TCF19 | SOX2 regulated genes in Keratinocyte Overexpression from PMID: 30772301 | SOX2 |
| 7074 | LZTS2 | SOX2 regulated genes in Keratinocyte Overexpression from PMID: 30772301 | SOX2 |

| | | | |
|---|---|---|---|
| 7075 | HSPB1 | SOX2 regulated genes in Keratinocyte Overexpression from PMID: 30772301 | SOX2 |
| 7076 | CPSF2 | SOX2 regulated genes in Keratinocyte Overexpression from PMID: 30772301 | SOX2 |
| 7077 | SHCBP1 | SOX2 regulated genes in Keratinocyte Overexpression from PMID: 30772301 | SOX2 |
| 7078 | DNLZ | SOX2 regulated genes in Keratinocyte Overexpression from PMID: 30772301 | SOX2 |
| 7079 | NSD2 | SOX2 regulated genes in Keratinocyte Overexpression from PMID: 30772301 | SOX2 |
| 7080 | ZFP1 | SOX2 regulated genes in Keratinocyte Overexpression from PMID: 30772301 | SOX2 |
| 7081 | GTPBP10 | SOX2 regulated genes in Keratinocyte Overexpression from PMID: 30772301 | SOX2 |
| 7082 | DEPDC1B | SOX2 regulated genes in Keratinocyte Overexpression from PMID: 30772301 | SOX2 |
| 7083 | SERPINB9 | SOX2 regulated genes in Keratinocyte Overexpression from PMID: 30772301 | SOX2 |
| 7084 | KIAA1549 | SOX2 regulated genes in Keratinocyte Overexpression from PMID: 30772301 | SOX2 |
| 7085 | MBLAC1 | SOX2 regulated genes in Keratinocyte Overexpression from PMID: 30772301 | SOX2 |
| 7086 | B3GALT6 | SOX2 regulated genes in Keratinocyte Overexpression from PMID: 30772301 | SOX2 |
| 7087 | IFRD1 | SOX2 regulated genes in Keratinocyte Overexpression from PMID: 30772301 | SOX2 |
| 7088 | KIT | SOX2 regulated genes in Keratinocyte Overexpression from PMID: 30772301 | SOX2 |
| 7089 | DIS3L | SOX2 regulated genes in Keratinocyte Overexpression from PMID: 30772301 | SOX2 |
| 7090 | S100PBP | SOX2 regulated genes in Keratinocyte Overexpression from PMID: 30772301 | SOX2 |
| 7091 | TDG | SOX2 regulated genes in Keratinocyte Overexpression from PMID: 30772301 | SOX2 |
| 7092 | POU2F1 | SOX2 regulated genes in Keratinocyte Overexpression from PMID: 30772301 | SOX2 |
| 7093 | ABLIM3 | SOX2 regulated genes in Keratinocyte Overexpression from PMID: 30772301 | SOX2 |
| 7094 | KCNK7 | SOX2 regulated genes in Keratinocyte Overexpression from PMID: 30772301 | SOX2 |
| 7095 | KCNC3 | SOX2 regulated genes in Keratinocyte Overexpression from PMID: 30772301 | SOX2 |
| 7096 | TXLNA | SOX2 regulated genes in Keratinocyte Overexpression from PMID: 30772301 | SOX2 |
| 7097 | DDX5 | SOX2 regulated genes in Keratinocyte Overexpression from PMID: 30772301 | SOX2 |
| 7098 | RITA1 | SOX2 regulated genes in Keratinocyte Overexpression from PMID: 30772301 | SOX2 |
| 7099 | RUNDC3A | SOX2 regulated genes in Keratinocyte Overexpression from PMID: 30772301 | SOX2 |
| 7100 | NLRX1 | SOX2 regulated genes in Keratinocyte Overexpression from PMID: 30772301 | SOX2 |
| 7101 | DDX59 | SOX2 regulated genes in Keratinocyte Overexpression from PMID: 30772301 | SOX2 |
| 7102 | ARHGAP29 | SOX2 regulated genes in Keratinocyte Overexpression from PMID: 30772301 | SOX2 |
| 7103 | FBXO25 | SOX2 regulated genes in Keratinocyte Overexpression from PMID: 30772301 | SOX2 |
| 7104 | SMPD4 | SOX2 regulated genes in Keratinocyte Overexpression from PMID: 30772301 | SOX2 |
| 7105 | ODF2 | SOX2 regulated genes in Keratinocyte Overexpression from PMID: 30772301 | SOX2 |
| 7106 | TXNL4B | SOX2 regulated genes in Keratinocyte Overexpression from PMID: 30772301 | SOX2 |
| 7107 | TIGD5 | SOX2 regulated genes in Keratinocyte Overexpression from PMID: 30772301 | SOX2 |
| 7108 | PLA2G4B | SOX2 regulated genes in Keratinocyte Overexpression from PMID: 30772301 | SOX2 |
| 7109 | HAUS8 | SOX2 regulated genes in Keratinocyte Overexpression from PMID: 30772301 | SOX2 |
| 7110 | C6orf163 | SOX2 regulated genes in Keratinocyte Overexpression from PMID: 30772301 | SOX2 |
| 7111 | FBXO38 | SOX2 regulated genes in Keratinocyte Overexpression from PMID: 30772301 | SOX2 |
| 7112 | UBE3B | SOX2 regulated genes in Keratinocyte Overexpression from PMID: 30772301 | SOX2 |
| 7113 | CACNA1H | SOX2 regulated genes in Keratinocyte Overexpression from PMID: 30772301 | SOX2 |
| 7114 | RFX1 | SOX2 regulated genes in Keratinocyte Overexpression from PMID: 30772301 | SOX2 |
| 7115 | TTC13 | SOX2 regulated genes in Keratinocyte Overexpression from PMID: 30772301 | SOX2 |
| 7116 | NGEF | SOX2 regulated genes in Keratinocyte Overexpression from PMID: 30772301 | SOX2 |
| 7117 | LRRC59 | SOX2 regulated genes in Keratinocyte Overexpression from PMID: 30772301 | SOX2 |
| 7118 | PDCD10 | SOX2 regulated genes in Keratinocyte Overexpression from PMID: 30772301 | SOX2 |
| 7119 | TOP2B | SOX2 regulated genes in Keratinocyte Overexpression from PMID: 30772301 | SOX2 |
| 7120 | OVCA2 | SOX2 regulated genes in Keratinocyte Overexpression from PMID: 30772301 | SOX2 |
| 7121 | COX7A2 | SOX2 regulated genes in Keratinocyte Overexpression from PMID: 30772301 | SOX2 |
| 7122 | SPSB2 | SOX2 regulated genes in Keratinocyte Overexpression from PMID: 30772301 | SOX2 |
| 7123 | SNAPC2 | SOX2 regulated genes in Keratinocyte Overexpression from PMID: 30772301 | SOX2 |
| 7124 | LETMD1 | SOX2 regulated genes in Keratinocyte Overexpression from PMID: 30772301 | SOX2 |
| 7125 | TRMT13 | SOX2 regulated genes in Keratinocyte Overexpression from PMID: 30772301 | SOX2 |
| 7126 | TRNAU1AP | SOX2 regulated genes in Keratinocyte Overexpression from PMID: 30772301 | SOX2 |
| 7127 | HEXIM2 | SOX2 regulated genes in Keratinocyte Overexpression from PMID: 30772301 | SOX2 |
| 7128 | C3orf62 | SOX2 regulated genes in Keratinocyte Overexpression from PMID: 30772301 | SOX2 |
| 7129 | KIFC1 | SOX2 regulated genes in Keratinocyte Overexpression from PMID: 30772301 | SOX2 |
| 7130 | PFDN6 | SOX2 regulated genes in Keratinocyte Overexpression from PMID: 30772301 | SOX2 |
| 7131 | PPM1D | SOX2 regulated genes in Keratinocyte Overexpression from PMID: 30772301 | SOX2 |
| 7132 | CD320 | SOX2 regulated genes in Keratinocyte Overexpression from PMID: 30772301 | SOX2 |
| 7133 | NUDC | SOX2 regulated genes in Keratinocyte Overexpression from PMID: 30772301 | SOX2 |
| 7134 | TRADD | SOX2 regulated genes in Keratinocyte Overexpression from PMID: 30772301 | SOX2 |
| 7135 | SSB | SOX2 regulated genes in Keratinocyte Overexpression from PMID: 30772301 | SOX2 |

| | | | |
|---|---|---|---|
| 7136 | MUL1 | SOX2 regulated genes in Keratinocyte Overexpression from PMID: 30772301 | SOX2 |
| 7137 | C16orf91 | SOX2 regulated genes in Keratinocyte Overexpression from PMID: 30772301 | SOX2 |
| 7138 | CHMP2B | SOX2 regulated genes in Keratinocyte Overexpression from PMID: 30772301 | SOX2 |
| 7139 | FAM169A | SOX2 regulated genes in Keratinocyte Overexpression from PMID: 30772301 | SOX2 |
| 7140 | TFPT | SOX2 regulated genes in Keratinocyte Overexpression from PMID: 30772301 | SOX2 |
| 7141 | ZFAND2B | SOX2 regulated genes in Keratinocyte Overexpression from PMID: 30772301 | SOX2 |
| 7142 | PHLDA3 | SOX2 regulated genes in Keratinocyte Overexpression from PMID: 30772301 | SOX2 |
| 7143 | ARAP2 | SOX2 regulated genes in Keratinocyte Overexpression from PMID: 30772301 | SOX2 |
| 7144 | ATP13A2 | SOX2 regulated genes in Keratinocyte Overexpression from PMID: 30772301 | SOX2 |
| 7145 | RAD9A | SOX2 regulated genes in Keratinocyte Overexpression from PMID: 30772301 | SOX2 |
| 7146 | PLEKHG6 | SOX2 regulated genes in Keratinocyte Overexpression from PMID: 30772301 | SOX2 |
| 7147 | AMT | SOX2 regulated genes in Keratinocyte Overexpression from PMID: 30772301 | SOX2 |
| 7148 | ERI2 | SOX2 regulated genes in Keratinocyte Overexpression from PMID: 30772301 | SOX2 |
| 7149 | CPNE4 | SOX2 regulated genes in Keratinocyte Overexpression from PMID: 30772301 | SOX2 |
| 7150 | KLK7 | SOX2 regulated genes in Keratinocyte Overexpression from PMID: 30772301 | SOX2 |
| 7151 | PYM1 | SOX2 regulated genes in Keratinocyte Overexpression from PMID: 30772301 | SOX2 |
| 7152 | ATG9B | SOX2 regulated genes in Keratinocyte Overexpression from PMID: 30772301 | SOX2 |
| 7153 | ECE1 | SOX2 regulated genes in Keratinocyte Overexpression from PMID: 30772301 | SOX2 |
| 7154 | ZNF768 | SOX2 regulated genes in Keratinocyte Overexpression from PMID: 30772301 | SOX2 |
| 7155 | NPLOC4 | SOX2 regulated genes in Keratinocyte Overexpression from PMID: 30772301 | SOX2 |
| 7156 | RAD54L | SOX2 regulated genes in Keratinocyte Overexpression from PMID: 30772301 | SOX2 |
| 7157 | SFSWAP | SOX2 regulated genes in Keratinocyte Overexpression from PMID: 30772301 | SOX2 |
| 7158 | ARF6 | SOX2 regulated genes in Keratinocyte Overexpression from PMID: 30772301 | SOX2 |
| 7159 | GSK3B | SOX2 regulated genes in Keratinocyte Overexpression from PMID: 30772301 | SOX2 |
| 7160 | SMYD3 | SOX2 regulated genes in Keratinocyte Overexpression from PMID: 30772301 | SOX2 |
| 7161 | VPS53 | SOX2 regulated genes in Keratinocyte Overexpression from PMID: 30772301 | SOX2 |
| 7162 | FBXO41 | SOX2 regulated genes in Keratinocyte Overexpression from PMID: 30772301 | SOX2 |
| 7163 | VPS26B | SOX2 regulated genes in Keratinocyte Overexpression from PMID: 30772301 | SOX2 |
| 7164 | CNOT3 | SOX2 regulated genes in Keratinocyte Overexpression from PMID: 30772301 | SOX2 |
| 7165 | SEC61A1 | SOX2 regulated genes in Keratinocyte Overexpression from PMID: 30772301 | SOX2 |
| 7166 | FBXW11 | SOX2 regulated genes in Keratinocyte Overexpression from PMID: 30772301 | SOX2 |
| 7167 | DUSP10 | SOX2 regulated genes in Keratinocyte Overexpression from PMID: 30772301 | SOX2 |
| 7168 | PPP1R13L | SOX2 regulated genes in Keratinocyte Overexpression from PMID: 30772301 | SOX2 |
| 7169 | ZFYVE27 | SOX2 regulated genes in Keratinocyte Overexpression from PMID: 30772301 | SOX2 |
| 7170 | BDP1 | SOX2 regulated genes in Keratinocyte Overexpression from PMID: 30772301 | SOX2 |
| 7171 | PDE6G | SOX2 regulated genes in Keratinocyte Overexpression from PMID: 30772301 | SOX2 |
| 7172 | KLF16 | SOX2 regulated genes in Keratinocyte Overexpression from PMID: 30772301 | SOX2 |
| 7173 | GPR19 | SOX2 regulated genes in Keratinocyte Overexpression from PMID: 30772301 | SOX2 |
| 7174 | RBM33 | SOX2 regulated genes in Keratinocyte Overexpression from PMID: 30772301 | SOX2 |
| 7175 | WDR55 | SOX2 regulated genes in Keratinocyte Overexpression from PMID: 30772301 | SOX2 |
| 7176 | ZBTB7A | SOX2 regulated genes in Keratinocyte Overexpression from PMID: 30772301 | SOX2 |
| 7177 | RBBP4 | SOX2 regulated genes in Keratinocyte Overexpression from PMID: 30772301 | SOX2 |
| 7178 | SNRPE | SOX2 regulated genes in Keratinocyte Overexpression from PMID: 30772301 | SOX2 |
| 7179 | MRPL24 | SOX2 regulated genes in Keratinocyte Overexpression from PMID: 30772301 | SOX2 |
| 7180 | PPHLN1 | SOX2 regulated genes in Keratinocyte Overexpression from PMID: 30772301 | SOX2 |
| 7181 | PCDH1 | SOX2 regulated genes in Keratinocyte Overexpression from PMID: 30772301 | SOX2 |
| 7182 | RUNDC1 | SOX2 regulated genes in Keratinocyte Overexpression from PMID: 30772301 | SOX2 |
| 7183 | YBX2 | SOX2 regulated genes in Keratinocyte Overexpression from PMID: 30772301 | SOX2 |
| 7184 | VPS9D1 | SOX2 regulated genes in Keratinocyte Overexpression from PMID: 30772301 | SOX2 |
| 7185 | RTCA | SOX2 regulated genes in Keratinocyte Overexpression from PMID: 30772301 | SOX2 |
| 7186 | HOXA5 | SOX2 regulated genes in Keratinocyte Overexpression from PMID: 30772301 | SOX2 |
| 7187 | RGL2 | SOX2 regulated genes in Keratinocyte Overexpression from PMID: 30772301 | SOX2 |
| 7188 | VIPAS39 | SOX2 regulated genes in Keratinocyte Overexpression from PMID: 30772301 | SOX2 |
| 7189 | SWT1 | SOX2 regulated genes in Keratinocyte Overexpression from PMID: 30772301 | SOX2 |
| 7190 | ATP5F1B | SOX2 regulated genes in Keratinocyte Overexpression from PMID: 30772301 | SOX2 |
| 7191 | DNMT1 | SOX2 regulated genes in Keratinocyte Overexpression from PMID: 30772301 | SOX2 |
| 7192 | SENP5 | SOX2 regulated genes in Keratinocyte Overexpression from PMID: 30772301 | SOX2 |
| 7193 | CLDN1 | SOX2 regulated genes in Keratinocyte Overexpression from PMID: 30772301 | SOX2 |
| 7194 | ABCB11 | SOX2 regulated genes in Keratinocyte Overexpression from PMID: 30772301 | SOX2 |
| 7195 | SCRN2 | SOX2 regulated genes in Keratinocyte Overexpression from PMID: 30772301 | SOX2 |
| 7196 | DLEC1 | SOX2 regulated genes in Keratinocyte Overexpression from PMID: 30772301 | SOX2 |

| | | | |
|---|---|---|---|
| 7197 | TEPSIN | SOX2 regulated genes in Keratinocyte Overexpression from PMID: 30772301 | SOX2 |
| 7198 | UBE2F | SOX2 regulated genes in Keratinocyte Overexpression from PMID: 30772301 | SOX2 |
| 7199 | ACVR2A | SOX2 regulated genes in Keratinocyte Overexpression from PMID: 30772301 | SOX2 |
| 7200 | BSDC1 | SOX2 regulated genes in Keratinocyte Overexpression from PMID: 30772301 | SOX2 |
| 7201 | KIF3C | SOX2 regulated genes in Keratinocyte Overexpression from PMID: 30772301 | SOX2 |
| 7202 | ZNF280D | SOX2 regulated genes in Keratinocyte Overexpression from PMID: 30772301 | SOX2 |
| 7203 | VPS25 | SOX2 regulated genes in Keratinocyte Overexpression from PMID: 30772301 | SOX2 |
| 7204 | BLM | SOX2 regulated genes in Keratinocyte Overexpression from PMID: 30772301 | SOX2 |
| 7205 | NMD3 | SOX2 regulated genes in Keratinocyte Overexpression from PMID: 30772301 | SOX2 |
| 7206 | NTF4 | SOX2 regulated genes in Keratinocyte Overexpression from PMID: 30772301 | SOX2 |
| 7207 | ZNF653 | SOX2 regulated genes in Keratinocyte Overexpression from PMID: 30772301 | SOX2 |
| 7208 | PRMT6 | SOX2 regulated genes in Keratinocyte Overexpression from PMID: 30772301 | SOX2 |
| 7209 | ATOX1 | SOX2 regulated genes in Keratinocyte Overexpression from PMID: 30772301 | SOX2 |
| 7210 | ZNF691 | SOX2 regulated genes in Keratinocyte Overexpression from PMID: 30772301 | SOX2 |
| 7211 | JUN | SOX2 regulated genes in Keratinocyte Overexpression from PMID: 30772301 | SOX2 |
| 7212 | MTMR4 | SOX2 regulated genes in Keratinocyte Overexpression from PMID: 30772301 | SOX2 |
| 7213 | MRPL4 | SOX2 regulated genes in Keratinocyte Overexpression from PMID: 30772301 | SOX2 |
| 7214 | USP5 | SOX2 regulated genes in Keratinocyte Overexpression from PMID: 30772301 | SOX2 |
| 7215 | MCOLN3 | SOX2 regulated genes in Keratinocyte Overexpression from PMID: 30772301 | SOX2 |
| 7216 | HNRNPR | SOX2 regulated genes in Keratinocyte Overexpression from PMID: 30772301 | SOX2 |
| 7217 | DAPK3 | SOX2 regulated genes in Keratinocyte Overexpression from PMID: 30772301 | SOX2 |
| 7218 | CTXN1 | SOX2 regulated genes in Keratinocyte Overexpression from PMID: 30772301 | SOX2 |
| 7219 | CACNB3 | SOX2 regulated genes in Keratinocyte Overexpression from PMID: 30772301 | SOX2 |
| 7220 | SLC45A3 | SOX2 regulated genes in Keratinocyte Overexpression from PMID: 30772301 | SOX2 |
| 7221 | TRIM45 | SOX2 regulated genes in Keratinocyte Overexpression from PMID: 30772301 | SOX2 |
| 7222 | PHYHIP | SOX2 regulated genes in Keratinocyte Overexpression from PMID: 30772301 | SOX2 |
| 7223 | TAF6 | SOX2 regulated genes in Keratinocyte Overexpression from PMID: 30772301 | SOX2 |
| 7224 | ZNF329 | SOX2 regulated genes in Keratinocyte Overexpression from PMID: 30772301 | SOX2 |
| 7225 | ERF | SOX2 regulated genes in Keratinocyte Overexpression from PMID: 30772301 | SOX2 |
| 7226 | NUP42 | SOX2 regulated genes in Keratinocyte Overexpression from PMID: 30772301 | SOX2 |
| 7227 | PIGF | SOX2 regulated genes in Keratinocyte Overexpression from PMID: 30772301 | SOX2 |
| 7228 | RIMS1 | SOX2 regulated genes in Keratinocyte Overexpression from PMID: 30772301 | SOX2 |
| 7229 | FAM20A | SOX2 regulated genes in Keratinocyte Overexpression from PMID: 30772301 | SOX2 |
| 7230 | EIF5B | SOX2 regulated genes in Keratinocyte Overexpression from PMID: 30772301 | SOX2 |
| 7231 | MXD1 | SOX2 regulated genes in Keratinocyte Overexpression from PMID: 30772301 | SOX2 |
| 7232 | RGMA | SOX2 regulated genes in Keratinocyte Overexpression from PMID: 30772301 | SOX2 |
| 7233 | RYBP | SOX2 regulated genes in Keratinocyte Overexpression from PMID: 30772301 | SOX2 |
| 7234 | NME7 | SOX2 regulated genes in Keratinocyte Overexpression from PMID: 30772301 | SOX2 |
| 7235 | NUP43 | SOX2 regulated genes in Keratinocyte Overexpression from PMID: 30772301 | SOX2 |
| 7236 | MLX | SOX2 regulated genes in Keratinocyte Overexpression from PMID: 30772301 | SOX2 |
| 7237 | CCS | SOX2 regulated genes in Keratinocyte Overexpression from PMID: 30772301 | SOX2 |
| 7238 | PSMB3 | SOX2 regulated genes in Keratinocyte Overexpression from PMID: 30772301 | SOX2 |
| 7239 | CBARP | SOX2 regulated genes in Keratinocyte Overexpression from PMID: 30772301 | SOX2 |
| 7240 | USP2 | SOX2 regulated genes in Keratinocyte Overexpression from PMID: 30772301 | SOX2 |
| 7241 | ITGA10 | SOX2 regulated genes in Keratinocyte Overexpression from PMID: 30772301 | SOX2 |
| 7242 | VPS72 | SOX2 regulated genes in Keratinocyte Overexpression from PMID: 30772301 | SOX2 |
| 7243 | PGBD1 | SOX2 regulated genes in Keratinocyte Overexpression from PMID: 30772301 | SOX2 |
| 7244 | SDE2 | SOX2 regulated genes in Keratinocyte Overexpression from PMID: 30772301 | SOX2 |
| 7245 | TSR3 | SOX2 regulated genes in Keratinocyte Overexpression from PMID: 30772301 | SOX2 |
| 7246 | KHDRBS1 | SOX2 regulated genes in Keratinocyte Overexpression from PMID: 30772301 | SOX2 |
| 7247 | COX10 | SOX2 regulated genes in Keratinocyte Overexpression from PMID: 30772301 | SOX2 |
| 7248 | EMP2 | SOX2 regulated genes in Keratinocyte Overexpression from PMID: 30772301 | SOX2 |
| 7249 | SIRT7 | SOX2 regulated genes in Keratinocyte Overexpression from PMID: 30772301 | SOX2 |
| 7250 | IFNLR1 | SOX2 regulated genes in Keratinocyte Overexpression from PMID: 30772301 | SOX2 |
| 7251 | LARP4B | SOX2 regulated genes in Keratinocyte Overexpression from PMID: 30772301 | SOX2 |
| 7252 | ANKRD11 | SOX2 regulated genes in Keratinocyte Overexpression from PMID: 30772301 | SOX2 |
| 7253 | FOXP1 | SOX2 regulated genes in Keratinocyte Overexpression from PMID: 30772301 | SOX2 |
| 7254 | WNT5B | SOX2 regulated genes in Keratinocyte Overexpression from PMID: 30772301 | SOX2 |
| 7255 | FAM214A | SOX2 regulated genes in Keratinocyte Overexpression from PMID: 30772301 | SOX2 |
| 7256 | STK19 | SOX2 regulated genes in Keratinocyte Overexpression from PMID: 30772301 | SOX2 |
| 7257 | EGR1 | SOX2 regulated genes in Keratinocyte Overexpression from PMID: 30772301 | SOX2 |

| | | | |
|---|---|---|---|
| 7258 | ZC3H8 | SOX2 regulated genes in Keratinocyte Overexpression from PMID: 30772301 | SOX2 |
| 7259 | GEMIN4 | SOX2 regulated genes in Keratinocyte Overexpression from PMID: 30772301 | SOX2 |
| 7260 | FABP12 | SOX2 regulated genes in Keratinocyte Overexpression from PMID: 30772301 | SOX2 |
| 7261 | MYH15 | SOX2 regulated genes in Keratinocyte Overexpression from PMID: 30772301 | SOX2 |
| 7262 | SERPINI1 | SOX2 regulated genes in Keratinocyte Overexpression from PMID: 30772301 | SOX2 |
| 7263 | DCTN1 | SOX2 regulated genes in Keratinocyte Overexpression from PMID: 30772301 | SOX2 |
| 7264 | LLPH | SOX2 regulated genes in Keratinocyte Overexpression from PMID: 30772301 | SOX2 |
| 7265 | SNAPC3 | SOX2 regulated genes in Keratinocyte Overexpression from PMID: 30772301 | SOX2 |
| 7266 | POU4F3 | SOX2 regulated genes in Keratinocyte Overexpression from PMID: 30772301 | SOX2 |
| 7267 | GRSF1 | SOX2 regulated genes in Keratinocyte Overexpression from PMID: 30772301 | SOX2 |
| 7268 | PRKAB1 | SOX2 regulated genes in Keratinocyte Overexpression from PMID: 30772301 | SOX2 |
| 7269 | ACTR8 | SOX2 regulated genes in Keratinocyte Overexpression from PMID: 30772301 | SOX2 |
| 7270 | CAMK2N1 | SOX2 regulated genes in Keratinocyte Overexpression from PMID: 30772301 | SOX2 |
| 7271 | SRSF4 | SOX2 regulated genes in Keratinocyte Overexpression from PMID: 30772301 | SOX2 |
| 7272 | ZNF512 | SOX2 regulated genes in Keratinocyte Overexpression from PMID: 30772301 | SOX2 |
| 7273 | GDPD1 | SOX2 regulated genes in Keratinocyte Overexpression from PMID: 30772301 | SOX2 |
| 7274 | SIRT1 | SOX2 regulated genes in Keratinocyte Overexpression from PMID: 30772301 | SOX2 |
| 7275 | NUP133 | SOX2 regulated genes in Keratinocyte Overexpression from PMID: 30772301 | SOX2 |
| 7276 | MKNK2 | SOX2 regulated genes in Keratinocyte Overexpression from PMID: 30772301 | SOX2 |
| 7277 | WDR53 | SOX2 regulated genes in Keratinocyte Overexpression from PMID: 30772301 | SOX2 |
| 7278 | MAPKBP1 | SOX2 regulated genes in Keratinocyte Overexpression from PMID: 30772301 | SOX2 |
| 7279 | SIRT4 | SOX2 regulated genes in Keratinocyte Overexpression from PMID: 30772301 | SOX2 |
| 7280 | PHB | SOX2 regulated genes in Keratinocyte Overexpression from PMID: 30772301 | SOX2 |
| 7281 | MACF1 | SOX2 regulated genes in Keratinocyte Overexpression from PMID: 30772301 | SOX2 |
| 7282 | TSG101 | SOX2 regulated genes in Keratinocyte Overexpression from PMID: 30772301 | SOX2 |
| 7283 | RNF208 | SOX2 regulated genes in Keratinocyte Overexpression from PMID: 30772301 | SOX2 |
| 7284 | ARHGAP22 | SOX2 regulated genes in Keratinocyte Overexpression from PMID: 30772301 | SOX2 |
| 7285 | GCDH | SOX2 regulated genes in Keratinocyte Overexpression from PMID: 30772301 | SOX2 |
| 7286 | SDF2 | SOX2 regulated genes in Keratinocyte Overexpression from PMID: 30772301 | SOX2 |
| 7287 | TKFC | SOX2 regulated genes in Keratinocyte Overexpression from PMID: 30772301 | SOX2 |
| 7288 | MZT2A | SOX2 regulated genes in Keratinocyte Overexpression from PMID: 30772301 | SOX2 |
| 7289 | MZT2B | SOX2 regulated genes in Keratinocyte Overexpression from PMID: 30772301 | SOX2 |
| 7290 | EPN1 | SOX2 regulated genes in Keratinocyte Overexpression from PMID: 30772301 | SOX2 |
| 7291 | DHODH | SOX2 regulated genes in Keratinocyte Overexpression from PMID: 30772301 | SOX2 |
| 7292 | TBCA | SOX2 regulated genes in Keratinocyte Overexpression from PMID: 30772301 | SOX2 |
| 7293 | MFSD2B | SOX2 regulated genes in Keratinocyte Overexpression from PMID: 30772301 | SOX2 |
| 7294 | MTX1 | SOX2 regulated genes in Keratinocyte Overexpression from PMID: 30772301 | SOX2 |
| 7295 | DCUN1D5 | SOX2 regulated genes in Keratinocyte Overexpression from PMID: 30772301 | SOX2 |
| 7296 | NINJ2 | SOX2 regulated genes in Keratinocyte Overexpression from PMID: 30772301 | SOX2 |
| 7297 | RHOU | SOX2 regulated genes in Keratinocyte Overexpression from PMID: 30772301 | SOX2 |
| 7298 | PPIP5K1 | SOX2 regulated genes in Keratinocyte Overexpression from PMID: 30772301 | SOX2 |
| 7299 | RPS15 | SOX2 regulated genes in Keratinocyte Overexpression from PMID: 30772301 | SOX2 |
| 7300 | FBXL18 | SOX2 regulated genes in Keratinocyte Overexpression from PMID: 30772301 | SOX2 |
| 7301 | FCAMR | SOX2 regulated genes in Keratinocyte Overexpression from PMID: 30772301 | SOX2 |
| 7302 | ALKBH7 | SOX2 regulated genes in Keratinocyte Overexpression from PMID: 30772301 | SOX2 |
| 7303 | C1orf109 | SOX2 regulated genes in Keratinocyte Overexpression from PMID: 30772301 | SOX2 |
| 7304 | ZNF800 | SOX2 regulated genes in Keratinocyte Overexpression from PMID: 30772301 | SOX2 |
| 7305 | USH2A | SOX2 regulated genes in Keratinocyte Overexpression from PMID: 30772301 | SOX2 |
| 7306 | AXDND1 | SOX2 regulated genes in Keratinocyte Overexpression from PMID: 30772301 | SOX2 |
| 7307 | TIMM22 | SOX2 regulated genes in Keratinocyte Overexpression from PMID: 30772301 | SOX2 |
| 7308 | C1orf35 | SOX2 regulated genes in Keratinocyte Overexpression from PMID: 30772301 | SOX2 |
| 7309 | PRKCQ | SOX2 regulated genes in Keratinocyte Overexpression from PMID: 30772301 | SOX2 |
| 7310 | MIGA1 | SOX2 regulated genes in Keratinocyte Overexpression from PMID: 30772301 | SOX2 |
| 7311 | PDXDC1 | SOX2 regulated genes in Keratinocyte Overexpression from PMID: 30772301 | SOX2 |
| 7312 | PTOV1 | SOX2 regulated genes in Keratinocyte Overexpression from PMID: 30772301 | SOX2 |
| 7313 | NFATC2IP | SOX2 regulated genes in Keratinocyte Overexpression from PMID: 30772301 | SOX2 |
| 7314 | STRIP1 | SOX2 regulated genes in Keratinocyte Overexpression from PMID: 30772301 | SOX2 |
| 7315 | MEX3D | SOX2 regulated genes in Keratinocyte Overexpression from PMID: 30772301 | SOX2 |
| 7316 | ASB7 | SOX2 regulated genes in Keratinocyte Overexpression from PMID: 30772301 | SOX2 |
| 7317 | PRSS22 | SOX2 regulated genes in Keratinocyte Overexpression from PMID: 30772301 | SOX2 |
| 7318 | HGS | SOX2 regulated genes in Keratinocyte Overexpression from PMID: 30772301 | SOX2 |

| | | | |
|---|---|---|---|
| 7319 | NKIRAS2 | SOX2 regulated genes in Keratinocyte Overexpression from PMID: 30772301 | SOX2 |
| 7320 | MGST3 | SOX2 regulated genes in Keratinocyte Overexpression from PMID: 30772301 | SOX2 |
| 7321 | CAPN10 | SOX2 regulated genes in Keratinocyte Overexpression from PMID: 30772301 | SOX2 |
| 7322 | EME1 | SOX2 regulated genes in Keratinocyte Overexpression from PMID: 30772301 | SOX2 |
| 7323 | DLG1 | SOX2 regulated genes in Keratinocyte Overexpression from PMID: 30772301 | SOX2 |
| 7324 | SHH | SOX2 regulated genes in Keratinocyte Overexpression from PMID: 30772301 | SOX2 |
| 7325 | PRPF3 | SOX2 regulated genes in Keratinocyte Overexpression from PMID: 30772301 | SOX2 |
| 7326 | DUS3L | SOX2 regulated genes in Keratinocyte Overexpression from PMID: 30772301 | SOX2 |
| 7327 | FAT2 | SOX2 regulated genes in Keratinocyte Overexpression from PMID: 30772301 | SOX2 |
| 7328 | PDHB | SOX2 regulated genes in Keratinocyte Overexpression from PMID: 30772301 | SOX2 |
| 7329 | TOMM40 | SOX2 regulated genes in Keratinocyte Overexpression from PMID: 30772301 | SOX2 |
| 7330 | ORC1 | SOX2 regulated genes in Keratinocyte Overexpression from PMID: 30772301 | SOX2 |
| 7331 | SLC44A2 | SOX2 regulated genes in Keratinocyte Overexpression from PMID: 30772301 | SOX2 |
| 7332 | MRPS5 | SOX2 regulated genes in Keratinocyte Overexpression from PMID: 30772301 | SOX2 |
| 7333 | TFAP2E | SOX2 regulated genes in Keratinocyte Overexpression from PMID: 30772301 | SOX2 |
| 7334 | KIF2A | SOX2 regulated genes in Keratinocyte Overexpression from PMID: 30772301 | SOX2 |
| 7335 | TADA3 | SOX2 regulated genes in Keratinocyte Overexpression from PMID: 30772301 | SOX2 |
| 7336 | RRP8 | SOX2 regulated genes in Keratinocyte Overexpression from PMID: 30772301 | SOX2 |
| 7337 | PFDN2 | SOX2 regulated genes in Keratinocyte Overexpression from PMID: 30772301 | SOX2 |
| 7338 | ZNF513 | SOX2 regulated genes in Keratinocyte Overexpression from PMID: 30772301 | SOX2 |
| 7339 | B3GNTL1 | SOX2 regulated genes in Keratinocyte Overexpression from PMID: 30772301 | SOX2 |
| 7340 | S100A2 | SOX2 regulated genes in Keratinocyte Overexpression from PMID: 30772301 | SOX2 |
| 7341 | LRIG3 | SOX2 regulated genes in Keratinocyte Overexpression from PMID: 30772301 | SOX2 |
| 7342 | LUZP1 | SOX2 regulated genes in Keratinocyte Overexpression from PMID: 30772301 | SOX2 |
| 7343 | C8orf58 | SOX2 regulated genes in Keratinocyte Overexpression from PMID: 30772301 | SOX2 |
| 7344 | SSR2 | SOX2 regulated genes in Keratinocyte Overexpression from PMID: 30772301 | SOX2 |
| 7345 | CDC42BPA | SOX2 regulated genes in Keratinocyte Overexpression from PMID: 30772301 | SOX2 |
| 7346 | USP40 | SOX2 regulated genes in Keratinocyte Overexpression from PMID: 30772301 | SOX2 |
| 7347 | SLC5A10 | SOX2 regulated genes in Keratinocyte Overexpression from PMID: 30772301 | SOX2 |
| 7348 | SMG6 | SOX2 regulated genes in Keratinocyte Overexpression from PMID: 30772301 | SOX2 |
| 7349 | EFCAB5 | SOX2 regulated genes in Keratinocyte Overexpression from PMID: 30772301 | SOX2 |
| 7350 | ATP1B1 | SOX2 regulated genes in Keratinocyte Overexpression from PMID: 30772301 | SOX2 |
| 7351 | DENND5B | SOX2 regulated genes in Keratinocyte Overexpression from PMID: 30772301 | SOX2 |
| 7352 | CNOT6L | SOX2 regulated genes in Keratinocyte Overexpression from PMID: 30772301 | SOX2 |
| 7353 | LIMA1 | SOX2 regulated genes in Keratinocyte Overexpression from PMID: 30772301 | SOX2 |
| 7354 | EFNA4 | SOX2 regulated genes in Keratinocyte Overexpression from PMID: 30772301 | SOX2 |
| 7355 | RAF1 | SOX2 regulated genes in Keratinocyte Overexpression from PMID: 30772301 | SOX2 |
| 7356 | CRIPT | SOX2 regulated genes in Keratinocyte Overexpression from PMID: 30772301 | SOX2 |
| 7357 | TBCEL | SOX2 regulated genes in Keratinocyte Overexpression from PMID: 30772301 | SOX2 |
| 7358 | NIPAL4 | SOX2 regulated genes in Keratinocyte Overexpression from PMID: 30772301 | SOX2 |
| 7359 | LRRC14 | SOX2 regulated genes in Keratinocyte Overexpression from PMID: 30772301 | SOX2 |
| 7360 | TMEM40 | SOX2 regulated genes in Keratinocyte Overexpression from PMID: 30772301 | SOX2 |
| 7361 | GRM3 | SOX2 regulated genes in Keratinocyte Overexpression from PMID: 30772301 | SOX2 |
| 7362 | DDX46 | SOX2 regulated genes in Keratinocyte Overexpression from PMID: 30772301 | SOX2 |
| 7363 | SLC16A10 | SOX2 regulated genes in Keratinocyte Overexpression from PMID: 30772301 | SOX2 |
| 7364 | AURKC | SOX2 regulated genes in Keratinocyte Overexpression from PMID: 30772301 | SOX2 |
| 7365 | PASK | SOX2 regulated genes in Keratinocyte Overexpression from PMID: 30772301 | SOX2 |
| 7366 | LARP4 | SOX2 regulated genes in Keratinocyte Overexpression from PMID: 30772301 | SOX2 |
| 7367 | PARD3B | SOX2 regulated genes in Keratinocyte Overexpression from PMID: 30772301 | SOX2 |
| 7368 | HJURP | SOX2 regulated genes in Keratinocyte Overexpression from PMID: 30772301 | SOX2 |
| 7369 | TBCB | SOX2 regulated genes in Keratinocyte Overexpression from PMID: 30772301 | SOX2 |
| 7370 | PGAP1 | SOX2 regulated genes in Keratinocyte Overexpression from PMID: 30772301 | SOX2 |
| 7371 | PSMA5 | SOX2 regulated genes in Keratinocyte Overexpression from PMID: 30772301 | SOX2 |
| 7372 | SLC27A6 | SOX2 regulated genes in Keratinocyte Overexpression from PMID: 30772301 | SOX2 |
| 7373 | ETV3 | SOX2 regulated genes in Keratinocyte Overexpression from PMID: 30772301 | SOX2 |
| 7374 | RBM8A | SOX2 regulated genes in Keratinocyte Overexpression from PMID: 30772301 | SOX2 |
| 7375 | MEGF11 | SOX2 regulated genes in Keratinocyte Overexpression from PMID: 30772301 | SOX2 |
| 7376 | IQGAP3 | SOX2 regulated genes in Keratinocyte Overexpression from PMID: 30772301 | SOX2 |
| 7377 | TSEN34 | SOX2 regulated genes in Keratinocyte Overexpression from PMID: 30772301 | SOX2 |
| 7378 | PHF13 | SOX2 regulated genes in Keratinocyte Overexpression from PMID: 30772301 | SOX2 |
| 7379 | TARS2 | SOX2 regulated genes in Keratinocyte Overexpression from PMID: 30772301 | SOX2 |

| | | | |
|---|---|---|---|
| 7380 | FYTTD1 | SOX2 regulated genes in Keratinocyte Overexpression from PMID: 30772301 | SOX2 |
| 7381 | ZKSCAN4 | SOX2 regulated genes in Keratinocyte Overexpression from PMID: 30772301 | SOX2 |
| 7382 | SNX16 | SOX2 regulated genes in Keratinocyte Overexpression from PMID: 30772301 | SOX2 |
| 7383 | PFN2 | SOX2 regulated genes in Keratinocyte Overexpression from PMID: 30772301 | SOX2 |
| 7384 | RNF32 | SOX2 regulated genes in Keratinocyte Overexpression from PMID: 30772301 | SOX2 |
| 7385 | DISP2 | SOX2 regulated genes in Keratinocyte Overexpression from PMID: 30772301 | SOX2 |
| 7386 | RIF1 | SOX2 regulated genes in Keratinocyte Overexpression from PMID: 30772301 | SOX2 |
| 7387 | SH2B1 | SOX2 regulated genes in Keratinocyte Overexpression from PMID: 30772301 | SOX2 |
| 7388 | TLK2 | SOX2 regulated genes in Keratinocyte Overexpression from PMID: 30772301 | SOX2 |
| 7389 | PSMB6 | SOX2 regulated genes in Keratinocyte Overexpression from PMID: 30772301 | SOX2 |
| 7390 | BEAN1 | SOX2 regulated genes in Keratinocyte Overexpression from PMID: 30772301 | SOX2 |
| 7391 | TRMT112 | SOX2 regulated genes in Keratinocyte Overexpression from PMID: 30772301 | SOX2 |
| 7392 | TARBP1 | SOX2 regulated genes in Keratinocyte Overexpression from PMID: 30772301 | SOX2 |
| 7393 | COQ10B | SOX2 regulated genes in Keratinocyte Overexpression from PMID: 30772301 | SOX2 |
| 7394 | SHF | SOX2 regulated genes in Keratinocyte Overexpression from PMID: 30772301 | SOX2 |
| 7395 | USF1 | SOX2 regulated genes in Keratinocyte Overexpression from PMID: 30772301 | SOX2 |
| 7396 | RPGRIP1L | SOX2 regulated genes in Keratinocyte Overexpression from PMID: 30772301 | SOX2 |
| 7397 | AXIN2 | SOX2 regulated genes in Keratinocyte Overexpression from PMID: 30772301 | SOX2 |
| 7398 | LIN37 | SOX2 regulated genes in Keratinocyte Overexpression from PMID: 30772301 | SOX2 |
| 7399 | INAFM2 | SOX2 regulated genes in Keratinocyte Overexpression from PMID: 30772301 | SOX2 |
| 7400 | PSMB4 | SOX2 regulated genes in Keratinocyte Overexpression from PMID: 30772301 | SOX2 |
| 7401 | ABCG4 | SOX2 regulated genes in Keratinocyte Overexpression from PMID: 30772301 | SOX2 |
| 7402 | NUCKS1 | SOX2 regulated genes in Keratinocyte Overexpression from PMID: 30772301 | SOX2 |
| 7403 | MKRN2OS | SOX2 regulated genes in Keratinocyte Overexpression from PMID: 30772301 | SOX2 |
| 7404 | HIP1R | SOX2 regulated genes in Keratinocyte Overexpression from PMID: 30772301 | SOX2 |
| 7405 | METTL8 | SOX2 regulated genes in Keratinocyte Overexpression from PMID: 30772301 | SOX2 |
| 7406 | IQCC | SOX2 regulated genes in Keratinocyte Overexpression from PMID: 30772301 | SOX2 |
| 7407 | CLDN7 | SOX2 regulated genes in Keratinocyte Overexpression from PMID: 30772301 | SOX2 |
| 7408 | TCEA1 | SOX2 regulated genes in Keratinocyte Overexpression from PMID: 30772301 | SOX2 |
| 7409 | KLC3 | SOX2 regulated genes in Keratinocyte Overexpression from PMID: 30772301 | SOX2 |
| 7410 | DPH5 | SOX2 regulated genes in Keratinocyte Overexpression from PMID: 30772301 | SOX2 |
| 7411 | PCBP2 | SOX2 regulated genes in Keratinocyte Overexpression from PMID: 30772301 | SOX2 |
| 7412 | NOL6 | SOX2 regulated genes in Keratinocyte Overexpression from PMID: 30772301 | SOX2 |
| 7413 | OOEP | SOX2 regulated genes in Keratinocyte Overexpression from PMID: 30772301 | SOX2 |
| 7414 | NMT1 | SOX2 regulated genes in Keratinocyte Overexpression from PMID: 30772301 | SOX2 |
| 7415 | HR | SOX2 regulated genes in Keratinocyte Overexpression from PMID: 30772301 | SOX2 |
| 7416 | DROSHA | SOX2 regulated genes in Keratinocyte Overexpression from PMID: 30772301 | SOX2 |
| 7417 | PELP1 | SOX2 regulated genes in Keratinocyte Overexpression from PMID: 30772301 | SOX2 |
| 7418 | SPRR4 | SOX2 regulated genes in Keratinocyte Overexpression from PMID: 30772301 | SOX2 |
| 7419 | SNAPC4 | SOX2 regulated genes in Keratinocyte Overexpression from PMID: 30772301 | SOX2 |
| 7420 | TAF1C | SOX2 regulated genes in Keratinocyte Overexpression from PMID: 30772301 | SOX2 |
| 7421 | CGNL1 | SOX2 regulated genes in Keratinocyte Overexpression from PMID: 30772301 | SOX2 |
| 7422 | WDR83OS | SOX2 regulated genes in Keratinocyte Overexpression from PMID: 30772301 | SOX2 |
| 7423 | DUSP16 | SOX2 regulated genes in Keratinocyte Overexpression from PMID: 30772301 | SOX2 |
| 7424 | THOP1 | SOX2 regulated genes in Keratinocyte Overexpression from PMID: 30772301 | SOX2 |
| 7425 | SUSD4 | SOX2 regulated genes in Keratinocyte Overexpression from PMID: 30772301 | SOX2 |
| 7426 | CDC42SE1 | SOX2 regulated genes in Keratinocyte Overexpression from PMID: 30772301 | SOX2 |
| 7427 | MFF | SOX2 regulated genes in Keratinocyte Overexpression from PMID: 30772301 | SOX2 |
| 7428 | IER5 | SOX2 regulated genes in Keratinocyte Overexpression from PMID: 30772301 | SOX2 |
| 7429 | RAD21 | SOX2 regulated genes in Keratinocyte Overexpression from PMID: 30772301 | SOX2 |
| 7430 | SETD5 | SOX2 regulated genes in Keratinocyte Overexpression from PMID: 30772301 | SOX2 |
| 7431 | MBTPS1 | SOX2 regulated genes in Keratinocyte Overexpression from PMID: 30772301 | SOX2 |
| 7432 | REXO1 | SOX2 regulated genes in Keratinocyte Overexpression from PMID: 30772301 | SOX2 |
| 7433 | SRPRB | SOX2 regulated genes in Keratinocyte Overexpression from PMID: 30772301 | SOX2 |
| 7434 | AKTIP | SOX2 regulated genes in Keratinocyte Overexpression from PMID: 30772301 | SOX2 |
| 7435 | RNF223 | SOX2 regulated genes in Keratinocyte Overexpression from PMID: 30772301 | SOX2 |
| 7436 | TIMM13 | SOX2 regulated genes in Keratinocyte Overexpression from PMID: 30772301 | SOX2 |
| 7437 | RABL3 | SOX2 regulated genes in Keratinocyte Overexpression from PMID: 30772301 | SOX2 |
| 7438 | XRRA1 | SOX2 regulated genes in Keratinocyte Overexpression from PMID: 30772301 | SOX2 |
| 7439 | PIGL | SOX2 regulated genes in Keratinocyte Overexpression from PMID: 30772301 | SOX2 |
| 7440 | MARCKSL1 | SOX2 regulated genes in Keratinocyte Overexpression from PMID: 30772301 | SOX2 |

| | | | |
|---|---|---|---|
| 7441 | PUS7L | SOX2 regulated genes in Keratinocyte Overexpression from PMID: 30772301 | SOX2 |
| 7442 | CCNE2 | SOX2 regulated genes in Keratinocyte Overexpression from PMID: 30772301 | SOX2 |
| 7443 | HOOK2 | SOX2 regulated genes in Keratinocyte Overexpression from PMID: 30772301 | SOX2 |
| 7444 | HNRNPH3 | SOX2 regulated genes in Keratinocyte Overexpression from PMID: 30772301 | SOX2 |
| 7445 | LONP1 | SOX2 regulated genes in Keratinocyte Overexpression from PMID: 30772301 | SOX2 |
| 7446 | MRPL53 | SOX2 regulated genes in Keratinocyte Overexpression from PMID: 30772301 | SOX2 |
| 7447 | ANXA9 | SOX2 regulated genes in Keratinocyte Overexpression from PMID: 30772301 | SOX2 |
| 7448 | GALR2 | SOX2 regulated genes in Keratinocyte Overexpression from PMID: 30772301 | SOX2 |
| 7449 | PRPF40A | SOX2 regulated genes in Keratinocyte Overexpression from PMID: 30772301 | SOX2 |
| 7450 | TFCP2 | SOX2 regulated genes in Keratinocyte Overexpression from PMID: 30772301 | SOX2 |
| 7451 | RING1 | SOX2 regulated genes in Keratinocyte Overexpression from PMID: 30772301 | SOX2 |
| 7452 | GBF1 | SOX2 regulated genes in Keratinocyte Overexpression from PMID: 30772301 | SOX2 |
| 7453 | CCDC51 | SOX2 regulated genes in Keratinocyte Overexpression from PMID: 30772301 | SOX2 |
| 7454 | PDGFA | SOX2 regulated genes in Keratinocyte Overexpression from PMID: 30772301 | SOX2 |
| 7455 | CEP72 | SOX2 regulated genes in Keratinocyte Overexpression from PMID: 30772301 | SOX2 |
| 7456 | HCN3 | SOX2 regulated genes in Keratinocyte Overexpression from PMID: 30772301 | SOX2 |
| 7457 | TMEM181 | SOX2 regulated genes in Keratinocyte Overexpression from PMID: 30772301 | SOX2 |
| 7458 | VSIG10L | SOX2 regulated genes in Keratinocyte Overexpression from PMID: 30772301 | SOX2 |
| 7459 | CUL2 | SOX2 regulated genes in Keratinocyte Overexpression from PMID: 30772301 | SOX2 |
| 7460 | TSN | SOX2 regulated genes in Keratinocyte Overexpression from PMID: 30772301 | SOX2 |
| 7461 | NKD1 | SOX2 regulated genes in Keratinocyte Overexpression from PMID: 30772301 | SOX2 |
| 7462 | SLC39A14 | SOX2 regulated genes in Keratinocyte Overexpression from PMID: 30772301 | SOX2 |
| 7463 | GRM5 | SOX2 regulated genes in Keratinocyte Overexpression from PMID: 30772301 | SOX2 |
| 7464 | NIT1 | SOX2 regulated genes in Keratinocyte Overexpression from PMID: 30772301 | SOX2 |
| 7465 | DND1 | SOX2 regulated genes in Keratinocyte Overexpression from PMID: 30772301 | SOX2 |
| 7466 | LMNTD1 | SOX2 regulated genes in Keratinocyte Overexpression from PMID: 30772301 | SOX2 |
| 7467 | CDH1 | SOX2 regulated genes in Keratinocyte Overexpression from PMID: 30772301 | SOX2 |
| 7468 | FRS2 | SOX2 regulated genes in Keratinocyte Overexpression from PMID: 30772301 | SOX2 |
| 7469 | AP4B1 | SOX2 regulated genes in Keratinocyte Overexpression from PMID: 30772301 | SOX2 |
| 7470 | SGK3 | SOX2 regulated genes in Keratinocyte Overexpression from PMID: 30772301 | SOX2 |
| 7471 | RAD23A | SOX2 regulated genes in Keratinocyte Overexpression from PMID: 30772301 | SOX2 |
| 7472 | POM121 | SOX2 regulated genes in Keratinocyte Overexpression from PMID: 30772301 | SOX2 |
| 7473 | POM121C | SOX2 regulated genes in Keratinocyte Overexpression from PMID: 30772301 | SOX2 |
| 7474 | BRICD5 | SOX2 regulated genes in Keratinocyte Overexpression from PMID: 30772301 | SOX2 |
| 7475 | TANC2 | SOX2 regulated genes in Keratinocyte Overexpression from PMID: 30772301 | SOX2 |
| 7476 | SLC41A1 | SOX2 regulated genes in Keratinocyte Overexpression from PMID: 30772301 | SOX2 |
| 7477 | CACNA2D2 | SOX2 regulated genes in Keratinocyte Overexpression from PMID: 30772301 | SOX2 |
| 7478 | RNF157 | SOX2 regulated genes in Keratinocyte Overexpression from PMID: 30772301 | SOX2 |
| 7479 | DAP3 | SOX2 regulated genes in Keratinocyte Overexpression from PMID: 30772301 | SOX2 |
| 7480 | MAP3K13 | SOX2 regulated genes in Keratinocyte Overexpression from PMID: 30772301 | SOX2 |
| 7481 | CNNM4 | SOX2 regulated genes in Keratinocyte Overexpression from PMID: 30772301 | SOX2 |
| 7482 | CWC27 | SOX2 regulated genes in Keratinocyte Overexpression from PMID: 30772301 | SOX2 |
| 7483 | MRPL55 | SOX2 regulated genes in Keratinocyte Overexpression from PMID: 30772301 | SOX2 |
| 7484 | RAI1 | SOX2 regulated genes in Keratinocyte Overexpression from PMID: 30772301 | SOX2 |
| 7485 | CCSAP | SOX2 regulated genes in Keratinocyte Overexpression from PMID: 30772301 | SOX2 |
| 7486 | EZH2 | SOX2 regulated genes in Keratinocyte Overexpression from PMID: 30772301 | SOX2 |
| 7487 | ARHGAP27 | SOX2 regulated genes in Keratinocyte Overexpression from PMID: 30772301 | SOX2 |
| 7488 | SRFBP1 | SOX2 regulated genes in Keratinocyte Overexpression from PMID: 30772301 | SOX2 |
| 7489 | TMEM41B | SOX2 regulated genes in Keratinocyte Overexpression from PMID: 30772301 | SOX2 |
| 7490 | QRICH2 | SOX2 regulated genes in Keratinocyte Overexpression from PMID: 30772301 | SOX2 |
| 7491 | TNNI1 | SOX2 regulated genes in Keratinocyte Overexpression from PMID: 30772301 | SOX2 |
| 7492 | TUBA4A | SOX2 regulated genes in Keratinocyte Overexpression from PMID: 30772301 | SOX2 |
| 7493 | SYNRG | SOX2 regulated genes in Keratinocyte Overexpression from PMID: 30772301 | SOX2 |
| 7494 | RARB | SOX2 regulated genes in Keratinocyte Overexpression from PMID: 30772301 | SOX2 |
| 7495 | HSF4 | SOX2 regulated genes in Keratinocyte Overexpression from PMID: 30772301 | SOX2 |
| 7496 | SRSF1 | SOX2 regulated genes in Keratinocyte Overexpression from PMID: 30772301 | SOX2 |
| 7497 | DMPK | SOX2 regulated genes in Keratinocyte Overexpression from PMID: 30772301 | SOX2 |
| 7498 | EIF4EBP3 | SOX2 regulated genes in Keratinocyte Overexpression from PMID: 30772301 | SOX2 |
| 7499 | C1orf131 | SOX2 regulated genes in Keratinocyte Overexpression from PMID: 30772301 | SOX2 |
| 7500 | IL11 | SOX2 regulated genes in Keratinocyte Overexpression from PMID: 30772301 | SOX2 |
| 7501 | SAXO2 | SOX2 regulated genes in Keratinocyte Overexpression from PMID: 30772301 | SOX2 |

| | | | |
|---|---|---|---|
| 7502 | PNKP | SOX2 regulated genes in Keratinocyte Overexpression from PMID: 30772301 | SOX2 |
| 7503 | C1orf74 | SOX2 regulated genes in Keratinocyte Overexpression from PMID: 30772301 | SOX2 |
| 7504 | DVL1 | SOX2 regulated genes in Keratinocyte Overexpression from PMID: 30772301 | SOX2 |
| 7505 | SLC19A3 | SOX2 regulated genes in Keratinocyte Overexpression from PMID: 30772301 | SOX2 |
| 7506 | PMS1 | SOX2 regulated genes in Keratinocyte Overexpression from PMID: 30772301 | SOX2 |
| 7507 | ECI2 | SOX2 regulated genes in Keratinocyte Overexpression from PMID: 30772301 | SOX2 |
| 7508 | CDC42EP5 | SOX2 regulated genes in Keratinocyte Overexpression from PMID: 30772301 | SOX2 |
| 7509 | DPP9 | SOX2 regulated genes in Keratinocyte Overexpression from PMID: 30772301 | SOX2 |
| 7510 | MYL6B | SOX2 regulated genes in Keratinocyte Overexpression from PMID: 30772301 | SOX2 |
| 7511 | DTL | SOX2 regulated genes in Keratinocyte Overexpression from PMID: 30772301 | SOX2 |
| 7512 | PCGF2 | SOX2 regulated genes in Keratinocyte Overexpression from PMID: 30772301 | SOX2 |
| 7513 | TAF10 | SOX2 regulated genes in Keratinocyte Overexpression from PMID: 30772301 | SOX2 |
| 7514 | PCF11 | SOX2 regulated genes in Keratinocyte Overexpression from PMID: 30772301 | SOX2 |
| 7515 | SLC23A3 | SOX2 regulated genes in Keratinocyte Overexpression from PMID: 30772301 | SOX2 |
| 7516 | PROSER3 | SOX2 regulated genes in Keratinocyte Overexpression from PMID: 30772301 | SOX2 |
| 7517 | SETD6 | SOX2 regulated genes in Keratinocyte Overexpression from PMID: 30772301 | SOX2 |
| 7518 | PARL | SOX2 regulated genes in Keratinocyte Overexpression from PMID: 30772301 | SOX2 |
| 7519 | IFRD2 | SOX2 regulated genes in Keratinocyte Overexpression from PMID: 30772301 | SOX2 |
| 7520 | RPL23A | SOX2 regulated genes in Keratinocyte Overexpression from PMID: 30772301 | SOX2 |
| 7521 | LANCL2 | SOX2 regulated genes in Keratinocyte Overexpression from PMID: 30772301 | SOX2 |
| 7522 | HBEGF | SOX2 regulated genes in Keratinocyte Overexpression from PMID: 30772301 | SOX2 |
| 7523 | YME1L1 | SOX2 regulated genes in Keratinocyte Overexpression from PMID: 30772301 | SOX2 |
| 7524 | THUMPD1 | SOX2 regulated genes in Keratinocyte Overexpression from PMID: 30772301 | SOX2 |
| 7525 | MRPL58 | SOX2 regulated genes in Keratinocyte Overexpression from PMID: 30772301 | SOX2 |
| 7526 | ANKRD39 | SOX2 regulated genes in Keratinocyte Overexpression from PMID: 30772301 | SOX2 |
| 7527 | AUNIP | SOX2 regulated genes in Keratinocyte Overexpression from PMID: 30772301 | SOX2 |
| 7528 | CNST | SOX2 regulated genes in Keratinocyte Overexpression from PMID: 30772301 | SOX2 |
| 7529 | ECSIT | SOX2 regulated genes in Keratinocyte Overexpression from PMID: 30772301 | SOX2 |
| 7530 | RPS27 | SOX2 regulated genes in Keratinocyte Overexpression from PMID: 30772301 | SOX2 |
| 7531 | GRIN2A | SOX2 regulated genes in Keratinocyte Overexpression from PMID: 30772301 | SOX2 |
| 7532 | HDGF | SOX2 regulated genes in Keratinocyte Overexpression from PMID: 30772301 | SOX2 |
| 7533 | F12 | SOX2 regulated genes in Keratinocyte Overexpression from PMID: 30772301 | SOX2 |
| 7534 | SIK3 | SOX2 regulated genes in Keratinocyte Overexpression from PMID: 30772301 | SOX2 |
| 7535 | GLOD4 | SOX2 regulated genes in Keratinocyte Overexpression from PMID: 30772301 | SOX2 |
| 7536 | NDUFAF7 | SOX2 regulated genes in Keratinocyte Overexpression from PMID: 30772301 | SOX2 |
| 7537 | MDH2 | SOX2 regulated genes in Keratinocyte Overexpression from PMID: 30772301 | SOX2 |
| 7538 | SBNO1 | SOX2 regulated genes in Keratinocyte Overexpression from PMID: 30772301 | SOX2 |
| 7539 | GPR42 | SOX2 regulated genes in Keratinocyte Overexpression from PMID: 30772301 | SOX2 |
| 7540 | SIN3B | SOX2 regulated genes in Keratinocyte Overexpression from PMID: 30772301 | SOX2 |
| 7541 | ESPNL | SOX2 regulated genes in Keratinocyte Overexpression from PMID: 30772301 | SOX2 |
| 7542 | DHRS9 | SOX2 regulated genes in Keratinocyte Overexpression from PMID: 30772301 | SOX2 |
| 7543 | CCDC78 | SOX2 regulated genes in Keratinocyte Overexpression from PMID: 30772301 | SOX2 |
| 7544 | GTF2F1 | SOX2 regulated genes in Keratinocyte Overexpression from PMID: 30772301 | SOX2 |
| 7545 | DVL2 | SOX2 regulated genes in Keratinocyte Overexpression from PMID: 30772301 | SOX2 |
| 7546 | RECQL5 | SOX2 regulated genes in Keratinocyte Overexpression from PMID: 30772301 | SOX2 |
| 7547 | TATDN1 | SOX2 regulated genes in Keratinocyte Overexpression from PMID: 30772301 | SOX2 |
| 7548 | ALOX12 | SOX2 regulated genes in Keratinocyte Overexpression from PMID: 30772301 | SOX2 |
| 7549 | MRPS27 | SOX2 regulated genes in Keratinocyte Overexpression from PMID: 30772301 | SOX2 |
| 7550 | LSG1 | SOX2 regulated genes in Keratinocyte Overexpression from PMID: 30772301 | SOX2 |
| 7551 | METTL23 | SOX2 regulated genes in Keratinocyte Overexpression from PMID: 30772301 | SOX2 |
| 7552 | POGK | SOX2 regulated genes in Keratinocyte Overexpression from PMID: 30772301 | SOX2 |
| 7553 | MRPS22 | SOX2 regulated genes in Keratinocyte Overexpression from PMID: 30772301 | SOX2 |
| 7554 | HP1BP3 | SOX2 regulated genes in Keratinocyte Overexpression from PMID: 30772301 | SOX2 |
| 7555 | CAPN5 | SOX2 regulated genes in Keratinocyte Overexpression from PMID: 30772301 | SOX2 |
| 7556 | SRCAP | SOX2 regulated genes in Keratinocyte Overexpression from PMID: 30772301 | SOX2 |
| 7557 | GTF3C2 | SOX2 regulated genes in Keratinocyte Overexpression from PMID: 30772301 | SOX2 |
| 7558 | TMA7 | SOX2 regulated genes in Keratinocyte Overexpression from PMID: 30772301 | SOX2 |
| 7559 | PPP1R16A | SOX2 regulated genes in Keratinocyte Overexpression from PMID: 30772301 | SOX2 |
| 7560 | HOXB5 | SOX2 regulated genes in Keratinocyte Overexpression from PMID: 30772301 | SOX2 |
| 7561 | ENOPH1 | SOX2 regulated genes in Keratinocyte Overexpression from PMID: 30772301 | SOX2 |
| 7562 | ZNF146 | SOX2 regulated genes in Keratinocyte Overexpression from PMID: 30772301 | SOX2 |

| | | | |
|---|---|---|---|
| 7563 | POLR2D | SOX2 regulated genes in Keratinocyte Overexpression from PMID: 30772301 | SOX2 |
| 7564 | DCAF15 | SOX2 regulated genes in Keratinocyte Overexpression from PMID: 30772301 | SOX2 |
| 7565 | FAT1 | SOX2 regulated genes in Keratinocyte Overexpression from PMID: 30772301 | SOX2 |
| 7566 | SDR16C5 | SOX2 regulated genes in Keratinocyte Overexpression from PMID: 30772301 | SOX2 |
| 7567 | TRNT1 | SOX2 regulated genes in Keratinocyte Overexpression from PMID: 30772301 | SOX2 |
| 7568 | BCL2L12 | SOX2 regulated genes in Keratinocyte Overexpression from PMID: 30772301 | SOX2 |
| 7569 | LMBR1L | SOX2 regulated genes in Keratinocyte Overexpression from PMID: 30772301 | SOX2 |
| 7570 | BLOC1S4 | SOX2 regulated genes in Keratinocyte Overexpression from PMID: 30772301 | SOX2 |
| 7571 | VANGL1 | SOX2 regulated genes in Keratinocyte Overexpression from PMID: 30772301 | SOX2 |
| 7572 | SAP30BP | SOX2 regulated genes in Keratinocyte Overexpression from PMID: 30772301 | SOX2 |
| 7573 | CCDC158 | SOX2 regulated genes in Keratinocyte Overexpression from PMID: 30772301 | SOX2 |
| 7574 | CCDC91 | SOX2 regulated genes in Keratinocyte Overexpression from PMID: 30772301 | SOX2 |
| 7575 | SPATA24 | SOX2 regulated genes in Keratinocyte Overexpression from PMID: 30772301 | SOX2 |
| 7576 | C8orf76 | SOX2 regulated genes in Keratinocyte Overexpression from PMID: 30772301 | SOX2 |
| 7577 | ZHX1-C8orf76 | SOX2 regulated genes in Keratinocyte Overexpression from PMID: 30772301 | SOX2 |
| 7578 | MAZ | SOX2 regulated genes in Keratinocyte Overexpression from PMID: 30772301 | SOX2 |
| 7579 | JUND | SOX2 regulated genes in Keratinocyte Overexpression from PMID: 30772301 | SOX2 |
| 7580 | POLR3A | SOX2 regulated genes in Keratinocyte Overexpression from PMID: 30772301 | SOX2 |
| 7581 | FOXH1 | SOX2 regulated genes in Keratinocyte Overexpression from PMID: 30772301 | SOX2 |
| 7582 | KIFC2 | SOX2 regulated genes in Keratinocyte Overexpression from PMID: 30772301 | SOX2 |
| 7583 | RGS16 | SOX2 regulated genes in Keratinocyte Overexpression from PMID: 30772301 | SOX2 |
| 7584 | SPIRE2 | SOX2 regulated genes in Keratinocyte Overexpression from PMID: 30772301 | SOX2 |
| 7585 | CFAP45 | SOX2 regulated genes in Keratinocyte Overexpression from PMID: 30772301 | SOX2 |
| 7586 | RNASEK-C17orf49 | SOX2 regulated genes in Keratinocyte Overexpression from PMID: 30772301 | SOX2 |
| 7587 | C17orf49 | SOX2 regulated genes in Keratinocyte Overexpression from PMID: 30772301 | SOX2 |
| 7588 | BTF3L4 | SOX2 regulated genes in Keratinocyte Overexpression from PMID: 30772301 | SOX2 |
| 7589 | ATG101 | SOX2 regulated genes in Keratinocyte Overexpression from PMID: 30772301 | SOX2 |
| 7590 | SOS1 | SOX2 regulated genes in Keratinocyte Overexpression from PMID: 30772301 | SOX2 |
| 7591 | TMEM101 | SOX2 regulated genes in Keratinocyte Overexpression from PMID: 30772301 | SOX2 |
| 7592 | BAP1 | SOX2 regulated genes in Keratinocyte Overexpression from PMID: 30772301 | SOX2 |
| 7593 | PIK3R2 | SOX2 regulated genes in Keratinocyte Overexpression from PMID: 30772301 | SOX2 |
| 7594 | ZMYM1 | SOX2 regulated genes in Keratinocyte Overexpression from PMID: 30772301 | SOX2 |
| 7595 | B4GALT3 | SOX2 regulated genes in Keratinocyte Overexpression from PMID: 30772301 | SOX2 |
| 7596 | KRR1 | SOX2 regulated genes in Keratinocyte Overexpression from PMID: 30772301 | SOX2 |
| 7597 | KLHDC3 | SOX2 regulated genes in Keratinocyte Overexpression from PMID: 30772301 | SOX2 |
| 7598 | ARHGAP35 | SOX2 regulated genes in Keratinocyte Overexpression from PMID: 30772301 | SOX2 |
| 7599 | CXCR5 | SOX2 regulated genes in Keratinocyte Overexpression from PMID: 30772301 | SOX2 |
| 7600 | NMRAL1 | SOX2 regulated genes in Keratinocyte Overexpression from PMID: 30772301 | SOX2 |
| 7601 | SUPT7L | SOX2 regulated genes in Keratinocyte Overexpression from PMID: 30772301 | SOX2 |
| 7602 | TMEM65 | SOX2 regulated genes in Keratinocyte Overexpression from PMID: 30772301 | SOX2 |
| 7603 | MCM2 | SOX2 regulated genes in Keratinocyte Overexpression from PMID: 30772301 | SOX2 |
| 7604 | HSPE1 | SOX2 regulated genes in Keratinocyte Overexpression from PMID: 30772301 | SOX2 |
| 7605 | MFSD2A | SOX2 regulated genes in Keratinocyte Overexpression from PMID: 30772301 | SOX2 |
| 7606 | HMBS | SOX2 regulated genes in Keratinocyte Overexpression from PMID: 30772301 | SOX2 |
| 7607 | SRPK2 | SOX2 regulated genes in Keratinocyte Overexpression from PMID: 30772301 | SOX2 |
| 7608 | SPC25 | SOX2 regulated genes in Keratinocyte Overexpression from PMID: 30772301 | SOX2 |
| 7609 | MAP4K4 | SOX2 regulated genes in Keratinocyte Overexpression from PMID: 30772301 | SOX2 |
| 7610 | UBA52 | SOX2 regulated genes in Keratinocyte Overexpression from PMID: 30772301 | SOX2 |
| 7611 | TTYH1 | SOX2 regulated genes in Keratinocyte Overexpression from PMID: 30772301 | SOX2 |
| 7612 | MMADHC | SOX2 regulated genes in Keratinocyte Overexpression from PMID: 30772301 | SOX2 |
| 7613 | ZNF598 | SOX2 regulated genes in Keratinocyte Overexpression from PMID: 30772301 | SOX2 |
| 7614 | PHLDB3 | SOX2 regulated genes in Keratinocyte Overexpression from PMID: 30772301 | SOX2 |
| 7615 | FBRSL1 | SOX2 regulated genes in Keratinocyte Overexpression from PMID: 30772301 | SOX2 |
| 7616 | USP7 | SOX2 regulated genes in Keratinocyte Overexpression from PMID: 30772301 | SOX2 |
| 7617 | OGG1 | SOX2 regulated genes in Keratinocyte Overexpression from PMID: 30772301 | SOX2 |
| 7618 | TEX19 | SOX2 regulated genes in Keratinocyte Overexpression from PMID: 30772301 | SOX2 |
| 7619 | SMARCD2 | SOX2 regulated genes in Keratinocyte Overexpression from PMID: 30772301 | SOX2 |
| 7620 | CHD1L | SOX2 regulated genes in Keratinocyte Overexpression from PMID: 30772301 | SOX2 |
| 7621 | MORN2 | SOX2 regulated genes in Keratinocyte Overexpression from PMID: 30772301 | SOX2 |
| 7622 | PWWP2A | SOX2 regulated genes in Keratinocyte Overexpression from PMID: 30772301 | SOX2 |
| 7623 | RPS13 | SOX2 regulated genes in Keratinocyte Overexpression from PMID: 30772301 | SOX2 |

| | | | |
|---|---|---|---|
| 7624 | OPRD1 | SOX2 regulated genes in Keratinocyte Overexpression from PMID: 30772301 | SOX2 |
| 7625 | ZBTB9 | SOX2 regulated genes in Keratinocyte Overexpression from PMID: 30772301 | SOX2 |
| 7626 | DDX54 | SOX2 regulated genes in Keratinocyte Overexpression from PMID: 30772301 | SOX2 |
| 7627 | UNG | SOX2 regulated genes in Keratinocyte Overexpression from PMID: 30772301 | SOX2 |
| 7628 | EIF2D | SOX2 regulated genes in Keratinocyte Overexpression from PMID: 30772301 | SOX2 |
| 7629 | CHMP1A | SOX2 regulated genes in Keratinocyte Overexpression from PMID: 30772301 | SOX2 |
| 7630 | ICAM4 | SOX2 regulated genes in Keratinocyte Overexpression from PMID: 30772301 | SOX2 |
| 7631 | ATRIP | SOX2 regulated genes in Keratinocyte Overexpression from PMID: 30772301 | SOX2 |
| 7632 | ATP2A2 | SOX2 regulated genes in Keratinocyte Overexpression from PMID: 30772301 | SOX2 |
| 7633 | ERBB3 | SOX2 regulated genes in Keratinocyte Overexpression from PMID: 30772301 | SOX2 |
| 7634 | HARS2 | SOX2 regulated genes in Keratinocyte Overexpression from PMID: 30772301 | SOX2 |
| 7635 | TP53BP2 | SOX2 regulated genes in Keratinocyte Overexpression from PMID: 30772301 | SOX2 |
| 7636 | PIGV | SOX2 regulated genes in Keratinocyte Overexpression from PMID: 30772301 | SOX2 |
| 7637 | IGSF8 | SOX2 regulated genes in Keratinocyte Overexpression from PMID: 30772301 | SOX2 |
| 7638 | FIGNL1 | SOX2 regulated genes in Keratinocyte Overexpression from PMID: 30772301 | SOX2 |
| 7639 | CENPL | SOX2 regulated genes in Keratinocyte Overexpression from PMID: 30772301 | SOX2 |
| 7640 | MON1A | SOX2 regulated genes in Keratinocyte Overexpression from PMID: 30772301 | SOX2 |
| 7641 | LRRC56 | SOX2 regulated genes in Keratinocyte Overexpression from PMID: 30772301 | SOX2 |
| 7642 | AGRN | SOX2 regulated genes in Keratinocyte Overexpression from PMID: 30772301 | SOX2 |
| 7643 | UBE2T | SOX2 regulated genes in Keratinocyte Overexpression from PMID: 30772301 | SOX2 |
| 7644 | C16orf96 | SOX2 regulated genes in Keratinocyte Overexpression from PMID: 30772301 | SOX2 |
| 7645 | SDS | SOX2 regulated genes in Keratinocyte Overexpression from PMID: 30772301 | SOX2 |
| 7646 | CRIP3 | SOX2 regulated genes in Keratinocyte Overexpression from PMID: 30772301 | SOX2 |
| 7647 | PRR36 | SOX2 regulated genes in Keratinocyte Overexpression from PMID: 30772301 | SOX2 |
| 7648 | LIN9 | SOX2 regulated genes in Keratinocyte Overexpression from PMID: 30772301 | SOX2 |
| 7649 | CYTH2 | SOX2 regulated genes in Keratinocyte Overexpression from PMID: 30772301 | SOX2 |
| 7650 | NPHP1 | SOX2 regulated genes in Keratinocyte Overexpression from PMID: 30772301 | SOX2 |
| 7651 | ACOT4 | SOX2 regulated genes in Keratinocyte Overexpression from PMID: 30772301 | SOX2 |
| 7652 | ADGRL2 | SOX2 regulated genes in Keratinocyte Overexpression from PMID: 30772301 | SOX2 |
| 7653 | C6orf141 | SOX2 regulated genes in Keratinocyte Overexpression from PMID: 30772301 | SOX2 |
| 7654 | HSPA4 | SOX2 regulated genes in Keratinocyte Overexpression from PMID: 30772301 | SOX2 |
| 7655 | XAB2 | SOX2 regulated genes in Keratinocyte Overexpression from PMID: 30772301 | SOX2 |
| 7656 | TRPM8 | SOX2 regulated genes in Keratinocyte Overexpression from PMID: 30772301 | SOX2 |
| 7657 | NOP16 | SOX2 regulated genes in Keratinocyte Overexpression from PMID: 30772301 | SOX2 |
| 7658 | PMF1-BGLAP | SOX2 regulated genes in Keratinocyte Overexpression from PMID: 30772301 | SOX2 |
| 7659 | PMF1 | SOX2 regulated genes in Keratinocyte Overexpression from PMID: 30772301 | SOX2 |
| 7660 | ZBTB11 | SOX2 regulated genes in Keratinocyte Overexpression from PMID: 30772301 | SOX2 |
| 7661 | TMEM120A | SOX2 regulated genes in Keratinocyte Overexpression from PMID: 30772301 | SOX2 |
| 7662 | CNPPD1 | SOX2 regulated genes in Keratinocyte Overexpression from PMID: 30772301 | SOX2 |
| 7663 | NEURL4 | SOX2 regulated genes in Keratinocyte Overexpression from PMID: 30772301 | SOX2 |
| 7664 | POLR3K | SOX2 regulated genes in Keratinocyte Overexpression from PMID: 30772301 | SOX2 |
| 7665 | DVL3 | SOX2 regulated genes in Keratinocyte Overexpression from PMID: 30772301 | SOX2 |
| 7666 | RAVER1 | SOX2 regulated genes in Keratinocyte Overexpression from PMID: 30772301 | SOX2 |
| 7667 | CHEK1 | SOX2 regulated genes in Keratinocyte Overexpression from PMID: 30772301 | SOX2 |
| 7668 | TMEM238 | SOX2 regulated genes in Keratinocyte Overexpression from PMID: 30772301 | SOX2 |
| 7669 | UCHL5 | SOX2 regulated genes in Keratinocyte Overexpression from PMID: 30772301 | SOX2 |
| 7670 | CHD1 | SOX2 regulated genes in Keratinocyte Overexpression from PMID: 30772301 | SOX2 |
| 7671 | SLC25A3 | SOX2 regulated genes in Keratinocyte Overexpression from PMID: 30772301 | SOX2 |
| 7672 | ZNF32 | SOX2 regulated genes in Keratinocyte Overexpression from PMID: 30772301 | SOX2 |
| 7673 | SENP8 | SOX2 regulated genes in Keratinocyte Overexpression from PMID: 30772301 | SOX2 |
| 7674 | METTL5 | SOX2 regulated genes in Keratinocyte Overexpression from PMID: 30772301 | SOX2 |
| 7675 | ZNF566 | SOX2 regulated genes in Keratinocyte Overexpression from PMID: 30772301 | SOX2 |
| 7676 | HAPLN3 | SOX2 regulated genes in Keratinocyte Overexpression from PMID: 30772301 | SOX2 |
| 7677 | SMC3 | SOX2 regulated genes in Keratinocyte Overexpression from PMID: 30772301 | SOX2 |
| 7678 | GADD45B | SOX2 regulated genes in Keratinocyte Overexpression from PMID: 30772301 | SOX2 |
| 7679 | MAP6 | SOX2 regulated genes in Keratinocyte Overexpression from PMID: 30772301 | SOX2 |
| 7680 | ANKRD52 | SOX2 regulated genes in Keratinocyte Overexpression from PMID: 30772301 | SOX2 |
| 7681 | PALMD | SOX2 regulated genes in Keratinocyte Overexpression from PMID: 30772301 | SOX2 |
| 7682 | EGR2 | SOX2 regulated genes in Keratinocyte Overexpression from PMID: 30772301 | SOX2 |
| 7683 | PRR11 | SOX2 regulated genes in Keratinocyte Overexpression from PMID: 30772301 | SOX2 |
| 7684 | CWC25 | SOX2 regulated genes in Keratinocyte Overexpression from PMID: 30772301 | SOX2 |

| | | | |
|---|---|---|---|
| 7685 | HDAC3 | SOX2 regulated genes in Keratinocyte Overexpression from PMID: 30772301 | SOX2 |
| 7686 | DHX37 | SOX2 regulated genes in Keratinocyte Overexpression from PMID: 30772301 | SOX2 |
| 7687 | NUDCD1 | SOX2 regulated genes in Keratinocyte Overexpression from PMID: 30772301 | SOX2 |
| 7688 | LIPH | SOX2 regulated genes in Keratinocyte Overexpression from PMID: 30772301 | SOX2 |
| 7689 | KIF21B | SOX2 regulated genes in Keratinocyte Overexpression from PMID: 30772301 | SOX2 |
| 7690 | DHPS | SOX2 regulated genes in Keratinocyte Overexpression from PMID: 30772301 | SOX2 |
| 7691 | TRNP1 | SOX2 regulated genes in Keratinocyte Overexpression from PMID: 30772301 | SOX2 |
| 7692 | C2CD2L | SOX2 regulated genes in Keratinocyte Overexpression from PMID: 30772301 | SOX2 |
| 7693 | AMBRA1 | SOX2 regulated genes in Keratinocyte Overexpression from PMID: 30772301 | SOX2 |
| 7694 | LCLAT1 | SOX2 regulated genes in Keratinocyte Overexpression from PMID: 30772301 | SOX2 |
| 7695 | MRPL28 | SOX2 regulated genes in Keratinocyte Overexpression from PMID: 30772301 | SOX2 |
| 7696 | ANGPTL4 | SOX2 regulated genes in Keratinocyte Overexpression from PMID: 30772301 | SOX2 |
| 7697 | PLEKHA1 | SOX2 regulated genes in Keratinocyte Overexpression from PMID: 30772301 | SOX2 |
| 7698 | UBXN1 | SOX2 regulated genes in Keratinocyte Overexpression from PMID: 30772301 | SOX2 |
| 7699 | SDHB | SOX2 regulated genes in Keratinocyte Overexpression from PMID: 30772301 | SOX2 |
| 7700 | GALE | SOX2 regulated genes in Keratinocyte Overexpression from PMID: 30772301 | SOX2 |
| 7701 | ORAI2 | SOX2 regulated genes in Keratinocyte Overexpression from PMID: 30772301 | SOX2 |
| 7702 | AMMECR1L | SOX2 regulated genes in Keratinocyte Overexpression from PMID: 30772301 | SOX2 |
| 7703 | FIGLA | SOX2 regulated genes in Keratinocyte Overexpression from PMID: 30772301 | SOX2 |
| 7704 | BTBD19 | SOX2 regulated genes in Keratinocyte Overexpression from PMID: 30772301 | SOX2 |
| 7705 | ZRANB2 | SOX2 regulated genes in Keratinocyte Overexpression from PMID: 30772301 | SOX2 |
| 7706 | PTCHD4 | SOX2 regulated genes in Keratinocyte Overexpression from PMID: 30772301 | SOX2 |
| 7707 | PRPF40B | SOX2 regulated genes in Keratinocyte Overexpression from PMID: 30772301 | SOX2 |
| 7708 | RTF1 | SOX2 regulated genes in Keratinocyte Overexpression from PMID: 30772301 | SOX2 |
| 7709 | GPAT4 | SOX2 regulated genes in Keratinocyte Overexpression from PMID: 30772301 | SOX2 |
| 7710 | SMC4 | SOX2 regulated genes in Keratinocyte Overexpression from PMID: 30772301 | SOX2 |
| 7711 | SPTY2D1 | SOX2 regulated genes in Keratinocyte Overexpression from PMID: 30772301 | SOX2 |
| 7712 | GATA3 | SOX2 regulated genes in Keratinocyte Overexpression from PMID: 30772301 | SOX2 |
| 7713 | ZNF296 | SOX2 regulated genes in Keratinocyte Overexpression from PMID: 30772301 | SOX2 |
| 7714 | KIF9 | SOX2 regulated genes in Keratinocyte Overexpression from PMID: 30772301 | SOX2 |
| 7715 | AZIN2 | SOX2 regulated genes in Keratinocyte Overexpression from PMID: 30772301 | SOX2 |
| 7716 | INTS5 | SOX2 regulated genes in Keratinocyte Overexpression from PMID: 30772301 | SOX2 |
| 7717 | SULT2B1 | SOX2 regulated genes in Keratinocyte Overexpression from PMID: 30772301 | SOX2 |
| 7718 | ALOX15B | SOX2 regulated genes in Keratinocyte Overexpression from PMID: 30772301 | SOX2 |
| 7719 | TIMM17A | SOX2 regulated genes in Keratinocyte Overexpression from PMID: 30772301 | SOX2 |
| 7720 | CCNL1 | SOX2 regulated genes in Keratinocyte Overexpression from PMID: 30772301 | SOX2 |
| 7721 | KLHL29 | SOX2 regulated genes in Keratinocyte Overexpression from PMID: 30772301 | SOX2 |
| 7722 | RAD54L2 | SOX2 regulated genes in Keratinocyte Overexpression from PMID: 30772301 | SOX2 |
| 7723 | LLGL2 | SOX2 regulated genes in Keratinocyte Overexpression from PMID: 30772301 | SOX2 |
| 7724 | CYBRD1 | SOX2 regulated genes in Keratinocyte Overexpression from PMID: 30772301 | SOX2 |
| 7725 | DPY30 | SOX2 regulated genes in Keratinocyte Overexpression from PMID: 30772301 | SOX2 |
| 7726 | C2orf69 | SOX2 regulated genes in Keratinocyte Overexpression from PMID: 30772301 | SOX2 |
| 7727 | CEP83 | SOX2 regulated genes in Keratinocyte Overexpression from PMID: 30772301 | SOX2 |
| 7728 | NCAPD3 | SOX2 regulated genes in Keratinocyte Overexpression from PMID: 30772301 | SOX2 |
| 7729 | RNF220 | SOX2 regulated genes in Keratinocyte Overexpression from PMID: 30772301 | SOX2 |
| 7730 | USP42 | SOX2 regulated genes in Keratinocyte Overexpression from PMID: 30772301 | SOX2 |
| 7731 | RFC4 | SOX2 regulated genes in Keratinocyte Overexpression from PMID: 30772301 | SOX2 |
| 7732 | KCTD15 | SOX2 regulated genes in Keratinocyte Overexpression from PMID: 30772301 | SOX2 |
| 7733 | ZNF239 | SOX2 regulated genes in Keratinocyte Overexpression from PMID: 30772301 | SOX2 |
| 7734 | RSRC2 | SOX2 regulated genes in Keratinocyte Overexpression from PMID: 30772301 | SOX2 |
| 7735 | RTKN | SOX2 regulated genes in Keratinocyte Overexpression from PMID: 30772301 | SOX2 |
| 7736 | ZNF780A | SOX2 regulated genes in Keratinocyte Overexpression from PMID: 30772301 | SOX2 |
| 7737 | NVL | SOX2 regulated genes in Keratinocyte Overexpression from PMID: 30772301 | SOX2 |
| 7738 | TUFT1 | SOX2 regulated genes in Keratinocyte Overexpression from PMID: 30772301 | SOX2 |
| 7739 | SRP19 | SOX2 regulated genes in Keratinocyte Overexpression from PMID: 30772301 | SOX2 |
| 7740 | YARS2 | SOX2 regulated genes in Keratinocyte Overexpression from PMID: 30772301 | SOX2 |
| 7741 | TDRD5 | SOX2 regulated genes in Keratinocyte Overexpression from PMID: 30772301 | SOX2 |
| 7742 | RPS25 | SOX2 regulated genes in Keratinocyte Overexpression from PMID: 30772301 | SOX2 |
| 7743 | ECT2 | SOX2 regulated genes in Keratinocyte Overexpression from PMID: 30772301 | SOX2 |
| 7744 | MARK4 | SOX2 regulated genes in Keratinocyte Overexpression from PMID: 30772301 | SOX2 |
| 7745 | GRM2 | SOX2 regulated genes in Keratinocyte Overexpression from PMID: 30772301 | SOX2 |

| | | | |
|---|---|---|---|
| 7746 | ZNF180 | SOX2 regulated genes in Keratinocyte Overexpression from PMID: 30772301 | SOX2 |
| 7747 | ACSM3 | SOX2 regulated genes in Keratinocyte Overexpression from PMID: 30772301 | SOX2 |
| 7748 | PURA | SOX2 regulated genes in Keratinocyte Overexpression from PMID: 30772301 | SOX2 |
| 7749 | SCAF1 | SOX2 regulated genes in Keratinocyte Overexpression from PMID: 30772301 | SOX2 |
| 7750 | ZBTB49 | SOX2 regulated genes in Keratinocyte Overexpression from PMID: 30772301 | SOX2 |
| 7751 | FBXL14 | SOX2 regulated genes in Keratinocyte Overexpression from PMID: 30772301 | SOX2 |
| 7752 | DDX41 | SOX2 regulated genes in Keratinocyte Overexpression from PMID: 30772301 | SOX2 |
| 7753 | SLC30A6 | SOX2 regulated genes in Keratinocyte Overexpression from PMID: 30772301 | SOX2 |
| 7754 | MRPL37 | SOX2 regulated genes in Keratinocyte Overexpression from PMID: 30772301 | SOX2 |
| 7755 | OTUD7B | SOX2 regulated genes in Keratinocyte Overexpression from PMID: 30772301 | SOX2 |
| 7756 | UBE2J2 | SOX2 regulated genes in Keratinocyte Overexpression from PMID: 30772301 | SOX2 |
| 7757 | HMGCS2 | SOX2 regulated genes in Keratinocyte Overexpression from PMID: 30772301 | SOX2 |
| 7758 | SLC23A1 | SOX2 regulated genes in Keratinocyte Overexpression from PMID: 30772301 | SOX2 |
| 7759 | STIL | SOX2 regulated genes in Keratinocyte Overexpression from PMID: 30772301 | SOX2 |
| 7760 | EIF3E | SOX2 regulated genes in Keratinocyte Overexpression from PMID: 30772301 | SOX2 |
| 7761 | COPS7B | SOX2 regulated genes in Keratinocyte Overexpression from PMID: 30772301 | SOX2 |
| 7762 | DPM3 | SOX2 regulated genes in Keratinocyte Overexpression from PMID: 30772301 | SOX2 |
| 7763 | SLC6A15 | SOX2 regulated genes in Keratinocyte Overexpression from PMID: 30772301 | SOX2 |
| 7764 | ENC1 | SOX2 regulated genes in Keratinocyte Overexpression from PMID: 30772301 | SOX2 |
| 7765 | TIMM44 | SOX2 regulated genes in Keratinocyte Overexpression from PMID: 30772301 | SOX2 |
| 7766 | CCDC106 | SOX2 regulated genes in Keratinocyte Overexpression from PMID: 30772301 | SOX2 |
| 7767 | ZNF131 | SOX2 regulated genes in Keratinocyte Overexpression from PMID: 30772301 | SOX2 |
| 7768 | PTP4A1 | SOX2 regulated genes in Keratinocyte Overexpression from PMID: 30772301 | SOX2 |
| 7769 | UBIAD1 | SOX2 regulated genes in Keratinocyte Overexpression from PMID: 30772301 | SOX2 |
| 7770 | SLC7A14 | SOX2 regulated genes in Keratinocyte Overexpression from PMID: 30772301 | SOX2 |
| 7771 | MRPL12 | SOX2 regulated genes in Keratinocyte Overexpression from PMID: 30772301 | SOX2 |
| 7772 | NUDT1 | SOX2 regulated genes in Keratinocyte Overexpression from PMID: 30772301 | SOX2 |
| 7773 | TMEM138 | SOX2 regulated genes in Keratinocyte Overexpression from PMID: 30772301 | SOX2 |
| 7774 | PBLD | SOX2 regulated genes in Keratinocyte Overexpression from PMID: 30772301 | SOX2 |
| 7775 | PRR14 | SOX2 regulated genes in Keratinocyte Overexpression from PMID: 30772301 | SOX2 |
| 7776 | GGPS1 | SOX2 regulated genes in Keratinocyte Overexpression from PMID: 30772301 | SOX2 |
| 7777 | SLC45A1 | SOX2 regulated genes in Keratinocyte Overexpression from PMID: 30772301 | SOX2 |
| 7778 | PTHLH | SOX2 regulated genes in Keratinocyte Overexpression from PMID: 30772301 | SOX2 |
| 7779 | FAM83E | SOX2 regulated genes in Keratinocyte Overexpression from PMID: 30772301 | SOX2 |
| 7780 | KCNJ13 | SOX2 regulated genes in Keratinocyte Overexpression from PMID: 30772301 | SOX2 |
| 7781 | COA7 | SOX2 regulated genes in Keratinocyte Overexpression from PMID: 30772301 | SOX2 |
| 7782 | TOMM7 | SOX2 regulated genes in Keratinocyte Overexpression from PMID: 30772301 | SOX2 |
| 7783 | KRAS | SOX2 regulated genes in Keratinocyte Overexpression from PMID: 30772301 | SOX2 |
| 7784 | HS2ST1 | SOX2 regulated genes in Keratinocyte Overexpression from PMID: 30772301 | SOX2 |
| 7785 | NEK8 | SOX2 regulated genes in Keratinocyte Overexpression from PMID: 30772301 | SOX2 |
| 7786 | ZNF23 | SOX2 regulated genes in Keratinocyte Overexpression from PMID: 30772301 | SOX2 |
| 7787 | EPOP | SOX2 regulated genes in Keratinocyte Overexpression from PMID: 30772301 | SOX2 |
| 7788 | NT5E | SOX2 regulated genes in Keratinocyte Overexpression from PMID: 30772301 | SOX2 |
| 7789 | S100A1 | SOX2 regulated genes in Keratinocyte Overexpression from PMID: 30772301 | SOX2 |
| 7790 | FUT1 | SOX2 regulated genes in Keratinocyte Overexpression from PMID: 30772301 | SOX2 |
| 7791 | POLR3D | SOX2 regulated genes in Keratinocyte Overexpression from PMID: 30772301 | SOX2 |
| 7792 | EGFR | SOX2 regulated genes in Keratinocyte Overexpression from PMID: 30772301 | SOX2 |
| 7793 | ZNF446 | SOX2 regulated genes in Keratinocyte Overexpression from PMID: 30772301 | SOX2 |
| 7794 | PIH1D2 | SOX2 regulated genes in Keratinocyte Overexpression from PMID: 30772301 | SOX2 |
| 7795 | CDK5RAP2 | SOX2 regulated genes in Keratinocyte Overexpression from PMID: 30772301 | SOX2 |
| 7796 | VPS37C | SOX2 regulated genes in Keratinocyte Overexpression from PMID: 30772301 | SOX2 |
| 7797 | GALK1 | SOX2 regulated genes in Keratinocyte Overexpression from PMID: 30772301 | SOX2 |
| 7798 | G3BP1 | SOX2 regulated genes in Keratinocyte Overexpression from PMID: 30772301 | SOX2 |
| 7799 | BBS2 | SOX2 regulated genes in Keratinocyte Overexpression from PMID: 30772301 | SOX2 |
| 7800 | OVGP1 | SOX2 regulated genes in Keratinocyte Overexpression from PMID: 30772301 | SOX2 |
| 7801 | FBXO33 | SOX2 regulated genes in Keratinocyte Overexpression from PMID: 30772301 | SOX2 |
| 7802 | EPN3 | SOX2 regulated genes in Keratinocyte Overexpression from PMID: 30772301 | SOX2 |
| 7803 | FOXO6 | SOX2 regulated genes in Keratinocyte Overexpression from PMID: 30772301 | SOX2 |
| 7804 | ZDHHC1 | SOX2 regulated genes in Keratinocyte Overexpression from PMID: 30772301 | SOX2 |
| 7805 | PHOSPHO2 | SOX2 regulated genes in Keratinocyte Overexpression from PMID: 30772301 | SOX2 |
| 7806 | LRPPRC | SOX2 regulated genes in Keratinocyte Overexpression from PMID: 30772301 | SOX2 |

| | | | |
|---|---|---|---|
| 7807 | RNF2 | SOX2 regulated genes in Keratinocyte Overexpression from PMID: 30772301 | SOX2 |
| 7808 | ALDH3B2 | SOX2 regulated genes in Keratinocyte Overexpression from PMID: 30772301 | SOX2 |
| 7809 | BACE1 | SOX2 regulated genes in Keratinocyte Overexpression from PMID: 30772301 | SOX2 |
| 7810 | ABCB6 | SOX2 regulated genes in Keratinocyte Overexpression from PMID: 30772301 | SOX2 |
| 7811 | IMP3 | SOX2 regulated genes in Keratinocyte Overexpression from PMID: 30772301 | SOX2 |
| 7812 | PSMD2 | SOX2 regulated genes in Keratinocyte Overexpression from PMID: 30772301 | SOX2 |
| 7813 | KCNJ16 | SOX2 regulated genes in Keratinocyte Overexpression from PMID: 30772301 | SOX2 |
| 7814 | LYG1 | SOX2 regulated genes in Keratinocyte Overexpression from PMID: 30772301 | SOX2 |
| 7815 | ZNF251 | SOX2 regulated genes in Keratinocyte Overexpression from PMID: 30772301 | SOX2 |
| 7816 | BANP | SOX2 regulated genes in Keratinocyte Overexpression from PMID: 30772301 | SOX2 |
| 7817 | TAMM41 | SOX2 regulated genes in Keratinocyte Overexpression from PMID: 30772301 | SOX2 |
| 7818 | MIER2 | SOX2 regulated genes in Keratinocyte Overexpression from PMID: 30772301 | SOX2 |
| 7819 | MAB21L3 | SOX2 regulated genes in Keratinocyte Overexpression from PMID: 30772301 | SOX2 |
| 7820 | DAXX | SOX2 regulated genes in Keratinocyte Overexpression from PMID: 30772301 | SOX2 |
| 7821 | GIPR | SOX2 regulated genes in Keratinocyte Overexpression from PMID: 30772301 | SOX2 |
| 7822 | HOXD4 | SOX2 regulated genes in Keratinocyte Overexpression from PMID: 30772301 | SOX2 |
| 7823 | DDX10 | SOX2 regulated genes in Keratinocyte Overexpression from PMID: 30772301 | SOX2 |
| 7824 | NSD1 | SOX2 regulated genes in Keratinocyte Overexpression from PMID: 30772301 | SOX2 |
| 7825 | ABHD14A-ACY1 | SOX2 regulated genes in Keratinocyte Overexpression from PMID: 30772301 | SOX2 |
| 7826 | ACY1 | SOX2 regulated genes in Keratinocyte Overexpression from PMID: 30772301 | SOX2 |
| 7827 | RAD9B | SOX2 regulated genes in Keratinocyte Overexpression from PMID: 30772301 | SOX2 |
| 7828 | CCDC71L | SOX2 regulated genes in Keratinocyte Overexpression from PMID: 30772301 | SOX2 |
| 7829 | MFN1 | SOX2 regulated genes in Keratinocyte Overexpression from PMID: 30772301 | SOX2 |
| 7830 | SLC16A11 | SOX2 regulated genes in Keratinocyte Overexpression from PMID: 30772301 | SOX2 |
| 7831 | CENPE | SOX2 regulated genes in Keratinocyte Overexpression from PMID: 30772301 | SOX2 |
| 7832 | PSMD6 | SOX2 regulated genes in Keratinocyte Overexpression from PMID: 30772301 | SOX2 |
| 7833 | RHBDF2 | SOX2 regulated genes in Keratinocyte Overexpression from PMID: 30772301 | SOX2 |
| 7834 | HMG20B | SOX2 regulated genes in Keratinocyte Overexpression from PMID: 30772301 | SOX2 |
| 7835 | ZSWIM4 | SOX2 regulated genes in Keratinocyte Overexpression from PMID: 30772301 | SOX2 |
| 7836 | IFT122 | SOX2 regulated genes in Keratinocyte Overexpression from PMID: 30772301 | SOX2 |
| 7837 | NOS1AP | SOX2 regulated genes in Keratinocyte Overexpression from PMID: 30772301 | SOX2 |
| 7838 | ANAPC13 | SOX2 regulated genes in Keratinocyte Overexpression from PMID: 30772301 | SOX2 |
| 7839 | CDC23 | SOX2 regulated genes in Keratinocyte Overexpression from PMID: 30772301 | SOX2 |
| 7840 | CLCN2 | SOX2 regulated genes in Keratinocyte Overexpression from PMID: 30772301 | SOX2 |
| 7841 | ZNF445 | SOX2 regulated genes in Keratinocyte Overexpression from PMID: 30772301 | SOX2 |
| 7842 | CDC73 | SOX2 regulated genes in Keratinocyte Overexpression from PMID: 30772301 | SOX2 |
| 7843 | ASAP3 | SOX2 regulated genes in Keratinocyte Overexpression from PMID: 30772301 | SOX2 |
| 7844 | RFNG | SOX2 regulated genes in Keratinocyte Overexpression from PMID: 30772301 | SOX2 |
| 7845 | LEMD3 | SOX2 regulated genes in Keratinocyte Overexpression from PMID: 30772301 | SOX2 |
| 7846 | MAPK6 | SOX2 regulated genes in Keratinocyte Overexpression from PMID: 30772301 | SOX2 |
| 7847 | ZNF628 | SOX2 regulated genes in Keratinocyte Overexpression from PMID: 30772301 | SOX2 |
| 7848 | IMMT | SOX2 regulated genes in Keratinocyte Overexpression from PMID: 30772301 | SOX2 |
| 7849 | AKT1S1 | SOX2 regulated genes in Keratinocyte Overexpression from PMID: 30772301 | SOX2 |
| 7850 | RPAP1 | SOX2 regulated genes in Keratinocyte Overexpression from PMID: 30772301 | SOX2 |
| 7851 | TAF12 | SOX2 regulated genes in Keratinocyte Overexpression from PMID: 30772301 | SOX2 |
| 7852 | MFSD3 | SOX2 regulated genes in Keratinocyte Overexpression from PMID: 30772301 | SOX2 |
| 7853 | KIAA0895L | SOX2 regulated genes in Keratinocyte Overexpression from PMID: 30772301 | SOX2 |
| 7854 | ERCC8 | SOX2 regulated genes in Keratinocyte Overexpression from PMID: 30772301 | SOX2 |
| 7855 | TRIM62 | SOX2 regulated genes in Keratinocyte Overexpression from PMID: 30772301 | SOX2 |
| 7856 | YAF2 | SOX2 regulated genes in Keratinocyte Overexpression from PMID: 30772301 | SOX2 |
| 7857 | GTF2H5 | SOX2 regulated genes in Keratinocyte Overexpression from PMID: 30772301 | SOX2 |
| 7858 | NDUFA7 | SOX2 regulated genes in Keratinocyte Overexpression from PMID: 30772301 | SOX2 |
| 7859 | MSI2 | SOX2 regulated genes in Keratinocyte Overexpression from PMID: 30772301 | SOX2 |
| 7860 | ABHD17A | SOX2 regulated genes in Keratinocyte Overexpression from PMID: 30772301 | SOX2 |
| 7861 | PLEKHJ1 | SOX2 regulated genes in Keratinocyte Overexpression from PMID: 30772301 | SOX2 |
| 7862 | TAF15 | SOX2 regulated genes in Keratinocyte Overexpression from PMID: 30772301 | SOX2 |
| 7863 | NMNAT3 | SOX2 regulated genes in Keratinocyte Overexpression from PMID: 30772301 | SOX2 |
| 7864 | ZNF575 | SOX2 regulated genes in Keratinocyte Overexpression from PMID: 30772301 | SOX2 |
| 7865 | UBXN8 | SOX2 regulated genes in Keratinocyte Overexpression from PMID: 30772301 | SOX2 |
| 7866 | GZMM | SOX2 regulated genes in Keratinocyte Overexpression from PMID: 30772301 | SOX2 |
| 7867 | ASPM | SOX2 regulated genes in Keratinocyte Overexpression from PMID: 30772301 | SOX2 |

| | | | |
|---|---|---|---|
| 7868 | ZNF700 | SOX2 regulated genes in Keratinocyte Overexpression from PMID: 30772301 | SOX2 |
| 7869 | ZNF878 | SOX2 regulated genes in Keratinocyte Overexpression from PMID: 30772301 | SOX2 |
| 7870 | WDR35 | SOX2 regulated genes in Keratinocyte Overexpression from PMID: 30772301 | SOX2 |
| 7871 | NME5 | SOX2 regulated genes in Keratinocyte Overexpression from PMID: 30772301 | SOX2 |
| 7872 | STXBP2 | SOX2 regulated genes in Keratinocyte Overexpression from PMID: 30772301 | SOX2 |
| 7873 | SZT2 | SOX2 regulated genes in Keratinocyte Overexpression from PMID: 30772301 | SOX2 |
| 7874 | MRFAP1L1 | SOX2 regulated genes in Keratinocyte Overexpression from PMID: 30772301 | SOX2 |
| 7875 | MRFAP1 | SOX2 regulated genes in Keratinocyte Overexpression from PMID: 30772301 | SOX2 |
| 7876 | CACTIN | SOX2 regulated genes in Keratinocyte Overexpression from PMID: 30772301 | SOX2 |
| 7877 | HPCA | SOX2 regulated genes in Keratinocyte Overexpression from PMID: 30772301 | SOX2 |
| 7878 | PGLS | SOX2 regulated genes in Keratinocyte Overexpression from PMID: 30772301 | SOX2 |
| 7879 | NAPA | SOX2 regulated genes in Keratinocyte Overexpression from PMID: 30772301 | SOX2 |
| 7880 | PRIM1 | SOX2 regulated genes in Keratinocyte Overexpression from PMID: 30772301 | SOX2 |
| 7881 | SLC20A1 | SOX2 regulated genes in Keratinocyte Overexpression from PMID: 30772301 | SOX2 |
| 7882 | BBS5 | SOX2 regulated genes in Keratinocyte Overexpression from PMID: 30772301 | SOX2 |
| 7883 | GPA33 | SOX2 regulated genes in Keratinocyte Overexpression from PMID: 30772301 | SOX2 |
| 7884 | NOTUM | SOX2 regulated genes in Keratinocyte Overexpression from PMID: 30772301 | SOX2 |
| 7885 | RPS9 | SOX2 regulated genes in Keratinocyte Overexpression from PMID: 30772301 | SOX2 |
| 7886 | PCYT2 | SOX2 regulated genes in Keratinocyte Overexpression from PMID: 30772301 | SOX2 |
| 7887 | ASCL4 | SOX2 regulated genes in Keratinocyte Overexpression from PMID: 30772301 | SOX2 |
| 7888 | DNAH9 | SOX2 regulated genes in Keratinocyte Overexpression from PMID: 30772301 | SOX2 |
| 7889 | CIB1 | SOX2 regulated genes in Keratinocyte Overexpression from PMID: 30772301 | SOX2 |
| 7890 | CAMSAP2 | SOX2 regulated genes in Keratinocyte Overexpression from PMID: 30772301 | SOX2 |
| 7891 | ATG4D | SOX2 regulated genes in Keratinocyte Overexpression from PMID: 30772301 | SOX2 |
| 7892 | ABCB10 | SOX2 regulated genes in Keratinocyte Overexpression from PMID: 30772301 | SOX2 |
| 7893 | ABTB1 | SOX2 regulated genes in Keratinocyte Overexpression from PMID: 30772301 | SOX2 |
| 7894 | SLC12A8 | SOX2 regulated genes in Keratinocyte Overexpression from PMID: 30772301 | SOX2 |
| 7895 | MAT2A | SOX2 regulated genes in Keratinocyte Overexpression from PMID: 30772301 | SOX2 |
| 7896 | CYP4F11 | SOX2 regulated genes in Keratinocyte Overexpression from PMID: 30772301 | SOX2 |
| 7897 | CYP4A11 | SOX2 regulated genes in Keratinocyte Overexpression from PMID: 30772301 | SOX2 |
| 7898 | GNB1 | SOX2 regulated genes in Keratinocyte Overexpression from PMID: 30772301 | SOX2 |
| 7899 | IER2 | SOX2 regulated genes in Keratinocyte Overexpression from PMID: 30772301 | SOX2 |
| 7900 | RBAK | SOX2 regulated genes in Keratinocyte Overexpression from PMID: 30772301 | SOX2 |
| 7901 | PODXL2 | SOX2 regulated genes in Keratinocyte Overexpression from PMID: 30772301 | SOX2 |
| 7902 | MARVELD2 | SOX2 regulated genes in Keratinocyte Overexpression from PMID: 30772301 | SOX2 |
| 7903 | POLD1 | SOX2 regulated genes in Keratinocyte Overexpression from PMID: 30772301 | SOX2 |
| 7904 | RPL28 | SOX2 regulated genes in Keratinocyte Overexpression from PMID: 30772301 | SOX2 |
| 7905 | BMPR2 | SOX2 regulated genes in Keratinocyte Overexpression from PMID: 30772301 | SOX2 |
| 7906 | LARP7 | SOX2 regulated genes in Keratinocyte Overexpression from PMID: 30772301 | SOX2 |
| 7907 | VIL1 | SOX2 regulated genes in Keratinocyte Overexpression from PMID: 30772301 | SOX2 |
| 7908 | TIMM29 | SOX2 regulated genes in Keratinocyte Overexpression from PMID: 30772301 | SOX2 |
| 7909 | PARPBP | SOX2 regulated genes in Keratinocyte Overexpression from PMID: 30772301 | SOX2 |
| 7910 | TMC6 | SOX2 regulated genes in Keratinocyte Overexpression from PMID: 30772301 | SOX2 |
| 7911 | FAM222B | SOX2 regulated genes in Keratinocyte Overexpression from PMID: 30772301 | SOX2 |
| 7912 | CUL1 | SOX2 regulated genes in Keratinocyte Overexpression from PMID: 30772301 | SOX2 |
| 7913 | GOLGA7 | SOX2 regulated genes in Keratinocyte Overexpression from PMID: 30772301 | SOX2 |
| 7914 | SRSF11 | SOX2 regulated genes in Keratinocyte Overexpression from PMID: 30772301 | SOX2 |
| 7915 | INO80B | SOX2 regulated genes in Keratinocyte Overexpression from PMID: 30772301 | SOX2 |
| 7916 | ZNF574 | SOX2 regulated genes in Keratinocyte Overexpression from PMID: 30772301 | SOX2 |
| 7917 | C19orf25 | SOX2 regulated genes in Keratinocyte Overexpression from PMID: 30772301 | SOX2 |
| 7918 | RASSF1 | SOX2 regulated genes in Keratinocyte Overexpression from PMID: 30772301 | SOX2 |
| 7919 | CIT | SOX2 regulated genes in Keratinocyte Overexpression from PMID: 30772301 | SOX2 |
| 7920 | CDC42EP3 | SOX2 regulated genes in Keratinocyte Overexpression from PMID: 30772301 | SOX2 |
| 7921 | DPYSL4 | SOX2 regulated genes in Keratinocyte Overexpression from PMID: 30772301 | SOX2 |
| 7922 | CCDC73 | SOX2 regulated genes in Keratinocyte Overexpression from PMID: 30772301 | SOX2 |
| 7923 | PRRT3 | SOX2 regulated genes in Keratinocyte Overexpression from PMID: 30772301 | SOX2 |
| 7924 | SPTBN2 | SOX2 regulated genes in Keratinocyte Overexpression from PMID: 30772301 | SOX2 |
| 7925 | RYK | SOX2 regulated genes in Keratinocyte Overexpression from PMID: 30772301 | SOX2 |
| 7926 | SPINT2 | SOX2 regulated genes in Keratinocyte Overexpression from PMID: 30772301 | SOX2 |
| 7927 | IFT20 | SOX2 regulated genes in Keratinocyte Overexpression from PMID: 30772301 | SOX2 |
| 7928 | PTRHD1 | SOX2 regulated genes in Keratinocyte Overexpression from PMID: 30772301 | SOX2 |

| | | | |
|---|---|---|---|
| 7929 | CCDC43 | SOX2 regulated genes in Keratinocyte Overexpression from PMID: 30772301 | SOX2 |
| 7930 | DCTN2 | SOX2 regulated genes in Keratinocyte Overexpression from PMID: 30772301 | SOX2 |
| 7931 | LRRC46 | SOX2 regulated genes in Keratinocyte Overexpression from PMID: 30772301 | SOX2 |
| 7932 | TAF1A | SOX2 regulated genes in Keratinocyte Overexpression from PMID: 30772301 | SOX2 |
| 7933 | AAMP | SOX2 regulated genes in Keratinocyte Overexpression from PMID: 30772301 | SOX2 |
| 7934 | ACVR1B | SOX2 regulated genes in Keratinocyte Overexpression from PMID: 30772301 | SOX2 |
| 7935 | CCDC74B | SOX2 regulated genes in Keratinocyte Overexpression from PMID: 30772301 | SOX2 |
| 7936 | CCDC74A | SOX2 regulated genes in Keratinocyte Overexpression from PMID: 30772301 | SOX2 |
| 7937 | SFTPC | SOX2 regulated genes in Keratinocyte Overexpression from PMID: 30772301 | SOX2 |
| 7938 | AHSA1 | SOX2 regulated genes in Keratinocyte Overexpression from PMID: 30772301 | SOX2 |
| 7939 | RNF145 | SOX2 regulated genes in Keratinocyte Overexpression from PMID: 30772301 | SOX2 |
| 7940 | CXCL12 | SOX2 regulated genes in Keratinocyte Overexpression from PMID: 30772301 | SOX2 |
| 7941 | SNCAIP | SOX2 regulated genes in Keratinocyte Overexpression from PMID: 30772301 | SOX2 |
| 7942 | TADA1 | SOX2 regulated genes in Keratinocyte Overexpression from PMID: 30772301 | SOX2 |
| 7943 | IGSF3 | SOX2 regulated genes in Keratinocyte Overexpression from PMID: 30772301 | SOX2 |
| 7944 | CENPO | SOX2 regulated genes in Keratinocyte Overexpression from PMID: 30772301 | SOX2 |
| 7945 | RBM12B | SOX2 regulated genes in Keratinocyte Overexpression from PMID: 30772301 | SOX2 |
| 7946 | APPBP2 | SOX2 regulated genes in Keratinocyte Overexpression from PMID: 30772301 | SOX2 |
| 7947 | TMEM186 | SOX2 regulated genes in Keratinocyte Overexpression from PMID: 30772301 | SOX2 |
| 7948 | EZR | SOX2 regulated genes in Keratinocyte Overexpression from PMID: 30772301 | SOX2 |
| 7949 | TTC21B | SOX2 regulated genes in Keratinocyte Overexpression from PMID: 30772301 | SOX2 |
| 7950 | FANCD2 | SOX2 regulated genes in Keratinocyte Overexpression from PMID: 30772301 | SOX2 |
| 7951 | ACOT1 | SOX2 regulated genes in Keratinocyte Overexpression from PMID: 30772301 | SOX2 |
| 7952 | ACOT2 | SOX2 regulated genes in Keratinocyte Overexpression from PMID: 30772301 | SOX2 |
| 7953 | SSX2IP | SOX2 regulated genes in Keratinocyte Overexpression from PMID: 30772301 | SOX2 |
| 7954 | TCOF1 | SOX2 regulated genes in Keratinocyte Overexpression from PMID: 30772301 | SOX2 |
| 7955 | ZNF205 | SOX2 regulated genes in Keratinocyte Overexpression from PMID: 30772301 | SOX2 |
| 7956 | MAFK | SOX2 regulated genes in Keratinocyte Overexpression from PMID: 30772301 | SOX2 |
| 7957 | LIN7B | SOX2 regulated genes in Keratinocyte Overexpression from PMID: 30772301 | SOX2 |
| 7958 | DUSP7 | SOX2 regulated genes in Keratinocyte Overexpression from PMID: 30772301 | SOX2 |
| 7959 | NOTCH3 | SOX2 regulated genes in Keratinocyte Overexpression from PMID: 30772301 | SOX2 |
| 7960 | CFHR2 | SOX2 regulated genes in Keratinocyte Overexpression from PMID: 30772301 | SOX2 |
| 7961 | CFAP36 | SOX2 regulated genes in Keratinocyte Overexpression from PMID: 30772301 | SOX2 |
| 7962 | ZNF845 | SOX2 regulated genes in Keratinocyte Overexpression from PMID: 30772301 | SOX2 |
| 7963 | CTNND1 | SOX2 regulated genes in Keratinocyte Overexpression from PMID: 30772301 | SOX2 |
| 7964 | ZNF777 | SOX2 regulated genes in Keratinocyte Overexpression from PMID: 30772301 | SOX2 |
| 7965 | APLP1 | SOX2 regulated genes in Keratinocyte Overexpression from PMID: 30772301 | SOX2 |
| 7966 | TMEM231 | SOX2 regulated genes in Keratinocyte Overexpression from PMID: 30772301 | SOX2 |
| 7967 | CRTC2 | SOX2 regulated genes in Keratinocyte Overexpression from PMID: 30772301 | SOX2 |
| 7968 | FAM8A1 | SOX2 regulated genes in Keratinocyte Overexpression from PMID: 30772301 | SOX2 |
| 7969 | TMEM54 | SOX2 regulated genes in Keratinocyte Overexpression from PMID: 30772301 | SOX2 |
| 7970 | HNRNPUL1 | SOX2 regulated genes in Keratinocyte Overexpression from PMID: 30772301 | SOX2 |
| 7971 | CYB5R2 | SOX2 regulated genes in Keratinocyte Overexpression from PMID: 30772301 | SOX2 |
| 7972 | BCL9 | SOX2 regulated genes in Keratinocyte Overexpression from PMID: 30772301 | SOX2 |
| 7973 | FAM189A1 | SOX2 regulated genes in Keratinocyte Overexpression from PMID: 30772301 | SOX2 |
| 7974 | OSCAR | SOX2 regulated genes in Keratinocyte Overexpression from PMID: 30772301 | SOX2 |
| 7975 | DYNC1I1 | SOX2 regulated genes in Keratinocyte Overexpression from PMID: 30772301 | SOX2 |
| 7976 | ZNF286A | SOX2 regulated genes in Keratinocyte Overexpression from PMID: 30772301 | SOX2 |
| 7977 | RHOB | SOX2 regulated genes in Keratinocyte Overexpression from PMID: 30772301 | SOX2 |
| 7978 | MRPL43 | SOX2 regulated genes in Keratinocyte Overexpression from PMID: 30772301 | SOX2 |
| 7979 | KMT2D | SOX2 regulated genes in Keratinocyte Overexpression from PMID: 30772301 | SOX2 |
| 7980 | SRRM1 | SOX2 regulated genes in Keratinocyte Overexpression from PMID: 30772301 | SOX2 |
| 7981 | SRSF10 | SOX2 regulated genes in Keratinocyte Overexpression from PMID: 30772301 | SOX2 |
| 7982 | GPBP1L1 | SOX2 regulated genes in Keratinocyte Overexpression from PMID: 30772301 | SOX2 |
| 7983 | ADGRB1 | SOX2 regulated genes in Keratinocyte Overexpression from PMID: 30772301 | SOX2 |
| 7984 | PYCR3 | SOX2 regulated genes in Keratinocyte Overexpression from PMID: 30772301 | SOX2 |
| 7985 | FAM98A | SOX2 regulated genes in Keratinocyte Overexpression from PMID: 30772301 | SOX2 |
| 7986 | STARD10 | SOX2 regulated genes in Keratinocyte Overexpression from PMID: 30772301 | SOX2 |
| 7987 | CSNK1G2 | SOX2 regulated genes in Keratinocyte Overexpression from PMID: 30772301 | SOX2 |
| 7988 | TRIP12 | SOX2 regulated genes in Keratinocyte Overexpression from PMID: 30772301 | SOX2 |
| 7989 | WWTR1 | SOX2 regulated genes in Keratinocyte Overexpression from PMID: 30772301 | SOX2 |

| | | | |
|---|---|---|---|
| 7990 | C2CD3 | SOX2 regulated genes in Keratinocyte Overexpression from PMID: 30772301 | SOX2 |
| 7991 | SLC25A39 | SOX2 regulated genes in Keratinocyte Overexpression from PMID: 30772301 | SOX2 |
| 7992 | NACA | SOX2 regulated genes in Keratinocyte Overexpression from PMID: 30772301 | SOX2 |
| 7993 | MAP2K2 | SOX2 regulated genes in Keratinocyte Overexpression from PMID: 30772301 | SOX2 |
| 7994 | WDR6 | SOX2 regulated genes in Keratinocyte Overexpression from PMID: 30772301 | SOX2 |
| 7995 | TMEM216 | SOX2 regulated genes in Keratinocyte Overexpression from PMID: 30772301 | SOX2 |
| 7996 | XPO1 | SOX2 regulated genes in Keratinocyte Overexpression from PMID: 30772301 | SOX2 |
| 7997 | FAM53C | SOX2 regulated genes in Keratinocyte Overexpression from PMID: 30772301 | SOX2 |
| 7998 | SHANK2 | SOX2 regulated genes in Keratinocyte Overexpression from PMID: 30772301 | SOX2 |
| 7999 | CEBPZ | SOX2 regulated genes in Keratinocyte Overexpression from PMID: 30772301 | SOX2 |
| 8000 | CNOT1 | SOX2 regulated genes in Keratinocyte Overexpression from PMID: 30772301 | SOX2 |
| 8001 | CTF1 | SOX2 regulated genes in Keratinocyte Overexpression from PMID: 30772301 | SOX2 |
| 8002 | RPL19 | SOX2 regulated genes in Keratinocyte Overexpression from PMID: 30772301 | SOX2 |
| 8003 | C19orf33 | SOX2 regulated genes in Keratinocyte Overexpression from PMID: 30772301 | SOX2 |
| 8004 | ELP6 | SOX2 regulated genes in Keratinocyte Overexpression from PMID: 30772301 | SOX2 |
| 8005 | NAA38 | SOX2 regulated genes in Keratinocyte Overexpression from PMID: 30772301 | SOX2 |
| 8006 | LRRN2 | SOX2 regulated genes in Keratinocyte Overexpression from PMID: 30772301 | SOX2 |
| 8007 | SCCPDH | SOX2 regulated genes in Keratinocyte Overexpression from PMID: 30772301 | SOX2 |
| 8008 | ARHGAP33 | SOX2 regulated genes in Keratinocyte Overexpression from PMID: 30772301 | SOX2 |
| 8009 | TMEM161B | SOX2 regulated genes in Keratinocyte Overexpression from PMID: 30772301 | SOX2 |
| 8010 | RELL2 | SOX2 regulated genes in Keratinocyte Overexpression from PMID: 30772301 | SOX2 |
| 8011 | KCNJ14 | SOX2 regulated genes in Keratinocyte Overexpression from PMID: 30772301 | SOX2 |
| 8012 | SPINK8 | SOX2 regulated genes in Keratinocyte Overexpression from PMID: 30772301 | SOX2 |
| 8013 | LSM14A | SOX2 regulated genes in Keratinocyte Overexpression from PMID: 30772301 | SOX2 |
| 8014 | METRNL | SOX2 regulated genes in Keratinocyte Overexpression from PMID: 30772301 | SOX2 |
| 8015 | HSD11B2 | SOX2 regulated genes in Keratinocyte Overexpression from PMID: 30772301 | SOX2 |
| 8016 | DCAF7 | SOX2 regulated genes in Keratinocyte Overexpression from PMID: 30772301 | SOX2 |
| 8017 | ANKRD22 | SOX2 regulated genes in Keratinocyte Overexpression from PMID: 30772301 | SOX2 |
| 8018 | ZCRB1 | SOX2 regulated genes in Keratinocyte Overexpression from PMID: 30772301 | SOX2 |
| 8019 | ACAN | SOX2 regulated genes in Keratinocyte Overexpression from PMID: 30772301 | SOX2 |
| 8020 | GADD45GIP1 | SOX2 regulated genes in Keratinocyte Overexpression from PMID: 30772301 | SOX2 |
| 8021 | NKPD1 | SOX2 regulated genes in Keratinocyte Overexpression from PMID: 30772301 | SOX2 |
| 8022 | HOXD8 | SOX2 regulated genes in Keratinocyte Overexpression from PMID: 30772301 | SOX2 |
| 8023 | ARHGAP20 | SOX2 regulated genes in Keratinocyte Overexpression from PMID: 30772301 | SOX2 |
| 8024 | WDR89 | SOX2 regulated genes in Keratinocyte Overexpression from PMID: 30772301 | SOX2 |
| 8025 | SENP2 | SOX2 regulated genes in Keratinocyte Overexpression from PMID: 30772301 | SOX2 |
| 8026 | ELMOD3 | SOX2 regulated genes in Keratinocyte Overexpression from PMID: 30772301 | SOX2 |
| 8027 | TCERG1 | SOX2 regulated genes in Keratinocyte Overexpression from PMID: 30772301 | SOX2 |
| 8028 | CCDC174 | SOX2 regulated genes in Keratinocyte Overexpression from PMID: 30772301 | SOX2 |
| 8029 | PRCC | SOX2 regulated genes in Keratinocyte Overexpression from PMID: 30772301 | SOX2 |
| 8030 | DBI | SOX2 regulated genes in Keratinocyte Overexpression from PMID: 30772301 | SOX2 |
| 8031 | NEMF | SOX2 regulated genes in Keratinocyte Overexpression from PMID: 30772301 | SOX2 |
| 8032 | ZNF326 | SOX2 regulated genes in Keratinocyte Overexpression from PMID: 30772301 | SOX2 |
| 8033 | ITGB3BP | SOX2 regulated genes in Keratinocyte Overexpression from PMID: 30772301 | SOX2 |
| 8034 | JAGN1 | SOX2 regulated genes in Keratinocyte Overexpression from PMID: 30772301 | SOX2 |
| 8035 | SIPA1L2 | SOX2 regulated genes in Keratinocyte Overexpression from PMID: 30772301 | SOX2 |
| 8036 | XPO7 | SOX2 regulated genes in Keratinocyte Overexpression from PMID: 30772301 | SOX2 |
| 8037 | ELOF1 | SOX2 regulated genes in Keratinocyte Overexpression from PMID: 30772301 | SOX2 |
| 8038 | SPG7 | SOX2 regulated genes in Keratinocyte Overexpression from PMID: 30772301 | SOX2 |
| 8039 | DIMT1 | SOX2 regulated genes in Keratinocyte Overexpression from PMID: 30772301 | SOX2 |
| 8040 | RPS23 | SOX2 regulated genes in Keratinocyte Overexpression from PMID: 30772301 | SOX2 |
| 8041 | ARL6IP6 | SOX2 regulated genes in Keratinocyte Overexpression from PMID: 30772301 | SOX2 |
| 8042 | WNT4 | SOX2 regulated genes in Keratinocyte Overexpression from PMID: 30772301 | SOX2 |
| 8043 | LMBRD2 | SOX2 regulated genes in Keratinocyte Overexpression from PMID: 30772301 | SOX2 |
| 8044 | ZNF358 | SOX2 regulated genes in Keratinocyte Overexpression from PMID: 30772301 | SOX2 |
| 8045 | ANKRD13C | SOX2 regulated genes in Keratinocyte Overexpression from PMID: 30772301 | SOX2 |
| 8046 | GNL1 | SOX2 regulated genes in Keratinocyte Overexpression from PMID: 30772301 | SOX2 |
| 8047 | PTCD3 | SOX2 regulated genes in Keratinocyte Overexpression from PMID: 30772301 | SOX2 |
| 8048 | PKP1 | SOX2 regulated genes in Keratinocyte Overexpression from PMID: 30772301 | SOX2 |
| 8049 | ZNF579 | SOX2 regulated genes in Keratinocyte Overexpression from PMID: 30772301 | SOX2 |
| 8050 | MME | SOX2 regulated genes in Keratinocyte Overexpression from PMID: 30772301 | SOX2 |

| | | | |
|---|---|---|---|
| 8051 | PUM2 | SOX2 regulated genes in Keratinocyte Overexpression from PMID: 30772301 | SOX2 |
| 8052 | DDX20 | SOX2 regulated genes in Keratinocyte Overexpression from PMID: 30772301 | SOX2 |
| 8053 | MLPH | SOX2 regulated genes in Keratinocyte Overexpression from PMID: 30772301 | SOX2 |
| 8054 | KRT20 | SOX2 regulated genes in Keratinocyte Overexpression from PMID: 30772301 | SOX2 |
| 8055 | MSTO1 | SOX2 regulated genes in Keratinocyte Overexpression from PMID: 30772301 | SOX2 |
| 8056 | DHX30 | SOX2 regulated genes in Keratinocyte Overexpression from PMID: 30772301 | SOX2 |
| 8057 | NECAB2 | SOX2 regulated genes in Keratinocyte Overexpression from PMID: 30772301 | SOX2 |
| 8058 | RNF126 | SOX2 regulated genes in Keratinocyte Overexpression from PMID: 30772301 | SOX2 |
| 8059 | NFYC | SOX2 regulated genes in Keratinocyte Overexpression from PMID: 30772301 | SOX2 |
| 8060 | ALKBH3 | SOX2 regulated genes in Keratinocyte Overexpression from PMID: 30772301 | SOX2 |
| 8061 | GUCY2D | SOX2 regulated genes in Keratinocyte Overexpression from PMID: 30772301 | SOX2 |
| 8062 | MARK2 | SOX2 regulated genes in Keratinocyte Overexpression from PMID: 30772301 | SOX2 |
| 8063 | SLC25A10 | SOX2 regulated genes in Keratinocyte Overexpression from PMID: 30772301 | SOX2 |
| 8064 | RNASET2 | SOX2 regulated genes in Keratinocyte Overexpression from PMID: 30772301 | SOX2 |
| 8065 | CEP97 | SOX2 regulated genes in Keratinocyte Overexpression from PMID: 30772301 | SOX2 |
| 8066 | RPL24 | SOX2 regulated genes in Keratinocyte Overexpression from PMID: 30772301 | SOX2 |
| 8067 | ORC4 | SOX2 regulated genes in Keratinocyte Overexpression from PMID: 30772301 | SOX2 |
| 8068 | SNRNP48 | SOX2 regulated genes in Keratinocyte Overexpression from PMID: 30772301 | SOX2 |
| 8069 | CYSTM1 | SOX2 regulated genes in Keratinocyte Overexpression from PMID: 30772301 | SOX2 |
| 8070 | ZNF623 | SOX2 regulated genes in Keratinocyte Overexpression from PMID: 30772301 | SOX2 |
| 8071 | ZNF781 | SOX2 regulated genes in Keratinocyte Overexpression from PMID: 30772301 | SOX2 |
| 8072 | ARFIP2 | SOX2 regulated genes in Keratinocyte Overexpression from PMID: 30772301 | SOX2 |
| 8073 | HNRNPH1 | SOX2 regulated genes in Keratinocyte Overexpression from PMID: 30772301 | SOX2 |
| 8074 | WBP1 | SOX2 regulated genes in Keratinocyte Overexpression from PMID: 30772301 | SOX2 |
| 8075 | EML2 | SOX2 regulated genes in Keratinocyte Overexpression from PMID: 30772301 | SOX2 |
| 8076 | BRD8 | SOX2 regulated genes in Keratinocyte Overexpression from PMID: 30772301 | SOX2 |
| 8077 | TUBA1C | SOX2 regulated genes in Keratinocyte Overexpression from PMID: 30772301 | SOX2 |
| 8078 | CMTM4 | SOX2 regulated genes in Keratinocyte Overexpression from PMID: 30772301 | SOX2 |
| 8079 | CD9 | SOX2 regulated genes in Keratinocyte Overexpression from PMID: 30772301 | SOX2 |
| 8080 | ZNF672 | SOX2 regulated genes in Keratinocyte Overexpression from PMID: 30772301 | SOX2 |
| 8081 | CHTOP | SOX2 regulated genes in Keratinocyte Overexpression from PMID: 30772301 | SOX2 |
| 8082 | TMEM218 | SOX2 regulated genes in Keratinocyte Overexpression from PMID: 30772301 | SOX2 |
| 8083 | PEF1 | SOX2 regulated genes in Keratinocyte Overexpression from PMID: 30772301 | SOX2 |
| 8084 | COG1 | SOX2 regulated genes in Keratinocyte Overexpression from PMID: 30772301 | SOX2 |
| 8085 | IL1F10 | SOX2 regulated genes in Keratinocyte Overexpression from PMID: 30772301 | SOX2 |
| 8086 | SAMD4B | SOX2 regulated genes in Keratinocyte Overexpression from PMID: 30772301 | SOX2 |
| 8087 | ERBB2 | SOX2 regulated genes in Keratinocyte Overexpression from PMID: 30772301 | SOX2 |
| 8088 | NOM1 | SOX2 regulated genes in Keratinocyte Overexpression from PMID: 30772301 | SOX2 |
| 8089 | VAV3 | SOX2 regulated genes in Keratinocyte Overexpression from PMID: 30772301 | SOX2 |
| 8090 | DAPL1 | SOX2 regulated genes in Keratinocyte Overexpression from PMID: 30772301 | SOX2 |
| 8091 | RHBDD2 | SOX2 regulated genes in Keratinocyte Overexpression from PMID: 30772301 | SOX2 |
| 8092 | SH3BP5L | SOX2 regulated genes in Keratinocyte Overexpression from PMID: 30772301 | SOX2 |
| 8093 | TPGS1 | SOX2 regulated genes in Keratinocyte Overexpression from PMID: 30772301 | SOX2 |
| 8094 | KRTAP2-3 | SOX2 regulated genes in Keratinocyte Overexpression from PMID: 30772301 | SOX2 |
| 8095 | KRTAP4-11 | SOX2 regulated genes in Keratinocyte Overexpression from PMID: 30772301 | SOX2 |
| 8096 | KRTAP4-8 | SOX2 regulated genes in Keratinocyte Overexpression from PMID: 30772301 | SOX2 |
| 8097 | NACC1 | SOX2 regulated genes in Keratinocyte Overexpression from PMID: 30772301 | SOX2 |
| 8098 | RNF26 | SOX2 regulated genes in Keratinocyte Overexpression from PMID: 30772301 | SOX2 |
| 8099 | WNT9A | SOX2 regulated genes in Keratinocyte Overexpression from PMID: 30772301 | SOX2 |
| 8100 | HCAR1 | SOX2 regulated genes in Keratinocyte Overexpression from PMID: 30772301 | SOX2 |
| 8101 | DHFR2 | SOX2 regulated genes in Keratinocyte Overexpression from PMID: 30772301 | SOX2 |
| 8102 | TBP | SOX2 regulated genes in Keratinocyte Overexpression from PMID: 30772301 | SOX2 |
| 8103 | CAMSAP3 | SOX2 regulated genes in Keratinocyte Overexpression from PMID: 30772301 | SOX2 |
| 8104 | MDM1 | SOX2 regulated genes in Keratinocyte Overexpression from PMID: 30772301 | SOX2 |
| 8105 | CDC42EP4 | SOX2 regulated genes in Keratinocyte Overexpression from PMID: 30772301 | SOX2 |
| 8106 | MOB2 | SOX2 regulated genes in Keratinocyte Overexpression from PMID: 30772301 | SOX2 |
| 8107 | C5orf22 | SOX2 regulated genes in Keratinocyte Overexpression from PMID: 30772301 | SOX2 |
| 8108 | JMJD6 | SOX2 regulated genes in Keratinocyte Overexpression from PMID: 30772301 | SOX2 |
| 8109 | ELL | SOX2 regulated genes in Keratinocyte Overexpression from PMID: 30772301 | SOX2 |
| 8110 | RCCD1 | SOX2 regulated genes in Keratinocyte Overexpression from PMID: 30772301 | SOX2 |
| 8111 | THOC7 | SOX2 regulated genes in Keratinocyte Overexpression from PMID: 30772301 | SOX2 |

| | | | |
|---|---|---|---|
| 8112 | UGT1A6 | SOX2 regulated genes in Keratinocyte Overexpression from PMID: 30772301 | SOX2 |
| 8113 | MED16 | SOX2 regulated genes in Keratinocyte Overexpression from PMID: 30772301 | SOX2 |
| 8114 | VEZT | SOX2 regulated genes in Keratinocyte Overexpression from PMID: 30772301 | SOX2 |
| 8115 | SNRNP25 | SOX2 regulated genes in Keratinocyte Overexpression from PMID: 30772301 | SOX2 |
| 8116 | RNF19B | SOX2 regulated genes in Keratinocyte Overexpression from PMID: 30772301 | SOX2 |
| 8117 | DDX49 | SOX2 regulated genes in Keratinocyte Overexpression from PMID: 30772301 | SOX2 |
| 8118 | BAZ2B | SOX2 regulated genes in Keratinocyte Overexpression from PMID: 30772301 | SOX2 |
| 8119 | MUTYH | SOX2 regulated genes in Keratinocyte Overexpression from PMID: 30772301 | SOX2 |
| 8120 | DDX55 | SOX2 regulated genes in Keratinocyte Overexpression from PMID: 30772301 | SOX2 |
| 8121 | EIF4G1 | SOX2 regulated genes in Keratinocyte Overexpression from PMID: 30772301 | SOX2 |
| 8122 | VPS37B | SOX2 regulated genes in Keratinocyte Overexpression from PMID: 30772301 | SOX2 |
| 8123 | ATP13A3 | SOX2 regulated genes in Keratinocyte Overexpression from PMID: 30772301 | SOX2 |
| 8124 | RPL26 | SOX2 regulated genes in Keratinocyte Overexpression from PMID: 30772301 | SOX2 |
| 8125 | PLK2 | SOX2 regulated genes in Keratinocyte Overexpression from PMID: 30772301 | SOX2 |
| 8126 | SPRY4 | SOX2 regulated genes in Keratinocyte Overexpression from PMID: 30772301 | SOX2 |
| 8127 | SPATA2L | SOX2 regulated genes in Keratinocyte Overexpression from PMID: 30772301 | SOX2 |
| 8128 | EXOC3L4 | SOX2 regulated genes in Keratinocyte Overexpression from PMID: 30772301 | SOX2 |
| 8129 | F2RL1 | SOX2 regulated genes in Keratinocyte Overexpression from PMID: 30772301 | SOX2 |
| 8130 | KRT23 | SOX2 regulated genes in Keratinocyte Overexpression from PMID: 30772301 | SOX2 |
| 8131 | ULK1 | SOX2 regulated genes in Keratinocyte Overexpression from PMID: 30772301 | SOX2 |
| 8132 | C19orf67 | SOX2 regulated genes in Keratinocyte Overexpression from PMID: 30772301 | SOX2 |
| 8133 | CCNJ | SOX2 regulated genes in Keratinocyte Overexpression from PMID: 30772301 | SOX2 |
| 8134 | CNTROB | SOX2 regulated genes in Keratinocyte Overexpression from PMID: 30772301 | SOX2 |
| 8135 | UXS1 | SOX2 regulated genes in Keratinocyte Overexpression from PMID: 30772301 | SOX2 |
| 8136 | MVK | SOX2 regulated genes in Keratinocyte Overexpression from PMID: 30772301 | SOX2 |
| 8137 | RASAL2 | SOX2 regulated genes in Keratinocyte Overexpression from PMID: 30772301 | SOX2 |
| 8138 | STK11 | SOX2 regulated genes in Keratinocyte Overexpression from PMID: 30772301 | SOX2 |
| 8139 | RAD17 | SOX2 regulated genes in Keratinocyte Overexpression from PMID: 30772301 | SOX2 |
| 8140 | NDUFC2 | SOX2 regulated genes in Keratinocyte Overexpression from PMID: 30772301 | SOX2 |
| 8141 | DUOXA1 | SOX2 regulated genes in Keratinocyte Overexpression from PMID: 30772301 | SOX2 |
| 8142 | PPP2R2B | SOX2 regulated genes in Keratinocyte Overexpression from PMID: 30772301 | SOX2 |
| 8143 | TNFAIP8L1 | SOX2 regulated genes in Keratinocyte Overexpression from PMID: 30772301 | SOX2 |
| 8144 | HDAC4 | SOX2 regulated genes in Keratinocyte Overexpression from PMID: 30772301 | SOX2 |
| 8145 | DPP3 | SOX2 regulated genes in Keratinocyte Overexpression from PMID: 30772301 | SOX2 |
| 8146 | DHRS13 | SOX2 regulated genes in Keratinocyte Overexpression from PMID: 30772301 | SOX2 |
| 8147 | DYNC2LI1 | SOX2 regulated genes in Keratinocyte Overexpression from PMID: 30772301 | SOX2 |
| 8148 | ZBTB2 | SOX2 regulated genes in Keratinocyte Overexpression from PMID: 30772301 | SOX2 |
| 8149 | EIF1 | SOX2 regulated genes in Keratinocyte Overexpression from PMID: 30772301 | SOX2 |
| 8150 | RPF1 | SOX2 regulated genes in Keratinocyte Overexpression from PMID: 30772301 | SOX2 |
| 8151 | IPO7 | SOX2 regulated genes in Keratinocyte Overexpression from PMID: 30772301 | SOX2 |
| 8152 | RASSF9 | SOX2 regulated genes in Keratinocyte Overexpression from PMID: 30772301 | SOX2 |
| 8153 | ZFAND1 | SOX2 regulated genes in Keratinocyte Overexpression from PMID: 30772301 | SOX2 |
| 8154 | RABIF | SOX2 regulated genes in Keratinocyte Overexpression from PMID: 30772301 | SOX2 |
| 8155 | MYT1L | SOX2 regulated genes in Keratinocyte Overexpression from PMID: 30772301 | SOX2 |
| 8156 | USP37 | SOX2 regulated genes in Keratinocyte Overexpression from PMID: 30772301 | SOX2 |
| 8157 | GPN1 | SOX2 regulated genes in Keratinocyte Overexpression from PMID: 30772301 | SOX2 |
| 8158 | MED19 | SOX2 regulated genes in Keratinocyte Overexpression from PMID: 30772301 | SOX2 |
| 8159 | ADGRL3 | SOX2 regulated genes in Keratinocyte Overexpression from PMID: 30772301 | SOX2 |
| 8160 | STK11IP | SOX2 regulated genes in Keratinocyte Overexpression from PMID: 30772301 | SOX2 |
| 8161 | PNMA1 | SOX2 regulated genes in Keratinocyte Overexpression from PMID: 30772301 | SOX2 |
| 8162 | CC2D1B | SOX2 regulated genes in Keratinocyte Overexpression from PMID: 30772301 | SOX2 |
| 8163 | FABP4 | SOX2 regulated genes in Keratinocyte Overexpression from PMID: 30772301 | SOX2 |
| 8164 | ABHD14A | SOX2 regulated genes in Keratinocyte Overexpression from PMID: 30772301 | SOX2 |
| 8165 | NR2F6 | SOX2 regulated genes in Keratinocyte Overexpression from PMID: 30772301 | SOX2 |
| 8166 | ARMT1 | SOX2 regulated genes in Keratinocyte Overexpression from PMID: 30772301 | SOX2 |
| 8167 | WDR33 | SOX2 regulated genes in Keratinocyte Overexpression from PMID: 30772301 | SOX2 |
| 8168 | HNRNPA3 | SOX2 regulated genes in Keratinocyte Overexpression from PMID: 30772301 | SOX2 |
| 8169 | LRIG1 | SOX2 regulated genes in Keratinocyte Overexpression from PMID: 30772301 | SOX2 |
| 8170 | ETV2 | SOX2 regulated genes in Keratinocyte Overexpression from PMID: 30772301 | SOX2 |
| 8171 | BNIPL | SOX2 regulated genes in Keratinocyte Overexpression from PMID: 30772301 | SOX2 |
| 8172 | INSIG2 | SOX2 regulated genes in Keratinocyte Overexpression from PMID: 30772301 | SOX2 |

| | | | |
|---|---|---|---|
| 8173 | CETN3 | SOX2 regulated genes in Keratinocyte Overexpression from PMID: 30772301 | SOX2 |
| 8174 | FAM98C | SOX2 regulated genes in Keratinocyte Overexpression from PMID: 30772301 | SOX2 |
| 8175 | HAS3 | SOX2 regulated genes in Keratinocyte Overexpression from PMID: 30772301 | SOX2 |
| 8176 | PAN2 | SOX2 regulated genes in Keratinocyte Overexpression from PMID: 30772301 | SOX2 |
| 8177 | SPNS2 | SOX2 regulated genes in Keratinocyte Overexpression from PMID: 30772301 | SOX2 |
| 8178 | TRIM33 | SOX2 regulated genes in Keratinocyte Overexpression from PMID: 30772301 | SOX2 |
| 8179 | SGMS2 | SOX2 regulated genes in Keratinocyte Overexpression from PMID: 30772301 | SOX2 |
| 8180 | ZNF771 | SOX2 regulated genes in Keratinocyte Overexpression from PMID: 30772301 | SOX2 |
| 8181 | CLP1 | SOX2 regulated genes in Keratinocyte Overexpression from PMID: 30772301 | SOX2 |
| 8182 | DCBLD2 | SOX2 regulated genes in Keratinocyte Overexpression from PMID: 30772301 | SOX2 |
| 8183 | RRP9 | SOX2 regulated genes in Keratinocyte Overexpression from PMID: 30772301 | SOX2 |
| 8184 | OTUD3 | SOX2 regulated genes in Keratinocyte Overexpression from PMID: 30772301 | SOX2 |
| 8185 | LRRC8E | SOX2 regulated genes in Keratinocyte Overexpression from PMID: 30772301 | SOX2 |
| 8186 | TONSL | SOX2 regulated genes in Keratinocyte Overexpression from PMID: 30772301 | SOX2 |
| 8187 | NRDC | SOX2 regulated genes in Keratinocyte Overexpression from PMID: 30772301 | SOX2 |
| 8188 | DCP1A | SOX2 regulated genes in Keratinocyte Overexpression from PMID: 30772301 | SOX2 |
| 8189 | ELMO3 | SOX2 regulated genes in Keratinocyte Overexpression from PMID: 30772301 | SOX2 |
| 8190 | ZNF12 | SOX2 regulated genes in Keratinocyte Overexpression from PMID: 30772301 | SOX2 |
| 8191 | FAM126B | SOX2 regulated genes in Keratinocyte Overexpression from PMID: 30772301 | SOX2 |
| 8192 | EDARADD | SOX2 regulated genes in Keratinocyte Overexpression from PMID: 30772301 | SOX2 |
| 8193 | NABP2 | SOX2 regulated genes in Keratinocyte Overexpression from PMID: 30772301 | SOX2 |
| 8194 | TKT | SOX2 regulated genes in Keratinocyte Overexpression from PMID: 30772301 | SOX2 |
| 8195 | MCUR1 | SOX2 regulated genes in Keratinocyte Overexpression from PMID: 30772301 | SOX2 |
| 8196 | GALNT3 | SOX2 regulated genes in Keratinocyte Overexpression from PMID: 30772301 | SOX2 |
| 8197 | WIF1 | SOX2 regulated genes in Keratinocyte Overexpression from PMID: 30772301 | SOX2 |
| 8198 | SGTA | SOX2 regulated genes in Keratinocyte Overexpression from PMID: 30772301 | SOX2 |
| 8199 | DGKE | SOX2 regulated genes in Keratinocyte Overexpression from PMID: 30772301 | SOX2 |
| 8200 | OCLN | SOX2 regulated genes in Keratinocyte Overexpression from PMID: 30772301 | SOX2 |
| 8201 | PDLIM7 | SOX2 regulated genes in Keratinocyte Overexpression from PMID: 30772301 | SOX2 |
| 8202 | INHA | SOX2 regulated genes in Keratinocyte Overexpression from PMID: 30772301 | SOX2 |
| 8203 | BOLA2 | SOX2 regulated genes in Keratinocyte Overexpression from PMID: 30772301 | SOX2 |
| 8204 | USH1G | SOX2 regulated genes in Keratinocyte Overexpression from PMID: 30772301 | SOX2 |
| 8205 | KLHL18 | SOX2 regulated genes in Keratinocyte Overexpression from PMID: 30772301 | SOX2 |
| 8206 | PFDN1 | SOX2 regulated genes in Keratinocyte Overexpression from PMID: 30772301 | SOX2 |
| 8207 | DSP | SOX2 regulated genes in Keratinocyte Overexpression from PMID: 30772301 | SOX2 |
| 8208 | TRAF3IP1 | SOX2 regulated genes in Keratinocyte Overexpression from PMID: 30772301 | SOX2 |
| 8209 | MRPS12 | SOX2 regulated genes in Keratinocyte Overexpression from PMID: 30772301 | SOX2 |
| 8210 | KNL1 | SOX2 regulated genes in Keratinocyte Overexpression from PMID: 30772301 | SOX2 |
| 8211 | CACYBP | SOX2 regulated genes in Keratinocyte Overexpression from PMID: 30772301 | SOX2 |
| 8212 | ARIH2 | SOX2 regulated genes in Keratinocyte Overexpression from PMID: 30772301 | SOX2 |
| 8213 | POLQ | SOX2 regulated genes in Keratinocyte Overexpression from PMID: 30772301 | SOX2 |
| 8214 | TEN1 | SOX2 regulated genes in Keratinocyte Overexpression from PMID: 30772301 | SOX2 |
| 8215 | GTF2E1 | SOX2 regulated genes in Keratinocyte Overexpression from PMID: 30772301 | SOX2 |
| 8216 | RAP1GAP | SOX2 regulated genes in Keratinocyte Overexpression from PMID: 30772301 | SOX2 |
| 8217 | TMCO6 | SOX2 regulated genes in Keratinocyte Overexpression from PMID: 30772301 | SOX2 |
| 8218 | SPTBN4 | SOX2 regulated genes in Keratinocyte Overexpression from PMID: 30772301 | SOX2 |
| 8219 | TMEM237 | SOX2 regulated genes in Keratinocyte Overexpression from PMID: 30772301 | SOX2 |
| 8220 | ASF1B | SOX2 regulated genes in Keratinocyte Overexpression from PMID: 30772301 | SOX2 |
| 8221 | SLC8A2 | SOX2 regulated genes in Keratinocyte Overexpression from PMID: 30772301 | SOX2 |
| 8222 | SNX11 | SOX2 regulated genes in Keratinocyte Overexpression from PMID: 30772301 | SOX2 |
| 8223 | CTU1 | SOX2 regulated genes in Keratinocyte Overexpression from PMID: 30772301 | SOX2 |
| 8224 | AKAP8 | SOX2 regulated genes in Keratinocyte Overexpression from PMID: 30772301 | SOX2 |
| 8225 | CCNL2 | SOX2 regulated genes in Keratinocyte Overexpression from PMID: 30772301 | SOX2 |
| 8226 | SPC24 | SOX2 regulated genes in Keratinocyte Overexpression from PMID: 30772301 | SOX2 |
| 8227 | SMIM5 | SOX2 regulated genes in Keratinocyte Overexpression from PMID: 30772301 | SOX2 |
| 8228 | GPATCH3 | SOX2 regulated genes in Keratinocyte Overexpression from PMID: 30772301 | SOX2 |
| 8229 | RBM4 | SOX2 regulated genes in Keratinocyte Overexpression from PMID: 30772301 | SOX2 |
| 8230 | NUF2 | SOX2 regulated genes in Keratinocyte Overexpression from PMID: 30772301 | SOX2 |
| 8231 | MRPS14 | SOX2 regulated genes in Keratinocyte Overexpression from PMID: 30772301 | SOX2 |
| 8232 | XPOT | SOX2 regulated genes in Keratinocyte Overexpression from PMID: 30772301 | SOX2 |
| 8233 | LZTFL1 | SOX2 regulated genes in Keratinocyte Overexpression from PMID: 30772301 | SOX2 |

| | | | |
|---|---|---|---|
| 8234 | TMEM221 | SOX2 regulated genes in Keratinocyte Overexpression from PMID: 30772301 | SOX2 |
| 8235 | KLK13 | SOX2 regulated genes in Keratinocyte Overexpression from PMID: 30772301 | SOX2 |
| 8236 | C7orf25 | SOX2 regulated genes in Keratinocyte Overexpression from PMID: 30772301 | SOX2 |
| 8237 | GLMN | SOX2 regulated genes in Keratinocyte Overexpression from PMID: 30772301 | SOX2 |
| 8238 | WDR83 | SOX2 regulated genes in Keratinocyte Overexpression from PMID: 30772301 | SOX2 |
| 8239 | LINGO1 | SOX2 regulated genes in Keratinocyte Overexpression from PMID: 30772301 | SOX2 |
| 8240 | CNFN | SOX2 regulated genes in Keratinocyte Overexpression from PMID: 30772301 | SOX2 |
| 8241 | CLN8 | SOX2 regulated genes in Keratinocyte Overexpression from PMID: 30772301 | SOX2 |
| 8242 | RTBDN | SOX2 regulated genes in Keratinocyte Overexpression from PMID: 30772301 | SOX2 |
| 8243 | ALG8 | SOX2 regulated genes in Keratinocyte Overexpression from PMID: 30772301 | SOX2 |
| 8244 | ABCF3 | SOX2 regulated genes in Keratinocyte Overexpression from PMID: 30772301 | SOX2 |
| 8245 | DDX23 | SOX2 regulated genes in Keratinocyte Overexpression from PMID: 30772301 | SOX2 |
| 8246 | SLC26A9 | SOX2 regulated genes in Keratinocyte Overexpression from PMID: 30772301 | SOX2 |
| 8247 | TARBP2 | SOX2 regulated genes in Keratinocyte Overexpression from PMID: 30772301 | SOX2 |
| 8248 | FUT4 | SOX2 regulated genes in Keratinocyte Overexpression from PMID: 30772301 | SOX2 |
| 8249 | COPZ1 | SOX2 regulated genes in Keratinocyte Overexpression from PMID: 30772301 | SOX2 |
| 8250 | ETHE1 | SOX2 regulated genes in Keratinocyte Overexpression from PMID: 30772301 | SOX2 |
| 8251 | CNBP | SOX2 regulated genes in Keratinocyte Overexpression from PMID: 30772301 | SOX2 |
| 8252 | TMEM9 | SOX2 regulated genes in Keratinocyte Overexpression from PMID: 30772301 | SOX2 |
| 8253 | ZNF233 | SOX2 regulated genes in Keratinocyte Overexpression from PMID: 30772301 | SOX2 |
| 8254 | RXRB | SOX2 regulated genes in Keratinocyte Overexpression from PMID: 30772301 | SOX2 |
| 8255 | AKAP9 | SOX2 regulated genes in Keratinocyte Overexpression from PMID: 30772301 | SOX2 |
| 8256 | SERTAD4 | SOX2 regulated genes in Keratinocyte Overexpression from PMID: 30772301 | SOX2 |
| 8257 | UBALD2 | SOX2 regulated genes in Keratinocyte Overexpression from PMID: 30772301 | SOX2 |
| 8258 | DNASE2 | SOX2 regulated genes in Keratinocyte Overexpression from PMID: 30772301 | SOX2 |
| 8259 | OMG | SOX2 regulated genes in Keratinocyte Overexpression from PMID: 30772301 | SOX2 |
| 8260 | MAN1A2 | SOX2 regulated genes in Keratinocyte Overexpression from PMID: 30772301 | SOX2 |
| 8261 | TMEM80 | SOX2 regulated genes in Keratinocyte Overexpression from PMID: 30772301 | SOX2 |
| 8262 | OLA1 | SOX2 regulated genes in Keratinocyte Overexpression from PMID: 30772301 | SOX2 |
| 8263 | TRAPPC6A | SOX2 regulated genes in Keratinocyte Overexpression from PMID: 30772301 | SOX2 |
| 8264 | SMIM7 | SOX2 regulated genes in Keratinocyte Overexpression from PMID: 30772301 | SOX2 |
| 8265 | TMEM234 | SOX2 regulated genes in Keratinocyte Overexpression from PMID: 30772301 | SOX2 |
| 8266 | PCNX3 | SOX2 regulated genes in Keratinocyte Overexpression from PMID: 30772301 | SOX2 |
| 8267 | STK36 | SOX2 regulated genes in Keratinocyte Overexpression from PMID: 30772301 | SOX2 |
| 8268 | NUMA1 | SOX2 regulated genes in Keratinocyte Overexpression from PMID: 30772301 | SOX2 |
| 8269 | FAM72C | SOX2 regulated genes in Keratinocyte Overexpression from PMID: 30772301 | SOX2 |
| 8270 | FAM72A | SOX2 regulated genes in Keratinocyte Overexpression from PMID: 30772301 | SOX2 |
| 8271 | FAM72D | SOX2 regulated genes in Keratinocyte Overexpression from PMID: 30772301 | SOX2 |
| 8272 | FAM72B | SOX2 regulated genes in Keratinocyte Overexpression from PMID: 30772301 | SOX2 |
| 8273 | MCPH1 | SOX2 regulated genes in Keratinocyte Overexpression from PMID: 30772301 | SOX2 |
| 8274 | RBM15 | SOX2 regulated genes in Keratinocyte Overexpression from PMID: 30772301 | SOX2 |
| 8275 | MYL6 | SOX2 regulated genes in Keratinocyte Overexpression from PMID: 30772301 | SOX2 |
| 8276 | GNB5 | SOX2 regulated genes in Keratinocyte Overexpression from PMID: 30772301 | SOX2 |
| 8277 | SLC4A1AP | SOX2 regulated genes in Keratinocyte Overexpression from PMID: 30772301 | SOX2 |
| 8278 | KRTCAP3 | SOX2 regulated genes in Keratinocyte Overexpression from PMID: 30772301 | SOX2 |
| 8279 | RSRC1 | SOX2 regulated genes in Keratinocyte Overexpression from PMID: 30772301 | SOX2 |
| 8280 | SP9 | SOX2 regulated genes in Keratinocyte Overexpression from PMID: 30772301 | SOX2 |
| 8281 | ASTE1 | SOX2 regulated genes in Keratinocyte Overexpression from PMID: 30772301 | SOX2 |
| 8282 | TTC1 | SOX2 regulated genes in Keratinocyte Overexpression from PMID: 30772301 | SOX2 |
| 8283 | FLYWCH1 | SOX2 regulated genes in Keratinocyte Overexpression from PMID: 30772301 | SOX2 |
| 8284 | MRPS35 | SOX2 regulated genes in Keratinocyte Overexpression from PMID: 30772301 | SOX2 |
| 8285 | CLEC2D | SOX2 regulated genes in Keratinocyte Overexpression from PMID: 30772301 | SOX2 |
| 8286 | GLI2 | SOX2 regulated genes in Keratinocyte Overexpression from PMID: 30772301 | SOX2 |
| 8287 | GAS8 | SOX2 regulated genes in Keratinocyte Overexpression from PMID: 30772301 | SOX2 |
| 8288 | DPP4 | SOX2 regulated genes in Keratinocyte Overexpression from PMID: 30772301 | SOX2 |
| 8289 | ARMC9 | SOX2 regulated genes in Keratinocyte Overexpression from PMID: 30772301 | SOX2 |
| 8290 | FKBP8 | SOX2 regulated genes in Keratinocyte Overexpression from PMID: 30772301 | SOX2 |
| 8291 | AGER | SOX2 regulated genes in Keratinocyte Overexpression from PMID: 30772301 | SOX2 |
| 8292 | USP36 | SOX2 regulated genes in Keratinocyte Overexpression from PMID: 30772301 | SOX2 |
| 8293 | TMEM128 | SOX2 regulated genes in Keratinocyte Overexpression from PMID: 30772301 | SOX2 |
| 8294 | KIF21A | SOX2 regulated genes in Keratinocyte Overexpression from PMID: 30772301 | SOX2 |

| | | | |
|---|---|---|---|
| 8295 | BRI3BP | SOX2 regulated genes in Keratinocyte Overexpression from PMID: 30772301 | SOX2 |
| 8296 | ADAT3 | SOX2 regulated genes in Keratinocyte Overexpression from PMID: 30772301 | SOX2 |
| 8297 | USP54 | SOX2 regulated genes in Keratinocyte Overexpression from PMID: 30772301 | SOX2 |
| 8298 | TIPIN | SOX2 regulated genes in Keratinocyte Overexpression from PMID: 30772301 | SOX2 |
| 8299 | RPTN | SOX2 regulated genes in Keratinocyte Overexpression from PMID: 30772301 | SOX2 |
| 8300 | RAB3GAP2 | SOX2 regulated genes in Keratinocyte Overexpression from PMID: 30772301 | SOX2 |
| 8301 | ULK4 | SOX2 regulated genes in Keratinocyte Overexpression from PMID: 30772301 | SOX2 |
| 8302 | RAB24 | SOX2 regulated genes in Keratinocyte Overexpression from PMID: 30772301 | SOX2 |
| 8303 | PFKFB2 | SOX2 regulated genes in Keratinocyte Overexpression from PMID: 30772301 | SOX2 |
| 8304 | MARVELD3 | SOX2 regulated genes in Keratinocyte Overexpression from PMID: 30772301 | SOX2 |
| 8305 | BAG6 | SOX2 regulated genes in Keratinocyte Overexpression from PMID: 30772301 | SOX2 |
| 8306 | DCTPP1 | SOX2 regulated genes in Keratinocyte Overexpression from PMID: 30772301 | SOX2 |
| 8307 | UBXN7 | SOX2 regulated genes in Keratinocyte Overexpression from PMID: 30772301 | SOX2 |
| 8308 | THADA | SOX2 regulated genes in Keratinocyte Overexpression from PMID: 30772301 | SOX2 |
| 8309 | LTBP4 | SOX2 regulated genes in Keratinocyte Overexpression from PMID: 30772301 | SOX2 |
| 8310 | ARHGEF28 | SOX2 regulated genes in Keratinocyte Overexpression from PMID: 30772301 | SOX2 |
| 8311 | HNRNPA1 | SOX2 regulated genes in Keratinocyte Overexpression from PMID: 30772301 | SOX2 |
| 8312 | DDR1 | SOX2 regulated genes in Keratinocyte Overexpression from PMID: 30772301 | SOX2 |
| 8313 | TYW3 | SOX2 regulated genes in Keratinocyte Overexpression from PMID: 30772301 | SOX2 |
| 8314 | TOPBP1 | SOX2 regulated genes in Keratinocyte Overexpression from PMID: 30772301 | SOX2 |
| 8315 | PRKCZ | SOX2 regulated genes in Keratinocyte Overexpression from PMID: 30772301 | SOX2 |
| 8316 | DEAF1 | SOX2 regulated genes in Keratinocyte Overexpression from PMID: 30772301 | SOX2 |
| 8317 | BRD4 | SOX2 regulated genes in Keratinocyte Overexpression from PMID: 30772301 | SOX2 |
| 8318 | ATP5F1D | SOX2 regulated genes in Keratinocyte Overexpression from PMID: 30772301 | SOX2 |
| 8319 | MMACHC | SOX2 regulated genes in Keratinocyte Overexpression from PMID: 30772301 | SOX2 |
| 8320 | CCNJL | SOX2 regulated genes in Keratinocyte Overexpression from PMID: 30772301 | SOX2 |
| 8321 | PHF23 | SOX2 regulated genes in Keratinocyte Overexpression from PMID: 30772301 | SOX2 |
| 8322 | KIFAP3 | SOX2 regulated genes in Keratinocyte Overexpression from PMID: 30772301 | SOX2 |
| 8323 | POLE | SOX2 regulated genes in Keratinocyte Overexpression from PMID: 30772301 | SOX2 |
| 8324 | DDB1 | SOX2 regulated genes in Keratinocyte Overexpression from PMID: 30772301 | SOX2 |
| 8325 | CNIH4 | SOX2 regulated genes in Keratinocyte Overexpression from PMID: 30772301 | SOX2 |
| 8326 | GNG4 | SOX2 regulated genes in Keratinocyte Overexpression from PMID: 30772301 | SOX2 |
| 8327 | GORASP1 | SOX2 regulated genes in Keratinocyte Overexpression from PMID: 30772301 | SOX2 |
| 8328 | ZNF316 | SOX2 regulated genes in Keratinocyte Overexpression from PMID: 30772301 | SOX2 |
| 8329 | YOD1 | SOX2 regulated genes in Keratinocyte Overexpression from PMID: 30772301 | SOX2 |
| 8330 | TTLL4 | SOX2 regulated genes in Keratinocyte Overexpression from PMID: 30772301 | SOX2 |
| 8331 | JUNB | SOX2 regulated genes in Keratinocyte Overexpression from PMID: 30772301 | SOX2 |
| 8332 | KIF1B | SOX2 regulated genes in Keratinocyte Overexpression from PMID: 30772301 | SOX2 |
| 8333 | PRPF4B | SOX2 regulated genes in Keratinocyte Overexpression from PMID: 30772301 | SOX2 |
| 8334 | GRXCR2 | SOX2 regulated genes in Keratinocyte Overexpression from PMID: 30772301 | SOX2 |
| 8335 | NDC1 | SOX2 regulated genes in Keratinocyte Overexpression from PMID: 30772301 | SOX2 |
| 8336 | RBBP5 | SOX2 regulated genes in Keratinocyte Overexpression from PMID: 30772301 | SOX2 |
| 8337 | VAMP8 | SOX2 regulated genes in Keratinocyte Overexpression from PMID: 30772301 | SOX2 |
| 8338 | KNTC1 | SOX2 regulated genes in Keratinocyte Overexpression from PMID: 30772301 | SOX2 |
| 8339 | SOX13 | SOX2 regulated genes in Keratinocyte Overexpression from PMID: 30772301 | SOX2 |
| 8340 | TMEM79 | SOX2 regulated genes in Keratinocyte Overexpression from PMID: 30772301 | SOX2 |
| 8341 | INTS2 | SOX2 regulated genes in Keratinocyte Overexpression from PMID: 30772301 | SOX2 |
| 8342 | RBM34 | SOX2 regulated genes in Keratinocyte Overexpression from PMID: 30772301 | SOX2 |
| 8343 | PSMG3 | SOX2 regulated genes in Keratinocyte Overexpression from PMID: 30772301 | SOX2 |
| 8344 | NMNAT2 | SOX2 regulated genes in Keratinocyte Overexpression from PMID: 30772301 | SOX2 |
| 8345 | FAM171A2 | SOX2 regulated genes in Keratinocyte Overexpression from PMID: 30772301 | SOX2 |
| 8346 | TEX38 | SOX2 regulated genes in Keratinocyte Overexpression from PMID: 30772301 | SOX2 |
| 8347 | TTC36 | SOX2 regulated genes in Keratinocyte Overexpression from PMID: 30772301 | SOX2 |
| 8348 | CELF1 | SOX2 regulated genes in Keratinocyte Overexpression from PMID: 30772301 | SOX2 |
| 8349 | MRPL9 | SOX2 regulated genes in Keratinocyte Overexpression from PMID: 30772301 | SOX2 |
| 8350 | CERS3 | SOX2 regulated genes in Keratinocyte Overexpression from PMID: 30772301 | SOX2 |
| 8351 | CIDEB | SOX2 regulated genes in Keratinocyte Overexpression from PMID: 30772301 | SOX2 |
| 8352 | METTL2A | SOX2 regulated genes in Keratinocyte Overexpression from PMID: 30772301 | SOX2 |
| 8353 | IL11RA | SOX2 regulated genes in Keratinocyte Overexpression from PMID: 30772301 | SOX2 |
| 8354 | BCL7C | SOX2 regulated genes in Keratinocyte Overexpression from PMID: 30772301 | SOX2 |
| 8355 | AZIN1 | SOX2 regulated genes in Keratinocyte Overexpression from PMID: 30772301 | SOX2 |

| | | | |
|---|---|---|---|
| 8356 | DNAJC9 | SOX2 regulated genes in Keratinocyte Overexpression from PMID: 30772301 | SOX2 |
| 8357 | IMPA1 | SOX2 regulated genes in Keratinocyte Overexpression from PMID: 30772301 | SOX2 |
| 8358 | PRRC2C | SOX2 regulated genes in Keratinocyte Overexpression from PMID: 30772301 | SOX2 |
| 8359 | TXNDC9 | SOX2 regulated genes in Keratinocyte Overexpression from PMID: 30772301 | SOX2 |
| 8360 | ODC1 | SOX2 regulated genes in Keratinocyte Overexpression from PMID: 30772301 | SOX2 |
| 8361 | ZNF142 | SOX2 regulated genes in Keratinocyte Overexpression from PMID: 30772301 | SOX2 |
| 8362 | FOXRED1 | SOX2 regulated genes in Keratinocyte Overexpression from PMID: 30772301 | SOX2 |
| 8363 | YEATS2 | SOX2 regulated genes in Keratinocyte Overexpression from PMID: 30772301 | SOX2 |
| 8364 | DEF8 | SOX2 regulated genes in Keratinocyte Overexpression from PMID: 30772301 | SOX2 |
| 8365 | TRMT10C | SOX2 regulated genes in Keratinocyte Overexpression from PMID: 30772301 | SOX2 |
| 8366 | EFCAB12 | SOX2 regulated genes in Keratinocyte Overexpression from PMID: 30772301 | SOX2 |
| 8367 | PSRC1 | SOX2 regulated genes in Keratinocyte Overexpression from PMID: 30772301 | SOX2 |
| 8368 | IRF3 | SOX2 regulated genes in Keratinocyte Overexpression from PMID: 30772301 | SOX2 |
| 8369 | CSNK1A1 | SOX2 regulated genes in Keratinocyte Overexpression from PMID: 30772301 | SOX2 |
| 8370 | KANSL1 | SOX2 regulated genes in Keratinocyte Overexpression from PMID: 30772301 | SOX2 |
| 8371 | BDH1 | SOX2 regulated genes in Keratinocyte Overexpression from PMID: 30772301 | SOX2 |
| 8372 | TIA1 | SOX2 regulated genes in Keratinocyte Overexpression from PMID: 30772301 | SOX2 |
| 8373 | C16orf72 | SOX2 regulated genes in Keratinocyte Overexpression from PMID: 30772301 | SOX2 |
| 8374 | DARS2 | SOX2 regulated genes in Keratinocyte Overexpression from PMID: 30772301 | SOX2 |
| 8375 | PANX1 | SOX2 regulated genes in Keratinocyte Overexpression from PMID: 30772301 | SOX2 |
| 8376 | IPO11 | SOX2 regulated genes in Keratinocyte Overexpression from PMID: 30772301 | SOX2 |
| 8377 | PLEKHH2 | SOX2 regulated genes in Keratinocyte Overexpression from PMID: 30772301 | SOX2 |
| 8378 | SMG8 | SOX2 regulated genes in Keratinocyte Overexpression from PMID: 30772301 | SOX2 |
| 8379 | HMCES | SOX2 regulated genes in Keratinocyte Overexpression from PMID: 30772301 | SOX2 |
| 8380 | C1orf56 | SOX2 regulated genes in Keratinocyte Overexpression from PMID: 30772301 | SOX2 |
| 8381 | CYB561D1 | SOX2 regulated genes in Keratinocyte Overexpression from PMID: 30772301 | SOX2 |
| 8382 | CBX5 | SOX2 regulated genes in Keratinocyte Overexpression from PMID: 30772301 | SOX2 |
| 8383 | KLK10 | SOX2 regulated genes in Keratinocyte Overexpression from PMID: 30772301 | SOX2 |
| 8384 | SS18L2 | SOX2 regulated genes in Keratinocyte Overexpression from PMID: 30772301 | SOX2 |
| 8385 | RNF40 | SOX2 regulated genes in Keratinocyte Overexpression from PMID: 30772301 | SOX2 |
| 8386 | CLPP | SOX2 regulated genes in Keratinocyte Overexpression from PMID: 30772301 | SOX2 |
| 8387 | ZNF608 | SOX2 regulated genes in Keratinocyte Overexpression from PMID: 30772301 | SOX2 |
| 8388 | MYO18A | SOX2 regulated genes in Keratinocyte Overexpression from PMID: 30772301 | SOX2 |
| 8389 | HOXD1 | SOX2 regulated genes in Keratinocyte Overexpression from PMID: 30772301 | SOX2 |
| 8390 | SKA2 | SOX2 regulated genes in Keratinocyte Overexpression from PMID: 30772301 | SOX2 |
| 8391 | ATF7IP2 | SOX2 regulated genes in Keratinocyte Overexpression from PMID: 30772301 | SOX2 |
| 8392 | AMH | SOX2 regulated genes in Keratinocyte Overexpression from PMID: 30772301 | SOX2 |
| 8393 | MVB12A | SOX2 regulated genes in Keratinocyte Overexpression from PMID: 30772301 | SOX2 |
| 8394 | METTL16 | SOX2 regulated genes in Keratinocyte Overexpression from PMID: 30772301 | SOX2 |
| 8395 | BUB1 | SOX2 regulated genes in Keratinocyte Overexpression from PMID: 30772301 | SOX2 |
| 8396 | HDAC5 | SOX2 regulated genes in Keratinocyte Overexpression from PMID: 30772301 | SOX2 |
| 8397 | DENND2C | SOX2 regulated genes in Keratinocyte Overexpression from PMID: 30772301 | SOX2 |
| 8398 | ATR | SOX2 regulated genes in Keratinocyte Overexpression from PMID: 30772301 | SOX2 |
| 8399 | TRAPPC4 | SOX2 regulated genes in Keratinocyte Overexpression from PMID: 30772301 | SOX2 |
| 8400 | DDX6 | SOX2 regulated genes in Keratinocyte Overexpression from PMID: 30772301 | SOX2 |
| 8401 | PSD3 | SOX2 regulated genes in Keratinocyte Overexpression from PMID: 30772301 | SOX2 |
| 8402 | PLBD1 | SOX2 regulated genes in Keratinocyte Overexpression from PMID: 30772301 | SOX2 |
| 8403 | CCDC97 | SOX2 regulated genes in Keratinocyte Overexpression from PMID: 30772301 | SOX2 |
| 8404 | KIAA0825 | SOX2 regulated genes in Keratinocyte Overexpression from PMID: 30772301 | SOX2 |
| 8405 | TSEN2 | SOX2 regulated genes in Keratinocyte Overexpression from PMID: 30772301 | SOX2 |
| 8406 | KLF1 | SOX2 regulated genes in Keratinocyte Overexpression from PMID: 30772301 | SOX2 |
| 8407 | COL11A2 | SOX2 regulated genes in Keratinocyte Overexpression from PMID: 30772301 | SOX2 |
| 8408 | TDRD1 | SOX2 regulated genes in Keratinocyte Overexpression from PMID: 30772301 | SOX2 |
| 8409 | IGSF9 | SOX2 regulated genes in Keratinocyte Overexpression from PMID: 30772301 | SOX2 |
| 8410 | EPS8L2 | SOX2 regulated genes in Keratinocyte Overexpression from PMID: 30772301 | SOX2 |
| 8411 | TERF1 | SOX2 regulated genes in Keratinocyte Overexpression from PMID: 30772301 | SOX2 |
| 8412 | MXD3 | SOX2 regulated genes in Keratinocyte Overexpression from PMID: 30772301 | SOX2 |
| 8413 | GABPB1 | SOX2 regulated genes in Keratinocyte Overexpression from PMID: 30772301 | SOX2 |
| 8414 | WWP2 | SOX2 regulated genes in Keratinocyte Overexpression from PMID: 30772301 | SOX2 |
| 8415 | CCAR2 | SOX2 regulated genes in Keratinocyte Overexpression from PMID: 30772301 | SOX2 |
| 8416 | SH3TC2 | SOX2 regulated genes in Keratinocyte Overexpression from PMID: 30772301 | SOX2 |

| | | | |
|---|---|---|---|
| 8417 | PRR22 | SOX2 regulated genes in Keratinocyte Overexpression from PMID: 30772301 | SOX2 |
| 8418 | C1orf52 | SOX2 regulated genes in Keratinocyte Overexpression from PMID: 30772301 | SOX2 |
| 8419 | SH3GL1 | SOX2 regulated genes in Keratinocyte Overexpression from PMID: 30772301 | SOX2 |
| 8420 | DALRD3 | SOX2 regulated genes in Keratinocyte Overexpression from PMID: 30772301 | SOX2 |
| 8421 | NDUFAF3 | SOX2 regulated genes in Keratinocyte Overexpression from PMID: 30772301 | SOX2 |
| 8422 | NOSIP | SOX2 regulated genes in Keratinocyte Overexpression from PMID: 30772301 | SOX2 |
| 8423 | CREB3L4 | SOX2 regulated genes in Keratinocyte Overexpression from PMID: 30772301 | SOX2 |
| 8424 | TRPV1 | SOX2 regulated genes in Keratinocyte Overexpression from PMID: 30772301 | SOX2 |
| 8425 | HYOU1 | SOX2 regulated genes in Keratinocyte Overexpression from PMID: 30772301 | SOX2 |
| 8426 | PSPN | SOX2 regulated genes in Keratinocyte Overexpression from PMID: 30772301 | SOX2 |
| 8427 | SLC35E2B | SOX2 regulated genes in Keratinocyte Overexpression from PMID: 30772301 | SOX2 |
| 8428 | TMEM198 | SOX2 regulated genes in Keratinocyte Overexpression from PMID: 30772301 | SOX2 |
| 8429 | RPL35A | SOX2 regulated genes in Keratinocyte Overexpression from PMID: 30772301 | SOX2 |
| 8430 | LSM6 | SOX2 regulated genes in Keratinocyte Overexpression from PMID: 30772301 | SOX2 |
| 8431 | TYW5 | SOX2 regulated genes in Keratinocyte Overexpression from PMID: 30772301 | SOX2 |
| 8432 | NFRKB | SOX2 regulated genes in Keratinocyte Overexpression from PMID: 30772301 | SOX2 |
| 8433 | ZC3H10 | SOX2 regulated genes in Keratinocyte Overexpression from PMID: 30772301 | SOX2 |
| 8434 | RCC1L | SOX2 regulated genes in Keratinocyte Overexpression from PMID: 30772301 | SOX2 |
| 8435 | AHCTF1 | SOX2 regulated genes in Keratinocyte Overexpression from PMID: 30772301 | SOX2 |
| 8436 | ENGASE | SOX2 regulated genes in Keratinocyte Overexpression from PMID: 30772301 | SOX2 |
| 8437 | PSMD4 | SOX2 regulated genes in Keratinocyte Overexpression from PMID: 30772301 | SOX2 |
| 8438 | SMG7 | SOX2 regulated genes in Keratinocyte Overexpression from PMID: 30772301 | SOX2 |
| 8439 | CDK10 | SOX2 regulated genes in Keratinocyte Overexpression from PMID: 30772301 | SOX2 |
| 8440 | KLHL21 | SOX2 regulated genes in Keratinocyte Overexpression from PMID: 30772301 | SOX2 |
| 8441 | EVI5L | SOX2 regulated genes in Keratinocyte Overexpression from PMID: 30772301 | SOX2 |
| 8442 | INTS11 | SOX2 regulated genes in Keratinocyte Overexpression from PMID: 30772301 | SOX2 |
| 8443 | SUPT6H | SOX2 regulated genes in Keratinocyte Overexpression from PMID: 30772301 | SOX2 |
| 8444 | FZD7 | SOX2 regulated genes in Keratinocyte Overexpression from PMID: 30772301 | SOX2 |
| 8445 | SPAST | SOX2 regulated genes in Keratinocyte Overexpression from PMID: 30772301 | SOX2 |
| 8446 | APOLD1 | SOX2 regulated genes in Keratinocyte Overexpression from PMID: 30772301 | SOX2 |
| 8447 | RAPGEFL1 | SOX2 regulated genes in Keratinocyte Overexpression from PMID: 30772301 | SOX2 |
| 8448 | RNFT2 | SOX2 regulated genes in Keratinocyte Overexpression from PMID: 30772301 | SOX2 |
| 8449 | ADGRL1 | SOX2 regulated genes in Keratinocyte Overexpression from PMID: 30772301 | SOX2 |
| 8450 | MOB4 | SOX2 regulated genes in Keratinocyte Overexpression from PMID: 30772301 | SOX2 |
| 8451 | S100A6 | SOX2 regulated genes in Keratinocyte Overexpression from PMID: 30772301 | SOX2 |
| 8452 | SCYL3 | SOX2 regulated genes in Keratinocyte Overexpression from PMID: 30772301 | SOX2 |
| 8453 | RAPGEF2 | SOX2 regulated genes in Keratinocyte Overexpression from PMID: 30772301 | SOX2 |
| 8454 | STAMBP | SOX2 regulated genes in Keratinocyte Overexpression from PMID: 30772301 | SOX2 |
| 8455 | LSR | SOX2 regulated genes in Keratinocyte Overexpression from PMID: 30772301 | SOX2 |
| 8456 | FBLN7 | SOX2 regulated genes in Keratinocyte Overexpression from PMID: 30772301 | SOX2 |
| 8457 | ATIC | SOX2 regulated genes in Keratinocyte Overexpression from PMID: 30772301 | SOX2 |
| 8458 | CARF | SOX2 regulated genes in Keratinocyte Overexpression from PMID: 30772301 | SOX2 |
| 8459 | NHSL1 | SOX2 regulated genes in Keratinocyte Overexpression from PMID: 30772301 | SOX2 |
| 8460 | CSRNP2 | SOX2 regulated genes in Keratinocyte Overexpression from PMID: 30772301 | SOX2 |
| 8461 | CCND1 | SOX2 regulated genes in Keratinocyte Overexpression from PMID: 30772301 | SOX2 |
| 8462 | UBE2G1 | SOX2 regulated genes in Keratinocyte Overexpression from PMID: 30772301 | SOX2 |
| 8463 | CDKN2D | SOX2 regulated genes in Keratinocyte Overexpression from PMID: 30772301 | SOX2 |
| 8464 | WDR46 | SOX2 regulated genes in Keratinocyte Overexpression from PMID: 30772301 | SOX2 |
| 8465 | SETD1B | SOX2 regulated genes in Keratinocyte Overexpression from PMID: 30772301 | SOX2 |
| 8466 | PUM1 | SOX2 regulated genes in Keratinocyte Overexpression from PMID: 30772301 | SOX2 |
| 8467 | NDUFV1 | SOX2 regulated genes in Keratinocyte Overexpression from PMID: 30772301 | SOX2 |
| 8468 | RABGGTB | SOX2 regulated genes in Keratinocyte Overexpression from PMID: 30772301 | SOX2 |
| 8469 | SLC35A1 | SOX2 regulated genes in Keratinocyte Overexpression from PMID: 30772301 | SOX2 |
| 8470 | CSPP1 | SOX2 regulated genes in Keratinocyte Overexpression from PMID: 30772301 | SOX2 |
| 8471 | COMMD1 | SOX2 regulated genes in Keratinocyte Overexpression from PMID: 30772301 | SOX2 |
| 8472 | LYPD3 | SOX2 regulated genes in Keratinocyte Overexpression from PMID: 30772301 | SOX2 |
| 8473 | ST3GAL5 | SOX2 regulated genes in Keratinocyte Overexpression from PMID: 30772301 | SOX2 |
| 8474 | VPS54 | SOX2 regulated genes in Keratinocyte Overexpression from PMID: 30772301 | SOX2 |
| 8475 | CTTNBP2 | SOX2 regulated genes in Keratinocyte Overexpression from PMID: 30772301 | SOX2 |
| 8476 | MAP2K7 | SOX2 regulated genes in Keratinocyte Overexpression from PMID: 30772301 | SOX2 |
| 8477 | MYO19 | SOX2 regulated genes in Keratinocyte Overexpression from PMID: 30772301 | SOX2 |

| | | | |
|---|---|---|---|
| 8478 | FMO4 | SOX2 regulated genes in Keratinocyte Overexpression from PMID: 30772301 | SOX2 |
| 8479 | VEPH1 | SOX2 regulated genes in Keratinocyte Overexpression from PMID: 30772301 | SOX2 |
| 8480 | PPL | SOX2 regulated genes in Keratinocyte Overexpression from PMID: 30772301 | SOX2 |
| 8481 | EFHD1 | SOX2 regulated genes in Keratinocyte Overexpression from PMID: 30772301 | SOX2 |
| 8482 | DYNC2H1 | SOX2 regulated genes in Keratinocyte Overexpression from PMID: 30772301 | SOX2 |
| 8483 | SNRNP70 | SOX2 regulated genes in Keratinocyte Overexpression from PMID: 30772301 | SOX2 |
| 8484 | KDF1 | SOX2 regulated genes in Keratinocyte Overexpression from PMID: 30772301 | SOX2 |
| 8485 | GLIS1 | SOX2 regulated genes in Keratinocyte Overexpression from PMID: 30772301 | SOX2 |
| 8486 | ERC2 | SOX2 regulated genes in Keratinocyte Overexpression from PMID: 30772301 | SOX2 |
| 8487 | LRRC40 | SOX2 regulated genes in Keratinocyte Overexpression from PMID: 30772301 | SOX2 |
| 8488 | TFDP2 | SOX2 regulated genes in Keratinocyte Overexpression from PMID: 30772301 | SOX2 |
| 8489 | NCDN | SOX2 regulated genes in Keratinocyte Overexpression from PMID: 30772301 | SOX2 |
| 8490 | SLC35A4 | SOX2 regulated genes in Keratinocyte Overexpression from PMID: 30772301 | SOX2 |
| 8491 | MRPL35 | SOX2 regulated genes in Keratinocyte Overexpression from PMID: 30772301 | SOX2 |

**Supplemental Table 6.** Module 7 genes known to be regulated by SOX2 and/or KLF5.

| | gene | source | regulator |
|---|---|---|---|
| 1 | APOL1 | SOX2 regulated genes in Wounding from PMID: 30772301 | SOX2 |
| 2 | APOL1 | SOX2 regulated genes in Keratinocyte Overexpression from PMID: 30772301 | SOX2 |
| 3 | APOL1 | Enrichr CHEA_2022: SOX2 20726797 ChIP-Seq SW620 Human | SOX2 |
| 4 | ATP8B1 | SOX2-KLF5 coregulated genes from PMID: 33972779 | SOX2 + KLF5 |
| 5 | BCL2L15 | SOX2-KLF5 coregulated genes from PMID: 33972779 | SOX2 + KLF5 |
| 6 | CA2 | SOX2 regulated genes in Wounding from PMID: 30772301 | SOX2 |
| 7 | CA2 | Enrichr CHEA_2022: SOX2 20726797 ChIP-Seq SW620 Human | SOX2 |
| 8 | CAMK1D | Enrichr TF Perturbations followed by expression: KLF5 20726797 ChIP-Seq SW620 Human | KLF5 |
| 9 | CDH3 | SOX2 regulated genes in Wounding from PMID: 30772301 | SOX2 |
| 10 | CLDN1 | SOX2 regulated genes in Wounding from PMID: 30772301 | SOX2 |
| 11 | CLDN1 | SOX2-KLF5 coregulated genes from PMID: 33972779 | SOX2 + KLF5 |
| 12 | CLDN1 | SOX2 regulated genes in Keratinocyte Overexpression from PMID: 30772301 | SOX2 |
| 13 | CLDN1 | Enrichr CHEA_2022: SOX2 20726797 ChIP-Seq SW620 Human | SOX2 |
| 14 | CYP2S1 | SOX2 regulated genes in Wounding from PMID: 30772301 | SOX2 |
| 15 | DPYD | Enrichr CHEA_2022: SOX2 20726797 ChIP-Seq SW620 Human | SOX2 |
| 16 | DSG3 | SOX2-KLF5 coregulated genes from PMID: 33972779 | SOX2 + KLF5 |
| 17 | DSG3 | SOX2 regulated genes in Wounding from PMID: 30772301 | SOX2 |
| 18 | DSG3 | SOX2 regulated genes in Keratinocyte Overexpression from PMID: 30772301 | SOX2 |
| 19 | DSG3 | Enrichr TF Perturbations followed by expression: KLF5 20726797 ChIP-Seq SW620 Human | KLF5 |
| 20 | DSP | SOX2-KLF5 coregulated genes from PMID: 33972779 | SOX2 + KLF5 |
| 21 | DSP | SOX2 regulated genes in Wounding from PMID: 30772301 | SOX2 |
| 22 | DSP | SOX2 regulated genes in Keratinocyte Overexpression from PMID: 30772301 | SOX2 |
| 23 | DSP | Enrichr CHEA_2022: SOX2 20726797 ChIP-Seq SW620 Human | SOX2 |
| 24 | DUOX1 | SOX2 regulated genes in Wounding from PMID: 30772301 | SOX2 |
| 25 | EPPK1 | SOX2 regulated genes in Wounding from PMID: 30772301 | SOX2 |
| 26 | EPPK1 | Enrichr CHEA_2022: SOX2 20726797 ChIP-Seq SW620 Human | SOX2 |
| 27 | GALNT5 | Enrichr CHEA_2022: SOX2 20726797 ChIP-Seq SW620 Human | SOX2 |
| 28 | GALNT5 | Enrichr TF Perturbations followed by expression: KLF5 20726797 ChIP-Seq SW620 Human | KLF5 |
| 29 | HAS3 | SOX2 regulated genes in Keratinocyte Overexpression from PMID: 30772301 | SOX2 |
| 30 | HAS3 | SOX2 regulated genes in Wounding from PMID: 30772301 | SOX2 |
| 31 | HAS3 | Enrichr CHEA_2022: SOX2 20726797 ChIP-Seq SW620 Human | SOX2 |
| 32 | HPGD | Enrichr CHEA_2022: SOX2 20726797 ChIP-Seq SW620 Human | SOX2 |
| 33 | HPGD | SOX2 regulated genes from PMID: 20726797 | SOX2 |
| 34 | KCNJ2 | SOX2-KLF5 coregulated genes from PMID: 33972779 | SOX2 + KLF5 |
| 35 | MPP7 | SOX2 regulated genes in Wounding from PMID: 30772301 | SOX2 |
| 36 | MX1 | Enrichr CHEA_2022: SOX2 20726797 ChIP-Seq SW620 Human | SOX2 |
| 37 | MYH9 | SOX2 regulated genes in Wounding from PMID: 30772301 | SOX2 |
| 38 | MYH9 | SOX2-KLF5 coregulated genes from PMID: 33972779 | SOX2 + KLF5 |
| 39 | MYH9 | SOX2 regulated genes in Keratinocyte Overexpression from PMID: 30772301 | SOX2 |
| 40 | NEFL | Enrichr TF Perturbations followed by expression: KLF5 20726797 ChIP-Seq SW620 Human | KLF5 |
| 41 | PDZK1IP1 | Enrichr CHEA_2022: SOX2 20726797 ChIP-Seq SW620 Human | SOX2 |
| 42 | PHLDB2 | Enrichr CHEA_2022: SOX2 20726797 ChIP-Seq SW620 Human | SOX2 |
| 43 | PHLDB2 | SOX2 regulated genes in Wounding from PMID: 30772301 | SOX2 |
| 44 | PHLDB2 | SOX2 regulated genes in Keratinocyte Overexpression from PMID: 30772301 | SOX2 |
| 45 | SERPINB3 | Enrichr CHEA_2022: SOX2 20726797 ChIP-Seq SW620 Human | SOX2 |
| 46 | SFRP1 | SOX2 regulated genes in Keratinocyte Overexpression from PMID: 30772301 | SOX2 |
| 47 | SFRP1 | SOX2 regulated genes in Wounding from PMID: 30772301 | SOX2 |
| 48 | SGK1 | SOX2-KLF5 coregulated genes from PMID: 33972779 | SOX2 + KLF5 |
| 49 | SGK1 | SOX2 regulated genes in Wounding from PMID: 30772301 | SOX2 |
| 50 | SH3RF2 | SOX2 regulated genes in Keratinocyte Overexpression from PMID: 30772301 | SOX2 |
| 51 | SH3RF2 | Enrichr TF Perturbations followed by expression: KLF5 20726797 ChIP-Seq SW620 Human | KLF5 |
| 52 | TFPI | Enrichr CHEA_2022: SOX2 20726797 ChIP-Seq SW620 Human | SOX2 |
| 53 | TNFAIP6 | SOX2 regulated genes in Wounding from PMID: 30772301 | SOX2 |

**Supplemental Table 7.** Gene expression profile of upregulated EoE DEGs known to be regulated by SOX2 and/or KLF5.

| | gene | Log2FC | Pval | FDR_Pval | pct.EoE_Biopsy | pct.Healthy_Control | Top_Changed_Compartment | Regulators |
|---|---|---|---|---|---|---|---|---|
| 1 | TNFAIP6 | 14.03276603 | 0.000195546 | 0.010489143 | 26.30969971 | 0.008333034 | Suprabasal | SOX2 |
| 2 | NRXN1 | 8.481843776 | 0.000599264 | 0.010733065 | 5.067165747 | 0.005928385 | Suprabasal | SOX2 + KLF5 |
| 3 | DPP4 | 7.164647328 | 0.004291437 | 0.022284502 | 7.110944705 | 0.042029564 | Suprabasal | SOX2 |
| 4 | TRPM6 | 6.351720177 | 0.000545165 | 0.007274975 | 12.68360569 | 0.06651167 | Superficial | SOX2 |
| 5 | LURAP1L | 6.213447206 | 0.000715353 | 0.007615337 | 18.34363791 | 0.132134256 | Superficial | SOX2 |
| 6 | GLDC | 5.812522131 | 9.15E-05 | 0.010489143 | 17.66430308 | 0.256528795 | Suprabasal | KLF5 |
| 7 | MYRFL | 5.249646268 | 0.000375835 | 0.007176147 | 6.520629319 | 0.195081078 | Superficial | SOX2 |
| 8 | NEFL | 5.137160698 | 0.000504805 | 0.007178523 | 23.20878673 | 1.038348856 | Superficial | KLF5 |
| 9 | RPTN | 4.987397171 | 9.26E-05 | 0.007176147 | 9.402714391 | 0.681967049 | Superficial | SOX2 |
| 10 | NFATC2 | 4.981012295 | 0.000989444 | 0.008042817 | 15.46639165 | 0.386585977 | Superficial | SOX2 + KLF5 |
| 11 | LBH | 4.82051517 | 0.000506425 | 0.010489143 | 15.18280651 | 0.367077206 | Suprabasal | SOX2 |
| 12 | ADAM28 | 4.740686733 | 0.003303641 | 0.024854712 | 5.530658274 | 0.161625895 | Basal | KLF5 |
| 13 | KCNJ2 | 4.535506071 | 0.000196466 | 0.010489143 | 42.41879274 | 3.237283542 | Suprabasal | SOX2 + KLF5 |
| 14 | GABRP | 4.346532147 | 5.55E-05 | 0.012455402 | 27.82034888 | 2.002158343 | Basal | SOX2, KLF5 |
| 15 | SFRP1 | 4.336773754 | 0.000153669 | 0.010489143 | 44.6669391 | 2.593848432 | Suprabasal | SOX2 |
| 16 | EDAR | 4.248568858 | 0.000401828 | 0.007176147 | 19.44942997 | 1.08545527 | Superficial | SOX2 |
| 17 | EPPK1 | 4.143182566 | 0.006166656 | 0.027303057 | 34.09283879 | 3.257315631 | Suprabasal | SOX2 |
| 18 | IFFO2 | 4.134503394 | 0.000250281 | 0.007176147 | 59.91842482 | 9.317539544 | Superficial | SOX2 |
| 19 | CD200R1 | 4.133533272 | 0.000506425 | 0.007178523 | 27.369393 | 0.983226601 | Superficial | SOX2 |
| 20 | APOL1 | 4.043383815 | 0.001232925 | 0.008679903 | 40.60842821 | 3.499022944 | Superficial | SOX2 |
| 21 | BMPR1B | 3.971683288 | 0.00591569 | 0.032981404 | 5.098860854 | 0.282051825 | Basal | SOX2 |
| 22 | IFI35 | 3.965447773 | 0.001232925 | 0.008679903 | 48.98584135 | 4.252996719 | Superficial | SOX2 |
| 23 | UPK1B | 3.886272489 | 0.000119764 | 0.007176147 | 30.00643469 | 3.383373143 | Superficial | KLF5, SOX2 |
| 24 | CFB | 3.852144876 | 0.000401828 | 0.007176147 | 33.99028752 | 3.320047752 | Superficial | SOX2 |
| 25 | LOXL4 | 3.819802832 | 0.000992312 | 0.014320968 | 17.36548658 | 1.18068853 | Basal | SOX2 |
| 26 | RUNX2 | 3.769129479 | 0.000504805 | 0.007178523 | 13.80876203 | 0.842100666 | Superficial | SOX2, SOX2 + KLF5 |
| 27 | CA2 | 3.736257496 | 7.20E-05 | 0.010489143 | 56.63468793 | 13.10997844 | Suprabasal | SOX2 |
| 28 | SLC28A3 | 3.717054301 | 0.000635975 | 0.007274975 | 22.8202256 | 2.11699801 | Superficial | SOX2 |
| 29 | LYPD6B | 3.712117954 | 0.004211401 | 0.016013117 | 23.48505282 | 1.455960635 | Superficial | SOX2 + KLF5, SOX2 |
| 30 | SGK1 | 3.680085691 | 0.000153669 | 0.007176147 | 61.49330151 | 14.372983 | Superficial | SOX2 + KLF5, SOX2 |
| 31 | SERPINB3 | 3.601168438 | 7.20E-05 | 0.012455402 | 61.04787159 | 17.17234047 | Basal | SOX2 |
| 32 | ST8SIA6 | 3.586099976 | 0.000634007 | 0.007274975 | 9.78899944 | 0.441575518 | Superficial | SOX2 + KLF5, SOX2 |
| 33 | CCDC68 | 3.570963328 | 0.000506425 | 0.007178523 | 25.08739932 | 1.949503374 | Superficial | SOX2 |
| 34 | CMYA5 | 3.552817397 | 0.001526458 | 0.009387421 | 11.51192042 | 0.649201685 | Superficial | SOX2 + KLF5 |
| 35 | PHLDB2 | 3.526366051 | 0.000795824 | 0.010838271 | 19.35094986 | 1.167949601 | Suprabasal | SOX2 |
| 36 | CYP2S1 | 3.520993063 | 0.000992312 | 0.008042817 | 38.06773576 | 4.106293048 | Superficial | SOX2 |
| 37 | HAS3 | 3.515910568 | 0.000635975 | 0.007274975 | 31.49524154 | 3.578664882 | Superficial | SOX2 |
| 38 | PCSK6 | 3.498236414 | 5.55E-05 | 0.012455402 | 10.72980733 | 1.09770805 | Basal | SOX2 |
| 39 | NFE2L3 | 3.298054351 | 0.000250281 | 0.007176147 | 27.78456634 | 3.028185524 | Superficial | SOX2 |
| 40 | BCL2L15 | 3.169348667 | 0.000401828 | 0.010489143 | 52.77173551 | 9.09037378 | Suprabasal | SOX2 + KLF5 |
| 41 | UPK3BL2 | 3.151048298 | 0.0033823 | 0.014108599 | 11.77819237 | 0.999182365 | Superficial | SOX2 |
| 42 | CADM4 | 3.111929462 | 0.000401828 | 0.007176147 | 34.66563546 | 5.040451809 | Superficial | SOX2 |
| 43 | PAPSS2 | 3.082313982 | 0.002315114 | 0.020052792 | 14.29165028 | 1.259098065 | Basal | SOX2 |
| 44 | MUC4 | 3.064276121 | 0.001883197 | 0.018415354 | 9.945660526 | 1.283194417 | Basal | KLF5 |
| 45 | NAV1 | 3.01803895 | 0.000795824 | 0.007615337 | 24.20119131 | 3.215183485 | Superficial | SOX2 |
| 46 | TNS4 | 2.993108132 | 0.001232925 | 0.008679903 | 32.53389208 | 3.314587495 | Superficial | SOX2, SOX2 + KLF5 |
| 47 | APOL4 | 2.985249452 | 0.001526458 | 0.009387421 | 13.88024072 | 1.439759571 | Superficial | SOX2 |
| 48 | FA2H | 2.981191994 | 7.20E-05 | 0.007176147 | 27.14904728 | 2.928368931 | Superficial | KLF5 |
| 49 | GALNT4 | 2.956378661 | 0.000963884 | 0.008042817 | 31.11337918 | 3.725274822 | Superficial | SOX2, KLF5 |
| 50 | GCNT4 | 2.929654443 | 0.000196466 | 0.007176147 | 10.75329782 | 1.615330968 | Superficial | SOX2 |
| 51 | KLK1 | 2.807174652 | 0.005105016 | 0.018023407 | 15.83369405 | 1.804143357 | Superficial | KLF5, SOX2 |
| 52 | RASGRP1 | 2.782742217 | 0.008904627 | 0.040392504 | 17.65106615 | 2.44746748 | Basal | SOX2 + KLF5 |
| 53 | PLA2G3 | 2.78085798 | 0.000989444 | 0.008042817 | 18.96022052 | 2.368144974 | Superficial | SOX2 |
| 54 | SERPINB13 | 2.770886827 | 0.000119764 | 0.012455402 | 64.24636718 | 20.44419918 | Basal | SOX2 |
| 55 | FOXE1 | 2.755248075 | 0.000795824 | 0.007615337 | 34.51116941 | 5.187653321 | Superficial | SOX2 |
| 56 | HRH1 | 2.7242374 | 0.001883197 | 0.014679007 | 24.54736841 | 3.982327992 | Suprabasal | SOX2 + KLF5 |
| 57 | APOL2 | 2.72256637 | 0.001232925 | 0.008679903 | 27.58021793 | 3.445414582 | Superficial | SOX2 |
| 58 | IFI27 | 2.685018167 | 0.002836074 | 0.0176927 | 78.44556802 | 38.98017695 | Suprabasal | SOX2 |
| 59 | KITLG | 2.677505538 | 7.20E-05 | 0.010489143 | 40.63911213 | 7.568671567 | Suprabasal | SOX2 |
| 60 | LY75 | 2.675558265 | 5.55E-05 | 0.007176147 | 16.56232468 | 1.694338305 | Superficial | SOX2, SOX2 + KLF5 |
| 61 | SULF1 | 2.665452586 | 0.008904627 | 0.034252903 | 2.866133536 | 0.322283713 | Suprabasal | SOX2 + KLF5 |
| 62 | CAMK1D | 2.657917619 | 0.000506425 | 0.007178523 | 49.15873668 | 9.200779378 | Superficial | KLF5 |
| 63 | SHF | 2.642479077 | 0.001883197 | 0.010234018 | 10.17290233 | 1.181406638 | Superficial | SOX2 |
| 64 | APOL3 | 2.616122791 | 0.000317696 | 0.007176147 | 14.03510257 | 2.124512865 | Superficial | SOX2 |
| 65 | PLSCR3 | 2.590344397 | 9.30E-05 | 0.007176147 | 51.28244557 | 9.30799657 | Superficial | SOX2 |
| 66 | DPYD | 2.582534489 | 0.000250281 | 0.007176147 | 56.95813881 | 13.81974657 | Superficial | SOX2 |
| 67 | C1orf74 | 2.569639155 | 0.000635975 | 0.007274975 | 18.30073531 | 2.487355771 | Superficial | SOX2 |
| 68 | HS3ST3A1 | 2.549605795 | 0.003462059 | 0.014108599 | 16.86965958 | 2.495265241 | Superficial | SOX2 + KLF5 |
| 69 | ADAT3 | 2.540027253 | 0.000196466 | 0.007176147 | 10.05232351 | 1.179084409 | Superficial | SOX2 |
| 70 | RARB | 2.530013416 | 0.000992312 | 0.008042817 | 22.96760446 | 3.024920953 | Superficial | SOX2 + KLF5, SOX2 |
| 71 | TSPAN1 | 2.529823549 | 0.008904627 | 0.025990329 | 11.37029984 | 1.026495801 | Superficial | SOX2 |

| | | | | | | | | |
|---|---|---|---|---|---|---|---|---|
| 72 | GALNT5 | 2.527879018 | 0.000153669 | 0.007176147 | 62.61494461 | 14.19470699 | Superficial | SOX2, KLF5 |
| 73 | HPGD | 2.517803476 | 9.30E-05 | 0.012455402 | 63.4370818 | 26.66092629 | Basal | SOX2 |
| 74 | NEK6 | 2.497806445 | 0.000196466 | 0.010489143 | 9.695906506 | 1.247626474 | Suprabasal | SOX2 |
| 75 | USP2 | 2.470920442 | 7.20E-05 | 0.010489143 | 20.35654798 | 3.795647067 | Suprabasal | SOX2 |
| 76 | CDH3 | 2.454629294 | 0.001883197 | 0.014679007 | 33.06720283 | 5.979505217 | Suprabasal | SOX2 |
| 77 | NRP2 | 2.4318478 | 0.000635975 | 0.010733065 | 10.71042608 | 1.788625 | Suprabasal | SOX2, SOX2 + KLF5 |
| 78 | CLDN1 | 2.418800238 | 0.000506425 | 0.007178523 | 59.84833475 | 16.1506835 | Superficial | SOX2, SOX2 + KLF5 |
| 79 | EHD2 | 2.365985362 | 0.000992312 | 0.008042817 | 22.15175338 | 3.977983815 | Superficial | SOX2 |
| 80 | EPHA4 | 2.363866254 | 0.002315114 | 0.011381143 | 23.97424291 | 3.646350234 | Superficial | SOX2, KLF5 |
| 81 | ATP11C | 2.357061891 | 0.006166656 | 0.020290909 | 9.252597911 | 1.160883516 | Superficial | SOX2 |
| 82 | TMTC3 | 2.351630508 | 0.000635975 | 0.007274975 | 45.10321587 | 9.259139716 | Superficial | SOX2 |
| 83 | VDR | 2.343310449 | 0.000506425 | 0.007178523 | 46.65513349 | 9.496180257 | Superficial | SOX2 |
| 84 | ILDR1 | 2.3418189 | 0.000635975 | 0.007274975 | 10.28288777 | 1.988042993 | Superficial | SOX2 |
| 85 | FZD10 | 2.335435136 | 0.000196466 | 0.007176147 | 38.87791107 | 6.491347125 | Superficial | SOX2 |
| 86 | GJB4 | 2.297127992 | 0.001883197 | 0.010234018 | 14.24072294 | 2.556640803 | Superficial | SOX2 |
| 87 | SOX21 | 2.296941678 | 0.000506425 | 0.007178523 | 41.19240615 | 8.728487216 | Superficial | SOX2 + KLF5, SOX2 |
| 88 | SERINC5 | 2.292352434 | 0.000992312 | 0.008042817 | 57.34016553 | 13.89975778 | Superficial | SOX2 |
| 89 | APOL6 | 2.290382729 | 0.000401828 | 0.007176147 | 47.34837516 | 9.465108027 | Superficial | SOX2 |
| 90 | SH3RF2 | 2.288890426 | 0.000401828 | 0.007176147 | 57.94436358 | 14.02325258 | Superficial | KLF5, SOX2 |
| 91 | LGALSL | 2.268271415 | 0.001526458 | 0.009387421 | 55.59902515 | 18.58782962 | Superficial | SOX2 |
| 92 | AADAC | 2.267827947 | 0.005105016 | 0.018023407 | 10.87524615 | 1.59110865 | Superficial | SOX2 |
| 93 | SGPP2 | 2.250143174 | 0.000119764 | 0.007176147 | 52.07167939 | 12.31623848 | Superficial | SOX2 |
| 94 | PDZK1IP1 | 2.242093901 | 0.001526458 | 0.016792744 | 64.0916668 | 22.57691274 | Basal | SOX2 |
| 95 | GGH | 2.23883892 | 0.000317696 | 0.007176147 | 60.15013769 | 13.53596753 | Superficial | SOX2, SOX2 + KLF5 |
| 96 | PKP2 | 2.222886641 | 0.000795824 | 0.007615337 | 26.63168494 | 2.617248693 | Superficial | SOX2 |
| 97 | UTRN | 2.217242242 | 0.000196466 | 0.007176147 | 50.01075484 | 10.51356248 | Superficial | SOX2, SOX2 + KLF5 |
| 98 | NUP210 | 2.213606148 | 0.000795824 | 0.010838271 | 9.800731136 | 2.142050625 | Suprabasal | SOX2 |
| 99 | GCNT2 | 2.205203227 | 0.000317696 | 0.007176147 | 24.98548601 | 4.185120601 | Superficial | SOX2 |
| 100 | TSPAN3 | 2.193440752 | 0.000250281 | 0.007176147 | 72.63979106 | 28.82128249 | Superficial | SOX2 |
| 101 | MYOF | 2.181332839 | 0.000795824 | 0.007615337 | 51.76574142 | 12.51822189 | Superficial | SOX2 + KLF5 |
| 102 | DAPK2 | 2.170784779 | 0.001232925 | 0.012246384 | 11.93187041 | 2.113989461 | Suprabasal | SOX2 |
| 103 | CEP72 | 2.167186654 | 0.000196466 | 0.010489143 | 5.579343935 | 1.139148869 | Suprabasal | SOX2 |
| 104 | GALR2 | 2.166969424 | 0.001232925 | 0.008679903 | 13.57405301 | 2.436665487 | Superficial | SOX2 |
| 105 | AMOTL1 | 2.165279249 | 0.000153669 | 0.007176147 | 53.80914283 | 12.90952452 | Superficial | SOX2 |
| 106 | FAM83E | 2.154161807 | 0.001526458 | 0.009387421 | 6.837893593 | 1.341925006 | Superficial | KLF5, SOX2 |
| 107 | HBEGF | 2.139981014 | 0.000401828 | 0.007176147 | 49.07344726 | 14.25840594 | Superficial | SOX2 |
| 108 | GIPC2 | 2.130796901 | 0.000119764 | 0.007176147 | 11.29330817 | 1.737078975 | Superficial | SOX2, SOX2 + KLF5 |
| 109 | PIK3CB | 2.116387008 | 0.000317696 | 0.007176147 | 30.49497273 | 5.37065142 | Superficial | SOX2 |
| 110 | FNTB | 2.113774756 | 0.000196466 | 0.007176147 | 38.35216549 | 9.331467725 | Superficial | SOX2 |
| 111 | CALML4 | 2.098383243 | 0.006116011 | 0.020290909 | 16.40768279 | 3.452763393 | Superficial | SOX2 |
| 112 | PPP2R5B | 2.096347506 | 0.000196466 | 0.007176147 | 8.853189121 | 1.585170099 | Superficial | SOX2 |
| 113 | RAI1 | 2.083551412 | 0.000401828 | 0.007176147 | 19.62176647 | 3.030777351 | Superficial | SOX2 |
| 114 | SOX2 | 2.081913146 | 0.000401828 | 0.007176147 | 64.78270526 | 23.45059511 | Superficial | SOX2, KLF5 |
| 115 | JDP2 | 2.080656788 | 0.000119764 | 0.007176147 | 38.95069837 | 7.653122801 | Superficial | SOX2 |
| 116 | KLHL5 | 2.07040443 | 0.001883197 | 0.014679007 | 18.03936572 | 3.182826682 | Suprabasal | SOX2 |
| 117 | PADI3 | 2.067623508 | 0.006166656 | 0.032981404 | 5.285130858 | 0.790356966 | Basal | SOX2 |
| 118 | HS3ST1 | 2.063319572 | 0.001232925 | 0.015721789 | 23.50845249 | 5.330860422 | Basal | SOX2, SOX2 + KLF5 |
| 119 | ARHGAP23 | 2.062876631 | 0.000992312 | 0.008042817 | 43.87809247 | 10.02607057 | Superficial | SOX2 |
| 120 | HECW2 | 2.058221439 | 9.30E-05 | 0.007176147 | 15.71587344 | 3.116907862 | Superficial | SOX2 |
| 121 | PRICKLE2 | 2.055478112 | 0.000795824 | 0.010838271 | 25.65292316 | 5.691109794 | Suprabasal | SOX2 |
| 122 | C9orf152 | 2.042243894 | 0.000317696 | 0.007176147 | 13.25621132 | 2.381023333 | Superficial | SOX2 |
| 123 | F2RL1 | 2.04202804 | 0.000795824 | 0.007615337 | 15.22355279 | 3.090463749 | Superficial | SOX2 |
| 124 | STK17B | 2.02769262 | 0.001526458 | 0.009387421 | 41.63652012 | 7.333922578 | Superficial | SOX2 |
| 125 | C9orf40 | 2.018582283 | 0.000401828 | 0.007176147 | 18.20397196 | 3.43412846 | Superficial | SOX2 |
| 126 | MYH9 | 2.016181739 | 0.000401828 | 0.007176147 | 61.10773592 | 18.88556925 | Superficial | SOX2 + KLF5, SOX2 |
| 127 | PLAUR | 2.002795365 | 0.002836074 | 0.012662188 | 15.93572912 | 3.786355668 | Superficial | SOX2 |
| 128 | CD44 | 1.998034871 | 0.000506425 | 0.007178523 | 61.11347514 | 19.65892181 | Superficial | SOX2 |
| 129 | PPARG | 1.995811597 | 0.000401828 | 0.007176147 | 27.13150116 | 5.448962677 | Superficial | SOX2, KLF5 |
| 130 | PDLIM4 | 1.989624172 | 0.000317696 | 0.007176147 | 69.2066852 | 25.82174452 | Superficial | SOX2, SOX2 + KLF5 |
| 131 | C16orf74 | 1.983715146 | 0.005105016 | 0.018023407 | 23.21339782 | 4.600880205 | Superficial | SOX2 |
| 132 | NUAK2 | 1.98293209 | 0.001232925 | 0.008679903 | 43.77931763 | 10.90822545 | Superficial | SOX2 + KLF5 |
| 133 | KLK7 | 1.977463828 | 0.001883197 | 0.010234018 | 66.73816902 | 34.01620165 | Superficial | SOX2 |
| 134 | LPCAT4 | 1.972938051 | 0.000795824 | 0.007615337 | 62.90111134 | 20.64436002 | Superficial | SOX2 + KLF5, SOX2 |
| 135 | RIPK2 | 1.966802897 | 0.000795824 | 0.007615337 | 24.08933789 | 4.963363477 | Superficial | SOX2 |
| 136 | VANGL2 | 1.960523499 | 0.005105016 | 0.018023407 | 9.102787321 | 1.461988544 | Superficial | SOX2 |
| 137 | KCTD12 | 1.956658379 | 0.000401828 | 0.010489143 | 15.00195718 | 2.947502003 | Suprabasal | SOX2 |
| 138 | PLXNA1 | 1.956245434 | 0.002315114 | 0.011381143 | 13.16078784 | 2.350686874 | Superficial | SOX2 |
| 139 | UBE2Z | 1.951186499 | 0.000506425 | 0.007178523 | 50.65988193 | 13.56189302 | Superficial | SOX2 |
| 140 | LRRC8B | 1.948757418 | 0.001526458 | 0.009387421 | 14.99469118 | 2.794121962 | Superficial | SOX2 + KLF5 |
| 141 | DUSP10 | 1.946963734 | 0.000992312 | 0.01142308 | 16.06853386 | 3.224143338 | Suprabasal | SOX2 |
| 142 | MVP | 1.941839212 | 0.000401828 | 0.007176147 | 51.13459459 | 14.41912597 | Superficial | SOX2 |
| 143 | TLE1 | 1.923101892 | 0.000119764 | 0.007176147 | 27.10808303 | 5.539944391 | Superficial | SOX2, SOX2 + KLF5 |
| 144 | CLMN | 1.900395402 | 0.000401828 | 0.010489143 | 10.25851781 | 1.920383559 | Suprabasal | SOX2 |

| | | | | | | | |
|---|---|---|---|---|---|---|---|
| 145 | BTBD11 | 1.881944659 | 0.000401828 | 0.007176147 | 32.29979612 | 7.660902713 | Superficial | SOX2 |
| 146 | UCK2 | 1.879818849 | 0.000506425 | 0.007178523 | 25.81261694 | 5.843574533 | Superficial | SOX2 |
| 147 | PRICKLE3 | 1.874661139 | 0.001232925 | 0.008679903 | 19.11973397 | 4.835371474 | Superficial | SOX2 |
| 148 | SGMS2 | 1.870786282 | 0.000401828 | 0.007176147 | 35.5703333 | 7.638278866 | Superficial | SOX2 + KLF5, SOX2 |
| 149 | FMN1 | 1.866940741 | 0.000992312 | 0.008042817 | 13.93481253 | 2.490917074 | Superficial | SOX2 + KLF5, SOX2 |
| 150 | MBOAT1 | 1.856378922 | 0.004211401 | 0.016013117 | 20.12833534 | 4.645925894 | Superficial | SOX2 |
| 151 | PLCXD2 | 1.855577703 | 0.001883197 | 0.018415354 | 10.25091971 | 1.496509635 | Basal | SOX2 |
| 152 | SLFN5 | 1.848849547 | 0.001232925 | 0.008679903 | 27.75619287 | 6.018649947 | Superficial | SOX2 |
| 153 | TRAK1 | 1.846069152 | 0.000401828 | 0.007176147 | 35.07851392 | 7.922600535 | Superficial | SOX2 |
| 154 | PLAG1 | 1.834621629 | 0.002315114 | 0.020052792 | 7.541644707 | 1.172226914 | Basal | SOX2 |
| 155 | PCGF2 | 1.829922147 | 0.000401828 | 0.007176147 | 18.9966338 | 3.843979558 | Superficial | SOX2 |
| 156 | CFH | 1.828796647 | 0.002315114 | 0.011381143 | 15.10224119 | 2.619784914 | Superficial | SOX2 |
| 157 | BPNT1 | 1.821261596 | 0.000119764 | 0.007176147 | 23.50791508 | 5.202532602 | Superficial | SOX2 |
| 158 | SERPINB5 | 1.819859014 | 0.000506425 | 0.007178523 | 67.98078257 | 28.24706043 | Superficial | SOX2, SOX2 + KLF5 |
| 159 | VPS9D1 | 1.8139499 | 0.004211401 | 0.016013117 | 32.5671341 | 9.788833771 | Superficial | SOX2 |
| 160 | PARP14 | 1.810369034 | 0.005105016 | 0.018023407 | 38.26737244 | 7.373411595 | Superficial | SOX2 |
| 161 | PPFIA3 | 1.808834433 | 0.000506425 | 0.007178523 | 14.42758209 | 3.603490342 | Superficial | SOX2 |
| 162 | HAPLN3 | 1.805854094 | 0.017811616 | 0.043066688 | 9.09934586 | 1.590747789 | Superficial | SOX2 |
| 163 | SLC26A2 | 1.800102188 | 0.001526458 | 0.009387421 | 45.30830537 | 11.39121798 | Superficial | SOX2 |
| 164 | RHBDF2 | 1.7980639 | 0.000317696 | 0.007176147 | 22.85167975 | 5.481160876 | Superficial | SOX2 |
| 165 | EPB41L1 | 1.793989519 | 0.001232925 | 0.008679903 | 42.29613711 | 10.90537302 | Superficial | SOX2 |
| 166 | LZTS2 | 1.792350541 | 0.000795824 | 0.007615337 | 21.57360734 | 4.784089227 | Superficial | SOX2 |
| 167 | DUOX1 | 1.791874816 | 0.000250281 | 0.007176147 | 78.09045313 | 40.85380185 | Superficial | SOX2 |
| 168 | RGS2 | 1.791283961 | 0.012681065 | 0.033478436 | 25.73956524 | 5.400301716 | Superficial | SOX2 |
| 169 | CDH1 | 1.790786868 | 0.000196466 | 0.007176147 | 66.43421356 | 23.36472372 | Superficial | SOX2 |
| 170 | FLNB | 1.789509468 | 0.000795824 | 0.007615337 | 41.12314191 | 11.05496167 | Superficial | SOX2 |
| 171 | ACSL5 | 1.783440086 | 0.006166656 | 0.027303057 | 11.74714312 | 2.411107733 | Suprabasal | SOX2 |
| 172 | EFNA3 | 1.780245834 | 0.000317696 | 0.007176147 | 27.72683451 | 6.791475493 | Superficial | SOX2 |
| 173 | MX1 | 1.775059796 | 0.000992312 | 0.008042817 | 52.23414703 | 14.14516091 | Superficial | SOX2 |
| 174 | HSPA4L | 1.774195024 | 0.000992312 | 0.008042817 | 44.30957301 | 11.98002916 | Superficial | SOX2 |
| 175 | NMI | 1.773761102 | 0.002315114 | 0.011381143 | 47.10510036 | 12.99264976 | Superficial | SOX2 |
| 176 | TMED8 | 1.76610339 | 0.000795824 | 0.007615337 | 9.847590848 | 1.882202548 | Superficial | SOX2 |
| 177 | ACTN1 | 1.757688537 | 0.000992312 | 0.008042817 | 49.01098628 | 14.15301096 | Superficial | SOX2 |
| 178 | ARHGAP26 | 1.751173085 | 7.20E-05 | 0.007176147 | 19.94818841 | 4.677088953 | Superficial | SOX2 + KLF5 |
| 179 | RAB40B | 1.750815018 | 0.002315114 | 0.011381143 | 29.68013769 | 6.615881291 | Superficial | SOX2 |
| 180 | LITAF | 1.746481223 | 0.000317696 | 0.010489143 | 65.22896326 | 21.81166448 | Suprabasal | SOX2, SOX2 + KLF5 |
| 181 | BIK | 1.744369223 | 0.001526458 | 0.016792744 | 17.82374359 | 4.38678935 | Basal | SOX2 |
| 182 | LMTK3 | 1.740611516 | 0.000401828 | 0.007176147 | 31.40700303 | 7.955807442 | Superficial | SOX2 |
| 183 | PCP4L1 | 1.738642273 | 0.002315114 | 0.020052792 | 8.98214398 | 2.87134725 | Basal | KLF5 |
| 184 | TRIM34 | 1.734596734 | 0.002257606 | 0.011381143 | 12.00649374 | 2.23866282 | Superficial | SOX2 |
| 185 | DUSP7 | 1.73342227 | 0.005105016 | 0.018023407 | 26.21992767 | 6.095908265 | Superficial | SOX2 |
| 186 | IGF2R | 1.719942816 | 0.000992312 | 0.008042817 | 48.48639949 | 13.65241974 | Superficial | SOX2 + KLF5, SOX2 |
| 187 | UBR1 | 1.719522138 | 0.000196466 | 0.007176147 | 14.06178488 | 2.530448228 | Superficial | SOX2 |
| 188 | SMAD7 | 1.715565935 | 0.000317696 | 0.010489143 | 23.33039354 | 6.328498368 | Suprabasal | SOX2 |
| 189 | CFAP45 | 1.713651152 | 0.008904627 | 0.034252903 | 3.768521822 | 1.162924672 | Suprabasal | SOX2 |
| 190 | PSMB9 | 1.713579321 | 0.001883197 | 0.014679007 | 46.76968644 | 13.20609663 | Suprabasal | SOX2 |
| 191 | RAD50 | 1.709565758 | 0.000250281 | 0.007176147 | 41.71108638 | 10.58867033 | Superficial | SOX2 |
| 192 | ARFGEF1 | 1.709561526 | 5.55E-05 | 0.007176147 | 59.31506671 | 18.23150087 | Superficial | SOX2 |
| 193 | PCCA | 1.708003858 | 0.000250281 | 0.007176147 | 25.73152384 | 5.309542158 | Superficial | SOX2 |
| 194 | TNIK | 1.706346895 | 0.017811616 | 0.043066688 | 5.392242583 | 0.752506829 | Superficial | SOX2 |
| 195 | ITGAV | 1.702941913 | 0.000317696 | 0.007176147 | 28.00302136 | 5.643803668 | Superficial | SOX2 |
| 196 | NAB2 | 1.702282164 | 0.000401828 | 0.007176147 | 9.914356536 | 2.392261266 | Superficial | SOX2 |
| 197 | PITRM1 | 1.702119527 | 0.002315114 | 0.011381143 | 39.3035774 | 9.553439545 | Superficial | SOX2 |
| 198 | GALNT7 | 1.696244002 | 0.000317696 | 0.007176147 | 53.0088514 | 14.34898878 | Superficial | KLF5 |
| 199 | FAM3C | 1.6929743 | 0.000992312 | 0.008042817 | 62.57202139 | 21.71212462 | Superficial | SOX2, SOX2 + KLF5 |
| 200 | CARNMT1 | 1.69008525 | 0.000635975 | 0.007274975 | 27.23772045 | 6.517738001 | Superficial | SOX2 |
| 201 | PLXNB1 | 1.689316796 | 0.000635975 | 0.007274975 | 28.19070091 | 8.442735153 | Superficial | SOX2 |
| 202 | AQP3 | 1.688886054 | 0.000635975 | 0.007274975 | 88.79367883 | 56.05032049 | Superficial | SOX2 |
| 203 | MAOA | 1.687062072 | 0.000250281 | 0.007176147 | 53.15988498 | 16.42076969 | Superficial | SOX2 |
| 204 | PLEKHA7 | 1.686755148 | 0.000635975 | 0.007274975 | 50.68256995 | 15.01051921 | Superficial | SOX2 |
| 205 | BHLHE40 | 1.683975933 | 0.000250281 | 0.012455402 | 66.70715515 | 29.07714623 | Basal | SOX2 |
| 206 | TMEM268 | 1.683413946 | 0.000795824 | 0.007615337 | 6.305050499 | 1.013843833 | Superficial | SOX2 |
| 207 | RGS12 | 1.679974365 | 0.000196466 | 0.007176147 | 52.44813675 | 16.05946172 | Superficial | SOX2 |
| 208 | FAM3B | 1.679634197 | 0.000401828 | 0.007176147 | 76.76679222 | 41.20933182 | Superficial | SOX2, KLF5 |
| 209 | FAM83C | 1.679177088 | 0.000992312 | 0.008042817 | 34.67295623 | 9.58995339 | Superficial | SOX2 |
| 210 | USP54 | 1.677511187 | 0.001883197 | 0.010234018 | 48.22455818 | 13.1742308 | Superficial | SOX2 |
| 211 | NRARP | 1.676776951 | 0.001526458 | 0.009387421 | 59.48866715 | 22.43678013 | Superficial | SOX2 |
| 212 | VANGL1 | 1.674308055 | 0.000992312 | 0.008042817 | 26.754429 | 5.911651905 | Superficial | SOX2 |
| 213 | CD9 | 1.664880927 | 0.000317696 | 0.007176147 | 95.7959489 | 87.17960747 | Superficial | SOX2 + KLF5, SOX2 |
| 214 | SIPA1L1 | 1.662110852 | 9.30E-05 | 0.007176147 | 45.53935844 | 12.70121283 | Superficial | SOX2 |
| 215 | FZD1 | 1.660406189 | 0.000401828 | 0.007176147 | 15.36367518 | 3.652475248 | Superficial | SOX2 |
| 216 | DSG3 | 1.659685097 | 0.000401828 | 0.007176147 | 76.2731689 | 37.72281451 | Superficial | KLF5, SOX2 + KLF5, SOX2 |
| 217 | FAM83G | 1.655884408 | 0.001883197 | 0.010234018 | 35.56417675 | 10.87087784 | Superficial | SOX2 |

| | | | | | | | |
|---|---|---|---|---|---|---|---|
| 218 | ADCY7 | 1.655709863 | 0.002836074 | 0.012662188 | 26.2365789 | 5.391710904 | Superficial | SOX2 |
| 219 | MDGA1 | 1.649359333 | 0.005105016 | 0.024262694 | 6.424994564 | 1.914752811 | Suprabasal | SOX2 |
| 220 | CEACAM6 | 1.648410896 | 0.001232925 | 0.015721789 | 14.47583323 | 4.150146377 | Basal | SOX2 |
| 221 | ATP2C1 | 1.646785429 | 0.000401828 | 0.007176147 | 34.44258344 | 8.519262101 | Superficial | SOX2 |
| 222 | TCF7L2 | 1.645495297 | 0.000635975 | 0.007274975 | 24.10556753 | 5.483411759 | Superficial | SOX2 |
| 223 | DEF6 | 1.643050311 | 0.000401828 | 0.007176147 | 22.74032432 | 5.806535233 | Superficial | SOX2 |
| 224 | ERI1 | 1.638743032 | 0.000795824 | 0.007615337 | 19.59835376 | 5.244473791 | Superficial | SOX2 |
| 225 | E2F8 | 1.632929528 | 0.002836074 | 0.012662188 | 19.05196528 | 4.16204622 | Superficial | SOX2 |
| 226 | PRDM1 | 1.628753891 | 0.005105016 | 0.029792864 | 9.74035703 | 2.23802724 | Basal | SOX2 |
| 227 | WNT5A | 1.626161756 | 0.000401828 | 0.007176147 | 40.61458636 | 11.46950904 | Superficial | SOX2 |
| 228 | ACSL1 | 1.625384316 | 0.006166656 | 0.020290909 | 45.16007735 | 13.44840278 | Superficial | SOX2 |
| 229 | IFIT2 | 1.625274549 | 0.021001058 | 0.04834985 | 7.148106854 | 1.150788534 | Superficial | SOX2 |
| 230 | AEBP2 | 1.622302638 | 0.000795824 | 0.007615337 | 39.97302843 | 9.842466958 | Superficial | SOX2 |
| 231 | STX6 | 1.612517544 | 0.000795824 | 0.007615337 | 41.35112964 | 11.00713868 | Superficial | SOX2 |
| 232 | TEF | 1.60997414 | 0.007423146 | 0.030449933 | 15.90490035 | 4.406701305 | Suprabasal | SOX2 |
| 233 | ZNF326 | 1.607653042 | 0.000119764 | 0.007176147 | 58.61505907 | 18.15590786 | Superficial | SOX2 |
| 234 | CASK | 1.607623668 | 0.000317696 | 0.007176147 | 25.64028894 | 6.432160107 | Superficial | SOX2 |
| 235 | STAT6 | 1.605855949 | 0.000992312 | 0.008042817 | 35.45387829 | 11.31713013 | Superficial | SOX2 |
| 236 | NECTIN1 | 1.600704419 | 0.000992312 | 0.008042817 | 62.49026247 | 22.13619408 | Superficial | SOX2 |
| 237 | LDLRAD3 | 1.598177175 | 0.000795824 | 0.007615337 | 13.15101248 | 2.23742001 | Superficial | SOX2 |
| 238 | SIRT6 | 1.593566613 | 0.003462059 | 0.014108599 | 11.50078903 | 3.946469396 | Superficial | SOX2 |
| 239 | ADGRG1 | 1.588484691 | 0.002315114 | 0.011381143 | 40.98278712 | 11.67161753 | Superficial | SOX2 |
| 240 | MACF1 | 1.585172836 | 0.001883197 | 0.010234018 | 45.42867422 | 13.42301982 | Superficial | SOX2 + KLF5, SOX2 |
| 241 | KALRN | 1.568401192 | 0.001232925 | 0.008679903 | 39.8906692 | 11.36810286 | Superficial | SOX2 |
| 242 | EFNA1 | 1.567624298 | 0.000250281 | 0.007176147 | 35.81170057 | 9.972419373 | Superficial | SOX2 |
| 243 | SIAH2 | 1.561358934 | 0.000506425 | 0.007178523 | 54.29178907 | 18.17855221 | Superficial | SOX2 |
| 244 | MPP7 | 1.555620236 | 0.000795824 | 0.007615337 | 74.5401543 | 35.82324735 | Superficial | SOX2 |
| 245 | LRIG3 | 1.554041648 | 0.001526458 | 0.009387421 | 16.87745803 | 4.27268611 | Superficial | SOX2 |
| 246 | CBLC | 1.550539974 | 0.000795824 | 0.007615337 | 54.08040173 | 16.73185142 | Superficial | SOX2 |
| 247 | RAPGEFL1 | 1.546741477 | 0.000635975 | 0.007274975 | 59.0466891 | 21.22229912 | Superficial | KLF5, SOX2 |
| 248 | NBEAL2 | 1.543643779 | 0.001526458 | 0.009387421 | 66.32498761 | 27.02085762 | Superficial | SOX2 |
| 249 | TMPRSS11D | 1.540082196 | 0.008904627 | 0.040392504 | 18.5208653 | 5.78738771 | Basal | SOX2 + KLF5 |
| 250 | CYB561D1 | 1.539648021 | 0.003462059 | 0.014108599 | 15.63686224 | 3.624462484 | Superficial | SOX2 |
| 251 | DYRK2 | 1.537828927 | 0.000317696 | 0.007176147 | 30.61055724 | 7.976776919 | Superficial | SOX2 |
| 252 | EGR2 | 1.532135907 | 0.010644787 | 0.029507311 | 8.930218254 | 2.317618094 | Superficial | SOX2 |
| 253 | UNC5B | 1.532021274 | 0.004211401 | 0.016013117 | 21.54159413 | 7.012623892 | Superficial | SOX2 |
| 254 | EGR1 | 1.531257162 | 0.000401828 | 0.007176147 | 77.3750348 | 37.25352246 | Superficial | SOX2 |
| 255 | ID4 | 1.529712713 | 0.005105016 | 0.018023407 | 61.2478735 | 25.59836537 | Superficial | SOX2 |
| 256 | NRIP1 | 1.527774608 | 0.000992312 | 0.008042817 | 31.56908623 | 8.02702205 | Superficial | SOX2, SOX2 + KLF5 |
| 257 | SETBP1 | 1.526392337 | 0.000795824 | 0.007615337 | 12.37924636 | 2.847820355 | Superficial | SOX2 + KLF5, SOX2 |
| 258 | CTTNBP2 | 1.523952442 | 0.001526458 | 0.009387421 | 38.23349606 | 9.953505753 | Superficial | SOX2 |
| 259 | ZNF750 | 1.521825816 | 0.000635975 | 0.007274975 | 71.74142611 | 34.19483123 | Superficial | SOX2 |
| 260 | ROBO1 | 1.521395177 | 0.002315114 | 0.011381143 | 20.09483419 | 4.223508011 | Superficial | SOX2 |
| 261 | SERPINB2 | 1.520402251 | 0.001232925 | 0.015721789 | 56.47425191 | 25.15403273 | Basal | SOX2 + KLF5 |
| 262 | DLG5 | 1.520094924 | 0.000795824 | 0.010838271 | 31.97247699 | 10.87101714 | Suprabasal | SOX2 |
| 263 | USP32 | 1.513329317 | 0.000992312 | 0.008042817 | 27.57071909 | 6.844546538 | Superficial | SOX2 |
| 264 | MBTPS1 | 1.5075199 | 0.000992312 | 0.008042817 | 40.77920392 | 12.36148303 | Superficial | SOX2 |
| 265 | SPTBN2 | 1.507423391 | 0.000635975 | 0.007274975 | 45.72760689 | 15.53689929 | Superficial | SOX2 |
| 266 | MYH14 | 1.506036602 | 0.000635975 | 0.007274975 | 65.02307798 | 27.62556019 | Superficial | SOX2, SOX2 + KLF5 |
| 267 | TTC39B | 1.504724353 | 0.000153669 | 0.007176147 | 31.35300257 | 8.277748331 | Superficial | SOX2 |
| 268 | PPFIBP2 | 1.502913882 | 0.000506425 | 0.007178523 | 38.08840061 | 10.11643122 | Superficial | SOX2 |
| 269 | PARP9 | 1.497471188 | 0.002836074 | 0.012662188 | 45.0973443 | 13.68164257 | Superficial | SOX2 |
| 270 | MCC | 1.495164205 | 0.001232925 | 0.008679903 | 33.17844453 | 9.480764163 | Superficial | SOX2, SOX2 + KLF5 |
| 271 | SLC16A7 | 1.492659835 | 0.000317696 | 0.007176147 | 36.68869929 | 10.26325143 | Superficial | SOX2 + KLF5 |
| 272 | KIAA1522 | 1.491037323 | 0.000795824 | 0.007615337 | 57.95200293 | 21.15180697 | Superficial | SOX2 |
| 273 | CTF1 | 1.489903466 | 0.015054848 | 0.038009884 | 6.497786284 | 2.532560526 | Superficial | SOX2 |
| 274 | ZFYVE28 | 1.489529705 | 0.004211401 | 0.016013117 | 11.28814489 | 2.205883735 | Superficial | SOX2 |
| 275 | MARK2 | 1.485849189 | 0.000795824 | 0.007615337 | 64.26837362 | 27.57208954 | Superficial | SOX2 |
| 276 | ZNF512B | 1.482940443 | 0.000153669 | 0.007176147 | 14.48764198 | 3.998364131 | Superficial | SOX2 |
| 277 | EEA1 | 1.479639763 | 0.000196466 | 0.007176147 | 68.01721549 | 26.67149732 | Superficial | SOX2 + KLF5, SOX2 |
| 278 | ANG | 1.470912351 | 0.017811616 | 0.043066688 | 10.58303921 | 2.925742081 | Superficial | SOX2 |
| 279 | PSTPIP2 | 1.470147182 | 0.000153669 | 0.007176147 | 20.22059631 | 3.435694389 | Superficial | SOX2 |
| 280 | GLS | 1.46994638 | 0.000119764 | 0.007176147 | 24.94966084 | 6.436039892 | Superficial | SOX2 |
| 281 | ACADSB | 1.46729936 | 0.000635975 | 0.007274975 | 33.22753084 | 7.725227649 | Superficial | SOX2 |
| 282 | TAPBP | 1.462687605 | 0.002315114 | 0.011381143 | 60.46527377 | 23.25742706 | Superficial | SOX2 |
| 283 | TMC6 | 1.459446866 | 0.003462059 | 0.014108599 | 33.19438017 | 10.41223051 | Superficial | SOX2 |
| 284 | ATF3 | 1.4580875 | 0.003462059 | 0.014108599 | 64.8069383 | 31.24154827 | Superficial | SOX2 |
| 285 | GPAT4 | 1.454502053 | 0.001232925 | 0.008679903 | 16.79760858 | 4.655621659 | Superficial | SOX2 |
| 286 | P2RY2 | 1.454448709 | 0.001883197 | 0.010234018 | 26.87639549 | 8.337861884 | Superficial | SOX2 |
| 287 | NR2F2 | 1.453940982 | 0.000250281 | 0.007176147 | 7.515066166 | 1.627785245 | Superficial | SOX2 |
| 288 | KLF4 | 1.451647249 | 0.000317696 | 0.007176147 | 74.16657085 | 38.12955866 | Superficial | SOX2 + KLF5 |
| 289 | PPARGC1B | 1.451020759 | 0.010644787 | 0.029507311 | 17.5730362 | 4.146815669 | Superficial | KLF5 |
| 290 | MARK1 | 1.450576993 | 0.005105016 | 0.018023407 | 11.31410423 | 2.61341336 | Superficial | SOX2 |

| | | | | | | | |
|---|---|---|---|---|---|---|---|
| 291 | ACVR2A | 1.449181971 | 0.000795824 | 0.007615337 | 30.56926601 | 7.030736758 | Superficial | SOX2 |
| 292 | EFS | 1.443933494 | 0.001526458 | 0.009387421 | 29.99052364 | 9.591608224 | Superficial | SOX2 |
| 293 | FOSL1 | 1.44377816 | 0.012681065 | 0.049425568 | 19.99797835 | 6.879080718 | Basal | SOX2 |
| 294 | SOX9 | 1.443054728 | 0.003462059 | 0.024854712 | 24.37891667 | 6.330952656 | Basal | SOX2 |
| 295 | RAB11FIP1 | 1.442200578 | 0.000317696 | 0.012455402 | 44.59727837 | 19.1411243 | Basal | SOX2, SOX2 + KLF5 |
| 296 | EXT1 | 1.436512243 | 0.002836074 | 0.012662188 | 13.54263902 | 3.548045055 | Superficial | SOX2 |
| 297 | GBP3 | 1.435078234 | 0.002836074 | 0.012662188 | 39.4589241 | 11.77543464 | Superficial | SOX2 |
| 298 | AGPAT5 | 1.429902007 | 0.001883197 | 0.010234018 | 27.61419053 | 6.879942189 | Superficial | SOX2 |
| 299 | TJP2 | 1.428786705 | 0.000250281 | 0.012455402 | 37.63135572 | 12.37153651 | Basal | SOX2, SOX2 + KLF5 |
| 300 | MYO1B | 1.424682138 | 0.000795824 | 0.0136425 | 35.06249235 | 12.49455715 | Basal | SOX2 |
| 301 | WRN | 1.422447463 | 0.000317696 | 0.007176147 | 12.03213883 | 2.82000383 | Superficial | SOX2 |
| 302 | MTSS1 | 1.414208577 | 7.20E-05 | 0.007176147 | 56.05422812 | 18.62672861 | Superficial | SOX2 |
| 303 | RARA | 1.414049556 | 0.007423146 | 0.022959455 | 10.0578091 | 1.361551578 | Superficial | SOX2 |
| 304 | CELSR2 | 1.412827235 | 0.001883197 | 0.010234018 | 32.83846129 | 8.471091898 | Superficial | SOX2 |
| 305 | TMEM62 | 1.41211041 | 0.000401828 | 0.007176147 | 15.85865511 | 4.494080284 | Superficial | SOX2 |
| 306 | ATP1A1 | 1.410743248 | 0.002836074 | 0.012662188 | 54.54542011 | 19.58899334 | Superficial | SOX2 |
| 307 | KDM6B | 1.410464515 | 0.000635975 | 0.007274975 | 26.18538702 | 7.047815 | Superficial | SOX2 |
| 308 | AHR | 1.409902398 | 0.000196466 | 0.010489143 | 37.6195811 | 12.92727326 | Suprabasal | SOX2 |
| 309 | FAM83B | 1.409210634 | 0.005105016 | 0.018023407 | 24.44070299 | 5.677640141 | Superficial | SOX2, SOX2 + KLF5 |
| 310 | IL15RA | 1.405856308 | 0.000992312 | 0.01142308 | 8.983045649 | 2.947652454 | Suprabasal | SOX2 |
| 311 | TRNP1 | 1.39874506 | 0.001526458 | 0.016792744 | 26.25894725 | 8.679371806 | Basal | SOX2 |
| 312 | IGSF9 | 1.397302481 | 0.015054848 | 0.038009884 | 10.74871348 | 2.239511216 | Superficial | SOX2 |
| 313 | B3GALNT2 | 1.394665908 | 0.000317696 | 0.007176147 | 23.68419665 | 6.990842078 | Superficial | SOX2 |
| 314 | ANGPTL4 | 1.390767663 | 9.30E-05 | 0.012455402 | 16.31586531 | 4.670732675 | Basal | SOX2 + KLF5, SOX2 |
| 315 | GFOD1 | 1.385270232 | 0.001883197 | 0.010234018 | 23.04950039 | 5.817350747 | Superficial | SOX2 |
| 316 | TPPP | 1.384548287 | 0.004211401 | 0.021879278 | 16.77886912 | 4.832901743 | Suprabasal | SOX2 |
| 317 | DUOXA1 | 1.37848433 | 0.000401828 | 0.012455402 | 48.21228319 | 19.71710377 | Basal | SOX2 |
| 318 | EDC4 | 1.373595403 | 0.000980874 | 0.008042817 | 19.45990268 | 5.084122399 | Superficial | SOX2 |
| 319 | PRCP | 1.372815391 | 0.000506425 | 0.007178523 | 51.28852932 | 17.1700345 | Superficial | SOX2 |
| 320 | KDM4A | 1.368537175 | 0.000506425 | 0.007178523 | 21.96859569 | 4.670725141 | Superficial | SOX2 |
| 321 | AHCTF1 | 1.364426399 | 0.000992312 | 0.008042817 | 20.42186335 | 4.885164396 | Superficial | SOX2 |
| 322 | CCDC51 | 1.364414193 | 0.006166656 | 0.020290909 | 30.08544728 | 9.187453014 | Superficial | SOX2 |
| 323 | GBF1 | 1.363385147 | 0.000992312 | 0.008042817 | 37.52272583 | 12.18132941 | Superficial | SOX2 |
| 324 | CIB1 | 1.359374292 | 0.000635975 | 0.007274975 | 71.89402045 | 37.61930816 | Superficial | SOX2 |
| 325 | ZBTB40 | 1.357638761 | 0.001526458 | 0.009387421 | 9.441108813 | 1.982079465 | Superficial | SOX2 |
| 326 | PDK3 | 1.356669527 | 0.001232925 | 0.008679903 | 15.30966875 | 4.188927431 | Superficial | SOX2 |
| 327 | UBE2L6 | 1.353073987 | 0.001526458 | 0.009387421 | 34.95959257 | 8.921171972 | Superficial | SOX2 |
| 328 | ANXA8L1 | 1.352486799 | 0.000506425 | 0.007178523 | 31.13710693 | 8.787587644 | Superficial | SOX2 |
| 329 | ETS2 | 1.348438538 | 0.000506425 | 0.010489143 | 70.5467627 | 37.00163852 | Suprabasal | SOX2, SOX2 + KLF5 |
| 330 | ZNF316 | 1.346975294 | 0.000506425 | 0.007178523 | 14.80715013 | 4.977715972 | Superficial | SOX2 |
| 331 | PPFIA1 | 1.345698612 | 5.55E-05 | 0.007176147 | 34.56770634 | 9.912383496 | Superficial | SOX2 |
| 332 | NUP205 | 1.342293498 | 0.000401828 | 0.007176147 | 14.31714078 | 4.145322997 | Superficial | SOX2 |
| 333 | TMEM129 | 1.341297541 | 0.003462059 | 0.014108599 | 15.54081075 | 4.998039805 | Superficial | SOX2 |
| 334 | KRT6A | 1.338300874 | 0.007423146 | 0.036548425 | 70.83206676 | 49.5782978 | Basal | SOX2 |
| 335 | UTP20 | 1.338154756 | 0.003462059 | 0.014108599 | 13.15682885 | 2.679443984 | Superficial | SOX2 |
| 336 | CERS3 | 1.336695002 | 0.000992312 | 0.008042817 | 66.75354368 | 27.89605679 | Superficial | KLF5, SOX2 |
| 337 | PHF2 | 1.333360461 | 0.000506425 | 0.007178523 | 18.51205223 | 4.924088628 | Superficial | SOX2 |
| 338 | SOWAHB | 1.330616268 | 0.000506425 | 0.010489143 | 19.25278672 | 6.539010872 | Suprabasal | SOX2 |
| 339 | UBE2O | 1.32923008 | 0.012681065 | 0.033478436 | 10.25200313 | 3.090639077 | Superficial | SOX2 |
| 340 | SEMA4B | 1.328875176 | 0.005105016 | 0.018023407 | 40.34899141 | 14.00945595 | Superficial | SOX2 |
| 341 | MUC15 | 1.328370967 | 0.000506425 | 0.007178523 | 59.25483221 | 20.34713306 | Superficial | SOX2 |
| 342 | TSHZ1 | 1.32797418 | 0.000795824 | 0.007615337 | 17.38245968 | 4.734116887 | Superficial | SOX2 |
| 343 | SLC30A4 | 1.324891857 | 0.000992312 | 0.008042817 | 15.57983981 | 4.129145692 | Superficial | SOX2 |
| 344 | PLAC8 | 1.321845779 | 0.008904627 | 0.040392504 | 23.19185551 | 9.186900909 | Basal | SOX2 |
| 345 | PROS1 | 1.320692718 | 0.001232925 | 0.008679903 | 16.76337057 | 4.722148576 | Superficial | SOX2 |
| 346 | QRSL1 | 1.319743783 | 0.000196466 | 0.007176147 | 15.55650242 | 4.687149578 | Superficial | SOX2 |
| 347 | OCIAD2 | 1.31691942 | 0.000795824 | 0.007615337 | 50.38690675 | 15.63470735 | Superficial | SOX2 + KLF5 |
| 348 | MUC1 | 1.316339356 | 0.008904627 | 0.040392504 | 16.0412245 | 4.959969675 | Basal | KLF5, SOX2 |
| 349 | PAK6 | 1.316197272 | 0.005105016 | 0.018023407 | 17.81476293 | 5.584040472 | Superficial | SOX2 |
| 350 | RNASE4 | 1.314834402 | 0.005094082 | 0.018023407 | 12.16976102 | 3.108359674 | Superficial | SOX2 |
| 351 | ARHGAP18 | 1.313208605 | 0.001526458 | 0.009387421 | 17.67087429 | 4.49131787 | Superficial | SOX2, SOX2 + KLF5 |
| 352 | TOP2B | 1.312446793 | 0.000795824 | 0.007615337 | 39.86008267 | 11.05700902 | Superficial | SOX2 |
| 353 | MARVELD2 | 1.312375904 | 0.000317696 | 0.007176147 | 28.05964149 | 9.039323031 | Superficial | SOX2 |
| 354 | ERC1 | 1.308852525 | 0.000992312 | 0.008042817 | 39.6157528 | 12.99023703 | Superficial | SOX2 + KLF5, SOX2 |
| 355 | TMEM132A | 1.306731913 | 0.000250281 | 0.007176147 | 36.14513057 | 9.791551434 | Superficial | SOX2 |
| 356 | TMEM125 | 1.305306728 | 0.000635975 | 0.007274975 | 31.74072791 | 10.27266171 | Superficial | SOX2 |
| 357 | ATP2A2 | 1.30393959 | 0.002836074 | 0.012662188 | 67.33126411 | 29.68867078 | Superficial | SOX2 |
| 358 | GSTO1 | 1.30389326 | 0.002315114 | 0.011381143 | 58.6348769 | 23.03531844 | Superficial | KLF5 |
| 359 | SOX13 | 1.301876207 | 0.004211401 | 0.016013117 | 13.72289931 | 3.118639101 | Superficial | SOX2 + KLF5, SOX2 |
| 360 | RCC1L | 1.30181624 | 0.000795824 | 0.007615337 | 28.93228558 | 8.302303983 | Superficial | SOX2 |
| 361 | PI4K2B | 1.301679821 | 0.004211401 | 0.016013117 | 19.88158248 | 4.693846417 | Superficial | SOX2 |
| 362 | NBN | 1.293606262 | 0.000317696 | 0.007176147 | 40.2244907 | 12.58433084 | Superficial | SOX2 |
| 363 | EPHA1 | 1.292228293 | 0.000795824 | 0.007615337 | 47.17389356 | 15.8337107 | Superficial | SOX2 |

| | | | | | | | |
|---|---|---|---|---|---|---|---|
| 364 | CBX6 | 1.29167696 | 0.002315114 | 0.011381143 | 31.83148599 | 10.1731285 | Superficial | SOX2 |
| 365 | TMPRSS4 | 1.291648313 | 0.000317696 | 0.012455402 | 52.64854085 | 22.60403832 | Basal | SOX2, KLF5, SOX2 + KLF5 |
| 366 | PDE4D | 1.288243 | 0.003462059 | 0.014108599 | 11.39595236 | 2.129328554 | Superficial | SOX2, SOX2 + KLF5 |
| 367 | DBNDD2 | 1.286934243 | 0.001219203 | 0.008679903 | 49.89250831 | 17.33137271 | Superficial | SOX2 |
| 368 | IRX2 | 1.284964767 | 5.55E-05 | 0.007176147 | 13.28747174 | 3.955746035 | Superficial | SOX2 |
| 369 | PTPRF | 1.283332755 | 0.001883197 | 0.010234018 | 54.86782032 | 19.86110899 | Superficial | SOX2 |
| 370 | RNGTT | 1.281199467 | 0.000250281 | 0.007176147 | 27.16286848 | 7.841618321 | Superficial | SOX2 |
| 371 | PLA2G4B | 1.280849172 | 0.017698983 | 0.043066688 | 29.65748348 | 11.18833209 | Superficial | SOX2 |
| 372 | CDC42EP1 | 1.280809566 | 0.000506425 | 0.012898798 | 49.93940725 | 20.79124099 | Basal | SOX2 |
| 373 | ARL5B | 1.2806538 | 0.002315114 | 0.011381143 | 38.32525881 | 10.54936115 | Superficial | SOX2 |
| 374 | FAM135A | 1.278999435 | 0.000992312 | 0.008042817 | 63.70652239 | 28.27070091 | Superficial | SOX2 |
| 375 | RAPGEF5 | 1.278312014 | 0.000153669 | 0.010489143 | 38.66857075 | 15.89877526 | Suprabasal | SOX2, SOX2 + KLF5 |
| 376 | PAK4 | 1.278029191 | 0.000153669 | 0.007176147 | 36.06503889 | 11.77443905 | Superficial | SOX2 |
| 377 | SPRYD3 | 1.277772516 | 0.000992312 | 0.008042817 | 24.20572055 | 6.994672203 | Superficial | SOX2 |
| 378 | FGFR3 | 1.277577146 | 0.004211401 | 0.016013117 | 43.50353723 | 15.34914819 | Superficial | SOX2, KLF5 |
| 379 | LSR | 1.271662219 | 0.001526458 | 0.009387421 | 59.20009736 | 22.67903464 | Superficial | SOX2 |
| 380 | PARG | 1.267231595 | 0.000153669 | 0.007176147 | 18.63111661 | 5.046069019 | Superficial | SOX2 |
| 381 | GSDMC | 1.265904005 | 0.001526458 | 0.009387421 | 51.45721865 | 18.99913397 | Superficial | KLF5 |
| 382 | RANGAP1 | 1.262746573 | 0.001883197 | 0.010234018 | 39.92014309 | 12.52755039 | Superficial | SOX2 |
| 383 | CTSD | 1.261128396 | 0.000401828 | 0.007176147 | 87.81296053 | 60.78044895 | Superficial | SOX2 |
| 384 | KDM1A | 1.259944271 | 0.001883197 | 0.010234018 | 24.42107785 | 6.395816716 | Superficial | SOX2 |
| 385 | SHROOM2 | 1.258271405 | 0.000635975 | 0.007274975 | 16.76833305 | 5.01205908 | Superficial | SOX2 + KLF5 |
| 386 | PTPN1 | 1.25765891 | 0.000506425 | 0.007178523 | 39.88181934 | 12.6414766 | Superficial | SOX2 |
| 387 | MAP7D1 | 1.256968222 | 0.000795824 | 0.007615337 | 56.70640296 | 22.14215924 | Superficial | SOX2 |
| 388 | FSCN1 | 1.256104721 | 0.001883197 | 0.010234018 | 24.7998747 | 6.980756334 | Superficial | SOX2 |
| 389 | MID1 | 1.254097172 | 0.006166656 | 0.020290909 | 15.17176327 | 4.455197887 | Superficial | SOX2 |
| 390 | LIN7C | 1.25095532 | 0.005105016 | 0.018023407 | 43.72459786 | 13.99481155 | Superficial | SOX2 |
| 391 | AP1S3 | 1.248619366 | 0.002836074 | 0.012662188 | 11.99405619 | 3.409620745 | Superficial | SOX2 |
| 392 | SP110 | 1.247917051 | 0.002315114 | 0.011381143 | 19.79106955 | 6.139409573 | Superficial | SOX2, KLF5 |
| 393 | DTX3L | 1.24559442 | 0.000795824 | 0.010838271 | 38.57229673 | 15.76772911 | Suprabasal | SOX2 |
| 394 | DAB2IP | 1.245422489 | 0.001232925 | 0.012246384 | 23.98713205 | 8.994272101 | Suprabasal | SOX2 |
| 395 | METRNL | 1.242786049 | 0.000317696 | 0.012455402 | 32.07582811 | 13.26662833 | Basal | SOX2 |
| 396 | FBLIM1 | 1.23859892 | 0.000992312 | 0.008042817 | 14.55913436 | 3.715759949 | Superficial | SOX2 |
| 397 | CDK5R1 | 1.23645138 | 0.000635975 | 0.007274975 | 13.61044702 | 4.337751879 | Superficial | SOX2 |
| 398 | BDP1 | 1.231393276 | 0.000250281 | 0.007176147 | 39.56516605 | 11.52677018 | Superficial | SOX2 |
| 399 | SMARCA5 | 1.229865827 | 0.000153669 | 0.007176147 | 44.63132421 | 14.69604831 | Superficial | SOX2 |
| 400 | ARHGEF1 | 1.222985993 | 0.002315114 | 0.011381143 | 18.69026155 | 5.468147191 | Superficial | SOX2 |
| 401 | ANO9 | 1.222957452 | 0.006166656 | 0.020290909 | 11.72755418 | 4.158654321 | Superficial | KLF5, SOX2 |
| 402 | TMEM30B | 1.222077703 | 0.001232925 | 0.008679903 | 42.12799768 | 12.74691984 | Superficial | SOX2 |
| 403 | EVL | 1.221452066 | 0.003462059 | 0.014108599 | 18.78014165 | 5.231162739 | Superficial | SOX2 |
| 404 | BTC | 1.219656775 | 0.001883197 | 0.010234018 | 23.18820738 | 7.421255007 | Superficial | SOX2 |
| 405 | HYOU1 | 1.21822226 | 0.003462059 | 0.014108599 | 28.78499902 | 11.00331255 | Superficial | SOX2 |
| 406 | MREG | 1.217754559 | 0.000401828 | 0.012455402 | 27.45165405 | 10.06667687 | Basal | SOX2 + KLF5, SOX2 |
| 407 | TFPI | 1.217749911 | 0.008904627 | 0.040392504 | 20.46071555 | 6.988812663 | Basal | SOX2 |
| 408 | TFCP2 | 1.214854752 | 0.001883197 | 0.010234018 | 14.82542175 | 4.280428251 | Superficial | SOX2 |
| 409 | NOTCH3 | 1.213496016 | 0.004211401 | 0.016013117 | 41.33700568 | 15.82539899 | Superficial | SOX2 |
| 410 | MBOAT2 | 1.21344202 | 0.000795824 | 0.0136425 | 18.88332959 | 7.348331732 | Basal | SOX2 + KLF5 |
| 411 | RGMA | 1.21235656 | 0.000795824 | 0.007615337 | 26.3261414 | 8.690737267 | Superficial | SOX2 |
| 412 | ROCK2 | 1.20861768 | 0.001232925 | 0.008679903 | 31.89008375 | 9.583988437 | Superficial | SOX2 + KLF5 |
| 413 | IL27RA | 1.208220342 | 0.000795824 | 0.010838271 | 9.158256842 | 3.247951193 | Suprabasal | SOX2 |
| 414 | STK11IP | 1.207708762 | 0.002836074 | 0.012662188 | 9.584264857 | 3.029777569 | Superficial | SOX2 |
| 415 | PPIP5K1 | 1.206931468 | 0.000317696 | 0.007176147 | 8.678543046 | 2.470109768 | Superficial | SOX2 |
| 416 | MXRA7 | 1.20590904 | 0.006166656 | 0.020290909 | 30.70137793 | 7.885702193 | Superficial | SOX2 |
| 417 | LIMA1 | 1.205072658 | 0.001232925 | 0.008679903 | 57.36614584 | 22.15407701 | Superficial | SOX2, SOX2 + KLF5 |
| 418 | P4HA2 | 1.203825583 | 0.002315114 | 0.011381143 | 29.25974961 | 9.077935764 | Superficial | SOX2 |
| 419 | SLC25A24 | 1.202476433 | 0.002836074 | 0.012662188 | 31.87004604 | 9.835333136 | Superficial | SOX2 + KLF5 |
| 420 | E2F2 | 1.202226402 | 0.000401828 | 0.012455402 | 25.69220688 | 10.11018419 | Basal | SOX2 |
| 421 | NCS1 | 1.199970243 | 0.010644787 | 0.029507311 | 18.84944464 | 5.393233849 | Superficial | SOX2 |
| 422 | ABCC5 | 1.199556541 | 0.000506425 | 0.010489143 | 47.79352163 | 22.35662642 | Suprabasal | SOX2 |
| 423 | VSNL1 | 1.198717733 | 0.002836074 | 0.012662188 | 41.90794306 | 13.72784784 | Superficial | SOX2 |
| 424 | MUC20 | 1.197371809 | 0.006166656 | 0.032981404 | 17.33627206 | 6.009769404 | Basal | SOX2, KLF5 |
| 425 | NIPAL4 | 1.196951548 | 0.015054848 | 0.038009884 | 5.624375738 | 1.930404675 | Superficial | SOX2 |
| 426 | CYP4F12 | 1.196520524 | 0.010644787 | 0.029507311 | 28.25141081 | 8.950303583 | Superficial | SOX2 |
| 427 | FHDC1 | 1.196319661 | 0.006166656 | 0.020290909 | 23.99616288 | 7.021853973 | Superficial | SOX2 |
| 428 | SCNN1A | 1.194828355 | 0.000992312 | 0.014320968 | 27.93955498 | 11.20588616 | Basal | SOX2, KLF5 |
| 429 | ATP8A1 | 1.192565378 | 0.000635975 | 0.010733065 | 6.400921197 | 2.31669173 | Suprabasal | SOX2 + KLF5 |
| 430 | MYO10 | 1.19168863 | 0.000635975 | 0.007274975 | 35.09234401 | 9.749988388 | Superficial | SOX2, SOX2 + KLF5 |
| 431 | AFF1 | 1.191217771 | 0.001526458 | 0.009387421 | 36.90037795 | 11.39818151 | Superficial | SOX2, SOX2 + KLF5 |
| 432 | MYCBP2 | 1.190959105 | 0.001232925 | 0.008679903 | 58.42803821 | 21.11847344 | Superficial | SOX2 |
| 433 | GCNT3 | 1.190903852 | 0.002315114 | 0.011381143 | 68.37598759 | 34.32034018 | Superficial | SOX2, SOX2 + KLF5 |
| 434 | ANXA4 | 1.188849498 | 0.000635975 | 0.007274975 | 55.84232817 | 19.17807815 | Superficial | SOX2 |
| 435 | MLLT3 | 1.18449493 | 0.002836074 | 0.012662188 | 15.62951055 | 4.585014268 | Superficial | SOX2, SOX2 + KLF5 |
| 436 | PTGS1 | 1.18264683 | 0.010644787 | 0.029507311 | 20.59203145 | 5.959216272 | Superficial | SOX2 |

| | | | | | | | | |
|---|---|---|---|---|---|---|---|---|
| 437 | DLG3 | 1.181702408 | 0.000506425 | 0.007178523 | 53.66370214 | 20.84782301 | Superficial | SOX2 |
| 438 | CXCL16 | 1.179795921 | 0.001883197 | 0.010234018 | 46.79771293 | 17.26782239 | Superficial | SOX2 |
| 439 | ATP11B | 1.178194803 | 0.000635975 | 0.007274975 | 65.46139314 | 27.9797841 | Superficial | SOX2 |
| 440 | CACFD1 | 1.177820059 | 0.002315114 | 0.011381143 | 14.49874791 | 4.530803054 | Superficial | SOX2 |
| 441 | MEIS1 | 1.17701975 | 0.000992312 | 0.008042817 | 34.91496587 | 11.38435718 | Superficial | SOX2 + KLF5 |
| 442 | PBX2 | 1.176236683 | 0.000153669 | 0.007176147 | 30.22501723 | 8.896086592 | Superficial | SOX2 |
| 443 | NSUN2 | 1.17513159 | 0.004211401 | 0.016013117 | 18.99430485 | 5.369107729 | Superficial | SOX2 |
| 444 | FUBP3 | 1.172413423 | 9.30E-05 | 0.007176147 | 27.44845694 | 8.374105466 | Superficial | SOX2 |
| 445 | ATP8B1 | 1.17101992 | 0.000401828 | 0.012455402 | 39.21041059 | 15.67056765 | Basal | SOX2 + KLF5 |
| 446 | PARP12 | 1.170055421 | 0.000635975 | 0.012925379 | 15.58978688 | 5.126299574 | Basal | SOX2, SOX2 + KLF5 |
| 447 | GBA2 | 1.169999924 | 0.005105016 | 0.018023407 | 14.65856801 | 4.196633117 | Superficial | SOX2 |
| 448 | EPHA2 | 1.169962509 | 0.004211401 | 0.027185186 | 30.15484264 | 11.81886947 | Basal | SOX2 |
| 449 | ZNF700 | 1.167879082 | 0.000196466 | 0.007176147 | 12.40076197 | 3.881582813 | Superficial | SOX2 |
| 450 | VPS54 | 1.167549799 | 0.004211401 | 0.016013117 | 17.16867299 | 4.649827082 | Superficial | SOX2 |
| 451 | LMBRD2 | 1.162448387 | 0.007423146 | 0.022959455 | 15.96772823 | 4.108028138 | Superficial | SOX2 |
| 452 | LRRC59 | 1.161487916 | 0.000635975 | 0.007274975 | 42.52666787 | 14.53896465 | Superficial | SOX2 |
| 453 | FASTKD3 | 1.16065057 | 0.017811616 | 0.043066688 | 10.31174552 | 3.498409326 | Superficial | SOX2 |
| 454 | TATDN2 | 1.159747117 | 0.001883197 | 0.010234018 | 15.92499475 | 5.709440714 | Superficial | SOX2 |
| 455 | ZNF827 | 1.158149884 | 0.010644787 | 0.029507311 | 6.442233256 | 1.5745672 | Superficial | SOX2 |
| 456 | ENDOD1 | 1.157892731 | 0.001526458 | 0.009387421 | 35.09992903 | 11.74305729 | Superficial | SOX2 |
| 457 | SLC30A7 | 1.157881703 | 0.006166656 | 0.020290909 | 13.96798379 | 3.84775842 | Superficial | SOX2 |
| 458 | XBP1 | 1.155508829 | 0.003462059 | 0.014108599 | 57.3569173 | 22.63957265 | Superficial | SOX2 |
| 459 | STK26 | 1.15468951 | 0.000992312 | 0.008042817 | 31.7416801 | 8.213308058 | Superficial | SOX2 |
| 460 | BARX2 | 1.154142342 | 0.000317696 | 0.007176147 | 60.46456916 | 23.96402767 | Superficial | SOX2 + KLF5, SOX2 |
| 461 | APOBEC3C | 1.150402917 | 0.002315114 | 0.011381143 | 31.44693812 | 10.14252514 | Superficial | SOX2 |
| 462 | TRIB1 | 1.147939362 | 0.001526458 | 0.016792744 | 28.23414165 | 9.735724133 | Basal | SOX2 |
| 463 | TP63 | 1.146853343 | 0.006166656 | 0.020290909 | 37.28071544 | 11.45048963 | Superficial | SOX2 + KLF5, SOX2 |
| 464 | SEMA3F | 1.146811766 | 0.001883197 | 0.010234018 | 49.11393328 | 19.38989025 | Superficial | SOX2 |
| 465 | B4GALNT3 | 1.142848614 | 0.012681065 | 0.033478436 | 9.131762208 | 2.77867419 | Superficial | SOX2 + KLF5, SOX2 |
| 466 | ZCCHC2 | 1.141906951 | 0.000992312 | 0.008042817 | 16.46931387 | 4.382144031 | Superficial | SOX2 |
| 467 | RPS6KA3 | 1.139760282 | 0.001232925 | 0.008679903 | 37.84514834 | 11.68629436 | Superficial | SOX2 + KLF5, SOX2 |
| 468 | CD24 | 1.136471416 | 0.005105016 | 0.029792864 | 54.26310106 | 31.8283184 | Basal | SOX2 |
| 469 | FAM98A | 1.136144885 | 0.000250281 | 0.007176147 | 32.20407289 | 10.52410277 | Superficial | SOX2 |
| 470 | PWWP2A | 1.135767663 | 0.000119764 | 0.007176147 | 23.25985825 | 7.532852069 | Superficial | SOX2 |
| 471 | ROBO2 | 1.132994815 | 0.007423146 | 0.022959455 | 10.96655989 | 2.722355349 | Superficial | SOX2 + KLF5 |
| 472 | LEPROTL1 | 1.130920154 | 0.002836074 | 0.012662188 | 37.83090472 | 12.64397208 | Superficial | SOX2 |
| 473 | USP14 | 1.129592738 | 0.001526458 | 0.009387421 | 41.12020623 | 13.37419131 | Superficial | SOX2 |
| 474 | LONRF1 | 1.128697791 | 0.003462059 | 0.014108599 | 16.98832356 | 3.932101419 | Superficial | SOX2 |
| 475 | PTPN11 | 1.127491694 | 0.000250281 | 0.007176147 | 45.38995352 | 15.60369219 | Superficial | SOX2 |
| 476 | SLC44A1 | 1.124630481 | 0.010644787 | 0.029507311 | 30.70360064 | 9.008386975 | Superficial | SOX2 |
| 477 | PAPSS1 | 1.123121593 | 0.002315114 | 0.011381143 | 42.22052091 | 14.95779843 | Superficial | SOX2 |
| 478 | AMMECR1 | 1.123103887 | 0.000506425 | 0.007178523 | 40.51025441 | 13.19490157 | Superficial | SOX2 |
| 479 | ATG4D | 1.120578518 | 0.001883197 | 0.010234018 | 42.99015922 | 16.54741472 | Superficial | SOX2 |
| 480 | FOXQ1 | 1.120308157 | 0.000795824 | 0.010838271 | 49.85377761 | 21.73787519 | Suprabasal | SOX2 + KLF5, SOX2 |
| 481 | DROSHA | 1.120211159 | 0.002315114 | 0.011381143 | 12.39509232 | 3.912492171 | Superficial | SOX2 |
| 482 | DGCR2 | 1.119753322 | 0.000992312 | 0.008042817 | 36.05458181 | 11.88605041 | Superficial | SOX2 |
| 483 | NAT1 | 1.11949899 | 0.005105016 | 0.018023407 | 9.560135729 | 2.733991516 | Superficial | SOX2 |
| 484 | SNX7 | 1.118765388 | 0.000506425 | 0.007178523 | 24.582421 | 8.291960086 | Superficial | SOX2 |
| 485 | BCL2L2 | 1.118547819 | 0.002315114 | 0.011381143 | 33.03297089 | 10.55302431 | Superficial | SOX2 |
| 486 | OXNAD1 | 1.115216096 | 0.001232925 | 0.008679903 | 12.00572046 | 3.740562131 | Superficial | SOX2 |
| 487 | KANK1 | 1.115157148 | 0.003462059 | 0.014108599 | 45.85522332 | 16.13807382 | Superficial | SOX2 |
| 488 | FBXL18 | 1.11490085 | 0.008904627 | 0.025990329 | 18.8479122 | 5.558665317 | Superficial | SOX2 |
| 489 | TRIM4 | 1.114744712 | 0.000119764 | 0.007176147 | 21.61232033 | 6.44415365 | Superficial | SOX2 |
| 490 | DCTD | 1.114599725 | 0.000992312 | 0.008042817 | 41.37982677 | 14.56407296 | Superficial | SOX2 |
| 491 | TMEM87A | 1.112172671 | 9.30E-05 | 0.007176147 | 62.99294426 | 28.05497565 | Superficial | SOX2 |
| 492 | ACVR1 | 1.11208563 | 0.010644787 | 0.029507311 | 7.565015766 | 1.571333265 | Superficial | SOX2, SOX2 + KLF5 |
| 493 | PRPF8 | 1.110632241 | 0.001232925 | 0.008679903 | 43.91749476 | 15.59100667 | Superficial | SOX2 |
| 494 | SLC5A3 | 1.108546111 | 0.017811616 | 0.043066688 | 15.68467168 | 4.283838368 | Superficial | SOX2 + KLF5 |
| 495 | KNOP1 | 1.107463007 | 0.000635975 | 0.007274975 | 26.61624999 | 8.346682897 | Superficial | SOX2 |
| 496 | RASEF | 1.106436035 | 0.002836074 | 0.012662188 | 33.89849056 | 10.29433174 | Superficial | SOX2 |
| 497 | ITPRIP | 1.103478092 | 0.000119764 | 0.012455402 | 20.07023032 | 8.016875374 | Basal | SOX2 |
| 498 | AGAP1 | 1.103049375 | 0.004211401 | 0.016013117 | 10.92596213 | 3.220511929 | Superficial | SOX2 |
| 499 | LYPD3 | 1.10226506 | 0.001526458 | 0.009387421 | 86.62604201 | 62.41967361 | Superficial | SOX2 |
| 500 | CEBPB | 1.100953174 | 0.001883197 | 0.010234018 | 59.35662314 | 25.73980246 | Superficial | SOX2 |
| 501 | EPHB6 | 1.098421093 | 0.002315114 | 0.011381143 | 23.20066096 | 6.510947152 | Superficial | SOX2, SOX2 + KLF5 |
| 502 | SMAD5 | 1.096440138 | 0.006166656 | 0.020290909 | 35.81411772 | 11.0476338 | Superficial | SOX2 |
| 503 | GBP6 | 1.096333706 | 0.000506425 | 0.007178523 | 71.41018052 | 33.62966856 | Superficial | SOX2 |
| 504 | ASCC3 | 1.096048273 | 0.000795824 | 0.007615337 | 35.83077768 | 11.24188818 | Superficial | SOX2 |
| 505 | FAT2 | 1.095564156 | 0.007423146 | 0.022959455 | 32.04126368 | 11.20764163 | Superficial | SOX2 |
| 506 | ECT2 | 1.095234017 | 0.004211401 | 0.021879278 | 15.76508345 | 5.983007448 | Suprabasal | SOX2 |
| 507 | ZBTB7C | 1.095229116 | 0.006166656 | 0.020290909 | 37.09729954 | 12.38960308 | Superficial | SOX2 |
| 508 | ELK3 | 1.093367327 | 0.001883197 | 0.018415354 | 28.56296451 | 10.92497033 | Basal | SOX2 |
| 509 | CRYZ | 1.092201992 | 0.001883197 | 0.010234018 | 21.73380901 | 5.586580444 | Superficial | SOX2 + KLF5 |

| | | | | | | | | |
|---|---|---|---|---|---|---|---|---|
| 510 | PITPNC1 | 1.091116348 | 0.004211401 | 0.021879278 | 21.70312809 | 8.978402459 | Suprabasal | SOX2 |
| 511 | MYO1E | 1.090802208 | 0.004211401 | 0.027185186 | 27.99278453 | 10.70128805 | Basal | SOX2 |
| 512 | LACTB2 | 1.090260768 | 0.001883197 | 0.010234018 | 20.41603957 | 5.557427606 | Superficial | SOX2 |
| 513 | ANXA8 | 1.088520498 | 0.006166656 | 0.020290909 | 37.31820819 | 12.19482701 | Superficial | SOX2 |
| 514 | PDIA3 | 1.087627963 | 0.000992312 | 0.008042817 | 71.74290452 | 35.69881232 | Superficial | SOX2 |
| 515 | EPAS1 | 1.087157497 | 0.007423146 | 0.030449933 | 48.81513341 | 24.82490815 | Suprabasal | SOX2 |
| 516 | EPHB4 | 1.086430382 | 0.002836074 | 0.012662188 | 8.61356527 | 3.03940643 | Superficial | SOX2 |
| 517 | FAM83A | 1.084655702 | 0.000992312 | 0.014320968 | 24.79974942 | 11.00318279 | Basal | SOX2 |
| 518 | PAQR5 | 1.084374053 | 0.000401828 | 0.012455402 | 11.68677813 | 4.523001064 | Basal | SOX2 |
| 519 | ACTR1B | 1.083564297 | 0.002836074 | 0.012662188 | 31.0276409 | 10.4311608 | Superficial | SOX2 |
| 520 | ID1 | 1.083067962 | 0.008904627 | 0.025990329 | 67.99885112 | 36.38589517 | Superficial | SOX2 |
| 521 | ITPR2 | 1.082561726 | 0.002315114 | 0.011381143 | 36.76971811 | 11.19914389 | Superficial | SOX2, KLF5 |
| 522 | TGFBR1 | 1.082552543 | 0.017811616 | 0.043066688 | 21.28035822 | 5.768770277 | Superficial | SOX2 + KLF5 |
| 523 | ELOVL5 | 1.081800374 | 0.004211401 | 0.021879278 | 35.05474557 | 15.29151468 | Suprabasal | SOX2 |
| 524 | HNRNPLL | 1.081470571 | 0.000119764 | 0.007176147 | 19.51931021 | 5.953028379 | Superficial | SOX2 |
| 525 | CKMT1A | 1.080797256 | 0.003462059 | 0.014108599 | 45.82714419 | 16.93959481 | Superficial | KLF5 |
| 526 | RTKN2 | 1.07996988 | 0.000401828 | 0.012455402 | 26.91170963 | 12.18065285 | Basal | SOX2 |
| 527 | IRF6 | 1.076699665 | 0.000635975 | 0.007274975 | 66.79817342 | 31.74169161 | Superficial | SOX2 |
| 528 | GJB6 | 1.074050502 | 0.010644787 | 0.029507311 | 72.67287901 | 34.07712441 | Superficial | SOX2 |
| 529 | ZNF276 | 1.073832933 | 0.004211401 | 0.016013117 | 21.6021014 | 8.143746 | Superficial | SOX2 |
| 530 | GJB3 | 1.073481421 | 0.001883197 | 0.010234018 | 52.3986983 | 20.43895374 | Superficial | SOX2 |
| 531 | ZDHHC24 | 1.073405826 | 0.005105016 | 0.018023407 | 31.33471846 | 10.87359273 | Superficial | SOX2 |
| 532 | TXLNA | 1.072929136 | 0.007423146 | 0.022959455 | 26.7723363 | 9.668176579 | Superficial | SOX2 |
| 533 | GNPDA2 | 1.072787336 | 0.003462059 | 0.014108599 | 8.553457117 | 2.209436035 | Superficial | SOX2 |
| 534 | ATP7A | 1.07034877 | 0.001526458 | 0.009387421 | 18.37821598 | 5.769978836 | Superficial | SOX2 |
| 535 | CDKN2B | 1.070294963 | 0.012681065 | 0.049425568 | 18.07850545 | 8.341417478 | Basal | SOX2 |
| 536 | PCNX1 | 1.068627785 | 0.000795824 | 0.007615337 | 13.87619405 | 3.845484788 | Superficial | SOX2 |
| 537 | CCDC60 | 1.068090748 | 0.004173922 | 0.016013117 | 5.07941827 | 1.732237896 | Superficial | SOX2 |
| 538 | EML4 | 1.066487042 | 0.001232925 | 0.008679903 | 29.48556773 | 8.712000848 | Superficial | SOX2 |
| 539 | ZNF219 | 1.066374966 | 0.002836074 | 0.012662188 | 38.58342754 | 13.5543592 | Superficial | SOX2 |
| 540 | RIF1 | 1.06443131 | 0.004211401 | 0.016013117 | 24.59154314 | 6.647944401 | Superficial | SOX2 |
| 541 | CDC42EP4 | 1.064259537 | 0.000506425 | 0.010489143 | 28.86197028 | 11.84880536 | Suprabasal | SOX2 |
| 542 | HR | 1.060194125 | 0.000635975 | 0.007274975 | 22.82513006 | 8.75455057 | Superficial | SOX2 |
| 543 | WWP1 | 1.05999757 | 7.20E-05 | 0.007176147 | 45.17265309 | 16.34281943 | Superficial | SOX2 |
| 544 | BOLA2 | 1.059889855 | 0.012659085 | 0.033478436 | 27.52597963 | 8.841058943 | Superficial | SOX2 |
| 545 | HES2 | 1.059617182 | 0.000196466 | 0.012455402 | 47.53905383 | 23.28976828 | Basal | SOX2 |
| 546 | CDC42BPG | 1.058014931 | 0.006166656 | 0.020290909 | 29.73744229 | 10.84726719 | Superficial | SOX2 |
| 547 | SQLE | 1.057507387 | 0.010644787 | 0.029507311 | 53.21453026 | 22.40711233 | Superficial | SOX2 |
| 548 | PIGO | 1.055646727 | 0.004211401 | 0.016013117 | 11.60956119 | 3.701016275 | Superficial | SOX2 |
| 549 | TADA3 | 1.055607235 | 0.000317696 | 0.007176147 | 40.42552055 | 15.3811476 | Superficial | SOX2 |
| 550 | ZNFX1 | 1.05419277 | 0.000992312 | 0.014320968 | 16.99829037 | 5.815432506 | Basal | SOX2 |
| 551 | SMG6 | 1.052156641 | 0.000506425 | 0.007178523 | 28.41227063 | 10.15996499 | Superficial | SOX2 |
| 552 | RAB11FIP2 | 1.051539922 | 0.004211401 | 0.016013117 | 12.72551013 | 2.863554896 | Superficial | SOX2 + KLF5, SOX2 |
| 553 | GALNT12 | 1.049379383 | 0.001232925 | 0.008679903 | 34.35833398 | 11.63233406 | Superficial | SOX2 |
| 554 | PDIK1L | 1.048431055 | 0.006166656 | 0.020290909 | 11.5882273 | 3.088217267 | Superficial | SOX2 |
| 555 | FOXC1 | 1.047638776 | 0.017811616 | 0.043066688 | 15.52853274 | 7.61471716 | Superficial | SOX2 |
| 556 | COG6 | 1.04676455 | 0.001526458 | 0.009387421 | 18.12524566 | 6.012127181 | Superficial | SOX2 |
| 557 | CLDN4 | 1.046614095 | 0.003462059 | 0.024854712 | 53.00129205 | 29.07079485 | Basal | SOX2, SOX2 + KLF5 |
| 558 | TBC1D5 | 1.045495453 | 0.000506425 | 0.007178523 | 21.15279137 | 6.383257151 | Superficial | SOX2 |
| 559 | GBP2 | 1.043537437 | 0.006166656 | 0.020290909 | 43.075913 | 16.48742557 | Superficial | SOX2, SOX2 + KLF5 |
| 560 | CHCHD4 | 1.041517737 | 0.005105016 | 0.018023407 | 13.20831071 | 4.101917452 | Superficial | SOX2 |
| 561 | PIAS3 | 1.039484217 | 0.005105016 | 0.018023407 | 21.87521411 | 7.021571866 | Superficial | SOX2 |
| 562 | PDZRN3 | 1.038845423 | 0.005105016 | 0.018023407 | 9.730521189 | 3.580156724 | Superficial | SOX2 + KLF5, SOX2 |
| 563 | EPB41L5 | 1.038189591 | 0.001883197 | 0.010234018 | 32.11329554 | 10.05839763 | Superficial | SOX2 |
| 564 | TNKS | 1.036998801 | 0.001883197 | 0.010234018 | 19.29376441 | 5.687144619 | Superficial | SOX2 |
| 565 | GJB5 | 1.036529192 | 0.012681065 | 0.033478436 | 50.72292565 | 17.65083088 | Superficial | SOX2 + KLF5 |
| 566 | MANF | 1.032784742 | 0.003462059 | 0.014108599 | 68.1159678 | 32.72827979 | Superficial | SOX2 |
| 567 | HMGXB3 | 1.031603042 | 0.005105016 | 0.018023407 | 13.16907487 | 4.247519548 | Superficial | SOX2 |
| 568 | DLG1 | 1.030612623 | 0.000250281 | 0.007176147 | 59.02506663 | 26.05889543 | Superficial | SOX2 |
| 569 | ATAD2B | 1.029425719 | 0.001526458 | 0.009387421 | 21.42384634 | 5.846551603 | Superficial | SOX2 |
| 570 | USP12 | 1.028326657 | 0.000506425 | 0.010489143 | 24.13745095 | 9.410233246 | Suprabasal | SOX2, SOX2 + KLF5 |
| 571 | USP42 | 1.027599754 | 0.001526458 | 0.009387421 | 11.22238109 | 3.728432416 | Superficial | SOX2 |
| 572 | APEX2 | 1.026096018 | 0.000992312 | 0.008042817 | 25.71451459 | 9.11992571 | Superficial | SOX2 |
| 573 | RAP2C | 1.025837921 | 0.000401828 | 0.007176147 | 31.69093352 | 10.06563417 | Superficial | SOX2 |
| 574 | CGN | 1.025734272 | 0.001526458 | 0.009387421 | 44.55844951 | 16.51730629 | Superficial | SOX2 |
| 575 | HCFC2 | 1.025545192 | 0.001526458 | 0.009387421 | 17.90303463 | 5.748783799 | Superficial | SOX2 |
| 576 | CLIP1 | 1.025361477 | 0.000250281 | 0.012455402 | 51.95265842 | 26.04386187 | Basal | SOX2 |
| 577 | SRFBP1 | 1.025231274 | 0.006166656 | 0.020290909 | 20.35835208 | 6.194081963 | Superficial | SOX2 |
| 578 | SFN | 1.024113903 | 0.003462059 | 0.014108599 | 86.40801124 | 59.03093976 | Superficial | SOX2 |
| 579 | CHMP7 | 1.023040049 | 0.004211401 | 0.016013117 | 14.31554793 | 5.15490207 | Superficial | SOX2 |
| 580 | SEMA4C | 1.020675853 | 0.000401828 | 0.012455402 | 13.60453169 | 4.878061834 | Basal | SOX2 |
| 581 | IMMT | 1.020403361 | 0.007423146 | 0.022959455 | 41.15591597 | 13.86769913 | Superficial | SOX2 |
| 582 | MED27 | 1.016751381 | 0.001883197 | 0.010234018 | 20.7375588 | 6.37266373 | Superficial | SOX2 |

| | | | | | | | |
|---|---|---|---|---|---|---|---|
| 583 | STK39 | 1.016669337 | 0.001883197 | 0.018415354 | 21.59722621 | 8.154177809 | Basal | SOX2 |
| 584 | VCL | 1.016433297 | 0.000250281 | 0.012455402 | 50.18358999 | 23.40486234 | Basal | SOX2 + KLF5, SOX2 |
| 585 | GLG1 | 1.014764777 | 0.000506425 | 0.007178523 | 38.66809769 | 13.84952322 | Superficial | SOX2 |
| 586 | DGKH | 1.014239019 | 0.012681065 | 0.033478436 | 16.90556525 | 3.927563317 | Superficial | SOX2 |
| 587 | SPIRE2 | 1.012985199 | 0.010644787 | 0.038255467 | 4.88784887 | 2.320639171 | Suprabasal | KLF5, SOX2 |
| 588 | PDIA4 | 1.011720547 | 0.006166656 | 0.020290909 | 44.57312376 | 18.01306715 | Superficial | SOX2 |
| 589 | IPO8 | 1.011278306 | 0.002315114 | 0.011381143 | 19.10865866 | 6.340994881 | Superficial | SOX2 |
| 590 | WRNIP1 | 1.01058191 | 0.000401828 | 0.007176147 | 31.36699098 | 10.96769113 | Superficial | SOX2 |
| 591 | MRPL35 | 1.009994966 | 0.001526458 | 0.009387421 | 44.0994244 | 15.84466649 | Superficial | SOX2 |
| 592 | CYP2J2 | 1.009979358 | 0.002836074 | 0.012662188 | 39.69669461 | 14.93332307 | Superficial | SOX2 |
| 593 | SSBP2 | 1.008614228 | 0.003462059 | 0.014108599 | 34.56102056 | 10.69519446 | Superficial | SOX2 |
| 594 | EMP2 | 1.008286992 | 0.000401828 | 0.007176147 | 85.15076119 | 65.60610945 | Superficial | SOX2 |
| 595 | ELOVL1 | 1.007657584 | 0.002836074 | 0.012662188 | 67.73851337 | 30.2888522 | Superficial | SOX2 |
| 596 | WIZ | 1.003348574 | 0.004211401 | 0.016013117 | 14.06167781 | 4.932187334 | Superficial | SOX2 |
| 597 | FLYWCH1 | 1.003246571 | 0.015054848 | 0.048140298 | 13.30169272 | 6.134460931 | Suprabasal | SOX2 |
| 598 | PANK3 | 1.002008835 | 0.005105016 | 0.018023407 | 42.2483754 | 14.84793779 | Superficial | SOX2 |
| 599 | ERBB2 | 1.000797318 | 0.003462059 | 0.014108599 | 64.90384468 | 29.36807898 | Superficial | SOX2 |
| 600 | GJB2 | 1.000707221 | 0.015054848 | 0.038009884 | 77.54819408 | 42.9288687 | Superficial | SOX2 |
| 601 | SREBF2 | 1.000004846 | 0.001526458 | 0.009387421 | 46.98351695 | 20.86267964 | Superficial | SOX2 |
| 602 | NAA35 | 0.999499739 | 0.000250281 | 0.007176147 | 24.88781225 | 7.308081533 | Superficial | SOX2 |
| 603 | OAS3 | 0.999411652 | 0.017811616 | 0.043066688 | 33.91694122 | 9.730762134 | Superficial | SOX2 |
| 604 | SLC4A2 | 0.998539121 | 0.000506425 | 0.007178523 | 17.02633306 | 5.480113383 | Superficial | SOX2 |
| 605 | RNF220 | 0.998062117 | 0.001526458 | 0.009387421 | 21.53668386 | 7.445293659 | Superficial | SOX2 |
| 606 | CYP3A5 | 0.997896563 | 0.008904627 | 0.025990329 | 33.40356 | 12.07692171 | Superficial | SOX2 |
| 607 | PREP | 0.997620789 | 0.000506425 | 0.007178523 | 18.8332859 | 5.754453574 | Superficial | SOX2 |
| 608 | ANKRD50 | 0.997212216 | 0.006166656 | 0.020290909 | 10.84753741 | 3.542220569 | Superficial | SOX2 |
| 609 | SMARCAD1 | 0.996268912 | 0.007423146 | 0.022959455 | 14.35571326 | 4.114308749 | Superficial | SOX2 |
| 610 | CLASP2 | 0.995469306 | 0.021001058 | 0.04834985 | 17.84008933 | 5.495820276 | Superficial | SOX2 |
| 611 | CREB3 | 0.993048401 | 0.021001058 | 0.04834985 | 16.08875673 | 6.82847754 | Superficial | SOX2 |
| 612 | GSN | 0.992984158 | 0.000250281 | 0.012455402 | 80.3245791 | 55.83838799 | Basal | SOX2, KLF5 |
| 613 | PLCH2 | 0.990452369 | 0.000317696 | 0.010489143 | 19.86664017 | 9.082930267 | Suprabasal | SOX2 |
| 614 | KDM1B | 0.988968052 | 0.001883197 | 0.010234018 | 17.13891304 | 5.515185538 | Superficial | SOX2 |
| 615 | MAPKAPK3 | 0.985942266 | 0.000250281 | 0.007176147 | 61.16829182 | 27.56571044 | Superficial | SOX2 |
| 616 | N4BP3 | 0.982741775 | 0.000317696 | 0.012455402 | 13.99363767 | 5.703652307 | Basal | SOX2 |
| 617 | RAB3GAP2 | 0.981611807 | 0.000795824 | 0.007615337 | 30.71255762 | 10.52537581 | Superficial | SOX2 |
| 618 | CDC40 | 0.979458465 | 0.001526458 | 0.009387421 | 42.10328835 | 14.46730754 | Superficial | SOX2 |
| 619 | APOBEC3B | 0.978993045 | 0.006166656 | 0.032981404 | 10.8000218 | 4.018199162 | Basal | SOX2 |
| 620 | ETV4 | 0.978127481 | 0.021001058 | 0.04834985 | 18.32034763 | 5.168240674 | Superficial | SOX2 |
| 621 | RBM27 | 0.976527696 | 0.000317696 | 0.007176147 | 41.41526646 | 14.85977794 | Superficial | SOX2 |
| 622 | VPS13A | 0.975658933 | 0.001883197 | 0.010234018 | 29.64566246 | 9.650092596 | Superficial | SOX2 |
| 623 | TRIM29 | 0.975559011 | 0.003462059 | 0.014108599 | 83.4744379 | 60.80594581 | Superficial | SOX2 |
| 624 | CLNS1A | 0.973230763 | 0.001526458 | 0.009387421 | 52.95854222 | 20.20355588 | Superficial | SOX2 |
| 625 | CLDN7 | 0.973223705 | 0.000119764 | 0.012455402 | 72.00569277 | 42.20990403 | Basal | SOX2 |
| 626 | NFIL3 | 0.973007118 | 0.000153669 | 0.010489143 | 31.80256845 | 14.76095525 | Suprabasal | SOX2 |
| 627 | OVOL1 | 0.972186252 | 0.004211401 | 0.027185186 | 36.62987761 | 17.67239326 | Basal | SOX2 |
| 628 | ARL6IP6 | 0.971918062 | 0.001883197 | 0.010234018 | 27.33317617 | 8.29542678 | Superficial | SOX2 |
| 629 | VLDLR | 0.971565055 | 0.005105016 | 0.018023407 | 34.62065303 | 10.68412266 | Superficial | SOX2 |
| 630 | EFNB2 | 0.971028932 | 0.002315114 | 0.011381143 | 34.08986984 | 10.68114608 | Superficial | SOX2 + KLF5, SOX2 |
| 631 | C5orf22 | 0.964610162 | 0.002315114 | 0.011381143 | 14.12842584 | 4.040622089 | Superficial | SOX2 |
| 632 | SLC35F6 | 0.963323494 | 0.004211401 | 0.016013117 | 36.32587527 | 13.4031697 | Superficial | SOX2 |
| 633 | SBF1 | 0.962933302 | 0.000401828 | 0.012455402 | 18.97321274 | 7.722577967 | Basal | SOX2 |
| 634 | HEATR3 | 0.962903361 | 0.010644787 | 0.029507311 | 10.24391514 | 3.623009779 | Superficial | SOX2 |
| 635 | GRHL2 | 0.962221635 | 0.002836074 | 0.012662188 | 26.28896804 | 9.49884602 | Superficial | SOX2 |
| 636 | CCDC88C | 0.960891309 | 0.008904627 | 0.025990329 | 14.82332483 | 4.120973742 | Superficial | SOX2 + KLF5 |
| 637 | OCLN | 0.95992656 | 0.000250281 | 0.012455402 | 16.5872587 | 6.29292647 | Basal | SOX2 |
| 638 | GK | 0.959674774 | 0.012681065 | 0.033478436 | 22.60421961 | 7.571280152 | Superficial | SOX2 |
| 639 | CASP1 | 0.957510724 | 0.000795824 | 0.007615337 | 26.91963947 | 8.607716575 | Superficial | KLF5 |
| 640 | PATJ | 0.954789173 | 0.000119764 | 0.007176147 | 62.19275094 | 28.5356374 | Superficial | SOX2 |
| 641 | RBM12B | 0.951788942 | 0.007423146 | 0.022959455 | 9.07536362 | 2.496305965 | Superficial | SOX2 |
| 642 | MBNL3 | 0.951778334 | 0.005105016 | 0.018023407 | 20.86518059 | 7.084354369 | Superficial | SOX2 + KLF5, SOX2 |
| 643 | GPHN | 0.950352459 | 0.005105016 | 0.018023407 | 10.12383445 | 2.84515677 | Superficial | SOX2 + KLF5 |
| 644 | CLASP1 | 0.948346748 | 0.003462059 | 0.019729589 | 23.72656834 | 10.93478622 | Suprabasal | SOX2 |
| 645 | TJP1 | 0.947613883 | 0.000196466 | 0.012455402 | 59.43942815 | 31.52598771 | Basal | SOX2 |
| 646 | NTF4 | 0.947273158 | 0.012681065 | 0.042921737 | 5.706436766 | 2.210792774 | Suprabasal | SOX2 |
| 647 | ITPKC | 0.947170762 | 0.002836074 | 0.022562695 | 44.35874301 | 23.17878921 | Basal | SOX2 |
| 648 | SNRNP200 | 0.945149806 | 0.000506425 | 0.007178523 | 38.43103539 | 14.67962995 | Superficial | SOX2 |
| 649 | LIMK2 | 0.943596112 | 0.002836074 | 0.012662188 | 38.02056013 | 14.07641227 | Superficial | SOX2 |
| 650 | IFI44 | 0.942373886 | 0.012681065 | 0.042921737 | 16.09657028 | 5.918626253 | Suprabasal | SOX2 |
| 651 | IFNGR1 | 0.941175231 | 0.004211401 | 0.016013117 | 33.31017628 | 11.24701455 | Superficial | SOX2 |
| 652 | TXNDC5 | 0.940628861 | 0.000795824 | 0.0136425 | 35.12375823 | 13.57082836 | Basal | SOX2 |
| 653 | PKP1 | 0.940526247 | 0.005105016 | 0.018023407 | 73.57168146 | 41.33040784 | Superficial | SOX2 + KLF5, SOX2 |
| 654 | TM7SF2 | 0.940277008 | 0.003462059 | 0.014108599 | 74.58981573 | 44.97928738 | Superficial | SOX2 |
| 655 | ATXN1 | 0.939973269 | 0.008904627 | 0.025990329 | 30.66077976 | 9.153560369 | Superficial | SOX2 + KLF5, SOX2 |

| | | | | | | | | |
|---|---|---|---|---|---|---|---|---|
| 656 | HYAL2 | 0.939193889 | 0.008904627 | 0.025990329 | 25.45374355 | 9.165485301 | Superficial | SOX2 |
| 657 | PIR | 0.938528782 | 0.010644787 | 0.029507311 | 42.51794111 | 15.60827055 | Superficial | SOX2 |
| 658 | SRCAP | 0.937080417 | 0.004211401 | 0.021879278 | 23.12213982 | 10.22666155 | Suprabasal | SOX2 |
| 659 | KDM5A | 0.936624009 | 0.000401828 | 0.007176147 | 52.87431596 | 22.545332 | Superficial | SOX2 |
| 660 | HIPK2 | 0.936277725 | 0.000795824 | 0.007615337 | 44.33080382 | 16.53588677 | Superficial | SOX2 |
| 661 | XPR1 | 0.935980881 | 0.008904627 | 0.034252903 | 13.83697012 | 5.461275977 | Suprabasal | SOX2 |
| 662 | C11orf80 | 0.934813283 | 0.001883197 | 0.010234018 | 19.61554888 | 7.757036142 | Superficial | SOX2 |
| 663 | TLCD1 | 0.934508989 | 0.008904627 | 0.025990329 | 26.94830138 | 9.425029458 | Superficial | SOX2 |
| 664 | IL15 | 0.931288218 | 0.002836074 | 0.0176927 | 10.39551987 | 3.705311357 | Suprabasal | SOX2, SOX2 + KLF5 |
| 665 | IGF1R | 0.930404525 | 0.003462059 | 0.019729589 | 35.5263818 | 16.47930112 | Suprabasal | SOX2 |
| 666 | VPS16 | 0.930332913 | 0.003462059 | 0.014108599 | 23.27652963 | 9.006523619 | Superficial | SOX2 |
| 667 | PARN | 0.930213959 | 0.015054848 | 0.038009884 | 13.26146873 | 4.826815714 | Superficial | SOX2 |
| 668 | IFIH1 | 0.930212135 | 0.007423146 | 0.030449933 | 37.56081844 | 17.67361083 | Suprabasal | SOX2 + KLF5 |
| 669 | MLKL | 0.929694873 | 0.007423146 | 0.022959455 | 13.79556589 | 5.302842768 | Superficial | SOX2 |
| 670 | BCL6 | 0.929155584 | 0.004211401 | 0.021879278 | 25.26670431 | 10.89606288 | Suprabasal | SOX2 |
| 671 | SULF2 | 0.926816247 | 0.004211401 | 0.021879278 | 56.00061141 | 30.06269751 | Suprabasal | SOX2 |
| 672 | SPRY2 | 0.92469581 | 0.003462059 | 0.019729589 | 9.968966492 | 3.716915077 | Suprabasal | SOX2 |
| 673 | RPUSD3 | 0.921826912 | 0.004211401 | 0.016013117 | 36.06544029 | 14.38430226 | Superficial | SOX2 |
| 674 | TNKS1BP1 | 0.921810736 | 0.000401828 | 0.007176147 | 43.2456963 | 17.19199187 | Superficial | SOX2 |
| 675 | METTL16 | 0.921425744 | 0.015054848 | 0.038009884 | 10.71836209 | 3.663641463 | Superficial | SOX2 |
| 676 | THUMPD1 | 0.921027525 | 0.000795824 | 0.007615337 | 35.43257965 | 11.99410572 | Superficial | SOX2 |
| 677 | MORC4 | 0.919114825 | 0.021001058 | 0.04834985 | 19.58222702 | 6.690165479 | Superficial | SOX2 |
| 678 | SMPD3 | 0.91883153 | 0.012681065 | 0.033478436 | 9.692248683 | 3.781776836 | Superficial | SOX2, KLF5, SOX2 + KLF5 |
| 679 | CTSS | 0.918754346 | 0.003462059 | 0.019729589 | 58.74895434 | 32.65659344 | Suprabasal | SOX2 |
| 680 | DDX23 | 0.918262459 | 0.000635975 | 0.007274975 | 33.28708232 | 12.49172321 | Superficial | SOX2 |
| 681 | DNAJC11 | 0.918087484 | 0.005105016 | 0.018023407 | 17.65883872 | 6.07592926 | Superficial | SOX2 |
| 682 | CHD1 | 0.916971655 | 0.002315114 | 0.011381143 | 29.52238981 | 10.04115794 | Superficial | SOX2 |
| 683 | NUP133 | 0.914397334 | 0.015054848 | 0.038009884 | 13.23558916 | 4.066194821 | Superficial | SOX2 |
| 684 | ZNF532 | 0.913554972 | 0.008904627 | 0.034252903 | 14.08271725 | 5.827776916 | Suprabasal | SOX2 |
| 685 | ANK3 | 0.913274645 | 0.012681065 | 0.033478436 | 32.92849736 | 10.82374859 | Superficial | SOX2 + KLF5 |
| 686 | PEX26 | 0.911924325 | 0.021001058 | 0.04834985 | 17.34632142 | 5.907570971 | Superficial | SOX2 |
| 687 | RTCB | 0.910029323 | 0.000506425 | 0.007178523 | 64.11821594 | 31.35594231 | Superficial | SOX2 |
| 688 | AFG3L2 | 0.909662689 | 0.001526458 | 0.009387421 | 45.0308695 | 18.25711922 | Superficial | SOX2 |
| 689 | UBE3C | 0.908984058 | 0.002315114 | 0.011381143 | 28.28995982 | 9.861957868 | Superficial | SOX2 |
| 690 | SCEL | 0.908585979 | 0.004211401 | 0.027185186 | 35.21521921 | 18.03409494 | Basal | SOX2 |
| 691 | CCAR1 | 0.908014534 | 0.008904627 | 0.025990329 | 41.8177982 | 14.38612398 | Superficial | SOX2 |
| 692 | TNFSF10 | 0.906889478 | 0.001526458 | 0.016792744 | 90.0032318 | 67.96705726 | Basal | SOX2, KLF5 |
| 693 | SLC45A4 | 0.906108969 | 0.008904627 | 0.040392504 | 8.764812014 | 2.929838771 | Basal | SOX2 |
| 694 | MAGED2 | 0.905063097 | 0.000635975 | 0.007274975 | 36.3342794 | 13.70758675 | Superficial | SOX2 |
| 695 | BCR | 0.904694009 | 0.000795824 | 0.0136425 | 21.22696209 | 8.213040475 | Basal | SOX2 |
| 696 | ITPR3 | 0.90283063 | 0.001526458 | 0.016792744 | 27.31887442 | 11.01249269 | Basal | SOX2 |
| 697 | ARHGEF5 | 0.902031503 | 0.004211401 | 0.021879278 | 49.08157174 | 27.73966165 | Suprabasal | SOX2 |
| 698 | TBC1D10A | 0.901080387 | 0.004211401 | 0.016013117 | 31.31177269 | 11.81774667 | Superficial | SOX2 |
| 699 | ANKIB1 | 0.900545456 | 0.000153669 | 0.007176147 | 37.04964394 | 13.30203144 | Superficial | SOX2 |
| 700 | SLC4A7 | 0.899999926 | 0.017811616 | 0.043066688 | 6.29061434 | 1.861275296 | Superficial | SOX2 + KLF5 |
| 701 | FXYD3 | 0.899656014 | 0.002315114 | 0.011381143 | 87.03589411 | 66.84367355 | Superficial | SOX2 |
| 702 | CDKAL1 | 0.89933994 | 0.000506425 | 0.007178523 | 11.61467749 | 3.791144178 | Superficial | SOX2 |
| 703 | FURIN | 0.899232364 | 0.000153669 | 0.012455402 | 17.94979085 | 6.915633525 | Basal | SOX2 |
| 704 | CAMSAP1 | 0.899201892 | 0.000119764 | 0.012455402 | 33.88678212 | 15.33563619 | Basal | SOX2 |
| 705 | PHC2 | 0.896625623 | 0.001883197 | 0.010234018 | 17.54207804 | 6.22869448 | Superficial | SOX2 |
| 706 | LRRC42 | 0.895968102 | 0.001232925 | 0.008679903 | 32.72943189 | 11.61962184 | Superficial | SOX2 |
| 707 | SLC39A6 | 0.895816619 | 0.006166656 | 0.020290909 | 16.16572683 | 4.572418678 | Superficial | SOX2 |
| 708 | ST14 | 0.89536654 | 7.20E-05 | 0.012455402 | 50.40009109 | 25.34790976 | Basal | SOX2, SOX2 + KLF5 |
| 709 | PLEKHG3 | 0.895015137 | 0.015054848 | 0.038009884 | 13.95511449 | 4.823953346 | Superficial | SOX2 |
| 710 | TIAM1 | 0.894450063 | 0.000992312 | 0.008042817 | 54.68463967 | 24.04975501 | Superficial | SOX2 |
| 711 | KIAA1217 | 0.893786401 | 0.000992312 | 0.008042817 | 53.56579201 | 21.83646765 | Superficial | SOX2 |
| 712 | SPECC1L | 0.893320652 | 0.001232925 | 0.015721789 | 24.77042245 | 9.815567676 | Basal | SOX2 |
| 713 | EPN2 | 0.893135704 | 0.000795824 | 0.007615337 | 63.41211342 | 30.86327485 | Superficial | SOX2 + KLF5 |
| 714 | BMPR2 | 0.892138295 | 0.001883197 | 0.010234018 | 38.46729266 | 14.761013 | Superficial | SOX2, SOX2 + KLF5 |
| 715 | CDCP1 | 0.890652929 | 0.000992312 | 0.008042817 | 66.76017312 | 32.69936178 | Superficial | SOX2 |
| 716 | TNRC18 | 0.889205782 | 0.017811616 | 0.043066688 | 24.24592989 | 8.527045348 | Superficial | SOX2 + KLF5, SOX2 |
| 717 | TOMM70 | 0.887267944 | 0.003462059 | 0.014108599 | 30.88206662 | 10.68652037 | Superficial | SOX2 |
| 718 | KCNK7 | 0.886984141 | 0.012681065 | 0.033478436 | 31.28106222 | 10.82219727 | Superficial | SOX2 |
| 719 | EPHB3 | 0.885669184 | 0.001526458 | 0.016792744 | 17.94114574 | 7.389341179 | Basal | SOX2 |
| 720 | BMPR1A | 0.882293181 | 0.003462059 | 0.014108599 | 12.01608827 | 3.634607464 | Superficial | SOX2 |
| 721 | TRAF3IP1 | 0.882212095 | 0.005105016 | 0.018023407 | 14.53927799 | 4.312759643 | Superficial | SOX2 |
| 722 | NOL10 | 0.882088665 | 0.015054848 | 0.038009884 | 17.02148532 | 6.188813952 | Superficial | SOX2 + KLF5, SOX2 |
| 723 | RNF19B | 0.881491342 | 0.000506425 | 0.012898798 | 25.32034955 | 10.32639443 | Basal | SOX2 + KLF5, SOX2 |
| 724 | UGCG | 0.880956022 | 0.000317696 | 0.012455402 | 41.6884798 | 18.93866914 | Basal | SOX2, SOX2 + KLF5 |
| 725 | ABLIM2 | 0.87843226 | 0.001232925 | 0.012246384 | 21.47830619 | 10.05916517 | Suprabasal | SOX2 |
| 726 | CEP170B | 0.878140221 | 0.000401828 | 0.012455402 | 19.88590991 | 7.313422792 | Basal | SOX2 |
| 727 | MORC2 | 0.87731019 | 0.001526458 | 0.009387421 | 11.36270155 | 3.478454545 | Superficial | SOX2 |
| 728 | FTSJ3 | 0.876905147 | 0.000635975 | 0.007274975 | 19.33425643 | 6.629415989 | Superficial | SOX2 |

| | | | | | | | | |
|---|---|---|---|---|---|---|---|---|
| 729 | NOTCH1 | 0.876227202 | 0.003462059 | 0.024854712 | 28.32956634 | 13.11437681 | Basal | SOX2 |
| 730 | ZDHHC16 | 0.875936954 | 0.008904627 | 0.025990329 | 18.0535891 | 6.514104604 | Superficial | SOX2 |
| 731 | IGSF3 | 0.875568181 | 0.004211401 | 0.016013117 | 20.64854284 | 7.060117599 | Superficial | SOX2 |
| 732 | METTL14 | 0.875376042 | 0.002836074 | 0.012662188 | 20.15456999 | 6.39679254 | Superficial | SOX2 |
| 733 | HDGF | 0.875145778 | 0.000506425 | 0.007178523 | 63.33322983 | 30.12978082 | Superficial | SOX2 |
| 734 | BTG1 | 0.874641303 | 7.20E-05 | 0.007176147 | 76.33575407 | 46.98814185 | Superficial | SOX2, SOX2 + KLF5 |
| 735 | CHRAC1 | 0.874329147 | 0.001526458 | 0.009387421 | 32.58122904 | 11.03633825 | Superficial | SOX2 |
| 736 | PPP1R26 | 0.874196498 | 0.000795824 | 0.010838271 | 9.010442668 | 3.864186864 | Suprabasal | SOX2 |
| 737 | CAMSAP3 | 0.873243075 | 0.001232925 | 0.015721789 | 12.6460338 | 5.306066422 | Basal | SOX2 |
| 738 | TP53RK | 0.873165537 | 0.001883197 | 0.010234018 | 34.69402433 | 13.34150254 | Superficial | SOX2 |
| 739 | LETM1 | 0.872109216 | 0.000635975 | 0.007274975 | 48.00505357 | 21.60553929 | Superficial | SOX2 |
| 740 | RBM19 | 0.870734356 | 0.008904627 | 0.025990329 | 9.455645202 | 3.177116214 | Superficial | SOX2 |
| 741 | NOP14 | 0.870040844 | 0.012681065 | 0.033478436 | 17.67069829 | 5.687473923 | Superficial | SOX2 |
| 742 | SLC25A46 | 0.869726759 | 0.010644787 | 0.029507311 | 27.18592165 | 9.768766573 | Superficial | SOX2 |
| 743 | NLRX1 | 0.869680205 | 0.006166656 | 0.020290909 | 44.2626525 | 19.65292128 | Superficial | SOX2 |
| 744 | ADCK2 | 0.869583124 | 0.007423146 | 0.022959455 | 21.60813141 | 8.531418818 | Superficial | SOX2 |
| 745 | RAB3D | 0.86952271 | 0.015054848 | 0.038009884 | 36.43417123 | 14.40115223 | Superficial | SOX2 |
| 746 | HNRNPAB | 0.868905205 | 0.002836074 | 0.012662188 | 55.3602438 | 22.22158232 | Superficial | SOX2 |
| 747 | CSNK1G3 | 0.868108155 | 0.000635975 | 0.007274975 | 25.97849208 | 8.715818884 | Superficial | SOX2 |
| 748 | PLXNA2 | 0.867985641 | 0.006166656 | 0.020290909 | 29.09724384 | 8.662493629 | Superficial | SOX2 |
| 749 | POLR3A | 0.867854544 | 0.017811616 | 0.043066688 | 9.674285364 | 2.890829878 | Superficial | SOX2 |
| 750 | PLOD2 | 0.867405121 | 0.003462059 | 0.014108599 | 24.23100976 | 7.329784954 | Superficial | SOX2 |
| 751 | CLOCK | 0.867335537 | 0.015054848 | 0.038009884 | 35.97663274 | 12.96369281 | Superficial | SOX2 |
| 752 | VRK2 | 0.866624796 | 0.002836074 | 0.012662188 | 27.63407787 | 9.880195058 | Superficial | SOX2 |
| 753 | MTA2 | 0.866164613 | 0.017811616 | 0.043066688 | 24.48027662 | 10.40104993 | Superficial | SOX2 |
| 754 | IRF1 | 0.864606905 | 7.20E-05 | 0.012455402 | 66.36951881 | 38.12897296 | Basal | SOX2 |
| 755 | SPSB2 | 0.863671948 | 0.021001058 | 0.04834985 | 12.26063796 | 4.552374875 | Superficial | SOX2 |
| 756 | UNC45A | 0.863347383 | 0.008904627 | 0.025990329 | 30.21492431 | 11.95655965 | Superficial | SOX2 |
| 757 | FAM110C | 0.863318044 | 0.000401828 | 0.010489143 | 49.1183436 | 27.19016149 | Suprabasal | SOX2, SOX2 + KLF5 |
| 758 | RNF214 | 0.860639186 | 0.000992312 | 0.008042817 | 20.37905803 | 7.210504233 | Superficial | SOX2 |
| 759 | NLN | 0.860495589 | 0.002836074 | 0.012662188 | 10.91799313 | 3.618501469 | Superficial | SOX2 |
| 760 | NSMAF | 0.858277222 | 0.012681065 | 0.033478436 | 11.88905994 | 4.845257228 | Superficial | SOX2 |
| 761 | TPST1 | 0.858189681 | 0.000992312 | 0.01142308 | 22.00115385 | 9.487918828 | Suprabasal | SOX2 |
| 762 | MCMBP | 0.856648759 | 0.000401828 | 0.007176147 | 28.42016649 | 10.86479165 | Superficial | SOX2 |
| 763 | OSBP | 0.855633877 | 0.006166656 | 0.020290909 | 40.09049547 | 16.42969957 | Superficial | SOX2 |
| 764 | FAS | 0.855309759 | 0.000196466 | 0.010489143 | 28.48938444 | 12.96714597 | Suprabasal | SOX2 |
| 765 | LUZP1 | 0.855211261 | 0.000196466 | 0.007176147 | 37.10983106 | 13.77168569 | Superficial | SOX2 |
| 766 | SLC10A3 | 0.855190282 | 0.008904627 | 0.025990329 | 24.99035341 | 9.877241285 | Superficial | SOX2 |
| 767 | PDCD4 | 0.852743281 | 0.005105016 | 0.018023407 | 76.60735903 | 42.69342999 | Superficial | SOX2 |
| 768 | PHF14 | 0.852521794 | 0.002836074 | 0.012662188 | 29.36397933 | 9.593443001 | Superficial | SOX2 |
| 769 | HDAC1 | 0.851318097 | 0.000992312 | 0.008042817 | 61.95501145 | 30.23757095 | Superficial | SOX2 |
| 770 | FCHSD2 | 0.850963554 | 0.002836074 | 0.0176927 | 8.91420517 | 4.048490828 | Suprabasal | SOX2 + KLF5 |
| 771 | ERCC3 | 0.849248528 | 0.008904627 | 0.025990329 | 12.5092704 | 4.311489353 | Superficial | SOX2 |
| 772 | DMRTA2 | 0.848884765 | 0.002315114 | 0.016074975 | 6.741522978 | 3.19563976 | Suprabasal | SOX2 |
| 773 | ALCAM | 0.848190649 | 0.007423146 | 0.022959455 | 11.32639601 | 3.871626927 | Superficial | SOX2 + KLF5, SOX2 |
| 774 | SKIV2L | 0.848108426 | 0.000506425 | 0.007178523 | 13.76146259 | 4.464769501 | Superficial | SOX2 |
| 775 | DSC2 | 0.846921441 | 0.017811616 | 0.043066688 | 72.26288159 | 40.41740992 | Superficial | SOX2 |
| 776 | DTNBP1 | 0.846794959 | 0.021001058 | 0.04834985 | 14.28105129 | 4.932464707 | Superficial | SOX2 + KLF5 |
| 777 | BORCS5 | 0.846268858 | 0.002836074 | 0.012662188 | 20.25345521 | 7.504594465 | Superficial | SOX2 |
| 778 | POLE | 0.846155403 | 0.002836074 | 0.0176927 | 7.470452375 | 3.782665953 | Suprabasal | SOX2 |
| 779 | MAP3K8 | 0.845745236 | 0.004211401 | 0.027185186 | 25.65143559 | 12.18378813 | Basal | SOX2, SOX2 + KLF5 |
| 780 | RUFY3 | 0.844831691 | 0.000635975 | 0.007274975 | 44.04485076 | 16.50861145 | Superficial | SOX2 |
| 781 | ITGA3 | 0.844554837 | 0.007423146 | 0.030449933 | 13.76692359 | 6.266478229 | Suprabasal | SOX2 |
| 782 | ARFGEF2 | 0.844127629 | 0.002315114 | 0.011381143 | 41.32971032 | 15.23832937 | Superficial | SOX2 |
| 783 | SPNS1 | 0.842942886 | 0.003462059 | 0.014108599 | 21.89951967 | 7.858241511 | Superficial | SOX2 |
| 784 | UBR5 | 0.842481048 | 0.006166656 | 0.020290909 | 41.02552479 | 15.33512244 | Superficial | SOX2 |
| 785 | ALG9 | 0.839853938 | 0.007423146 | 0.022959455 | 10.62872224 | 4.26325388 | Superficial | SOX2 |
| 786 | MTHFD1L | 0.839822411 | 0.007423146 | 0.036548425 | 14.53056637 | 6.246920981 | Basal | SOX2 |
| 787 | CHD4 | 0.839801259 | 0.000635975 | 0.007274975 | 57.16082184 | 26.07656657 | Superficial | SOX2 |
| 788 | OAF | 0.839505259 | 0.003462059 | 0.019729589 | 13.15030459 | 5.79206902 | Suprabasal | SOX2 |
| 789 | MAP4 | 0.839469094 | 0.000506425 | 0.007178523 | 56.55506459 | 24.597045 | Superficial | SOX2 |
| 790 | GLE1 | 0.835935781 | 0.008904627 | 0.025990329 | 16.91032054 | 5.336535038 | Superficial | SOX2 |
| 791 | NECTIN4 | 0.835610283 | 0.000635975 | 0.012925379 | 30.87073029 | 15.59987036 | Basal | SOX2 |
| 792 | IWS1 | 0.83526127 | 0.001526458 | 0.009387421 | 33.72332735 | 12.23744966 | Superficial | SOX2 |
| 793 | RBM12 | 0.834785213 | 0.001232925 | 0.008679903 | 23.96776883 | 8.159807832 | Superficial | SOX2 |
| 794 | CCDC71L | 0.834591944 | 0.006166656 | 0.020290909 | 18.69169873 | 6.190759114 | Superficial | SOX2 |
| 795 | MACC1 | 0.834417685 | 0.001883197 | 0.018415354 | 24.81749513 | 10.86425508 | Basal | SOX2 + KLF5, SOX2 |
| 796 | HES1 | 0.834394053 | 0.015054848 | 0.038009884 | 72.77617354 | 40.31242283 | Superficial | SOX2 |
| 797 | DNMBP | 0.834004899 | 0.017811616 | 0.043066688 | 12.19837134 | 4.683669686 | Superficial | SOX2 + KLF5 |
| 798 | EFL1 | 0.833887149 | 0.002836074 | 0.012662188 | 12.54418382 | 4.708387837 | Superficial | SOX2 |
| 799 | YLPM1 | 0.833877819 | 0.004211401 | 0.016013117 | 21.20338158 | 7.420526934 | Superficial | SOX2 |
| 800 | PHC3 | 0.833132444 | 0.021001058 | 0.04834985 | 15.7324662 | 4.172159185 | Superficial | SOX2 |
| 801 | ARHGAP1 | 0.832801399 | 0.001526458 | 0.009387421 | 49.16523241 | 21.6705731 | Superficial | SOX2 |

| | | | | | | | |
|---|---|---|---|---|---|---|---|
| 802 | MTMR4 | 0.832773168 | 0.012681065 | 0.033478436 | 12.61282616 | 4.384877816 Superficial | SOX2 |
| 803 | USP34 | 0.831927479 | 0.000250281 | 0.007176147 | 48.76179144 | 19.59301972 Superficial | SOX2 |
| 804 | LTN1 | 0.831844928 | 0.003462059 | 0.014108599 | 32.27681339 | 10.69037507 Superficial | SOX2 |
| 805 | SAMD8 | 0.83084174 | 0.007423146 | 0.022959455 | 30.77720131 | 12.43241064 Superficial | SOX2 |
| 806 | LTBP4 | 0.830027435 | 0.000795824 | 0.010838271 | 42.65305007 | 22.45104814 Suprabasal | SOX2 |
| 807 | EFNB1 | 0.828770052 | 0.007423146 | 0.022959455 | 12.44857866 | 4.371969674 Superficial | SOX2 |
| 808 | TAF15 | 0.827829113 | 0.002836074 | 0.012662188 | 46.64494557 | 17.66507047 Superficial | SOX2 |
| 809 | ZNF205 | 0.827619182 | 0.006166656 | 0.020290909 | 9.321767583 | 3.501849539 Superficial | SOX2 |
| 810 | DSC3 | 0.82685136 | 0.021001058 | 0.04834985 | 60.40379599 | 27.05844558 Superficial | SOX2 + KLF5, SOX2 |
| 811 | TDG | 0.826493209 | 0.005105016 | 0.018023407 | 45.1714113 | 18.47106601 Superficial | SOX2 |
| 812 | PPA2 | 0.825614967 | 0.000795824 | 0.007615337 | 45.85643088 | 18.36050404 Superficial | SOX2 |
| 813 | EPB41L2 | 0.825187272 | 0.012681065 | 0.042921737 | 13.2310882 | 5.383598174 Suprabasal | SOX2 |
| 814 | RHOV | 0.824745801 | 0.000506425 | 0.012898798 | 55.32474773 | 31.36023103 Basal | SOX2 |
| 815 | THOC3 | 0.823897476 | 0.010644787 | 0.029507311 | 59.47882876 | 28.67750392 Superficial | SOX2 |
| 816 | TWF2 | 0.823549219 | 0.000992312 | 0.008042817 | 58.15254953 | 27.21826086 Superficial | SOX2 + KLF5 |
| 817 | ATRN | 0.823194799 | 0.021001058 | 0.04834985 | 17.37967553 | 6.135836441 Superficial | SOX2 |
| 818 | PPL | 0.822801611 | 0.000317696 | 0.012455402 | 57.78159745 | 33.11426547 Basal | SOX2, SOX2 + KLF5 |
| 819 | FBXO38 | 0.822800065 | 0.000992312 | 0.008042817 | 19.93874301 | 6.63278819 Superficial | SOX2 |
| 820 | MRPS35 | 0.82271128 | 0.012681065 | 0.033478436 | 41.47677578 | 15.25102828 Superficial | SOX2 |
| 821 | ACAD9 | 0.822605788 | 0.000401828 | 0.007176147 | 36.21095613 | 14.39931058 Superficial | SOX2 + KLF5 |
| 822 | SNX12 | 0.822585602 | 0.006166656 | 0.020290909 | 25.06938895 | 11.27638862 Superficial | SOX2 |
| 823 | TFAP2A | 0.820450997 | 0.002315114 | 0.011381143 | 42.82071234 | 17.82890683 Superficial | SOX2 |
| 824 | ATN1 | 0.819507951 | 0.000992312 | 0.008042817 | 32.18520156 | 12.76600372 Superficial | SOX2 |
| 825 | CPT2 | 0.818968522 | 0.015054848 | 0.038009884 | 23.64381963 | 8.697933431 Superficial | SOX2 |
| 826 | SPTLC2 | 0.817816947 | 0.008904627 | 0.025990329 | 58.58796534 | 29.63327216 Superficial | SOX2 |
| 827 | HACE1 | 0.816745618 | 0.008904627 | 0.034252903 | 13.23797985 | 6.170834204 Suprabasal | SOX2 |
| 828 | PIM1 | 0.816682613 | 0.000506425 | 0.012898798 | 35.4644167 | 17.60863066 Basal | SOX2 + KLF5, SOX2 |
| 829 | DHX16 | 0.816455155 | 0.000992312 | 0.008042817 | 20.49134581 | 7.234232442 Superficial | SOX2 |
| 830 | DNAAF2 | 0.815990058 | 0.008904627 | 0.025990329 | 12.88862781 | 5.011421676 Superficial | SOX2 |
| 831 | SDC1 | 0.813719129 | 0.002315114 | 0.011381143 | 87.75694466 | 66.95145226 Superficial | SOX2 |
| 832 | ELL3 | 0.813317704 | 0.015054848 | 0.048140298 | 7.192728655 | 3.329046936 Suprabasal | SOX2 |
| 833 | CCNE1 | 0.812722988 | 0.001883197 | 0.018415354 | 11.8773326 | 5.413782807 Basal | SOX2 |
| 834 | PLXNB2 | 0.812108846 | 0.001232925 | 0.015721789 | 46.65187659 | 22.95899987 Basal | SOX2 |
| 835 | WDR47 | 0.811734977 | 5.55E-05 | 0.012455402 | 35.21788058 | 15.80573817 Basal | SOX2 |
| 836 | MID1IP1 | 0.810782697 | 0.002836074 | 0.012662188 | 33.06994348 | 13.45685532 Superficial | SOX2 |
| 837 | MFN1 | 0.810406977 | 0.000795824 | 0.007615337 | 37.4800731 | 14.78617492 Superficial | SOX2 |
| 838 | EFNA4 | 0.809859537 | 0.003462059 | 0.014108599 | 21.55207442 | 7.257920127 Superficial | SOX2 |
| 839 | ARMCX3 | 0.808889184 | 0.001883197 | 0.010234018 | 22.49931016 | 7.20222225 Superficial | SOX2 |
| 840 | DDX46 | 0.808617792 | 0.001883197 | 0.010234018 | 55.8352134 | 23.18074829 Superficial | SOX2 |
| 841 | NXF1 | 0.808612993 | 0.021001058 | 0.04834985 | 18.32360737 | 7.616265295 Superficial | SOX2 |
| 842 | NDUFV1 | 0.808605636 | 0.008904627 | 0.025990329 | 58.69079218 | 25.16361058 Superficial | SOX2 |
| 843 | KIF1C | 0.808132244 | 0.000401828 | 0.012455402 | 35.31714962 | 16.56280017 Basal | SOX2 |
| 844 | CDKN2D | 0.807747808 | 0.017811616 | 0.043066688 | 31.18476328 | 12.50599597 Superficial | SOX2 |
| 845 | TMED5 | 0.807701462 | 0.001883197 | 0.010234018 | 40.71109999 | 16.55487528 Superficial | SOX2 |
| 846 | ANKRD22 | 0.804461386 | 0.001526458 | 0.009387421 | 53.90767745 | 23.26610733 Superficial | SOX2 |
| 847 | MCPH1 | 0.800090408 | 0.012681065 | 0.033478436 | 17.43680128 | 6.513088304 Superficial | SOX2 |
| 848 | FNDC3A | 0.800001038 | 0.004211401 | 0.016013117 | 20.19240284 | 7.23045575 Superficial | SOX2 |
| 849 | SIRT1 | 0.799938477 | 0.007423146 | 0.022959455 | 14.80977543 | 4.692575729 Superficial | SOX2 |
| 850 | LRPPRC | 0.798944589 | 0.021001058 | 0.04834985 | 31.40078505 | 9.816116635 Superficial | SOX2 |
| 851 | ATP5F1A | 0.798738249 | 0.006166656 | 0.020290909 | 71.23746796 | 38.86356897 Superficial | SOX2 |
| 852 | CPSF2 | 0.798423626 | 0.000992312 | 0.008042817 | 34.48820073 | 13.36730562 Superficial | SOX2 |
| 853 | UBALD2 | 0.798313434 | 0.000250281 | 0.012455402 | 58.45055428 | 32.53669687 Basal | SOX2 |
| 854 | SFPQ | 0.797818023 | 0.001232925 | 0.008679903 | 61.97537887 | 29.57785022 Superficial | SOX2 |
| 855 | SLC52A2 | 0.797173775 | 0.010644787 | 0.029507311 | 28.79074487 | 10.90435535 Superficial | SOX2 |
| 856 | NFRKB | 0.797033244 | 0.004211401 | 0.016013117 | 12.0529207 | 4.844985206 Superficial | SOX2 |
| 857 | ABHD13 | 0.796556468 | 0.000992312 | 0.008042817 | 22.41406526 | 8.264951864 Superficial | SOX2 |
| 858 | ARHGAP21 | 0.795336566 | 0.004211401 | 0.021879278 | 36.03350368 | 18.17611891 Suprabasal | SOX2 |
| 859 | EVPL | 0.793544001 | 0.000795824 | 0.0136425 | 44.99753962 | 23.63771042 Basal | SOX2 |
| 860 | MSH3 | 0.793458301 | 0.003462059 | 0.014108599 | 22.98398105 | 7.918764414 Superficial | SOX2 |
| 861 | ARL1 | 0.79214833 | 0.001232925 | 0.008679903 | 71.80733536 | 42.39486522 Superficial | SOX2 |
| 862 | KCTD1 | 0.790819901 | 0.000795824 | 0.007615337 | 50.90920202 | 21.19738831 Superficial | SOX2, SOX2 + KLF5 |
| 863 | FAM199X | 0.790489698 | 0.000992312 | 0.008042817 | 33.68678037 | 12.41133265 Superficial | SOX2 |
| 864 | IPPK | 0.790086212 | 0.021001058 | 0.04834985 | 32.36939208 | 11.4416446 Superficial | SOX2 |
| 865 | GRIPAP1 | 0.790033602 | 0.000992312 | 0.008042817 | 32.41525033 | 11.48465582 Superficial | SOX2 |
| 866 | IGSF8 | 0.789619705 | 0.002315114 | 0.011381143 | 14.91047128 | 5.385316998 Superficial | SOX2 |
| 867 | SMARCE1 | 0.788128386 | 0.000401828 | 0.007176147 | 53.98102131 | 23.8797572 Superficial | SOX2 |
| 868 | PAK1 | 0.786995191 | 0.000635975 | 0.007274975 | 29.64145241 | 11.08264459 Superficial | SOX2 |
| 869 | SLC22A18 | 0.786650694 | 0.000992312 | 0.008042817 | 27.90101417 | 9.786049759 Superficial | SOX2 |
| 870 | SCRIB | 0.785180723 | 0.000317696 | 0.012455402 | 24.07048471 | 10.71812052 Basal | SOX2 |
| 871 | PPP2R1B | 0.785024498 | 0.002836074 | 0.012662188 | 28.21036033 | 10.75997223 Superficial | SOX2 |
| 872 | KLK10 | 0.784915262 | 0.001883197 | 0.018415354 | 49.63143813 | 30.40373894 Basal | SOX2 |
| 873 | ESRP2 | 0.784736975 | 0.003462059 | 0.014108599 | 49.49809927 | 23.21995429 Superficial | SOX2 |
| 874 | DSP | 0.78419116 | 0.000506425 | 0.012898798 | 95.57120634 | 83.16967388 Basal | SOX2, SOX2 + KLF5 |

| | | | | | | | | |
|---|---|---|---|---|---|---|---|---|
| 875 | SUPT6H | 0.783663486 | 0.001883197 | 0.010234018 | 39.09967722 | 17.04241278 | Superficial | SOX2 |
| 876 | NSD1 | 0.782623544 | 0.000250281 | 0.007176147 | 44.34785098 | 18.99005533 | Superficial | SOX2 |
| 877 | RARG | 0.781083073 | 0.000401828 | 0.007176147 | 76.26745981 | 42.47873787 | Superficial | SOX2 |
| 878 | RASSF6 | 0.780974466 | 0.021001058 | 0.04834985 | 19.09196449 | 5.374914478 | Superficial | SOX2, SOX2 + KLF5 |
| 879 | TRIM5 | 0.780767742 | 0.021001058 | 0.04834985 | 14.72009644 | 4.341097256 | Superficial | SOX2 |
| 880 | ARHGAP35 | 0.779858557 | 0.000795824 | 0.007615337 | 30.72874296 | 11.10752697 | Superficial | SOX2 |
| 881 | TBRG4 | 0.777140895 | 0.017811616 | 0.043066688 | 23.73424314 | 8.528389369 | Superficial | SOX2 |
| 882 | TRABD | 0.776314911 | 0.006166656 | 0.020290909 | 44.64577932 | 19.65405638 | Superficial | SOX2 |
| 883 | ARMCX6 | 0.776127211 | 0.001883197 | 0.010234018 | 31.53888816 | 12.22350712 | Superficial | SOX2 |
| 884 | NCOA5 | 0.775988711 | 0.008904627 | 0.025990329 | 9.113392929 | 3.455771311 | Superficial | SOX2 |
| 885 | FRMD4B | 0.772823978 | 0.001526458 | 0.009387421 | 72.27208121 | 40.58876961 | Superficial | SOX2 |
| 886 | LAP3 | 0.771547699 | 0.001883197 | 0.010234018 | 40.8294852 | 15.63526895 | Superficial | SOX2 |
| 887 | CDC42BPB | 0.770069085 | 0.000153669 | 0.012455402 | 39.33363482 | 18.95615945 | Basal | SOX2 |
| 888 | SLC7A8 | 0.768503851 | 0.003462059 | 0.014108599 | 23.18110177 | 9.569188016 | Superficial | SOX2 |
| 889 | TMEM154 | 0.766978768 | 0.010644787 | 0.029507311 | 73.6246979 | 44.77279121 | Superficial | SOX2 |
| 890 | TXN | 0.766068419 | 0.017811616 | 0.043066688 | 94.92987078 | 90.59413398 | Superficial | SOX2 |
| 891 | CAAP1 | 0.765982227 | 0.001232925 | 0.008679903 | 27.8715948 | 10.35073339 | Superficial | SOX2 |
| 892 | ZNF407 | 0.765275288 | 0.005105016 | 0.018023407 | 9.565013872 | 2.98368203 | Superficial | SOX2 |
| 893 | TAB1 | 0.765243129 | 0.008904627 | 0.025990329 | 7.66350226 | 2.519377628 | Superficial | SOX2 |
| 894 | TPM4 | 0.765005693 | 0.002836074 | 0.022562695 | 83.37863428 | 60.677827 | Basal | SOX2 |
| 895 | UTP6 | 0.764059543 | 0.001883197 | 0.010234018 | 24.32650647 | 9.796163309 | Superficial | SOX2 |
| 896 | ZNF142 | 0.763854846 | 0.000795824 | 0.010838271 | 7.199945873 | 3.709384109 | Suprabasal | SOX2 |
| 897 | USP7 | 0.762927097 | 0.000506425 | 0.007178523 | 57.37381352 | 26.69855543 | Superficial | SOX2 |
| 898 | TKFC | 0.762788105 | 0.021001058 | 0.04834985 | 15.27432299 | 5.605121704 | Superficial | SOX2 |
| 899 | TTC17 | 0.761819095 | 0.008904627 | 0.025990329 | 24.36359453 | 8.844420798 | Superficial | SOX2 |
| 900 | PXK | 0.758536609 | 0.005105016 | 0.018023407 | 14.76121213 | 4.158985349 | Superficial | SOX2 |
| 901 | NMB | 0.757007026 | 0.003462059 | 0.024854712 | 19.49023009 | 8.735687673 | Basal | SOX2 |
| 902 | PIGK | 0.75685835 | 0.010644787 | 0.029507311 | 10.61173633 | 3.827762891 | Superficial | SOX2 |
| 903 | PRPF40A | 0.755048603 | 0.000795824 | 0.007615337 | 58.42226805 | 27.22883402 | Superficial | SOX2 |
| 904 | RIPK4 | 0.753786917 | 0.000506425 | 0.010489143 | 56.92429997 | 34.29619525 | Suprabasal | SOX2 |
| 905 | MARCKSL1 | 0.753039523 | 0.005105016 | 0.029792864 | 19.57397271 | 7.984022926 | Basal | SOX2 |
| 906 | COX10 | 0.75216747 | 0.017811616 | 0.043066688 | 15.53587995 | 5.187538716 | Superficial | SOX2 |
| 907 | XRN2 | 0.751931606 | 0.008904627 | 0.025990329 | 38.10142195 | 14.27251888 | Superficial | SOX2 |
| 908 | POR | 0.751814117 | 0.000992312 | 0.008042817 | 66.781586 | 35.01139862 | Superficial | KLF5, SOX2 |
| 909 | MECOM | 0.750699729 | 0.002315114 | 0.016074975 | 56.52334109 | 32.74640448 | Suprabasal | KLF5 |
| 910 | PLP2 | 0.749774242 | 0.010644787 | 0.029507311 | 77.34020689 | 49.58391191 | Superficial | SOX2 |
| 911 | RAB11FIP4 | 0.74958562 | 0.002836074 | 0.022562695 | 11.93939559 | 4.465728883 | Basal | SOX2 |
| 912 | CRADD | 0.748667913 | 0.007423146 | 0.022959455 | 14.26033086 | 4.924503899 | Superficial | SOX2 + KLF5 |
| 913 | ATXN7L3B | 0.748484548 | 0.000153669 | 0.007176147 | 44.03819642 | 18.55042579 | Superficial | SOX2 |
| 914 | GRIP1 | 0.747835615 | 0.003462059 | 0.019729589 | 7.134486236 | 3.06073658 | Suprabasal | SOX2 + KLF5 |
| 915 | NIPAL1 | 0.747293766 | 0.001883197 | 0.010234018 | 58.7054824 | 29.04320719 | Superficial | SOX2 |
| 916 | RASA1 | 0.747100481 | 0.003462059 | 0.014108599 | 25.67635873 | 8.934911271 | Superficial | SOX2 |
| 917 | MFSD6 | 0.746746359 | 0.012681065 | 0.033478436 | 53.60128769 | 24.41723396 | Superficial | SOX2 |
| 918 | INTS2 | 0.746668773 | 0.010644787 | 0.029507311 | 8.778939305 | 2.442925882 | Superficial | SOX2 |
| 919 | CCDC34 | 0.746367735 | 0.001526458 | 0.013290882 | 40.02681251 | 19.40977035 | Suprabasal | SOX2 |
| 920 | UBR7 | 0.74631484 | 0.012681065 | 0.033478436 | 16.05246902 | 5.706209921 | Superficial | SOX2 |
| 921 | SNTB2 | 0.745802206 | 0.001883197 | 0.010234018 | 27.06126241 | 9.939822852 | Superficial | SOX2 |
| 922 | MAN1A2 | 0.744280933 | 0.001883197 | 0.010234018 | 50.53089819 | 20.95246014 | Superficial | SOX2 |
| 923 | NDUFA8 | 0.743813982 | 0.004211401 | 0.016013117 | 56.18648971 | 26.12503468 | Superficial | SOX2 |
| 924 | EIF3B | 0.74245088 | 0.005105016 | 0.018023407 | 43.13425729 | 17.85244483 | Superficial | SOX2 |
| 925 | POLR3B | 0.74062025 | 0.000506425 | 0.007178523 | 21.82467954 | 8.464220903 | Superficial | SOX2 |
| 926 | WNK4 | 0.739616678 | 0.008904627 | 0.025990329 | 16.96339688 | 6.600647357 | Superficial | SOX2 |
| 927 | MRPS30 | 0.739498814 | 0.010644787 | 0.029507311 | 26.85606255 | 9.562664138 | Superficial | SOX2 |
| 928 | RBBP8NL | 0.738379695 | 0.008904627 | 0.025990329 | 18.58587441 | 7.808829489 | Superficial | SOX2 |
| 929 | ARFGAP3 | 0.737414745 | 0.004211401 | 0.016013117 | 34.43334593 | 13.93679235 | Superficial | SOX2 + KLF5 |
| 930 | PRRC2B | 0.737288087 | 0.004211401 | 0.016013117 | 37.54429978 | 15.16190471 | Superficial | SOX2 |
| 931 | CLIC4 | 0.737162028 | 0.007423146 | 0.036548425 | 18.68317984 | 8.50586959 | Basal | SOX2 |
| 932 | AFDN | 0.736654672 | 5.55E-05 | 0.012455402 | 48.38939108 | 25.14959746 | Basal | SOX2 |
| 933 | PHF13 | 0.73612879 | 0.000635975 | 0.012925379 | 18.94767185 | 8.609546677 | Basal | SOX2 |
| 934 | ZYX | 0.735779973 | 0.015054848 | 0.038009884 | 36.28202793 | 16.07075639 | Superficial | SOX2 |
| 935 | TMEM98 | 0.735476257 | 0.008904627 | 0.034252903 | 30.33653877 | 14.52821983 | Suprabasal | SOX2 |
| 936 | BICDL2 | 0.735059923 | 0.002315114 | 0.020052792 | 22.07254004 | 10.57748777 | Basal | SOX2 |
| 937 | PRC1 | 0.734220942 | 0.002836074 | 0.022562695 | 44.06968575 | 25.03313541 | Basal | SOX2 |
| 938 | ZNF131 | 0.733472982 | 0.002315114 | 0.011381143 | 25.08109627 | 8.963884002 | Superficial | SOX2 |
| 939 | PIK3R1 | 0.733066436 | 0.007423146 | 0.022959455 | 19.32332041 | 6.742500994 | Superficial | SOX2 |
| 940 | BST2 | 0.732636317 | 0.012681065 | 0.049425568 | 41.46014877 | 20.02884409 | Basal | SOX2 |
| 941 | MGST3 | 0.732351335 | 0.010644787 | 0.029507311 | 65.51241176 | 32.75084606 | Superficial | SOX2 |
| 942 | HTATIP2 | 0.732198546 | 0.005105016 | 0.018023407 | 54.11311985 | 24.77233011 | Superficial | SOX2 |
| 943 | ZPR1 | 0.730854063 | 0.017811616 | 0.043066688 | 28.17455576 | 10.73635346 | Superficial | SOX2 |
| 944 | URI1 | 0.730252363 | 0.000795824 | 0.007615337 | 48.99660551 | 20.34154253 | Superficial | SOX2 |
| 945 | EZR | 0.72742657 | 0.002315114 | 0.020052792 | 82.97696886 | 59.68243158 | Basal | SOX2 + KLF5, SOX2 |
| 946 | TMEM184A | 0.726810155 | 0.006166656 | 0.032981404 | 16.76278201 | 7.803146974 | Basal | SOX2 |
| 947 | GRB7 | 0.726375743 | 0.000317696 | 0.012455402 | 39.18241856 | 19.55517725 | Basal | SOX2 |

| | | | | | | | | |
|---|---|---|---|---|---|---|---|---|
| 948 | MRPL4 | 0.726193947 | 0.008904627 | 0.025990329 | 43.49259063 | 17.67791629 | Superficial | SOX2 |
| 949 | AKAP13 | 0.72614015 | 0.000635975 | 0.007274975 | 69.84387281 | 38.37719104 | Superficial | SOX2 |
| 950 | ICMT | 0.726052153 | 0.000992312 | 0.014320968 | 39.13219758 | 20.3780701 | Basal | SOX2 |
| 951 | G3BP1 | 0.72578975 | 0.017811616 | 0.043066688 | 45.8882534 | 18.82497802 | Superficial | SOX2 |
| 952 | INPP5F | 0.725043383 | 0.007423146 | 0.022959455 | 13.93767328 | 4.990127193 | Superficial | SOX2 |
| 953 | RNF138 | 0.724834032 | 0.017811616 | 0.043066688 | 16.12579921 | 5.149135029 | Superficial | SOX2 |
| 954 | ETV3 | 0.724257442 | 0.006166656 | 0.027303057 | 36.51993618 | 18.36824247 | Suprabasal | SOX2 |
| 955 | PBDC1 | 0.724096304 | 0.007423146 | 0.022959455 | 47.58340403 | 19.4017906 | Superficial | SOX2 |
| 956 | TBC1D4 | 0.722873499 | 0.017811616 | 0.043066688 | 9.618053519 | 3.152159836 | Superficial | SOX2 |
| 957 | WWC2 | 0.722704636 | 0.000635975 | 0.012925379 | 14.52289953 | 5.932481265 | Basal | SOX2 |
| 958 | CD82 | 0.721546364 | 0.012681065 | 0.033478436 | 57.52904438 | 27.31531055 | Superficial | SOX2 |
| 959 | PFKP | 0.720566334 | 0.002836074 | 0.022562695 | 42.15295301 | 21.9314053 | Basal | SOX2 |
| 960 | GTF3C5 | 0.72007063 | 0.001526458 | 0.009387421 | 28.57071781 | 12.03709938 | Superficial | SOX2 |
| 961 | SUGT1 | 0.719687512 | 0.001232925 | 0.008679903 | 60.55400256 | 28.61191437 | Superficial | SOX2 |
| 962 | EIF4G1 | 0.718208604 | 0.000635975 | 0.012925379 | 63.74988687 | 36.77598904 | Basal | SOX2 |
| 963 | RHBDD2 | 0.717960761 | 0.000992312 | 0.01142308 | 43.30584464 | 25.55995337 | Suprabasal | SOX2 |
| 964 | RAB3IP | 0.717948673 | 0.001232925 | 0.012246384 | 18.14985456 | 8.781444715 | Suprabasal | SOX2 |
| 965 | GOPC | 0.717192669 | 0.000506425 | 0.007178523 | 40.99698046 | 17.8396565 | Superficial | SOX2 |
| 966 | CA13 | 0.716964043 | 0.012681065 | 0.033478436 | 26.84068309 | 9.406202184 | Superficial | SOX2 |
| 967 | FOXP4 | 0.716400668 | 0.000635975 | 0.012925379 | 10.1266356 | 4.367339639 | Basal | SOX2 |
| 968 | DIAPH1 | 0.715897124 | 0.000992312 | 0.014320968 | 49.98329195 | 26.85333151 | Basal | SOX2 |
| 969 | MPHOSPH10 | 0.715736333 | 0.006166656 | 0.020290909 | 41.49656213 | 16.38485875 | Superficial | SOX2 |
| 970 | PPIC | 0.715013805 | 0.015054848 | 0.038009884 | 68.41924217 | 34.96201968 | Superficial | SOX2 |
| 971 | KHDRBS1 | 0.714588018 | 0.000401828 | 0.007176147 | 62.73550413 | 29.47083304 | Superficial | SOX2 |
| 972 | UTP3 | 0.714515084 | 0.006166656 | 0.020290909 | 25.70999785 | 9.52559203 | Superficial | SOX2 |
| 973 | SPEN | 0.713127731 | 0.002836074 | 0.012662188 | 48.98677362 | 21.73061439 | Superficial | SOX2 |
| 974 | KREMEN1 | 0.713062714 | 0.000401828 | 0.012455402 | 51.12077132 | 27.52790114 | Basal | SOX2 |
| 975 | TMEM147 | 0.712669208 | 0.007423146 | 0.022959455 | 72.89427386 | 49.57190643 | Superficial | SOX2 |
| 976 | TOM1L1 | 0.712554624 | 0.004211401 | 0.016013117 | 23.61587939 | 8.747457616 | Superficial | SOX2 |
| 977 | TECPR2 | 0.712470979 | 9.30E-05 | 0.010489143 | 13.29861592 | 6.763670291 | Suprabasal | SOX2 |
| 978 | NR6A1 | 0.711946773 | 0.002315114 | 0.016074975 | 14.65826514 | 7.063050314 | Suprabasal | SOX2 |
| 979 | KHDRBS3 | 0.710487783 | 0.000635975 | 0.012925379 | 11.98724433 | 5.379939887 | Basal | SOX2 + KLF5 |
| 980 | GTF3C2 | 0.70899392 | 0.002836074 | 0.012662188 | 27.90357716 | 11.75661752 | Superficial | SOX2 |
| 981 | SETD7 | 0.708207417 | 0.000635975 | 0.007274975 | 27.99014587 | 13.201659 | Superficial | SOX2 |
| 982 | PARD6G | 0.708078019 | 0.004211401 | 0.016013117 | 44.41779238 | 18.59663141 | Superficial | SOX2 |
| 983 | LASP1 | 0.70727919 | 0.001526458 | 0.016792744 | 36.89944035 | 16.6859833 | Basal | SOX2 |
| 984 | MIPEP | 0.707166754 | 0.021001058 | 0.04834985 | 10.893853 | 3.702076167 | Superficial | SOX2 + KLF5, SOX2 |
| 985 | LEMD3 | 0.706801908 | 0.006166656 | 0.020290909 | 18.31855904 | 5.567001364 | Superficial | SOX2 |
| 986 | IER3 | 0.705746954 | 0.005105016 | 0.029792864 | 91.0662064 | 71.33517237 | Basal | SOX2 |
| 987 | TM9SF3 | 0.705468786 | 0.000317696 | 0.007176147 | 75.11455963 | 44.74854753 | Superficial | SOX2 |
| 988 | TPST2 | 0.705368445 | 0.008904627 | 0.025990329 | 29.58094075 | 12.14561785 | Superficial | SOX2 |
| 989 | MAST4 | 0.70518553 | 0.015054848 | 0.038009884 | 60.88267442 | 33.03507244 | Superficial | SOX2 |
| 990 | PPP1R3D | 0.705122846 | 0.001883197 | 0.010234018 | 23.83506292 | 9.135189693 | Superficial | SOX2 |
| 991 | SENP6 | 0.703318284 | 0.001526458 | 0.009387421 | 39.02975419 | 15.42974016 | Superficial | SOX2 |
| 992 | FAM83H | 0.702860491 | 0.002315114 | 0.020052792 | 35.23918804 | 17.10164149 | Basal | SOX2 |
| 993 | HIP1R | 0.702694987 | 0.001526458 | 0.016792744 | 11.04972321 | 4.742867003 | Basal | SOX2 |
| 994 | USP40 | 0.702333154 | 0.006166656 | 0.027303057 | 29.51910976 | 16.18770834 | Suprabasal | SOX2 |
| 995 | PPARA | 0.702123681 | 0.000153669 | 0.010489143 | 30.47842317 | 16.4040569 | Suprabasal | SOX2 |
| 996 | NECTIN2 | 0.701897768 | 0.000250281 | 0.012455402 | 33.17085744 | 15.91725556 | Basal | SOX2 |
| 997 | XPO7 | 0.701674727 | 0.012681065 | 0.033478436 | 17.37725907 | 6.268661514 | Superficial | SOX2 |
| 998 | MUS81 | 0.701670445 | 0.008904627 | 0.025990329 | 10.32278069 | 4.818304548 | Superficial | SOX2 |
| 999 | YRDC | 0.69921272 | 0.003462059 | 0.014108599 | 21.51501832 | 8.558992813 | Superficial | SOX2 |
| 1000 | BRD1 | 0.699059598 | 0.002315114 | 0.016074975 | 42.85777072 | 23.81696503 | Suprabasal | SOX2 |
| 1001 | SEMA4D | 0.699049431 | 0.006166656 | 0.020290909 | 29.50955164 | 12.78839383 | Superficial | SOX2 |
| 1002 | TRUB2 | 0.698692417 | 0.017811616 | 0.043066688 | 28.49174933 | 11.26687257 | Superficial | SOX2 |
| 1003 | SEC63 | 0.698476952 | 0.012681065 | 0.033478436 | 36.80728048 | 14.45232384 | Superficial | SOX2 |
| 1004 | ATP5F1D | 0.698358337 | 0.007423146 | 0.022959455 | 82.5012794 | 63.14211755 | Superficial | SOX2 |
| 1005 | AJUBA | 0.697980755 | 0.015054848 | 0.048140298 | 24.81183555 | 12.03369669 | Suprabasal | SOX2 |
| 1006 | RWDD4 | 0.696875045 | 0.000401828 | 0.007176147 | 51.77094435 | 24.11983066 | Superficial | SOX2 |
| 1007 | NACC1 | 0.696463116 | 0.000506425 | 0.012898798 | 20.19965077 | 9.054914034 | Basal | SOX2 |
| 1008 | FLNA | 0.695574276 | 0.001883197 | 0.018415354 | 68.91808084 | 43.431091 | Basal | SOX2 |
| 1009 | SHROOM3 | 0.695335913 | 0.007423146 | 0.036548425 | 9.574191785 | 3.812745015 | Basal | SOX2 + KLF5, SOX2 |
| 1010 | SLTM | 0.695313462 | 0.000506425 | 0.007178523 | 63.31126801 | 31.76210791 | Superficial | SOX2 |
| 1011 | SCAPER | 0.694072236 | 0.007423146 | 0.022959455 | 27.74667976 | 10.29388181 | Superficial | SOX2 |
| 1012 | MON1B | 0.693303248 | 0.006166656 | 0.020290909 | 27.61617344 | 11.3071686 | Superficial | SOX2 |
| 1013 | NPC1 | 0.692962549 | 0.004211401 | 0.027185186 | 15.36013339 | 6.703777517 | Basal | SOX2 |
| 1014 | TAX1BP3 | 0.692447302 | 0.005105016 | 0.018023407 | 78.13399476 | 51.10089037 | Superficial | SOX2 |
| 1015 | SGMS1 | 0.691297817 | 0.000635975 | 0.010733065 | 29.86271925 | 15.19705323 | Suprabasal | SOX2 + KLF5 |
| 1016 | SDC4 | 0.690969578 | 0.001883197 | 0.018415354 | 63.26780819 | 36.20162181 | Basal | SOX2 |
| 1017 | RING1 | 0.690902998 | 0.002836074 | 0.012662188 | 22.65387288 | 9.21827708 | Superficial | SOX2 |
| 1018 | SUSD4 | 0.688905232 | 0.006166656 | 0.020290909 | 48.73984961 | 21.34803969 | Superficial | KLF5, SOX2 |
| 1019 | POLR3D | 0.687357327 | 0.003462059 | 0.014108599 | 20.8707821 | 7.43613414 | Superficial | SOX2 |
| 1020 | PSMB4 | 0.685299892 | 0.001526458 | 0.009387421 | 58.61299947 | 27.96010638 | Superficial | SOX2 |

| | | | | | | | | |
|---|---|---|---|---|---|---|---|---|
| 1021 | HSPB1 | 0.684800626 | 0.012681065 | 0.033478436 | 98.45290194 | 93.43259814 | Superficial | SOX2 |
| 1022 | EGFR | 0.684370469 | 0.010644787 | 0.029507311 | 41.9502083 | 15.63246924 | Superficial | SOX2 |
| 1023 | GTF3C1 | 0.683517925 | 0.004211401 | 0.016013117 | 60.9092851 | 33.78517881 | Superficial | SOX2 |
| 1024 | ADGRL2 | 0.683192397 | 0.008904627 | 0.040392504 | 10.7417921 | 4.139408272 | Basal | SOX2 |
| 1025 | HDAC4 | 0.682742832 | 0.015054848 | 0.038009884 | 13.30421243 | 4.220842783 | Superficial | SOX2 |
| 1026 | CEP55 | 0.682070144 | 0.001232925 | 0.015721789 | 33.14551567 | 19.35039483 | Basal | SOX2 |
| 1027 | ASH2L | 0.681737225 | 0.001883197 | 0.010234018 | 23.09281134 | 8.409669845 | Superficial | SOX2 |
| 1028 | USP37 | 0.681305269 | 0.000992312 | 0.01142308 | 15.85613317 | 8.095477402 | Suprabasal | SOX2 |
| 1029 | COBL | 0.681239795 | 0.015054848 | 0.038009884 | 31.42195234 | 12.03903453 | Superficial | SOX2 |
| 1030 | CDC42SE2 | 0.679917135 | 0.017811616 | 0.043066688 | 13.52002582 | 5.114852508 | Superficial | SOX2 |
| 1031 | LSM14B | 0.678316094 | 0.012681065 | 0.033478436 | 21.66978744 | 8.709179062 | Superficial | SOX2 |
| 1032 | KLHL21 | 0.677983908 | 0.000401828 | 0.012455402 | 38.21633848 | 18.43710045 | Basal | SOX2 |
| 1033 | CASP7 | 0.677572835 | 0.004211401 | 0.021879278 | 32.86386227 | 16.70000183 | Suprabasal | SOX2 |
| 1034 | CCSAP | 0.675473422 | 0.000250281 | 0.012455402 | 15.54470514 | 7.578541078 | Basal | SOX2 |
| 1035 | BRI3BP | 0.675241931 | 0.017811616 | 0.043066688 | 27.70199565 | 10.31740369 | Superficial | SOX2 |
| 1036 | ELP3 | 0.674993585 | 0.007423146 | 0.022959455 | 12.44484451 | 4.645805987 | Superficial | SOX2 |
| 1037 | XPNPEP1 | 0.673875063 | 0.012681065 | 0.033478436 | 22.6153257 | 8.807703763 | Superficial | SOX2 |
| 1038 | TSPAN31 | 0.673329211 | 0.007423146 | 0.022959455 | 32.32370625 | 14.57475959 | Superficial | SOX2 |
| 1039 | GATAD1 | 0.672889383 | 9.30E-05 | 0.007176147 | 33.34992132 | 13.42729982 | Superficial | SOX2 |
| 1040 | SBNO1 | 0.67149385 | 0.001232925 | 0.008679903 | 39.14640306 | 15.55429681 | Superficial | SOX2 |
| 1041 | TTC23 | 0.670929937 | 0.004211401 | 0.016013117 | 11.357851 | 5.074134562 | Superficial | SOX2 |
| 1042 | TBRG1 | 0.670360458 | 9.30E-05 | 0.007176147 | 35.02731125 | 14.25893381 | Superficial | SOX2 |
| 1043 | MYO1D | 0.670223138 | 0.001883197 | 0.010234018 | 48.98233231 | 23.24023778 | Superficial | SOX2 |
| 1044 | AGAP3 | 0.669761063 | 0.001232925 | 0.008679903 | 41.32602591 | 18.75107029 | Superficial | SOX2 |
| 1045 | DAXX | 0.668789997 | 0.003462059 | 0.014108599 | 30.21577035 | 12.06403782 | Superficial | SOX2 |
| 1046 | YKT6 | 0.668035629 | 0.001526458 | 0.016792744 | 26.64004822 | 12.79324033 | Basal | SOX2 |
| 1047 | CDK17 | 0.668015169 | 0.003462059 | 0.019729589 | 16.41360549 | 8.270330179 | Suprabasal | SOX2 |
| 1048 | MAL2 | 0.666971971 | 0.000992312 | 0.014320968 | 67.6656544 | 46.069602 | Basal | SOX2, SOX2 + KLF5 |
| 1049 | LGALS8 | 0.666036557 | 0.005105016 | 0.018023407 | 31.07031235 | 15.28461671 | Superficial | SOX2 |
| 1050 | SLC44A2 | 0.665545253 | 0.007423146 | 0.022959455 | 61.18924802 | 32.86938981 | Superficial | SOX2 |
| 1051 | ZBTB17 | 0.665396487 | 0.000795824 | 0.007615337 | 15.53672061 | 6.058259512 | Superficial | SOX2 |
| 1052 | RABAC1 | 0.665008765 | 0.006166656 | 0.020290909 | 73.60564023 | 45.0711054 | Superficial | SOX2 |
| 1053 | TNFRSF10B | 0.663928537 | 0.012681065 | 0.042921737 | 8.142564324 | 4.107895104 | Suprabasal | SOX2 |
| 1054 | PHF10 | 0.663225937 | 0.001232925 | 0.012246384 | 57.30647758 | 35.85848239 | Suprabasal | SOX2 |
| 1055 | CDC25A | 0.662365282 | 0.003462059 | 0.024854712 | 11.09681846 | 5.301737689 | Basal | SOX2 |
| 1056 | JOSD2 | 0.661599184 | 0.001883197 | 0.010234018 | 65.70995211 | 35.94558676 | Superficial | SOX2 |
| 1057 | TAF6 | 0.661561026 | 0.001526458 | 0.009387421 | 16.25357545 | 6.858573155 | Superficial | SOX2 |
| 1058 | PSMC2 | 0.661474033 | 0.008904627 | 0.025990329 | 48.13192169 | 20.69490945 | Superficial | SOX2 |
| 1059 | INTS10 | 0.661149836 | 0.021001058 | 0.04834985 | 25.09640797 | 9.976829546 | Superficial | SOX2 |
| 1060 | NUP50 | 0.660882124 | 0.010644787 | 0.038255467 | 53.95810736 | 32.51693783 | Suprabasal | SOX2 |
| 1061 | B3GNT2 | 0.659308899 | 0.015054848 | 0.038009884 | 48.76608833 | 22.08346533 | Superficial | SOX2 + KLF5, SOX2 |
| 1062 | TEAD4 | 0.65922919 | 0.000795824 | 0.0136425 | 17.22245281 | 8.132829946 | Basal | SOX2 |
| 1063 | MIDN | 0.658466962 | 0.010644787 | 0.029507311 | 76.35570508 | 51.43854435 | Superficial | SOX2 |
| 1064 | IL13RA1 | 0.658415331 | 0.000506425 | 0.012898798 | 29.98096386 | 15.03877761 | Basal | SOX2 |
| 1065 | PRR14L | 0.658376186 | 0.000506425 | 0.007178523 | 42.93223762 | 17.25211076 | Superficial | SOX2 |
| 1066 | AARS2 | 0.658110216 | 0.010644787 | 0.029507311 | 10.38037111 | 3.661738738 | Superficial | SOX2 |
| 1067 | LRP6 | 0.656261512 | 0.001883197 | 0.014679007 | 10.47197382 | 5.225409838 | Suprabasal | SOX2 |
| 1068 | SLC2A1 | 0.655554397 | 0.007423146 | 0.022959455 | 54.5759334 | 27.05487701 | Superficial | SOX2 |
| 1069 | RBMXL1 | 0.655488128 | 0.021001058 | 0.04834985 | 21.07186616 | 8.056498063 | Superficial | SOX2 |
| 1070 | ARHGEF3 | 0.655422067 | 0.005105016 | 0.018023407 | 22.4596444 | 8.620999995 | Superficial | SOX2 |
| 1071 | TPP2 | 0.654015025 | 0.004211401 | 0.016013117 | 29.24394806 | 12.0423832 | Superficial | SOX2 |
| 1072 | AKAP11 | 0.653966626 | 0.017811616 | 0.043066688 | 21.36105886 | 7.919796639 | Superficial | SOX2 |
| 1073 | MYO6 | 0.653027843 | 0.001526458 | 0.016792744 | 43.91930761 | 23.10721262 | Basal | SOX2 |
| 1074 | PHF6 | 0.652696533 | 0.005105016 | 0.018023407 | 25.04223587 | 10.11666217 | Superficial | SOX2 |
| 1075 | MAFK | 0.650414578 | 0.010644787 | 0.044687249 | 14.57536945 | 6.225943242 | Basal | SOX2 |
| 1076 | LLGL2 | 0.649878814 | 0.000401828 | 0.012455402 | 35.39044093 | 19.03596924 | Basal | SOX2 |
| 1077 | ZNF146 | 0.649786015 | 0.021001058 | 0.04834985 | 16.61304701 | 5.79814609 | Superficial | SOX2 |
| 1078 | SLC9A3R1 | 0.649284456 | 0.000317696 | 0.012455402 | 70.56397011 | 47.30720979 | Basal | SOX2 |
| 1079 | COPB1 | 0.649171708 | 0.000635975 | 0.007274975 | 59.26939583 | 29.49107963 | Superficial | SOX2 |
| 1080 | CXXC5 | 0.649012341 | 0.000635975 | 0.012925379 | 21.53093653 | 10.79801587 | Basal | SOX2 |
| 1081 | NFIA | 0.64874555 | 0.002836074 | 0.012662188 | 53.7574233 | 26.49796931 | Superficial | SOX2 |
| 1082 | SNAI2 | 0.648425085 | 0.015054848 | 0.048140298 | 12.43176037 | 6.318331755 | Suprabasal | SOX2 |
| 1083 | PIP4K2C | 0.648192568 | 0.000401828 | 0.012455402 | 18.10247521 | 8.037186698 | Basal | SOX2 |
| 1084 | ILVBL | 0.648089774 | 0.021001058 | 0.04834985 | 48.24008432 | 20.18812239 | Superficial | SOX2 + KLF5, SOX2 |
| 1085 | PRKD2 | 0.647914132 | 0.000992312 | 0.014320968 | 23.08315078 | 10.53299073 | Basal | SOX2 |
| 1086 | GRHL1 | 0.647828984 | 0.002315114 | 0.020052792 | 17.11618921 | 8.766363269 | Basal | SOX2 |
| 1087 | CTR9 | 0.647808145 | 0.001883197 | 0.010234018 | 46.14068978 | 21.288081 | Superficial | SOX2 |
| 1088 | MRPS5 | 0.647420793 | 0.000795824 | 0.007615337 | 49.35732999 | 23.18354163 | Superficial | SOX2 |
| 1089 | GCH1 | 0.647126938 | 0.002836074 | 0.022562695 | 14.58129743 | 6.129531195 | Basal | SOX2 |
| 1090 | DNAJC2 | 0.646987291 | 0.021001058 | 0.04834985 | 25.67749226 | 9.788446426 | Superficial | SOX2 |
| 1091 | CRTC2 | 0.646003988 | 0.004211401 | 0.027185186 | 14.0923151 | 5.928055588 | Basal | SOX2 |
| 1092 | BCL7C | 0.64482131 | 0.007423146 | 0.022959455 | 43.10442673 | 18.3022768 | Superficial | SOX2 |
| 1093 | KLF16 | 0.644492518 | 0.010644787 | 0.044687249 | 26.12696008 | 12.3925702 | Basal | SOX2 |

| | | | | | | | |
|---|---|---|---|---|---|---|---|
| 1094 | RWDD1 | 0.644476403 | 0.000153669 | 0.007176147 | 73.7233654 | 44.41614732 | Superficial | SOX2 |
| 1095 | REXO1 | 0.644393123 | 0.000506425 | 0.010489143 | 21.29491628 | 11.63250148 | Suprabasal | SOX2 |
| 1096 | ALKBH6 | 0.643264176 | 0.015054848 | 0.038009884 | 8.987903727 | 4.148990879 | Superficial | SOX2 |
| 1097 | TP53BP2 | 0.642931517 | 0.004211401 | 0.027185186 | 27.76129219 | 12.84269071 | Basal | SOX2 |
| 1098 | ETV6 | 0.642798297 | 0.001883197 | 0.014679007 | 37.04574122 | 20.39227825 | Suprabasal | SOX2 |
| 1099 | RPAIN | 0.640544324 | 0.002836074 | 0.012662188 | 38.56017222 | 15.87384888 | Superficial | SOX2 |
| 1100 | NPLOC4 | 0.638617174 | 0.000635975 | 0.012925379 | 29.33666899 | 14.58457346 | Basal | SOX2 |
| 1101 | CLN6 | 0.636965297 | 0.010644787 | 0.038255467 | 17.08599633 | 8.317263013 | Suprabasal | SOX2 |
| 1102 | GDAP1 | 0.635852454 | 0.007423146 | 0.030449933 | 10.90194566 | 5.623306424 | Suprabasal | SOX2 |
| 1103 | SEC23IP | 0.634659651 | 0.007423146 | 0.022959455 | 22.1392895 | 8.593233906 | Superficial | SOX2 |
| 1104 | CDR2 | 0.634532008 | 0.001883197 | 0.018415354 | 10.22859731 | 4.830933407 | Basal | SOX2 |
| 1105 | B3GNT5 | 0.634212695 | 0.010644787 | 0.029507311 | 51.9158176 | 27.30802555 | Superficial | SOX2 |
| 1106 | SPPL2A | 0.633852245 | 0.004211401 | 0.016013117 | 43.33985558 | 18.32567339 | Superficial | SOX2 |
| 1107 | GSK3B | 0.633816567 | 0.000506425 | 0.007178523 | 40.39832657 | 17.8837765 | Superficial | SOX2 |
| 1108 | PUS7L | 0.633488025 | 0.002836074 | 0.012662188 | 29.01936478 | 12.46560073 | Superficial | SOX2 |
| 1109 | LINS1 | 0.631907647 | 0.004211401 | 0.016013117 | 16.64845294 | 5.985100765 | Superficial | SOX2 |
| 1110 | APPL2 | 0.631652725 | 0.008904627 | 0.034252903 | 42.9805544 | 24.81626319 | Suprabasal | SOX2 |
| 1111 | TACSTD2 | 0.631602355 | 0.000795824 | 0.0136425 | 98.55844464 | 89.75262691 | Basal | SOX2, KLF5 |
| 1112 | RALB | 0.631027731 | 0.001526458 | 0.009387421 | 66.92308937 | 38.04879059 | Superficial | SOX2 |
| 1113 | NUS1 | 0.630748992 | 0.001526458 | 0.009387421 | 26.18152527 | 10.93146073 | Superficial | SOX2 |
| 1114 | PLEKHG6 | 0.630732547 | 0.010644787 | 0.044687249 | 10.86442646 | 4.748218178 | Basal | SOX2 |
| 1115 | CTDP1 | 0.630691088 | 0.001883197 | 0.018415354 | 12.37768396 | 5.680913332 | Basal | SOX2 |
| 1116 | HCFC1 | 0.62946212 | 0.010644787 | 0.038255467 | 25.31814262 | 13.62847611 | Suprabasal | SOX2 |
| 1117 | TPBG | 0.628615415 | 0.008904627 | 0.025990329 | 12.37015338 | 4.658716986 | Superficial | SOX2 |
| 1118 | PSMD12 | 0.628161006 | 0.001883197 | 0.010234018 | 60.76021059 | 32.83275084 | Superficial | SOX2 |
| 1119 | VPS53 | 0.627854593 | 0.005105016 | 0.018023407 | 31.16917285 | 11.9110685 | Superficial | SOX2 |
| 1120 | PHRF1 | 0.627348754 | 0.000635975 | 0.010733065 | 28.08723807 | 15.10110419 | Suprabasal | SOX2 |
| 1121 | WDFY2 | 0.627045805 | 0.001232925 | 0.008679903 | 37.26846613 | 15.89920156 | Superficial | SOX2 |
| 1122 | DDX54 | 0.626076966 | 0.015054848 | 0.038009884 | 27.37361913 | 9.95378725 | Superficial | SOX2 |
| 1123 | GNA15 | 0.625857909 | 0.001526458 | 0.016792744 | 34.12656535 | 17.38627815 | Basal | SOX2 |
| 1124 | PACS1 | 0.624993882 | 0.000635975 | 0.012925379 | 32.06671086 | 15.46331705 | Basal | SOX2 + KLF5 |
| 1125 | KIFAP3 | 0.624614681 | 0.010644787 | 0.029507311 | 24.60834527 | 9.675495534 | Superficial | SOX2 |
| 1126 | TMBIM1 | 0.624550933 | 0.001883197 | 0.018415354 | 47.7802457 | 26.01769816 | Basal | SOX2 |
| 1127 | RPAP3 | 0.62447027 | 0.021001058 | 0.04834985 | 22.6548561 | 8.653170341 | Superficial | SOX2 |
| 1128 | PKIA | 0.624035874 | 0.003462059 | 0.019729589 | 17.9108499 | 9.009216803 | Suprabasal | SOX2 |
| 1129 | TTLL4 | 0.622221452 | 0.010644787 | 0.038255467 | 6.490223942 | 3.11242573 | Suprabasal | SOX2 |
| 1130 | TMEM51 | 0.621004095 | 0.000506425 | 0.012898798 | 13.93565344 | 6.630753718 | Basal | SOX2 |
| 1131 | CAPN15 | 0.620456443 | 0.008904627 | 0.040392504 | 15.5871956 | 6.756952252 | Basal | SOX2 |
| 1132 | TTC39C | 0.619828384 | 0.017811616 | 0.043066688 | 15.32945915 | 7.118245549 | Superficial | SOX2 + KLF5 |
| 1133 | LMNB1 | 0.619073998 | 0.001526458 | 0.016792744 | 46.9676545 | 27.82506765 | Basal | SOX2 |
| 1134 | NDST1 | 0.618998721 | 0.001232925 | 0.015721789 | 16.85826686 | 7.854229583 | Basal | SOX2 |
| 1135 | HS6ST1 | 0.618991894 | 0.002315114 | 0.016074975 | 26.59241943 | 14.78961537 | Suprabasal | SOX2 + KLF5, SOX2 |
| 1136 | NSD2 | 0.618825078 | 0.002836074 | 0.022562695 | 34.05755097 | 18.57604866 | Basal | SOX2 |
| 1137 | NDUFA4 | 0.617621009 | 0.007423146 | 0.022959455 | 87.20515658 | 75.61760295 | Superficial | SOX2 + KLF5 |
| 1138 | TTL | 0.616625565 | 0.015054848 | 0.038009884 | 9.39597726 | 3.866933503 | Superficial | SOX2 |
| 1139 | SCAF1 | 0.616556744 | 0.000401828 | 0.012455402 | 26.69961579 | 12.46298288 | Basal | SOX2 |
| 1140 | LMNA | 0.616183225 | 0.002315114 | 0.011381143 | 80.38720529 | 57.80369745 | Superficial | SOX2, SOX2 + KLF5 |
| 1141 | PARL | 0.615975765 | 0.006166656 | 0.020290909 | 34.42614282 | 14.25106949 | Superficial | SOX2 |
| 1142 | CBX4 | 0.614355706 | 0.006166656 | 0.032981404 | 24.91951176 | 12.4972595 | Basal | SOX2 |
| 1143 | BRWD3 | 0.613658138 | 0.008904627 | 0.025990329 | 34.96651289 | 13.44089262 | Superficial | SOX2 |
| 1144 | RIN2 | 0.613619312 | 0.003462059 | 0.019729589 | 48.89795167 | 27.70980442 | Suprabasal | SOX2 + KLF5 |
| 1145 | AXIN1 | 0.613476955 | 9.30E-05 | 0.010489143 | 14.62137054 | 7.716630072 | Suprabasal | SOX2 |
| 1146 | JUN | 0.612848095 | 0.010644787 | 0.029507311 | 77.53502385 | 50.74961947 | Superficial | SOX2, SOX2 + KLF5 |
| 1147 | MTOR | 0.612490458 | 0.000119764 | 0.012455402 | 22.77006362 | 11.23841573 | Basal | SOX2 |
| 1148 | PTK7 | 0.612100471 | 0.007423146 | 0.030449933 | 17.046313 | 8.949674957 | Suprabasal | SOX2 |
| 1149 | HNRNPU | 0.611217861 | 0.005105016 | 0.018023407 | 71.0978732 | 39.92723052 | Superficial | SOX2 |
| 1150 | ZBTB7A | 0.611118396 | 0.000635975 | 0.012925379 | 72.61060878 | 45.60543903 | Basal | SOX2 |
| 1151 | LMNB2 | 0.609316573 | 0.001526458 | 0.016792744 | 41.27761467 | 23.83188838 | Basal | SOX2 |
| 1152 | STAM | 0.608060929 | 0.015054848 | 0.038009884 | 33.32040064 | 13.39469361 | Superficial | SOX2 |
| 1153 | PSPC1 | 0.607965342 | 0.006166656 | 0.020290909 | 26.2929197 | 9.608926425 | Superficial | SOX2 |
| 1154 | UBA2 | 0.607402761 | 0.004211401 | 0.016013117 | 38.70500567 | 16.10997115 | Superficial | SOX2 |
| 1155 | REEP5 | 0.606826435 | 0.001232925 | 0.008679903 | 66.34652048 | 37.75642278 | Superficial | SOX2 |
| 1156 | ST3GAL1 | 0.606535423 | 0.017811616 | 0.043066688 | 24.29650378 | 9.987123775 | Superficial | SOX2 |
| 1157 | USP47 | 0.60533587 | 0.002836074 | 0.012662188 | 45.99539819 | 18.77180015 | Superficial | SOX2 |
| 1158 | DICER1 | 0.605163275 | 0.007423146 | 0.022959455 | 54.5955466 | 26.41130541 | Superficial | SOX2 |
| 1159 | STIL | 0.605032204 | 0.012681065 | 0.049425568 | 17.41909562 | 9.114647452 | Basal | SOX2 |
| 1160 | MEX3C | 0.604743821 | 0.002836074 | 0.0176927 | 19.5438782 | 10.34685587 | Suprabasal | SOX2 |
| 1161 | EBNA1BP2 | 0.604051292 | 0.008904627 | 0.025990329 | 40.02189028 | 17.49491332 | Superficial | SOX2 |
| 1162 | PKMYT1 | 0.603866657 | 0.005105016 | 0.029792864 | 26.85050168 | 15.06546933 | Basal | SOX2 |
| 1163 | DNTTIP2 | 0.60369074 | 0.007423146 | 0.022959455 | 38.13670698 | 15.82482952 | Superficial | SOX2 |
| 1164 | RABEP1 | 0.603633062 | 0.002315114 | 0.011381143 | 24.08573739 | 9.127485293 | Superficial | SOX2 |
| 1165 | PSMB2 | 0.603597338 | 0.007423146 | 0.022959455 | 59.36204562 | 29.71514402 | Superficial | SOX2 |
| 1166 | ATP11A | 0.602712245 | 0.007423146 | 0.036548425 | 16.71750869 | 7.367259194 | Basal | SOX2 |

| | | | | | | | | |
|---|---|---|---|---|---|---|---|---|
| 1167 | HSD17B1 | 0.602155405 | 0.004211401 | 0.016013117 | 24.56695092 | 11.3566743 | Superficial | SOX2 |
| 1168 | DDX1 | 0.601574847 | 0.008904627 | 0.025990329 | 44.5230924 | 18.97647767 | Superficial | SOX2 |
| 1169 | CCDC43 | 0.601502898 | 0.012681065 | 0.033478436 | 28.05317601 | 11.88733009 | Superficial | SOX2 |
| 1170 | F11R | 0.601020271 | 0.001526458 | 0.016792744 | 63.0512991 | 36.17533407 | Basal | SOX2 |
| 1171 | RRM2 | 0.600475257 | 0.010644787 | 0.044687249 | 39.28345524 | 25.80666442 | Basal | SOX2 |
| 1172 | ZNF367 | 0.600262693 | 0.000317696 | 0.012455402 | 15.13660037 | 7.800981888 | Basal | SOX2 |
| 1173 | IMP3 | 0.60004906 | 0.006166656 | 0.020290909 | 64.98953505 | 33.25115252 | Superficial | SOX2 |
| 1174 | FUT2 | 0.599158969 | 0.010644787 | 0.044687249 | 11.62944402 | 5.532194569 | Basal | SOX2 |
| 1175 | DAAM1 | 0.59915439 | 0.000992312 | 0.008042817 | 76.38738117 | 49.14984258 | Superficial | SOX2 |
| 1176 | SYPL1 | 0.598514626 | 0.008904627 | 0.025990329 | 76.69146901 | 50.8731247 | Superficial | SOX2 |
| 1177 | PDIA6 | 0.598308542 | 0.021001058 | 0.04834985 | 67.6873498 | 34.83698894 | Superficial | SOX2 |
| 1178 | FRMD8 | 0.598257819 | 0.003462059 | 0.024854712 | 21.81927065 | 10.05410138 | Basal | SOX2 |
| 1179 | ZNF286A | 0.598000696 | 0.002315114 | 0.016074975 | 10.4340277 | 5.156451451 | Suprabasal | SOX2 |
| 1180 | DDX5 | 0.596734835 | 0.002315114 | 0.011381143 | 80.40466764 | 53.89941309 | Superficial | SOX2 |
| 1181 | BRD3 | 0.596508411 | 0.001883197 | 0.014679007 | 26.70160161 | 14.51964849 | Suprabasal | SOX2 |
| 1182 | BACE2 | 0.594567261 | 0.005105016 | 0.029792864 | 44.63772598 | 24.73032218 | Basal | SOX2 + KLF5 |
| 1183 | TNIP1 | 0.594212161 | 0.000635975 | 0.012925379 | 39.40134427 | 20.60852556 | Basal | SOX2 + KLF5 |
| 1184 | PALB2 | 0.593794395 | 0.002836074 | 0.0176927 | 16.12236284 | 8.209849083 | Suprabasal | SOX2 |
| 1185 | KLHL2 | 0.592070357 | 0.000795824 | 0.0136425 | 12.37145745 | 5.614726525 | Basal | SOX2 |
| 1186 | KMT5A | 0.59128364 | 0.006166656 | 0.020290909 | 65.0270615 | 35.7780276 | Superficial | SOX2 |
| 1187 | FBXO45 | 0.588969119 | 0.017811616 | 0.043066688 | 19.16361532 | 8.241544775 | Superficial | SOX2 |
| 1188 | S100PBP | 0.588772465 | 0.003462059 | 0.019729589 | 15.51095779 | 7.592054591 | Suprabasal | SOX2 |
| 1189 | DDB1 | 0.588606432 | 0.021001058 | 0.04834985 | 36.33819273 | 15.45625643 | Superficial | SOX2 |
| 1190 | CCDC25 | 0.587148492 | 0.021001058 | 0.04834985 | 47.68870804 | 21.05364584 | Superficial | SOX2 |
| 1191 | DGKZ | 0.587067796 | 0.002836074 | 0.0176927 | 30.55586994 | 17.20076353 | Suprabasal | SOX2 |
| 1192 | KLF6 | 0.586976239 | 0.001526458 | 0.009387421 | 83.35824169 | 60.69685533 | Superficial | SOX2 |
| 1193 | CEBPD | 0.585393294 | 0.004211401 | 0.021879278 | 85.52303744 | 65.28861982 | Suprabasal | SOX2 |
| 1194 | ANKRD9 | 0.584905296 | 0.005105016 | 0.029792864 | 35.83613769 | 18.98583165 | Basal | SOX2 |
| 1195 | SPAST | 0.584604471 | 0.017811616 | 0.043066688 | 24.12051452 | 10.02080618 | Superficial | SOX2 |
| 1196 | NT5C3B | 0.584186592 | 0.021001058 | 0.04834985 | 18.38476757 | 7.533026674 | Superficial | SOX2 |
| 1197 | CDC123 | 0.583905789 | 0.010644787 | 0.029507311 | 42.59845803 | 18.30831463 | Superficial | SOX2 |
| 1198 | ADAM10 | 0.583902874 | 0.012681065 | 0.049425568 | 46.92135914 | 25.26441665 | Basal | SOX2 |
| 1199 | TBCB | 0.583533491 | 0.005105016 | 0.018023407 | 57.99479646 | 30.40976353 | Superficial | SOX2 |
| 1200 | TTC9 | 0.583380961 | 0.012681065 | 0.049425568 | 20.57073371 | 10.38099781 | Basal | SOX2 |
| 1201 | HMCES | 0.583210817 | 0.002836074 | 0.012662188 | 35.52203737 | 16.2197152 | Superficial | SOX2 |
| 1202 | PEX11B | 0.583019182 | 0.015054848 | 0.038009884 | 23.01143191 | 9.454768488 | Superficial | SOX2 |
| 1203 | RABGAP1 | 0.582347637 | 0.012681065 | 0.033478436 | 52.92135628 | 25.18508645 | Superficial | SOX2 |
| 1204 | PAIP2B | 0.580739387 | 0.021001058 | 0.04834985 | 44.41576247 | 20.3766825 | Superficial | SOX2 + KLF5, SOX2 |
| 1205 | CHD2 | 0.580357251 | 0.000795824 | 0.010838271 | 54.26481291 | 33.87147191 | Suprabasal | SOX2 |
| 1206 | MYO1C | 0.579266015 | 0.004211401 | 0.027185186 | 50.3517943 | 28.31410955 | Basal | SOX2 |
| 1207 | ACOT7 | 0.579261072 | 0.017811616 | 0.043066688 | 21.06320546 | 8.444328094 | Superficial | SOX2 |
| 1208 | ORC1 | 0.578777928 | 0.006166656 | 0.032981404 | 13.08983666 | 6.574679554 | Basal | SOX2 |
| 1209 | GNL3L | 0.577846754 | 0.012681065 | 0.042921737 | 27.7323956 | 15.59629515 | Suprabasal | SOX2 |
| 1210 | MICALL2 | 0.577590978 | 0.000795824 | 0.0136425 | 24.54449984 | 12.34438818 | Basal | SOX2 |
| 1211 | CDC5L | 0.577367904 | 0.010644787 | 0.029507311 | 46.85050398 | 20.28831503 | Superficial | SOX2 |
| 1212 | ZNF503 | 0.575857212 | 0.010644787 | 0.029507311 | 27.20047124 | 12.22366228 | Superficial | SOX2 |
| 1213 | MYO5B | 0.575610396 | 0.001883197 | 0.018415354 | 12.41517352 | 5.794956349 | Basal | SOX2 |
| 1214 | SHPK | 0.574491073 | 0.008904627 | 0.025990329 | 9.471988201 | 3.753573693 | Superficial | SOX2 |
| 1215 | BZW2 | 0.57447235 | 0.017811616 | 0.043066688 | 40.84455152 | 16.52980784 | Superficial | SOX2 |
| 1216 | SPPL3 | 0.573634654 | 0.003462059 | 0.014108599 | 47.28374064 | 22.03450031 | Superficial | SOX2 + KLF5, SOX2 |
| 1217 | CUL3 | 0.573016092 | 0.000401828 | 0.012455402 | 59.14956542 | 35.1104698 | Basal | SOX2 |
| 1218 | RTN4R | 0.573006335 | 0.010644787 | 0.044687249 | 10.07206455 | 4.963485412 | Basal | SOX2 |
| 1219 | KATNA1 | 0.572676717 | 0.000250281 | 0.007176147 | 25.18839815 | 11.23271671 | Superficial | SOX2 |
| 1220 | CHFR | 0.572376262 | 0.005105016 | 0.029792864 | 15.92621312 | 7.523139889 | Basal | SOX2 |
| 1221 | ABCB10 | 0.571942293 | 0.004211401 | 0.027185186 | 15.32551643 | 7.592237018 | Basal | SOX2 |
| 1222 | TSTD2 | 0.571866552 | 0.017811616 | 0.043066688 | 10.47451837 | 4.636528888 | Superficial | SOX2 |
| 1223 | STK11 | 0.570912857 | 0.004211401 | 0.027185186 | 39.54364401 | 21.58271748 | Basal | SOX2 |
| 1224 | CNOT6 | 0.570895163 | 0.010644787 | 0.029507311 | 27.29005571 | 10.21974886 | Superficial | SOX2 |
| 1225 | SMARCC1 | 0.570863674 | 0.010644787 | 0.029507311 | 41.37989477 | 19.06307239 | Superficial | SOX2 |
| 1226 | STRBP | 0.570780625 | 0.002315114 | 0.011381143 | 59.67198685 | 30.93992246 | Superficial | SOX2 |
| 1227 | ATXN7L3 | 0.569461391 | 0.000635975 | 0.012925379 | 22.89580349 | 10.95539312 | Basal | SOX2 |
| 1228 | MAP3K1 | 0.569441306 | 0.015054848 | 0.038009884 | 45.13891115 | 19.58848491 | Superficial | SOX2 |
| 1229 | KLF10 | 0.568633025 | 0.008904627 | 0.034252903 | 36.85873202 | 21.70693085 | Suprabasal | SOX2 |
| 1230 | MYLIP | 0.568414442 | 0.001232925 | 0.012246384 | 41.7823653 | 24.48742287 | Suprabasal | SOX2 |
| 1231 | PDXK | 0.567977931 | 0.000992312 | 0.014320968 | 57.7535651 | 33.79410889 | Basal | SOX2 |
| 1232 | ZMYM1 | 0.56738358 | 0.000506425 | 0.010489143 | 25.27509558 | 13.49142163 | Suprabasal | SOX2 |
| 1233 | UBLCP1 | 0.566918222 | 0.004211401 | 0.016013117 | 39.65019314 | 17.19327528 | Superficial | SOX2 |
| 1234 | MYBBP1A | 0.56688332 | 0.003462059 | 0.024854712 | 22.06185652 | 10.56027538 | Basal | SOX2 |
| 1235 | FBRSL1 | 0.566139832 | 0.003462059 | 0.024854712 | 23.29403516 | 10.99696911 | Basal | SOX2 |
| 1236 | OPA1 | 0.565910951 | 0.015054848 | 0.038009884 | 40.1444524 | 16.26258589 | Superficial | SOX2 |
| 1237 | MCM4 | 0.565800435 | 0.003462059 | 0.024854712 | 57.40336068 | 35.47954179 | Basal | SOX2 |
| 1238 | PLAGL2 | 0.565398167 | 0.008904627 | 0.040392504 | 19.64283433 | 9.125336889 | Basal | SOX2 |
| 1239 | TNFAIP8L1 | 0.564537093 | 0.007423146 | 0.036548425 | 18.10017498 | 10.5457503 | Basal | SOX2 |

| | | | | | | | | |
|---|---|---|---|---|---|---|---|---|
| 1240 | LARS2 | 0.563555094 | 0.006166656 | 0.032981404 | 12.02626978 | 5.845680963 | Basal | SOX2 |
| 1241 | ALYREF | 0.563301502 | 0.021001058 | 0.04834985 | 41.48176973 | 18.83980691 | Superficial | SOX2 |
| 1242 | LRP10 | 0.563132611 | 0.000153669 | 0.012455402 | 60.99423501 | 37.67474411 | Basal | SOX2 |
| 1243 | SLC12A2 | 0.562474231 | 0.003462059 | 0.019729589 | 10.77812199 | 5.408597712 | Suprabasal | SOX2 |
| 1244 | DDR1 | 0.562373943 | 0.000795824 | 0.010838271 | 68.0993919 | 45.28934753 | Suprabasal | SOX2 |
| 1245 | CNOT3 | 0.561689481 | 0.000992312 | 0.01142308 | 28.71801011 | 17.27564704 | Suprabasal | SOX2 |
| 1246 | NUP188 | 0.561464204 | 0.002836074 | 0.022562695 | 30.2577921 | 15.1332891 | Basal | SOX2 |
| 1247 | RGP1 | 0.560435385 | 0.003462059 | 0.019729589 | 13.42346619 | 7.532436965 | Suprabasal | SOX2 |
| 1248 | ZNF768 | 0.559358288 | 0.001883197 | 0.014679007 | 12.46714038 | 6.490597492 | Suprabasal | SOX2 |
| 1249 | INTS1 | 0.55918513 | 0.006166656 | 0.032981404 | 30.31020082 | 16.19867459 | Basal | SOX2 |
| 1250 | BAP1 | 0.558420415 | 0.015054848 | 0.048140298 | 30.97902085 | 18.53894319 | Suprabasal | SOX2 |
| 1251 | USP25 | 0.557295031 | 0.000196466 | 0.012455402 | 29.0726618 | 14.66598057 | Basal | SOX2 |
| 1252 | GUF1 | 0.5567566 | 0.007423146 | 0.022959455 | 23.83669527 | 8.827857569 | Superficial | SOX2 |
| 1253 | FAF2 | 0.556593152 | 0.001526458 | 0.009387421 | 36.29272897 | 14.54763219 | Superficial | SOX2 |
| 1254 | EGFL6 | 0.556147345 | 0.006166656 | 0.027303057 | 33.72800327 | 18.89160666 | Suprabasal | SOX2 |
| 1255 | ATP13A2 | 0.555814137 | 0.000401828 | 0.012455402 | 17.53077521 | 9.000378518 | Basal | SOX2 |
| 1256 | RAB27B | 0.555355604 | 0.002315114 | 0.011381143 | 59.79240219 | 31.73912783 | Superficial | SOX2, SOX2 + KLF5 |
| 1257 | XPO5 | 0.554728734 | 0.015054848 | 0.038009884 | 12.41895402 | 4.453709417 | Superficial | SOX2 |
| 1258 | TARBP1 | 0.553824775 | 0.003462059 | 0.019729589 | 19.92559264 | 11.6501114 | Suprabasal | SOX2 |
| 1259 | RRP12 | 0.553439503 | 0.001232925 | 0.015721789 | 12.13356295 | 5.922857206 | Basal | SOX2 + KLF5, SOX2 |
| 1260 | MCM2 | 0.553145931 | 0.008904627 | 0.040392504 | 26.87734726 | 14.5281405 | Basal | SOX2 |
| 1261 | TES | 0.551859738 | 0.006166656 | 0.020290909 | 40.8340672 | 18.31834669 | Superficial | SOX2 |
| 1262 | SLC25A10 | 0.551805139 | 0.010644787 | 0.044687249 | 38.67628151 | 21.72036806 | Basal | SOX2 + KLF5, SOX2 |
| 1263 | TMEM33 | 0.550352753 | 0.003462059 | 0.014108599 | 43.15784579 | 21.71450811 | Superficial | SOX2 |
| 1264 | ECD | 0.549038951 | 0.017811616 | 0.043066688 | 19.31874557 | 9.430919656 | Superficial | SOX2 |
| 1265 | TBCD | 0.548662953 | 0.003462059 | 0.024854712 | 27.13218841 | 14.47561619 | Basal | SOX2 |
| 1266 | JUP | 0.54864085 | 0.000795824 | 0.0136425 | 86.39745245 | 66.15826609 | Basal | SOX2, SOX2 + KLF5 |
| 1267 | FAM102A | 0.547362144 | 0.000635975 | 0.012925379 | 31.91566041 | 16.70012751 | Basal | SOX2 |
| 1268 | TRIP12 | 0.545203053 | 0.007423146 | 0.036548425 | 49.7497347 | 27.85387132 | Basal | SOX2 |
| 1269 | ATP6V0A2 | 0.544362438 | 0.015054848 | 0.048140298 | 13.39822462 | 7.244591033 | Suprabasal | SOX2 |
| 1270 | SYTL1 | 0.544315471 | 0.021001058 | 0.04834985 | 73.97048156 | 45.83813186 | Superficial | SOX2 |
| 1271 | HNRNPM | 0.543720788 | 0.012681065 | 0.033478436 | 63.11040245 | 32.54516521 | Superficial | SOX2 |
| 1272 | ELF3 | 0.54136574 | 0.012681065 | 0.049425568 | 56.54352937 | 38.3781648 | Basal | SOX2, KLF5, SOX2 + KLF5 |
| 1273 | ERBB3 | 0.541330501 | 0.000795824 | 0.0136425 | 42.43033253 | 24.05521405 | Basal | SOX2 |
| 1274 | KLC2 | 0.54054988 | 0.004211401 | 0.021879278 | 11.36024429 | 6.507917261 | Suprabasal | SOX2 |
| 1275 | CBFB | 0.539190443 | 0.001526458 | 0.016792744 | 30.44720682 | 14.98475264 | Basal | SOX2 |
| 1276 | MICALL1 | 0.53903016 | 0.001526458 | 0.016792744 | 26.62948582 | 13.60714347 | Basal | SOX2 |
| 1277 | ATP13A3 | 0.538620588 | 0.006166656 | 0.032981404 | 33.65402979 | 16.85817377 | Basal | SOX2 |
| 1278 | BMP2K | 0.5381026 | 0.010644787 | 0.029507311 | 30.86873589 | 12.87664539 | Superficial | SOX2 |
| 1279 | IRAK1 | 0.538095381 | 0.002315114 | 0.020052792 | 38.96317903 | 21.4747565 | Basal | SOX2 |
| 1280 | TM7SF3 | 0.537123275 | 0.003462059 | 0.014108599 | 39.04521885 | 18.4880171 | Superficial | SOX2 |
| 1281 | PSMB3 | 0.535779065 | 0.002315114 | 0.011381143 | 73.69962452 | 44.93950696 | Superficial | SOX2 |
| 1282 | CRYBG2 | 0.535424694 | 0.003462059 | 0.024854712 | 13.47297953 | 6.898201952 | Basal | KLF5 |
| 1283 | SOD2 | 0.535167327 | 0.008904627 | 0.040392504 | 74.53929203 | 49.21507383 | Basal | SOX2 |
| 1284 | SNN | 0.534874669 | 0.000401828 | 0.012455402 | 12.75407637 | 6.481786994 | Basal | SOX2 |
| 1285 | CWC27 | 0.534707173 | 0.004211401 | 0.016013117 | 26.12936858 | 11.5878165 | Superficial | SOX2 |
| 1286 | ARSJ | 0.533709462 | 0.012681065 | 0.042921737 | 7.698190606 | 3.763901651 | Suprabasal | SOX2 |
| 1287 | NUDT3 | 0.533337388 | 0.012681065 | 0.033478436 | 25.04808303 | 9.578946621 | Superficial | SOX2 |
| 1288 | ANKRD52 | 0.53277344 | 0.004211401 | 0.021879278 | 10.11138008 | 5.40810124 | Suprabasal | SOX2 |
| 1289 | S100A11 | 0.532661764 | 0.004211401 | 0.027185186 | 99.86119542 | 98.82193269 | Basal | SOX2 |
| 1290 | CNOT11 | 0.5325964 | 0.002315114 | 0.011381143 | 38.1402724 | 17.44697014 | Superficial | SOX2 |
| 1291 | CUL2 | 0.532346494 | 0.012681065 | 0.033478436 | 31.02835804 | 13.27069577 | Superficial | SOX2 |
| 1292 | EIF2S1 | 0.532152229 | 0.006166656 | 0.020290909 | 47.45304071 | 21.71808627 | Superficial | SOX2 |
| 1293 | ERCC6 | 0.531874264 | 0.002836074 | 0.0176927 | 24.24188941 | 14.03843851 | Suprabasal | SOX2 |
| 1294 | NKAP | 0.531109713 | 0.002315114 | 0.011381143 | 36.56583353 | 15.56036051 | Superficial | SOX2 |
| 1295 | ITPKB | 0.530289527 | 0.004211401 | 0.021879278 | 8.021108444 | 4.822059891 | Suprabasal | SOX2 |
| 1296 | TGOLN2 | 0.529440537 | 0.006166656 | 0.020290909 | 45.81425303 | 23.33559263 | Superficial | SOX2 |
| 1297 | TRIP10 | 0.527179997 | 0.000795824 | 0.0136425 | 26.74185502 | 13.52408073 | Basal | SOX2 + KLF5 |
| 1298 | WAPL | 0.527163417 | 0.017811616 | 0.043066688 | 45.72441968 | 20.94921423 | Superficial | SOX2 |
| 1299 | ANLN | 0.526520833 | 0.008904627 | 0.040392504 | 36.530004 | 22.85027331 | Basal | SOX2 |
| 1300 | ANXA1 | 0.526354665 | 0.010644787 | 0.044687249 | 99.04728296 | 97.15687469 | Basal | SOX2 + KLF5, SOX2 |
| 1301 | CTNNA1 | 0.525896828 | 0.002315114 | 0.020052792 | 77.18314236 | 53.75795767 | Basal | SOX2 |
| 1302 | NXN | 0.52540313 | 0.012681065 | 0.042921737 | 43.88886404 | 24.88549412 | Suprabasal | SOX2 |
| 1303 | KLHL8 | 0.523147525 | 0.010644787 | 0.038255467 | 15.65540226 | 8.980403041 | Suprabasal | SOX2 + KLF5 |
| 1304 | JAG1 | 0.522042648 | 0.002836074 | 0.022562695 | 60.931408 | 37.7428955 | Basal | SOX2, SOX2 + KLF5 |
| 1305 | CHD7 | 0.521984316 | 0.007423146 | 0.022959455 | 33.38086425 | 14.66102246 | Superficial | SOX2 |
| 1306 | RAD23A | 0.521616714 | 0.010644787 | 0.029507311 | 49.6591592 | 21.30608421 | Superficial | SOX2 |
| 1307 | ACBD3 | 0.521467682 | 0.001526458 | 0.013290882 | 80.56801461 | 59.80486031 | Suprabasal | SOX2 |
| 1308 | NUMA1 | 0.521181726 | 0.005105016 | 0.029792864 | 53.1244713 | 30.20781378 | Basal | SOX2 |
| 1309 | ATRIP | 0.520900406 | 0.002315114 | 0.020052792 | 14.17842467 | 7.484290553 | Basal | SOX2 |
| 1310 | NFATC2IP | 0.52049701 | 0.012681065 | 0.033478436 | 26.51865935 | 10.89065796 | Superficial | SOX2 |
| 1311 | GRSF1 | 0.519672061 | 0.000506425 | 0.007178523 | 47.21126079 | 22.01541314 | Superficial | SOX2 |
| 1312 | ARID2 | 0.518042147 | 0.015054848 | 0.038009884 | 26.3492912 | 11.03379219 | Superficial | SOX2 |

| | | | | | | | | |
|---|---|---|---|---|---|---|---|---|
| 1313 | CUEDC1 | 0.517765462 | 0.002315114 | 0.020052792 | 16.95961156 | 8.258408028 | Basal | SOX2 |
| 1314 | KLF3 | 0.517679593 | 0.007423146 | 0.030449933 | 83.92570547 | 65.51864617 | Suprabasal | SOX2 + KLF5 |
| 1315 | MTRF1L | 0.51714331 | 0.010644787 | 0.029507311 | 27.58706894 | 13.04922908 | Superficial | SOX2 |
| 1316 | ELP4 | 0.516927801 | 0.010644787 | 0.029507311 | 14.78220504 | 6.588135394 | Superficial | SOX2 |
| 1317 | CDC42EP5 | 0.515557091 | 0.008904627 | 0.040392504 | 51.64386602 | 28.76256482 | Basal | SOX2 |
| 1318 | RFWD3 | 0.515309713 | 0.004211401 | 0.021879278 | 11.70454255 | 6.736320405 | Suprabasal | SOX2 |
| 1319 | AQR | 0.514022203 | 0.015054848 | 0.038009884 | 31.19422048 | 12.82419177 | Superficial | SOX2 |
| 1320 | SDE2 | 0.512741406 | 0.006166656 | 0.027303057 | 23.7675373 | 12.72793341 | Suprabasal | SOX2 |
| 1321 | PNKP | 0.511632792 | 0.008904627 | 0.025990329 | 23.42211961 | 10.10092883 | Superficial | SOX2 |
| 1322 | UPF1 | 0.511286401 | 0.003462059 | 0.024854712 | 37.07981162 | 19.42343512 | Basal | SOX2 |
| 1323 | PTK6 | 0.510135249 | 0.012681065 | 0.049425568 | 19.27144799 | 9.872191405 | Basal | SOX2 |
| 1324 | USP6NL | 0.509593969 | 0.008904627 | 0.040392504 | 17.68897498 | 8.50089317 | Basal | SOX2 |
| 1325 | MPZL1 | 0.508728407 | 0.008904627 | 0.025990329 | 43.3369847 | 22.76864965 | Superficial | SOX2 + KLF5, SOX2 |
| 1326 | BUB3 | 0.508714222 | 0.007423146 | 0.022959455 | 45.69659583 | 21.03149772 | Superficial | SOX2 |
| 1327 | PURA | 0.507065933 | 0.000401828 | 0.007176147 | 44.46355188 | 21.71574662 | Superficial | SOX2 |
| 1328 | FRS2 | 0.506244634 | 0.021001058 | 0.04834985 | 18.7799487 | 7.366259053 | Superficial | SOX2 |
| 1329 | FRMD6 | 0.505534122 | 0.002315114 | 0.011381143 | 72.31695849 | 41.61355295 | Superficial | SOX2 |
| 1330 | CCND1 | 0.505169811 | 0.021001058 | 0.04834985 | 55.07589057 | 29.22761701 | Superficial | SOX2 |
| 1331 | GNB2 | 0.50420714 | 0.005105016 | 0.029792864 | 78.3157134 | 55.61594237 | Basal | SOX2 |
| 1332 | SH2B1 | 0.503568083 | 0.015054848 | 0.048140298 | 17.07369447 | 9.923961829 | Suprabasal | SOX2 |
| 1333 | ASPH | 0.501403718 | 0.004211401 | 0.021879278 | 70.82171402 | 48.84771551 | Suprabasal | SOX2 |
| 1334 | PPP6R1 | 0.5009654 | 0.001232925 | 0.015721789 | 29.47347937 | 15.12279427 | Basal | SOX2 |
| 1335 | NAPG | 0.499670728 | 0.003462059 | 0.014108599 | 57.91755322 | 29.49803192 | Superficial | SOX2 |
| 1336 | MNAT1 | 0.499035232 | 0.001526458 | 0.009387421 | 33.82642552 | 15.08818925 | Superficial | SOX2 |
| 1337 | HNRNPA2B1 | 0.497371459 | 0.015054848 | 0.038009884 | 74.51684124 | 46.06121545 | Superficial | SOX2 |
| 1338 | CEP44 | 0.497034882 | 0.021001058 | 0.04834985 | 13.30912046 | 5.031562544 | Superficial | SOX2 |
| 1339 | FAM111A | 0.496459518 | 0.005105016 | 0.024262694 | 17.72632172 | 10.20689672 | Suprabasal | SOX2 |
| 1340 | GUCD1 | 0.49643982 | 0.001883197 | 0.018415354 | 31.74971695 | 16.31569375 | Basal | SOX2 |
| 1341 | CLUAP1 | 0.496336728 | 0.008904627 | 0.034252903 | 21.28641366 | 11.54826649 | Suprabasal | SOX2 |
| 1342 | KCTD3 | 0.494652574 | 0.000992312 | 0.014320968 | 39.57678112 | 21.59878832 | Basal | SOX2 |
| 1343 | SAP30 | 0.493979071 | 0.007423146 | 0.036548425 | 45.00566094 | 26.41014799 | Basal | SOX2 |
| 1344 | HNRNPR | 0.491063799 | 0.017811616 | 0.043066688 | 47.55427304 | 22.21570082 | Superficial | SOX2 |
| 1345 | NRDC | 0.490991214 | 0.021001058 | 0.04834985 | 55.72038228 | 30.60547076 | Superficial | SOX2 |
| 1346 | TRIM33 | 0.489450207 | 0.017811616 | 0.043066688 | 39.7606089 | 17.51831737 | Superficial | SOX2 + KLF5, SOX2 |
| 1347 | SEC61A1 | 0.488628097 | 0.015054848 | 0.038009884 | 45.15063283 | 22.414793 | Superficial | SOX2 |
| 1348 | WDHD1 | 0.488116912 | 0.001526458 | 0.016792744 | 24.01800884 | 13.26388537 | Basal | SOX2 |
| 1349 | SPAG9 | 0.48675745 | 0.000795824 | 0.0136425 | 51.34410616 | 30.18405598 | Basal | SOX2 |
| 1350 | SEC14L2 | 0.486212927 | 0.001883197 | 0.018415354 | 12.00086909 | 5.227538256 | Basal | SOX2 |
| 1351 | MRPS31 | 0.486202719 | 0.017811616 | 0.043066688 | 38.70154943 | 17.3136704 | Superficial | SOX2 |
| 1352 | BDKRB2 | 0.485062961 | 0.006166656 | 0.032981404 | 17.36183784 | 8.801648887 | Basal | SOX2 |
| 1353 | SRPK2 | 0.482498645 | 0.003462059 | 0.014108599 | 44.41141015 | 20.7636501 | Superficial | SOX2 |
| 1354 | PSMA1 | 0.482165624 | 0.008904627 | 0.025990329 | 65.97719582 | 36.27990299 | Superficial | SOX2 |
| 1355 | TPGS1 | 0.481782703 | 0.017811616 | 0.043066688 | 41.59724017 | 19.97718233 | Superficial | SOX2 |
| 1356 | CACTIN | 0.481764839 | 0.010644787 | 0.038255467 | 9.797951605 | 5.480509165 | Suprabasal | SOX2 |
| 1357 | CTTN | 0.481760114 | 0.002315114 | 0.020052792 | 68.34703483 | 44.04458385 | Basal | SOX2 |
| 1358 | ZDHHC5 | 0.481638674 | 0.002836074 | 0.022562695 | 45.41187793 | 26.26191115 | Basal | SOX2 |
| 1359 | SH3GL1 | 0.481509796 | 0.000317696 | 0.012455402 | 40.9722635 | 22.65329046 | Basal | SOX2 |
| 1360 | FMR1 | 0.481142069 | 0.005105016 | 0.024262694 | 43.2968248 | 26.86475631 | Suprabasal | SOX2 |
| 1361 | NADK | 0.480684707 | 0.010644787 | 0.044687249 | 20.85555924 | 10.89633778 | Basal | SOX2 |
| 1362 | RALY | 0.479004374 | 0.017811616 | 0.043066688 | 71.36994745 | 45.26189074 | Superficial | SOX2 |
| 1363 | SPECC1 | 0.478790004 | 0.010644787 | 0.044687249 | 16.64152372 | 8.452011622 | Basal | SOX2 |
| 1364 | DNAJB14 | 0.47843695 | 0.007423146 | 0.022959455 | 35.23391723 | 15.120162 | Superficial | SOX2 |
| 1365 | TRIM16 | 0.475968725 | 0.006166656 | 0.032981404 | 45.63964355 | 28.50255283 | Basal | SOX2 |
| 1366 | DVL3 | 0.475846201 | 0.000250281 | 0.010489143 | 34.21125811 | 21.10756577 | Suprabasal | SOX2 |
| 1367 | ARMC1 | 0.474807714 | 0.008904627 | 0.025990329 | 27.30242392 | 11.60866805 | Superficial | SOX2 |
| 1368 | R3HDM4 | 0.474476837 | 0.000992312 | 0.014320968 | 34.1889281 | 18.11556037 | Basal | SOX2 |
| 1369 | TWF1 | 0.473939191 | 0.001883197 | 0.018415354 | 57.49152249 | 35.50918611 | Basal | SOX2 |
| 1370 | DENND2D | 0.47280977 | 0.000317696 | 0.012455402 | 36.61631128 | 20.5686023 | Basal | SOX2 |
| 1371 | AHNAK | 0.472580773 | 0.000992312 | 0.014320968 | 94.70051847 | 83.04168059 | Basal | SOX2, SOX2 + KLF5 |
| 1372 | SIK3 | 0.471835245 | 0.003462059 | 0.024854712 | 16.02132565 | 7.698144507 | Basal | SOX2 |
| 1373 | EHD1 | 0.471129417 | 0.006166656 | 0.032981404 | 18.87141849 | 9.712745193 | Basal | SOX2 |
| 1374 | SCO1 | 0.469811368 | 0.002315114 | 0.011381143 | 27.81739673 | 11.84774497 | Superficial | SOX2 |
| 1375 | RCN1 | 0.469437498 | 0.007423146 | 0.030449933 | 39.25339792 | 22.71572912 | Suprabasal | SOX2 |
| 1376 | THUMPD3 | 0.469358263 | 0.021001058 | 0.04834985 | 30.94173887 | 12.9790936 | Superficial | SOX2 |
| 1377 | RYK | 0.469142284 | 0.002315114 | 0.011381143 | 30.19491679 | 13.27920261 | Superficial | SOX2 |
| 1378 | PHF20L1 | 0.468437348 | 0.007423146 | 0.022959455 | 61.30332804 | 34.85572051 | Superficial | SOX2 |
| 1379 | SHCBP1 | 0.467837807 | 0.005105016 | 0.029792864 | 22.36528606 | 12.86186587 | Basal | SOX2 |
| 1380 | DENND1A | 0.467812365 | 0.012681065 | 0.042921737 | 6.96267468 | 4.220814957 | Suprabasal | SOX2 + KLF5 |
| 1381 | PPM1D | 0.467316307 | 0.002836074 | 0.022562695 | 13.17120466 | 6.699147999 | Basal | SOX2 |
| 1382 | ZNF598 | 0.466855393 | 0.002315114 | 0.020052792 | 20.47352294 | 10.30751396 | Basal | SOX2 |
| 1383 | PGAP2 | 0.466709668 | 0.012681065 | 0.033478436 | 35.43820831 | 16.82219589 | Superficial | SOX2 |
| 1384 | PELP1 | 0.466601935 | 0.005105016 | 0.024262694 | 27.07148126 | 15.08417818 | Suprabasal | SOX2 |
| 1385 | MAP4K4 | 0.464946588 | 0.002315114 | 0.020052792 | 29.0491676 | 15.70885276 | Basal | SOX2 |

| | | | | | | | | |
|---|---|---|---|---|---|---|---|---|
| 1386 | USP22 | 0.463671294 | 0.012681065 | 0.042921737 | 44.5022348 | 27.78089775 | Suprabasal | SOX2 |
| 1387 | MGAT4B | 0.463578307 | 0.012681065 | 0.033478436 | 63.79064886 | 35.34931688 | Superficial | SOX2 |
| 1388 | DUSP16 | 0.463416326 | 0.010644787 | 0.044687249 | 12.4917934 | 5.563776748 | Basal | SOX2 |
| 1389 | FAAP100 | 0.462850974 | 0.003462059 | 0.024854712 | 14.96658424 | 7.878325253 | Basal | SOX2 |
| 1390 | ERICH1 | 0.462223974 | 0.015054848 | 0.038009884 | 25.89510034 | 10.06784504 | Superficial | SOX2 |
| 1391 | SLC22A23 | 0.461757582 | 0.006166656 | 0.027303057 | 39.49224725 | 24.87529787 | Suprabasal | SOX2 |
| 1392 | STAM2 | 0.459897509 | 0.002836074 | 0.0176927 | 43.15189679 | 27.25445502 | Suprabasal | SOX2 |
| 1393 | STRN4 | 0.459425666 | 0.000506425 | 0.012898798 | 25.0584803 | 12.69442914 | Basal | SOX2 |
| 1394 | PRKCI | 0.459071509 | 0.012681065 | 0.042921737 | 51.41691073 | 33.04730749 | Suprabasal | SOX2 |
| 1395 | GATAD2A | 0.458863107 | 0.003462059 | 0.024854712 | 35.79161866 | 18.9186545 | Basal | SOX2 |
| 1396 | TMEM68 | 0.458247895 | 0.015054848 | 0.038009884 | 16.45128655 | 6.283049534 | Superficial | SOX2 |
| 1397 | EIF5B | 0.45771753 | 0.017811616 | 0.043066688 | 63.11019127 | 36.34802617 | Superficial | SOX2 |
| 1398 | NUP98 | 0.457010854 | 0.000992312 | 0.014320968 | 32.6907407 | 17.7336952 | Basal | SOX2 |
| 1399 | DYNC1LI2 | 0.456769397 | 0.000635975 | 0.007274975 | 61.99864339 | 35.22299681 | Superficial | SOX2 |
| 1400 | CAND1 | 0.454883616 | 0.015054848 | 0.048140298 | 51.67717794 | 32.39796423 | Suprabasal | SOX2 |
| 1401 | SEC16A | 0.454482179 | 0.010644787 | 0.044687249 | 21.18626829 | 10.61205209 | Basal | SOX2 |
| 1402 | ADAM17 | 0.453611455 | 0.010644787 | 0.029507311 | 30.19171051 | 11.82663939 | Superficial | SOX2 |
| 1403 | THUMPD2 | 0.45263715 | 0.021001058 | 0.04834985 | 15.97784169 | 6.56066906 | Superficial | SOX2 + KLF5 |
| 1404 | ACACA | 0.452417245 | 0.008904627 | 0.040392504 | 29.6471181 | 15.65104834 | Basal | SOX2 |
| 1405 | PPP2R5E | 0.452064751 | 0.021001058 | 0.04834985 | 55.92045393 | 30.5051335 | Superficial | SOX2 |
| 1406 | SLK | 0.451135179 | 0.000401828 | 0.012455402 | 60.34261778 | 38.60359737 | Basal | SOX2 |
| 1407 | MOCOS | 0.450507041 | 0.007423146 | 0.036548425 | 11.66780025 | 6.100869399 | Basal | SOX2 |
| 1408 | TCOF1 | 0.449688868 | 0.001526458 | 0.016792744 | 41.03802722 | 24.1069045 | Basal | SOX2 |
| 1409 | PANX1 | 0.449051913 | 0.012681065 | 0.049425568 | 9.645531418 | 4.874082636 | Basal | SOX2 |
| 1410 | ZFP90 | 0.448202743 | 0.002315114 | 0.016074975 | 15.3130742 | 8.801039518 | Suprabasal | SOX2 |
| 1411 | FBXO46 | 0.447624189 | 0.000992312 | 0.014320968 | 16.59204452 | 8.127326468 | Basal | SOX2 |
| 1412 | STRN | 0.447432023 | 0.010644787 | 0.044687249 | 38.84829394 | 21.44867213 | Basal | SOX2 |
| 1413 | NOL6 | 0.446672582 | 0.010644787 | 0.044687249 | 13.11910862 | 6.729699791 | Basal | SOX2 |
| 1414 | MDN1 | 0.446383845 | 0.005105016 | 0.029792864 | 24.44692351 | 12.62374257 | Basal | SOX2 |
| 1415 | CNN2 | 0.445861687 | 0.006166656 | 0.032981404 | 56.8142339 | 32.70563775 | Basal | SOX2 |
| 1416 | ADA | 0.445206613 | 0.005105016 | 0.029792864 | 22.03751847 | 11.78920222 | Basal | SOX2 |
| 1417 | FBRS | 0.444705702 | 0.012681065 | 0.049425568 | 30.07899109 | 15.75416998 | Basal | SOX2 |
| 1418 | STON2 | 0.443577293 | 0.008904627 | 0.034252903 | 31.50370257 | 18.35266099 | Suprabasal | SOX2 |
| 1419 | ACOX1 | 0.442951284 | 0.004211401 | 0.027185186 | 30.32328526 | 15.83888351 | Basal | SOX2 + KLF5 |
| 1420 | NEDD1 | 0.442012152 | 0.008904627 | 0.034252903 | 14.81115727 | 7.965198757 | Suprabasal | SOX2 |
| 1421 | ARAF | 0.44188733 | 0.001883197 | 0.018415354 | 23.96647628 | 12.83259466 | Basal | SOX2 |
| 1422 | GTF3C4 | 0.438299723 | 0.010644787 | 0.038255467 | 20.14127445 | 12.16362485 | Suprabasal | SOX2 |
| 1423 | ARMT1 | 0.437711806 | 0.010644787 | 0.029507311 | 27.61166468 | 11.64950814 | Superficial | SOX2 |
| 1424 | ZNF513 | 0.437149719 | 0.002836074 | 0.0176927 | 13.08879921 | 8.269687283 | Suprabasal | SOX2 |
| 1425 | GPC1 | 0.4335995 | 0.005105016 | 0.024262694 | 58.84720048 | 40.39056084 | Suprabasal | SOX2 |
| 1426 | PLS1 | 0.433501833 | 0.007423146 | 0.036548425 | 23.59769785 | 12.01671039 | Basal | SOX2 + KLF5 |
| 1427 | TOPBP1 | 0.432977178 | 0.003462059 | 0.019729589 | 26.22138818 | 15.40008064 | Suprabasal | SOX2 |
| 1428 | CTSB | 0.43282805 | 0.002315114 | 0.020052792 | 79.26851537 | 55.67357313 | Basal | SOX2 |
| 1429 | CRABP2 | 0.432650393 | 0.008904627 | 0.040392504 | 83.33089204 | 72.13197537 | Basal | SOX2 |
| 1430 | RAD54L2 | 0.431128164 | 0.006166656 | 0.027303057 | 13.54171258 | 7.65361492 | Suprabasal | SOX2 |
| 1431 | PCBP4 | 0.430029248 | 0.012681065 | 0.042921737 | 12.17578092 | 7.218205687 | Suprabasal | SOX2 |
| 1432 | SMCHD1 | 0.429808244 | 0.006166656 | 0.027303057 | 37.02295778 | 22.63417127 | Suprabasal | SOX2 |
| 1433 | SNX1 | 0.429491131 | 0.002315114 | 0.016074975 | 63.4365984 | 43.72910637 | Suprabasal | SOX2 |
| 1434 | DHX32 | 0.427564306 | 0.005105016 | 0.029792864 | 29.98898918 | 16.15208417 | Basal | SOX2 |
| 1435 | PTPN9 | 0.426795876 | 0.015054848 | 0.048140298 | 12.96039252 | 7.098054443 | Suprabasal | SOX2 |
| 1436 | SRPRA | 0.426396935 | 0.002315114 | 0.020052792 | 47.1307841 | 26.94631004 | Basal | SOX2 |
| 1437 | MAPRE3 | 0.426209732 | 0.012681065 | 0.042921737 | 6.598992634 | 3.77396755 | Suprabasal | SOX2 |
| 1438 | STK24 | 0.423981945 | 0.004211401 | 0.027185186 | 66.7561237 | 42.88101471 | Basal | SOX2 |
| 1439 | PPIL4 | 0.423935678 | 0.001526458 | 0.009387421 | 33.29502682 | 14.07107106 | Superficial | SOX2 |
| 1440 | HDAC2 | 0.423736835 | 0.012681065 | 0.033478436 | 46.77046386 | 22.13070123 | Superficial | SOX2 |
| 1441 | DONSON | 0.422694654 | 0.002315114 | 0.020052792 | 18.01339736 | 10.3784105 | Basal | SOX2 |
| 1442 | ATAD2 | 0.422335996 | 0.007423146 | 0.036548425 | 52.37566144 | 34.75413718 | Basal | SOX2 |
| 1443 | EIF4G3 | 0.421871801 | 0.012681065 | 0.033478436 | 41.23540469 | 19.80500389 | Superficial | SOX2 + KLF5 |
| 1444 | KIF1B | 0.421858334 | 0.001232925 | 0.015721789 | 28.04042735 | 15.11331966 | Basal | SOX2 |
| 1445 | CSNK1G2 | 0.421806566 | 0.001526458 | 0.016792744 | 41.06864299 | 23.48149592 | Basal | SOX2 |
| 1446 | AP1S1 | 0.421228877 | 0.015054848 | 0.038009884 | 48.96038672 | 26.48981535 | Superficial | SOX2 |
| 1447 | TAF13 | 0.420823645 | 0.007423146 | 0.036548425 | 21.98425883 | 10.86907487 | Basal | SOX2 |
| 1448 | RPRD1B | 0.420555262 | 0.004211401 | 0.027185186 | 24.94256673 | 13.10438258 | Basal | SOX2 |
| 1449 | PSME3 | 0.419754849 | 0.003462059 | 0.024854712 | 36.95658387 | 21.39527241 | Basal | SOX2 |
| 1450 | EIF4G2 | 0.419134212 | 0.005105016 | 0.029792864 | 89.21513224 | 71.25553618 | Basal | SOX2 |
| 1451 | ABHD17C | 0.418557758 | 0.007423146 | 0.036548425 | 29.43653927 | 15.59754586 | Basal | SOX2 |
| 1452 | BAG3 | 0.418424075 | 0.001526458 | 0.016792744 | 57.19094076 | 35.78013283 | Basal | SOX2 |
| 1453 | DCTN1 | 0.418072859 | 0.002315114 | 0.016074975 | 51.97888715 | 35.8322889 | Suprabasal | SOX2 |
| 1454 | SOS1 | 0.416701949 | 0.004211401 | 0.021879278 | 38.80402609 | 24.64701172 | Suprabasal | SOX2 |
| 1455 | PLD1 | 0.416621216 | 0.010644787 | 0.044687249 | 26.55414004 | 14.95657525 | Basal | SOX2 + KLF5 |
| 1456 | CSNK2A2 | 0.416250629 | 0.007423146 | 0.030449933 | 49.98222445 | 31.77717767 | Suprabasal | SOX2 |
| 1457 | SIGMAR1 | 0.415664039 | 0.010644787 | 0.044687249 | 45.53288418 | 27.00160704 | Basal | SOX2 |
| 1458 | BAIAP2L1 | 0.414981604 | 0.000992312 | 0.014320968 | 23.04935728 | 12.37712073 | Basal | SOX2, SOX2 + KLF5 |

| | | | | | | | | |
|---|---|---|---|---|---|---|---|---|
| 1459 | RRAGD | 0.411940318 | 0.008904627 | 0.025990329 | 53.16936183 | 28.17200598 | Superficial | SOX2 |
| 1460 | LARP4B | 0.411512699 | 0.015054848 | 0.048140298 | 44.28576258 | 28.30531636 | Suprabasal | SOX2 |
| 1461 | RRP1B | 0.411164292 | 0.000795824 | 0.010838271 | 32.97235215 | 19.66653476 | Suprabasal | SOX2 |
| 1462 | MAPK1 | 0.410899477 | 0.005105016 | 0.029792864 | 48.74884732 | 28.8759107 | Basal | SOX2 |
| 1463 | OXSR1 | 0.410887757 | 0.012681065 | 0.049425568 | 31.81485499 | 17.55439172 | Basal | SOX2 |
| 1464 | PPP4R3A | 0.409837121 | 0.012681065 | 0.042921737 | 46.70037185 | 29.32635926 | Suprabasal | SOX2 |
| 1465 | ABR | 0.40963197 | 0.008904627 | 0.040392504 | 28.23171008 | 14.95736728 | Basal | SOX2 + KLF5 |
| 1466 | TULP4 | 0.409453896 | 0.000992312 | 0.01142308 | 35.80588971 | 22.82858622 | Suprabasal | SOX2 + KLF5 |
| 1467 | DARS2 | 0.408800324 | 0.007423146 | 0.036548425 | 16.0927049 | 8.816252156 | Basal | SOX2 |
| 1468 | TULP3 | 0.40868924 | 0.004211401 | 0.021879278 | 33.2829504 | 21.53384324 | Suprabasal | SOX2 + KLF5, SOX2 |
| 1469 | RAB6A | 0.408482401 | 0.012681065 | 0.033478436 | 64.19858162 | 38.33939519 | Superficial | SOX2 |
| 1470 | NF2 | 0.407599037 | 0.000401828 | 0.012455402 | 24.32252414 | 13.08445788 | Basal | SOX2 |
| 1471 | HNRNPUL1 | 0.404922839 | 0.000795824 | 0.010838271 | 60.58384221 | 41.805875 | Suprabasal | SOX2 |
| 1472 | UBE3A | 0.404126921 | 0.008904627 | 0.025990329 | 52.68022712 | 27.47703569 | Superficial | SOX2 |
| 1473 | DNAJC21 | 0.40361818 | 0.000992312 | 0.008042817 | 59.66645697 | 34.72447459 | Superficial | SOX2 + KLF5, SOX2 |
| 1474 | CREBBP | 0.402571645 | 0.015054848 | 0.048140298 | 39.79409462 | 25.00942118 | Suprabasal | SOX2 |
| 1475 | SMYD4 | 0.402425291 | 0.008904627 | 0.034252903 | 7.546562152 | 4.396569188 | Suprabasal | SOX2 |
| 1476 | FAM204A | 0.399857747 | 0.003462059 | 0.014108599 | 44.29048694 | 20.2946133 | Superficial | SOX2 |
| 1477 | SRBD1 | 0.399456503 | 0.007423146 | 0.030449933 | 14.79261982 | 9.197075729 | Suprabasal | SOX2 |
| 1478 | HLA-E | 0.399386809 | 0.002315114 | 0.020052792 | 91.46666117 | 75.79158387 | Basal | SOX2 |
| 1479 | CUL1 | 0.398192813 | 0.007423146 | 0.030449933 | 40.65802972 | 25.80310235 | Suprabasal | SOX2 |
| 1480 | PSMA5 | 0.398145724 | 0.021001058 | 0.04834985 | 65.47278703 | 39.48031861 | Superficial | SOX2 |
| 1481 | KAT5 | 0.396068142 | 0.005105016 | 0.018023407 | 28.64432152 | 13.29728154 | Superficial | SOX2 |
| 1482 | HDLBP | 0.394837607 | 0.000401828 | 0.012455402 | 79.95998073 | 57.10289635 | Basal | SOX2 |
| 1483 | ABCB7 | 0.392337334 | 0.003462059 | 0.014108599 | 21.58574896 | 8.963636351 | Superficial | SOX2 |
| 1484 | LRRFIP1 | 0.391779281 | 0.000401828 | 0.012455402 | 76.39416407 | 54.39210723 | Basal | SOX2 |
| 1485 | CAB39 | 0.391085547 | 0.008904627 | 0.040392504 | 38.52456553 | 21.76720296 | Basal | SOX2 |
| 1486 | ZNF740 | 0.387806449 | 0.003462059 | 0.019729589 | 11.0501489 | 6.429114307 | Suprabasal | SOX2 |
| 1487 | DNAJC5 | 0.387002103 | 0.006166656 | 0.032981404 | 30.47029786 | 16.50597108 | Basal | SOX2 |
| 1488 | VBP1 | 0.386752618 | 0.021001058 | 0.04834985 | 48.30199504 | 24.9791918 | Superficial | SOX2 |
| 1489 | VGLL4 | 0.386286071 | 0.006166656 | 0.027303057 | 69.20123638 | 49.11019076 | Suprabasal | SOX2 |
| 1490 | PPARD | 0.385509275 | 0.005105016 | 0.029792864 | 13.50772511 | 7.152754309 | Basal | SOX2 |
| 1491 | RAVER2 | 0.385020215 | 0.015054848 | 0.048140298 | 11.98122369 | 7.225062747 | Suprabasal | SOX2 |
| 1492 | IQCK | 0.384096874 | 0.015054848 | 0.048140298 | 7.59743852 | 4.233400157 | Suprabasal | SOX2 |
| 1493 | CAP1 | 0.383283453 | 0.004211401 | 0.027185186 | 70.38203023 | 48.7778019 | Basal | SOX2 |
| 1494 | PRCC | 0.382064623 | 0.001883197 | 0.010234018 | 33.12701812 | 15.6260195 | Superficial | SOX2 |
| 1495 | SPINT2 | 0.381507478 | 0.017811616 | 0.043066688 | 89.89260612 | 77.91907541 | Superficial | SOX2 |
| 1496 | ATP6V1C1 | 0.38106856 | 0.004211401 | 0.021879278 | 29.29032455 | 18.23094114 | Suprabasal | SOX2 |
| 1497 | HNRNPD | 0.379637837 | 0.008904627 | 0.025990329 | 66.22805206 | 39.57404479 | Superficial | SOX2 |
| 1498 | EZH2 | 0.379576368 | 0.004211401 | 0.027185186 | 40.42867394 | 25.49765124 | Basal | SOX2 |
| 1499 | ZFP64 | 0.37948406 | 0.010644787 | 0.038255467 | 12.40083125 | 7.455865285 | Suprabasal | SOX2 |
| 1500 | GNL1 | 0.37843941 | 0.006166656 | 0.020290909 | 31.11718831 | 14.43086912 | Superficial | SOX2 |
| 1501 | CDK12 | 0.378351456 | 0.001526458 | 0.013290882 | 48.53275386 | 31.62280627 | Suprabasal | SOX2 |
| 1502 | NCKAP1 | 0.378170379 | 0.010644787 | 0.044687249 | 58.11956911 | 37.04242786 | Basal | SOX2 |
| 1503 | ZNF672 | 0.376681299 | 0.010644787 | 0.044687249 | 19.56555888 | 10.59380964 | Basal | SOX2 |
| 1504 | POLR2B | 0.37636934 | 0.012681065 | 0.042921737 | 41.3802777 | 24.66201886 | Suprabasal | SOX2 |
| 1505 | GMCL1 | 0.375873197 | 0.007423146 | 0.022959455 | 31.99316097 | 14.72130798 | Superficial | SOX2 |
| 1506 | KCTD5 | 0.375183782 | 0.004211401 | 0.027185186 | 23.89015549 | 13.41836054 | Basal | SOX2 |
| 1507 | GDE1 | 0.372793116 | 0.021001058 | 0.04834985 | 46.2836854 | 23.83445075 | Superficial | SOX2 |
| 1508 | HGS | 0.372464258 | 0.002836074 | 0.022562695 | 37.66991891 | 21.6422198 | Basal | SOX2 |
| 1509 | ACTR1A | 0.372227417 | 0.004211401 | 0.027185186 | 36.13204977 | 21.11663527 | Basal | SOX2 |
| 1510 | SRRT | 0.371491622 | 0.001883197 | 0.014679007 | 37.76365529 | 23.85917939 | Suprabasal | SOX2 |
| 1511 | TCFL5 | 0.371359147 | 0.000506425 | 0.010489143 | 8.353413253 | 4.994880006 | Suprabasal | SOX2 |
| 1512 | ASB6 | 0.367944847 | 0.003462059 | 0.024854712 | 12.57515056 | 6.43489909 | Basal | SOX2 |
| 1513 | DSN1 | 0.367401907 | 0.006166656 | 0.027303057 | 14.87864595 | 9.062953013 | Suprabasal | SOX2 |
| 1514 | KLF5 | 0.366989473 | 0.004211401 | 0.016013117 | 90.73767968 | 78.42717769 | Superficial | KLF5, SOX2 + KLF5 |
| 1515 | UBE3B | 0.365018038 | 0.010644787 | 0.044687249 | 22.00039008 | 11.50400949 | Basal | SOX2 |
| 1516 | CLSPN | 0.364721605 | 0.012681065 | 0.049425568 | 28.84436189 | 17.50717111 | Basal | SOX2 |
| 1517 | EIF4ENIF1 | 0.364297387 | 0.000635975 | 0.010733065 | 22.31296889 | 13.97106774 | Suprabasal | SOX2 |
| 1518 | C6orf132 | 0.363808744 | 0.001232925 | 0.015721789 | 47.99793227 | 30.24246367 | Basal | SOX2 |
| 1519 | TMCC1 | 0.361299072 | 0.002836074 | 0.022562695 | 15.70166472 | 8.402344171 | Basal | SOX2 |
| 1520 | SPATS2L | 0.360680694 | 0.000992312 | 0.01142308 | 80.43917619 | 62.40270925 | Suprabasal | KLF5 |
| 1521 | CDC26 | 0.359851754 | 0.005105016 | 0.018023407 | 53.12997972 | 29.84803495 | Superficial | SOX2 |
| 1522 | HOOK2 | 0.359210707 | 0.021001058 | 0.04834985 | 48.76542349 | 25.87326508 | Superficial | SOX2 |
| 1523 | RASAL2 | 0.357863099 | 0.012681065 | 0.049425568 | 28.13204675 | 15.36324284 | Basal | SOX2 |
| 1524 | GALE | 0.357551358 | 0.010644787 | 0.044687249 | 38.86560774 | 23.09276026 | Basal | SOX2 |
| 1525 | MASTL | 0.355928529 | 0.010644787 | 0.044687249 | 22.26375576 | 12.5982032 | Basal | SOX2 |
| 1526 | CUL4A | 0.354110326 | 0.003462059 | 0.024854712 | 38.8403561 | 23.05650822 | Basal | SOX2 |
| 1527 | TMOD3 | 0.353471008 | 0.010644787 | 0.044687249 | 73.12027102 | 52.13714682 | Basal | SOX2 |
| 1528 | NUDT4 | 0.353268938 | 0.008904627 | 0.034252903 | 30.43480114 | 18.89608606 | Suprabasal | SOX2, SOX2 + KLF5 |
| 1529 | XPO1 | 0.352059571 | 0.015054848 | 0.048140298 | 60.17473282 | 39.24143377 | Suprabasal | SOX2 |
| 1530 | ZNF330 | 0.352054942 | 0.021001058 | 0.04834985 | 33.66922418 | 16.39634589 | Superficial | SOX2 |
| 1531 | TLK2 | 0.351292132 | 0.007423146 | 0.030449933 | 37.89732037 | 23.84071245 | Suprabasal | SOX2 |

| | | | | | | | | |
|---|---|---|---|---|---|---|---|---|
| 1532 | LONP1 | 0.351176435 | 0.010644787 | 0.044687249 | 27.96313576 | 15.97710597 | Basal | SOX2 |
| 1533 | TADA2A | 0.350900882 | 0.004211401 | 0.021879278 | 16.73317655 | 10.39759773 | Suprabasal | SOX2 + KLF5, SOX2 |
| 1534 | UBAP1 | 0.348148108 | 0.012681065 | 0.049425568 | 28.3949319 | 15.61827932 | Basal | SOX2 |
| 1535 | ETF1 | 0.347930763 | 0.012681065 | 0.049425568 | 64.85346179 | 43.8817305 | Basal | SOX2 |
| 1536 | SIN3B | 0.345189095 | 0.010644787 | 0.038255467 | 21.68181576 | 14.24138181 | Suprabasal | SOX2 |
| 1537 | EHMT1 | 0.344749701 | 0.001232925 | 0.015721789 | 34.67135042 | 19.58732315 | Basal | SOX2 |
| 1538 | NOM1 | 0.341602586 | 0.004211401 | 0.027185186 | 29.52543453 | 17.88515075 | Basal | SOX2 |
| 1539 | DCAF7 | 0.341384602 | 0.021001058 | 0.04834985 | 34.97799652 | 16.46057478 | Superficial | SOX2 |
| 1540 | HECTD1 | 0.341197856 | 0.008904627 | 0.040392504 | 55.19918194 | 35.45256668 | Basal | SOX2 |
| 1541 | ETAA1 | 0.34086341 | 0.002836074 | 0.0176927 | 18.78995792 | 11.89426174 | Suprabasal | SOX2 |
| 1542 | FOXK2 | 0.340257957 | 0.000635975 | 0.012925379 | 28.52080745 | 15.8799137 | Basal | SOX2 |
| 1543 | MPRIP | 0.339903944 | 0.012681065 | 0.049425568 | 29.41008776 | 16.52158519 | Basal | SOX2 |
| 1544 | ACTN4 | 0.337577641 | 0.010644787 | 0.044687249 | 82.9768807 | 62.57568151 | Basal | SOX2 |
| 1545 | UBQLN1 | 0.337087049 | 0.007423146 | 0.036548425 | 47.23144812 | 28.48757739 | Basal | SOX2 |
| 1546 | ZNF398 | 0.335352651 | 0.012681065 | 0.049425568 | 11.78243869 | 6.596489406 | Basal | SOX2 |
| 1547 | PGRMC2 | 0.335234595 | 0.021001058 | 0.04834985 | 52.81530241 | 27.35480933 | Superficial | SOX2 |
| 1548 | BAG6 | 0.334098072 | 0.000635975 | 0.012925379 | 45.49164076 | 27.55833727 | Basal | SOX2 |
| 1549 | CD2AP | 0.328979616 | 0.007423146 | 0.036548425 | 55.36500458 | 35.22370908 | Basal | SOX2 |
| 1550 | MARK3 | 0.328532891 | 0.001232925 | 0.015721789 | 39.36693965 | 23.67234087 | Basal | SOX2 |
| 1551 | BTBD10 | 0.324843477 | 0.012681065 | 0.049425568 | 37.09246296 | 21.95821711 | Basal | SOX2 |
| 1552 | RAF1 | 0.324610208 | 0.010644787 | 0.044687249 | 37.95147784 | 22.23152954 | Basal | SOX2 |
| 1553 | SNX9 | 0.323080903 | 0.003462059 | 0.024854712 | 31.57020056 | 18.17767224 | Basal | SOX2 |
| 1554 | RNF126 | 0.321893513 | 0.012681065 | 0.049425568 | 56.94156687 | 36.51439612 | Basal | SOX2 |
| 1555 | NCAPD3 | 0.319375719 | 0.005105016 | 0.029792864 | 37.33730953 | 23.54175813 | Basal | SOX2 |
| 1556 | TPD52L1 | 0.319203489 | 0.012681065 | 0.049425568 | 70.8735422 | 49.83127217 | Basal | SOX2 |
| 1557 | TADA2B | 0.318377235 | 0.007423146 | 0.036548425 | 11.44702194 | 6.08819351 | Basal | SOX2 |
| 1558 | RAD21 | 0.317255801 | 0.007423146 | 0.036548425 | 72.37467016 | 51.73760027 | Basal | SOX2 |
| 1559 | DPP9 | 0.316281208 | 0.005105016 | 0.029792864 | 24.5146002 | 13.78317029 | Basal | SOX2 |
| 1560 | DHX15 | 0.315489463 | 0.010644787 | 0.038255467 | 50.30688361 | 33.87821156 | Suprabasal | SOX2 |
| 1561 | PRRC2C | 0.314883273 | 0.007423146 | 0.030449933 | 87.62866402 | 72.69746995 | Suprabasal | SOX2 |
| 1562 | AKAP8 | 0.314009175 | 0.005105016 | 0.024262694 | 26.62854161 | 16.76378779 | Suprabasal | SOX2 |
| 1563 | DNAAF5 | 0.313393642 | 0.004211401 | 0.027185186 | 25.39436291 | 14.69899341 | Basal | SOX2 |
| 1564 | CSNK1A1 | 0.311878468 | 0.002836074 | 0.0176927 | 93.53449428 | 84.71976112 | Suprabasal | SOX2 |
| 1565 | MIS18BP1 | 0.311634917 | 0.005105016 | 0.024262694 | 39.65562411 | 24.88374556 | Suprabasal | SOX2 |
| 1566 | ANKRD11 | 0.311535163 | 0.005105016 | 0.024262694 | 77.75687683 | 59.74175098 | Suprabasal | SOX2 |
| 1567 | SUPT5H | 0.311462565 | 0.010644787 | 0.044687249 | 41.30614259 | 23.88312125 | Basal | SOX2 |
| 1568 | PRKCH | 0.310775043 | 0.010644787 | 0.044687249 | 12.78922278 | 7.133312113 | Basal | SOX2, SOX2 + KLF5 |
| 1569 | ANXA11 | 0.309510653 | 0.007423146 | 0.036548425 | 80.1395934 | 59.90618267 | Basal | SOX2 + KLF5 |
| 1570 | SMARCA2 | 0.30736207 | 0.012681065 | 0.042921737 | 50.47896445 | 33.01055583 | Suprabasal | SOX2 |
| 1571 | RBM26 | 0.306807667 | 0.004211401 | 0.021879278 | 45.24969731 | 28.33994468 | Suprabasal | SOX2 |
| 1572 | EDC3 | 0.305267304 | 0.003462059 | 0.024854712 | 13.40354087 | 7.180247317 | Basal | SOX2 |
| 1573 | ACTB | 0.304849735 | 0.004211401 | 0.027185186 | 99.71378693 | 97.52395914 | Basal | SOX2 |
| 1574 | RNASEL | 0.304812706 | 0.008904627 | 0.034252903 | 20.4978733 | 13.22398589 | Suprabasal | SOX2 |
| 1575 | PRRG4 | 0.303894969 | 0.006166656 | 0.027303057 | 83.56882003 | 69.16067751 | Suprabasal | SOX2, SOX2 + KLF5 |
| 1576 | CNDP2 | 0.301775633 | 0.007423146 | 0.036548425 | 41.87890185 | 25.30966965 | Basal | SOX2 |
| 1577 | PPM1A | 0.297552231 | 0.010644787 | 0.044687249 | 31.80766749 | 17.79182567 | Basal | SOX2 |
| 1578 | PCM1 | 0.29663167 | 0.012681065 | 0.042921737 | 62.41064601 | 43.03776245 | Suprabasal | SOX2 |
| 1579 | XRCC1 | 0.296049831 | 0.012681065 | 0.042921737 | 16.1532907 | 10.76767722 | Suprabasal | SOX2 |
| 1580 | PTBP1 | 0.294116815 | 0.005105016 | 0.029792864 | 65.2690628 | 44.63874797 | Basal | SOX2 |
| 1581 | RNF40 | 0.292210922 | 0.002836074 | 0.022562695 | 23.01616376 | 12.95846387 | Basal | SOX2 |
| 1582 | INTS14 | 0.291042279 | 0.012681065 | 0.042921737 | 21.96252873 | 13.69680143 | Suprabasal | SOX2 |
| 1583 | UBTF | 0.289426243 | 0.004211401 | 0.021879278 | 55.33188696 | 37.74718139 | Suprabasal | SOX2 |
| 1584 | BDH1 | 0.287800194 | 0.001526458 | 0.016792744 | 37.68427329 | 23.60581804 | Basal | SOX2 |
| 1585 | ZNF174 | 0.285899464 | 0.006166656 | 0.027303057 | 6.81185006 | 4.65851935 | Suprabasal | SOX2 |
| 1586 | TAX1BP1 | 0.28569555 | 0.006166656 | 0.032981404 | 70.90543279 | 49.84024475 | Basal | SOX2 |
| 1587 | OPA3 | 0.285098991 | 0.015054848 | 0.048140298 | 18.78897384 | 12.76558836 | Suprabasal | SOX2 |
| 1588 | KPNA4 | 0.284385351 | 0.008904627 | 0.040392504 | 53.03150064 | 33.71170895 | Basal | SOX2 |
| 1589 | CSNK1D | 0.282246278 | 0.006166656 | 0.032981404 | 38.6253149 | 23.71317908 | Basal | SOX2 |
| 1590 | C6orf136 | 0.282069153 | 0.003462059 | 0.024854712 | 25.12174217 | 14.50978536 | Basal | SOX2 + KLF5 |
| 1591 | KDM2A | 0.281821164 | 0.005105016 | 0.029792864 | 44.65559244 | 26.94707233 | Basal | SOX2 |
| 1592 | NFX1 | 0.281778885 | 0.008904627 | 0.034252903 | 30.94818377 | 19.69308165 | Suprabasal | SOX2 |
| 1593 | KAT7 | 0.280513646 | 0.003462059 | 0.019729589 | 31.96355226 | 20.50682651 | Suprabasal | SOX2 |
| 1594 | ZNF623 | 0.278474992 | 0.005105016 | 0.024262694 | 10.25246344 | 6.504611864 | Suprabasal | SOX2 |
| 1595 | PSMD2 | 0.278455513 | 0.017811616 | 0.043066688 | 64.95891017 | 40.28973587 | Superficial | SOX2 |
| 1596 | SYNRG | 0.277856083 | 0.008904627 | 0.040392504 | 27.18815891 | 15.59211998 | Basal | SOX2 |
| 1597 | CDKN2AIP | 0.27405597 | 0.003462059 | 0.024854712 | 48.78322567 | 32.15435898 | Basal | SOX2 |
| 1598 | APLP2 | 0.270292897 | 0.012681065 | 0.042921737 | 73.82533826 | 57.93670449 | Suprabasal | SOX2 |
| 1599 | SF3A1 | 0.268950282 | 0.002315114 | 0.020052792 | 40.45425161 | 24.11803561 | Basal | SOX2 |
| 1600 | DNAJC9 | 0.267541983 | 0.008904627 | 0.034252903 | 36.74656251 | 23.59342056 | Suprabasal | SOX2 |
| 1601 | PAWR | 0.266458622 | 0.010644787 | 0.044687249 | 63.09732871 | 43.42629148 | Basal | SOX2 |
| 1602 | CTCF | 0.266102688 | 0.010644787 | 0.038255467 | 50.85548206 | 33.85027116 | Suprabasal | SOX2 |
| 1603 | SMC3 | 0.265657661 | 0.010644787 | 0.038255467 | 66.24412926 | 45.72828955 | Suprabasal | SOX2 |
| 1604 | PAFAH1B2 | 0.265099423 | 0.006166656 | 0.032981404 | 49.89534049 | 31.20887734 | Basal | SOX2 |

| | | | | | | | | |
|---|---|---|---|---|---|---|---|---|
| 1605 | DDA1 | 0.264148216 | 0.004211401 | 0.027185186 | 39.59917761 | 24.07286753 | Basal | SOX2 |
| 1606 | BRD4 | 0.260587942 | 0.010644787 | 0.044687249 | 59.06233998 | 38.69743774 | Basal | SOX2 |
| 1607 | EPN1 | 0.259066852 | 0.004211401 | 0.027185186 | 52.19066592 | 33.714443 | Basal | SOX2 |
| 1608 | NFKBIB | 0.258046044 | 0.010644787 | 0.044687249 | 27.74931509 | 16.39772077 | Basal | SOX2 |
| 1609 | ELF1 | 0.257772111 | 0.006166656 | 0.027303057 | 78.78518903 | 61.89883688 | Suprabasal | SOX2 |
| 1610 | SAFB2 | 0.256603266 | 0.006166656 | 0.027303057 | 33.63209571 | 21.6889824 | Suprabasal | SOX2 |
| 1611 | SPINT1 | 0.251117707 | 0.008904627 | 0.040392504 | 62.93750767 | 43.57976501 | Basal | SOX2 |
| 1612 | LSM12 | 0.248991693 | 0.002836074 | 0.022562695 | 57.64303137 | 37.74858485 | Basal | SOX2 |
| 1613 | RAD23B | 0.233829178 | 0.012681065 | 0.049425568 | 69.23893558 | 49.60189286 | Basal | SOX2 |
| 1614 | ANP32E | 0.232016051 | 0.012681065 | 0.049425568 | 57.77117902 | 40.06006791 | Basal | SOX2 |
| 1615 | RBM15B | 0.231504327 | 0.008904627 | 0.040392504 | 28.6956174 | 17.30885728 | Basal | SOX2 |
| 1616 | RIOK1 | 0.229649382 | 0.012681065 | 0.049425568 | 45.19160897 | 29.09861844 | Basal | SOX2 |
| 1617 | BSG | 0.229322399 | 0.008904627 | 0.040392504 | 86.2205218 | 68.15708431 | Basal | SOX2 |
| 1618 | TCERG1 | 0.229183326 | 0.010644787 | 0.044687249 | 47.17707217 | 30.39663149 | Basal | SOX2 |
| 1619 | BAZ1A | 0.226783923 | 0.005105016 | 0.029792864 | 67.80402219 | 47.92001143 | Basal | SOX2 |
| 1620 | RAP2A | 0.219584981 | 0.012681065 | 0.049425568 | 20.63551863 | 12.4165553 | Basal | SOX2, SOX2 + KLF5 |
| 1621 | SEC23B | 0.206323525 | 0.008904627 | 0.040392504 | 26.53475501 | 15.84923426 | Basal | SOX2 |
| 1622 | USP16 | 0.202143273 | 0.006166656 | 0.020290909 | 55.11025916 | 31.92112904 | Superficial | SOX2 |
| 1623 | LUC7L | 0.193345812 | 0.017811616 | 0.043066688 | 24.87083467 | 12.73663129 | Superficial | SOX2 |
| 1624 | FBXO7 | 0.178143875 | 0.010644787 | 0.038255467 | 43.93005609 | 30.99583959 | Suprabasal | SOX2 |
| 1625 | CNOT2 | 0.164714044 | 0.015054848 | 0.048140298 | 57.84791185 | 42.11150241 | Suprabasal | SOX2 |

**Supplemental Table 8.** Gene expression profile of EoE DEGs known to be regulated by the SOX2-KLF5 interaction.

| | gene | Log2FC | Pval | FDR_Pval | Compartment |
|---|---|---|---|---|---|
| 1 | NRXN1 | 5.011065 | 0.000184 | 0.010398 | Suprabasal |
| 2 | NFATC2 | 4.981012 | 0.000989 | 0.008043 | Superficial |
| 3 | KCNJ2 | 4.494243 | 0.000506 | 0.010398 | Suprabasal |
| 4 | BCL2L15 | 4.350032 | 0.000154 | 0.010398 | Suprabasal |
| 5 | RUNX2 | 3.769129 | 0.000505 | 0.007179 | Superficial |
| 6 | LYPD6B | 3.712118 | 0.004211 | 0.016013 | Superficial |
| 7 | SGK1 | 3.680086 | 0.000154 | 0.007176 | Superficial |
| 8 | ST8SIA6 | 3.5861 | 0.000634 | 0.007275 | Superficial |
| 9 | CMYA5 | 3.552817 | 0.001526 | 0.009387 | Superficial |
| 10 | HRH1 | 3.502635 | 0.000992 | 0.011395 | Suprabasal |
| 11 | TNS4 | 2.993108 | 0.001233 | 0.00868 | Superficial |
| 12 | RASGRP1 | 2.869247 | 0.000506 | 0.010398 | Suprabasal |
| 13 | LY75 | 2.675558 | 5.55E-05 | 0.007176 | Superficial |
| 14 | HS3ST3A1 | 2.549606 | 0.003462 | 0.014109 | Superficial |
| 15 | HS3ST1 | 2.54417 | 0.000154 | 0.010398 | Suprabasal |
| 16 | RARB | 2.530013 | 0.000992 | 0.008043 | Superficial |
| 17 | CLDN1 | 2.4188 | 0.000506 | 0.007179 | Superficial |
| 18 | LITAF | 2.418478 | 0.000402 | 0.010398 | Suprabasal |
| 19 | NRP2 | 2.366104 | 0.001233 | 0.012403 | Suprabasal |
| 20 | SOX21 | 2.296942 | 0.000506 | 0.007179 | Superficial |
| 21 | GGH | 2.238839 | 0.000318 | 0.007176 | Superficial |
| 22 | UTRN | 2.217242 | 0.000196 | 0.007176 | Superficial |
| 23 | MYOF | 2.181333 | 0.000796 | 0.007615 | Superficial |
| 24 | GIPC2 | 2.130797 | 0.00012 | 0.007176 | Superficial |
| 25 | MYH9 | 2.016182 | 0.000402 | 0.007176 | Superficial |
| 26 | PDLIM4 | 1.989624 | 0.000318 | 0.007176 | Superficial |
| 27 | NUAK2 | 1.982932 | 0.001233 | 0.00868 | Superficial |
| 28 | LPCAT4 | 1.972938 | 0.000796 | 0.007615 | Superficial |
| 29 | LRRC8B | 1.948757 | 0.001526 | 0.009387 | Superficial |
| 30 | TLE1 | 1.923102 | 0.00012 | 0.007176 | Superficial |
| 31 | SGMS2 | 1.870786 | 0.000402 | 0.007176 | Superficial |
| 32 | FMN1 | 1.866941 | 0.000992 | 0.008043 | Superficial |
| 33 | GCNT3 | 1.859067 | 0.002315 | 0.016077 | Suprabasal |
| 34 | SERPINB5 | 1.819859 | 0.000506 | 0.007179 | Superficial |
| 35 | ARHGAP26 | 1.751173 | 7.20E-05 | 0.007176 | Superficial |
| 36 | ADAM9 | 1.728379 | 0.001233 | 0.012403 | Suprabasal |
| 37 | IGF2R | 1.719943 | 0.000992 | 0.008043 | Superficial |
| 38 | ATP8B1 | 1.711098 | 9.30E-05 | 0.010398 | Suprabasal |
| 39 | FAM3C | 1.692974 | 0.000992 | 0.008043 | Superficial |
| 40 | CD9 | 1.664881 | 0.000318 | 0.007176 | Superficial |
| 41 | DSG3 | 1.659685 | 0.000402 | 0.007176 | Superficial |
| 42 | SERPINB2 | 1.591511 | 0.000636 | 0.010421 | Suprabasal |
| 43 | MACF1 | 1.585173 | 0.001883 | 0.010234 | Superficial |
| 44 | TMPRSS11I | 1.540082 | 0.008905 | 0.040393 | Basal |
| 45 | NRIP1 | 1.527775 | 0.000992 | 0.008043 | Superficial |
| 46 | SETBP1 | 1.526392 | 0.000796 | 0.007615 | Superficial |
| 47 | MYH14 | 1.506037 | 0.000636 | 0.007275 | Superficial |
| 48 | MCC | 1.495164 | 0.001233 | 0.00868 | Superficial |
| 49 | SLC16A7 | 1.49266 | 0.000318 | 0.007176 | Superficial |
| 50 | EEA1 | 1.47964 | 0.000196 | 0.007176 | Superficial |

| | | | | | |
|---|---|---|---|---|---|
| 51 | KLF4 | 1.451647 | 0.000318 | 0.007176 | Superficial |
| 52 | RAB11FIP1 | 1.442201 | 0.000318 | 0.012455 | Basal |
| 53 | ETS2 | 1.44046 | 0.000318 | 0.010398 | Suprabasal |
| 54 | TJP2 | 1.428787 | 0.00025 | 0.012455 | Basal |
| 55 | FAM83B | 1.409211 | 0.005105 | 0.018023 | Superficial |
| 56 | ANGPTL4 | 1.390768 | 9.30E-05 | 0.012455 | Basal |
| 57 | OCIAD2 | 1.316919 | 0.000796 | 0.007615 | Superficial |
| 58 | ARHGAP18 | 1.313209 | 0.001526 | 0.009387 | Superficial |
| 59 | ERC1 | 1.308853 | 0.000992 | 0.008043 | Superficial |
| 60 | SOX13 | 1.301876 | 0.004211 | 0.016013 | Superficial |
| 61 | TMPRSS4 | 1.291648 | 0.000318 | 0.012455 | Basal |
| 62 | PDE4D | 1.288243 | 0.003462 | 0.014109 | Superficial |
| 63 | SHROOM2 | 1.258271 | 0.000636 | 0.007275 | Superficial |
| 64 | MREG | 1.217755 | 0.000402 | 0.012455 | Basal |
| 65 | MBOAT2 | 1.213442 | 0.000796 | 0.013643 | Basal |
| 66 | ROCK2 | 1.208618 | 0.001233 | 0.00868 | Superficial |
| 67 | LIMA1 | 1.205073 | 0.001233 | 0.00868 | Superficial |
| 68 | SLC25A24 | 1.202476 | 0.002836 | 0.012662 | Superficial |
| 69 | MYO10 | 1.191689 | 0.000636 | 0.007275 | Superficial |
| 70 | AFF1 | 1.191218 | 0.001526 | 0.009387 | Superficial |
| 71 | MLLT3 | 1.184495 | 0.002836 | 0.012662 | Superficial |
| 72 | MEIS1 | 1.17702 | 0.000992 | 0.008043 | Superficial |
| 73 | PARP12 | 1.170055 | 0.000636 | 0.012925 | Basal |
| 74 | RAPGEF5 | 1.16116 | 0.000402 | 0.007176 | Superficial |
| 75 | BARX2 | 1.154142 | 0.000318 | 0.007176 | Superficial |
| 76 | TP63 | 1.146853 | 0.006167 | 0.020291 | Superficial |
| 77 | B4GALNT3 | 1.142849 | 0.012681 | 0.033478 | Superficial |
| 78 | RPS6KA3 | 1.13976 | 0.001233 | 0.00868 | Superficial |
| 79 | ROBO2 | 1.132995 | 0.007423 | 0.022959 | Superficial |
| 80 | FOXQ1 | 1.123032 | 0.000992 | 0.011395 | Suprabasal |
| 81 | ACVR1 | 1.112086 | 0.010645 | 0.029507 | Superficial |
| 82 | SLC5A3 | 1.108546 | 0.017812 | 0.043067 | Superficial |
| 83 | EPHB6 | 1.098421 | 0.002315 | 0.011381 | Superficial |
| 84 | CRYZ | 1.092202 | 0.001883 | 0.010234 | Superficial |
| 85 | TGFBR1 | 1.082553 | 0.017812 | 0.043067 | Superficial |
| 86 | VCL | 1.068577 | 0.000402 | 0.010398 | Suprabasal |
| 87 | IFIH1 | 1.055273 | 0.002315 | 0.016077 | Suprabasal |
| 88 | RAB11FIP2 | 1.05154 | 0.004211 | 0.016013 | Superficial |
| 89 | CLDN4 | 1.046614 | 0.003462 | 0.024855 | Basal |
| 90 | GBP2 | 1.043537 | 0.006167 | 0.020291 | Superficial |
| 91 | PDZRN3 | 1.038845 | 0.005105 | 0.018023 | Superficial |
| 92 | GJB5 | 1.036529 | 0.012681 | 0.033478 | Superficial |
| 93 | USP12 | 1.026388 | 0.001233 | 0.012403 | Suprabasal |
| 94 | LMTK2 | 0.992633 | 0.002315 | 0.016077 | Suprabasal |
| 95 | ITGB8 | 0.982346 | 0.012681 | 0.043322 | Suprabasal |
| 96 | RNF19A | 0.979825 | 0.00025 | 0.010398 | Suprabasal |
| 97 | EFNB2 | 0.971029 | 0.002315 | 0.011381 | Superficial |
| 98 | CCDC88C | 0.960891 | 0.008905 | 0.02599 | Superficial |
| 99 | MBNL3 | 0.951778 | 0.005105 | 0.018023 | Superficial |
| 100 | GPHN | 0.950352 | 0.005105 | 0.018023 | Superficial |
| 101 | PKP1 | 0.940526 | 0.005105 | 0.018023 | Superficial |
| 102 | ATXN1 | 0.939973 | 0.008905 | 0.02599 | Superficial |

| | | | | | |
|---|---|---|---|---|---|
| 103 | TNK2 | 0.921896 | 0.005105 | 0.02473 | Suprabasal |
| 104 | SMPD3 | 0.918832 | 0.012681 | 0.033478 | Superficial |
| 105 | ANK3 | 0.913275 | 0.012681 | 0.033478 | Superficial |
| 106 | SLC4A7 | 0.9 | 0.017812 | 0.043067 | Superficial |
| 107 | ST14 | 0.895367 | 7.20E-05 | 0.012455 | Basal |
| 108 | EPN2 | 0.893136 | 0.000796 | 0.007615 | Superficial |
| 109 | KRT19 | 0.89218 | 0.012681 | 0.043322 | Suprabasal |
| 110 | BMPR2 | 0.892138 | 0.001883 | 0.010234 | Superficial |
| 111 | TNRC18 | 0.889206 | 0.017812 | 0.043067 | Superficial |
| 112 | FAM110C | 0.886645 | 0.000506 | 0.010398 | Suprabasal |
| 113 | LMO7 | 0.886093 | 0.002315 | 0.016077 | Suprabasal |
| 114 | NOL10 | 0.882089 | 0.015055 | 0.03801 | Superficial |
| 115 | RNF19B | 0.881491 | 0.000506 | 0.012899 | Basal |
| 116 | UGCG | 0.880956 | 0.000318 | 0.012455 | Basal |
| 117 | BTG1 | 0.874641 | 7.20E-05 | 0.007176 | Superficial |
| 118 | ALCAM | 0.848191 | 0.007423 | 0.022959 | Superficial |
| 119 | DTNBP1 | 0.846795 | 0.021001 | 0.04835 | Superficial |
| 120 | MAP3K8 | 0.845745 | 0.004211 | 0.027185 | Basal |
| 121 | MACC1 | 0.834418 | 0.001883 | 0.018415 | Basal |
| 122 | DNMBP | 0.834005 | 0.017812 | 0.043067 | Superficial |
| 123 | NDFIP2 | 0.829957 | 0.000636 | 0.010421 | Suprabasal |
| 124 | DSC3 | 0.826851 | 0.021001 | 0.04835 | Superficial |
| 125 | TWF2 | 0.823549 | 0.000992 | 0.008043 | Superficial |
| 126 | PPL | 0.822802 | 0.000318 | 0.012455 | Basal |
| 127 | ACAD9 | 0.822606 | 0.000402 | 0.007176 | Superficial |
| 128 | PIM1 | 0.816683 | 0.000506 | 0.012899 | Basal |
| 129 | TNIP1 | 0.799214 | 0.000506 | 0.010398 | Suprabasal |
| 130 | BACE2 | 0.792111 | 0.000506 | 0.010398 | Suprabasal |
| 131 | PLS1 | 0.792064 | 0.002836 | 0.018003 | Suprabasal |
| 132 | KCTD1 | 0.79082 | 0.000796 | 0.007615 | Superficial |
| 133 | KLF3 | 0.788074 | 0.000402 | 0.010398 | Suprabasal |
| 134 | DSP | 0.784191 | 0.000506 | 0.012899 | Basal |
| 135 | RASSF6 | 0.780974 | 0.021001 | 0.04835 | Superficial |
| 136 | IL15 | 0.76698 | 0.007423 | 0.030891 | Suprabasal |
| 137 | CRADD | 0.748668 | 0.007423 | 0.022959 | Superficial |
| 138 | FAM53B | 0.742226 | 0.002836 | 0.018003 | Suprabasal |
| 139 | ACOX1 | 0.738899 | 0.007423 | 0.030891 | Suprabasal |
| 140 | ELF3 | 0.738836 | 0.003462 | 0.019935 | Suprabasal |
| 141 | ARFGAP3 | 0.737415 | 0.004211 | 0.016013 | Superficial |
| 142 | ZFAND5 | 0.732781 | 0.000636 | 0.010421 | Suprabasal |
| 143 | EZR | 0.727427 | 0.002315 | 0.020053 | Basal |
| 144 | KHDRBS3 | 0.710488 | 0.000636 | 0.012925 | Basal |
| 145 | MIPEP | 0.707167 | 0.021001 | 0.04835 | Superficial |
| 146 | SHROOM3 | 0.695336 | 0.007423 | 0.036548 | Basal |
| 147 | FCHSD2 | 0.691176 | 0.007423 | 0.030891 | Suprabasal |
| 148 | TXNRD1 | 0.681193 | 0.004211 | 0.022366 | Suprabasal |
| 149 | SGMS1 | 0.679916 | 0.00025 | 0.010398 | Suprabasal |
| 150 | ANXA11 | 0.668315 | 0.000402 | 0.010398 | Suprabasal |
| 151 | MAL2 | 0.666972 | 0.000992 | 0.014321 | Basal |
| 152 | B3GNT2 | 0.659309 | 0.015055 | 0.03801 | Superficial |
| 153 | ILVBL | 0.64809 | 0.021001 | 0.04835 | Superficial |
| 154 | PACS1 | 0.624994 | 0.000636 | 0.012925 | Basal |

| | | | | | |
|---|---|---|---|---|---|
| 155 | RIN2 | 0.623406 | 0.002836 | 0.018003 | Suprabasal |
| 156 | TTC39C | 0.619828 | 0.017812 | 0.043067 | Superficial |
| 157 | NDUFA4 | 0.617621 | 0.007423 | 0.022959 | Superficial |
| 158 | LMNA | 0.616183 | 0.002315 | 0.011381 | Superficial |
| 159 | JUN | 0.612848 | 0.010645 | 0.029507 | Superficial |
| 160 | NCOR2 | 0.603984 | 0.005105 | 0.02473 | Suprabasal |
| 161 | RAP2B | 0.602605 | 0.012681 | 0.043322 | Suprabasal |
| 162 | MGAT4A | 0.589002 | 0.00025 | 0.010398 | Suprabasal |
| 163 | HS6ST1 | 0.587523 | 0.001883 | 0.01459 | Suprabasal |
| 164 | IPMK | 0.585907 | 0.008905 | 0.034888 | Suprabasal |
| 165 | PAIP2B | 0.580739 | 0.021001 | 0.04835 | Superficial |
| 166 | SPPL3 | 0.573635 | 0.003462 | 0.014109 | Superficial |
| 167 | TULP3 | 0.561441 | 0.000154 | 0.010398 | Suprabasal |
| 168 | RAB27B | 0.555356 | 0.002315 | 0.011381 | Superficial |
| 169 | RERE | 0.55374 | 0.002315 | 0.016077 | Suprabasal |
| 170 | RRP12 | 0.55344 | 0.001233 | 0.015722 | Basal |
| 171 | JUP | 0.548641 | 0.000796 | 0.013643 | Basal |
| 172 | TRIP10 | 0.52718 | 0.000796 | 0.013643 | Basal |
| 173 | ANXA1 | 0.526355 | 0.010645 | 0.044687 | Basal |
| 174 | MPZL1 | 0.508728 | 0.008905 | 0.02599 | Superficial |
| 175 | EHF | 0.492196 | 0.010645 | 0.03891 | Suprabasal |
| 176 | TRIM33 | 0.48945 | 0.017812 | 0.043067 | Superficial |
| 177 | AHNAK | 0.472581 | 0.000992 | 0.014321 | Basal |
| 178 | THUMPD2 | 0.452637 | 0.021001 | 0.04835 | Superficial |
| 179 | STK38 | 0.448032 | 0.001526 | 0.013166 | Suprabasal |
| 180 | FNBP1 | 0.44158 | 0.003462 | 0.019935 | Suprabasal |
| 181 | EIF4G3 | 0.421872 | 0.012681 | 0.033478 | Superficial |
| 182 | PLD1 | 0.416621 | 0.010645 | 0.044687 | Basal |
| 183 | BAIAP2L1 | 0.414982 | 0.000992 | 0.014321 | Basal |
| 184 | SMG7 | 0.408191 | 0.015055 | 0.048244 | Suprabasal |
| 185 | DNAJC21 | 0.403618 | 0.000992 | 0.008043 | Superficial |
| 186 | N4BP1 | 0.398064 | 0.000992 | 0.011395 | Suprabasal |
| 187 | TULP4 | 0.387329 | 0.001526 | 0.013166 | Suprabasal |
| 188 | SETD5 | 0.37761 | 0.005105 | 0.02473 | Suprabasal |
| 189 | CAPN2 | 0.368913 | 0.015055 | 0.048244 | Suprabasal |
| 190 | KLF5 | 0.366989 | 0.004211 | 0.016013 | Superficial |
| 191 | PARD3 | 0.323073 | 0.002836 | 0.018003 | Suprabasal |
| 192 | PHACTR4 | 0.321553 | 0.015055 | 0.048244 | Suprabasal |
| 193 | PRKCH | 0.310775 | 0.010645 | 0.044687 | Basal |
| 194 | C6orf136 | 0.282069 | 0.003462 | 0.024855 | Basal |
| 195 | ASAH1 | -0.26554 | 0.008905 | 0.040393 | Basal |
| 196 | IL20RB | -0.32366 | 0.012681 | 0.043322 | Suprabasal |
| 197 | CTSZ | -0.33085 | 0.010645 | 0.044687 | Basal |
| 198 | SLC1A4 | -0.54385 | 0.004211 | 0.027185 | Basal |
| 199 | MPPED2 | -0.68819 | 0.012681 | 0.049426 | Basal |
| 200 | PTGR1 | -0.71596 | 0.001883 | 0.018415 | Basal |
| 201 | TGFBI | -0.82612 | 0.001526 | 0.016793 | Basal |
| 202 | SPRY1 | -0.83185 | 0.008905 | 0.034888 | Suprabasal |
| 203 | FHL2 | -0.95479 | 0.000318 | 0.012455 | Basal |
| 204 | EPCAM | -1.00287 | 0.021001 | 0.04835 | Superficial |
| 205 | LFNG | -1.0368 | 0.000992 | 0.014321 | Basal |
| 206 | ANKH | -1.06814 | 0.00012 | 0.012455 | Basal |

| | | | | | |
|---|---|---|---|---|---|
| 207 | ARMCX1 | -1.10077 | 0.000318 | 0.012455 | Basal |
| 208 | SORBS2 | -1.12594 | 0.000636 | 0.012925 | Basal |
| 209 | CYYR1 | -1.22211 | 0.001883 | 0.018415 | Basal |
| 210 | PFN2 | -1.24968 | 0.000796 | 0.013643 | Basal |
| 211 | EFEMP1 | -1.26834 | 0.000196 | 0.012455 | Basal |
| 212 | TGM3 | -1.39805 | 0.001883 | 0.01459 | Suprabasal |
| 213 | SYT17 | -1.41193 | 0.008905 | 0.02599 | Superficial |
| 214 | GHR | -1.42445 | 0.00025 | 0.012455 | Basal |
| 215 | STXBP1 | -1.60505 | 0.006167 | 0.020291 | Superficial |
| 216 | ALOX12 | -1.64626 | 0.012681 | 0.043322 | Suprabasal |
| 217 | NGEF | -1.69712 | 0.000992 | 0.011395 | Suprabasal |
| 218 | ZNF365 | -1.72312 | 0.008905 | 0.02599 | Superficial |
| 219 | PLB1 | -1.80572 | 0.001883 | 0.010234 | Superficial |
| 220 | DHRS9 | -2.3798 | 0.00025 | 0.007176 | Superficial |
| 221 | TGM7 | -2.41928 | 0.010171 | 0.029507 | Superficial |
| 222 | IL1A | -2.49769 | 0.003462 | 0.014109 | Superficial |
| 223 | RAI14 | -2.77967 | 0.000318 | 0.007176 | Superficial |
| 224 | SULT1B1 | -3.33906 | 0.000183 | 0.007176 | Superficial |

**Supplemental Table 9.** List of antibodies used in the study.

| Antibody | Company | Catalog # | Assay | Concentration | Secondary / Conjugate |
|---|---|---|---|---|---|
| anti-KI-67 | Abcam | 16667 | IHC | 1:100 | goat anti-rabbit |
| anti-SOX2 | Cell Signaling | 14962S | IHC | 1:500 | goat anti-rabbit |
| anti-KLF5 | Sigma-Aldrich | HPA040398 | IHC | 1:150 | goat anti-rabbit |
| anti-KRT14 | Cell Signaling | 48020 | Multispectral IF | 1:200 | Opal 540 |
| anti-P63 | ABclonal | A19652 | Multispectral IF | 1:100 | Opal 690 |
| anti-IVL | Santa Cruz Biotechnology | sc-21748 | Multispectral IF | 1:500 | Opal 520 |
| anti-CNFN | Novus Biologicals | NBP2-14668 | Multispectral IF | 1:200 | Opal 570 |
